# Investigation of the Fasciola Cinereum, Absent in BTBR mice, and Comparison with the Hippocampal Area CA2

**DOI:** 10.1101/2024.03.21.586108

**Authors:** Su Hyun Lee, Michaela E Cooke, Kai Zheng Duan, Sarah K Williams Avram, June Song, Abdel G Elkahloun, George McGrady, NISC Comparative Sequencing Program, Austin Howley, Babru Samal, W. Scott Young

## Abstract

The arginine vasopressin 1b receptor (Avpr1b) plays an important role in social behaviors including social learning, memory, and aggression, and is known to be a specific marker for the cornu ammonis area 2 (CA2) regions of the hippocampus. The fasciola cinereum (FC) is an anatomical region in which Avpr1b expressing neurons are prominent, but the functional roles of the FC have yet to be investigated. Surprisingly, the FC is absent in the inbred BTBR T+tf/J (BTBR) mouse strain used to study core behavioral deficits of autism. Here, we characterized and compared transcriptomic expression profiles using single nucleus RNA sequencing and identified 7 different subpopulations and heterogeneity within the dorsal CA2 (dCA2) and FC. *Mef2c,* involved in autism spectrum disorder, is more highly expressed in the FC. Using Hiplex *in situ* hybridization, we examined the neuroanatomical locations of these subpopulations in the proximal and distal regions of the hippocampus. Anterograde tracing of Avpr1b neurons specific for the FC showed projections to the IG, dCA2, lacunosum molecular layer of CA1, dorsal fornix, septofibrial nuclei, and intermediate lateral septum (iLS). In contrast to the dCA2, inhibition of Avpr1b neurons in the FC by the inhibitory DREADD system during behavioral testing did not impair social memory. We performed single nucleus RNA sequencing in the dCA2 region and compared between wildtype (WT) and BTBR mice. We found that transcriptomic profiles of dCA2 neurons between BTBR and WT mice are very similar as they did not form any unique clusters; yet, we found there were differentially expressed genes between the dCA2s of BTBR and WT mice. Overall, this is a comprehensive study of the comparison of Avpr1b neuronal subpopulations between the FC and dCA2. The fact that FC is absent in BTBR mice, a mouse model for autism spectrum disorder, suggests that the FC may play a role in understanding neuropsychiatric disease.

## Introduction

The arginine vasopressin 1b receptor (Avpr1b) plays an important role in social behaviors including social learning, memory, and aggression (Wersinger et al., 2002), and is known to be a specific marker for the cornu ammonis area 2 (CA2) regions of the hippocampus (Young et al., 2006). In mice, inhibiting dorsal CA2 (dCA2) pyramidal neurons causes loss of ability to remember a conspecific (Hitti & Siegelbaum, 2014; Stevenson & Caldwell, 2014), while enhancing vasopressin presynaptic projection to dCA2 results in enhancement of social memory (Smith et al., 2016).

The fasciola cinereum (FC) is an anatomical region in which Avpr1b expressing neurons are prominent, but the functional roles of the FC have yet to be investigated. The FC appears to be a remnant of the original dorsally located hippocampal anlage: a continuation of the dorsal taenia tecta and induseum griseum (IG) and an anterior and dorsal hippocampal rudiment.

Surprisingly, the FC is absent in the inbred BTBR T+tf/J (BTBR) mouse strain used to study core behavioral deficits of autism (Young and Song, unpublished). Originally bred to study insulin-resistance, it displays strong behavioral characteristics of autism spectrum disorder which include high levels of repetitive self-grooming behavior (Silverman et al., 2010), deficits in social interactions (Bolivar et al., 2007; Endo et al., 2019; Meyza & Blanchard, 2017; Moy et al., 2007) and an unusual repertoire of vocalizations (Scattoni et al., 2008).

Here, we characterized and compared expression profiles and identified subpopulations and heterogeneity within the dCA2 and FC. We examined and compared their transcriptional and cell type profiles using RNA sequencing from single nuclei. We were interested in unique genes expressed in the FC vs. dCA2 and how their expression profiles relate to the subpopulation of Avpr1b neurons. Using Hiplex *in situ* hybridization, we examined the neuroanatomical locations of these subpopulations in the proximal and distal regions of the dorsal hippocampus. We also further investigated the anterior projection of Avpr1b neurons of the FC as well as their functional roles in memory formation and social behavior. In addition, as the FC is absent in BTBR mice, we compared the transcriptional profiles of Avpr1b neurons in the dCA2 between BTBR and wildtype (WT) mice.

## Methods

### Mouse Housing Conditions

All housing and procedures were approved by the Animal Care and Use Committee of the National Institute of Mental Health. Mice were housed in an AAALAC accredited, specific pathogen-free vivarium kept at constant temperature and humidity (∼21°C, 50%), in plastic micro-isolator cages (12”×6.5”×5.5”) containing wood chip bedding (Nepco Beta-chips) and cotton nestlets. All cages were maintained on high-density ventilated racks (Super Mouse 750, Lab Products Inc.). Mice were maintained on a 12-h light cycle (lights off at 1500h) with *ad libitum* access to standard mouse chow (Purina Lab Diet, Product #5R31) and water bottles. Cages were changed on a bi-weekly basis primarily by the same animal caretaker. All breeding pairs were fed a high-fat diet (Purina Lab Diet, Product #5058) for healthier pregnancies and pups. All offspring were weaned at ∼21 days of age into cages with their same-sex littermates. No animals that were singly housed during adolescence were used in behavioral testing.

### Generation of the *Avpr1b-cre x* B6;129GT(ROSA)26Sor^tm5(CAG-SUN1/sGFP)Nat/^J mice mouse line

8-week old Avpr1bCre^Avpr1b/cre^(Avpr1b-cre) knock-in female mice generated in house (Williams Avram et al., 2019) were bred with homozygous B6;129GT(ROSA)26Sor^tm5(CAG-SUN1/sGFP)Nat/^J (SUN1GFP) male mice (Strain #021039, The Jackson Laboratory, Bar Harbor, ME, USA) (Mo et al., 2015) to generate Avpr1b-cre x SUN1GFP mice. In these mice, GFP is expressed in the nuclei of Avpr1b-expressing neurons (Supplemental Figure 1).

### Nuclei isolation and preparation for single nucleus-RNA sequencing

We isolated single nuclei from FC and dCA2 of Avpr1b-cre x SUN1GFP transgenic mice whose nuclear membranes of Avpr1b neurons are labeled with GFP fluorescence. Each mouse was anesthetized with isoflurane and its brain quickly removed and placed in a petri dish. It was rinsed with sterile PBS (Gibco), then placed on a brain matrix (RBMS-205C, Kent Scientific Co., Torrington, CT) with a 0.5mm coronal slicer. The dCA2 sections generated were placed on a rubber stopper. Using a dissecting microscope, bilateral dCA2 regions (Avpr1b-cre x SUN1GFP male n=8, female n=8; BTBR mice male n=10, female n=10; WT mice male n=10, female n=10) were punched out using a 2 mm diameter Palkovits punch (Palkovits & Jacobowitz, 1974), and snap-frozen by placing the tissue in pre-chilled low bind 2 mL microfuge tubes on dry ice (#022431048 Eppendorf). FC sections (male n = 27, female n=24) were generated and FC regions punched out using 1.5 mm diameter Palkovits punch and placed directly into pre-chilled 2 mL Lobind tubes. 1 mL of lysis buffer [1mL of low sucrose buffer (2.75g sucrose, 250µL of 1M HEPES, 125µL of 1M CaCl2, 75µL of 1M MgAC, 5 µL of 0.5 EDTA, 25 µL of 15.5ug/100µL of DTT, final volume with 25mL with ultrapure DNA/RNAse free water), 10 µL of 10% NP-40, (#L8896, Sigma-Aldrich)] was added directly to each Lobind tube containing tissue and transferred to the pre-chilled 2 mL douncer (#D8938-1SET, Sigma) using 1 mL serological pipettes. Samples were dounced 10 times each with an A pestle followed by a B pestle. Each mL of nuclei was transferred to a pre-chilled 50 mL Falcon tube with 40 µM cell strainer with a pre-wetted filter. Dounce tubes were rinsed with 1.5 mL low sucrose buffer to gather remaining lysed nuclear contents and passed through the filter. Filtered solutions were transferred into 2 mL low bind microfuge tubes and centrifuged for 5 min at speed of 300 g at 4 °C. Supernatants were removed, and 2 mL of low sucrose buffer were added. The pellets were broken by gently pipetting 3-5 times, followed by centrifugation for 5 min at speed of 300 g at 4 °C. Supernatants were removed and 250 µL of nuclei wash and resuspension buffer were added [1xPBS (#10010023, ThermoFisher), 1% BSA (#130-091-376, MACS BSA stock solution), 0.2U/µL RNAse inhibitors (#N8080119, ThermoFisher)]. Replicate tubes were pooled into one tube. In separate 2 mL Lobind tubes, 900 µL of sucrose cushion buffer I [2.7mL Sigma PURE 2M Sucrose Cushion Solution + 300 µL Sigma PURE Sucrose Cushion Buffer (#NUC201-1KT Nuclei PURE Prep, Sigma)] were added and mixed by pipetting 10 times. The mixed nuclei suspension was carefully layered on top of the sucrose cushion without mixing, followed by centrifugation at 13,000 x g for 45 mins at 4°C. All but 100 µL of the supernatant was removed from each tube and the nuclei pellets were resuspended in an additional 150 µL to 500 µL volume of nuclei resuspension buffer containing ∼0.15 - 0.2M sucrose to achieve a nuclei concentration of 1000 nuclei/µL. Nuclei were counted using trypan blue solution 0.4% (#15250061, ThermoFisher) to visualize on a hemocytometer. 10 µL of 1:1 nuclei and trypan blue dye were added to hemocytometer and 5 squares of the hemocytometer were counted and averaged for determining the concentration. 2.5 µL of 5mM DRAQ5 fluorescent probe solution (#62251, ThermoFisher) were added to achieve a 5-10 µM final concentration. Nuclei were passed through 40 µm cell strainers (#08-771-1, Fisher Scientific). Avpr1b x SUN1GFP samples were sorted on Sony SH800 sorter (sorter purity mode, target event rate of 2500 - 5000 events per second with target flow setting of 7) (Supplemental Figure 2). A maximum of 20,000 nuclei per well were loaded onto the 10x Genomics Chromium Controller chip following the manufacturer’s instructions (Chromium Single Cell 3’ Reagent Kits V2, 10x Genomics). Gel bead emulsions (GEM) were generated using microfluidics and nuclei were barcoded with UMI and cell barcode. cDNAs were generated by reverse transcription, followed by post-GEM RT clean up and cDNA amplification. After cDNA amplification, 3’Gen Expression library (GEX) construction was performed by involving steps of fragmentation of cDNA, end pair and A-tailing, post-fragmentation and sample index, and Illumina adaptor ligation. 10x 3’GEX libraries were sequenced on an Illumina Novaseq 6000 at sequencing depth of 100,000 reads/cell. Binary base call (BCL) files were generated and demultiplexed and then FASTQ files were generated. FASTQ files were aligned to Mus musculus. GRCm38.102.chr.gtf.gz with custom references added for the CRE, GFP and Avpr1b transgenes enabled the 10x Genomics Cell Ranger pipeline (V2) to generate filtered gene matrices files for downstream analysis.

### Downstream data analysis

We used the Seurat V3 analysis package for downstream single nucleus RNA sequencing data analysis. This code is available in Supplemental files and GitHub https://github.com/sulee85/Fasciola-Cinereum (SuLee_FCproject_code_example.html, FC_dCA2_all_vector_integration_SUN1GFP_MYC_20231006.html, BTBR_all_vector_Integration_FINAL_20230924.html). Some of the cell type classification methods that were previously described (Hashikawa et al., 2020) were modified and utilized to define cell populations. The groups were integrated using canonical correlation analysis (Butler et al., 2018) and mutual nearest neighbor analysis (Haghverdi et al., 2018) as previously described. We filtered out cells that have unique feature counts over 15,000 and less than 700, and cells with more than 20% of mitochondrial counts. The feature expression measurement for each cell by the total expression was normalized by scale factor of 10,000 and log-transformed. 5,000 highly variable genes were selected for the Avpr1b-cre x SUN1GFP data set and 2,000 highly variable genes were selected for the BTBR dataset. Anchors between individual datasets were identified using FindIntegrationAnchors function and integrated the gene expression matrix. Integrated gene expression matrices were scaled and visualized in principle component analysis (PCA) dimension (Macosko et al., 2015). The numbers of the principal component (PC) value of 1 to 30 were chosen based on elbow plot where the data flattens. Data was visualized by generating Uniform Manifold Approximation and Projections (UMAP) (Becht et al., 2019), and clustering was performed by Louvain clustering (resolution = 0.3 for Avpr1bcre x SUN1GFP dataset and resolution = 0.1 for BTBR dataset). For cluster annotation, feature plot with gene expression was used and cell type annotation was classified in supervised manner. We constructed phylogenic trees to compare distance between the cell types to determine transcriptional similarities as described in https://www.rdocumentation.org/packages/ape/versions/5.2/topics/plot.phylo. For the Avpr1b-cre x SUN1GFP dataset, we calculated the top 50 differentially expressed gene (DEG) markers in 7 different cell types using FindALLMarkers function, corrected with 5% false discovery rate [FDR] and log fold change >0.25, followed by listing the top 50 genes with avg_log2FC. Gene Ontology (GO) was performed on enrich R to query hundreds of thousands of annotated gene sets for both dCA2 and FC regions (see https://maayanlab.cloud/Enrichr) (Chen et al., 2013; Hashikawa et al., 2020; Kuleshov et al., 2016; Xie et al., 2021). GO Biological Process 2023, GO Molecular Function 2023, and GO Cellular Component 2023 were used as a reference.

### Hiplex assay

The neuroanatomical distributions and colocalizations of the expressions of selected genes in the dCA2 and FC regions were analyzed using the ACD Hiplex RNAscope system. Two sets of 12 genes (T1-T12 each set) were created by ACD. Sections (12 µm) from a single WT mouse brain were processed per the manufacturer’s instructions at several levels. Sections were scanned by the NIMH Systems Neuroscience Imaging Resource using the Zeiss AxioScan Z1 widefield slide scanner (software version v3.1) and implementing autofocus, online stitching and shading correction. The AxionScan used a Zeiss plan-apochromat 20X/0.8NA M27 objective and a Hamamatsu Orcu Flash (sCMOS, 2048X2048; pixel size = 6.5 um; qE 82%) camera. The scaled pixel dimension of images was 0.325 µm. All images were collected as 16-bit. The illumination source was from an LED (HXP-120, all wavelengths set to 100%, 3-12 W/mm) with exposure time adjusted automatically to minimize saturation (range 5-300 ms) in areas of brightest signal on the section. The beam splitters were at 395, 570, 660 nm. The excitation filter sets used were 375-395, 455-483, 538-562, 620-643 nm. The emission filters were 410-440, 499-529, 570-640, and 659-759 nm.

### Stereotaxic surgery, viral tracing, and optogenetic surgical implantation

For anterograde viral tracing, Avpr1b-cre mice (n=2) were injected with 150 nL of a Cre-dependent AAV virus expressing YFP [pAAV-CAG-DIO-ChR2(H134R)-eYFP] (#127090-PHPeB, Addgene) into the FC region at 1 x 10^%*^ transducing units per ml (FC coordinates of the bilateral injections at 15 degrees were ML: +/-0.17mm, AP: -2.18mm, DV: -1.83mm) with survival for 4 weeks.

For silencing of Avpr1b neurons in FC for behavioral effects, 7- to 9-week-old male Avpr1b-cre mice and WT mice (n= 24) were injected with 150 nL of the Cre-dependent AAV2-hSyn-DREADD-Gi-mCherry (#44362-AAV2, Addgene) at 5 x 10^%+^ transducing units per ml at the same coordinates.

### Social recognition memory behavioral assay

Behavioral tests were conducted 6 to 8 weeks after the viral injection. All behavioral tests were conducted during the light-cycle between 9:00 AM and 2:00 PM. Mice were injected with either of 3mg/kg of clozapine N-oxide (CNO) or saline control and habituated to the testing room for 30 mins prior to behavioral testing. Mice were placed in new mouse cage for 5 mins prior to behavioral testing. Behaviors were recorded using Ethovision XT (version 13; Noldus Information Technology, Leesburg, VA). During acquisition trials, mice were exposed to a freely moving unfamiliar OVX mouse in a cage for 5 min. After 30-min retention interval in the absence of the stimulus female, retrieval trials were performed in which mice were exposed to the same familiar OVX female for 5 min. A decrease in the sniffing ratio (calculated by Retrieval sniffing duration − Acquisition sniffing duration_:Acquisition sniffing duration x10_0) between acquisition to retrievals of >30% indicates that the test mouse remembers the OVX stimulus mouse.

### Hippocampal slice preparation

Coronal dorsal hippocampal slices were prepared from 8- to 12-week-old Avpr1b-cre (+) Ai9-tdTomato mice (Lee et al., 2023; Williams Avram et al., 2019). Animals were anesthetized with isoflurane and killed by decapitation in accordance with NIH institutional regulations. Brains were quickly removed and dorsal hippocampal slices (350 μm) were cut on a vibratome (Leica VT 1200S) in ice-cold NMDG-HEPES solution containing 93mM NMDG (N-methyl-D-glucamine diatrizoate), 2.5mM KCl, 1.2mM NaH2PO4, 30mM NaHCO3, 20mM HEPES, 25mM D-glucose, 5mM sodium ascorbate, 2mM thiourea, 3mM sodium pyruvate, 10mM MgSO4 and 0.5mM CaCl2 (adjusted to pH 7.3–7.4 with 10N HCl, bubbled with 95% O2/ 5% CO2). Brain slices were then incubated in NMDG-HEPES solution (32°C) for 12 min, and transferred to an immersed-type chamber in the standard ACSF containing 124mM NaCl, 2.5mM KCl, 1.2mM NaH2PO4, 24mM NaHCO3, 5mM HEPES, 12.5 D-glucose, 2 MgSO4 and 2 CaCl2 (adjusted to pH 7.3–7.4, bubbled with 95% O2/ 5% CO2) at room temperature. The slices were stabilized for at least 1.5 h before transfer to the recording chamber. All electrophysiological recordings were performed at 30-32°C.

### Electrophysiological recordings

Whole-cell patch clamp recordings were obtained from the Avpr1b-tdTomato positive neurons in FC or dCA2 by using live imaging system of a Nikon Eclipse FN1 microscope (Supplemental Figure 3). The glass electrode (4∼6 MΩ) was pulled by P-97 (Sutter Instrument) and filled with a patch pipette solution containing 125mM K-gluconate, 0.1mM EGTA, 5mM KCl, 2mM NaCl, 5mM MgATP, 0.4mM Na2GTP, 10mM Na2-Phosphocreatine, 10mM HEPES (adjusted to pH 7.3, 280 mOsm). Resting membrane potential was measured immediately after membrane puncture. Spontaneous and current evoked action potentials were recorded in current-clamp mode. Current injections lasted for 500 ms at 100-pA increments for evoked action potentials. Recording data were obtained using an Axon Multiclamp 700B amplifier and digitized using an Axon Digidata 1550B board. Data were sampled at 10 kHz and analyzed with Clampfit 11.2.

## Results

### Single nucleus RNA sequencing datasets for transcriptomics profiling

We isolated singlet nuclei from the FC and dCA2 using the sucrose gradient method from Avpr1b-cre x SUN1GFP transgenic mice in which the nuclear membranes of Avpr1b neurons are labeled with GFP fluorescence. 3’gene expression libraries were created as described in the methods, and then normalized and sequenced at a sequencing depth of 100,000 reads/cell (Figure 1).

**Figure 1.**
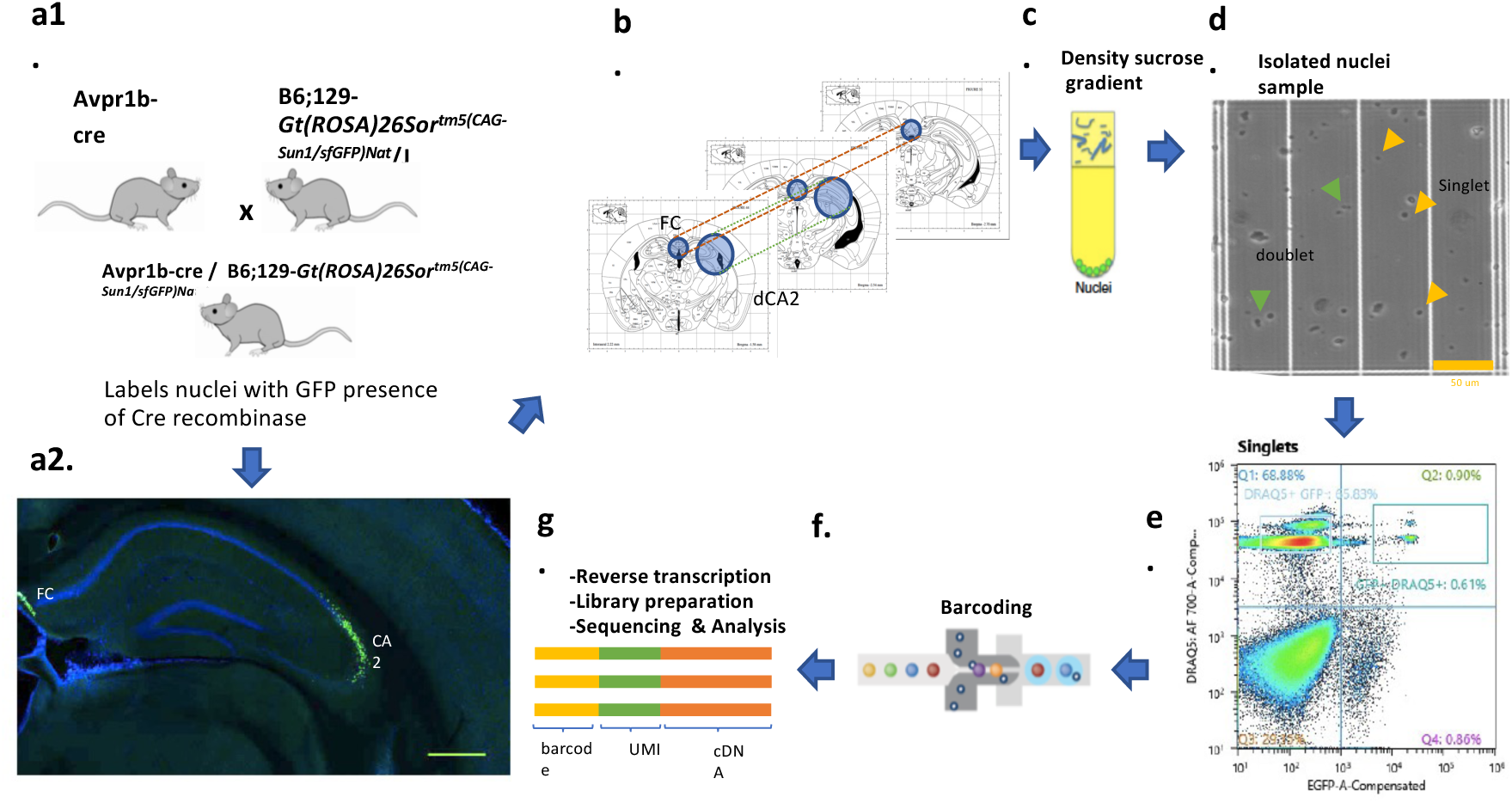
Shows experiment procedure for single nuclei RNA-sequencing. (a1) Transgenic mouse model showing Avpr1b-cre/B6;129-*Gt(ROSA)26Sor^tm5(CAG-Sun1/sfGFP)Nat^*/J to label (a2) Avpr1b nuclei with GFP in the presence of Cre recombinase. (b) FC and dCA2 regions were removed using the Palkovits punch technique. (c, d) Nuclei were isolated using a density sucrose gradient method and singlet/doublets were visualized. (e) The GFP+ DRAQ5+ nuclei populations were selectively isolated using FACS. (f-g) Selected nuclei were barcoded using 10x Genomics Chromium Controller System and sequenced with an Illumina NovaSeq 6000.

Overall, 10 different groups with different conditions were sequenced, integrated and analyzed. These groups included GFP-positive (GFP+) nuclei from the dCA2 of male mice (dCA2_MI_GFPve_pos) with 4541 median UMI reads/cell and 2507 mean genes/cell in total of 2000 nuclei; GFP+ nuclei from the dCA2 of female mice (dCA2_FI_GFPve_pos) with 3974 median UMI reads/cell and 2293 mean genes/cell in total of 1142 nuclei; GFP-negative (GFP–) nuclei from the dCA2 of male mice (dCA2_MI_GFPve_neg) with 2293 median UMI reads/cell and 1611 mean genes/cell in total of 5437 nuclei; GFP+ nuclei from the FC of male mice (FC_MI_GFPve_pos) with 3561 median UMI reads/cell and 2203 mean genes/cell in total of 1247 nuclei; GFP+ from the FC of female mice (FC_FI_GFPve_pos) with 2902 median UMI/cell and 1916 mean genes/cell in total of 1575 nuclei; and GFP– nuclei from the FC of male mice (FC_FI_GFPve_neg) with 2800 median UMI reads/cell and 1989 mean genes/cell in total of 4954 nuclei. For BTBR vs WT mice dCA2 control comparisons, dataset groups included BTBR female mice (BTBR_F) with 3995 median UMI reads/cell and 2252 mean genes/cell in total of 8719 nuclei; BTBR male mice (BTBR_M) with 4005 median UMI reads/cell and 2281 mean genes/cell in total of 9790 nuclei; WT control female mice (WT_F) with 4083 median UMI reads/cell and 2345 mean genes/cell in total of 9764 nuclei; and WT control male mice (WT_M) with 4251 median UMI reads/cell and 2341 mean genes/cell in total of 9625 nuclei.

### Avpr1b-cre x SUN1GFP mice single nucleus RNA sequencing data integration and cluster generation among different cell types based on transcriptional profiling

Nuclei from different groups were integrated by implementing both canonical correlation analysis (CCA) and mutual nearest neighbor analysis (MNN). After integration, we had 3072 median UMI reads/cell and 2023 mean genes/cells in 16102 of total nuclei. The dimensions of the integrated data were reduced by principal-component analysis (PCA) followed by graph-based clustering and visualization using the Uniform Manifold Approximation and Projection (UMAP) algorithm. We identified 15 different clusters and these included neuronal and non-neuronal cell types by their expression of canonical gene markers (Figures 2a, b and f). We identified 10 neuronal cell types and further defined neuronal clusters that express dCA2 genes including Rgs14, Avpr1b, and Amigo2 on featured plots (Figures 2c, d and e), as well as MAPK15 (Supplemental Figure 4). We visually determined, based on the feature plot, that the dCA2 neuronal clusters are clusters 1 and 2 and that they also include most of the GFP+ neurons of the dCA2 and FC.

**Figure 2.**
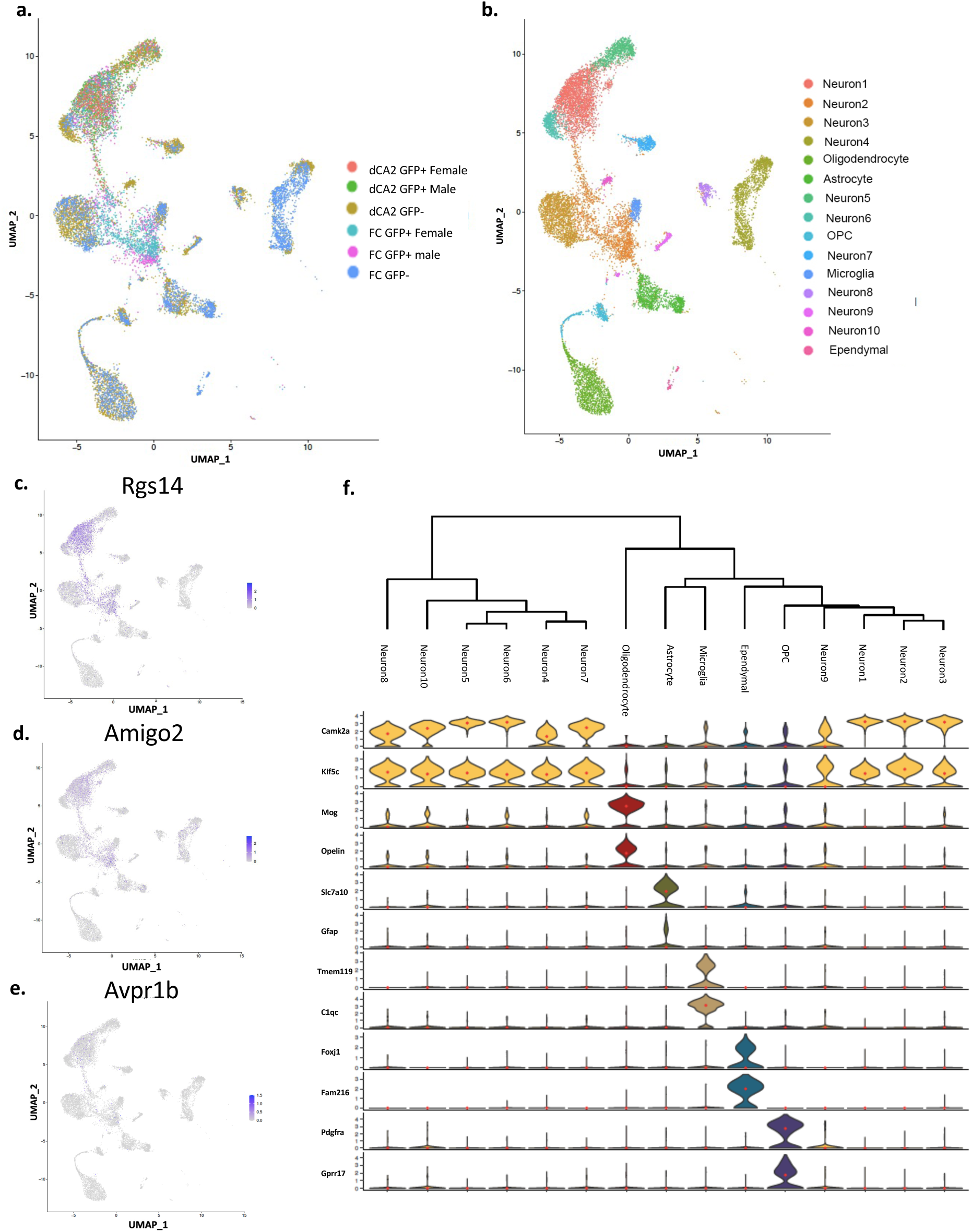
(a) UMAP dimensional reduction and visualization of integrated transcriptional profiles. Groups include dCA2 region of GFP-positive (GFP+) female, dCA2 region of GFP+ male, dCA2 region of GFP-negative (GFP–) control, FC region of GFP+ female, FC region of GFP+ male, FC region of GFP– control. (b) Nuclei classified by different cell type clusters. (c-e) Expression plots in UMAP space illustrating normalized expression values of dCA2 marker Rgs14 (c), Amigo2 (d), Avpr1b (e). (f) Dendrogram showing relationship between cell types and violin plots showing levels of expression of canonical marker genes of different cell types.

### Single nucleus RNA sequencing transcriptomic analysis for subset data of Avpr1b neurons in the FC and dCA2 only

We further analyzed the subsets of data by integrating Avpr1b GFP+ neurons in FC and dCA2 only that included dCA2_MI_GFPve_pos, dCA2_FI_GFPve_pos, FC_MI_GFPve_pos and FC_FI_GFPve_pos groups. After integration, 3795 median UMI reads/cell and 2257 mean genes/cell were identified in a total of 5928 nuclei. Within the integrated data set, 3119 nuclei in the dCA2 and 2809 nuclei in FC were identified (Figure 3a). Differentially expressed genes (DEGs) in the dCA2 and FCs were compared. 1185 genes are differentially expressed between the dCA2 and FCs (corrected with a 5% false discovery rate [FDR] and a log fold change of >0.25, (Supplemental Table 1). Some of the significantly highly expressing genes in the FC are *Mef2c* (average log fold change of 1.674, expressed in 64.2% of the FC neurons and 18.3% of the dCA2 neurons), *Zfp536* (average log fold change of 1.061, expressed in 32.8% of the FC neurons and 1.4% of the dCA2 neurons), *Notch2* (average log fold change of 1.55, expressed in 47.3% of FC neurons and 7.1% of the dCA2 neurons), and *Igfbp4* (average log fold change of 2.134, expressed in 67% of the FC neurons and 9.4% of dCA2 neurons) (Figure 3b-e, Supplemental Figure 5). Some of significantly highly expressing genes in dCA2 are *Cpne7* (average log fold change of 2.79, expressed in 90.1% of the dCA2 neurons and 18.9% of FC neurons), Ttr (average log fold change of 1.48, expressed in 36.7% of the dCA2 neurons and 3.9% of the FC neurons), *Cpne6* (average log fold change of 1.93, expressed in 88.2% of the dCA2 neurons and 30.2% of the FC neurons), and *Scn1a* (average log fold change of 0.845, expressed in 80.1% of the dCA2 neurons and 43.6% of the FC neurons) (Figure 3f-i). Gene Ontology (GO) was performed on “enrich R” for involvement of biological, molecular and cellular process and both dCA2 and FCs are involved in essential roles for the hippocampus including nervous system development and post-synaptic density. Interestingly, one distinct functional involvement of the dCA2 is for amyloid-beta binding (*Lrp1*, *Gria2*, and *Grin2a*) (Figure 4, Supplemental Figures 6-7).

**Figure 3.**
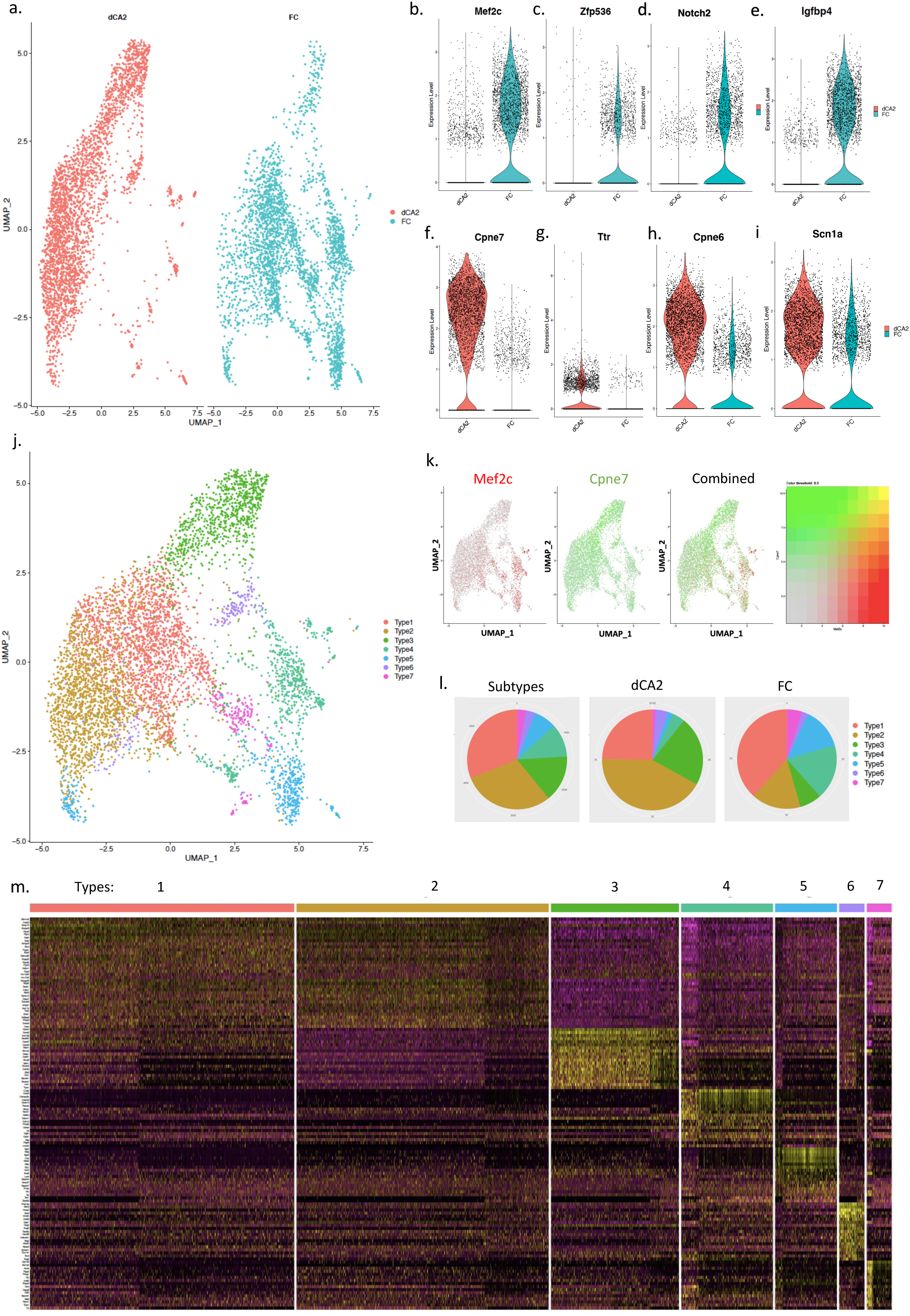
(a) UMAP dimensional reduction separated in group by region of interest comparing between dCA2 and FC groups. (b-i) Violin plot comparing gene expression levels of FC and dCA2 regions. (b) Mef2c, (c) Zfp536, (d) Notch2, (e) Igfbp4, (f) Cpne7, (g) Ttr, (h) Cpne6, and (i) Scn1a. (j) UMAP visualization of Avpr1b GFP+ neuronal clusters. Clusters less than 0.1% of nuclei population were removed. (k) UMAP visualization of gene expression overlap of Mef2c and Cpne7. (l) (left) Proportion of nuclei in each cell subtype cluster both dCA2 and FC combined, (middle) within dCA2 and (right) within FC region. (m) Heatmap showing scaled expression of marker genes in each subtype.

**Figure 4.**
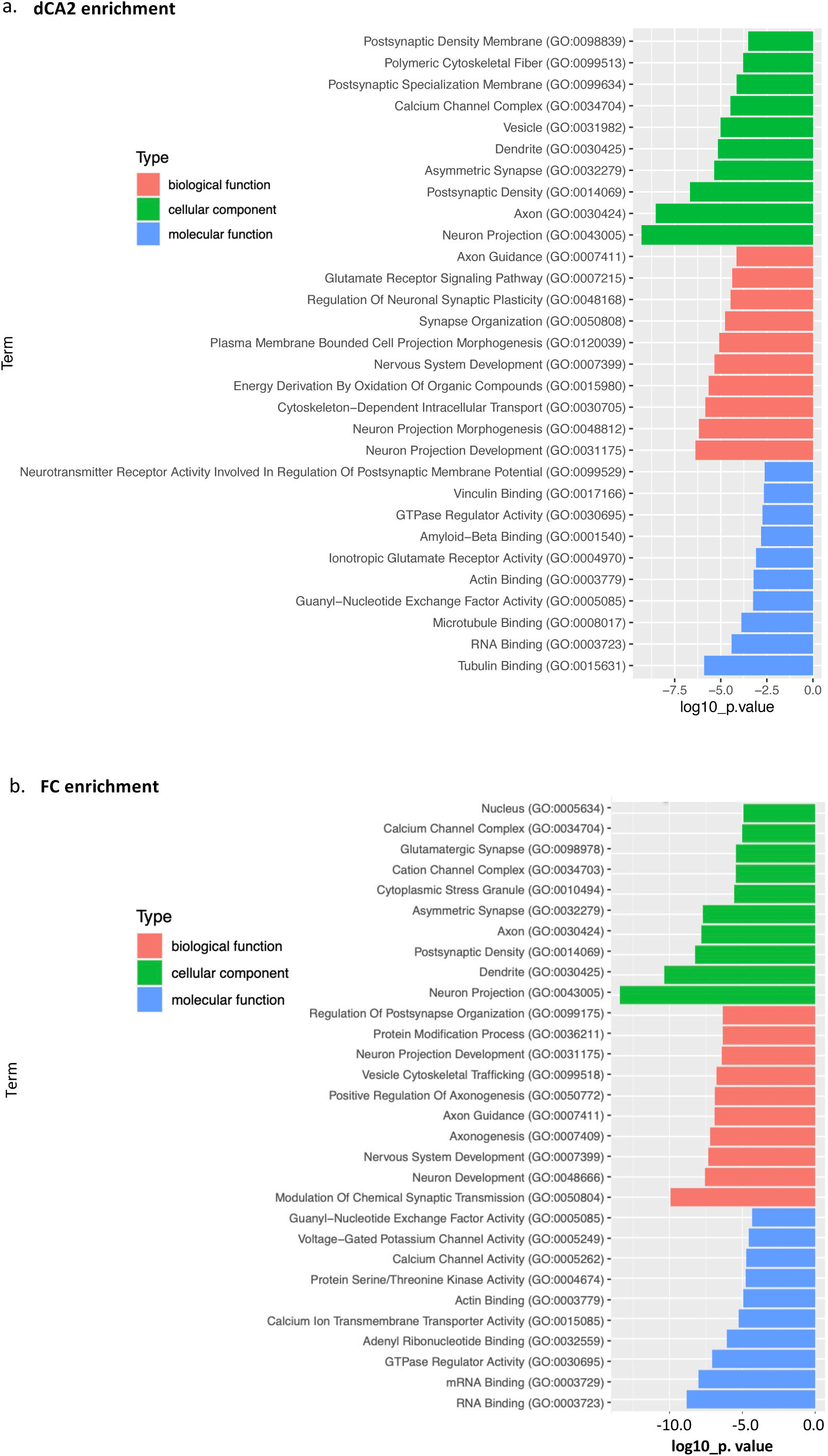
Enriched gene ontology terms and fraction listed for all genes in (a) dCA2 and (b) FC region.

### 7 different cell types are identified in Avpr1b neurons of the FC and dCA2 with subtypes 4, 5 and 7 more prominent in the FC

After the integration of Avpr1b positive neurons in the FC and dCA2, dimensions of integrated data were reduced by PCA followed by graphics-based clustering and visualization on UMAP space. Initially, 9 different subtype clusters were obtained (subtype 1:1830 cells, subtype 2: 1746 cells, subtype 3: 886 cells, subtype 4: 635 cells, subtype 5: 428 cells, subtype 6: 176 cells, subtype 7:173 cells, subtype 8: 35 cells, and subtype 9: 19 cells). We only included cluster cell types that represent more than 1% of cells in that overall cluster. Therefore, only cell subtypes 1-7 were included in the analysis (Figures 3j, k, l and 5b). Overall, dCA2 neurons are represented by 24.6% of subtype 1, 42.4% of subtype 2, 22.3% subtype 3, 4% of type 4, 1.7% of subtype 5, 3.9% of subtype 6, and 1.1% of subtype 7 (figure 3l, middle panel). FC neurons are represented by 38.3% of subtype 1, 16% of subtype 2, 7.1% of subtype 3, 18% of subtype 4, 13.4% of subtype 5, 2% of subtype 6, and 5% of subtype 7 (figure 3l, right panel).

We calculated the top 50 differentially expressed gene (DEG) markers in the 7 different cell subtypes (corrected with a 5% false discovery rate [FDR] and log fold change >0.25) and visualized in the heatmap (Figures 3m and 5a and Supplemental Table 2). Top DEG markers for cell subtype 1 include *Man1a2, Kcnk2,* and *Akap13*; subtype 2 *Stxbp5l, Rapgef5,* and *Zbtb20*; subtype 3 *Opcml, Ccnd2,* and *Rnf182*; subtype 4 *C1ql3, Cntn5,* and *Stxbp6*; subtype 5 *Gjc3, Ermn,* and *Ugt8a*; subtype 6 *Pde10a, Ntrk2,* and *Nr4a2*; and subtype 7 *Ednrb, Sox6,* and *Gjb6*.

**Figure 5.**
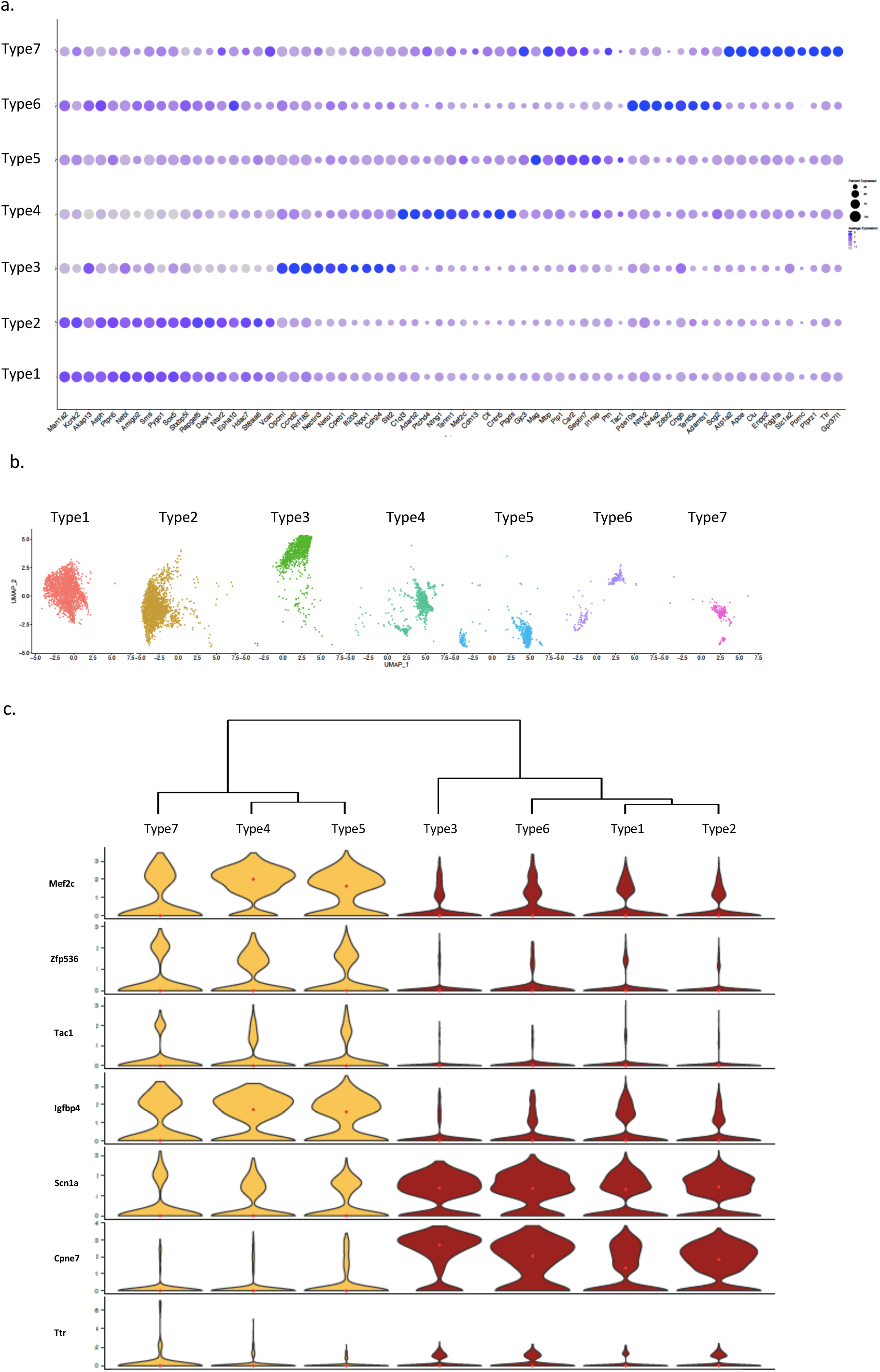
(a) Dot plot showing scaled expression level (color) and the proportion of expressing cells (dot size) of top 10 marker genes for each cell subtype of Avpr1b GFP+ neurons. (b) UMAP dimensional reduction plot separated by cell subtypes (c) Dendrogram showing relationship between each subtype and violin plots show levels of expression of marker genes for each subtypes for both FC vs dCA2 regions.

We constructed phylogenic trees to compare distances between the cell types to determine transcriptional similarities (Figure 5c). This resulted in two main clusters, one cluster including cell subtypes 4, 5 and 7 and another cluster including cell subtypes 1, 2, 3 and 6. Next, top DEG markers identified in FC and dCA2 prior were visualized within each of the cell types by violin plots (Figure 5c, Supplemental Table 3). Overall, based on visualization, DEG markers initially identified in the FC are abundantly expressed in cell subtypes 4, 5 and 7 and those initially identified in the dCA2 are abundantly expressed in cell subtypes 1, 2, 3 and 6.

Mef2c is expressed significantly higher in cell subtype 4 (average log fold change of 1.05; 82.8% in cell subtype 4 vs 45.8% in other cell types) and in cell subtype 5 (average log fold change of 0.522; 65.7% in cell subtype 5 vs 48.5% in other cell subtypes). Zfp536 is expressed higher in cell subtype 4 (average log fold change of 0.538; 46.5% in cell subtype 4 vs 14.7% in other cell types), cell subtype 5 (average log fold change of 0.5; 36.7% in cell subtype 5 vs 16.7% in other cell subtypes), and cell subtype 7 (average log fold change of 0.61; 27.7% in cell subtype 7 vs 17.8% in other cell subtypes). Tac1 is expressed higher in cell subtype 4 (average log fold change of 0.34; 30.9% in cell subtype 4 and 25.4% in other cell subtypes), cell subtype 5 (average log fold change of 0.601; 31.3% in cell subtype 5 and 25.6% in other cell subtypes), and cell subtype 7 (average log fold change of 0.31 higher in lower subpopulation of 16.2% in cell subtype 7 comparison to 26.3% in other cell subtypes). Cpne7 is expressed significantly higher in cell subtype 3 (average log fold change of 0.8; 98.6% in cell subtype 3 and 95.2% in other cell subtypes). Although Ttr is expressed higher in the dCA2 and has higher representation in cell subtypes 2 and 3 than in the FC, Ttr expression is significantly higher in cell subtype 7 (average log fold change of 3.13, 91.3% in cell subtype 7 and 78.9% in other cell subtypes).

### *In situ* hybridization localization of different subtypes in proximal and distal regions of the hippocampus

Although the top 50 differentially expressed gene (DEG) markers in 7 different cell subtypes were ranked by fold change and p-value, the gene expression of many were low and not easily detected by *in situ* hybridization histochemistry. Therefore, in order to minimize artifacts arising from the background expression, we extracted raw RNA counts suset@assays$RNA@counts for DEGs and we ranked the highest to lowest expression per cell type based on RNA counts for raw counts per cell type/total sum of raw RNA counts of all cell types. For example, RNA counts for a gene expressed in cell type1 = RNA counts cell type1/(RNAcounts cell type2 + RNAcounts cell type3 + RNAcounts cell type 4 + RNAcounts cell type 5 + RNAcounts cell type 6 + RNAcounts cell type 7). Furthermore, we visualized the list of genes on the feature plot to confirm that they are uniquely expressed in a specific cell subtype. We then performed HiPlex fluorescence *in situ* hybridization (FISH) multipex assays on mouse brain sections containing anterior to posterior sections of the dorsal hippocampus in order to spatially confirm the DEGs from our single nucleus RNA sequencing data (Figure 6). As a reference, the dCA2 was localized by *Rgs14* and *Amigo2* expression on hippocampal sections (Figure 6). The CA1 was identified by *Wfs1* gene expression and the CA3 by *Bok* gene expression.

**Figure 6.**
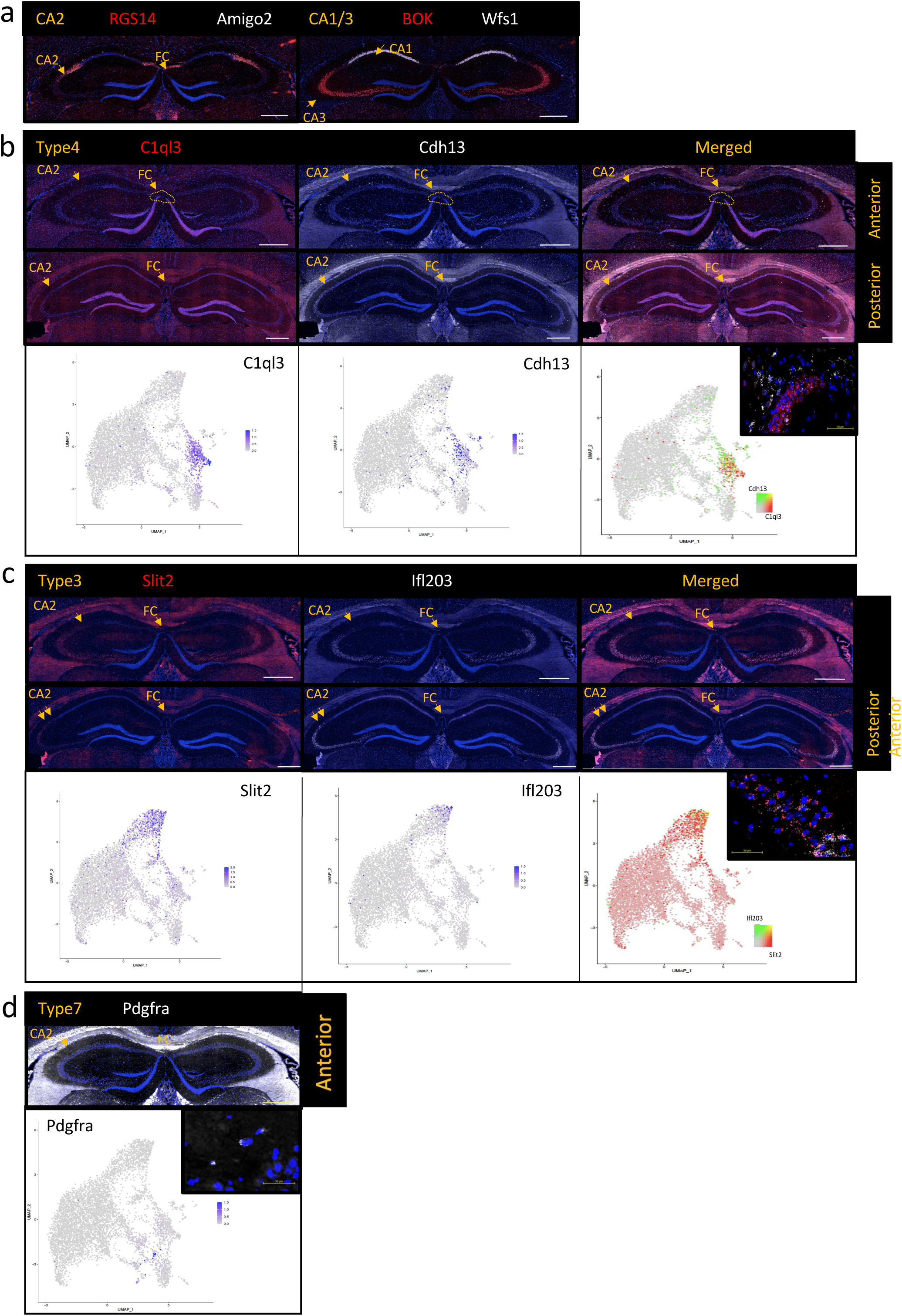
Hippocampal *in situ* hybridization localization of gene expression. (a) CA2 marker genes Rgs14 and Amigo2 in dCA2 and FC (left), CA1 marker gene Wfs1 and CA3 marker gene Bok (right). (b) Expression of type 4 cell subtype genes C1ql3 (left), Cdh13 (middle), and merged (right) in anterior and posterior levels of the dorsal hippocampus. Insert is higher magnification of merged expression from within the FC. These type 4 markers are also shown in UMAP plots (bottom). (c) Expression of type 3 cell subtype genes Slit2 (left), Ifl203 (middle), and merged (right) in anterior and posterior levels of the dorsal hippocampus. Insert is higher magnification of merged expression from within the dCA2. These type 3 markers are also shown in UMAP plots (bottom). (d) Expression of type 7 cell subtype Pgdfa gene. Scale bar represents 500 um. Higher magnification of expression from within the dCA2 region of stratum Orien layer of the non-pyramidal neurons. Scale bar represents 50um. This type 7 marker is also shown in a UMAP plot (bottom).

We found that cell subtype 4 markers, including *C1ql3* and *Cdh13*, are present in distal hippocampus and prominently represented in the FC in continuation with the dentate gyrus. On the other hand, cell type 3 markers, including *Slit2* and *Ifl203*, were present in the proximal part of the hippocampus with continuation into the CA3 region. Cell type 7 *Pdgfra* had very sparse expression in the stratum oriens layer of the dCA2 region in the non-pyramidal neurons.

### Anterior projection of Avpr1b neurons in the FC and inhibition of Avpr1b neurons in the FC

Anterior projections of Avpr1b neurons in the FC were traced by injecting Cre-dependent AAV virus expressing YFP [pAAV-CAG-DIO-ChR2(H134R)-eYFP] into the FC of Avpr1b-cre mice. Anterograde tracing of Avpr1b neurons specific for the FC showed projections to the IG, lateral septum intermediate (iLS), and lacunosum molecular layer of CA1 (Figures 7a-c). Other regions included the dCA2, dorsal fornix and septofibrial nuclei.

**Figure 7.**
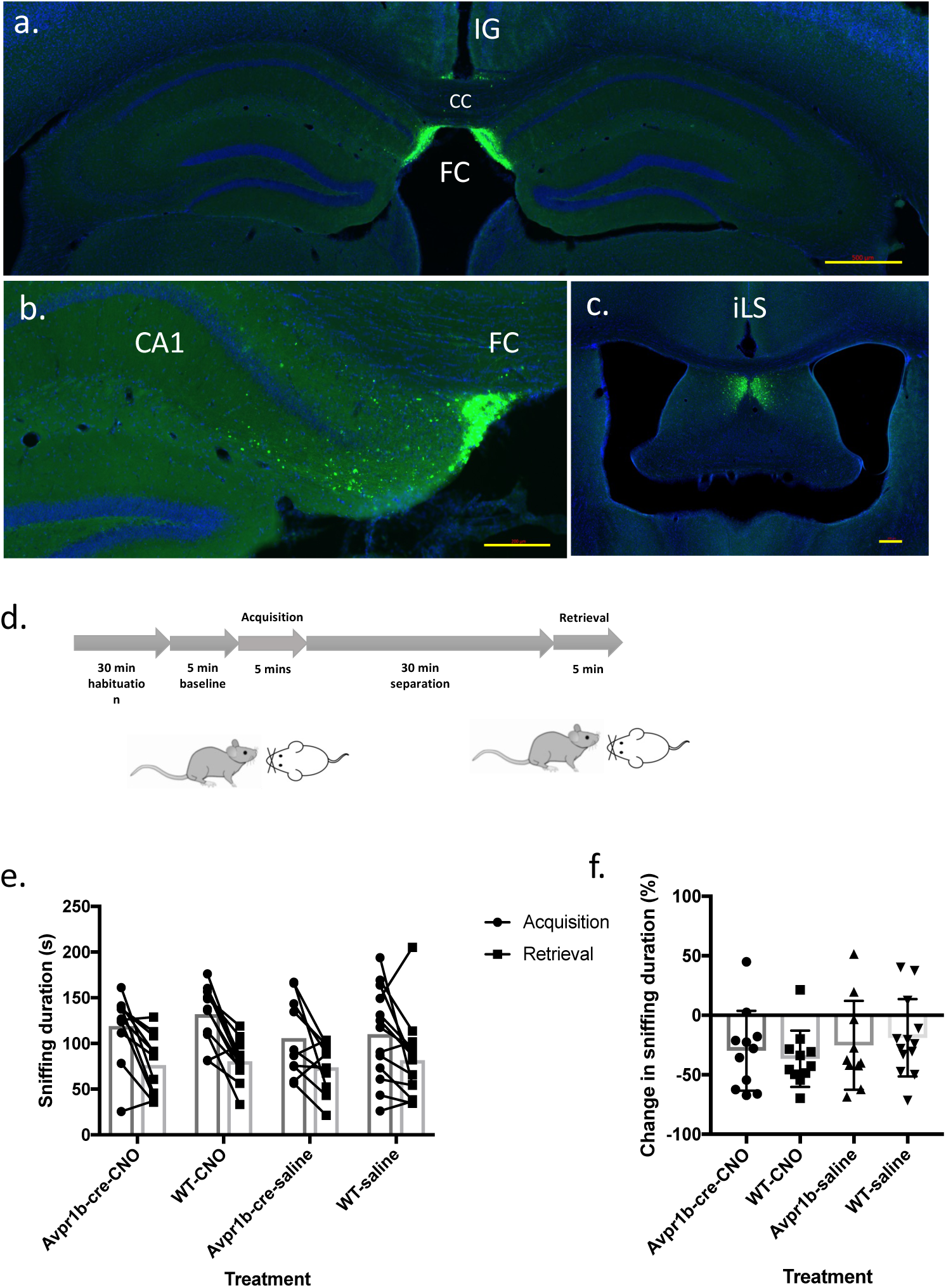
Avpr1b neurons (insert) in the FC, labeled with YFP, project to (a) the induseum griseum (IG; cc=corpus callosum), (b) lacunosum molecular layer of CA1, and (c) intermediate lateral septum (iLS). (d) Behavioral paradigm to study the role of Avpr1b neurons in FC by silencing FC region after injecting AAV2-hSyn-DREADD-Gi-mCherry into Avpr1b-cre and WT mice. 30 mins after CNO injection, the subject interacts with an ovariectomized Balb/C female for 5 minutes and after a 30-minute interval, is exposed to the same mouse (familiar test). Animals without viral expression were excluded from the analysis. Avpr1b-cre-CNO = Avpr1b-cre group that received CNO (N=11); WT-CNO = WT group that received CNO (N=11); Avpr1b-cre-saline = Avpr1b-cre group that received saline (N= 10); WT-saline = WT group that received saline (N= 13). (e) Avpr1b-cre mice and WT mice sniffed significantly less during the retrieval period when compared to the acquisition period (two-way repeated measures of ANOVA: genotype x trial interaction F (3, 21) = 0.9464, *p* = 0.4361; trial F (1, 21) = 40.66, *p* <0.001; genotype F (3, 21) = 0.507, *p* =0.681). Social recognition memory is unaffected. (f) Also, no significant differences were observed between treatment and genotypes in terms of the change in sniffing duration. Change in sniffing duration calculated by (Retrieval sniffing duration − Acquisition sniffing duration)_7Acquisition sniffing durationx_ 100. Avpr1b-cre-CNO= 11, average change in sniffing duration = -29.81; WT-CNO= 11, average change in sniffing duration = -36.58, Avpr1b-saline = 10, average change in sniffing duration = -25.25; WT-saline = 13, average change in sniffing duration = -18.88.

The behavioral role of Avpr1b neurons in FC was examined by silencing them with CNO after injections of the AAV2-hSyn-DREADD-Gi-mCherry virus into Avpr1b-cre (and control WT) mice 6-8 weeks earlier (Figures 7d-f). Thirty minutes after a CNO injection, the interaction of the experimental mouse with an ovariectomized Balb/C female for 5 minutes (acquisition) and again with the same female after a 30-minute interval (retrieval, familiar test) was studied. A decrease in the sniffing ratio (calculated by _[_ (Retrieval sniffing duration − Acquisition sniffing duration)_=Acquisition sniffing duration]_ x100) between acquisition to retrieval trials of >30% indicates that the test mouse remembers the OVX stimulus mouse. Two-way repeated measures of ANOVA (interaction of genotype x trial) and post-hoc Sidak’s multiple comparison tests were performed. None of the groups showed significant impairment in social memory [(Avpr1b-cre group received CNO (N=11); WT group received CNO (N=11); Avpr1b-cre group received saline (N= 10); WT group received saline (N= 13); not significant, p >0.05)] (Figure 7e). Furthermore, there was no significant difference in change in sniffing duration between the groups (p >0.05) (Figure 7f).

### Comparison of transcriptional profiles in subsets of neurons in dCA2 between BTBR mice and WT mice

All of the different groups from single nucleus RNA sequencing described above were integrated and clustering of different cell subtypes were generated. This resulted in 3523 median UMI reads/cell and 2167 mean genes/cells in 52,387 total nuclei. Dimensions of the integrated data were reduced by principal-component analysis followed by graph-based clustering and visualized using the Uniform Manifold Approximation and Projection (UMAP) algorithm. We identified 15 clusters and these included neuronal and non-neuronal cell types by their canonical gene markers (Figure 8, Supplemental Figures 8-11). We further defined dCA2 clusters as expressing *Amigo2*, *Rgs14* and *Map3k15* (Figure 8c-e). CA3 clusters expressed *Bok* and CA1 pyramidal neuronal clusters expressed *Wfs1* (Supplemental Figure 11).

**Figure 8.**
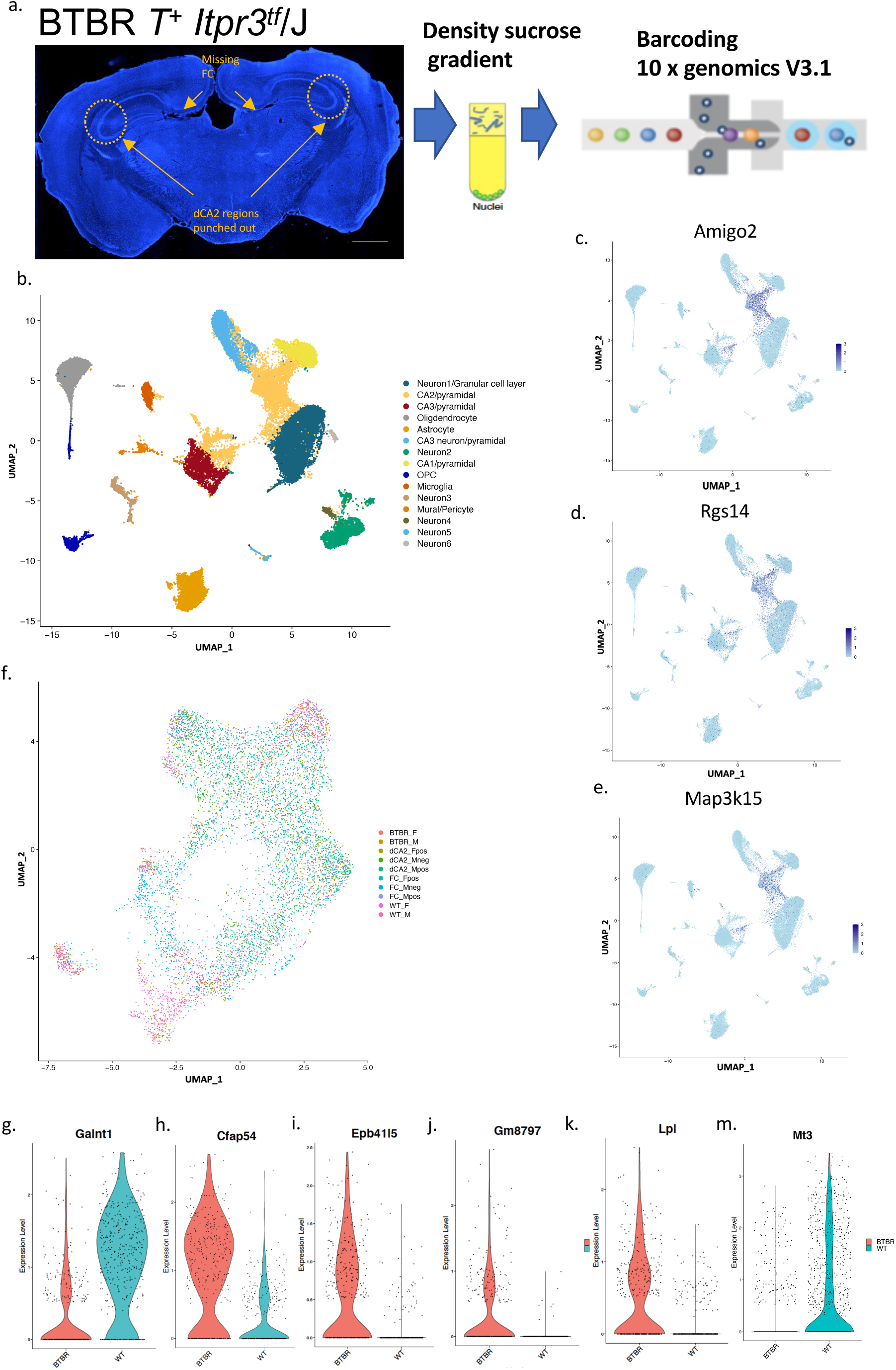
(a) Image shows FC is absent in BTBR *T*^+^ *Itpr3^tf^*/J mouse brain. CA2 samples were isolated using the Pakovits punch and isolated with a sucrose gradient method. Isolated nuclei were barcoded using the 10x Genomics Chromium single cell RNAsequencing system and sequenced with NovaSeq 6000. (b) UMAP dimensional reduction and visualization of integrated transcriptional profiles. UMAP plot shows different cell types in hippocampus based on a transcriptional profile. (c-e) Expression plots in UMAP space illustrating normalized expression values of CA2 marker genes (c) Amigo2 (d) Rgs14 (e) Map3k15 (f) UMAP dimensional reduction of selected subset of CA2 neurons based on CA2 gene markers. (g-m) Violin plot comparing top 10 marker gene expression levels of BTBR in comparison to WT mice in dCA2 subset of neurons. (g) Galnt1, (h) Cfap54, (i) Epb41l5, (j) Gm8797, (k) Lpl, and (m) Mt3.

We further defined a subset of dCA2 clusters (Figure 8f), and compared the DEGs between BTBR and WT groups only. Between these two groups, 2979 median UMI/cell and 2261 mean genes/cell were identified in 733 nuclei in the BTBR subset and 1044 nuclei in the WT subset (Supplemental Table 4). Some of the genes that are highly expressed in the dCA2 of BTBR mice compared to WT are *Cfap54* (average log fold change of 2.82, 46% of neurons in BTBR group and 14.6% of neurons in WT group), *Epb41l5* (average log fold change of 3.09, 34.7% of neurons in BTBR group and 5.6% neurons in WT group), *Gm8797* (average log fold change of 4.67, 31.2% of neurons in BTBR group and 1.3% WT group), and *Lpl* (average log fold change of 2.21, 34.9% of neurons BTBR group and 9.6% WT group). On the other hand, *Galnt1* (average log fold change of 1.6, 23% of neurons BTBR group and 43.5% WT group) and *Mt3* (average log fold change of 2.13, 16% of neurons BTBR group and 42.9% WT group) were expressed lower in the dCA2 of BTBR mice compared to WT mice.

## Discussion

The arginine vasopressin 1b receptor (Avpr1b) plays an important role in social behaviors including social learning, memory, and aggression (Wersinger et al., 2002), and is known to be a specific marker for the CA2 (Young et al., 2006). The fasciola cinereum (FC) is an anatomical region in which Avpr1b expressing neurons are prominent, but the functional roles of the FC have yet to be investigated. Surprisingly, the FC is absent in the BTBR T+tf/J (BTBR) mouse strain used to study core behavioral deficits of autism (Young and Song, unpublished). The FC appears to be a remnant of the original dorsally located hippocampal anlage: a continuation of the dorsal taenia tecta and IG and an anterior and dorsal hippocampal rudiment.

### FC anterogradely projects to the intermediate lateral septum while dCA2 projects to dorsal lateral septum

Anterograde tracing of Avpr1b neurons specific for the FC showed projections to the IG, dCA2, lacunosum molecular layer of CA1, dorsal fornix, septofibrial nuclei, and intermediate lateral septum (iLS) (Figure 7). Recent studies showed that a projection to the ventral CA1 from dCA2 is implicated in encoding, consolidation and recall phases of social memory (Meira et al., 2018); while a projection from the dCA2 to the lateral septum plays a role in disinhibiting social aggression (Leroy et al., 2018). This latter study also described, using monosynaptic retrograde tracing, projections from both the FC and dCA2 to the dorsal lateral septum. In agreement, our results shows that the FC projects to lateral septum, although mainly to the intermediate lateral septum, and then continues posteriorly to the septofibrial nuclei and dorsal fornix region.

### Single nucleus RNA sequencing of FC and dCA2

In this study, we used mice with specific fluorescent labeling of the nuclear membranes of Avpr1b neurons in the dCA2, vCA2, FC and indusium griseum regions (Supplemental Figure 1). We performed single nucleus RNA sequencing on RNA extracted from the labeled nuclei of these Avpr1b neurons from the FC and dCA2 of both female and male mice. This allowed us to understand how FC neurons compare to dCA2 neurons and learn the differences in the transcriptomic profiles that might provide insight into their roles in social memory formation and social behavior. We examined both GFP+ and GFP– groups and identified 15 different clusters. CA2 marker genes, including *Rgs14*, *Amigo2* and *Avpr1b,* and *MAPK15* are concentrated in neuronal clusters 1 and 2 where, as expected, the majority of GFP+ cells are located (Supplemental Figure 4). Other cell type clusters in GFP– groups included oligodendrocytes (and their precursors), astrocytes, and ependymal cells.

Next, we investigated subsets of Avpr1b-positive neurons from only FC or dCA2 to compare DEGs between those two regions. Some of the highly expressing genes in the FC are *Mef2c*, *Zfp536*, *Notch2*, and *Igfbp4* (Figure 3b-e). The *Mef2c* gene is known to be linked to autism in the Mef2c haploinsufficiency syndrome (MCHS) and *Mef2c* heterozygous mice display autism-related behaviors (Harrington et al., 2020). *Zfp536* is expressed abundantly in the developing central nervous system and is known to negatively regulate neuronal differentiation by repressing retinoic acid-induced gene transcription (Qin et al., 2009). *Notch2* signaling is shown to promote cortical neurogenesis (Suzuki et al., 2018).

Some of the highly expressing genes in the dCA2 are *Ttr*, *Cpne6, Cpne,7* and *Scn1a (*Figure 3f-l). Transthyretin (*Ttr*) binds the amyloid-beta (Aβ) fibrils that accumulate in the plaques of Alzheimer’s disease (Li et al., 2011). Gene ontology indicates both dCA2 and FC are involved in essential roles in the hippocampus including nervous system development and post synaptic density. Of course, the involvement of dCA2 in memory is indicated by the presence of *Lrp1* (also known as apolipoprotein E receptor), and the glutamate receptors, *Gria2* and *Grin2a*.

Interestingly, the *Scn1a* gene is significantly more highly expressed in the dCA2 compared to the FC. *Scn1a* encodes the brain voltage-gated sodium channel Nav1.1. It was previously shown that deletion of *Scn1a* causes selective reduction of excitability of inhibitory neurons in hippocampus, evoked seizures, and spatial learning deficits but no abnormalities in social interaction (Silvennoinen et al., 2022; Stein et al., 2019). In conjunction with the transcriptomic data, slice electrophysiology recordings suggested that a higher threshold of stimuli was required to induce action potentials in Avpr1b neurons in FC compared to dCA2 (Supplemental Figure 3). However, in this study, we only examined 15 FC neurons and 6 dCA2 neurons. Therefore, further investigations are necessary to assess this finding.

### 7 subtypes of neurons are identified in Avpr1b neurons of FC and dCA2 and subtypes 4, 5 and 7 are more prominent in FC

After the integration of Avpr1b neurons of FC and dCA2 only, we obtained 7 different cell type clusters that represent more than 1% of cells in the overall cluster. We calculated the top 50 differentially expressed gene markers in those 7 different clusters. Top DEG markers for cell type 1 included *Man1a2, Kcnk2, Akap13*; cell type 2 included *Stxbp5l, Rapgef5, Zbtb20*; cell type 3 included *Opcml, Ccnd2, Rnf182*; cell type 4 included *C1ql3, Cntn5, Stxbp6*; cell type 5 included *Gjc3, Ermn, Ugt8a*; cell type 6 included *Pde10a, Ntrk2, Nr4a2*; and cell type 7 included *Ednrb, Sox6, Gjb6*.

Next phylogenic trees were constructed to compare distances between the cell types to determine transcriptional similarities. This resulted in two main clusters, the first cluster including cell types 4, 5 and 7 found prominently in the FC and second cluster including cell types 1, 2, 3 and 6 found more prominently in the dCA2. Expression of DEG markers *Mef2c*, *Zfp536* and *Tac1* are expressed significantly higher in FC subtypes 4, 5 and 7 in comparison to the other cell types (Figure 5c). Therefore, although there are similarities in the transcriptomic profiles of FC and dCA2, there were subtypes that were represented more prominently by FC.

### *In situ* hybridization labels cell subtype 3 in the proximal hippocampus and subtype 4 in the distal hippocampus and FC

From the single cell RNA sequencing data, we found that cell types 3, 4 and 7 can be labelled by unique gene markers. Using Hiplex *in situ* hybridization histochemistry to determine heterogeneity in Avpr1b neurons, we have found that cell subtype 4 gene markers *C1ql3* and *Cdh13* (has lower expression) are present in the anterior section of distal hippocampus, prominently in the FC and in continuation ventrally in the dentate gyrus. On the other hand, cell type 3 markers *Slit2* and *Lfi203* are present in the both anterior and posterior sections of the proximal part of the hippocampus in CA3 neurons. Cell type 7 marker *Pdgfra* has very sparse expression within non-pyramidal neurons.

### Inhibiting Avpr1b neurons in FC does not impair social recognition memory

The dCA2 is essential for social behaviors, especially social aggression and social memory. Inhibiting dCA2 pyramidal neurons causes reduces the ability for mice to remember a conspecific (Hitti & Siegelbaum, 2014; Stevenson & Caldwell, 2014), while enhancing vasopressin synaptic release onto dCA2 neurons enhances social memory (Smith et al 2016). Therefore, we were interested to learn if the FC plays a role in social memory as well. We found that unlike dCA2, silencing Avpr1b neurons in the FC using the AAV2-hSyn-DREADD-Gi-mCherry virus and CNO does not affect short-term social recognition memory. Unlike the dCA2, the preliminary electrophysiological data suggesting a potentially less excitable nature of FC neurons and significantly lower Scn1a gene expression in FC region may indicate a more regulatory of central nervous system development and maintenance than a mnemonic role. On the other hand, a recent study showed that the FC is important for the acquisition of visual contextual memory in rats (Park et al., 2022), a memory that we did not examine. The abundant projections of FC neurons to intermediate lateral septum, dorsal fornix, and septofibrial nuclei invites further study to investigate potential behavioral roles.

### FC is absent in BTBR mice prompting comparisons of dCA2 transcriptional profiles between BTBR and WT mice

The FC is absent in the BTBR T+tf/J (BTBR) mouse strain (Young and Song, unpublished) used to study core behavioral deficits of autism. At the genomic level, advances in long-read whole genome sequencing have shown that BTBR mice have an 8bp deletion at the 3’ end of exon 2 of Draxin that encodes the chemo-repulsive axon guidance molecule. This mutation contributes to the absence of its corpus collosum (Arslan et al., 2023). Also, it was recently discovered that a disturbed epigenetic silencing mechanism leads to a hyperactive endogenous retrovirus that increases *de novo* copy number variations in BTBR mice (Lin et al., 2023). In order to determine transcriptomic changes in the dCA2 of BTBR mice, we compared DEGs between dCA2s of BTBR and WT mice. As there is no transgenic mouse model that would fluorescently labels Avpr1b neurons within BTBR mouse, we used a sub-setting approach to select dCA2 genes using known dCA2 gene markers after sequencing was performed. We found that transcriptomic profiles of dCA2 neurons between BTBR and WT mice are very similar as they do not form any unique clusters. Some of top DEGs include *Cfap54, Epb41I5, Gm8797,* and *Lpl* with higher expression in the dCA2 of BTBR mice; and *Galnt1* and *Mt3* with higher expression in the dCA2 of WT mice compared to BTBR mice. The roles of these genes in the dCA2 should be further explored.

## Summary

Our data provide comprehensive transcriptomic profiles of subpopulations of neurons in the FC and dCA2 as well as some preliminary proximal-distal spatial distributions of cell types. Both transcriptomic profiles of dCA2 and FC are involved in essential roles for hippocampus including nervous system development and post-synaptic density. We show that *Mef2c,* involved in autism spectrum disorder, is more highly expressed in the FC. *Ttr* is more highly expressed in the dCA2, binds amyloid-beta, and may have an important in role of social memory. The FC shows lower gene expression of the sodium voltage-gated channel 1 alpha subunit (*Scn1a*) compared to the dCA2. Although only a suggestion at this point, slice electrophysiology recordings demonstrated that a higher threshold of stimuli was required to induce action potentials in Avpr1b neurons in FC compared to dCA2. This should be further investigated in more detail and with regard to cell subtype. Recent advances in whole mouse brain spatial transcriptomics using MERFISH had established a benchmark reference atlas for spatial cell types across brain (Yao et al., 2023). From MERFISH whole brain reference, we observe that both FC and dCA2 regions express glutamatergic neurons abundantly. Interestingly, it shows that vascular leptomeningeal cells (VLMC) and choroid plexus (CHOR) co-localized clusters (5296-5299) are specifically located in pia close to ventricles where FC is located (Yao et al., 2023). Therefore, it would be interesting to merge our transcriptomic data and integrate with this spatial transcriptomic reference to see multimodally where these cells are located in FC and dCA2 to further investigate the functional role of FC region.

Anterograde tracing shows that both the FC and dCA2 project to the lateral septum, with dCA2 previously shown to project to the dorsal lateral septum and FC projecting to the intermediate lateral septum. Additionally, in contrast to the dCA2, inhibition of Avpr1b neurons in the FC by the inhibitory DREADD system during behavior testing does not impair social memory.

Surprisingly, we found that the FC is absent in the BTBR T+tf/J (BTBR) mouse, a strain used to study core behavioral deficits of autism and, yet, the dCA2 is still intact. From a single nuclei RNA sequencing study, we found that transcriptomic profiles of dCA2 neurons between BTBR and WT mice are very similar as they did not form any unique clusters, yet we found there were differentially expressed genes between the dCA2s of BTBR and WT mice. Recent advances in long-read sequencing allows one to determine copy number variants at whole genome level without a 3’ or 5’ sequencing bias. Previous studies have observed an increase of copy number variation in BTBR mice. Furthermore, this approach allows one to determine isoforms of transcripts at the single cell level. It would be interesting to determine if there are any isoform variants at the transcriptomic level in the dCA2 region between BTBR and WT mice using long read sequencing. Overall, this is a comprehensive study of the comparison of Avpr1b neuronal subpopulations between the FC and dCA2. The fact that FC is absent in BTBR mice, a mouse model for autism spectrum disorder, suggests that the FC may play a role in understanding neuropsychiatric disease.

## Supporting information

BTBR vs WT Integration

7_Characterization of fasciola cinereum, all group Integration

8_Characterization of Fasciola cinerum, Avpr1b neurons only

NIH Publishing Agreement

## Acknowledgement

A special thank you to everyone in the Section on Neural Gene Expression for their contributions and hard work. We would like to acknowledge the contributions of Jonathan Kuo and Dr. Ted Usdin and the rest of the staff in the Systems Neuroscience Imaging Resource within the National Institute of Mental Health Intramural Program. Thank you also to Dr. Timothy Petros and Dr. Christopher Rhodes (NICHD) of the Unit on Cellular and Molecular Neuro Development for advice on nuclei isolation techniques. Thank you to Drs. Kory Johnson and Fahad Almsned at NINDS Bioinformatics core facility for advice on single nuclei RNAsequencing coding. Dr. Lee Eiden (NIMH) provided administrative support after the retirement of WSY.

## Data availability statement

All datasets generated for this study are included in the article and Supplemental Material.

## Ethics statement

The animal study was reviewed and approved by National Institute of Mental Health Animal Care and Use Committee.

## Supplemental Figure Legends

Supplemental Figure 1. (a-h) Fluorescent labelling of Avpr1b neuronal nuclei in dCA2, FC, and vCA2 with GFP from Avpr1b-Cre; *R26-CAG-LSL-sfGFP-myc* mice. (a) Avpr1b neurons in anterior dCA2 region. (b) Avpr1b neurons in FC regions in anterior hippocampus. (c) Avpr1b neurons in ventral CA2. (d-f) Avpr1b neurons in posterior FC and dCA2 regions. (g-h) Avpr1b neurons in FC region in the posterior hippocampus.

Supplemental Figure 2. Sorting gating for GFP positive and DRAQ5+ nuclei in both (a) dCA2 and (b) FC regions.

Supplemental Figure 3. FC neurons have the lower excitability and unique characteristics compared with dCA2 neurons. (A) Alive images show the brain slice containing dorsal hippocampus on DIV and tdTomato channels from an Avpr1b-cre-Ai9tdTomato mouse (4-fold objective, left side). Alive images show the Avpr1b positive neurons in FC region (60-fold objective, right side). A star indicates the recording electrode was patching on a Avpr1b positive neuron in FC region. (B) Spontaneous action potentials were recorded in FC and dCA2. (C) Summary of the different types of evoked potentials recording on Avpr1b positive neurons in FC and dCA2. Type III red trace show the subthreshold potential which is unique in FC neurons. (D) Statistical analysis of resting membrane potentials in FC and dCA2 neurons. The solid color square box is the mean. *p*= 0.134.

Supplemental Figure 4. Expression plots in UMAP space illustrating normalized expression values of dCA2 marker (a) MAP3k15, (b) GFP and (c) Cre in continuation of main figure 2.

Supplemental Figure 5. UMAP dimensional reduction and visualization of neurons comparing sex differences within (a) dCA2 (b) FC region. (c-f) Proportion of nuclei in each subtype cluster, (c) dCA2 GFP+ female, (d) dCA2 GFP+ male, (e) FC GFP+ female, (f) FC GFP+ male groups.

Supplemental Figures 6. Gene Ontology performed using enrich R specific for different cell types. (a) type 1, (b) type 2, (c) type 3, (d) type 4.

Supplemental Figures 7. Gene Ontology performed using enrich R specific for different cell types. (a) type 5, (b) type 6, (c) type 7.

Supplemental Figures 8. (a) UMAP dimensional reduction and visualization of integrated transcriptional profiles grouped by different groups which includes BTBR_F, BTBR_M, dCA2_ Fpos, dCA2_Mneg, dCA2_Mpos, FC_Fpos, FC_Mneg, FC_Mpos, WT_F and WT_M. (b) UMAP dimension reduction and visualization showing only subset of dCA2 cluster. (c-e) Expression plots in UMAP space illustrating normalized expression values (c) Gfp, (d) Cre, (e) Avpr1b.

Supplemental Figure 9. Expression plots in UMAP space illustrating normalized expression values genes characterized for neuronal types in integrated transcriptional profiles which include BTBR, WT control, and all other groups. (a) Tmem119, (b) C1qc, (c) Mog, (d) Opalin, (e) Pdgfra, and (f) Gpr17.

Supplemental Figure 10. Continuation of expression plots in UMAP space illustrating normalized expression values genes characterized for neuronal types in integrated transcriptional profiles which include BTBR, WT control, and all other groups. (a) Cldn5, (b) Flt1, (c) Slc7a10, (d) Gfap, (e) Stmn2, and (f) Thy1.

Supplemental Figure 11. Continuation of expression plots in UMAP space illustrating normalized expression values genes characterized for neuronal types in integrated transcriptional profiles which include BTBR, WT control, and all other groups. (a) Camk2a, (b) Kif5c, (c) Wfs1, and (d) Bok.

