## Supplemental figures for "Investigation of the Fasciola Cinereum, Absent in BTBR mice, and Comparison with the Hippocampal Area CA2"

Supplemental Figure 1

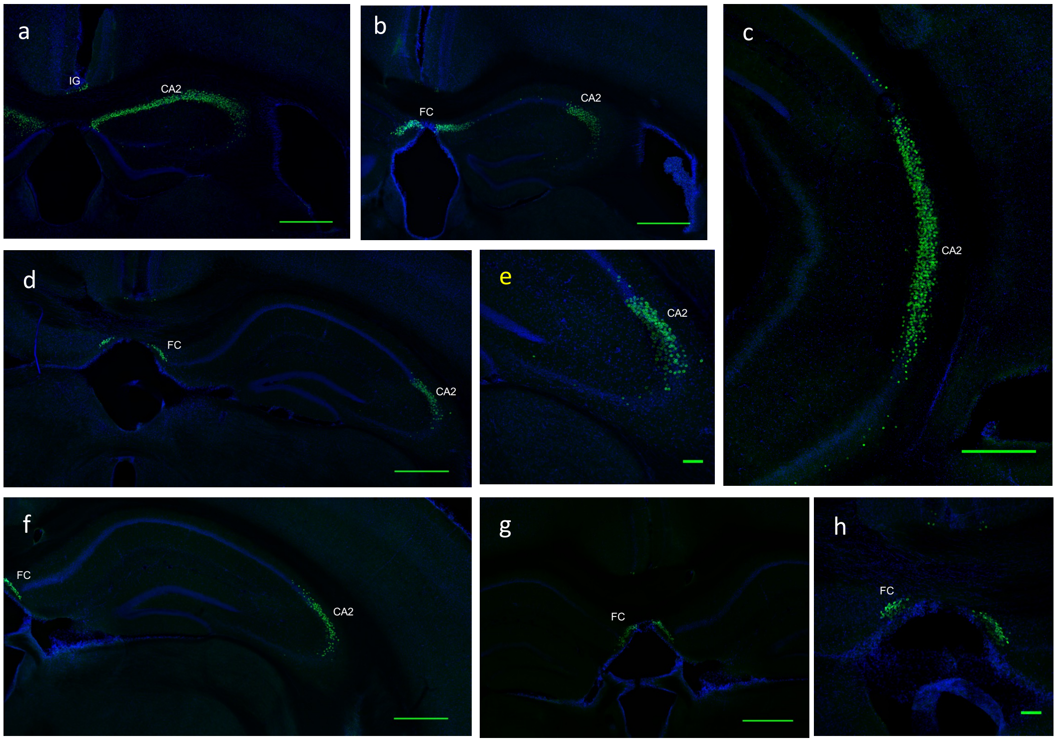

Supplemental Figure 2

### dCA2 sort gating

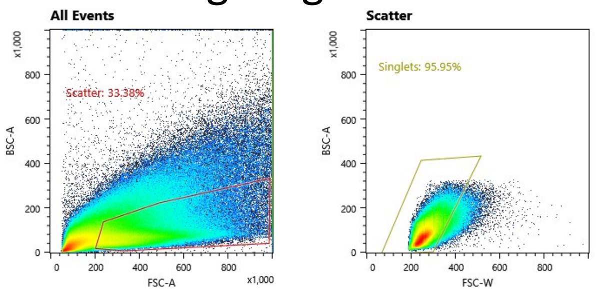


**Gates and Statistics**

| Name | Events | %Parent | %Total |
|---|---|---|---|
| All Events | 1,000,000 | 0.00% | 100.00% |
| Scatter | 333,795 | 33.38% | 33.38% |
| Singlets | 320,283 | 95.95% | 32.03% |
| Q1 | 220,620 | 68.88% | 22.06% |
| Q2 | 2,886 | 0.90% | 0.29% |
| Q3 | 94,019 | 29.35% | 9.40% |
| Q4 | 2,758 | 0.86% | 0.28% |
| GFP+ DRAQ5+ | 1,965 | 0.61% | 0.20% |
| DRAQ5+ GFP- | 210,828 | 65.83% | 21.08% |

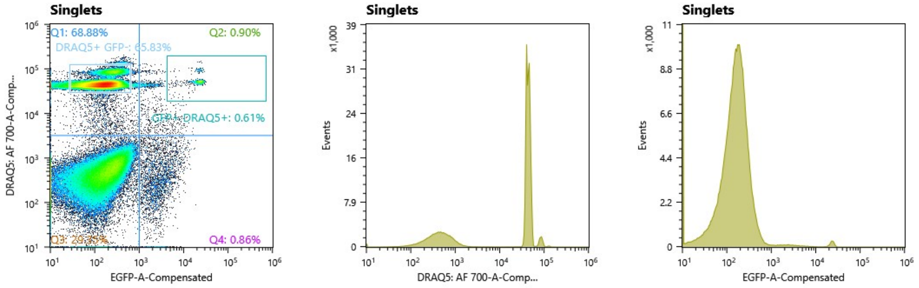


| Collection Tube | Sort Gate | Sort Mode | Elapsed Time | Total Event | Target Ratio | Sorted Count | Sort Rate | Sort Efficiency | Aborted Count | Abort Rate |
|---|---|---|---|---|---|---|---|---|---|---|
| Left | DRAQ5+ GFP- | Ultra Purity | 00:56:54 | 5,574,607 | 19.77% | 877,382 | 256.97eps | 79.62% | 224,621 | 65.79eps |
| Right | GFP+ DRAQ5+ | Ultra Purity | 00:56:54 | 5,574,607 | 0.20% | 9,252 | 2.71eps | 84.76% | 1,663 | 0.49eps |

### FC sort gating

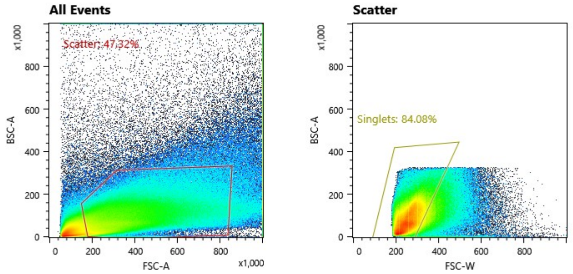


**Gates and Statistics**

| Name | Events | %Parent | %Total |
|---|---|---|---|
| All Events | 1,000,000 | 0.00% | 100.00% |
| Scatter | 473,169 | 47.32% | 47.32% |
| Singlets | 397,860 | 84.08% | 39.79% |
| Q1 | 125,100 | 31.44% | 12.51% |
| Q2 | 1,881 | 0.47% | 0.19% |
| Q3 | 268,747 | 67.55% | 26.87% |
| Q4 | 2,132 | 0.54% | 0.21% |
| GFP+ DRAQ5+ | 1,622 | 0.41% | 0.16% |
| DRAQ5+ GFP- | 123,219 | 30.97% | 12.32% |

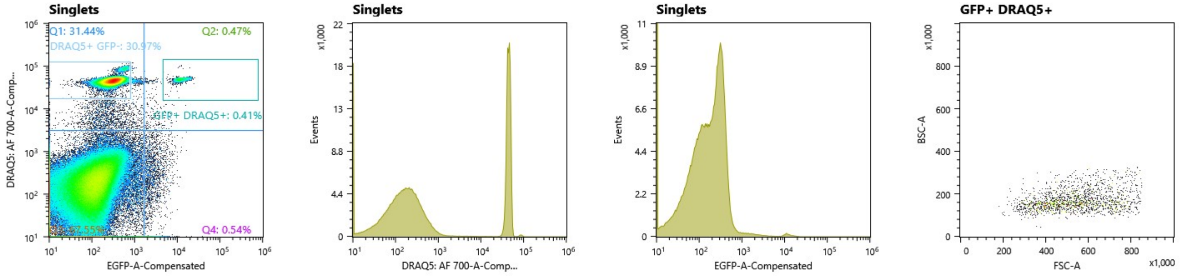


| Collection Tube | Sort Gate | Sort Mode | Elapsed Time | Total Event | Target Ratio | Sorted Count | Sort Rate | Sort Efficiency | Aborted Count | Abort Rate |
|---|---|---|---|---|---|---|---|---|---|---|
| Left | DRAQ5+ GFP- | Ultra Purity | 00:48:20 | 2,632,377 | 11.82% | 279,858 | 96.48eps | 89.92% | 31,364 | 10.81eps |
| Right | GFP+ DRAQ5+ | Ultra Purity | 00:48:20 | 2,632,377 | 0.16% | 3,921 | 1.35eps | 91.61% | 359 | 0.12eps |

Supplemental Figure 3

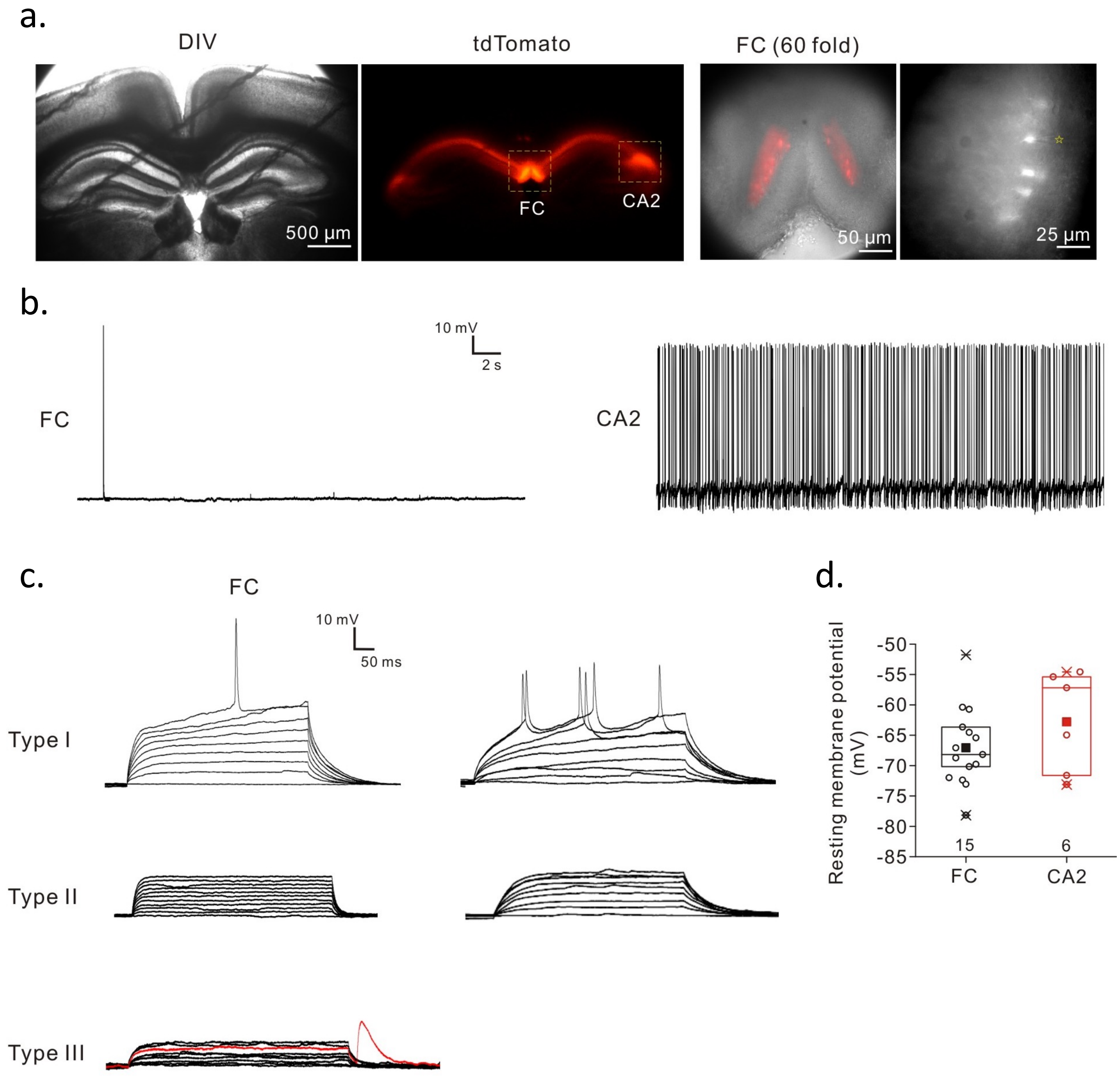

### Supplemental Figure 4

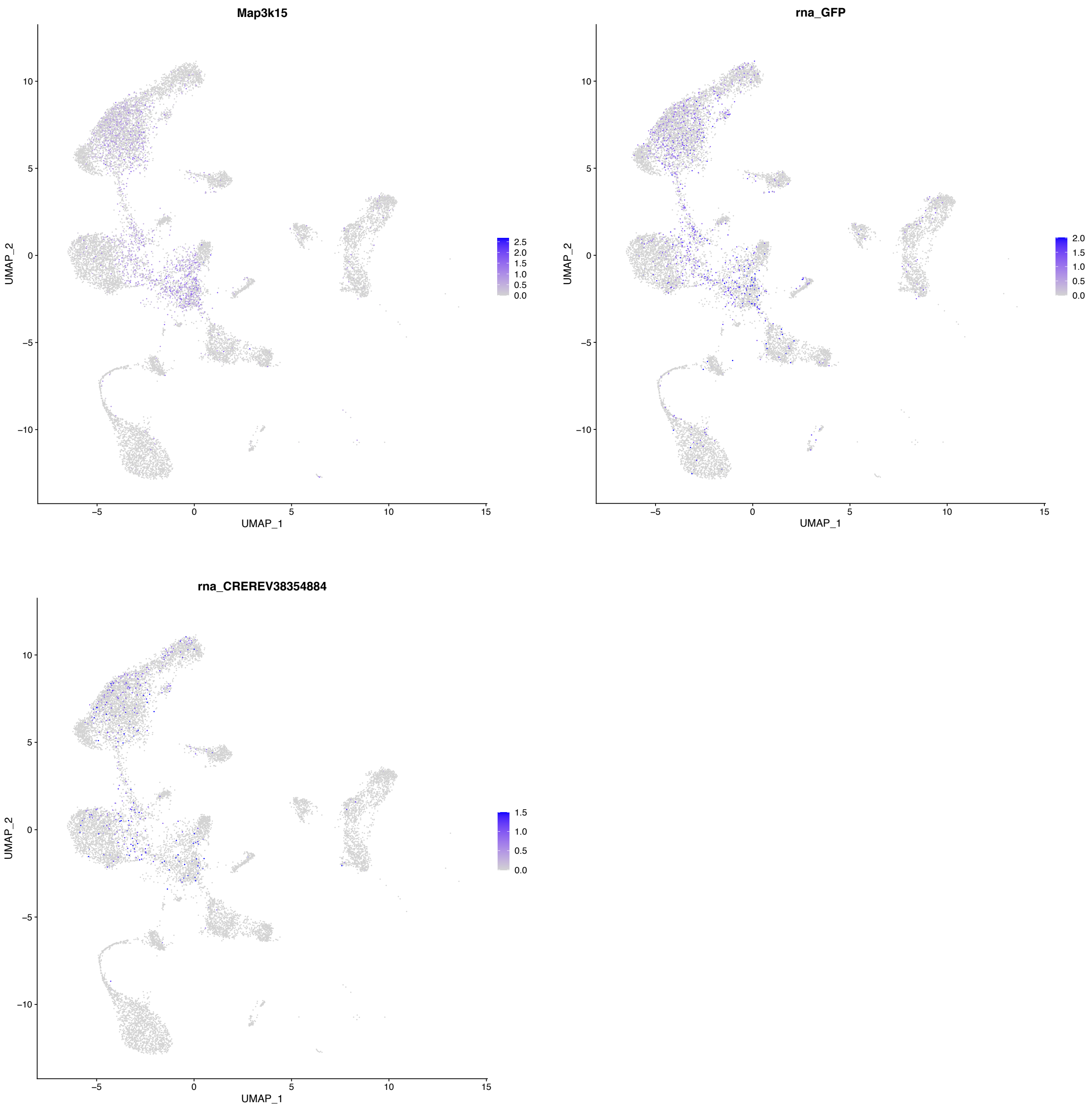

Supplemental Figure 5

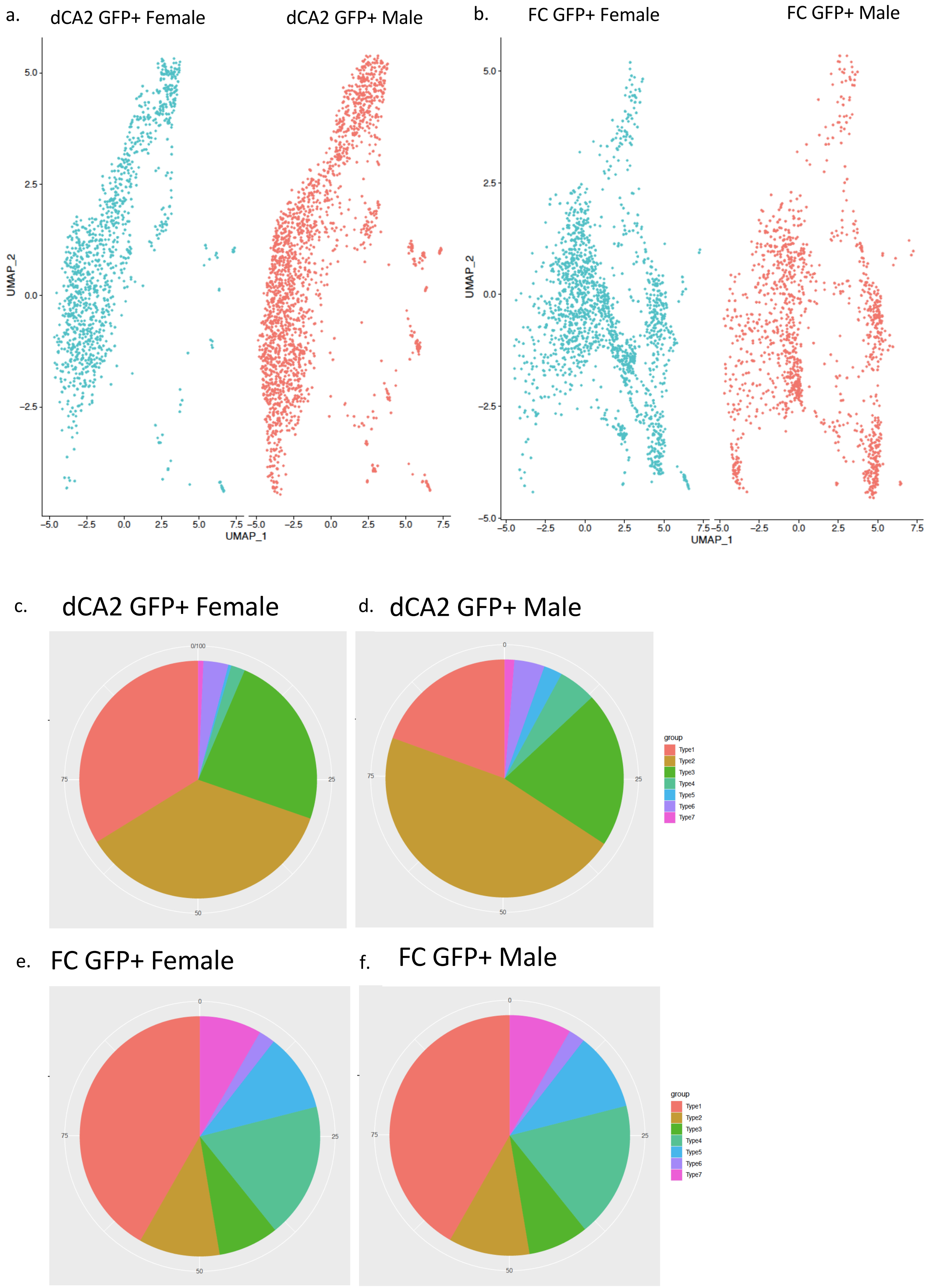

### Supplemental Figure 6

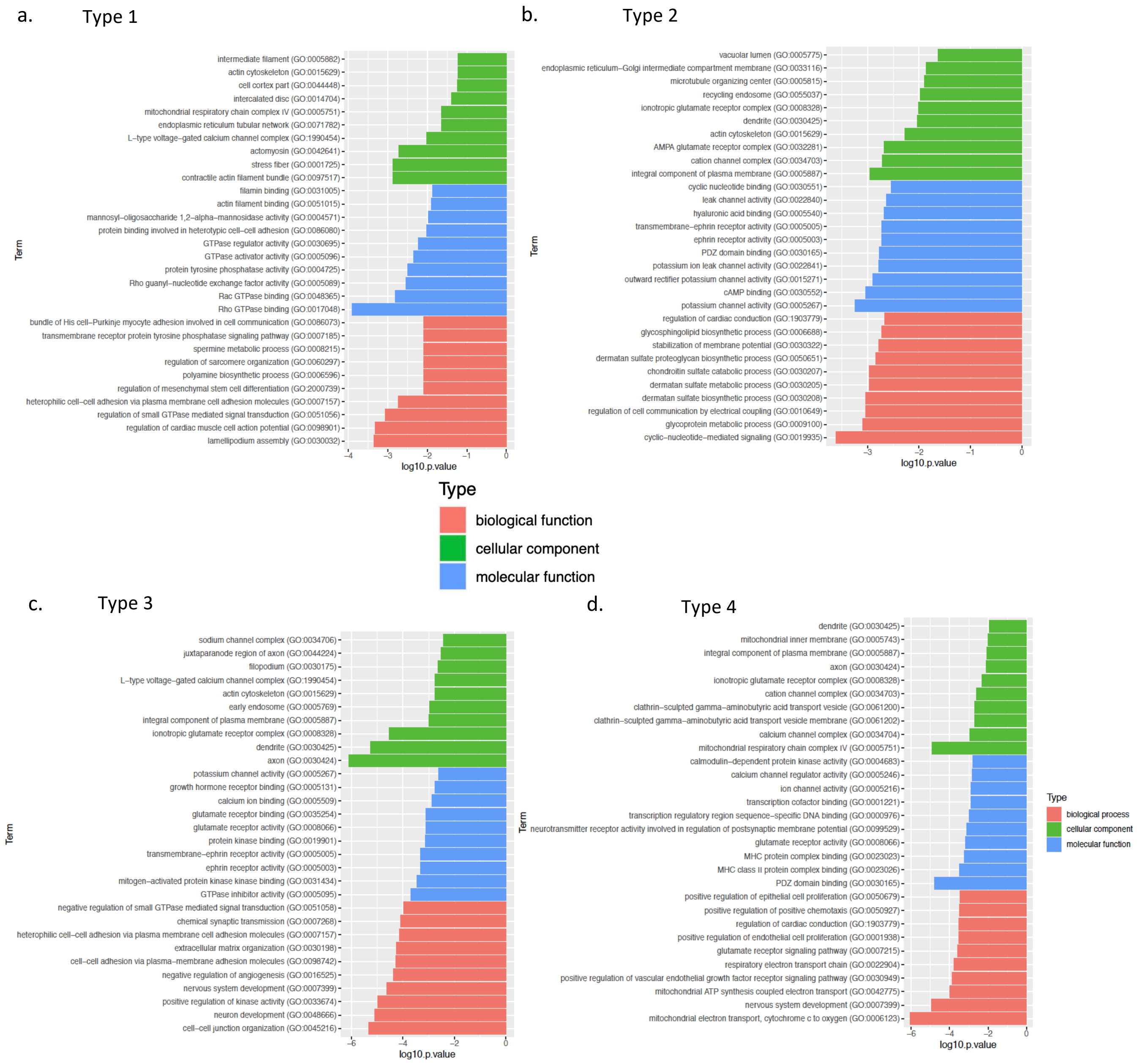

Supplemental Figure 7

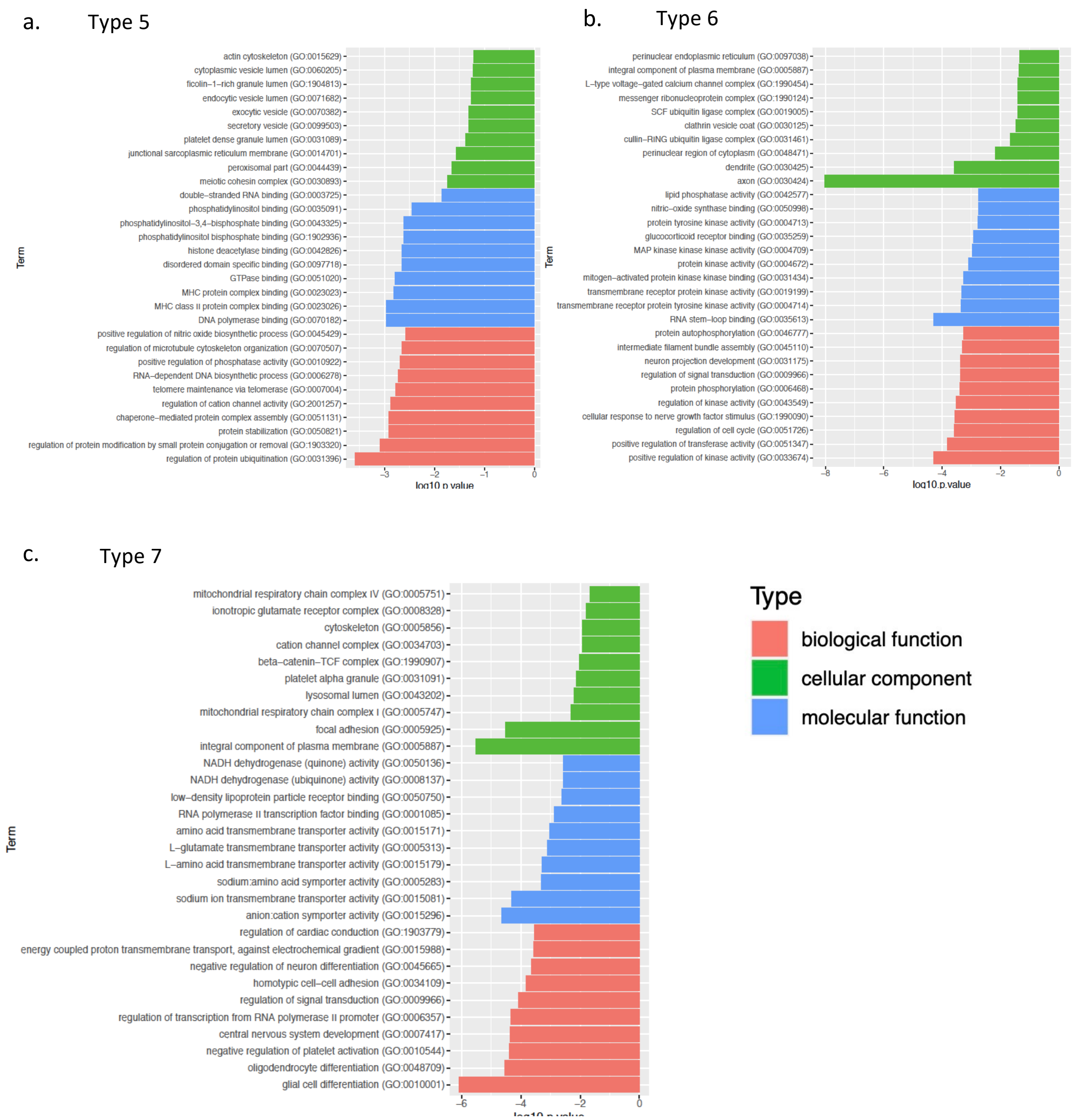

Supplemental Figure 8

a.

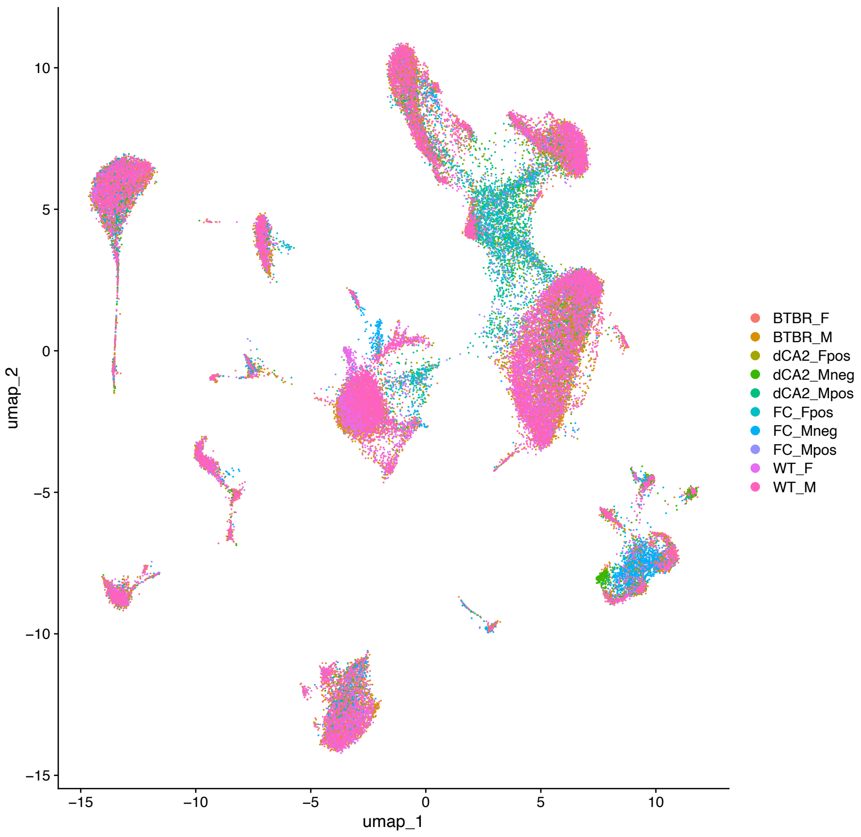


b.

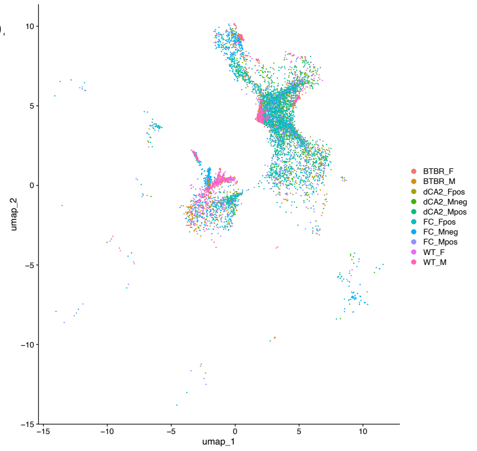


c. Gfp

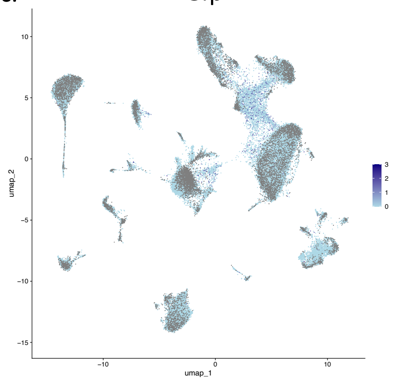


d. Cre

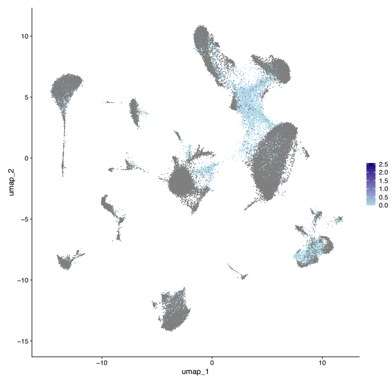


e. Avpr1b

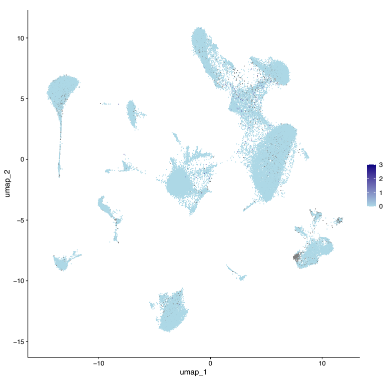

### Supplemental Figure 9

#### Microglia

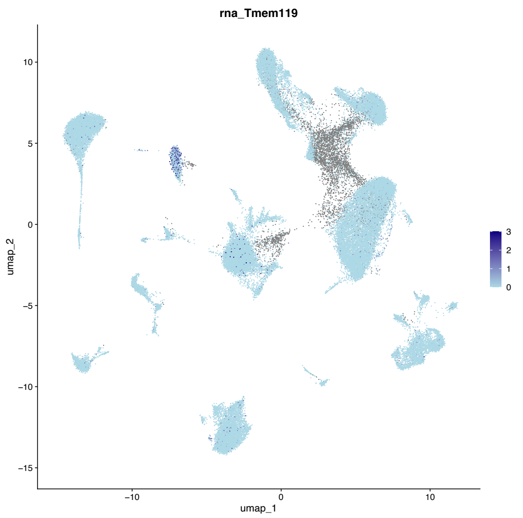


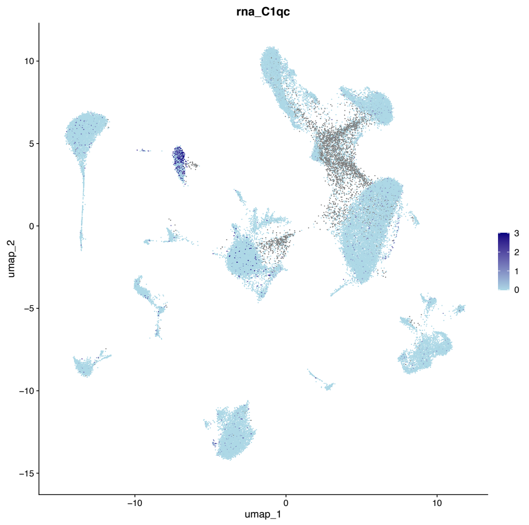


#### Oligodendrocytes

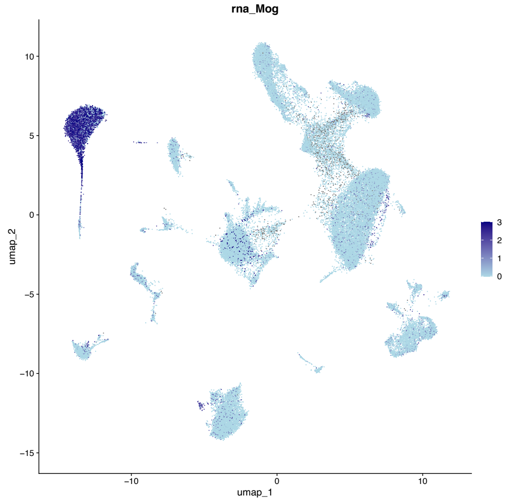


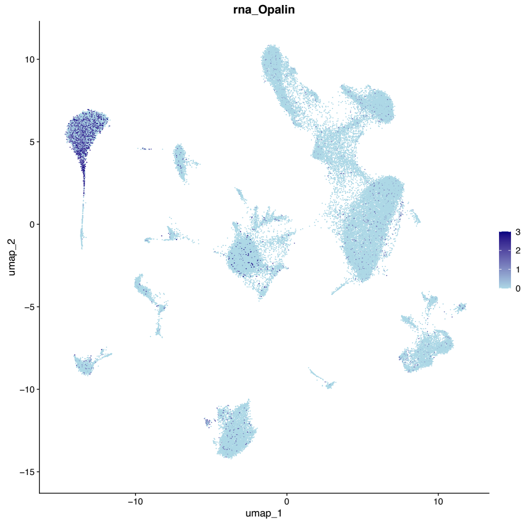


#### Oligodendrocyte Precursor Cell (OPC)

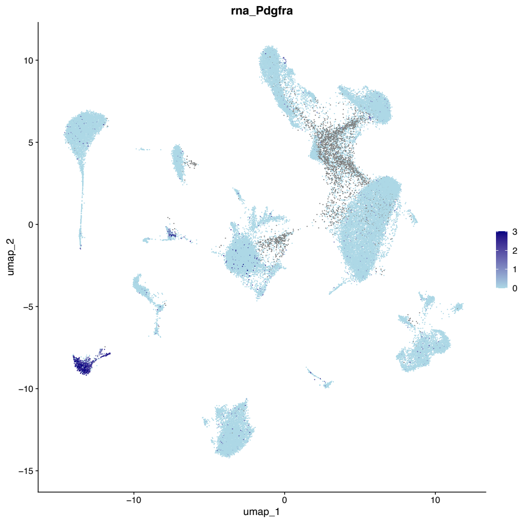


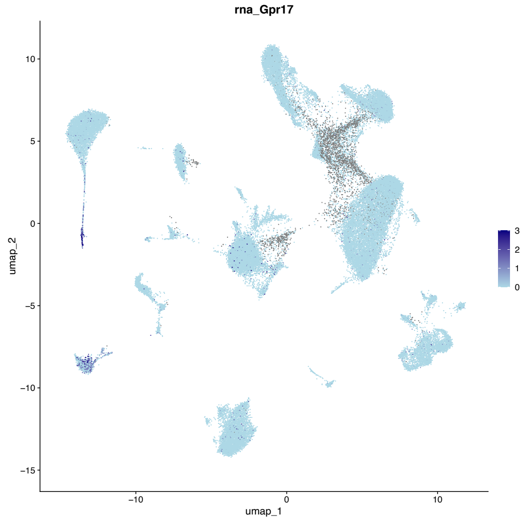

### Supplemental Figure 10

#### Pericytes

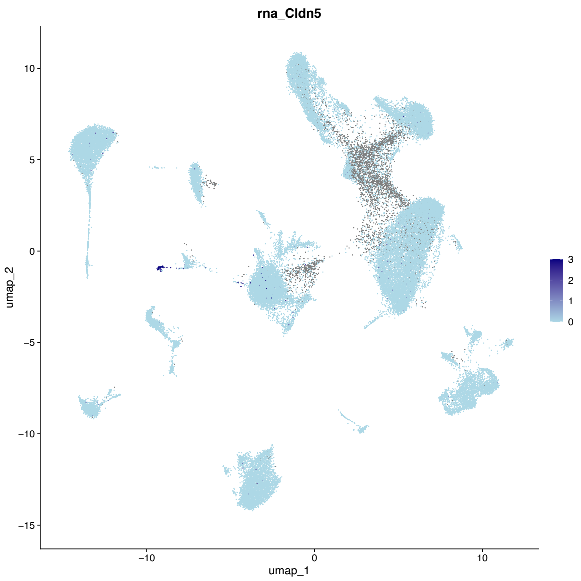


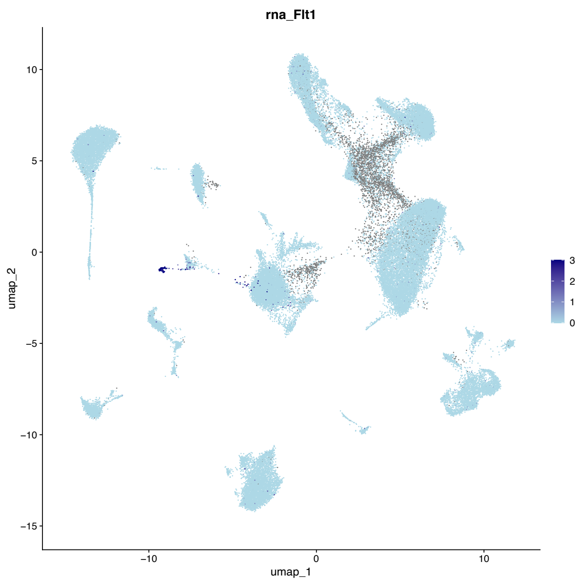


#### Astrocyte

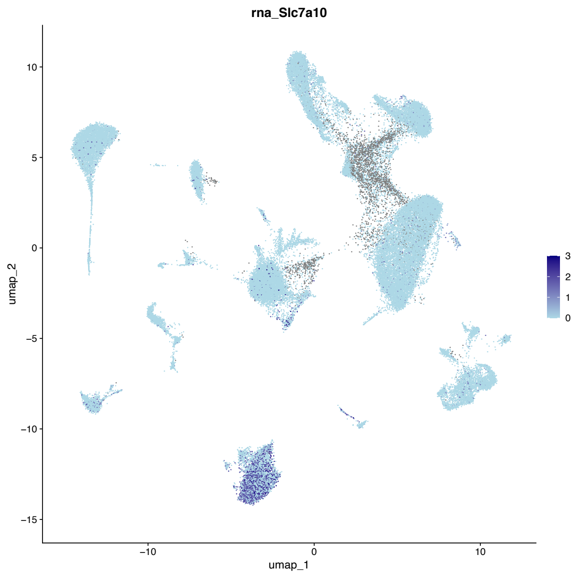


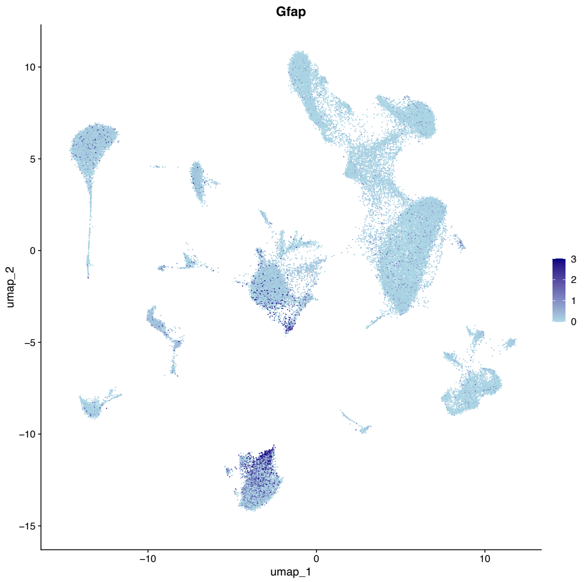


#### Neuronal

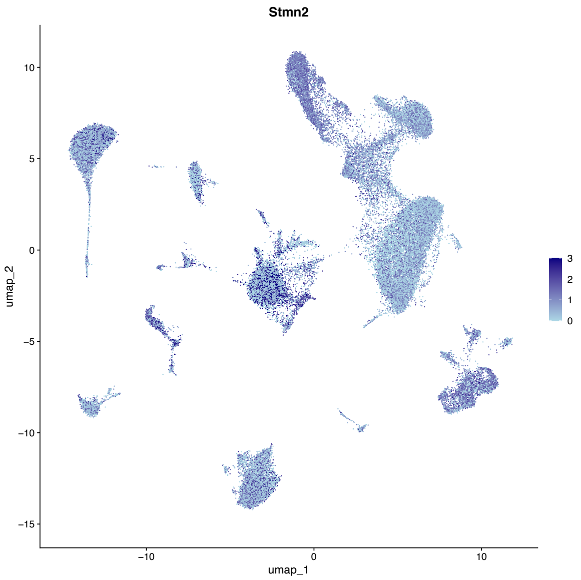


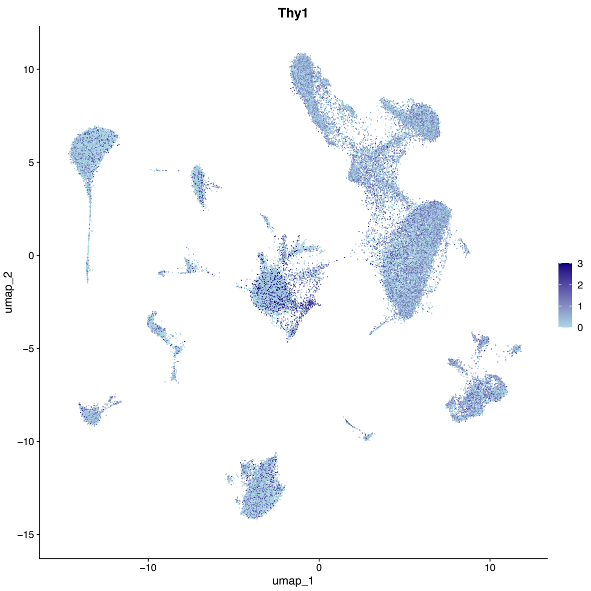

### Supplemental Figure 11

#### Neuronal

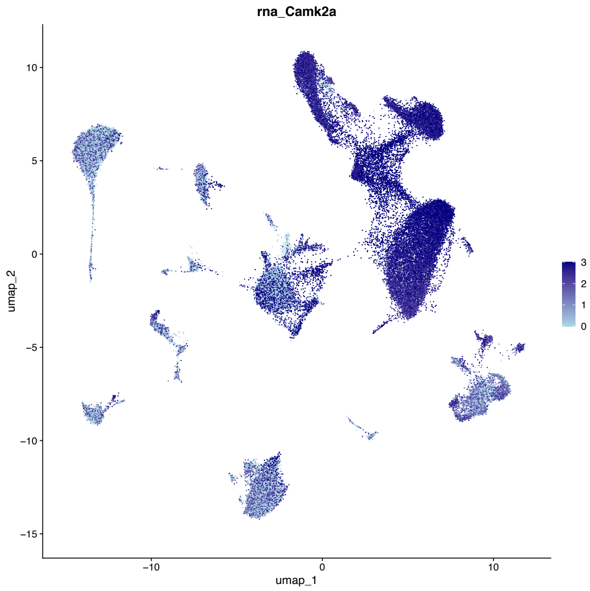


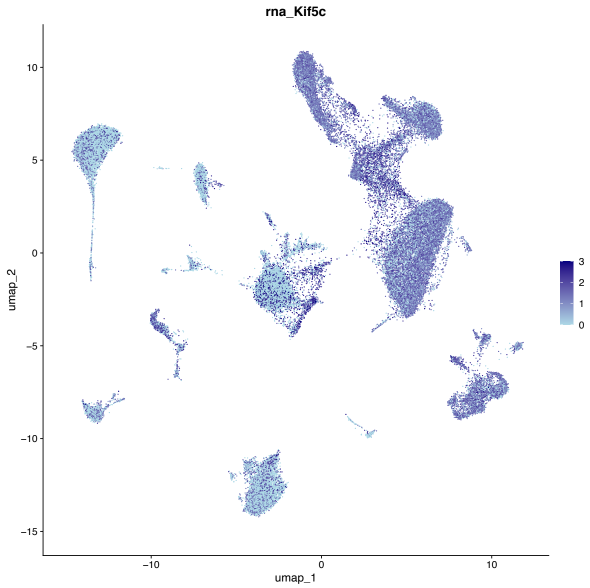


## CA1

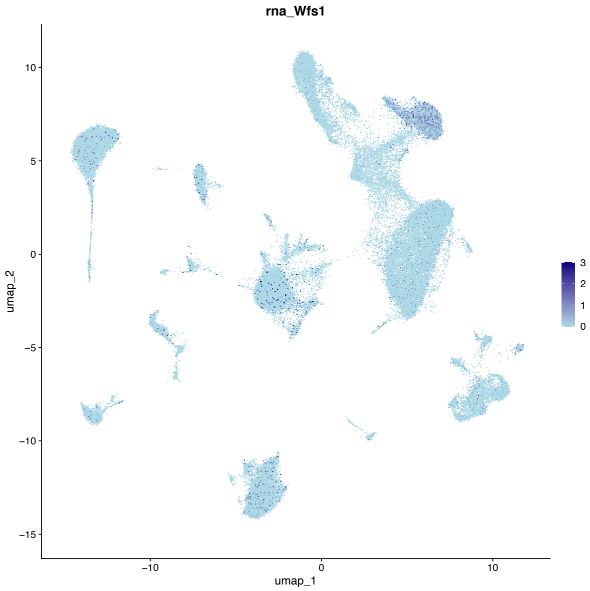


## CA3

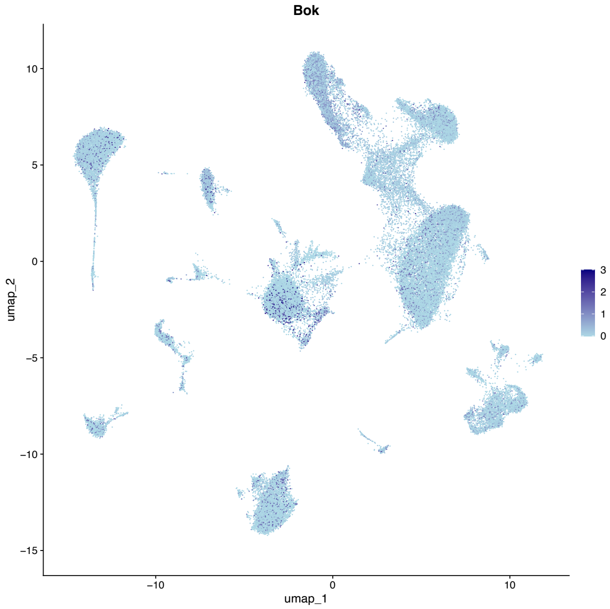
