## Supplementary material for "Investigation of the Fasciola Cinereum, Absent in BTBR mice, and Comparison with the Hippocampal Area CA2": BTBR vs WT Integration

This is an R Markdown (http://rmarkdown.rstudio.com) Notebook. When you execute code within the notebook, the results appear beneath the code.

Try executing this chunk by clicking the *Run* button within the chunk or by placing your cursor inside it and pressing *Cmd+Shift+Enter*.

```
library(Seurat)
library(dplyr)
library(magrittr)
library("ggplot2")
library(grid)
library(reshape2)
library("scales")
```

dCA2 Male GFP positive sample

```
#attach data for Male GFP positive sample (sample_1_dCA2_M)
#get working directory
getwd()
#Set working directory where data file located
setwd("/Users/lees44/R_project/SUN1GFP_MYC_CREREV38354884_OUTPUT/sample_1_dCA2_M")
#check folder and list files
list.files("/Users/lees44/R_project/SUN1GFP_MYC_CREREV38354884_OUTPUT/sample_1_dCA2_
M")
#Load M1_GFPve_pos data
dCA2_M1_GFPve_pos_barcodes.data <- Read10X(data.dir = "/Users/lees44/R_project/SUN1GF
P_MYC_CREREV38354884_OUTPUT/sample_1_dCA2_M")
colnames(dCA2_M1_GFPve_pos_barcodes.data) = paste0(colnames(dCA2_M1_GFPve_pos_barcode
s.data),"dCA2_MI_GFPve_pos")
#check dimension for data
dim(dCA2_M1_GFPve_pos_barcodes.data)
#Initialize the Seurat object with the raw (non-normalized data).
dCA2_MI_GFPve_pos <- CreateSeuratObject(counts = dCA2_M1_GFPve_pos_barcodes.data, pro
ject = "dCA2", min.cells = 3, min.features = 200)
#only genes that are are expressed in 3 or more cells and cells with complexity of 20
0 genes or more
dCA2_MI_GFPve_pos@meta.data$protocol<- "dCA2_Mpos"
```

dCA2 Female GFP positive sample

```
#attach data for Female GFP positive sample (sample_2_dCA2_F)
#get working directory
getwd()
#Set working directory where data file located
setwd("/Users/lees44/R_project/SUN1GFP_MYC_CREREV38354884_OUTPUT/sample_2_dCA2_F")
#check folder and list files
list.files("/Users/lees44/R_project/SUN1GFP_MYC_CREREV38354884_OUTPUT/sample_2_dCA2_
F")
#Load M1_GFPve_pos data
dCA2_FI_GFPve_pos_barcodes.data <- Read10X(data.dir = "/Users/lees44/R_project/SUN1GF
P_MYC_CREREV38354884_OUTPUT/sample_2_dCA2_F")
colnames(dCA2_FI_GFPve_pos_barcodes.data) = paste0(colnames(dCA2_FI_GFPve_pos_barcode
s.data),"dCA2_FI_GFPve_pos")
#check dimension for data
dim(dCA2_FI_GFPve_pos_barcodes.data)
#Initialize the Seurat object with the raw (non-normalized data).
dCA2_FI_GFPve_pos <- CreateSeuratObject(counts = dCA2_FI_GFPve_pos_barcodes.data, pro
ject = "dCA2", min.cells = 3, min.features = 200)
dCA2_FI_GFPve_pos@meta.data$protocol <- "dCA2_Fpos"
```

dCA2 GFP negative sample

```
#attach data for GFP negative sample (sample_3_GFP_negative)
#get working directory
getwd()
#Set working directory where data file located
setwd("/Users/lees44/R_project/SUN1GFP_MYC_CREREV38354884_OUTPUT/sample_3_GFP_ve")
#check folder and list files
list.files("/Users/lees44/R_project/SUN1GFP_MYC_CREREV38354884_OUTPUT/sample_3_GFP_v
e")
#Load M1_GFPve_pos data
dCA2_MI_GFPve_neg_barcodes.data <- Read10X(data.dir = "/Users/lees44/R_project/SUN1GF
P_MYC_CREREV38354884_OUTPUT/sample_3_GFP_ve")
#check dimension for data
dim(dCA2_MI_GFPve_neg_barcodes.data)
colnames(dCA2_MI_GFPve_neg_barcodes.data) = paste0(colnames(dCA2_MI_GFPve_neg_barcode
s.data),"dCA2_MI_GFPve_neg_barcodes")
#Initialize the Seurat object with the raw (non-normalized data).
dCA2_MI_GFPve_neg <- CreateSeuratObject(counts = dCA2_MI_GFPve_neg_barcodes.data, pro
ject = "dCA2", min.cells = 3, min.features = 200)
dCA2_MI_GFPve_neg@meta.data$protocol<-"dCA2_Mneg"
```

FC male GFP positive

```
#attach data for Male GFP positive sample (sample_1_FC_M)
#get working directory
getwd()
#Set working directory where data file located
setwd("/Users/lees44/R_project/FC_SUN1GFP_MYC_042021_OUTPUT/sample_1_FC_M")
#check folder and list files
list.files("/Users/lees44/R_project/FC_SUN1GFP_MYC_042021_OUTPUT/sample_1_FC_M")
#Load M1_GFPve_pos data
FC_M1_GFPve_pos_barcodes.data <- Read10X(data.dir = "/Users/lees44/R_project/FC_SUN1G
FP_MYC_042021_OUTPUT/sample_1_FC_M")
colnames(FC_M1_GFPve_pos_barcodes.data) = paste0(colnames(FC_M1_GFPve_pos_barcodes.da
ta),"FC_MI_GFPve_pos")
#check dimension for data
dim(FC_M1_GFPve_pos_barcodes.data)
#Initialize the Seurat object with the raw (non-normalized data).
FC_MI_GFPve_pos <- CreateSeuratObject(counts = FC_M1_GFPve_pos_barcodes.data, project
= "dCA2", min.cells = 3, min.features = 200)
#only genes that are are expressed in 3 or more cells and cells with complexity of 20
0 genes or more
FC_MI_GFPve_pos@meta.data$protocol<- "FC_Mpos"
```

```
#How many genes are you left with? How many cells?
dim(FC_MI_GFPve_pos)
```

```
#How much memory does a sparse matrix take up relative to a dense matrix
object.size(FC_M1_GFPve_pos_barcodes.data)
object.size(as.matrix(FC_M1_GFPve_pos_barcodes.data))
```

```
#Filtering low-quality cells
#Look at the summary counts for genes and cells
counts_per_cell <- Matrix::colSums(FC_M1_GFPve_pos_barcodes.data)
cat("counts per cell: ", counts_per_cell[1:5], "\n") ## counts for first 5 cells
```

```
counts_per_gene <- Matrix::rowSums(FC_M1_GFPve_pos_barcodes.data)
cat("counts per gene: ", counts_per_gene[1:5], "\n")  ## counts for first 5 genes
```

```
genes_per_cell <- Matrix::colSums(FC_M1_GFPve_pos_barcodes.data > 0) # count gene onl
y if it has non-zero reads mapped.
cat("counts for non-zero genes: ", genes_per_cell[1:5])  ## counts for first 5 genes
```

```
hist(log10(counts_per_cell+1),main='counts per cell',col='wheat')
```

```
hist(log10(genes_per_cell+1), main='genes per cell', col='wheat')
```

```
plot(counts_per_cell, genes_per_cell, log='xy', col='wheat')
title('counts vs genes per cell')
```

```
#plot cells ranked by their number of detected genes
#Here we rank each cell by its library complexity, ie the number of genes detected pe
r cell. This is a very useful plot as it shows the distribution of library complexity
in the sequencing run. One can use this plot to investigate observations (potential c
ells) that are actually failed libraries (lower end outliers) or observations that ar
e cell doublets (higher end outliers).
plot(sort(genes_per_cell), xlab='cell', log='y', main='genes per cell (ordered)')
```

BTBR female sample

```
#attach data for BTBR female sample (sample BTBR_F)
#get working directory
getwd()
#Set working directory where data file located
setwd("/Users/lees44/R_project/BTBR/1_BTBR_F/filtered_feature_bc_matrix")
#check folder and list files
list.files("/Users/lees44/R_project/BTBR/1_BTBR_F")
#Load M1_GFPve_pos data
BTBR_F_barcodes.data <- Read10X(data.dir = "/Users/lees44/R_project/BTBR/1_BTBR_F/fil
tered_feature_bc_matrix/")
colnames(BTBR_F_barcodes.data) = paste0(colnames(BTBR_F_barcodes.data),"BTBR_F")
#check dimension for data
dim(BTBR_F_barcodes.data)
#Initialize the Seurat object with the raw (non-normalized data).
BTBR_F <- CreateSeuratObject(counts = BTBR_F_barcodes.data, project = "dCA2", min.cel
ls = 3, min.features = 200)
#only genes that are are expressed in 3 or more cells and cells with complexity of 20
0 genes or more
BTBR_F@meta.data$protocol<- "BTBR_F"
```

BTBR male sample

```
#attach data for BTBR female sample (sample BTBR_M)
#get working directory
getwd()
#Set working directory where data file located
setwd("/Users/lees44/R_project/BTBR/2_BTBR_M/filtered_feature_bc_matrix")
#check folder and list files
list.files("/Users/lees44/R_project/BTBR/2_BTBR_M")
#Load M1_GFPve_pos data
BTBR_M_barcodes.data <- Read10X(data.dir = "/Users/lees44/R_project/BTBR/2_BTBR_M/fil
tered_feature_bc_matrix/")
colnames(BTBR_M_barcodes.data) = paste0(colnames(BTBR_M_barcodes.data),"BTBR_M")
#check dimension for data
dim(BTBR_M_barcodes.data)
#Initialize the Seurat object with the raw (non-normalized data).
BTBR_M <- CreateSeuratObject(counts = BTBR_M_barcodes.data, project = "dCA2", min.cel
ls = 3, min.features = 200)
#only genes that are are expressed in 3 or more cells and cells with complexity of 20
0 genes or more
BTBR_M@meta.data$protocol<- "BTBR_M"
```

WT female sample

```
#attach data for BTBR female sample (sample WT_F)
#get working directory
getwd()
#Set working directory where data file located
setwd("/Users/lees44/R_project/BTBR/3_WT_F/filtered_feature_bc_matrix")
#check folder and list files
list.files("/Users/lees44/R_project/BTBR/3_WT_F")
#Load M1_GFPve_pos data
WT_F_barcodes.data <- Read10X(data.dir = "/Users/lees44/R_project/BTBR/3_WT_F/filtere
d_feature_bc_matrix/")
colnames(WT_F_barcodes.data) = paste0(colnames(WT_F_barcodes.data),"WT_F")
#check dimension for data
dim(WT_F_barcodes.data)
#Initialize the Seurat object with the raw (non-normalized data).
WT_F <- CreateSeuratObject(counts = WT_F_barcodes.data, project = "dCA2", min.cells =
3, min.features = 200)
#only genes that are are expressed in 3 or more cells and cells with complexity of 20
0 genes or more
WT_F@meta.data$protocol<- "WT_F"
```

WT male sample

```
#attach data for WT male sample (sample WT_M)
#get working directory
getwd()
#Set working directory where data file located
setwd("/Users/lees44/R_project/BTBR/4_WT_M/filtered_feature_bc_matrix")
#check folder and list files
list.files("/Users/lees44/R_project/BTBR/4_WT_M")
#Load M1_GFPve_pos data
WT_M_barcodes.data <- Read10X(data.dir = "/Users/lees44/R_project/BTBR/4_WT_M/filtere
d_feature_bc_matrix/")
colnames(WT_M_barcodes.data) = paste0(colnames(WT_M_barcodes.data),"WT_M")
#check dimension for data
dim(WT_M_barcodes.data)
#Initialize the Seurat object with the raw (non-normalized data).
WT_M <- CreateSeuratObject(counts = WT_M_barcodes.data, project = "dCA2", min.cells =
3, min.features = 200)
#only genes that are are expressed in 3 or more cells and cells with complexity of 20
0 genes or more
WT_M@meta.data$protocol<- "WT_M"
```

FC Female GFP positive

```
#attach data for Female GFP positive sample (sample_2_FC_F)
#get working directory
getwd()
#Set working directory where data file located
setwd("/Users/lees44/R_project/FC_SUN1GFP_MYC_042021_OUTPUT/sample_2_FC_F")
#check folder and list files
list.files("/Users/lees44/R_project/FC_SUN1GFP_MYC_042021_OUTPUT/sample_2_FC_F")
#Load M1_GFPve_pos data
FC_FI_GFPve_pos_barcodes.data <- Read10X(data.dir = "/Users/lees44/R_project/FC_SUN1G
FP_MYC_042021_OUTPUT/sample_2_FC_F")
colnames(FC_FI_GFPve_pos_barcodes.data) = paste0(colnames(FC_FI_GFPve_pos_barcodes.da
ta),"FC_FI_GFPve_pos")
#check dimension for data
dim(FC_FI_GFPve_pos_barcodes.data)
#Initialize the Seurat object with the raw (non-normalized data).
FC_FI_GFPve_pos <- CreateSeuratObject(counts = FC_FI_GFPve_pos_barcodes.data, project
= "dCA2", min.cells = 3, min.features = 200)
FC_FI_GFPve_pos@meta.data$protocol <- "FC_Fpos"
```

FC GFP negative

```
#attach data for GFP negative sample (FC_sample_3_GFP_negative)
#get working directory
getwd()
#Set working directory where data file located
setwd("/Users/lees44/R_project/FC_SUN1GFP_MYC_042021_OUTPUT/sample_3_FC_GFP_ve")
#check folder and list files
list.files("/Users/lees44/R_project/FC_SUN1GFP_MYC_042021_OUTPUT/sample_3_FC_GFP_ve")
#Load M1_GFPve_pos data
FC_MI_GFPve_neg_barcodes.data <- Read10X(data.dir = "/Users/lees44/R_project/FC_SUN1G
FP_MYC_042021_OUTPUT/sample_3_FC_GFP_ve")
#check dimension for data
dim(FC_MI_GFPve_neg_barcodes.data)
colnames(FC_MI_GFPve_neg_barcodes.data) = paste0(colnames(FC_MI_GFPve_neg_barcodes.da
ta),"FC_MI_GFPve_neg_barcodes")
#Initialize the Seurat object with the raw (non-normalized data).
FC_MI_GFPve_neg <- CreateSeuratObject(counts = FC_MI_GFPve_neg_barcodes.data, project
= "dCA2", min.cells = 3, min.features = 200)
FC_MI_GFPve_neg@meta.data$protocol<-"FC_Mneg"
```

```
mito.features <- grep(pattern = "^mt-", x = rownames(x = dCA2_MI_GFPve_pos), value =
TRUE)
percent.mito <- Matrix::colSums(x = GetAssayData(object = dCA2_MI_GFPve_pos, slot = '
counts')[mito.features, ]) / Matrix::colSums(x = GetAssayData(object = dCA2_MI_GFPve_
pos, slot = 'counts'))
dCA2_MI_GFPve_pos[['percent.mito']] <- percent.mito
```

```
mito.features <- grep(pattern = "^mt-", x = rownames(x = dCA2_FI_GFPve_pos), value =
TRUE)
percent.mito <- Matrix::colSums(x = GetAssayData(object = dCA2_FI_GFPve_pos, slot = '
counts')[mito.features, ]) / Matrix::colSums(x = GetAssayData(object = dCA2_FI_GFPve_
pos, slot = 'counts'))
dCA2_FI_GFPve_pos[['percent.mito']] <- percent.mito
```

```
mito.features <- grep(pattern = "^mt-", x = rownames(x = dCA2_MI_GFPve_neg), value =
TRUE)
percent.mito <- Matrix::colSums(x = GetAssayData(object = dCA2_MI_GFPve_neg, slot = '
counts')[mito.features, ]) / Matrix::colSums(x = GetAssayData(object = dCA2_MI_GFPve_
neg, slot = 'counts'))
dCA2_MI_GFPve_neg[['percent.mito']] <- percent.mito
```

```
mito.features <- grep(pattern = "^mt-", x = rownames(x = FC_MI_GFPve_pos), value = TR
UE)
percent.mito <- Matrix::colSums(x = GetAssayData(object = FC_MI_GFPve_pos, slot = 'co
unts')[mito.features, ]) / Matrix::colSums(x = GetAssayData(object = FC_MI_GFPve_pos,
slot = 'counts'))
FC_MI_GFPve_pos[['percent.mito']] <- percent.mito
```

```
mito.features <- grep(pattern = "^mt-", x = rownames(x = FC_FI_GFPve_pos), value = TR
UE)
percent.mito <- Matrix::colSums(x = GetAssayData(object = FC_FI_GFPve_pos, slot = 'co
unts')[mito.features, ]) / Matrix::colSums(x = GetAssayData(object = FC_FI_GFPve_pos,
slot = 'counts'))
FC_FI_GFPve_pos[['percent.mito']] <- percent.mito
```

```
mito.features <- grep(pattern = "^mt-", x = rownames(x = FC_MI_GFPve_neg), value = TR
UE)
percent.mito <- Matrix::colSums(x = GetAssayData(object = FC_MI_GFPve_neg, slot = 'co
unts')[mito.features, ]) / Matrix::colSums(x = GetAssayData(object = FC_MI_GFPve_neg,
slot = 'counts'))
FC_MI_GFPve_neg[['percent.mito']] <- percent.mito
```

```
mito.features <- grep(pattern = "^mt-", x = rownames(x = BTBR_F), value = TRUE)
percent.mito <- Matrix::colSums(x = GetAssayData(object = BTBR_F, slot = 'counts')[mi
to.features, ]) / Matrix::colSums(x = GetAssayData(object = BTBR_F, slot = 'counts'))
BTBR_F[['percent.mito']] <- percent.mito
```

```
mito.features <- grep(pattern = "^mt-", x = rownames(x = BTBR_M), value = TRUE)
percent.mito <- Matrix::colSums(x = GetAssayData(object = BTBR_M, slot = 'counts')[mi
to.features, ]) / Matrix::colSums(x = GetAssayData(object = BTBR_M, slot = 'counts'))
BTBR_M[['percent.mito']] <- percent.mito
```

```
mito.features <- grep(pattern = "^mt-", x = rownames(x = WT_F), value = TRUE)
percent.mito <- Matrix::colSums(x = GetAssayData(object = WT_F, slot = 'counts')[mit
o.features, ]) / Matrix::colSums(x = GetAssayData(object = WT_F, slot = 'counts'))
WT_F[['percent.mito']] <- percent.mito
```

```
mito.features <- grep(pattern = "^mt-", x = rownames(x = WT_M), value = TRUE)
percent.mito <- Matrix::colSums(x = GetAssayData(object = WT_M, slot = 'counts')[mit
o.features, ]) / Matrix::colSums(x = GetAssayData(object = WT_M, slot = 'counts'))
WT_M[['percent.mito']] <- percent.mito
```

#########filtering and feature genes###############

```
dCA2_MI_GFPve_pos <- subset(x = dCA2_MI_GFPve_pos, subset = nCount_RNA > 700 & nCount
_RNA < 15000 & percent.mito < 0.20)
dCA2_FI_GFPve_pos <- subset(x = dCA2_FI_GFPve_pos, subset = nCount_RNA > 700 & nCount
_RNA < 15000 & percent.mito < 0.20)
dCA2_MI_GFPve_neg <- subset(x = dCA2_MI_GFPve_neg, subset = nCount_RNA > 700 & nCount
_RNA < 15000 & percent.mito < 0.20)
FC_MI_GFPve_pos <- subset(x = FC_MI_GFPve_pos, subset = nCount_RNA > 700 & nCount_RNA
< 15000 & percent.mito < 0.20)
FC_FI_GFPve_pos <- subset(x = FC_FI_GFPve_pos, subset = nCount_RNA > 700 & nCount_RNA
< 15000 & percent.mito < 0.20)
FC_MI_GFPve_neg <- subset(x = FC_MI_GFPve_neg, subset = nCount_RNA > 700 & nCount_RNA
< 15000 & percent.mito < 0.20)
BTBR_F <- subset(x = BTBR_F, subset = nCount_RNA > 700 & nCount_RNA < 15000 & percen
t.mito < 0.20)
BTBR_M <- subset(x = BTBR_M, subset = nCount_RNA > 700 & nCount_RNA < 15000 & percen
t.mito < 0.20)
WT_F <- subset(x = WT_F, subset = nCount_RNA > 700 & nCount_RNA < 15000 & percent.mit
o < 0.20)
BTBR_M <- subset(x = BTBR_M, subset = nCount_RNA > 700 & nCount_RNA < 15000 & percen
t.mito < 0.20)
```

```
dCA2_MI_GFPve_pos<- NormalizeData(object = dCA2_MI_GFPve_pos,verbose = FALSE)
dCA2_FI_GFPve_pos<- NormalizeData(object = dCA2_FI_GFPve_pos,verbose = FALSE)
dCA2_MI_GFPve_neg<- NormalizeData(object = dCA2_MI_GFPve_neg,verbose = FALSE)
FC_MI_GFPve_pos<- NormalizeData(object = FC_MI_GFPve_pos,verbose = FALSE)
FC_FI_GFPve_pos<- NormalizeData(object = FC_FI_GFPve_pos,verbose = FALSE)
FC_MI_GFPve_neg<- NormalizeData(object = FC_MI_GFPve_neg,verbose = FALSE)
BTBR_F<- NormalizeData(object = BTBR_F,verbose = FALSE)
BTBR_M<- NormalizeData(object = BTBR_M,verbose = FALSE)
WT_F<- NormalizeData(object = WT_F,verbose = FALSE)
WT_M<- NormalizeData(object = WT_M,verbose = FALSE)
```

###########nfeatures#######

```
dCA2_MI_GFPve_pos<- FindVariableFeatures(object =dCA2_MI_GFPve_pos,selection.method =
"vst", nfeatures = 2000, verbose = FALSE)
dCA2_FI_GFPve_pos<- FindVariableFeatures(object =dCA2_FI_GFPve_pos,selection.method =
"vst", nfeatures = 2000, verbose = FALSE)
dCA2_MI_GFPve_neg<- FindVariableFeatures(object =dCA2_MI_GFPve_neg,selection.method =
"vst", nfeatures = 2000, verbose = FALSE)
FC_MI_GFPve_pos<- FindVariableFeatures(object =FC_MI_GFPve_pos,selection.method = "vs
t", nfeatures = 2000, verbose = FALSE)
FC_FI_GFPve_pos<- FindVariableFeatures(object =FC_FI_GFPve_pos,selection.method = "vs
t", nfeatures = 2000, verbose = FALSE)
FC_MI_GFPve_neg<- FindVariableFeatures(object =FC_MI_GFPve_neg,selection.method = "vs
t", nfeatures = 2000, verbose = FALSE)
BTBR_F<- FindVariableFeatures(object =BTBR_F,selection.method = "vst", nfeatures = 20
00, verbose = FALSE)
BTBR_M<- FindVariableFeatures(object =BTBR_M,selection.method = "vst", nfeatures = 20
00, verbose = FALSE)
WT_F<- FindVariableFeatures(object =WT_F,selection.method = "vst", nfeatures = 2000,
verbose = FALSE)
WT_M<- FindVariableFeatures(object =WT_M,selection.method = "vst", nfeatures = 2000,
verbose = FALSE)
```

#########integration of groups##########################

```
#integration of multiple groups using vector (x, y=c(1,2) and split by group.
dCA2.combined = merge(dCA2_MI_GFPve_pos, y = c(dCA2_FI_GFPve_pos,  dCA2_MI_GFPve_neg,
FC_MI_GFPve_pos, FC_FI_GFPve_pos, FC_MI_GFPve_neg, BTBR_F, BTBR_M, WT_F, WT_M), add.c
ell.ids = c( "dCA2_Mpos", "dCA2_Fpos", "dCA2_Mneg", "FC_Mpos", "FC_Fpos", "FC_Mneg",
"BTBR_F", "BTBR_M", "WT_F", "WT_M"), project = "protocol")
```

```
#integration of multiple groups using vector (x, y=c(1,2) and split by group.
dCA2.combined = merge(BTBR_F, y = c(dCA2_FI_GFPve_pos, dCA2_MI_GFPve_pos, dCA2_MI_GFP
ve_neg, FC_MI_GFPve_pos, FC_FI_GFPve_pos, FC_MI_GFPve_neg,BTBR_M, WT_F, WT_M), add.ce
ll.ids = c("BTBR_F","dCA2_Fpos", "dCA2_Mpos", "dCA2_Mneg", "FC_Mpos", "FC_Fpos", "FC_
Mneg","BTBR_M", "WT_F", "WT_M"), project ="protocol")
```

```
dCA2.list <-SplitObject(dCA2.combined, split.by = "protocol")
reference.list <-dCA2.list[c("BTBR_F","dCA2_Fpos", "dCA2_Mpos", "dCA2_Mneg", "FC_Mpo
s", "FC_Fpos", "FC_Mneg","BTBR_M", "WT_F", "WT_M")]
dCA2.anchors <- FindIntegrationAnchors(object.list = reference.list, dims = 1:30)
```

############integrate data##############

```
#intgrated analysis
dCA2_FC_integrated.combined <-IntegrateData(anchorset = dCA2.anchors, dims = 1:30)
```

```
library(usethis)
usethis::edit_r_environ()
```

```
library(ggplot2)
library(cowplot)
DefaultAssay(dCA2_FC_integrated.combined) <- "integrated"
```

```
set.seed(1234)
```

```
dCA2_FC_integrated.combined <- ScaleData(dCA2_FC_integrated.combined, verbose = FALS
E)
```

```
dCA2_FC_integrated.combined <- RunPCA(dCA2_FC_integrated.combined, npcs = 50, verbose
= FALSE)
```

Elbow plot to determine pc value

```
mat <- GetAssayData(dCA2_FC_integrated.combined, slot = "scale.data")
pca <- dCA2_FC_integrated.combined[["pca"]]
total_variance <- sum(matrixStats::rowVars(mat))
eigValues = (pca@stdev)^2  ## EigenValues
varExplained = (eigValues / total_variance)*100
plot(varExplained,type='l')
```

```
dCA2_FC_integrated.combined <- RunPCA(dCA2_FC_integrated.combined, npcs = 30, verbose
= FALSE)
```

```
dCA2_FC_integrated.combined <- RunUMAP(dCA2_FC_integrated.combined, reduction = "pc
a", dims = 1:30)
```

```
set.seed(1234)
```

```
dim(dCA2_FC_integrated.combined)
```

```
p1<-DimPlot(dCA2_FC_integrated.combined, reduction = "umap", group.by = "protocol")
plot_grid(p1)
ggsave(file="/Users/lees44/R_project/BTBR/All_UMAP.pdf",width=10,height=10)
```

```
p1<-DimPlot(dCA2_FC_integrated.combined, reduction = "umap", split.by = "protocol")
plot_grid(p1)
ggsave(file="/Users/lees44/R_project/BTBR/all_UMAP_split_group.pdf",width=10,height=10)
```

```
dCA2_FC_integrated.combined <- FindNeighbors(dCA2_FC_integrated.combined, reduction = "pca", dims = 1:30)
```

```
dCA2_FC_integrated.combined <- FindClusters(dCA2_FC_integrated.combined, resolution = 0.1)
```

```
#clustering by population
p1<-DimPlot(object = dCA2_FC_integrated.combined, reduction = "umap", group.by = "integrated_snn_res.0.1", label = TRUE, repel = TRUE)
plot_grid(p1)
ggsave(file="/Users/lees44/R_project/BTBR/all_Clustering.pdf",width=10,height=10)
```

```
p1<-DimPlot(dCA2_FC_integrated.combined, reduction = "umap", split.by = "protocol")
plot_grid(p1)
ggsave(file="/Users/lees44/R_project/BTBR/all_Clustering_splited_group.pdf",width=10,height=10)
```

##################defining cell types #####################################################

Feature plot is used and cell type was classified in supervised manner .

```
# CA2 markers
F<-FeaturePlot(object =dCA2_FC_integrated.combined, features =  c("Avpr1b","Amigo2", "Map3k15", "Rgs14"), blend.threshold = 0.5, min.cutoff = 0, max.cutoff = 3, cols = c("lightblue", "navyblue"))
ggsave(file="/Users/lees44/R_project/BTBR/all_dCA2.pdf",width=10,height=10)
plot(F)
```

```
#Avpr1b
F<-FeaturePlot(object =dCA2_FC_integrated.combined, features =  c("Avpr1b"), blend.threshold = 0.5, min.cutoff = 0, max.cutoff = 3, cols = c("lightblue", "navyblue"))
ggsave(file="/Users/lees44/R_project/BTBR/all_dCA2_Avpr1b.pdf",width=10,height=10)
plot(F)
```

```
#Amigo2
F<-FeaturePlot(object =dCA2_FC_integrated.combined, features =  c("Amigo2"), blend.threshold = 0.5, min.cutoff = 0, max.cutoff = 3, cols = c("lightblue", "navyblue"))
ggsave(file="/Users/lees44/R_project/BTBR/all_dCA2_Amigo2.pdf",width=10,height=10)
plot(F)
```

```
# Map3k15
F<-FeaturePlot(object =dCA2_FC_integrated.combined, features =  c("Map3k15"), blend.threshold = 0.5, min.cutoff = 0, max.cutoff = 3, cols = c("lightblue", "navyblue"))
ggsave(file="/Users/lees44/R_project/BTBR/all_dCA2_Map3k15.pdf",width=10,height=10)
plot(F)
```

```
# Rgs14
F<-FeaturePlot(object =dCA2_FC_integrated.combined, features =  c("Rgs14"), blend.threshold = 0.5, min.cutoff = 0, max.cutoff = 3, cols = c("lightblue", "navyblue"))
ggsave(file="/Users/lees44/R_project/BTBR/all_dCA2_Rgs14.pdf",width=10,height=10)
plot(F)
```

```
# CRE
F<-FeaturePlot(object =dCA2_FC_integrated.combined, features =  c("CREREV38354884"), blend.threshold = 0.5, min.cutoff = 0, max.cutoff = 3, cols = c("lightblue", "navyblue"))
ggsave(file="/Users/lees44/R_project/BTBR/all_dCA2_CRE.pdf",width=10,height=10)
plot(F)
```

```
# GFP
F<-FeaturePlot(object =dCA2_FC_integrated.combined, features =  c("GFP"), blend.threshold = 0.5, min.cutoff = 0, max.cutoff = 3, cols = c("lightblue", "navyblue"))
ggsave(file="/Users/lees44/R_project/BTBR/all_dCA2_GFP.pdf",width=10,height=10)
plot(F)
```

Assign clusters to cell types

```
#celltypes asigned based on that gene marker expression
new.ident <- c("CA3/Granular cell", "CA3 neurons", "CA2 neurons", "Oligdendrocyte", "Astrocyte", "CA3 neuron/non-granular", "CA1/non-granular", "Neuron1", "Microglia", "OPC", "Neuron2", "Pericyte", "Neuron3", "Neuron4", "Neuron5", "Neuron6", "Neuron7", "Neuron8", "Neuron9", "Neuron10")
names(x = new.ident) <- levels(x =dCA2_FC_integrated.combined)
dCA2_FC_integrated.combined<- RenameIdents(object =dCA2_FC_integrated.combined, new.ident)
```

```
color<-c("#19647e","#ffc857","#9B0C1E","#999999","#E69F00","#56B4E9","#009E73","#F0E4
42","#0007B2","#D55E00","#C79A70","#E67F00","#676833","#56B4E9","#b7b7b7","#E8C6C7","#
4b3f72","#ffc857","#19647e","#B99A69", "#009E73", "#F04421", "#0072B2", "#D55E00","#C
C9A70", "#CC6666", "#9999CC", "#66CC99", "#4bbb70")
```

```
DimPlot(object = dCA2_FC_integrated.combined, reduction = "umap", label = FALSE, repe
l = TRUE,cols=color,label.size = 3,pt.size=0.5)
ggsave(file="/Users/lees44/R_project/BTBR/Neurotype_All.pdf",width=10,height=7)
```

```
# microglia and oligo clusters, (Tmem119, C1qc = microglia)
F<-FeaturePlot(object =dCA2_FC_integrated.combined, features =  c("Tmem119","C1qc"),
blend.threshold = 0.5, min.cutoff = 0, max.cutoff = 3, cols = c("lightblue", "navyblu
e"))
ggsave(file="/Users/lees44/R_project/BTBR/Microglia_clusters.pdf",width=10,height=10)
plot(F)
```


```
# microglia and oligo clusters, (Tmem119, C1qc = microglia)
F<-FeaturePlot(object =dCA2_FC_integrated.combined, features =  c("Tmem119","C1qc"),
blend.threshold = 0.5, min.cutoff = 0, max.cutoff = 3, cols = c("lightblue", "navyblu
e"))
ggsave(file="/Users/lees44/R_project/BTBR/Microglia_clusters.pdf",width=10,height=10)
plot(F)
```


```
# microglia and oligo clusters, (Tmem119 = microglia)
F<-FeaturePlot(object =dCA2_FC_integrated.combined, features =  c("Tmem119"), blend.t
hreshold = 0.5, min.cutoff = 0, max.cutoff = 3, cols = c("lightblue", "navyblue"))
ggsave(file="/Users/lees44/R_project/BTBR/Tmem119_clusters.pdf",width=10,height=10)
plot(F)
```

```
# microglia and oligo clusters, (C1qc = microglia)
F<-FeaturePlot(object =dCA2_FC_integrated.combined, features =  c("C1qc"), blend.thre
shold = 0.5, min.cutoff = 0, max.cutoff = 3, cols = c("lightblue", "navyblue"))
ggsave(file="/Users/lees44/R_project/BTBR/C1qc_clusters.pdf",width=10,height=10)
plot(F)
```

```
#oligo cluster (Mog, Opalin = Oligodendrocyte)
F<-FeaturePlot(object =dCA2_FC_integrated.combined, features =  c("Mog","Opalin"), bl
end.threshold = 0.5, min.cutoff = 0, max.cutoff = 3, cols = c("lightblue", "navyblu
e"))
ggsave(file="/Users/lees44/R_project/BTBR/Oligo_clusters.pdf",width=10,height=10)
plot(F)
```

```
#oligo cluster (Mog = Oligodendrocyte)
F<-FeaturePlot(object =dCA2_FC_integrated.combined, features =  c("Mog"), blend.thres
hold = 0.5, min.cutoff = 0, max.cutoff = 3, cols = c("lightblue", "navyblue"))
ggsave(file="/Users/lees44/R_project/BTBR/Mog_clusters.pdf",width=10,height=10)
plot(F)
```

```
#oligo cluster (Opalin = Oligodendrocyte)
F<-FeaturePlot(object =dCA2_FC_integrated.combined, features =  c("Opalin"), blend.th
reshold = 0.5, min.cutoff = 0, max.cutoff = 3, cols = c("lightblue", "navyblue"))
ggsave(file="/Users/lees44/R_project/BTBR/Opalin_clusters.pdf",width=10,height=10)
plot(F)
```

```
#OPC (OPC=pdgfra, gpr17)
F<-FeaturePlot(object =dCA2_FC_integrated.combined, features =  c("Pdgfra","Gpr17"),
blend.threshold = 0.5, min.cutoff = 0, max.cutoff = 3, cols = c("lightblue", "navyblu
e"))
ggsave(file="/Users/lees44/R_project/BTBR/OPC.pdf",width=10,height=10)
plot(F)
```

```
#OPC (OPC=pdgfra)
F<-FeaturePlot(object =dCA2_FC_integrated.combined, features =  c("Pdgfra"), blend.th
reshold = 0.5, min.cutoff = 0, max.cutoff = 3, cols = c("lightblue", "navyblue"))
ggsave(file="/Users/lees44/R_project/BTBR/Pdgfra_Cluster.pdf",width=10,height=10)
plot(F)
```

```
#OPC (OPC= gpr17)
F<-FeaturePlot(object =dCA2_FC_integrated.combined, features =  c("Gpr17"), blend.thr
eshold = 0.5, min.cutoff = 0, max.cutoff = 3, cols = c("lightblue", "navyblue"))
ggsave(file="/Users/lees44/R_project/BTBR/Gpr17_cluster.pdf",width=10,height=10)
plot(F)
```

```
#mural (mural=Tagln, Vtn)
F<-FeaturePlot(object =dCA2_FC_integrated.combined, features =  c("Tagln","Vtn"), ble
nd.threshold = 0.5, min.cutoff = 0, max.cutoff = 3, cols = c("lightblue", "navyblu
e"))
ggsave(file="/Users/lees44/R_project/BTBR/Mural.pdf",width=10,height=10)
plot(F)
```

```
#mural (mural=Tagln)
F<-FeaturePlot(object =dCA2_FC_integrated.combined, features =  c("Tagln"), blend.thr
eshold = 0.5, min.cutoff = 0, max.cutoff = 3, cols = c("lightblue", "navyblue"))
ggsave(file="/Users/lees44/R_project/BTBR/Tagln_cluster.pdf",width=10,height=10)
plot(F)
```

```
#mural (mural=Vtn)
F<-FeaturePlot(object =dCA2_FC_integrated.combined, features =  c("Vtn"), blend.thres
hold = 0.5, min.cutoff = 0, max.cutoff = 3, cols = c("lightblue", "navyblue"))
ggsave(file="/Users/lees44/R_project/BTBR/Vtn_cluster.pdf",width=10,height=10)
plot(F)
```

```
##pericytes (Pericyte=Cldn5, Flt1)
F<-FeaturePlot(object =dCA2_FC_integrated.combined, features =  c("Cldn5","Flt1"), bl
end.threshold = 0.5, min.cutoff = 0, max.cutoff = 3, cols = c("lightblue", "navyblu
e"))
ggsave(file="/Users/lees44/R_project/BTBR/Pericyte.pdf",width=10,height=10)
plot(F)
```

```
##pericytes (Pericyte=Cldn5)
F<-FeaturePlot(object =dCA2_FC_integrated.combined, features =  c("Cldn5"), blend.thr
eshold = 0.5, min.cutoff = 0, max.cutoff = 3, cols = c("lightblue", "navyblue"))
ggsave(file="/Users/lees44/R_project/BTBR/Cldn5_cluster.pdf",width=10,height=10)
plot(F)
```

```
##pericytes (Pericyte= Flt1)
F<-FeaturePlot(object =dCA2_FC_integrated.combined, features =  c("Flt1"), blend.thre
shold = 0.5, min.cutoff = 0, max.cutoff = 3, cols = c("lightblue", "navyblue"))
ggsave(file="/Users/lees44/R_project/BTBR/Flt1_cluster.pdf",width=10,height=10)
plot(F)
```

```
##eppen (Foxj1, Fam216b)
F<-FeaturePlot(object =dCA2_FC_integrated.combined, features =  c("Foxj1","Fam216b"),
blend.threshold = 0.5, min.cutoff = 0, max.cutoff = 3, cols = c("lightblue", "navyblu
e"))
ggsave(file="/Users/lees44/R_project/BTBR/Eppen.pdf",width=10,height=10)
plot(F)
```

```
##eppen (Foxj1)
F<-FeaturePlot(object =dCA2_FC_integrated.combined, features =  c("Foxj1"), blend.threshold = 0.5, min.cutoff = 0, max.cutoff = 3, cols = c("lightblue", "navyblue"))
ggsave(file="/Users/lees44/R_project/BTBR/Foxj1_cluster.pdf",width=10,height=10)
plot(F)
```

```
##eppen (Fam216b)
F<-FeaturePlot(object =dCA2_FC_integrated.combined, features =  c("Fam216b"), blend.threshold = 0.5, min.cutoff = 0, max.cutoff = 3, cols = c("lightblue", "navyblue"))
ggsave(file="/Users/lees44/R_project/BTBR/Fam216b_cluster.pdf",width=10,height=10)
plot(F)
```

```
##Astrocyte (Slc7a10, Gfap)
F<-FeaturePlot(object =dCA2_FC_integrated.combined, features =  c("Slca7a10","Gfap"), blend.threshold = 0.5, min.cutoff = 0, max.cutoff = 3, cols = c("lightblue", "navyblue"))
ggsave(file="/Users/lees44/R_project/BTBR/Astrocyte.pdf",width=10,height=10)
plot(F)
```

```
##Astrocyte (Slc7a10)
F<-FeaturePlot(object =dCA2_FC_integrated.combined, features =  c("Slc7a10"), blend.threshold = 0.5, min.cutoff = 0, max.cutoff = 3, cols = c("lightblue", "navyblue"))
ggsave(file="/Users/lees44/R_project/BTBR/Slc7a10_cluster.pdf",width=10,height=10)
plot(F)
```

```
##Astrocyte (Slc7a10, Gfap)
F<-FeaturePlot(object =dCA2_FC_integrated.combined, features =  c("Gfap"), blend.threshold = 0.5, min.cutoff = 0, max.cutoff = 3, cols = c("lightblue", "navyblue"))
ggsave(file="/Users/lees44/R_project/BTBR/Gfap_clusters.pdf",width=10,height=10)
plot(F)
```

```
#Neuronal
F<-FeaturePlot(object =dCA2_FC_integrated.combined, features =  c("Stmn2","Thy1","Camk2a", "Kif5c"), blend.threshold = 0.5, min.cutoff = 0, max.cutoff = 3, cols = c("lightblue", "navyblue"))
ggsave(file="/Users/lees44/R_project/BTBR/Neuronal.pdf",width=10,height=10)
plot(F)
```

```
#Neuronal
F<-FeaturePlot(object =dCA2_FC_integrated.combined, features =  c("Stmn2"), blend.thr
eshold = 0.5, min.cutoff = 0, max.cutoff = 3, cols = c("lightblue", "navyblue"))
ggsave(file="/Users/lees44/R_project/BTBR/Stmn2_cluster.pdf",width=10,height=10)
plot(F)
```

```
#Neuronal
F<-FeaturePlot(object =dCA2_FC_integrated.combined, features =  c("Thy1"), blend.thre
shold = 0.5, min.cutoff = 0, max.cutoff = 3, cols = c("lightblue", "navyblue"))
ggsave(file="/Users/lees44/R_project/BTBR/Thy1_cluster.pdf",width=10,height=10)
plot(F)
```

```
#Neuronal
F<-FeaturePlot(object =dCA2_FC_integrated.combined, features =  c("Camk2a"), blend.th
reshold = 0.5, min.cutoff = 0, max.cutoff = 3, cols = c("lightblue", "navyblue"))
ggsave(file="/Users/lees44/R_project/BTBR/Camk2a_cluster.pdf",width=10,height=10)
plot(F)
```

```
#Neuronal
F<-FeaturePlot(object =dCA2_FC_integrated.combined, features =  c("Kif5c"), blend.thr
eshold = 0.5, min.cutoff = 0, max.cutoff = 3, cols = c("lightblue", "navyblue"))
ggsave(file="/Users/lees44/R_project/BTBR/kif5c_cluster.pdf",width=10,height=10)
plot(F)
```

```
#CA1(Wfs1 and Trpc4)
F<-FeaturePlot(object =dCA2_FC_integrated.combined, features =  c("Wfs1", "Trpc4"), b
lend.threshold = 0.5, min.cutoff = 0, max.cutoff = 3, cols = c("lightblue", "navyblu
e"))
ggsave(file="/Users/lees44/R_project/BTBR/CA1.pdf",width=10,height=10)
plot(F)
```

```
#CA1(Wfs1)
F<-FeaturePlot(object =dCA2_FC_integrated.combined, features =  c("Wfs1"), blend.thre
shold = 0.5, min.cutoff = 0, max.cutoff = 3, cols = c("lightblue", "navyblue"))
ggsave(file="/Users/lees44/R_project/BTBR/Wfs1_cluster.pdf",width=10,height=10)
plot(F)
```

```
#CA1(Trpc4)
F<-FeaturePlot(object =dCA2_FC_integrated.combined, features =  c("Trpc4"), blend.thr
eshold = 0.5, min.cutoff = 0, max.cutoff = 3, cols = c("lightblue", "navyblue"))
ggsave(file="/Users/lees44/R_project/BTBR/Trpc4_cluster.pdf",width=10,height=10)
plot(F)
```

```
#CA3(Bok3 Iyd)
F<-FeaturePlot(object =dCA2_FC_integrated.combined, features =  c("Bok", "Iyd"), blen
d.threshold = 0.5, min.cutoff = 0, max.cutoff = 3, cols = c("lightblue", "navyblue"))
ggsave(file="/Users/lees44/R_project/BTBR/CA3.pdf",width=10,height=10)
plot(F)
```

```
#CA3(Bok3)
F<-FeaturePlot(object =dCA2_FC_integrated.combined, features =  c("Bok"), blend.thres
hold = 0.5, min.cutoff = 0, max.cutoff = 3, cols = c("lightblue", "navyblue"))
ggsave(file="/Users/lees44/R_project/BTBR/Bok_cluster.pdf",width=10,height=10)
plot(F)
```

```
#CA3(Iyd)
F<-FeaturePlot(object =dCA2_FC_integrated.combined, features =  c("Iyd"), blend.thres
hold = 0.5, min.cutoff = 0, max.cutoff = 3, cols = c("lightblue", "navyblue"))
ggsave(file="/Users/lees44/R_project/BTBR/Iyd_cluster.pdf",width=10,height=10)
plot(F)
```

```
#Granule cells (Prox1)
F<-FeaturePlot(object =dCA2_FC_integrated.combined, features =  c("Prox1"), blend.thr
eshold = 0.5, min.cutoff = 0, max.cutoff = 3, cols = c("lightblue", "navyblue"))
ggsave(file="/Users/lees44/R_project/BTBR/Granule_cells.pdf",width=10,height=10)
plot(F)
```

```
#Non-granule cells (Dkk3)
F<-FeaturePlot(object =dCA2_FC_integrated.combined, features =  c("Dkk3"), blend.thre
shold = 0.5, min.cutoff = 0, max.cutoff = 3, cols = c("lightblue", "navyblue"))
ggsave(file="/Users/lees44/R_project/BTBR/Non_granule.pdf",width=10,height=10)
plot(F)
```

```
#Mossy cells (Calb2)
F<-FeaturePlot(object =dCA2_FC_integrated.combined, features =  c("Calb2"), blend.thr
eshold = 0.5, min.cutoff = 0, max.cutoff = 3, cols = c("lightblue", "navyblue"))
ggsave(file="/Users/lees44/R_project/BTBR/Mossy_cells.pdf",width=10,height=10)
plot(F)
```

```
#All_pyramids
F<-FeaturePlot(object =dCA2_FC_integrated.combined, features =  c("Ociad2"), blend.th
reshold = 0.5, min.cutoff = 0, max.cutoff = 3, cols = c("lightblue", "navyblue"))
ggsave(file="/Users/lees44/R_project/BTBR/All_pyramids.pdf",width=10,height=10)
plot(F)
```

```
#CA2_pyramids
F<-FeaturePlot(object =dCA2_FC_integrated.combined, features =  c("Cacng5"), blend.th
reshold = 0.5, min.cutoff = 0, max.cutoff = 3, cols = c("lightblue", "navyblue"))
ggsave(file="/Users/lees44/R_project/BTBR/CA2_pyramids.pdf",width=10,height=10)
plot(F)
```

```
#CA1_pyramids
F<-FeaturePlot(object =dCA2_FC_integrated.combined, features =  c("Fibcd1"), blend.th
reshold = 0.5, min.cutoff = 0, max.cutoff = 3, cols = c("lightblue", "navyblue"))
ggsave(file="/Users/lees44/R_project/BTBR/CA1_pyramids.pdf",width=10,height=10)
plot(F)
```

######02022023####################### Subset for CA2 neurons only (cluster2)

```
CA2_subset<-subset(x= dCA2_FC_integrated.combined, idents ="CA2")
```

```
p2<-DimPlot(CA2_subset, reduction = "umap", group.by = "protocol")
plot_grid(p2)
ggsave(file="/Users/lees44/R_project/BTBR/CA2subset_UMAP.pdf",width=10,height=10)
```

```
set.seed(1234)
```

```
CA2_subset_combined <- RunPCA(CA2_subset, npcs = 30, verbose = FALSE)
```

```
CA2_subset_combined <- RunUMAP(CA2_subset, reduction = "pca", dims = 1:30)
```

```
p2<-DimPlot(CA2_subset_combined, reduction = "umap", group.by = "protocol")
plot_grid(p2)
ggsave(file="/Users/lees44/R_project/BTBR/CA2subset_PCA_UMAP.pdf",width=10,height=10)
```

```
p1<-DimPlot(CA2_subset_combined, reduction = "umap", split.by = "protocol")
plot_grid(p1)
ggsave(file="/Users/lees44/R_project/BTBR/CA2subset_splited_group.pdf",width=10,heigh
t=10)
```

###################################BTBR vs WT gene expression comparison. First top markers then violin plot##############

```
DefaultAssay(CA2_subset_combined) <- "RNA"
```

Generalized BTBR vs WT

Differential expression testing: see https://satijalab.org/seurat/archive/v3.1/de_vignette.html (https://satijalab.org/seurat/archive/v3.1/de_vignette.html) https://satijalab.org/seurat/reference/findmarkers (https://satijalab.org/seurat/reference/findmarkers)

```
CA2_subset_combined@meta.data$Group = "no"
CA2_subset_combined@meta.data$Group[CA2_subset_combined@meta.data$protocol == "BTBR_
M" | CA2_subset_combined@meta.data$protocol == "BTBR_F"] = "BTBR"
CA2_subset_combined@meta.data$Group[CA2_subset_combined@meta.data$protocol == "WT_M"
| CA2_subset_combined@meta.data$protocol == "WT_F"] = "WT"

table(CA2_subset_combined@meta.data$Group)

Idents(object = CA2_subset_combined) <- "Group"

BTBRvsWT.markers <- FindMarkers(CA2_subset_combined, ident.1 = "BTBR" , ident.2 = "W
T", min.pct = 0.25)
head(BTBRvsWT.markers, n = 10)
write.csv(BTBRvsWT.markers, "/Users/lees44/R_project/BTBR/BTBRvsWTtop_10markers_integ
rated.csv")
```

```
CA2_subset_combined_only <- subset(x = CA2_subset_combined, idents = c("BTBR", "WT"))
```

```
p4<-DimPlot(CA2_subset_combined_only, reduction = "umap", group.by = "Group",  pt=1)
plot_grid(p4)
ggsave(file="/Users/lees44/R_project/BTBR/BTBRvsWT_Group_split.pdf",width=10,height=1
0)
```

Violin plot for genes in BTBR vs WT see https://ggplot2.tidyverse.org/reference/theme.html (https://ggplot2.tidyverse.org/reference/theme.html) changing graph using ggplot theme

```
VlnPlot(object = CA2_subset_combined, features = "Galnt1", pt.size = 0.5) +theme(titl
e = element_text(size = 30), axis.title.x = element_text(size =20), axis.title.y = el
ement_text(size = 20), axis.text.x = element_text(size = 20), legend.text = element_t
ext(size =20))
ggsave(file="/Users/lees44/R_project/BTBR/Galnt1_BTBRvsWT.pdf", width=7,height=10)
```

```
VlnPlot(object = CA2_subset_combined_only, features = "Galnt1", pt.size = 0.5) +them
e(title = element_text(size = 30), axis.title.x = element_text(size =20), axis.title.
y = element_text(size = 20), axis.text.x = element_text(size = 20), legend.text = ele
ment_text(size =20))
ggsave(file="/Users/lees44/R_project/BTBR/Galnt1_BTBRvsWT_only.pdf", width=7,height=1
0)
```

```
VlnPlot(object = CA2_subset_combined, features = "Cfap54", pt.size = 0.5) +theme(titl
e = element_text(size = 30), axis.title.x = element_text(size =20), axis.title.y = el
ement_text(size = 20), axis.text.x = element_text(size = 20), legend.text = element_t
ext(size =20))
ggsave(file="/Users/lees44/R_project/BTBR/Cfap54_BTBRvsWT.pdf", width=7,height=10)
```

```
VlnPlot(object = CA2_subset_combined_only, features = "Cfap54", pt.size = 0.5) +them
e(title = element_text(size = 30), axis.title.x = element_text(size =20), axis.title.
y = element_text(size = 20), axis.text.x = element_text(size = 20), legend.text = ele
ment_text(size =20))
ggsave(file="/Users/lees44/R_project/BTBR/Cfap54_BTBRvsWT_only.pdf", width=7,height=1
0)
```

```
VlnPlot(object = CA2_subset_combined, features = "Epb41l5", pt.size = 0.5) +theme(tit
le = element_text(size = 30), axis.title.x = element_text(size =20), axis.title.y = e
lement_text(size = 20), axis.text.x = element_text(size = 20), legend.text = element_
text(size =20))
ggsave(file="/Users/lees44/R_project/BTBR/Epb41l5_BTBRvsWT.pdf", width=7,height=10)
```

```
VlnPlot(object = CA2_subset_combined_only, features = "Epb41l5", pt.size = 0.5) +them
e(title = element_text(size = 30), axis.title.x = element_text(size =20), axis.title.
y = element_text(size = 20), axis.text.x = element_text(size = 20), legend.text = ele
ment_text(size =20))
ggsave(file="/Users/lees44/R_project/BTBR/Epb41l5_BTBRvsWT_only.pdf", width=7,height=
10)
```

```
VlnPlot(object = CA2_subset_combined, features = "Snhg11", pt.size = 0.5) +theme(titl
e = element_text(size = 30), axis.title.x = element_text(size =20), axis.title.y = el
ement_text(size = 20), axis.text.x = element_text(size = 20), legend.text = element_t
ext(size =20))
ggsave(file="/Users/lees44/R_project/BTBR/Snhg11_BTBRvsWT.pdf", width=7,height=10)
```

```
VlnPlot(object = CA2_subset_combined_only, features = "Snhg11", pt.size = 0.5) +them
e(title = element_text(size = 30), axis.title.x = element_text(size =20), axis.title.
y = element_text(size = 20), axis.text.x = element_text(size = 20), legend.text = ele
ment_text(size =20))
ggsave(file="/Users/lees44/R_project/BTBR/Snhg11_BTBRvsWT_only.pdf", width=7,height=1
0)
```

```
VlnPlot(object = CA2_subset_combined, features = "Dchs2", pt.size = 0.5) +theme(title
= element_text(size = 30), axis.title.x = element_text(size =20), axis.title.y = elem
ent_text(size = 20), axis.text.x = element_text(size = 20), legend.text = element_tex
t(size =20))
ggsave(file="/Users/lees44/R_project/BTBR/Dchs2_BTBRvsWT.pdf", width=7,height=10)
```

```
VlnPlot(object = CA2_subset_combined_only, features = "Dchs2", pt.size = 0.5) +theme(
title = element_text(size = 30), axis.title.x = element_text(size =20), axis.title.y
= element_text(size = 20), axis.text.x = element_text(size = 20), legend.text = eleme
nt_text(size =20))
ggsave(file="/Users/lees44/R_project/BTBR/Dchs2_BTBRvsWT_only.pdf", width=7,height=1
0)
```

```
VlnPlot(object = CA2_subset_combined, features = "Sav1", pt.size = 0.5) +theme(title
= element_text(size = 30), axis.title.x = element_text(size =20), axis.title.y = elem
ent_text(size = 20), axis.text.x = element_text(size = 20), legend.text = element_tex
t(size =20))
ggsave(file="/Users/lees44/R_project/BTBR/Sav1_BTBRvsWT.pdf", width=7,height=10)
```

```
VlnPlot(object = CA2_subset_combined, features = "Kcnip4", pt.size = 0.5) +theme(titl
e = element_text(size = 30), axis.title.x = element_text(size =20), axis.title.y = el
ement_text(size = 20), axis.text.x = element_text(size = 20), legend.text = element_t
ext(size =20))
ggsave(file="/Users/lees44/R_project/BTBR/Kcnip4_BTBRvsWT.pdf", width=7,height=10)
```

```
VlnPlot(object = CA2_subset_combined, features = "Gm8797", pt.size = 0.5) +theme(titl
e = element_text(size = 30), axis.title.x = element_text(size =20), axis.title.y = el
ement_text(size = 20), axis.text.x = element_text(size = 20), legend.text = element_t
ext(size =20))
ggsave(file="/Users/lees44/R_project/BTBR/Gm8797_BTBRvsWT.pdf", width=7,height=10)
```

```
VlnPlot(object = CA2_subset_combined_only, features = "Gm8797", pt.size = 0.5) +them
e(title = element_text(size = 30), axis.title.x = element_text(size =20), axis.title.
y = element_text(size = 20), axis.text.x = element_text(size = 20), legend.text = ele
ment_text(size =20))
ggsave(file="/Users/lees44/R_project/BTBR/Gm8797_BTBRvsWT_only.pdf", width=7,height=1
0)
```

```
VlnPlot(object = CA2_subset_combined, features = "Heg1", pt.size = 0.5) +theme(title
= element_text(size = 30), axis.title.x = element_text(size =20), axis.title.y = elem
ent_text(size = 20), axis.text.x = element_text(size = 20), legend.text = element_tex
t(size =20))
ggsave(file="/Users/lees44/R_project/BTBR/Heg1_BTBRvsWT.pdf", width=7,height=10)
```

```
VlnPlot(object = CA2_subset_combined_only, features = "Heg1", pt.size = 0.5) +theme(t
itle = element_text(size = 30), axis.title.x = element_text(size =20), axis.title.y =
element_text(size = 20), axis.text.x = element_text(size = 20), legend.text = element
_text(size =20))
ggsave(file="/Users/lees44/R_project/BTBR/Heg1_BTBRvsWT.pdf", width=7,height=10)
```

```
VlnPlot(object = CA2_subset_combined, features = "Amph", pt.size = 0.5) +theme(title
= element_text(size = 30), axis.title.x = element_text(size =20), axis.title.y = elem
ent_text(size = 20), axis.text.x = element_text(size = 20), legend.text = element_tex
t(size =20))
ggsave(file="/Users/lees44/R_project/BTBR/Amph_BTBRvsWT.pdf", width=7,height=10)
```

```
VlnPlot(object = CA2_subset_combined, features = "Lpl", pt.size = 0.5) +theme(title =
element_text(size = 30), axis.title.x = element_text(size =20), axis.title.y = elemen
t_text(size = 20), axis.text.x = element_text(size = 20), legend.text = element_text(
size =20))
ggsave(file="/Users/lees44/R_project/BTBR/Lpl_BTBRvsWT.pdf", width=7,height=10)
```

```
VlnPlot(object = CA2_subset_combined_only, features = "Lpl", pt.size = 0.5) +theme(ti
tle = element_text(size = 30), axis.title.x = element_text(size =20), axis.title.y =
element_text(size = 20), axis.text.x = element_text(size = 20), legend.text = element
_text(size =20))
ggsave(file="/Users/lees44/R_project/BTBR/Lpl_BTBRvsWT_only.pdf", width=7,height=10)
```

cell number for BTBR vs WT CA2 subset cluster

```
dim(CA2_subset_combined_only)
```

```
BTBR_subset<-subset(CA2_subset_combined_only, idents = "BTBR")
```

```
dim(BTBR_subset)
```

```
WT_subset<-subset(CA2_subset_combined_only, idents = "WT")
```

```
dim(WT_subset)
```

########the end ##########
