## Supplementary material for "Investigation of the Fasciola Cinereum, Absent in BTBR mice, and Comparison with the Hippocampal Area CA2": 7_Characterization of fasciola cinereum, all group Integration

**Su Hyun Lee**

This is an R Markdown (http://rmarkdown.rstudio.com) Notebook. When you execute code within the notebook, the results appear beneath the code.

Try executing this chunk by clicking the *Run* button within the chunk or by placing your cursor inside it and pressing *Cmd+Shift+Enter*.

```
knitr::opts_chunk$set(echo = TRUE, eval=FALSE)
```

```
library(Seurat)
library(dplyr)
library(magrittr)
library("ggplot2")
library(grid)
library(reshape2)
library("scales")
```

nfeatures increaed from 2000 to 5000 in order to detect rarely expressed genes. default = 2000

```
dCA2_MI_GFPve_pos<- FindVariableFeatures(object =dCA2_MI_GFPve_pos,selection.method =
"vst", nfeatures = 5000, verbose = FALSE)
dCA2_FI_GFPve_pos<- FindVariableFeatures(object =dCA2_FI_GFPve_pos,selection.method =
"vst", nfeatures = 5000, verbose = FALSE)
dCA2_MI_GFPve_neg<- FindVariableFeatures(object =dCA2_MI_GFPve_neg,selection.method =
"vst", nfeatures = 5000, verbose = FALSE)
FC_MI_GFPve_pos<- FindVariableFeatures(object =FC_MI_GFPve_pos,selection.method = "vs
t", nfeatures = 5000, verbose = FALSE)
FC_FI_GFPve_pos<- FindVariableFeatures(object =FC_FI_GFPve_pos,selection.method = "vs
t", nfeatures = 5000, verbose = FALSE)
FC_MI_GFPve_neg<- FindVariableFeatures(object =FC_MI_GFPve_neg,selection.method = "vs
t", nfeatures = 5000, verbose = FALSE)
```

################integration 050321##########################

```
#integration of multiple groups using vector (x, y=c(1,2) and split by group.
dCA2.combined = merge(dCA2_MI_GFPve_pos, y = c(dCA2_FI_GFPve_pos,  dCA2_MI_GFPve_neg,
FC_MI_GFPve_pos, FC_FI_GFPve_pos, FC_MI_GFPve_neg), add.cell.ids = c( "dCA2_Mpos", "d
CA2_Fpos", "dCA2_Mneg", "FC_Mpos", "FC_Fpos", "FC_Mneg"), project = "protocol")
```

```
dCA2.list <-SplitObject(dCA2.combined, split.by = "protocol")
reference.list <-dCA2.list[c("dCA2_Mpos", "dCA2_Fpos", "dCA2_Mneg", "FC_Mpos", "FC_Fp
os", "FC_Mneg")]
dCA2.anchors <- FindIntegrationAnchors(object.list = reference.list, dims = 1:30)
```

```
#intgrated analysis
dCA2_FC_integrated.combined <-IntegrateData(anchorset = dCA2.anchors, dims = 1:30)
```

```
library(ggplot2)
library(cowplot)
DefaultAssay(dCA2_FC_integrated.combined) <- "integrated"
```

```
set.seed(1234)
```

```
dCA2_FC_integrated.combined <- ScaleData(dCA2_FC_integrated.combined, verbose = FALS
E)
```

```
dCA2_FC_integrated.combined <- RunPCA(dCA2_FC_integrated.combined, npcs = 50, verbose
= FALSE)
```

Elbow plot to determine pc value

```
mat <- GetAssayData(dCA2_FC_integrated.combined, slot = "scale.data")
pca <- dCA2_FC_integrated.combined[["pca"]]
total_variance <- sum(matrixStats::rowVars(mat))
eigValues = (pca@stdev)^2  ## EigenValues
varExplained = (eigValues / total_variance)*100
plot(varExplained,type='l')
```

```
dCA2_FC_integrated.combined <- RunPCA(dCA2_FC_integrated.combined, npcs = 30, verbose
= FALSE)
```

```
dCA2_FC_integrated.combined <- RunUMAP(dCA2_FC_integrated.combined, reduction = "pc
a", dims = 1:30)
```

```
set.seed(1234)
```

```
p1<-DimPlot(dCA2_FC_integrated.combined, reduction = "umap", group.by = "protocol")
plot_grid(p1)
ggsave(file="/Users/lees44/R_project/FC_dCA2_all_vector_Integration_SUN1GFP_MYC_05032
1/UMAP.pdf",width=10,height=10)
```

```
p1<-DimPlot(dCA2_FC_integrated.combined, reduction = "umap", split.by = "protocol")
plot_grid(p1)
ggsave(file="/Users/lees44/R_project/FC_dCA2_all_vector_Integration_SUN1GFP_MYC_05032
1/UMAP_split_group.pdf",width=10,height=10)
```

```
dCA2_FC_integrated.combined <- FindNeighbors(dCA2_FC_integrated.combined, reduction =
"pca", dims = 1:30)
```

```
dCA2_FC_integrated.combined <- FindClusters(dCA2_FC_integrated.combined, resolution =
0.3)
```

```
#clustering by population
p1<-DimPlot(object = dCA2_FC_integrated.combined, reduction = "umap", group.by = "int
egrated_snn_res.0.3", label = TRUE, repel = TRUE)
plot_grid(p1)
ggsave(file="/Users/lees44/R_project/FC_dCA2_all_vector_Integration_SUN1GFP_MYC_05032
1/Clustering.pdf",width=10,height=10)
```

```
p1<-DimPlot(dCA2_FC_integrated.combined, reduction = "umap", split.by = "protocol")
plot_grid(p1)
ggsave(file="/Users/lees44/R_project/FC_dCA2_all_vector_Integration_SUN1GFP_MYC_05032
1/Clustering_splited_group.pdf",width=10,height=10)
```

###################defining cell types ##################################################

Feature plot is used and cell type was classified in supervised manner.

```
# microglia and oligo clusters, (Tmem119, C1qc = microglia)
F<-FeaturePlot(object =dCA2_FC_integrated.combined, features =  c("Tmem119","C1qc"))
ggsave(file="/Users/lees44/R_project/FC_dCA2_all_vector_Integration_SUN1GFP_MYC_05032
1/Microglia_clusters.pdf",width=10,height=10)
plot(F)
```

```
#oligo cluster (Mog, Opalin = Oligodendrocyte)
F<-FeaturePlot(object =dCA2_FC_integrated.combined, features =  c("Mog","Opalin"))
ggsave(file="/Users/lees44/R_project/FC_dCA2_all_vector_Integration_SUN1GFP_MYC_05032
1/Oligodendrocyte_clusters.pdf",width=10,height=10)
plot(F)
```

```
#OPC (OPC=pdgfra, gpr17)
F<-FeaturePlot(object =dCA2_FC_integrated.combined, features = c("Pdgfra","Gpr17"))
ggsave(file="/Users/lees44/R_project/FC_dCA2_all_vector_Integration_SUN1GFP_MYC_05032
1/OPC.pdf",width=10,height=10)
plot(F)
```

```
#mural (mural=Tagln, Vtn)
F<-FeaturePlot(object =dCA2_FC_integrated.combined, features = c("Tagln", "Vtn"))
ggsave(file="/Users/lees44/R_project/FC_dCA2_all_vector_Integration_SUN1GFP_MYC_05032
1/Mural.pdf",width=10,height=10)
plot(F)
```

```
#pericytes (Pericyte=Cldn5, Flt1)
F<-FeaturePlot(object =dCA2_FC_integrated.combined, features = c("Cldn5","Flt1"))
ggsave(file="/Users/lees44/R_project/FC_dCA2_all_vector_Integration_SUN1GFP_MYC_05032
1/Pericyte.pdf",width=10,height=10)
plot(F)
```

```
#eppen (Foxj1, Fam216b)
F<-FeaturePlot(object =dCA2_FC_integrated.combined, features = c("Foxj1","Fam216b"))
ggsave(file="/Users/lees44/R_project/FC_dCA2_all_vector_Integration_SUN1GFP_MYC_05032
1/Eppen.pdf",width=10,height=10)
plot(F)
```

```
#  Astrocyte
F<-FeaturePlot(object =dCA2_FC_integrated.combined, features = c("Slc7a10", "Gfap"))
ggsave(file="/Users/lees44/R_project/FC_dCA2_all_vector_Integration_SUN1GFP_MYC_05032
1/Astrocyte.pdf",width=10,height=10)
plot(F)
```

```
#  Neuronal
F<-FeaturePlot(object =dCA2_FC_integrated.combined, features = c("Stmn2","Thy1","Camk
2a", "Kif5c"))
ggsave(file="/Users/lees44/R_project/FC_dCA2_all_vector_Integration_SUN1GFP_MYC_05032
1/Neuron.pdf",width=10,height=10)
plot(F)
```

#####################doublet code##################################

```
install.packages("scDblfinder")
```

converting seurat object to sce object https://satijalab.org/seurat/archive/v3.0/conversion_vignette.html (https://satijalab.org/seurat/archive/v3.0/conversion_vignette.html)

Doublet removal

```
BiocManager::install("scDblFinder")
library(scDblFinder)
library(scater)
library(SingleCellExperiment)
library(Seurat)
library(scDblFinder)
library(scuttle)
```

Doublet detection with clusters

The doubletCluster() function will identify clusters that have intermediate expression profiles of two other clusters (Bach et al. 2017). Specifically, it will examine every possible triplet of clusters consisting of a query cluster and its two “parents”. It will then compute a number of statistics:

The number of genes (N) that are differentially expressed in the same direction in the query cluster compared to both of the parent clusters. Such genes would be unique markers for the query cluster and provide evidence against the null hypothesis, i.e., that the query cluster consists of doublets from the two parents. Clusters with few unique genes are more likely to be doublets. The ratio of the median library size in each parent to the median library size in the query (lib.size*). Doublet libraries are generated from a larger initial pool of RNA compared to libraries for single cells, and thus the former should have larger library sizes. Library size ratios much greater than unity are inconsistent with a doublet identity for the query. The proportion of cells in the query cluster should also be reasonable - typically less than 5% of all cells, depending on how many cells were loaded onto the 10X Genomics device.

```
# prepare object to inspect for doublet clusters
DefaultAssay(dCA2.integrated) <- "RNA"
sce.object <- as.SingleCellExperiment(dCA2.integrated)
colLabels(sce.object) <- colData(sce.object)$ident
dbl.out <- findDoubletClusters(sce.object,colLabels(sce.object))
```

```
# outlier clusters by number of significant genes (num.de)
rownames(dbl.out)[isOutlier(dbl.out$num.de, log=TRUE, type="lower")]
```

```
# outlier clusters by number of significant genes (num.de)
rownames(dbl.out)[dbl.out$lib.size1 < 1 & dbl.out$lib.size2 < 1]
```

```
# be sure to reset your defualt assay back
DefaultAssay(dCA2.integrated) <- "integrated"
```

########################################dCA2 markers##################################################################

Feature plot highlighting dCA2 markers in all population Expression pots in UMAP space illustrating normalized expression values of dCA2 markers

“CREREV38354884”, “GFP”,“Rgs14”, “Avpr1b”, “Amigo2”, “Map3k15”, “Actn2”, “Ptpn5”, “Cacng5”, “Oxtr”, “Pcp4”

```
install.packages("RColorBrewer")
```

```
library(RColorBrewer)
```

changing color palettes in ggplot F<-FeaturePlot(object =dCA2_FC_integrated.combined, features = c(“Rgs14”)) +scale_colour_gradient(colours = rev(brewer.pal(n = 4, name =“RdBu”))) https://github.com/satijalab/seurat/issues/2400 (https://github.com/satijalab/seurat/issues/2400)

```
#Feature plot highlighting dCA2 markers in all population
F<-FeaturePlot(object =dCA2_FC_integrated.combined, features = c("CREREV38354884"), b
lend.threshold = 0.5, min.cutoff = 0, max.cutoff = 1.5, cols = c("lightblue", "navybl
ue"))
plot(F)
ggsave(file="/Users/lees44/R_project/FC_dCA2_all_vector_Integration_SUN1GFP_MYC_05032
1/CREREV38354884.pdf",width=10,height=10)
```

```
#Feature plot highlighting dCA2 markers in all population
F<-FeaturePlot(object =dCA2_FC_integrated.combined, features = c("GFP"), blend.thresh
old = 0.5, min.cutoff = 0, max.cutoff = 3, cols = c("lightblue", "navyblue"))
plot(F)
ggsave(file="/Users/lees44/R_project/FC_dCA2_all_vector_Integration_SUN1GFP_MYC_05032
1/GFP.pdf",width=10,height=10)
```

```
#Feature plot highlighting dCA2 markers in all population
F<-FeaturePlot(object =dCA2_FC_integrated.combined, features = c("Rgs14"), blend.thre
shold = 0.5, min.cutoff = 0, max.cutoff = 3, cols = c("lightblue", "navyblue"))
plot(F)
ggsave(file="/Users/lees44/R_project/FC_dCA2_all_vector_Integration_SUN1GFP_MYC_05032
1/Rgs14.pdf",width=10,height=10)
```

```
#Feature plot highlighting dCA2 markers in all population
F<-FeaturePlot(object =dCA2_FC_integrated.combined, features = c("Avpr1b"), blend.thr
eshold = 0.5, min.cutoff = 0, max.cutoff = 1.5, cols = c("lightblue", "navyblue"))
plot(F)
ggsave(file="/Users/lees44/R_project/FC_dCA2_all_vector_Integration_SUN1GFP_MYC_05032
1/Avpr1b.pdf",width=10,height=10)
```

```
#Feature plot highlighting dCA2 markers in all population
F<-FeaturePlot(object =dCA2_FC_integrated.combined, features = c("Amigo2"), blend.thr
eshold = 0.2, min.cutoff = 0, max.cutoff = 3, cols = c("lightblue", "navyblue"))
plot(F)
ggsave(file="/Users/lees44/R_project/FC_dCA2_all_vector_Integration_SUN1GFP_MYC_05032
1/Amigo2.pdf",width=10,height=10)
```

“CREREV38354884”, “GFP”,“Rgs14”, “Avpr1b”, “Amigo2”, “Map3k15”, “Actn2”, “Ptpn5”, “Cacng5”, “Oxtr”, “Pcp4”

```
#Feature plot highlighting dCA2 markers in all population
F<-FeaturePlot(object =dCA2_FC_integrated.combined, features = c("Map3k15"), blend.th
reshold = 0.1, min.cutoff = 0, max.cutoff = 3, cols = c("lightblue", "navyblue"))
plot(F)
ggsave(file="/Users/lees44/R_project/FC_dCA2_all_vector_Integration_SUN1GFP_MYC_05032
1/Map3k15.pdf",width=10,height=10)
```

```
#Feature plot highlighting dCA2 markers in all population
F<-FeaturePlot(object =dCA2_FC_integrated.combined, features = c("Actn2"), blend.thre
shold = 0.1, min.cutoff = 0, max.cutoff = 2, cols = c("lightblue", "navyblue"))
plot(F)
ggsave(file="/Users/lees44/R_project/FC_dCA2_all_vector_Integration_SUN1GFP_MYC_05032
1/Actn2.pdf",width=10,height=10)
```

```
#Feature plot highlighting dCA2 markers in all population
F<-FeaturePlot(object =dCA2_FC_integrated.combined, features = c("Actn2"), blend.thre
shold = 0.2, min.cutoff = 0, max.cutoff = 2, cols = c("lightblue", "navyblue"))
plot(F)
ggsave(file="/Users/lees44/R_project/FC_dCA2_all_vector_Integration_SUN1GFP_MYC_05032
1/Actn2_blue.pdf",width=10,height=10)
```

```
#Feature plot highlighting dCA2 markers in all population
F<-FeaturePlot(object =dCA2_FC_integrated.combined, features = c("Ptpn5"), blend.thre
shold = 0.5, min.cutoff = 0, max.cutoff = 3, cols = c("lightblue", "navyblue"))
plot(F)
ggsave(file="/Users/lees44/R_project/FC_dCA2_all_vector_Integration_SUN1GFP_MYC_05032
1/Ptpn5.pdf",width=10,height=10)
```

```
#Feature plot highlighting dCA2 markers in all population
F<-FeaturePlot(object =dCA2_FC_integrated.combined, features = c("Cacng5"), blend.thr
eshold = 0.5, min.cutoff = 0, max.cutoff = 3, cols = c("lightblue", "navyblue"))
plot(F)
ggsave(file="/Users/lees44/R_project/FC_dCA2_all_vector_Integration_SUN1GFP_MYC_05032
1/Cacng5.pdf",width=10,height=10)
```

```
#Feature plot highlighting dCA2 markers in all population
F<-FeaturePlot(object =dCA2_FC_integrated.combined, features = c("Oxtr"), blend.thres
hold = 0.5, min.cutoff = 0, max.cutoff = 1.5, cols = c("lightblue", "navyblue"))
plot(F)
ggsave(file="/Users/lees44/R_project/FC_dCA2_all_vector_Integration_SUN1GFP_MYC_05032
1/Oxtr.pdf",width=10,height=10)
```

```
#Feature plot highlighting dCA2 markers in all population
F<-FeaturePlot(object =dCA2_FC_integrated.combined, features = c("Pcp4"), blend.thres
hold = 0.5, min.cutoff = 0, max.cutoff = 3, cols = c("lightblue", "navyblue"))
plot(F)
ggsave(file="/Users/lees44/R_project/FC_dCA2_all_vector_Integration_SUN1GFP_MYC_05032
1/Pcp4.pdf",width=10,height=10)
```

Assign clusters to cell types

```
#celltypes asigned based on that gene marker expression
new.ident <- c("Neuron1", "Neuron2", "Neuron3", "Neuron4", "Oligodendrocyte", "Astroc
yte", "Neuron5", "Neuron6", "NC4", "OPC", "Neuron7", "Microglia",  "NC1", "Neuron8",
"Neuron9", "NC2", "Neuron10", "Eppendymal", "NC3", "NC5")
names(x = new.ident) <- levels(x =dCA2_FC_integrated.combined)
dCA2_FC_integrated.combined<- RenameIdents(object =dCA2_FC_integrated.combined, new.i
dent)
```

```
color<-c("#19647e","#ffc857","#9B0C1E","#9B0C1E","#9B0C1E","#9B0C1E","#9B0C1E","#9B0C
1E","#4b3f72","#4b3f72","#ffc857","#9B0C1E","#676833","#9B0C1E","#b7b7b7","#E8C6C7","#
4b3f72","#ffc857","#19647e","#B99A69")
```

```
DimPlot(object = dCA2_FC_integrated.combined, reduction = "umap", label = FALSE, repel = TRUE,cols=color,label.size = 12,pt.size=0.5)
ggsave(file="/Users/lees44/R_project/FC_dCA2_all_vector_Integration_SUN1GFP_MYC_050321/Neuronaltype.pdf",width=10,height=7)
```

```
DimPlot(object = dCA2_FC_integrated.combined, reduction = "umap", label = TRUE, repel = TRUE)
ggsave(file="/Users/lees44/R_project/FC_dCA2_all_vector_Integration_SUN1GFP_MYC_050321/Neuronaltype.pdf",width=10,height=10)
```

################Work from here 07/13/21###########################

Cell type and violin plot based on cell type

```
#cluster tree to go with the violin plot based on cell type
pdf(file="/Users/lees44/R_project/FC_dCA2_all_vector_Integration_SUN1GFP_MYC_050321/cluster_tree.pdf",width=20,height=8,paper='special')
#https://www.rdocumentation.org/packages/ape/versions/5.2/topics/plot.phylo
dCA2.integrated <- BuildClusterTree(dCA2_FC_integrated.combined, verbose = FALSE, reorder = FALSE)
plot(dCA2.integrated@tools$BuildClusterTree, type = "phylogram", edge.color = "black", edge.width = 4,edge.lty = 1, srt = 0, label.offset = 10, direction = "downwards", tip.color = "black")
dev.off()
```

Violinplot for canonical markers, figure goes with cluster tree

```
Cell_type<-rev(c("Neuron1", "Neuron2", "Neuron3", "Neuron4", "Oligodendrocyte", "Astrocyte", "Neuron5", "Neuron6", "NC4", "OPC", "Neuron7", "Microglia",  "NC1", "Neuron8", "Neuron9", "NC2", "Neuron10", "Eppendymal", "NC3", "NC5"))
gene_list<-c("Mog" ,"Opalin","Camk2a", "Kif5c","Slc7a10", "Gfap", "Ccnd1","Pdgfra", "Gpr17", "Cldn5", "Tagln", "Flt1", "Vtn", "Tmem119", "C1qc", "Foxj1", "Fam216b")
```

```
Cell_number <- NULL
for (i in 1:length(Cell_type)){
                temp1 <- data.frame(type=factor(rep(Cell_type[i],sum(Cell_type[i]==Idents(dCA2_FC_integrated.combined)))))
                temp2 <- as.data.frame(dCA2_FC_integrated.combined@assays$RNA@data[gene_list,Idents(dCA2_FC_integrated.combined)==Cell_type[i]])
                temp3 <- cbind(temp1,t(temp2))
                Cell_number <- rbind(temp3,Cell_number)
}
rownames(Cell_number) <- NULL
```

```
colors<-c("#ffc857","#ffc857","#ffc857","#9B0C1E","#9B0C1E","#9B0C1E","#9B0C1E","#9B0C1E","#9B0C1E","#9B0C1E","#9B0C1E","#676833","#B99A69","#19647e","#19647e","#4b3f72","#4b3f72","#4b3f72","#b7b7b7","#E8C6C7")
```

```
for (k in 1:length(gene_list))
    {
if (k==length(gene_list)){
assign(paste("P",k,sep=""),ggplot(Cell_number,aes_string(x="type",y=gene_list[k],fill="type"))+geom_violin(scale = "width")+scale_fill_manual(values=colors)
+stat_summary(fun=median, geom="point", size=0.6, color="red")+ylab(gene_list[k])+ theme(axis.title.x=element_blank(),
axis.text.y=element_text(size=5),axis.title.y=element_text(size=6,angle=0,face="bold",margin = margin(t = 10, r = 8, b = 0, l = 10),vjust=0.5),axis.text.x=element_text(size=5,face="bold")
,axis.title=element_text(size=4,face="bold"),panel.grid.major = element_blank(), panel.grid.minor = element_blank(),
panel.background = element_blank(), axis.line = element_line(colour = "black"),legend.position="none",plot.margin = unit(c(0, 0,0, 0), "cm")))}
else{
assign(paste("P",k,sep=""),ggplot(Cell_number,aes_string(x="type",y=gene_list[k],fill="type"))+geom_violin(scale = "width")+scale_fill_manual(values=colors)
+stat_summary(fun=median, geom="point", size=0.6, color="red")+ylab(gene_list[k])+ theme(axis.title.x=element_blank(),
axis.text.x=element_blank(),axis.text.y=element_text(size=5),axis.title.y=element_text(size=6,angle=0,face="bold",margin = margin(t = 10, r = 8, b = 0, l = 10),vjust=0.5),axis.ticks.x=element_blank(),axis.text=element_text(size=5)
,axis.title=element_text(size=4,face="bold"),panel.grid.major = element_blank(), panel.grid.minor = element_blank(),
panel.background = element_blank(), axis.line = element_line(colour = "black"),legend.position="none",plot.margin = unit(c(0, 0,0, 0), "cm")))}

}
```

```
P1
P2
P3
P4
P5
P6
P7
P8
P9
P10
P11
P12
P13
```

```
merge<-list()
for (i in length(gene_list):1){
       if (length(merge)==0){
        merge<-ggplotGrob(eval(parse(text=paste("P",i,sep = ""))))}else{merge<-rbind(
ggplotGrob(eval(parse(text=paste("P",i,sep = "")))),merge,size = "last")}}

 #all genes
pdf(paste("/Users/lees44/R_project/FC_dCA2_all_vector_Integration_SUN1GFP_MYC_050321/
Cell_Type_Gene_Expression",".pdf",sep=""),height=6, width=50 , paper = "letter")


grid.newpage()
grid.draw(merge)
```

################SubsetData: Return a subset of the Seurat object################################ Subset data https://www.rdocumentation.org/packages/Seurat/versions/3.1.4/topics/SubsetData (https://www.rdocumentation.org/packages/Seurat/versions/3.1.4/topics/SubsetData) Look at hashikawa codes

```
for (i in 1:length(new.ident)){assign(paste(new.ident[i],"_barcode",sep=""),colnames(
dCA2_FC_integrated.combined@assays$RNA@data[,which(Idents(object=dCA2_FC_integrated.c
ombined) %in% new.ident[i])]))}
```

```
dCA2_FC_integrated.combined_clean<-subset(x=dCA2_FC_integrated.combined,cells=c(Neuro
n1_barcode, Neuron2_barcode, Neuron3_barcode, Neuron4_barcode, Neuron5_barcode, Neuro
n6_barcode, Neuron7_barcode, Neuron8_barcode, Neuron9_barcode, Neuron10_barcode, Olig
odendrocyte_barcode, Astrocyte_barcode, OPC_barcode, Microglia_barcode, Eppendymal_ba
rcode))
```

```
saveRDS(dCA2_FC_integrated.combined_clean, file = "/Users/lees44/R_project/FC_dCA2_al
l_vector_Integration_SUN1GFP_MYC_050321/dCA2_FC_integrated.combined_clean.rds")
```

```
DimPlot(object = dCA2_FC_integrated.combined_clean, reduction = "umap", label = FALS
E, repel = TRUE) & theme(legend.text=element_text(size=18))
ggsave(file="/Users/lees44/R_project/FC_dCA2_all_vector_Integration_SUN1GFP_MYC_05032
1/Neuronaltype_clean.pdf",width=10,height=10)
```

```
p2<-DimPlot(dCA2_FC_integrated.combined_clean, reduction = "umap", group.by = "protoc
ol")& theme(legend.text=element_text(size=18))
plot_grid(p2)
ggsave(file="/Users/lees44/R_project/FC_dCA2_all_vector_Integration_SUN1GFP_MYC_05032
1/UMAP_clean.pdf",width=10,height=10)
```

#################Cleaned data set 07.14.21########################## object = dCA2_FC_integrated.combined_clean

Cluster tree

```
#cluster tree to go with the violin plot based on cell type
pdf(file="/Users/lees44/R_project/FC_dCA2_all_vector_Integration_SUN1GFP_MYC_050321/c
luster_tree_clean.pdf",width=20,height=8,paper='special')
#https://www.rdocumentation.org/packages/ape/versions/5.2/topics/plot.phylo
dCA2_FC_integrated.combined_clean <- BuildClusterTree(dCA2_FC_integrated.combined_cle
an, verbose = FALSE, reorder = FALSE)
plot(dCA2_FC_integrated.combined_clean@tools$BuildClusterTree, type = "phylogram", ed
ge.color = "black", edge.width = 4,edge.lty = 1, srt = 0, label.offset = 10, directio
n = "downwards", tip.color = "black", font = 2)
dev.off()
```

Cell type and violin plot based on cell type

Violinplot for canonical markers, figure goes with cluster tree

```
Cell_type<-rev(c("Neuron8", "Neuron10", "Neuron5", "Neuron6",  "Neuron4", "Neuron7",
"Oligodendrocyte", "Astrocyte", "Microglia",  "Eppendymal", "OPC", "Neuron9", "Neuron
1","Neuron2",  "Neuron3"))
gene_list<-c("Camk2a", "Kif5c", "Mog" ,"Opalin", "Slc7a10", "Gfap", "Tmem119", "C1q
c", "Foxj1", "Fam216b", "Pdgfra", "Gpr17")
```

```
Cell_number <- NULL
for (i in 1:length(Cell_type)){
                temp1 <- data.frame(type=factor(rep(Cell_type[i],sum(Cell_type[i]==Id
ents(dCA2_FC_integrated.combined_clean)))))
                temp2 <- as.data.frame(dCA2_FC_integrated.combined_clean@assays$RNA@d
ata[gene_list,Idents(dCA2_FC_integrated.combined_clean)==Cell_type[i]])
                temp3 <- cbind(temp1,t(temp2))
                Cell_number <- rbind(temp3,Cell_number)
}
rownames(Cell_number) <- NULL
```

```
colors<-c("#ffc857","#ffc857","#ffc857","#ffc857","#ffc857","#ffc857","#9B0C1E","#676
833","#B99A69","#19647e","#4b3f72","#ffc857","#ffc857","#ffc857","#ffc857")
```

```
for (k in 1:length(gene_list))
    {
if (k==length(gene_list)){
assign(paste("P",k,sep=""),ggplot(Cell_number,aes_string(x="type",y=gene_list[k],fill
="type"))+geom_violin(scale = "width")+scale_fill_manual(values=colors)
+stat_summary(fun=median, geom="point", size=0.6, color="red")+ylab(gene_list[k])+ th
eme(axis.title.x=element_blank(),
axis.text.y=element_text(size=5),axis.title.y=element_text(size=6,angle=0,face="bold",
margin = margin(t = 10, r = 8, b = 0, l = 10),vjust=0.5),axis.text.x=element_text(siz
e=5,face="bold")
,axis.title=element_text(size=4,face="bold"),panel.grid.major = element_blank(), pane
l.grid.minor = element_blank(),
panel.background = element_blank(), axis.line = element_line(colour = "black"),legen
d.position="none",plot.margin = unit(c(0, 0,0, 0), "cm")))}
else{
assign(paste("P",k,sep=""),ggplot(Cell_number,aes_string(x="type",y=gene_list[k],fill
="type"))+geom_violin(scale = "width")+scale_fill_manual(values=colors)
+stat_summary(fun=median, geom="point", size=0.6, color="red")+ylab(gene_list[k])+ th
eme(axis.title.x=element_blank(),
axis.text.x=element_blank(),axis.text.y=element_text(size=5),axis.title.y=element_tex
t(size=6,angle=0,face="bold",margin = margin(t = 10, r = 8, b = 0, l = 10),vjust=0.
5),axis.ticks.x=element_blank(),axis.text=element_text(size=5)
,axis.title=element_text(size=4,face="bold"),panel.grid.major = element_blank(), pane
l.grid.minor = element_blank(),
panel.background = element_blank(), axis.line = element_line(colour = "black"),legen
d.position="none",plot.margin = unit(c(0, 0,0, 0), "cm")))}

}
```

```
P1
P2
P3
P4
P5
P6
P7
P8
P9
P10
P11
P12
P13
```

```
merge<-list()
for (i in length(gene_list):1){
       if (length(merge)==0){
        merge<-ggplotGrob(eval(parse(text=paste("P",i,sep = ""))))}else{merge<-rbind(
ggplotGrob(eval(parse(text=paste("P",i,sep = "")))),merge,size = "last")}}

 #all genes
pdf(paste("/Users/lees44/R_project/FC_dCA2_all_vector_Integration_SUN1GFP_MYC_050321/
Cell_Type_Gene_Expression_clean",".pdf",sep=""),height=6, width=50 , paper = "lette
r")


grid.newpage()
grid.draw(merge)
```

######################################072021##################subset dCA2 markers################################

Feature plot highlighting dCA2 markers in all population Expression pots in UMAP space illustrating normalized expression values of dCA2 markers in clean dataset

“CREREV38354884”, “GFP”,“Rgs14”, “Avpr1b”, “Amigo2”, “Map3k15”, “Actn2”, “Ptpn5”, “Cacng5”, “Oxtr”, “Pcp4”

```
#Feature plot highlighting dCA2 markers in all population
F<-FeaturePlot(object =dCA2_FC_integrated.combined_clean, features = c("CREREV3835488
4"), blend.threshold = 0.5, min.cutoff = 0, max.cutoff = 1.5) + scale_fill_continuou
s(type= "viridis")

plot(F)
ggsave(file="/Users/lees44/R_project/FC_dCA2_all_vector_Integration_SUN1GFP_MYC_05032
1/CREREV38354884_clean.pdf",width=10,height=10)
```

```
#Feature plot highlighting dCA2 markers in all population
F<-FeaturePlot(object =dCA2_FC_integrated.combined_clean, features = c("GFP"), blend.
threshold = 0.5, min.cutoff = 0, max.cutoff = 2) + scale_fill_continuous(type= "virid
is")
plot(F)
ggsave(file="/Users/lees44/R_project/FC_dCA2_all_vector_Integration_SUN1GFP_MYC_05032
1/GFP_clean.pdf",width=10,height=10)
```

```
#Feature plot highlighting dCA2 markers in all population
F<-FeaturePlot(object =dCA2_FC_integrated.combined_clean, features = c("Rgs14"), blen
d.threshold = 0.5, min.cutoff = 0, max.cutoff = 3) + scale_fill_continuous(type= "vir
idis")
plot(F)
ggsave(file="/Users/lees44/R_project/FC_dCA2_all_vector_Integration_SUN1GFP_MYC_05032
1/Rgs14_clean.pdf",width=10,height=10)
```

```
#Feature plot highlighting dCA2 markers in all population
F<-FeaturePlot(object =dCA2_FC_integrated.combined_clean, features = c("Avpr1b"), ble
nd.threshold = 0.5, min.cutoff = 0, max.cutoff = 1.5) + scale_fill_continuous(type= "
viridis")
plot(F)
ggsave(file="/Users/lees44/R_project/FC_dCA2_all_vector_Integration_SUN1GFP_MYC_05032
1/Avpr1b_clean.pdf",width=10,height=10)
```

```
#Feature plot highlighting dCA2 markers in all population
F<-FeaturePlot(object =dCA2_FC_integrated.combined_clean, features = c("Amigo2"), ble
nd.threshold = 0.2, min.cutoff = 0, max.cutoff = 3) + scale_fill_continuous(type= "vi
ridis")
plot(F)
ggsave(file="/Users/lees44/R_project/FC_dCA2_all_vector_Integration_SUN1GFP_MYC_05032
1/Amigo2_clean.pdf",width=10,height=10)
```

```
#Feature plot highlighting dCA2 markers in all population
F<-FeaturePlot(object =dCA2_FC_integrated.combined_clean, features = c("Map3k15"), bl
end.threshold = 0.1, min.cutoff = 0, max.cutoff = 3) + scale_fill_continuous(type= "v
iridis")
plot(F)
ggsave(file="/Users/lees44/R_project/FC_dCA2_all_vector_Integration_SUN1GFP_MYC_05032
1/Map3k15_clean.pdf",width=10,height=10)
```

```
#Feature plot highlighting dCA2 markers in all population
F<-FeaturePlot(object =dCA2_FC_integrated.combined_clean, features = c("Actn2"), blen
d.threshold = 0.1, min.cutoff = 0, max.cutoff = 2)  + scale_fill_continuous(type= "vi
ridis")
plot(F)
ggsave(file="/Users/lees44/R_project/FC_dCA2_all_vector_Integration_SUN1GFP_MYC_05032
1/Actn2_clean.pdf",width=10,height=10)
```

```
#Feature plot highlighting dCA2 markers in all population
F<-FeaturePlot(object =dCA2_FC_integrated.combined_clean, features = c("Ptpn5"), blen
d.threshold = 0.5, min.cutoff = 0, max.cutoff = 3) + scale_fill_continuous(type= "vir
idis")
plot(F)
ggsave(file="/Users/lees44/R_project/FC_dCA2_all_vector_Integration_SUN1GFP_MYC_05032
1/Ptpn5_clean.pdf",width=10,height=10)
```

```
#Feature plot highlighting dCA2 markers in all population
F<-FeaturePlot(object =dCA2_FC_integrated.combined_clean, features = c("Cacng5"), ble
nd.threshold = 0.5, min.cutoff = 0, max.cutoff = 3) + scale_fill_continuous(type= "vi
ridis")
plot(F)
ggsave(file="/Users/lees44/R_project/FC_dCA2_all_vector_Integration_SUN1GFP_MYC_05032
1/Cacng5_clean.pdf",width=10,height=10)
```

```
#Feature plot highlighting dCA2 markers in all population
F<-FeaturePlot(object =dCA2_FC_integrated.combined_clean, features = c("Oxtr"), blen
d.threshold = 0.5, min.cutoff = 0, max.cutoff = 1.5) + scale_fill_continuous(type= "v
iridis")
plot(F)
ggsave(file="/Users/lees44/R_project/FC_dCA2_all_vector_Integration_SUN1GFP_MYC_05032
1/Oxtr_clean.pdf",width=10,height=10)
```

```
#Feature plot highlighting dCA2 markers in all population
F<-FeaturePlot(object =dCA2_FC_integrated.combined_clean, features = c("Pcp4"), blen
d.threshold = 0.5, min.cutoff = 0, max.cutoff = 3) + scale_fill_continuous(type= "vir
idis")
plot(F)
ggsave(file="/Users/lees44/R_project/FC_dCA2_all_vector_Integration_SUN1GFP_MYC_05032
1/Pcp4_clean.pdf",width=10,height=10)
```
