## Supplementary material for "Investigation of the Fasciola Cinereum, Absent in BTBR mice, and Comparison with the Hippocampal Area CA2": 8_Characterization of Fasciola cinerum, Avpr1b neurons only

**Su Hyun Lee**

This is an R Markdown (http://rmarkdown.rstudio.com) Notebook. When you execute code within the notebook, the results appear beneath the code.

Try executing this chunk by clicking the *Run* button within the chunk or by placing your cursor inside it and pressing *Cmd+Shift+Enter*.

```
knitr::opts_chunk$set(error = TRUE) # do not interrupt in case of errors
```

Upload all the packages

```
library(Seurat)
```

```
## Attaching SeuratObject
```

```
library(dplyr)
```

```
##
## Attaching package: 'dplyr'
```

```
## The following objects are masked from 'package:stats':
##
##     filter, lag
```

```
## The following objects are masked from 'package:base':
##
##     intersect, setdiff, setequal, union
```

```
library(magrittr)
library("ggplot2")
library(grid)
library(reshape2)
library("scales")
```

Next step describes creating object for each group and integration of all the GFP positive samples. dCA2 Male GFP positive sample

```
#attach data for Male GFP positive sample (sample_1_dCA2_M)
#get working directory
getwd()
```

```
## [1] "/Users/lees44/R_project"
```

```
#Set working directory where data file located
setwd("/Users/lees44/R_project/SUN1GFP_MYC_CREREV38354884_OUTPUT/sample_1_dCA2_M")
#check folder and list files
list.files("/Users/lees44/R_project/SUN1GFP_MYC_CREREV38354884_OUTPUT/sample_1_dCA2_M")
```

```
## [1] "barcodes.tsv.gz" "features.tsv.gz" "matrix.mtx.gz"
```

```
#Load M1_GFPve_pos data
dCA2_M1_GFPve_pos_barcodes.data <- Read10X(data.dir = "/Users/lees44/R_project/SUN1GFP_MYC_CREREV38354884_OUTPUT/
sample_1_dCA2_M")
colnames(dCA2_M1_GFPve_pos_barcodes.data) = paste0(colnames(dCA2_M1_GFPve_pos_barcodes.data),"dCA2_MI_GFPve_pos")
#check dimension for data
dim(dCA2_M1_GFPve_pos_barcodes.data)
```

```
## [1] 21861  2000
```

```
#Initialize the Seurat object with the raw (non-normalized data).
dCA2_MI_GFPve_pos <- CreateSeuratObject(counts = dCA2_M1_GFPve_pos_barcodes.data, project = "dCA2", min.cells =
3, min.features = 200)
#only genes that are are expressed in 3 or more cells and cells with complexity of 200 genes or more
dCA2_MI_GFPve_pos@meta.data$protocol<- "dCA2_Mpos"
```

dCA2 Female GFP positive sample

```
#attach data for Female GFP positive sample (sample_2_dCA2_F)
#get working directory
getwd()
```

```
## [1] "/Users/lees44/R_project"
```

```
#Set working directory where data file located
setwd("/Users/lees44/R_project/SUN1GFP_MYC_CREREV38354884_OUTPUT/sample_2_dCA2_F")
#check folder and list files
list.files("/Users/lees44/R_project/SUN1GFP_MYC_CREREV38354884_OUTPUT/sample_2_dCA2_F")
```

```
## [1] "barcodes.tsv.gz" "features.tsv.gz" "matrix.mtx.gz"
```

```
#Load M1_GFPve_pos data
dCA2_FI_GFPve_pos_barcodes.data <- Read10X(data.dir = "/Users/lees44/R_project/SUN1GFP_MYC_CREREV38354884_OUTPUT/
sample_2_dCA2_F")
colnames(dCA2_FI_GFPve_pos_barcodes.data) = paste0(colnames(dCA2_FI_GFPve_pos_barcodes.data),"dCA2_FI_GFPve_pos")
#check dimension for data
dim(dCA2_FI_GFPve_pos_barcodes.data)
```

```
## [1] 21861  1142
```

```
#Initialize the Seurat object with the raw (non-normalized data).
dCA2_FI_GFPve_pos <- CreateSeuratObject(counts = dCA2_FI_GFPve_pos_barcodes.data, project = "dCA2", min.cells =
3, min.features = 200)
dCA2_FI_GFPve_pos@meta.data$protocol <- "dCA2_Fpos"
```

FC male GFP positive

```
#attach data for Male GFP positive sample (sample_1_FC_M)
#get working directory
getwd()
```

```
## [1] "/Users/lees44/R_project"
```

```
#Set working directory where data file located
setwd("/Users/lees44/R_project/FC_SUN1GFP_MYC_042021_OUTPUT/sample_1_FC_M")
#check folder and list files
list.files("/Users/lees44/R_project/FC_SUN1GFP_MYC_042021_OUTPUT/sample_1_FC_M")
```

```
## [1] "barcodes.tsv.gz" "features.tsv.gz" "matrix.mtx.gz"
```

```
#Load M1_GFPve_pos data
FC_M1_GFPve_pos_barcodes.data <- Read10X(data.dir = "/Users/lees44/R_project/FC_SUN1GFP_MYC_042021_OUTPUT/sample_
1_FC_M")
colnames(FC_M1_GFPve_pos_barcodes.data) = paste0(colnames(FC_M1_GFPve_pos_barcodes.data),"FC_MI_GFPve_pos")
#check dimension for data
dim(FC_M1_GFPve_pos_barcodes.data)
```

```
## [1] 21861  1247
```

```
#Initialize the Seurat object with the raw (non-normalized data).
FC_MI_GFPve_pos <- CreateSeuratObject(counts = FC_M1_GFPve_pos_barcodes.data, project = "dCA2", min.cells = 3, mi
n.features = 200)
#only genes that are are expressed in 3 or more cells and cells with complexity of 200 genes or more
FC_MI_GFPve_pos@meta.data$protocol<- "FC_Mpos"
```

Checking data for each group

```
#How many genes are you left with? How many cells?
dim(FC_MI_GFPve_pos)
```

```
## [1] 12899  1247
```

```
#How much memory does a sparse matrix take up relative to a dense matrix
object.size(FC_M1_GFPve_pos_barcodes.data)
```

```
## 34546008 bytes
```

```
object.size(as.matrix(FC_M1_GFPve_pos_barcodes.data))
```

```
## 219631392 bytes
```

```
#Filtering low-quality cells
#Look at the summary counts for genes and cells
counts_per_cell <- Matrix::colSums(FC_M1_GFPve_pos_barcodes.data)
cat("counts per cell: ", counts_per_cell[1:5], "\n") ## counts for first 5 cells
```

```
## counts per cell:  3663 4798 2670 3773 3434
```

```
counts_per_gene <- Matrix::rowSums(FC_M1_GFPve_pos_barcodes.data)
cat("counts per gene: ", counts_per_gene[1:5], "\n")  ## counts for first 5 genes
```

```
## counts per gene:  375 0 0 329 156
```

```
genes_per_cell <- Matrix::colSums(FC_M1_GFPve_pos_barcodes.data > 0) # count gene only if it has non-zero reads m
apped.
cat("counts for non-zero genes: ", genes_per_cell[1:5])  ## counts for first 5 genes
```

```
## counts for non-zero genes:  2186 2653 1748 2250 1972
```

```
hist(log10(counts_per_cell+1),main='counts per cell',col='wheat')
```

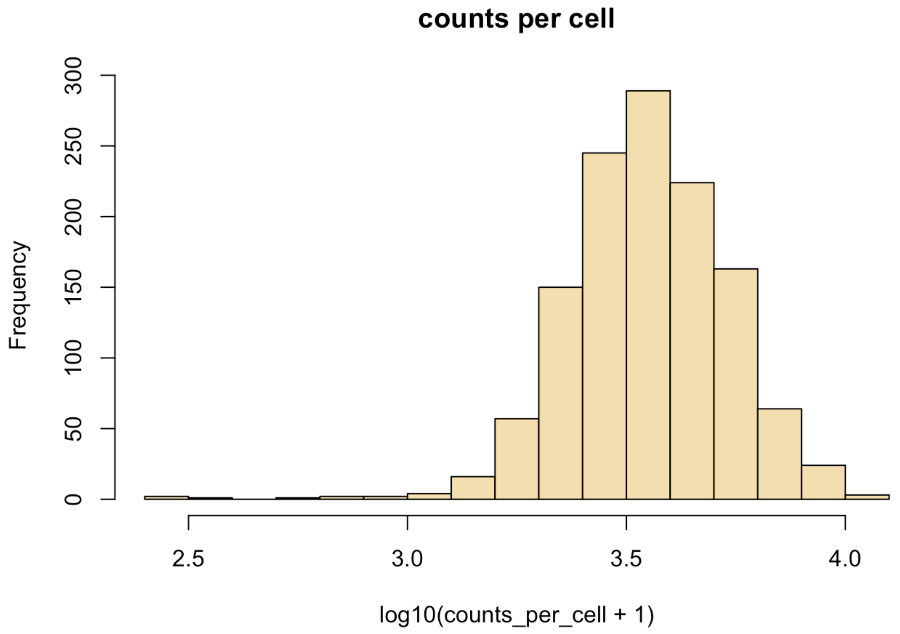


```
hist(log10(genes_per_cell+1), main='genes per cell', col='wheat')
```

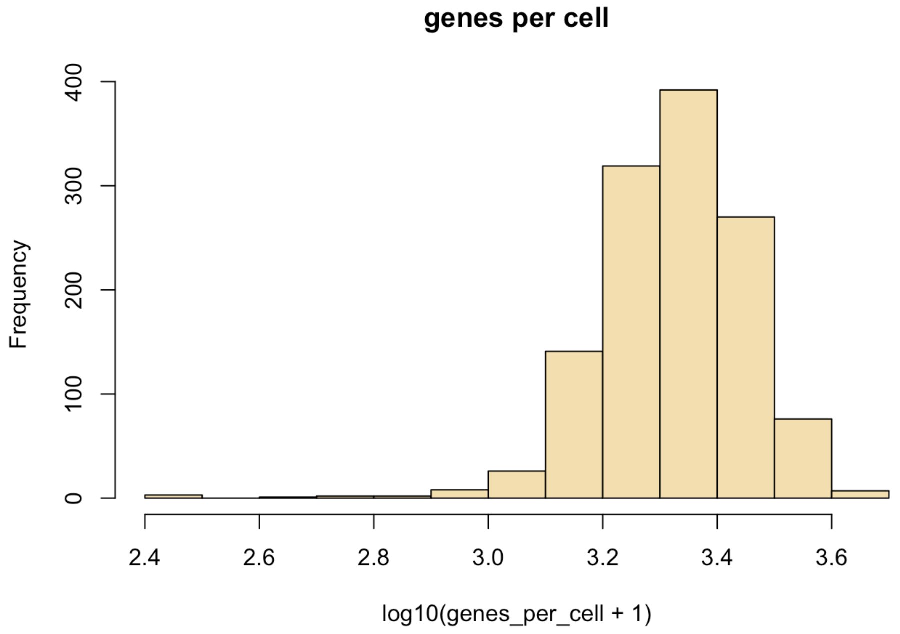


```
plot(counts_per_cell, genes_per_cell, log='xy', col='wheat')
title('counts vs genes per cell')
```

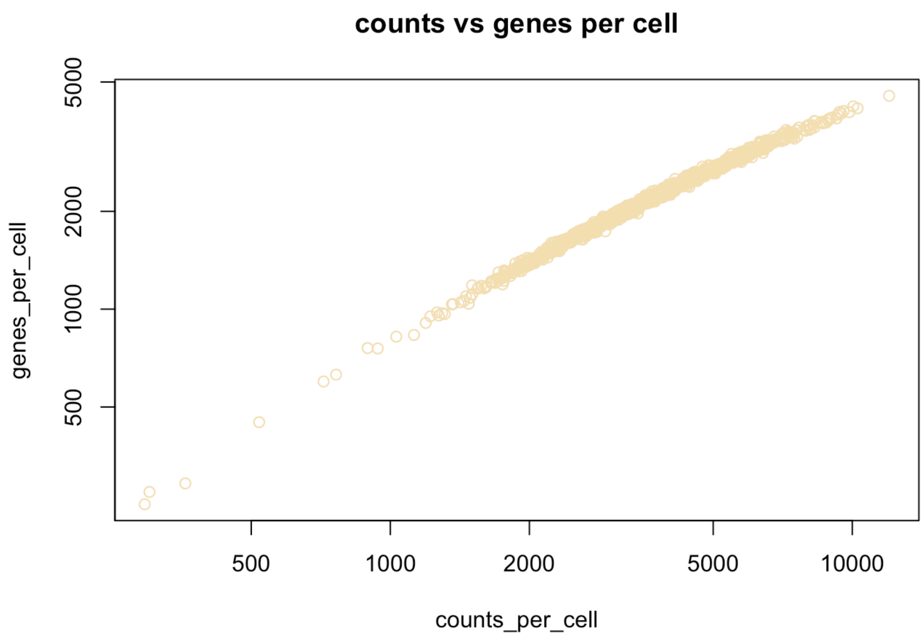


```
#plot cells ranked by their number of detected genes
#Here we rank each cell by its library complexity, ie the number of genes detected per cell. This is a very usefu
l plot as it shows the distribution of library complexity in the sequencing run. One can use this plot to investi
gate observations (potential cells) that are actually failed libraries (lower end outliers) or observations that
are cell doublets (higher end outliers).
plot(sort(genes_per_cell), xlab='cell', log='y', main='genes per cell (ordered)')
```

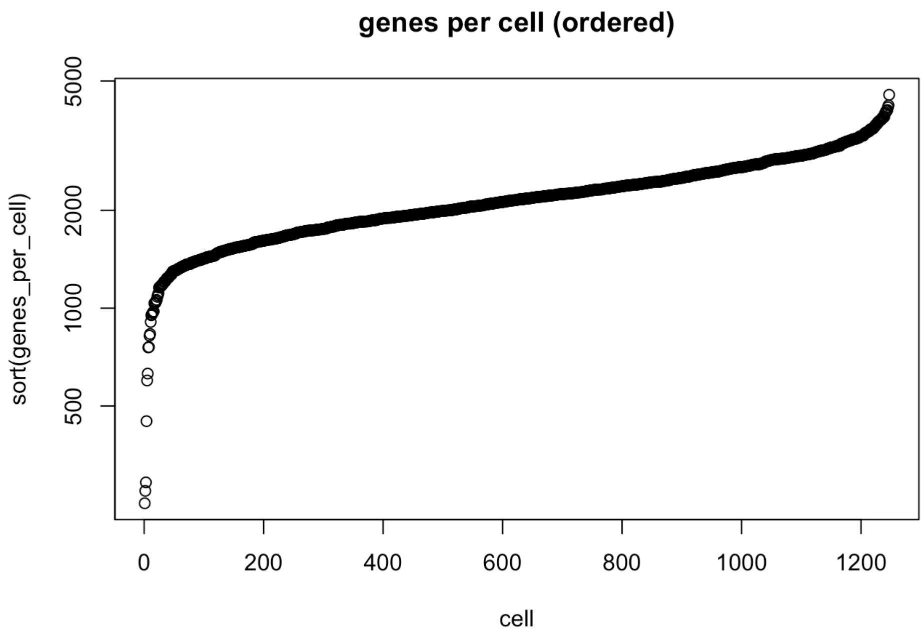


FC Female GFP positive

```
#attach data for Female GFP positive sample (sample_2_FC_F)
#get working directory
getwd()
```

```
## [1] "/Users/lees44/R_project"
```

```
#Set working directory where data file located
setwd("/Users/lees44/R_project/FC_SUN1GFP_MYC_042021_OUTPUT/sample_2_FC_F")
#check folder and list files
list.files("/Users/lees44/R_project/FC_SUN1GFP_MYC_042021_OUTPUT/sample_2_FC_F")
```

```
## [1] "barcodes.tsv.gz" "features.tsv.gz" "matrix.mtx.gz"
```

```
#Load M1_GFPve_pos data
FC_FI_GFPve_pos_barcodes.data <- Read10X(data.dir = "/Users/lees44/R_project/FC_SUN1GFP_MYC_042021_OUTPUT/sample_
2_FC_F")
colnames(FC_FI_GFPve_pos_barcodes.data) = paste0(colnames(FC_FI_GFPve_pos_barcodes.data),"FC_FI_GFPve_pos")
#check dimension for data
dim(FC_FI_GFPve_pos_barcodes.data)
```

```
## [1] 21861  1575
```

```
#Initialize the Seurat object with the raw (non-normalized data).
FC_FI_GFPve_pos <- CreateSeuratObject(counts = FC_FI_GFPve_pos_barcodes.data, project = "dCA2", min.cells = 3, mi
n.features = 200)
FC_FI_GFPve_pos@meta.data$protocol <- "FC_Fpos"
```

QC and normalization prior to integration

```
mito.features <- grep(pattern = "^mt-", x = rownames(x = dCA2_MI_GFPve_pos), value = TRUE)
percent.mito <- Matrix::colSums(x = GetAssayData(object = dCA2_MI_GFPve_pos, slot = 'counts')[mito.features, ]) /
Matrix::colSums(x = GetAssayData(object = dCA2_MI_GFPve_pos, slot = 'counts'))
dCA2_MI_GFPve_pos[['percent.mito']] <- percent.mito
```

```
mito.features <- grep(pattern = "^mt-", x = rownames(x = dCA2_FI_GFPve_pos), value = TRUE)
percent.mito <- Matrix::colSums(x = GetAssayData(object = dCA2_FI_GFPve_pos, slot = 'counts')[mito.features, ]) /
Matrix::colSums(x = GetAssayData(object = dCA2_FI_GFPve_pos, slot = 'counts'))
dCA2_FI_GFPve_pos[['percent.mito']] <- percent.mito
```

```
mito.features <- grep(pattern = "^mt-", x = rownames(x = FC_MI_GFPve_pos), value = TRUE)
percent.mito <- Matrix::colSums(x = GetAssayData(object = FC_MI_GFPve_pos, slot = 'counts')[mito.features, ]) / M
atrix::colSums(x = GetAssayData(object = FC_MI_GFPve_pos, slot = 'counts'))
FC_MI_GFPve_pos[['percent.mito']] <- percent.mito
```

```
mito.features <- grep(pattern = "^mt-", x = rownames(x = FC_FI_GFPve_pos), value = TRUE)
percent.mito <- Matrix::colSums(x = GetAssayData(object = FC_FI_GFPve_pos, slot = 'counts')[mito.features, ]) / M
atrix::colSums(x = GetAssayData(object = FC_FI_GFPve_pos, slot = 'counts'))
FC_FI_GFPve_pos[['percent.mito']] <- percent.mito
```

```
dCA2_MI_GFPve_pos <- subset(x = dCA2_MI_GFPve_pos, subset = nCount_RNA > 700 & nCount_RNA < 15000 & percent.mito
< 0.20)
dCA2_FI_GFPve_pos <- subset(x = dCA2_FI_GFPve_pos, subset = nCount_RNA > 700 & nCount_RNA < 15000 & percent.mito
< 0.20)
FC_MI_GFPve_pos <- subset(x = FC_MI_GFPve_pos, subset = nCount_RNA > 700 & nCount_RNA < 15000 & percent.mito < 0.
20)
FC_FI_GFPve_pos <- subset(x = FC_FI_GFPve_pos, subset = nCount_RNA > 700 & nCount_RNA < 15000 & percent.mito < 0.
20)
```

```
dCA2_MI_GFPve_pos<- NormalizeData(object = dCA2_MI_GFPve_pos,verbose = FALSE)
dCA2_FI_GFPve_pos<- NormalizeData(object = dCA2_FI_GFPve_pos,verbose = FALSE)
FC_MI_GFPve_pos<- NormalizeData(object = FC_MI_GFPve_pos,verbose = FALSE)
FC_FI_GFPve_pos<- NormalizeData(object = FC_FI_GFPve_pos,verbose = FALSE)
```

nfeatures increaed from 2000 to 5000 in order to detect rarely expressed genes. default = 2000. Avpr1b and Oxtr transcripts expression are specific for CA2 but expression.

```
dCA2_MI_GFPve_pos<- FindVariableFeatures(object =dCA2_MI_GFPve_pos,selection.method = "vst", nfeatures = 5000, ve
rbose = FALSE)
dCA2_FI_GFPve_pos<- FindVariableFeatures(object =dCA2_FI_GFPve_pos,selection.method = "vst", nfeatures = 5000, ve
rbose = FALSE)
FC_MI_GFPve_pos<- FindVariableFeatures(object =FC_MI_GFPve_pos,selection.method = "vst", nfeatures = 5000, verbos
e = FALSE)
FC_FI_GFPve_pos<- FindVariableFeatures(object =FC_FI_GFPve_pos,selection.method = "vst", nfeatures = 5000, verbos
e = FALSE)
```

Integration of multiple dataset

We have multiple groups of 4 GFP positive groups. Here we merge and integrate samples by sing vector (x, y=c(1,2)). If only two samples you can follow stim vs control example from Satija et al seurat integration method.

```
#integration of multiple groups using vector (x, y=c(1,2) and split by group.
dCA2.combined = merge(dCA2_MI_GFPve_pos, y = c(dCA2_FI_GFPve_pos, FC_MI_GFPve_pos, FC_FI_GFPve_pos), add.cell.ids
= c("dCA2_Mpos", "dCA2_Fpos", "FC_Mpos", "FC_Fpos"), project = "protocol" )
```

Here we are splitting group by “protocol” and the groups include “dCA2_Mpos”, “dCA2_Fpos”,“FC_Mpos”, “FC_Fpos” from metadata we have created prior. Find a set of anchors between a list of Seurat objects. These anchors can later be used to integrate the objects using the IntegrateData function.

```
dCA2.list <-SplitObject(dCA2.combined, split.by = "protocol")
reference.list <-dCA2.list[c("dCA2_Mpos", "dCA2_Fpos","FC_Mpos", "FC_Fpos")]
dCA2.anchors <- FindIntegrationAnchors(object.list = reference.list, dims = 1:30)
```

```
## Computing 2000 integration features
```

```
## No variable features found for object1 in the object.list. Running FindVariableFeatures ...
```

```
## No variable features found for object2 in the object.list. Running FindVariableFeatures ...
```

```
## No variable features found for object3 in the object.list. Running FindVariableFeatures ...
```

```
## No variable features found for object4 in the object.list. Running FindVariableFeatures ...
```

```
## Scaling features for provided objects
```

```
## Finding all pairwise anchors
```

```
## Running CCA
```

```
## Merging objects
```

```
## Finding neighborhoods
```

```
## Finding anchors
```

```
##  Found 4981 anchors
```

```
## Filtering anchors
```

```
##  Retained 2194 anchors
```

```
## Running CCA
```

```
## Merging objects
```

```
## Finding neighborhoods
```

```
## Finding anchors
```

```
##  Found 5407 anchors
```

```
## Filtering anchors
```

```
##  Retained 1646 anchors
```

```
## Running CCA
```

```
## Merging objects
```

```
## Finding neighborhoods
```

```
## Finding anchors
```

```
##  Found 4373 anchors
```

```
## Filtering anchors
```

```
##  Retained 1291 anchors
```

```
## Running CCA
```

```
## Merging objects
```

```
## Finding neighborhoods
```

```
## Finding anchors
```

```
##  Found 6242 anchors
```

```
## Filtering anchors
```

```
##  Retained 1485 anchors
```

```
## Running CCA
```

```
## Merging objects
```

```
## Finding neighborhoods
```

```
## Finding anchors
```

```
##  Found 4753 anchors
```

```
## Filtering anchors
```

```
##  Retained 1184 anchors
```

```
## Running CCA
```

```
## Merging objects
```

```
## Finding neighborhoods
```

```
## Finding anchors
```

```
##  Found 5090 anchors
```

```
## Filtering anchors
```

```
##  Retained 1554 anchors
```

Integration https://satijalab.org/seurat/reference/integratedata (https://satijalab.org/seurat/reference/integratedata)

```
#intgrated analysis
dCA2.integrated <-IntegrateData(anchorset = dCA2.anchors, dims = 1:30)
```

```
## Merging dataset 2 into 1
```

```
## Extracting anchors for merged samples
```

```
## Finding integration vectors
```

```
## Finding integration vector weights
```

```
## Integrating data
```

```
## Merging dataset 3 into 4
```

```
## Extracting anchors for merged samples
```

```
## Finding integration vectors
```

```
## Finding integration vector weights
```

```
## Integrating data
```

```
## Merging dataset 4 3 into 1 2
```

```
## Extracting anchors for merged samples
```

```
## Finding integration vectors
```

```
## Finding integration vector weights
```

```
## Integrating data
```

```
str(dCA2.integrated)
```

```
## Formal class 'Seurat' [package "SeuratObject"] with 13 slots
##   ..@ assays      :List of 2
##   .. ..$ RNA       :Formal class 'Assay' [package "SeuratObject"] with 8 slots
##   .. .. .. ..@ counts       :Formal class 'dgCMatrix' [package "Matrix"] with 6 slots
##   .. .. .. .. .. ..@ i       : int [1:13380726] 1 2 3 7 9 10 13 20 21 22 ...
##   .. .. .. .. .. ..@ p       : int [1:5929] 0 3052 6083 7585 9263 14024 16028 17807 21209 23193 ...
##   .. .. .. .. .. ..@ Dim     : int [1:2] 14756 5928
##   .. .. .. .. .. ..@ Dimnames:List of 2
##   .. .. .. .. .. .. ..$ : chr [1:14756] "Xkr4" "Mrpl15" "Lypla1" "Tcea1" ...
##   .. .. .. .. .. .. ..$ : chr [1:5928] "dCA2_Mpos_AAACCCATCTGAGGCC-1dCA2_MI_GFPve_pos" "dCA2_Mpos_AAACGAAGTCCA
AAGG-1dCA2_MI_GFPve_pos" "dCA2_Mpos_AAAGAACCAATCACGT-1dCA2_MI_GFPve_pos" "dCA2_Mpos_AAAGAACCATGTTCGA-1dCA2_MI_GFP
ve_pos" ...
##   .. .. .. .. .. ..@ x       : num [1:13380726] 2 2 1 1 2 1 1 2 4 1 ...
##   .. .. .. .. .. ..@ factors : list()
##   .. .. .. ..@ data         :Formal class 'dgCMatrix' [package "Matrix"] with 6 slots
##   .. .. .. .. .. ..@ i       : int [1:13380726] 1 2 3 7 9 10 13 20 21 22 ...
##   .. .. .. .. .. ..@ p       : int [1:5929] 0 3052 6083 7585 9263 14024 16028 17807 21209 23193 ...
##   .. .. .. .. .. ..@ Dim     : int [1:2] 14756 5928
##   .. .. .. .. .. ..@ Dimnames:List of 2
##   .. .. .. .. .. .. ..$ : chr [1:14756] "Xkr4" "Mrpl15" "Lypla1" "Tcea1" ...
##   .. .. .. .. .. .. ..$ : chr [1:5928] "dCA2_Mpos_AAACCCATCTGAGGCC-1dCA2_MI_GFPve_pos" "dCA2_Mpos_AAACGAAGTCCA
AAGG-1dCA2_MI_GFPve_pos" "dCA2_Mpos_AAAGAACCAATCACGT-1dCA2_MI_GFPve_pos" "dCA2_Mpos_AAAGAACCATGTTCGA-1dCA2_MI_GFP
ve_pos" ...
##   .. .. .. .. .. ..@ x       : num [1:13380726] 1.442 1.442 0.961 0.961 1.442 ...
##   .. .. .. .. .. ..@ factors : list()
##   .. .. .. ..@ scale.data   : num[0 , 0 ]
##   .. .. .. ..@ key          : Named chr "rna_"
##   .. .. .. .. ..- attr(*, "names")= chr ""
##   .. .. .. ..@ assay.orig   : NULL
##   .. .. .. ..@ var.features : logi(0)
##   .. .. .. ..@ meta.features:'data.frame':   14756 obs. of  0 variables
##   .. .. .. ..@ misc         : list()
##   .. ..$ integrated:Formal class 'Assay' [package "SeuratObject"] with 8 slots
##   .. .. .. ..@ counts       :Formal class 'dgCMatrix' [package "Matrix"] with 6 slots
##   .. .. .. .. .. ..@ i       : int(0)
##   .. .. .. .. .. ..@ p       : int 0
##   .. .. .. .. .. ..@ Dim     : int [1:2] 0 0
##   .. .. .. .. .. ..@ Dimnames:List of 2
##   .. .. .. .. .. .. ..$ : NULL
##   .. .. .. .. .. .. ..$ : NULL
##   .. .. .. .. .. ..@ x       : num(0)
##   .. .. .. .. .. ..@ factors : list()
##   .. .. .. ..@ data         :Formal class 'dgCMatrix' [package "Matrix"] with 6 slots
##   .. .. .. .. .. ..@ i       : int [1:8459499] 3 5 9 11 14 22 35 38 39 43 ...
##   .. .. .. .. .. ..@ p       : int [1:5929] 0 539 1066 1332 1622 2434 2796 3124 3683 4043 ...
##   .. .. .. .. .. ..@ Dim     : int [1:2] 2000 5928
##   .. .. .. .. .. ..@ Dimnames:List of 2
##   .. .. .. .. .. .. ..$ : chr [1:2000] "Plp1" "Mbp" "mt-Co1" "mt-Co3" ...
##   .. .. .. .. .. .. ..$ : chr [1:5928] "dCA2_Mpos_AAACCCATCTGAGGCC-1dCA2_MI_GFPve_pos" "dCA2_Mpos_AAACGAAGTCCA
AAGG-1dCA2_MI_GFPve_pos" "dCA2_Mpos_AAAGAACCAATCACGT-1dCA2_MI_GFPve_pos" "dCA2_Mpos_AAAGAACCATGTTCGA-1dCA2_MI_GFP
ve_pos" ...
##   .. .. .. .. .. ..@ x       : num [1:8459499] 1.442 0.961 0.961 5.546 0.961 ...
##   .. .. .. .. .. ..@ factors : list()
##   .. .. .. ..@ scale.data   : logi [1, 1] NA
##   .. .. .. ..@ key          : chr "integrated_"
##   .. .. .. ..@ assay.orig   : NULL
##   .. .. .. ..@ var.features : chr [1:2000] "Plp1" "Mbp" "mt-Co1" "mt-Co3" ...
##   .. .. .. ..@ meta.features:'data.frame':   2000 obs. of  0 variables
##   .. .. .. ..@ misc         : NULL
##   ..@ meta.data   :'data.frame': 5928 obs. of  5 variables:
##   .. ..$ orig.ident  : chr [1:5928] "dCA2" "dCA2" "dCA2" "dCA2" ...
##   .. ..$ nCount_RNA  : num [1:5928] 6192 6050 2320 2360 12474 ...
##   .. ..$ nFeature_RNA: int [1:5928] 3052 3031 1502 1678 4761 2004 1779 3402 1984 2281 ...
```

```
##   .. ..$ protocol    : chr [1:5928] "dCA2_Mpos" "dCA2_Mpos" "dCA2_Mpos" "dCA2_Mpos" ...
##   .. ..$ percent.mito: num [1:5928] 0.000484 0.001818 0 0.002966 0.000561 ...
##   ..@ active.assay: chr "integrated"
##   ..@ active.ident: Factor w/ 1 level "dCA2": 1 1 1 1 1 1 1 1 1 1 ...
##   .. ..- attr(*, "names")= chr [1:5928] "dCA2_Mpos_AAACCCATCTGAGGCC-1dCA2_MI_GFPve_pos" "dCA2_Mpos_AAACGAAGTCC
AAAGG-1dCA2_MI_GFPve_pos" "dCA2_Mpos_AAAGAACCAATCACGT-1dCA2_MI_GFPve_pos" "dCA2_Mpos_AAAGAACCATGTTCGA-1dCA2_MI_GF
Pve_pos" ...
##   ..@ graphs      : list()
##   ..@ neighbors   : list()
##   ..@ reductions  : list()
##   ..@ images      : list()
##   ..@ project.name: chr "SeuratProject"
##   ..@ misc        : list()
##   ..@ version     :Classes 'package_version', 'numeric_version'  hidden list of 1
##   .. ..$ : int [1:3] 4 0 1
##   ..@ commands    :List of 2
##   .. ..$ FindIntegrationAnchors:Formal class 'SeuratCommand' [package "SeuratObject"] with 5 slots
##   .. .. .. ..@ name       : chr "FindIntegrationAnchors"
##   .. .. .. ..@ time.stamp : POSIXct[1:1], format: "2021-11-18 13:46:34"
##   .. .. .. ..@ assay.used : Named chr [1:4] "RNA" "RNA" "RNA" "RNA"
##   .. .. .. .. ..- attr(*, "names")= chr [1:4] "dCA2_Mpos" "dCA2_Fpos" "FC_Mpos" "FC_Fpos"
##   .. .. .. ..@ call.string: chr "FindIntegrationAnchors(object.list = reference.list, dims = 1:30)"
##   .. .. .. ..@ params     :List of 15
##   .. .. .. .. ..$ assay               : Named chr [1:4] "RNA" "RNA" "RNA" "RNA"
##   .. .. .. .. .. ..- attr(*, "names")= chr [1:4] "dCA2_Mpos" "dCA2_Fpos" "FC_Mpos" "FC_Fpos"
##   .. .. .. .. ..$ anchor.features     : chr [1:2000] "Plp1" "Mbp" "mt-Co1" "mt-Co3" ...
##   .. .. .. .. ..$ scale               : logi TRUE
##   .. .. .. .. ..$ normalization.method: chr "LogNormalize"
##   .. .. .. .. ..$ reduction           : chr "cca"
##   .. .. .. .. ..$ l2.norm             : logi TRUE
##   .. .. .. .. ..$ dims                : int [1:30] 1 2 3 4 5 6 7 8 9 10 ...
##   .. .. .. .. ..$ k.anchor            : num 5
##   .. .. .. .. ..$ k.filter            : num 200
##   .. .. .. .. ..$ k.score             : num 30
##   .. .. .. .. ..$ max.features        : num 200
##   .. .. .. .. ..$ nn.method           : chr "annoy"
##   .. .. .. .. ..$ n.trees             : num 50
##   .. .. .. .. ..$ eps                 : num 0
##   .. .. .. .. ..$ verbose             : logi TRUE
##   .. ..$ withCallingHandlers   :Formal class 'SeuratCommand' [package "SeuratObject"] with 5 slots
##   .. .. .. ..@ name       : chr "withCallingHandlers"
##   .. .. .. ..@ time.stamp : POSIXct[1:1], format: "2021-11-18 13:47:07"
##   .. .. .. ..@ assay.used : NULL
##   .. .. .. ..@ call.string: chr [1:2] "withCallingHandlers(expr, warning = function(w) if (inherits(w, " "
classes)) tryInvokeRestart(\"muffleWarning\"))"
##   .. .. .. ..@ params     :List of 9
##   .. .. .. .. ..$ new.assay.name      : chr "integrated"
##   .. .. .. .. ..$ normalization.method: chr "LogNormalize"
##   .. .. .. .. ..$ features            : chr [1:2000] "Plp1" "Mbp" "mt-Co1" "mt-Co3" ...
##   .. .. .. .. ..$ dims                : int [1:30] 1 2 3 4 5 6 7 8 9 10 ...
##   .. .. .. .. ..$ k.weight            : num 100
##   .. .. .. .. ..$ sd.weight           : num 1
##   .. .. .. .. ..$ preserve.order      : logi FALSE
##   .. .. .. .. ..$ eps                 : num 0
##   .. .. .. .. ..$ verbose             : logi TRUE
##   ..@ tools       :List of 1
##   .. ..$ Integration:Formal class 'IntegrationData' [package "Seurat"] with 7 slots
##   .. .. .. ..@ neighbors         : NULL
##   .. .. .. ..@ weights           : NULL
##   .. .. .. ..@ integration.matrix: NULL
##   .. .. .. ..@ anchors           :'data.frame':  18708 obs. of  5 variables:
##   .. .. .. .. ..$ cell1   : num [1:18708] 2 2 6 8 9 10 11 11 11 12 ...
##   .. .. .. .. ..$ cell2   : num [1:18708] 1005 85 100 1112 772 ...
##   .. .. .. .. ..$ score   : num [1:18708] 0.385 0.231 0.462 0.423 0.231 ...
##   .. .. .. .. ..$ dataset1: int [1:18708] 1 1 1 1 1 1 1 1 1 1 ...
##   .. .. .. .. ..$ dataset2: int [1:18708] 2 2 2 2 2 2 2 2 2 2 ...
##   .. .. .. ..@ offsets           : NULL
##   .. .. .. ..@ objects.ncell     : NULL
##   .. .. .. ..@ sample.tree       : num [1:3, 1:2] -1 -4 1 -2 -3 2
```

```
str(MI_GFPve_pos)
```

```
## Error in str(MI_GFPve_pos): object 'MI_GFPve_pos' not found
```

```
#median genes/cell
median(dCA2.integrated@meta.data$nFeature_RNA)
```

```
## [1] 2239
```

```
dim(dCA2.integrated)
```

```
## [1] 2000 5928
```

Upload library for plot visualization

```
library(ggplot2)
library(cowplot)
```

For integrated data or visualization on UMAP space and clustering, use integrated vs for DEG use RNA

```
DefaultAssay(dCA2.integrated) <- "integrated" # you can switch back and forth with RNA
```

Setseed function generates random number and stablizes the UMAP, if this is not set, there won't be stablilty when data is visualized on UMAP space.

```
set.seed(1234)
```

Scale data

```
dCA2.integrated <- ScaleData(object = dCA2.integrated, vars.to.regress = c("nCount_RNA", "percent.mito"),verbose
= FALSE)
```

Vizualize on PCA dimension

```
dCA2.integrated <- RunPCA(object = dCA2.integrated, npcs = 50, verbose = FALSE)# npcs 50 default
```

You must define and justify the PC values that are chosen. Here we plot elbow plot and determine where the data flattens.

```
mat <- GetAssayData(dCA2.integrated, slot = "scale.data")
pca <- dCA2.integrated[["pca"]]
total_variance <- sum(matrixStats::rowVars(mat))
eigValues = (pca@stdev)^2  ## EigenValues
varExplained = (eigValues / total_variance)*100
plot(varExplained,type='l')
```

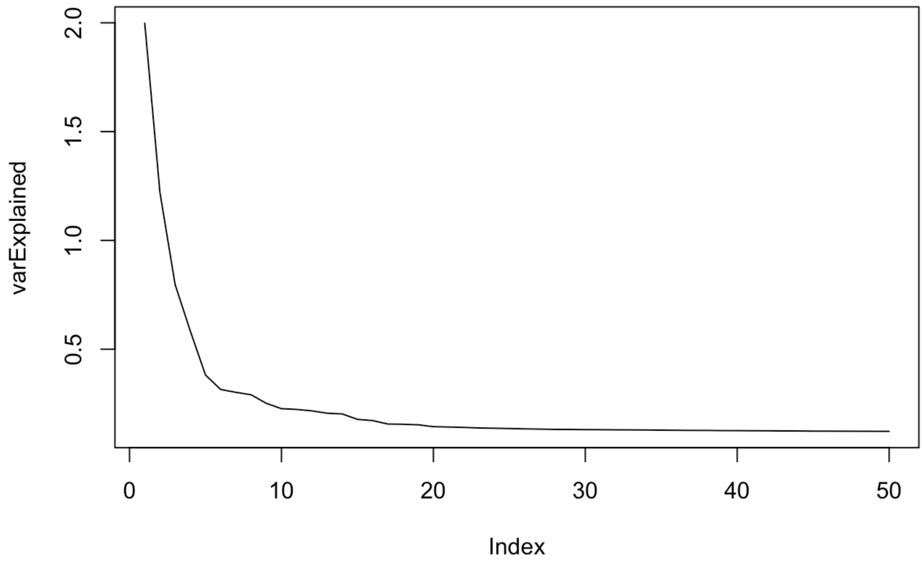


We have chosen pc value of 30 based on elbow plot and ran PCA.

```
dCA2.integrated <- RunPCA(object = dCA2.integrated, npcs = 30, verbose = FALSE)# npcs 50 default
```

RunUMAP

```
dCA2.integrated <- RunUMAP(object = dCA2.integrated, reduction = "pca", dims = 1:30)
```

```
## Warning: The default method for RunUMAP has changed from calling Python UMAP via reticulate to the R-native UW
OT using the cosine metric
## To use Python UMAP via reticulate, set umap.method to 'umap-learn' and metric to 'correlation'
## This message will be shown once per session
```

```
## 13:49:29 UMAP embedding parameters a = 0.9922 b = 1.112
```

```
## 13:49:29 Read 5928 rows and found 30 numeric columns
```

```
## 13:49:29 Using Annoy for neighbor search, n_neighbors = 30
```

```
## 13:49:29 Building Annoy index with metric = cosine, n_trees = 50
```

```
## 0%   10   20   30   40   50   60   70   80   90   100%
```

```
## [----|----|----|----|----|----|----|----|----|----|
```

```
## **************************************************|
## 13:49:30 Writing NN index file to temp file /var/folders/9w/_lzbttfd7nv1_y4jd_50rlmj7j_gxg/T//RtmpJyR1ZK/file1
3cbb698e4e49
## 13:49:30 Searching Annoy index using 1 thread, search_k = 3000
## 13:49:32 Annoy recall = 100%
## 13:49:32 Commencing smooth kNN distance calibration using 1 thread
## 13:49:33 Initializing from normalized Laplacian + noise
## 13:49:33 Commencing optimization for 500 epochs, with 259982 positive edges
## 13:49:41 Optimization finished
```

```
#highlight GFP positive male dCA2 population
p1 <- DimPlot(object = dCA2.integrated, reduction = "umap", group.by = "protocol")
plot_grid(p1)
```

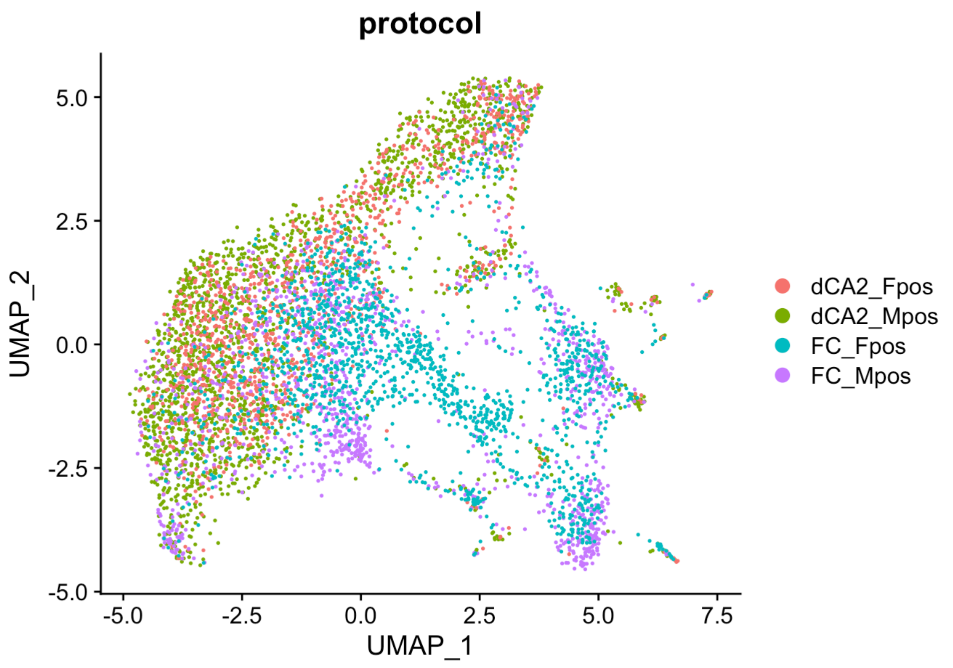


```
ggsave(file="/Users/lees44/R_project/FC_vs_dCA2_comparison_vector_integration_042821/UMAP.pdf",width=10,height=1
0)
```

```
p1<-DimPlot(dCA2.integrated, reduction = "umap", split.by = "protocol")
plot_grid(p1)
```

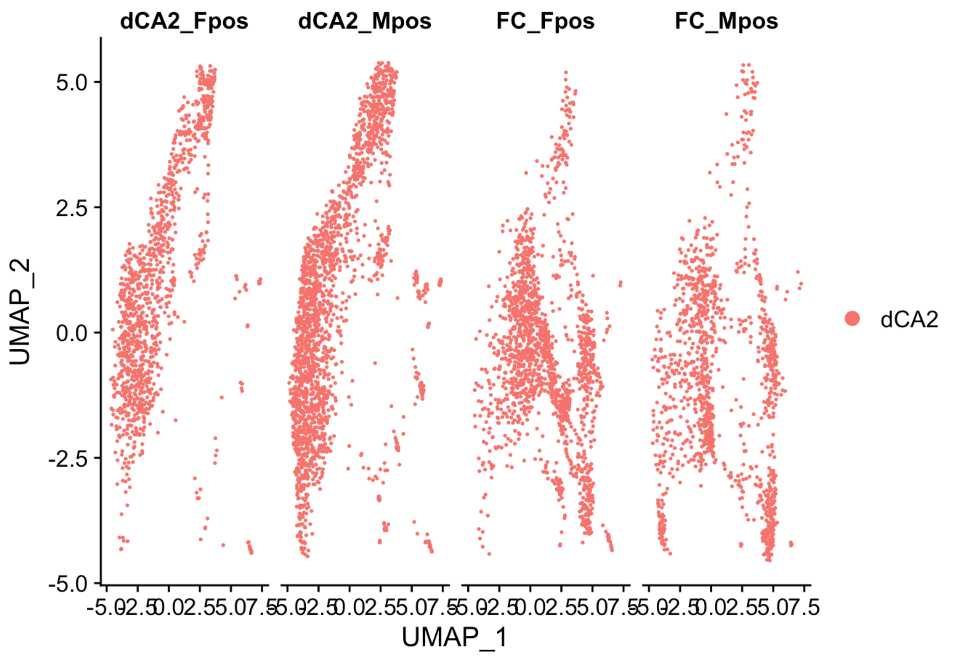


```
ggsave(file="/Users/lees44/R_project/FC_vs_dCA2_comparison_vector_integration_042821/UMAP_split.pdf",width=10,hei
ght=10)
```

(Shared) Nearest-neighbor graph construction.Computes the k.param nearest neighbors for a given dataset. Can also optionally (via compute.SNN), construct a shared nearest neighbor graph by calculating the neighborhood overlap (Jaccard index) between every cell and its k.param nearest neighbors.

```
#integrated analysis
dCA2.integrated <- FindNeighbors(object = dCA2.integrated)
```

```
## Computing nearest neighbor graph
```

```
## Computing SNN
```

find clusters within the groups. Identify clusters of cells by a shared nearest neighbor (SNN) modularity optimization based clustering algorithm. First calculate k-nearest neighbors and construct the SNN graph. Then optimize the modularity function to determine clusters.

```
dCA2.integrated <- FindClusters(dCA2.integrated, resolution = 0.3, dims.use = 1:30)
```

```
## Warning: The following arguments are not used: dims.use
```

```
## Suggested parameter: dims instead of dims.use
```

```
## Warning: The following arguments are not used: dims.use
```

```
## Suggested parameter: dims instead of dims.use
```

```
## Modularity Optimizer version 1.3.0 by Ludo Waltman and Nees Jan van Eck
##
## Number of nodes: 5928
## Number of edges: 192862
##
## Running Louvain algorithm...
## Maximum modularity in 10 random starts: 0.8578
## Number of communities: 9
## Elapsed time: 0 seconds
```

```
#clustering by population
p1<-DimPlot(object = dCA2.integrated, reduction = "umap", group.by = "integrated_snn_res.0.3", label = TRUE, repe
l = TRUE)
plot_grid(p1)
```

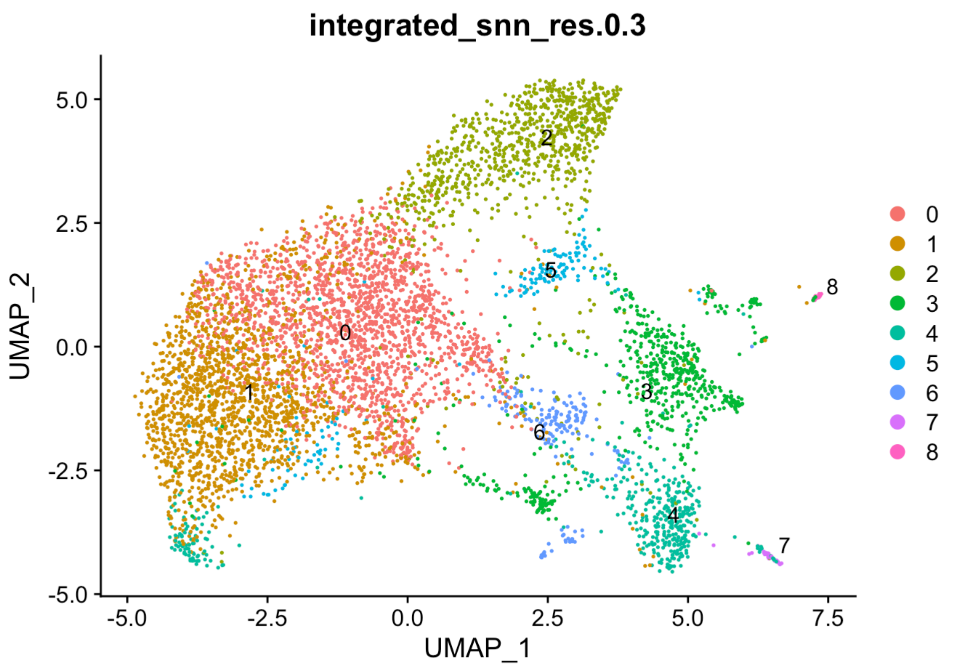


```
ggsave(file="/Users/lees44/R_project/FC_vs_dCA2_comparison_vector_integration_042821/clustering.pdf",width=10,hei
ght=10)
```

```
#clustering by population
p1<-DimPlot(object = dCA2.integrated, reduction = "umap", group.by = "integrated_snn_res.0.3", label = TRUE, repe
l = TRUE, split.by = "protocol")
plot_grid(p1)
```

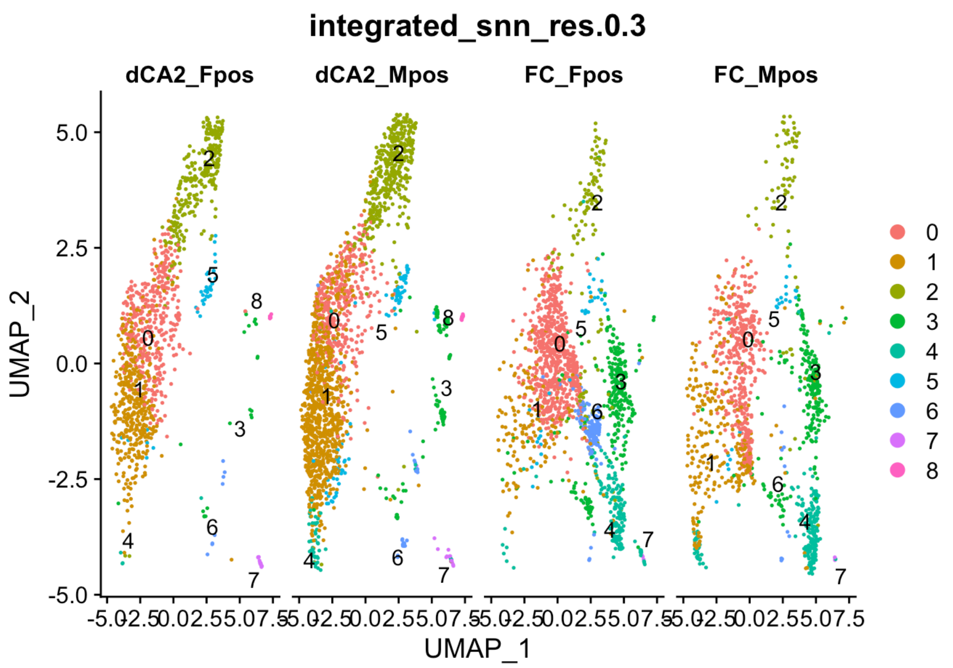


```
ggsave(file="/Users/lees44/R_project/FC_vs_dCA2_comparison_vector_integration_042821/clustering_split.pdf",width=
10,height=10)
```

#######################cluster tree code################################################################################### Building a cluster tree for the clusters

```
install.packages('ape')
```

```
## Installing package into '/Users/lees44/Library/R/4.0/library'
## (as 'lib' is unspecified)
```

```
## Error in contrib.url(repos, "source"): trying to use CRAN without setting a mirror
```

```
library('ape')
```

```
#cluster tree to go with the violin plot based on cell type
pdf(file="/Users/lees44/R_project/FC_vs_dCA2_comparison_vector_integration_042821/cluster_tree.pdf",width=20,heig
ht=8,paper='special')
#https://www.rdocumentation.org/packages/ape/versions/5.2/topics/plot.phylo
dCA2.integrated <- BuildClusterTree(dCA2.integrated, verbose = FALSE, reorder = FALSE)
plot(dCA2.integrated@tools$BuildClusterTree, type = "phylogram", edge.color = "black", edge.width = 4,edge.lty =
1, srt = 0, label.offset = 10, direction = "downwards", tip.color = "black")
dev.off()
```

```
## quartz_off_screen
##                 2
```

##########################doublet code###################################################################################

Remove doublet before sample integration.

```
install.packages("scDblfinder")
```

```
## Installing package into '/Users/lees44/Library/R/4.0/library'
## (as 'lib' is unspecified)
```

```
## Error in contrib.url(repos, "source"): trying to use CRAN without setting a mirror
```

converting seurat object to sce object https://satijalab.org/seurat/archive/v3.0/conversion_vignette.html (https://satijalab.org/seurat/archive/v3.0/conversion_vignette.html)

Doublet removal

```
BiocManager::install("scDblFinder")
```

```
## Bioconductor version 3.12 (BiocManager 1.30.14), R 4.0.4 (2021-02-15)
```

```
## Warning: package(s) not installed when version(s) same as current; use `force = TRUE` to
##   re-install: 'scDblFinder'
```

```
## Installation paths not writeable, unable to update packages
##   path: /Library/Frameworks/R.framework/Versions/4.0/Resources/library
##   packages:
##     boot, class, cluster, KernSmooth, lattice, MASS, mgcv, nlme, nnet, spatial,
##     survival
```

```
## Old packages: 'backports', 'beeswarm', 'BiocManager', 'blob', 'broom', 'bslib',
##   'cachem', 'cli', 'colorspace', 'conquer', 'cpp11', 'crayon', 'crosstalk',
##   'curl', 'data.table', 'deldir', 'desc', 'diffobj', 'digest', 'dplyr',
##   'fansi', 'fitdistrplus', 'formatR', 'future', 'future.apply', 'gargle',
##   'generics', 'ggplot2', 'glue', 'goftest', 'googledrive', 'googlesheets4',
##   'gtools', 'haven', 'hms', 'htmltools', 'htmlwidgets', 'httpuv', 'igraph',
##   'isoband', 'knitr', 'later', 'leiden', 'lifecycle', 'lmtest', 'lubridate',
##   'Matrix', 'matrixStats', 'metap', 'mime', 'mvtnorm', 'openssl', 'parallelly',
##   'pbapply', 'pillar', 'pkgload', 'plotly', 'plotrix', 'quantreg', 'R6',
##   'rbibutils', 'Rcpp', 'RcppAnnoy', 'RcppArmadillo', 'RCurl', 'Rdpack',
##   'readr', 'reprex', 'reticulate', 'rJava', 'rlang', 'rmarkdown', 'roxygen2',
##   'rvest', 'sandwich', 'sass', 'Seurat', 'SeuratObject', 'shiny', 'sitmo',
##   'snow', 'spatstat.core', 'spatstat.geom', 'spatstat.utils', 'statmod',
##   'stringi', 'testthat', 'TH.data', 'tibble', 'tidyr', 'tinytex', 'utf8',
##   'uuid', 'viridis', 'waldo', 'xfun', 'xgboost', 'zip'
```

```
library(scDblFinder)
library(scater)
```

```
## Loading required package: SingleCellExperiment
```

```
## Loading required package: SummarizedExperiment
```

```
## Loading required package: MatrixGenerics
```

```
## Loading required package: matrixStats
```

```
##
## Attaching package: 'matrixStats'
```

```
## The following object is masked from 'package:dplyr':
##
##     count
```

```
##
## Attaching package: 'MatrixGenerics'
```

```
## The following objects are masked from 'package:matrixStats':
##
##     colAlls, colAnyNAs, colAnys, colAvgsPerRowSet, colCollapse,
##     colCounts, colCummaxs, colCummins, colCumprods, colCumsums,
##     colDiffs, colIQRDiffs, colIQRs, colLogSumExps, colMadDiffs,
##     colMads, colMaxs, colMeans2, colMedians, colMins, colOrderStats,
##     colProds, colQuantiles, colRanges, colRanks, colSdDiffs, colSds,
##     colSums2, colTabulates, colVarDiffs, colVars, colWeightedMads,
##     colWeightedMeans, colWeightedMedians, colWeightedSds,
##     colWeightedVars, rowAlls, rowAnyNAs, rowAnys, rowAvgsPerColSet,
##     rowCollapse, rowCounts, rowCummaxs, rowCummins, rowCumprods,
##     rowCumsums, rowDiffs, rowIQRDiffs, rowIQRs, rowLogSumExps,
##     rowMadDiffs, rowMads, rowMaxs, rowMeans2, rowMedians, rowMins,
##     rowOrderStats, rowProds, rowQuantiles, rowRanges, rowRanks,
##     rowSdDiffs, rowSds, rowSums2, rowTabulates, rowVarDiffs, rowVars,
##     rowWeightedMads, rowWeightedMeans, rowWeightedMedians,
##     rowWeightedSds, rowWeightedVars
```

```
## Loading required package: GenomicRanges
```

```
## Loading required package: stats4
```

```
## Loading required package: BiocGenerics
```

```
## Warning: package 'BiocGenerics' was built under R version 4.0.5
```

```
## Loading required package: parallel
```

```
##
## Attaching package: 'BiocGenerics'
```

```
## The following objects are masked from 'package:parallel':
##
##     clusterApply, clusterApplyLB, clusterCall, clusterEvalQ,
##     clusterExport, clusterMap, parApply, parCapply, parLapply,
##     parLapplyLB, parRapply, parSapply, parSapplyLB
```

```
## The following objects are masked from 'package:dplyr':
##
##     combine, intersect, setdiff, union
```

```
## The following objects are masked from 'package:stats':
##
##     IQR, mad, sd, var, xtabs
```

```
## The following objects are masked from 'package:base':
##
##     anyDuplicated, append, as.data.frame, basename, cbind, colnames,
##     dirname, do.call, duplicated, eval, evalq, Filter, Find, get, grep,
##     grepl, intersect, is.unsorted, lapply, Map, mapply, match, mget,
##     order, paste, pmax, pmax.int, pmin, pmin.int, Position, rank,
##     rbind, Reduce, rownames, sapply, setdiff, sort, table, tapply,
##     union, unique, unsplit, which.max, which.min
```

```
## Loading required package: S4Vectors
```

```
##
## Attaching package: 'S4Vectors'
```

```
## The following objects are masked from 'package:dplyr':
##
##     first, rename
```

```
## The following object is masked from 'package:base':
##
##     expand.grid
```

```
## Loading required package: IRanges
```

```
##
## Attaching package: 'IRanges'
```

```
## The following objects are masked from 'package:dplyr':
##
##     collapse, desc, slice
```

```
## Loading required package: GenomeInfoDb
```

```
## Warning: package 'GenomeInfoDb' was built under R version 4.0.5
```

```
## Loading required package: Biobase
```

```
## Welcome to Bioconductor
##
##     Vignettes contain introductory material; view with
##     'browseVignettes()'. To cite Bioconductor, see
##     'citation("Biobase")', and for packages 'citation("pkgname")'.
```

```
##
## Attaching package: 'Biobase'
```

```
## The following object is masked from 'package:MatrixGenerics':
##
##     rowMedians
```

```
## The following objects are masked from 'package:matrixStats':
##
##     anyMissing, rowMedians
```

```
##
## Attaching package: 'SummarizedExperiment'
```

```
## The following object is masked from 'package:SeuratObject':
##
##     Assays
```

```
## The following object is masked from 'package:Seurat':
##
##     Assays
```

```
library(SingleCellExperiment)
library(Seurat)
library(scDblFinder)
library(scuttle)
```

```
# prepare object to inspect for doublet clusters
DefaultAssay(dCA2.integrated) <- "RNA"
sce.object <- as.SingleCellExperiment(dCA2.integrated)
colLabels(sce.object) <- colData(sce.object)$ident
dbl.out <- findDoubletClusters(sce.object,colLabels(sce.object))
```

```
# outlier clusters by number of significant genes (num.de)
rownames(dbl.out)[isOutlier(dbl.out$num.de, log=TRUE, type="lower")]
```

```
## [1] "0"
```

```
# outlier clusters by number of significant genes (num.de)
rownames(dbl.out)[dbl.out$lib.size1 < 1 & dbl.out$lib.size2 < 1]
```

```
## [1] "3" "1" "2"
```

```
# be sure to reset your defualt assay back
DefaultAssay(dCA2.integrated) <- "integrated"
```

################################################dCA2 Markers#####################################################################

Feature plot highlighting dCA2 markers in all population Expression pots in UMAP space illustrating normalized expression values of dCA2 markers

“CREREV38354884”, “GFP”,“Rgs14”, “Avpr1b”, “Amigo2”, “Map3k15”, “Actn2”, “Ptpn5”, “Cacng5”, “Oxtr”, “Pcp4”

```
#Feature plot highlighting dCA2 markers in all population
p1<-FeaturePlot(object =dCA2.integrated, features = c("Rgs14"), min.cutoff = 0)
plot_grid(p1)
```

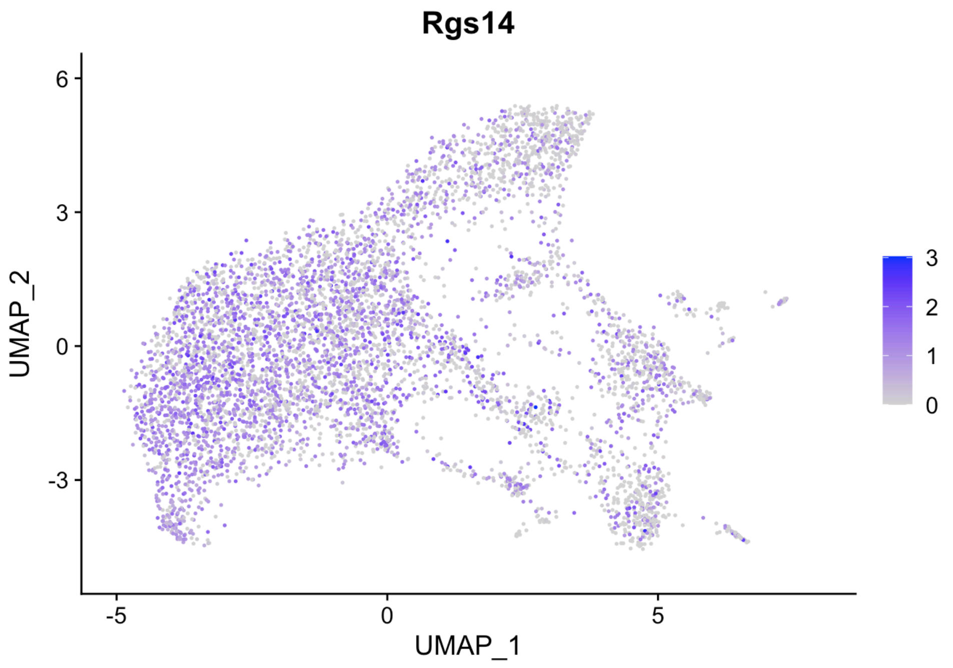


```
ggsave(file="/Users/lees44/R_project/FC_vs_dCA2_comparison_vector_integration_042821/Rgs14.pdf",width=10,height=1
0)
```

```
#Feature plot highlighting dCA2 markers in all population
p1<-FeaturePlot(object =dCA2.integrated, features = c("CREREV38354884"), min.cutoff = 0, max.cutoff = 1.5 )
plot_grid(p1)
```

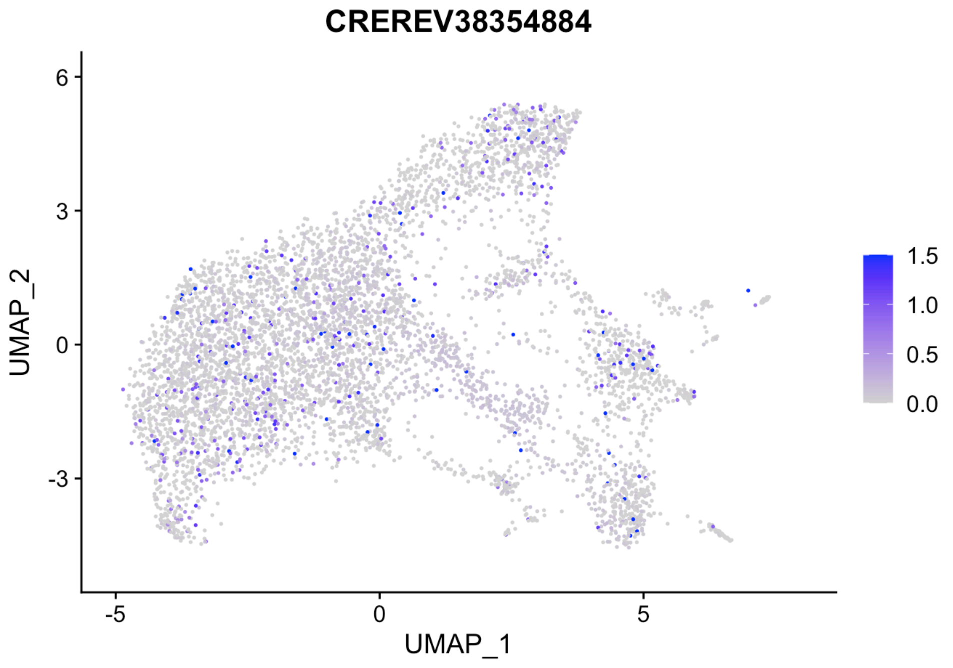

```
ggsave(file="/Users/lees44/R_project/FC_vs_dCA2_comparison_vector_integration_042821/CREREV38354884.pdf",width=10,
height=10)
```

```
#Feature plot highlighting dCA2 markers in all population
p1<-FeaturePlot(object =dCA2.integrated, features = c("GFP"), min.cutoff = 0, max.cutoff = 3)
```

```
## Warning: Could not find GFP in the default search locations, found in RNA assay
## instead
```

```
plot_grid(p1)
```

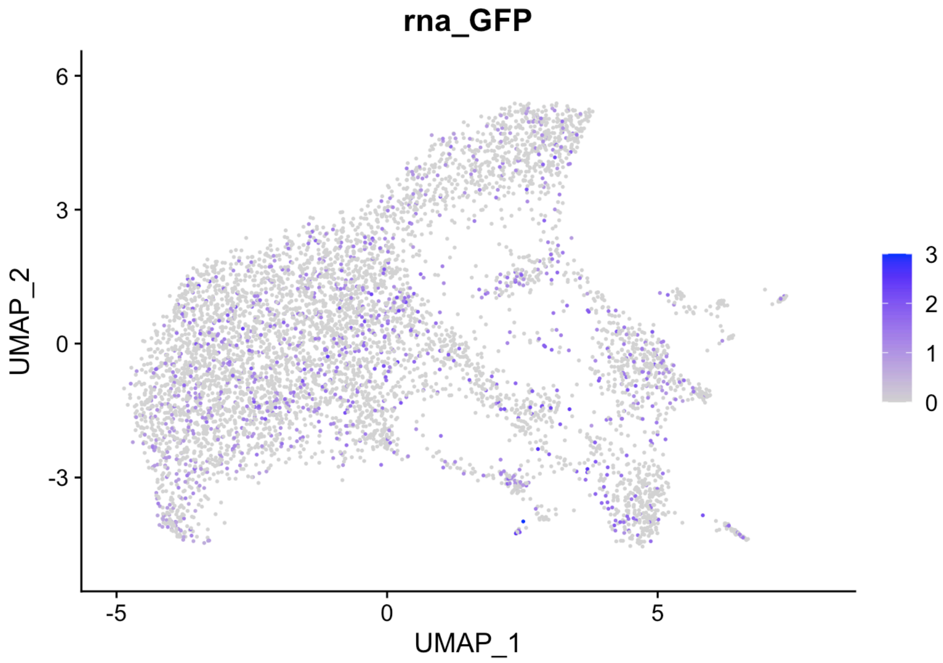


```
ggsave(file="/Users/lees44/R_project/FC_vs_dCA2_comparison_vector_integration_042821/GFP.pdf",width=10,height=10)
```

“CREREV38354884”, “GFP”,“Rgs14”, “Avpr1b”, “Amigo2”, “Map3k15”, “Actn2”, “Ptpn5”, “Cacng5”, “Oxtr”, “Pcp4”

```
#Feature plot highlighting dCA2 markers in all population
p1<-FeaturePlot(object =dCA2.integrated, features = c("Avpr1b"), min.cutoff = 0, max.cutoff = 1.5)
```

```
## Warning: Could not find Avpr1b in the default search locations, found in RNA
## assay instead
```

```
plot_grid(p1)
```

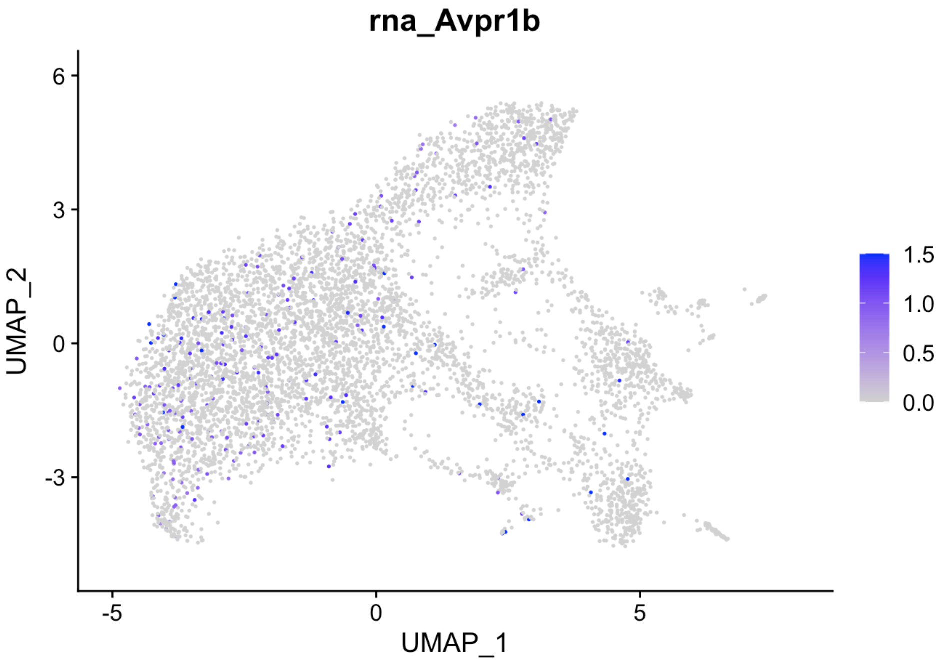


```
ggsave(file="/Users/lees44/R_project/FC_vs_dCA2_comparison_vector_integration_042821/Avpr1b.pdf",width=10,height=10)
```

```
#Feature plot highlighting dCA2 markers in all population
p1<-FeaturePlot(object =dCA2.integrated, features = c("Amigo2"), min.cutoff = 0, max.cutoff = 3)
plot_grid(p1)
```

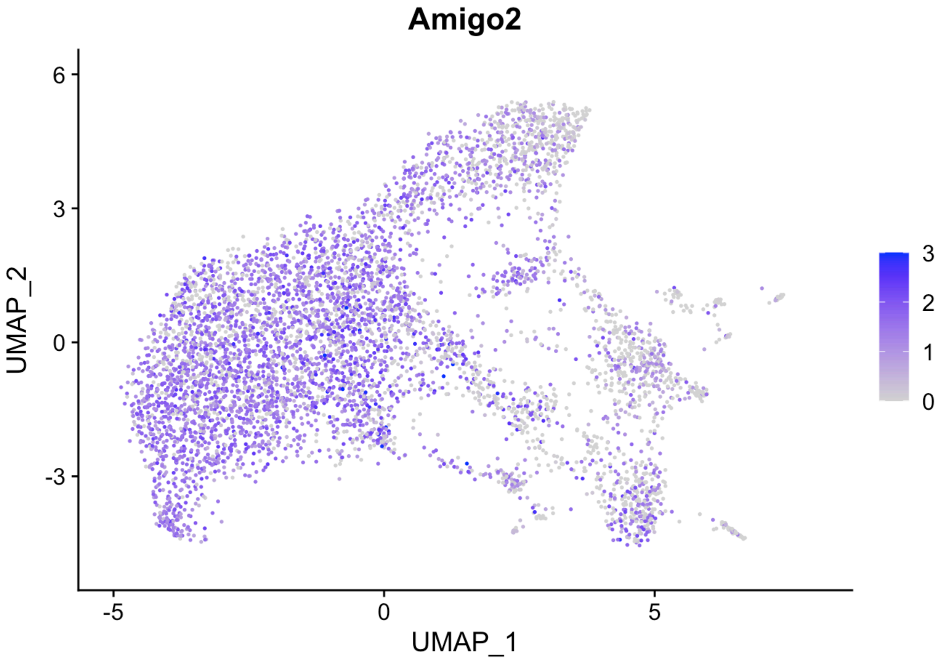


```
ggsave(file="/Users/lees44/R_project/FC_vs_dCA2_comparison_vector_integration_042821/Amigo2.pdf",width=10,height=10)
```

“CREREV38354884”, “GFP”,“Rgs14”, “Avpr1b”, “Amigo2”, “Map3k15”, “Actn2”, “Ptpn5”, “Cacng5”, “Oxtr”, “Pcp4”

```
#Feature plot highlighting dCA2 markers in all population
p1<-FeaturePlot(object =dCA2.integrated, features = c("Map3k15"), min.cutoff = 0, max.cutoff = 3)
plot_grid(p1)
```

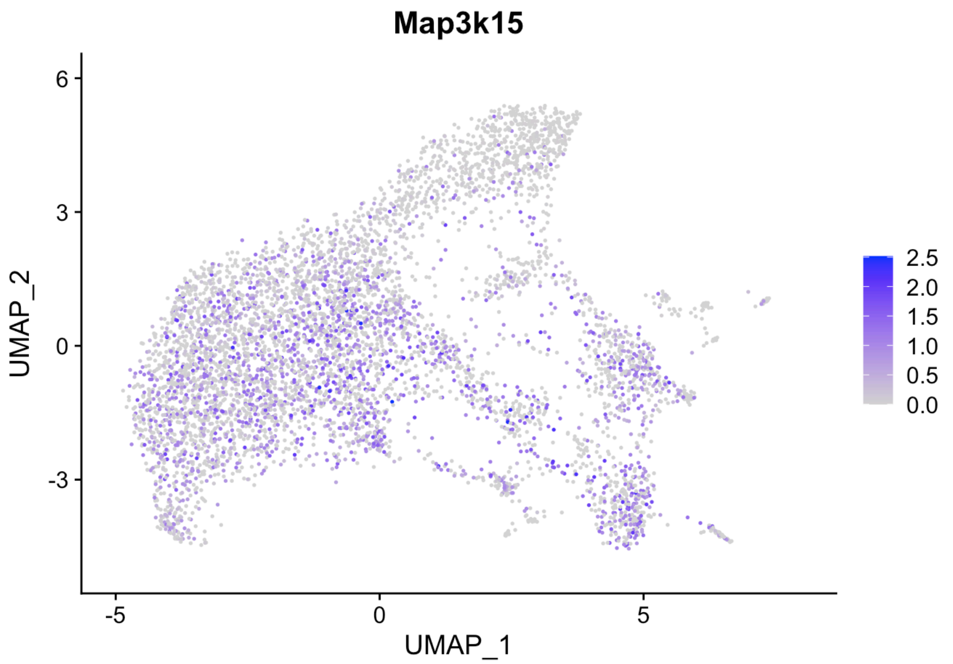


```
ggsave(file="/Users/lees44/R_project/FC_vs_dCA2_comparison_vector_integration_042821/Map3k15.pdf",width=10,height
=10)
```

```
#Feature plot highlighting dCA2 markers in all population
p1<-FeaturePlot(object =dCA2.integrated, features = c("Actn2"), min.cutoff = 0, max.cutoff = 3)
```

```
## Warning: Could not find Actn2 in the default search locations, found in RNA
## assay instead
```

```
plot_grid(p1)
```

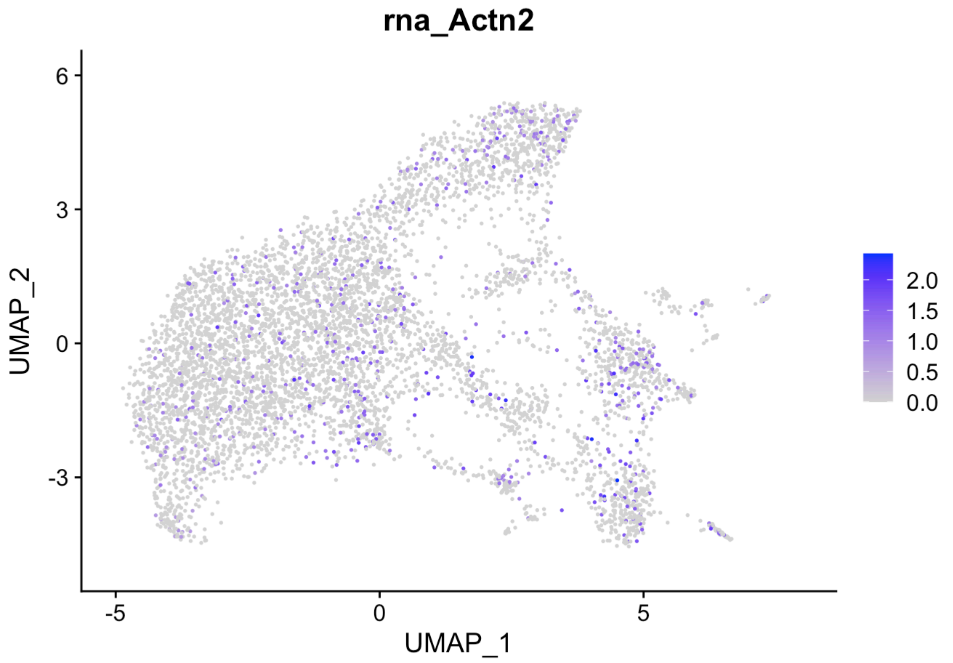


```
ggsave(file="/Users/lees44/R_project/FC_vs_dCA2_comparison_vector_integration_042821/Actn2.pdf",width=10,height=1
0)
```

“CREREV38354884”, “GFP”,“Rgs14”, “Avpr1b”, “Amigo2”, “Map3k15”, “Actn2”, “Ptpn5”, “Cacng5”, “Oxtr”, “Pcp4”

```
#Feature plot highlighting dCA2 markers in all population
p1<-FeaturePlot(object =dCA2.integrated, features = c("Ptpn5"), min.cutoff = 0,max.cutoff = 3)
plot_grid(p1)
```

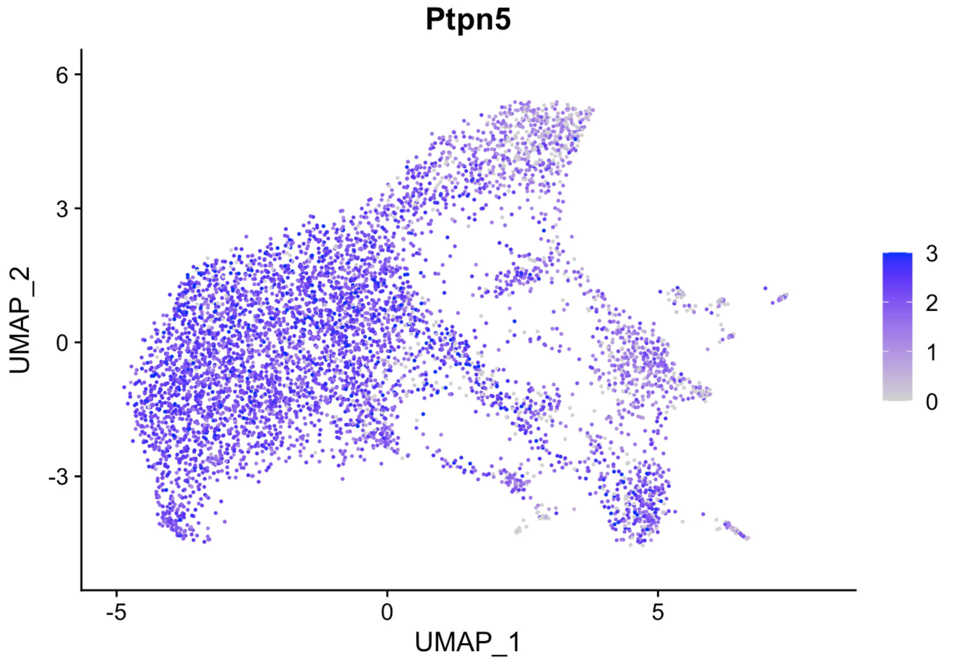


```
ggsave(file="/Users/lees44/R_project/FC_vs_dCA2_comparison_vector_integration_042821/Ptpn5.pdf",width=10,height=1
0)
```

```
#Feature plot highlighting dCA2 markers in all population
p1<-FeaturePlot(object =dCA2.integrated, features = c("Cacng5"), min.cutoff = 0, max.cutoff = 3)
plot_grid(p1)
```

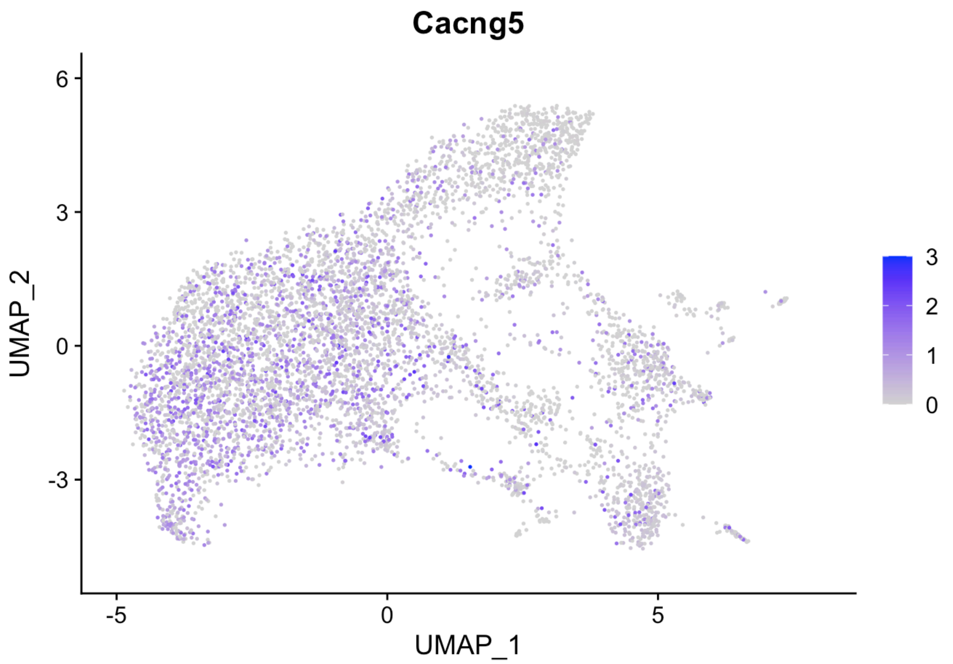


```
ggsave(file="/Users/lees44/R_project/FC_vs_dCA2_comparison_vector_integration_042821/Cacng5.pdf",width=10,height=
10)
```

```
#Feature plot highlighting dCA2 markers in all population
p1<-FeaturePlot(object =dCA2.integrated, features = c("Oxtr"), min.cutoff = 0, max.cutoff = 1.5)
```

```
## Warning: Could not find Oxtr in the default search locations, found in RNA assay
## instead
```

```
plot_grid(p1)
```

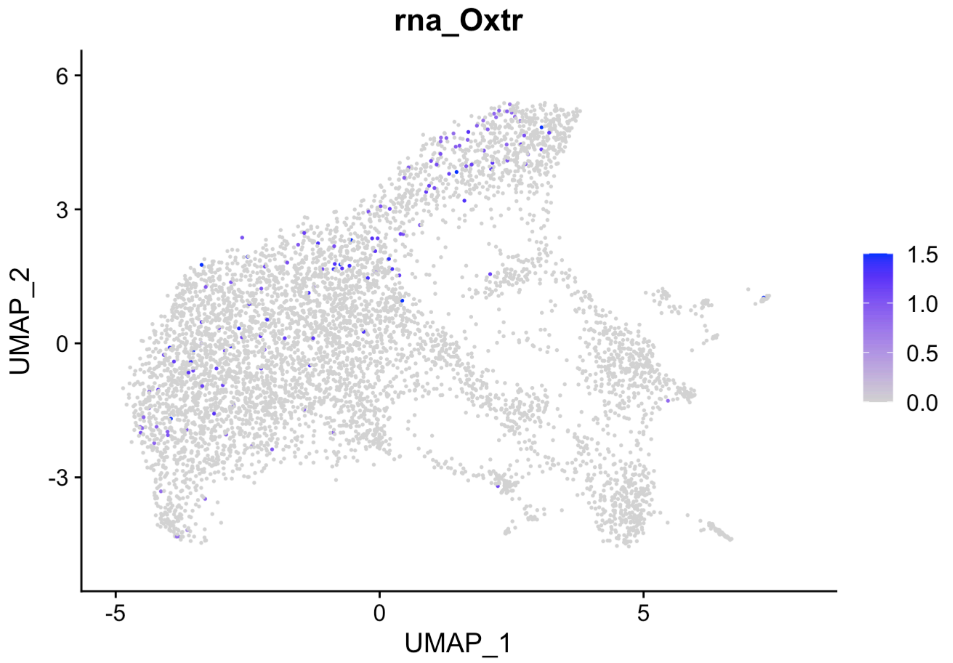


```
ggsave(file="/Users/lees44/R_project/FC_vs_dCA2_comparison_vector_integration_042821/Oxtr.pdf",width=10,height=10)
```

```
#Feature plot highlighting dCA2 markers in all population
p1<-FeaturePlot(object =dCA2.integrated, features = c("Pcp4"), min.cutoff = 0, max.cutoff = 3)
plot_grid(p1)
```

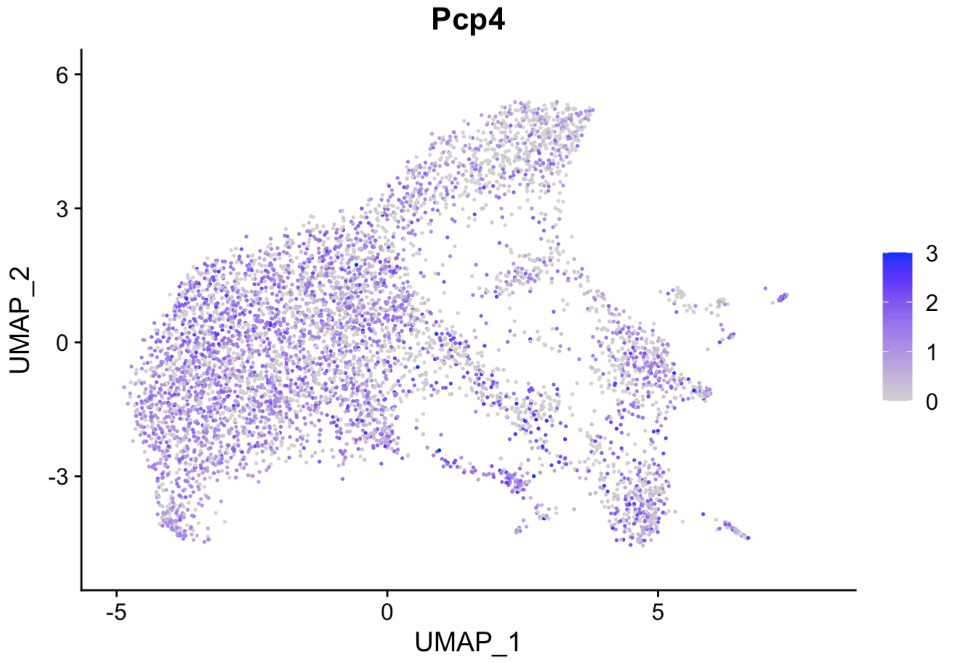


```
ggsave(file="/Users/lees44/R_project/FC_vs_dCA2_comparison_vector_integration_042821/Pcp4.pdf",width=10,height=10)
```

sodium channel

```
#Feature plot highlighting dCA2 markers in all population
p1<-FeaturePlot(object =dCA2.integrated, features = c("Scn1a"), min.cutoff = 0)
plot_grid(p1)
```

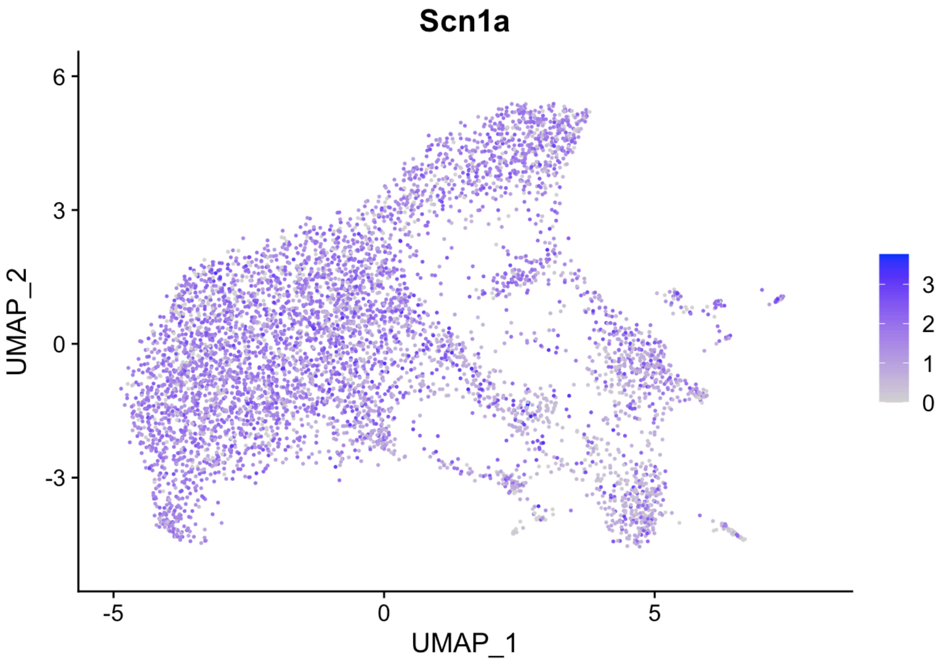


```
ggsave(file="/Users/lees44/R_project/FC_vs_dCA2_comparison_vector_integration_042821/SCN1A.pdf",width=10,height=1
0)
```

```
#Feature plot highlighting dCA2 markers in all population
p1<-FeaturePlot(object =dCA2.integrated, features = c("Scn2a"), min.cutoff = 0, max.cutoff = 3)
```

```
## Warning: Could not find Scn2a in the default search locations, found in RNA
## assay instead
```

```
plot_grid(p1)
```

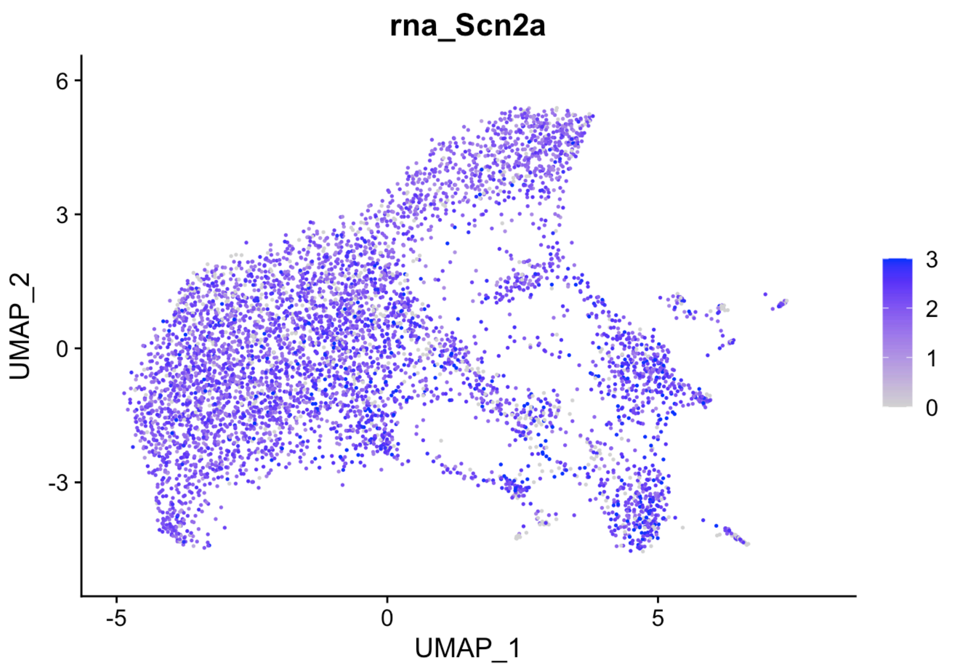


```
ggsave(file="/Users/lees44/R_project/FC_vs_dCA2_comparison_vector_integration_042821/SCN2A.pdf",width=10,height=1
0)
```

```
#Feature plot highlighting dCA2 markers in all population
p1<-FeaturePlot(object =dCA2.integrated, features = c("Scn3a"), min.cutoff = 0)
plot_grid(p1)
```

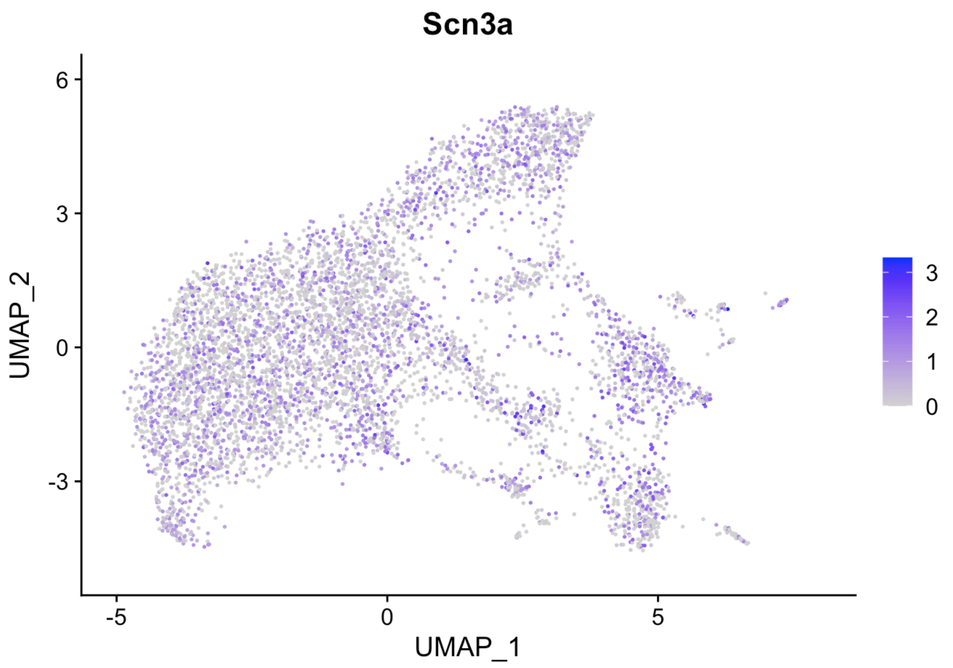


```
ggsave(file="/Users/lees44/R_project/FC_vs_dCA2_comparison_vector_integration_042821/SCN3A.pdf",width=10,height=1
0)
```

```
#Feature plot highlighting dCA2 markers in all population
p1<-FeaturePlot(object =dCA2.integrated, features = c("Scn4a"), min.cutoff = 0, max.cutoff = 3)
```

```
## Warning in FetchData(object = object, vars = c(dims, "ident", features), : The
## following requested variables were not found: Scn4a
```

```
## Error: None of the requested features were found: Scn4a in slot data
```

```
plot_grid(p1)
```

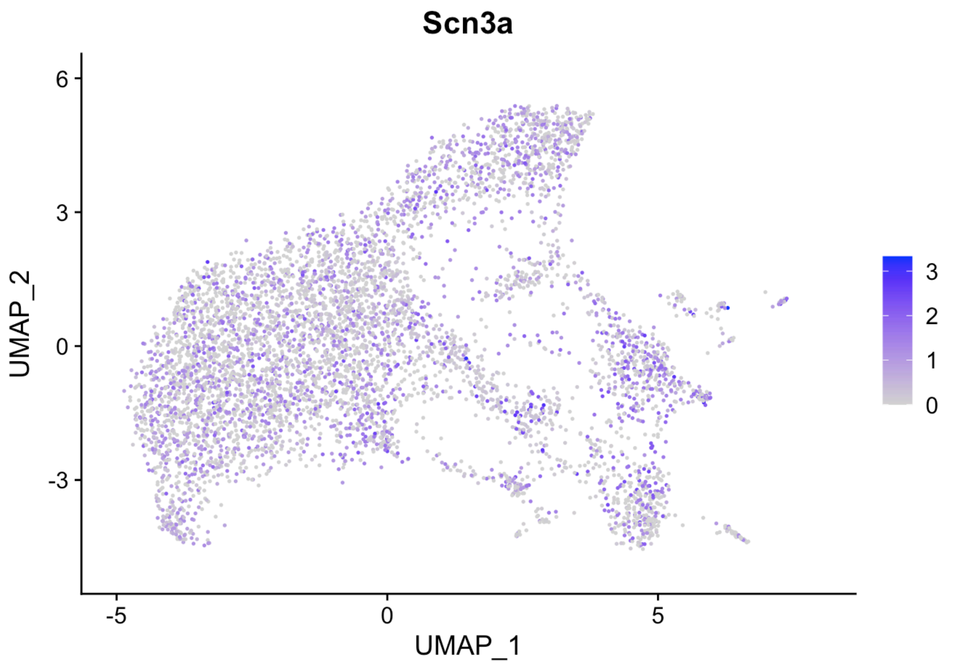


```
ggsave(file="/Users/lees44/R_project/FC_vs_dCA2_comparison_vector_integration_042821/SCN4A.pdf",width=10,height=1
0)
```

```
#Feature plot highlighting dCA2 markers in all population
p1<-FeaturePlot(object =dCA2.integrated, features = c("Scn8a"), min.cutoff = 0)
```

```
## Warning: Could not find Scn8a in the default search locations, found in RNA
## assay instead
```

```
plot_grid(p1)
```

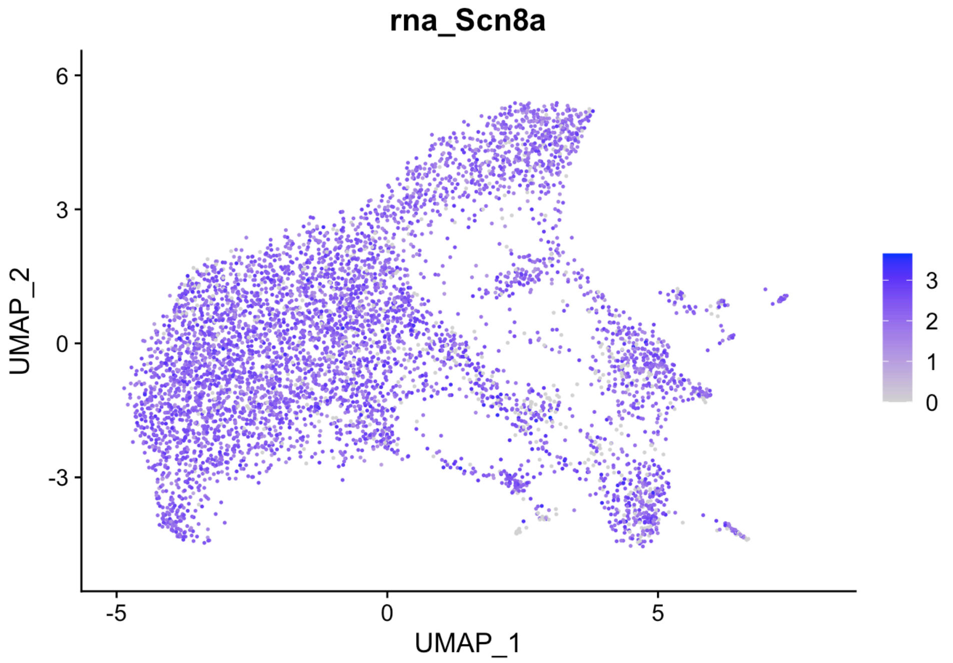


```
ggsave(file="/Users/lees44/R_project/FC_vs_dCA2_comparison_vector_integration_042821/SCN8A.pdf",width=10,height=1
0)
```

```
#Feature plot highlighting dCA2 markers in all population
p1<-FeaturePlot(object =dCA2.integrated, features = c("Scn9a"), min.cutoff = 0, max.cutoff = 3)
plot_grid(p1)
```

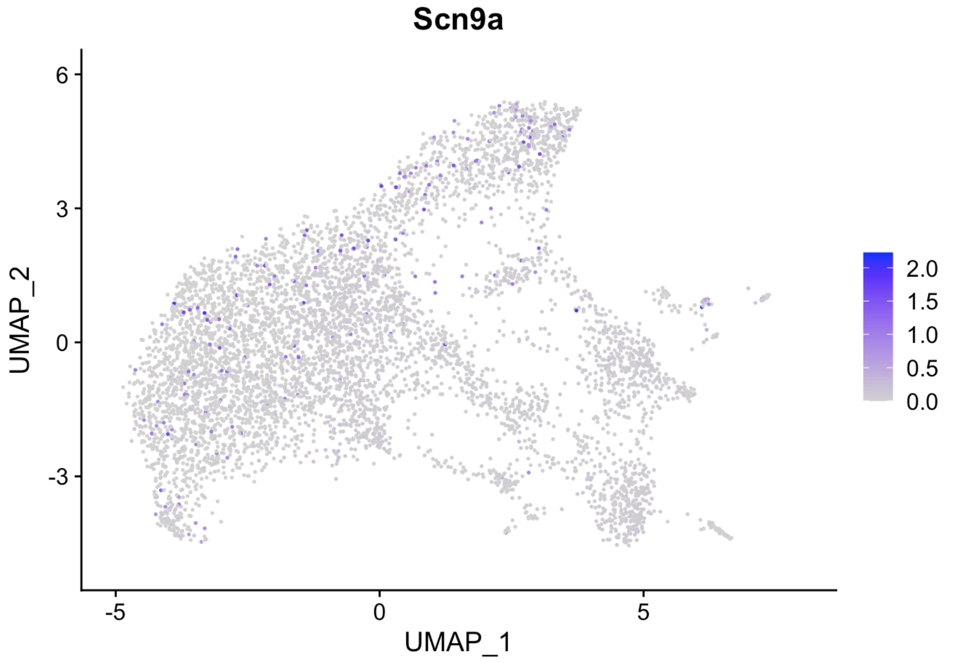


```
ggsave(file="/Users/lees44/R_project/FC_vs_dCA2_comparison_vector_integration_042821/SCN9A.pdf",width=10,height=1
0)
```

```
#Feature plot highlighting dCA2 markers in all population
p1<-FeaturePlot(object =dCA2.integrated, features = c("Scn5a"), min.cutoff = 0, max.cutoff = 3)
```

```
## Warning in FetchData(object = object, vars = c(dims, "ident", features), : The
## following requested variables were not found: Scn5a
```

```
## Error: None of the requested features were found: Scn5a in slot data
```

```
plot_grid(p1)
```

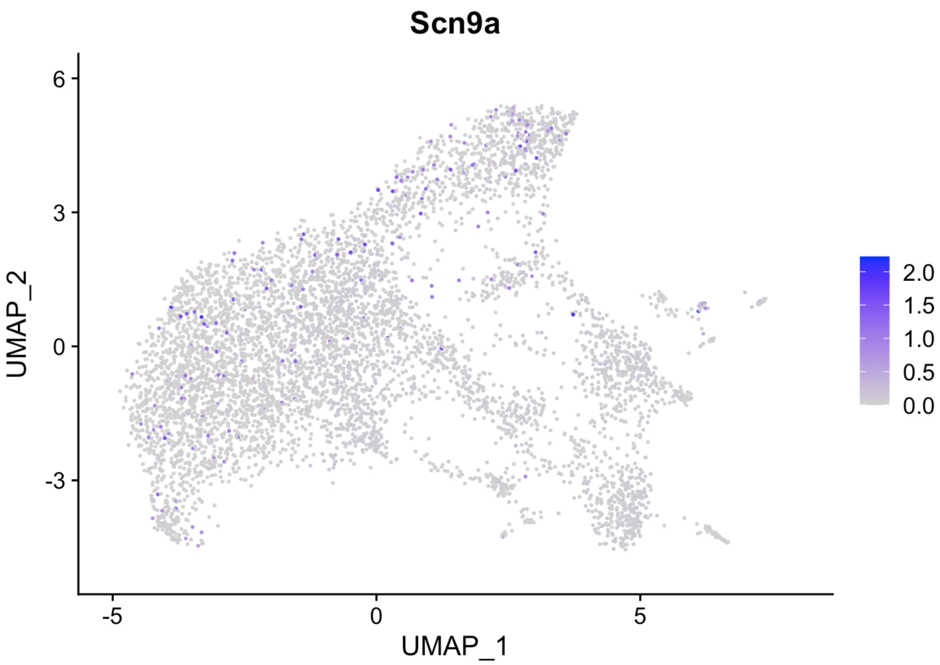


```
ggsave(file="/Users/lees44/R_project/FC_vs_dCA2_comparison_vector_integration_042821/SCN5A.pdf",width=10,height=1
0)
```

```
#Feature plot highlighting dCA2 markers in all population
p1<-FeaturePlot(object =dCA2.integrated, features = c("Scn10a"), min.cutoff = 0, max.cutoff = 3)
```

```
## Warning in FetchData(object = object, vars = c(dims, "ident", features), : The
## following requested variables were not found: Scn10a
```

```
## Error: None of the requested features were found: Scn10a in slot data
```

```
plot_grid(p1)
```

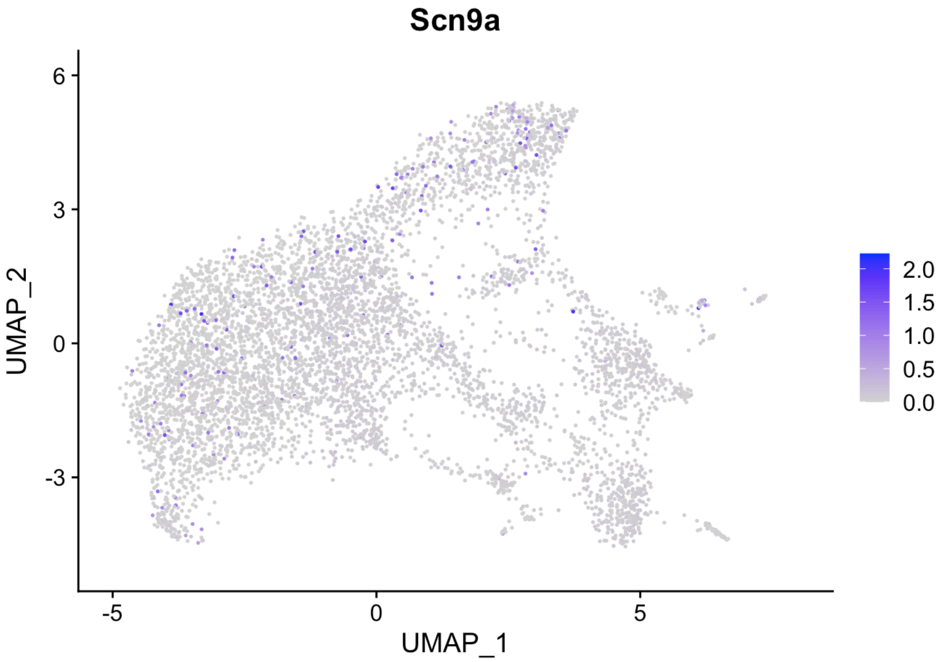


```
ggsave(file="/Users/lees44/R_project/FC_vs_dCA2_comparison_vector_integration_042821/SCN10A.pdf",width=10,height=
10)
```

```
#Feature plot highlighting dCA2 markers in all population
p1<-FeaturePlot(object =dCA2.integrated, features = c("Scn11a"), min.cutoff = 0, max.cutoff = 3)
```

```
## Warning in FetchData(object = object, vars = c(dims, "ident", features), : The
## following requested variables were not found: Scn11a
```

```
## Error: None of the requested features were found: Scn11a in slot data
```

```
plot_grid(p1)
```

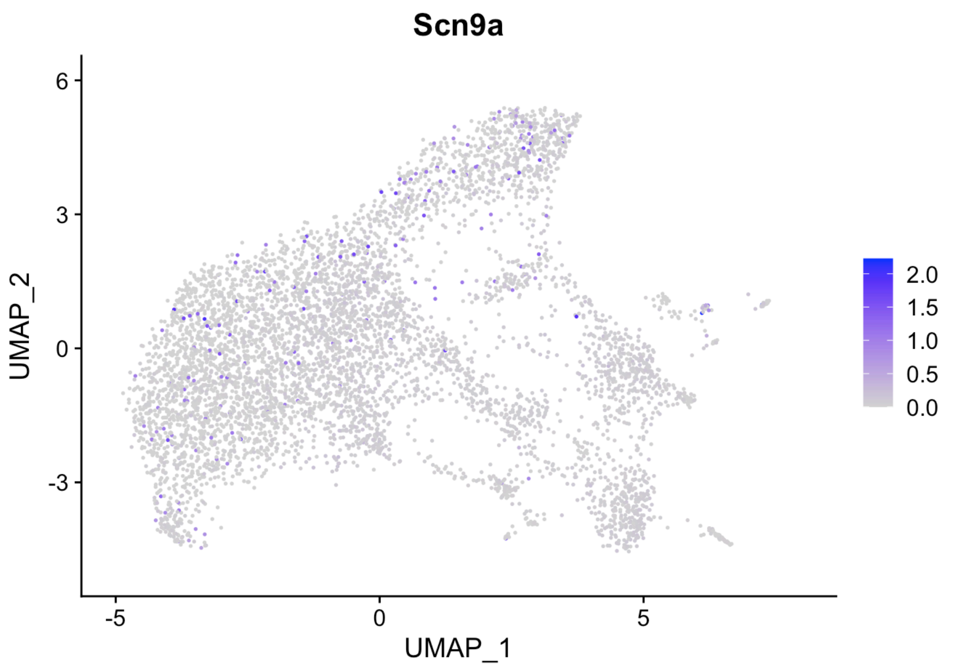


```
ggsave(file="/Users/lees44/R_project/FC_vs_dCA2_comparison_vector_integration_042821/SCN11A.pdf",width=10,height=10)
```

############################potassium channel#########################

```
#Feature plot highlighting potassium channel in all population
p1<-FeaturePlot(object =dCA2.integrated, features = c("Kcna1"), min.cutoff = 0)
plot_grid(p1)
```

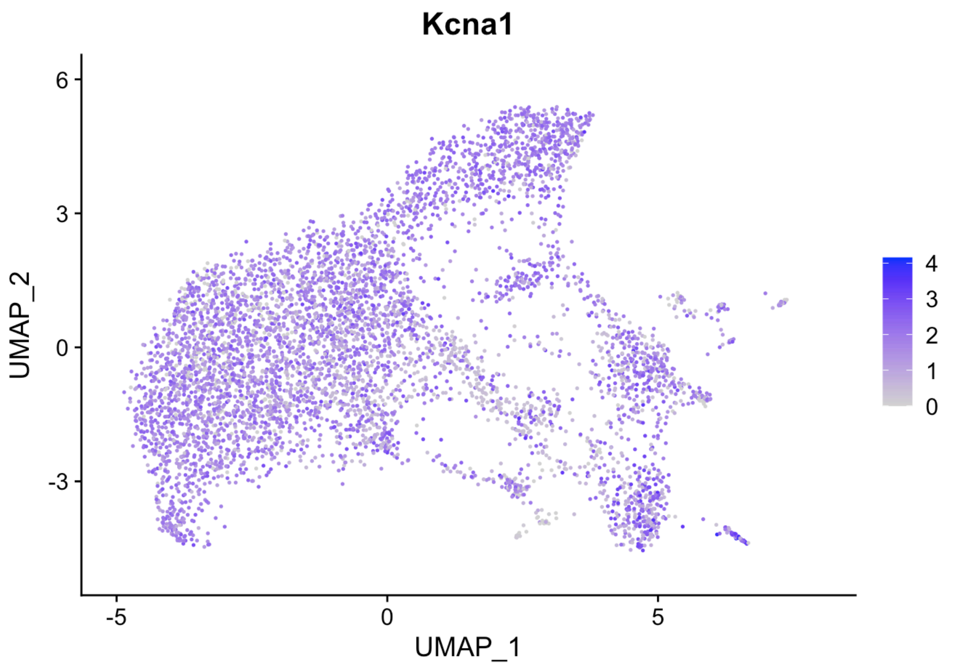


```
ggsave(file="/Users/lees44/R_project/FC_vs_dCA2_comparison_vector_integration_042821/Kcna1.pdf",width=10,height=10)
```

```
#Feature plot highlighting potassium channel in all population
p1<-FeaturePlot(object =dCA2.integrated, features = c("Kcna2"), min.cutoff = 0)
```

```
## Warning: Could not find Kcna2 in the default search locations, found in RNA
## assay instead
```

```
plot_grid(p1)
```

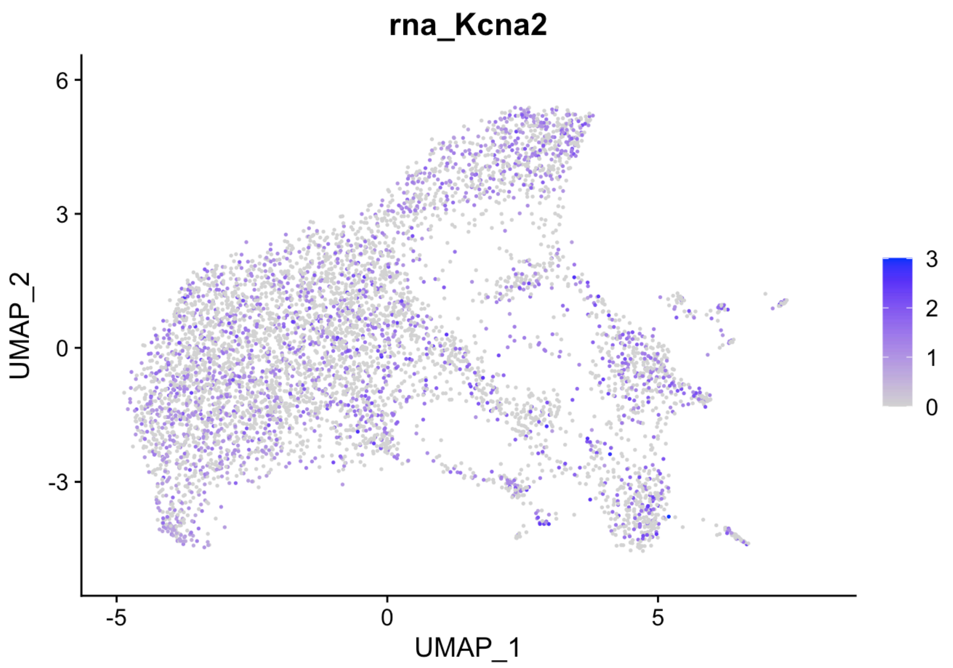


```
ggsave(file="/Users/lees44/R_project/FC_vs_dCA2_comparison_vector_integration_042821/Kcna2.pdf",width=10,height=1
0)
```

```
#Feature plot highlighting potassium channel in all population
p1<-FeaturePlot(object =dCA2.integrated, features = c("Kcna3"), min.cutoff = 0)
```

```
## Warning: Could not find Kcna3 in the default search locations, found in RNA
## assay instead
```

```
plot_grid(p1)
```

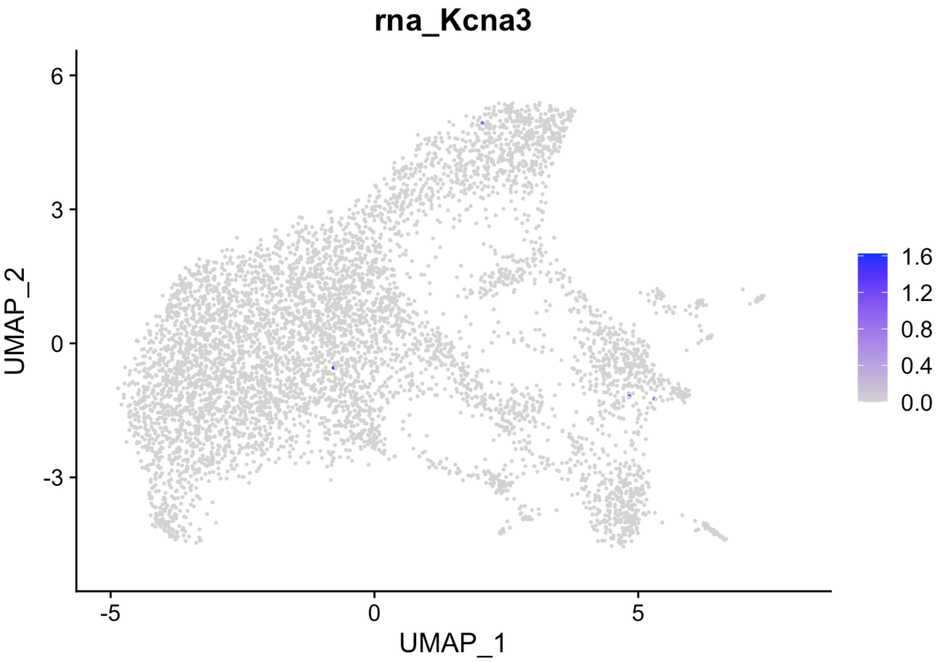


```
ggsave(file="/Users/lees44/R_project/FC_vs_dCA2_comparison_vector_integration_042821/Kcna3.pdf",width=10,height=1
0)
```

```
#Feature plot highlighting potassium channel in all population
p1<-FeaturePlot(object =dCA2.integrated, features = c("Kcna4"), min.cutoff = 0)
plot_grid(p1)
```

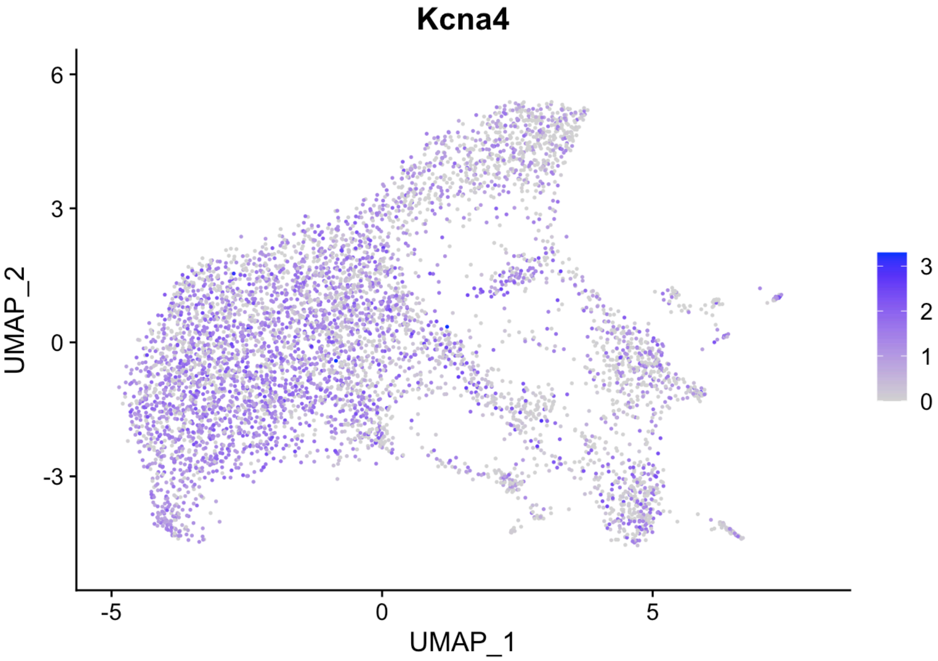


```
ggsave(file="/Users/lees44/R_project/FC_vs_dCA2_comparison_vector_integration_042821/Kcna4.pdf",width=10,height=1
0)
```

```
#Feature plot highlighting potassium channel in all population
p1<-FeaturePlot(object =dCA2.integrated, features = c("Kcna6"), min.cutoff = 0)
```

```
## Warning: Could not find Kcna6 in the default search locations, found in RNA
## assay instead
```

```
plot_grid(p1)
```

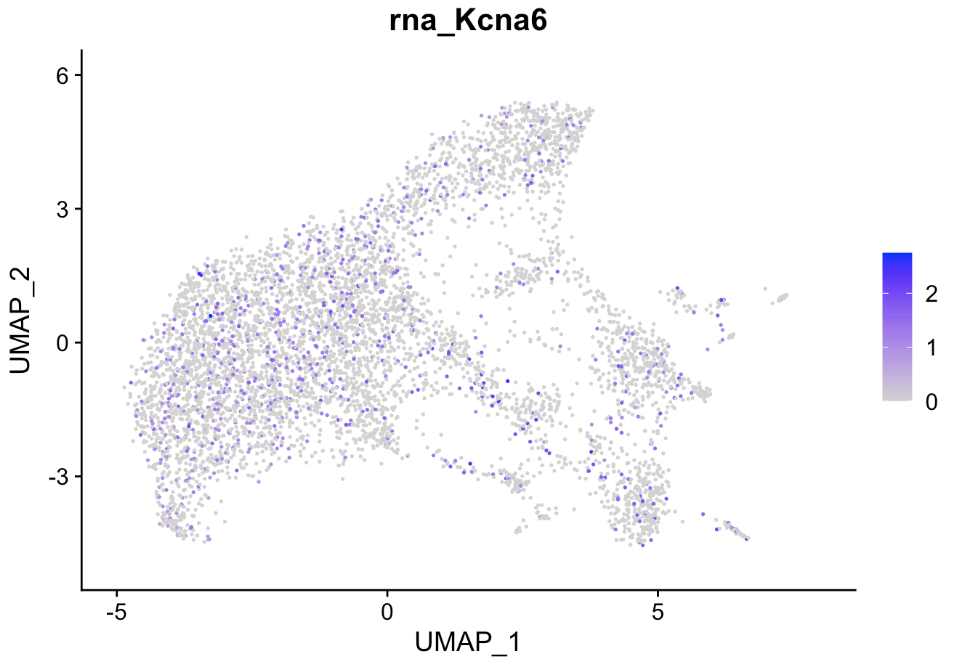


```
ggsave(file="/Users/lees44/R_project/FC_vs_dCA2_comparison_vector_integration_042821/Kcna6.pdf",width=10,height=1
0)
```

```
#Feature plot highlighting potassium channel in all population
p1<-FeaturePlot(object =dCA2.integrated, features = c("Kcnb1"), min.cutoff = 0)
plot_grid(p1)
```

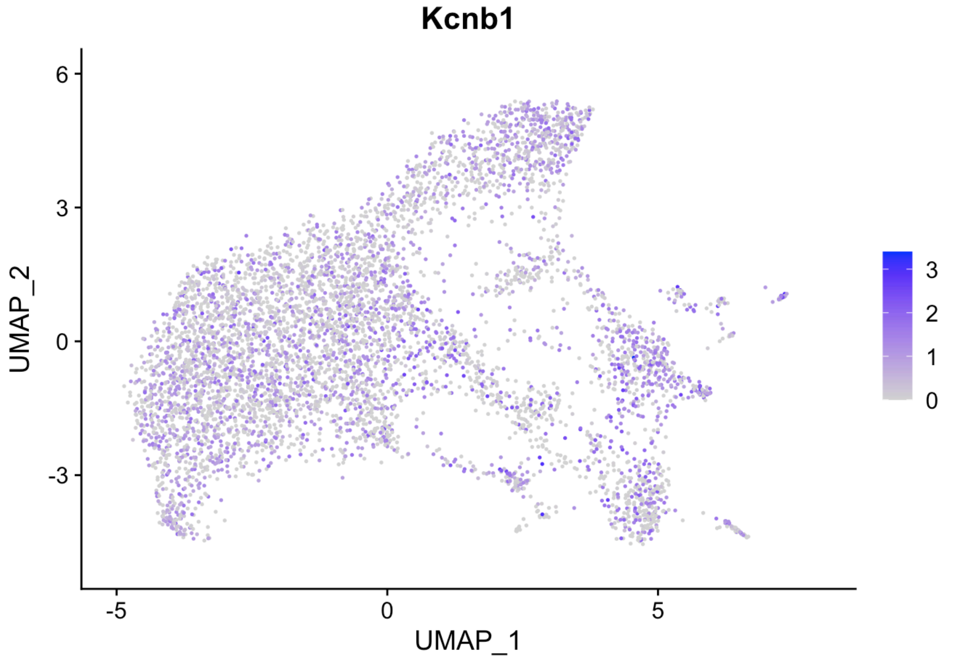

```
ggsave(file="/Users/lees44/R_project/FC_vs_dCA2_comparison_vector_integration_042821/Kcnb1.pdf",width=10,height=1
0)
```

```
#Feature plot highlighting potassium channel in all population
p1<-FeaturePlot(object =dCA2.integrated, features = c("Kcnb2"), min.cutoff = 0)
plot_grid(p1)
```

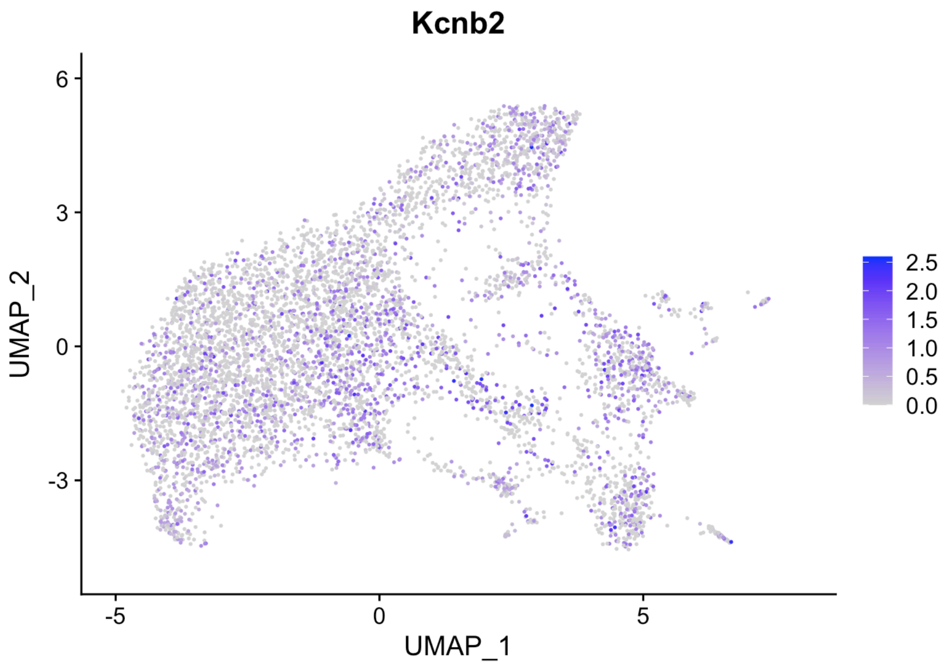


```
ggsave(file="/Users/lees44/R_project/FC_vs_dCA2_comparison_vector_integration_042821/Kcnb2.pdf",width=10,height=1
0)
```

```
#Feature plot highlighting potassium channel in all population
p1<-FeaturePlot(object =dCA2.integrated, features = c("Kcnc1"), min.cutoff = 0)
plot_grid(p1)
```

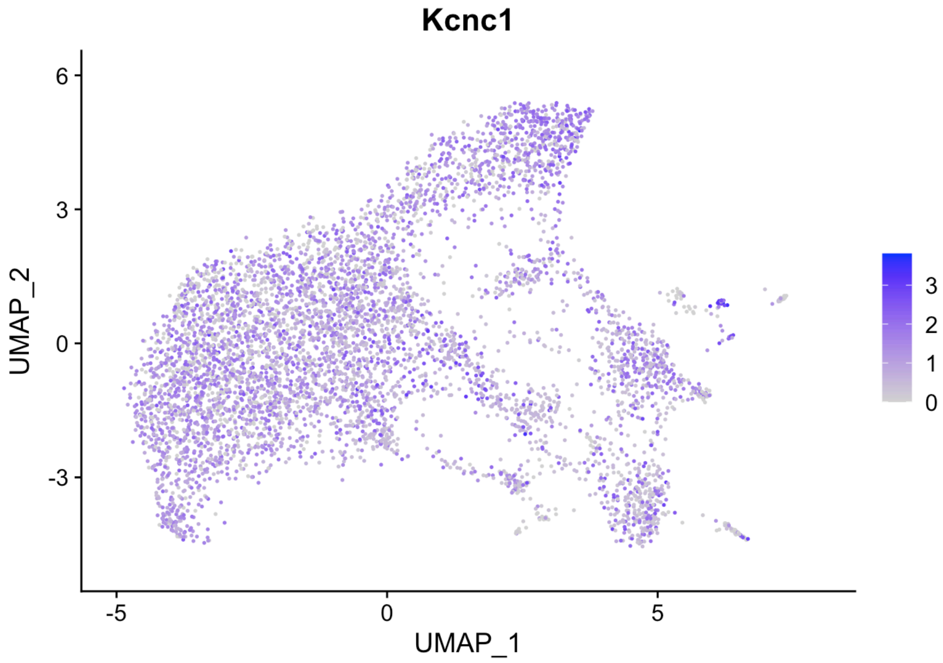


```
ggsave(file="/Users/lees44/R_project/FC_vs_dCA2_comparison_vector_integration_042821/Kcnc1.pdf",width=10,height=1
0)
```

```
#Feature plot highlighting potassium channel in all population
p1<-FeaturePlot(object =dCA2.integrated, features = c("Kcnc2"), min.cutoff = 0)
plot_grid(p1)
```

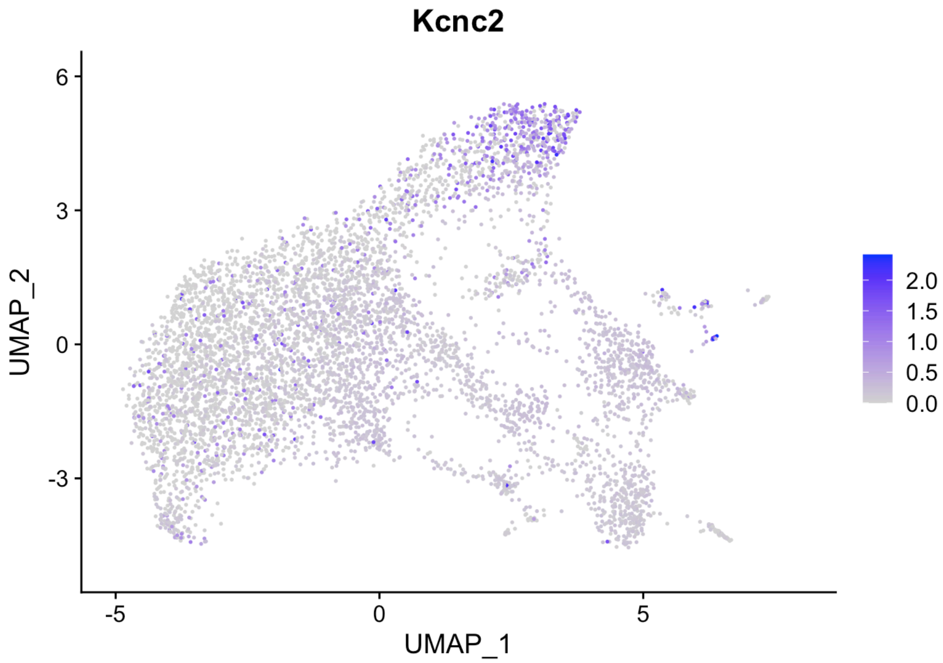


```
ggsave(file="/Users/lees44/R_project/FC_vs_dCA2_comparison_vector_integration_042821/Kcnc2.pdf",width=10,height=1
0)
```

```
#Feature plot highlighting potassium channel in all population
p1<-FeaturePlot(object =dCA2.integrated, features = c("Kcnc3"), min.cutoff = 0)
```

```
## Warning: Could not find Kcnc3 in the default search locations, found in RNA
## assay instead
```

```
plot_grid(p1)
```

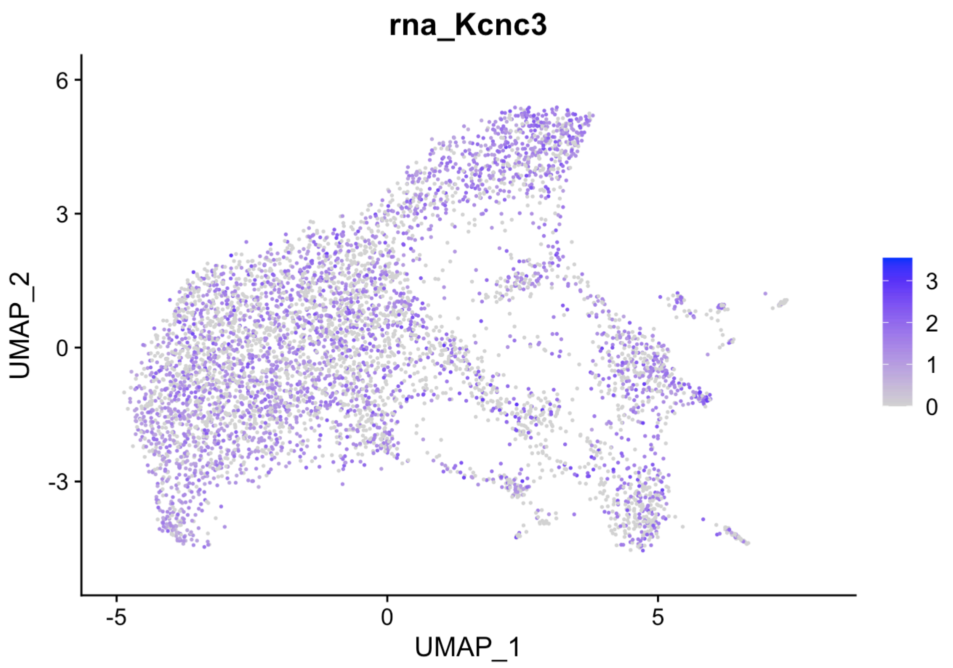


```
ggsave(file="/Users/lees44/R_project/FC_vs_dCA2_comparison_vector_integration_042821/Kcnc3.pdf",width=10,height=1
0)
```

```
#Feature plot highlighting potassium channel in all population
p1<-FeaturePlot(object =dCA2.integrated, features = c("Kcnc4"), min.cutoff = 0)
```

```
## Warning: Could not find Kcnc4 in the default search locations, found in RNA
## assay instead
```

```
plot_grid(p1)
```

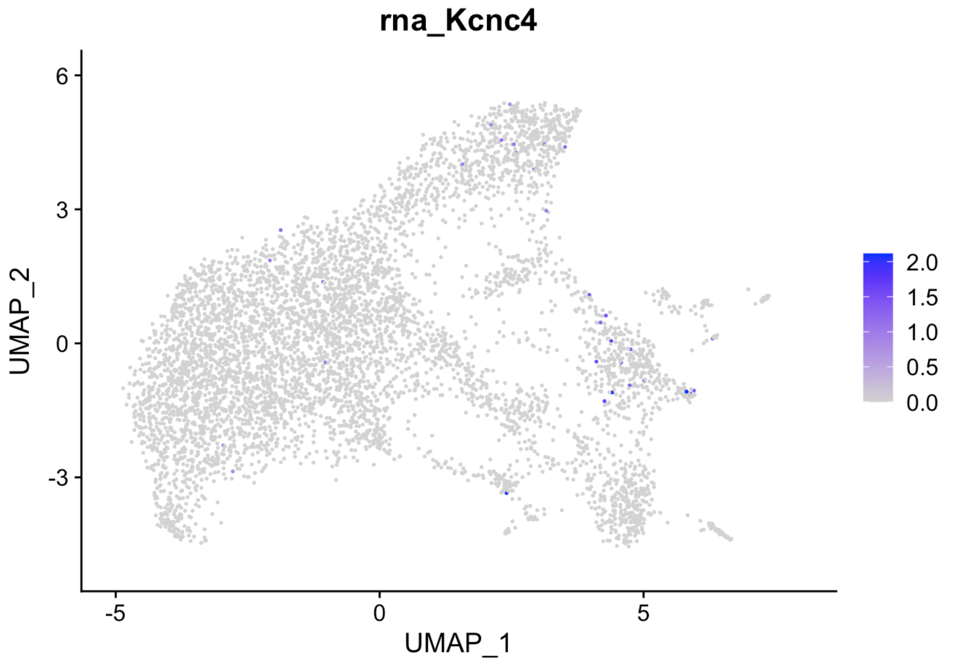


```
ggsave(file="/Users/lees44/R_project/FC_vs_dCA2_comparison_vector_integration_042821/Kcnc4.pdf",width=10,height=1
0)
```

```
#Feature plot highlighting potassium channel in all population
p1<-FeaturePlot(object =dCA2.integrated, features = c("Kcnd1"), min.cutoff = 0)
```

```
## Warning: Could not find Kcnd1 in the default search locations, found in RNA
## assay instead
```

```
plot_grid(p1)
```

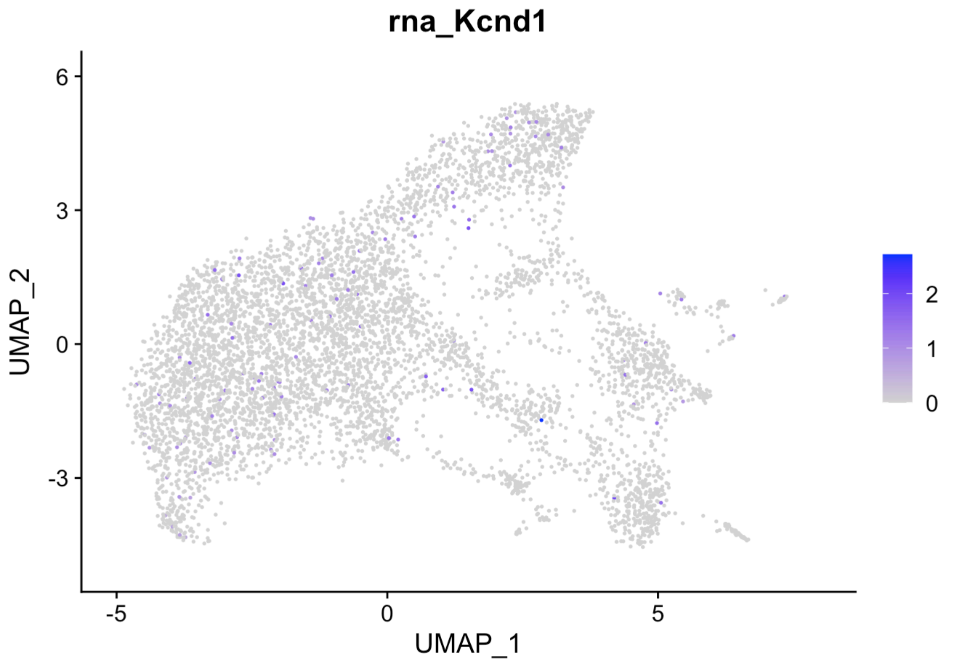

```
ggsave(file="/Users/lees44/R_project/FC_vs_dCA2_comparison_vector_integration_042821/Kcnd1.pdf",width=10,height=1
0)
```

```
#Feature plot highlighting potassium channel in all population
p1<-FeaturePlot(object =dCA2.integrated, features = c("Kcnd2"), min.cutoff = 0)
plot_grid(p1)
```

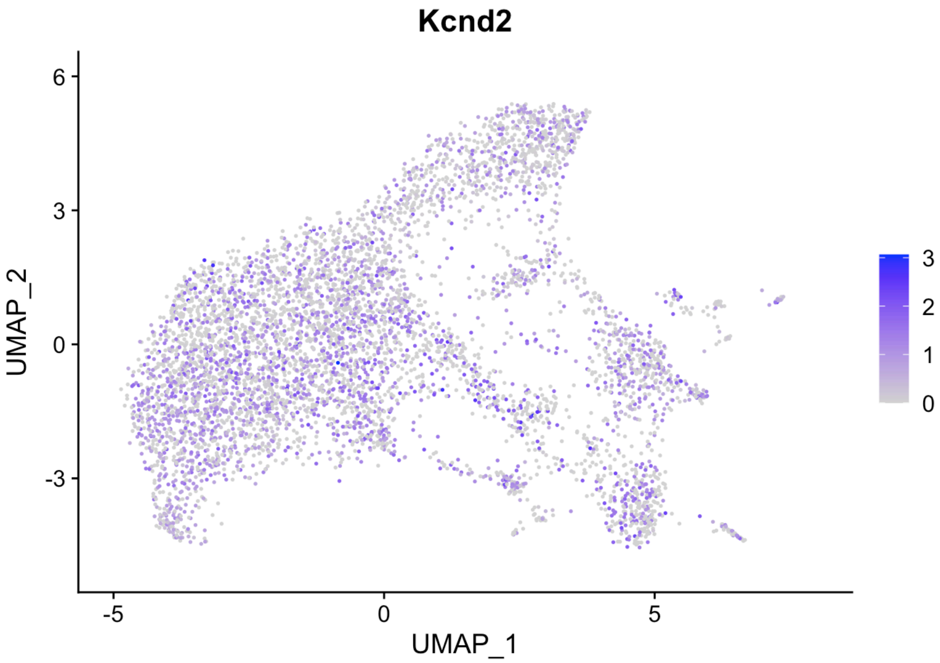


```
ggsave(file="/Users/lees44/R_project/FC_vs_dCA2_comparison_vector_integration_042821/Kcnd2.pdf",width=10,height=1
0)
```

```
#Feature plot highlighting potassium channel in all population
p1<-FeaturePlot(object =dCA2.integrated, features = c("Kcnd3"), min.cutoff = 0)
```

```
## Warning: Could not find Kcnd3 in the default search locations, found in RNA
## assay instead
```

```
plot_grid(p1)
```

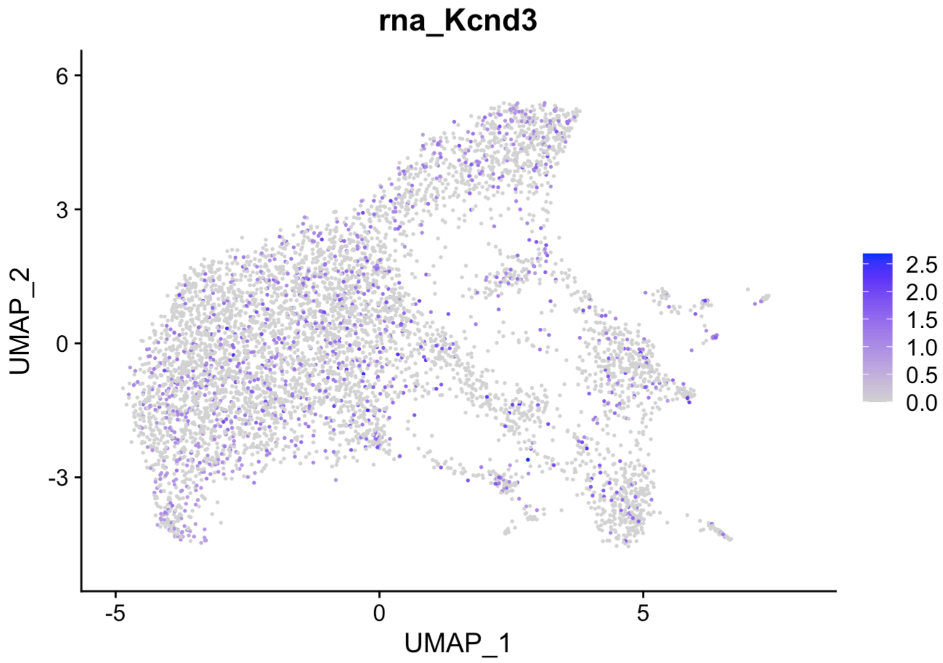


```
ggsave(file="/Users/lees44/R_project/FC_vs_dCA2_comparison_vector_integration_042821/Kcnd3.pdf",width=10,height=1
0)
```

```
#Feature plot highlighting potassium channel in all population
p1<-FeaturePlot(object =dCA2.integrated, features = c("Kcnq2"), min.cutoff = 0)
```

```
## Warning: Could not find Kcnq2 in the default search locations, found in RNA
## assay instead
```

```
plot_grid(p1)
```

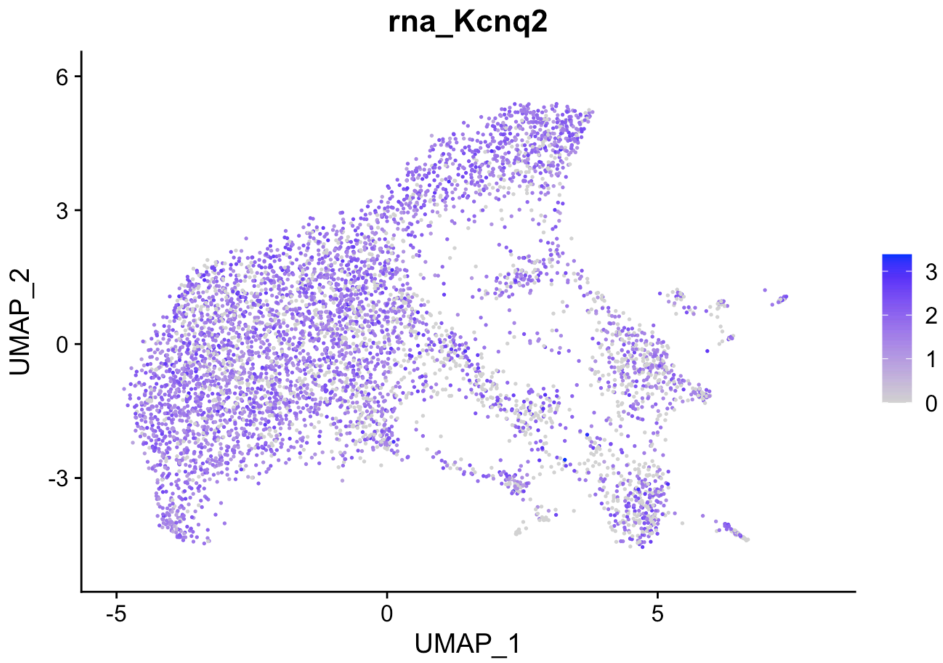

```
ggsave(file="/Users/lees44/R_project/FC_vs_dCA2_comparison_vector_integration_042821/Kcnq2.pdf",width=10,height=1
0)
```

```
#Feature plot highlighting potassium channel in all population
p1<-FeaturePlot(object =dCA2.integrated, features = c("Kcnq3"), min.cutoff = 0)
plot_grid(p1)
```

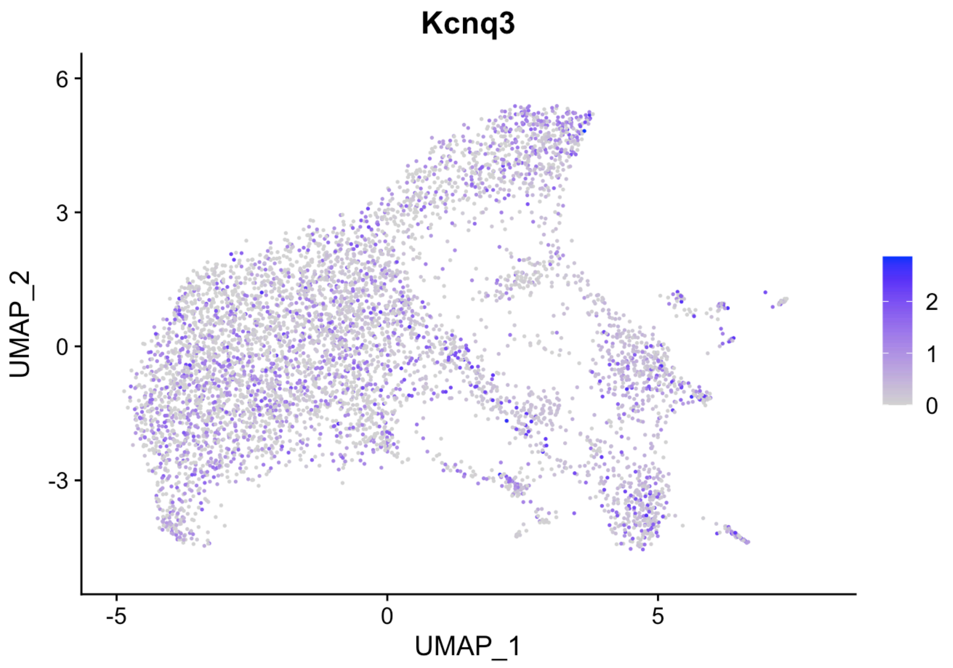


```
ggsave(file="/Users/lees44/R_project/FC_vs_dCA2_comparison_vector_integration_042821/Kcnq3.pdf",width=10,height=1
0)
```

```
#Feature plot highlighting potassium channel in all population
p1<-FeaturePlot(object =dCA2.integrated, features = c("Kcnq5"), min.cutoff = 0)
plot_grid(p1)
```

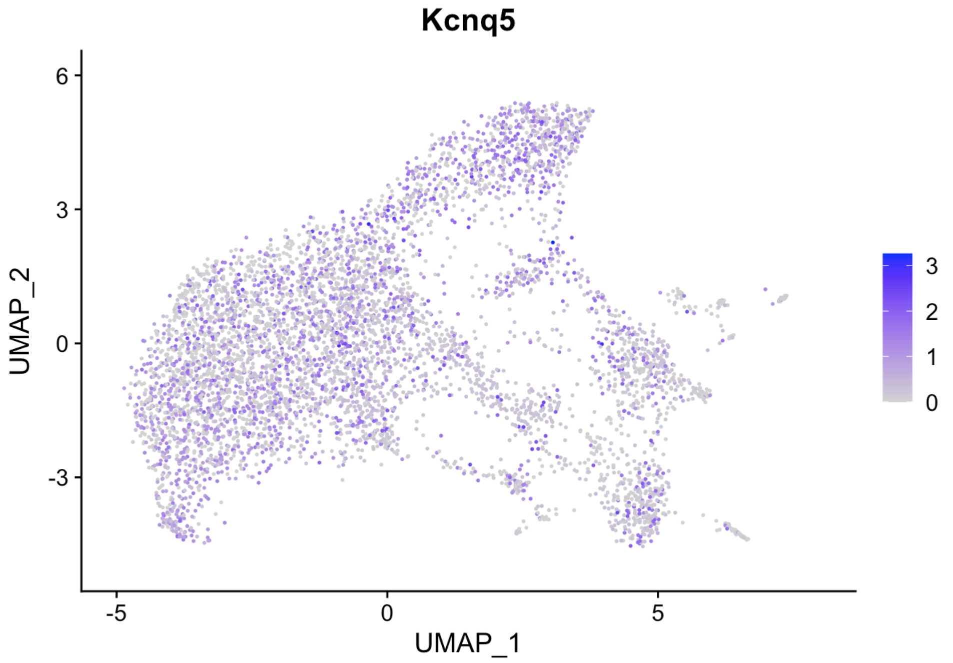


```
ggsave(file="/Users/lees44/R_project/FC_vs_dCA2_comparison_vector_integration_042821/Kcnq5.pdf",width=10,height=1
0)
```

Assign clusters types cell types

```
#celltypes asigned based on that gene marker expression
new.ident <- c("Type1", "Type2", "Type3", "Type4", "Type5", "Type6", "Type7", "Type8", "Type9")
names(x = new.ident) <- levels(x =dCA2.integrated)
dCA2.integrated<- RenameIdents(object =dCA2.integrated, new.ident)
```

```
color<-c("#19647e","#ffc857","#9B0C1E","#9B0C1E","#9B0C1E","#9B0C1E","#9B0C1E","#9B0C1E","#4b3f72")
```

```
DimPlot(object = dCA2.integrated, reduction = "umap", label = FALSE, repel = TRUE,cols=color,label.size = 12,pt.s
ize=0.5)
```

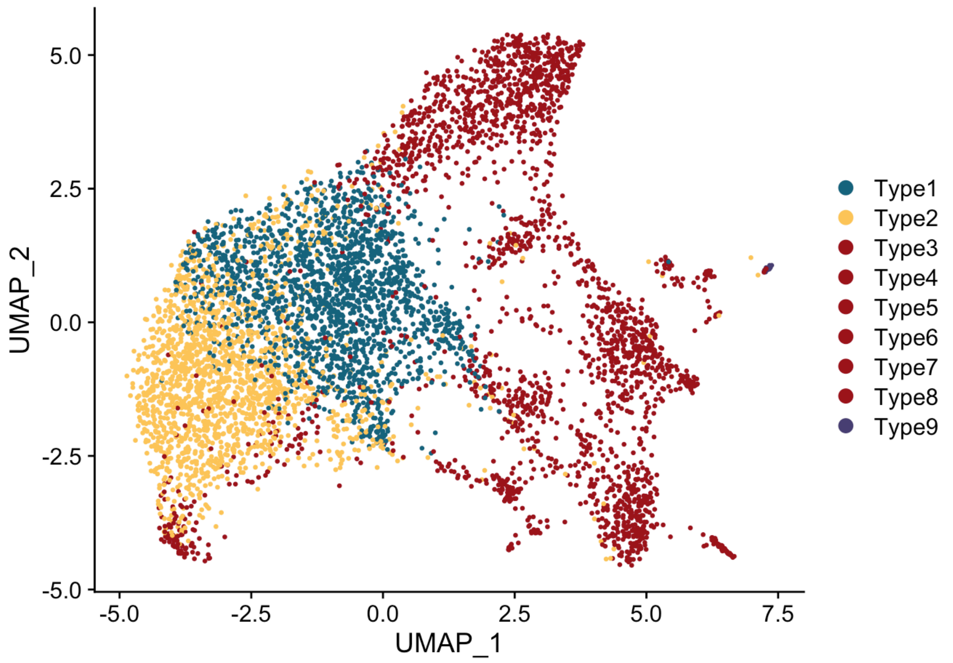


```
ggsave(file="/Users/lees44/R_project/FC_vs_dCA2_comparison_vector_integration_042821/Neuronaltype.pdf",width=10,h
eight=7)
```

```
DimPlot(object = dCA2.integrated, reduction = "umap", label = TRUE, repel = TRUE)
```

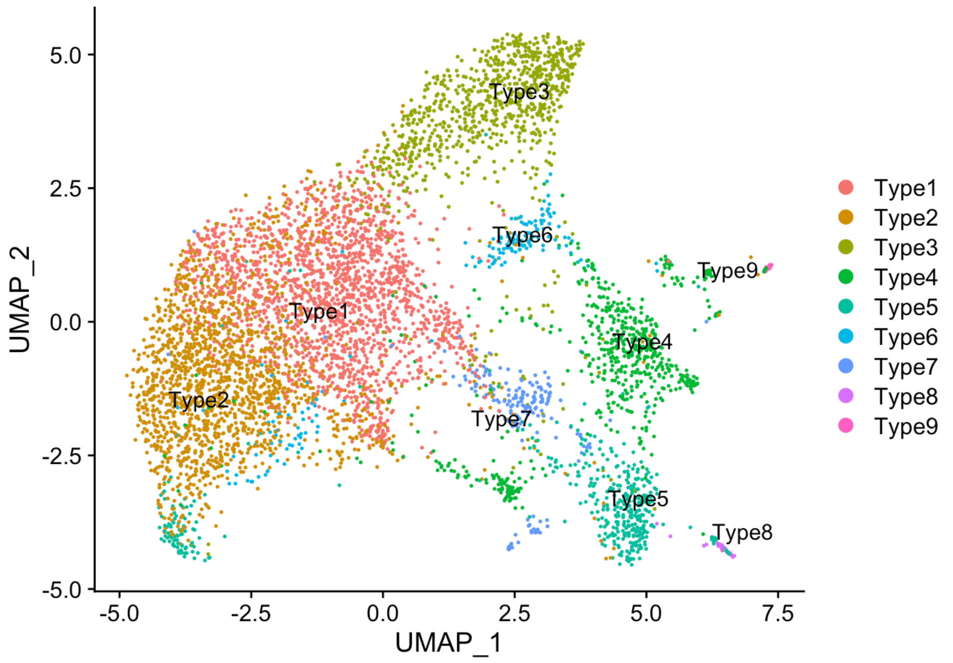


```
ggsave(file="/Users/lees44/R_project/FC_vs_dCA2_comparison_vector_integration_042821/Neuronaltype.pdf",width=10,h
eight=10)
```

Identification of all markers for each cluster see https://hbctraining.github.io/scRNA-seq/lessons/sc_exercises_integ_marker_identification.html (https://hbctraining.github.io/scRNA-seq/lessons/sc_exercises_integ_marker_identification.html)

This analysis compares each cluster against all others and outputs the genes that are differentially expressed/present using the FindAllMarkers() function.

For different DEG functions read

https://www.biostars.org/p/409790/ (https://www.biostars.org/p/409790/)

```
# Find markers for every cluster compared to all remaining cells, report only the positive ones
combined_markers <- FindAllMarkers(object = dCA2.integrated,
                          only.pos = TRUE,
                          logfc.threshold = 0.25)
```

```
## Calculating cluster Type1
```

```
## Calculating cluster Type2
```

```
## Calculating cluster Type3
```

```
## Calculating cluster Type4
```

```
## Calculating cluster Type5
```

```
## Calculating cluster Type6
```

```
## Calculating cluster Type7
```

```
## Calculating cluster Type8
```

```
## Calculating cluster Type9
```

```
View(combined_markers)

# Save markers to file
write.csv(combined_markers,
          file = "/Users/lees44/R_project/FC_vs_dCA2_comparison_vector_integration_042821/combined_all_markers1.c
sv",
          quote = FALSE,
          row.names = FALSE)
```

We can also output the top 5 markers by log2 fold change for each cluster for a quick perusal.

```
# Extract top 50 markers per cluster
top50_comb <- combined_markers %>%
        group_by(cluster) %>%
        top_n(n = 50,
              wt = avg_log2FC)

# Visualize top 50 markers per cluster
View(top50_comb)

# Save markers to file
write.csv(combined_markers,
          file = "/Users/lees44/R_project/FC_vs_dCA2_comparison_vector_integration_042821/top50_markers1.csv",
          quote = FALSE,
          row.names = FALSE)
```

Heatmap for different cell types from combined markers

DoHeatmap generates an expression heatmap for given cells and features. In this case, we are plotting the top 20 markers (or all markers if less than 20) for each cluster. https://satijalab.org/seurat/archive/v3.2/pbmc3k_tutorial.html (https://satijalab.org/seurat/archive/v3.2/pbmc3k_tutorial.html)

```
combined_markers %>%
    group_by(cluster) %>%
    top_n(n = 10, wt = avg_log2FC) -> top10
DoHeatmap(dCA2.integrated, features = top10$gene) + NoLegend()
```

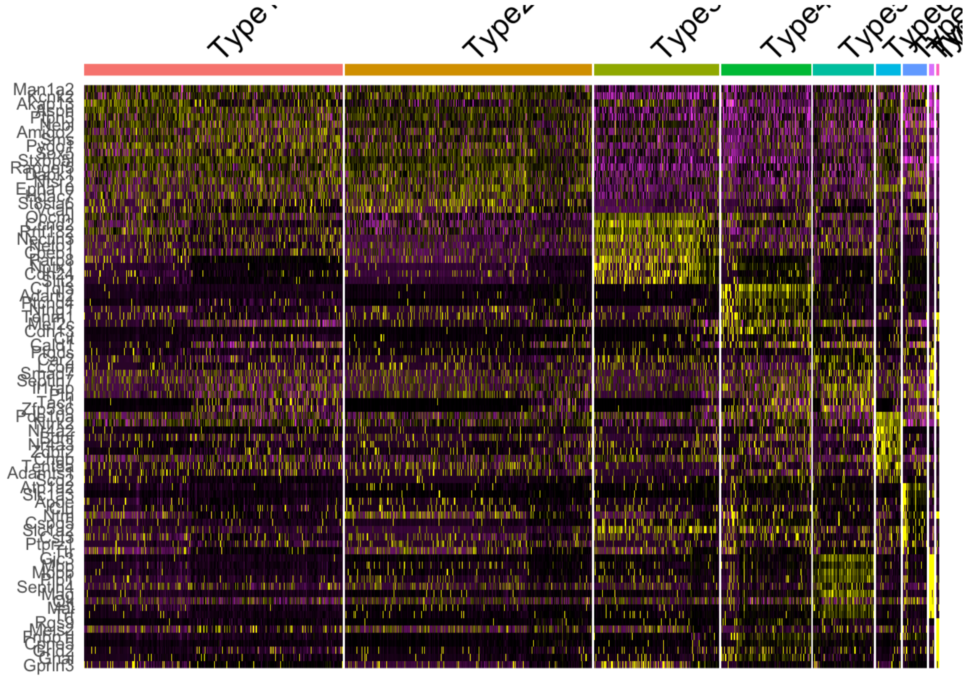

```
ggsave(file="/Users/lees44/R_project/FC_vs_dCA2_comparison_vector_integration_042821/heatmap_top10_celltype1.pdf",
width=50,height=25, limitsize = FALSE)
```

Dot plot for Top 10 markers from combined markers

https://blog.dolomite-bio.com/visualization-of-single-cell-data-from-seurat-object-to-umap/ (https://blog.dolomite-bio.com/visualization-of-single-cell-data-from-seurat-object-to-umap/)

Change font and dot size

https://satijalab.org/seurat/reference/seurattheme (https://satijalab.org/seurat/reference/seurattheme)

```
features <- c("Man1a2", "Kcnk2", "Akap13", "Asph", "Ptpn5", "Nebl", "Amigo2", "Sms", "Pygo1", "Sox5", "Stxbp5l",
"Rapgef5", "Dapk1", "Ntsr2", "Epha10", "Hdac7", "St8sia6", "Vcan", "Opcml", "Ccnd2", "Rnf182", "Nectin3", "Neto
1", "Cpeb1", "Parp8", "Nptx1", "Cdh24", "Slit2", "C1ql3", "Adarb2", "Ptchd4", "Ntng1", "Tenm1", "Mef2c", "Cdh13",
"Cit", "Cald1", "Ptgds", "Car2", "Lcorl", "Smad7", "Septin7", "Il1rap", "Ptn", "Tac1", "Zfp536", "Pde10a", "Ntrk
2", "Nr4a2", "Bdnf", "Nr4a3", "Zdbf2", "Chgb", "Tent5a", "Adamts1", "Scg2", "Atp1a2", "Slc1a3", "Apoe", "Clu", "N
tm", "Cspg5", "Slc1a2", "Cst3", "Ptprz1", "Ttr", "Gjc3", "Mbp", "Mobp", "Plp1", "Septin4", "Mag", "Qk", "Mal", "T
rf", "Rgs9", "Meis2", "Fnbp1l", "Cpne5", "Grid2", "Gnal", "Gprin3")
```

```
DotPlot(dCA2.integrated, features = features) + RotatedAxis() + scale_size(range = c(1, 20))  + FontSize(x.text =
30, y.text = 30)
```

```
## Scale for 'size' is already present. Adding another scale for 'size', which
## will replace the existing scale.
```

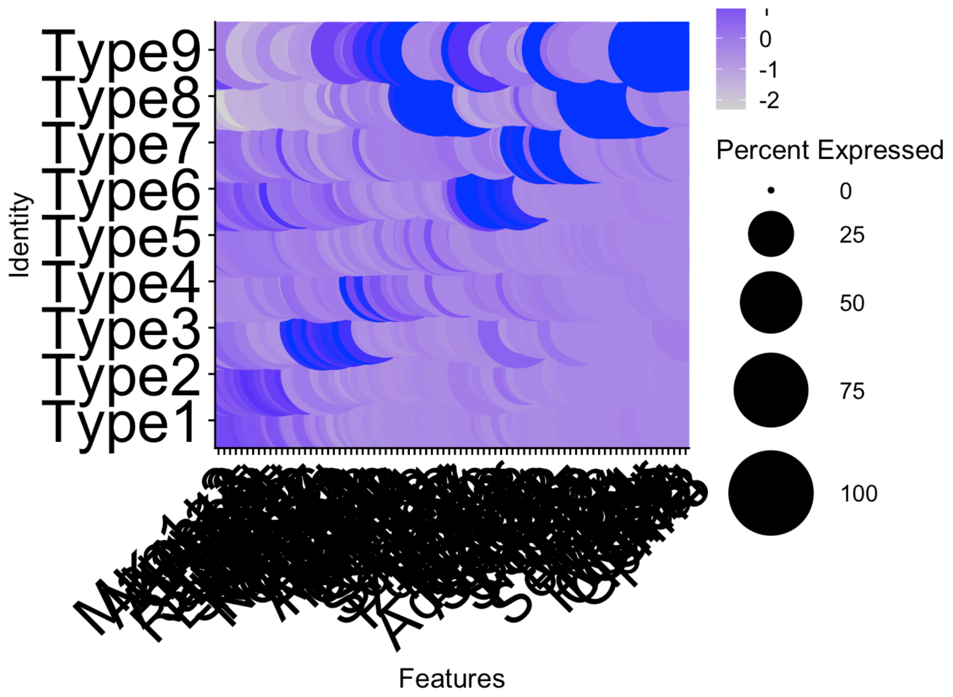


```
ggsave(file="/Users/lees44/R_project/FC_vs_dCA2_comparison_vector_integration_042821/dotplot_top10_celltype.pdf",
width=50, height=25, limitsize = FALSE)
```

Tabulate cells by cluster ID, replicate, or both. see https://satijalab.org/seurat/archive/v3.0/interaction_vignette.html (https://satijalab.org/seurat/archive/v3.0/interaction_vignette.html)

```
# How many cells are in each cluster
table(Idents(dCA2.integrated))
```

```
##
## Type1 Type2 Type3 Type4 Type5 Type6 Type7 Type8 Type9
##  1830  1746   886   635   428   176   173    35    19
```

```
df <-data.frame(
  group = c("Type1", "Type2", "Type3", "Type4", "Type5", "Type6","Type7", "Type8", "Type9"),
  value = c(1830, 1746, 886, 635, 428, 176, 173, 35, 19)
)
head(df)
```

```
##   group value
## 1 Type1  1830
## 2 Type2  1746
## 3 Type3   886
## 4 Type4   635
## 5 Type5   428
## 6 Type6   176
```

```
bp<- ggplot(df, aes(x="", y=value, fill=group))+
geom_bar(width = 1, stat = "identity")
bp
```

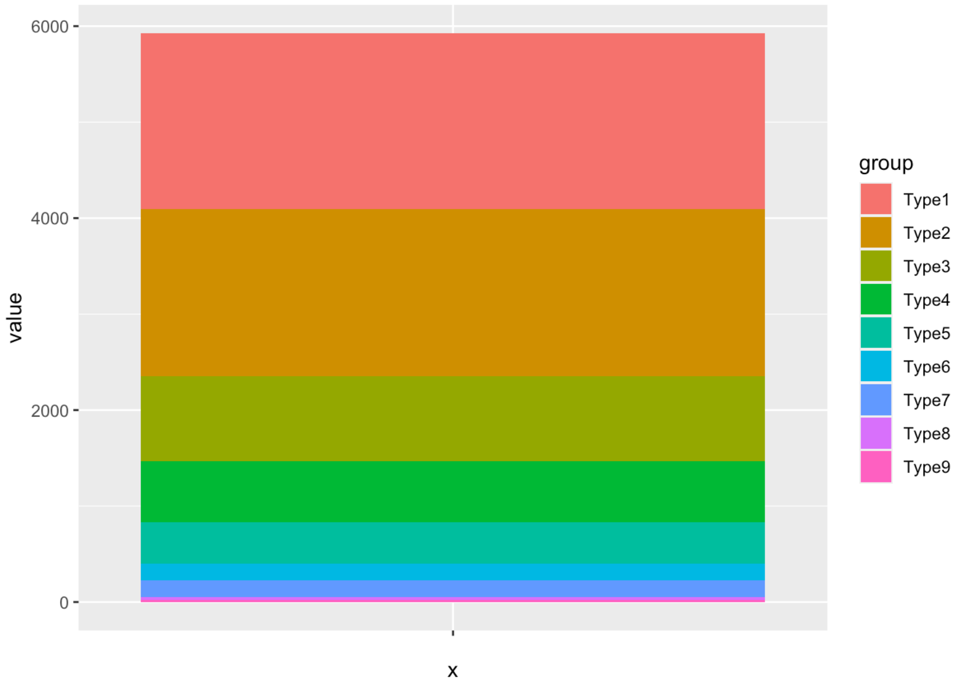


```
pie <- bp + coord_polar("y", start=0)
pie
```

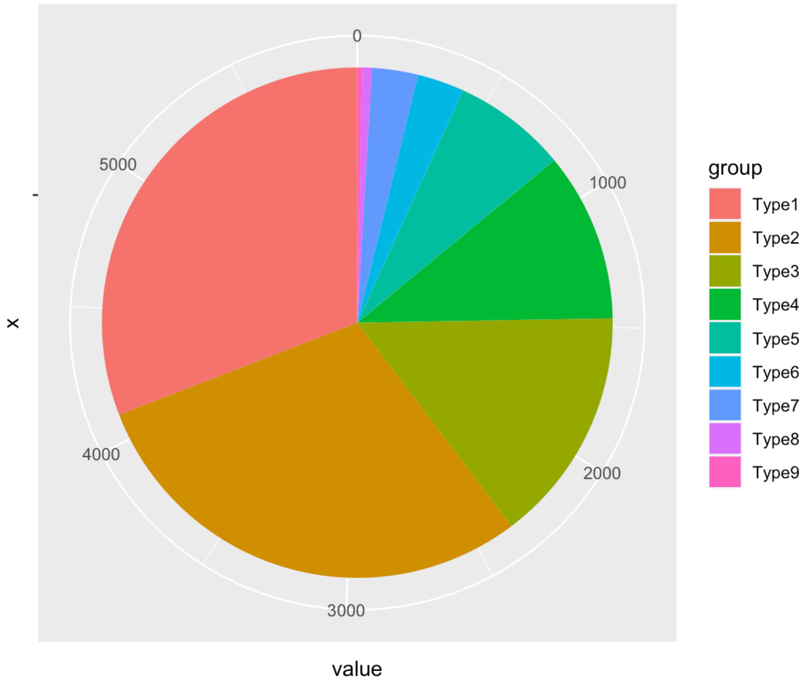


```
ggsave(file="/Users/lees44/R_project/FC_vs_dCA2_comparison_vector_integration_042821/cell_number_cluster_pie.pdf",
width=10,height=7)
```

```
# How many cells are in each group?
table(dCA2.integrated$protocol)
```

```
##
## dCA2_Fpos dCA2_Mpos   FC_Fpos   FC_Mpos
##      1138      1981      1566      1243
```

```
df <-data.frame(
  group = c("dCA2_Fpos", "dCA2_Mpos", "FC_Fpos", "FC_Mpos"),
  value = c(1138, 1981, 1566, 1243)
)
head(df)
```

```
##       group value
## 1 dCA2_Fpos  1138
## 2 dCA2_Mpos  1981
## 3   FC_Fpos  1566
## 4   FC_Mpos  1243
```

```
bp<- ggplot(df, aes(x="", y=value, fill=group))+
geom_bar(width = 1, stat = "identity")
bp
```

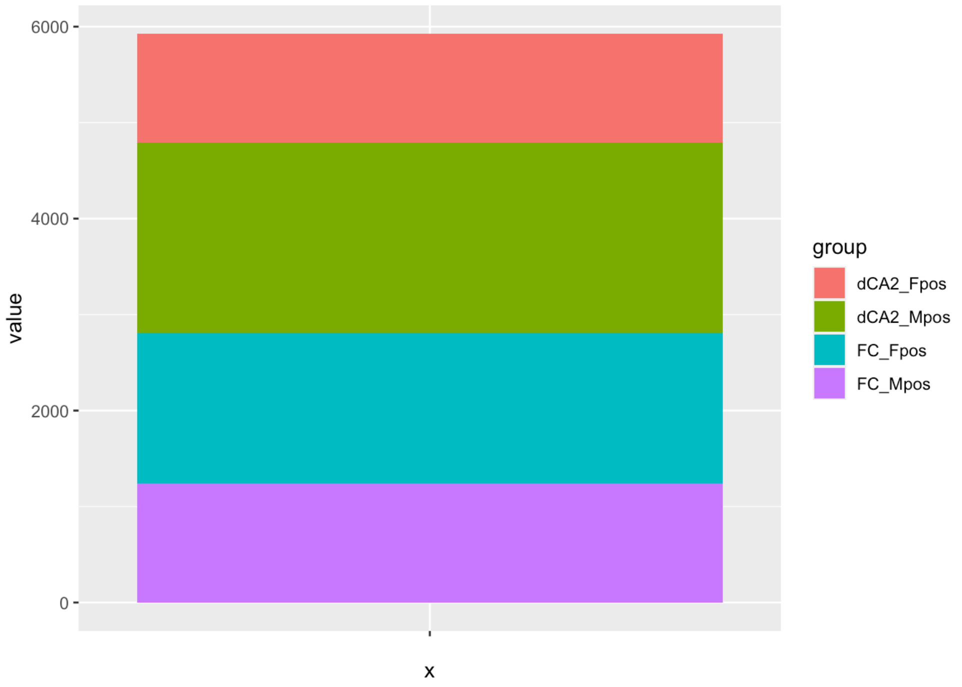


```
pie <- bp + coord_polar("y", start=0)
pie
```

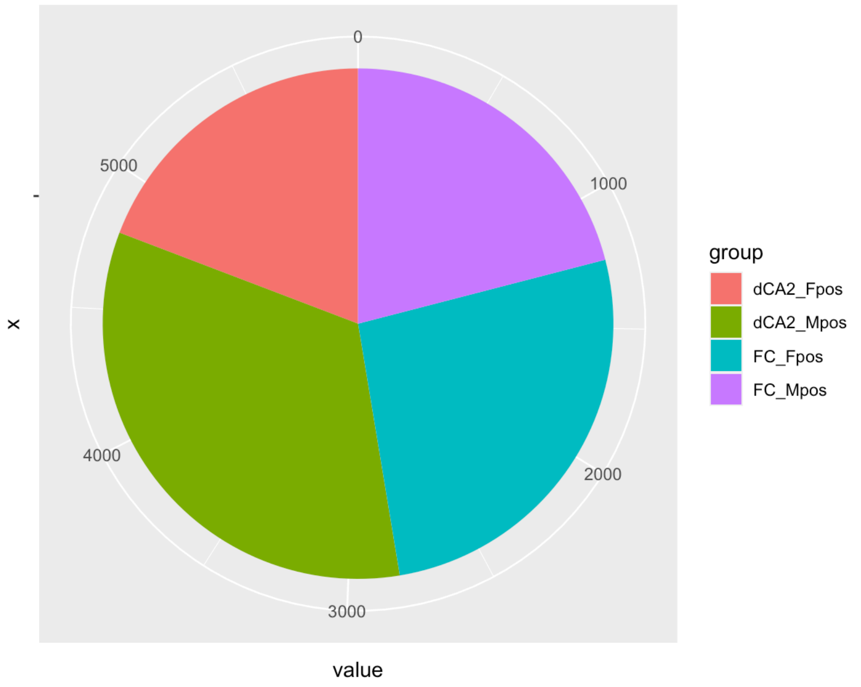


```
ggsave(file="/Users/lees44/R_project/FC_vs_dCA2_comparison_vector_integration_042821/cell_number_group_pie.pdf",w
idth=10,height=7)
```

```
# What proportion of cells are in each cluster?
prop.table(table(Idents(dCA2.integrated)))
```

```
##
##       Type1       Type2       Type3       Type4       Type5       Type6
## 0.308704453 0.294534413 0.149460189 0.107118758 0.072199730 0.029689609
##       Type7       Type8       Type9
## 0.029183536 0.005904184 0.003205128
```

```
# How does cluster membership vary by group?
table(Idents(dCA2.integrated), dCA2.integrated$protocol)
```

```
##
##         dCA2_Fpos dCA2_Mpos FC_Fpos FC_Mpos
##   Type1       378       378     652     422
##   Type2       402       900     171     273
##   Type3       268       418     127      73
##   Type4        21       101     285     228
##   Type5         4        49     164     211
##   Type6        38        82      34      22
##   Type7         8        25     129      11
##   Type8        12        16       4       3
##   Type9         7        12       0       0
```

pie graph for different cell types

```
#type1
df <-data.frame(
  group = c("dCA2_Fpos", "dCA2_Mpos", "FC_Fpos", "FC_Mpos"),
  value = c(378, 378, 652, 422)
)
head(df)
```

```
##        group value
## 1 dCA2_Fpos   378
## 2 dCA2_Mpos   378
## 3   FC_Fpos   652
## 4   FC_Mpos   422
```

```
bp<- ggplot(df, aes(x="", y=value, fill=group))+
geom_bar(width = 1, stat = "identity")
bp
```

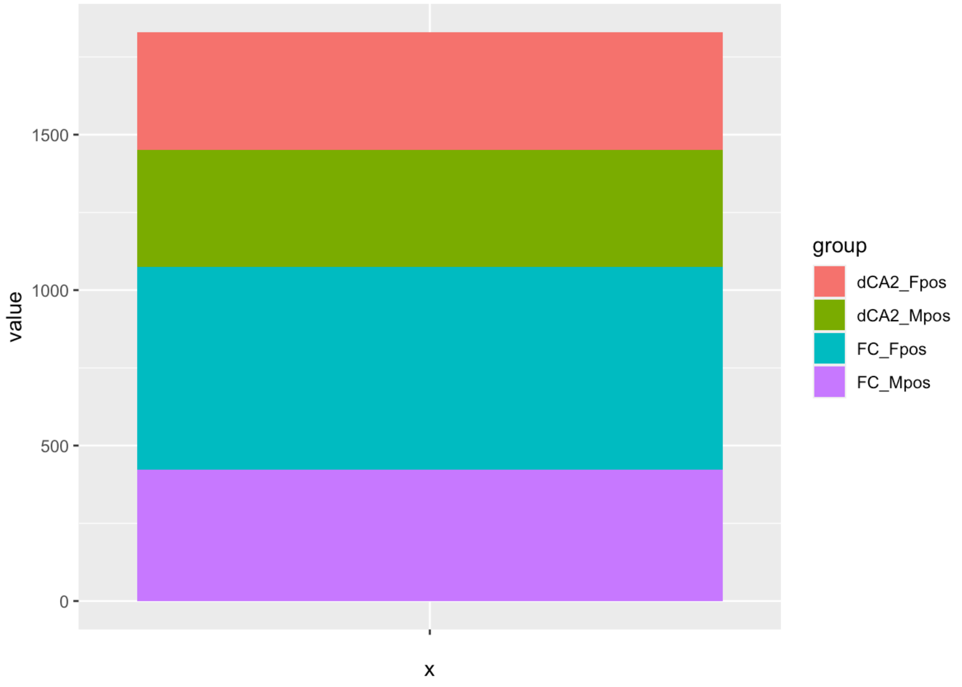


```
pie <- bp + coord_polar("y", start=0)
pie
```

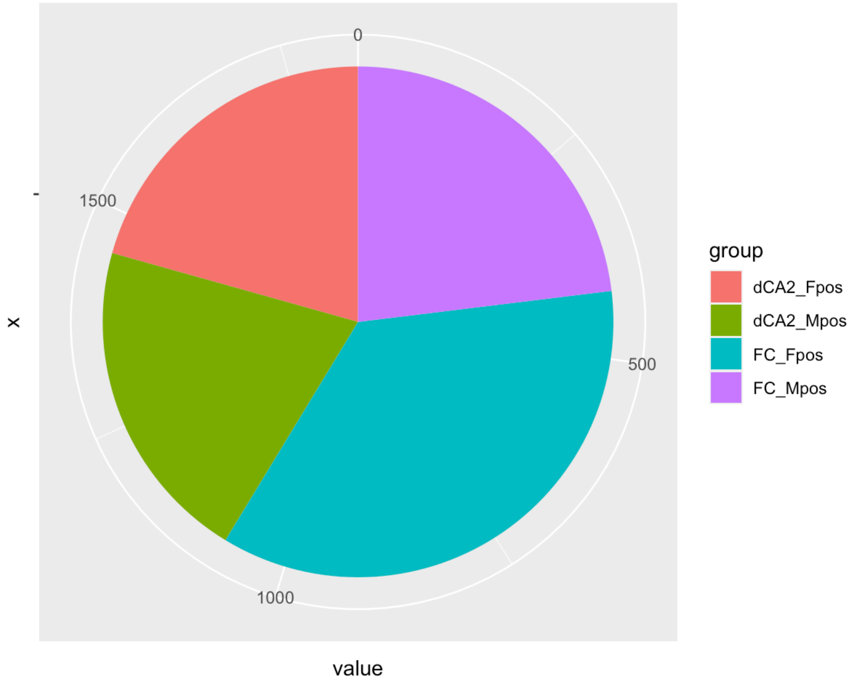

```
ggsave(file="/Users/lees44/R_project/FC_vs_dCA2_comparison_vector_integration_042821/cell_number_group_type1_pie.
pdf",width=10,height=7)
```

```
#type2
df <-data.frame(
  group = c("dCA2_Fpos", "dCA2_Mpos", "FC_Fpos", "FC_Mpos"),
  value = c(402, 900, 171, 273)
)
head(df)
```

```
##       group value
## 1 dCA2_Fpos   402
## 2 dCA2_Mpos   900
## 3   FC_Fpos   171
## 4   FC_Mpos   273
```

```
bp<- ggplot(df, aes(x="", y=value, fill=group))+
geom_bar(width = 1, stat = "identity")
bp
```

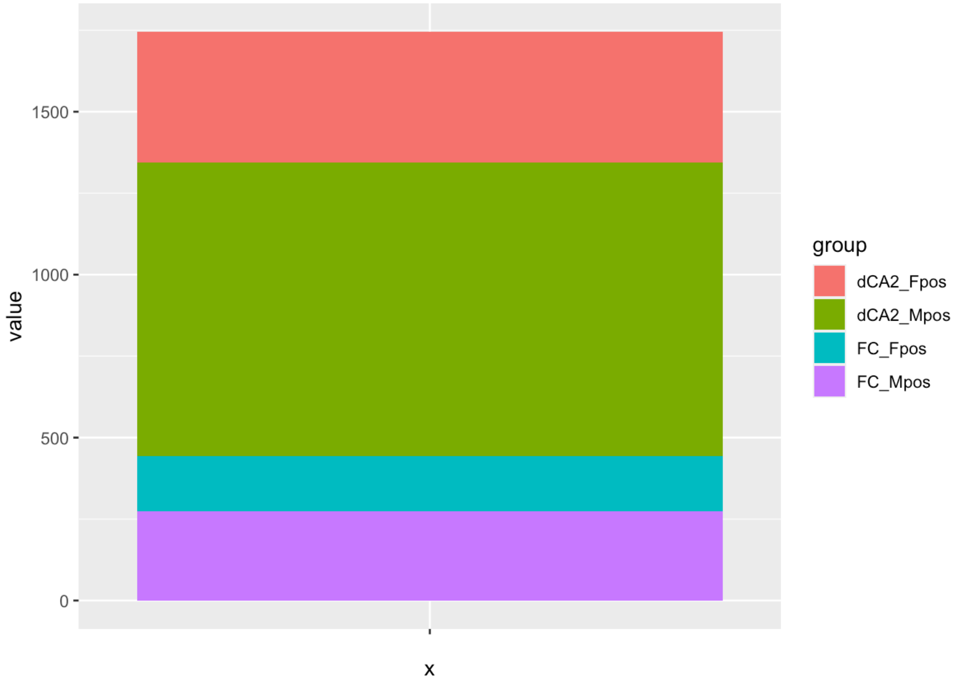


```
pie <- bp + coord_polar("y", start=0)
pie
```

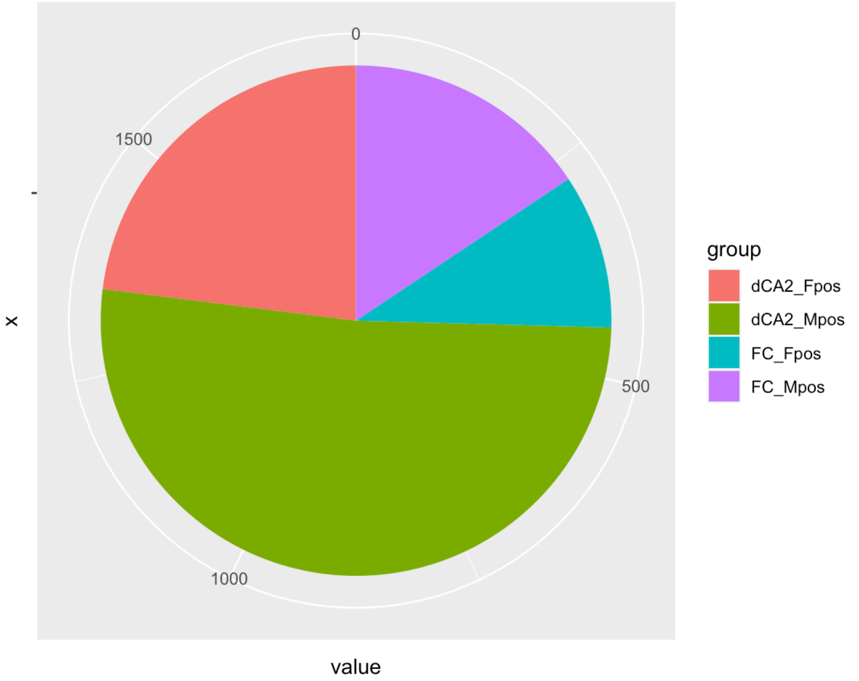


```
ggsave(file="/Users/lees44/R_project/FC_vs_dCA2_comparison_vector_integration_042821/cell_number_group_type2_pie.
pdf",width=10,height=7)
```

```
#type3
df <-data.frame(
  group = c("dCA2_Fpos", "dCA2_Mpos", "FC_Fpos", "FC_Mpos"),
  value = c(268, 418, 127, 73)
)
head(df)
```

```
##       group value
## 1 dCA2_Fpos   268
## 2 dCA2_Mpos   418
## 3   FC_Fpos   127
## 4   FC_Mpos    73
```

```
bp<- ggplot(df, aes(x="", y=value, fill=group))+
geom_bar(width = 1, stat = "identity")
bp
```

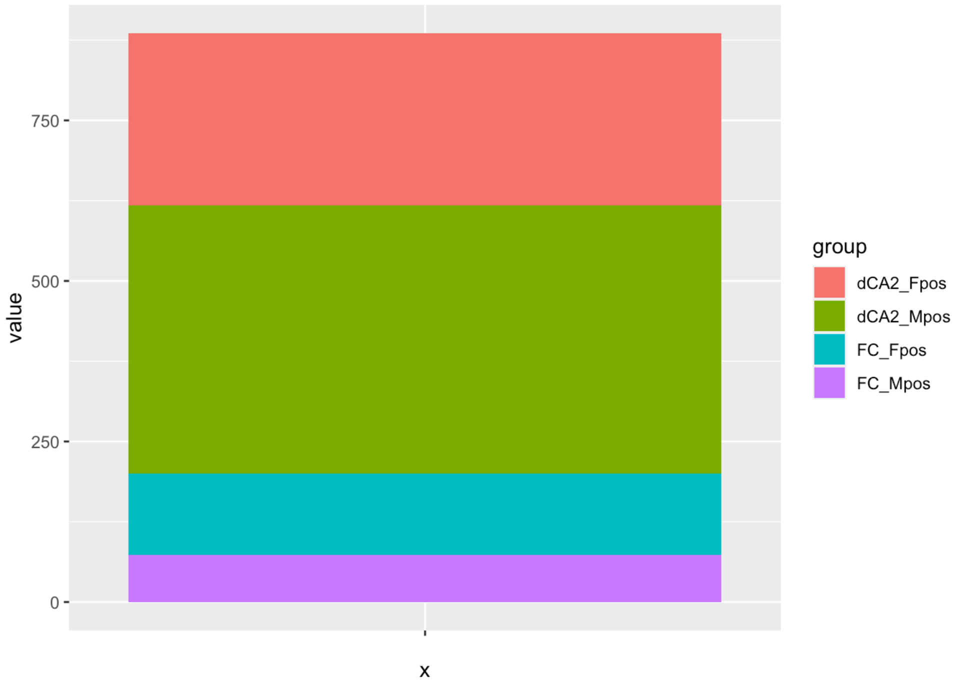


```
pie <- bp + coord_polar("y", start=0)
pie
```

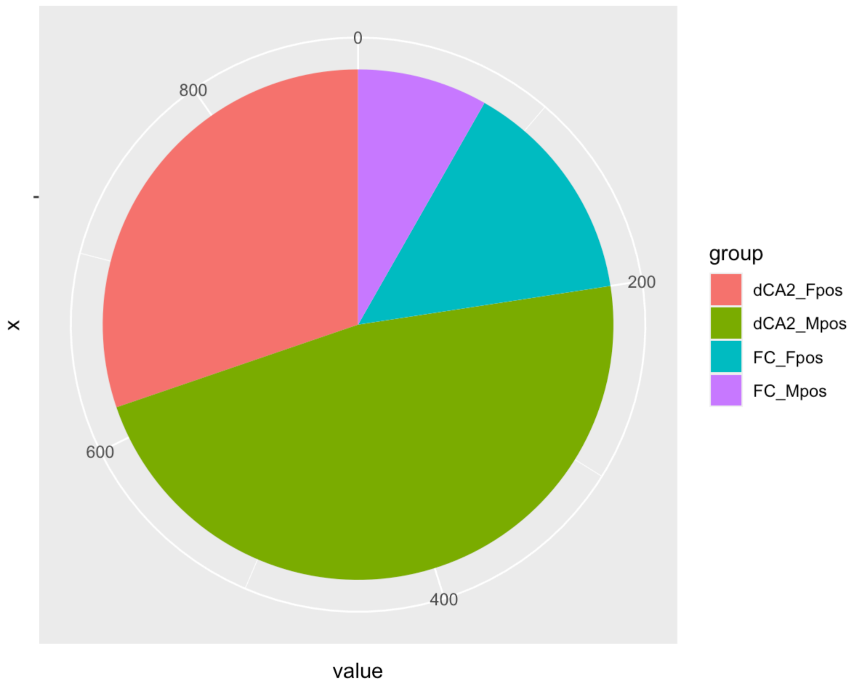


```
ggsave(file="/Users/lees44/R_project/FC_vs_dCA2_comparison_vector_integration_042821/cell_number_group_type3_pie.
pdf",width=10,height=7)
```

```
#type4
df <-data.frame(
  group = c("dCA2_Fpos", "dCA2_Mpos", "FC_Fpos", "FC_Mpos"),
  value = c(21, 101, 285, 228)
)
head(df)
```

```
##       group value
## 1 dCA2_Fpos    21
## 2 dCA2_Mpos   101
## 3   FC_Fpos   285
## 4   FC_Mpos   228
```

```
bp<- ggplot(df, aes(x="", y=value, fill=group))+
geom_bar(width = 1, stat = "identity")
bp
```

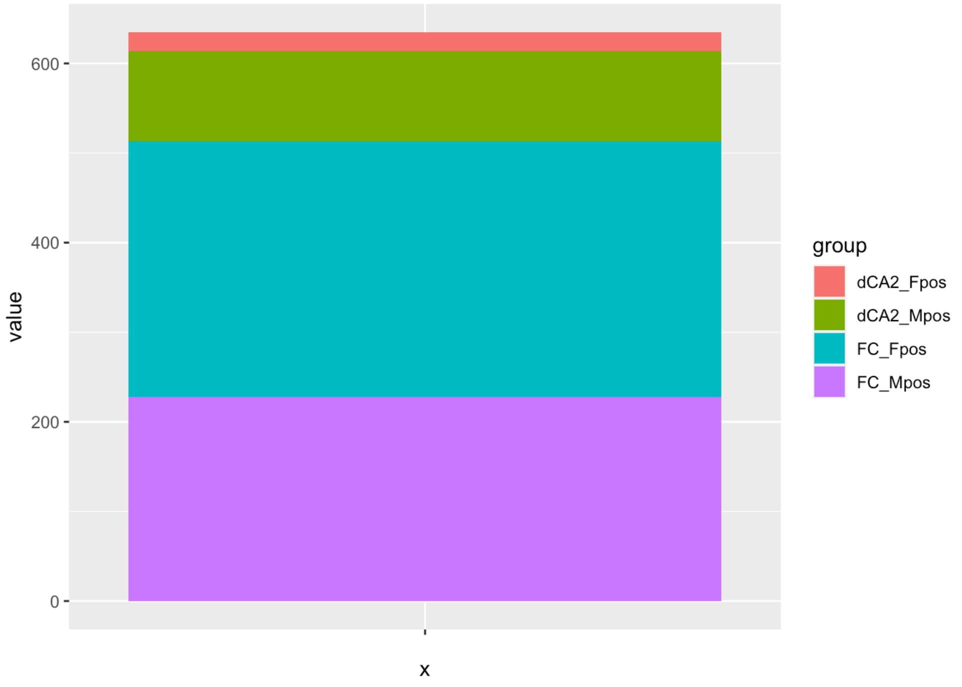


```
pie <- bp + coord_polar("y", start=0)
pie
```

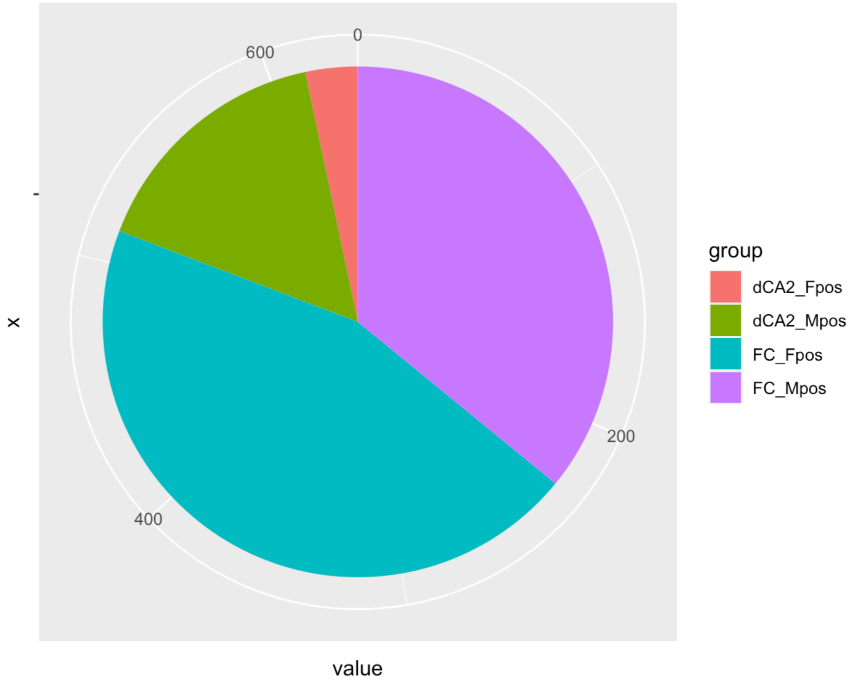

```
ggsave(file="/Users/lees44/R_project/FC_vs_dCA2_comparison_vector_integration_042821/cell_number_group_type4_pie.
pdf",width=10,height=7)
```

```
#type5
df <-data.frame(
  group = c("dCA2_Fpos", "dCA2_Mpos", "FC_Fpos", "FC_Mpos"),
  value = c(4, 49, 164, 211)
)
head(df)
```

```
##       group value
## 1 dCA2_Fpos     4
## 2 dCA2_Mpos    49
## 3   FC_Fpos   164
## 4   FC_Mpos   211
```

```
bp<- ggplot(df, aes(x="", y=value, fill=group))+
geom_bar(width = 1, stat = "identity")
bp
```

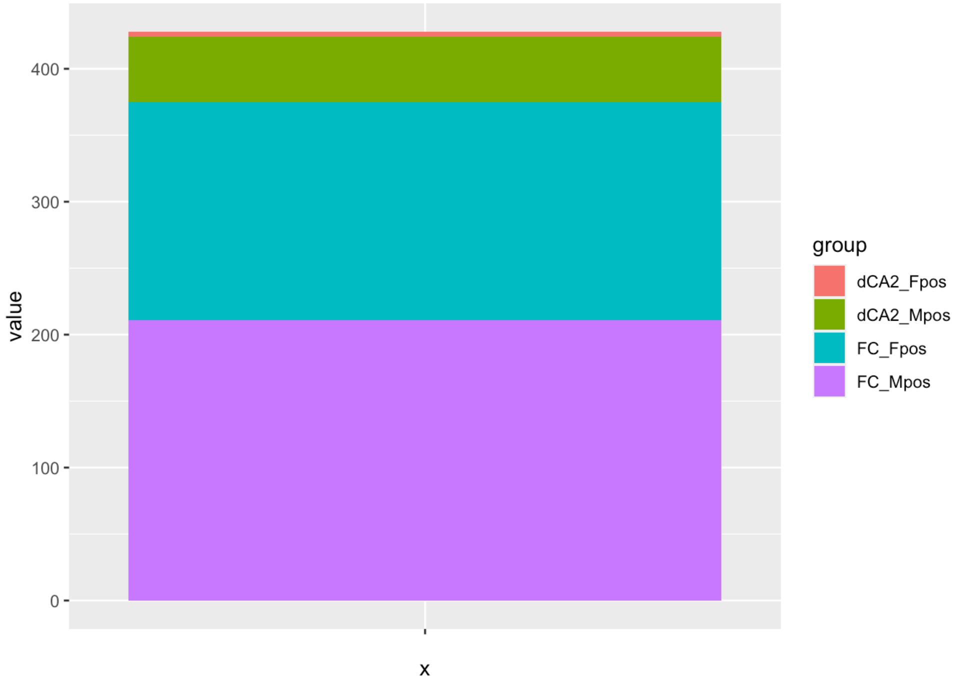


```
pie <- bp + coord_polar("y", start=0)
pie
```

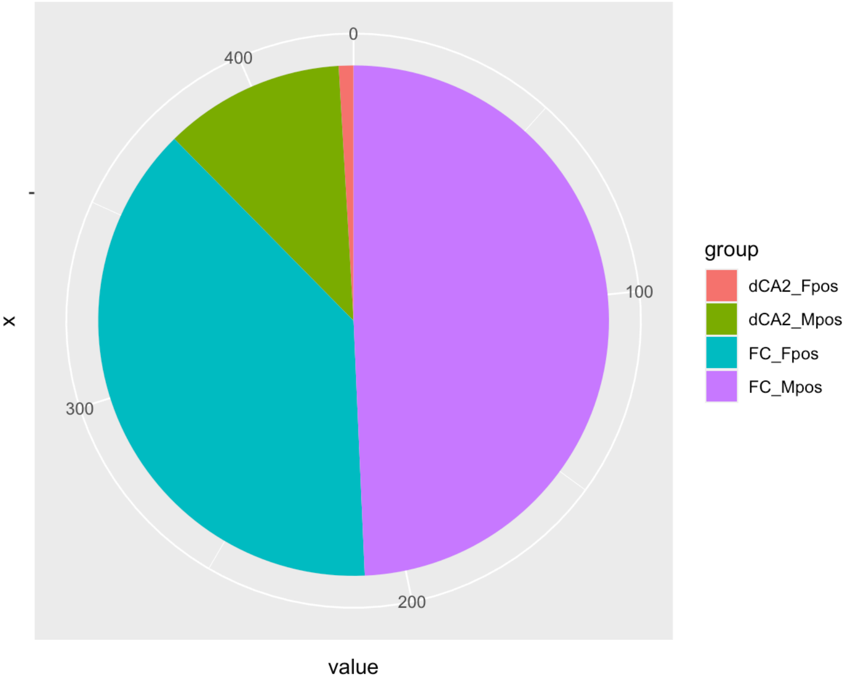


```
ggsave(file="/Users/lees44/R_project/FC_vs_dCA2_comparison_vector_integration_042821/cell_number_group_type3_pie.
pdf",width=10,height=7)
```

```
#type6
df <-data.frame(
  group = c("dCA2_Fpos", "dCA2_Mpos", "FC_Fpos", "FC_Mpos"),
  value = c(38, 82, 34, 22)
)
head(df)
```

```
##       group value
## 1 dCA2_Fpos    38
## 2 dCA2_Mpos    82
## 3   FC_Fpos    34
## 4   FC_Mpos    22
```

```
bp<- ggplot(df, aes(x="", y=value, fill=group))+
geom_bar(width = 1, stat = "identity")
bp
```

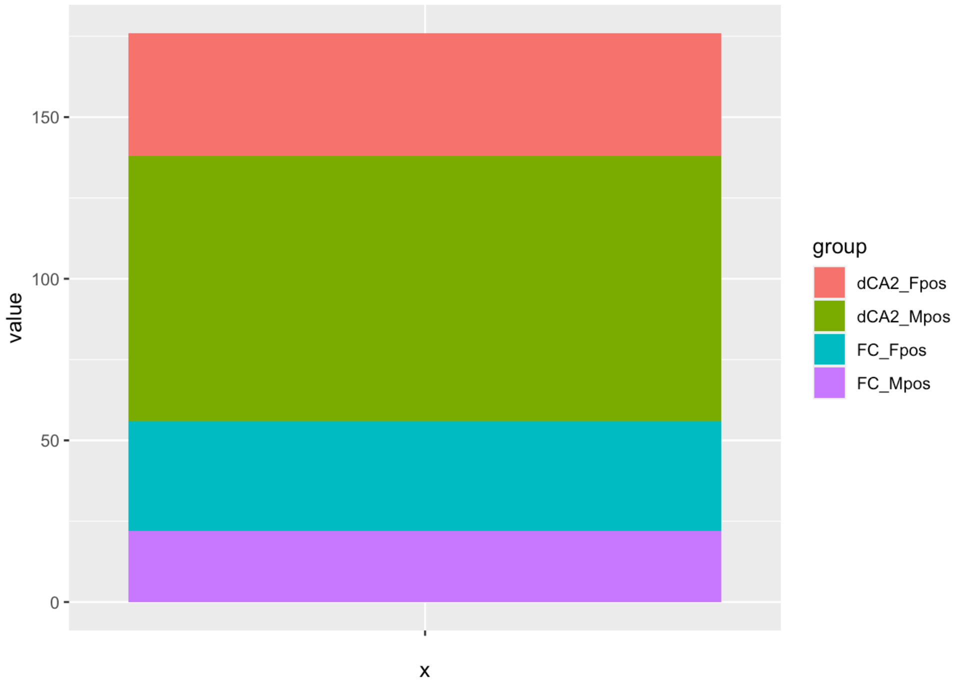


```
pie <- bp + coord_polar("y", start=0)
pie
```

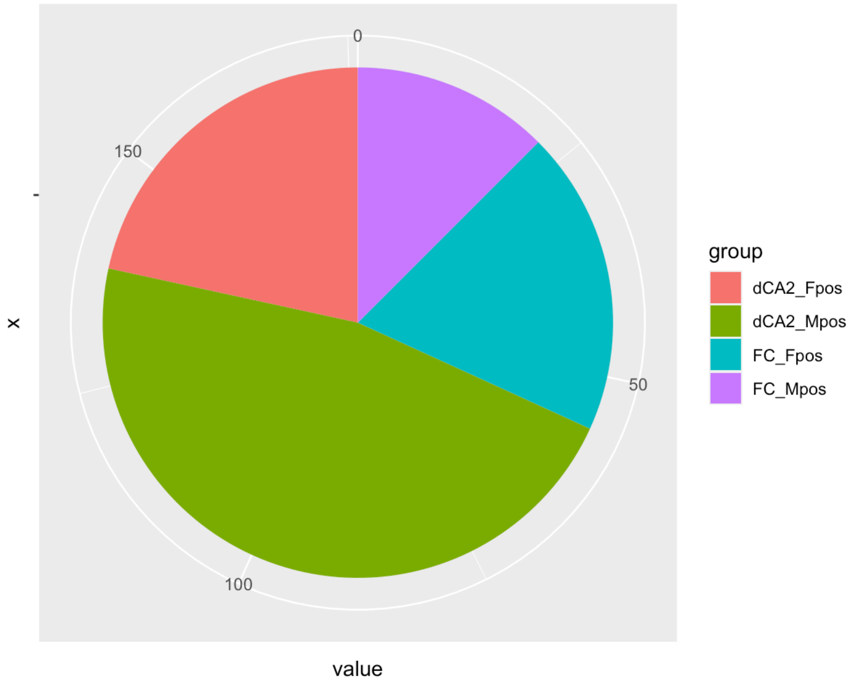


```
ggsave(file="/Users/lees44/R_project/FC_vs_dCA2_comparison_vector_integration_042821/cell_number_group_type6_pie.
pdf",width=10,height=7)
```

```
#type7
df <-data.frame(
  group = c("dCA2_Fpos", "dCA2_Mpos", "FC_Fpos", "FC_Mpos"),
  value = c(8, 25, 129, 11)
)
head(df)
```

```
##        group value
## 1 dCA2_Fpos     8
## 2 dCA2_Mpos    25
## 3   FC_Fpos   129
## 4   FC_Mpos    11
```

```
bp<- ggplot(df, aes(x="", y=value, fill=group))+
geom_bar(width = 1, stat = "identity")
bp
```

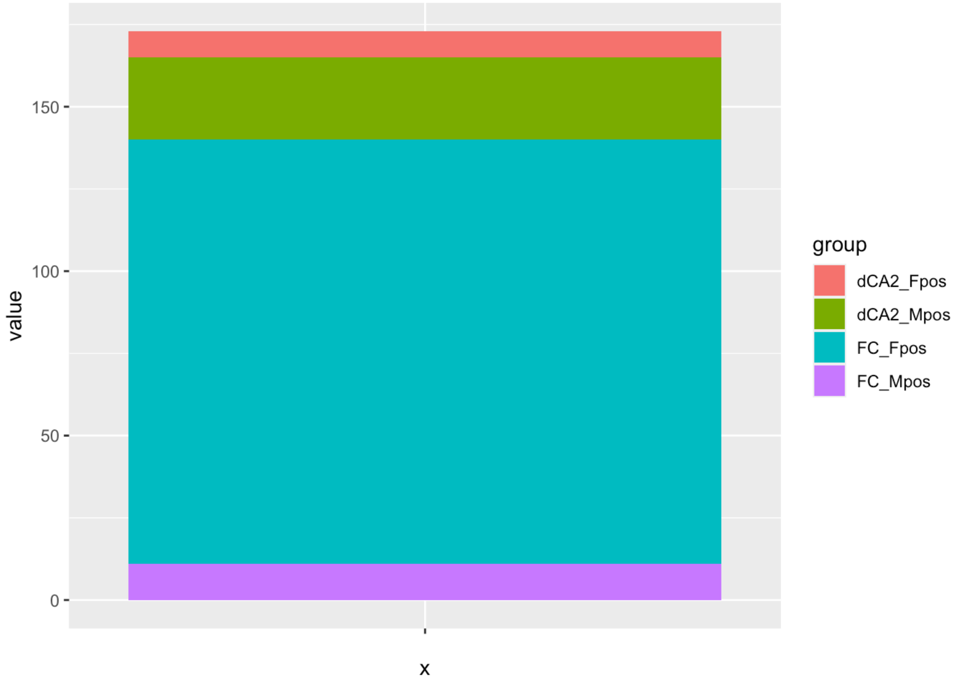


```
pie <- bp + coord_polar("y", start=0)
pie
```

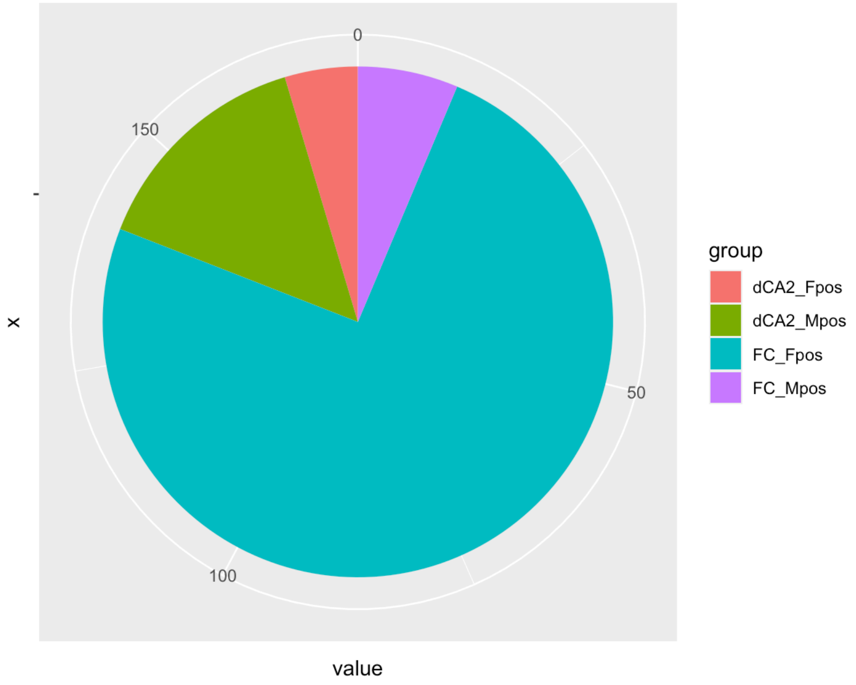

```
ggsave(file="/Users/lees44/R_project/FC_vs_dCA2_comparison_vector_integration_042821/cell_number_group_type7_pie.
pdf",width=10,height=7)
```

```
#type8
df <-data.frame(
  group = c("dCA2_Fpos", "dCA2_Mpos", "FC_Fpos", "FC_Mpos"),
  value = c(12, 16, 4, 3)
)
head(df)
```

```
##       group value
## 1 dCA2_Fpos    12
## 2 dCA2_Mpos    16
## 3   FC_Fpos     4
## 4   FC_Mpos     3
```

```
bp<- ggplot(df, aes(x="", y=value, fill=group))+
geom_bar(width = 1, stat = "identity")
bp
```

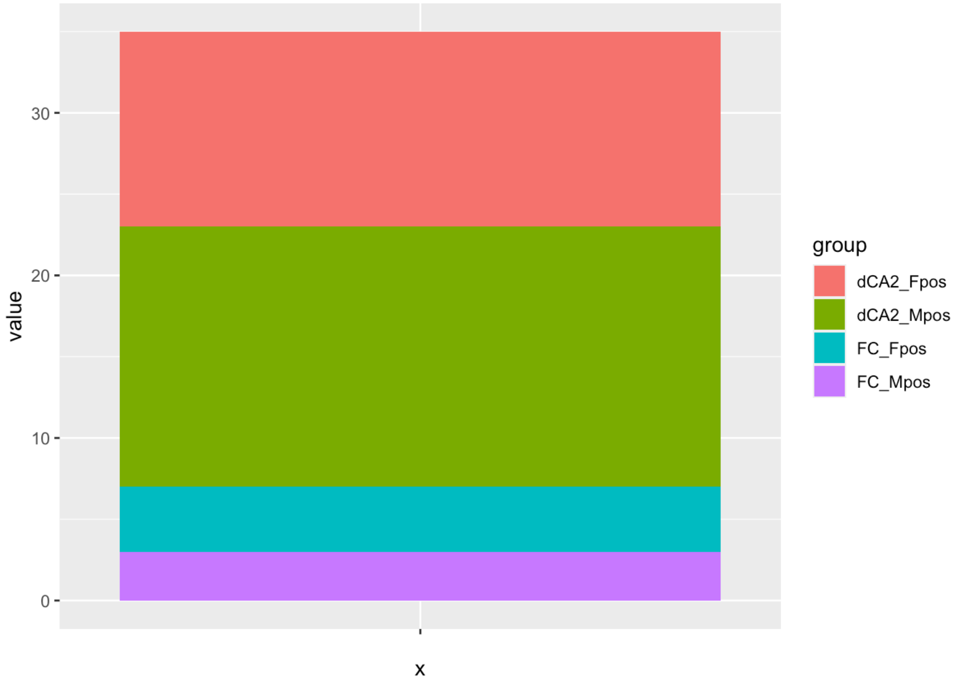


```
pie <- bp + coord_polar("y", start=0)
pie
```

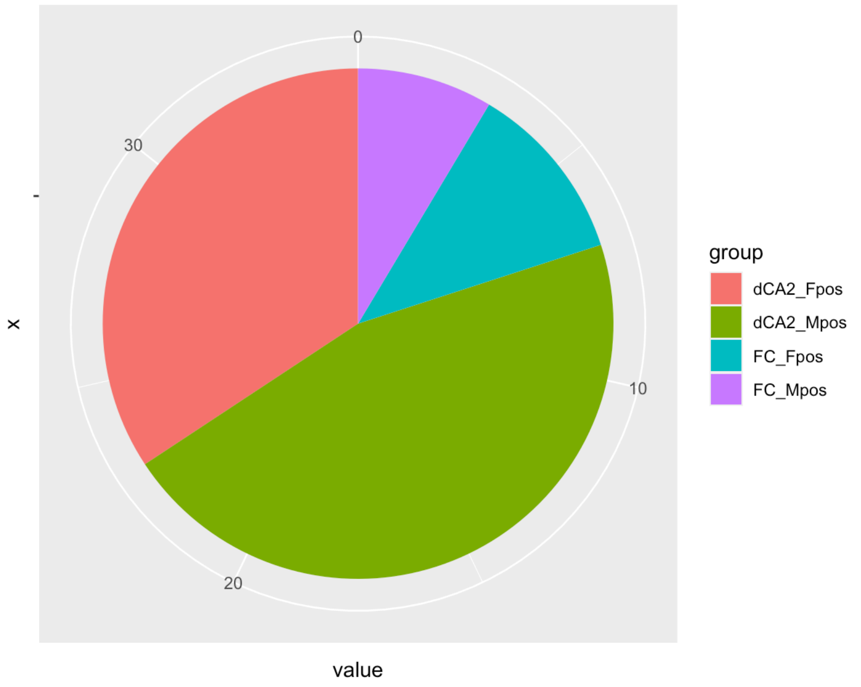


```
ggsave(file="/Users/lees44/R_project/FC_vs_dCA2_comparison_vector_integration_042821/cell_number_group_type8_pie.
pdf",width=10,height=7)
```

```
#type9
df <-data.frame(
  group = c("dCA2_Fpos", "dCA2_Mpos", "FC_Fpos", "FC_Mpos"),
  value = c(7, 12, 0, 0)
)
head(df)
```

```
##       group value
## 1 dCA2_Fpos     7
## 2 dCA2_Mpos    12
## 3   FC_Fpos     0
## 4   FC_Mpos     0
```

```
bp<- ggplot(df, aes(x="", y=value, fill=group))+
geom_bar(width = 1, stat = "identity")
bp
```

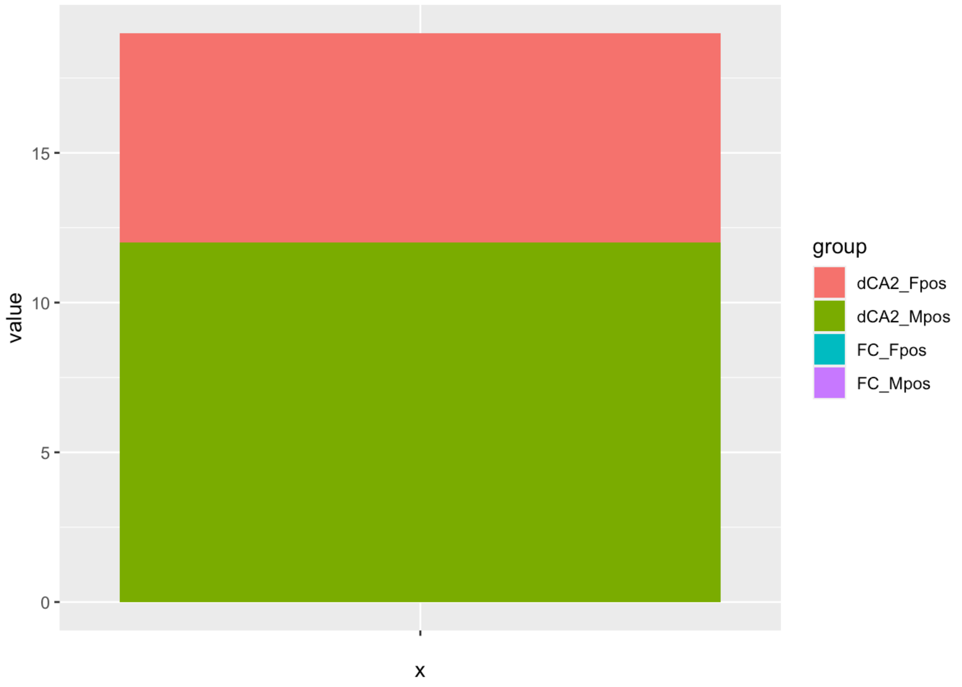


```
pie <- bp + coord_polar("y", start=0)
pie
```

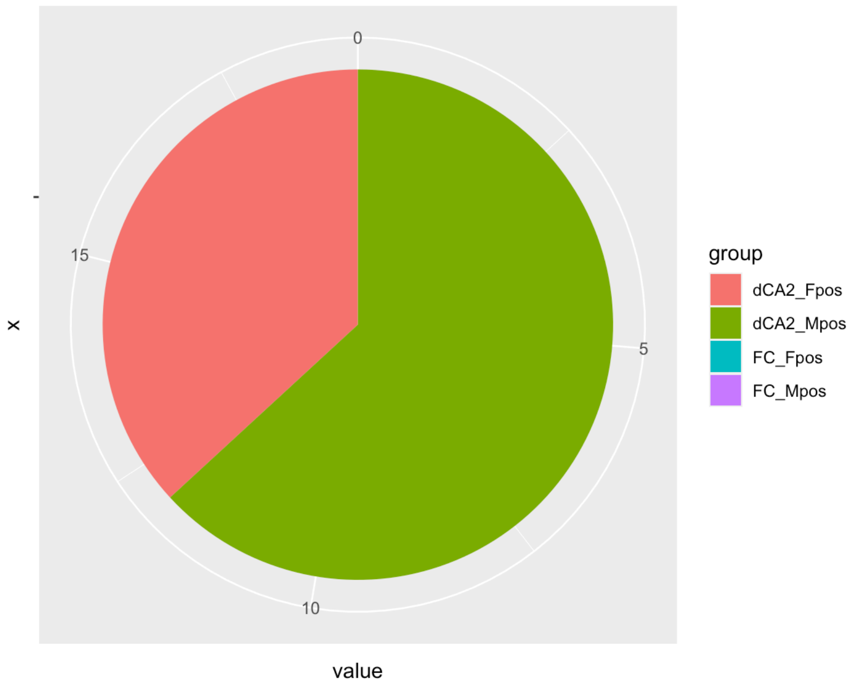


```
ggsave(file="/Users/lees44/R_project/FC_vs_dCA2_comparison_vector_integration_042821/cell_number_group_type9_pie.
pdf",width=10,height=7)
```

```
prop.table(table(Idents(dCA2.integrated), dCA2.integrated$protocol), margin = 2)
```

```
##
##          dCA2_Fpos   dCA2_Mpos      FC_Fpos      FC_Mpos
##   Type1 0.332161687 0.190812721 0.416347382 0.339501207
##   Type2 0.353251318 0.454316002 0.109195402 0.219629928
##   Type3 0.235500879 0.211004543 0.081098340 0.058728882
##   Type4 0.018453427 0.050984351 0.181992337 0.183427192
##   Type5 0.003514938 0.024734982 0.104725415 0.169750603
##   Type6 0.033391916 0.041393236 0.021711367 0.017699115
##   Type7 0.007029877 0.012619889 0.082375479 0.008849558
##   Type8 0.010544815 0.008076729 0.002554278 0.002413516
##   Type9 0.006151142 0.006057547 0.000000000 0.000000000
```

########percentage within the group###################

```
#percentage of nuclei dCA2_Fpos
df <-data.frame(
  group = c("Type1", "Type2", "Type3", "Type4", "Type5", "Type6", "Type7", "Type8", "Type9"),
  value = c(33.2, 35.3, 23.5, 1.84, 0.35, 3.3, 0.7, 1, 0.6)
)
head(df)
```

```
##   group value
## 1 Type1 33.20
## 2 Type2 35.30
## 3 Type3 23.50
## 4 Type4  1.84
## 5 Type5  0.35
## 6 Type6  3.30
```

```
bp<- ggplot(df, aes(x="", y=value, fill=group))+
geom_bar(width = 1, stat = "identity")+ ggtitle("dCA2_Fpos")
bp
```

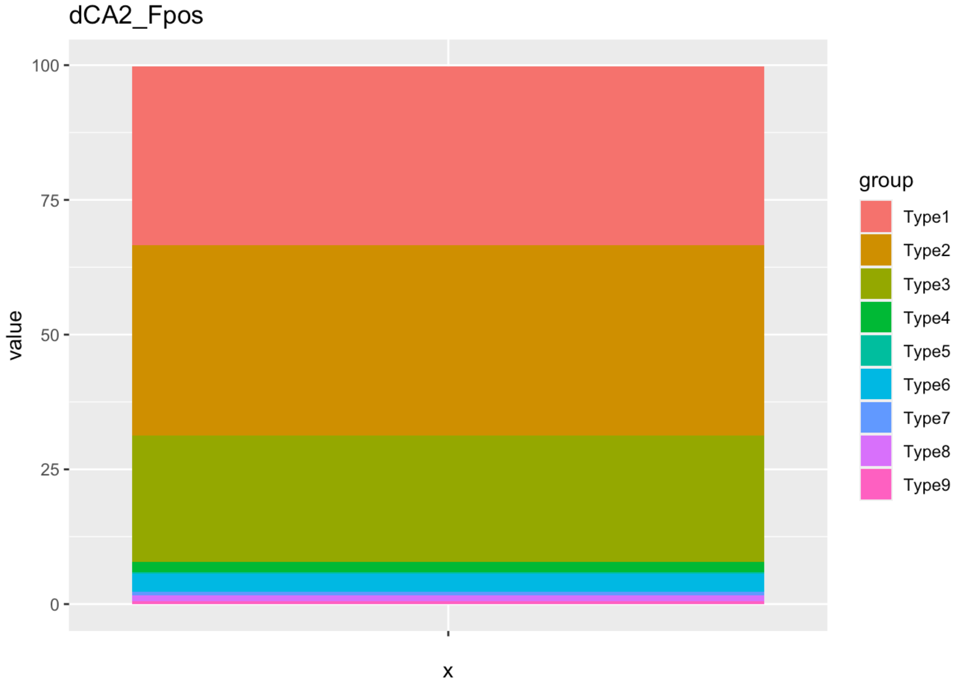


```
pie <- bp + coord_polar("y", start=0)
pie
```

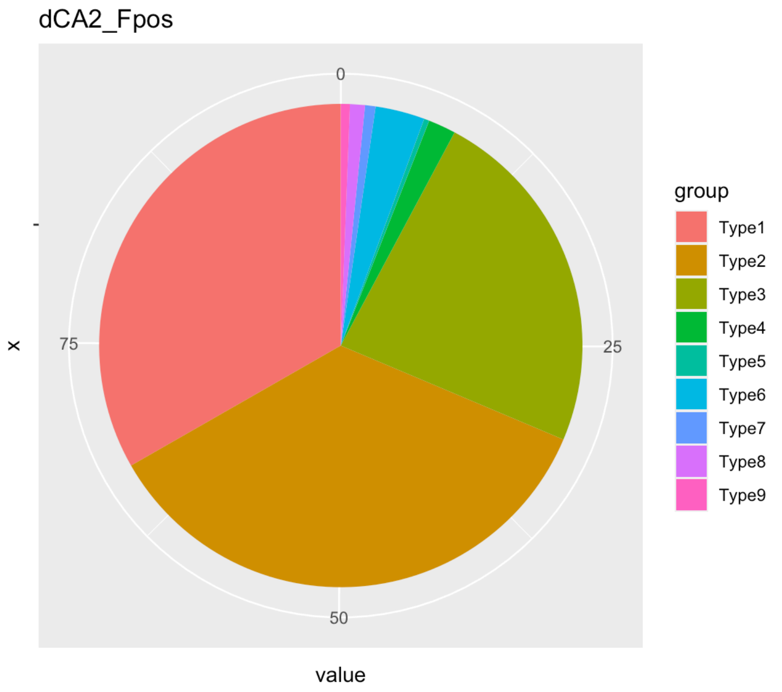


```
ggsave(file="/Users/lees44/R_project/FC_vs_dCA2_comparison_vector_integration_042821/cell_type_percentage_dCA2_Fp
os.pdf",width=10,height=7)
```

```
#percentage of nuclei dCA2_Mpos
df <-data.frame(
  group = c("Type1", "Type2", "Type3", "Type4", "Type5", "Type6", "Type7", "Type8", "Type9"),
  value = c(19.1, 45.4, 21.1, 5.1, 2.5, 4.1, 1.3, 0.8, 0.6)
)
head(df)
```

```
##   group value
## 1 Type1  19.1
## 2 Type2  45.4
## 3 Type3  21.1
## 4 Type4   5.1
## 5 Type5   2.5
## 6 Type6   4.1
```

```
bp<- ggplot(df, aes(x="", y=value, fill=group))+
geom_bar(width = 1, stat = "identity")+ ggtitle("dCA2_Mpos")
bp
```

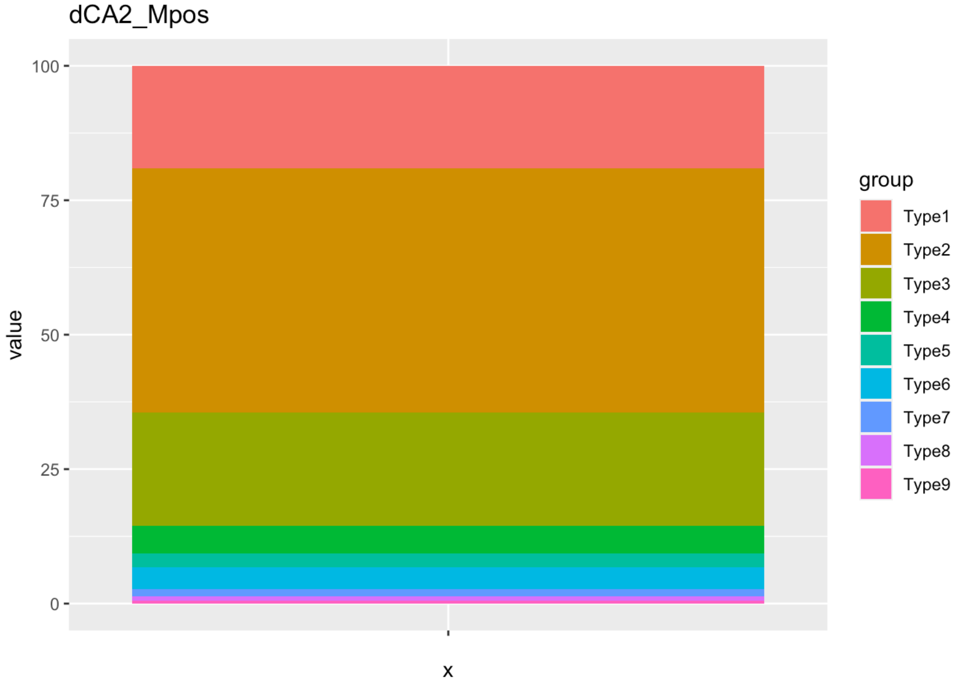


```
pie <- bp + coord_polar("y", start=0)
pie
```

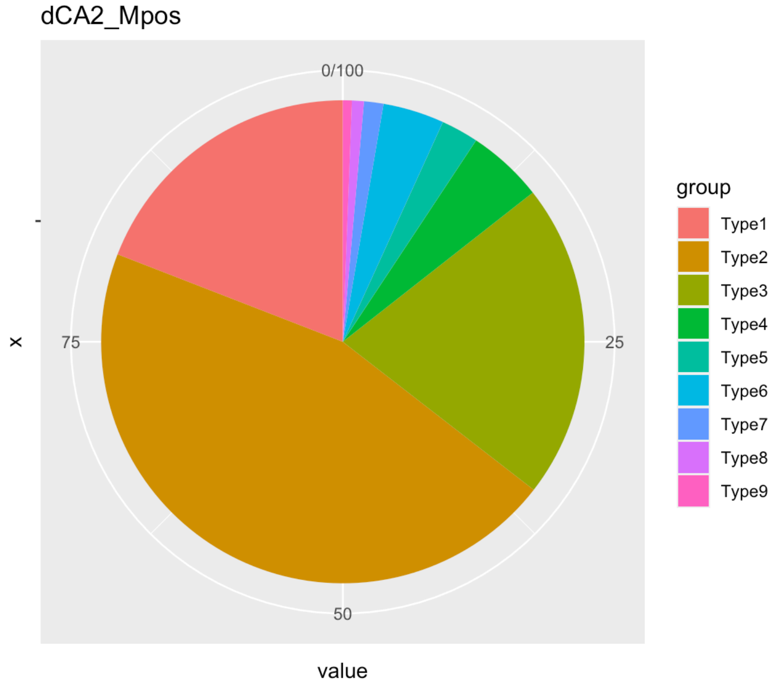


```
ggsave(file="/Users/lees44/R_project/FC_vs_dCA2_comparison_vector_integration_042821/cell_type_percentage_dCA2_Mp
os.pdf",width=10,height=7)
```

```
#percentage of nuclei FC_Fpos
df <-data.frame(
  group = c("Type1", "Type2", "Type3", "Type4", "Type5", "Type6", "Type7", "Type8", "Type9"),
  value = c(41.6, 10.9, 8.1, 18.2, 10.5, 2.2, 8.2, 0.3, 0)
)
head(df)
```

```
##   group value
## 1 Type1  41.6
## 2 Type2  10.9
## 3 Type3   8.1
## 4 Type4  18.2
## 5 Type5  10.5
## 6 Type6   2.2
```

```
bp<- ggplot(df, aes(x="", y=value, fill=group))+
geom_bar(width = 1, stat = "identity")+ ggtitle("FC_Fpos")
bp
```

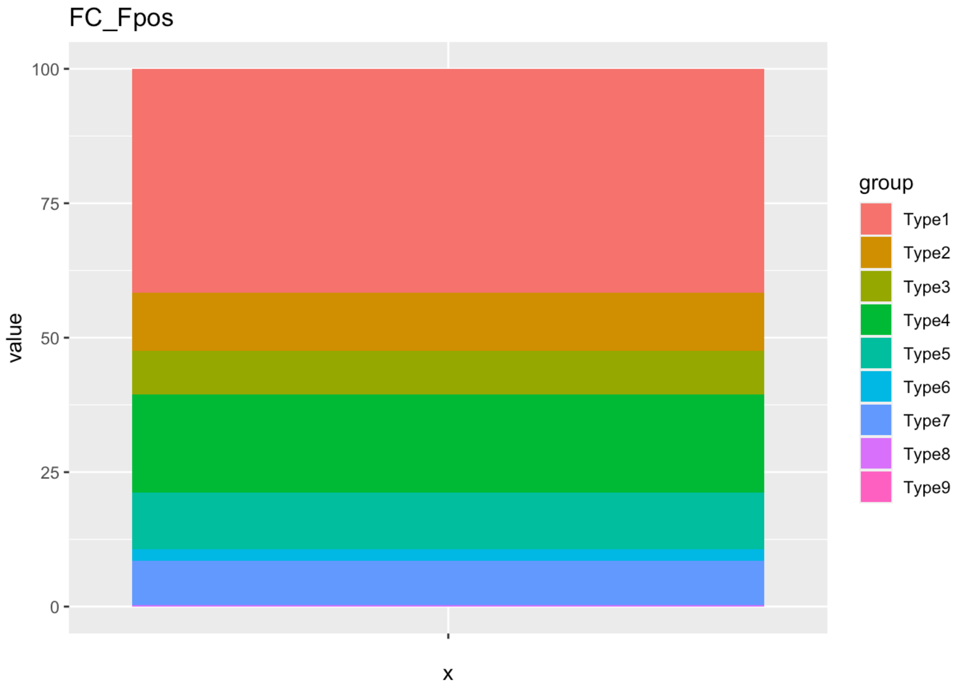


```
pie <- bp + coord_polar("y", start=0)
pie
```

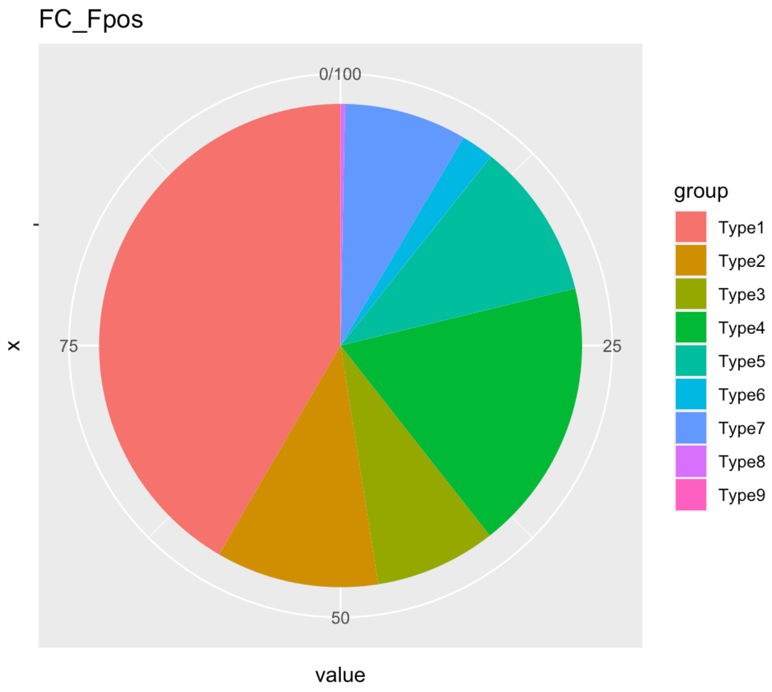


```
ggsave(file="/Users/lees44/R_project/FC_vs_dCA2_comparison_vector_integration_042821/cell_type_percentage_FC_Fpo
s.pdf",width=10,height=7)
```

```
#percentage of nuclei FC_Mpos
df <-data.frame(
  group = c("Type1", "Type2", "Type3", "Type4", "Type5", "Type6", "Type7", "Type8", "Type9"),
  value = c(34.2, 22.1, 5.9, 18.5, 17, 1.8, 0.9, 0.2, 0)
)
head(df)
```

```
##   group value
## 1 Type1  34.2
## 2 Type2  22.1
## 3 Type3   5.9
## 4 Type4  18.5
## 5 Type5  17.0
## 6 Type6   1.8
```

```
bp<- ggplot(df, aes(x="", y=value, fill=group))+
geom_bar(width = 1, stat = "identity")+ ggtitle("FC_Mpos")
bp
```

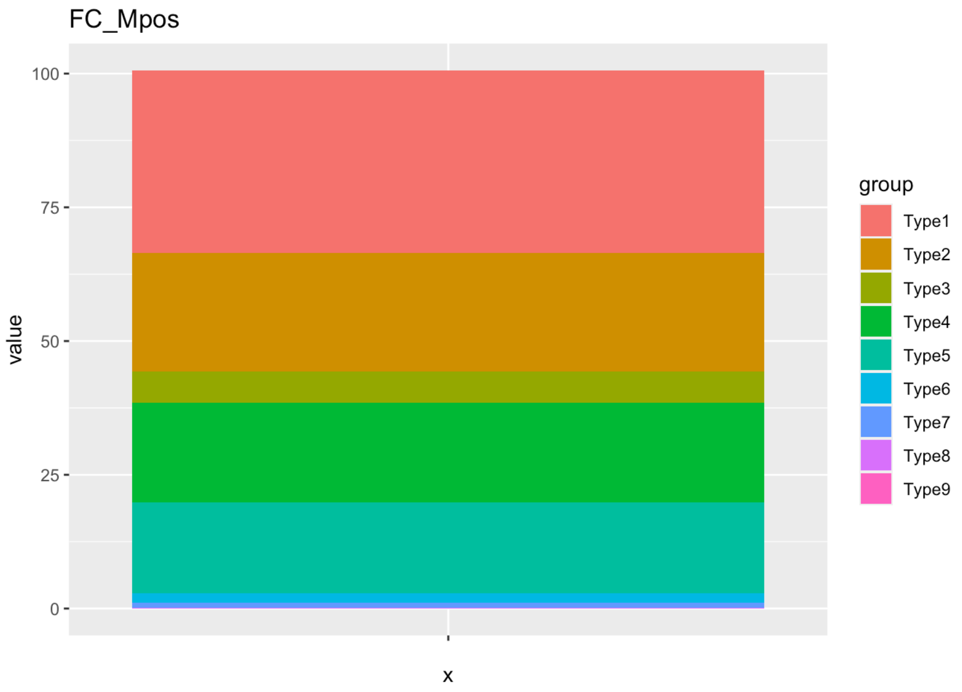


```
pie <- bp + coord_polar("y", start=0)
pie
```

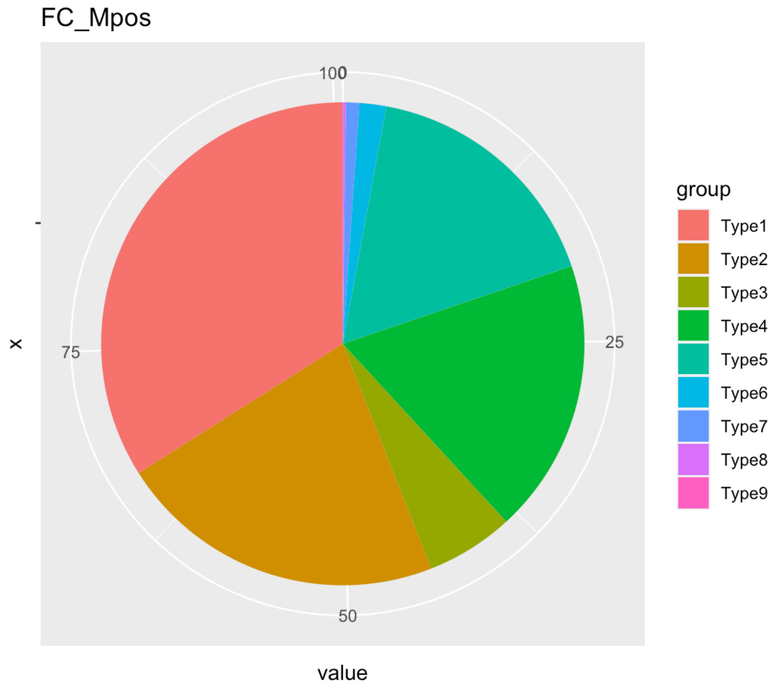


```
ggsave(file="/Users/lees44/R_project/FC_vs_dCA2_comparison_vector_integration_042821/cell_type_percentage_FC_Mpo
s.pdf",width=10,height=7)
```

```
#percentage of nuclei dCA2
df <-data.frame(
  group = c("Type1", "Type2", "Type3", "Type4", "Type5", "Type6", "Type7", "Type8", "Type9"),
  value = c(24.2, 41.7, 22, 4, 1.7, 3.8, 1, 0.9, 0.6)
)
head(df)
```

```
##   group value
## 1 Type1  24.2
## 2 Type2  41.7
## 3 Type3  22.0
## 4 Type4   4.0
## 5 Type5   1.7
## 6 Type6   3.8
```

```
bp<- ggplot(df, aes(x="", y=value, fill=group))+
geom_bar(width = 1, stat = "identity")+ ggtitle("dCA2")
bp
```

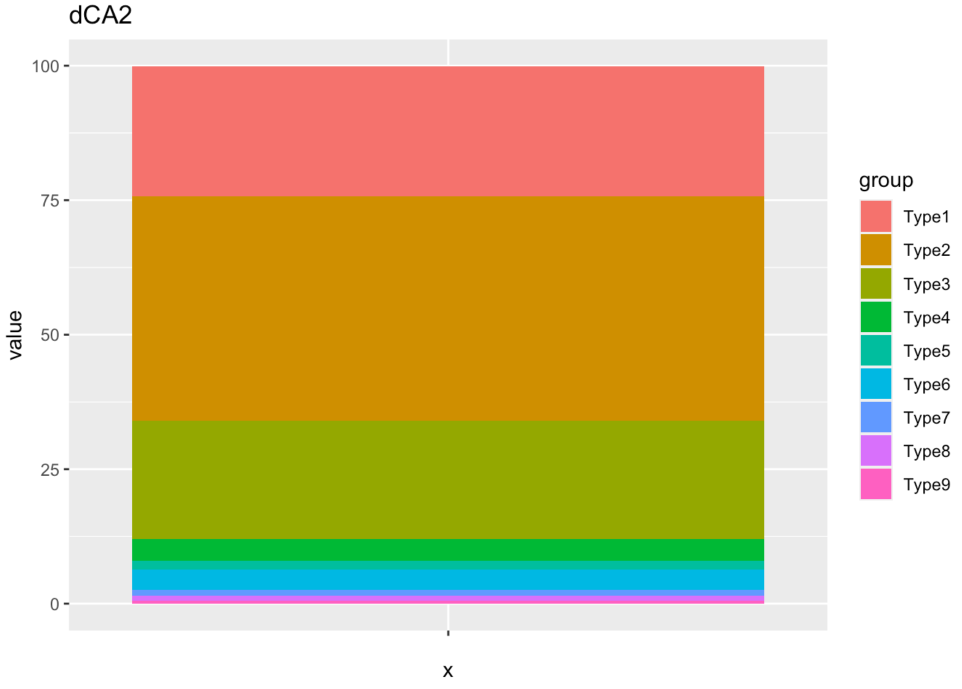


```
pie <- bp + coord_polar("y", start=0)
pie
```

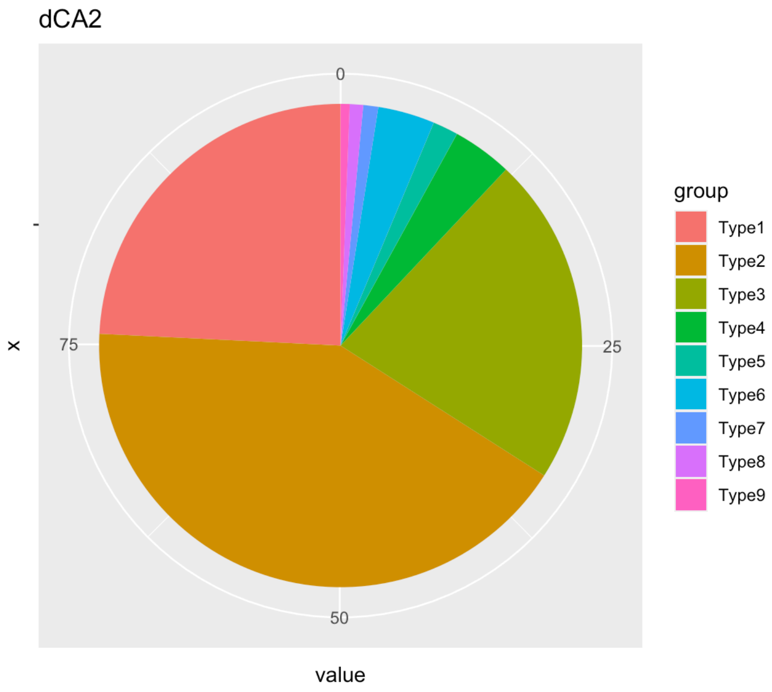


```
ggsave(file="/Users/lees44/R_project/FC_vs_dCA2_comparison_vector_integration_042821/cell_type_percentage_dCA2.pd
f",width=10,height=7)
```

```
#percentage of nuclei FC
df <-data.frame(
  group = c("Type1", "Type2", "Type3", "Type4", "Type5", "Type6", "Type7", "Type8", "Type9"),
  value = c(38, 16, 7.1, 18, 13, 2, 5, 0.3, 0)
)
head(df)
```

```
##   group value
## 1 Type1  38.0
## 2 Type2  16.0
## 3 Type3   7.1
## 4 Type4  18.0
## 5 Type5  13.0
## 6 Type6   2.0
```

```
bp<- ggplot(df, aes(x="", y=value, fill=group))+
geom_bar(width = 1, stat = "identity")+ ggtitle("FC")
bp
```

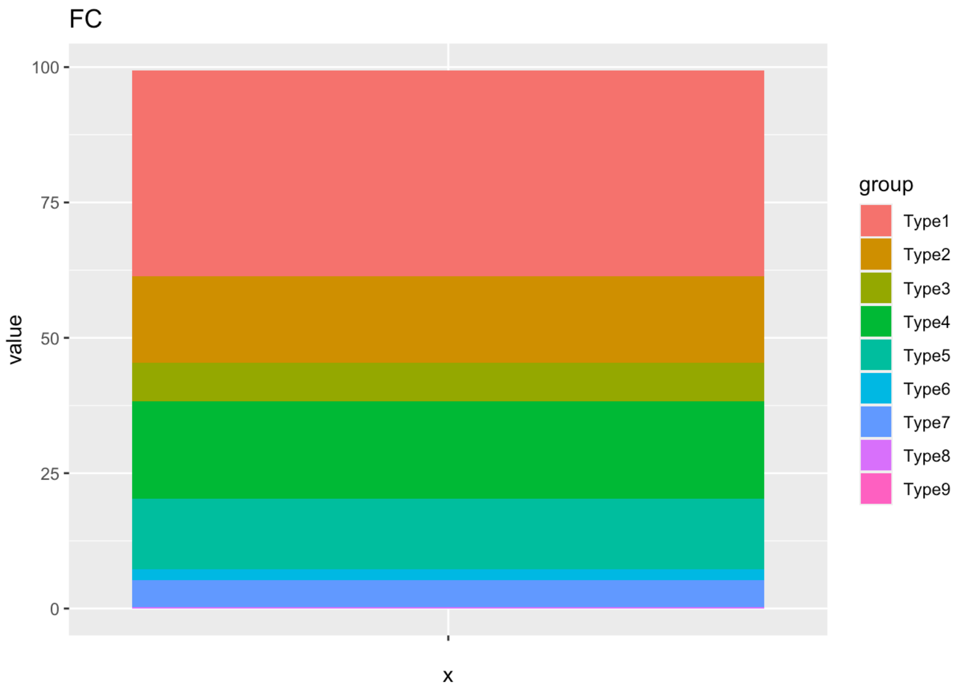


```
pie <- bp + coord_polar("y", start=0)
pie
```

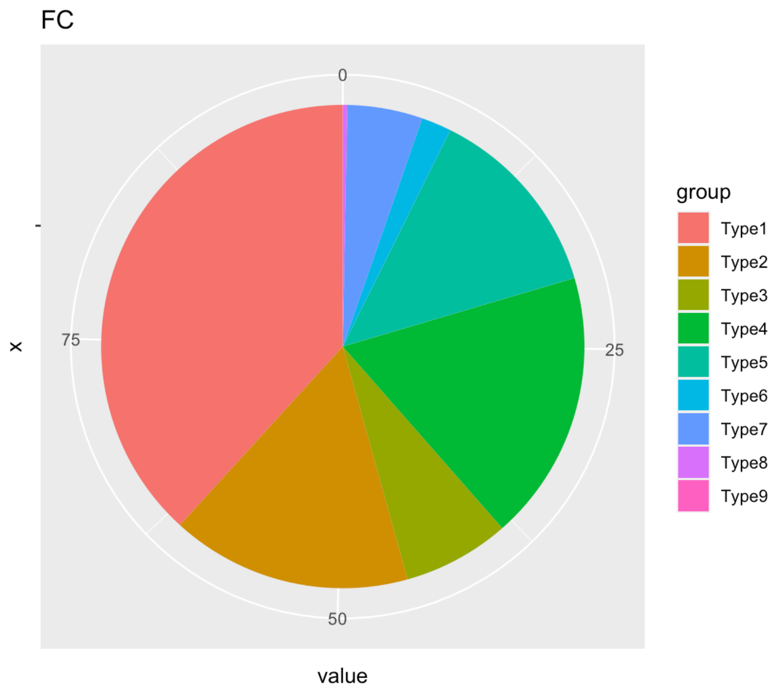


```
ggsave(file="/Users/lees44/R_project/FC_vs_dCA2_comparison_vector_integration_042821/cell_type_percentage_FC.pdf",
width=10,height=7)
```

“CREREV38354884”, “GFP”,“Rgs14”, “Avpr1b”, “Amigo2”, “Map3k15”, “Actn2”, “Ptpn5”, “Cacng5”, “Oxtr”, “Pcp4”

Violinplot for dCA2 markers, figure goes with cluster tree

```
Cell_type<-rev(c("Type1", "Type2", "Type3", "Type4", "Type5", "Type6", "Type7", "Type8", "Type9"))
gene_list<-c("Rgs14" ,"Avpr1b","Amigo2", "Map3k15","Actn2", "Ptpn5", "Cacng5","Oxtr", "Pcp4")
```

```
Cell_number <- NULL
for (i in 1:length(Cell_type)){
                temp1 <- data.frame(type=factor(rep(Cell_type[i],sum(Cell_type[i]==Idents(dCA2.integrated)))))
                temp2 <- as.data.frame(dCA2.integrated@assays$RNA@data[gene_list,Idents(dCA2.integrated)==Cell_ty
pe[i]])
                temp3 <- cbind(temp1,t(temp2))
                Cell_number <- rbind(temp3,Cell_number)
}
rownames(Cell_number) <- NULL
```

```
colors<-c("#ffc857","#ffc857","#ffc857","#9B0C1E","#9B0C1E","#9B0C1E","#9B0C1E","#9B0C1E","#9B0C1E")
```

```
for (k in 1:length(gene_list))
    {
if (k==length(gene_list)){
assign(paste("P",k,sep=""),ggplot(Cell_number,aes_string(x="type",y=gene_list[k],fill="type"))+geom_violin(scale
= "width")+scale_fill_manual(values=colors)
+stat_summary(fun=median, geom="point", size=0.6, color="red")+ylab(gene_list[k])+ theme(axis.title.x=element_bla
nk(),
axis.text.y=element_text(size=5),axis.title.y=element_text(size=6,angle=0,face="bold",margin = margin(t = 10, r =
8, b = 0, l = 10),vjust=0.5),axis.text.x=element_text(size=5,face="bold")
,axis.title=element_text(size=4,face="bold"),panel.grid.major = element_blank(), panel.grid.minor = element_blan
k(),
panel.background = element_blank(), axis.line = element_line(colour = "black"),legend.position="none",plot.margin
= unit(c(0, 0,0, 0), "cm")))}
else{
assign(paste("P",k,sep=""),ggplot(Cell_number,aes_string(x="type",y=gene_list[k],fill="type"))+geom_violin(scale
= "width")+scale_fill_manual(values=colors)
+stat_summary(fun=median, geom="point", size=0.6, color="red")+ylab(gene_list[k])+ theme(axis.title.x=element_bla
nk(),
axis.text.x=element_blank(),axis.text.y=element_text(size=5),axis.title.y=element_text(size=6,angle=0,face="bold",
margin = margin(t = 10, r = 8, b = 0, l = 10),vjust=0.5),axis.ticks.x=element_blank(),axis.text=element_text(size
=5)
,axis.title=element_text(size=4,face="bold"),panel.grid.major = element_blank(), panel.grid.minor = element_blan
k(),
panel.background = element_blank(), axis.line = element_line(colour = "black"),legend.position="none",plot.margin
= unit(c(0, 0,0, 0), "cm")))}

}
```

```
P1
```

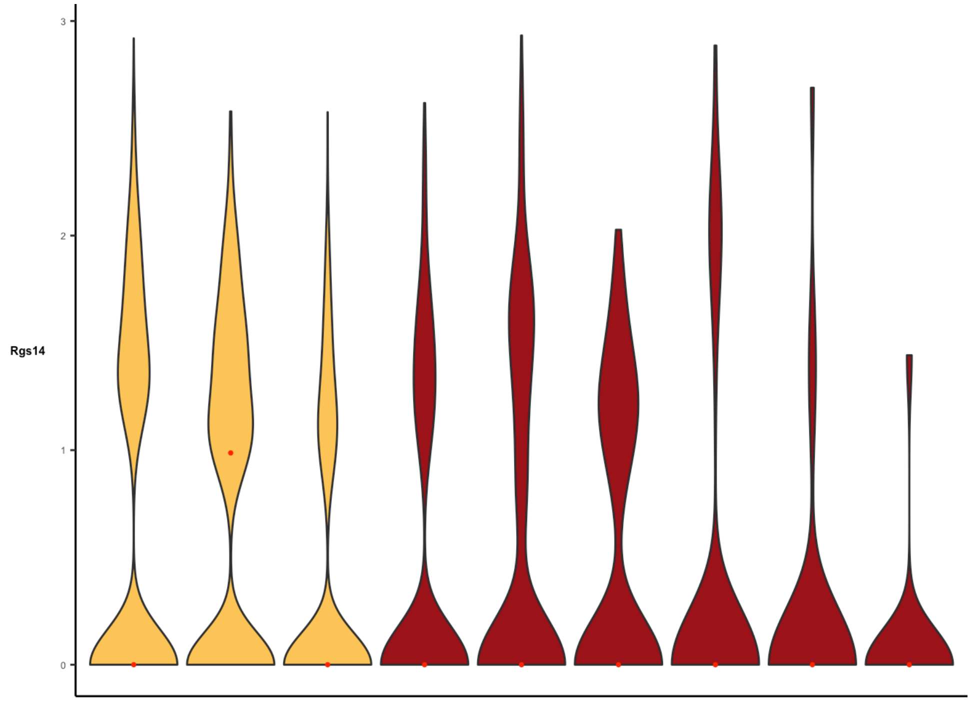


```
P2
```

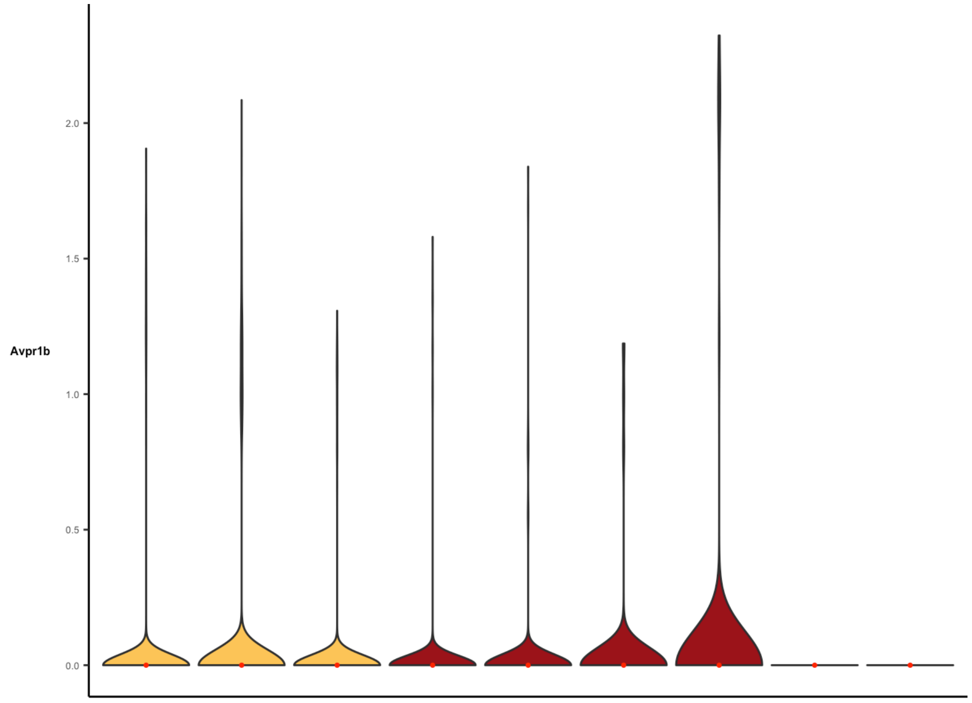


```
P3
```

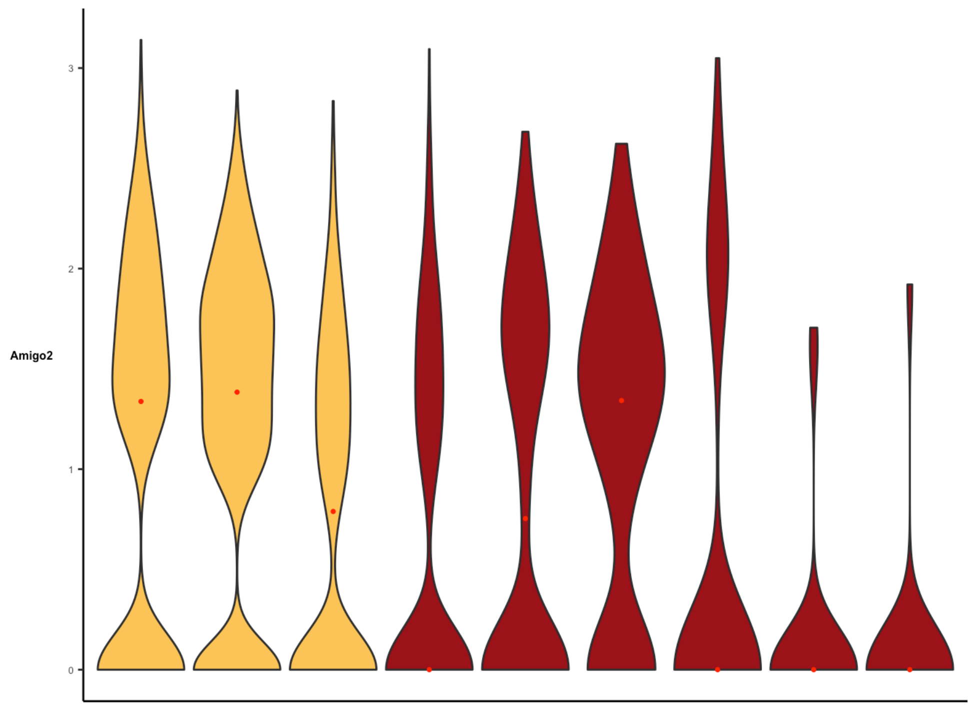


```
P4
```

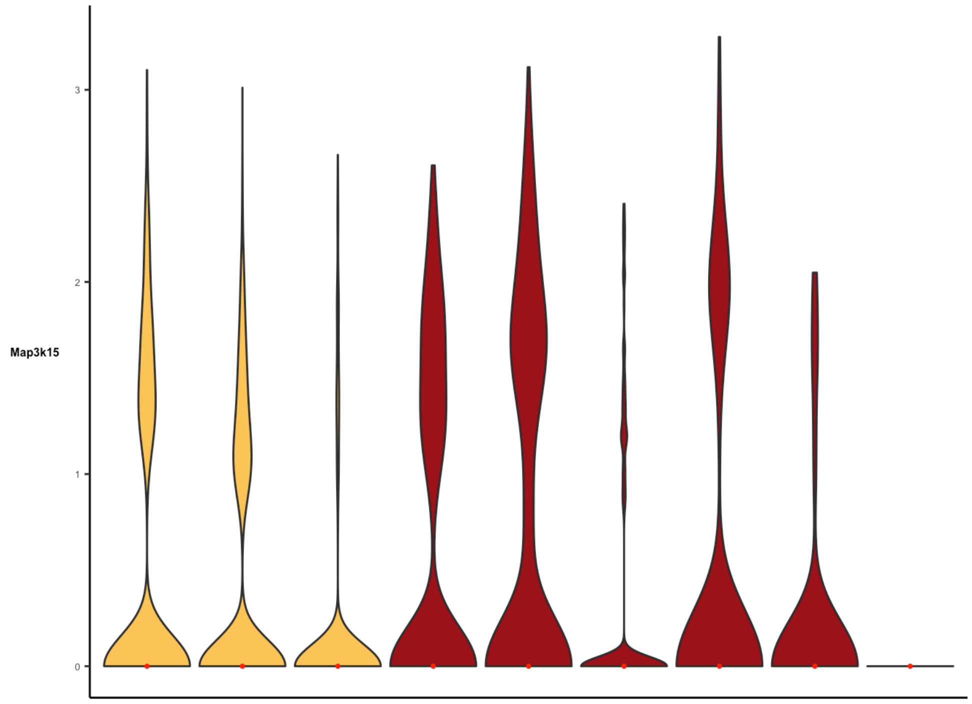


```
P5
```

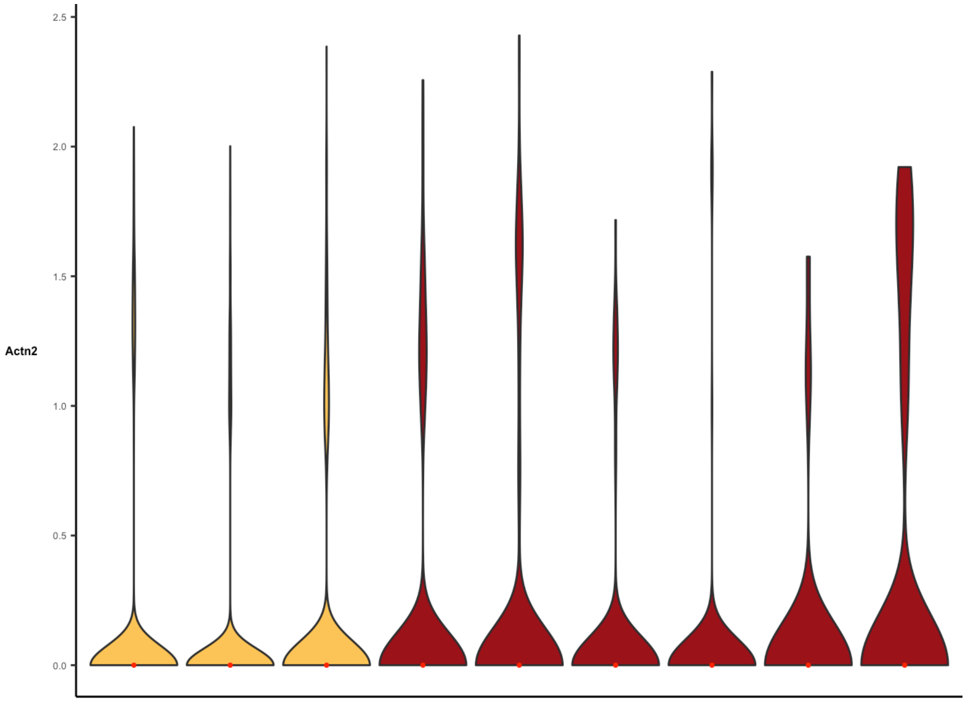


```
P6
```

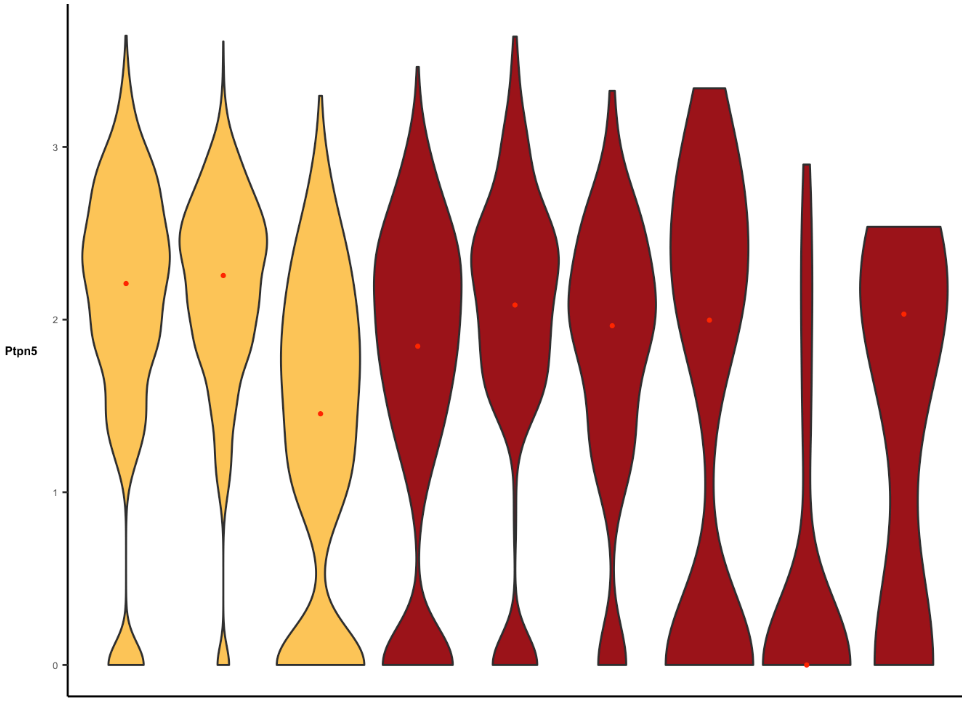


```
P7
```

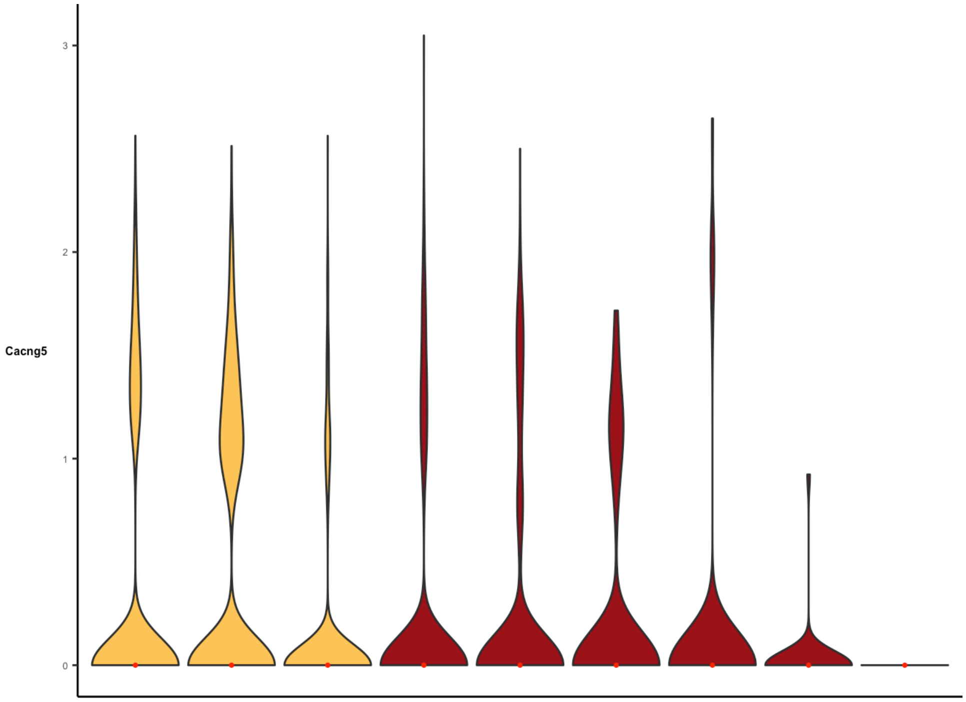


```
P8
```

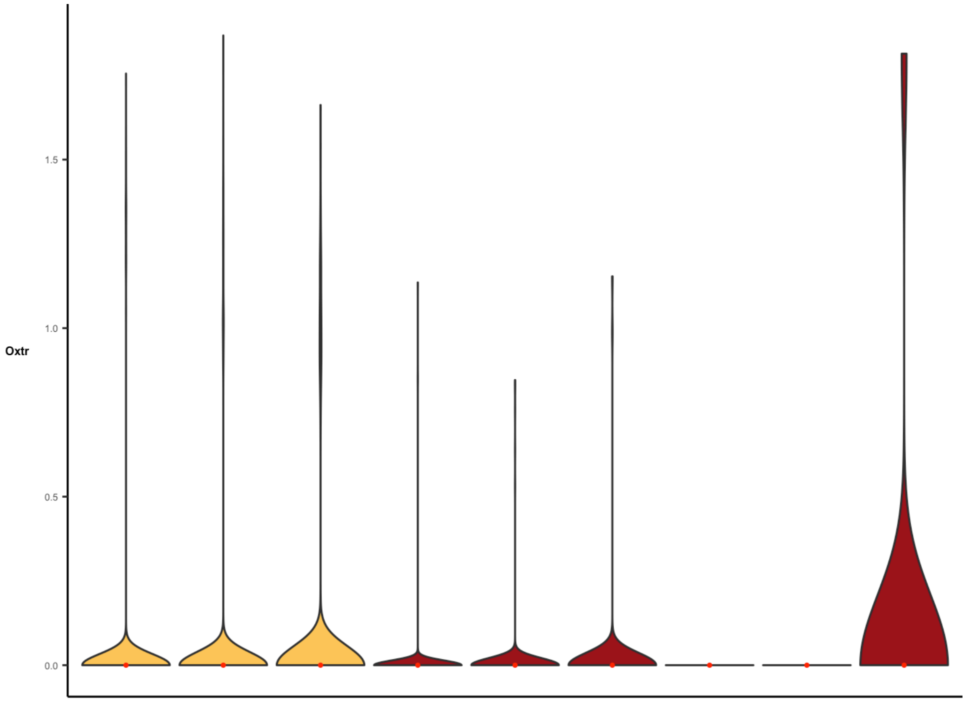


```
P9
```

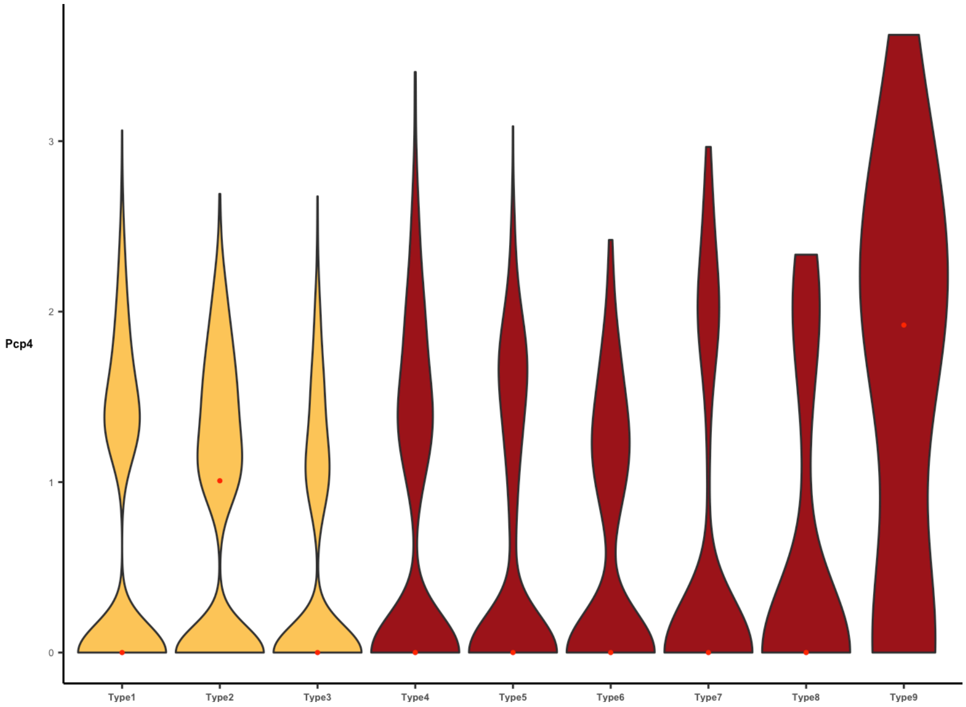


```
merge<-list()
for (i in length(gene_list):1){
       if (length(merge)==0){
        merge<-ggplotGrob(eval(parse(text=paste("P",i,sep = ""))))}else{merge<-rbind(ggplotGrob(eval(parse(text=p
aste("P",i,sep = ""))),merge,size = "last")}}

 #all genes
pdf(paste("/Users/lees44/R_project/FC_vs_dCA2_comparison_vector_integration_042821/cluster_tree_CJ.pdf",".png",se
p=""),height=6, width=50 , paper = "letter")


grid.newpage()
grid.draw(merge)
```

#########violin plot for Sodium channels ###################### Violinplot for sodium channels, figure goes with cluster tree

```
Cell_type<-rev(c("Type1", "Type2", "Type3", "Type4", "Type5", "Type6", "Type7", "Type8", "Type9"))
gene_list<-c("Scn1a" ,"Scn2a","Scn3a", "Scn8a","Scn9a")
```

```
Cell_number <- NULL
for (i in 1:length(Cell_type)){
                temp1 <- data.frame(type=factor(rep(Cell_type[i],sum(Cell_type[i]==Idents(dCA2.integrated)))))
                temp2 <- as.data.frame(dCA2.integrated@assays$RNA@data[gene_list,Idents(dCA2.integrated)==Cell_ty
pe[i]])
                temp3 <- cbind(temp1,t(temp2))
                Cell_number <- rbind(temp3,Cell_number)
}
rownames(Cell_number) <- NULL
```

```
colors<-c("#ffc857","#ffc857","#ffc857","#9B0C1E","#9B0C1E","#9B0C1E","#9B0C1E","#9B0C1E","#9B0C1E")
```

```
for (k in 1:length(gene_list))
    {
if (k==length(gene_list)){
assign(paste("P",k,sep=""),ggplot(Cell_number,aes_string(x="type",y=gene_list[k],fill="type"))+geom_violin(scale
= "width")+scale_fill_manual(values=colors)
+stat_summary(fun=median, geom="point", size=0.6, color="red")+ylab(gene_list[k])+ theme(axis.title.x=element_bla
nk(),
axis.text.y=element_text(size=5),axis.title.y=element_text(size=6,angle=0,face="bold",margin = margin(t = 10, r =
8, b = 0, l = 10),vjust=0.5),axis.text.x=element_text(size=5,face="bold")
,axis.title=element_text(size=4,face="bold"),panel.grid.major = element_blank(), panel.grid.minor = element_blan
k(),
panel.background = element_blank(), axis.line = element_line(colour = "black"),legend.position="none",plot.margin
= unit(c(0, 0,0, 0), "cm")))}
else{
assign(paste("P",k,sep=""),ggplot(Cell_number,aes_string(x="type",y=gene_list[k],fill="type"))+geom_violin(scale
= "width")+scale_fill_manual(values=colors)
+stat_summary(fun=median, geom="point", size=0.6, color="red")+ylab(gene_list[k])+ theme(axis.title.x=element_bla
nk(),
axis.text.x=element_blank(),axis.text.y=element_text(size=5),axis.title.y=element_text(size=6,angle=0,face="bold",
margin = margin(t = 10, r = 8, b = 0, l = 10),vjust=0.5),axis.ticks.x=element_blank(),axis.text=element_text(size
=5)
,axis.title=element_text(size=4,face="bold"),panel.grid.major = element_blank(), panel.grid.minor = element_blan
k(),
panel.background = element_blank(), axis.line = element_line(colour = "black"),legend.position="none",plot.margin
= unit(c(0, 0,0, 0), "cm")))}

}
```

```
P1
```

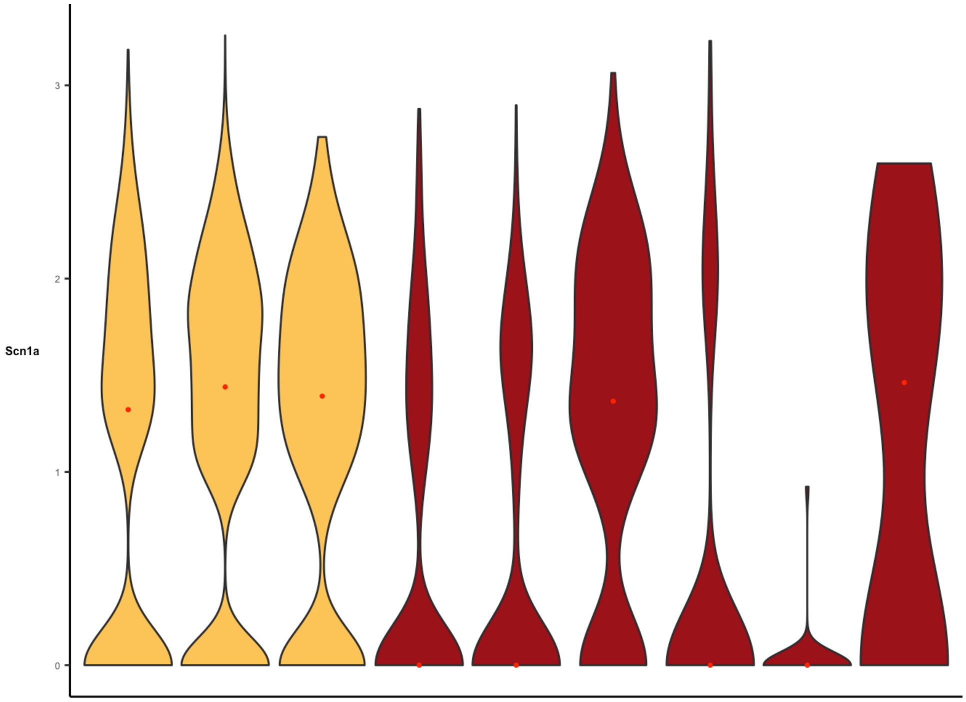


```
P2
```

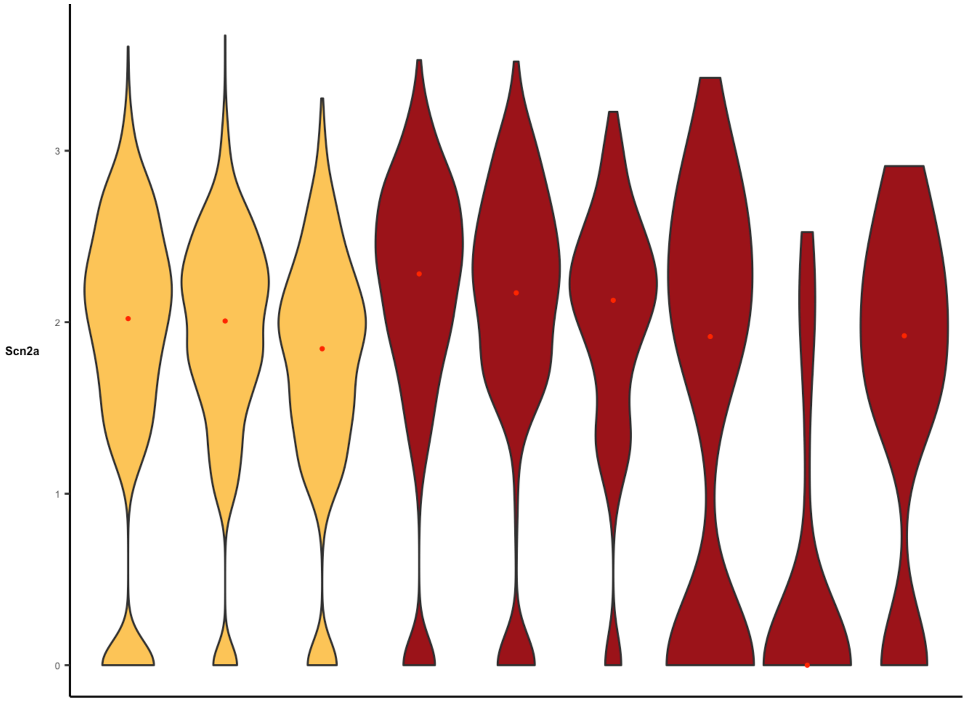


```
P3
```

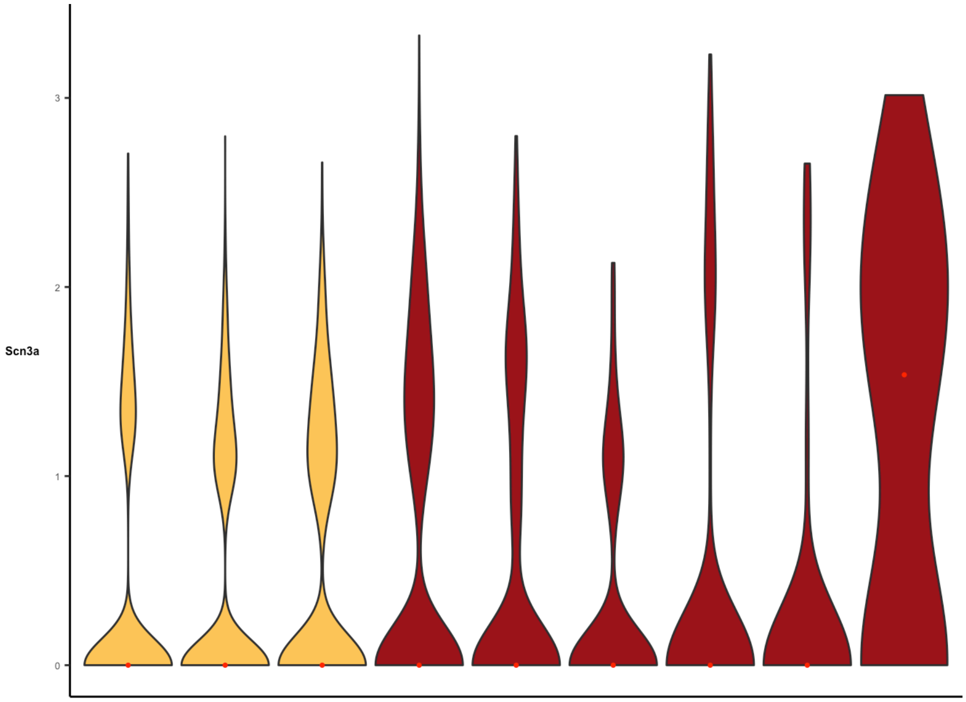


```
P4
```

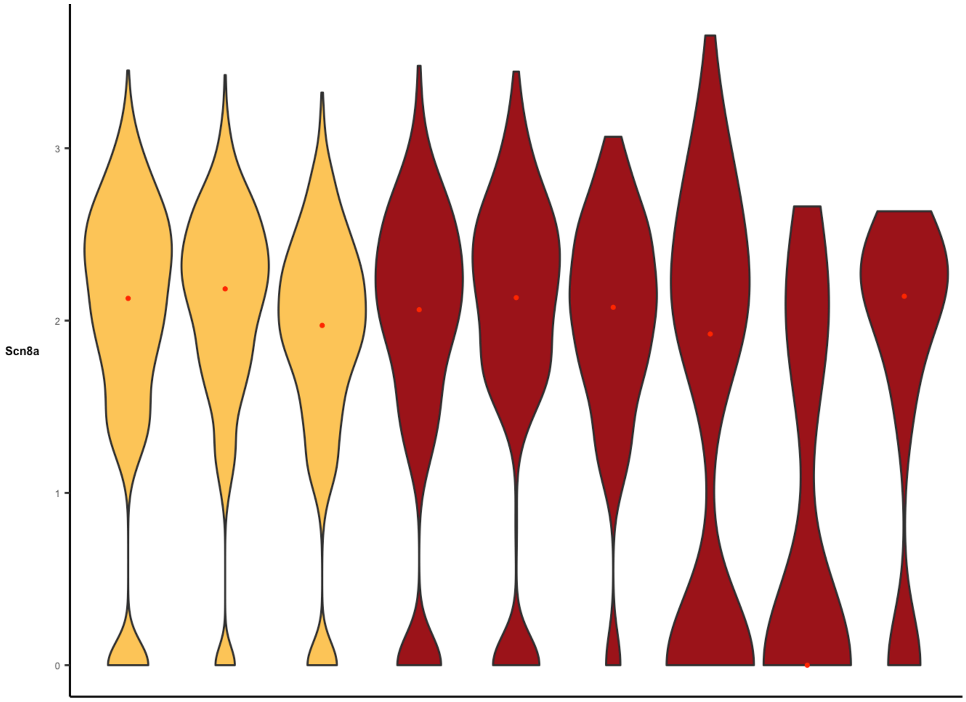
Scn8a
0
1
2
3

```
P5
```

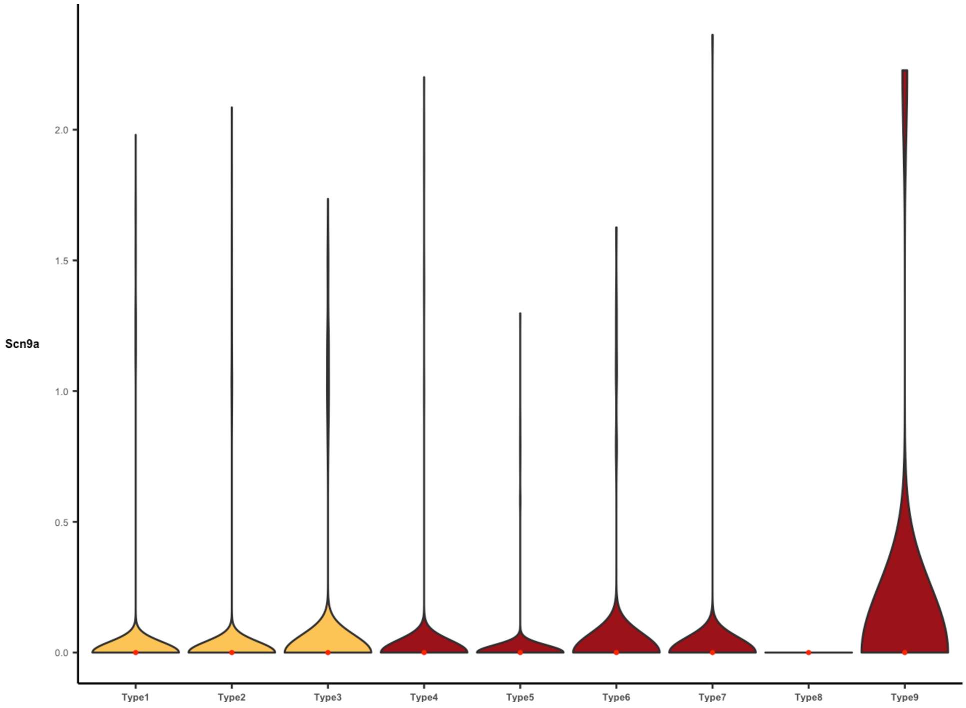
Scn9a
0.0
0.5
1.0
1.5
2.0
Type1
Type2
Type3
Type4
Type5
Type6
Type7
Type8
Type9

```
merge<-list()
for (i in length(gene_list):1){
       if (length(merge)==0){
        merge<-ggplotGrob(eval(parse(text=paste("P",i,sep = ""))))}else{merge<-rbind(ggplotGrob(eval(parse(text=p
aste("P",i,sep = "")))),merge,size = "last")}}

 #all genes
pdf(paste("/Users/lees44/R_project/FC_vs_dCA2_comparison_vector_integration_042821/cluster_tree_sodium_channel.pd
f",".png",sep=""),height=6, width=50 , paper = "letter")


grid.newpage()
grid.draw(merge)
```

#########violin plot for Potassium channels ##################### Violinplot for potassium channels, figure goes with cluster tree Kcna and Kcnb potassium channel

```
Cell_type<-rev(c("Type1", "Type2", "Type3", "Type4", "Type5", "Type6", "Type7", "Type8", "Type9"))
gene_list<-c("Kcna1","Kcna2", "Kcna3", "Kcna4", "Kcna6", "Kcnb1", "Kcnb2")
```

```
Cell_number <- NULL
for (i in 1:length(Cell_type)){
                temp1 <- data.frame(type=factor(rep(Cell_type[i],sum(Cell_type[i]==Idents(dCA2.integrated)))))
                temp2 <- as.data.frame(dCA2.integrated@assays$RNA@data[gene_list,Idents(dCA2.integrated)==Cell_ty
pe[i]])
                temp3 <- cbind(temp1,t(temp2))
                Cell_number <- rbind(temp3,Cell_number)
}
rownames(Cell_number) <- NULL
```

```
colors<-c("#ffc857","#ffc857","#ffc857","#9B0C1E","#9B0C1E","#9B0C1E","#9B0C1E","#9B0C1E","#9B0C1E")
```

```
for (k in 1:length(gene_list))
    {
if (k==length(gene_list)){
assign(paste("P",k,sep=""),ggplot(Cell_number,aes_string(x="type",y=gene_list[k],fill="type"))+geom_violin(scale
= "width")+scale_fill_manual(values=colors)
+stat_summary(fun=median, geom="point", size=0.6, color="red")+ylab(gene_list[k])+ theme(axis.title.x=element_bla
nk(),
axis.text.y=element_text(size=5),axis.title.y=element_text(size=6,angle=0,face="bold",margin = margin(t = 10, r =
8, b = 0, l = 10),vjust=0.5),axis.text.x=element_text(size=5,face="bold")
,axis.title=element_text(size=4,face="bold"),panel.grid.major = element_blank(), panel.grid.minor = element_blan
k(),
panel.background = element_blank(), axis.line = element_line(colour = "black"),legend.position="none",plot.margin
= unit(c(0, 0,0, 0), "cm")))}
else{
assign(paste("P",k,sep=""),ggplot(Cell_number,aes_string(x="type",y=gene_list[k],fill="type"))+geom_violin(scale
= "width")+scale_fill_manual(values=colors)
+stat_summary(fun=median, geom="point", size=0.6, color="red")+ylab(gene_list[k])+ theme(axis.title.x=element_bla
nk(),
axis.text.x=element_blank(),axis.text.y=element_text(size=5),axis.title.y=element_text(size=6,angle=0,face="bold",
margin = margin(t = 10, r = 8, b = 0, l = 10),vjust=0.5),axis.ticks.x=element_blank(),axis.text=element_text(size
=5)
,axis.title=element_text(size=4,face="bold"),panel.grid.major = element_blank(), panel.grid.minor = element_blan
k(),
panel.background = element_blank(), axis.line = element_line(colour = "black"),legend.position="none",plot.margin
= unit(c(0, 0,0, 0), "cm")))}

}
```

```
P1
```

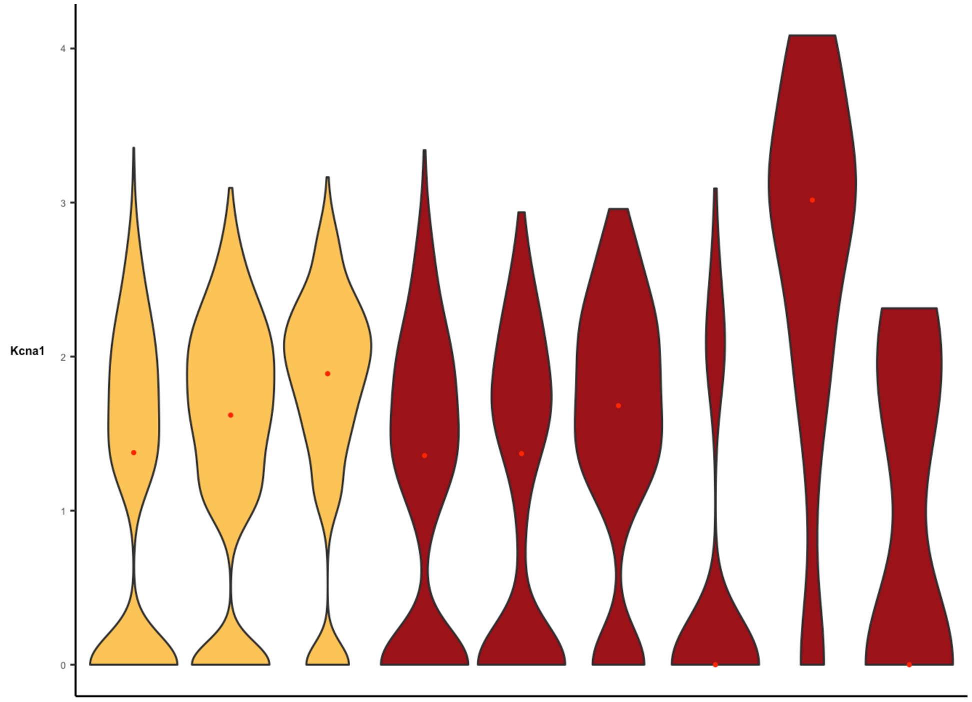


```
P2
```

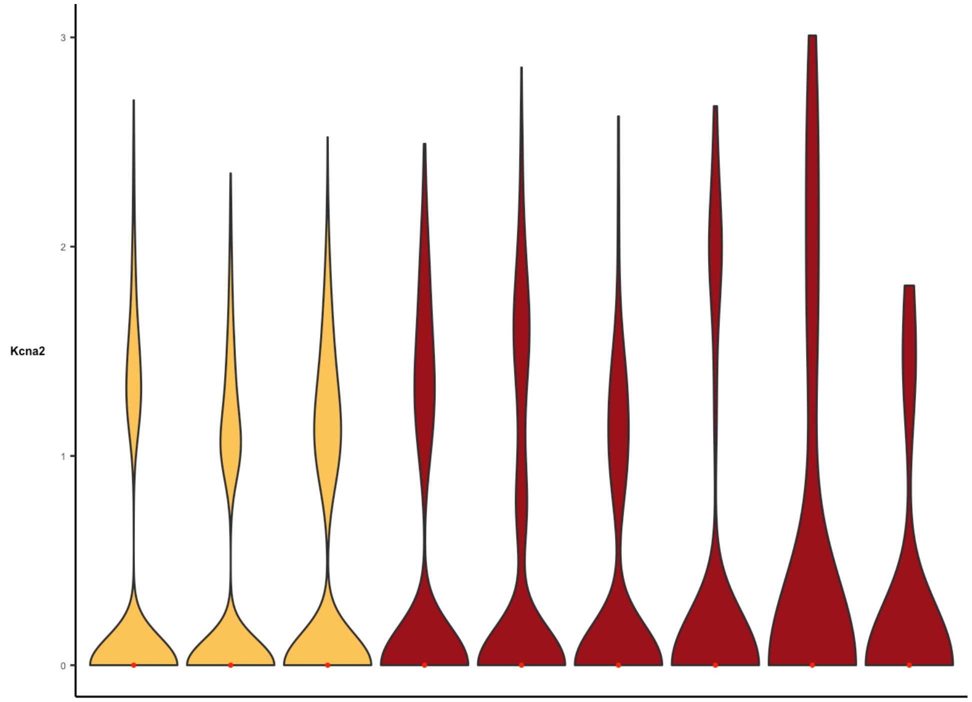


```
P3
```

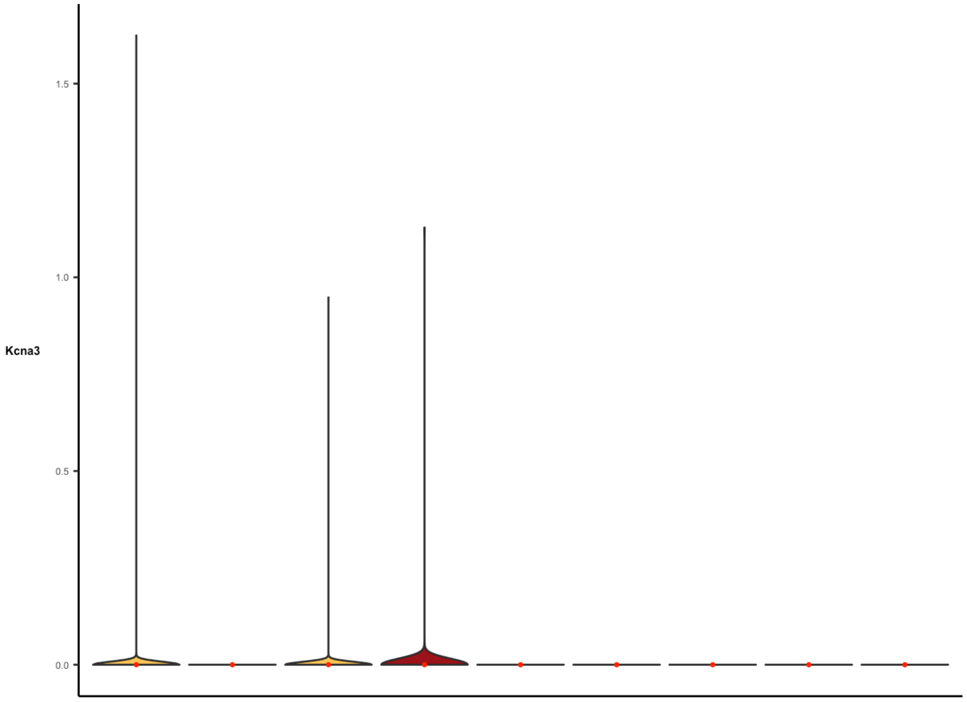


```
P4
```

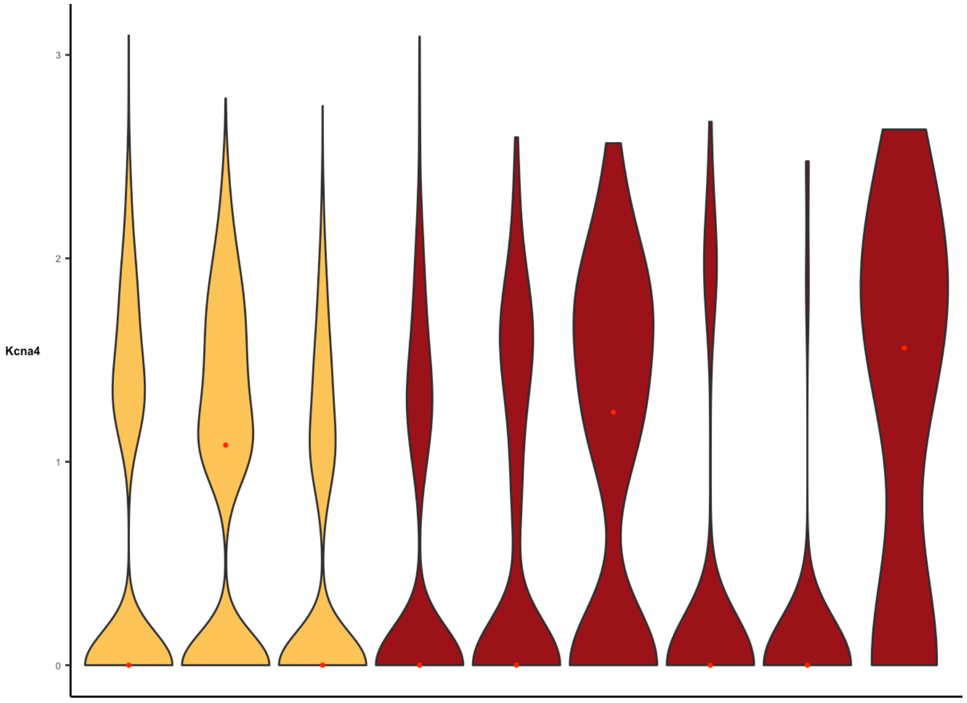


```
P5
```

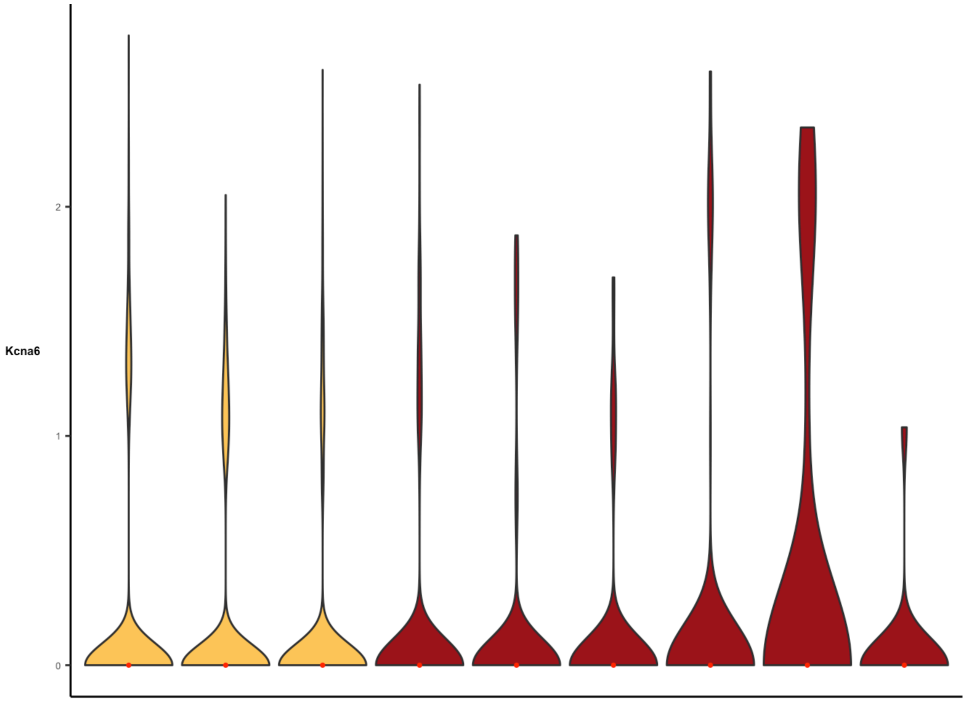


P6

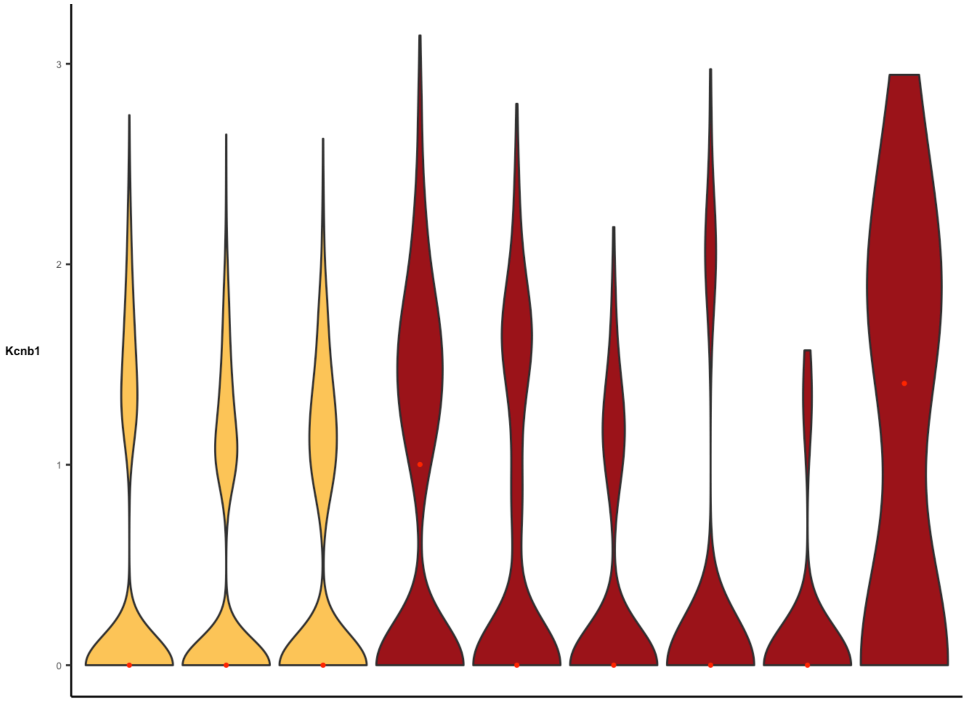


P7

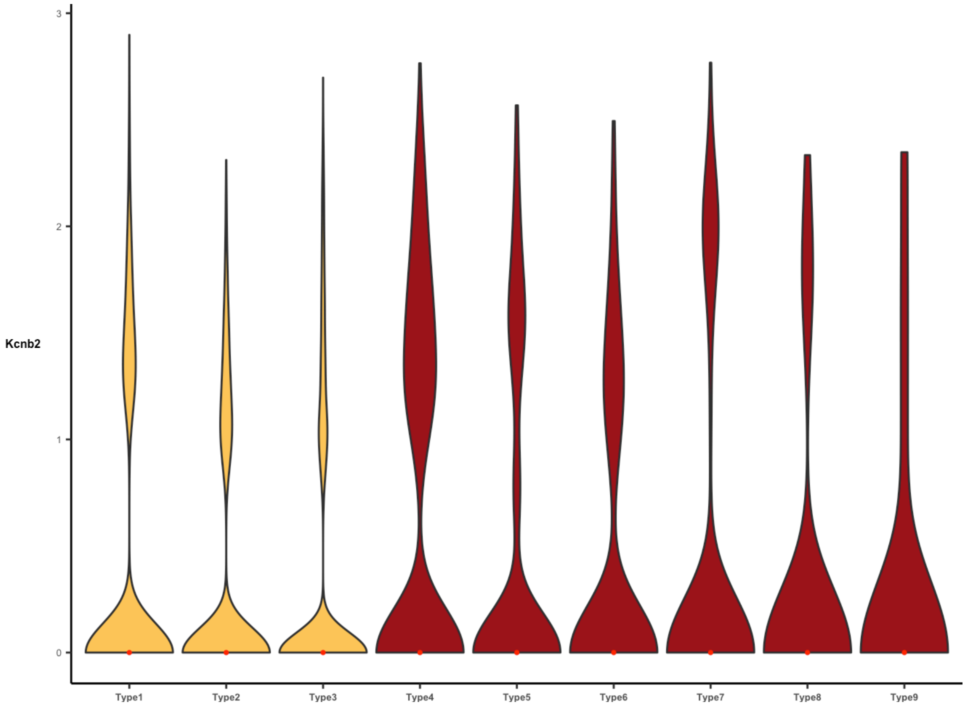


```
merge<-list()
for (i in length(gene_list):1){
       if (length(merge)==0){
        merge<-ggplotGrob(eval(parse(text=paste("P",i,sep = ""))))}else{merge<-rbind(ggplotGrob(eval(parse(text=p
aste("P",i,sep = "")))),merge,size = "last")}}

 #all genes
pdf(paste("/Users/lees44/R_project/FC_vs_dCA2_comparison_vector_integration_042821/cluster_tree_potassium_channel
_kcna_kcnb.pdf",".png",sep=""),height=6, width=50 , paper = "letter")


grid.newpage()
grid.draw(merge)
```

KCNC and KCND postassium channels

Violinplot for potassium channels, figure goes with cluster tree

```
Cell_type<-rev(c("Type1", "Type2", "Type3", "Type4", "Type5", "Type6", "Type7", "Type8", "Type9"))
gene_list<-c("Kcnc1","Kcnc2", "Kcnc3", "Kcnc4", "Kcnd1", "Kcnd2", "Kcnd3")
```

```
Cell_number <- NULL
for (i in 1:length(Cell_type)){
                temp1 <- data.frame(type=factor(rep(Cell_type[i],sum(Cell_type[i]==Idents(dCA2.integrated)))))
                temp2 <- as.data.frame(dCA2.integrated@assays$RNA@data[gene_list,Idents(dCA2.integrated)==Cell_ty
pe[i]])
                temp3 <- cbind(temp1,t(temp2))
                Cell_number <- rbind(temp3,Cell_number)
}
rownames(Cell_number) <- NULL
```

```
colors<-c("#ffc857","#ffc857","#ffc857","#9B0C1E","#9B0C1E","#9B0C1E","#9B0C1E","#9B0C1E","#9B0C1E")
```

```
for (k in 1:length(gene_list))
    {
if (k==length(gene_list)){
assign(paste("P",k,sep=""),ggplot(Cell_number,aes_string(x="type",y=gene_list[k],fill="type"))+geom_violin(scale
= "width")+scale_fill_manual(values=colors)
+stat_summary(fun=median, geom="point", size=0.6, color="red")+ylab(gene_list[k])+ theme(axis.title.x=element_bla
nk(),
axis.text.y=element_text(size=5),axis.title.y=element_text(size=6,angle=0,face="bold",margin = margin(t = 10, r =
8, b = 0, l = 10),vjust=0.5),axis.text.x=element_text(size=5,face="bold")
,axis.title=element_text(size=4,face="bold"),panel.grid.major = element_blank(), panel.grid.minor = element_blan
k(),
panel.background = element_blank(), axis.line = element_line(colour = "black"),legend.position="none",plot.margin
= unit(c(0, 0,0, 0), "cm")))}
else{
assign(paste("P",k,sep=""),ggplot(Cell_number,aes_string(x="type",y=gene_list[k],fill="type"))+geom_violin(scale
= "width")+scale_fill_manual(values=colors)
+stat_summary(fun=median, geom="point", size=0.6, color="red")+ylab(gene_list[k])+ theme(axis.title.x=element_bla
nk(),
axis.text.x=element_blank(),axis.text.y=element_text(size=5),axis.title.y=element_text(size=6,angle=0,face="bold",
margin = margin(t = 10, r = 8, b = 0, l = 10),vjust=0.5),axis.ticks.x=element_blank(),axis.text=element_text(size
=5)
,axis.title=element_text(size=4,face="bold"),panel.grid.major = element_blank(), panel.grid.minor = element_blan
k(),
panel.background = element_blank(), axis.line = element_line(colour = "black"),legend.position="none",plot.margin
= unit(c(0, 0,0, 0), "cm")))}

}
```

```
P1
```

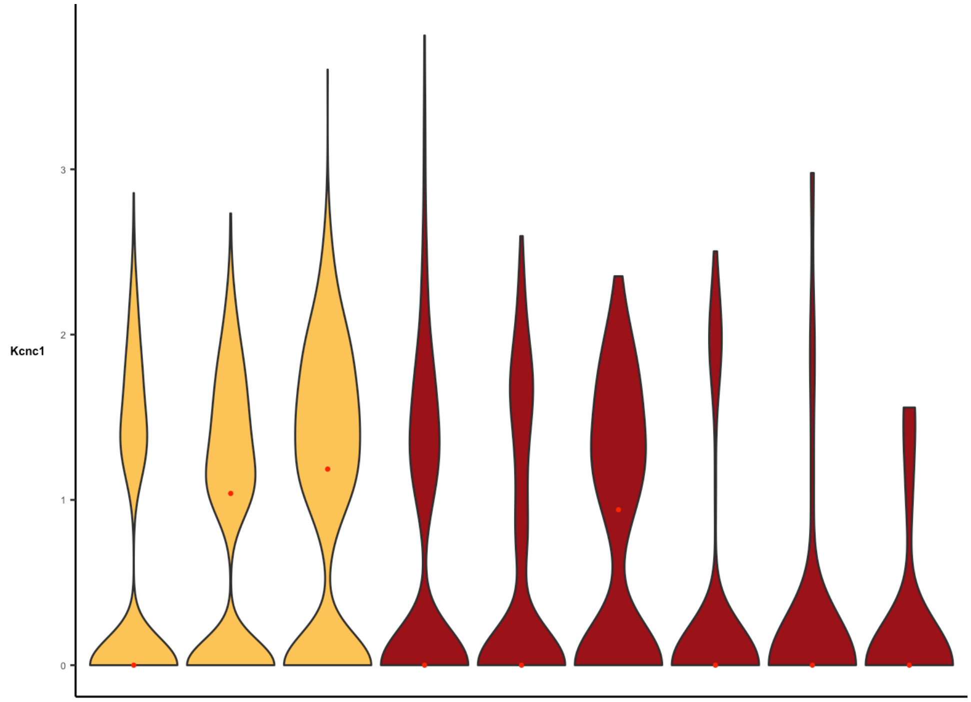


```
P2
```

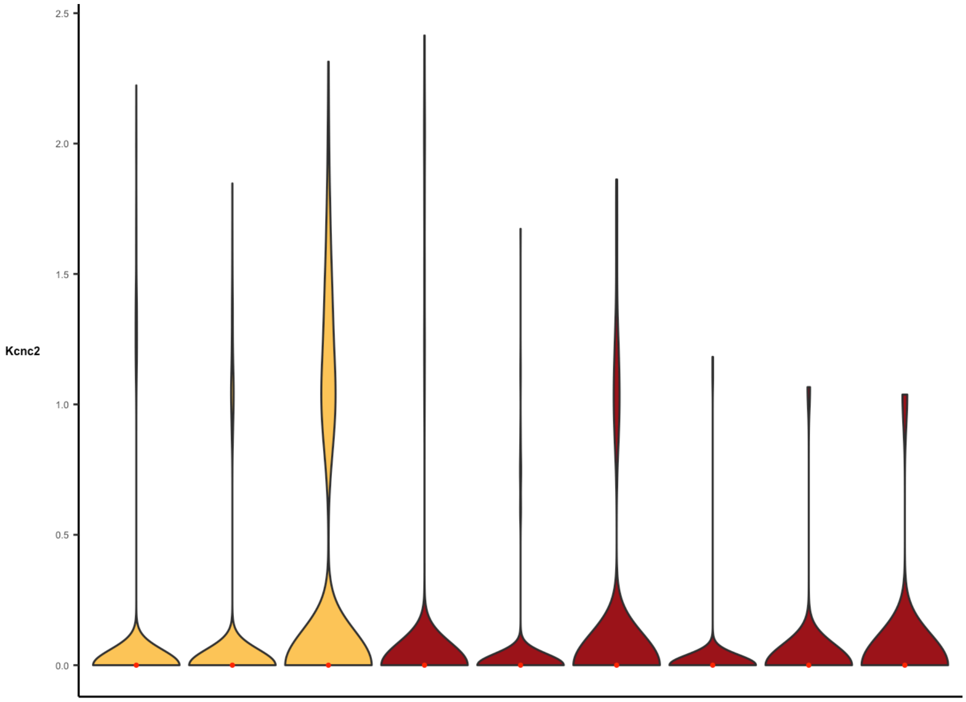


```
P3
```

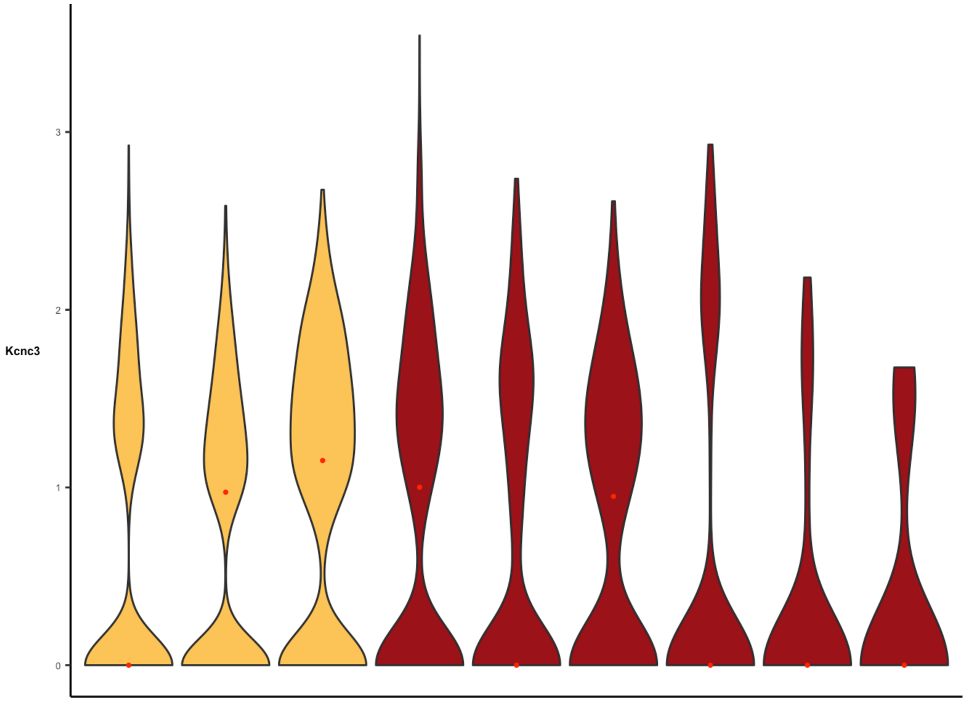


```
P4
```

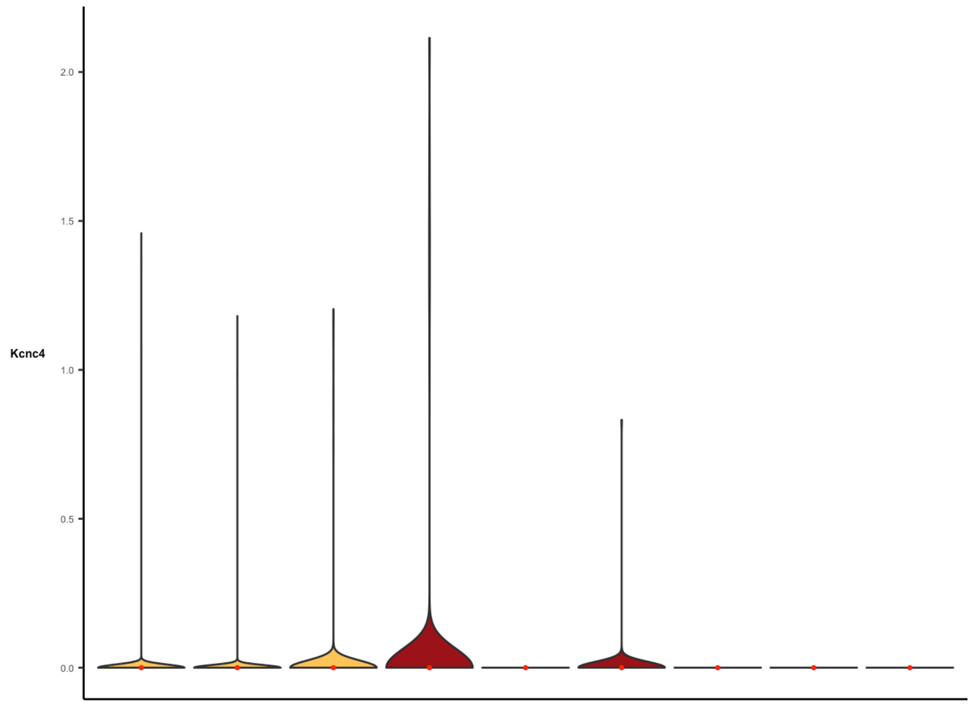


```
P5
```

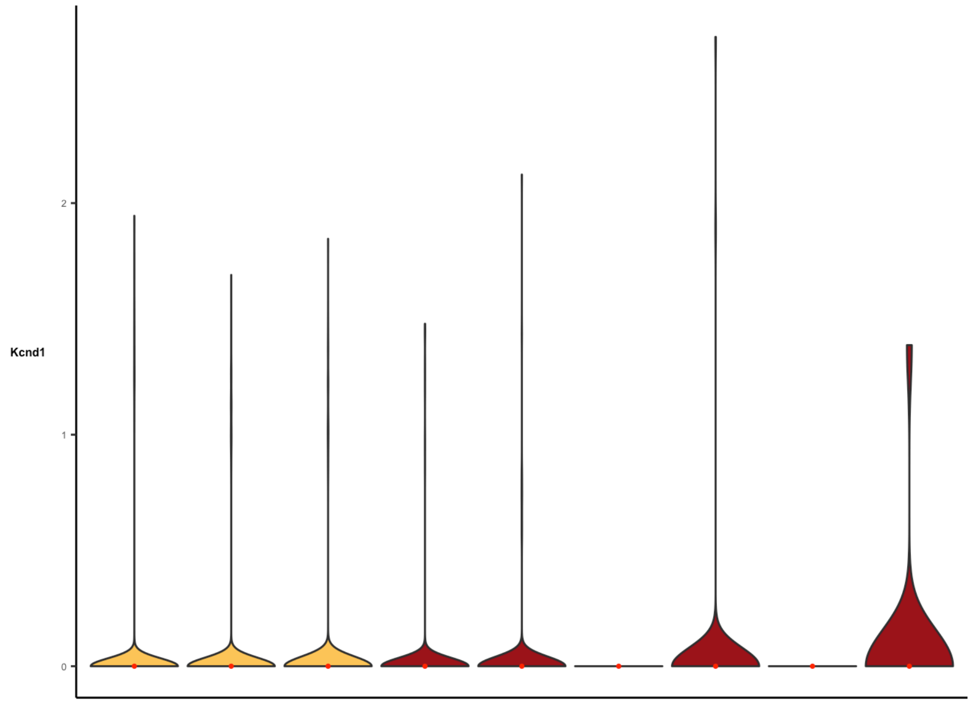


```
P6
```

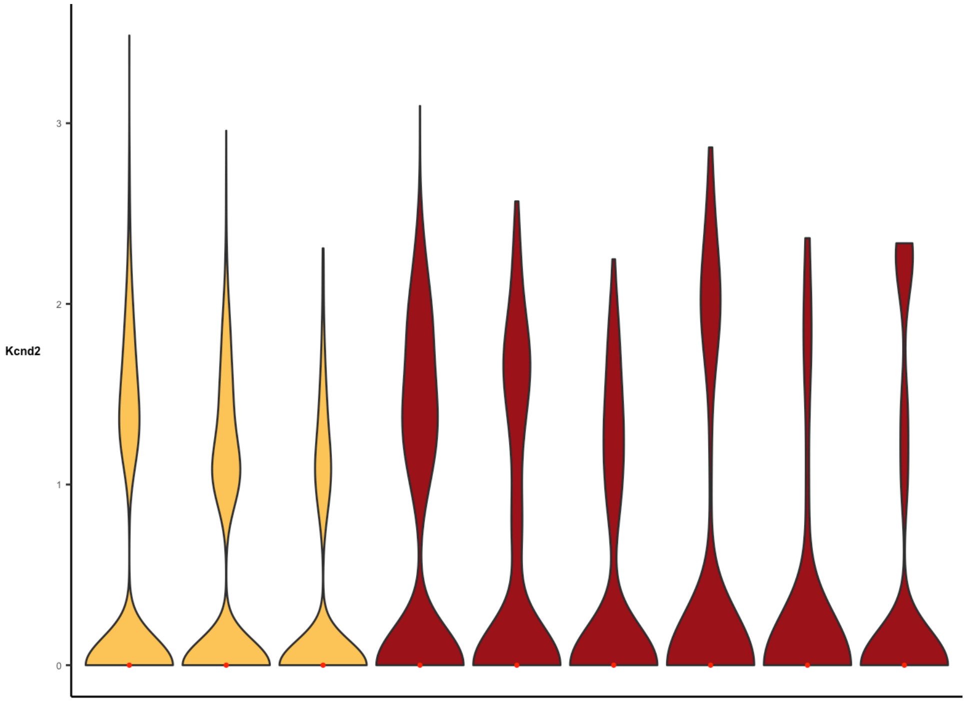
Kcnd2
0
1
2
3

```
P7
```

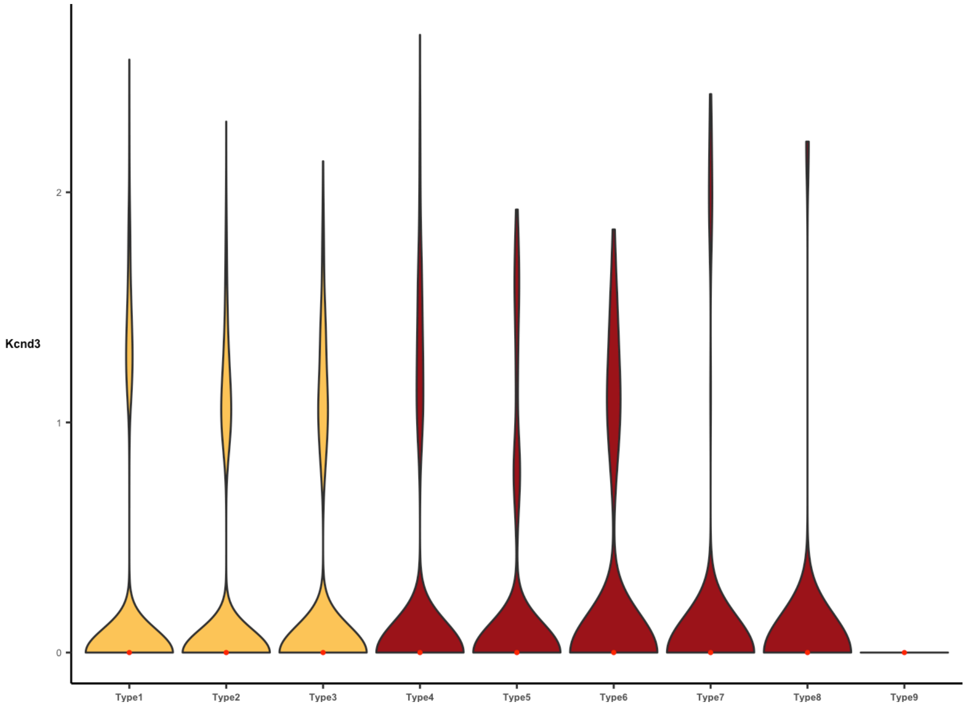
Kcnd3
0
1
2
Type1
Type2
Type3
Type4
Type5
Type6
Type7
Type8
Type9

```
merge<-list()
for (i in length(gene_list):1){
       if (length(merge)==0){
        merge<-ggplotGrob(eval(parse(text=paste("P",i,sep = ""))))}else{merge<-rbind(ggplotGrob(eval(parse(text=p
aste("P",i,sep = "")))),merge,size = "last")}}

 #all genes
pdf(paste("/Users/lees44/R_project/FC_vs_dCA2_comparison_vector_integration_042821/cluster_tree_potassium_channel
_kcnc_kcnd.pdf",".png",sep=""),height=6, width=50 , paper = "letter")


grid.newpage()
grid.draw(merge)
```

Kcnq Violinplot for potassium channels, figure goes with cluster tree

```
Cell_type<-rev(c("Type1", "Type2", "Type3", "Type4", "Type5", "Type6", "Type7", "Type8", "Type9"))
gene_list<-c("Kcnq2","Kcnq3", "Kcnq5")
```

```
Cell_number <- NULL
for (i in 1:length(Cell_type)){
                temp1 <- data.frame(type=factor(rep(Cell_type[i],sum(Cell_type[i]==Idents(dCA2.integrated)))))
                temp2 <- as.data.frame(dCA2.integrated@assays$RNA@data[gene_list,Idents(dCA2.integrated)==Cell_ty
pe[i]])
                temp3 <- cbind(temp1,t(temp2))
                Cell_number <- rbind(temp3,Cell_number)
}
rownames(Cell_number) <- NULL
```

```
colors<-c("#ffc857","#ffc857","#ffc857","#9B0C1E","#9B0C1E","#9B0C1E","#9B0C1E","#9B0C1E","#9B0C1E")
```

```
for (k in 1:length(gene_list))
    {
if (k==length(gene_list)){
assign(paste("P",k,sep=""),ggplot(Cell_number,aes_string(x="type",y=gene_list[k],fill="type"))+geom_violin(scale
= "width")+scale_fill_manual(values=colors)
+stat_summary(fun=median, geom="point", size=0.6, color="red")+ylab(gene_list[k])+ theme(axis.title.x=element_bla
nk(),
axis.text.y=element_text(size=5),axis.title.y=element_text(size=6,angle=0,face="bold",margin = margin(t = 10, r =
8, b = 0, l = 10),vjust=0.5),axis.text.x=element_text(size=5,face="bold")
,axis.title=element_text(size=4,face="bold"),panel.grid.major = element_blank(), panel.grid.minor = element_blan
k(),
panel.background = element_blank(), axis.line = element_line(colour = "black"),legend.position="none",plot.margin
= unit(c(0, 0,0, 0), "cm")))}
else{
assign(paste("P",k,sep=""),ggplot(Cell_number,aes_string(x="type",y=gene_list[k],fill="type"))+geom_violin(scale
= "width")+scale_fill_manual(values=colors)
+stat_summary(fun=median, geom="point", size=0.6, color="red")+ylab(gene_list[k])+ theme(axis.title.x=element_bla
nk(),
axis.text.x=element_blank(),axis.text.y=element_text(size=5),axis.title.y=element_text(size=6,angle=0,face="bold",
margin = margin(t = 10, r = 8, b = 0, l = 10),vjust=0.5),axis.ticks.x=element_blank(),axis.text=element_text(size
=5)
,axis.title=element_text(size=4,face="bold"),panel.grid.major = element_blank(), panel.grid.minor = element_blan
k(),
panel.background = element_blank(), axis.line = element_line(colour = "black"),legend.position="none",plot.margin
= unit(c(0, 0,0, 0), "cm")))}

}
```

```
P1
```

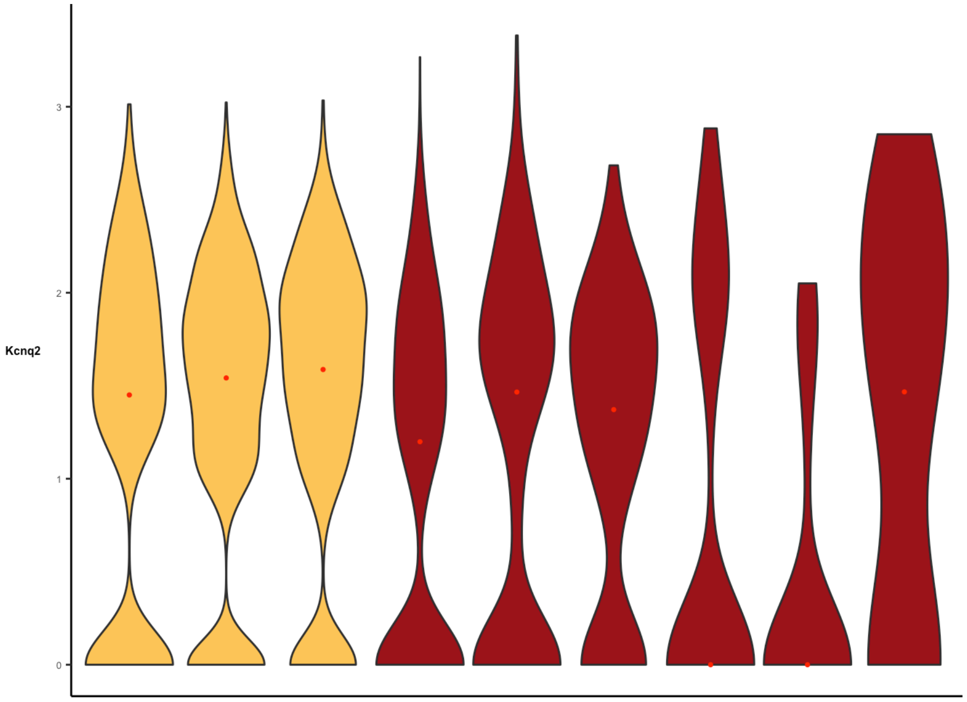


```
P2
```

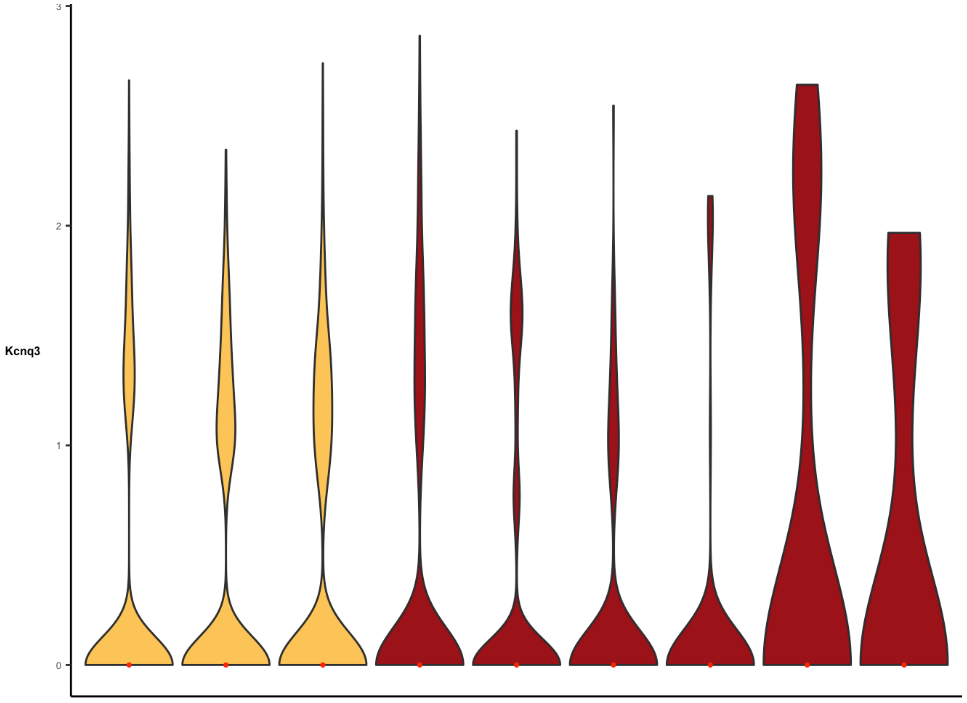


```
P3
```

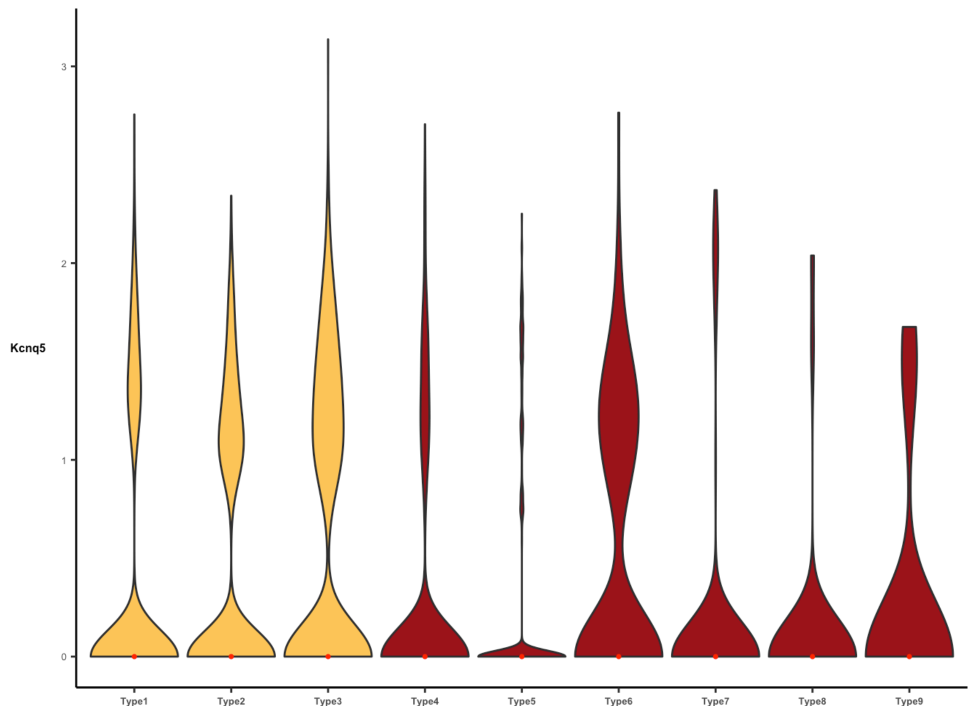


```
merge<-list()
for (i in length(gene_list):1){
       if (length(merge)==0){
        merge<-ggplotGrob(eval(parse(text=paste("P",i,sep = ""))))}else{merge<-rbind(ggplotGrob(eval(parse(text=p
aste("P",i,sep = "")))),merge,size = "last")}}

 #all genes
pdf(paste("/Users/lees44/R_project/FC_vs_dCA2_comparison_vector_integration_042821/cluster_tree_potassium_channel
_kcnq.pdf",".png",sep=""),height=6, width=50 , paper = "letter")


grid.newpage()
grid.draw(merge)
```

“CREREV38354884”, “GFP”,“Rgs14”, “Avpr1b”, “Amigo2”, “Map3k15”, “Actn2”, “Ptpn5”, “Cacng5”, “Oxtr”, “Pcp4”

Violinplot for dCA2 markers, figure goes with cluster tree

```
Cell_type<-rev(c("Type1", "Type2", "Type3", "Type4", "Type5", "Type6", "Type7", "Type8", "Type9"))
gene_list<-c("C1ql3" ,"Adarb2","Ptchd4", "Mef2c","Cit")
```

```
Cell_number <- NULL
for (i in 1:length(Cell_type)){
                temp1 <- data.frame(type=factor(rep(Cell_type[i],sum(Cell_type[i]==Idents(dCA2.integrated)))))
                temp2 <- as.data.frame(dCA2.integrated@assays$RNA@data[gene_list,Idents(dCA2.integrated)==Cell_ty
pe[i]])
                temp3 <- cbind(temp1,t(temp2))
                Cell_number <- rbind(temp3,Cell_number)
}
rownames(Cell_number) <- NULL
```

```
colors<-c("#ffc857","#ffc857","#ffc857","#9B0C1E","#9B0C1E","#9B0C1E","#9B0C1E","#9B0C1E","#9B0C1E")
```

```
for (k in 1:length(gene_list))
    {
if (k==length(gene_list)){
assign(paste("P",k,sep=""),ggplot(Cell_number,aes_string(x="type",y=gene_list[k],fill="type"))+geom_violin(scale
= "width")+scale_fill_manual(values=colors)
+stat_summary(fun=median, geom="point", size=0.6, color="red")+ylab(gene_list[k])+ theme(axis.title.x=element_bla
nk(),
axis.text.y=element_text(size=5),axis.title.y=element_text(size=6,angle=0,face="bold",margin = margin(t = 10, r =
8, b = 0, l = 10),vjust=0.5),axis.text.x=element_text(size=5,face="bold")
,axis.title=element_text(size=4,face="bold"),panel.grid.major = element_blank(), panel.grid.minor = element_blan
k(),
panel.background = element_blank(), axis.line = element_line(colour = "black"),legend.position="none",plot.margin
= unit(c(0, 0,0, 0), "cm")))}
else{
assign(paste("P",k,sep=""),ggplot(Cell_number,aes_string(x="type",y=gene_list[k],fill="type"))+geom_violin(scale
= "width")+scale_fill_manual(values=colors)
+stat_summary(fun=median, geom="point", size=0.6, color="red")+ylab(gene_list[k])+ theme(axis.title.x=element_bla
nk(),
axis.text.x=element_blank(),axis.text.y=element_text(size=5),axis.title.y=element_text(size=6,angle=0,face="bold",
margin = margin(t = 10, r = 8, b = 0, l = 10),vjust=0.5),axis.ticks.x=element_blank(),axis.text=element_text(size
=5)
,axis.title=element_text(size=4,face="bold"),panel.grid.major = element_blank(), panel.grid.minor = element_blan
k(),
panel.background = element_blank(), axis.line = element_line(colour = "black"),legend.position="none",plot.margin
= unit(c(0, 0,0, 0), "cm")))}

}
```

```
P1
```

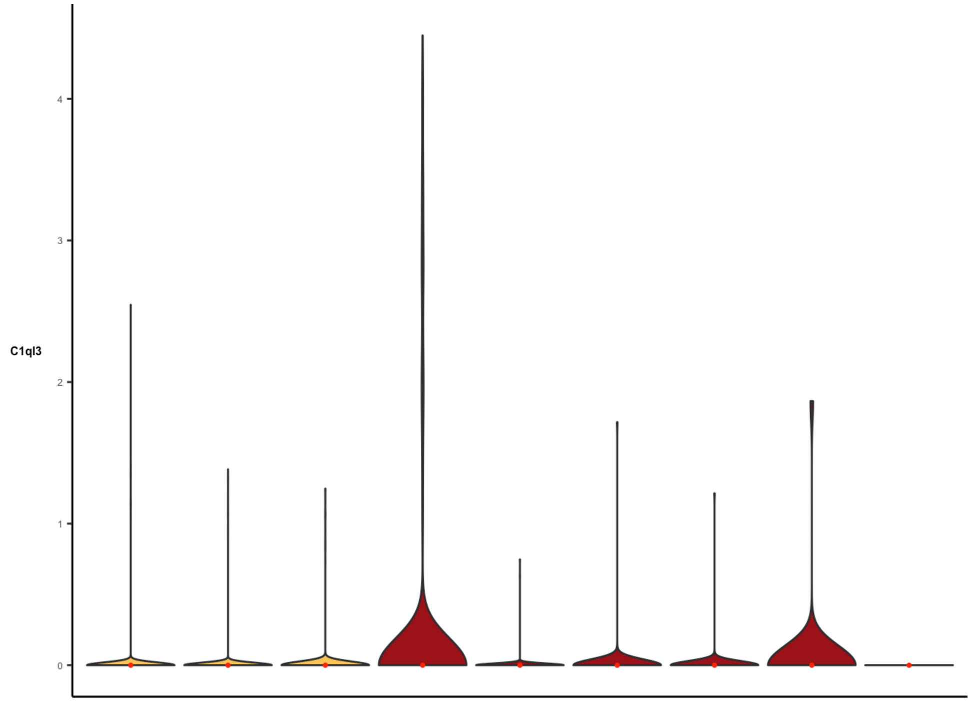


```
P2
```

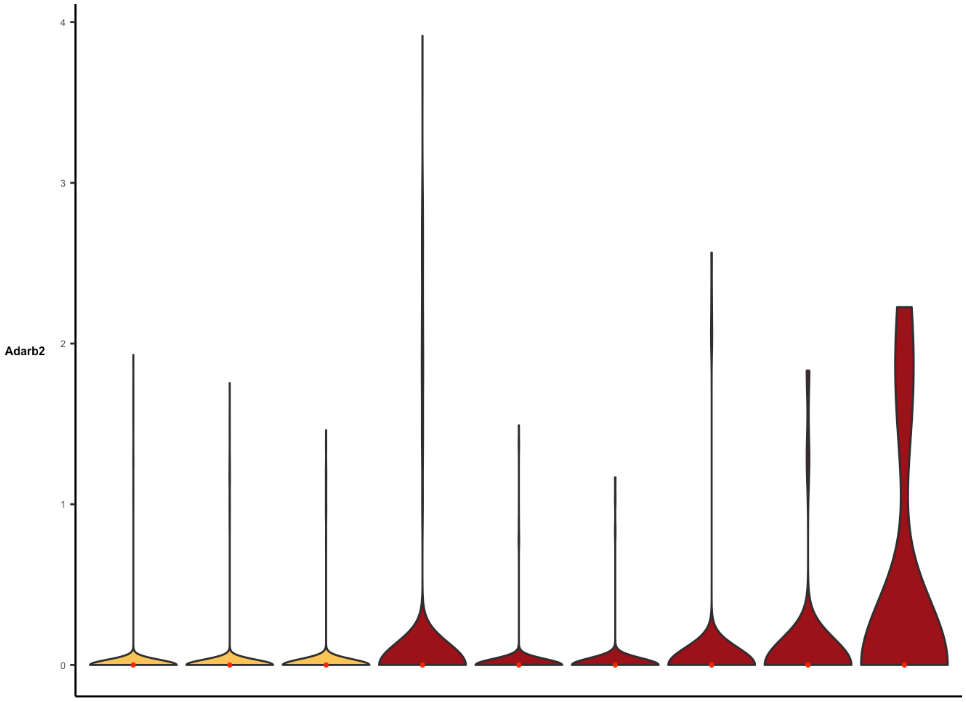


```
P3
```

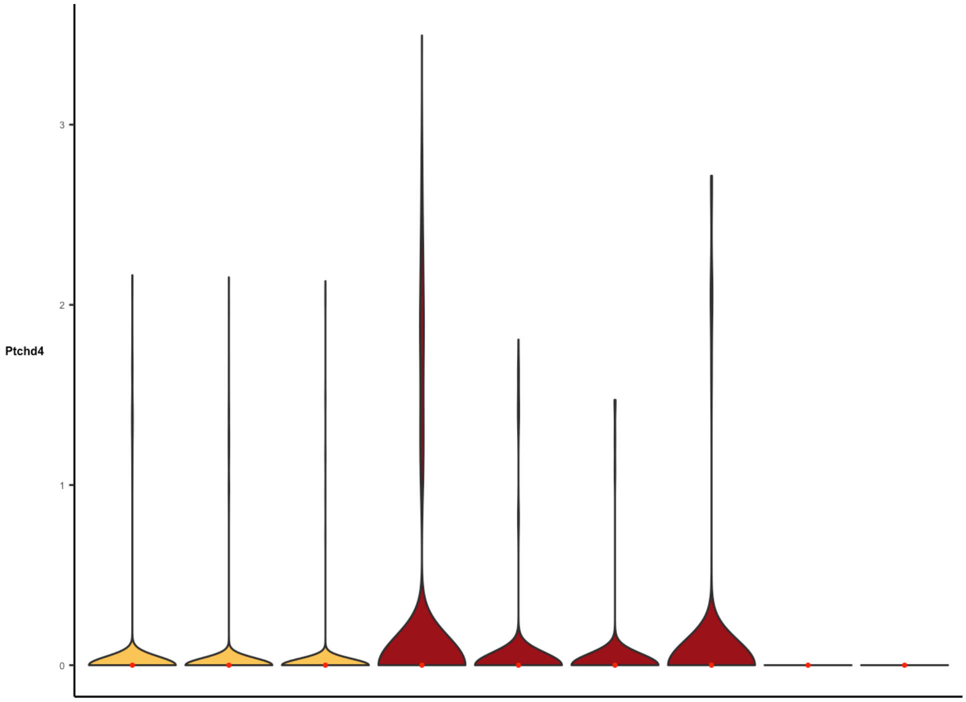


```
P4
```

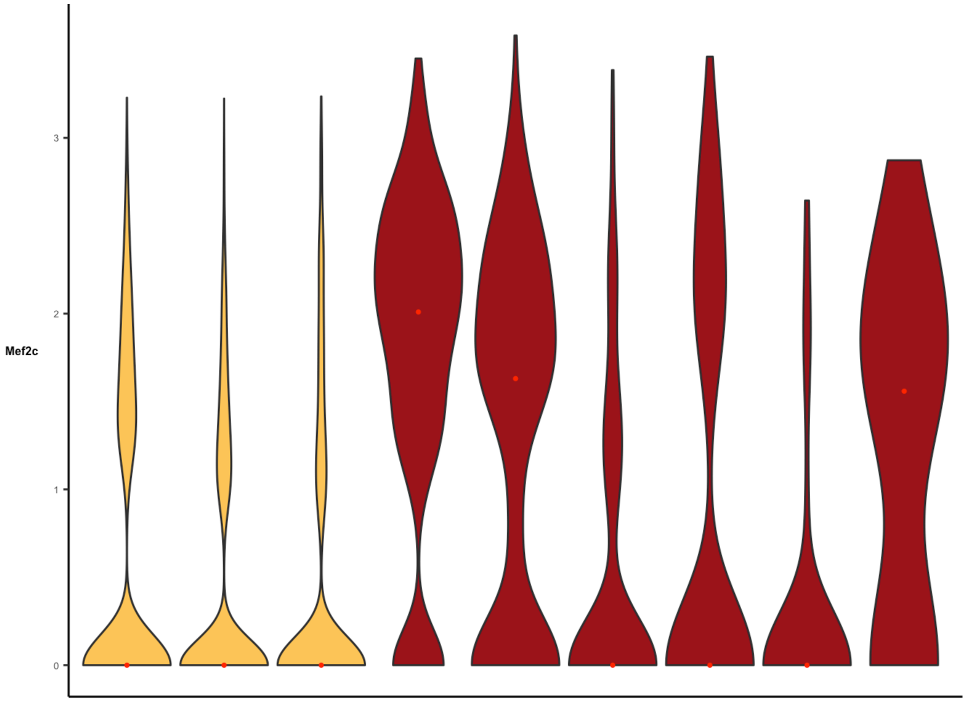
Mef2c
0
1
2
3

```
P5
```

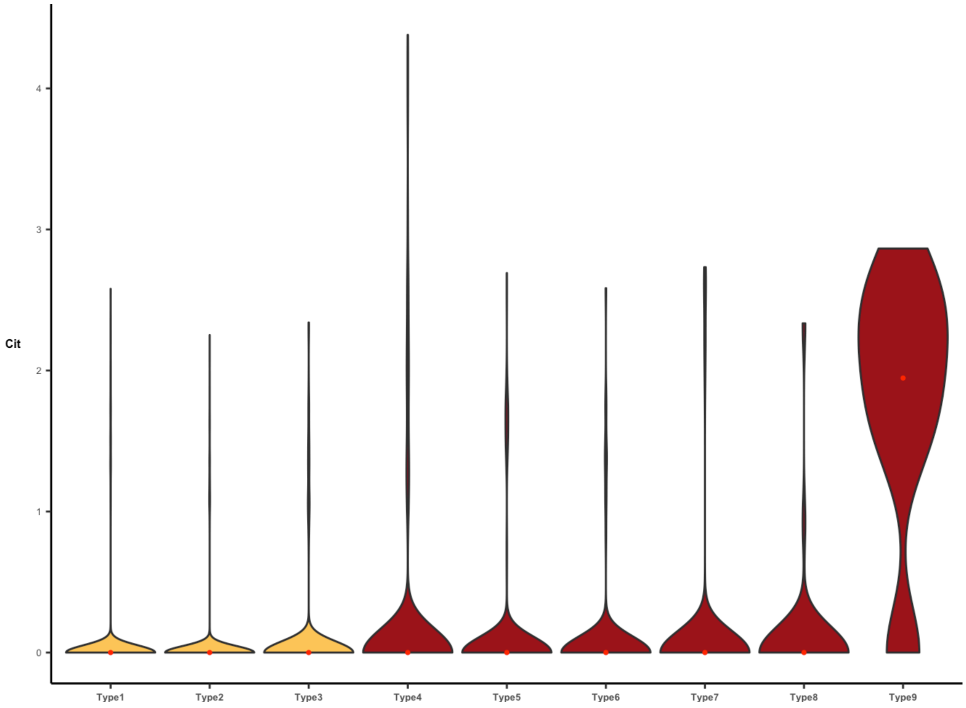
Cit
0
1
2
3
4
Type1
Type2
Type3
Type4
Type5
Type6
Type7
Type8
Type9

```
merge<-list()
for (i in length(gene_list):1){
       if (length(merge)==0){
        merge<-ggplotGrob(eval(parse(text=paste("P",i,sep = ""))))}else{merge<-rbind(ggplotGrob(eval(parse(text=p
aste("P",i,sep = "")))),merge,size = "last")}}

 #all genes
pdf(paste("/Users/lees44/R_project/FC_vs_dCA2_comparison_vector_integration_042821/cluster_tree_MEF4c.pdf",".png",
sep=""),height=6, width=50 , paper = "letter")


grid.newpage()
grid.draw(merge)
```

UMI plot for each cluster Identifying conserved markers for each clusters

```
install.packages("xlsx")
```

```
## Installing package into '/Users/lees44/Library/R/4.0/library'
## (as 'lib' is unspecified)
```

```
## Error in contrib.url(repos, "source"): trying to use CRAN without setting a mirror
```

```
install.packages("rJava",type='source')
```

```
## Installing package into '/Users/lees44/Library/R/4.0/library'
## (as 'lib' is unspecified)
```

```
## Error in contrib.url(repos, type): trying to use CRAN without setting a mirror
```

```
library("rJava")
```

```
library("xlsx")
```

```
install.packages("metap")
```

```
## Installing package into '/Users/lees44/Library/R/4.0/library'
## (as 'lib' is unspecified)
```

```
## Error in contrib.url(repos, "source"): trying to use CRAN without setting a mirror
```

```
BiocManager::install('multtest')
```

```
## Bioconductor version 3.12 (BiocManager 1.30.14), R 4.0.4 (2021-02-15)
```

```
## Warning: package(s) not installed when version(s) same as current; use `force = TRUE` to
##   re-install: 'multtest'
```

```
## Installation paths not writeable, unable to update packages
##   path: /Library/Frameworks/R.framework/Versions/4.0/Resources/library
##   packages:
##     boot, class, cluster, KernSmooth, lattice, MASS, mgcv, nlme, nnet, spatial,
##     survival
```

```
## Old packages: 'backports', 'beeswarm', 'BiocManager', 'blob', 'broom', 'bslib',
##   'cachem', 'cli', 'colorspace', 'conquer', 'cpp11', 'crayon', 'crosstalk',
##   'curl', 'data.table', 'deldir', 'desc', 'diffobj', 'digest', 'dplyr',
##   'fansi', 'fitdistrplus', 'formatR', 'future', 'future.apply', 'gargle',
##   'generics', 'ggplot2', 'glue', 'goftest', 'googledrive', 'googlesheets4',
##   'gtools', 'haven', 'hms', 'htmltools', 'htmlwidgets', 'httpuv', 'igraph',
##   'isoband', 'knitr', 'later', 'leiden', 'lifecycle', 'lmtest', 'lubridate',
##   'Matrix', 'matrixStats', 'metap', 'mime', 'mvtnorm', 'openssl', 'parallelly',
##   'pbapply', 'pillar', 'pkgload', 'plotly', 'plotrix', 'quantreg', 'R6',
##   'rbibutils', 'Rcpp', 'RcppAnnoy', 'RcppArmadillo', 'RCurl', 'Rdpack',
##   'readr', 'reprex', 'reticulate', 'rJava', 'rlang', 'rmarkdown', 'roxygen2',
##   'rvest', 'sandwich', 'sass', 'Seurat', 'SeuratObject', 'shiny', 'sitmo',
##   'snow', 'spatstat.core', 'spatstat.geom', 'spatstat.utils', 'statmod',
##   'stringi', 'testthat', 'TH.data', 'tibble', 'tidyr', 'tinytex', 'utf8',
##   'uuid', 'viridis', 'waldo', 'xfun', 'xgboost', 'zip'
```

```
library("metap")
```

```
#divide by avarage read number
length(Type1_barcode)/10000
```

```
## Error in eval(expr, envir, enclos): object 'Type1_barcode' not found
```

```
length(Type2_barcode)/10000
```

```
## Error in eval(expr, envir, enclos): object 'Type2_barcode' not found
```

```
length(Type3_barcode)/10000
```

```
## Error in eval(expr, envir, enclos): object 'Type3_barcode' not found
```

```
length(Type4_barcode)/10000
```

```
## Error in eval(expr, envir, enclos): object 'Type4_barcode' not found
```

```
length(Type5_barcode)/10000
```

```
## Error in eval(expr, envir, enclos): object 'Type5_barcode' not found
```

```
length(Type6_barcode)/10000
```

```
## Error in eval(expr, envir, enclos): object 'Type6_barcode' not found
```

```
length(Type7_barcode)/10000
```

```
## Error in eval(expr, envir, enclos): object 'Type7_barcode' not found
```

```
length(Type8_barcode)/10000
```

```
## Error in eval(expr, envir, enclos): object 'Type8_barcode' not found
```

```
length(Type9_barcode)/10000
```

```
## Error in eval(expr, envir, enclos): object 'Type9_barcode' not found
```

```
type<-numeric()
for (i in 1:dim(dCA2.integrated@meta.data)[1]){
if(rownames(dCA2.integrated@meta.data)[i] %in% Type1_barcode){type[i]<-"Type1"}
else if(rownames(dCA2.integrated@meta.data)[i] %in% Type2_barcode){type[i]<-"Type2"}
else if(rownames(dCA2.integrated@meta.data)[i] %in% Type3_barcode){type[i]<-"Type3"}
else if(rownames(dCA2.integrated@meta.data)[i] %in% Type4_barcode){type[i]<-"Type4"}
else if(rownames(dCA2.integrated@meta.data)[i] %in% Type5_barcode){type[i]<-"Type5"}
else if(rownames(dCA2.integrated@meta.data)[i] %in% Type6_barcode){type[i]<-"Type6"}
else if(rownames(dCA2.integrated@meta.data)[i] %in% Type7_barcode){type[i]<-"Type7"}
else if(rownames(dCA2.integrated@meta.data)[i] %in% Type8_barcode){type[i]<-"Type8"}
else if(rownames(dCA2.integrated@meta.data)[i] %in% Type9_barcode){type[i]<-"Type9"}
else{type[i]<-"NA"}
}
```

```
## Error in h(simpleError(msg, call)): error in evaluating the argument 'table' in selecting a method for functio
n '%in%': object 'Type1_barcode' not found
```

```
dCA2.integrated@meta.data$type<-type
```

```
## Error in `$<-.data.frame`(`*tmp*`, type, value = numeric(0)): replacement has 0 rows, data has 5928
```

```
colors<-c("Type1"="#19647e","Type2"="#ffc857","Type3"="#9B0C1E","Type4"="#4b3f72","Type5"="#ffc857","Type6"="#676
833", "Type7"="#19647e","Type8"="#ffc857", "Type9"="#9B0C1E")
```

```
subset_meta<-subset(dCA2.integrated@meta.data,type=="Type1"|type=="Type2"|type=="Type3"|type=="Type4"|type=="Type
5"|type=="Type6"|type=="Type7"|type=="Type8"|type=="Type9")
ggplot(subset_meta,aes_string(x="type",y="nCount_RNA",fill="type"))+geom_violin(scale = "width")+scale_fill_manua
l(values=colors)+
stat_summary(fun=median, geom="point", size=0.6, color="red")+ylab("UMI")+
theme(axis.title.x=element_blank(),axis.text.y=element_text(size=10),axis.title.y=element_text(size=10,angle=0,fa
ce="bold",margin = margin(t = 10, r = 8, b = 0, l = 10),vjust=0.5),axis.text.x=element_text(size=10,face="bold"),
axis.title=element_text(size=15,face="bold"),panel.grid.major = element_blank(), panel.grid.minor = element_blan
k(),panel.background = element_blank(), axis.line = element_line(colour = "black"),legend.position="none",plot.ma
rgin = unit(c(0,
0,0, 0), "cm"))
```

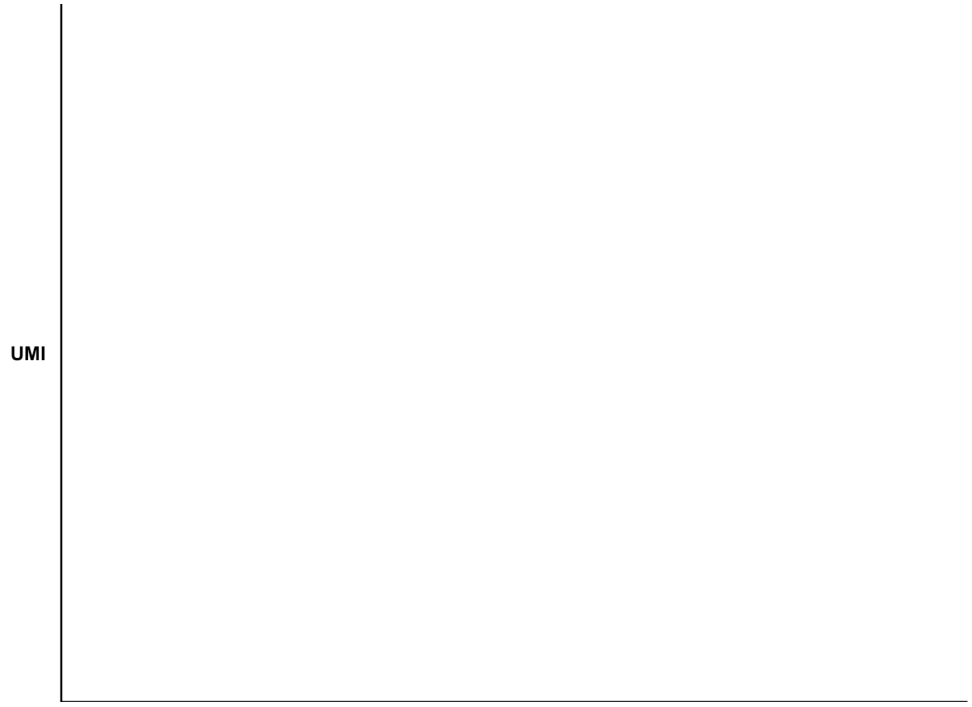

```
ggsave(file="/Users/lees44/R_project/FC_vs_dCA2_comparison_vector_integration_042821/cell_type_UMI_count.pdf",width=10,height=7)
```

#######################save image .PDF#######################

########################subset data clean with 7 cell types only 08.03.2021###################################### We have only included cell types that represent more than 0.1%. Cell type 8 and 9 therefore did not qualify. Here we are analyzing subset data of 7 different cell types. We are using subset function to subset the clean data.

```
for (i in 1:length(new.ident)){assign(paste(new.ident[i],"_barcode",sep=""),colnames(dCA2.integrated@assays$RNA@data[,which(Idents(object=dCA2.integrated) %in% new.ident[i])]))}
```

```
dCA2.integrated_clean_7celltype<-subset(x=dCA2.integrated,cells=c(Type1_barcode, Type2_barcode, Type3_barcode, Type4_barcode, Type5_barcode, Type6_barcode, Type7_barcode))
```

```
saveRDS(dCA2.integrated_clean_7celltype, file = "/Users/lees44/R_project/FC_vs_dCA2_comparison_vector_integration_042821/dCA2_integrated_clean_7celltype.rds")
```

```
DimPlot(object = dCA2.integrated_clean_7celltype, reduction = "umap", label = FALSE, repel = TRUE, pt=1)
```

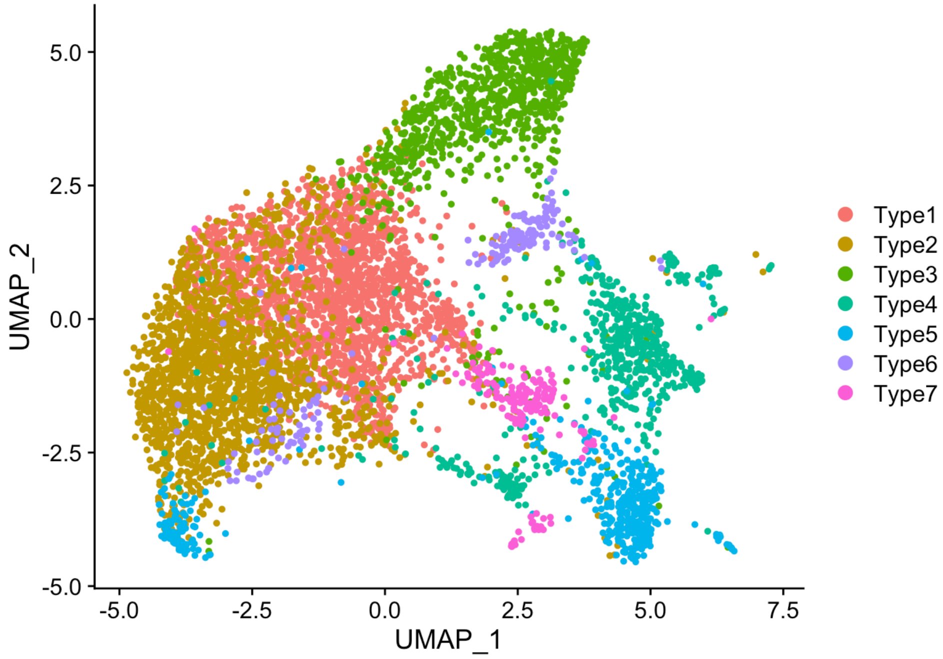


```
ggsave(file="/Users/lees44/R_project/FC_vs_dCA2_comparison_vector_integration_042821/FC_dCA2_Neuronaltype_clean_7celltype.pdf",width=10,height=10)
```

split by identity see https://satijalab.org/seurat/reference/splitobject (https://satijalab.org/seurat/reference/splitobject)

Changed width and height to make sure it fits. Check other split by plots

```
DimPlot(object = dCA2.integrated_clean_7celltype, reduction = "umap", split.by = "ident", label = FALSE, repel = TRUE, pt=1)
```

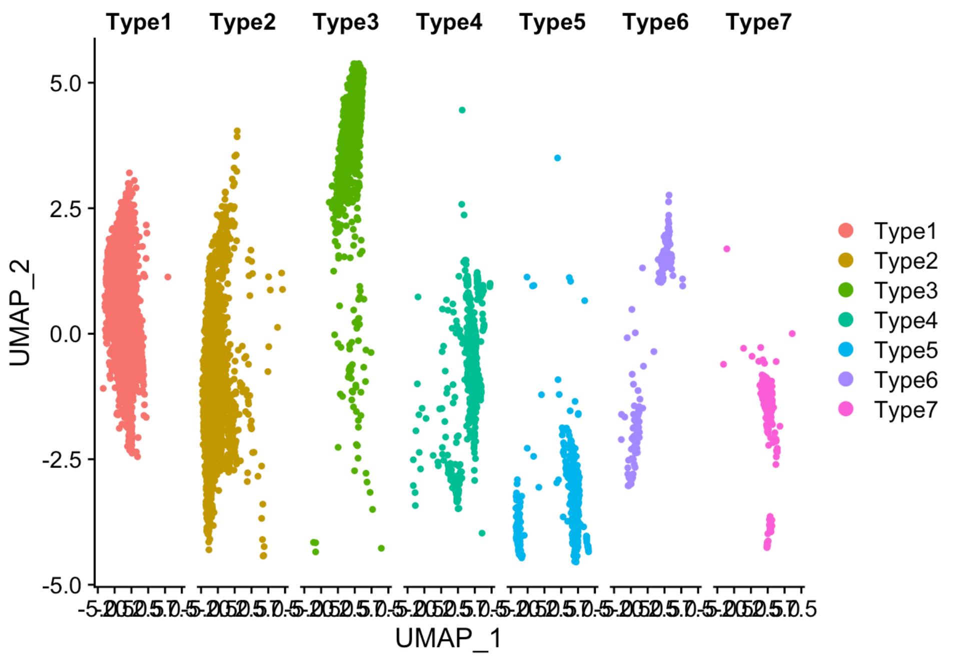


```
ggsave(file="/Users/lees44/R_project/FC_vs_dCA2_comparison_vector_integration_042821/FC_dCA2_Neuronaltype_clean_7celltype_splitby.pdf", width = 25, height = 5)
```

```
p2<-DimPlot(dCA2.integrated_clean_7celltype, reduction = "umap", group.by = "protocol", pt=1)
plot_grid(p2)
```

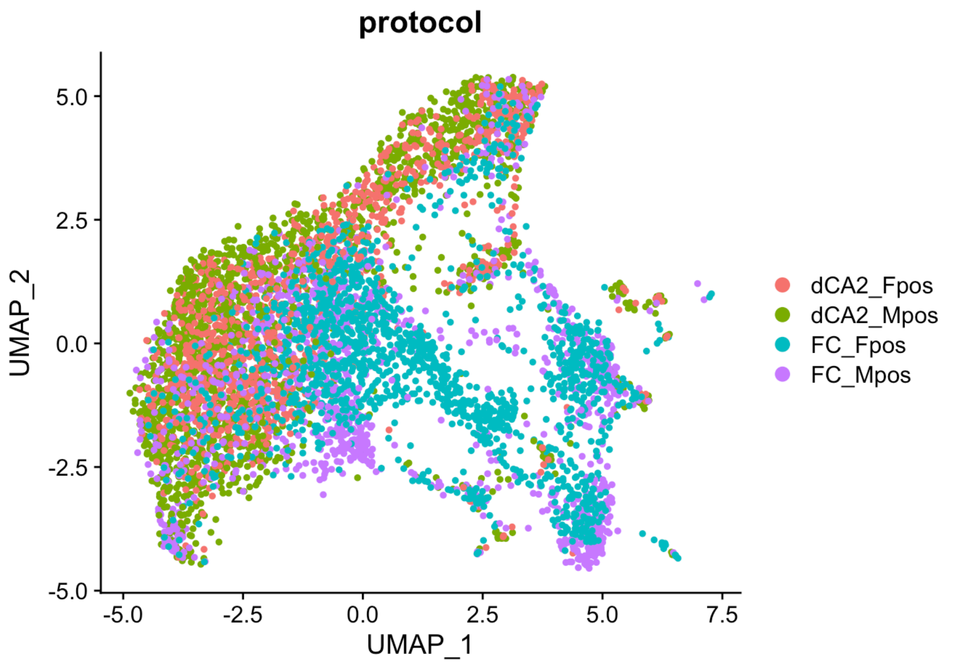


```
ggsave(file="/Users/lees44/R_project/FC_vs_dCA2_comparison_vector_integration_042821/FC_dCA2_UMAP_clean_7celltypes.pdf",width=10,height=10)
```

```
p2<-DimPlot(dCA2.integrated_clean_7celltype, reduction = "umap", split.by = "protocol", pt=1)
plot_grid(p2)
```

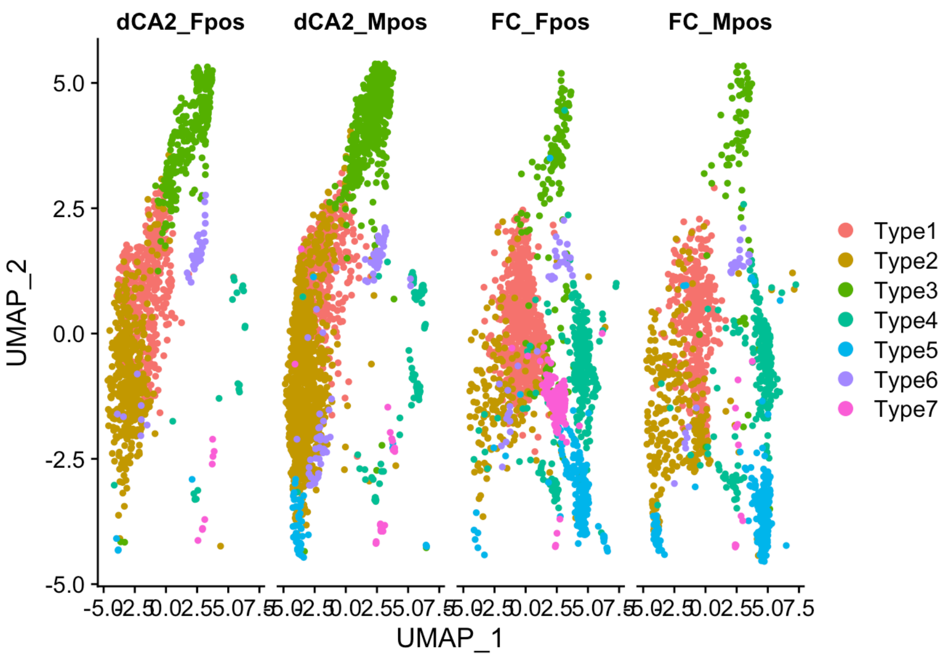


```
ggsave(file="/Users/lees44/R_project/FC_vs_dCA2_comparison_vector_integration_042821/FC_dCA2_UMAP_clean_7celltype
s_splot.pdf",width=10,height=10)
```

3D plot using plotly for vizualization of 3D UMAP for fun! https://github.com/Dragonmasterx87/Interactive-3D-Plotting-in-Seurat-3.0.0/blob/master/3D%20UMAP%20Plotting%20v1.3.R (https://github.com/Dragonmasterx87/Interactive-3D-Plotting-in-Seurat-3.0.0/blob/master/3D%20UMAP%20Plotting%20v1.3.R)

```
# Construct a dataframe using data from your pre-clustered Seurat v3.1.1 object
# Here 'seurat_clusters' is list of numeric cluster identities, you can find it here: yourseuratobject[["seurat_c
luster"]],
# or yourseuratobject$seurat_clusters, where 'yourseuratobject' is a Seurat object created with Seurat v3.1.1 (wo
rks for v3.0.0 as well)
yourseuratobject <- dCA2.integrated_clean_7celltype

# Re-run UMAPs that you have accurate calculations for all UMAP(s)
yourseuratobject <- RunUMAP(dCA2.integrated_clean_7celltype,
                            dims = 1:10,
                            n.components = 3L)
```

```
## 13:54:12 UMAP embedding parameters a = 0.9922 b = 1.112
```

```
## 13:54:12 Read 5874 rows and found 10 numeric columns
```

```
## 13:54:12 Using Annoy for neighbor search, n_neighbors = 30
```

```
## 13:54:12 Building Annoy index with metric = cosine, n_trees = 50
```

```
## 0%   10   20   30   40   50   60   70   80   90   100%
```

```
## [----|----|----|----|----|----|----|----|----|----|
```

```
## **************************************************|
## 13:54:13 Writing NN index file to temp file /var/folders/9w/_lzbttfd7nv1_y4jd_50rlmj7j_gxg/T//RtmpJyR1ZK/file1
3cbb60cdc221
## 13:54:13 Searching Annoy index using 1 thread, search_k = 3000
## 13:54:15 Annoy recall = 100%
## 13:54:16 Commencing smooth kNN distance calibration using 1 thread
## 13:54:17 Initializing from normalized Laplacian + noise
## 13:54:17 Commencing optimization for 500 epochs, with 247942 positive edges
## 13:54:26 Optimization finished
```

```
# Extract tSNE information from Seurat Object
umap_1 <- yourseuratobject[["umap"]]@cell.embeddings[,1]
umap_2 <- yourseuratobject[["umap"]]@cell.embeddings[,2]
umap_3 <- yourseuratobject[["umap"]]@cell.embeddings[,3]

# Visualize what headings are called so that you can extract them to form a dataframe
Embeddings(object = yourseuratobject, reduction = "umap")
```

```
##                                                      UMAP_1        UMAP_2
## dCA2_Mpos_AAAGGTATCTAGAACC-1dCA2_MI_GFPve_pos -0.4275558717  0.7304168068
## dCA2_Mpos_AACAGGGAGACCTCAT-1dCA2_MI_GFPve_pos  2.1587267093  3.0438065849
## dCA2_Mpos_AACCATGGTACCAGAG-1dCA2_MI_GFPve_pos -0.6158885486  1.6330867134
## dCA2_Mpos_AACCCAAAGCGTATAA-1dCA2_MI_GFPve_pos  1.9863319568  3.1878486000
## dCA2_Mpos_AACGTCATCACGGACC-1dCA2_MI_GFPve_pos  1.0036353044  2.2135806404
## dCA2_Mpos_AAGACAATCTGACGCG-1dCA2_MI_GFPve_pos  0.7382008605  2.4206981979
## dCA2_Mpos_AAGGAATCATATCGGT-1dCA2_MI_GFPve_pos -0.9841944404  1.7300467811
## dCA2_Mpos_AAGTTCGAGGATATAC-1dCA2_MI_GFPve_pos -0.8072851725  2.2656932197
## dCA2_Mpos_AATCACGCAGAGTAAT-1dCA2_MI_GFPve_pos  1.7082556658  3.3141889892
## dCA2_Mpos_AATGAAGAGAGTGAAG-1dCA2_MI_GFPve_pos -1.8012481756  1.0820875488
## dCA2_Mpos_ACACTGAGTAATGATG-1dCA2_MI_GFPve_pos -1.5951852627  1.0675683342
## dCA2_Mpos_ACAGCCGGTTATAGAG-1dCA2_MI_GFPve_pos  0.8555886917  2.1946978889
## dCA2_Mpos_ACATGCATCGCTTGCT-1dCA2_MI_GFPve_pos  1.1394607000  3.4866857849
## dCA2_Mpos_ACCAAACTCTCGTGGG-1dCA2_MI_GFPve_pos -1.1431969233  2.0519881569
## dCA2_Mpos_ACCATTTTCAAGTCTG-1dCA2_MI_GFPve_pos  0.4345435076  2.9047389351
## dCA2_Mpos_ACCCTTGTCGTGTCAA-1dCA2_MI_GFPve_pos -1.7413452692  2.0008159004
## dCA2_Mpos_ACCGTTCAGGTACAAT-1dCA2_MI_GFPve_pos  1.9108350448  2.0151682220
## dCA2_Mpos_ACCTGTCGTAGGTTTC-1dCA2_MI_GFPve_pos  0.7532432847  2.8343224846
## dCA2_Mpos_ACGATGTCACCGGTCA-1dCA2_MI_GFPve_pos  1.2327791624  2.3663945518
## dCA2_Mpos_ACGGTCGTCCCGTTGT-1dCA2_MI_GFPve_pos  1.5544850998  2.6744385086
## dCA2_Mpos_ACGTAACCATGTTCAG-1dCA2_MI_GFPve_pos  0.9197045736  2.6167507492
## dCA2_Mpos_ACGTACAAGGTCGTAG-1dCA2_MI_GFPve_pos  0.8943735533  3.0903234802
## dCA2_Mpos_ACGTCCTCACGCGGTT-1dCA2_MI_GFPve_pos  1.1243298225  2.3308801971
## dCA2_Mpos_ACGTCCTTCCGGACTG-1dCA2_MI_GFPve_pos -0.9759363361  1.3793297134
## dCA2_Mpos_ACTACGATCGGCTATA-1dCA2_MI_GFPve_pos  1.1724549703  2.0651030861
## dCA2_Mpos_ACTACGATCTTGAGCA-1dCA2_MI_GFPve_pos  1.0533690624  1.9654026352
## dCA2_Mpos_ACTATGGGTTCGCGTG-1dCA2_MI_GFPve_pos  0.5990601353  3.5104818664
## dCA2_Mpos_ACTCTCGAGCCTGAGA-1dCA2_MI_GFPve_pos  1.3291417532  2.3669367157
## dCA2_Mpos_ACTTCCGCAGCTTTGA-1dCA2_MI_GFPve_pos -1.2535745926  1.8084359489
## dCA2_Mpos_ACTTCGCTCGAGAACG-1dCA2_MI_GFPve_pos -0.6006376870  0.8792252861
## dCA2_Mpos_ACTTCGCTCTGGTGGC-1dCA2_MI_GFPve_pos  0.9331752233  2.5069051109
## dCA2_Mpos_AGAAATGGTCTTGCTC-1dCA2_MI_GFPve_pos -1.4720359869 -0.0201053299
## dCA2_Mpos_AGAACCTAGAAGGCTC-1dCA2_MI_GFPve_pos  1.7355178766  2.1879282318
## dCA2_Mpos_AGACTCACACGGGTAA-1dCA2_MI_GFPve_pos  1.6731039457  2.2544279419
## dCA2_Mpos_AGAGAGCCAAAGGAGA-1dCA2_MI_GFPve_pos  1.3581266575  3.1699309669
## dCA2_Mpos_AGAGAGCCAGAATTCC-1dCA2_MI_GFPve_pos -1.4926651545  0.0135550819
## dCA2_Mpos_AGATGAAGTCTTTCTA-1dCA2_MI_GFPve_pos  1.8254244499  2.7636947952
## dCA2_Mpos_AGATGCTGTGCCTACG-1dCA2_MI_GFPve_pos  1.0207883291  1.1395068489
## dCA2_Mpos_AGCGATTAGACTTCCA-1dCA2_MI_GFPve_pos -0.3359729923  1.9075470291
## dCA2_Mpos_AGCGCCAAGGTAGCAC-1dCA2_MI_GFPve_pos  1.9361622505  1.9773860298
## dCA2_Mpos_AGCGTCGCAGTTGGTT-1dCA2_MI_GFPve_pos -1.3229795523  2.0131712280
## dCA2_Mpos_AGCGTCGCATCCTGTC-1dCA2_MI_GFPve_pos -0.8830491133  3.3874044739
## dCA2_Mpos_AGCGTCGGTCAAATCC-1dCA2_MI_GFPve_pos  0.6622066669  2.2938781105
## dCA2_Mpos_AGGACTTAGTTTCGAC-1dCA2_MI_GFPve_pos  0.2777117960  2.2570414863
## dCA2_Mpos_AGGAGGTAGACCTGGA-1dCA2_MI_GFPve_pos  0.0156611525  1.7579999290
## dCA2_Mpos_AGGATAACATCTCAAG-1dCA2_MI_GFPve_pos  1.9088773899  3.0795879684
## dCA2_Mpos_AGGCATTAGGTGGCTA-1dCA2_MI_GFPve_pos -0.0938132457  2.3952017151
## dCA2_Mpos_AGGTCATTCGATACTG-1dCA2_MI_GFPve_pos  1.1030092411  2.9802270256
```

```
## dCA2_Mpos_AGTACCAAGCGCCGTT-1dCA2_MI_GFPve_pos  0.2378240101  2.3692479454
## dCA2_Mpos_AGTAGCTTCCTCTTTC-1dCA2_MI_GFPve_pos -0.3197393544  2.4408030830
## dCA2_Mpos_AGTAGTCAGCGTCGAA-1dCA2_MI_GFPve_pos  0.6449605994  2.0885110222
## dCA2_Mpos_AGTCATGTCTGTAAGC-1dCA2_MI_GFPve_pos -1.7924576826  3.2353692375
## dCA2_Mpos_AGTCTCCAGCTAGAAT-1dCA2_MI_GFPve_pos  1.2252743416  2.5581150375
## dCA2_Mpos_AGTCTCCAGTTGCTCA-1dCA2_MI_GFPve_pos  0.6669757538  1.6378283821
## dCA2_Mpos_AGTCTCCCAATGACCT-1dCA2_MI_GFPve_pos -0.2724454082  1.5097384773
## dCA2_Mpos_AGTGCCGGTGTAAACA-1dCA2_MI_GFPve_pos -0.5774147041  2.3272023521
## dCA2_Mpos_AGTGTTGGTACGCTAT-1dCA2_MI_GFPve_pos  1.3563869171  1.6675463043
## dCA2_Mpos_AGTTAGCTCTGGACTA-1dCA2_MI_GFPve_pos  0.1828210823  2.4841385208
## dCA2_Mpos_ATACCGAAGGCGACAT-1dCA2_MI_GFPve_pos -2.8100025482  0.8400340400
## dCA2_Mpos_ATACCGATCATCGACA-1dCA2_MI_GFPve_pos -0.9614112206  3.1241817795
## dCA2_Mpos_ATACCTTCACAAGGTG-1dCA2_MI_GFPve_pos  0.6704341941  2.2523589454
## dCA2_Mpos_ATAGGCTAGAAACCCG-1dCA2_MI_GFPve_pos  0.1014084689  1.0046916328
## dCA2_Mpos_ATCACGACACCAGCGT-1dCA2_MI_GFPve_pos -0.9959953256  1.7906279884
## dCA2_Mpos_ATCAGGTAGCGTGAAC-1dCA2_MI_GFPve_pos -0.2683494441  2.4778647743
## dCA2_Mpos_ATCCACCGTGGGACAT-1dCA2_MI_GFPve_pos -0.4520399101  1.0467324577
## dCA2_Mpos_ATCCATTGTAGTACGG-1dCA2_MI_GFPve_pos  1.5984997921  3.2365565620
## dCA2_Mpos_ATCCTATGTCGAATTC-1dCA2_MI_GFPve_pos  1.4248490505  2.5391993843
## dCA2_Mpos_ATCGATGCACCCAATA-1dCA2_MI_GFPve_pos  1.7849137955  3.0832691513
## dCA2_Mpos_ATCGGATGTCCACAGC-1dCA2_MI_GFPve_pos -0.9029792376  2.5027199112
## dCA2_Mpos_ATCGTAGAGCCTGTCG-1dCA2_MI_GFPve_pos  1.6523443632  2.3016553245
## dCA2_Mpos_ATGAAAGTCTACCCAC-1dCA2_MI_GFPve_pos -0.6949447222  3.3606148086
## dCA2_Mpos_ATGACCACATGCCGCA-1dCA2_MI_GFPve_pos  1.4358203583  2.2952761970
## dCA2_Mpos_ATGATCGCATGTGGCC-1dCA2_MI_GFPve_pos  0.3171466164  2.8870988212
## dCA2_Mpos_ATGCATGTCACACGAT-1dCA2_MI_GFPve_pos -0.5428244300  0.7581410728
## dCA2_Mpos_ATGTCCCTCTGTGCAA-1dCA2_MI_GFPve_pos  0.6661664419  1.8979535423
## dCA2_Mpos_ATGTCTTTCGCTGTCT-1dCA2_MI_GFPve_pos  1.6656500273  3.0210638366
## dCA2_Mpos_ATTATCCAGCTGTCCG-1dCA2_MI_GFPve_pos -0.0972423606  2.3256492935
## dCA2_Mpos_ATTCACTTCACCGGTG-1dCA2_MI_GFPve_pos -0.1964615654  2.9600215278
## dCA2_Mpos_ATTCCTATCCAAGCTA-1dCA2_MI_GFPve_pos  0.7141116433  2.4076776825
## dCA2_Mpos_ATTCTTGGTCAGTCTA-1dCA2_MI_GFPve_pos  0.0852690511  1.9818730675
## dCA2_Mpos_ATTCTTGTCTTTGCGC-1dCA2_MI_GFPve_pos  0.9392521553  1.1486273132
## dCA2_Mpos_ATTGGGTTCTGACGCG-1dCA2_MI_GFPve_pos -0.6110603638  1.3023276649
## dCA2_Mpos_ATTGTTCAGCCAAGCA-1dCA2_MI_GFPve_pos  1.9606940918  2.2242160164
## dCA2_Mpos_ATTTACCAGAAGCGAA-1dCA2_MI_GFPve_pos  1.7055046730  2.3191209160
## dCA2_Mpos_ATTTCACCAAGTCGTT-1dCA2_MI_GFPve_pos -1.6218659706  1.9368467651
## dCA2_Mpos_ATTTCACTCGTCGCTT-1dCA2_MI_GFPve_pos  0.5533706717  3.5457096420
## dCA2_Mpos_CAACAGTGTCCCGGTA-1dCA2_MI_GFPve_pos  2.3576641254  2.4867611252
## dCA2_Mpos_CAACCTCTCTAGAGCT-1dCA2_MI_GFPve_pos -0.2313633092  2.1092129074
## dCA2_Mpos_CAACGATCAGGCATTT-1dCA2_MI_GFPve_pos -0.7120454736  1.6433272682
## dCA2_Mpos_CAAGCTAAGGCGTTAG-1dCA2_MI_GFPve_pos  2.2697496586  2.0377540909
## dCA2_Mpos_CAAGGGAGTGCAACGA-1dCA2_MI_GFPve_pos  1.6280749016  3.2374368034
## dCA2_Mpos_CAATCGAAGAGCCCAA-1dCA2_MI_GFPve_pos -1.6385464497  2.9993868194
## dCA2_Mpos_CACCAAAAGACCAAGC-1dCA2_MI_GFPve_pos  0.6868375711  2.2321448646
## dCA2_Mpos_CACGGGTAGCGCTTCG-1dCA2_MI_GFPve_pos  0.8738973312  2.6220598541
## dCA2_Mpos_CACTGAAGTCAAGGCA-1dCA2_MI_GFPve_pos -0.3535494782  1.6887765251
## dCA2_Mpos_CAGATACCACCGGAAA-1dCA2_MI_GFPve_pos  2.2118482761  2.5761294685
## dCA2_Mpos_CAGATCAAGTCGAATA-1dCA2_MI_GFPve_pos -0.3651628770  2.4740944229
## dCA2_Mpos_CAGATTGAGATGGTCG-1dCA2_MI_GFPve_pos  0.5103722386  2.1681566559
## dCA2_Mpos_CAGCGTGAGGGATGTC-1dCA2_MI_GFPve_pos -1.5170283146  2.1769738518
## dCA2_Mpos_CAGTGCGAGGAGCAAA-1dCA2_MI_GFPve_pos -0.2835911639  1.2693653427
## dCA2_Mpos_CATACAGGTCTTGCGG-1dCA2_MI_GFPve_pos -0.7296464272  3.4005632721
## dCA2_Mpos_CATCGCTTCTTTCGAT-1dCA2_MI_GFPve_pos  0.2987339966  2.2511039100
## dCA2_Mpos_CATTCATGTAGGACTG-1dCA2_MI_GFPve_pos  0.8305701785  2.6820698105
## dCA2_Mpos_CATTCATGTCTCGGAC-1dCA2_MI_GFPve_pos  0.7982234530  1.9862404190
## dCA2_Mpos_CATTCCGTCGGACAAG-1dCA2_MI_GFPve_pos -1.0923182793  2.9866376243
## dCA2_Mpos_CCAAGCGCACGTATAC-1dCA2_MI_GFPve_pos -0.1280375101  0.7461867653
## dCA2_Mpos_CCAATTTTCGTGTTCC-1dCA2_MI_GFPve_pos -0.9016626902  2.4036736809
## dCA2_Mpos_CCACGAGAGACTCCGC-1dCA2_MI_GFPve_pos  1.1424666815  1.1048431717
## dCA2_Mpos_CCACTTGTCCTAGAGT-1dCA2_MI_GFPve_pos  1.6001482181  1.1481113754
## dCA2_Mpos_CCCGAAGTCATGGAGG-1dCA2_MI_GFPve_pos  1.8899111919  2.1611671768
## dCA2_Mpos_CCCGAAGTCCGGTAAT-1dCA2_MI_GFPve_pos -0.2201844171  1.0189991318
## dCA2_Mpos_CCCTCTCAGTAAGAGG-1dCA2_MI_GFPve_pos -0.9229177065  1.0540671669
## dCA2_Mpos_CCGATCTTCAGCAGAG-1dCA2_MI_GFPve_pos -1.6131166048  2.0152511917
## dCA2_Mpos_CCGATGGGTATTTCCT-1dCA2_MI_GFPve_pos  0.1913468115  2.7884774528
## dCA2_Mpos_CCGATGGTCCAAGCCG-1dCA2_MI_GFPve_pos  0.7766636185  1.6829510055
## dCA2_Mpos_CCGCAAGTCGGCTGAC-1dCA2_MI_GFPve_pos  1.7672988348  3.1458196960
## dCA2_Mpos_CCGGACAAGTGCAACG-1dCA2_MI_GFPve_pos  2.5402524643  2.8336148582
## dCA2_Mpos_CCGTGAGCATGTGGTT-1dCA2_MI_GFPve_pos -0.9446797795  3.2974605881
```

```
## dCA2_Mpos_CCTCACAAGTATAGGT-1dCA2_MI_GFPve_pos  1.4305035047  1.7572021805
## dCA2_Mpos_CCTCAGTGTAGTCACT-1dCA2_MI_GFPve_pos -1.4300203152  3.2926988922
## dCA2_Mpos_CCTCCTCCAGTAGTGG-1dCA2_MI_GFPve_pos -0.8279872961  2.0237732254
## dCA2_Mpos_CCTCTAGTCTTCGTAT-1dCA2_MI_GFPve_pos -1.4860292502  1.9776449524
## dCA2_Mpos_CCTCTCCAGGTCTTTG-1dCA2_MI_GFPve_pos -1.3841557331  1.5727310501
## dCA2_Mpos_CCTGCATTCGGTCACG-1dCA2_MI_GFPve_pos -1.0331213303  2.5519576393
## dCA2_Mpos_CCTTCAGAGTCACGCC-1dCA2_MI_GFPve_pos  0.4799041920  2.2411914192
## dCA2_Mpos_CCTTGTGGTATAATGG-1dCA2_MI_GFPve_pos  0.1124094002  2.3719115577
## dCA2_Mpos_CCTTGTGTCACGATAC-1dCA2_MI_GFPve_pos  1.1508746319  3.4221077286
## dCA2_Mpos_CGACAGCTCGCTAGCG-1dCA2_MI_GFPve_pos  1.1279972963  2.7310452781
## dCA2_Mpos_CGAGTGCGTAGGTCAG-1dCA2_MI_GFPve_pos  1.2632640772  2.2743068062
## dCA2_Mpos_CGATCGGGTGTGCCTG-1dCA2_MI_GFPve_pos  0.9119259290  2.6584921203
## dCA2_Mpos_CGATGCGAGCGATGGT-1dCA2_MI_GFPve_pos -0.5840417273  2.7541423164
## dCA2_Mpos_CGATGGCCAACAAAGT-1dCA2_MI_GFPve_pos  2.2041447334  2.4472976051
## dCA2_Mpos_CGCAGGTCAAGTCGTT-1dCA2_MI_GFPve_pos -1.4446179695  2.3588538490
## dCA2_Mpos_CGCATGGTCGGTGTTA-1dCA2_MI_GFPve_pos -0.8164712019  2.7085247360
## dCA2_Mpos_CGCCAGATCGCCGATG-1dCA2_MI_GFPve_pos  0.9030521087 -1.5240909733
## dCA2_Mpos_CGCGTGAAGAGTATAC-1dCA2_MI_GFPve_pos -0.1818650575  2.2319922767
## dCA2_Mpos_CGGGACTGTAGTTAGA-1dCA2_MI_GFPve_pos  0.2380846612  3.2592201553
## dCA2_Mpos_CGGGTGTTCTGATGGT-1dCA2_MI_GFPve_pos -1.7257140703  1.0920772873
## dCA2_Mpos_CGGTCAGAGATTGCGG-1dCA2_MI_GFPve_pos -0.2065891787  3.5254421554
## dCA2_Mpos_CGGTCAGGTTACGATC-1dCA2_MI_GFPve_pos  0.2744628362  2.4732590042
## dCA2_Mpos_CGTAGTAGTGCTCTTC-1dCA2_MI_GFPve_pos  1.0770470075  3.0159120880
## dCA2_Mpos_CGTCAAACACGACAAG-1dCA2_MI_GFPve_pos  0.4740573339  1.3469925247
## dCA2_Mpos_CGTGTCTAGTGGCCTC-1dCA2_MI_GFPve_pos  0.6333958678  2.8826642357
## dCA2_Mpos_CGTTGGGGTAGACGTG-1dCA2_MI_GFPve_pos  0.1924515240  3.1917887054
## dCA2_Mpos_CTAAGTGGTGTAAATG-1dCA2_MI_GFPve_pos -0.2476120420  2.3874802910
## dCA2_Mpos_CTACGGGTCTAGAGCT-1dCA2_MI_GFPve_pos -1.1972112484  1.8295092903
## dCA2_Mpos_CTAGACAAGTGATAAC-1dCA2_MI_GFPve_pos  0.2537261717  0.7172346435
## dCA2_Mpos_CTCAACCAGGGATCTG-1dCA2_MI_GFPve_pos  0.9628873043  1.7694139801
## dCA2_Mpos_CTCAAGAGTCGTGATT-1dCA2_MI_GFPve_pos -0.0880052499  2.5597901665
## dCA2_Mpos_CTCATCGGTACGAGCA-1dCA2_MI_GFPve_pos  1.4364857845  2.7452373825
## dCA2_Mpos_CTCATGCTCTCATAGG-1dCA2_MI_GFPve_pos  0.2462584250  1.6275916420
## dCA2_Mpos_CTCGAGGCATGAAAGT-1dCA2_MI_GFPve_pos -1.2202244826  2.9966936432
## dCA2_Mpos_CTGAGCGAGGAGAATG-1dCA2_MI_GFPve_pos  1.7986263208  3.1868362747
## dCA2_Mpos_CTGCAGGCAATTGCGT-1dCA2_MI_GFPve_pos  0.6145917945  1.6508479439
## dCA2_Mpos_CTGCATCAGGACTAAT-1dCA2_MI_GFPve_pos -2.2869465180  1.7735042892
## dCA2_Mpos_CTGCCATAGGAGATAG-1dCA2_MI_GFPve_pos  2.1723666363  2.7429762207
## dCA2_Mpos_CTGCCATTCCTCGATC-1dCA2_MI_GFPve_pos -1.6022212334  2.1953106247
## dCA2_Mpos_CTGCGAGAGCCTGCCA-1dCA2_MI_GFPve_pos  0.0277112745  2.8706202827
## dCA2_Mpos_CTGGACGCACGTGAGA-1dCA2_MI_GFPve_pos  0.8558945827  2.6299200378
## dCA2_Mpos_CTGGACGGTGACTAAA-1dCA2_MI_GFPve_pos -1.3059091397  1.7674088798
## dCA2_Mpos_CTGGACGTCAGACCTA-1dCA2_MI_GFPve_pos  1.5462538175  3.0645838104
## dCA2_Mpos_CTGGCAGCATGCGTGC-1dCA2_MI_GFPve_pos -0.7448153324  1.6206016861
## dCA2_Mpos_CTGGTCTTCCAACCGG-1dCA2_MI_GFPve_pos  2.2013964824  3.1504369102
## dCA2_Mpos_CTGGTCTTCGAGATAA-1dCA2_MI_GFPve_pos -0.4917137749  2.1529589020
## dCA2_Mpos_CTGTAGACATGAGATA-1dCA2_MI_GFPve_pos -0.3151337929  0.7673831306
## dCA2_Mpos_CTGTGAATCCGGACGT-1dCA2_MI_GFPve_pos  0.1641362183  3.3575344406
## dCA2_Mpos_CTGTGGGCATGCAGGA-1dCA2_MI_GFPve_pos  2.1230216198  2.3574462257
## dCA2_Mpos_CTGTGGGTCATGCCCT-1dCA2_MI_GFPve_pos -0.2651353679  2.1699905716
## dCA2_Mpos_CTTAGGATCTCCGAGG-1dCA2_MI_GFPve_pos  1.0086621218  2.2015900932
## dCA2_Mpos_CTTCGGTCAGGACTTT-1dCA2_MI_GFPve_pos -0.9577593632  1.8248176895
## dCA2_Mpos_CTTGAGAGTTCAGCGC-1dCA2_MI_GFPve_pos  0.9686693125  2.8021631561
## dCA2_Mpos_CTTTCAAAGGAGTCTG-1dCA2_MI_GFPve_pos -0.8230698533  1.7931270920
## dCA2_Mpos_GAAACCTCACATGGTT-1dCA2_MI_GFPve_pos  1.1369747095  2.9736781441
## dCA2_Mpos_GAAGAATTCGGCTATA-1dCA2_MI_GFPve_pos -0.7148886152  3.4212494217
## dCA2_Mpos_GAAGCGAAGACTTAAG-1dCA2_MI_GFPve_pos  1.2467869692  3.2672796570
## dCA2_Mpos_GAATAGACAGATACCT-1dCA2_MI_GFPve_pos  0.6737420730  2.2089362465
## dCA2_Mpos_GAATAGAGTGCGACAA-1dCA2_MI_GFPve_pos -0.9991464443  2.7619538627
## dCA2_Mpos_GAATCACAGGCATGCA-1dCA2_MI_GFPve_pos  0.8170558386  2.7884292923
## dCA2_Mpos_GAATCGTTCGCGGACT-1dCA2_MI_GFPve_pos -0.8470173307  2.8557362877
## dCA2_Mpos_GACCCAGCAAGAGGCT-1dCA2_MI_GFPve_pos -1.2304861374  0.6576233230
## dCA2_Mpos_GACCCTTCACCTAAAC-1dCA2_MI_GFPve_pos  0.4800890856  1.0847044311
## dCA2_Mpos_GACTCTCGTAAGGCTG-1dCA2_MI_GFPve_pos  0.0853661768  2.2133875213
## dCA2_Mpos_GACTGATTCGCGTAGC-1dCA2_MI_GFPve_pos -0.5490646191  1.5379667602
## dCA2_Mpos_GAGACCCCACTGGCGT-1dCA2_MI_GFPve_pos -1.5282832213  2.0870404564
## dCA2_Mpos_GAGAGGTAGTTGCATC-1dCA2_MI_GFPve_pos  1.8929374389  3.1845617615
## dCA2_Mpos_GAGAGGTCAAGATGTA-1dCA2_MI_GFPve_pos  1.2583222561  1.7180309616
## dCA2_Mpos_GAGATGGTCTCCTGTG-1dCA2_MI_GFPve_pos  0.1601999872  2.8910184227
## dCA2_Mpos_GAGCTGCCAGCTAACT-1dCA2_MI_GFPve_pos -2.2688369580  0.5267501198
```

```
## dCA2_Mpos_GAGCTGCGTAGCTTAC-1dCA2_MI_GFPve_pos -1.2463875838  1.5804658256
## dCA2_Mpos_GAGCTGCTCCTATTGT-1dCA2_MI_GFPve_pos  1.2204916649  2.9040751777
## dCA2_Mpos_GAGGGTAAGTAGACCG-1dCA2_MI_GFPve_pos -1.3069994278  1.6421656929
## dCA2_Mpos_GAGTCTAGTGATACCT-1dCA2_MI_GFPve_pos  0.4682159595  3.3619699798
## dCA2_Mpos_GAGTGAGAGTCAGCGA-1dCA2_MI_GFPve_pos -0.0683278493  2.5873184524
## dCA2_Mpos_GAGTGTTAGGGCGAGA-1dCA2_MI_GFPve_pos  1.3384316139  3.4847336136
## dCA2_Mpos_GAGTTACAGAGCTGAC-1dCA2_MI_GFPve_pos  1.0148543291  2.1500230156
## dCA2_Mpos_GAGTTTGGTTGGTACT-1dCA2_MI_GFPve_pos  0.9147936277  3.2397709213
## dCA2_Mpos_GATCAGTAGCGAGAAA-1dCA2_MI_GFPve_pos -0.3271219410  2.5044060074
## dCA2_Mpos_GATCAGTAGGCAGGTT-1dCA2_MI_GFPve_pos  1.3627995424  2.7208605133
## dCA2_Mpos_GATCAGTAGTTCGGTT-1dCA2_MI_GFPve_pos  0.1075022988  2.4363780342
## dCA2_Mpos_GATTCTTTCAGAATAG-1dCA2_MI_GFPve_pos -1.3640818424  2.4338288627
## dCA2_Mpos_GCAGTTATCGGAATGG-1dCA2_MI_GFPve_pos  2.4425065689  2.6479182564
## dCA2_Mpos_GCATCGGGTTGCTCCT-1dCA2_MI_GFPve_pos  1.2577260904  3.0401459060
## dCA2_Mpos_GCCATGGCAAGGCGTA-1dCA2_MI_GFPve_pos -1.4163347311  2.5007839523
## dCA2_Mpos_GCCATGGGTTCCTACC-1dCA2_MI_GFPve_pos -0.4964864381  3.3958807312
## dCA2_Mpos_GCCGATGTCGCGGACT-1dCA2_MI_GFPve_pos  2.4064274006  3.0041828476
## dCA2_Mpos_GCGATCGAGAGCTGCA-1dCA2_MI_GFPve_pos  2.3935804539  3.0011234604
## dCA2_Mpos_GCGGATCAGAGAATCT-1dCA2_MI_GFPve_pos  1.9465394192  3.0000095688
## dCA2_Mpos_GCGGATCTCAACGTGT-1dCA2_MI_GFPve_pos  0.7300108843  2.5165977798
## dCA2_Mpos_GCGTGCAAGGTTCCAT-1dCA2_MI_GFPve_pos -0.8949123450  3.0674639068
## dCA2_Mpos_GCGTGCAAGTTTGGCT-1dCA2_MI_GFPve_pos -0.2000209369  1.7954836212
## dCA2_Mpos_GCTTTCGGTAGTGGCA-1dCA2_MI_GFPve_pos -0.0898565181  1.3404026352
## dCA2_Mpos_GGAACCCCAACAGCCC-1dCA2_MI_GFPve_pos -0.7815670796  1.6490011535
## dCA2_Mpos_GGAACCCGTAGTCTGT-1dCA2_MI_GFPve_pos  0.7943177991  1.6613178573
## dCA2_Mpos_GGAACCCTCTCTGGTC-1dCA2_MI_GFPve_pos  0.1658548884  3.0085230194
## dCA2_Mpos_GGAATCTCAGGGCTTC-1dCA2_MI_GFPve_pos  1.6705688410  3.0461626373
## dCA2_Mpos_GGAATGGGTGTTAGCT-1dCA2_MI_GFPve_pos  2.0323295765  1.9050364815
## dCA2_Mpos_GGAGGTATCATCGGGC-1dCA2_MI_GFPve_pos  0.5620199613  2.0241022430
## dCA2_Mpos_GGATCTAGTCGTGTTA-1dCA2_MI_GFPve_pos -1.1774998732  0.1216502510
## dCA2_Mpos_GGATGTTAGGAAGTCC-1dCA2_MI_GFPve_pos  0.4947732858  2.0941996895
## dCA2_Mpos_GGCGTCAGTGACCTGC-1dCA2_MI_GFPve_pos -0.5071403809  1.7074256264
## dCA2_Mpos_GGGACCTAGTTACGGG-1dCA2_MI_GFPve_pos  1.2193219833  3.2055278145
## dCA2_Mpos_GGGAGTAGTTCGAGCC-1dCA2_MI_GFPve_pos -0.0662406064  1.8453712784
## dCA2_Mpos_GGGAGTATCAGCGTCG-1dCA2_MI_GFPve_pos -1.5012001820  2.3810577713
## dCA2_Mpos_GGGCGTTTCTTACCAT-1dCA2_MI_GFPve_pos  0.8937337570  2.9402790390
## dCA2_Mpos_GGGCTCAGTTCGGTAT-1dCA2_MI_GFPve_pos -1.3967503138  1.6777754150
## dCA2_Mpos_GGGTAGAAGGAGAATG-1dCA2_MI_GFPve_pos  0.3642910890  2.7253842674
## dCA2_Mpos_GGGTCTGAGGGATCTG-1dCA2_MI_GFPve_pos -0.2987504728  3.3917637192
## dCA2_Mpos_GGGTCTGAGTAGACCG-1dCA2_MI_GFPve_pos -0.1030493133  2.4469662033
## dCA2_Mpos_GGGTCTGGTCTACATG-1dCA2_MI_GFPve_pos -1.3335740395  1.1317944847
## dCA2_Mpos_GGTAGAGAGCCTCATA-1dCA2_MI_GFPve_pos  1.9690287285  2.9303407989
## dCA2_Mpos_GGTGAAGGTAACGATA-1dCA2_MI_GFPve_pos  0.5610341005  3.0965633713
## dCA2_Mpos_GGTGTCGGTTCTCCAC-1dCA2_MI_GFPve_pos -0.3704013802  2.1557412468
## dCA2_Mpos_GGTGTCGGTTTACCTT-1dCA2_MI_GFPve_pos  1.5882630520  3.1705246292
## dCA2_Mpos_GGTGTCGTCGCGTCGA-1dCA2_MI_GFPve_pos  0.1662804060  2.7455864273
## dCA2_Mpos_GGTTAACAGGAAAGTG-1dCA2_MI_GFPve_pos  0.9506499939  3.3336296402
## dCA2_Mpos_GGTTCTCTCGCCAGTG-1dCA2_MI_GFPve_pos  0.6109788589  3.1349883400
## dCA2_Mpos_GTAACCACATAGCTGT-1dCA2_MI_GFPve_pos  1.3999204807  1.7138138138
## dCA2_Mpos_GTAGAGGAGGAAAGAC-1dCA2_MI_GFPve_pos  2.0637550525  2.3033066116
## dCA2_Mpos_GTAGATCAGTCATACC-1dCA2_MI_GFPve_pos -0.5789604910  2.4030022941
## dCA2_Mpos_GTAGCTAGTAATTGGA-1dCA2_MI_GFPve_pos -0.0092714853  2.9402361236
## dCA2_Mpos_GTAGGTTCATTGTAGC-1dCA2_MI_GFPve_pos  1.2882974319  2.5417194687
## dCA2_Mpos_GTCACTCTCGGTAGGA-1dCA2_MI_GFPve_pos -0.3723603465  2.6419549309
## dCA2_Mpos_GTCCCATGTAGGGAGG-1dCA2_MI_GFPve_pos -0.4413870402  3.1259084068
## dCA2_Mpos_GTCGAATGTACAAACA-1dCA2_MI_GFPve_pos  0.3125959806  2.0692186676
## dCA2_Mpos_GTCGTTCGTGTTCCTC-1dCA2_MI_GFPve_pos  0.2542637460  2.7447686516
## dCA2_Mpos_GTCTAGAGTGATTCTG-1dCA2_MI_GFPve_pos  0.9633282594  2.8549542747
## dCA2_Mpos_GTCTCACCATCGAAGG-1dCA2_MI_GFPve_pos  2.0656268768  3.0720348678
## dCA2_Mpos_GTCTCACGTACCTAGT-1dCA2_MI_GFPve_pos -0.6127579637  0.6249184929
## dCA2_Mpos_GTGACGCAGTACGAGC-1dCA2_MI_GFPve_pos  0.7014766507  2.4180927597
## dCA2_Mpos_GTGAGGATCGTTCCTG-1dCA2_MI_GFPve_pos  1.1028717928  1.2872639023
## dCA2_Mpos_GTGCTTCGTTAGGACG-1dCA2_MI_GFPve_pos  0.3248904221  2.6875863395
## dCA2_Mpos_GTGGAGACAAACCACT-1dCA2_MI_GFPve_pos -2.0429186650  2.6415777527
## dCA2_Mpos_GTGTAACCAGATTTCG-1dCA2_MI_GFPve_pos -1.3983944483  2.0151710830
## dCA2_Mpos_GTTACCCGTTCGGTAT-1dCA2_MI_GFPve_pos  1.8978834324  3.0283251129
## dCA2_Mpos_GTTACGACACCGTCGA-1dCA2_MI_GFPve_pos  1.4064380102  2.6457553230
## dCA2_Mpos_GTTAGACGTCTTGAAC-1dCA2_MI_GFPve_pos  1.4049691133  2.0570412002
## dCA2_Mpos_GTTGAACAGTCACGCC-1dCA2_MI_GFPve_pos -0.9065477081  2.9215460144
## dCA2_Mpos_GTTTACTCATGTCTAG-1dCA2_MI_GFPve_pos  0.8713149004  2.4451589905
```

```
## dCA2_Mpos_TAACACGTCGTTCTCG-1dCA2_MI_GFPve_pos -1.4324453898  1.9037890754
## dCA2_Mpos_TAACGACAGGAAACGA-1dCA2_MI_GFPve_pos  1.4553680591  2.0375480972
## dCA2_Mpos_TAAGTCGGTAACTTCG-1dCA2_MI_GFPve_pos  2.1128797703  3.0728140198
## dCA2_Mpos_TACAACGGTGTTAACC-1dCA2_MI_GFPve_pos -1.1311118431  1.2287693344
## dCA2_Mpos_TACCCACGTTCTCCCA-1dCA2_MI_GFPve_pos -1.1063187547  2.8238787971
## dCA2_Mpos_TACCTGCGTGTCATGT-1dCA2_MI_GFPve_pos  1.7157102995  3.1724200569
## dCA2_Mpos_TACTTACAGCTCTGTA-1dCA2_MI_GFPve_pos -1.2539410420  1.8455434166
## dCA2_Mpos_TACTTACCAGAAATTG-1dCA2_MI_GFPve_pos  1.7312396936  2.4273429237
## dCA2_Mpos_TAGAGTCCAATGGGTG-1dCA2_MI_GFPve_pos  1.5942757301  2.7473969780
## dCA2_Mpos_TAGAGTCCAGTCCGTG-1dCA2_MI_GFPve_pos -0.3562304057  2.8866763435
## dCA2_Mpos_TAGAGTCGTACGTGAG-1dCA2_MI_GFPve_pos  1.5849793129  3.3286848388
## dCA2_Mpos_TAGCACACAACCTATG-1dCA2_MI_GFPve_pos -0.8303390570  1.7162957512
## dCA2_Mpos_TAGGAGGAGATTGCGG-1dCA2_MI_GFPve_pos  0.9619578295  3.4722338043
## dCA2_Mpos_TAGTGCAGTTAGGCCC-1dCA2_MI_GFPve_pos  0.0194009565  2.6885018669
## dCA2_Mpos_TAGTGCAGTTTCACTT-1dCA2_MI_GFPve_pos  1.0514538460  1.7020616852
## dCA2_Mpos_TATATCCAGGTGAGAA-1dCA2_MI_GFPve_pos  0.6348685198  3.1620989166
## dCA2_Mpos_TATCCTAAGCTCTTCC-1dCA2_MI_GFPve_pos  1.3854152136  2.0759945236
## dCA2_Mpos_TATCCTACAGAGAAAG-1dCA2_MI_GFPve_pos -1.2486290760 -0.5150055565
## dCA2_Mpos_TATCTGTCATAGCACT-1dCA2_MI_GFPve_pos  0.3663417868  2.4597287498
## dCA2_Mpos_TATCTTGGTCGGATTT-1dCA2_MI_GFPve_pos  1.7401739530  2.8964624725
## dCA2_Mpos_TATGTTCGTTAGGACG-1dCA2_MI_GFPve_pos -1.0902579255  1.9121823631
## dCA2_Mpos_TATTGGGTCTTCCAGC-1dCA2_MI_GFPve_pos -2.0038914509  0.2063627563
## dCA2_Mpos_TCAAGACAGTTTGAGA-1dCA2_MI_GFPve_pos  0.5043047838  1.8343510948
## dCA2_Mpos_TCAAGCAGTTGGCCGT-1dCA2_MI_GFPve_pos  0.9761696033  1.3033123337
## dCA2_Mpos_TCACATTGTTAGGGAC-1dCA2_MI_GFPve_pos  1.3269908600  3.2031722389
## dCA2_Mpos_TCACGCTCAATAGGGC-1dCA2_MI_GFPve_pos  2.2324969940  2.4900999389
## dCA2_Mpos_TCACTATTCCTTGAAG-1dCA2_MI_GFPve_pos  0.3553717904  1.0818639122
## dCA2_Mpos_TCAGCCTGTGTTCCTC-1dCA2_MI_GFPve_pos -1.5631397791  2.0026307426
## dCA2_Mpos_TCAGGGCAGGTGATAT-1dCA2_MI_GFPve_pos -2.0277229376  1.7645054184
## dCA2_Mpos_TCAGTGAGTTAATGAG-1dCA2_MI_GFPve_pos  0.8590003423  2.9816994987
## dCA2_Mpos_TCATTACCAATACGCT-1dCA2_MI_GFPve_pos  0.3954121642  2.9547906242
## dCA2_Mpos_TCATTACTCTACGCGG-1dCA2_MI_GFPve_pos -0.1132463045  0.7809062324
## dCA2_Mpos_TCATTCAGTCGTTGCG-1dCA2_MI_GFPve_pos  0.9605201654  2.6325526558
## dCA2_Mpos_TCATTTGAGATGATTG-1dCA2_MI_GFPve_pos -0.6343398221  1.1520638786
## dCA2_Mpos_TCATTTGTCGCCAACG-1dCA2_MI_GFPve_pos  0.4620165400  1.5680995308
## dCA2_Mpos_TCCACGTCATATCTGG-1dCA2_MI_GFPve_pos  0.0299703799  2.4058504425
## dCA2_Mpos_TCCAGAAAGTATTAGG-1dCA2_MI_GFPve_pos -1.8530541487  2.0200853668
## dCA2_Mpos_TCCCAGTGTGGGCTCT-1dCA2_MI_GFPve_pos  1.8320602350  2.5325379692
## dCA2_Mpos_TCCGATCAGTAGACAT-1dCA2_MI_GFPve_pos  0.8968781404  3.4281797729
## dCA2_Mpos_TCCGTGTTCCCAGCGA-1dCA2_MI_GFPve_pos  1.7268420629  3.1161632858
## dCA2_Mpos_TCCTGCAGTACAGCGA-1dCA2_MI_GFPve_pos  0.2446624808  2.3267655693
## dCA2_Mpos_TCCTTCTGTACGGTTT-1dCA2_MI_GFPve_pos  0.5595320635  1.3524289451
## dCA2_Mpos_TCCTTTCCAGCGTATT-1dCA2_MI_GFPve_pos  0.5638249688  2.9363651596
## dCA2_Mpos_TCGACCTGTAAGTCAA-1dCA2_MI_GFPve_pos  0.0763918929  2.5753126465
## dCA2_Mpos_TCGATTTAGGTGGTTG-1dCA2_MI_GFPve_pos -0.2767137490  2.4384765945
## dCA2_Mpos_TCGCAGGAGCAGGGAG-1dCA2_MI_GFPve_pos  0.8506460361  2.4464774452
## dCA2_Mpos_TCGCAGGTCGGCGATC-1dCA2_MI_GFPve_pos  1.4878040724  2.9857087456
## dCA2_Mpos_TCGCTTGGTAGTCGGA-1dCA2_MI_GFPve_pos  0.4129812293  2.5412364326
## dCA2_Mpos_TCGGGACCAGTAACCT-1dCA2_MI_GFPve_pos -1.0158436723  3.3281059585
## dCA2_Mpos_TCGGGACTCGGAGATG-1dCA2_MI_GFPve_pos  0.9311294727  1.7964020096
## dCA2_Mpos_TCGGTCTAGGTCGTGA-1dCA2_MI_GFPve_pos  0.0032736234  2.9424076401
## dCA2_Mpos_TCGTAGATCATGAGAA-1dCA2_MI_GFPve_pos  0.1306024067  2.5240836464
## dCA2_Mpos_TCGTGCTGTTCTAAGC-1dCA2_MI_GFPve_pos -1.1967450209  2.0675306640
## dCA2_Mpos_TCGTGCTTCAAACGTC-1dCA2_MI_GFPve_pos -1.3564065523  1.7705893837
## dCA2_Mpos_TCGTGGGTCAGTCAGT-1dCA2_MI_GFPve_pos  0.4400238566  1.4726600967
## dCA2_Mpos_TCTATCACATGAATCC-1dCA2_MI_GFPve_pos -0.2588600270  1.9672808967
## dCA2_Mpos_TCTCAGCAGAACGCGT-1dCA2_MI_GFPve_pos -1.2665376492  1.6318106972
## dCA2_Mpos_TCTTCCTAGAGTCTGG-1dCA2_MI_GFPve_pos  0.7782482080  2.5830278717
## dCA2_Mpos_TCTTCCTTCTCGAGTA-1dCA2_MI_GFPve_pos -1.3606217928  2.3522224747
## dCA2_Mpos_TGAATCGCAAGACGAC-1dCA2_MI_GFPve_pos -1.0500126906  1.2787061058
## dCA2_Mpos_TGAATGCTCGTCGACG-1dCA2_MI_GFPve_pos -0.0123870052  0.8642530762
## dCA2_Mpos_TGAGCATCACTGCGTG-1dCA2_MI_GFPve_pos  1.2947175674  1.5832176529
## dCA2_Mpos_TGAGCGCGTTTCGTGA-1dCA2_MI_GFPve_pos  0.4906066828  2.9093427978
## dCA2_Mpos_TGAGGTTAGAGTCCGA-1dCA2_MI_GFPve_pos -1.2048172779  1.6473207794
## dCA2_Mpos_TGAGGTTAGGTTCAGG-1dCA2_MI_GFPve_pos -1.0966844983 -0.8946025051
## dCA2_Mpos_TGAGGTTGTAAGATAC-1dCA2_MI_GFPve_pos  0.8200556211  2.2952957474
## dCA2_Mpos_TGATTCTCACATTGTG-1dCA2_MI_GFPve_pos -0.1549367109  1.9231744133
## dCA2_Mpos_TGATTTCAGCACCGTC-1dCA2_MI_GFPve_pos  0.2055962555  2.1227202736
## dCA2_Mpos_TGCAGATCAAGTCATC-1dCA2_MI_GFPve_pos  0.9106859140  1.6595473610
## dCA2_Mpos_TGCGACGCATAAGCGG-1dCA2_MI_GFPve_pos  1.6348755531  2.2130837761
```

```
## dCA2_Mpos_TGCGGCAAGCTCCATA-1dCA2_MI_GFPve_pos -2.1119823284  1.1488089882
## dCA2_Mpos_TGCGGCAGTCAGATTC-1dCA2_MI_GFPve_pos  0.3617867999  2.4651317917
## dCA2_Mpos_TGCGGCAGTCTACAGT-1dCA2_MI_GFPve_pos  1.7540164165  2.0478186928
## dCA2_Mpos_TGCGGGTGTACTAGCT-1dCA2_MI_GFPve_pos -1.6334342785  2.9363704048
## dCA2_Mpos_TGCTCCAAGTGCCGAA-1dCA2_MI_GFPve_pos -0.0919761784  3.0820670448
## dCA2_Mpos_TGCTTGCGTGATAGAT-1dCA2_MI_GFPve_pos  1.8328461818  2.2382321678
## dCA2_Mpos_TGGAGGATCTAAGAAG-1dCA2_MI_GFPve_pos -0.8357866235  2.7360987984
## dCA2_Mpos_TGGGAAGGTGTCCCTT-1dCA2_MI_GFPve_pos -1.2009733744  2.6948128067
## dCA2_Mpos_TGGGAAGTCGTAGGAG-1dCA2_MI_GFPve_pos  1.7443102770  2.8208461128
## dCA2_Mpos_TGGGAGAAGAAGTATC-1dCA2_MI_GFPve_pos -0.7096770234  2.0845313392
## dCA2_Mpos_TGGGCGTCAAGCACAG-1dCA2_MI_GFPve_pos -0.1707535291  1.9263763748
## dCA2_Mpos_TGGGCTGGTAAGCTCT-1dCA2_MI_GFPve_pos  0.2005693190  2.7016754471
## dCA2_Mpos_TGGTGATAGTAAGCAT-1dCA2_MI_GFPve_pos -0.0716765501  1.1034479461
## dCA2_Mpos_TGTACAGAGTCGGGAT-1dCA2_MI_GFPve_pos -1.3972993918  1.9147773109
## dCA2_Mpos_TGTCCCATCCACGTCT-1dCA2_MI_GFPve_pos -0.6500235326  2.3010969482
## dCA2_Mpos_TGTCCTGGTAACATGA-1dCA2_MI_GFPve_pos -0.3482754059  1.3914490066
## dCA2_Mpos_TGTGAGTAGTTGCGAG-1dCA2_MI_GFPve_pos  1.6033396892  3.1714082084
## dCA2_Mpos_TGTGTGATCTAGCAAC-1dCA2_MI_GFPve_pos  1.6848056488  1.7829122864
## dCA2_Mpos_TGTTCCGCAGCCGGTT-1dCA2_MI_GFPve_pos  1.2742736511  2.7688598953
## dCA2_Mpos_TGTTCTAGTACGCTAT-1dCA2_MI_GFPve_pos  1.6166471414  2.7169585548
## dCA2_Mpos_TGTTCTAGTCAGTCGC-1dCA2_MI_GFPve_pos -0.8737717099  0.9401388489
## dCA2_Mpos_TGTTCTAGTTAGGCTT-1dCA2_MI_GFPve_pos -1.3188226052  1.1892075859
## dCA2_Mpos_TGTTTGTCACAACCGC-1dCA2_MI_GFPve_pos -2.3562209435  2.7595043503
## dCA2_Mpos_TTACGTTAGGATACAT-1dCA2_MI_GFPve_pos -1.1137074657  1.8647413574
## dCA2_Mpos_TTACTGTAGGCTCAAG-1dCA2_MI_GFPve_pos -0.2899832107  2.3292732559
## dCA2_Mpos_TTAGTCTAGGTCATCT-1dCA2_MI_GFPve_pos -1.8574603625  1.7249722801
## dCA2_Mpos_TTCAATCGTTGCCGCA-1dCA2_MI_GFPve_pos -0.0959510259  1.3208899818
## dCA2_Mpos_TTCCGGTAGCCTATTG-1dCA2_MI_GFPve_pos  0.7912820153  2.4898524605
## dCA2_Mpos_TTCCTAAGTTTCGATG-1dCA2_MI_GFPve_pos -1.0603937335  2.3866224609
## dCA2_Mpos_TTCCTTCAGAGCAAGA-1dCA2_MI_GFPve_pos -1.3796539135  1.5618410431
## dCA2_Mpos_TTCGATTGTTGGAGGT-1dCA2_MI_GFPve_pos -0.0390802599  3.2690439544
## dCA2_Mpos_TTCGATTTCCGATGCG-1dCA2_MI_GFPve_pos -1.5916502304  1.6046648346
## dCA2_Mpos_TTCTAACAGGAAGTGA-1dCA2_MI_GFPve_pos -0.1022606544  2.4435515724
## dCA2_Mpos_TTCTGTATCGGACGTC-1dCA2_MI_GFPve_pos  0.8111490779  3.2207351051
## dCA2_Mpos_TTCTTGAAGCAAGTGC-1dCA2_MI_GFPve_pos -1.5046825237  2.4809360824
## dCA2_Mpos_TTCTTGACACATCATG-1dCA2_MI_GFPve_pos  1.4000190668  1.8393917404
## dCA2_Mpos_TTGAGTGGTAAGGAGA-1dCA2_MI_GFPve_pos -1.4519249029  1.5855532013
## dCA2_Mpos_TTGATGGAGTAACCTC-1dCA2_MI_GFPve_pos -0.9184409805  2.9130635582
## dCA2_Mpos_TTGATGGAGTCGCTAT-1dCA2_MI_GFPve_pos  0.8624904327  2.4655585609
## dCA2_Mpos_TTGCCTGCACGTCGGT-1dCA2_MI_GFPve_pos  0.0369911812  2.8579702698
## dCA2_Mpos_TTGCGTCGTTGAAGTA-1dCA2_MI_GFPve_pos -2.2277874775  0.9369154297
## dCA2_Mpos_TTGCTGCCATCTCATT-1dCA2_MI_GFPve_pos -1.1961008139  2.3878259979
## dCA2_Mpos_TTGGGCGCACCTGAAT-1dCA2_MI_GFPve_pos -0.0981940456  3.3292303406
## dCA2_Mpos_TTGTGGACACCTGCTT-1dCA2_MI_GFPve_pos  0.9843452148  1.6389198623
## dCA2_Mpos_TTGTGGATCCATCGTC-1dCA2_MI_GFPve_pos -0.7030714579  3.1932387672
## dCA2_Mpos_TTGTTTGAGGCTGAAC-1dCA2_MI_GFPve_pos  0.3163336389  2.1968751274
## dCA2_Mpos_TTTCCTCAGCATGCGA-1dCA2_MI_GFPve_pos  0.5204143696  2.0022282921
## dCA2_Mpos_TTTCGATCAGTGGGTA-1dCA2_MI_GFPve_pos  0.5237417989  3.1516609512
## dCA2_Mpos_TTTGACTAGACCCTTA-1dCA2_MI_GFPve_pos -0.2432162888  2.8418293320
## dCA2_Mpos_TTTGTTGCATGTGTCA-1dCA2_MI_GFPve_pos -2.2462293930  2.9439325653
## dCA2_Fpos_AAACGAACAATTCTCT-1dCA2_FI_GFPve_pos  0.0150269292  2.5331082664
## dCA2_Fpos_AAAGGATTCCGGTAAT-1dCA2_FI_GFPve_pos  1.5413668327  1.7341842972
## dCA2_Fpos_AACAAGACAGAGCTAG-1dCA2_FI_GFPve_pos -0.7778423853  2.0522661529
## dCA2_Fpos_AACCATGCAACTCATG-1dCA2_FI_GFPve_pos -0.0372792758  2.1476350151
## dCA2_Fpos_AACCATGGTCCGATCG-1dCA2_FI_GFPve_pos -1.2214510269  3.0060091339
## dCA2_Fpos_AACCTGACAATCCTTT-1dCA2_FI_GFPve_pos -2.0982085295  0.7528605781
## dCA2_Fpos_AACGAAAAGGGCAGTT-1dCA2_FI_GFPve_pos -1.9459740944  2.4484825455
## dCA2_Fpos_AACTTCTTCCAGCACG-1dCA2_FI_GFPve_pos  0.8274549417  3.1025681816
## dCA2_Fpos_AAGCATCAGGGCCTCT-1dCA2_FI_GFPve_pos -0.9042025633  3.3144941650
## dCA2_Fpos_AAGCATCTCCTATTTG-1dCA2_FI_GFPve_pos -0.0564852260  0.9287677131
## dCA2_Fpos_AAGCGAGAGTCATTGC-1dCA2_FI_GFPve_pos -0.2855511062  2.3639073692
## dCA2_Fpos_AAGCGAGGTAACCCGC-1dCA2_FI_GFPve_pos  1.6002530031  3.1997628532
## dCA2_Fpos_AAGTGAAAGGAGCAAA-1dCA2_FI_GFPve_pos  1.2421395950  3.0175915085
## dCA2_Fpos_AAGTTCGAGACAGTCG-1dCA2_FI_GFPve_pos  0.6815461688  2.2211938225
## dCA2_Fpos_AAGTTCGGTCCTCCAT-1dCA2_FI_GFPve_pos  0.3336216383  2.7058096252
## dCA2_Fpos_AATAGAGGTAACGGTG-1dCA2_FI_GFPve_pos -1.8910319634  3.2428541504
## dCA2_Fpos_AATGAAGAGGACAACC-1dCA2_FI_GFPve_pos  1.4183781318  3.2038798652
## dCA2_Fpos_AATGCCAGTTCATCTT-1dCA2_FI_GFPve_pos -0.0575750299  3.4769735657
## dCA2_Fpos_AATGGAAAGGCAGGGA-1dCA2_FI_GFPve_pos  1.7328328066  2.4179287277
## dCA2_Fpos_ACAAAGAGTTAAGGAT-1dCA2_FI_GFPve_pos  1.4309802227  2.7046194397
```

```
## dCA2_Fpos_ACAACCATCACAATGC-1dCA2_FI_GFPve_pos -0.9076064654  1.4850268684
## dCA2_Fpos_ACACGCGCATGAATAG-1dCA2_FI_GFPve_pos  0.2029673629  3.5507974945
## dCA2_Fpos_ACACTGACATTACGGT-1dCA2_FI_GFPve_pos -1.2384482451  1.5748482071
## dCA2_Fpos_ACCAACAGTTGTGTTG-1dCA2_FI_GFPve_pos -0.6007691450  3.6228132568
## dCA2_Fpos_ACCCTTGAGGAACTCG-1dCA2_FI_GFPve_pos  0.8681588344  3.4263887726
## dCA2_Fpos_ACCTGAAAGCTGAAGC-1dCA2_FI_GFPve_pos -1.7700896092  1.2142014824
## dCA2_Fpos_ACGATGTAGGGAGGTG-1dCA2_FI_GFPve_pos  1.3053416185  2.9649730049
## dCA2_Fpos_ACGATGTAGTAGTGCG-1dCA2_FI_GFPve_pos -0.3016145952  1.8888583504
## dCA2_Fpos_ACGATGTGTCTGGTTA-1dCA2_FI_GFPve_pos  1.4260742836  2.9291272484
## dCA2_Fpos_ACGCACGCAAAGTGTA-1dCA2_FI_GFPve_pos  2.6849413090  2.4148784004
## dCA2_Fpos_ACGGAAGAGTTTCGAC-1dCA2_FI_GFPve_pos  1.3084912472  2.0075121246
## dCA2_Fpos_ACGGGTCGTTGCTCGG-1dCA2_FI_GFPve_pos -1.7501888104  1.9301233612
## dCA2_Fpos_ACGGTCGTCCTCACTG-1dCA2_FI_GFPve_pos -0.6928295918  2.2344317756
## dCA2_Fpos_ACTATCTCAAACAGGC-1dCA2_FI_GFPve_pos -1.2166099377  1.2206592880
## dCA2_Fpos_ACTCTCGAGCGCCATC-1dCA2_FI_GFPve_pos -1.3599937983  0.0248246513
## dCA2_Fpos_ACTGCAAGTGACAGCA-1dCA2_FI_GFPve_pos  1.1708690100  2.8102684341
## dCA2_Fpos_ACTTAGGTCGAAGCCC-1dCA2_FI_GFPve_pos  1.3755799703  2.6020174347
## dCA2_Fpos_AGAAATGCACGCAAAG-1dCA2_FI_GFPve_pos -1.0035603471  1.6769748054
## dCA2_Fpos_AGACAAATCGCGATCG-1dCA2_FI_GFPve_pos -2.2372307606  2.2674737297
## dCA2_Fpos_AGACCATTCTGGGTCG-1dCA2_FI_GFPve_pos  1.3833958082  3.2702589355
## dCA2_Fpos_AGACCCGCAACCACAT-1dCA2_FI_GFPve_pos  1.9600227051  2.8774381004
## dCA2_Fpos_AGACCCGCAATCGTCA-1dCA2_FI_GFPve_pos  1.0152380638  2.4472704254
## dCA2_Fpos_AGACCCGTCGTGCGAC-1dCA2_FI_GFPve_pos -0.6113032229  2.3341131531
## dCA2_Fpos_AGAGAATCATCAGCTA-1dCA2_FI_GFPve_pos -1.6785971947  1.6081877075
## dCA2_Fpos_AGAGAATGTAACGATA-1dCA2_FI_GFPve_pos  1.0133792333  1.8601708732
## dCA2_Fpos_AGAGCCCCAGACGGAT-1dCA2_FI_GFPve_pos  0.2424751394  2.0366802536
## dCA2_Fpos_AGAGCCCGTTAGAGTA-1dCA2_FI_GFPve_pos  1.0652529173  3.1615796409
## dCA2_Fpos_AGATCCAGTCCGGATC-1dCA2_FI_GFPve_pos -1.2347745724  1.9426222168
## dCA2_Fpos_AGATCGTTCCGCAAAT-1dCA2_FI_GFPve_pos -0.4275605209  2.1090703331
## dCA2_Fpos_AGATGCTAGGTTTGAA-1dCA2_FI_GFPve_pos  0.0461621605  2.9685726486
## dCA2_Fpos_AGATGCTCATTCGATG-1dCA2_FI_GFPve_pos  2.1255994015  2.8896389328
## dCA2_Fpos_AGCCAATCAGAATCGG-1dCA2_FI_GFPve_pos  0.5367328696  3.6353245102
## dCA2_Fpos_AGCGTATGTAGAGTTA-1dCA2_FI_GFPve_pos  0.0340885095  2.1307487808
## dCA2_Fpos_AGCTCAAAGCGCATCC-1dCA2_FI_GFPve_pos -2.1846451588  2.9834704719
## dCA2_Fpos_AGCTCAAGTACTGTTG-1dCA2_FI_GFPve_pos -0.6592787631  1.3008222900
## dCA2_Fpos_AGCTCAATCCATTTCA-1dCA2_FI_GFPve_pos  1.3605339699  2.2368188225
## dCA2_Fpos_AGGAATAAGGAATTAC-1dCA2_FI_GFPve_pos  0.4701037578  3.3304882370
## dCA2_Fpos_AGGAGGTCACAAAGCG-1dCA2_FI_GFPve_pos  1.7436665468  3.1250653587
## dCA2_Fpos_AGGGAGTCATCGGCCA-1dCA2_FI_GFPve_pos  1.1562787466  3.5895042740
## dCA2_Fpos_AGGGCTCGTTCGAACT-1dCA2_FI_GFPve_pos  1.0961617403  2.6233849846
## dCA2_Fpos_AGGGTCCGTCATAACC-1dCA2_FI_GFPve_pos  0.2423805468  2.8088913284
## dCA2_Fpos_AGGTTACAGATCACTC-1dCA2_FI_GFPve_pos -0.5984775849  2.1941442810
## dCA2_Fpos_AGGTTACGTCTATGAC-1dCA2_FI_GFPve_pos -1.6743621655  2.2891817413
## dCA2_Fpos_AGGTTGTAGGCGTTGA-1dCA2_FI_GFPve_pos  0.2459314041  1.1545291267
## dCA2_Fpos_AGTAACCTCCGCAACG-1dCA2_FI_GFPve_pos  0.5088964276  2.2890587173
## dCA2_Fpos_AGTGACTAGAACCGCA-1dCA2_FI_GFPve_pos -0.3346107282  2.2832713447
## dCA2_Fpos_AGTTAGCGTCGTAATC-1dCA2_FI_GFPve_pos -0.9471042581  2.4693432174
## dCA2_Fpos_ATACCTTGTCCAGCCA-1dCA2_FI_GFPve_pos -0.3500261433  1.7631154381
## dCA2_Fpos_ATACTTCAGTGGCGAT-1dCA2_FI_GFPve_pos  0.2611166708  2.3563399635
## dCA2_Fpos_ATAGGCTGTACGTGAG-1dCA2_FI_GFPve_pos  1.4901027851  3.3486538254
## dCA2_Fpos_ATCAGGTTCCGTGGTG-1dCA2_FI_GFPve_pos  0.8851691656  2.4395408951
## dCA2_Fpos_ATCATTCAGGTTGAGC-1dCA2_FI_GFPve_pos -1.3654185362  2.2852650009
## dCA2_Fpos_ATCCACCAGCAGAAAG-1dCA2_FI_GFPve_pos  1.6851675682  2.4626555763
## dCA2_Fpos_ATCGATGCAACACGAG-1dCA2_FI_GFPve_pos -1.4634219237  2.0696912132
## dCA2_Fpos_ATCGGATAGCATTTCG-1dCA2_FI_GFPve_pos  1.4473799639  3.1489372574
## dCA2_Fpos_ATCGGATGTCAACCAT-1dCA2_FI_GFPve_pos  1.2669711284  3.2935915314
## dCA2_Fpos_ATGAAAGTCCCGTGTT-1dCA2_FI_GFPve_pos  1.4812643699  3.1218099914
## dCA2_Fpos_ATGAAAGTCTATACTC-1dCA2_FI_GFPve_pos  0.6963357501  3.1079831444
## dCA2_Fpos_ATGAGGGTCAGAGCGA-1dCA2_FI_GFPve_pos  1.1065502338  3.0083885513
## dCA2_Fpos_ATGATCGAGCGAGAAA-1dCA2_FI_GFPve_pos -1.6456416674  2.4729924522
## dCA2_Fpos_ATGCATGCATGTTTGG-1dCA2_FI_GFPve_pos -1.2113016672  3.3002977691
## dCA2_Fpos_ATGGAGGGTGGACCTC-1dCA2_FI_GFPve_pos  0.0420689605  2.7964463554
## dCA2_Fpos_ATGGGTTCAAGTTCGT-1dCA2_FI_GFPve_pos  1.8414519005  2.9517932258
## dCA2_Fpos_ATGGTTGGTAAGATTG-1dCA2_FI_GFPve_pos  1.1001682453  3.1888590179
## dCA2_Fpos_ATTACCTCAGCAGATG-1dCA2_FI_GFPve_pos -0.9207382627  2.2348738037
## dCA2_Fpos_ATTCCATAGGTTGACG-1dCA2_FI_GFPve_pos  0.4815226369  3.1816644989
## dCA2_Fpos_ATTCCATCATAAGCAA-1dCA2_FI_GFPve_pos -0.8699585028  1.7376265846
## dCA2_Fpos_ATTCTACAGGTTCCGC-1dCA2_FI_GFPve_pos  0.4057944588  1.2030186973
## dCA2_Fpos_ATTCTTGCAAGAGTGC-1dCA2_FI_GFPve_pos -0.6928721495  3.4532909714
## dCA2_Fpos_ATTGTTCAGCGTGAGT-1dCA2_FI_GFPve_pos  0.2088935249  3.1681175552
```

```
## dCA2_Fpos_CAAAGAATCCGCACTT-1dCA2_FI_GFPve_pos -0.6464678772  2.5899586998
## dCA2_Fpos_CAAAGAATCGATACAC-1dCA2_FI_GFPve_pos -0.2704630770  2.8731222473
## dCA2_Fpos_CAAGACTTCGGTGAAG-1dCA2_FI_GFPve_pos  0.9012251072  2.2644071899
## dCA2_Fpos_CAAGAGGTCTAGCCAA-1dCA2_FI_GFPve_pos -0.9995758362  1.4711628280
## dCA2_Fpos_CAAGCTAAGGTTGACG-1dCA2_FI_GFPve_pos  2.1381871872  2.5394888244
## dCA2_Fpos_CAAGCTAGTAGTTAGA-1dCA2_FI_GFPve_pos -1.1009797282  3.1350136123
## dCA2_Fpos_CAAGCTATCCATGATG-1dCA2_FI_GFPve_pos  1.7294823103  2.6420669876
## dCA2_Fpos_CAATACGGTACCTAAC-1dCA2_FI_GFPve_pos -1.0757202335  2.0665850960
## dCA2_Fpos_CAATCGATCGCGTAGC-1dCA2_FI_GFPve_pos  1.1620305948  2.8712044082
## dCA2_Fpos_CACAACAGTTCCTAGA-1dCA2_FI_GFPve_pos  0.1734472983  2.2740712486
## dCA2_Fpos_CACAGATAGCGCACAA-1dCA2_FI_GFPve_pos -0.2496411554  2.8404002510
## dCA2_Fpos_CACAGATAGGTTACAA-1dCA2_FI_GFPve_pos  1.7550277881  2.3854976021
## dCA2_Fpos_CACATGACAAGCACCC-1dCA2_FI_GFPve_pos  1.0208449535  2.5261035286
## dCA2_Fpos_CACGTGGTCGAGAAAT-1dCA2_FI_GFPve_pos  1.9671642952  2.9521475158
## dCA2_Fpos_CACGTGGTCGTTCGCT-1dCA2_FI_GFPve_pos  1.1209480934  2.9303245865
## dCA2_Fpos_CACTGGGAGATGGCAC-1dCA2_FI_GFPve_pos  1.5209205322  3.4780207001
## dCA2_Fpos_CACTGTCAGCGTGAGT-1dCA2_FI_GFPve_pos -0.5447160132  3.2351599060
## dCA2_Fpos_CATAAGCCATGTGCCG-1dCA2_FI_GFPve_pos  0.5323057942  2.5040245376
## dCA2_Fpos_CATCGGGTCCTAGCTC-1dCA2_FI_GFPve_pos  0.0680246525  2.8763656936
## dCA2_Fpos_CATCGTCTCGGAGATG-1dCA2_FI_GFPve_pos -1.6005725689  2.2029986702
## dCA2_Fpos_CATGCGGAGGATCACG-1dCA2_FI_GFPve_pos  1.2908734255  2.8248653732
## dCA2_Fpos_CATTCTACATGAAGCG-1dCA2_FI_GFPve_pos -2.2534341641  2.9702682815
## dCA2_Fpos_CATTGTTTCCCGTGAG-1dCA2_FI_GFPve_pos -0.0261133589  2.3943291031
## dCA2_Fpos_CATTTCATCGCTACAA-1dCA2_FI_GFPve_pos -0.2091711514  3.0470853172
## dCA2_Fpos_CCAATGAGTGTCCATA-1dCA2_FI_GFPve_pos  0.5570334368  2.5426278435
## dCA2_Fpos_CCCATTGGTGTGCTTA-1dCA2_FI_GFPve_pos  0.2172049157  3.0143266044
## dCA2_Fpos_CCCTAACTCACCCTCA-1dCA2_FI_GFPve_pos -2.1818990536  3.0891890846
## dCA2_Fpos_CCCTTAGAGACTTCCA-1dCA2_FI_GFPve_pos  2.4069767170  2.2736091934
## dCA2_Fpos_CCCTTAGTCTGGCTGG-1dCA2_FI_GFPve_pos  0.3233146541  2.9739909492
## dCA2_Fpos_CCGATCTCACTGGCGT-1dCA2_FI_GFPve_pos -1.5002075262  1.9180126510
## dCA2_Fpos_CCGGTAGAGGCATCAG-1dCA2_FI_GFPve_pos  0.9230702095  1.9493203483
## dCA2_Fpos_CCGGTAGAGTGATAGT-1dCA2_FI_GFPve_pos  0.4975894861  2.8161125503
## dCA2_Fpos_CCGGTAGGTTGGAGAC-1dCA2_FI_GFPve_pos -0.5257600970  1.8027687393
## dCA2_Fpos_CCGGTGAGTCCAATCA-1dCA2_FI_GFPve_pos  1.2954014711  1.9899683319
## dCA2_Fpos_CCTAACCCATGACAAA-1dCA2_FI_GFPve_pos  0.3169418447  1.4280696235
## dCA2_Fpos_CCTACGTAGTCACGAG-1dCA2_FI_GFPve_pos -1.2637752123  1.4593735061
## dCA2_Fpos_CCTCCAATCCGTAATG-1dCA2_FI_GFPve_pos  1.0205990248  1.8016181312
## dCA2_Fpos_CCTCCTCAGGTGCTAG-1dCA2_FI_GFPve_pos  0.6102395229  2.3583026253
## dCA2_Fpos_CCTTGTGTCCTCAGAA-1dCA2_FI_GFPve_pos -0.2614425547  2.8347106300
## dCA2_Fpos_CCTTTGGCAGCTAACT-1dCA2_FI_GFPve_pos  1.4297577314  1.8129449211
## dCA2_Fpos_CCTTTGGTCGGTTCAA-1dCA2_FI_GFPve_pos -1.9091493912  2.6146998726
## dCA2_Fpos_CGACAGCTCAGAGCAG-1dCA2_FI_GFPve_pos  0.9367226534  3.5322399460
## dCA2_Fpos_CGATCGGAGACCAGAC-1dCA2_FI_GFPve_pos  0.7934151463  1.7952232681
## dCA2_Fpos_CGCATGGGTAGGTGCA-1dCA2_FI_GFPve_pos  1.4319789581  3.2902555786
## dCA2_Fpos_CGCCAGATCGGTCGAC-1dCA2_FI_GFPve_pos -1.5333536692  1.2616839729
## dCA2_Fpos_CGCCATTAGCTGTCCG-1dCA2_FI_GFPve_pos  0.6265424065  3.0058961235
## dCA2_Fpos_CGCGTGAGTTTCTATC-1dCA2_FI_GFPve_pos -1.4011835880  1.8418164573
## dCA2_Fpos_CGGAACCGTTACCTTT-1dCA2_FI_GFPve_pos -0.4714668222  2.3461580597
## dCA2_Fpos_CGGAACCTCCCTCTAG-1dCA2_FI_GFPve_pos  1.3329744510  2.8233766876
## dCA2_Fpos_CGGAGAATCGAGATAA-1dCA2_FI_GFPve_pos -1.7447477408  3.3327112518
## dCA2_Fpos_CGGCAGTAGGAGTACC-1dCA2_FI_GFPve_pos  0.0372789823  3.1662621818
## dCA2_Fpos_CGGGTGTCACTACGGC-1dCA2_FI_GFPve_pos  0.5013055377  2.1899462066
## dCA2_Fpos_CGTCCATTCAGTCTTT-1dCA2_FI_GFPve_pos -0.0365610845  3.4229498230
## dCA2_Fpos_CGTGAATTCTCCACTG-1dCA2_FI_GFPve_pos -1.1881427593  3.2421326958
## dCA2_Fpos_CGTGATAAGCTGCCTG-1dCA2_FI_GFPve_pos  0.6109945349  2.7015405021
## dCA2_Fpos_CGTGCTTTCCGGTAAT-1dCA2_FI_GFPve_pos -1.4840357132  1.7332368217
## dCA2_Fpos_CGTTCTGTCGAAGAAT-1dCA2_FI_GFPve_pos  1.2519817524  2.7770295463
## dCA2_Fpos_CTAAGTGCAAGAGCTG-1dCA2_FI_GFPve_pos -0.1725041085  1.4858303390
## dCA2_Fpos_CTACAGATCACTGATG-1dCA2_FI_GFPve_pos  1.1631248169  2.3884244285
## dCA2_Fpos_CTACATTCACAAAGCG-1dCA2_FI_GFPve_pos  1.8666825466  1.8469171844
## dCA2_Fpos_CTACTATCACGTCGGT-1dCA2_FI_GFPve_pos  1.3847698145  3.2941131912
## dCA2_Fpos_CTATAGGAGTGACCTT-1dCA2_FI_GFPve_pos  0.8345961742  2.1882577263
## dCA2_Fpos_CTCAAGACAGAGTGAC-1dCA2_FI_GFPve_pos  1.3434060983  2.1840877853
## dCA2_Fpos_CTCACTGTCAGGAAGC-1dCA2_FI_GFPve_pos -1.4976775475  2.3856563888
## dCA2_Fpos_CTCAGGGTCCCAGCGA-1dCA2_FI_GFPve_pos  0.9818535023  1.8025293670
## dCA2_Fpos_CTCATCGGTAATGCTC-1dCA2_FI_GFPve_pos  0.8070956640  2.8952703796
## dCA2_Fpos_CTCCAACTCACAACCA-1dCA2_FI_GFPve_pos  0.9872696571  1.0340175949
## dCA2_Fpos_CTCCATGAGGCAGCTA-1dCA2_FI_GFPve_pos -1.6333142348  1.5009141289
## dCA2_Fpos_CTCCCAATCGAAGGAC-1dCA2_FI_GFPve_pos  0.9756817989  2.4955830894
## dCA2_Fpos_CTCCCTCCACTGAATC-1dCA2_FI_GFPve_pos  0.0439662062  1.9083791099
```

```
## dCA2_Fpos_CTCTGGTAGCCGATAG-1dCA2_FI_GFPve_pos  1.4374059610  2.6574201904
## dCA2_Fpos_CTGAGGCTCTAGGCCG-1dCA2_FI_GFPve_pos  1.5024852924  2.1597919784
## dCA2_Fpos_CTGATCCAGAGCTTTC-1dCA2_FI_GFPve_pos -1.9478413887  0.8323688827
## dCA2_Fpos_CTGATCCCAGACCCGT-1dCA2_FI_GFPve_pos -0.8182108827  1.7197480522
## dCA2_Fpos_CTGCAGGGTAGCGCCT-1dCA2_FI_GFPve_pos -1.9006763525  3.2019086204
## dCA2_Fpos_CTGCATCAGCGGGTTA-1dCA2_FI_GFPve_pos  2.2180843525  2.7838883720
## dCA2_Fpos_CTGCATCAGGCTCAAG-1dCA2_FI_GFPve_pos  0.8263412885  2.6387834869
## dCA2_Fpos_CTGCATCCAAGTGACG-1dCA2_FI_GFPve_pos  1.1470145159  1.7967820488
## dCA2_Fpos_CTGCATCGTACGTGTT-1dCA2_FI_GFPve_pos -0.3540188201  1.5298438392
## dCA2_Fpos_CTGCATCGTGACTCTA-1dCA2_FI_GFPve_pos -0.8320930429  3.0034494720
## dCA2_Fpos_CTGCCTACAAGGTTGG-1dCA2_FI_GFPve_pos -0.6590277679  1.9385090195
## dCA2_Fpos_CTGCGAGCAGCTGCCA-1dCA2_FI_GFPve_pos  1.0437075071  2.5855889641
## dCA2_Fpos_CTGCTCAGTTAGGGAC-1dCA2_FI_GFPve_pos -0.1321372218  1.6224203430
## dCA2_Fpos_CTGGTCTCACGGTAGA-1dCA2_FI_GFPve_pos  0.1883552842  1.2071323715
## dCA2_Fpos_CTGGTCTTCTCGTCGT-1dCA2_FI_GFPve_pos -1.6250042744  2.7006383262
## dCA2_Fpos_CTGTAGATCTCAGTCC-1dCA2_FI_GFPve_pos  1.5759359770  1.9175062500
## dCA2_Fpos_CTGTCGTAGGATGAGA-1dCA2_FI_GFPve_pos -0.4436044522  2.4481945358
## dCA2_Fpos_CTGTCGTAGTAAACGT-1dCA2_FI_GFPve_pos  0.2554361098  2.8752150856
## dCA2_Fpos_CTGTCGTCATAAGATG-1dCA2_FI_GFPve_pos  0.6998495035  2.5314107261
## dCA2_Fpos_CTTAGGATCTACCCAC-1dCA2_FI_GFPve_pos -0.4879954286  2.3381467186
## dCA2_Fpos_CTTCCTTCATCATTGG-1dCA2_FI_GFPve_pos  0.5758278541  1.9360986076
## dCA2_Fpos_CTTCTCTTCGATGGAG-1dCA2_FI_GFPve_pos -1.5760497875  0.9868188225
## dCA2_Fpos_CTTGATTTCATGCATG-1dCA2_FI_GFPve_pos -1.5609164067  1.8638801895
## dCA2_Fpos_GAACACTGTATGAGGC-1dCA2_FI_GFPve_pos -1.2484215327  2.2687449775
## dCA2_Fpos_GAACTGTAGAGAAGGT-1dCA2_FI_GFPve_pos  0.3090959363  2.6272631012
## dCA2_Fpos_GAATAGAGTACTGAGG-1dCA2_FI_GFPve_pos  1.3323878221  2.6494279228
## dCA2_Fpos_GAATCACCAAGCGAGT-1dCA2_FI_GFPve_pos  0.0460665844  2.8198404632
## dCA2_Fpos_GACACGCAGATAGTCA-1dCA2_FI_GFPve_pos  0.6978057198  1.9537878357
## dCA2_Fpos_GACCCAGGTTCGTTCC-1dCA2_FI_GFPve_pos  0.5771578722  2.3541221939
## dCA2_Fpos_GACCGTGAGCGCCGTT-1dCA2_FI_GFPve_pos  1.3994307689  2.0062227569
## dCA2_Fpos_GACCGTGTCCAAGAGG-1dCA2_FI_GFPve_pos  0.7345423274  2.5132255874
## dCA2_Fpos_GACGTTACATCGGAAG-1dCA2_FI_GFPve_pos  0.5305705838  3.1740413032
## dCA2_Fpos_GAGACCCGTAGAGTTA-1dCA2_FI_GFPve_pos  0.2440625303  2.4514108024
## dCA2_Fpos_GAGACTTGTTGTGCCG-1dCA2_FI_GFPve_pos -0.0706241481  2.3336482368
## dCA2_Fpos_GAGGCAAAGATCGCTT-1dCA2_FI_GFPve_pos -1.6740987129  3.1828251205
## dCA2_Fpos_GAGGCAACAGGAGGTT-1dCA2_FI_GFPve_pos  0.9871196918  2.9239335380
## dCA2_Fpos_GAGGCCTGTTCCAGGC-1dCA2_FI_GFPve_pos  1.1590465479  1.9836182915
## dCA2_Fpos_GAGGGTACAGTTAAAG-1dCA2_FI_GFPve_pos -1.1535689659  1.4701772056
## dCA2_Fpos_GAGGGTATCCCGATCT-1dCA2_FI_GFPve_pos  1.3121823244  3.4425888382
## dCA2_Fpos_GAGTGTTTCCTTCTTC-1dCA2_FI_GFPve_pos -0.3013835676  3.4523973785
## dCA2_Fpos_GATCAGTTCATTGCGA-1dCA2_FI_GFPve_pos  0.0736608379  3.6866560303
## dCA2_Fpos_GATCATGGTGGTTTAC-1dCA2_FI_GFPve_pos  0.3867204361  3.1764188133
## dCA2_Fpos_GATGTTGGTGCATGTT-1dCA2_FI_GFPve_pos  0.2130800657  2.6047692619
## dCA2_Fpos_GATTCTTGTTCGGGTC-1dCA2_FI_GFPve_pos  0.1579295866  3.0898256622
## dCA2_Fpos_GCAACATAGGAAGTGA-1dCA2_FI_GFPve_pos -1.5630569287  2.9536538444
## dCA2_Fpos_GCAACCGAGTCAGAGC-1dCA2_FI_GFPve_pos  0.2567331366  1.5439844452
## dCA2_Fpos_GCACGGTGTAATCAGA-1dCA2_FI_GFPve_pos  1.7091443710  2.5910220466
## dCA2_Fpos_GCACGTGTCTTGCGCT-1dCA2_FI_GFPve_pos  0.3737088613  3.3814611755
## dCA2_Fpos_GCAGCCACACTTGGCG-1dCA2_FI_GFPve_pos  0.3863046341  3.1091213547
## dCA2_Fpos_GCAGGCTGTCTGTCAA-1dCA2_FI_GFPve_pos  0.8985827141  3.0332813583
## dCA2_Fpos_GCATCGGAGGGTTTCT-1dCA2_FI_GFPve_pos  0.2083785407  1.7254691444
## dCA2_Fpos_GCCAGCAGTCTCACGG-1dCA2_FI_GFPve_pos -0.7813521452  2.0025539718
## dCA2_Fpos_GCCAGGTGTTCTCTAT-1dCA2_FI_GFPve_pos -0.7027062006  3.5102949463
## dCA2_Fpos_GCCAGGTTCAACACCA-1dCA2_FI_GFPve_pos  0.1503759496  2.1869073234
## dCA2_Fpos_GCCAGTGAGCAAATGT-1dCA2_FI_GFPve_pos  0.5998960905  1.0452394806
## dCA2_Fpos_GCCATGGTCAGCTCTC-1dCA2_FI_GFPve_pos  0.4301469497  2.1031799637
## dCA2_Fpos_GCGAGAAGTATAGGGC-1dCA2_FI_GFPve_pos -0.9816551633  2.1366930328
## dCA2_Fpos_GCGTGCAAGACTCATC-1dCA2_FI_GFPve_pos -1.8863603897  2.4818229996
## dCA2_Fpos_GCGTGCACAGAATGTA-1dCA2_FI_GFPve_pos  0.8046178989  2.9262676559
## dCA2_Fpos_GCGTGCAGTTGAGAGC-1dCA2_FI_GFPve_pos -0.6649391480  2.9605207763
## dCA2_Fpos_GCTCAAAAGTCCTGCG-1dCA2_FI_GFPve_pos -0.7101378865  1.5623975120
## dCA2_Fpos_GCTCAAATCGTAATGC-1dCA2_FI_GFPve_pos  0.8167734318  3.4955311142
## dCA2_Fpos_GCTGCAGAGTCCGCGT-1dCA2_FI_GFPve_pos -0.4011852570  2.8552255951
## dCA2_Fpos_GGAAGTGGTGTGTCGC-1dCA2_FI_GFPve_pos -2.2027413674  3.0587687813
## dCA2_Fpos_GGAATGGCACATTCTT-1dCA2_FI_GFPve_pos -0.5245680340  1.3042216621
## dCA2_Fpos_GGAGCAATCACTGTTT-1dCA2_FI_GFPve_pos -0.8765077419  2.9083309494
## dCA2_Fpos_GGATCTACAATCAAGA-1dCA2_FI_GFPve_pos -1.2414463587  1.8672771774
## dCA2_Fpos_GGATGTTAGACCTCAT-1dCA2_FI_GFPve_pos  0.1086260252  2.7763920151
## dCA2_Fpos_GGCGTCATCTCGTGGG-1dCA2_FI_GFPve_pos -0.2631073795  1.8114824615
## dCA2_Fpos_GGCTTGGGTCTCACGG-1dCA2_FI_GFPve_pos -1.5256839819  3.2845049225
```

```
## dCA2_Fpos_GGCTTTCAGGTTAGTA-1dCA2_FI_GFPve_pos  0.1380657725  2.4038744293
## dCA2_Fpos_GGGACAAAGTACTGGG-1dCA2_FI_GFPve_pos -0.3647446461  3.1341467224
## dCA2_Fpos_GGGACCTAGCGACAGT-1dCA2_FI_GFPve_pos -1.7051863499  2.4664392792
## dCA2_Fpos_GGGAGATCACTTGAAC-1dCA2_FI_GFPve_pos  0.9179226332  2.9730334602
## dCA2_Fpos_GGGATGACAACTGAAA-1dCA2_FI_GFPve_pos -1.7532015867  1.9923997246
## dCA2_Fpos_GGGCTACAGAAAGTCT-1dCA2_FI_GFPve_pos -1.3868167229  1.6169901214
## dCA2_Fpos_GGGCTACGTCAGTTTG-1dCA2_FI_GFPve_pos -2.2164218254  2.9377727829
## dCA2_Fpos_GGGCTCAGTAGCGTTT-1dCA2_FI_GFPve_pos  0.5101084285  2.1942277275
## dCA2_Fpos_GGGTGTCCAAGTGCTT-1dCA2_FI_GFPve_pos  0.9997059278  3.4180088363
## dCA2_Fpos_GGGTTATAGTTGCATC-1dCA2_FI_GFPve_pos  0.9471545391  2.4089784942
## dCA2_Fpos_GGGTTTACAATCAAGA-1dCA2_FI_GFPve_pos -1.6642018385  1.9922833763
## dCA2_Fpos_GGTAACTAGCTACGTT-1dCA2_FI_GFPve_pos -0.8971490092  1.5289802872
## dCA2_Fpos_GGTGTCGGTGCCTTCT-1dCA2_FI_GFPve_pos  1.5723999910  2.2569718681
## dCA2_Fpos_GTAACACAGTGGCAGT-1dCA2_FI_GFPve_pos -0.2257001489  0.7144790016
## dCA2_Fpos_GTAATGCCATGCCGCA-1dCA2_FI_GFPve_pos -0.9304713435  2.3097224556
## dCA2_Fpos_GTACAGTGTTCGGCGT-1dCA2_FI_GFPve_pos -0.4698055215  1.9059658371
## dCA2_Fpos_GTAGAGGGTCAAGCGA-1dCA2_FI_GFPve_pos  1.2838651113  2.6204505287
## dCA2_Fpos_GTAGATCCAGTCTTCC-1dCA2_FI_GFPve_pos  0.8007514052  1.1638083778
## dCA2_Fpos_GTAGGTTGTGTGTGTT-1dCA2_FI_GFPve_pos  0.2038026624  2.6053448044
## dCA2_Fpos_GTAGTACAGATACCAA-1dCA2_FI_GFPve_pos  1.4254201822  3.0344829880
## dCA2_Fpos_GTATTGGAGTTATGGA-1dCA2_FI_GFPve_pos  1.6103917532  2.3240795456
## dCA2_Fpos_GTATTGGTCGAACGGA-1dCA2_FI_GFPve_pos -1.4082019158  1.9700856529
## dCA2_Fpos_GTATTTCTCCGTGCGA-1dCA2_FI_GFPve_pos -0.5802343018  2.0697059952
## dCA2_Fpos_GTCAGCGGTATGGAAT-1dCA2_FI_GFPve_pos  1.1006691627  1.4554996811
## dCA2_Fpos_GTCGTAAGTATGGAAT-1dCA2_FI_GFPve_pos -0.2785906918  1.6840687118
## dCA2_Fpos_GTCTACCAGGGAGTTC-1dCA2_FI_GFPve_pos -0.7240986057  1.7931185089
## dCA2_Fpos_GTCTTTATCGCTACGG-1dCA2_FI_GFPve_pos -1.1533037849  1.9325843178
## dCA2_Fpos_GTGCGTGTCGCGTCGA-1dCA2_FI_GFPve_pos  0.1830130093  2.8835759483
## dCA2_Fpos_GTGCTTCGTGAGTCAG-1dCA2_FI_GFPve_pos -2.5648493595 -0.8265196957
## dCA2_Fpos_GTGGAAGCAACTAGAA-1dCA2_FI_GFPve_pos  1.1048504286  3.4850454651
## dCA2_Fpos_GTGGAAGGTCGTGGTC-1dCA2_FI_GFPve_pos -1.7492958136  1.8213062607
## dCA2_Fpos_GTGTGATGTCTCAGGC-1dCA2_FI_GFPve_pos  0.4792386465  2.0413275085
## dCA2_Fpos_GTGTGGCAGCGGTAGT-1dCA2_FI_GFPve_pos -1.7912932463  3.0581174217
## dCA2_Fpos_GTGTTAGTCTTCACGC-1dCA2_FI_GFPve_pos -0.7394778557  2.8803868614
## dCA2_Fpos_GTGTTCCGTTCTCAGA-1dCA2_FI_GFPve_pos -0.0442221068  2.9308548294
## dCA2_Fpos_GTTAGTGGTCGCTCGA-1dCA2_FI_GFPve_pos  0.3628487758  2.9982772194
## dCA2_Fpos_GTTCATTTCTTCGACC-1dCA2_FI_GFPve_pos  0.1381418280  2.6302404724
## dCA2_Fpos_GTTGAACAGTCGAAGC-1dCA2_FI_GFPve_pos  1.5328016453  2.7892713867
## dCA2_Fpos_GTTGCGGCAAAGCACG-1dCA2_FI_GFPve_pos  0.1405280106  2.2559242569
## dCA2_Fpos_GTTGCGGCAGCTATTG-1dCA2_FI_GFPve_pos -0.0054427661  2.1840257965
## dCA2_Fpos_GTTGTAGAGGAAAGGT-1dCA2_FI_GFPve_pos  1.3356321029  2.6679087005
## dCA2_Fpos_TAACCAGCAGTTAGGG-1dCA2_FI_GFPve_pos  1.6467615537  2.7795839630
## dCA2_Fpos_TAAGCGTCAATCGCAT-1dCA2_FI_GFPve_pos -1.2342194147  1.9748254143
## dCA2_Fpos_TAATTCCGTCAAAGCG-1dCA2_FI_GFPve_pos  0.2810287647  1.7823276840
## dCA2_Fpos_TACACCCTCACGGTCG-1dCA2_FI_GFPve_pos  0.9701634817  3.5728092514
## dCA2_Fpos_TACACCCTCTGAGGCC-1dCA2_FI_GFPve_pos  0.4352951221  2.5954766594
## dCA2_Fpos_TACACCCTCTGGTGGC-1dCA2_FI_GFPve_pos -2.3110589810  1.1023288093
## dCA2_Fpos_TACAGGTGTACAATAG-1dCA2_FI_GFPve_pos -1.4574916191  2.4519992195
## dCA2_Fpos_TACCCACCAACTCCAA-1dCA2_FI_GFPve_pos -1.7929938860  1.7415819488
## dCA2_Fpos_TACCTCGAGGTTCTTG-1dCA2_FI_GFPve_pos  0.9737880401  1.8580127083
## dCA2_Fpos_TACCTCGCATCCTGTC-1dCA2_FI_GFPve_pos -0.9849479027  3.2196851097
## dCA2_Fpos_TACGCTCGTAGGTTTC-1dCA2_FI_GFPve_pos  0.2086288802  3.3863830887
## dCA2_Fpos_TACGGTACACCTCAGG-1dCA2_FI_GFPve_pos -1.2609122820  1.7265982948
## dCA2_Fpos_TACTTGTTCGTCTACC-1dCA2_FI_GFPve_pos  1.0736221247  2.2081032120
## dCA2_Fpos_TAGACCAGTCTTTATC-1dCA2_FI_GFPve_pos -2.3303992577  2.4517708145
## dCA2_Fpos_TAGATCGAGTTAGAAC-1dCA2_FI_GFPve_pos  0.8079219036  2.1610737167
## dCA2_Fpos_TAGATCGTCAGGGATG-1dCA2_FI_GFPve_pos -1.5256869621  1.5891185127
## dCA2_Fpos_TAGATCGTCCCTCTCC-1dCA2_FI_GFPve_pos  1.4389788084  2.8979788147
## dCA2_Fpos_TAGCACAAGAAGCTCG-1dCA2_FI_GFPve_pos  0.4640984707  3.3538055740
## dCA2_Fpos_TAGGTTGCATCTAACG-1dCA2_FI_GFPve_pos -1.1927947827  3.0434995018
## dCA2_Fpos_TATCCTAGTAGACGGT-1dCA2_FI_GFPve_pos  1.8121946029  2.1156421028
## dCA2_Fpos_TATCTGTTCCATCAGA-1dCA2_FI_GFPve_pos  1.4284765892  1.5490680061
## dCA2_Fpos_TATTTCGGTTTCGGCG-1dCA2_FI_GFPve_pos  0.5634191804  3.0562286697
## dCA2_Fpos_TCACACCAGAGCCGTA-1dCA2_FI_GFPve_pos  0.0358278476  3.2672767959
## dCA2_Fpos_TCACACCAGGTTGGAC-1dCA2_FI_GFPve_pos -0.6443613596  3.0290975891
## dCA2_Fpos_TCACGCTTCATCAGTG-1dCA2_FI_GFPve_pos  1.4566747122  2.0965366684
## dCA2_Fpos_TCAGGGCAGTAAATGC-1dCA2_FI_GFPve_pos -0.0214489080  1.7872181259
## dCA2_Fpos_TCAGGGCGTCCTGTTC-1dCA2_FI_GFPve_pos -0.0740032472  3.4629469238
## dCA2_Fpos_TCAGTTTGTGAGCAGT-1dCA2_FI_GFPve_pos  0.4607640676  2.3196344696
## dCA2_Fpos_TCATGCCAGTGTTCAC-1dCA2_FI_GFPve_pos -0.0388091035  1.5013656936
```

```
## dCA2_Fpos_TCATTACCAGGAAGTC-1dCA2_FI_GFPve_pos  1.7508711986  3.0638499580
## dCA2_Fpos_TCATTCAGTAGCGTCC-1dCA2_FI_GFPve_pos -2.0626970358  1.3275404297
## dCA2_Fpos_TCATTGTCAGCACAGA-1dCA2_FI_GFPve_pos -1.0293221302  3.3035908066
## dCA2_Fpos_TCATTTGCAGACCAGA-1dCA2_FI_GFPve_pos  0.0958495014  1.7971816383
## dCA2_Fpos_TCCATCGGTGGGACAT-1dCA2_FI_GFPve_pos  1.2253186874  3.3222093902
## dCA2_Fpos_TCCATCGTCTCGGTAA-1dCA2_FI_GFPve_pos  0.1802179687  2.6468348823
## dCA2_Fpos_TCCCACAAGACAACAT-1dCA2_FI_GFPve_pos  1.2330746822  3.0902800880
## dCA2_Fpos_TCCGAAAGTAACCAGG-1dCA2_FI_GFPve_pos -0.4025921650  3.2255730949
## dCA2_Fpos_TCCTTCTAGTATGATG-1dCA2_FI_GFPve_pos -1.8751553126  3.0533361755
## dCA2_Fpos_TCCTTCTCATCTCCCA-1dCA2_FI_GFPve_pos  1.0050078802  1.7626562439
## dCA2_Fpos_TCGAACAAGCGCTGCT-1dCA2_FI_GFPve_pos -0.4383737571  1.7464886032
## dCA2_Fpos_TCGACGGGTACATACC-1dCA2_FI_GFPve_pos -1.0603902168  1.5289478622
## dCA2_Fpos_TCGATTTAGTTACGAA-1dCA2_FI_GFPve_pos -0.7003421016  1.5290627800
## dCA2_Fpos_TCGCTCAAGCGGTATG-1dCA2_FI_GFPve_pos -1.0135294624  1.5094452225
## dCA2_Fpos_TCGCTTGGTACGACAG-1dCA2_FI_GFPve_pos  0.8678296976  2.3400984131
## dCA2_Fpos_TCGGATATCGAGAACG-1dCA2_FI_GFPve_pos  1.1933753662  3.3026972137
## dCA2_Fpos_TCGGATATCTAGATCG-1dCA2_FI_GFPve_pos -2.2934601135  2.9503531776
## dCA2_Fpos_TCGGTCTCATAACAGA-1dCA2_FI_GFPve_pos  0.3008020394  2.8927626930
## dCA2_Fpos_TCGTAGAGTTGATGTC-1dCA2_FI_GFPve_pos  0.0517334811  2.2708449684
## dCA2_Fpos_TCGTCCAGTAACGCGA-1dCA2_FI_GFPve_pos  1.1759439878  3.2874870621
## dCA2_Fpos_TCGTGCTCACTCCTGT-1dCA2_FI_GFPve_pos  0.2564446800  2.4363432251
## dCA2_Fpos_TCGTGCTGTATCTTCT-1dCA2_FI_GFPve_pos -0.4504575260  1.6820264183
## dCA2_Fpos_TCTCAGCGTGTCCACG-1dCA2_FI_GFPve_pos  1.3374369316  3.2341256462
## dCA2_Fpos_TCTCTGGAGAGTATAC-1dCA2_FI_GFPve_pos  0.1437266640  1.7747116409
## dCA2_Fpos_TCTGCCAAGGCGAACT-1dCA2_FI_GFPve_pos -1.5642468758  2.6547556244
## dCA2_Fpos_TCTGGCTTCGTGGACC-1dCA2_FI_GFPve_pos  0.6594374947  3.1765184723
## dCA2_Fpos_TCTTCCTTCCGTACGG-1dCA2_FI_GFPve_pos  0.1894148760  1.6523790680
## dCA2_Fpos_TCTTGCGCATCGAAGG-1dCA2_FI_GFPve_pos  1.6720105581  2.1059871040
## dCA2_Fpos_TCTTGCGGTTCCTACC-1dCA2_FI_GFPve_pos  1.0773625545  3.0159111343
## dCA2_Fpos_TGACCCTAGAGTCCGA-1dCA2_FI_GFPve_pos  0.9831320219  1.8896293960
## dCA2_Fpos_TGACGCGTCGGCACTG-1dCA2_FI_GFPve_pos  0.5814209752  2.6651621185
## dCA2_Fpos_TGAGACTGTCGCAGTC-1dCA2_FI_GFPve_pos -0.3234256782  2.0038843475
## dCA2_Fpos_TGAGGAGAGTTGTCAC-1dCA2_FI_GFPve_pos -2.3329062290  2.8778329216
## dCA2_Fpos_TGAGGAGTCCGGTAAT-1dCA2_FI_GFPve_pos  0.6333078913  2.2246256195
## dCA2_Fpos_TGATCAGGTTAAGACA-1dCA2_FI_GFPve_pos -1.1088725753  1.8890943847
## dCA2_Fpos_TGCACGGTCTGCTCTG-1dCA2_FI_GFPve_pos  0.6610108785  2.6372599922
## dCA2_Fpos_TGCGACGAGCTTGTGT-1dCA2_FI_GFPve_pos  1.4197787218  2.5990887008
## dCA2_Fpos_TGCGACGGTGTCCTAA-1dCA2_FI_GFPve_pos  0.5619077258  2.5076494537
## dCA2_Fpos_TGCTCCACATCAGCTA-1dCA2_FI_GFPve_pos -1.4077955313  2.1467171036
## dCA2_Fpos_TGCTCGTGTTCTAAGC-1dCA2_FI_GFPve_pos -0.7267693110  1.0262346588
## dCA2_Fpos_TGCTCGTTCGTCAACA-1dCA2_FI_GFPve_pos -0.9588348098  2.6930985771
## dCA2_Fpos_TGCTGAATCCGAGCTG-1dCA2_FI_GFPve_pos -0.8065689035  2.3981895767
## dCA2_Fpos_TGGAGGAAGATGGCGT-1dCA2_FI_GFPve_pos  1.0828184061  1.9955430351
## dCA2_Fpos_TGGGTTAGTAAGCGGT-1dCA2_FI_GFPve_pos -0.8147499986  1.9458709083
## dCA2_Fpos_TGGTACATCAGACCGC-1dCA2_FI_GFPve_pos -0.9337510891  1.6331010185
## dCA2_Fpos_TGGTGATGTAGGAGGG-1dCA2_FI_GFPve_pos  0.9489973955  2.3628831230
## dCA2_Fpos_TGGTTAGTCGGTCAGC-1dCA2_FI_GFPve_pos  1.0879547767  2.8466615997
## dCA2_Fpos_TGTCCCATCAAGCGTT-1dCA2_FI_GFPve_pos -2.1134818621  1.1245365463
## dCA2_Fpos_TGTCCTGTCGCTAGCG-1dCA2_FI_GFPve_pos -1.3206851311  2.9655862175
## dCA2_Fpos_TGTGGCGGTAGACGTG-1dCA2_FI_GFPve_pos -0.1418094072  2.8002348266
## dCA2_Fpos_TGTTCATTCGACCACG-1dCA2_FI_GFPve_pos  1.1835103206  2.3572631202
## dCA2_Fpos_TGTTTGTCAAACTCTG-1dCA2_FI_GFPve_pos  2.4039333038  2.4665580116
## dCA2_Fpos_TTAATCCCAAGTACCT-1dCA2_FI_GFPve_pos  0.1067990832  3.0881820045
## dCA2_Fpos_TTACAGGGTCCAAATC-1dCA2_FI_GFPve_pos  0.1154396109  2.2413235031
## dCA2_Fpos_TTACAGGGTTAAAGTG-1dCA2_FI_GFPve_pos -1.8385011978  2.8342304550
## dCA2_Fpos_TTAGGCAAGCACTCCG-1dCA2_FI_GFPve_pos  0.4781637363  2.3300872169
## dCA2_Fpos_TTAGGCAGTATGGAAT-1dCA2_FI_GFPve_pos  1.0840813093  3.1716733299
## dCA2_Fpos_TTAGGGTCAGAGTAAT-1dCA2_FI_GFPve_pos -0.3043600298 -0.0668058075
## dCA2_Fpos_TTCAATCCAAACCACT-1dCA2_FI_GFPve_pos  0.0526024573  3.1209321342
## dCA2_Fpos_TTCAGGACAAATGGAT-1dCA2_FI_GFPve_pos -0.9592299886  2.8487906776
## dCA2_Fpos_TTCCGTGAGCTGTGCC-1dCA2_FI_GFPve_pos  0.6263136439  3.5019169174
## dCA2_Fpos_TTCCTCTAGGTAGTAT-1dCA2_FI_GFPve_pos  0.8585486584  1.1574755035
## dCA2_Fpos_TTCTAACCAGGCAATG-1dCA2_FI_GFPve_pos  1.2295651607  1.3311019264
## dCA2_Fpos_TTCTAGTAGGCTCTAT-1dCA2_FI_GFPve_pos -0.0012862213  2.8217439972
## dCA2_Fpos_TTCTGTAAGTGATAAC-1dCA2_FI_GFPve_pos  0.3500471883  3.3558140121
## dCA2_Fpos_TTCTGTAGTCTCTCCA-1dCA2_FI_GFPve_pos -0.4590492077  1.9803376518
## dCA2_Fpos_TTGGATGAGACTTCAC-1dCA2_FI_GFPve_pos  0.9990333490  0.6750655494
## dCA2_Fpos_TTGGGTACACTCCGGA-1dCA2_FI_GFPve_pos -1.3896990843  2.5417147003
## dCA2_Fpos_TTGGGTATCAGAGTGG-1dCA2_FI_GFPve_pos -0.0541043110  2.8090100609
## dCA2_Fpos_TTGGTTTAGGCTCTAT-1dCA2_FI_GFPve_pos  0.0428074174  1.5565710388
```

```
## dCA2_Fpos_TTGTGGACATGCACTA-1dCA2_FI_GFPve_pos  0.3167801075  3.3464689575
## dCA2_Fpos_TTGTTGTCAGTCCGTG-1dCA2_FI_GFPve_pos  1.6656129532  1.3826580368
## dCA2_Fpos_TTTACGTAGCTCGGCT-1dCA2_FI_GFPve_pos  0.7645385437  2.0572367035
## dCA2_Fpos_TTTACTGAGTCGAAGC-1dCA2_FI_GFPve_pos  1.2648587398  3.1948943458
## dCA2_Fpos_TTTCATGCACAAATGA-1dCA2_FI_GFPve_pos  1.0181507044  2.3579974495
## dCA2_Fpos_TTTCATGGTATCCCTC-1dCA2_FI_GFPve_pos  1.1801262073  1.7605486236
## dCA2_Fpos_TTTGACTCATCGTGCG-1dCA2_FI_GFPve_pos -0.3329522438  3.5929727875
## dCA2_Fpos_TTTGATCAGTGCCTCG-1dCA2_FI_GFPve_pos  0.0903652184  2.5055833183
## FC_Mpos_AAAGGTATCTCTGCCA-1FC_MI_GFPve_pos      0.2757373922  2.4181642853
## FC_Mpos_AAAGTGAAGCTTTCTT-1FC_MI_GFPve_pos     -0.4980940349  2.0649853073
## FC_Mpos_AAAGTGAAGTTGGCTT-1FC_MI_GFPve_pos      0.5850191288  2.8584881149
## FC_Mpos_AAAGTGAGTCGAATTC-1FC_MI_GFPve_pos     -0.6610841580  2.8837361656
## FC_Mpos_AACAAAGTCTGGTCAA-1FC_MI_GFPve_pos     -1.8003836699  1.8959613167
## FC_Mpos_AACCAACTCGGCCAAC-1FC_MI_GFPve_pos      0.0588362239  1.7387576423
## FC_Mpos_AACCTGATCTCGACCT-1FC_MI_GFPve_pos     -0.5374028809  2.7216191612
## FC_Mpos_AAGTACCAGTAGGAAG-1FC_MI_GFPve_pos     -0.5743304320  3.4765358291
## FC_Mpos_AAGTACCGTCCTACGG-1FC_MI_GFPve_pos      0.4726630621  2.0953097664
## FC_Mpos_AAGTACCTCCACCTGT-1FC_MI_GFPve_pos     -0.6119999714  1.2013454757
## FC_Mpos_AAGTGAAGTTCAAGGG-1FC_MI_GFPve_pos     -1.9397580452  0.9727793060
## FC_Mpos_AATAGAGGTGTGTCGC-1FC_MI_GFPve_pos     -0.7060590811  1.9730234466
## FC_Mpos_AATCGTGAGGCATCAG-1FC_MI_GFPve_pos      1.0368436746  2.2695045791
## FC_Mpos_AATCGTGCACCAAAGG-1FC_MI_GFPve_pos      0.4400241546  0.5351839386
## FC_Mpos_AATGAAGGTACTAAGA-1FC_MI_GFPve_pos     -0.2265223130  3.3017063461
## FC_Mpos_AATGCCACAGAAGCGT-1FC_MI_GFPve_pos     -1.4158544369  2.9543924652
## FC_Mpos_AATGGCTGTAGCACGA-1FC_MI_GFPve_pos     -1.4423519201  2.4164190613
## FC_Mpos_ACAAAGAGTTAGCGGA-1FC_MI_GFPve_pos      0.9195997886  1.7415666900
## FC_Mpos_ACACAGTGTGCACGCT-1FC_MI_GFPve_pos     -1.1139297910  1.7442589126
## FC_Mpos_ACACCAAGTCGATGCC-1FC_MI_GFPve_pos     -0.1385172844  2.7250991188
## FC_Mpos_ACAGAAACAGCTATAC-1FC_MI_GFPve_pos     -1.4532174893  1.6863828026
## FC_Mpos_ACCCTCAGTAGGTACG-1FC_MI_GFPve_pos     -0.9433752127 -0.1124901451
## FC_Mpos_ACCCTTGCACTGTTCC-1FC_MI_GFPve_pos      1.8655080967  1.9937701546
## FC_Mpos_ACCGTTCGTTCCGGTG-1FC_MI_GFPve_pos      0.0443454109  2.8439164482
## FC_Mpos_ACCTGAACAGCGCGTT-1FC_MI_GFPve_pos      0.3897726707  2.6543240867
## FC_Mpos_ACGATCACAGTTCTAG-1FC_MI_GFPve_pos      1.2476924829  3.1102905594
## FC_Mpos_ACGATCACATCGTTCC-1FC_MI_GFPve_pos     -0.5635888405  1.3999252640
## FC_Mpos_ACGGAAGGTATCACCA-1FC_MI_GFPve_pos     -2.2519807644  2.8149419151
## FC_Mpos_ACGGTCGTCGCATGAT-1FC_MI_GFPve_pos      1.1729033165  2.5258612953
## FC_Mpos_ACGTACATCTTCTGTA-1FC_MI_GFPve_pos      0.8743136101  1.2520232521
## FC_Mpos_ACTTAGGTCGTTCCTG-1FC_MI_GFPve_pos      0.5024228864  0.6915784202
## FC_Mpos_ACTTATCGTAGGAGGG-1FC_MI_GFPve_pos     -0.8930664487  3.4948578201
## FC_Mpos_ACTTCCGCAATAGGAT-1FC_MI_GFPve_pos     -1.8496277161  2.6174955688
## FC_Mpos_AGAACAAGTACAAAGT-1FC_MI_GFPve_pos     -1.0043542810  1.9256677948
## FC_Mpos_AGAAGCGAGAGGATGA-1FC_MI_GFPve_pos      2.1331274681  2.7541981064
## FC_Mpos_AGACCCGAGAGCCCAA-1FC_MI_GFPve_pos     -0.4211749263  1.5306930862
## FC_Mpos_AGACTCATCATTATCC-1FC_MI_GFPve_pos     -1.5222729273  2.2843938194
## FC_Mpos_AGAGCAGGTGCTTATG-1FC_MI_GFPve_pos     -1.7844819851  2.9807105385
## FC_Mpos_AGAGCCCTCTTGCAGA-1FC_MI_GFPve_pos     -1.1731768675  3.3123722397
## FC_Mpos_AGATGAAAGCGATTCT-1FC_MI_GFPve_pos     -2.0915559359  2.7486696564
## FC_Mpos_AGATGAAGTTAAGGAT-1FC_MI_GFPve_pos     -0.4629370995  2.6199755989
## FC_Mpos_AGCATCATCACTCGAA-1FC_MI_GFPve_pos      1.3166766338  2.7999954544
## FC_Mpos_AGCGTCGCAGGGTCTC-1FC_MI_GFPve_pos     -0.3471378960  2.3516269050
## FC_Mpos_AGCTTCCGTTAGAAGT-1FC_MI_GFPve_pos     -0.5118142195  2.0346184097
## FC_Mpos_AGGAATATCGATACGT-1FC_MI_GFPve_pos      0.1684835427  2.1110844932
## FC_Mpos_AGGCATTAGGGTAATT-1FC_MI_GFPve_pos      0.9610489778  0.5613799415
## FC_Mpos_AGGCATTCATCAGCGC-1FC_MI_GFPve_pos     -0.4837877877  1.2593140922
## FC_Mpos_AGGCATTGTGGCGTAA-1FC_MI_GFPve_pos     -1.2551247902  2.5468545280
## FC_Mpos_AGGGCCTGTGTCACAT-1FC_MI_GFPve_pos     -1.4820577927  2.4327903114
## FC_Mpos_AGGGTCCGTTTATGCG-1FC_MI_GFPve_pos      0.6267716460  2.0415826164
## FC_Mpos_AGGGTTTCATACAGAA-1FC_MI_GFPve_pos      0.6715620928  0.2489781700
## FC_Mpos_AGGTCTATCCTCTAAT-1FC_MI_GFPve_pos      0.5932718091  2.2575273834
## FC_Mpos_AGGTTGTCAATACCTG-1FC_MI_GFPve_pos      0.2953232102  1.4570641838
## FC_Mpos_AGTACTGCAGCTACTA-1FC_MI_GFPve_pos      0.5200373106  3.2813673340
## FC_Mpos_AGTAGCTAGTTCATCG-1FC_MI_GFPve_pos      1.1832348280  1.9756808601
## FC_Mpos_AGTAGTCAGTGAGTTA-1FC_MI_GFPve_pos     -0.1998436727  2.6820369087
## FC_Mpos_AGTAGTCCACTCTAGA-1FC_MI_GFPve_pos     -0.4279499777  2.8541779838
## FC_Mpos_AGTCATGAGTTAACGA-1FC_MI_GFPve_pos     -0.2149399571  3.0307464920
## FC_Mpos_AGTCTCCGTAGATGTA-1FC_MI_GFPve_pos     -0.4548913963  3.4832506500
## FC_Mpos_AGTGTTGAGTGTGTTC-1FC_MI_GFPve_pos     -0.2820774518  2.8769059501
## FC_Mpos_AGTTCGACAGCTGAGA-1FC_MI_GFPve_pos     -0.2361006044  3.4767179809
## FC_Mpos_ATACCGACACTCAGAT-1FC_MI_GFPve_pos      0.1405710749  2.5755830131
```

```
## FC_Mpos_ATACCTTAGTTCATCG-1FC_MI_GFPve_pos     -1.0452468820  2.1139345489
## FC_Mpos_ATAGACCCATAGATCC-1FC_MI_GFPve_pos      0.1943020098  2.6548948608
## FC_Mpos_ATAGACCTCCATCCGT-1FC_MI_GFPve_pos     -1.1119956203  3.2353868805
## FC_Mpos_ATCCACCTCGGTAGGA-1FC_MI_GFPve_pos     -0.1332788408  2.4590235077
## FC_Mpos_ATCCCTGAGTGCTACT-1FC_MI_GFPve_pos      1.2958222799  1.6232557617
## FC_Mpos_ATCGATGCACCTGCAG-1FC_MI_GFPve_pos     -1.0592774577  1.3610177360
## FC_Mpos_ATCGATGGTGGAGGTT-1FC_MI_GFPve_pos      0.4373770528  2.5711417518
## FC_Mpos_ATCGGATCACATTCGA-1FC_MI_GFPve_pos     -0.7528101630  1.5136952720
## FC_Mpos_ATCGGCGTCTTCCTAA-1FC_MI_GFPve_pos     -1.3224159308  2.5513892494
## FC_Mpos_ATCTCTATCGTGCAGC-1FC_MI_GFPve_pos      0.1580432229  2.3643417679
## FC_Mpos_ATGAAAGGTGTCCCTT-1FC_MI_GFPve_pos     -1.1784883566  2.7987637840
## FC_Mpos_ATGACCATCCACTAGA-1FC_MI_GFPve_pos     -0.5545566507  0.4465542160
## FC_Mpos_ATGAGTCTCGTTCCCA-1FC_MI_GFPve_pos     -1.6651986666  2.4953656517
## FC_Mpos_ATGATCGGTCTACAAC-1FC_MI_GFPve_pos      0.2659535281  2.4400062881
## FC_Mpos_ATGATCGTCGCTTACC-1FC_MI_GFPve_pos     -2.1455083914  2.6276760422
## FC_Mpos_ATGCCTCTCACCCTTG-1FC_MI_GFPve_pos     -2.0404758282  3.1224956833
## FC_Mpos_ATGCGATTCGGCATCG-1FC_MI_GFPve_pos      1.0302220754  2.0540447555
## FC_Mpos_ATGGAGGAGATAGTCA-1FC_MI_GFPve_pos     -0.5482473500  3.1049199425
## FC_Mpos_ATGGGAGTCGACATAC-1FC_MI_GFPve_pos      0.6256335549  2.7896781288
## FC_Mpos_ATGGGTTCACATAGCT-1FC_MI_GFPve_pos     -0.4851060338  1.9259896599
## FC_Mpos_ATGGTTGAGCAGGTCA-1FC_MI_GFPve_pos     -2.0259704418  2.0157623611
## FC_Mpos_ATGGTTGGTAGTACGG-1FC_MI_GFPve_pos      0.4817269020  1.8366098724
## FC_Mpos_ATGGTTGTCTAAGAAG-1FC_MI_GFPve_pos     -0.7622555919  2.2282853447
## FC_Mpos_ATTACCTCACTCCTGT-1FC_MI_GFPve_pos     -0.6824216671  2.5870762192
## FC_Mpos_ATTACTCGTTGTGTTG-1FC_MI_GFPve_pos     -1.6314785309  2.0013385139
## FC_Mpos_ATTCACTGTGCTGTCG-1FC_MI_GFPve_pos     -1.5907372065  2.5694885574
## FC_Mpos_ATTCAGGCAATTCACG-1FC_MI_GFPve_pos     -0.8365617223  3.0027194343
## FC_Mpos_ATTCATCGTGTCTTGA-1FC_MI_GFPve_pos     -0.1345398813  2.4356355987
## FC_Mpos_ATTCCATCAGGTGTGA-1FC_MI_GFPve_pos      0.4277169280  2.9363694511
## FC_Mpos_ATTCTACTCCTCTTTC-1FC_MI_GFPve_pos     -0.7986825533  2.5557070099
## FC_Mpos_ATTCTTGTCGATAACC-1FC_MI_GFPve_pos      0.6444819145  1.1832280479
## FC_Mpos_CAACAGTTCGCAGTGC-1FC_MI_GFPve_pos     -1.0854034252  2.8461141907
## FC_Mpos_CAACCTCGTTGATGTC-1FC_MI_GFPve_pos     -0.2505698629  3.0205188118
## FC_Mpos_CAAGAGGAGTACAACA-1FC_MI_GFPve_pos      1.0844034128  3.3452720962
## FC_Mpos_CAAGAGGCAGGTAGTG-1FC_MI_GFPve_pos     -0.8134065695  2.9438348137
## FC_Mpos_CAAGGGACACAAATGA-1FC_MI_GFPve_pos     -1.5108036823  2.8167839371
## FC_Mpos_CAAGGGATCAATGTCG-1FC_MI_GFPve_pos      1.3047095709  3.2446050964
## FC_Mpos_CAATACGTCTGGCCTT-1FC_MI_GFPve_pos      0.9187240772  2.5652466141
## FC_Mpos_CAATCGACATGCCATA-1FC_MI_GFPve_pos     -2.2433969803  2.6729083381
## FC_Mpos_CAATTTCAGCCGTCGT-1FC_MI_GFPve_pos     -2.6680090256  1.1699629150
## FC_Mpos_CAATTTCGTTGGGCCT-1FC_MI_GFPve_pos      0.9573526554  1.3361850105
## FC_Mpos_CACAGGCCAGCCCAGT-1FC_MI_GFPve_pos      0.8436483316  1.3138466202
## FC_Mpos_CACATGACAAGCAGGT-1FC_MI_GFPve_pos     -0.3282183088  2.3992042862
## FC_Mpos_CACCAAACATGACGTT-1FC_MI_GFPve_pos     -1.4607705898  1.9809055649
## FC_Mpos_CACGTGGTCTGTCGCT-1FC_MI_GFPve_pos      1.1048076324  2.6844387375
## FC_Mpos_CAGCACGTCCACTGGG-1FC_MI_GFPve_pos      0.9897991590  2.1536383949
## FC_Mpos_CAGCAGCGTTGCTTGA-1FC_MI_GFPve_pos     -0.8222658463  1.0939922653
## FC_Mpos_CAGCCAGCAAGAGTTA-1FC_MI_GFPve_pos      1.2887163334  2.0311317764
## FC_Mpos_CAGCGTGAGATAGTGT-1FC_MI_GFPve_pos      1.1723588877  3.2916174255
## FC_Mpos_CAGGTATAGCATGAAT-1FC_MI_GFPve_pos      0.6409467034  3.2848930679
## FC_Mpos_CAGTGCGCATAACTCG-1FC_MI_GFPve_pos     -0.4864781029  3.4024820648
## FC_Mpos_CATCGGGTCACATCAG-1FC_MI_GFPve_pos      1.3360210829  3.0776195846
## FC_Mpos_CATGCGGAGGATGGCT-1FC_MI_GFPve_pos      1.0883754663  3.2476425491
## FC_Mpos_CATTCATGTCATCTAG-1FC_MI_GFPve_pos     -0.6028387971  3.1204533897
## FC_Mpos_CATTGAGCAGTGACCC-1FC_MI_GFPve_pos     -0.8486441798  2.3080511413
## FC_Mpos_CCAATGACAATGAGCG-1FC_MI_GFPve_pos      1.8569247894  2.4069910370
## FC_Mpos_CCAATGAGTTCTTAGG-1FC_MI_GFPve_pos      1.9935052566  2.3634248100
## FC_Mpos_CCATAAGTCCAAAGGG-1FC_MI_GFPve_pos      0.4700312786  2.2088895164
## FC_Mpos_CCCTAACTCAGCGTCG-1FC_MI_GFPve_pos     -1.6759868689  2.2699766479
## FC_Mpos_CCCTCAAAGGTCTACT-1FC_MI_GFPve_pos     -0.5111743636  1.4969044052
## FC_Mpos_CCCTCAACAATCGAAA-1FC_MI_GFPve_pos     -0.4838267989  2.8595547996
## FC_Mpos_CCCTCTCTCTACCAGA-1FC_MI_GFPve_pos     -2.2699949570  2.7310347877
## FC_Mpos_CCGAACGAGGTGAGCT-1FC_MI_GFPve_pos      0.0681075029  2.7316346489
## FC_Mpos_CCGATGGAGTGTACCT-1FC_MI_GFPve_pos     -0.6649664171  3.4892254196
## FC_Mpos_CCGCAAGCATTCCTCG-1FC_MI_GFPve_pos      0.8400056534  2.2252011619
## FC_Mpos_CCGGACAAGCAGCCCT-1FC_MI_GFPve_pos     -0.1041237242  1.8108425461
## FC_Mpos_CCGGACAAGCCGAATG-1FC_MI_GFPve_pos     -0.0113747693  3.1392250381
## FC_Mpos_CCGGTAGCACCTGTCT-1FC_MI_GFPve_pos      0.2952932887  2.6397657715
## FC_Mpos_CCGTAGGGTCGGCTAC-1FC_MI_GFPve_pos     -0.4338799246  1.5778074585
## FC_Mpos_CCGTAGGGTTATCTGG-1FC_MI_GFPve_pos     -1.6522680350  1.9015574775
```

```
## FC_Mpos_CCGTGAGCACTGGATT-1FC_MI_GFPve_pos      0.2713364832  2.6346898399
## FC_Mpos_CCTAACCAGTTGCCCG-1FC_MI_GFPve_pos     -1.3399766512  3.1636228882
## FC_Mpos_CCTACGTAGCAGGTCA-1FC_MI_GFPve_pos      0.3646724396  2.3633399330
## FC_Mpos_CCTCACAGTCAGCGTC-1FC_MI_GFPve_pos     -0.9919681378  2.8151736580
## FC_Mpos_CCTCCAAAGAGCAACC-1FC_MI_GFPve_pos     -0.1568000657  2.6002512298
## FC_Mpos_CCTTGTGGTTGGACTT-1FC_MI_GFPve_pos      0.1604449324  0.6755023323
## FC_Mpos_CGAAGGATCCATATGG-1FC_MI_GFPve_pos      0.1795211308  3.3747568451
## FC_Mpos_CGACAGCCAGTTACCA-1FC_MI_GFPve_pos     -0.7310610838  2.0956430755
## FC_Mpos_CGAGGAACAAGTTTGC-1FC_MI_GFPve_pos     -0.6184187658  2.8950000129
## FC_Mpos_CGATCGGTCGAACACT-1FC_MI_GFPve_pos     -1.6477867193  3.1593528114
## FC_Mpos_CGCATAACAGAGGTTG-1FC_MI_GFPve_pos     -1.1559333034  1.9654703460
## FC_Mpos_CGCCAGAAGCCATTTG-1FC_MI_GFPve_pos      0.4834408335  1.7224426590
## FC_Mpos_CGGAACCAGAGTCAAT-1FC_MI_GFPve_pos     -1.0916802116  3.3175602279
## FC_Mpos_CGGGCATAGACGACGT-1FC_MI_GFPve_pos      0.4411401920  2.5446653686
## FC_Mpos_CGGTCAGCAAACGAGC-1FC_MI_GFPve_pos      0.1572898202  2.8502149902
## FC_Mpos_CGGTCAGTCGGAATGG-1FC_MI_GFPve_pos     -0.1452045958  2.6702008568
## FC_Mpos_CGTAGTAAGCTAAATG-1FC_MI_GFPve_pos     -1.9897392817  2.9873357139
## FC_Mpos_CGTCCATCAGCAGACA-1FC_MI_GFPve_pos     -1.8301734753  2.8669600807
## FC_Mpos_CGTGAATAGCCGGATA-1FC_MI_GFPve_pos      0.6458546095  2.2086329780
## FC_Mpos_CGTGATAGTGTGAATA-1FC_MI_GFPve_pos     -0.5918282993  1.9964685760
## FC_Mpos_CGTTCTGCATCCTCAC-1FC_MI_GFPve_pos      0.3064335458  2.4308348022
## FC_Mpos_CGTTGGGTCCACTGGG-1FC_MI_GFPve_pos     -1.2838631697  2.2911811195
## FC_Mpos_CGTTGGGTCTACCAGA-1FC_MI_GFPve_pos     -1.2370700665  2.1216469131
## FC_Mpos_CTATCCGCAAAGGAGA-1FC_MI_GFPve_pos      1.2070813350  2.4636631332
## FC_Mpos_CTATCCGTCGCACGAC-1FC_MI_GFPve_pos     -0.6034703679  1.0988178573
## FC_Mpos_CTCAACCCATCCTTGC-1FC_MI_GFPve_pos     -0.2496494256  0.9575991951
## FC_Mpos_CTCCATGGTCATCGCG-1FC_MI_GFPve_pos     -0.1957495209  3.0898113571
## FC_Mpos_CTCCCAAAGACGGTTG-1FC_MI_GFPve_pos     -0.0057897843  2.6136808716
## FC_Mpos_CTCGAGGAGCCAGAGT-1FC_MI_GFPve_pos      1.2750516109  2.7047591530
## FC_Mpos_CTCTGGTAGAACCCGA-1FC_MI_GFPve_pos     -1.5240810938  2.2920193992
## FC_Mpos_CTGAGGCAGACTTAAG-1FC_MI_GFPve_pos     -1.0834283657  3.3857250534
## FC_Mpos_CTGAGGCGTCGGATTT-1FC_MI_GFPve_pos      0.7994842105  2.5950637184
## FC_Mpos_CTGCAGGAGCGCTTCG-1FC_MI_GFPve_pos      0.2217439883  1.9626078926
## FC_Mpos_CTGCAGGTCAGCCTCT-1FC_MI_GFPve_pos      1.0085421972  1.8268485390
## FC_Mpos_CTGCATCCAACGACTT-1FC_MI_GFPve_pos     -1.5758477278  2.6049256645
## FC_Mpos_CTGCATCCACGCTGCA-1FC_MI_GFPve_pos     -1.7395087309  2.9705195747
## FC_Mpos_CTGGCAGGTTTGGGAG-1FC_MI_GFPve_pos      0.6053564720  2.2067652069
## FC_Mpos_CTGTAGATCCCGATCT-1FC_MI_GFPve_pos      1.1414896421  2.3948626838
## FC_Mpos_CTGTCGTGTCACGACC-1FC_MI_GFPve_pos     -0.0493156887  2.5134067856
## FC_Mpos_CTGTGAACAGGATGAC-1FC_MI_GFPve_pos     -1.0779664941  2.2718515716
## FC_Mpos_CTTACCGTCAGATGCT-1FC_MI_GFPve_pos     -0.0774314083  2.1801328979
## FC_Mpos_CTTACCGTCTTCGATT-1FC_MI_GFPve_pos     -0.9469686575  1.4254098259
## FC_Mpos_CTTCAATAGATGTTAG-1FC_MI_GFPve_pos      1.4070159368  3.1411424003
## FC_Mpos_CTTCAATTCAGCTTCC-1FC_MI_GFPve_pos     -0.0073127426  2.6899462066
## FC_Mpos_CTTCCGAGTTTGGCTA-1FC_MI_GFPve_pos     -0.6771243043  1.8115868889
## FC_Mpos_CTTCCGATCTGACGCG-1FC_MI_GFPve_pos      0.8364733629  1.4066276871
## FC_Mpos_CTTTCAAGTAGCTCGC-1FC_MI_GFPve_pos      0.8400818281  2.3592086159
## FC_Mpos_GAAATGATCATGGGAG-1FC_MI_GFPve_pos     -0.4818644352  0.3989434562
## FC_Mpos_GAACGTTAGGTAACTA-1FC_MI_GFPve_pos     -1.8955447502  1.2835431419
## FC_Mpos_GAACGTTTCTCGTGGG-1FC_MI_GFPve_pos      0.6503830128  2.6471786819
## FC_Mpos_GAAGAATTCTTGCGCT-1FC_MI_GFPve_pos     -0.6442486770  3.2560162864
## FC_Mpos_GAAGCGAGTCGTTGCG-1FC_MI_GFPve_pos      0.2934106164  2.5598297439
## FC_Mpos_GAATCACAGTACAACA-1FC_MI_GFPve_pos     -0.9546669550  2.1676297508
## FC_Mpos_GACAGCCCAAGCAATA-1FC_MI_GFPve_pos      0.0952450506  2.6212377868
## FC_Mpos_GACCCTTTCAGCCTCT-1FC_MI_GFPve_pos     -0.2540151156  3.0609259926
## FC_Mpos_GACCTTCAGAGTTGAT-1FC_MI_GFPve_pos     -2.3381628819  2.0845299087
## FC_Mpos_GACTCTCGTCGCCTAG-1FC_MI_GFPve_pos     -0.2265507184  2.2668605171
## FC_Mpos_GACTTCCGTGACCGAA-1FC_MI_GFPve_pos      1.4825064115  2.7213917099
## FC_Mpos_GAGAAATAGGTTGGAC-1FC_MI_GFPve_pos      1.9244444542  3.2153292023
## FC_Mpos_GAGCCTGGTAACACGG-1FC_MI_GFPve_pos      0.5824613147  3.1618299804
## FC_Mpos_GAGCTGCTCAGGAGAC-1FC_MI_GFPve_pos      0.5323808961  1.4449839912
## FC_Mpos_GAGGCCTGTTCTCTAT-1FC_MI_GFPve_pos     -1.2091079779  3.4704866730
## FC_Mpos_GAGGGATCAGAAGTGC-1FC_MI_GFPve_pos     -1.5283393688  2.6502943359
## FC_Mpos_GAGGGATGTTGCATTG-1FC_MI_GFPve_pos     -1.5269331761  3.2280078254
## FC_Mpos_GAGTCTAGTGGCAGAT-1FC_MI_GFPve_pos     -1.8504054375  3.1111579262
## FC_Mpos_GAGTTACCAAAGTATG-1FC_MI_GFPve_pos     -0.0192013807  1.6879110657
## FC_Mpos_GAGTTACCACGCCACA-1FC_MI_GFPve_pos     -0.4731901415  3.1238785110
## FC_Mpos_GAGTTGTGTACGCTAT-1FC_MI_GFPve_pos     -2.2706701584  2.9611807190
## FC_Mpos_GAGTTGTGTTGAGTCT-1FC_MI_GFPve_pos     -0.2450784646  3.2010455452
## FC_Mpos_GATCGTAAGCTCCATA-1FC_MI_GFPve_pos     -0.5926347501  1.9235163055
```

```
## FC_Mpos_GATGATCGTCATAGTC-1FC_MI_GFPve_pos     -1.0700752564  3.0246486984
## FC_Mpos_GATGTTGGTGACTATC-1FC_MI_GFPve_pos     -0.6088065036  3.2767505966
## FC_Mpos_GATTTCTCAAACTAGA-1FC_MI_GFPve_pos      1.5198114090  3.1059065185
## FC_Mpos_GATTTCTTCACCTCGT-1FC_MI_GFPve_pos     -0.1735588387  1.0852456413
## FC_Mpos_GCAACATAGGGTAATT-1FC_MI_GFPve_pos     -2.2858953304  2.1824994407
## FC_Mpos_GCAACATTCCGATCTC-1FC_MI_GFPve_pos      1.3059653215  1.4953847252
## FC_Mpos_GCAACCGTCATTGTTC-1FC_MI_GFPve_pos      1.7560585909  2.7254739128
## FC_Mpos_GCACATATCATGAAAG-1FC_MI_GFPve_pos      1.2088675670  1.6810889564
## FC_Mpos_GCACATATCTGTGCAA-1FC_MI_GFPve_pos      0.2259172671  1.9600158058
## FC_Mpos_GCACTAATCCTACGAA-1FC_MI_GFPve_pos      0.3092248611  2.7674460731
## FC_Mpos_GCAGTTAAGGAGCTGT-1FC_MI_GFPve_pos     -2.2296056576  2.2725510917
## FC_Mpos_GCAGTTACAAGAGGTC-1FC_MI_GFPve_pos     -0.7728134222  3.3828487716
## FC_Mpos_GCATGATGTCTGTCAA-1FC_MI_GFPve_pos     -0.5518216021  2.1654382072
## FC_Mpos_GCATTAGAGCCACCGT-1FC_MI_GFPve_pos      1.3949178629  2.9321608864
## FC_Mpos_GCCATGGCACTGCGAC-1FC_MI_GFPve_pos     -1.0225505061  2.2755751930
## FC_Mpos_GCCCAGAAGAGTGTGC-1FC_MI_GFPve_pos     -0.6433937676  2.5353446327
## FC_Mpos_GCCCAGACATTGAGCT-1FC_MI_GFPve_pos      1.0717513733  1.6254101120
## FC_Mpos_GCCCGAAGTTGGATCT-1FC_MI_GFPve_pos      0.1710096769  2.1562481247
## FC_Mpos_GCCCGAATCAGGGATG-1FC_MI_GFPve_pos     -1.8163856096  2.9995999656
## FC_Mpos_GCGATCGAGTTGCCCG-1FC_MI_GFPve_pos      0.4064892106  2.8285146080
## FC_Mpos_GCGTGCATCCAGGACC-1FC_MI_GFPve_pos     -1.2164680786  2.5310054146
## FC_Mpos_GCTCAAAGTTAGGGTG-1FC_MI_GFPve_pos     -0.8296217151  2.7003450714
## FC_Mpos_GCTGAATAGGGCAGGA-1FC_MI_GFPve_pos     -0.5752163656  2.6408267341
## FC_Mpos_GCTTCACCAGTCTACA-1FC_MI_GFPve_pos     -0.1126436084  1.4997711502
## FC_Mpos_GCTTGGGGTGTTGCCG-1FC_MI_GFPve_pos     -1.7016159125  2.1322369896
## FC_Mpos_GGAACCCCATCACGGC-1FC_MI_GFPve_pos      0.6070509724  2.7621512733
## FC_Mpos_GGAATCTGTTGCCGCA-1FC_MI_GFPve_pos     -1.7845978565  1.8606324516
## FC_Mpos_GGACGTCCATCCTAAG-1FC_MI_GFPve_pos     -2.3531422444  2.3823170982
## FC_Mpos_GGAGAACAGAGCAGAA-1FC_MI_GFPve_pos     -0.1123334519  0.6687283836
## FC_Mpos_GGAGCAAGTATCCTCC-1FC_MI_GFPve_pos      0.9317211084  2.6075692497
## FC_Mpos_GGCAGTCAGTTAACGA-1FC_MI_GFPve_pos      1.1014807396  1.8533468567
## FC_Mpos_GGCAGTCCATGATGCT-1FC_MI_GFPve_pos      0.6598273687  1.0929909073
## FC_Mpos_GGCGTCAGTCCGGCAT-1FC_MI_GFPve_pos     -1.9279255696  1.6815467201
## FC_Mpos_GGGACAAAGGTTGTTC-1FC_MI_GFPve_pos     -1.2184918947  2.4285993896
## FC_Mpos_GGGACTCGTTATGTGC-1FC_MI_GFPve_pos      0.7342178993  1.9346309028
## FC_Mpos_GGGAGATGTGAGACCA-1FC_MI_GFPve_pos      0.9751050644  2.1761050545
## FC_Mpos_GGGAGATTCGACATCA-1FC_MI_GFPve_pos      0.2612990610  2.5367656074
## FC_Mpos_GGGATCCTCATTCATC-1FC_MI_GFPve_pos      0.8667823248  2.3154564224
## FC_Mpos_GGGCCATAGATGGCGT-1FC_MI_GFPve_pos      1.1433147363  3.3704233490
## FC_Mpos_GGGTATTAGTAGCATA-1FC_MI_GFPve_pos      1.9052627258  2.4041457496
## FC_Mpos_GGGTCACGTTTCCCAC-1FC_MI_GFPve_pos     -1.0747297831  1.8764582001
## FC_Mpos_GGGTGTCAGTCGAAGC-1FC_MI_GFPve_pos     -0.8488630481  1.4279785476
## FC_Mpos_GGGTGTCAGTTTCTTC-1FC_MI_GFPve_pos     -0.5210700162  2.3626585327
## FC_Mpos_GGGTGTCCATGTTTGG-1FC_MI_GFPve_pos     -0.3184948839  1.3387027107
## FC_Mpos_GGTTGTAAGGTTCACT-1FC_MI_GFPve_pos     -0.4013099350  2.7099528633
## FC_Mpos_GTAAGTCTCGCACGGT-1FC_MI_GFPve_pos      0.3386063151  3.2051048599
## FC_Mpos_GTAGAGGCAGAGCCCT-1FC_MI_GFPve_pos     -1.8526574202  1.7997908912
## FC_Mpos_GTAGGAGAGGAACTCG-1FC_MI_GFPve_pos     -2.7189044781  0.9823017441
## FC_Mpos_GTAGGAGCAACGGTAG-1FC_MI_GFPve_pos     -2.3056289978  1.6342392288
## FC_Mpos_GTAGGTTCAATTCTTC-1FC_MI_GFPve_pos      0.4830036931  2.3990998588
## FC_Mpos_GTATTTCAGCAGGGAG-1FC_MI_GFPve_pos     -1.4970068760  3.2816181503
## FC_Mpos_GTATTTCGTCAACCTA-1FC_MI_GFPve_pos      0.3776221447  2.0914202057
## FC_Mpos_GTCATGATCGGATACT-1FC_MI_GFPve_pos     -0.9111826844  3.2169113479
## FC_Mpos_GTCCTCATCGTAACTG-1FC_MI_GFPve_pos     -1.0238511391  0.7886977516
## FC_Mpos_GTCTAGACATGACAGG-1FC_MI_GFPve_pos     -0.6537122555  1.3683758102
## FC_Mpos_GTCTCACGTAGGTCAG-1FC_MI_GFPve_pos     -0.5126883514  1.7154865585
## FC_Mpos_GTCTGTCCAAGGAGTC-1FC_MI_GFPve_pos     -0.3411162771  3.1945748649
## FC_Mpos_GTCTTTAAGGACTTCT-1FC_MI_GFPve_pos     -1.5552706547  2.7829571090
## FC_Mpos_GTGACGCGTAATTGGA-1FC_MI_GFPve_pos      0.5712150507  2.5111961685
## FC_Mpos_GTGACGCGTTCTAAGC-1FC_MI_GFPve_pos      0.4665513210  2.4470429741
## FC_Mpos_GTGACGCTCAGCGCGT-1FC_MI_GFPve_pos     -0.3473449327  0.7786779724
## FC_Mpos_GTGATGTTCACGAGGA-1FC_MI_GFPve_pos     -1.2446770497  2.2704911552
## FC_Mpos_GTGCACGTCCACACCT-1FC_MI_GFPve_pos     -1.0024847813  2.6501679741
## FC_Mpos_GTGCAGCCAGTGTACT-1FC_MI_GFPve_pos     -0.4122547783  1.3278103195
## FC_Mpos_GTGCAGCTCCAGCAAT-1FC_MI_GFPve_pos     -1.0620481200  2.9769192062
## FC_Mpos_GTGCGTGAGAACTGAT-1FC_MI_GFPve_pos      1.3004080229  2.1633029304
## FC_Mpos_GTGCGTGAGTACAGAT-1FC_MI_GFPve_pos     -1.2340402432  2.4612164818
## FC_Mpos_GTGCGTGCACATTACG-1FC_MI_GFPve_pos     -0.8448109455  3.5624256454
## FC_Mpos_GTGCTGGAGTAAGAGG-1FC_MI_GFPve_pos     -0.5558285542  3.1291570984
## FC_Mpos_GTGCTTCGTTGCACGC-1FC_MI_GFPve_pos     -1.5670739241  2.4471168838
```

```
## FC_Mpos_GTGGAAGAGAGGTTTA-1FC_MI_GFPve_pos     -0.3290371425  0.6561265312
## FC_Mpos_GTGGAAGAGGATGCGT-1FC_MI_GFPve_pos     -0.6679136582  2.0245385490
## FC_Mpos_GTGGCGTAGGTCCCTG-1FC_MI_GFPve_pos     -0.5007060297  3.2021780334
## FC_Mpos_GTGGCGTCAGAACGCA-1FC_MI_GFPve_pos      0.2636649184  1.6819210373
## FC_Mpos_GTGGGAACAGCAAGAC-1FC_MI_GFPve_pos      1.3546220236  3.1894116722
## FC_Mpos_GTGGGAATCAGGGTAG-1FC_MI_GFPve_pos      1.7892352514  2.0452900253
## FC_Mpos_GTGTAACCACCAGCCA-1FC_MI_GFPve_pos      0.2334109120  2.2833557449
## FC_Mpos_GTGTCCTAGAAGCTGC-1FC_MI_GFPve_pos     -0.2804649971  3.1527424179
## FC_Mpos_GTGTCCTCATCGGTTA-1FC_MI_GFPve_pos     -0.1795675313  1.6024523102
## FC_Mpos_GTGTTCCCAACTGTGT-1FC_MI_GFPve_pos      0.7734610014  2.1884670578
## FC_Mpos_GTGTTCCCAGTGTGGA-1FC_MI_GFPve_pos     -1.8821316786  3.0621300064
## FC_Mpos_GTGTTCCGTTCTCGCT-1FC_MI_GFPve_pos     -1.3656415768  2.2512460075
## FC_Mpos_GTTACAGCAATGAAAC-1FC_MI_GFPve_pos      1.2791743450  1.3413486801
## FC_Mpos_GTTATGGAGCAACTCT-1FC_MI_GFPve_pos     -0.7139314838  3.3635654770
## FC_Mpos_GTTCATTAGACATCCT-1FC_MI_GFPve_pos      0.6571459942  3.3653240524
## FC_Mpos_GTTCTATAGAGCCGTA-1FC_MI_GFPve_pos     -0.8755453057  2.9030256592
## FC_Mpos_GTTCTATTCCAGTTCC-1FC_MI_GFPve_pos     -1.3089518376  2.3529830299
## FC_Mpos_GTTGTAGGTGGGATTG-1FC_MI_GFPve_pos     -0.7097644634  2.1899462066
## FC_Mpos_GTTGTGAGTCAACACT-1FC_MI_GFPve_pos     -0.2626578711  2.8442201935
## FC_Mpos_GTTTACTCAGAGTGAC-1FC_MI_GFPve_pos      0.2640116982  2.8425965629
## FC_Mpos_TAAGCCATCGGAGTGA-1FC_MI_GFPve_pos     -2.0895677633  1.6574282966
## FC_Mpos_TAAGTCGGTCGTGTTA-1FC_MI_GFPve_pos     -1.1759056874  3.4579353653
## FC_Mpos_TAAGTCGGTTAAAGTG-1FC_MI_GFPve_pos     -0.7547959514  3.0053110443
## FC_Mpos_TACACCCCACTAGGTT-1FC_MI_GFPve_pos     -0.7681054898  3.1180219971
## FC_Mpos_TACCGGGTCTTCCAGC-1FC_MI_GFPve_pos     -1.2949330635  3.2591657959
## FC_Mpos_TACCTCGTCCGCGAGT-1FC_MI_GFPve_pos      2.3543157749  2.3669662796
## FC_Mpos_TACGCTCGTTCTTGCC-1FC_MI_GFPve_pos     -0.7144449182  1.8531933151
## FC_Mpos_TACTTACTCGGTCGAC-1FC_MI_GFPve_pos     -1.3035650082  1.6279945694
## FC_Mpos_TAGACCAGTTGCGGAA-1FC_MI_GFPve_pos     -0.8124170132  3.3082819305
## FC_Mpos_TAGAGTCGTAACTTCG-1FC_MI_GFPve_pos     -0.3341028668  1.1186557136
## FC_Mpos_TAGGAGGGTCATCACA-1FC_MI_GFPve_pos     -2.2014953919  1.3767829262
## FC_Mpos_TAGGAGGGTCGAATTC-1FC_MI_GFPve_pos      0.7544380002  2.5712943397
## FC_Mpos_TAGGTACAGTTCACTG-1FC_MI_GFPve_pos      1.5445318394  1.5668735824
## FC_Mpos_TAGTGCACATCGAGCC-1FC_MI_GFPve_pos     -0.7890707679  1.6469111763
## FC_Mpos_TATATCCGTCGTCAGC-1FC_MI_GFPve_pos     -1.0349807568  2.2785425506
## FC_Mpos_TATATCCTCCCGTGTT-1FC_MI_GFPve_pos     -0.0731507219  3.3059397064
## FC_Mpos_TATCCTAGTTCCTTGC-1FC_MI_GFPve_pos     -1.2596330471  1.9852595650
## FC_Mpos_TATCGCCGTATCCCTC-1FC_MI_GFPve_pos      0.1453994982  1.1835875831
## FC_Mpos_TATCTGTAGTGGTGGT-1FC_MI_GFPve_pos      0.2549476199  2.5220280014
## FC_Mpos_TCAAGCATCGCCTATC-1FC_MI_GFPve_pos      0.6735469274  2.3383546196
## FC_Mpos_TCACACCCACTATGTG-1FC_MI_GFPve_pos     -1.3836003370  1.4747028671
## FC_Mpos_TCACGCTTCCTTGGAA-1FC_MI_GFPve_pos      0.2384717040  2.1074233375
## FC_Mpos_TCACGGGTCCGGCAAC-1FC_MI_GFPve_pos     -0.3576211907  2.4921584450
## FC_Mpos_TCACTATAGCGTCAAG-1FC_MI_GFPve_pos      0.2787671857  2.6947331749
## FC_Mpos_TCACTCGCAGGACTAG-1FC_MI_GFPve_pos      0.4004978113  3.3137474380
## FC_Mpos_TCATCATAGACCTCCG-1FC_MI_GFPve_pos      0.2256789379  2.3084545456
## FC_Mpos_TCATCATCAGATTAAG-1FC_MI_GFPve_pos      0.9908200674  2.6006431900
## FC_Mpos_TCATCCGAGGCCTGAA-1FC_MI_GFPve_pos      0.2328576021  2.4638634048
## FC_Mpos_TCATGAGAGTTAGTAG-1FC_MI_GFPve_pos     -1.3710056372  2.0106821380
## FC_Mpos_TCATGAGTCGATGCTA-1FC_MI_GFPve_pos     -0.1001925968  2.7252378784
## FC_Mpos_TCATGGACAAATACAG-1FC_MI_GFPve_pos      1.2119621210  1.7292704902
## FC_Mpos_TCCATGCAGTCTGGAG-1FC_MI_GFPve_pos     -0.7728301711  3.3353405319
## FC_Mpos_TCCATGCTCATACGGT-1FC_MI_GFPve_pos      0.1714348666  2.0938124977
## FC_Mpos_TCCCACAAGCGATTCT-1FC_MI_GFPve_pos     -0.2305589966  1.4258141838
## FC_Mpos_TCCCACAGTGGAGAAA-1FC_MI_GFPve_pos     -1.6315911837  2.1644463859
## FC_Mpos_TCCCATGTCGTTCTCG-1FC_MI_GFPve_pos     -1.3438963719  2.0146389328
## FC_Mpos_TCCGATCAGTTGTAGA-1FC_MI_GFPve_pos     -1.8092588015  2.4688783012
## FC_Mpos_TCCGATCCAAAGGTTA-1FC_MI_GFPve_pos     -1.6275961228  2.6198673569
## FC_Mpos_TCCGTGTCACTTCCTG-1FC_MI_GFPve_pos     -0.3052121974  2.6397166572
## FC_Mpos_TCCTCTTCAGATCCAT-1FC_MI_GFPve_pos      0.2350122206  1.8286757789
## FC_Mpos_TCGACCTAGCTTACGT-1FC_MI_GFPve_pos     -1.1891170569  2.3594718300
## FC_Mpos_TCGACGGGTAACGCGA-1FC_MI_GFPve_pos      0.4279125504  2.7294373833
## FC_Mpos_TCGATTTTCCCGAATA-1FC_MI_GFPve_pos      1.0952753000  2.0112495743
## FC_Mpos_TCGGATATCGCGTTTC-1FC_MI_GFPve_pos     -1.6485558577  2.5550504051
## FC_Mpos_TCGGTCTAGACTTGTC-1FC_MI_GFPve_pos     -0.4352588482  2.1337385498
## FC_Mpos_TCGGTCTCATTGAGCT-1FC_MI_GFPve_pos     -1.7779027052  2.3508248649
## FC_Mpos_TCGTGCTCAGTTAAAG-1FC_MI_GFPve_pos     -1.6665997334  2.8606396042
## FC_Mpos_TCGTGCTGTGATACAA-1FC_MI_GFPve_pos     -0.6606825121  2.4353914581
## FC_Mpos_TCTAACTCAGGTCAAG-1FC_MI_GFPve_pos     -0.3199444540  0.4690943085
## FC_Mpos_TCTACCGAGCGAGGAG-1FC_MI_GFPve_pos     -0.3980768450  3.1434417091
```

```
## FC_Mpos_TCTACCGGTACTGAGG-1FC_MI_GFPve_pos      0.5976522736  2.2310858093
## FC_Mpos_TCTCAGCCACGTGTGC-1FC_MI_GFPve_pos     -1.4881951637  2.7664747558
## FC_Mpos_TCTCAGCGTATCGATC-1FC_MI_GFPve_pos      1.0264010601  2.5638294540
## FC_Mpos_TCTCTGGTCTCGTGAA-1FC_MI_GFPve_pos      1.1701209717  2.5114202820
## FC_Mpos_TCTGCCAAGAGCAGCT-1FC_MI_GFPve_pos     -0.1178892791  2.0280008636
## FC_Mpos_TCTGGCTCATACCGTA-1FC_MI_GFPve_pos     -0.2563540913  1.7551851593
## FC_Mpos_TCTGGCTGTCTGTAGT-1FC_MI_GFPve_pos     -1.2940587826  2.0304937683
## FC_Mpos_TCTTCCTTCAAAGAAC-1FC_MI_GFPve_pos      0.2634393446  3.0431738220
## FC_Mpos_TCTTGCGAGTTGAAGT-1FC_MI_GFPve_pos     -0.0318353750  3.3076029144
## FC_Mpos_TGAACGTTCGCTCATC-1FC_MI_GFPve_pos     -0.4556782670  2.9837670646
## FC_Mpos_TGACAGTCAGTCAACT-1FC_MI_GFPve_pos     -1.5104989834  1.8527674995
## FC_Mpos_TGACAGTTCCCTCTTT-1FC_MI_GFPve_pos     -0.1364413571  1.3847427688
## FC_Mpos_TGACGCGCAGCTGTAT-1FC_MI_GFPve_pos      0.3843211703  2.5688925110
## FC_Mpos_TGAGACTTCATTACGG-1FC_MI_GFPve_pos     -1.9478752442  3.2007556282
## FC_Mpos_TGAGGTTCAGTCGGAA-1FC_MI_GFPve_pos     -0.2160551041  3.1598692261
## FC_Mpos_TGAGGTTGTAACGATA-1FC_MI_GFPve_pos     -1.8381110258  2.2398801170
## FC_Mpos_TGATCAGTCGTAGCCG-1FC_MI_GFPve_pos     -0.8361230917  1.1989984833
## FC_Mpos_TGATGCACAGGTGTGA-1FC_MI_GFPve_pos     -0.2591834344  2.1590171181
## FC_Mpos_TGATGCACATCTTCGC-1FC_MI_GFPve_pos     -1.0598587818  2.5042491279
## FC_Mpos_TGATGCATCACTGTTT-1FC_MI_GFPve_pos      1.0361566715  2.6589751564
## FC_Mpos_TGATGGTAGTTTGAGA-1FC_MI_GFPve_pos     -0.5221598871  3.1125746094
## FC_Mpos_TGATTTCCATACCACA-1FC_MI_GFPve_pos      0.7432853751  2.9135980926
## FC_Mpos_TGCAGATGTACCGGAA-1FC_MI_GFPve_pos     -0.9795318074  3.1129131637
## FC_Mpos_TGCATGACAGCACGAA-1FC_MI_GFPve_pos     -0.8707265086  1.9710111938
## FC_Mpos_TGCCGAGCAGACGCTC-1FC_MI_GFPve_pos      0.3633389644  2.6469130836
## FC_Mpos_TGCGGGTTCTCGACCT-1FC_MI_GFPve_pos     -1.1431129403  2.2215614639
## FC_Mpos_TGCTCGTCAGGTTTAC-1FC_MI_GFPve_pos     -1.4040925331  1.9586081825
## FC_Mpos_TGGATCACATTGCCTC-1FC_MI_GFPve_pos     -2.1003660984  2.9919267021
## FC_Mpos_TGGATGTTCCAGCTCT-1FC_MI_GFPve_pos     -0.4123718418  1.4212456070
## FC_Mpos_TGGGCTGTCAGGTGTT-1FC_MI_GFPve_pos     -1.3646100588  3.1188035332
## FC_Mpos_TGGGTTATCACAATGC-1FC_MI_GFPve_pos     -0.8882146902  2.0027704559
## FC_Mpos_TGTAAGCTCCACGTGG-1FC_MI_GFPve_pos      0.1819431059  2.9392052017
## FC_Mpos_TGTCAGAGTAATGCGG-1FC_MI_GFPve_pos     -0.1502828576  2.6405635200
## FC_Mpos_TGTCCACTCTGCCTCA-1FC_MI_GFPve_pos     -0.1135487571  1.0600920044
## FC_Mpos_TGTCCTGTCGTTATCT-1FC_MI_GFPve_pos     -0.7580666371  1.9622268997
## FC_Mpos_TGTGATGGTGCCCTTT-1FC_MI_GFPve_pos     -1.1693414278  3.1879129730
## FC_Mpos_TGTGCGGCATGCCGGT-1FC_MI_GFPve_pos     -1.8429921694  2.0083670936
## FC_Mpos_TGTGGCGTCGCAACAT-1FC_MI_GFPve_pos     -1.9536942072  3.1100435577
## FC_Mpos_TGTGTGACAAGCAGGT-1FC_MI_GFPve_pos      1.5585693292  2.9513297401
## FC_Mpos_TGTTCCGTCATATGGC-1FC_MI_GFPve_pos     -0.4569602139  2.6478910766
## FC_Mpos_TGTTGAGGTCAGTCCG-1FC_MI_GFPve_pos      0.3600387745  1.7322554909
## FC_Mpos_TGTTTGTCAATACGAA-1FC_MI_GFPve_pos     -0.7871055432  2.8225460373
## FC_Mpos_TTAATCCTCCACTTTA-1FC_MI_GFPve_pos     -1.3271390028  2.7846370064
## FC_Mpos_TTACAGGGTGGAACAC-1FC_MI_GFPve_pos     -0.1766589280  2.9280619942
## FC_Mpos_TTACAGGTCTACTCAT-1FC_MI_GFPve_pos      0.6324185900  1.6978350006
## FC_Mpos_TTACCATGTACGGATG-1FC_MI_GFPve_pos      0.3044806890  1.8786278091
## FC_Mpos_TTACCGCTCCTCATAT-1FC_MI_GFPve_pos      0.2772751324  2.9448585830
## FC_Mpos_TTACGCCCATGGGCAA-1FC_MI_GFPve_pos      1.3480409555  1.9872513138
## FC_Mpos_TTAGGCACACCAGGTC-1FC_MI_GFPve_pos     -0.6770327516  1.0348382316
## FC_Mpos_TTAGTCTTCAATCGGT-1FC_MI_GFPve_pos      0.1619279854  0.0050330482
## FC_Mpos_TTATTGCAGGTTATAG-1FC_MI_GFPve_pos     -0.5617884226  2.7297277771
## FC_Mpos_TTATTGCAGTCACACT-1FC_MI_GFPve_pos     -0.0382362880  3.4220209442
## FC_Mpos_TTATTGCCAGGACATG-1FC_MI_GFPve_pos     -0.4712106414  2.2073746048
## FC_Mpos_TTCAATCCACACGCCA-1FC_MI_GFPve_pos     -0.5802343316  2.3688922249
## FC_Mpos_TTCACGCAGACGACGT-1FC_MI_GFPve_pos     -0.1824948365  1.9197130523
## FC_Mpos_TTCACGCTCTGTAACG-1FC_MI_GFPve_pos      0.3380735092  2.2750368439
## FC_Mpos_TTCCAATAGAGTTGAT-1FC_MI_GFPve_pos     -1.2076527901  3.2069211326
## FC_Mpos_TTCCGGTGTAGCGCTC-1FC_MI_GFPve_pos      1.0102467708  1.9494510017
## FC_Mpos_TTCCGTGAGACGATAT-1FC_MI_GFPve_pos     -1.3707924910  2.3654256187
## FC_Mpos_TTCCTAAAGGCCCACT-1FC_MI_GFPve_pos     -0.0678206734  3.1425538383
## FC_Mpos_TTCGCTGCAGGTCAGA-1FC_MI_GFPve_pos     -0.9100345798  2.2329340301
## FC_Mpos_TTCGGTCTCCTCTCGA-1FC_MI_GFPve_pos      0.5957339577  2.0038833938
## FC_Mpos_TTCTAACGTCCAGAAG-1FC_MI_GFPve_pos     -1.8392147846  2.7502065025
## FC_Mpos_TTCTTCCAGGCGTCCT-1FC_MI_GFPve_pos     -0.5941559203  3.1442809425
## FC_Mpos_TTCTTGACAACACGTT-1FC_MI_GFPve_pos     -1.4537025519  2.6906128250
## FC_Mpos_TTGAACGAGGCGAAGG-1FC_MI_GFPve_pos      0.2972100549  2.0860820137
## FC_Mpos_TTGAGTGAGAAGCGGG-1FC_MI_GFPve_pos     -0.0311043687  2.7404475532
## FC_Mpos_TTGAGTGAGGTGCTTT-1FC_MI_GFPve_pos      0.4885560446  2.8851609550
## FC_Mpos_TTGAGTGCAATAGGAT-1FC_MI_GFPve_pos     -1.1006707378  2.7321577392
## FC_Mpos_TTGAGTGTCGACGACC-1FC_MI_GFPve_pos     -1.6761616297  1.5770278297
```

```
## FC_Mpos_TTGCTGCAGCACTTTG-1FC_MI_GFPve_pos      0.1112963729  2.3218746506
## FC_Mpos_TTGGGCGAGACCTCAT-1FC_MI_GFPve_pos     -0.6235948630  0.8992791496
## FC_Mpos_TTGTTCATCACTCACC-1FC_MI_GFPve_pos     -1.8018206186  2.7403827034
## FC_Mpos_TTTACGTTCAAGTGGG-1FC_MI_GFPve_pos     -2.3838465042  1.7021613441
## FC_Mpos_TTTCACACATCGGAGA-1FC_MI_GFPve_pos      0.2605767720  2.6200776420
## FC_Mpos_TTTCACATCGTCCATC-1FC_MI_GFPve_pos     -1.1593688555  1.8716435753
## FC_Mpos_TTTCATGTCGGAACTT-1FC_MI_GFPve_pos     -0.8068082161  2.6076035820
## FC_Mpos_TTTCATGTCTACCACC-1FC_MI_GFPve_pos     -0.3984380938  2.8430576645
## FC_Mpos_TTTCGATGTATGGAAT-1FC_MI_GFPve_pos      1.2783147268  2.4730201088
## FC_Mpos_TTTGTTGAGCATGATA-1FC_MI_GFPve_pos      0.6494701676  2.8894329391
## FC_Fpos_AAACGCTAGGTTCAGG-1FC_FI_GFPve_pos     -1.8254207201  3.2088470779
## FC_Fpos_AAAGGATCAGTTTCGA-1FC_FI_GFPve_pos     -0.1939418927  3.0591135345
## FC_Fpos_AAAGGGCCAGTCTGGC-1FC_FI_GFPve_pos     -0.9751936026  3.1767735801
## FC_Fpos_AAAGGTACAGAGTCTT-1FC_FI_GFPve_pos      2.5186593704  2.7354941688
## FC_Fpos_AACACACTCGTTTACT-1FC_FI_GFPve_pos     -1.1966609783  1.2347793899
## FC_Fpos_AACAGGGAGGAACTCG-1FC_FI_GFPve_pos     -0.2714924715  2.7802868210
## FC_Fpos_AACCACAAGTTCTCTT-1FC_FI_GFPve_pos     -0.4039281733  2.0227179848
## FC_Fpos_AACCACAGTTGCTGAT-1FC_FI_GFPve_pos     -0.7146510310  2.1999883972
## FC_Fpos_AACCTGATCATTTACC-1FC_FI_GFPve_pos     -1.2810514994  2.3763509117
## FC_Fpos_AACCTTTGTTCGGGTC-1FC_FI_GFPve_pos     -1.5672513075  2.1458301864
## FC_Fpos_AACGTCATCTAATTCC-1FC_FI_GFPve_pos     -1.9939365216  0.3049269043
## FC_Fpos_AACTTCTTCGCTGCGA-1FC_FI_GFPve_pos     -1.4223158188  2.2572832428
## FC_Fpos_AAGAACACACAGTCGC-1FC_FI_GFPve_pos     -0.5263964660  2.3052759491
## FC_Fpos_AAGAACAGTCGGCTAC-1FC_FI_GFPve_pos     -1.0054930277  2.7280774437
## FC_Fpos_AAGACTCTCAGCCCAG-1FC_FI_GFPve_pos      0.6040911250  3.3564739548
## FC_Fpos_AAGCCATAGTATGTAG-1FC_FI_GFPve_pos      0.2075200550  2.5781951271
## FC_Fpos_AAGCCATCACAGTGTT-1FC_FI_GFPve_pos     -0.0634328850  2.0692406021
## FC_Fpos_AAGCCATGTCGAGCAA-1FC_FI_GFPve_pos     -1.3881453104  2.2784686409
## FC_Fpos_AAGCGAGCACTTCAAG-1FC_FI_GFPve_pos     -1.2240278549  1.9389515243
## FC_Fpos_AAGGAATCATTAAGCC-1FC_FI_GFPve_pos      2.2769477539  2.7074442230
## FC_Fpos_AAGGTAACACAACATC-1FC_FI_GFPve_pos     -0.9376278944  1.9975123726
## FC_Fpos_AAGGTAACACTACCGG-1FC_FI_GFPve_pos     -1.5826325245  1.9617748581
## FC_Fpos_AAGTACCAGATGGGCT-1FC_FI_GFPve_pos     -1.0567929573  1.4688411079
## FC_Fpos_AAGTACCGTTAAGACA-1FC_FI_GFPve_pos     -0.0234668381  0.9635038696
## FC_Fpos_AAGTACCTCCCGGTAG-1FC_FI_GFPve_pos     -1.5274247952  2.3507600151
## FC_Fpos_AAGTCGTAGGAAAGAC-1FC_FI_GFPve_pos     -2.0555478163  2.9113931976
## FC_Fpos_AAGTCGTTCCATCTCG-1FC_FI_GFPve_pos     -2.2376746960  2.0789490066
## FC_Fpos_AAGTCGTTCTCTCGCA-1FC_FI_GFPve_pos      0.0608855210  2.4607329689
## FC_Fpos_AAGTGAAGTTCCGCAG-1FC_FI_GFPve_pos     -0.4447646566  2.2551040970
## FC_Fpos_AAGTTCGCAACCCTAA-1FC_FI_GFPve_pos      0.3737566643  2.9079428039
## FC_Fpos_AATCGTGCACGGAAGT-1FC_FI_GFPve_pos      1.6339298658  3.0097985588
## FC_Fpos_AATGACCGTACCTAGT-1FC_FI_GFPve_pos     -0.9521500297  3.1115312897
## FC_Fpos_AATGCCATCCATCTCG-1FC_FI_GFPve_pos      0.7777806692  2.4299702965
## FC_Fpos_ACAAAGAGTGCCAAGA-1FC_FI_GFPve_pos      0.6743284039  1.7284202896
## FC_Fpos_ACAACCACAAGATCCT-1FC_FI_GFPve_pos     -1.5852994747  3.0809922539
## FC_Fpos_ACACAGTAGCTAGAGC-1FC_FI_GFPve_pos      1.6973766260  2.2755008064
## FC_Fpos_ACACAGTTCGACATCA-1FC_FI_GFPve_pos      0.1157553665 -0.1125974335
## FC_Fpos_ACACCAAAGCACGATG-1FC_FI_GFPve_pos     -0.9790220089  1.8759808861
## FC_Fpos_ACACGCGAGTAACAGT-1FC_FI_GFPve_pos      0.7489003711  2.7003355346
## FC_Fpos_ACACGCGGTACGGTTT-1FC_FI_GFPve_pos     -0.5076741703  2.2969866119
## FC_Fpos_ACACGCGTCGAATCCA-1FC_FI_GFPve_pos     -1.6499432154  2.5029707275
## FC_Fpos_ACAGCCGCATTGACCA-1FC_FI_GFPve_pos     -0.1871964630  2.7971043907
## FC_Fpos_ACAGCCGTCGCCTTTG-1FC_FI_GFPve_pos     -0.9956486054  1.4094853721
## FC_Fpos_ACAGGGACAGGCTTGC-1FC_FI_GFPve_pos     -1.3196732826  2.3366580330
## FC_Fpos_ACAGGGATCGGTGTAT-1FC_FI_GFPve_pos     -0.0556015871  3.0955400787
## FC_Fpos_ACCAACAGTCGTCATA-1FC_FI_GFPve_pos     -0.2570961840  3.0994816147
## FC_Fpos_ACCCAAAGTCTTGCTC-1FC_FI_GFPve_pos     -2.0744790859  2.9803581558
## FC_Fpos_ACCCTCACAATTAGGA-1FC_FI_GFPve_pos      0.0197699837  1.0930519424
## FC_Fpos_ACCGTTCGTAGAATAC-1FC_FI_GFPve_pos     -0.0123068221  1.8881178222
## FC_Fpos_ACCTACCAGCGGTATG-1FC_FI_GFPve_pos     -0.8748683162  2.2437138878
## FC_Fpos_ACCTGAACAGACGGAT-1FC_FI_GFPve_pos     -0.4009829737  2.4934416137
## FC_Fpos_ACCTGAATCTGGGCCA-1FC_FI_GFPve_pos     -0.7171432204  2.2629399620
## FC_Fpos_ACCTGTCTCCGGACGT-1FC_FI_GFPve_pos      1.4946806602  2.6207366310
## FC_Fpos_ACGCACGTCCAGTGCG-1FC_FI_GFPve_pos     -2.1385212965  1.9379806839
## FC_Fpos_ACGGAAGAGACTTCAC-1FC_FI_GFPve_pos     -0.9776491947  3.2118978821
## FC_Fpos_ACGGGTCCAAATGCGG-1FC_FI_GFPve_pos      0.5402705841  2.1557331405
## FC_Fpos_ACGGGTCCAGAACTTC-1FC_FI_GFPve_pos     -0.8173775501  2.9283829056
## FC_Fpos_ACGGGTCGTATGAAAC-1FC_FI_GFPve_pos     -0.6509292550  2.2017321907
## FC_Fpos_ACGGTCGCATCTTTCA-1FC_FI_GFPve_pos     -0.4237606652  2.9098630272
## FC_Fpos_ACGTACAAGTGCCCGT-1FC_FI_GFPve_pos     -0.5048927791  2.0039234482
```

```
## FC_Fpos_ACGTACAGTATGGTAA-1FC_FI_GFPve_pos     -1.9657181330  2.2890706383
## FC_Fpos_ACGTAGTAGACTCTAC-1FC_FI_GFPve_pos     -0.2618297763  2.3849840484
## FC_Fpos_ACGTAGTGTAGCGCCT-1FC_FI_GFPve_pos     -0.7512329645  3.4696092926
## FC_Fpos_ACGTCCTAGCTGAGCA-1FC_FI_GFPve_pos     -1.6753670998  1.9140224777
## FC_Fpos_ACGTCCTCAAGTCGTT-1FC_FI_GFPve_pos      1.2131455116  2.6061225258
## FC_Fpos_ACGTTCCCAGCACAAG-1FC_FI_GFPve_pos     -2.3297638722  2.2754330955
## FC_Fpos_ACGTTCCGTCGCGTCA-1FC_FI_GFPve_pos     -0.5870955117  2.1460542999
## FC_Fpos_ACTATCTAGCATCAAA-1FC_FI_GFPve_pos     -1.5978663988  2.7402983032
## FC_Fpos_ACTATGGTCAGTGGGA-1FC_FI_GFPve_pos      1.0087500982  2.9818287216
## FC_Fpos_ACTGATGTCGCAGATT-1FC_FI_GFPve_pos     -2.1200494595  2.6282553993
## FC_Fpos_ACTGCAAAGATACGAT-1FC_FI_GFPve_pos     -1.1189055867  2.7819609962
## FC_Fpos_ACTGCAAGTTTACTGG-1FC_FI_GFPve_pos     -2.1116109915  2.4454856239
## FC_Fpos_ACTGTCCAGCGGATCA-1FC_FI_GFPve_pos      0.5228619151  2.2867245994
## FC_Fpos_ACTTAGGCAAAGGTTA-1FC_FI_GFPve_pos      0.2013848595  3.0031309448
## FC_Fpos_ACTTCGCTCGGCTGAC-1FC_FI_GFPve_pos     -2.3119063206  2.8900285087
## FC_Fpos_ACTTTCAAGGAGAGGC-1FC_FI_GFPve_pos     -0.3695844985  3.3591938339
## FC_Fpos_ACTTTCATCTAGTCAG-1FC_FI_GFPve_pos     -0.3919201173  2.7023921333
## FC_Fpos_ACTTTGTCAGAAGTGC-1FC_FI_GFPve_pos     -1.9684120245  2.5827327095
## FC_Fpos_ACTTTGTGTTTGAACC-1FC_FI_GFPve_pos      0.7494124346  2.5901689850
## FC_Fpos_AGAACAACAATAAGGT-1FC_FI_GFPve_pos      0.3230366580  3.2505874954
## FC_Fpos_AGAACAAGTTAGAAAC-1FC_FI_GFPve_pos     -1.6429894991  1.8330612503
## FC_Fpos_AGAACCTCAGTAACCT-1FC_FI_GFPve_pos      0.6126676969  2.3956361137
## FC_Fpos_AGAAGCGGTAGGTCAG-1FC_FI_GFPve_pos     -2.4091989823  2.0385046325
## FC_Fpos_AGACAAATCTATACGG-1FC_FI_GFPve_pos      0.0490655474  2.1907735191
## FC_Fpos_AGACACTGTAACTGCT-1FC_FI_GFPve_pos     -0.2316756748  1.8087182365
## FC_Fpos_AGACACTGTTATGTGC-1FC_FI_GFPve_pos     -0.8480987377  1.5478654228
## FC_Fpos_AGACAGGTCCTCAGGG-1FC_FI_GFPve_pos     -0.9728477426  3.0513768516
## FC_Fpos_AGACCATAGATTGAGT-1FC_FI_GFPve_pos     -0.0181993224  2.8305368744
## FC_Fpos_AGACCATCACCCTGAG-1FC_FI_GFPve_pos      0.7852937155  2.5680780731
## FC_Fpos_AGAGAATCATGACTCA-1FC_FI_GFPve_pos     -1.9544427223  2.8815045677
## FC_Fpos_AGAGAATGTACTGGGA-1FC_FI_GFPve_pos     -1.5074546166  3.0781565033
## FC_Fpos_AGAGCCCGTGAACGGT-1FC_FI_GFPve_pos     -1.1807746716  2.7702913605
## FC_Fpos_AGAGCCCGTGAGACCA-1FC_FI_GFPve_pos     -1.6960372753  2.6039681755
## FC_Fpos_AGATAGAGTCGTAATC-1FC_FI_GFPve_pos     -0.0425063573  2.1787019096
## FC_Fpos_AGATGCTAGTAGCTCT-1FC_FI_GFPve_pos      0.0336934947  2.4625749908
## FC_Fpos_AGCCAATGTCGATGCC-1FC_FI_GFPve_pos     -0.7651658483  3.0493803345
## FC_Fpos_AGCCAATTCTATTCGT-1FC_FI_GFPve_pos      1.7996066980  2.7250075661
## FC_Fpos_AGCCACGCAGATTAAG-1FC_FI_GFPve_pos     -0.0309617795  2.4522705398
## FC_Fpos_AGCCAGCGTTTGTGGT-1FC_FI_GFPve_pos     -1.7308028765  1.9974971138
## FC_Fpos_AGCGATTTCGTGAGAG-1FC_FI_GFPve_pos     -2.3947341271  2.3072419487
## FC_Fpos_AGCGTATCAACGTATC-1FC_FI_GFPve_pos     -0.0158428229  3.0356874786
## FC_Fpos_AGCTACAGTTCGTAAC-1FC_FI_GFPve_pos     -1.7983494826  1.7640700661
## FC_Fpos_AGCTCAAAGGCTAGCA-1FC_FI_GFPve_pos     -0.5850399561  2.3316798530
## FC_Fpos_AGGAATACACGTACAT-1FC_FI_GFPve_pos     -0.4182315893  1.3233480774
## FC_Fpos_AGGAATAGTCTCAGGC-1FC_FI_GFPve_pos     -1.4143602677  2.1470151268
## FC_Fpos_AGGACTTGTCATCAGT-1FC_FI_GFPve_pos     -0.2091966696  2.8146190964
## FC_Fpos_AGGAGGTAGGACCCAA-1FC_FI_GFPve_pos     -0.5939896233  3.0135259949
## FC_Fpos_AGGAGGTGTCGGATTT-1FC_FI_GFPve_pos      2.0790288620  3.0721683822
## FC_Fpos_AGGATAAGTTTCCAAG-1FC_FI_GFPve_pos     -0.6607750483  3.2773008667
## FC_Fpos_AGGCATTAGGCCACCT-1FC_FI_GFPve_pos     -0.2750824965  2.8760500274
## FC_Fpos_AGGCATTGTGGCTACC-1FC_FI_GFPve_pos     -1.9627942867  2.4841018043
## FC_Fpos_AGGCCACCACATACTG-1FC_FI_GFPve_pos     -2.2248551674  2.8889012657
## FC_Fpos_AGGCCACTCTGCGAGC-1FC_FI_GFPve_pos      0.3548768930  1.9622321449
## FC_Fpos_AGGGAGTCAACCCTCT-1FC_FI_GFPve_pos      0.0857020788  2.5933394752
## FC_Fpos_AGGGCCTCAACCGTAT-1FC_FI_GFPve_pos      1.0072721414  2.9409895263
## FC_Fpos_AGGGTTTCAGGAATAT-1FC_FI_GFPve_pos      0.4436649852  2.3706612907
## FC_Fpos_AGGGTTTGTCCCTAAA-1FC_FI_GFPve_pos     -0.4804114230  1.3667769752
## FC_Fpos_AGGTCATAGATTCGCT-1FC_FI_GFPve_pos     -1.2336497135  2.1076498352
## FC_Fpos_AGGTCATTCGACGCTG-1FC_FI_GFPve_pos      0.8814350538  2.7134318672
## FC_Fpos_AGGTGTTTCCACAGGC-1FC_FI_GFPve_pos      0.7867353134  3.0768404327
## FC_Fpos_AGGTTACAGCATCCCG-1FC_FI_GFPve_pos     -0.1110942826  0.8278084122
## FC_Fpos_AGGTTACGTCTTAGTG-1FC_FI_GFPve_pos      2.0876059704  2.9734316192
## FC_Fpos_AGTAACCCACTCCCTA-1FC_FI_GFPve_pos     -0.0229961850  2.9230432831
## FC_Fpos_AGTACCAAGGCTCTAT-1FC_FI_GFPve_pos     -0.4675866194  1.5329351745
## FC_Fpos_AGTACCAGTTTGGCTA-1FC_FI_GFPve_pos     -1.1534723468  1.9318838440
## FC_Fpos_AGTACTGAGTCCTACA-1FC_FI_GFPve_pos     -1.5005098410  2.1038337074
## FC_Fpos_AGTACTGCACACACTA-1FC_FI_GFPve_pos      0.2050741188  2.1609540306
## FC_Fpos_AGTACTGTCGAACACT-1FC_FI_GFPve_pos     -1.4663094111  2.2017913185
## FC_Fpos_AGTAGCTCACGGCTAC-1FC_FI_GFPve_pos     -0.6721770592  0.6532268844
## FC_Fpos_AGTAGTCGTTCCTTGC-1FC_FI_GFPve_pos     -2.2181293793  1.8498630844
```

```
## FC_Fpos_AGTAGTCGTTGTGCCG-1FC_FI_GFPve_pos     -1.0747520156  2.9369602524
## FC_Fpos_AGTCATGCAGCGTTGC-1FC_FI_GFPve_pos     -1.1981259413  2.1490044914
## FC_Fpos_AGTCTCCCACGGTCTG-1FC_FI_GFPve_pos      0.7142398529  3.3182173096
## FC_Fpos_AGTGACTAGCTAGAAT-1FC_FI_GFPve_pos     -1.4644466467  2.2807321869
## FC_Fpos_AGTGTTGCAGAAGTGC-1FC_FI_GFPve_pos     -0.7018532582  0.5633073173
## FC_Fpos_ATACCGAAGGAAAGGT-1FC_FI_GFPve_pos      0.2628008299  2.7903528534
## FC_Fpos_ATACCTTCAGCGCGTT-1FC_FI_GFPve_pos      1.1826222114  3.2029629074
## FC_Fpos_ATACTTCTCTCACTCG-1FC_FI_GFPve_pos     -0.9124398656  2.0062485061
## FC_Fpos_ATAGACCTCGGAGTGA-1FC_FI_GFPve_pos     -1.3203970022  2.8765535675
## FC_Fpos_ATAGAGAGTATGCTAC-1FC_FI_GFPve_pos      1.0126222544  3.3244924866
## FC_Fpos_ATAGGCTGTTGCATAC-1FC_FI_GFPve_pos     -0.3317656048  3.1747055374
## FC_Fpos_ATCACGATCGGTCAGC-1FC_FI_GFPve_pos     -1.8086325951  2.1680302940
## FC_Fpos_ATCCCTGAGCTGCGAA-1FC_FI_GFPve_pos     -0.2107191287  2.9673467003
## FC_Fpos_ATCCTATAGATGGGCT-1FC_FI_GFPve_pos     -0.1858759873  0.6472316109
## FC_Fpos_ATCGATGAGAGGTATT-1FC_FI_GFPve_pos      0.5415005855  3.3830204330
## FC_Fpos_ATCGATGTCTAGTTCT-1FC_FI_GFPve_pos      0.6280377679  2.8212619148
## FC_Fpos_ATCGCCTCACTAGTAC-1FC_FI_GFPve_pos     -1.1085198946  2.8714971863
## FC_Fpos_ATCGGATAGTACTGGG-1FC_FI_GFPve_pos     -1.9506785698  2.4704685531
## FC_Fpos_ATCGGATGTATCACGT-1FC_FI_GFPve_pos     -0.1133704126  2.7416076980
## FC_Fpos_ATCGGCGTCTGTGCGG-1FC_FI_GFPve_pos     -0.7256805725  1.1099810921
## FC_Fpos_ATCTCTAAGTCAACAA-1FC_FI_GFPve_pos     -0.4241563804  2.2305808388
## FC_Fpos_ATGCGATAGTGATGGC-1FC_FI_GFPve_pos     -0.5816795953  2.8764515243
## FC_Fpos_ATGGGAGTCGGCTGAC-1FC_FI_GFPve_pos     -0.5148673780  2.6526346527
## FC_Fpos_ATGGTTGCACCTGAAT-1FC_FI_GFPve_pos      0.2873924844  1.7527108513
## FC_Fpos_ATTATCCTCGTTCGCT-1FC_FI_GFPve_pos     -0.9752694793  2.0695143066
## FC_Fpos_ATTCATCGTGAGGCAT-1FC_FI_GFPve_pos     -1.9892498083  2.4296202980
## FC_Fpos_ATTCCCGCACATAACC-1FC_FI_GFPve_pos     -0.2777474202  0.6116104446
## FC_Fpos_ATTCCTACATATCGGT-1FC_FI_GFPve_pos      0.6725354962  2.8341927849
## FC_Fpos_ATTCGTTAGAATCGTA-1FC_FI_GFPve_pos     -1.1022232599  0.7230806671
## FC_Fpos_ATTCTTGCAACCGCCA-1FC_FI_GFPve_pos     -0.3161387421  2.1113782249
## FC_Fpos_ATTCTTGTCTTGGTGA-1FC_FI_GFPve_pos     -1.1083458491  2.2202716194
## FC_Fpos_ATTGGGTTCTGTAACG-1FC_FI_GFPve_pos      0.0134745412  2.2453136764
## FC_Fpos_ATTGTTCTCACTACTT-1FC_FI_GFPve_pos     -0.3642185159  2.7020659767
## FC_Fpos_ATTTACCAGGATCATA-1FC_FI_GFPve_pos     -1.0176531501  1.4219622932
## FC_Fpos_CAACAACCAGCGGTCT-1FC_FI_GFPve_pos     -1.0029477902  2.8246507965
## FC_Fpos_CAACAGTGTACGTGTT-1FC_FI_GFPve_pos      0.7670583896  2.2189631782
## FC_Fpos_CAACAGTTCCTTTAGT-1FC_FI_GFPve_pos     -1.2919629641  1.1904754959
## FC_Fpos_CAACCTCGTAACTAAG-1FC_FI_GFPve_pos      0.1249136500  1.5639720283
## FC_Fpos_CAACCTCTCAAACGAA-1FC_FI_GFPve_pos      0.2240096502  1.8705206238
## FC_Fpos_CAAGACTTCAACGCTA-1FC_FI_GFPve_pos     -0.4377856977  1.6953716598
## FC_Fpos_CAAGAGGCAGCTCATA-1FC_FI_GFPve_pos     -1.3338278361  2.9548435531
## FC_Fpos_CAAGGGACACAGCATT-1FC_FI_GFPve_pos     -1.1404732294  3.2600202881
## FC_Fpos_CAAGGGACAGAGGGTT-1FC_FI_GFPve_pos     -1.0828754730  2.5197263084
## FC_Fpos_CAATACGTCCAGTGCG-1FC_FI_GFPve_pos     -1.6290864773  3.2500748955
## FC_Fpos_CAATCGAGTAACTGCT-1FC_FI_GFPve_pos     -0.6697458453  1.8278551422
## FC_Fpos_CAATCGAGTCGCACGT-1FC_FI_GFPve_pos     -1.9238601752  3.2182708107
## FC_Fpos_CACAACATCGCTTAAG-1FC_FI_GFPve_pos     -1.0638415642  0.6218118988
## FC_Fpos_CACACAAGTAGAGGAA-1FC_FI_GFPve_pos     -0.2635329269  3.3959331833
## FC_Fpos_CACATGAAGACGAAGA-1FC_FI_GFPve_pos     -0.6812242694  2.5293164573
## FC_Fpos_CACATGATCCTTATGT-1FC_FI_GFPve_pos      0.5169539027  1.1173596702
## FC_Fpos_CACCAAAGTGTGACCC-1FC_FI_GFPve_pos      0.9493991308  2.9387555443
## FC_Fpos_CACCGTTCATCCTCAC-1FC_FI_GFPve_pos      0.2314616911  1.1537075363
## FC_Fpos_CACGAATAGCCTCAGC-1FC_FI_GFPve_pos     -1.0463222690  3.2892389618
## FC_Fpos_CACGGGTGTACAGTAA-1FC_FI_GFPve_pos     -1.5184459515  3.0750418030
## FC_Fpos_CACGGGTGTCCCGTGA-1FC_FI_GFPve_pos      1.8309154682  2.0500436149
## FC_Fpos_CAGAGCCCACATTACG-1FC_FI_GFPve_pos     -0.4889995284  3.0011520706
## FC_Fpos_CAGCAGCCATCATCTT-1FC_FI_GFPve_pos      1.7268579178  2.6541128479
## FC_Fpos_CAGCCAGTCTAGGCCG-1FC_FI_GFPve_pos     -1.9660519190  3.1005759559
## FC_Fpos_CAGGCCAAGAAACCCG-1FC_FI_GFPve_pos     -0.2686184250  2.8451943718
## FC_Fpos_CAGGCCATCGTACCTC-1FC_FI_GFPve_pos      0.5369729571  2.6245646797
## FC_Fpos_CAGGGCTAGCGTCGAA-1FC_FI_GFPve_pos     -0.7591022082  3.5254860245
## FC_Fpos_CAGTTAGCATGATCTG-1FC_FI_GFPve_pos     -0.3649268873  1.5222726188
## FC_Fpos_CAGTTCCCATTGTCGA-1FC_FI_GFPve_pos     -0.3361269213  2.3526101433
## FC_Fpos_CATACAGGTGAATGAT-1FC_FI_GFPve_pos     -1.3799840040  2.2335086189
## FC_Fpos_CATAGACGTGGCTAGA-1FC_FI_GFPve_pos      0.6476432853  2.5263734184
## FC_Fpos_CATCAAGTCAGAGCGA-1FC_FI_GFPve_pos     -0.4553583987  1.3369422279
## FC_Fpos_CATCCACGTCACTGAT-1FC_FI_GFPve_pos     -0.2261312582  2.8673849426
## FC_Fpos_CATCCACGTCTACACA-1FC_FI_GFPve_pos     -1.9126726217  1.3656349502
## FC_Fpos_CATCCACGTTGAAGTA-1FC_FI_GFPve_pos     -0.4514046140  2.0988135658
## FC_Fpos_CATCCCATCTGGGTCG-1FC_FI_GFPve_pos     -2.0257867403  2.1070294700
```

```
## FC_Fpos_CATCGCTAGGCATTTC-1FC_FI_GFPve_pos      1.7707116775  1.8655166946
## FC_Fpos_CATGCGGTCGTAGGAG-1FC_FI_GFPve_pos      0.6722025447  2.0292930923
## FC_Fpos_CATGGATAGTGGTGGT-1FC_FI_GFPve_pos     -1.7748269863  2.6155844055
## FC_Fpos_CATTCATCAATCTAGC-1FC_FI_GFPve_pos     -1.7573833294  1.6135177933
## FC_Fpos_CATTGAGCACGACTAT-1FC_FI_GFPve_pos     -0.7730758615  1.2205143295
## FC_Fpos_CATTGCCCAGGACTAG-1FC_FI_GFPve_pos     -1.5833643503  2.2746033989
## FC_Fpos_CATTGCCGTGGACTGA-1FC_FI_GFPve_pos      0.9021340780  2.1684708916
## FC_Fpos_CCACACTGTTCTCAGA-1FC_FI_GFPve_pos     -1.6823214121  2.3335795723
## FC_Fpos_CCACGAGGTGACAACG-1FC_FI_GFPve_pos     -2.2830898590 -0.0954532303
## FC_Fpos_CCACTTGCAAGTGGCA-1FC_FI_GFPve_pos     -2.2224612065  2.2392640434
## FC_Fpos_CCACTTGCAGCACAAG-1FC_FI_GFPve_pos     -1.5958952732  2.1654401146
## FC_Fpos_CCATCACAGAGTGACC-1FC_FI_GFPve_pos     -1.6662393637  2.8687753998
## FC_Fpos_CCCATTGGTCAAGCCC-1FC_FI_GFPve_pos     -2.1619157620  0.6097984634
## FC_Fpos_CCCATTGGTCGTGATT-1FC_FI_GFPve_pos     -0.5096466072  2.5000181518
## FC_Fpos_CCCGGAACACGCGCAT-1FC_FI_GFPve_pos      1.7802360229  2.2376265846
## FC_Fpos_CCCGGAAGTCAAGGCA-1FC_FI_GFPve_pos      0.7083524041  2.1055565201
## FC_Fpos_CCCTAACTCCACTTTA-1FC_FI_GFPve_pos      0.8939438038  2.0740785919
## FC_Fpos_CCCTCTCGTAACCAGG-1FC_FI_GFPve_pos     -2.1254598923  2.2109155975
## FC_Fpos_CCGAACGCAAGGACAC-1FC_FI_GFPve_pos     -0.9207913704  2.1268868767
## FC_Fpos_CCGAACGGTCTTGCTC-1FC_FI_GFPve_pos     -1.2713830300  3.3353977524
## FC_Fpos_CCGATCTCAGAGGACT-1FC_FI_GFPve_pos      0.4307491355  2.5985446296
## FC_Fpos_CCGGACAGTCAAGCCC-1FC_FI_GFPve_pos     -1.3569043703  0.6239700638
## FC_Fpos_CCGGGTAAGATGGGCT-1FC_FI_GFPve_pos      0.7372713856  2.9292807899
## FC_Fpos_CCGGTAGGTCTTACTT-1FC_FI_GFPve_pos     -0.6858227678  2.3505969368
## FC_Fpos_CCGTTCATCGTGAGAG-1FC_FI_GFPve_pos     -1.9845690556  2.1089330040
## FC_Fpos_CCGTTCATCTGACGCG-1FC_FI_GFPve_pos      0.2343132369  2.5786590896
## FC_Fpos_CCTACGTGTCTAACTG-1FC_FI_GFPve_pos     -0.7806587644  2.2069416366
## FC_Fpos_CCTATCGGTATGGGAC-1FC_FI_GFPve_pos     -1.3143343754  2.9951963745
## FC_Fpos_CCTCATGAGGCCTAGA-1FC_FI_GFPve_pos     -1.2609755821  2.6332445465
## FC_Fpos_CCTCTAGCAAATGGCG-1FC_FI_GFPve_pos     -1.6963714190  2.5628357254
## FC_Fpos_CCTGTTGTCGTCCATC-1FC_FI_GFPve_pos      0.0876768403  3.1217384659
## FC_Fpos_CCTTTGGAGTATGTAG-1FC_FI_GFPve_pos     -0.7931817599  1.8683128677
## FC_Fpos_CGAAGGAAGATCCCGC-1FC_FI_GFPve_pos     -1.0837278791  3.1512241684
## FC_Fpos_CGAAGGATCGGCTATA-1FC_FI_GFPve_pos     -2.2000312634  2.1681347214
## FC_Fpos_CGAAGTTCATCGATCA-1FC_FI_GFPve_pos     -2.1305173464  1.9846206032
## FC_Fpos_CGACAGCCAGAGCCCT-1FC_FI_GFPve_pos     -0.9514655895  2.3255915962
## FC_Fpos_CGAGGAACATGAGTAA-1FC_FI_GFPve_pos     -1.8045493193  2.8963523231
## FC_Fpos_CGATCGGTCTGTCCGT-1FC_FI_GFPve_pos      0.5058447294  3.4185982071
## FC_Fpos_CGATGCGCAAACTCGT-1FC_FI_GFPve_pos      0.8896583490  2.2122140251
## FC_Fpos_CGCAGGTCAACCGACC-1FC_FI_GFPve_pos     -0.9078318901  2.9295640312
## FC_Fpos_CGCAGGTTCATTTACC-1FC_FI_GFPve_pos     -1.4130186863  1.8663292251
## FC_Fpos_CGCATAACACGCCACA-1FC_FI_GFPve_pos     -2.1133639641  2.1337671600
## FC_Fpos_CGCATAATCTGTCAGA-1FC_FI_GFPve_pos      0.8222563438  2.1843271576
## FC_Fpos_CGCATGGTCGAGTCCG-1FC_FI_GFPve_pos     -2.2778165169  2.0016670547
## FC_Fpos_CGCCAGAAGTCTTCCC-1FC_FI_GFPve_pos     -0.2034385666  2.7763552986
## FC_Fpos_CGCCAGAGTTACCGTA-1FC_FI_GFPve_pos     -1.2004266806  2.5643968902
## FC_Fpos_CGGACACAGCATCCCG-1FC_FI_GFPve_pos     -1.9726383515  2.4754877411
## FC_Fpos_CGGACACCAGTCGGTC-1FC_FI_GFPve_pos     -0.9273539610  2.6333203636
## FC_Fpos_CGGACACTCGGTAAGG-1FC_FI_GFPve_pos      0.5315948896  2.9964528404
## FC_Fpos_CGGCAGTTCCATCGTC-1FC_FI_GFPve_pos      0.1233197026  3.2239485107
## FC_Fpos_CGGGACTGTACTGCCG-1FC_FI_GFPve_pos     -1.2963574953  2.1133699737
## FC_Fpos_CGGGCATAGCTACAAA-1FC_FI_GFPve_pos     -0.3171680547  1.8937139831
## FC_Fpos_CGGGTGTAGCGGACAT-1FC_FI_GFPve_pos     -1.3319083281  2.6917057358
## FC_Fpos_CGTAATGAGGTATTGA-1FC_FI_GFPve_pos     -2.0286774464  1.6734047256
## FC_Fpos_CGTAATGTCGGTCTAA-1FC_FI_GFPve_pos     -0.4256490894  3.5624466263
## FC_Fpos_CGTAGTATCGCAGTTA-1FC_FI_GFPve_pos     -1.0820902415  2.4473843895
## FC_Fpos_CGTCAAACACCTCAGG-1FC_FI_GFPve_pos     -0.9883529968  0.8175149284
## FC_Fpos_CGTCCATAGGTTCTAC-1FC_FI_GFPve_pos     -0.3308109917  2.4814453445
## FC_Fpos_CGTCCATTCCCGTAAA-1FC_FI_GFPve_pos     -1.7998343535  2.9839086853
## FC_Fpos_CGTGAATAGTACAGAT-1FC_FI_GFPve_pos     -0.1773209792  2.8370814644
## FC_Fpos_CGTGAATCAACACGAG-1FC_FI_GFPve_pos      0.0797738604  1.3711643539
## FC_Fpos_CGTGAATCATGGACAG-1FC_FI_GFPve_pos     -0.3639281966  2.3157763801
## FC_Fpos_CGTGCTTGTCTTCTAT-1FC_FI_GFPve_pos      0.0749058001  3.4930658661
## FC_Fpos_CGTGCTTTCTCGTCAC-1FC_FI_GFPve_pos     -1.3474666901  1.4803510032
## FC_Fpos_CGTTCTGTCAGGACGA-1FC_FI_GFPve_pos     -1.6813690491  3.1947169624
## FC_Fpos_CGTTGGGCAATGGGTG-1FC_FI_GFPve_pos     -1.0949777789  2.6216912590
## FC_Fpos_CGTTGGGTCCTTTAGT-1FC_FI_GFPve_pos      1.4639501743  2.8541245781
## FC_Fpos_CTAACCCGTCGTATTG-1FC_FI_GFPve_pos     -1.2896058388  1.1712899528
## FC_Fpos_CTAAGTGAGAGGCGTT-1FC_FI_GFPve_pos      0.0229312800  2.4730806671
## FC_Fpos_CTACATTAGGGTCAAC-1FC_FI_GFPve_pos     -1.1725443430  2.5450325332
```

```
## FC_Fpos_CTACCTGGTTAAGGGC-1FC_FI_GFPve_pos     -0.5236894317  2.8787141166
## FC_Fpos_CTACTATGTGTCTTAG-1FC_FI_GFPve_pos     -1.0412463374  1.2080297790
## FC_Fpos_CTATAGGGTTCAACGT-1FC_FI_GFPve_pos     -0.3488572068  2.5510721527
## FC_Fpos_CTCAATTCAGGTACGA-1FC_FI_GFPve_pos     -2.0584243365  1.6787686668
## FC_Fpos_CTCAGGGCAATAAGGT-1FC_FI_GFPve_pos     -0.1929927688  1.1951337181
## FC_Fpos_CTCAGGGGTGGAACAC-1FC_FI_GFPve_pos     -0.5796700247  1.3552680336
## FC_Fpos_CTCCCAAGTAGGACCA-1FC_FI_GFPve_pos     -1.7179100342  2.5851884208
## FC_Fpos_CTCCCTCCATCCGTGG-1FC_FI_GFPve_pos     -0.3820634611  2.6217727981
## FC_Fpos_CTCCCTCCATTGCTTT-1FC_FI_GFPve_pos     -1.1697088309  1.2977528892
## FC_Fpos_CTCCCTCGTATACCTG-1FC_FI_GFPve_pos      0.9934258632  2.8973894439
## FC_Fpos_CTCCTCCGTGACTGAG-1FC_FI_GFPve_pos     -0.7539915271  2.4822345100
## FC_Fpos_CTCCTTTAGCGTCAAG-1FC_FI_GFPve_pos     -0.7722044416  2.7991919838
## FC_Fpos_CTCTCAGAGGATTCAA-1FC_FI_GFPve_pos     -1.6069599219  2.8777475677
## FC_Fpos_CTCTGGTGTCCTTAAG-1FC_FI_GFPve_pos     -0.1194711007  2.7555594765
## FC_Fpos_CTGAATGAGTCGAGGT-1FC_FI_GFPve_pos     -0.0258893348  2.3519602142
## FC_Fpos_CTGATCCTCTAAGCCA-1FC_FI_GFPve_pos     -1.3338609762  2.3563065849
## FC_Fpos_CTGCAGGGTCACGTGC-1FC_FI_GFPve_pos     -0.5503068395  2.0598874412
## FC_Fpos_CTGCATCAGTAACAGT-1FC_FI_GFPve_pos     -1.6100760527  3.2077489219
## FC_Fpos_CTGGCAGCATCATTTC-1FC_FI_GFPve_pos     -0.1164581530  0.2865982376
## FC_Fpos_CTGGCAGTCCCTCTAG-1FC_FI_GFPve_pos     -0.1375830896  1.2027287803
## FC_Fpos_CTGGTCTGTGGCTAGA-1FC_FI_GFPve_pos     -0.0696063243  2.9205847107
## FC_Fpos_CTGTAGACAGATAAAC-1FC_FI_GFPve_pos     -0.0254075207  2.9754529320
## FC_Fpos_CTGTAGAGTACAAGTA-1FC_FI_GFPve_pos     -0.5688844152  1.2731957756
## FC_Fpos_CTGTGGGTCCCGAGAC-1FC_FI_GFPve_pos     -0.1601620024  2.3189373337
## FC_Fpos_CTTAGGACAAATAAGC-1FC_FI_GFPve_pos      0.8033963613  1.5996241890
## FC_Fpos_CTTCAATGTCATCAGT-1FC_FI_GFPve_pos      1.7610900574  2.0163240753
## FC_Fpos_CTTCGGTTCGCTTGCT-1FC_FI_GFPve_pos      0.1147610717  3.0573950134
## FC_Fpos_CTTCGGTTCGGTCGAC-1FC_FI_GFPve_pos     -1.3883700199  1.3256526314
## FC_Fpos_CTTCGGTTCTGAACGT-1FC_FI_GFPve_pos     -0.9739917345  2.2934127174
## FC_Fpos_CTTCTAATCTCCTGAC-1FC_FI_GFPve_pos     -1.1185455747  2.5057468735
## FC_Fpos_CTTGAGATCAGAGCAG-1FC_FI_GFPve_pos     -0.6042546756  2.7002735458
## FC_Fpos_CTTTCGGCAAGCAGGT-1FC_FI_GFPve_pos     -0.2110832907  3.0914063774
## FC_Fpos_GAAACCTCATCGTTCC-1FC_FI_GFPve_pos     -2.2465431519  1.3056965195
## FC_Fpos_GAAATGATCAATCTTC-1FC_FI_GFPve_pos     -1.7311545439  2.5269313179
## FC_Fpos_GAACACTAGTAGCCAG-1FC_FI_GFPve_pos     -0.9738315172  0.8086452804
## FC_Fpos_GAACACTCAATGGCAG-1FC_FI_GFPve_pos     -0.0858495511  3.2423496567
## FC_Fpos_GAACGTTGTGTCTAAC-1FC_FI_GFPve_pos      0.4912704401  2.6969065986
## FC_Fpos_GAAGCCCTCTACTTCA-1FC_FI_GFPve_pos     -1.2612914868  3.2823505722
## FC_Fpos_GAAGGACGTACCCGAC-1FC_FI_GFPve_pos      0.5008881860  2.1678042732
## FC_Fpos_GAAGGGTAGGTCCGAA-1FC_FI_GFPve_pos     -1.2691204376  2.5164275490
## FC_Fpos_GAAGTAACAACATACC-1FC_FI_GFPve_pos      0.7630322747  2.1001038872
## FC_Fpos_GAAGTAACAGGCACAA-1FC_FI_GFPve_pos     -1.3271158762  1.8033099495
## FC_Fpos_GAATAGATCCTCACTG-1FC_FI_GFPve_pos     -1.7102138825  3.1247530304
## FC_Fpos_GAATCGTAGTCATGAA-1FC_FI_GFPve_pos     -0.2291199580  2.8630557380
## FC_Fpos_GACAGCCGTACCTGTA-1FC_FI_GFPve_pos      0.5493287616  1.5624466263
## FC_Fpos_GACATCACAGCGACAA-1FC_FI_GFPve_pos     -0.3330441691  1.9073710762
## FC_Fpos_GACATCAGTCCTGAAT-1FC_FI_GFPve_pos     -0.8143423624  2.3302893959
## FC_Fpos_GACATCATCTTTACAC-1FC_FI_GFPve_pos     -0.4607555099  2.4581809364
## FC_Fpos_GACCAATGTCGGAAAC-1FC_FI_GFPve_pos      0.9741715364  1.8708482109
## FC_Fpos_GACCCAGAGGTATAGT-1FC_FI_GFPve_pos      0.1438449793  2.8241668068
## FC_Fpos_GACCCTTAGCGCCTAC-1FC_FI_GFPve_pos     -1.1006502338  3.2381291710
## FC_Fpos_GACCGTGAGTGACCTT-1FC_FI_GFPve_pos     -0.7675199933  2.6348762832
## FC_Fpos_GACGTTACACTCCTGT-1FC_FI_GFPve_pos      0.9430678062  2.5623560272
## FC_Fpos_GACGTTATCAACACCA-1FC_FI_GFPve_pos     -0.3920995720  2.0894484840
## FC_Fpos_GACTCAAAGTCGCGAA-1FC_FI_GFPve_pos      0.1972351544  2.4764976822
## FC_Fpos_GACTCAACATAATGAG-1FC_FI_GFPve_pos      0.1684512965  2.5987429939
## FC_Fpos_GACTCTCAGTATGAAC-1FC_FI_GFPve_pos     -0.5916725226  0.6592355095
## FC_Fpos_GAGACCCAGCGAGTAC-1FC_FI_GFPve_pos      0.4492083959  3.1840653740
## FC_Fpos_GAGACTTAGCTGAAGC-1FC_FI_GFPve_pos      0.2515023820  3.4133825622
## FC_Fpos_GAGACTTCACTACGGC-1FC_FI_GFPve_pos      0.2968059950  3.0639129005
## FC_Fpos_GAGCCTGTCGAAACAA-1FC_FI_GFPve_pos     -2.3521590061  1.9538960777
## FC_Fpos_GAGCTGCAGTTGGCTT-1FC_FI_GFPve_pos     -0.5151557453  1.6779489837
## FC_Fpos_GAGGGTACAGCTGTTA-1FC_FI_GFPve_pos     -1.7685172386  1.9634442650
## FC_Fpos_GAGTGTTAGTAGGAAG-1FC_FI_GFPve_pos      0.5612946324  3.4012050949
## FC_Fpos_GAGTGTTCAGTTGTTG-1FC_FI_GFPve_pos     -0.1528055534  1.9422269188
## FC_Fpos_GAGTGTTTCGTAGAGG-1FC_FI_GFPve_pos     -1.1263145037  2.8598656975
## FC_Fpos_GAGTTACGTGGGAGAG-1FC_FI_GFPve_pos     -2.0708601303  1.9630427681
## FC_Fpos_GATAGCTAGTCTAACC-1FC_FI_GFPve_pos     -0.4194800980  1.7241216026
## FC_Fpos_GATCATGTCACACCCT-1FC_FI_GFPve_pos      1.0184751921  1.4086385093
## FC_Fpos_GATCCCTTCTATCGTT-1FC_FI_GFPve_pos      0.0456403814  2.5137048088
```

```
## FC_Fpos_GATGAGGCACGTCGTG-1FC_FI_GFPve_pos     -0.7249345012  1.5788188301
## FC_Fpos_GATGGAGCATAATCCG-1FC_FI_GFPve_pos      0.4319130354  3.0652475677
## FC_Fpos_GATGGAGCATCCCGTT-1FC_FI_GFPve_pos     -0.3700802006  2.3322973572
## FC_Fpos_GATGTTGAGAGAGCGG-1FC_FI_GFPve_pos     -0.5275306530  1.2891321502
## FC_Fpos_GATTCGATCATCGTAG-1FC_FI_GFPve_pos     -0.8659381099  2.5376811348
## FC_Fpos_GATTCGATCCCAATAG-1FC_FI_GFPve_pos      0.6650080852  3.2455954872
## FC_Fpos_GATTTCTTCCTGGCTT-1FC_FI_GFPve_pos     -0.5682645269  2.5579595886
## FC_Fpos_GCAACCGTCGAGAGCA-1FC_FI_GFPve_pos      0.1272174589  2.1654811226
## FC_Fpos_GCAACCGTCTACACAG-1FC_FI_GFPve_pos     -1.1109958120  1.9413147293
## FC_Fpos_GCACGGTCACCGGTCA-1FC_FI_GFPve_pos     -2.0622742005  1.2844605766
## FC_Fpos_GCACGTGCATCACGGC-1FC_FI_GFPve_pos     -0.5741947420  2.8931732498
## FC_Fpos_GCACTAAAGACTACCT-1FC_FI_GFPve_pos     -1.1178608365  2.0884051643
## FC_Fpos_GCAGGCTGTATTGCCA-1FC_FI_GFPve_pos     -1.4069577284  2.3269596420
## FC_Fpos_GCATCGGGTACCTATG-1FC_FI_GFPve_pos     -0.6727604695  3.4777889572
## FC_Fpos_GCATGATTCGACACCG-1FC_FI_GFPve_pos      1.1020343475  0.0716338478
## FC_Fpos_GCCAGCACATGGAACG-1FC_FI_GFPve_pos     -1.9580184050  1.3513803802
## FC_Fpos_GCCAGGTTCACGATAC-1FC_FI_GFPve_pos     -1.3986012764  2.5569176994
## FC_Fpos_GCCGATGTCAAATAGG-1FC_FI_GFPve_pos     -0.2774316572  3.3092823349
## FC_Fpos_GCGGATCCATACGCAT-1FC_FI_GFPve_pos     -0.8138221331  3.1732497536
## FC_Fpos_GCGTGCACAAGGCTTT-1FC_FI_GFPve_pos     -1.4500071831  3.0816703163
## FC_Fpos_GCGTTTCTCGTGCGAC-1FC_FI_GFPve_pos     -0.9130599327  2.9157572113
## FC_Fpos_GCTACAACACGCTATA-1FC_FI_GFPve_pos     -0.2667529188  1.7995272003
## FC_Fpos_GCTACCTCATTGTCGA-1FC_FI_GFPve_pos     -1.6623227425  2.2210188232
## FC_Fpos_GCTCAAACACGTCGGT-1FC_FI_GFPve_pos      1.7318399124  2.3259988151
## FC_Fpos_GCTCAAAGTCTTGCTC-1FC_FI_GFPve_pos     -1.5538970061  2.0703368507
## FC_Fpos_GCTGAATGTAGCTTAC-1FC_FI_GFPve_pos      0.1815644674  2.8037958465
## FC_Fpos_GCTGGGTAGATAGCTA-1FC_FI_GFPve_pos     -0.4301295407  1.8661580406
## FC_Fpos_GCTGGGTCACGTTGGC-1FC_FI_GFPve_pos      0.6369356684  2.6684165321
## FC_Fpos_GCTGGGTGTTACGGAG-1FC_FI_GFPve_pos     -1.4751907416  2.3895464264
## FC_Fpos_GCTGGGTTCTCGGTAA-1FC_FI_GFPve_pos     -1.6642825432  2.7713785492
## FC_Fpos_GCTTTCGGTCCGGATC-1FC_FI_GFPve_pos      0.4197998218  2.4441571556
## FC_Fpos_GGAAGTGAGTTCATGC-1FC_FI_GFPve_pos     -1.2557351418  2.4731855713
## FC_Fpos_GGAATCTAGATAGTGT-1FC_FI_GFPve_pos     -1.3739464112  2.6010465942
## FC_Fpos_GGAGATGTCCGATAAC-1FC_FI_GFPve_pos     -1.6950645275  2.7106767021
## FC_Fpos_GGAGGATGTTCTCAGA-1FC_FI_GFPve_pos      0.4905699067  2.1100120865
## FC_Fpos_GGCACGTTCCACTGAA-1FC_FI_GFPve_pos     -1.3940894432  2.4303574882
## FC_Fpos_GGCAGTCCATGTGGTT-1FC_FI_GFPve_pos     -0.3287369735  3.0417199455
## FC_Fpos_GGCAGTCTCACCGCTT-1FC_FI_GFPve_pos     -0.8937126823  2.3794765793
## FC_Fpos_GGCTTTCAGAAGGCTC-1FC_FI_GFPve_pos      0.0098476730  2.9673648201
## FC_Fpos_GGGACCTTCGTAATGC-1FC_FI_GFPve_pos     -1.9011612959  2.3569827400
## FC_Fpos_GGGACTCCACAAGGTG-1FC_FI_GFPve_pos     -0.7907250352  1.9302339874
## FC_Fpos_GGGAGATTCATTTGTC-1FC_FI_GFPve_pos     -0.9642541237  1.9401941620
## FC_Fpos_GGGATGAAGTATGATG-1FC_FI_GFPve_pos      0.8324257903  1.9721761070
## FC_Fpos_GGGATGACAAGAATAC-1FC_FI_GFPve_pos     -0.7146763034  2.3033662163
## FC_Fpos_GGGCCATCATGTAACC-1FC_FI_GFPve_pos     -0.7631745763  2.2886033378
## FC_Fpos_GGGCCATCATTCTCCG-1FC_FI_GFPve_pos     -0.9422393866  2.3008709274
## FC_Fpos_GGGCTCACATCCGTTC-1FC_FI_GFPve_pos     -1.3327904768  2.1361055694
## FC_Fpos_GGGTATTGTTAGTTCG-1FC_FI_GFPve_pos     -1.3261222668  2.6258044563
## FC_Fpos_GGGTCACGTAGTTACC-1FC_FI_GFPve_pos      0.1483120494  2.1805467926
## FC_Fpos_GGGTGAACACATAGCT-1FC_FI_GFPve_pos     -0.8929658957  2.8823638283
## FC_Fpos_GGGTTATGTGTTTACG-1FC_FI_GFPve_pos     -0.9377933569  1.7749963127
## FC_Fpos_GGGTTATTCCATTCGC-1FC_FI_GFPve_pos      0.0492733888  1.4183206878
## FC_Fpos_GGGTTTAAGCAACTCT-1FC_FI_GFPve_pos     -1.8658791609  2.9354930244
## FC_Fpos_GGGTTTAGTCCTGAAT-1FC_FI_GFPve_pos     -0.1525413493  2.2340584122
## FC_Fpos_GGTAACTAGAGCCTGA-1FC_FI_GFPve_pos     -0.7034094758  2.1487932525
## FC_Fpos_GGTAATCCACCATATG-1FC_FI_GFPve_pos     -0.3480783082  2.3279481254
## FC_Fpos_GGTAATCTCGAGATGG-1FC_FI_GFPve_pos      1.1091731958  2.0507445656
## FC_Fpos_GGTCACGAGTTTCGGT-1FC_FI_GFPve_pos     -2.2354540653  2.8307352386
## FC_Fpos_GGTGAAGTCATAGGCT-1FC_FI_GFPve_pos     -0.8658199139  2.5256076179
## FC_Fpos_GGTGGCTGTAGGACTG-1FC_FI_GFPve_pos      0.6715871267  3.4912472091
## FC_Fpos_GGTGGCTTCATTGCTT-1FC_FI_GFPve_pos      0.2335018389  2.7789893471
## FC_Fpos_GGTGGCTTCCACTAGA-1FC_FI_GFPve_pos     -1.0708256192  1.8013081871
## FC_Fpos_GGTTAACCAGGTCAGA-1FC_FI_GFPve_pos      1.0156995230  1.4870443664
## FC_Fpos_GGTTAACGTAGCGTCC-1FC_FI_GFPve_pos     -1.1002294846  1.1477847420
## FC_Fpos_GGTTGTAAGATGCGAC-1FC_FI_GFPve_pos     -1.3284125157  2.0095677696
## FC_Fpos_GGTTGTAGTGAATGAT-1FC_FI_GFPve_pos     -2.1709203549  1.5234170280
## FC_Fpos_GTAACACAGCTAGAGC-1FC_FI_GFPve_pos     -0.6946993299  3.0807004295
## FC_Fpos_GTAACACCACTGAGTT-1FC_FI_GFPve_pos     -1.3413398094  1.9144354187
## FC_Fpos_GTAAGTCTCATTACGG-1FC_FI_GFPve_pos     -0.3894285835  2.4518351875
## FC_Fpos_GTAAGTCTCTGAGGTT-1FC_FI_GFPve_pos     -0.6617469318  3.0939150177
```

```
## FC_Fpos_GTAATGCAGTACAACA-1FC_FI_GFPve_pos     -1.3678550549  2.5850701652
## FC_Fpos_GTACAACAGTAACAGT-1FC_FI_GFPve_pos     -0.5919829436  2.6108308159
## FC_Fpos_GTACAACTCTCGCTCA-1FC_FI_GFPve_pos     -1.0115703769  2.5970812164
## FC_Fpos_GTACAGTGTGTCCTAA-1FC_FI_GFPve_pos     -1.2447122164  3.0092845283
## FC_Fpos_GTAGAAACATGCAGCC-1FC_FI_GFPve_pos     -0.0301108040  2.5234461151
## FC_Fpos_GTAGAGGTCAACTCTT-1FC_FI_GFPve_pos      0.1940469317  3.0367441498
## FC_Fpos_GTAGCTAAGCAAGTCG-1FC_FI_GFPve_pos     -0.8375639744  3.2688551269
## FC_Fpos_GTAGGTTGTTATCTGG-1FC_FI_GFPve_pos     -0.5301181205  1.0766673408
## FC_Fpos_GTAGTACGTTCGTGCG-1FC_FI_GFPve_pos     -1.6548067160  1.7816858612
## FC_Fpos_GTATTTCTCGTAGAGG-1FC_FI_GFPve_pos      1.3852933817  2.7030825935
## FC_Fpos_GTCAAACTCATATGGC-1FC_FI_GFPve_pos     -1.9202520676  1.8826933227
## FC_Fpos_GTCAAGTAGCCGAATG-1FC_FI_GFPve_pos      0.2972099356  1.1664018951
## FC_Fpos_GTCACGGAGCAATTCC-1FC_FI_GFPve_pos     -1.4843263455  2.5037575088
## FC_Fpos_GTCACGGCATGACTCA-1FC_FI_GFPve_pos      1.1141560249  2.3278427444
## FC_Fpos_GTCACTCAGTCAGCCC-1FC_FI_GFPve_pos     -0.6423557527  1.0295620285
## FC_Fpos_GTCATCCAGCTGACAG-1FC_FI_GFPve_pos     -1.1451901026  3.1927347503
## FC_Fpos_GTCATTTGTTACGGAG-1FC_FI_GFPve_pos     -2.1719136067  2.5138759933
## FC_Fpos_GTCCTCACATGACTGT-1FC_FI_GFPve_pos      0.8910615616  3.3924112640
## FC_Fpos_GTCGAATCATTAAAGG-1FC_FI_GFPve_pos     -1.5319465227  1.8490104996
## FC_Fpos_GTCGCGAGTATGATCC-1FC_FI_GFPve_pos     -1.2349221535  1.7486358009
## FC_Fpos_GTCGTAATCCAGCCTT-1FC_FI_GFPve_pos     -1.5205410547  2.2462797485
## FC_Fpos_GTCTACCAGGTCCCGT-1FC_FI_GFPve_pos      1.8387081795  2.7639918648
## FC_Fpos_GTCTCACGTAGAATGT-1FC_FI_GFPve_pos     -0.4917023606  2.5988083206
## FC_Fpos_GTGACGCCAACGGGTA-1FC_FI_GFPve_pos      0.5681729488  0.8380117737
## FC_Fpos_GTGACGCCACTGGATT-1FC_FI_GFPve_pos     -1.0488485522  3.1670127235
## FC_Fpos_GTGAGGATCTCAACGA-1FC_FI_GFPve_pos     -0.8172392674  2.8841557823
## FC_Fpos_GTGAGTTCAAATCGTC-1FC_FI_GFPve_pos     -0.9813389607  3.0008597694
## FC_Fpos_GTGCACGTCAAGCCGC-1FC_FI_GFPve_pos     -0.9164442249  3.1710768066
## FC_Fpos_GTGCACGTCTGCGATA-1FC_FI_GFPve_pos     -0.4121888854  1.7452025734
## FC_Fpos_GTGCAGCAGGAAAGGT-1FC_FI_GFPve_pos     -1.4971235819  2.0300126396
## FC_Fpos_GTGCGTGGTTCTTGTT-1FC_FI_GFPve_pos     -1.3362207241  2.4789924942
## FC_Fpos_GTGCTGGCATTCAGCA-1FC_FI_GFPve_pos     -1.4483413525  2.6004553161
## FC_Fpos_GTGCTGGGTAGCTTTG-1FC_FI_GFPve_pos     -1.8605670757  1.8978648506
## FC_Fpos_GTGCTTCCACTAGTAC-1FC_FI_GFPve_pos     -2.0089392491  2.3263297401
## FC_Fpos_GTGCTTCTCTGTGCAA-1FC_FI_GFPve_pos     -1.1346837707  1.2413692795
## FC_Fpos_GTGGGAACACAGAGAC-1FC_FI_GFPve_pos     -1.4947436876  2.2962446533
## FC_Fpos_GTGGGAAGTGATACAA-1FC_FI_GFPve_pos      1.1483956747  2.6396580063
## FC_Fpos_GTGGGAATCTAGCCAA-1FC_FI_GFPve_pos      1.2412299089 -1.6331531681
## FC_Fpos_GTGGTTAAGCAGCCTC-1FC_FI_GFPve_pos     -0.5195275374  1.3905153595
## FC_Fpos_GTGTTCCTCATCACAG-1FC_FI_GFPve_pos     -0.5109177954  2.0310492836
## FC_Fpos_GTTACAGGTCGCTGCA-1FC_FI_GFPve_pos     -1.3126711674  2.6099090896
## FC_Fpos_GTTACAGGTGTTACAC-1FC_FI_GFPve_pos     -1.2337474652  2.1247668587
## FC_Fpos_GTTACGATCATAAGGA-1FC_FI_GFPve_pos     -1.6755069323  1.6559949241
## FC_Fpos_GTTAGACGTATCGAGG-1FC_FI_GFPve_pos      1.1521170311  1.9096541725
## FC_Fpos_GTTAGTGAGTAGTCTC-1FC_FI_GFPve_pos     -1.2703654595  2.8741464935
## FC_Fpos_GTTATGGAGGTACTGG-1FC_FI_GFPve_pos     -0.9767474361  3.4747686706
## FC_Fpos_GTTATGGTCAAATGAG-1FC_FI_GFPve_pos     -1.9789717026  1.6583638511
## FC_Fpos_GTTCATTAGGTACAGC-1FC_FI_GFPve_pos     -0.5265655048  3.1270623527
## FC_Fpos_GTTCATTGTCTCTCTG-1FC_FI_GFPve_pos     -0.9039147325  0.6166296325
## FC_Fpos_GTTCCGTCAGAGAATT-1FC_FI_GFPve_pos     -0.9371066518  1.4353561722
## FC_Fpos_GTTCGCTGTTTCGTAG-1FC_FI_GFPve_pos      0.6901235156  1.8737354599
## FC_Fpos_GTTCTATGTATGGGAC-1FC_FI_GFPve_pos     -1.6592866011  2.0512075744
## FC_Fpos_GTTCTATGTTACCCAA-1FC_FI_GFPve_pos      0.0627450071  2.9263058029
## FC_Fpos_GTTGCGGTCAGTGATC-1FC_FI_GFPve_pos     -1.0678631492  2.1501040779
## FC_Fpos_GTTGCTCCATCAGCAT-1FC_FI_GFPve_pos     -0.6002115138  1.9990120254
## FC_Fpos_GTTGTAGGTTGGGATG-1FC_FI_GFPve_pos     -1.4960630961  2.9628620468
## FC_Fpos_GTTGTCCAGAACAGGA-1FC_FI_GFPve_pos     -0.2742220215  1.9618749939
## FC_Fpos_GTTGTCCCATGGCGCT-1FC_FI_GFPve_pos     -1.3765739031  1.5385070167
## FC_Fpos_GTTGTGAAGGTGATAT-1FC_FI_GFPve_pos     -0.6203862257  1.7048731170
## FC_Fpos_GTTGTGAAGTAGCAAT-1FC_FI_GFPve_pos      1.0471730404  2.3175235115
## FC_Fpos_GTTGTGACAAGAGTTA-1FC_FI_GFPve_pos     -1.3927047081  2.3481374107
## FC_Fpos_GTTTACTGTTGCATGT-1FC_FI_GFPve_pos      0.0730141275  3.0619173370
## FC_Fpos_TAACACGGTATGAAAC-1FC_FI_GFPve_pos      0.2806924753  2.6726169906
## FC_Fpos_TAACCAGCAGCGTACC-1FC_FI_GFPve_pos     -1.1891764231  1.6964116417
## FC_Fpos_TAACGACTCGACGCTG-1FC_FI_GFPve_pos     -1.0237484999  2.2730040870
## FC_Fpos_TAACTTCAGCATGGGT-1FC_FI_GFPve_pos     -0.3709223546  3.3608217560
## FC_Fpos_TAAGCCAAGCGACTTT-1FC_FI_GFPve_pos     -1.6794004269  2.6935186706
## FC_Fpos_TAATTCCAGCACTCTA-1FC_FI_GFPve_pos     -0.3025683440  2.7575497947
## FC_Fpos_TAATTCCAGCTCCGAC-1FC_FI_GFPve_pos     -0.1326425686 -0.2297386803
## FC_Fpos_TACAGGTAGTAATTGG-1FC_FI_GFPve_pos     -0.6865012474  3.2545028053
```

```
## FC_Fpos_TACAGGTTCACTGGGC-1FC_FI_GFPve_pos     -0.9576706715  3.2129216514
## FC_Fpos_TACATTCGTAGTGATA-1FC_FI_GFPve_pos     -1.7742135353  3.0556817375
## FC_Fpos_TACCGAAGTGATATAG-1FC_FI_GFPve_pos     -1.2684323616  2.0300655685
## FC_Fpos_TACCGGGAGACATAGT-1FC_FI_GFPve_pos     -1.2227269240  3.2662725769
## FC_Fpos_TACCGGGGTGTGTTTG-1FC_FI_GFPve_pos     -1.4205901451  3.0170717560
## FC_Fpos_TACCTGCGTGTAGCAG-1FC_FI_GFPve_pos      0.4983564787  2.0091991745
## FC_Fpos_TACCTGCGTTTACTGG-1FC_FI_GFPve_pos     -1.2058554716  2.5963511787
## FC_Fpos_TACGCTCTCATTCCTA-1FC_FI_GFPve_pos      0.1589976423  2.5519624077
## FC_Fpos_TACGCTCTCCTGGCTT-1FC_FI_GFPve_pos      0.1239128880  1.8394222580
## FC_Fpos_TACTGCCAGTCCCGGT-1FC_FI_GFPve_pos     -0.3899464883  2.1817756019
## FC_Fpos_TACTTACGTAGCCCTG-1FC_FI_GFPve_pos      0.1642989688  1.2698135696
## FC_Fpos_TACTTCAAGGTGATCG-1FC_FI_GFPve_pos      0.0105282538  1.9936824165
## FC_Fpos_TAGACTGTCTTAGTTC-1FC_FI_GFPve_pos     -1.7439398594  2.1549406372
## FC_Fpos_TAGAGTCAGTTACGGG-1FC_FI_GFPve_pos     -1.3267885275  2.6407885872
## FC_Fpos_TAGGTACGTCGCGGTT-1FC_FI_GFPve_pos     -0.9616263337  2.6764617286
## FC_Fpos_TAGTGCATCGCAATTG-1FC_FI_GFPve_pos     -0.9507402606  0.2336621605
## FC_Fpos_TATACCTGTCAAGCCC-1FC_FI_GFPve_pos     -0.0135515459  0.5176730476
## FC_Fpos_TATCAGGCATAGCACT-1FC_FI_GFPve_pos      0.8397665195  2.6953521095
## FC_Fpos_TATCTTGTCCTAAGTG-1FC_FI_GFPve_pos     -0.5895314343  2.5364246689
## FC_Fpos_TATTCCAAGGTTTGAA-1FC_FI_GFPve_pos      0.0851913564  3.1506014190
## FC_Fpos_TATTCCACATGGAACG-1FC_FI_GFPve_pos     -1.2703591414  2.8216395698
## FC_Fpos_TATTGCTCAAAGCACG-1FC_FI_GFPve_pos     -0.1414015718  2.4750948273
## FC_Fpos_TATTTCGAGCAATTCC-1FC_FI_GFPve_pos     -1.3008235522  1.8691721283
## FC_Fpos_TATTTCGGTCTTGAGT-1FC_FI_GFPve_pos     -1.1201590128  2.0303173385
## FC_Fpos_TCAAGCATCGTCGCTT-1FC_FI_GFPve_pos     -1.2341586180  2.4076152168
## FC_Fpos_TCAAGTGCACGCGTCA-1FC_FI_GFPve_pos     -0.9560754724  3.3837747894
## FC_Fpos_TCAATCTGTGGATTTC-1FC_FI_GFPve_pos     -2.2420360871  2.1166181885
## FC_Fpos_TCACGGGAGGTACTGG-1FC_FI_GFPve_pos     -0.8408647604  3.3539629303
## FC_Fpos_TCACGGGGTTGATGTC-1FC_FI_GFPve_pos     -2.1666379996  1.7387657486
## FC_Fpos_TCAGGTAGTGCCGTAC-1FC_FI_GFPve_pos     -1.6145983763  1.4732361160
## FC_Fpos_TCAGGTATCTATTCGT-1FC_FI_GFPve_pos     -0.8166208692  3.0273080192
## FC_Fpos_TCAGTGAGTGCCGTAC-1FC_FI_GFPve_pos     -0.0994961239  2.3959679924
## FC_Fpos_TCAGTTTTCTGTCGCT-1FC_FI_GFPve_pos     -0.4118049718  1.7503133140
## FC_Fpos_TCATCATCACTTGGCG-1FC_FI_GFPve_pos     -0.4293308385  3.2539992653
## FC_Fpos_TCATGAGGTTTGCAGT-1FC_FI_GFPve_pos     -0.4704849549  2.3707809768
## FC_Fpos_TCATGCCAGCGTCTGC-1FC_FI_GFPve_pos     -0.6211182900  3.2851562820
## FC_Fpos_TCATGGAAGCCGATTT-1FC_FI_GFPve_pos     -1.6641393728  2.1169948898
## FC_Fpos_TCATGTTCACATATCG-1FC_FI_GFPve_pos     -1.7770926781  1.5311923347
## FC_Fpos_TCATGTTGTAAGCTCT-1FC_FI_GFPve_pos     -0.9356584974  3.0117206894
## FC_Fpos_TCATGTTGTACGATCT-1FC_FI_GFPve_pos     -0.2194041401  1.8305902801
## FC_Fpos_TCATTACAGGCATGCA-1FC_FI_GFPve_pos     -0.1935877852  2.0530977569
## FC_Fpos_TCATTACCATCTTCGC-1FC_FI_GFPve_pos     -1.4139911957  2.2528095566
## FC_Fpos_TCATTGTTCTAGCAAC-1FC_FI_GFPve_pos     -0.2233779482  1.1222730003
## FC_Fpos_TCCACCACAGAAATTG-1FC_FI_GFPve_pos      0.7849422269  2.3046331726
## FC_Fpos_TCCACCATCTTACTGT-1FC_FI_GFPve_pos     -1.3873752184  1.7456384025
## FC_Fpos_TCCAGAACAACGCCCA-1FC_FI_GFPve_pos     -1.0363274045  2.7341561638
## FC_Fpos_TCCATCGGTAAGATCA-1FC_FI_GFPve_pos     -1.3295682497  2.0906954132
## FC_Fpos_TCCCATGCACTTGTCC-1FC_FI_GFPve_pos     -1.3660857506  2.2377567611
## FC_Fpos_TCCGAAAGTGGAGGTT-1FC_FI_GFPve_pos     -1.2366417475  2.8386383377
## FC_Fpos_TCCGATCAGGGTTAGC-1FC_FI_GFPve_pos     -1.5536435671  2.9374995552
## FC_Fpos_TCCTAATAGACTCGAG-1FC_FI_GFPve_pos     -0.9479659028  2.8102522217
## FC_Fpos_TCCTCCCAGTTGCGAG-1FC_FI_GFPve_pos     -0.6618384547  1.7695389114
## FC_Fpos_TCCTCGAAGATTGAGT-1FC_FI_GFPve_pos     -0.4788356848  2.5153742157
## FC_Fpos_TCCTCTTAGCTCCACG-1FC_FI_GFPve_pos      0.0724505060  1.9651236854
## FC_Fpos_TCCTCTTTCGAGTACT-1FC_FI_GFPve_pos     -1.4071304627  2.3427610718
## FC_Fpos_TCCTGCAAGCTCCACG-1FC_FI_GFPve_pos     -1.4211982317  2.8105779014
## FC_Fpos_TCCTGCAGTGGCTAGA-1FC_FI_GFPve_pos     -0.7091719337  2.7423086487
## FC_Fpos_TCCTGCATCCATCCGT-1FC_FI_GFPve_pos     -0.4638207681  2.5611138664
## FC_Fpos_TCCTTCTCAGGAATCG-1FC_FI_GFPve_pos     -1.1115061469  2.9389787041
## FC_Fpos_TCCTTTCAGCTCATAC-1FC_FI_GFPve_pos      1.0734967165  2.3199529968
## FC_Fpos_TCCTTTCCAGTTTGGT-1FC_FI_GFPve_pos     -0.0163993813  1.7319164596
## FC_Fpos_TCGAACAAGGTAAGTT-1FC_FI_GFPve_pos     -2.1723115273  2.9371571861
## FC_Fpos_TCGACCTTCAGGTAAA-1FC_FI_GFPve_pos     -1.7305601664  1.4691024147
## FC_Fpos_TCGCACTAGGCCGCTT-1FC_FI_GFPve_pos     -1.6624684162  0.2537045799
## FC_Fpos_TCGCACTAGGCGTTAG-1FC_FI_GFPve_pos      0.6132229738  3.0313387237
## FC_Fpos_TCGCACTAGGGCTTCC-1FC_FI_GFPve_pos     -1.0167987175  2.1038613640
## FC_Fpos_TCGCACTGTACACGCC-1FC_FI_GFPve_pos     -1.2868493624  2.6527648292
## FC_Fpos_TCGGATAAGTAGGGTC-1FC_FI_GFPve_pos     -2.0525302716  2.5528607689
## FC_Fpos_TCGGATAGTTCCAGGC-1FC_FI_GFPve_pos     -0.1194069289  2.1739769302
## FC_Fpos_TCGGGCAAGAGCCTGA-1FC_FI_GFPve_pos      1.2201899462  3.5136080108
```

```
## FC_Fpos_TCGGTCTCATGTGCTA-1FC_FI_GFPve_pos     -1.2730351515  1.9358644806
## FC_Fpos_TCGTGCTAGACTCTTG-1FC_FI_GFPve_pos      0.4866174631  0.2684779487
## FC_Fpos_TCGTGCTCAAGGACAC-1FC_FI_GFPve_pos     -0.3956583865  2.8074117027
## FC_Fpos_TCGTGCTCACGGTAGA-1FC_FI_GFPve_pos     -0.2661062531  1.1261849724
## FC_Fpos_TCGTGGGAGTAGCAAT-1FC_FI_GFPve_pos     -0.5321460791  2.9987678848
## FC_Fpos_TCTAACTCATAGAGGC-1FC_FI_GFPve_pos     -0.4697570033  3.4058752380
## FC_Fpos_TCTAACTGTTATTCTC-1FC_FI_GFPve_pos      0.8829661779  2.2253003441
## FC_Fpos_TCTATCACAGAGTTCT-1FC_FI_GFPve_pos      1.9597675972  1.8730807625
## FC_Fpos_TCTCACGGTGTTCCTC-1FC_FI_GFPve_pos      0.1557441823  1.4350567184
## FC_Fpos_TCTCACGTCCTACTGC-1FC_FI_GFPve_pos     -1.0350326724  2.0083017669
## FC_Fpos_TCTCACGTCTCGTCAC-1FC_FI_GFPve_pos     -1.6982859202  2.1905184112
## FC_Fpos_TCTCAGCTCCGCAACG-1FC_FI_GFPve_pos     -0.9174986906  2.0576873146
## FC_Fpos_TCTCCGAAGTGCCCGT-1FC_FI_GFPve_pos      1.2863986664  3.2756071411
## FC_Fpos_TCTTAGTGTACCAATC-1FC_FI_GFPve_pos     -1.5065393276  2.8907904945
## FC_Fpos_TCTTAGTGTAGACACG-1FC_FI_GFPve_pos     -1.6934721298  3.1248760544
## FC_Fpos_TCTTCCTCATCTCAAG-1FC_FI_GFPve_pos     -1.0607471891  2.1440725647
## FC_Fpos_TCTTGCGGTAGGAAAG-1FC_FI_GFPve_pos     -0.5055653281  0.1010155998
## FC_Fpos_TCTTTGAAGTGCACAG-1FC_FI_GFPve_pos     -0.1095416225  2.9273944221
## FC_Fpos_TGAACGTCAGAAGCTG-1FC_FI_GFPve_pos      0.3604800277  2.9553628288
## FC_Fpos_TGAATCGCAATACGCT-1FC_FI_GFPve_pos      1.1074090175  1.5537348114
## FC_Fpos_TGAATCGTCTAAGAAG-1FC_FI_GFPve_pos     -0.8739036150  1.5931558929
## FC_Fpos_TGACAGTTCAACTCTT-1FC_FI_GFPve_pos      0.3474169545  2.6599893890
## FC_Fpos_TGACTCCCAGCTTCGG-1FC_FI_GFPve_pos     -1.5623698063  1.9758735023
## FC_Fpos_TGAGCGCGTTTGGGAG-1FC_FI_GFPve_pos     -0.4626692958  2.7351279579
## FC_Fpos_TGAGGAGAGCCTGAGA-1FC_FI_GFPve_pos      0.3836071067  2.5416050278
## FC_Fpos_TGAGGAGTCAAGTAAG-1FC_FI_GFPve_pos     -1.3073721953 -0.3552493729
## FC_Fpos_TGAGGTTCAGTAGAAT-1FC_FI_GFPve_pos     -0.7838397451  2.1159844719
## FC_Fpos_TGAGGTTGTCAGTTTG-1FC_FI_GFPve_pos     -0.9251834579  2.2720146499
## FC_Fpos_TGAGTCAAGAGCACTG-1FC_FI_GFPve_pos     -0.6805746980  1.2434282623
## FC_Fpos_TGATCTTGTCTAACGT-1FC_FI_GFPve_pos     -0.6337869294  3.3176632248
## FC_Fpos_TGATGCAGTTTAGTCG-1FC_FI_GFPve_pos     -1.4524206944  2.4756398521
## FC_Fpos_TGATTCTCAAAGTATG-1FC_FI_GFPve_pos      0.8709360533  2.5920367561
## FC_Fpos_TGATTCTGTCATAACC-1FC_FI_GFPve_pos     -1.3773581810  2.1458535515
## FC_Fpos_TGATTTCCAGTAACGG-1FC_FI_GFPve_pos     -1.5471198387  0.2844887100
## FC_Fpos_TGCAGATAGACCTCCG-1FC_FI_GFPve_pos     -1.2272279091  1.9797849975
## FC_Fpos_TGCAGATTCTCTTCAA-1FC_FI_GFPve_pos     -0.7222086020  3.1614723526
## FC_Fpos_TGCGATAAGTGCTCGC-1FC_FI_GFPve_pos     -1.0296931095  3.3553123794
## FC_Fpos_TGCGGGTCAATTGAAG-1FC_FI_GFPve_pos     -0.4805234797  3.0855732284
## FC_Fpos_TGCTCCAAGGTTCCGC-1FC_FI_GFPve_pos     -1.2154216595  2.7033362709
## FC_Fpos_TGCTCCACAGTGACCC-1FC_FI_GFPve_pos      1.0689294510  1.7456608139
## FC_Fpos_TGCTCGTGTCTCGACG-1FC_FI_GFPve_pos      0.1654544048  3.5047016464
## FC_Fpos_TGCTTGCGTAGATTAG-1FC_FI_GFPve_pos     -0.7193333931  2.7560401283
## FC_Fpos_TGGCGTGTCGAGTGGA-1FC_FI_GFPve_pos      1.1370570831  3.3199806534
## FC_Fpos_TGGGAAGTCCGGCTTT-1FC_FI_GFPve_pos     -1.1419602461  2.4765987716
## FC_Fpos_TGGGCGTGTACAGAGC-1FC_FI_GFPve_pos      0.9177222423  2.9310117088
## FC_Fpos_TGGTGATCACGCCACA-1FC_FI_GFPve_pos     -1.2597528525  2.2005248390
## FC_Fpos_TGGTGATTCACACCCT-1FC_FI_GFPve_pos     -0.1436292637  2.0021257721
## FC_Fpos_TGGTTAGCAAGCGAGT-1FC_FI_GFPve_pos      0.1607335381  3.0049748741
## FC_Fpos_TGTCAGACAGCAGTTT-1FC_FI_GFPve_pos     -0.3077517278  2.1100750290
## FC_Fpos_TGTGCGGTCACATTGG-1FC_FI_GFPve_pos     -1.2682231493  1.7651152931
## FC_Fpos_TGTGGCGAGATTGCGG-1FC_FI_GFPve_pos     -1.4721333809  3.1980829559
## FC_Fpos_TGTGTGACATTACTCT-1FC_FI_GFPve_pos     -0.4782765038  2.5748458229
## FC_Fpos_TGTGTGAGTGACGTCC-1FC_FI_GFPve_pos     -0.3239531644  2.9210353218
## FC_Fpos_TGTTCCGTCCAATCTT-1FC_FI_GFPve_pos     -1.0458862610  1.4906797729
## FC_Fpos_TGTTCTACAGTTACCA-1FC_FI_GFPve_pos     -1.0659486480  1.8717432342
## FC_Fpos_TGTTCTATCTATACTC-1FC_FI_GFPve_pos     -1.0112678833  3.0121178947
## FC_Fpos_TGTTGAGCAAGTGCTT-1FC_FI_GFPve_pos      0.2436550133  2.2593064628
## FC_Fpos_TGTTGGACATGACGTT-1FC_FI_GFPve_pos     -0.3329936094  2.9671736084
## FC_Fpos_TGTTTGTCACCTCAGG-1FC_FI_GFPve_pos     -0.2102103509  3.1046681724
## FC_Fpos_TTACCATTCAATGCAC-1FC_FI_GFPve_pos      0.2464474909  2.3265133224
## FC_Fpos_TTACCGCCATTCCTAT-1FC_FI_GFPve_pos     -1.2070191927  1.9822554909
## FC_Fpos_TTACGCCCATTGCAAC-1FC_FI_GFPve_pos     -2.3026838131  2.2537226997
## FC_Fpos_TTACGTTTCCGAGTGC-1FC_FI_GFPve_pos     -0.6071311064  2.4326873146
## FC_Fpos_TTAGGCAGTCGTCAGC-1FC_FI_GFPve_pos     -1.6522234507  2.0703425728
## FC_Fpos_TTAGGGTCAAGTCGTT-1FC_FI_GFPve_pos     -0.2890252225  2.7749257408
## FC_Fpos_TTAGTCTCAGTCCGTG-1FC_FI_GFPve_pos     -2.0570384093  2.2236280762
## FC_Fpos_TTAGTCTGTCAAAGTA-1FC_FI_GFPve_pos      0.3147806399  1.4705663048
## FC_Fpos_TTCACGCAGGCCTAGA-1FC_FI_GFPve_pos      1.6302777700  1.8939314209
## FC_Fpos_TTCACGCTCCACGAAT-1FC_FI_GFPve_pos      0.7437845640  1.7127075516
## FC_Fpos_TTCCAATAGTAGGCCA-1FC_FI_GFPve_pos     -1.1077080198  2.3898082100
```

```
## FC_Fpos_TTCCACGAGCCATTGT-1FC_FI_GFPve_pos      1.4474637680  2.5758767448
## FC_Fpos_TTCCACGAGGCGTTAG-1FC_FI_GFPve_pos     -1.4630539246  2.4307375274
## FC_Fpos_TTCCACGAGTGCGTCC-1FC_FI_GFPve_pos     -1.4382785387  3.1638007484
## FC_Fpos_TTCCACGCATCGGAAG-1FC_FI_GFPve_pos     -1.2718740530  2.4607191406
## FC_Fpos_TTCCTAAGTTCACGAT-1FC_FI_GFPve_pos      1.6915110521  1.5415506683
## FC_Fpos_TTCCTAAGTTGATGTC-1FC_FI_GFPve_pos     -0.2954165019  2.7969818435
## FC_Fpos_TTCCTCTTCAAGAATG-1FC_FI_GFPve_pos      0.8704961710  2.8764324508
## FC_Fpos_TTCCTTCAGACCTCCG-1FC_FI_GFPve_pos      1.2250710897  2.2451424919
## FC_Fpos_TTCCTTCGTGAGTAGC-1FC_FI_GFPve_pos      0.5832659178  2.5098653160
## FC_Fpos_TTCTCTCAGTACTGGG-1FC_FI_GFPve_pos     -1.0314580031  1.4566688858
## FC_Fpos_TTCTGTAAGCCTTTCC-1FC_FI_GFPve_pos      0.6646216087  1.9727359138
## FC_Fpos_TTCTGTAGTCTTACAG-1FC_FI_GFPve_pos     -0.5465183683  3.1254072510
## FC_Fpos_TTGAGTGCATCCTCAC-1FC_FI_GFPve_pos     -0.7168445416  3.0550747238
## FC_Fpos_TTGATGGTCCCGTAAA-1FC_FI_GFPve_pos     -1.9068449564  2.5046339355
## FC_Fpos_TTGCGTCCAGACGGAT-1FC_FI_GFPve_pos      1.7975512914  1.3098831497
## FC_Fpos_TTGGATGAGATAGCAT-1FC_FI_GFPve_pos     -0.3233338572  2.9207721077
## FC_Fpos_TTGGATGGTCCAGAAG-1FC_FI_GFPve_pos     -1.7361574002  3.1738314949
## FC_Fpos_TTGGGTAGTCTGCCTT-1FC_FI_GFPve_pos     -1.2217616863  2.6669917427
## FC_Fpos_TTGGGTAGTTACAGCT-1FC_FI_GFPve_pos     -1.2274601288  1.5247202240
## FC_Fpos_TTGGTTTTCCACGTAA-1FC_FI_GFPve_pos     -1.4997750349  1.1689906441
## FC_Fpos_TTGTGTTAGAATCCCT-1FC_FI_GFPve_pos      0.9428420238  3.3572607361
## FC_Fpos_TTGTTCAAGGATTTAG-1FC_FI_GFPve_pos      0.8716551237  1.6164641701
## FC_Fpos_TTGTTGTAGAATTCAG-1FC_FI_GFPve_pos     -0.6302582748  1.4606829010
## FC_Fpos_TTGTTTGAGACATATG-1FC_FI_GFPve_pos     -0.2990577944  2.8125896774
## FC_Fpos_TTTACCACATTCGGGC-1FC_FI_GFPve_pos     -0.2125238977  1.1349158607
## FC_Fpos_TTTACTGGTGGAGGTT-1FC_FI_GFPve_pos     -0.7852479644  2.2138715111
## FC_Fpos_TTTATGCCACCGAATT-1FC_FI_GFPve_pos      0.6132115893  2.8924436889
## FC_Fpos_TTTCACAAGTCATACC-1FC_FI_GFPve_pos      0.2121519022  3.2567296348
## FC_Fpos_TTTCATGCAGGCAATG-1FC_FI_GFPve_pos     -0.8243380614  1.5787086807
## FC_Fpos_TTTGGAGCAAGAGGTC-1FC_FI_GFPve_pos     -0.9539805479  1.3129411064
## FC_Fpos_TTTGGAGCACGCGCAT-1FC_FI_GFPve_pos      0.1006764941  2.6951432548
## FC_Fpos_TTTGGTTCAGATCCAT-1FC_FI_GFPve_pos     -1.7589030094  3.2181158386
## dCA2_Mpos_AAACCCATCTGAGGCC-1dCA2_MI_GFPve_pos  3.4907934837  0.1956572853
## dCA2_Mpos_AAACGAAGTCCAAAGG-1dCA2_MI_GFPve_pos  3.2473638229  0.3684335075
## dCA2_Mpos_AAAGGATAGCAACTCT-1dCA2_MI_GFPve_pos  2.2646272354  2.9275861107
## dCA2_Mpos_AAAGGATCAATGTCTG-1dCA2_MI_GFPve_pos  2.0207953625  1.9306459747
## dCA2_Mpos_AAAGGGCCAGCTACTA-1dCA2_MI_GFPve_pos  2.7314570122 -1.3682591595
## dCA2_Mpos_AAAGGTAAGTTGAATG-1dCA2_MI_GFPve_pos  2.3877413444  2.2073932014
## dCA2_Mpos_AAAGGTACAGAGTAAT-1dCA2_MI_GFPve_pos  2.5807016067  2.8297620140
## dCA2_Mpos_AAAGGTATCTTGCGCT-1dCA2_MI_GFPve_pos  3.2662782841 -0.5540895142
## dCA2_Mpos_AAAGTCCTCGAGTCTA-1dCA2_MI_GFPve_pos  2.3134777717 -0.9550542511
## dCA2_Mpos_AAATGGAGTTAATCGC-1dCA2_MI_GFPve_pos  2.6102373771 -0.8054580368
## dCA2_Mpos_AACAAAGCACGCGCTA-1dCA2_MI_GFPve_pos  1.1328180008 -0.6499037422
## dCA2_Mpos_AACAAAGTCACTTGGA-1dCA2_MI_GFPve_pos  2.6163733177 -0.7573930897
## dCA2_Mpos_AACAAGACATATCTGG-1dCA2_MI_GFPve_pos  2.6898191147  1.3066521011
## dCA2_Mpos_AACAGGGCAGCGATTT-1dCA2_MI_GFPve_pos  1.8347885780  0.4090032898
## dCA2_Mpos_AACAGGGGTTTCCAAG-1dCA2_MI_GFPve_pos  2.8575382404 -0.7065801300
## dCA2_Mpos_AACCAACCACGACGCT-1dCA2_MI_GFPve_pos  2.3570256405  0.3072138153
## dCA2_Mpos_AACCACACACATCCCT-1dCA2_MI_GFPve_pos  2.2765057258  0.7521944366
## dCA2_Mpos_AACCTGACACCCTTGT-1dCA2_MI_GFPve_pos  2.4595494442  1.6598029457
## dCA2_Mpos_AACCTGACACTTGTGA-1dCA2_MI_GFPve_pos  3.3924710922 -0.2684416451
## dCA2_Mpos_AACGAAACATCCGTGG-1dCA2_MI_GFPve_pos  2.8091669254  1.8877501808
## dCA2_Mpos_AACGGGAAGTAGTCCT-1dCA2_MI_GFPve_pos  2.5137913399  1.6329703651
## dCA2_Mpos_AACGTCAGTCCACTTC-1dCA2_MI_GFPve_pos  3.3619237118  1.8889985405
## dCA2_Mpos_AACGTCAGTCCTGGGT-1dCA2_MI_GFPve_pos  1.5892256670  1.4783892952
## dCA2_Mpos_AAGAACAGTCCAATCA-1dCA2_MI_GFPve_pos  0.3364518456  1.6080813728
## dCA2_Mpos_AAGACAAAGACTGGGT-1dCA2_MI_GFPve_pos  1.8657181434  1.2431435905
## dCA2_Mpos_AAGACAACAGTTTCAG-1dCA2_MI_GFPve_pos  1.9698951416  1.1156454407
## dCA2_Mpos_AAGCGAGAGCGCCATC-1dCA2_MI_GFPve_pos  3.3802259140  1.2241134964
## dCA2_Mpos_AAGCGAGCAGCAGACA-1dCA2_MI_GFPve_pos  1.9945266418  1.8140001617
## dCA2_Mpos_AAGGAATAGTTAGTGA-1dCA2_MI_GFPve_pos  3.5766911678  1.4192533813
## dCA2_Mpos_AAGTACCAGACTACCT-1dCA2_MI_GFPve_pos  2.9955170326 -1.2585739769
## dCA2_Mpos_AAGTACCCACACTTAG-1dCA2_MI_GFPve_pos  3.0587413482 -0.8701193012
## dCA2_Mpos_AAGTACCTCTCGCGTT-1dCA2_MI_GFPve_pos  2.0611105137  0.7801165901
## dCA2_Mpos_AAGTCGTGTAACTTCG-1dCA2_MI_GFPve_pos  1.5631202631  3.1002712570
## dCA2_Mpos_AAGTTCGGTCGTCAGC-1dCA2_MI_GFPve_pos  2.7242131405  0.7199068390
## dCA2_Mpos_AATCACGAGTTGCGCC-1dCA2_MI_GFPve_pos  1.8522916012  1.1172004066
## dCA2_Mpos_AATGACCTCATCGTAG-1dCA2_MI_GFPve_pos  3.1870248489  1.1681228004
## dCA2_Mpos_AATGCCAAGTGGACGT-1dCA2_MI_GFPve_pos  2.5787317924  1.1550226532
## dCA2_Mpos_AATGGCTAGTGCTCAT-1dCA2_MI_GFPve_pos  2.0653233700  3.1413507782
```

```
## dCA2_Mpos_AATTTCCCAATTCTCT-1dCA2_MI_GFPve_pos  3.2479829960 -0.4403814949
## dCA2_Mpos_ACAACCAAGCGCAATG-1dCA2_MI_GFPve_pos  2.7283303909 -0.7561709561
## dCA2_Mpos_ACAACCAAGTGTCATC-1dCA2_MI_GFPve_pos  2.0170498066  1.9231276832
## dCA2_Mpos_ACAACCAGTGACAGCA-1dCA2_MI_GFPve_pos  2.5997746162 -0.6041946091
## dCA2_Mpos_ACAAGCTAGTCACACT-1dCA2_MI_GFPve_pos  3.1703591518 -0.5745472588
## dCA2_Mpos_ACAAGCTCAGCATACT-1dCA2_MI_GFPve_pos  3.0841281585 -0.5999140419
## dCA2_Mpos_ACAAGCTTCTACGGGC-1dCA2_MI_GFPve_pos  3.2274651699  0.8854461036
## dCA2_Mpos_ACACAGTGTCCACTCT-1dCA2_MI_GFPve_pos  2.0136766605  1.0712552391
## dCA2_Mpos_ACACGCGAGCACACAG-1dCA2_MI_GFPve_pos  1.9286966495 -1.7614934124
## dCA2_Mpos_ACACGCGAGTATTGCC-1dCA2_MI_GFPve_pos  2.4670975380  0.8662171684
## dCA2_Mpos_ACAGAAATCATTGCGA-1dCA2_MI_GFPve_pos  3.4354012184  1.4465704284
## dCA2_Mpos_ACATCCCTCCGGACGT-1dCA2_MI_GFPve_pos  2.9349024467  2.2270083748
## dCA2_Mpos_ACATGCAGTCCATACA-1dCA2_MI_GFPve_pos  2.9405496292  0.3041110359
## dCA2_Mpos_ACCAAACGTTGCCGCA-1dCA2_MI_GFPve_pos  2.9132118397 -1.1089729943
## dCA2_Mpos_ACCAACACACCGTCTT-1dCA2_MI_GFPve_pos  2.8088505440 -0.1111745514
## dCA2_Mpos_ACCAACATCGGAAACG-1dCA2_MI_GFPve_pos  3.2736811809  0.1190080963
## dCA2_Mpos_ACCACAAAGAGATCGC-1dCA2_MI_GFPve_pos  2.5384979420  2.4135742508
## dCA2_Mpos_ACCCTCAAGTTGTCAC-1dCA2_MI_GFPve_pos -0.8084639735  0.4858408294
## dCA2_Mpos_ACCCTCACAAGCTGCC-1dCA2_MI_GFPve_pos  2.9595482521 -1.2150180020
## dCA2_Mpos_ACCTACCGTATCGATC-1dCA2_MI_GFPve_pos  3.2121465378 -0.9619590916
## dCA2_Mpos_ACCTGAAAGATCCGAG-1dCA2_MI_GFPve_pos  3.2892050915  1.6821513496
## dCA2_Mpos_ACCTGAACACAACCGC-1dCA2_MI_GFPve_pos  2.5444371872 -1.2912010826
## dCA2_Mpos_ACGATCAAGATGTTGA-1dCA2_MI_GFPve_pos  2.5510368519  2.0037527405
## dCA2_Mpos_ACGATCATCCCATGGG-1dCA2_MI_GFPve_pos  3.4707832508 -0.1348161377
## dCA2_Mpos_ACGATGTAGCAGAAAG-1dCA2_MI_GFPve_pos  2.8853779011  1.6023903213
## dCA2_Mpos_ACGATGTCAGGATTCT-1dCA2_MI_GFPve_pos  1.9007582836  1.9185719810
## dCA2_Mpos_ACGCACGGTGAGTTTC-1dCA2_MI_GFPve_pos  1.0045442753 -0.2489199318
## dCA2_Mpos_ACGGGTCGTACGACTT-1dCA2_MI_GFPve_pos  2.3374316864 -0.9723789372
## dCA2_Mpos_ACGGTTAAGCCATCCG-1dCA2_MI_GFPve_pos  1.4117535524 -1.6056797184
## dCA2_Mpos_ACGGTTAAGTGTCATC-1dCA2_MI_GFPve_pos  3.3205971889 -0.2158908524
## dCA2_Mpos_ACGGTTACATAGAATG-1dCA2_MI_GFPve_pos  2.4698958568  2.0425896965
## dCA2_Mpos_ACGTAACCACCTGTCT-1dCA2_MI_GFPve_pos  3.4171331100 -0.2773484863
## dCA2_Mpos_ACGTACATCATTGTTC-1dCA2_MI_GFPve_pos  1.9381926231  1.6216550194
## dCA2_Mpos_ACGTTCCAGCGTATGG-1dCA2_MI_GFPve_pos  1.8125295811  0.8755550705
## dCA2_Mpos_ACTATCTCAGAAATCA-1dCA2_MI_GFPve_pos  0.5497636370  0.9069471679
## dCA2_Mpos_ACTATGGCATGCGTGC-1dCA2_MI_GFPve_pos  2.4668998890  0.9751930557
## dCA2_Mpos_ACTATGGTCCAAGCTA-1dCA2_MI_GFPve_pos  2.7885220222  1.3412256561
## dCA2_Mpos_ACTATTCAGAAACCCG-1dCA2_MI_GFPve_pos  3.0778923206 -0.6244053520
## dCA2_Mpos_ACTCCCAAGATCCGAG-1dCA2_MI_GFPve_pos  2.5634701423  0.9664230667
## dCA2_Mpos_ACTCCCAAGGGTGGGA-1dCA2_MI_GFPve_pos  3.0348029308  1.8551102005
## dCA2_Mpos_ACTGCAAGTACTGCGC-1dCA2_MI_GFPve_pos  3.1568041019  1.4334016166
## dCA2_Mpos_ACTGTCCCAATTGAGA-1dCA2_MI_GFPve_pos  1.8038909607  1.4295039497
## dCA2_Mpos_ACTGTGACAACCAATC-1dCA2_MI_GFPve_pos  1.2773397140 -1.6856488861
## dCA2_Mpos_ACTTATCTCCCATGGG-1dCA2_MI_GFPve_pos  3.5439734630 -0.1476621308
## dCA2_Mpos_ACTTTCACACCGTACG-1dCA2_MI_GFPve_pos  3.4420333080  0.5661835991
## dCA2_Mpos_ACTTTCAGTAGGCTCC-1dCA2_MI_GFPve_pos  1.6605439358 -0.3617114701
## dCA2_Mpos_AGAAATGTCAGCGCAC-1dCA2_MI_GFPve_pos  2.3193669491 -1.6617719807
## dCA2_Mpos_AGAAGCGCAACGATCT-1dCA2_MI_GFPve_pos  2.1542637520  2.5687957130
## dCA2_Mpos_AGACAAAAGCCATATC-1dCA2_MI_GFPve_pos  2.4905278854 -0.4440431275
## dCA2_Mpos_AGACAAAAGCGCCGTT-1dCA2_MI_GFPve_pos  1.3371749096  2.3068127952
## dCA2_Mpos_AGACAAACAGCTGTTA-1dCA2_MI_GFPve_pos  3.4350912742  0.7299709640
## dCA2_Mpos_AGACACTAGACGACTG-1dCA2_MI_GFPve_pos  2.5899937324  1.9514260612
## dCA2_Mpos_AGACACTAGGGTATAT-1dCA2_MI_GFPve_pos  3.0969650917 -0.5894078888
## dCA2_Mpos_AGACAGGAGGTTTACC-1dCA2_MI_GFPve_pos  2.8159167938  0.5733557068
## dCA2_Mpos_AGACCCGCATATCTCT-1dCA2_MI_GFPve_pos  3.1138804130 -1.0502729096
## dCA2_Mpos_AGACTCATCGGAGCAA-1dCA2_MI_GFPve_pos  2.8501458339  0.9561190925
## dCA2_Mpos_AGAGAGCGTGATCGTT-1dCA2_MI_GFPve_pos  2.8075132541  2.7643351875
## dCA2_Mpos_AGAGCAGCAAATCGGG-1dCA2_MI_GFPve_pos  2.3112757377  2.1416764580
## dCA2_Mpos_AGAGCAGCACTCGATA-1dCA2_MI_GFPve_pos  2.2233948879 -0.0304159798
## dCA2_Mpos_AGAGCCCAGACCAAAT-1dCA2_MI_GFPve_pos  2.3767659836 -0.7226786293
## dCA2_Mpos_AGAGCCCTCTCACTCG-1dCA2_MI_GFPve_pos  3.1046722107 -0.8596582092
## dCA2_Mpos_AGATCCAAGGGTCACA-1dCA2_MI_GFPve_pos  2.4901662044  0.1025581680
## dCA2_Mpos_AGATCGTCAACAGTGG-1dCA2_MI_GFPve_pos  2.4251320534  1.7429547630
## dCA2_Mpos_AGATGAACAGCTGTAT-1dCA2_MI_GFPve_pos  3.1177754574  1.2092480980
## dCA2_Mpos_AGATGCTCATCATTTC-1dCA2_MI_GFPve_pos  2.4846053295  1.9177222572
## dCA2_Mpos_AGATGCTGTTCTTGTT-1dCA2_MI_GFPve_pos  3.2942285709  1.8763594948
## dCA2_Mpos_AGCATCATCACGGACC-1dCA2_MI_GFPve_pos  2.9150009327  1.4698987327
## dCA2_Mpos_AGCCAGCCATACCAGT-1dCA2_MI_GFPve_pos  2.3958645038 -0.3180870690
## dCA2_Mpos_AGCGATTGTGAGTAAT-1dCA2_MI_GFPve_pos  2.5646999054 -0.2138018288
## dCA2_Mpos_AGCGATTTCCGAGTGC-1dCA2_MI_GFPve_pos  2.5994730167  2.5275254570
```

```
## dCA2_Mpos_AGCGCCATCACCATAG-1dCA2_MI_GFPve_pos  2.5499761276 -1.5673896946
## dCA2_Mpos_AGCGCTGAGTGAGTTA-1dCA2_MI_GFPve_pos  2.5252523594  2.4208097778
## dCA2_Mpos_AGCGTATAGCCGTCGT-1dCA2_MI_GFPve_pos  2.5058267288  1.6839976631
## dCA2_Mpos_AGCTACAGTCATCACA-1dCA2_MI_GFPve_pos  2.4129941635  1.4637079559
## dCA2_Mpos_AGCTACAGTCGCAACC-1dCA2_MI_GFPve_pos  2.9361665420 -0.9237336792
## dCA2_Mpos_AGCTTCCCAAGCCTGC-1dCA2_MI_GFPve_pos  2.1752269439  2.1252875648
## dCA2_Mpos_AGCTTCCCAGTAGGAC-1dCA2_MI_GFPve_pos -0.9255258866 -0.3313941635
## dCA2_Mpos_AGCTTCCGTGATTGGG-1dCA2_MI_GFPve_pos  1.8120043449 -1.1800450958
## dCA2_Mpos_AGGAGGTAGATGGTAT-1dCA2_MI_GFPve_pos  2.6300170593 -0.8150694050
## dCA2_Mpos_AGGATAAAGAGTTGAT-1dCA2_MI_GFPve_pos  3.2843668632  1.2831693016
## dCA2_Mpos_AGGCTGCGTCACCCTT-1dCA2_MI_GFPve_pos  3.3001384907  1.0668249450
## dCA2_Mpos_AGGCTGCGTTACGATC-1dCA2_MI_GFPve_pos  3.1345169716 -0.0884971298
## dCA2_Mpos_AGGGAGTGTTTAGAGA-1dCA2_MI_GFPve_pos  2.1956761055  2.0936341606
## dCA2_Mpos_AGGGCCTAGATGCAGC-1dCA2_MI_GFPve_pos  3.2632012539 -0.9154464878
## dCA2_Mpos_AGGGTCCAGCATCAAA-1dCA2_MI_GFPve_pos  3.4032194786  0.1891546570
## dCA2_Mpos_AGGGTCCAGGTAGCAC-1dCA2_MI_GFPve_pos  3.5275302105 -0.0908546128
## dCA2_Mpos_AGGGTGACATGGAACG-1dCA2_MI_GFPve_pos  1.2977605037  2.8548412643
## dCA2_Mpos_AGGTCATCACTGGAAG-1dCA2_MI_GFPve_pos  2.1523523502  0.6379108749
## dCA2_Mpos_AGGTCTAGTCGATTTG-1dCA2_MI_GFPve_pos  2.7317154579  2.4933271728
## dCA2_Mpos_AGGTGTTTCCTCTGCA-1dCA2_MI_GFPve_pos  2.1067502670 -0.6384262718
## dCA2_Mpos_AGGTTACAGCGATTCT-1dCA2_MI_GFPve_pos  2.9578118496  1.7348637901
## dCA2_Mpos_AGGTTGTCAAATTGCC-1dCA2_MI_GFPve_pos  2.8817084007 -1.1606223263
## dCA2_Mpos_AGGTTGTGTACGCTTA-1dCA2_MI_GFPve_pos  2.4163942508  0.4270425163
## dCA2_Mpos_AGTAACCTCGCCTAGG-1dCA2_MI_GFPve_pos  1.4968456201  1.6111531578
## dCA2_Mpos_AGTACCAGTGGTTTGT-1dCA2_MI_GFPve_pos  2.7404658966  0.6796655975
## dCA2_Mpos_AGTACTGGTGTTTACG-1dCA2_MI_GFPve_pos  2.9391942196  0.9055095039
## dCA2_Mpos_AGTCAACTCTCATAGG-1dCA2_MI_GFPve_pos  3.1468565635 -0.8360988774
## dCA2_Mpos_AGTCATGAGACGCCAA-1dCA2_MI_GFPve_pos  1.7373055391  2.3384046875
## dCA2_Mpos_AGTCATGAGGGAGGTG-1dCA2_MI_GFPve_pos  2.6252489261  0.3394642196
## dCA2_Mpos_AGTCATGGTGTGTCGC-1dCA2_MI_GFPve_pos  1.9577753715  1.7906442009
## dCA2_Mpos_AGTCATGTCTAAGCCA-1dCA2_MI_GFPve_pos  2.1526279621 -0.0194720902
## dCA2_Mpos_AGTCTCCCACATGGTT-1dCA2_MI_GFPve_pos  0.8251276784  1.0361109100
## dCA2_Mpos_AGTCTCCTCCGACATA-1dCA2_MI_GFPve_pos  2.8733923607  1.7969594322
## dCA2_Mpos_AGTGACTGTGGATGAC-1dCA2_MI_GFPve_pos  1.2849590950  1.7357116066
## dCA2_Mpos_AGTGACTGTTTCCATT-1dCA2_MI_GFPve_pos  2.7645909958  1.6184931122
## dCA2_Mpos_AGTGATCAGCGGCTCT-1dCA2_MI_GFPve_pos  1.3132062130  0.6996693931
## dCA2_Mpos_AGTGATCCACGGCCAT-1dCA2_MI_GFPve_pos  2.2485804729  3.0774965606
## dCA2_Mpos_AGTTCCCGTAGCCCTG-1dCA2_MI_GFPve_pos  2.2244310550  1.3632383667
## dCA2_Mpos_ATACCGAGTTCTTCAT-1dCA2_MI_GFPve_pos  2.4219293766 -0.5906614937
## dCA2_Mpos_ATACTTCCAAGTGGTG-1dCA2_MI_GFPve_pos  2.8741142921  1.8631353699
## dCA2_Mpos_ATAGACCCATAAGCGG-1dCA2_MI_GFPve_pos -0.6815764733 -0.0649332680
## dCA2_Mpos_ATAGACCCATCGTTCC-1dCA2_MI_GFPve_pos  3.3893203907  1.1431374870
## dCA2_Mpos_ATAGAGATCTCTGCTG-1dCA2_MI_GFPve_pos  2.1468317680  2.3584885917
## dCA2_Mpos_ATAGGCTGTGACATCT-1dCA2_MI_GFPve_pos  2.9298155479  1.6033315979
## dCA2_Mpos_ATAGGCTTCCTTATCA-1dCA2_MI_GFPve_pos  1.9181521110  1.3717379890
## dCA2_Mpos_ATCACAGAGATGGGCT-1dCA2_MI_GFPve_pos  2.8518808059  0.9771962486
## dCA2_Mpos_ATCACAGTCCCATTTA-1dCA2_MI_GFPve_pos  2.8987000160  0.7454095207
## dCA2_Mpos_ATCACAGTCGCTGATA-1dCA2_MI_GFPve_pos  1.2633028202  1.4185996376
## dCA2_Mpos_ATCACGAGTGTTCGTA-1dCA2_MI_GFPve_pos  2.4584589176 -0.6428045906
## dCA2_Mpos_ATCATTCGTCGTGATT-1dCA2_MI_GFPve_pos  2.8159218006  1.0891723953
## dCA2_Mpos_ATCCACCAGAACAAGG-1dCA2_MI_GFPve_pos  3.1344592743 -0.5968336739
## dCA2_Mpos_ATCCACCCATCGAGCC-1dCA2_MI_GFPve_pos  1.8097008638 -1.7527305760
## dCA2_Mpos_ATCCACCGTACTGTTG-1dCA2_MI_GFPve_pos  0.3426921539 -2.2430758156
## dCA2_Mpos_ATCCCTGAGCAAGCCA-1dCA2_MI_GFPve_pos  0.7331758313  1.7252145134
## dCA2_Mpos_ATCCCTGCAATCAAGA-1dCA2_MI_GFPve_pos  3.0121774845  2.3466925941
## dCA2_Mpos_ATCCCTGCAGAGAAAG-1dCA2_MI_GFPve_pos  2.8550896816 -0.1678785958
## dCA2_Mpos_ATCCTATCATGGAATA-1dCA2_MI_GFPve_pos  1.5739512615  0.5287485443
## dCA2_Mpos_ATCCTATTCCGACAGC-1dCA2_MI_GFPve_pos  2.2022006683  0.2084851585
## dCA2_Mpos_ATCGATGTCCAACACA-1dCA2_MI_GFPve_pos  3.1953673534 -0.8227359928
## dCA2_Mpos_ATCGATGTCGCACTCT-1dCA2_MI_GFPve_pos  3.0996580295  0.5975208603
## dCA2_Mpos_ATCGATGTCTAGACAC-1dCA2_MI_GFPve_pos  3.3274741344 -0.4921679177
## dCA2_Mpos_ATCGCCTCAGATGCGA-1dCA2_MI_GFPve_pos  2.5684800319 -1.1430022396
## dCA2_Mpos_ATCGGATTCGGCTGGT-1dCA2_MI_GFPve_pos  2.6986725502  1.1455445610
## dCA2_Mpos_ATCGTAGCAGAGCCCT-1dCA2_MI_GFPve_pos  2.4530623131  1.6432080589
## dCA2_Mpos_ATCGTAGGTTCGTACA-1dCA2_MI_GFPve_pos  1.9107535057  0.9638481460
## dCA2_Mpos_ATCGTCCGTTGTATGC-1dCA2_MI_GFPve_pos  1.9282267265  1.9242153488
## dCA2_Mpos_ATCGTGACACTTGAAC-1dCA2_MI_GFPve_pos  2.9788091354  1.1788812004
## dCA2_Mpos_ATCTCTACATACTGTG-1dCA2_MI_GFPve_pos  2.4293060474  0.8244738899
## dCA2_Mpos_ATCTTCAAGTCCCGAC-1dCA2_MI_GFPve_pos  2.1378343277  0.9115615211
## dCA2_Mpos_ATCTTCACAGTTCTAG-1dCA2_MI_GFPve_pos  2.9755885773  1.0868435226
```

```
## dCA2_Mpos_ATGAAAGTCCATAAGC-1dCA2_MI_GFPve_pos  3.1057050400 -0.0818113960
## dCA2_Mpos_ATGACCAAGTACTGGG-1dCA2_MI_GFPve_pos  2.4717037849  0.0918469749
## dCA2_Mpos_ATGACCATCCCGAAAT-1dCA2_MI_GFPve_pos  1.2472039633  1.7504139267
## dCA2_Mpos_ATGAGTCAGCGCTTCG-1dCA2_MI_GFPve_pos  1.2921474151  1.2026367508
## dCA2_Mpos_ATGATCGGTCCTACAA-1dCA2_MI_GFPve_pos  2.2165439300  1.9931917511
## dCA2_Mpos_ATGCATGCACAATTCG-1dCA2_MI_GFPve_pos  2.1088032894  0.8639431320
## dCA2_Mpos_ATGCATGGTCTTACTT-1dCA2_MI_GFPve_pos  2.5961699657  0.1440420471
## dCA2_Mpos_ATGCCTCGTCGCAACC-1dCA2_MI_GFPve_pos  1.9721892051  0.3795748077
## dCA2_Mpos_ATGCGATAGGGTTGCA-1dCA2_MI_GFPve_pos  2.0956182651 -1.0238365807
## dCA2_Mpos_ATGCGATCAAGTGCAG-1dCA2_MI_GFPve_pos  2.3777239494  1.4857435547
## dCA2_Mpos_ATGCGATGTGGAGAAA-1dCA2_MI_GFPve_pos  3.5434477501  1.0394645057
## dCA2_Mpos_ATGGAGGCAAGACCTT-1dCA2_MI_GFPve_pos  1.6527345352  1.7354421936
## dCA2_Mpos_ATGGGTTCATCCGCGA-1dCA2_MI_GFPve_pos  3.0492034130 -0.9294573940
## dCA2_Mpos_ATGTCCCCATAGAAAC-1dCA2_MI_GFPve_pos  2.0234646969  2.9798832260
## dCA2_Mpos_ATGTCCCGTCACCGCA-1dCA2_MI_GFPve_pos  3.1929438286 -0.0552072205
## dCA2_Mpos_ATGTCTTAGCAACCAG-1dCA2_MI_GFPve_pos  2.4681687527  1.7920818649
## dCA2_Mpos_ATTACTCCAGTAGAAT-1dCA2_MI_GFPve_pos  2.0196657352  0.7270880066
## dCA2_Mpos_ATTATCCGTTTACCAG-1dCA2_MI_GFPve_pos  2.9779679947 -0.9121842064
## dCA2_Mpos_ATTATCCTCCACTTTA-1dCA2_MI_GFPve_pos  2.7477271728 -0.6558146157
## dCA2_Mpos_ATTCACTCACCTGCAG-1dCA2_MI_GFPve_pos  3.3235728912  1.1188278518
## dCA2_Mpos_ATTCATCTCCGCGATG-1dCA2_MI_GFPve_pos  2.5901303463 -0.6866340317
## dCA2_Mpos_ATTCCTACATTCTCCG-1dCA2_MI_GFPve_pos  2.1128773861 -0.3857583679
## dCA2_Mpos_ATTCCTAGTTAGCTAC-1dCA2_MI_GFPve_pos  1.2825975590  1.7404589973
## dCA2_Mpos_ATTCGTTAGCGTGCCT-1dCA2_MI_GFPve_pos  2.3797500305 -0.7107987084
## dCA2_Mpos_ATTCGTTGTATGACAA-1dCA2_MI_GFPve_pos  2.0723049812  1.1927895866
## dCA2_Mpos_ATTCGTTTCTCGACGG-1dCA2_MI_GFPve_pos  1.8729889564  0.6078434311
## dCA2_Mpos_ATTCTACCAAATGATG-1dCA2_MI_GFPve_pos  2.5859229736 -0.0880508103
## dCA2_Mpos_ATTGGGTCAACCGCCA-1dCA2_MI_GFPve_pos  2.1291122608 -1.3002056755
## dCA2_Mpos_ATTTACCTCAAACGAA-1dCA2_MI_GFPve_pos  1.8239332609  0.9713769279
## dCA2_Mpos_ATTTCACCAGTGAGCA-1dCA2_MI_GFPve_pos  3.0034265690 -0.2914342560
## dCA2_Mpos_ATTTCTGGTGTTCAGT-1dCA2_MI_GFPve_pos  2.8923447304  1.2975831352
## dCA2_Mpos_ATTTCTGTCTTGTGCC-1dCA2_MI_GFPve_pos  1.5151152782  1.4390869461
## dCA2_Mpos_CAACAACCACAATGTC-1dCA2_MI_GFPve_pos  1.2960187368  0.8336792312
## dCA2_Mpos_CAACCAACACTGCGTG-1dCA2_MI_GFPve_pos  2.0997369461  2.5087247215
## dCA2_Mpos_CAACCTCCATTACGGT-1dCA2_MI_GFPve_pos  3.1004986934 -1.2020914234
## dCA2_Mpos_CAACGGCGTGGATCGA-1dCA2_MI_GFPve_pos  3.3316345386 -0.3884834923
## dCA2_Mpos_CAACGGCGTGTGAGCA-1dCA2_MI_GFPve_pos  0.9937439136  1.7330613457
## dCA2_Mpos_CAAGCTAGTGCACATT-1dCA2_MI_GFPve_pos  3.0021915607  2.0753670059
## dCA2_Mpos_CAATCGAAGTATGCAA-1dCA2_MI_GFPve_pos  3.1193235092  0.1971211754
## dCA2_Mpos_CAATCGATCACGTAGT-1dCA2_MI_GFPve_pos  2.9858732395  0.9664550148
## dCA2_Mpos_CAATGACAGGTCCTGC-1dCA2_MI_GFPve_pos  2.3409476452  1.0616765343
## dCA2_Mpos_CAATTTCAGACTTCAC-1dCA2_MI_GFPve_pos  2.3015670948 -0.9328105129
## dCA2_Mpos_CAATTTCAGCATCTTG-1dCA2_MI_GFPve_pos  0.7594185643 -0.6924461998
## dCA2_Mpos_CAATTTCGTCTTTATC-1dCA2_MI_GFPve_pos  1.0670076780  0.8978247963
## dCA2_Mpos_CACACAAAGTGGCGAT-1dCA2_MI_GFPve_pos  2.9518039398  2.1087589584
## dCA2_Mpos_CACACAAAGTTTCGAC-1dCA2_MI_GFPve_pos  2.8267190628 -0.0850801148
## dCA2_Mpos_CACAGATCAAGTAGTA-1dCA2_MI_GFPve_pos  2.3234071903 -0.8897959866
## dCA2_Mpos_CACCAAACAAGTTCCA-1dCA2_MI_GFPve_pos  2.5605848007 -1.5237626709
## dCA2_Mpos_CACGAATCAAGAGTAT-1dCA2_MI_GFPve_pos  2.0047176056  0.5947633110
## dCA2_Mpos_CACGAATCAGTTGGTT-1dCA2_MI_GFPve_pos -0.2299341760 -2.8851358570
## dCA2_Mpos_CACGAATGTGCTGATT-1dCA2_MI_GFPve_pos  2.1394004993  0.4049415909
## dCA2_Mpos_CACGGGTTCTTTGCAT-1dCA2_MI_GFPve_pos  1.9508001976  2.0635714851
## dCA2_Mpos_CACGTGGCACCGTGAC-1dCA2_MI_GFPve_pos  3.3793246918 -0.3505501427
## dCA2_Mpos_CACGTGGGTTCTCACC-1dCA2_MI_GFPve_pos  2.5206940346 -0.2622542061
## dCA2_Mpos_CACGTGGTCCATTGGA-1dCA2_MI_GFPve_pos  1.8999405079  1.0528235756
## dCA2_Mpos_CACGTGGTCGATTTCT-1dCA2_MI_GFPve_pos  2.2401614361 -0.4182843842
## dCA2_Mpos_CACGTTCTCGCTTGAA-1dCA2_MI_GFPve_pos  2.1527519398  0.8942537628
## dCA2_Mpos_CACTAAGTCAGCTAGT-1dCA2_MI_GFPve_pos  2.7155785732  0.9200411163
## dCA2_Mpos_CACTAAGTCTCACGAA-1dCA2_MI_GFPve_pos  2.6902442150  2.6909447037
## dCA2_Mpos_CACTGGGCAAGAGAGA-1dCA2_MI_GFPve_pos  2.6953983478 -0.2370667137
## dCA2_Mpos_CACTGTCGTATCAGGG-1dCA2_MI_GFPve_pos  2.8518157177 -0.9159750618
## dCA2_Mpos_CACTGTCGTCATAGTC-1dCA2_MI_GFPve_pos  2.6430468731  1.6834063850
## dCA2_Mpos_CACTTCGAGACCTCCG-1dCA2_MI_GFPve_pos  2.5563924484 -1.2822966255
## dCA2_Mpos_CAGATCAGTATGAAAC-1dCA2_MI_GFPve_pos  3.3176131420 -0.2962221779
## dCA2_Mpos_CAGCACGAGTCTGCAT-1dCA2_MI_GFPve_pos  2.4979190998  1.7033596359
## dCA2_Mpos_CAGCACGTCATTGGTG-1dCA2_MI_GFPve_pos  1.9094202690  2.1683702789
## dCA2_Mpos_CAGCAGCCAAGCTGCC-1dCA2_MI_GFPve_pos  2.3443515472  0.3080988251
## dCA2_Mpos_CAGCGTGAGTGAGGCT-1dCA2_MI_GFPve_pos  3.0754597359  1.8184323631
## dCA2_Mpos_CAGCGTGCACATGTTG-1dCA2_MI_GFPve_pos  2.1711232834  1.5539923034
## dCA2_Mpos_CAGGCCACATGGTACT-1dCA2_MI_GFPve_pos  3.3316035442 -0.2490000405
```

```
## dCA2_Mpos_CAGGGCTCACCCAACG-1dCA2_MI_GFPve_pos  3.1455974750  0.5834651314
## dCA2_Mpos_CAGGGCTGTCACGTGC-1dCA2_MI_GFPve_pos  1.6829294138  0.3453293167
## dCA2_Mpos_CAGGTATTCATCCCGT-1dCA2_MI_GFPve_pos  3.2153487377 -0.8381743111
## dCA2_Mpos_CAGGTATTCTCTATGT-1dCA2_MI_GFPve_pos  2.1198156051  0.0390644394
## dCA2_Mpos_CAGTGCGAGTGATAGT-1dCA2_MI_GFPve_pos  1.7755428724  1.9202056251
## dCA2_Mpos_CAGTGCGCAGAACGCA-1dCA2_MI_GFPve_pos  2.0695579223  0.1229105316
## dCA2_Mpos_CAGTTAGAGAAGCGGG-1dCA2_MI_GFPve_pos  2.2888176613  2.4200167976
## dCA2_Mpos_CAGTTAGCAACACAGG-1dCA2_MI_GFPve_pos  0.0932980053  1.6611962639
## dCA2_Mpos_CAGTTAGGTAATACCC-1dCA2_MI_GFPve_pos  2.8063221149  1.2053938232
## dCA2_Mpos_CAGTTCCAGAGAGTTT-1dCA2_MI_GFPve_pos  3.0263662510  2.0626545273
## dCA2_Mpos_CAGTTCCTCATCCCGT-1dCA2_MI_GFPve_pos  1.9588012867  0.3594570480
## dCA2_Mpos_CATACCCCATACAGAA-1dCA2_MI_GFPve_pos  2.4877216987 -0.2480091728
## dCA2_Mpos_CATACTTGTAGAGACC-1dCA2_MI_GFPve_pos  2.1245515518  1.1187081657
## dCA2_Mpos_CATCCCAAGTGGTTCT-1dCA2_MI_GFPve_pos  2.4799604587  1.0827389084
## dCA2_Mpos_CATCCCAGTCACCTTC-1dCA2_MI_GFPve_pos  2.9669556789 -1.2651378788
## dCA2_Mpos_CATCCCATCGTTAGAC-1dCA2_MI_GFPve_pos  1.6491570644 -0.2741875328
## dCA2_Mpos_CATCGGGGTGAAAGTT-1dCA2_MI_GFPve_pos  3.1454284363 -0.6004857697
## dCA2_Mpos_CATCGTCCACTCCTGT-1dCA2_MI_GFPve_pos  2.9736619167 -0.7237152733
## dCA2_Mpos_CATGCCTAGTATGTAG-1dCA2_MI_GFPve_pos  2.7537167244  2.5448122345
## dCA2_Mpos_CATGCGGAGGGACTGT-1dCA2_MI_GFPve_pos  1.0903290682  1.0855570160
## dCA2_Mpos_CATGCGGAGGTGCCAA-1dCA2_MI_GFPve_pos  3.5641882591 -0.1050071396
## dCA2_Mpos_CATGCGGCATAGGTAA-1dCA2_MI_GFPve_pos  2.6514620952  1.5583901726
## dCA2_Mpos_CATGCTCCATCCGAAT-1dCA2_MI_GFPve_pos  2.1436767750  0.2676105820
## dCA2_Mpos_CATGGTAGTAATTAGG-1dCA2_MI_GFPve_pos  3.3671245746  1.5207186065
## dCA2_Mpos_CATGGTAGTCAAATCC-1dCA2_MI_GFPve_pos  3.2002072506  0.9900045715
## dCA2_Mpos_CATTCCGAGGGTGAGG-1dCA2_MI_GFPve_pos  2.6543977432  2.3131337486
## dCA2_Mpos_CATTGAGGTGATTCTG-1dCA2_MI_GFPve_pos  2.3764562778  0.9996056877
## dCA2_Mpos_CATTGCCGTTAGCTAC-1dCA2_MI_GFPve_pos  0.7383230500  1.1031394325
## dCA2_Mpos_CATTGTTGTTTGGGTT-1dCA2_MI_GFPve_pos  1.3342837267  1.3957419716
## dCA2_Mpos_CATTTCATCTGTTCAT-1dCA2_MI_GFPve_pos  0.8968287878 -0.9981937088
## dCA2_Mpos_CCAATGACAGCTACCG-1dCA2_MI_GFPve_pos  1.1939050131  1.1004643760
## dCA2_Mpos_CCACACTAGAGAGTGA-1dCA2_MI_GFPve_pos  3.5516371898  1.3152609192
## dCA2_Mpos_CCACACTCAATGGCCC-1dCA2_MI_GFPve_pos  3.0075724296 -1.0571260132
## dCA2_Mpos_CCACCATTCGTAGTCA-1dCA2_MI_GFPve_pos  1.0952915125  1.0662885032
## dCA2_Mpos_CCACGAGAGCTAAACA-1dCA2_MI_GFPve_pos -0.4806073733 -1.6442732491
## dCA2_Mpos_CCACGAGAGTATGCAA-1dCA2_MI_GFPve_pos  0.8503291778  0.8199763618
## dCA2_Mpos_CCACGAGCATATGAAG-1dCA2_MI_GFPve_pos  1.6678824596 -1.4567150749
## dCA2_Mpos_CCACGAGGTATGGGAC-1dCA2_MI_GFPve_pos  3.1530347042  0.1734686218
## dCA2_Mpos_CCACGTTCACCCTAGG-1dCA2_MI_GFPve_pos  2.0502147846 -1.2709831871
## dCA2_Mpos_CCACGTTCACGACTAT-1dCA2_MI_GFPve_pos  2.5501511268 -0.7341966309
## dCA2_Mpos_CCACGTTTCTTTCAGT-1dCA2_MI_GFPve_pos  3.2970938854  0.3085966430
## dCA2_Mpos_CCATAAGGTCGACTGC-1dCA2_MI_GFPve_pos  3.4700601272 -0.5074734368
## dCA2_Mpos_CCCAACTAGGCTTAGG-1dCA2_MI_GFPve_pos  2.0743818455  0.9781518303
## dCA2_Mpos_CCCATTGAGCTAATGA-1dCA2_MI_GFPve_pos  2.3951587848  2.2007575355
## dCA2_Mpos_CCCATTGGTTCTCTCG-1dCA2_MI_GFPve_pos  1.3420814447 -0.3361277260
## dCA2_Mpos_CCCGAAGAGACCCGCT-1dCA2_MI_GFPve_pos  2.3091166191  0.0777607284
## dCA2_Mpos_CCCGAAGAGTGTAGAT-1dCA2_MI_GFPve_pos  2.8737847976  0.6166010223
## dCA2_Mpos_CCCGAAGGTGAAAGTT-1dCA2_MI_GFPve_pos  3.5673453979 -0.0116367020
## dCA2_Mpos_CCCGGAAGTGCACAAG-1dCA2_MI_GFPve_pos  2.3222370319  2.1454191528
## dCA2_Mpos_CCCTGATCAATCTGCA-1dCA2_MI_GFPve_pos  1.8348741703  0.8628244720
## dCA2_Mpos_CCCTGATTCTGTGTGA-1dCA2_MI_GFPve_pos  2.9234845810  0.6177468620
## dCA2_Mpos_CCCTTAGAGACCATTC-1dCA2_MI_GFPve_pos -0.6824821062  0.4384169899
## dCA2_Mpos_CCCTTAGAGCGACATG-1dCA2_MI_GFPve_pos  2.6237735920  2.6887364708
## dCA2_Mpos_CCCTTAGAGGATTCCT-1dCA2_MI_GFPve_pos  2.2887723617 -1.3226864018
## dCA2_Mpos_CCCTTAGCACACACGC-1dCA2_MI_GFPve_pos  1.7967649870 -0.3865751900
## dCA2_Mpos_CCGATGGCACTGGCGT-1dCA2_MI_GFPve_pos  2.8009259872  1.6773071609
## dCA2_Mpos_CCGCAAGCAAATCAGA-1dCA2_MI_GFPve_pos  1.9628865890  1.7271056495
## dCA2_Mpos_CCGGACAGTGTAGGAC-1dCA2_MI_GFPve_pos  2.6812875442  1.0611996971
## dCA2_Mpos_CCGGTAGAGTTACGGG-1dCA2_MI_GFPve_pos  2.5312988930  1.2747445427
## dCA2_Mpos_CCGGTGATCCGGCAAC-1dCA2_MI_GFPve_pos  2.9102583103  1.7377724968
## dCA2_Mpos_CCGTAGGTCTGGAGAG-1dCA2_MI_GFPve_pos  2.8264932804  1.6655736290
## dCA2_Mpos_CCGTGAGCACGCGTGT-1dCA2_MI_GFPve_pos  1.7393111162  2.4253216110
## dCA2_Mpos_CCGTGAGCATCGGCCA-1dCA2_MI_GFPve_pos  2.1259598903 -0.0490922608
## dCA2_Mpos_CCGTTCAGTGTAACGG-1dCA2_MI_GFPve_pos  2.9413225822  1.2680421196
## dCA2_Mpos_CCTAACCCAACGTAAA-1dCA2_MI_GFPve_pos  3.3612098865 -0.6635632195
## dCA2_Mpos_CCTATCGTCTCTATGT-1dCA2_MI_GFPve_pos  3.2750322990 -0.8734614529
## dCA2_Mpos_CCTCATGCAACGATTC-1dCA2_MI_GFPve_pos  0.9900590353 -1.9211628117
## dCA2_Mpos_CCTCATGTCTCGAGTA-1dCA2_MI_GFPve_pos  3.3111505680  2.0604300819
## dCA2_Mpos_CCTCTCCCAAGTGGAC-1dCA2_MI_GFPve_pos  2.8328168564 -0.1157655396
## dCA2_Mpos_CCTCTCCGTATGTCCA-1dCA2_MI_GFPve_pos  1.6380512886 -0.7100228943
```

```
## dCA2_Mpos_CCTGCATAGCATCAGG-1dCA2_MI_GFPve_pos  2.0362320118 -1.8293986000
## dCA2_Mpos_CCTGCATGTGTCCAAT-1dCA2_MI_GFPve_pos -0.4415662117 -0.0322994866
## dCA2_Mpos_CCTTCAGCAAGTATAG-1dCA2_MI_GFPve_pos  1.5794050865  1.7328977905
## dCA2_Mpos_CCTTCAGCAAGTGCAG-1dCA2_MI_GFPve_pos  2.4887774162  2.0046730362
## dCA2_Mpos_CCTTCAGCATGGCTGC-1dCA2_MI_GFPve_pos  3.4413120918  0.1427336059
## dCA2_Mpos_CCTTGTGCAGACAAGC-1dCA2_MI_GFPve_pos  3.2101512127  1.7191520057
## dCA2_Mpos_CGAAGGAAGAGCATAT-1dCA2_MI_GFPve_pos  2.2275433712  2.3566766105
## dCA2_Mpos_CGAAGTTCAGCAGGAT-1dCA2_MI_GFPve_pos  2.7930600815 -0.1174063362
## dCA2_Mpos_CGAATTGTCAACCCGG-1dCA2_MI_GFPve_pos  1.6927253179  1.0268740974
## dCA2_Mpos_CGAGAAGCAATGACCT-1dCA2_MI_GFPve_pos  2.6579363518  0.5791769348
## dCA2_Mpos_CGAGGAAAGCTGCGAA-1dCA2_MI_GFPve_pos  2.4173491173  2.7357931457
## dCA2_Mpos_CGAGGAATCGCTGACG-1dCA2_MI_GFPve_pos  2.2454407387  1.4600272499
## dCA2_Mpos_CGAGGCTCACATCATG-1dCA2_MI_GFPve_pos  2.0332322292  0.8699708305
## dCA2_Mpos_CGAGTTATCTTTGCAT-1dCA2_MI_GFPve_pos  2.4223158531  1.1939683281
## dCA2_Mpos_CGATCGGTCTTCTGTA-1dCA2_MI_GFPve_pos  3.3071499042  1.1773777328
## dCA2_Mpos_CGATGCGGTTAGGGAC-1dCA2_MI_GFPve_pos  2.8060746364  1.1876921974
## dCA2_Mpos_CGCAGGTGTGTCATCA-1dCA2_MI_GFPve_pos  2.9642875366  0.4277153335
## dCA2_Mpos_CGCATAACAGAGGCTA-1dCA2_MI_GFPve_pos  2.4010551147  1.6090798698
## dCA2_Mpos_CGCATAATCCAAGCTA-1dCA2_MI_GFPve_pos  2.8202526741  1.0054445587
## dCA2_Mpos_CGCATGGAGACCATAA-1dCA2_MI_GFPve_pos  0.8172239237  1.0502114616
## dCA2_Mpos_CGCATGGCACAATGCT-1dCA2_MI_GFPve_pos  2.0065696411 -2.3397066273
## dCA2_Mpos_CGCATGGCATGTGCTA-1dCA2_MI_GFPve_pos  0.9250295334  0.2520938240
## dCA2_Mpos_CGCCAGACAAACGGCA-1dCA2_MI_GFPve_pos  2.7811885052 -1.1390256561
## dCA2_Mpos_CGCCATTAGGCTGTAG-1dCA2_MI_GFPve_pos  2.9484939747 -0.2037114777
## dCA2_Mpos_CGCCATTAGGTAGACC-1dCA2_MI_GFPve_pos  1.8999281101  0.6634922348
## dCA2_Mpos_CGCCATTGTGTAACGG-1dCA2_MI_GFPve_pos  2.5601499252  1.0913915954
## dCA2_Mpos_CGCGTGAGTAGTATAG-1dCA2_MI_GFPve_pos  2.6481132679  0.9000931106
## dCA2_Mpos_CGGAACCTCTAAGGAA-1dCA2_MI_GFPve_pos  2.8736710720 -0.3114194550
## dCA2_Mpos_CGGAGAAAGCACCAGA-1dCA2_MI_GFPve_pos  2.0525827579  1.9092760406
## dCA2_Mpos_CGGCAGTAGATCCGAG-1dCA2_MI_GFPve_pos  2.9329493217  1.8669901214
## dCA2_Mpos_CGGGACTAGAATAACC-1dCA2_MI_GFPve_pos  3.2148597412 -0.0096988358
## dCA2_Mpos_CGGGACTAGACTTCAC-1dCA2_MI_GFPve_pos  3.0910539799  0.5312218986
## dCA2_Mpos_CGGTCAGCAATCCAGT-1dCA2_MI_GFPve_pos  3.1391649418  1.7816949211
## dCA2_Mpos_CGTAAGTGTGGCTGCT-1dCA2_MI_GFPve_pos  2.9945004158  1.3449192367
## dCA2_Mpos_CGTAAGTTCTCTGGTC-1dCA2_MI_GFPve_pos  2.8533599548  0.7919040046
## dCA2_Mpos_CGTCAAAAGCCTCTCT-1dCA2_MI_GFPve_pos  1.0676611833  0.6424970947
## dCA2_Mpos_CGTCAAACAATGTCTG-1dCA2_MI_GFPve_pos  1.9357993774 -0.1661219277
## dCA2_Mpos_CGTCAAACATCCGAGC-1dCA2_MI_GFPve_pos  3.2133534126 -0.4437532105
## dCA2_Mpos_CGTCAAACATCCTGTC-1dCA2_MI_GFPve_pos  2.7252230816 -1.3208214916
## dCA2_Mpos_CGTCAAAGTCGACTGC-1dCA2_MI_GFPve_pos  2.6360984020  1.4784341179
## dCA2_Mpos_CGTCAAATCAACCCGG-1dCA2_MI_GFPve_pos  2.9105477505  0.6131749473
## dCA2_Mpos_CGTGCTTAGACCTGGA-1dCA2_MI_GFPve_pos  2.1970987491  2.8388328872
## dCA2_Mpos_CGTGCTTCAGGCCCTA-1dCA2_MI_GFPve_pos  2.2890148334  0.9467702232
## dCA2_Mpos_CGTTAGAAGGTCGTCC-1dCA2_MI_GFPve_pos  3.1065483265  1.9000463806
## dCA2_Mpos_CGTTGGGAGATACATG-1dCA2_MI_GFPve_pos  2.8146703415  2.0237379394
## dCA2_Mpos_CGTTGGGCATTGTACG-1dCA2_MI_GFPve_pos  2.9602415733  1.0824246727
## dCA2_Mpos_CGTTGGGGTATAGCTC-1dCA2_MI_GFPve_pos  1.7594254188  1.2433548294
## dCA2_Mpos_CTAACCCTCAAGGACG-1dCA2_MI_GFPve_pos  2.8093044929  2.2355985962
## dCA2_Mpos_CTAACTTCACGTACAT-1dCA2_MI_GFPve_pos  2.6432376079  2.4294381462
## dCA2_Mpos_CTACATTTCAAGCCTA-1dCA2_MI_GFPve_pos  2.8032341175 -0.9451324619
## dCA2_Mpos_CTACCTGCATAAGCAA-1dCA2_MI_GFPve_pos  2.3348500900  1.6609444938
## dCA2_Mpos_CTACCTGGTAGCGATG-1dCA2_MI_GFPve_pos  2.6416223221  2.4010439239
## dCA2_Mpos_CTACCTGTCCGTATGA-1dCA2_MI_GFPve_pos  2.8889086418 -0.0360250153
## dCA2_Mpos_CTACTATAGAGATTCA-1dCA2_MI_GFPve_pos  0.4508620672  1.4323091827
## dCA2_Mpos_CTAGACATCGTCAAAC-1dCA2_MI_GFPve_pos  1.9899747543  0.3888655029
## dCA2_Mpos_CTAGGTAGTCCTCAGG-1dCA2_MI_GFPve_pos  2.0473580532  1.7678017937
## dCA2_Mpos_CTATAGGCAACACACT-1dCA2_MI_GFPve_pos  3.0223634414 -0.0485615410
## dCA2_Mpos_CTCAACCAGACGCATG-1dCA2_MI_GFPve_pos  1.5660385065  1.4272642456
## dCA2_Mpos_CTCAACCCAACTTCTT-1dCA2_MI_GFPve_pos  2.8173737697 -0.9710874237
## dCA2_Mpos_CTCAACCGTAATGTGA-1dCA2_MI_GFPve_pos  2.0929725342 -2.1093558945
## dCA2_Mpos_CTCAAGAGTGTTGATC-1dCA2_MI_GFPve_pos  2.2418232136  3.0446529709
## dCA2_Mpos_CTCACTGAGATCACTC-1dCA2_MI_GFPve_pos  2.6328332596 -0.6071142830
## dCA2_Mpos_CTCAGAACAGGTTTAC-1dCA2_MI_GFPve_pos  3.1431555919  0.3330736481
## dCA2_Mpos_CTCAGGGAGCTACTAC-1dCA2_MI_GFPve_pos  1.9648175411 -2.0499865689
## dCA2_Mpos_CTCAGTCAGTCACACT-1dCA2_MI_GFPve_pos  2.8896241360  1.4780965172
## dCA2_Mpos_CTCAGTCTCCGCGATG-1dCA2_MI_GFPve_pos  1.0773474150  0.9600210510
## dCA2_Mpos_CTCATCGCATGTGCCG-1dCA2_MI_GFPve_pos  2.6304440670  0.1715307556
## dCA2_Mpos_CTCATGCTCATGCCAA-1dCA2_MI_GFPve_pos  2.1467421226  2.6519885383
## dCA2_Mpos_CTCATTAAGTGACACG-1dCA2_MI_GFPve_pos  2.7775643043  0.0544805847
## dCA2_Mpos_CTCCAACAGCTAATGA-1dCA2_MI_GFPve_pos  2.8610508613 -0.6561417259
```

```
## dCA2_Mpos_CTCCACACATCGATGT-1dCA2_MI_GFPve_pos  1.4397776298  2.6112156235
## dCA2_Mpos_CTCCATGCATCCTATT-1dCA2_MI_GFPve_pos  2.7999150924  1.3628626190
## dCA2_Mpos_CTCCCTCAGGAAAGAC-1dCA2_MI_GFPve_pos  2.5799789600  0.0769572578
## dCA2_Mpos_CTCCTCCGTTACCTTT-1dCA2_MI_GFPve_pos  2.4190638237 -0.4105467476
## dCA2_Mpos_CTCTCAGAGGAGAATG-1dCA2_MI_GFPve_pos  3.3573193722 -0.4280466713
## dCA2_Mpos_CTCTCAGCATTGGCAT-1dCA2_MI_GFPve_pos  2.2534129791 -0.8885299839
## dCA2_Mpos_CTCTCGAGTTGTGTAC-1dCA2_MI_GFPve_pos  2.2768101864  2.1837311111
## dCA2_Mpos_CTGAGGCTCGCCGAAC-1dCA2_MI_GFPve_pos  1.9443614654  2.0430822693
## dCA2_Mpos_CTGCAGGGTATCTTCT-1dCA2_MI_GFPve_pos  2.5305307083  1.0552707039
## dCA2_Mpos_CTGCAGGTCTAGTGTG-1dCA2_MI_GFPve_pos  3.1189699344  1.0149832092
## dCA2_Mpos_CTGCATCGTGAATTGA-1dCA2_MI_GFPve_pos  2.7764299087  2.0078592621
## dCA2_Mpos_CTGCATCTCCCTCATG-1dCA2_MI_GFPve_pos  1.7535109691 -1.8091218151
## dCA2_Mpos_CTGCTCAGTTGCCAAT-1dCA2_MI_GFPve_pos  1.8956093960 -0.3842205681
## dCA2_Mpos_CTGGCAGCAAGTGGTG-1dCA2_MI_GFPve_pos  1.8665876560  2.4942179046
## dCA2_Mpos_CTGGCAGGTAGTGGCA-1dCA2_MI_GFPve_pos  2.1789157562 -1.0339102425
## dCA2_Mpos_CTGGTCTAGAAATTCG-1dCA2_MI_GFPve_pos  2.4751939945  0.9282980285
## dCA2_Mpos_CTGTATTAGAGGTCAC-1dCA2_MI_GFPve_pos  2.9907446079  0.3492541633
## dCA2_Mpos_CTTACCGAGGGATCGT-1dCA2_MI_GFPve_pos  2.7021277122  1.6455970131
## dCA2_Mpos_CTTACCGGTGTCATGT-1dCA2_MI_GFPve_pos  2.1062996559  0.4424109779
## dCA2_Mpos_CTTAGGAGTAGACAAT-1dCA2_MI_GFPve_pos  1.1656943731 -0.5542568840
## dCA2_Mpos_CTTCCTTTCTATTGTC-1dCA2_MI_GFPve_pos  1.4771050386  1.6209597908
## dCA2_Mpos_CTTCTAAGTACTCGTA-1dCA2_MI_GFPve_pos  2.5878398590  1.8253650986
## dCA2_Mpos_CTTGAGAAGGTAGCCA-1dCA2_MI_GFPve_pos  2.9874563389  2.1285763107
## dCA2_Mpos_CTTTCGGAGGCGATAC-1dCA2_MI_GFPve_pos  1.6908174925  2.4377384506
## dCA2_Mpos_GAAACCTCACAACCGC-1dCA2_MI_GFPve_pos  1.5244962148  1.1911797844
## dCA2_Mpos_GAAACCTTCCGAGAAG-1dCA2_MI_GFPve_pos  2.6861696415 -1.5160400547
## dCA2_Mpos_GAAATGAAGATGAACT-1dCA2_MI_GFPve_pos  2.5497818165  0.7848129593
## dCA2_Mpos_GAAATGAGTCGTGCCA-1dCA2_MI_GFPve_pos  2.2844586544  2.9893293701
## dCA2_Mpos_GAAGAATGTTGGTGTT-1dCA2_MI_GFPve_pos  3.0774822407  2.2807045303
## dCA2_Mpos_GAAGCCCCAAAGGCTG-1dCA2_MI_GFPve_pos  3.2503955536  1.4947057091
## dCA2_Mpos_GAAGCGAGTTTCTATC-1dCA2_MI_GFPve_pos  0.8563184910  1.3956647239
## dCA2_Mpos_GAAGGACTCTCCATAT-1dCA2_MI_GFPve_pos  3.2130046062 -0.3270582833
## dCA2_Mpos_GAAGGGTGTACCCGCA-1dCA2_MI_GFPve_pos  3.3161118202  0.3454461418
## dCA2_Mpos_GAAGGGTTCTGGCTGG-1dCA2_MI_GFPve_pos  2.8292691879 -0.9209909119
## dCA2_Mpos_GAATAGATCCACAGGC-1dCA2_MI_GFPve_pos  0.2850423925  1.0518999420
## dCA2_Mpos_GAATAGATCCTCTTTC-1dCA2_MI_GFPve_pos  3.0194578342 -0.7613744415
## dCA2_Mpos_GAATCACTCGAGAGCA-1dCA2_MI_GFPve_pos  2.4760739975  2.7200231872
## dCA2_Mpos_GACAGCCGTACCTATG-1dCA2_MI_GFPve_pos  3.0317921810 -1.0386552490
## dCA2_Mpos_GACATCATCCGGTAAT-1dCA2_MI_GFPve_pos  2.8189056091 -1.2216849007
## dCA2_Mpos_GACCCAGGTGAAGCTG-1dCA2_MI_GFPve_pos  2.7927620582  0.8632531486
## dCA2_Mpos_GACCCTTCAGGTGACA-1dCA2_MI_GFPve_pos  2.0732400589 -0.4173998512
## dCA2_Mpos_GACCGTGAGCAGAAAG-1dCA2_MI_GFPve_pos  2.2871003322  0.3744049393
## dCA2_Mpos_GACTATGTCAGCCTTC-1dCA2_MI_GFPve_pos  1.7652912311 -0.2645597138
## dCA2_Mpos_GACTATGTCCGACAGC-1dCA2_MI_GFPve_pos  2.6787502937 -0.6609158196
## dCA2_Mpos_GACTCAAAGGCACCAA-1dCA2_MI_GFPve_pos  2.5348646812  2.4468508087
## dCA2_Mpos_GACTCTCGTATCGATC-1dCA2_MI_GFPve_pos  2.6312568359  0.3918695770
## dCA2_Mpos_GACTGATTCGAGCTGC-1dCA2_MI_GFPve_pos -0.1034696005 -1.1430897392
## dCA2_Mpos_GAGAAATAGATACCAA-1dCA2_MI_GFPve_pos  2.8427231483  0.0149789177
## dCA2_Mpos_GAGACCCCATCCCGTT-1dCA2_MI_GFPve_pos  3.1617696457 -0.9768011250
## dCA2_Mpos_GAGAGGTCAGAGTAAT-1dCA2_MI_GFPve_pos  3.1436751060  2.3132806145
## dCA2_Mpos_GAGAGGTCAGGTCAAG-1dCA2_MI_GFPve_pos  2.4905786686  2.6647854172
## dCA2_Mpos_GAGAGGTGTGCCTATA-1dCA2_MI_GFPve_pos  1.1820225887  0.6057396255
## dCA2_Mpos_GAGATGGTCAGTGATC-1dCA2_MI_GFPve_pos  2.0239732437 -0.9330799259
## dCA2_Mpos_GAGATGGTCTCGCGTT-1dCA2_MI_GFPve_pos  2.4021210842 -0.9517047085
## dCA2_Mpos_GAGCCTGGTGGGAGAG-1dCA2_MI_GFPve_pos  2.2522695236  2.0954470955
## dCA2_Mpos_GAGCTGCTCAGACCCG-1dCA2_MI_GFPve_pos  3.0548188858 -1.0641157307
## dCA2_Mpos_GAGGCAAAGGAAGAAC-1dCA2_MI_GFPve_pos  2.4832253628  2.2762322746
## dCA2_Mpos_GAGGCAATCAAACCTG-1dCA2_MI_GFPve_pos  3.1354036503 -0.4444846787
## dCA2_Mpos_GAGGCCTGTTCATCGA-1dCA2_MI_GFPve_pos  3.5187103920  0.0549578987
## dCA2_Mpos_GAGGGTAAGGACAGTC-1dCA2_MI_GFPve_pos  3.2823803596  1.8631687485
## dCA2_Mpos_GAGGGTACAAGAGTAT-1dCA2_MI_GFPve_pos  2.9947202377  1.7307191215
## dCA2_Mpos_GAGGGTACAAGGCTTT-1dCA2_MI_GFPve_pos  2.6606114082 -0.2375721611
## dCA2_Mpos_GAGGGTAGTACTAGCT-1dCA2_MI_GFPve_pos  1.9660322838 -0.3647427239
## dCA2_Mpos_GAGTCATTCCATCTGC-1dCA2_MI_GFPve_pos  2.9814329319  0.3307881676
## dCA2_Mpos_GAGTCTACATACGCAT-1dCA2_MI_GFPve_pos  2.2884199791 -0.0221242584
## dCA2_Mpos_GAGTGAGTCAACGCTA-1dCA2_MI_GFPve_pos  3.3989563160 -0.7785329498
## dCA2_Mpos_GAGTGAGTCTTCTCAA-1dCA2_MI_GFPve_pos  3.4103694134  0.3403535209
## dCA2_Mpos_GAGTTACGTTAGAGAT-1dCA2_MI_GFPve_pos  1.3795567922 -2.9678367056
## dCA2_Mpos_GAGTTGTAGTGATAAC-1dCA2_MI_GFPve_pos  1.5031521492  1.4354696594
## dCA2_Mpos_GAGTTGTCACTCTGCT-1dCA2_MI_GFPve_pos  2.0876350574  0.4050956093
```

```
## dCA2_Mpos_GAGTTTGTCGCTTGAA-1dCA2_MI_GFPve_pos  2.5707214050  0.0318975769
## dCA2_Mpos_GATAGAAGTGAAGCGT-1dCA2_MI_GFPve_pos  1.9110577278  1.1197500549
## dCA2_Mpos_GATAGCTAGGTAACTA-1dCA2_MI_GFPve_pos  3.3389020138 -0.7740471043
## dCA2_Mpos_GATAGCTCAACGGTAG-1dCA2_MI_GFPve_pos  3.0422208481 -0.7400254883
## dCA2_Mpos_GATAGCTCAGCTTTGA-1dCA2_MI_GFPve_pos  1.9197764568  1.0521598182
## dCA2_Mpos_GATCACAGTAGTGTGG-1dCA2_MI_GFPve_pos  3.2413993053  2.1274857841
## dCA2_Mpos_GATCATGAGATTAGAC-1dCA2_MI_GFPve_pos  2.9158015423 -0.7608174957
## dCA2_Mpos_GATCATGCAGCGGATA-1dCA2_MI_GFPve_pos  1.7075295620  1.8017754875
## dCA2_Mpos_GATCCCTAGCGGCTCT-1dCA2_MI_GFPve_pos  2.6325521641 -1.1688248791
## dCA2_Mpos_GATCGTATCATGCCAA-1dCA2_MI_GFPve_pos  1.2454359703  2.1212029777
## dCA2_Mpos_GATCGTATCCTACGGG-1dCA2_MI_GFPve_pos  3.2324307136 -0.7384247460
## dCA2_Mpos_GATGATCCAAGAAACT-1dCA2_MI_GFPve_pos  1.5014631920 -1.6654944100
## dCA2_Mpos_GATGCTAAGCACTCTA-1dCA2_MI_GFPve_pos  3.2499447040  0.4872827850
## dCA2_Mpos_GATGGAGAGTGCGCTC-1dCA2_MI_GFPve_pos  2.4016034775 -0.6708488144
## dCA2_Mpos_GATGGAGCAGGCCTGT-1dCA2_MI_GFPve_pos  1.5688592367  0.9994511925
## dCA2_Mpos_GATGTTGGTCATGCAT-1dCA2_MI_GFPve_pos  0.9692796640  0.5668869339
## dCA2_Mpos_GATTCGAAGCGTCGAA-1dCA2_MI_GFPve_pos  2.5009811096  2.4277820907
## dCA2_Mpos_GATTCGACACCTGTCT-1dCA2_MI_GFPve_pos  2.6549878292 -0.4941754021
## dCA2_Mpos_GATTCTTGTGCCTGAC-1dCA2_MI_GFPve_pos  2.9889607601  0.7573867164
## dCA2_Mpos_GATTCTTTCTTGGGCG-1dCA2_MI_GFPve_pos  1.9795708828  0.9185700737
## dCA2_Mpos_GCAACCGCAAGGGTCA-1dCA2_MI_GFPve_pos  2.8075049095  1.1705131851
## dCA2_Mpos_GCACATAGTACTCGCG-1dCA2_MI_GFPve_pos  2.4290218525  0.1800127350
## dCA2_Mpos_GCACGTGCAAGCTGCC-1dCA2_MI_GFPve_pos  2.3496470623 -0.1235651650
## dCA2_Mpos_GCACGTGCAGATGCGA-1dCA2_MI_GFPve_pos  1.9816172294  1.1663804374
## dCA2_Mpos_GCACGTGTCCCGAAAT-1dCA2_MI_GFPve_pos  1.2483568363 -1.7570357002
## dCA2_Mpos_GCACTAATCACGGACC-1dCA2_MI_GFPve_pos  2.8289330177 -0.8853540100
## dCA2_Mpos_GCAGCCAGTCGATTAC-1dCA2_MI_GFPve_pos  2.8511884384  1.9023928963
## dCA2_Mpos_GCAGCCATCATCACTT-1dCA2_MI_GFPve_pos  2.6672122650 -0.1430854477
## dCA2_Mpos_GCAGCTGAGTCCCGGT-1dCA2_MI_GFPve_pos  1.1118838959 -1.1162299790
## dCA2_Mpos_GCAGGCTTCCATCAGA-1dCA2_MI_GFPve_pos  2.4467990570  2.3156237922
## dCA2_Mpos_GCAGTTATCTGGGCAC-1dCA2_MI_GFPve_pos  2.3560197525  0.7764869057
## dCA2_Mpos_GCATCGGAGGCACTAG-1dCA2_MI_GFPve_pos  1.2825008802  2.0895486198
## dCA2_Mpos_GCATCGGCATGAGAAT-1dCA2_MI_GFPve_pos  3.1396293812 -0.5613689102
## dCA2_Mpos_GCATCGGGTAGGACTG-1dCA2_MI_GFPve_pos  2.2372605972  3.0757375084
## dCA2_Mpos_GCATCGGGTCTAGGTT-1dCA2_MI_GFPve_pos  1.6255964212  1.6176996551
## dCA2_Mpos_GCATCTCCAGAAACCG-1dCA2_MI_GFPve_pos  2.4533324413  0.9354806267
## dCA2_Mpos_GCATGATTCAGTGTTG-1dCA2_MI_GFPve_pos  2.2649946384 -0.1646475472
## dCA2_Mpos_GCCAGGTAGGTTAAAC-1dCA2_MI_GFPve_pos  3.0052113704 -0.7845487274
## dCA2_Mpos_GCCAGGTGTGTCCACG-1dCA2_MI_GFPve_pos  1.1571983270  2.4619107567
## dCA2_Mpos_GCCATGGGTCTACGAT-1dCA2_MI_GFPve_pos  2.3895659618  2.4573312126
## dCA2_Mpos_GCCATTCAGCCTTTGA-1dCA2_MI_GFPve_pos  2.8912701778 -1.1485247292
## dCA2_Mpos_GCCATTCTCCTTGACC-1dCA2_MI_GFPve_pos  2.3750553303  0.8037886940
## dCA2_Mpos_GCCCAGACACAACGAG-1dCA2_MI_GFPve_pos  2.6131458454  0.7885518394
## dCA2_Mpos_GCCCAGACATTCTGTT-1dCA2_MI_GFPve_pos  1.8193178348  1.9445176445
## dCA2_Mpos_GCCCAGAGTAGCGAGT-1dCA2_MI_GFPve_pos  3.2191429310 -0.0267119087
## dCA2_Mpos_GCCGATGCACGCGCAT-1dCA2_MI_GFPve_pos  2.7938089542  0.5584383331
## dCA2_Mpos_GCCGTGATCAACTCTT-1dCA2_MI_GFPve_pos  3.1336047821  0.8352952324
## dCA2_Mpos_GCCTGTTGTGGTCCCA-1dCA2_MI_GFPve_pos  1.9808967285  0.5253072105
## dCA2_Mpos_GCGAGAAAGAGGGTGG-1dCA2_MI_GFPve_pos  2.3083014660  1.9585776649
## dCA2_Mpos_GCGAGAATCCTTCACG-1dCA2_MI_GFPve_pos  1.1992540531 -1.7258081116
## dCA2_Mpos_GCTGAATAGACATCAA-1dCA2_MI_GFPve_pos  0.9407854252  1.1169214569
## dCA2_Mpos_GCTGAATTCGTGGAAG-1dCA2_MI_GFPve_pos  2.3616287880  0.0824365936
## dCA2_Mpos_GCTGCAGCAAATCGTC-1dCA2_MI_GFPve_pos  2.6793725662 -0.3763107933
## dCA2_Mpos_GCTGCAGTCACGACTA-1dCA2_MI_GFPve_pos  2.7584858112 -0.5047440209
## dCA2_Mpos_GCTGGGTCAAGCCATT-1dCA2_MI_GFPve_pos  2.6955564194 -1.3790578522
## dCA2_Mpos_GCTGGGTGTCACCCTT-1dCA2_MI_GFPve_pos  3.2841963939 -0.7032150902
## dCA2_Mpos_GGAACCCTCGCACGGT-1dCA2_MI_GFPve_pos  3.0342526607 -0.5637955345
## dCA2_Mpos_GGAAGTGAGTCCGTCG-1dCA2_MI_GFPve_pos  2.9723735027  1.9395013176
## dCA2_Mpos_GGAATCTCAGGAAGTC-1dCA2_MI_GFPve_pos  2.4014105968  2.3540764175
## dCA2_Mpos_GGAGAACGTCTGTGCG-1dCA2_MI_GFPve_pos  0.6728277378  1.1532435737
## dCA2_Mpos_GGAGCAAAGCTCACTA-1dCA2_MI_GFPve_pos  3.2611847095 -0.3885063805
## dCA2_Mpos_GGAGGATAGTAGTCCT-1dCA2_MI_GFPve_pos  1.9679188900 -1.8794612564
## dCA2_Mpos_GGAGGTACAGTATACC-1dCA2_MI_GFPve_pos  2.1346068554 -1.8225383438
## dCA2_Mpos_GGATCTAAGCACCTGC-1dCA2_MI_GFPve_pos  2.7221644096  0.4506278358
## dCA2_Mpos_GGCAGTCAGGGTTGCA-1dCA2_MI_GFPve_pos  2.8281181030 -0.5079273857
## dCA2_Mpos_GGCTGTGCAGTGGGTA-1dCA2_MI_GFPve_pos  2.2194144897  0.7414927803
## dCA2_Mpos_GGCTGTGCATAGGCGA-1dCA2_MI_GFPve_pos  3.2493076496 -0.3081030525
## dCA2_Mpos_GGCTGTGGTACCCACG-1dCA2_MI_GFPve_pos  1.9600088768  1.1397948585
## dCA2_Mpos_GGCTGTGGTCAAAGCG-1dCA2_MI_GFPve_pos  3.1495361500  0.6319871269
## dCA2_Mpos_GGCTGTGGTCTGTGGC-1dCA2_MI_GFPve_pos  1.0691958837  1.9145746551
```

```
## dCA2_Mpos_GGCTGTGTCGGCCCAA-1dCA2_MI_GFPve_pos  3.3838107757  1.0922541939
## dCA2_Mpos_GGCTTGGAGTTCACTG-1dCA2_MI_GFPve_pos  1.5815555029  0.2988996826
## dCA2_Mpos_GGCTTGGCAACAACAA-1dCA2_MI_GFPve_pos  2.3692104988 -1.1516239323
## dCA2_Mpos_GGCTTGGTCAGGAAAT-1dCA2_MI_GFPve_pos  2.1592695885  1.2980127655
## dCA2_Mpos_GGGAAGTAGTCTAGCT-1dCA2_MI_GFPve_pos  2.9129705601  0.5418916069
## dCA2_Mpos_GGGAAGTGTATTTCCT-1dCA2_MI_GFPve_pos  2.8707466297  0.9891825043
## dCA2_Mpos_GGGACAATCAACACGT-1dCA2_MI_GFPve_pos  2.3827941589 -1.5107068695
## dCA2_Mpos_GGGACCTCATGCCATA-1dCA2_MI_GFPve_pos  1.9161296062 -0.3376445450
## dCA2_Mpos_GGGACCTGTACTGTTG-1dCA2_MI_GFPve_pos  0.9473759107 -2.8466535486
## dCA2_Mpos_GGGACTCTCCTCACGT-1dCA2_MI_GFPve_pos  2.4823386841  0.0954752289
## dCA2_Mpos_GGGAGATTCGACGCGT-1dCA2_MI_GFPve_pos  2.1683886223  2.4241681419
## dCA2_Mpos_GGGAGTACACCATAAC-1dCA2_MI_GFPve_pos  2.4187877350  0.8121819816
## dCA2_Mpos_GGGAGTATCGACGTCG-1dCA2_MI_GFPve_pos  1.7512864046  1.6649327598
## dCA2_Mpos_GGGCTACCACAGAGAC-1dCA2_MI_GFPve_pos  2.7236828975  1.7497992836
## dCA2_Mpos_GGGCTCAGTACGAGCA-1dCA2_MI_GFPve_pos  3.5276873283  0.3206606231
## dCA2_Mpos_GGGCTCATCCCAAGTA-1dCA2_MI_GFPve_pos  2.5309176616  2.2523217521
## dCA2_Mpos_GGGTAGAGTCGAAACG-1dCA2_MI_GFPve_pos  2.5626039676  2.3637113891
## dCA2_Mpos_GGGTCACTCGCTCATC-1dCA2_MI_GFPve_pos  2.8463869266 -0.8853723683
## dCA2_Mpos_GGGTCTGAGCCTGAAG-1dCA2_MI_GFPve_pos  3.2217965297  0.4124474846
## dCA2_Mpos_GGGTGAATCACTTTGT-1dCA2_MI_GFPve_pos  2.2839472465 -0.0842461266
## dCA2_Mpos_GGGTTATGTCACGTGC-1dCA2_MI_GFPve_pos  1.0089321308  0.8461494766
## dCA2_Mpos_GGGTTATTCATGGAGG-1dCA2_MI_GFPve_pos  3.2148418598  0.8632264458
## dCA2_Mpos_GGTAACTTCCATCTGC-1dCA2_MI_GFPve_pos  2.3600073032  0.7717695556
## dCA2_Mpos_GGTAATCAGCCGTAAG-1dCA2_MI_GFPve_pos  1.6526080541  1.8177471481
## dCA2_Mpos_GGTAATCAGGCCCAAA-1dCA2_MI_GFPve_pos  3.6716687851  0.4803405128
## dCA2_Mpos_GGTAATCCAAGCGGAT-1dCA2_MI_GFPve_pos  1.9302425556  1.7563095413
## dCA2_Mpos_GGTAATCGTAGCTTTG-1dCA2_MI_GFPve_pos  2.5257103615  2.5155306183
## dCA2_Mpos_GGTAATCGTTCGGCTG-1dCA2_MI_GFPve_pos  1.5109948091  0.6755934082
## dCA2_Mpos_GGTAGAGAGTGCAACG-1dCA2_MI_GFPve_pos  2.3036143951  2.4725332580
## dCA2_Mpos_GGTAGAGCATTCACAG-1dCA2_MI_GFPve_pos  3.3056256943  0.3965502106
## dCA2_Mpos_GGTAGAGTCCATCAGA-1dCA2_MI_GFPve_pos  2.9116814308  1.6361260734
## dCA2_Mpos_GGTGAAGTCCATCTCG-1dCA2_MI_GFPve_pos  2.7580120735  0.1070547424
## dCA2_Mpos_GGTGATTGTAATCAGA-1dCA2_MI_GFPve_pos  3.0151820354 -0.0196828522
## dCA2_Mpos_GGTGGCTTCTCTAAGG-1dCA2_MI_GFPve_pos  2.3116185837 -1.0670966782
## dCA2_Mpos_GGTGTCGTCAACCCGG-1dCA2_MI_GFPve_pos  2.7513125114 -0.4342183747
## dCA2_Mpos_GGTGTTACAATGGCAG-1dCA2_MI_GFPve_pos  3.3242917232  1.1625619255
## dCA2_Mpos_GGTTAACAGGTTAGTA-1dCA2_MI_GFPve_pos  2.3010227852 -0.0969667114
## dCA2_Mpos_GGTTAACGTCATCCCT-1dCA2_MI_GFPve_pos  2.8453529053 -1.2444209732
## dCA2_Mpos_GGTTAACTCATGGATC-1dCA2_MI_GFPve_pos  2.3825013809  0.0824265800
## dCA2_Mpos_GGTTAACTCTTACCAT-1dCA2_MI_GFPve_pos  1.3470860891  1.6981773697
## dCA2_Mpos_GGTTCTCAGAAGAGCA-1dCA2_MI_GFPve_pos  2.6344998054  2.0731988273
## dCA2_Mpos_GGTTCTCAGCACTCCG-1dCA2_MI_GFPve_pos  2.8445575409 -0.1829771675
## dCA2_Mpos_GGTTCTCAGTGTACAA-1dCA2_MI_GFPve_pos  3.1191861801  1.9421549164
## dCA2_Mpos_GGTTCTCGTTTAGAGA-1dCA2_MI_GFPve_pos  3.2433848553  1.6351733528
## dCA2_Mpos_GTAACACGTATCCTCC-1dCA2_MI_GFPve_pos  2.2800052337  1.5488586746
## dCA2_Mpos_GTAACACTCAAGAGTA-1dCA2_MI_GFPve_pos  3.0593247585 -0.6026549019
## dCA2_Mpos_GTAACCAAGATCGACG-1dCA2_MI_GFPve_pos  2.3131022625  2.4468908630
## dCA2_Mpos_GTAACCAGTGGGCTCT-1dCA2_MI_GFPve_pos  2.0363197498  1.2313423477
## dCA2_Mpos_GTAAGTCGTATACCTG-1dCA2_MI_GFPve_pos  1.5849108867  1.2921142898
## dCA2_Mpos_GTAAGTCTCTACCACC-1dCA2_MI_GFPve_pos  1.7147953682 -0.4143557228
## dCA2_Mpos_GTAATCGAGCAAGCCA-1dCA2_MI_GFPve_pos  1.9700141125  1.3770218216
## dCA2_Mpos_GTAATCGCAATCGCCG-1dCA2_MI_GFPve_pos  2.1294169597  2.1256566368
## dCA2_Mpos_GTAATGCAGATGGTAT-1dCA2_MI_GFPve_pos  2.8690626793  2.3454089485
## dCA2_Mpos_GTAATGCCACAGTCCG-1dCA2_MI_GFPve_pos  2.5287852459  0.1486091934
## dCA2_Mpos_GTACAACGTATCGTAC-1dCA2_MI_GFPve_pos  1.7993819885  1.8702450119
## dCA2_Mpos_GTACAGTGTGTTGATC-1dCA2_MI_GFPve_pos  2.0671906643  0.6697898231
## dCA2_Mpos_GTAGAAAGTGGATTTC-1dCA2_MI_GFPve_pos  0.6935653262  1.1560187660
## dCA2_Mpos_GTAGAGGGTCTCCCTA-1dCA2_MI_GFPve_pos  2.2268967800  2.6477947555
## dCA2_Mpos_GTAGATCTCTGTACAG-1dCA2_MI_GFPve_pos  2.0914132767  2.3114366852
## dCA2_Mpos_GTCAAACGTCCAGGTC-1dCA2_MI_GFPve_pos  2.8682236843  0.7786684356
## dCA2_Mpos_GTCACTCAGACTGTTC-1dCA2_MI_GFPve_pos  2.9843485527  1.6694302879
## dCA2_Mpos_GTCACTCTCTTGGTGA-1dCA2_MI_GFPve_pos  1.5596567564  0.7232232414
## dCA2_Mpos_GTCAGCGGTTTAGACC-1dCA2_MI_GFPve_pos  2.6716239624  2.0428910576
## dCA2_Mpos_GTCATCCCACTTGTCC-1dCA2_MI_GFPve_pos  2.5987143688 -0.2814097084
## dCA2_Mpos_GTCATTTAGCTGACTT-1dCA2_MI_GFPve_pos  1.4390968256  1.3400950752
## dCA2_Mpos_GTCATTTGTAAGATTG-1dCA2_MI_GFPve_pos  1.8855581455  1.2867603622
## dCA2_Mpos_GTCCACTCAGCCATTA-1dCA2_MI_GFPve_pos  1.8286914997 -0.6548347153
## dCA2_Mpos_GTCCACTGTACTCAAC-1dCA2_MI_GFPve_pos  2.5069163017  0.9292497955
## dCA2_Mpos_GTCCACTTCGCTTGAA-1dCA2_MI_GFPve_pos  2.2698557548 -2.0085732617
## dCA2_Mpos_GTCCCATTCAAGTGTC-1dCA2_MI_GFPve_pos  2.8075554542  0.7185101829
```

```
## dCA2_Mpos_GTCCTCAGTTTCGGCG-1dCA2_MI_GFPve_pos  0.8949901037  1.7690816246
## dCA2_Mpos_GTCGCGACAGCCCAGT-1dCA2_MI_GFPve_pos  1.6093046837  2.0810838066
## dCA2_Mpos_GTCGCGAGTTTACTGG-1dCA2_MI_GFPve_pos  2.4173405342  2.9014206253
## dCA2_Mpos_GTCTACCAGCATCGAG-1dCA2_MI_GFPve_pos  2.1471555404  1.4608764969
## dCA2_Mpos_GTCTAGACAGGCTATT-1dCA2_MI_GFPve_pos  2.8198170833 -0.0658878960
## dCA2_Mpos_GTCTAGAGTCTTGAAC-1dCA2_MI_GFPve_pos  2.4310019188  2.6262288414
## dCA2_Mpos_GTCTCACGTCTCAGGC-1dCA2_MI_GFPve_pos  2.3390064411  0.9835791908
## dCA2_Mpos_GTCTGTCCAGCTCGGT-1dCA2_MI_GFPve_pos  1.6125966482  1.0358100258
## dCA2_Mpos_GTCTGTCGTCAGTCTA-1dCA2_MI_GFPve_pos  2.2925710850  2.8795576416
## dCA2_Mpos_GTCTGTCTCCTGTAAG-1dCA2_MI_GFPve_pos  2.7115390472  1.1155486427
## dCA2_Mpos_GTGAGCCGTATCGCGC-1dCA2_MI_GFPve_pos  0.4569081836 -3.8725400843
## dCA2_Mpos_GTGAGCCGTGTCCATA-1dCA2_MI_GFPve_pos  2.1623654537  1.3816037498
## dCA2_Mpos_GTGAGTTAGTAACGAT-1dCA2_MI_GFPve_pos  1.8661616020 -1.6578726448
## dCA2_Mpos_GTGATGTTCGTTGTTT-1dCA2_MI_GFPve_pos  3.1933822803  0.9402084671
## dCA2_Mpos_GTGCGTGAGCGGTAGT-1dCA2_MI_GFPve_pos  3.4306149654 -0.1658715882
## dCA2_Mpos_GTGCGTGCAACCGCCA-1dCA2_MI_GFPve_pos  1.0879297428 -1.3496429600
## dCA2_Mpos_GTGCGTGCAATTGGTC-1dCA2_MI_GFPve_pos  1.0543840103 -3.1997155823
## dCA2_Mpos_GTGCTGGTCACCTCAC-1dCA2_MI_GFPve_pos  2.4681232147  1.8719101272
## dCA2_Mpos_GTGCTTCAGTGCACCC-1dCA2_MI_GFPve_pos  3.1773209743 -0.5941986717
## dCA2_Mpos_GTGCTTCCAAGAGTAT-1dCA2_MI_GFPve_pos  0.0118854068  1.6384158455
## dCA2_Mpos_GTGCTTCCAGCTGAGA-1dCA2_MI_GFPve_pos  3.0091319256  1.4209242187
## dCA2_Mpos_GTGGAAGCATGCCGCA-1dCA2_MI_GFPve_pos  2.4271626644  2.8733802162
## dCA2_Mpos_GTGGCGTCACCAAATC-1dCA2_MI_GFPve_pos  1.9927249126  1.4126820885
## dCA2_Mpos_GTGGCGTTCAGGACGA-1dCA2_MI_GFPve_pos  3.2636089496  0.4373507820
## dCA2_Mpos_GTGGGAAAGACCTCCG-1dCA2_MI_GFPve_pos  2.6192724876  0.0168357216
## dCA2_Mpos_GTGGGAAGTAAGGAGA-1dCA2_MI_GFPve_pos  2.8096261196  1.5467973076
## dCA2_Mpos_GTGGTTACACACCTGG-1dCA2_MI_GFPve_pos  2.6261596851 -0.0243196167
## dCA2_Mpos_GTGGTTACATAAGCAA-1dCA2_MI_GFPve_pos  2.6573016815 -0.5792307533
## dCA2_Mpos_GTGTAACCAACGCATT-1dCA2_MI_GFPve_pos  3.0099139385  2.1784882866
## dCA2_Mpos_GTGTCCTCAGCACGAA-1dCA2_MI_GFPve_pos  2.3930912189  0.7260670982
## dCA2_Mpos_GTGTGATTCGCACGGT-1dCA2_MI_GFPve_pos  2.8472662144  0.3099375091
## dCA2_Mpos_GTGTTAGCATGATAGA-1dCA2_MI_GFPve_pos  2.1129768066  2.3680405937
## dCA2_Mpos_GTGTTAGTCTGAGTCA-1dCA2_MI_GFPve_pos  3.2046246700  1.5882702194
## dCA2_Mpos_GTGTTCCCACTCCACT-1dCA2_MI_GFPve_pos  3.2335758381  0.9371004425
## dCA2_Mpos_GTTACGAAGGAAACGA-1dCA2_MI_GFPve_pos  2.8395981960  2.5271854721
## dCA2_Mpos_GTTACGATCCCGAAAT-1dCA2_MI_GFPve_pos  0.1113772862  1.1712117515
## dCA2_Mpos_GTTAGACCAGATGCGA-1dCA2_MI_GFPve_pos  0.9942286186  1.4458427749
## dCA2_Mpos_GTTCATTCAAGGACAC-1dCA2_MI_GFPve_pos  2.7502517872 -0.4736699738
## dCA2_Mpos_GTTCTATCATAGATCC-1dCA2_MI_GFPve_pos  2.9828968220 -0.8156823792
## dCA2_Mpos_GTTCTATGTGTCATCA-1dCA2_MI_GFPve_pos  3.2916219406 -0.6321940102
## dCA2_Mpos_GTTGAACTCAAATAGG-1dCA2_MI_GFPve_pos  2.9542207889  0.1292753540
## dCA2_Mpos_GTTGCTCTCATCGCAA-1dCA2_MI_GFPve_pos  3.0442621879  0.6476526581
## dCA2_Mpos_GTTGTCCAGTATCCTG-1dCA2_MI_GFPve_pos  2.0176034145  0.4278321586
## dCA2_Mpos_GTTGTGACACAGCATT-1dCA2_MI_GFPve_pos  2.9582483940 -0.5866522469
## dCA2_Mpos_GTTGTGACACCATTCC-1dCA2_MI_GFPve_pos  1.5578972273  2.0460534416
## dCA2_Mpos_GTTTACTCATAGTCGT-1dCA2_MI_GFPve_pos  3.2224431209  2.0380082451
## dCA2_Mpos_GTTTACTGTGAATTGA-1dCA2_MI_GFPve_pos  1.6793743543  0.4895592056
## dCA2_Mpos_TAACGACAGTTCTACG-1dCA2_MI_GFPve_pos  2.7330522709 -0.5423283257
## dCA2_Mpos_TAACGACCAGAGATGC-1dCA2_MI_GFPve_pos -0.1792616067  0.6826753937
## dCA2_Mpos_TAACTTCCATATGGCT-1dCA2_MI_GFPve_pos  2.1571273975  0.9273901306
## dCA2_Mpos_TAAGCACCAGCCGTTG-1dCA2_MI_GFPve_pos  3.1232357197  1.5110068641
## dCA2_Mpos_TAAGCACGTACACGTT-1dCA2_MI_GFPve_pos  2.3044135742  0.6921658836
## dCA2_Mpos_TAAGCGTAGGGACACT-1dCA2_MI_GFPve_pos  3.1578922443 -0.6742162384
## dCA2_Mpos_TAATCTCGTCCGAAGA-1dCA2_MI_GFPve_pos  3.0026230984  0.2328090988
## dCA2_Mpos_TAATCTCGTGGGCTTC-1dCA2_MI_GFPve_pos  0.5858394675  0.9099517189
## dCA2_Mpos_TAATTCCGTACCATAC-1dCA2_MI_GFPve_pos  2.0573516063  1.2056055389
## dCA2_Mpos_TACATTCAGGAGAATG-1dCA2_MI_GFPve_pos  2.5165512733  1.1323871933
## dCA2_Mpos_TACATTCCACAAGTGG-1dCA2_MI_GFPve_pos  2.3875606231 -0.6530365624
## dCA2_Mpos_TACATTCCACCCTCTA-1dCA2_MI_GFPve_pos  3.0468123131 -1.1400308289
## dCA2_Mpos_TACATTCCATCTCGTC-1dCA2_MI_GFPve_pos  1.8276149206  1.7021603905
## dCA2_Mpos_TACATTCGTCGTTCAA-1dCA2_MI_GFPve_pos  2.2777710132 -1.7142817654
## dCA2_Mpos_TACCCACGTGGTACAG-1dCA2_MI_GFPve_pos  1.9730141334  2.8344803177
## dCA2_Mpos_TACCGGGAGCGTCAGA-1dCA2_MI_GFPve_pos  2.1424172096  2.5046959244
## dCA2_Mpos_TACCGGGGTTCTCCTG-1dCA2_MI_GFPve_pos  1.9126558475  0.9759435974
## dCA2_Mpos_TACCTCGTCATTACGG-1dCA2_MI_GFPve_pos  2.3418095283  1.1711678825
## dCA2_Mpos_TACCTGCTCATAGAGA-1dCA2_MI_GFPve_pos  1.2836891584  1.2649984680
## dCA2_Mpos_TACCTGCTCTACTCAT-1dCA2_MI_GFPve_pos  1.9265780620 -1.7466592468
## dCA2_Mpos_TACCTGCTCTGGTGGC-1dCA2_MI_GFPve_pos  1.9934904747  0.9308843933
## dCA2_Mpos_TACGGGCAGACTCATC-1dCA2_MI_GFPve_pos  3.2452418976 -0.5315193810
## dCA2_Mpos_TACGGTATCTGCGGGT-1dCA2_MI_GFPve_pos  2.9619023971  0.8634806000
```

```
## dCA2_Mpos_TACTGCCTCCTCAGAA-1dCA2_MI_GFPve_pos  2.0296673946  1.3061714493
## dCA2_Mpos_TACTTCAAGTCCCTAA-1dCA2_MI_GFPve_pos  3.2859642677 -0.5853428520
## dCA2_Mpos_TAGACCAGTTGCTGAT-1dCA2_MI_GFPve_pos  2.9581053428  0.2953515373
## dCA2_Mpos_TAGACTGCAGGATTCT-1dCA2_MI_GFPve_pos  3.1359136276 -0.9434652008
## dCA2_Mpos_TAGACTGCAGTGCGCT-1dCA2_MI_GFPve_pos  2.4713545017  1.6481604896
## dCA2_Mpos_TAGACTGGTGCTCGTG-1dCA2_MI_GFPve_pos  1.1998359136  1.4873237930
## dCA2_Mpos_TAGAGTCAGCTCTGTA-1dCA2_MI_GFPve_pos  2.8452584915 -0.5230140366
## dCA2_Mpos_TAGCACATCGGAACTT-1dCA2_MI_GFPve_pos  1.5761977606  1.6887984596
## dCA2_Mpos_TAGGAGGAGTTCGGTT-1dCA2_MI_GFPve_pos  2.4849300556 -0.3061532654
## dCA2_Mpos_TAGGGTTTCTATCGGA-1dCA2_MI_GFPve_pos  3.3716397457  0.0663194977
## dCA2_Mpos_TAGGTTGGTCTTTCTA-1dCA2_MI_GFPve_pos  3.1420552902 -0.4528627075
## dCA2_Mpos_TAGTGCAAGCGAATGC-1dCA2_MI_GFPve_pos  3.5285809212  0.4106517158
## dCA2_Mpos_TATACCTCAGCTGTAT-1dCA2_MI_GFPve_pos  2.3542888336 -0.1868843712
## dCA2_Mpos_TATCCTACAACAGATA-1dCA2_MI_GFPve_pos  2.6429431610  1.5929270111
## dCA2_Mpos_TATCGCCGTAAGTTGA-1dCA2_MI_GFPve_pos -0.3069895781  1.3517093979
## dCA2_Mpos_TATCTTGGTTCTTCAT-1dCA2_MI_GFPve_pos  1.2963348799  0.7859120689
## dCA2_Mpos_TATGTTCAGTGTGTTC-1dCA2_MI_GFPve_pos  1.5116173200  1.9416013084
## dCA2_Mpos_TATGTTCGTGAAAGTT-1dCA2_MI_GFPve_pos  1.2018281393  1.0504522644
## dCA2_Mpos_TATTCCAGTTATGTCG-1dCA2_MI_GFPve_pos  2.7359957866 -1.2296230473
## dCA2_Mpos_TATTCCATCTGGCCAG-1dCA2_MI_GFPve_pos  1.5995660000  0.5299678169
## dCA2_Mpos_TATTGGGAGTAAACTG-1dCA2_MI_GFPve_pos  2.5037129097  1.9655046783
## dCA2_Mpos_TCAAGCACAAGTAGTA-1dCA2_MI_GFPve_pos  1.4651463203  2.3182754837
## dCA2_Mpos_TCAAGTGCAGCGTACC-1dCA2_MI_GFPve_pos  2.2408302002  2.9475059830
## dCA2_Mpos_TCAATCTTCGCAGTCG-1dCA2_MI_GFPve_pos  2.7604839973  2.1913848244
## dCA2_Mpos_TCACAAGAGATAGGGA-1dCA2_MI_GFPve_pos  2.3851242237  0.5170584045
## dCA2_Mpos_TCACATTCAAAGCAAT-1dCA2_MI_GFPve_pos  3.3396928482 -0.7733299412
## dCA2_Mpos_TCACATTCAATACGCT-1dCA2_MI_GFPve_pos  2.1240611248  0.3041339241
## dCA2_Mpos_TCACATTTCAGCGCAC-1dCA2_MI_GFPve_pos  2.5379257374  1.5291538559
## dCA2_Mpos_TCACATTTCATCGCAA-1dCA2_MI_GFPve_pos  3.2819233112 -0.6026739754
## dCA2_Mpos_TCACGCTTCAGGAGAC-1dCA2_MI_GFPve_pos  2.4101057224 -1.0033018269
## dCA2_Mpos_TCACTCGTCCTACTGC-1dCA2_MI_GFPve_pos  2.1405496769  0.0445127807
## dCA2_Mpos_TCAGTCCCAACCGCTG-1dCA2_MI_GFPve_pos  2.8543608360 -0.9822079815
## dCA2_Mpos_TCAGTTTGTCAGCGTC-1dCA2_MI_GFPve_pos  2.0168118648  1.6414623581
## dCA2_Mpos_TCAGTTTTCCTTCAGC-1dCA2_MI_GFPve_pos  2.4933781795 -1.6670493759
## dCA2_Mpos_TCATACTGTCTTTCAT-1dCA2_MI_GFPve_pos  2.7413380317  1.2783422790
## dCA2_Mpos_TCATACTTCCTTCTAA-1dCA2_MI_GFPve_pos  1.7105164699  1.4459843956
## dCA2_Mpos_TCATATCAGGACCCAA-1dCA2_MI_GFPve_pos  2.8061568908 -0.7757582344
## dCA2_Mpos_TCATATCGTGGTACAG-1dCA2_MI_GFPve_pos  1.8448357754  2.2723436676
## dCA2_Mpos_TCATATCTCACAAGGG-1dCA2_MI_GFPve_pos  2.7798123531  2.4187689148
## dCA2_Mpos_TCATCATCACAGCTTA-1dCA2_MI_GFPve_pos  3.2119956188  0.6396732651
## dCA2_Mpos_TCATGCCCATCCGGTG-1dCA2_MI_GFPve_pos  1.2863266640  2.2441821419
## dCA2_Mpos_TCATGTTGTCAAAGAT-1dCA2_MI_GFPve_pos  2.8166268043 -0.4275822319
## dCA2_Mpos_TCATGTTGTTCGGTCG-1dCA2_MI_GFPve_pos  2.7285463981  1.3746686302
## dCA2_Mpos_TCATTACCAGGGTCTC-1dCA2_MI_GFPve_pos  0.2239014380  1.6285310112
## dCA2_Mpos_TCATTCATCATGGATC-1dCA2_MI_GFPve_pos  2.7235000305  2.4455952965
## dCA2_Mpos_TCATTGTGTATCGTAC-1dCA2_MI_GFPve_pos  2.1330609493  3.0700302444
## dCA2_Mpos_TCATTGTTCACTGTCC-1dCA2_MI_GFPve_pos  2.8041708641 -0.2947926201
## dCA2_Mpos_TCATTTGCAACTCGTA-1dCA2_MI_GFPve_pos  1.9016513996  2.5063634239
## dCA2_Mpos_TCATTTGGTTGAGGAC-1dCA2_MI_GFPve_pos  2.1089763813  2.2749896370
## dCA2_Mpos_TCCACGTGTCGACTGC-1dCA2_MI_GFPve_pos  1.8804595642  1.6025352798
## dCA2_Mpos_TCCACGTTCGTAGCTA-1dCA2_MI_GFPve_pos  1.9095387630  0.8737879120
## dCA2_Mpos_TCCAGAACAAAGGGCT-1dCA2_MI_GFPve_pos  1.8614897899  0.7829027496
## dCA2_Mpos_TCCATCGCAAGGGCAT-1dCA2_MI_GFPve_pos  1.5108546190 -0.0331020035
## dCA2_Mpos_TCCATCGCAGGGCTTC-1dCA2_MI_GFPve_pos  2.8008771114  0.7530780159
## dCA2_Mpos_TCCATGCCAGACCAGA-1dCA2_MI_GFPve_pos  3.2158937626 -0.3826164879
## dCA2_Mpos_TCCCACATCTTTGCAT-1dCA2_MI_GFPve_pos  3.3485124283  0.9811454140
## dCA2_Mpos_TCCCAGTAGCACAAAT-1dCA2_MI_GFPve_pos  2.0471570663 -0.4506883301
## dCA2_Mpos_TCCCAGTGTAGGAAAG-1dCA2_MI_GFPve_pos  3.3738053016 -0.4814343132
## dCA2_Mpos_TCCCAGTGTGGCTTAT-1dCA2_MI_GFPve_pos  1.9494140320  0.1099901520
## dCA2_Mpos_TCCCAGTTCTACTGAG-1dCA2_MI_GFPve_pos  2.1634752922  1.7393918358
## dCA2_Mpos_TCCCATGAGGGAGTTC-1dCA2_MI_GFPve_pos  2.8561227493 -1.3138518013
## dCA2_Mpos_TCCCATGTCCTACGAA-1dCA2_MI_GFPve_pos  2.0894015007  1.4569468818
## dCA2_Mpos_TCCGAAAAGAACGCGT-1dCA2_MI_GFPve_pos  2.9571347408 -0.0928554215
## dCA2_Mpos_TCCGATCCAGTTTGGT-1dCA2_MI_GFPve_pos  2.6929857902  0.8298297249
## dCA2_Mpos_TCCGATCTCGGTGAAG-1dCA2_MI_GFPve_pos  1.3593621425  1.3623791061
## dCA2_Mpos_TCCGTGTTCTAACGGT-1dCA2_MI_GFPve_pos  2.0290558510  1.6429014526
## dCA2_Mpos_TCCTCGAGTCCCGGTA-1dCA2_MI_GFPve_pos  3.0981366806 -0.2260160126
## dCA2_Mpos_TCCTCTTCAGATCACT-1dCA2_MI_GFPve_pos  3.2120928936  1.2815323196
## dCA2_Mpos_TCCTCTTTCACGATAC-1dCA2_MI_GFPve_pos  1.8745432071  0.2770967804
## dCA2_Mpos_TCCTGCAAGCCAAGCA-1dCA2_MI_GFPve_pos  1.3867913656  1.1555996261
```

```
## dCA2_Mpos_TCCTTTCCATGCCATA-1dCA2_MI_GFPve_pos  2.3040087395  0.7800321899
## dCA2_Mpos_TCGAACAAGGCAGCTA-1dCA2_MI_GFPve_pos  2.8176782303  0.2443642937
## dCA2_Mpos_TCGCACTCATCGGCCA-1dCA2_MI_GFPve_pos  3.2095935516  0.4651251159
## dCA2_Mpos_TCGCACTGTGATTAGA-1dCA2_MI_GFPve_pos  2.2856769733  0.9582500778
## dCA2_Mpos_TCGCAGGCAGATACTC-1dCA2_MI_GFPve_pos  2.2588257961  1.5970831238
## dCA2_Mpos_TCGCAGGGTGACACAG-1dCA2_MI_GFPve_pos  2.5459084682 -1.3205532707
## dCA2_Mpos_TCGGATAAGAGTTCGG-1dCA2_MI_GFPve_pos  2.2061004810  2.9253526054
## dCA2_Mpos_TCGGATATCGAACTCA-1dCA2_MI_GFPve_pos  3.5096867256  0.3246531807
## dCA2_Mpos_TCGGGACCACAAAGTA-1dCA2_MI_GFPve_pos  1.2034064703  2.1891985260
## dCA2_Mpos_TCGGGCAGTGCCCGTA-1dCA2_MI_GFPve_pos  2.4044504337 -0.7351126351
## dCA2_Mpos_TCGGGTGAGAAAGCGA-1dCA2_MI_GFPve_pos  1.5089887552  1.9803004585
## dCA2_Mpos_TCGTCCAAGTTGTCAC-1dCA2_MI_GFPve_pos  2.2049546413  1.7337146126
## dCA2_Mpos_TCGTGGGCAAGTCATC-1dCA2_MI_GFPve_pos  2.1937685184  1.9256015144
## dCA2_Mpos_TCTAACTGTACTCGCG-1dCA2_MI_GFPve_pos  2.4349017315  2.2313824020
## dCA2_Mpos_TCTACATTCACTCTTA-1dCA2_MI_GFPve_pos  2.5374963455  0.4807501159
## dCA2_Mpos_TCTACCGCAATCGCGC-1dCA2_MI_GFPve_pos  2.7472269707  0.6902394615
## dCA2_Mpos_TCTATACTCGAAGCAG-1dCA2_MI_GFPve_pos  1.9141440563  1.8094654403
## dCA2_Mpos_TCTATCAAGAGCCTGA-1dCA2_MI_GFPve_pos  3.2611103229 -0.0471172013
## dCA2_Mpos_TCTCAGCAGAACAGGA-1dCA2_MI_GFPve_pos  2.3378181629 -0.0439858116
## dCA2_Mpos_TCTCAGCCAACTCCCT-1dCA2_MI_GFPve_pos  1.1008906536  1.0822873436
## dCA2_Mpos_TCTCAGCCAGCAATTC-1dCA2_MI_GFPve_pos  2.6488065891  0.3212166153
## dCA2_Mpos_TCTCCGATCTCGACCT-1dCA2_MI_GFPve_pos  3.2446670704  0.8730025612
## dCA2_Mpos_TCTCCGATCTGGGCAC-1dCA2_MI_GFPve_pos  2.5140962772  1.8989696823
## dCA2_Mpos_TCTGCCAAGAGTTGTA-1dCA2_MI_GFPve_pos  2.8330488376  2.3454785667
## dCA2_Mpos_TCTGTCGCACGTTGGC-1dCA2_MI_GFPve_pos  1.7548712663  2.2439232193
## dCA2_Mpos_TCTGTCGGTGGTCTCG-1dCA2_MI_GFPve_pos  3.0773530178  1.2318444572
## dCA2_Mpos_TCTGTCGTCGGTTGTA-1dCA2_MI_GFPve_pos  3.0094759636  1.8195672355
## dCA2_Mpos_TCTTAGTAGGAATTAC-1dCA2_MI_GFPve_pos  2.0012664966  2.6313209854
## dCA2_Mpos_TCTTAGTAGGTTCCAT-1dCA2_MI_GFPve_pos  1.3174259834  0.8344235740
## dCA2_Mpos_TCTTTGATCCCAAGCG-1dCA2_MI_GFPve_pos  1.7637803726  1.9029279075
## dCA2_Mpos_TGAACGTGTCGCCTAG-1dCA2_MI_GFPve_pos  2.3904433422  2.6961484276
## dCA2_Mpos_TGAACGTTCCGTCAAA-1dCA2_MI_GFPve_pos  2.9505210094 -0.2568306603
## dCA2_Mpos_TGACGCGGTACAAACA-1dCA2_MI_GFPve_pos  2.2866046600  1.5477733932
## dCA2_Mpos_TGACTCCGTATGACAA-1dCA2_MI_GFPve_pos  2.9551885299 -0.2306651749
## dCA2_Mpos_TGAGCGCAGGTGGTTG-1dCA2_MI_GFPve_pos  1.0785155468  1.6079087578
## dCA2_Mpos_TGAGCGCGTTGCCGAC-1dCA2_MI_GFPve_pos  2.4721312694  2.6117301307
## dCA2_Mpos_TGAGGTTGTCACGACC-1dCA2_MI_GFPve_pos  2.4188709430  1.7786441170
## dCA2_Mpos_TGAGTCATCTAGAACC-1dCA2_MI_GFPve_pos  2.3374128513  2.2132120453
## dCA2_Mpos_TGATCAGAGACTGAGC-1dCA2_MI_GFPve_pos  2.3649757080  2.0977673851
## dCA2_Mpos_TGATCAGGTCTGCCTT-1dCA2_MI_GFPve_pos  2.2117381267  1.2433829628
## dCA2_Mpos_TGATCAGTCAGCACCG-1dCA2_MI_GFPve_pos  2.5108840637  2.7116380058
## dCA2_Mpos_TGATCAGTCGCGGACT-1dCA2_MI_GFPve_pos  0.5078604393  2.2156811081
## dCA2_Mpos_TGATCAGTCGTGAGAG-1dCA2_MI_GFPve_pos  2.6813757591  1.5657949768
## dCA2_Mpos_TGATCTTCACCAGCCA-1dCA2_MI_GFPve_pos  1.1964788608  1.8684821449
## dCA2_Mpos_TGATGGTCAGTCGTTA-1dCA2_MI_GFPve_pos  2.4517584019  1.1103163086
## dCA2_Mpos_TGATGGTTCATCCCGT-1dCA2_MI_GFPve_pos  2.6032767467  1.8160753570
## dCA2_Mpos_TGATTTCCAGTCGGAA-1dCA2_MI_GFPve_pos  1.5060693197  3.2820825897
## dCA2_Mpos_TGCACGGTCTCCGATC-1dCA2_MI_GFPve_pos  2.4131202869  2.4804373108
## dCA2_Mpos_TGCAGATTCCATCTCG-1dCA2_MI_GFPve_pos  0.7337541752 -1.6126198448
## dCA2_Mpos_TGCAGGCAGGAACATT-1dCA2_MI_GFPve_pos  2.5169270210 -1.6969296612
## dCA2_Mpos_TGCAGGCTCTCAACCC-1dCA2_MI_GFPve_pos  3.2188270263 -0.9097866692
## dCA2_Mpos_TGCAGTAAGACCACGA-1dCA2_MI_GFPve_pos  3.3536994629  1.1948318802
## dCA2_Mpos_TGCAGTAAGAGCCATG-1dCA2_MI_GFPve_pos  0.8381010465  1.2812009178
## dCA2_Mpos_TGCATCCCACAAATCC-1dCA2_MI_GFPve_pos  3.4957652263 -0.1172370590
## dCA2_Mpos_TGCGACGGTACGTTCA-1dCA2_MI_GFPve_pos  2.9957819156  2.0313725792
## dCA2_Mpos_TGCGATATCTTGATTC-1dCA2_MI_GFPve_pos  2.8733143978 -1.0630290188
## dCA2_Mpos_TGCGGCATCACTAGCA-1dCA2_MI_GFPve_pos  3.5013282471  0.2951932274
## dCA2_Mpos_TGCGGCATCAGAATAG-1dCA2_MI_GFPve_pos  2.7554018669 -1.1257602848
## dCA2_Mpos_TGCTGAACACACCAGC-1dCA2_MI_GFPve_pos -0.3915487803  0.2853207908
## dCA2_Mpos_TGCTTCGTCCATATGG-1dCA2_MI_GFPve_pos  1.1621717386  0.8839564644
## dCA2_Mpos_TGGAACTTCAAGCGTT-1dCA2_MI_GFPve_pos  0.5984112792  1.3107767425
## dCA2_Mpos_TGGAGGACAATGAACA-1dCA2_MI_GFPve_pos  2.7463724785  0.4706907593
## dCA2_Mpos_TGGATCACATGTCGTA-1dCA2_MI_GFPve_pos  1.3372883968  2.3489480339
## dCA2_Mpos_TGGCGTGAGGCTAAAT-1dCA2_MI_GFPve_pos  2.3415322475  2.9072761856
## dCA2_Mpos_TGGGATTTCAACGTGT-1dCA2_MI_GFPve_pos  2.3647909336  1.5462670646
## dCA2_Mpos_TGGGCGTTCCTCTCGA-1dCA2_MI_GFPve_pos  2.4508111648  2.2116122566
## dCA2_Mpos_TGGGCTGCACCGTCTT-1dCA2_MI_GFPve_pos  2.2244265251  1.6600346885
## dCA2_Mpos_TGGGCTGGTGACCGAA-1dCA2_MI_GFPve_pos  2.6168966465  1.5018453918
## dCA2_Mpos_TGGGCTGGTTCCTTGC-1dCA2_MI_GFPve_pos  2.7952833347  1.2821560226
## dCA2_Mpos_TGGGTTAGTCCCTAAA-1dCA2_MI_GFPve_pos  3.3068854980 -0.8793096222
```

```
## dCA2_Mpos_TGGTAGTGTCCACAGC-1dCA2_MI_GFPve_pos  2.6048708133  2.5917477928
## dCA2_Mpos_TGGTTAGCACGCGCAT-1dCA2_MI_GFPve_pos  2.0423901253 -0.1713299431
## dCA2_Mpos_TGGTTAGCAGGCGAAT-1dCA2_MI_GFPve_pos  2.4307773285  1.5354280792
## dCA2_Mpos_TGGTTAGGTTAGAAAC-1dCA2_MI_GFPve_pos  3.4779899292  1.0562711082
## dCA2_Mpos_TGGTTAGTCCTTGGAA-1dCA2_MI_GFPve_pos  2.0056750946  1.9750700317
## dCA2_Mpos_TGGTTAGTCGTGGAAG-1dCA2_MI_GFPve_pos  2.5746171646  2.1420445762
## dCA2_Mpos_TGTAACGCATATCTCT-1dCA2_MI_GFPve_pos  2.5743694477 -0.3568806328
## dCA2_Mpos_TGTAACGGTGGATCAG-1dCA2_MI_GFPve_pos  1.8378162555 -1.8709885754
## dCA2_Mpos_TGTAAGCAGTAGTCTC-1dCA2_MI_GFPve_pos  2.9475276642  1.2355242096
## dCA2_Mpos_TGTAAGCGTAGCCCTG-1dCA2_MI_GFPve_pos  2.3401453666  0.8729124389
## dCA2_Mpos_TGTACAGCATGGAGAC-1dCA2_MI_GFPve_pos  1.9262151890  2.1272864662
## dCA2_Mpos_TGTAGACGTCGCATGC-1dCA2_MI_GFPve_pos  3.4242031746  0.6859016739
## dCA2_Mpos_TGTCAGAGTTTACTTC-1dCA2_MI_GFPve_pos  1.6717462711  3.3032369934
## dCA2_Mpos_TGTCCACGTCATATGC-1dCA2_MI_GFPve_pos  2.0296619110  2.1848712288
## dCA2_Mpos_TGTCCACTCTTGGTCC-1dCA2_MI_GFPve_pos  1.8430640869 -0.2100667633
## dCA2_Mpos_TGTCCTGAGAATTCAG-1dCA2_MI_GFPve_pos  2.5142707996  0.8247514091
## dCA2_Mpos_TGTCCTGCACCATTCC-1dCA2_MI_GFPve_pos  2.3967924290  0.4407821022
## dCA2_Mpos_TGTCCTGGTACCCGCA-1dCA2_MI_GFPve_pos  2.2112016849  1.9418645225
## dCA2_Mpos_TGTGAGTAGCATCTTG-1dCA2_MI_GFPve_pos  3.1936981849 -0.4438337959
## dCA2_Mpos_TGTGAGTCACATACGT-1dCA2_MI_GFPve_pos  1.6111186914  1.5340662323
## dCA2_Mpos_TGTGCGGGTCATATGC-1dCA2_MI_GFPve_pos  1.1469323807  2.0477142654
## dCA2_Mpos_TGTGGCGTCTGTGCGG-1dCA2_MI_GFPve_pos  2.3071360760  2.9689050041
## dCA2_Mpos_TGTTACTTCTATTCGT-1dCA2_MI_GFPve_pos  1.5196165018  2.2181969009
## dCA2_Mpos_TGTTCCGGTACTCGCG-1dCA2_MI_GFPve_pos  2.9682312183  1.2470808349
## dCA2_Mpos_TGTTCCGTCCATTGGA-1dCA2_MI_GFPve_pos  0.2652167551  2.4192676864
## dCA2_Mpos_TTACAGGTCTATTGTC-1dCA2_MI_GFPve_pos  2.4666955643 -1.1519055046
## dCA2_Mpos_TTACCGCCACACTTAG-1dCA2_MI_GFPve_pos  2.6277399235 -1.2448630013
## dCA2_Mpos_TTACCGCTCGGAATGG-1dCA2_MI_GFPve_pos  2.3749096565  0.2363548599
## dCA2_Mpos_TTACGCCTCCCGAACG-1dCA2_MI_GFPve_pos  0.8253157906  1.1814561210
## dCA2_Mpos_TTACGTTAGAAAGTCT-1dCA2_MI_GFPve_pos  1.5836955480  2.6527567230
## dCA2_Mpos_TTAGGCAAGCATGCAG-1dCA2_MI_GFPve_pos  3.3077092342 -0.4348897614
## dCA2_Mpos_TTAGGCAGTGACAACG-1dCA2_MI_GFPve_pos  2.9050386123  0.3116164528
## dCA2_Mpos_TTAGGGTGTAGTCTTG-1dCA2_MI_GFPve_pos  3.5011770897  1.4093709312
## dCA2_Mpos_TTAGGGTTCTGAGGCC-1dCA2_MI_GFPve_pos  2.0550854378  0.5734510742
## dCA2_Mpos_TTAGTCTAGTCAGCCC-1dCA2_MI_GFPve_pos  2.5476827793 -0.9199340023
## dCA2_Mpos_TTATTGCCAACACACT-1dCA2_MI_GFPve_pos  2.9569935970 -0.7637042679
## dCA2_Mpos_TTCAATCCAGGAAGTC-1dCA2_MI_GFPve_pos  2.9693923168  0.8784389816
## dCA2_Mpos_TTCAATCGTGGCATCC-1dCA2_MI_GFPve_pos  2.8615529709  1.8602156959
## dCA2_Mpos_TTCACGCAGACCAAGC-1dCA2_MI_GFPve_pos  2.0287888222  0.0836778007
## dCA2_Mpos_TTCACGCCAACAGCTT-1dCA2_MI_GFPve_pos  2.0891966991  0.8825078331
## dCA2_Mpos_TTCACGCGTTATGTGC-1dCA2_MI_GFPve_pos  3.2783427410  0.9756555877
## dCA2_Mpos_TTCACGCTCCGATAGT-1dCA2_MI_GFPve_pos  3.2810225658 -0.4152884163
## dCA2_Mpos_TTCATGTCAATCACGT-1dCA2_MI_GFPve_pos  2.8009014301 -0.2661242165
## dCA2_Mpos_TTCATGTTCCGCTGTT-1dCA2_MI_GFPve_pos  2.4397180252  1.8180408798
## dCA2_Mpos_TTCCAATCAGCGTTGC-1dCA2_MI_GFPve_pos  2.3184735946  2.6372371040
## dCA2_Mpos_TTCCAATTCATGAGAA-1dCA2_MI_GFPve_pos  2.9087984733  1.7151418052
## dCA2_Mpos_TTCCACGAGAGACAAG-1dCA2_MI_GFPve_pos  1.9065540008  2.0652842842
## dCA2_Mpos_TTCCACGTCCAGGACC-1dCA2_MI_GFPve_pos  2.7950914078  0.5214529358
## dCA2_Mpos_TTCCGGTTCACGATCA-1dCA2_MI_GFPve_pos  2.6095879249 -0.4446944870
## dCA2_Mpos_TTCCTTCAGAGAGCGG-1dCA2_MI_GFPve_pos  2.2954187565  2.8062682472
## dCA2_Mpos_TTCGCTGAGCCATTCA-1dCA2_MI_GFPve_pos  2.6807029419  2.3753962837
## dCA2_Mpos_TTCGGTCCAAAGACTA-1dCA2_MI_GFPve_pos  2.2892210655 -1.2402252831
## dCA2_Mpos_TTCGGTCCACGACCTG-1dCA2_MI_GFPve_pos  3.2100305729  0.8253193222
## dCA2_Mpos_TTCTAACTCCATTCGC-1dCA2_MI_GFPve_pos  2.2130541973  0.8908696495
## dCA2_Mpos_TTCTCTCTCGTAATGC-1dCA2_MI_GFPve_pos  2.6809013061 -1.4381374993
## dCA2_Mpos_TTCTTCCCAACCTAAC-1dCA2_MI_GFPve_pos  2.2318957023  1.9874987922
## dCA2_Mpos_TTCTTCCTCGGCATCG-1dCA2_MI_GFPve_pos  0.9522875719  1.1289525352
## dCA2_Mpos_TTGAACGGTTACGCCG-1dCA2_MI_GFPve_pos  1.9199619465  1.6886487327
## dCA2_Mpos_TTGAGTGCACAGAGCA-1dCA2_MI_GFPve_pos  2.7866077595  0.1610861145
## dCA2_Mpos_TTGATGGAGTTGCGCC-1dCA2_MI_GFPve_pos  2.8054948025  1.3231606804
## dCA2_Mpos_TTGCATTAGAAGCCAC-1dCA2_MI_GFPve_pos  1.9428236656 -0.3532247223
## dCA2_Mpos_TTGCCTGAGCTCCACG-1dCA2_MI_GFPve_pos  1.5364617281  2.7258334480
## dCA2_Mpos_TTGCCTGCACCGAATT-1dCA2_MI_GFPve_pos  2.9491293602 -1.1655952610
## dCA2_Mpos_TTGCGTCAGACCATGG-1dCA2_MI_GFPve_pos  1.8968105488  1.5522475563
## dCA2_Mpos_TTGGATGAGTCGAAAT-1dCA2_MI_GFPve_pos  3.1694626980  1.6093321167
## dCA2_Mpos_TTGGGATCAATCGAAA-1dCA2_MI_GFPve_pos  2.1004479103  3.0601992927
## dCA2_Mpos_TTGGGATTCAAGTCGT-1dCA2_MI_GFPve_pos  2.8957643680  2.2450986229
## dCA2_Mpos_TTGGGCGAGTCACGAG-1dCA2_MI_GFPve_pos  2.4657604866  2.4768667541
## dCA2_Mpos_TTGGGTACAAGAGGCT-1dCA2_MI_GFPve_pos  2.3169875316  2.6481142364
## dCA2_Mpos_TTGGGTAGTAACCCGC-1dCA2_MI_GFPve_pos  3.5231695347  1.2152448021
```

```
## dCA2_Mpos_TTGGGTAGTCTACGTA-1dCA2_MI_GFPve_pos  2.6819994621  2.3445840202
## dCA2_Mpos_TTGGTTTCATACCAGT-1dCA2_MI_GFPve_pos  1.6747330599  1.4810567222
## dCA2_Mpos_TTGGTTTGTAGCGCCT-1dCA2_MI_GFPve_pos  2.1827621631  2.2509250961
## dCA2_Mpos_TTGTGGATCGTGTCAA-1dCA2_MI_GFPve_pos  2.8627014332  1.1815467201
## dCA2_Mpos_TTGTTGTAGACATCAA-1dCA2_MI_GFPve_pos  2.4907252960  2.1737313591
## dCA2_Mpos_TTTACCAAGGTTAAAC-1dCA2_MI_GFPve_pos  1.8896923237  0.0853581749
## dCA2_Mpos_TTTACCAGTCGATTAC-1dCA2_MI_GFPve_pos  2.3914036922  0.3861551605
## dCA2_Mpos_TTTACGTTCGCACGAC-1dCA2_MI_GFPve_pos  1.5969605617 -2.6445910610
## dCA2_Mpos_TTTACTGTCCATAGGT-1dCA2_MI_GFPve_pos  2.9928505592 -0.0359463371
## dCA2_Mpos_TTTAGTCCACATATGC-1dCA2_MI_GFPve_pos  2.6452870540 -0.0323576607
## dCA2_Mpos_TTTCACAAGCTGAGCA-1dCA2_MI_GFPve_pos  3.1048922710  1.8645687424
## dCA2_Mpos_TTTCACAAGTTAGTGA-1dCA2_MI_GFPve_pos  2.7124908142  0.1929750763
## dCA2_Mpos_TTTCACACAAACCACT-1dCA2_MI_GFPve_pos  2.4590187244  1.8055191360
## dCA2_Mpos_TTTCAGTGTATTGGCT-1dCA2_MI_GFPve_pos  1.4097788505  1.9995165191
## dCA2_Mpos_TTTCAGTTCCCTCATG-1dCA2_MI_GFPve_pos  2.0155909233 -0.4829296746
## dCA2_Mpos_TTTCATGAGCACGTCC-1dCA2_MI_GFPve_pos  1.2955149584  1.5210252128
## dCA2_Mpos_TTTCATGCAACGATTC-1dCA2_MI_GFPve_pos  3.2021949463  0.8778653465
## dCA2_Mpos_TTTCATGCAGCATACT-1dCA2_MI_GFPve_pos  2.0589115791 -1.6540600933
## dCA2_Mpos_TTTCATGCATCGCCTT-1dCA2_MI_GFPve_pos  2.3236608677  1.4322233520
## dCA2_Mpos_TTTCCTCCAGAGCGTA-1dCA2_MI_GFPve_pos  2.2992394142  2.8428283058
## dCA2_Mpos_TTTCGATGTCTGTAAC-1dCA2_MI_GFPve_pos  1.6874505453  2.9583587967
## dCA2_Mpos_TTTCGATTCTACAGGT-1dCA2_MI_GFPve_pos  2.1937313251  1.8461199127
## dCA2_Fpos_AAAGGTATCCGATAAC-1dCA2_FI_GFPve_pos  0.9277416639  1.3655681930
## dCA2_Fpos_AAAGTGACACAACGCC-1dCA2_FI_GFPve_pos  1.7409376077  2.7181849800
## dCA2_Fpos_AAATGGACAATACGCT-1dCA2_FI_GFPve_pos  3.2033126526  0.8282184921
## dCA2_Fpos_AAATGGACAGCTATAC-1dCA2_FI_GFPve_pos  2.6838081054 -0.8344294705
## dCA2_Fpos_AAATGGATCCTACTGC-1dCA2_FI_GFPve_pos  2.6913688354 -0.1544398941
## dCA2_Fpos_AACAAAGCATGCACTA-1dCA2_FI_GFPve_pos  2.2315776520 -0.0152187027
## dCA2_Fpos_AACAAAGTCGCGGTAC-1dCA2_FI_GFPve_pos  2.5099797420  2.6234093033
## dCA2_Fpos_AACACACGTCAAGCGA-1dCA2_FI_GFPve_pos  3.3124334984  1.4808860145
## dCA2_Fpos_AACACACGTCATAAAG-1dCA2_FI_GFPve_pos  2.5487864189 -0.8526303448
## dCA2_Fpos_AACCATGCAATAGAGT-1dCA2_FI_GFPve_pos  2.8299906425  2.2020893417
## dCA2_Fpos_AACCATGGTGTTGACT-1dCA2_FI_GFPve_pos  2.5628178291  1.7801676117
## dCA2_Fpos_AACCTTTTCTGCTTAT-1dCA2_FI_GFPve_pos  1.5712942056  1.9866209350
## dCA2_Fpos_AACGAAAGTTCAGTAC-1dCA2_FI_GFPve_pos  2.2589745693  1.4606705032
## dCA2_Fpos_AAGACAAGTATCGCTA-1dCA2_FI_GFPve_pos -0.6445424385  1.5124965034
## dCA2_Fpos_AAGACTCGTGTCTTCC-1dCA2_FI_GFPve_pos -0.3027651586  1.8383756004
## dCA2_Fpos_AAGCGTTGTTGCATGT-1dCA2_FI_GFPve_pos  1.6718249492  1.4594750725
## dCA2_Fpos_AATCGACGTTAGTTCG-1dCA2_FI_GFPve_pos  2.2571971588  1.8648901306
## dCA2_Fpos_AATTCCTGTCTCGACG-1dCA2_FI_GFPve_pos  2.4280982189  1.3646765075
## dCA2_Fpos_AATTTCCGTACTAACC-1dCA2_FI_GFPve_pos  2.7364261322  2.4617657982
## dCA2_Fpos_ACAAAGACAAGAATGT-1dCA2_FI_GFPve_pos  2.8968851738 -0.1524476685
## dCA2_Fpos_ACAAGCTCAATCTCTT-1dCA2_FI_GFPve_pos  3.1328504257  1.2584143005
## dCA2_Fpos_ACACAGTAGACTCTAC-1dCA2_FI_GFPve_pos  2.0401365928  1.2706075035
## dCA2_Fpos_ACACTGAGTCTGCGCA-1dCA2_FI_GFPve_pos  2.8859968357  0.4498153053
## dCA2_Fpos_ACAGAAACAGAAATCA-1dCA2_FI_GFPve_pos  0.2476703696  2.0075407348
## dCA2_Fpos_ACAGAAAGTTGCGAAG-1dCA2_FI_GFPve_pos  3.3470952682 -0.6721267380
## dCA2_Fpos_ACAGCCGGTCATTCCC-1dCA2_FI_GFPve_pos  2.8578968220 -0.3336410202
## dCA2_Fpos_ACATCCCTCGCAAGAG-1dCA2_FI_GFPve_pos  1.7234622412  0.7822385155
## dCA2_Fpos_ACCACAAGTGTTCGAT-1dCA2_FI_GFPve_pos  2.3582246475  0.6018939338
## dCA2_Fpos_ACCACAAGTTTCCAAG-1dCA2_FI_GFPve_pos  2.9297633343 -1.0075368561
## dCA2_Fpos_ACCCAAACAGTCTACA-1dCA2_FI_GFPve_pos  2.1754198246  0.2137794815
## dCA2_Fpos_ACCCAAACATTGCTTT-1dCA2_FI_GFPve_pos  3.1294357948  0.7930741630
## dCA2_Fpos_ACCCTTGGTCAAAGTA-1dCA2_FI_GFPve_pos  1.3347346954  1.0404424988
## dCA2_Fpos_ACCTACCAGTAACAGT-1dCA2_FI_GFPve_pos  3.2075960808  1.2685752235
## dCA2_Fpos_ACCTGAACACGCACCA-1dCA2_FI_GFPve_pos  1.8650074177 -0.0123686470
## dCA2_Fpos_ACGATGTCAGGTTTAC-1dCA2_FI_GFPve_pos  2.5426778965  1.7153282486
## dCA2_Fpos_ACGTTCCTCTTCGGAA-1dCA2_FI_GFPve_pos  2.7223920994  2.1392207466
## dCA2_Fpos_ACTATCTGTACTGCGC-1dCA2_FI_GFPve_pos  2.7647497825  0.5983987175
## dCA2_Fpos_ACTCCCAAGAATCGTA-1dCA2_FI_GFPve_pos  3.4303202801  1.4717660270
## dCA2_Fpos_ACTGATGCAACGATTC-1dCA2_FI_GFPve_pos  2.0879504852  1.0609174095
## dCA2_Fpos_ACTGCAAGTTGAAGTA-1dCA2_FI_GFPve_pos  3.0451131038  1.2534842811
## dCA2_Fpos_ACTGTGATCCGTTGAA-1dCA2_FI_GFPve_pos  1.1049498491  1.8205943428
## dCA2_Fpos_ACTTATCAGGAGGTTC-1dCA2_FI_GFPve_pos  3.0538985901  1.5438027702
## dCA2_Fpos_ACTTCCGGTTGCAACT-1dCA2_FI_GFPve_pos  3.4738192730  0.5064544998
## dCA2_Fpos_ACTTTCAGTGTTGCCG-1dCA2_FI_GFPve_pos  1.3189863138 -0.0641269364
## dCA2_Fpos_ACTTTGTCAGGAGGTT-1dCA2_FI_GFPve_pos  2.2261495762  2.9104318939
## dCA2_Fpos_AGAACAAAGTCGGCAA-1dCA2_FI_GFPve_pos  2.9019191437  0.6504674278
## dCA2_Fpos_AGACAAAAGACATAAC-1dCA2_FI_GFPve_pos  2.6532683544  1.1723637901
## dCA2_Fpos_AGACAGGCACAGCCTG-1dCA2_FI_GFPve_pos  2.6564524345  1.4592204414
```

```
## dCA2_Fpos_AGACAGGGTCCCTGTT-1dCA2_FI_GFPve_pos  2.8319316082  0.0992703758
## dCA2_Fpos_AGATCCAGTTTGGGTT-1dCA2_FI_GFPve_pos  1.4534154110  0.6755743347
## dCA2_Fpos_AGATGAAAGTATGATG-1dCA2_FI_GFPve_pos  1.2079131775  2.5802483879
## dCA2_Fpos_AGCATCACAGGAATCG-1dCA2_FI_GFPve_pos  2.4050300293  1.3300009094
## dCA2_Fpos_AGCGCTGAGTTATGGA-1dCA2_FI_GFPve_pos  0.4761809282  1.6003175102
## dCA2_Fpos_AGCTCAAAGGTTGGTG-1dCA2_FI_GFPve_pos  1.8227980308  2.6605887733
## dCA2_Fpos_AGGACGAGTCCTACAA-1dCA2_FI_GFPve_pos  1.5952016287  1.4412989937
## dCA2_Fpos_AGGCTGCTCCCATTCG-1dCA2_FI_GFPve_pos  2.2326014213 -0.9592392124
## dCA2_Fpos_AGGGAGTTCTTACCAT-1dCA2_FI_GFPve_pos  2.4964285068  0.8983245216
## dCA2_Fpos_AGGGCCTTCCGCCTAT-1dCA2_FI_GFPve_pos  1.2586001329  0.3313598953
## dCA2_Fpos_AGGTAGGTCAAGAGTA-1dCA2_FI_GFPve_pos  2.0130310230  2.2746749244
## dCA2_Fpos_AGGTCTACAGCTACTA-1dCA2_FI_GFPve_pos  3.2548666172  1.2828998886
## dCA2_Fpos_AGGTGTTGTAGACAGC-1dCA2_FI_GFPve_pos  2.7663266830  2.4477591835
## dCA2_Fpos_AGGTTACAGTATTAGG-1dCA2_FI_GFPve_pos  2.4736168556  2.1155972801
## dCA2_Fpos_AGTACCATCCTACCGT-1dCA2_FI_GFPve_pos  2.8346908264  2.0597796760
## dCA2_Fpos_AGTACTGCATATACCG-1dCA2_FI_GFPve_pos  2.1446390323  2.2095260940
## dCA2_Fpos_AGTACTGTCTTGGCTC-1dCA2_FI_GFPve_pos  2.4563388996 -0.1559562363
## dCA2_Fpos_AGTCTCCCAAGACCGA-1dCA2_FI_GFPve_pos  0.6480754189 -3.3695079960
## dCA2_Fpos_AGTGACTAGGAGATAG-1dCA2_FI_GFPve_pos  2.4460611515  1.4723344169
## dCA2_Fpos_AGTGTTGCAACTCCCT-1dCA2_FI_GFPve_pos  0.8932912521  1.3462848984
## dCA2_Fpos_AGTTAGCGTAGAGGAA-1dCA2_FI_GFPve_pos  2.5106325321  2.3398204170
## dCA2_Fpos_AGTTCCCCACACGGAA-1dCA2_FI_GFPve_pos  1.3341281585  2.0843043648
## dCA2_Fpos_AGTTCGAGTTTGGAGG-1dCA2_FI_GFPve_pos  2.3232655697  2.1985841118
## dCA2_Fpos_ATACCTTAGGCACGAT-1dCA2_FI_GFPve_pos  2.5450809173  0.7250891052
## dCA2_Fpos_ATACTTCGTACGTGTT-1dCA2_FI_GFPve_pos  1.6311417990  2.1439986549
## dCA2_Fpos_ATCCTATAGGTTCTTG-1dCA2_FI_GFPve_pos  2.3096854858  0.3441162430
## dCA2_Fpos_ATCGATGCATGCCGAC-1dCA2_FI_GFPve_pos  2.0768354111  0.1985712372
## dCA2_Fpos_ATCGCCTCAATAGTCC-1dCA2_FI_GFPve_pos  1.3357035093  1.1890888534
## dCA2_Fpos_ATCTCTAAGGACATCG-1dCA2_FI_GFPve_pos  2.0509736709  2.4179988228
## dCA2_Fpos_ATGAAAGGTACTGGGA-1dCA2_FI_GFPve_pos  2.4374351673  1.5261440597
## dCA2_Fpos_ATGACCACACCATAAC-1dCA2_FI_GFPve_pos  1.4134782724  1.5375972114
## dCA2_Fpos_ATGAGTCTCCATTGTT-1dCA2_FI_GFPve_pos  1.0392432384  1.5512290321
## dCA2_Fpos_ATGGATCAGTCCTGTA-1dCA2_FI_GFPve_pos  1.2291828565  2.1636543594
## dCA2_Fpos_ATGTCTTTCATTACGG-1dCA2_FI_GFPve_pos  2.4528625183  0.5023608528
## dCA2_Fpos_ATTACCTCACTTGTCC-1dCA2_FI_GFPve_pos  0.9568499498  1.7365885101
## dCA2_Fpos_ATTCTACAGGGTTAGC-1dCA2_FI_GFPve_pos -0.3522459544  0.5537200294
## dCA2_Fpos_ATTGTTCCACGCGCAT-1dCA2_FI_GFPve_pos  2.0232072048  2.1425390564
## dCA2_Fpos_ATTTACCAGGTAGTCG-1dCA2_FI_GFPve_pos  2.1112425499  1.9978022896
## dCA2_Fpos_ATTTCACGTCTTACTT-1dCA2_FI_GFPve_pos  1.4112606220 -1.9476742424
## dCA2_Fpos_ATTTCTGTCAACCTTT-1dCA2_FI_GFPve_pos  3.1016125850 -1.0011665501
## dCA2_Fpos_CAACAGTTCTCATAGG-1dCA2_FI_GFPve_pos  1.1792893581  1.2634439789
## dCA2_Fpos_CAACCTCAGACTAAGT-1dCA2_FI_GFPve_pos  2.9543953113 -1.1696490921
## dCA2_Fpos_CAACCTCAGTAGACCG-1dCA2_FI_GFPve_pos  2.7545674019 -0.0661673226
## dCA2_Fpos_CAAGGGAGTCGTTCAA-1dCA2_FI_GFPve_pos  1.9736972027  2.5581932388
## dCA2_Fpos_CAATGACTCTGGGCAC-1dCA2_FI_GFPve_pos  0.7513826661  1.7344928108
## dCA2_Fpos_CAATTTCAGCCTGTGC-1dCA2_FI_GFPve_pos  3.2840242557  1.1415224396
## dCA2_Fpos_CACACAATCCAACACA-1dCA2_FI_GFPve_pos  1.1755358152  2.1214723907
## dCA2_Fpos_CACTAAGAGGCTCAAG-1dCA2_FI_GFPve_pos  3.0401506595  1.2818384491
## dCA2_Fpos_CACTAAGCACTTGTGA-1dCA2_FI_GFPve_pos  2.0545849972 -0.8870832600
## dCA2_Fpos_CACTGGGGTGGATTTC-1dCA2_FI_GFPve_pos  2.0435517006  1.5321908317
## dCA2_Fpos_CACTGTCGTATGGAAT-1dCA2_FI_GFPve_pos  3.0817134552  2.1336565338
## dCA2_Fpos_CACTGTCTCTACCCAC-1dCA2_FI_GFPve_pos  2.1809239559  0.6150842033
## dCA2_Fpos_CAGATACCATGGCGCT-1dCA2_FI_GFPve_pos  2.4235027008  1.8090586983
## dCA2_Fpos_CAGATTGCACTCCCTA-1dCA2_FI_GFPve_pos  2.0254059009 -0.8018011727
## dCA2_Fpos_CAGCACGCATACAGCT-1dCA2_FI_GFPve_pos  1.3899064235  1.4503431640
## dCA2_Fpos_CAGGCCAGTGTGCTTA-1dCA2_FI_GFPve_pos  2.1870367698  1.0662741981
## dCA2_Fpos_CAGGCCATCGCGGTAC-1dCA2_FI_GFPve_pos  2.8286979370  2.1112375580
## dCA2_Fpos_CAGGTATCACCCAATA-1dCA2_FI_GFPve_pos  2.7048373394  1.4428100906
## dCA2_Fpos_CAGGTATTCTACACTT-1dCA2_FI_GFPve_pos  2.3363597564  1.9308715187
## dCA2_Fpos_CAGTTAGAGGCCTGAA-1dCA2_FI_GFPve_pos  2.6971085243  2.6002407394
## dCA2_Fpos_CAGTTCCAGCCAGTAG-1dCA2_FI_GFPve_pos  2.9329233341  2.0968976341
## dCA2_Fpos_CAGTTCCGTGAAGCGT-1dCA2_FI_GFPve_pos  2.6203806572 -0.6518134751
## dCA2_Fpos_CATCAAGAGAGTTGCG-1dCA2_FI_GFPve_pos  1.8281002216  1.5378542267
## dCA2_Fpos_CATCAAGAGATGGTAT-1dCA2_FI_GFPve_pos  1.3499719076  2.7596869789
## dCA2_Fpos_CATCCACCAGTCTACA-1dCA2_FI_GFPve_pos  2.2742176227  1.8181567512
## dCA2_Fpos_CATCCACCATAGTCAC-1dCA2_FI_GFPve_pos  1.2985609941  0.0574202858
## dCA2_Fpos_CATCCGTTCAAGTCGT-1dCA2_FI_GFPve_pos  1.9618418388  1.3810940109
## dCA2_Fpos_CATGAGTCACCATATG-1dCA2_FI_GFPve_pos  3.3571605854  0.8898125015
## dCA2_Fpos_CATTCCGCATTGCCGG-1dCA2_FI_GFPve_pos  2.4284548931  1.2611861549
## dCA2_Fpos_CATTGCCAGAACAAGG-1dCA2_FI_GFPve_pos  3.3415835075  1.6988544784
```

```
## dCA2_Fpos_CATTGCCTCCGATGTA-1dCA2_FI_GFPve_pos  3.2754328422 -0.5774993576
## dCA2_Fpos_CATTGTTAGGCGACAT-1dCA2_FI_GFPve_pos  1.6748150759  2.4914226852
## dCA2_Fpos_CCACCATAGTAGCATA-1dCA2_FI_GFPve_pos  1.8434403114  1.6437077843
## dCA2_Fpos_CCACGAGAGACGAAGA-1dCA2_FI_GFPve_pos  1.6336827450  2.4381146751
## dCA2_Fpos_CCCGGAAGTTAGAAGT-1dCA2_FI_GFPve_pos  2.6927361660  1.9981174789
## dCA2_Fpos_CCCTAACCACCTTCGT-1dCA2_FI_GFPve_pos  2.5944624119 -0.8548695721
## dCA2_Fpos_CCCTTAGAGCAAACAT-1dCA2_FI_GFPve_pos  2.5745361023  0.7042527519
## dCA2_Fpos_CCGAACGGTGGCAGAT-1dCA2_FI_GFPve_pos  1.7898000650  0.8344016395
## dCA2_Fpos_CCGATCTTCTAGTGTG-1dCA2_FI_GFPve_pos  1.1729842596  2.0422487579
## dCA2_Fpos_CCGGTGACACCACTGG-1dCA2_FI_GFPve_pos  1.3052997761  1.6569958053
## dCA2_Fpos_CCGGTGATCGTCCATC-1dCA2_FI_GFPve_pos  2.0350401573  1.7750478111
## dCA2_Fpos_CCGTAGGAGTTGCCCG-1dCA2_FI_GFPve_pos  3.3921609096  0.4795737587
## dCA2_Fpos_CCTAACCAGACCATTC-1dCA2_FI_GFPve_pos  3.0498531036 -1.0365032829
## dCA2_Fpos_CCTATCGAGACTGAGC-1dCA2_FI_GFPve_pos  1.0259410314  2.1050496422
## dCA2_Fpos_CCTATCGTCCTATTTG-1dCA2_FI_GFPve_pos  1.5324893169  1.6366439186
## dCA2_Fpos_CCTCAACCATTATGCG-1dCA2_FI_GFPve_pos  2.7990577392  2.1076398216
## dCA2_Fpos_CCTCATGTCAGAGTGG-1dCA2_FI_GFPve_pos  2.5916266613  1.7537679992
## dCA2_Fpos_CCTTGTGGTAACATCC-1dCA2_FI_GFPve_pos  0.5643590264  0.7389307342
## dCA2_Fpos_CCTTGTGTCTGTCCGT-1dCA2_FI_GFPve_pos  3.2808852367  0.2439990364
## dCA2_Fpos_CCTTTGGTCAAAGACA-1dCA2_FI_GFPve_pos  1.7473795585  1.0964541755
## dCA2_Fpos_CGAAGTTAGGGATCAC-1dCA2_FI_GFPve_pos  2.7351343803  2.0158124290
## dCA2_Fpos_CGAATTGAGCACGTCC-1dCA2_FI_GFPve_pos  1.3543015890  2.4302015625
## dCA2_Fpos_CGAGGCTAGCATGGGT-1dCA2_FI_GFPve_pos  1.7345653944  1.6544313751
## dCA2_Fpos_CGAGTTAAGAGGTCAC-1dCA2_FI_GFPve_pos  1.7768967323  0.9047432266
## dCA2_Fpos_CGATGGCCAGAGATGC-1dCA2_FI_GFPve_pos  2.0275755100  0.9457116447
## dCA2_Fpos_CGATGGCTCTTCTGTA-1dCA2_FI_GFPve_pos  2.2668798141  1.9390149437
## dCA2_Fpos_CGCATAAGTTGGCTAT-1dCA2_FI_GFPve_pos  2.7243085079 -0.3033327736
## dCA2_Fpos_CGCCAGATCTTGGATG-1dCA2_FI_GFPve_pos  2.7685825996  1.8266973816
## dCA2_Fpos_CGGACACGTATCCTCC-1dCA2_FI_GFPve_pos  3.3832013778  0.1631112419
## dCA2_Fpos_CGGACACTCTCTATGT-1dCA2_FI_GFPve_pos  2.9445102386 -0.0814590134
## dCA2_Fpos_CGGGTCAAGTCTAGCT-1dCA2_FI_GFPve_pos  2.4075153045  2.3099360786
## dCA2_Fpos_CGTAATGAGCCAGACA-1dCA2_FI_GFPve_pos  1.7348412447  1.9895320259
## dCA2_Fpos_CGTCCATAGGCTCCCA-1dCA2_FI_GFPve_pos  1.3943354063  1.5080447517
## dCA2_Fpos_CGTCCATTCGAACTCA-1dCA2_FI_GFPve_pos  2.2596135311  0.7052955948
## dCA2_Fpos_CGTGTCTTCAATCAGC-1dCA2_FI_GFPve_pos  2.0253215008  3.1353674255
## dCA2_Fpos_CGTTGGGCACAGCCAC-1dCA2_FI_GFPve_pos  2.6621277504  2.3709144913
## dCA2_Fpos_CTAACCCAGGGTTAGC-1dCA2_FI_GFPve_pos  2.8774240188 -0.0459942497
## dCA2_Fpos_CTAACCCGTCGGCTAC-1dCA2_FI_GFPve_pos  2.0971231632  0.7113447510
## dCA2_Fpos_CTAACCCGTTTCTTAC-1dCA2_FI_GFPve_pos  2.8938267402  0.6367535911
## dCA2_Fpos_CTAACTTTCAGCTTCC-1dCA2_FI_GFPve_pos  1.0730855636  0.7521004997
## dCA2_Fpos_CTAAGTGCAAAGACGC-1dCA2_FI_GFPve_pos  1.8550613098  2.1659350715
## dCA2_Fpos_CTACAGATCATTGCGA-1dCA2_FI_GFPve_pos  2.8372218780  1.3082003914
## dCA2_Fpos_CTACATTGTTATGACC-1dCA2_FI_GFPve_pos  1.7453297548 -0.0065769829
## dCA2_Fpos_CTACCCACATCGCTGG-1dCA2_FI_GFPve_pos  2.6744852238  1.4496460281
## dCA2_Fpos_CTAGGTAAGCATATGA-1dCA2_FI_GFPve_pos  2.2747841053  1.7626886688
## dCA2_Fpos_CTATAGGAGTAGGGTC-1dCA2_FI_GFPve_pos  2.5949521236  2.1516352020
## dCA2_Fpos_CTCAACCAGGCACAAC-1dCA2_FI_GFPve_pos  2.7860508137  2.2010059677
## dCA2_Fpos_CTCAATTGTGAGTTGG-1dCA2_FI_GFPve_pos  1.0498566799  2.3069773041
## dCA2_Fpos_CTCAGTCGTGCCGTAC-1dCA2_FI_GFPve_pos  2.7682573967  0.6509728752
## dCA2_Fpos_CTCATCGTCTAGAACC-1dCA2_FI_GFPve_pos  2.0188365154  2.0028400741
## dCA2_Fpos_CTCATGCTCCAATCCC-1dCA2_FI_GFPve_pos  2.5677428417  0.0095515571
## dCA2_Fpos_CTCCAACAGGTTACAA-1dCA2_FI_GFPve_pos  2.6333868675  1.7403273903
## dCA2_Fpos_CTCCCTCTCAGCCTTC-1dCA2_FI_GFPve_pos  1.2240712814  1.0446577392
## dCA2_Fpos_CTCCGATAGGGACACT-1dCA2_FI_GFPve_pos  3.0219695740  1.1932459198
## dCA2_Fpos_CTCGAGGGTCTTGCTC-1dCA2_FI_GFPve_pos  1.6896437578  0.8730273567
## dCA2_Fpos_CTCGAGGTCCGTCACT-1dCA2_FI_GFPve_pos  2.6623609237  1.3543606125
## dCA2_Fpos_CTCTCGATCATTGCTT-1dCA2_FI_GFPve_pos  2.8525960617 -0.6822938599
## dCA2_Fpos_CTCTGGTAGTCGTTAC-1dCA2_FI_GFPve_pos  2.2818014793  1.7591104828
## dCA2_Fpos_CTGGCAGGTATCTCTT-1dCA2_FI_GFPve_pos  1.4920007162 -0.3986472763
## dCA2_Fpos_CTGTAGACAAGTTCCA-1dCA2_FI_GFPve_pos  1.0456941299  1.9379468284
## dCA2_Fpos_CTGTAGACACGGCTAC-1dCA2_FI_GFPve_pos  1.3945896797  1.1093130432
## dCA2_Fpos_CTGTATTGTCGTTGCG-1dCA2_FI_GFPve_pos  2.2028792076  1.5279269539
## dCA2_Fpos_CTTACCGCAGCTCTGG-1dCA2_FI_GFPve_pos  3.3664024048  0.1976933800
## dCA2_Fpos_CTTCAATGTATGATCC-1dCA2_FI_GFPve_pos  2.6779995136  1.3571181618
## dCA2_Fpos_CTTCTCTGTGCCGGTT-1dCA2_FI_GFPve_pos  1.3596177273  1.1587729774
## dCA2_Fpos_CTTGAGAGTGTTACAC-1dCA2_FI_GFPve_pos  1.3376412563  2.1506638847
## dCA2_Fpos_CTTGATTCACACCTTC-1dCA2_FI_GFPve_pos  1.8858564071  0.7102003418
## dCA2_Fpos_GAAATGAGTACTCAAC-1dCA2_FI_GFPve_pos  2.5124972038  0.8950648628
## dCA2_Fpos_GAACTGTAGACTGTTC-1dCA2_FI_GFPve_pos  2.4071684055  0.8435483299
## dCA2_Fpos_GAACTGTTCATTTCCA-1dCA2_FI_GFPve_pos  1.2319511347  1.2972121559
```

```
## dCA2_Fpos_GAAGGACTCAGCGTCG-1dCA2_FI_GFPve_pos  0.8383698635  2.0884266220
## dCA2_Fpos_GAAGGGTCAAAGCTAA-1dCA2_FI_GFPve_pos  2.7757663898  0.4438648544
## dCA2_Fpos_GACAGCCTCTGCGGAC-1dCA2_FI_GFPve_pos  1.2335219555  0.0206985794
## dCA2_Fpos_GACCCAGCATACCAGT-1dCA2_FI_GFPve_pos  1.9876718693  0.7368259750
## dCA2_Fpos_GACCTTCAGCAGCCCT-1dCA2_FI_GFPve_pos  3.1157927685  0.6233659111
## dCA2_Fpos_GACTATGCACGTCGGT-1dCA2_FI_GFPve_pos  1.8934631519  2.0898433052
## dCA2_Fpos_GACTCTCGTCTTCATT-1dCA2_FI_GFPve_pos  2.1169283562 -0.2413506188
## dCA2_Fpos_GACTTCCTCAATGCAC-1dCA2_FI_GFPve_pos  2.1499152355  0.3843999229
## dCA2_Fpos_GAGAAATGTTTGTGGT-1dCA2_FI_GFPve_pos  0.8258195691  1.8135953270
## dCA2_Fpos_GAGACCCTCAGCTTGA-1dCA2_FI_GFPve_pos  3.5368814640  0.1599726997
## dCA2_Fpos_GAGACTTGTCTATGAC-1dCA2_FI_GFPve_pos  3.1996939354  1.0275931679
## dCA2_Fpos_GAGGCCTAGGCTATCT-1dCA2_FI_GFPve_pos  3.0777301960  1.4498095833
## dCA2_Fpos_GAGTCATAGCCGTTAT-1dCA2_FI_GFPve_pos  0.2395007782  1.0661931358
## dCA2_Fpos_GAGTCTAGTTAGAAAC-1dCA2_FI_GFPve_pos  2.3901081257  1.7281289421
## dCA2_Fpos_GATAGAACACGGCACT-1dCA2_FI_GFPve_pos  3.2997911148  0.8645763717
## dCA2_Fpos_GATAGCTCATTCATCT-1dCA2_FI_GFPve_pos  1.1833906345  0.2387724243
## dCA2_Fpos_GATCACAGTTTCTTAC-1dCA2_FI_GFPve_pos  2.1159722976  1.1841903053
## dCA2_Fpos_GATCATGCAGGCTACC-1dCA2_FI_GFPve_pos  1.8745227032  0.3204474769
## dCA2_Fpos_GATCCCTAGCGGGTAT-1dCA2_FI_GFPve_pos  2.8697717361  1.0341430031
## dCA2_Fpos_GATGACTCATGTCTAG-1dCA2_FI_GFPve_pos  2.1086487942  2.1837458931
## dCA2_Fpos_GATGCTAAGAGGCGTT-1dCA2_FI_GFPve_pos  2.2208030395  1.6827178322
## dCA2_Fpos_GATGCTAAGTGTGTTC-1dCA2_FI_GFPve_pos  3.2609243564 -0.1918582596
## dCA2_Fpos_GATGGAGGTCAACGCC-1dCA2_FI_GFPve_pos  2.1692650490  2.5750985466
## dCA2_Fpos_GCACATAAGCCTCAAT-1dCA2_FI_GFPve_pos  2.5456648044  0.1784830414
## dCA2_Fpos_GCACATACAATCCTAG-1dCA2_FI_GFPve_pos  2.8354706936 -0.8338610806
## dCA2_Fpos_GCCAGCAAGCCTAACT-1dCA2_FI_GFPve_pos  1.5626470022 -1.9198186077
## dCA2_Fpos_GCCAGGTTCGAGCCTG-1dCA2_FI_GFPve_pos  2.9674377613  0.1636815391
## dCA2_Fpos_GCCAGGTTCGATCCAA-1dCA2_FI_GFPve_pos  3.0694682770  1.1977444015
## dCA2_Fpos_GCCATGGTCCTGCCAT-1dCA2_FI_GFPve_pos  2.2661566906  0.0513382278
## dCA2_Fpos_GCCCGAAGTAACGATA-1dCA2_FI_GFPve_pos  1.6764450245  0.4031782471
## dCA2_Fpos_GCCGATGTCTGTCCCA-1dCA2_FI_GFPve_pos  2.6523125343  0.5223126732
## dCA2_Fpos_GCCTGTTTCCGGACGT-1dCA2_FI_GFPve_pos  1.9129131012  1.0026231132
## dCA2_Fpos_GCCTGTTTCGCTCTCA-1dCA2_FI_GFPve_pos  2.2376089268  0.0694599472
## dCA2_Fpos_GCGAGAAAGTTGTCAC-1dCA2_FI_GFPve_pos  2.2633867435  2.5409317337
## dCA2_Fpos_GCTACCTCAAATGCTC-1dCA2_FI_GFPve_pos  1.8051896267  2.4715814911
## dCA2_Fpos_GCTCAAAAGAGTTGCG-1dCA2_FI_GFPve_pos  2.5770871811  2.3902445160
## dCA2_Fpos_GCTGGGTCACTCCGAG-1dCA2_FI_GFPve_pos  1.6077525787  0.7177887283
## dCA2_Fpos_GGAGATGAGTAACGTA-1dCA2_FI_GFPve_pos  2.8158879452  1.3223715149
## dCA2_Fpos_GGATCTAAGGGCCTCT-1dCA2_FI_GFPve_pos  2.2506439857  1.8653593384
## dCA2_Fpos_GGATCTAGTCACTTAG-1dCA2_FI_GFPve_pos  2.3213303261  1.3444776855
## dCA2_Fpos_GGCGTCACAGTGTGCC-1dCA2_FI_GFPve_pos  2.3325820141  2.3313832603
## dCA2_Fpos_GGCTGTGCATAACAGA-1dCA2_FI_GFPve_pos  3.4951431923  0.8971371971
## dCA2_Fpos_GGGACCTTCAAACCTG-1dCA2_FI_GFPve_pos  2.9898340874  0.2004561744
## dCA2_Fpos_GGGACTCTCCTCTTTC-1dCA2_FI_GFPve_pos  1.4512150459  0.7943368278
## dCA2_Fpos_GGGAGATGTCTTGCTC-1dCA2_FI_GFPve_pos  1.8932395153  3.0018391929
## dCA2_Fpos_GGGATCCGTCCGGATC-1dCA2_FI_GFPve_pos  3.3261227779  0.9106197677
## dCA2_Fpos_GGGATGACAGTCAGCC-1dCA2_FI_GFPve_pos  2.0702166729  1.1134195648
## dCA2_Fpos_GGGCGTTAGCGGTATG-1dCA2_FI_GFPve_pos  1.6234253816 -0.1212591805
## dCA2_Fpos_GGGTAGAAGCTGACTT-1dCA2_FI_GFPve_pos  2.3618805580  2.4923658691
## dCA2_Fpos_GGGTCACCAGACAAAT-1dCA2_FI_GFPve_pos  2.6537273102  1.0441065155
## dCA2_Fpos_GGTAGAGAGACTTAAG-1dCA2_FI_GFPve_pos  3.2227320843  1.5606627785
## dCA2_Fpos_GGTCTGGTCGAGAGAC-1dCA2_FI_GFPve_pos  2.2497968845  2.2509141289
## dCA2_Fpos_GGTGATTAGGGTTAAT-1dCA2_FI_GFPve_pos  1.3718239240  1.3728504501
## dCA2_Fpos_GGTGATTGTCAGTCCG-1dCA2_FI_GFPve_pos  1.9696624450  1.9724297844
## dCA2_Fpos_GGTGGCTTCTGACCCT-1dCA2_FI_GFPve_pos  2.6741790943 -0.5950212158
## dCA2_Fpos_GGTGTTAAGCTTTGTG-1dCA2_FI_GFPve_pos  1.1031100921  2.1876035057
## dCA2_Fpos_GGTTAACTCGAACGCC-1dCA2_FI_GFPve_pos  1.7591557674  1.2885341965
## dCA2_Fpos_GGTTCTCGTTTCGTGA-1dCA2_FI_GFPve_pos  1.8529882602  1.7048058830
## dCA2_Fpos_GTAACACCAACAGTGG-1dCA2_FI_GFPve_pos  2.3079366855  2.5152569137
## dCA2_Fpos_GTAATCGGTCTTGTCC-1dCA2_FI_GFPve_pos  2.1350960903  2.2300873123
## dCA2_Fpos_GTAATCGTCGACGCTG-1dCA2_FI_GFPve_pos  2.2798688583  2.8119321190
## dCA2_Fpos_GTAATGCAGTTTCGAC-1dCA2_FI_GFPve_pos  1.9962363415  2.3281231247
## dCA2_Fpos_GTAGGAGGTATTGAGA-1dCA2_FI_GFPve_pos  3.3128833942  1.5283432327
## dCA2_Fpos_GTAGGTTAGCATGATA-1dCA2_FI_GFPve_pos  3.1021318607  1.2622199379
## dCA2_Fpos_GTAGTACAGCGAAACC-1dCA2_FI_GFPve_pos  2.9345138244  1.7976832710
## dCA2_Fpos_GTAGTACGTCATCGCG-1dCA2_FI_GFPve_pos  1.9634614162  2.4200911842
## dCA2_Fpos_GTAGTACGTTGGGAAC-1dCA2_FI_GFPve_pos  3.3381467037  0.4851203285
## dCA2_Fpos_GTCAAACGTTGTCCCT-1dCA2_FI_GFPve_pos  1.1991962366  2.7813673340
## dCA2_Fpos_GTCAAACTCTCTCGCA-1dCA2_FI_GFPve_pos -0.3564230479  0.3122215591
## dCA2_Fpos_GTCAGCGAGTTCACTG-1dCA2_FI_GFPve_pos  2.3077190094  2.7716908775
```

```
## dCA2_Fpos_GTCCTCATCTGAATGC-1dCA2_FI_GFPve_pos  1.6248638801 -0.4228763260
## dCA2_Fpos_GTCTAGAAGGGAGGCA-1dCA2_FI_GFPve_pos  1.8304094248  1.7143173538
## dCA2_Fpos_GTCTAGATCCGAAGGA-1dCA2_FI_GFPve_pos  2.1179816894  2.4008865677
## dCA2_Fpos_GTCTGTCTCATGAGTC-1dCA2_FI_GFPve_pos  1.8473632507  2.0846395813
## dCA2_Fpos_GTGATGTAGTAATTGG-1dCA2_FI_GFPve_pos  2.3087485008  2.2137923561
## dCA2_Fpos_GTGCTTCCAGAGTAAT-1dCA2_FI_GFPve_pos  3.2445650272  1.0184645973
## dCA2_Fpos_GTGGAGATCAGTGGGA-1dCA2_FI_GFPve_pos  1.6663320236  1.8324947677
## dCA2_Fpos_GTGTAACAGACCAAGC-1dCA2_FI_GFPve_pos  2.8041014843  2.2134876571
## dCA2_Fpos_GTGTGGCAGGTCTTTG-1dCA2_FI_GFPve_pos  1.2265877895  1.0728478752
## dCA2_Fpos_GTTCTATAGGACATCG-1dCA2_FI_GFPve_pos  2.1004593544  1.4772215210
## dCA2_Fpos_GTTGCTCGTCCAGTTA-1dCA2_FI_GFPve_pos  2.3638172321  0.7999797188
## dCA2_Fpos_GTTGTAGAGCATCCTA-1dCA2_FI_GFPve_pos  1.3066413574  0.3720603309
## dCA2_Fpos_GTTGTAGCAGGACGAT-1dCA2_FI_GFPve_pos  2.1469769649 -1.2723991551
## dCA2_Fpos_GTTTACTAGTTGCTGT-1dCA2_FI_GFPve_pos  2.2422306709  2.9254961334
## dCA2_Fpos_GTTTACTTCCATGCAA-1dCA2_FI_GFPve_pos  2.4033239059  1.4182648979
## dCA2_Fpos_TAACCAGTCGTGGAAG-1dCA2_FI_GFPve_pos  0.4907983117  2.4119124733
## dCA2_Fpos_TAAGCCAAGAAAGTCT-1dCA2_FI_GFPve_pos  3.1541168861  1.3009934746
## dCA2_Fpos_TAAGCCACACATGGTT-1dCA2_FI_GFPve_pos  2.2190566234 -0.1903295197
## dCA2_Fpos_TAAGCCATCATTGCTT-1dCA2_FI_GFPve_pos  0.4632308655  1.5224080406
## dCA2_Fpos_TAAGTCGTCTCGTTTA-1dCA2_FI_GFPve_pos  2.4688673191  2.6779013000
## dCA2_Fpos_TAATTCCGTCCAAGAG-1dCA2_FI_GFPve_pos  2.0104034118  0.6718106590
## dCA2_Fpos_TAATTCCTCTCAACGA-1dCA2_FI_GFPve_pos  2.2642455272 -0.4362029709
## dCA2_Fpos_TACAGGTAGGGTCACA-1dCA2_FI_GFPve_pos  2.4277248554  1.6874389969
## dCA2_Fpos_TACCCGTAGCCTAGGA-1dCA2_FI_GFPve_pos  2.6889581852  1.3263240181
## dCA2_Fpos_TACCCGTTCACTTCTA-1dCA2_FI_GFPve_pos  2.2177927666  1.4310164772
## dCA2_Fpos_TACCGAAGTCCGGTGT-1dCA2_FI_GFPve_pos  3.0726375751  2.1090803467
## dCA2_Fpos_TACCGGGAGCGATGGT-1dCA2_FI_GFPve_pos  2.9913847618  1.8871984802
## dCA2_Fpos_TACCTGCAGCGGTATG-1dCA2_FI_GFPve_pos  2.2367892437  1.9166217171
## dCA2_Fpos_TACCTGCCAAAGTGTA-1dCA2_FI_GFPve_pos  2.0109570198  1.9825859390
## dCA2_Fpos_TACGGTACAGAGAATT-1dCA2_FI_GFPve_pos  2.8499288730  0.7570672355
## dCA2_Fpos_TACTTACTCAACTGGT-1dCA2_FI_GFPve_pos  1.4927255087  1.6679096542
## dCA2_Fpos_TACTTGTAGCCATCCG-1dCA2_FI_GFPve_pos  2.9495208435  0.9223275505
## dCA2_Fpos_TAGACCATCATTTACC-1dCA2_FI_GFPve_pos  1.6795146637  0.7043843590
## dCA2_Fpos_TAGGTACAGATTCGAA-1dCA2_FI_GFPve_pos  1.5339279346  2.6601095520
## dCA2_Fpos_TAGGTTGTCGACGATT-1dCA2_FI_GFPve_pos  1.2023271494  1.4662733398
## dCA2_Fpos_TAGTGCAAGTAGCATA-1dCA2_FI_GFPve_pos  2.5444662742 -0.0353631653
## dCA2_Fpos_TATACCTAGCAGCGAT-1dCA2_FI_GFPve_pos  0.9715976887  0.3180165611
## dCA2_Fpos_TATACCTTCATCTGTT-1dCA2_FI_GFPve_pos  3.0267980270 -0.9674379505
## dCA2_Fpos_TATCTTGCAGCAGTCC-1dCA2_FI_GFPve_pos  2.2285530739  1.4297175728
## dCA2_Fpos_TATCTTGGTTATTCCT-1dCA2_FI_GFPve_pos  2.6115336590  1.4459362350
## dCA2_Fpos_TATGTTCCATCGTCCT-1dCA2_FI_GFPve_pos  2.7897586994 -0.7379526772
## dCA2_Fpos_TATTGCTCATCGCTGG-1dCA2_FI_GFPve_pos  1.8422579937  1.1481175743
## dCA2_Fpos_TATTGCTTCTCCCATG-1dCA2_FI_GFPve_pos  1.0553947859  1.0215149246
## dCA2_Fpos_TATTGGGCAGTTGTTG-1dCA2_FI_GFPve_pos  0.8462867908  1.6581344925
## dCA2_Fpos_TATTTCGAGCAATAAC-1dCA2_FI_GFPve_pos  0.4352120333 -0.1267909683
## dCA2_Fpos_TATTTCGTCAGCTTGA-1dCA2_FI_GFPve_pos  2.6955509357  0.5479698501
## dCA2_Fpos_TCACAAGCAAGCTGCC-1dCA2_FI_GFPve_pos  3.0997805767  1.9565491997
## dCA2_Fpos_TCACACCCAACAGCTT-1dCA2_FI_GFPve_pos  2.1417567901 -1.5198487915
## dCA2_Fpos_TCACGGGCAACCAACT-1dCA2_FI_GFPve_pos  1.6896923952  0.8215880714
## dCA2_Fpos_TCAGCCTAGCCGTAAG-1dCA2_FI_GFPve_pos  2.5677316360  2.2180466972
## dCA2_Fpos_TCAGCCTAGGCACGAT-1dCA2_FI_GFPve_pos  2.6398029499  1.0124287926
## dCA2_Fpos_TCAGCCTAGTCAGCGA-1dCA2_FI_GFPve_pos  2.4379406147 -1.1247362770
## dCA2_Fpos_TCAGGTAAGTGATTCC-1dCA2_FI_GFPve_pos  2.5855233841  1.4492755256
## dCA2_Fpos_TCATCCGAGATTAGAC-1dCA2_FI_GFPve_pos  2.4922073059  0.1791735016
## dCA2_Fpos_TCATCCGTCTCAATCT-1dCA2_FI_GFPve_pos  3.0750272445 -0.7955810703
## dCA2_Fpos_TCATGAGTCTGTGTGA-1dCA2_FI_GFPve_pos  2.4895794563  1.7168526970
## dCA2_Fpos_TCATGCCAGGTCCCTG-1dCA2_FI_GFPve_pos  1.4683259897  1.1407990776
## dCA2_Fpos_TCATTCACATGGTGGA-1dCA2_FI_GFPve_pos  1.7724611931  1.7419801079
## dCA2_Fpos_TCCAGAAAGACTCTAC-1dCA2_FI_GFPve_pos  1.7749440841  1.6647076927
## dCA2_Fpos_TCCAGAAAGTAGCAAT-1dCA2_FI_GFPve_pos  2.2091977768  1.1798782669
## dCA2_Fpos_TCCAGAACACTGCTTC-1dCA2_FI_GFPve_pos  1.8614366226  2.0869131408
## dCA2_Fpos_TCCATGCTCGGTAACT-1dCA2_FI_GFPve_pos  2.1069033317  1.4918175064
## dCA2_Fpos_TCCGAAATCTCGACGG-1dCA2_FI_GFPve_pos  2.3462667637  0.3791051231
## dCA2_Fpos_TCCTAATAGTCGGGAT-1dCA2_FI_GFPve_pos  1.4087507896  1.9045977913
## dCA2_Fpos_TCCTCGAAGTCTAACC-1dCA2_FI_GFPve_pos  1.8552415542 -1.0552043595
## dCA2_Fpos_TCCTCGAGTGTTCCAA-1dCA2_FI_GFPve_pos  1.6244684629  2.4706249557
## dCA2_Fpos_TCCTCGATCCTTGACC-1dCA2_FI_GFPve_pos  1.1663335733  0.7730961166
## dCA2_Fpos_TCCTTCTAGAACTTCC-1dCA2_FI_GFPve_pos  0.9224777393  1.6899004303
## dCA2_Fpos_TCGACCTAGATGACAT-1dCA2_FI_GFPve_pos  0.6755432658 -0.7948815502
## dCA2_Fpos_TCGACGGTCTCACCCA-1dCA2_FI_GFPve_pos  1.9421909027  2.7791996322
```

```
## dCA2_Fpos_TCGCAGGGTGGTCTTA-1dCA2_FI_GFPve_pos  2.1965024643  0.7352776848
## dCA2_Fpos_TCGGATACATGTCGTA-1dCA2_FI_GFPve_pos  2.8346786670  0.3835916839
## dCA2_Fpos_TCGGATATCGATTGAC-1dCA2_FI_GFPve_pos  2.0445208721  1.7657404266
## dCA2_Fpos_TCGGGTGGTGAGCTCC-1dCA2_FI_GFPve_pos  2.1842715912  2.5654869400
## dCA2_Fpos_TCGGTCTCACTCCCTA-1dCA2_FI_GFPve_pos -0.3020080306  0.8258452736
## dCA2_Fpos_TCTACCGCAATCTAGC-1dCA2_FI_GFPve_pos  1.4745979481  0.7916493736
## dCA2_Fpos_TCTATACAGGAATGTT-1dCA2_FI_GFPve_pos  1.2376089268  1.2641578041
## dCA2_Fpos_TCTATCAGTGAGTAAT-1dCA2_FI_GFPve_pos  2.3351767235  1.8823013626
## dCA2_Fpos_TCTCAGCTCTAACGGT-1dCA2_FI_GFPve_pos  1.1955036096  1.7351250969
## dCA2_Fpos_TCTCCGATCACGACTA-1dCA2_FI_GFPve_pos  2.2040033512  0.6190486275
## dCA2_Fpos_TCTGCCAGTCAAAGAT-1dCA2_FI_GFPve_pos  2.4780841045  1.4578748070
## dCA2_Fpos_TCTGGCTGTGGAGGTT-1dCA2_FI_GFPve_pos  3.1499517135  1.6956529938
## dCA2_Fpos_TCTTAGTCACAATGCT-1dCA2_FI_GFPve_pos  0.9068784885  1.9764333092
## dCA2_Fpos_TCTTAGTGTCGCTGCA-1dCA2_FI_GFPve_pos  1.3424782924  1.7427730881
## dCA2_Fpos_TCTTTGAAGCCTCTGG-1dCA2_FI_GFPve_pos  1.6461823158  1.6217742286
## dCA2_Fpos_TGAACGTAGTCCCGAC-1dCA2_FI_GFPve_pos  2.7418182068  1.5994134270
## dCA2_Fpos_TGAATCGAGTGCACCC-1dCA2_FI_GFPve_pos  2.5108258896  0.7826557480
## dCA2_Fpos_TGACAGTTCGGTCTGG-1dCA2_FI_GFPve_pos  0.7929806285  2.2739501320
## dCA2_Fpos_TGACGCGGTTGGGAAC-1dCA2_FI_GFPve_pos  2.1136791878  1.2111625992
## dCA2_Fpos_TGAGGAGTCAAGTGGG-1dCA2_FI_GFPve_pos  3.2321834736 -0.0683746018
## dCA2_Fpos_TGAGGGAGTATCGAAA-1dCA2_FI_GFPve_pos  1.6928945951  1.5205078445
## dCA2_Fpos_TGATCAGCAGGTCTCG-1dCA2_FI_GFPve_pos  2.2915177517  2.7398519836
## dCA2_Fpos_TGATGCAAGTACTGGG-1dCA2_FI_GFPve_pos  2.9304957561  2.0467748962
## dCA2_Fpos_TGATGCACAAGCGCAA-1dCA2_FI_GFPve_pos  1.2562681608  2.1899357162
## dCA2_Fpos_TGATGGTGTTTACGAC-1dCA2_FI_GFPve_pos  2.5650422745  1.8441272102
## dCA2_Fpos_TGATTCTTCGAAACAA-1dCA2_FI_GFPve_pos  2.2560975723  1.8891101204
## dCA2_Fpos_TGCACGGTCCACTAGA-1dCA2_FI_GFPve_pos  1.4423841409 -1.2409073986
## dCA2_Fpos_TGCAGATAGGTGCAGT-1dCA2_FI_GFPve_pos  2.2709648781  1.3667078338
## dCA2_Fpos_TGCAGATTCAGGGTAG-1dCA2_FI_GFPve_pos  3.0565047436  0.9741955124
## dCA2_Fpos_TGCAGATTCTCAACCC-1dCA2_FI_GFPve_pos  2.2401647739  0.1220412575
## dCA2_Fpos_TGCAGGCCAGCGAGTA-1dCA2_FI_GFPve_pos -1.3920634814  0.3658562027
## dCA2_Fpos_TGCGATACACTAGGTT-1dCA2_FI_GFPve_pos  2.6474101715  1.1584515892
## dCA2_Fpos_TGCGGGTGTGTTTACG-1dCA2_FI_GFPve_pos  1.2310263090  2.5574031196
## dCA2_Fpos_TGCTCCATCACGATAC-1dCA2_FI_GFPve_pos  1.4882606439  1.5749378525
## dCA2_Fpos_TGCTGAAAGTTCCATG-1dCA2_FI_GFPve_pos  2.0625732116 -0.3412069954
## dCA2_Fpos_TGCTTCGAGGTGGCTA-1dCA2_FI_GFPve_pos  2.3603644542  1.1736808143
## dCA2_Fpos_TGCTTGCTCGGATAAA-1dCA2_FI_GFPve_pos  1.9918563537 -0.5390972771
## dCA2_Fpos_TGGAACTGTATCGCGC-1dCA2_FI_GFPve_pos  2.6771686249  2.0548639618
## dCA2_Fpos_TGGATCAAGAGCAGAA-1dCA2_FI_GFPve_pos  1.5393644743  2.3457517944
## dCA2_Fpos_TGGATCATCAGAATAG-1dCA2_FI_GFPve_pos  2.3410468273  1.8438725792
## dCA2_Fpos_TGGATGTCATGACGTT-1dCA2_FI_GFPve_pos  1.1358506851  2.5786424003
## dCA2_Fpos_TGGGAGAGTCCACATA-1dCA2_FI_GFPve_pos  2.4258625679  1.5241055809
## dCA2_Fpos_TGGGTTAGTGGAAATT-1dCA2_FI_GFPve_pos  2.6649046116  1.8828745209
## dCA2_Fpos_TGTACAGGTTGTCCCT-1dCA2_FI_GFPve_pos  1.3843660526  1.1572995506
## dCA2_Fpos_TGTAGACAGTATGTAG-1dCA2_FI_GFPve_pos  2.7817602329  1.9431367241
## dCA2_Fpos_TGTCCTGGTATCACCA-1dCA2_FI_GFPve_pos  2.5629623108  0.4381132446
## dCA2_Fpos_TGTGATGCACGTAGTT-1dCA2_FI_GFPve_pos  1.1742966347  1.9750476204
## dCA2_Fpos_TGTGGCGAGCATCTTG-1dCA2_FI_GFPve_pos  3.2428279094 -0.3098811783
## dCA2_Fpos_TGTGGCGTCCATTGGA-1dCA2_FI_GFPve_pos  2.8356950455  1.9017472587
## dCA2_Fpos_TGTGTGATCCTCAGGG-1dCA2_FI_GFPve_pos  1.3335197143  1.7025995575
## dCA2_Fpos_TGTTCCGTCACAGAGG-1dCA2_FI_GFPve_pos  3.1466877632  1.3101864181
## dCA2_Fpos_TGTTCTAAGCTCGTGC-1dCA2_FI_GFPve_pos  2.8175120525  2.1716990791
## dCA2_Fpos_TTACGCCCATCCTAAG-1dCA2_FI_GFPve_pos  2.3437211685  1.8311872803
## dCA2_Fpos_TTACGTTAGCGGCTCT-1dCA2_FI_GFPve_pos  2.0761375599  1.3899112068
## dCA2_Fpos_TTACTGTTCTGCCTGT-1dCA2_FI_GFPve_pos  0.4019820862  1.4375067077
## dCA2_Fpos_TTAGGCACAGGAACCA-1dCA2_FI_GFPve_pos  1.7291513853  1.4084205948
## dCA2_Fpos_TTCAATCGTCACATTG-1dCA2_FI_GFPve_pos  0.6628288798  2.0642805420
## dCA2_Fpos_TTCATTGCAACTTGGT-1dCA2_FI_GFPve_pos  1.7468937807  1.2466526352
## dCA2_Fpos_TTCCAATCAACATACC-1dCA2_FI_GFPve_pos  1.3089046650  0.8577580772
## dCA2_Fpos_TTCCTCTAGGCCCAAA-1dCA2_FI_GFPve_pos  1.8183044367  1.0473280273
## dCA2_Fpos_TTCGATTTCTCTGCTG-1dCA2_FI_GFPve_pos  2.0089175873  0.4599099479
## dCA2_Fpos_TTCGCTGGTTCTCCTG-1dCA2_FI_GFPve_pos  3.1082477741  0.5230017029
## dCA2_Fpos_TTCTAACCACTCTCGT-1dCA2_FI_GFPve_pos  2.5876629524  0.6850948654
## dCA2_Fpos_TTCTAGTTCAAGCTGT-1dCA2_FI_GFPve_pos  2.4436524086  2.3107066475
## dCA2_Fpos_TTCTTCCCATCTAACG-1dCA2_FI_GFPve_pos  2.7949209385 -0.4781961121
## dCA2_Fpos_TTCTTCCTCATGAGGG-1dCA2_FI_GFPve_pos  3.2333052330 -0.5026135124
## dCA2_Fpos_TTGACCCAGGAACGTC-1dCA2_FI_GFPve_pos  1.8452212982  0.8480148636
## dCA2_Fpos_TTGACCCGTCGCAGTC-1dCA2_FI_GFPve_pos -0.6223780043  0.6284218155
## dCA2_Fpos_TTGCATTTCGTCTAAG-1dCA2_FI_GFPve_pos  2.2566738300  2.2388739906
## dCA2_Fpos_TTGCTGCTCATTGCTT-1dCA2_FI_GFPve_pos  2.2725973301 -0.5687493958
```

```
## dCA2_Fpos_TTGCTGCTCCTACTGC-1dCA2_FI_GFPve_pos  2.4232402019  0.0661344848
## dCA2_Fpos_TTTCCTCAGTCGCCAC-1dCA2_FI_GFPve_pos  2.3883204632  0.3293385826
## dCA2_Fpos_TTTGACTAGGATTCAA-1dCA2_FI_GFPve_pos  3.2009635143  0.2430777870
## dCA2_Fpos_TTTGGTTGTTCTCACC-1dCA2_FI_GFPve_pos  1.4349464350  1.2773175560
## FC_Mpos_AAACGCTAGAAACTAC-1FC_MI_GFPve_pos      3.3227694206 -0.6177663483
## FC_Mpos_AAAGGATCACCTGCGA-1FC_MI_GFPve_pos      2.1029229336  1.1672258697
## FC_Mpos_AAAGGATTCATTGCTT-1FC_MI_GFPve_pos      1.8189438753 -0.0860495247
## FC_Mpos_AAAGGGCAGCTCTATG-1FC_MI_GFPve_pos      2.4030518703  2.1062555633
## FC_Mpos_AAAGTCCTCAGTGATC-1FC_MI_GFPve_pos      0.4831180744  2.1191025101
## FC_Mpos_AAATGGAAGAGGGTGG-1FC_MI_GFPve_pos      1.5393395595 -0.5392880120
## FC_Mpos_AAATGGACAAGCAATA-1FC_MI_GFPve_pos     -0.0118306108 -3.6308417000
## FC_Mpos_AACAACCAGGAATGTT-1FC_MI_GFPve_pos      1.0422443323  2.5754700027
## FC_Mpos_AACACACCACTTGGGC-1FC_MI_GFPve_pos      2.2055688076  0.0337152801
## FC_Mpos_AACGAAAGTCGACTTA-1FC_MI_GFPve_pos      1.5967531376 -1.8522557892
## FC_Mpos_AACGTCATCGCCGAAC-1FC_MI_GFPve_pos      1.6105090551  1.5960951172
## FC_Mpos_AAGAACACAATGTGGG-1FC_MI_GFPve_pos      2.6870424919 -0.6716184296
## FC_Mpos_AAGATAGAGCCATTTG-1FC_MI_GFPve_pos      2.0957956486  0.1713700615
## FC_Mpos_AAGCGTTAGAGAGGGC-1FC_MI_GFPve_pos      2.7428484134 -0.5009617485
## FC_Mpos_AAGCGTTCACAGCGCT-1FC_MI_GFPve_pos      0.4532604389  0.8573289238
## FC_Mpos_AATGCCACAGAGGTAC-1FC_MI_GFPve_pos      1.6419565849  2.4976859413
## FC_Mpos_AATTCCTAGCTCTGTA-1FC_MI_GFPve_pos      2.0539925270  0.5904059730
## FC_Mpos_ACACAGTAGTGCACCC-1FC_MI_GFPve_pos      0.5048895650  1.3537493072
## FC_Mpos_ACACTGAAGGCCCAAA-1FC_MI_GFPve_pos      1.6330973081  1.7109303795
## FC_Mpos_ACATTTCAGTCCCTAA-1FC_MI_GFPve_pos      3.2131931953 -0.7462269940
## FC_Mpos_ACCACAAAGCTCCCTT-1FC_MI_GFPve_pos     -0.0650779135  0.4589748703
## FC_Mpos_ACCACAACAGGTCTCG-1FC_MI_GFPve_pos      0.8866496258  1.3192544304
## FC_Mpos_ACCATTTGTTCCAAAC-1FC_MI_GFPve_pos      2.6507933311 -1.3005036988
## FC_Mpos_ACCCAAAGTTGTGTAC-1FC_MI_GFPve_pos      2.3009634189 -1.2095048107
## FC_Mpos_ACCCTTGTCTCGGTCT-1FC_MI_GFPve_pos      2.4958093338  0.3427934967
## FC_Mpos_ACCTGAAAGATGCTAA-1FC_MI_GFPve_pos      2.4555483036  0.6939010940
## FC_Mpos_ACCTGAACACGCTGAC-1FC_MI_GFPve_pos      1.2195207052  0.9586778007
## FC_Mpos_ACGCACGCATATAGCC-1FC_MI_GFPve_pos     -0.8987679906 -3.4666707195
## FC_Mpos_ACGGTCGCACATATGC-1FC_MI_GFPve_pos      0.2012642793 -0.1308350243
## FC_Mpos_ACGTAGTTCCGTTTCG-1FC_MI_GFPve_pos      0.5978745394 -0.9698748268
## FC_Mpos_ACTTAGGAGGACATCG-1FC_MI_GFPve_pos      2.0194680862  2.0402822815
## FC_Mpos_ACTTATCTCACATACG-1FC_MI_GFPve_pos      2.8874674015 -0.1131519951
## FC_Mpos_AGACCATAGAATAACC-1FC_MI_GFPve_pos      2.7178964786  0.3814325653
## FC_Mpos_AGAGCAGCATAATCGC-1FC_MI_GFPve_pos      3.6385739021  1.0233898483
## FC_Mpos_AGATCGTGTAATTAGG-1FC_MI_GFPve_pos      1.7169295721  1.1756892525
## FC_Mpos_AGCCAATGTAATGATG-1FC_MI_GFPve_pos      0.0542863779  1.7512855850
## FC_Mpos_AGCGCCAAGTATTGCC-1FC_MI_GFPve_pos      1.6413206034  1.4079037033
## FC_Mpos_AGCTTCCGTCTGTTAG-1FC_MI_GFPve_pos      2.7560382061 -0.0306768097
## FC_Mpos_AGGAATAGTCTTAGTG-1FC_MI_GFPve_pos      2.2953860931  1.3613634430
## FC_Mpos_AGGACGACAGAGGAAA-1FC_MI_GFPve_pos      3.1920473747 -0.4619388260
## FC_Mpos_AGGAGGTTCGTGGACC-1FC_MI_GFPve_pos      0.5614447765 -0.7407087959
## FC_Mpos_AGGCATTCACGCGCAT-1FC_MI_GFPve_pos      2.6780736618  1.7514829956
## FC_Mpos_AGGCCACCAACCCTAA-1FC_MI_GFPve_pos      2.9919204883  0.4176087700
## FC_Mpos_AGGTTACAGAGGTTTA-1FC_MI_GFPve_pos      2.8704641037 -0.4252500214
## FC_Mpos_AGTCTCCTCGACGTCG-1FC_MI_GFPve_pos      3.4589557819  1.4109397255
## FC_Mpos_AGTGATCCAGCAAGAC-1FC_MI_GFPve_pos      3.3073072605 -0.8695652165
## FC_Mpos_AGTGATCGTGTGTACT-1FC_MI_GFPve_pos      2.8167672329  1.2007913910
## FC_Mpos_ATACCGACACGCCACA-1FC_MI_GFPve_pos      2.1573882274 -0.3839864411
## FC_Mpos_ATACTTCTCGAACGGA-1FC_MI_GFPve_pos      2.0282485657  0.5162620865
## FC_Mpos_ATATCCTCACGAAGAC-1FC_MI_GFPve_pos      0.9156227283 -0.4725918450
## FC_Mpos_ATCACAGTCCACGGAC-1FC_MI_GFPve_pos      1.8821659260 -1.2247464337
## FC_Mpos_ATCCACCTCCGTGTCT-1FC_MI_GFPve_pos      2.8784945183 -0.9694671311
## FC_Mpos_ATCTCTAGTATACGGG-1FC_MI_GFPve_pos      3.1714360885  1.3190460525
## FC_Mpos_ATGAGTCAGGAACGAA-1FC_MI_GFPve_pos      2.2365191154  1.0853233658
## FC_Mpos_ATGATCGGTGCATCTA-1FC_MI_GFPve_pos      2.1918656997  0.3766203247
## FC_Mpos_ATGCATGCACCCTGTT-1FC_MI_GFPve_pos      2.0015464001 -1.2447864689
## FC_Mpos_ATGGGTTAGTTGCGCC-1FC_MI_GFPve_pos      1.7056253128  2.3650732361
## FC_Mpos_ATGGGTTCAAATTGCC-1FC_MI_GFPve_pos      1.4340428285  1.0026717506
## FC_Mpos_ATGTCCCCACAAAGCG-1FC_MI_GFPve_pos     -0.5319377310  0.4018416725
## FC_Mpos_ATGTCCCCACATTGTG-1FC_MI_GFPve_pos      1.5443635158 -1.8799926914
## FC_Mpos_ATTACCTTCGCTTACC-1FC_MI_GFPve_pos      0.7441619806  1.4514131866
## FC_Mpos_ATTCACTGTACTAAGA-1FC_MI_GFPve_pos      2.6346864872 -0.9822976269
## FC_Mpos_ATTCCCGGTGGCAGAT-1FC_MI_GFPve_pos      0.9457066231  1.5164008461
## FC_Mpos_ATTTCACGTCTCAAGT-1FC_MI_GFPve_pos      3.0364394359  0.9604058586
## FC_Mpos_ATTTCACTCGACGAGA-1FC_MI_GFPve_pos      3.1012888126 -0.2878322281
## FC_Mpos_CAACCTCAGATCCCAT-1FC_MI_GFPve_pos      1.9525306396 -1.5333826222
```

```
## FC_Mpos_CAATTTCAGATGTTAG-1FC_MI_GFPve_pos      2.5955333881  2.3883591018
## FC_Mpos_CACACAACATTGTAGC-1FC_MI_GFPve_pos      0.5762745313 -0.7039994873
## FC_Mpos_CACGTGGGTCCCTAAA-1FC_MI_GFPve_pos      1.2617045812  1.4007115684
## FC_Mpos_CACTGGGCAGGTTCAT-1FC_MI_GFPve_pos      3.1130068474 -0.3270172753
## FC_Mpos_CAGATCACAAGTATCC-1FC_MI_GFPve_pos      0.1788937919  1.2846961342
## FC_Mpos_CAGATCAGTTAAGTCC-1FC_MI_GFPve_pos      2.1333918743 -1.7992183842
## FC_Mpos_CAGATCAGTTCACCGG-1FC_MI_GFPve_pos      2.9025802784  0.4643121086
## FC_Mpos_CAGCAATGTGATAGTA-1FC_MI_GFPve_pos      0.0405320101  1.0278663955
## FC_Mpos_CAGCACGCAACATCGT-1FC_MI_GFPve_pos      2.3763582878 -1.8008696713
## FC_Mpos_CAGTTCCCAGTCAGAG-1FC_MI_GFPve_pos      2.5646519832 -1.2264711537
## FC_Mpos_CATACCCGTGCCCACA-1FC_MI_GFPve_pos      2.8605222873  0.8651609741
## FC_Mpos_CATACTTGTAGACACG-1FC_MI_GFPve_pos      2.1511690788  0.4111695610
## FC_Mpos_CATGCGGAGATTGCGG-1FC_MI_GFPve_pos     -0.2549864925  1.5499773346
## FC_Mpos_CATGCGGTCTCGAGTA-1FC_MI_GFPve_pos     -0.5897790022 -4.2434408046
## FC_Mpos_CATTCCGCAACTCCAA-1FC_MI_GFPve_pos      2.5718133621  0.1152172409
## FC_Mpos_CATTGAGGTTGTGGAG-1FC_MI_GFPve_pos      2.9197149448  1.7387957893
## FC_Mpos_CATTGCCAGTCTTCGA-1FC_MI_GFPve_pos      3.2846768074 -0.0398544945
## FC_Mpos_CATTGTTTCTACTTCA-1FC_MI_GFPve_pos      1.4110931330  0.5984444938
## FC_Mpos_CCAATGATCTGTAAGC-1FC_MI_GFPve_pos      1.7979780607  1.7497043930
## FC_Mpos_CCACACTGTGACAACG-1FC_MI_GFPve_pos      2.3890762501  1.5943713508
## FC_Mpos_CCACTTGCACCCATAA-1FC_MI_GFPve_pos     -0.6359858937  0.1150646530
## FC_Mpos_CCACTTGCACGACGCT-1FC_MI_GFPve_pos      2.6288321190  1.8189239822
## FC_Mpos_CCCTAACGTGTTCAGT-1FC_MI_GFPve_pos      2.4078738861  1.1465940796
## FC_Mpos_CCGAACGCAAGTGATA-1FC_MI_GFPve_pos      1.9246998005  1.8343749367
## FC_Mpos_CCGTAGGAGTGTCATC-1FC_MI_GFPve_pos      0.8641807012  0.6998544060
## FC_Mpos_CCGTAGGTCTATCCAT-1FC_MI_GFPve_pos      3.0844776802  1.3947105728
## FC_Mpos_CCTAAGACAGTCTTCC-1FC_MI_GFPve_pos      2.3618607693 -1.7187430538
## FC_Mpos_CCTAAGATCTCGGGAC-1FC_MI_GFPve_pos      2.3438012771  1.8332109772
## FC_Mpos_CCTCACACAGCAGTGA-1FC_MI_GFPve_pos      0.6831754617 -0.3913178124
## FC_Mpos_CCTCAGTTCCATACTT-1FC_MI_GFPve_pos     -0.4181857832  1.3485608421
## FC_Mpos_CCTCATGGTGTCTCCT-1FC_MI_GFPve_pos      2.8473234348 -1.1162092365
## FC_Mpos_CCTCCTCCACCAAATC-1FC_MI_GFPve_pos      2.9686834984 -0.6822490372
## FC_Mpos_CCTTCAGTCCTATGGA-1FC_MI_GFPve_pos      2.4525299244  2.5143652283
## FC_Mpos_CGCATAAGTGTCCTAA-1FC_MI_GFPve_pos      3.2664907150  1.4469943367
## FC_Mpos_CGCCAGAGTTGGGCCT-1FC_MI_GFPve_pos      3.2328417473  1.5638985954
## FC_Mpos_CGGCAGTTCTGGGCCA-1FC_MI_GFPve_pos      2.9443152122 -0.9023675598
## FC_Mpos_CGGGCATAGAATTTGG-1FC_MI_GFPve_pos      1.2666241102  2.1439543091
## FC_Mpos_CGTCCATAGTATCCTG-1FC_MI_GFPve_pos      0.8886171751 -0.2072043099
## FC_Mpos_CGTGCTTCACGTCTCT-1FC_MI_GFPve_pos      2.2012291126 -1.7254998364
## FC_Mpos_CGTTCTGGTTCAAACC-1FC_MI_GFPve_pos     -0.1292587422  0.4476008735
## FC_Mpos_CTAACCCGTGGCCTCA-1FC_MI_GFPve_pos     -0.2064898215 -3.8333412089
## FC_Mpos_CTAGGTACAGTGAGCA-1FC_MI_GFPve_pos      2.1359875374 -1.6104941048
## FC_Mpos_CTCAACCTCCAGTGTA-1FC_MI_GFPve_pos      1.2598220281  1.6275043808
## FC_Mpos_CTCACTGGTCTTGAGT-1FC_MI_GFPve_pos      2.5188570194  1.5705662094
## FC_Mpos_CTCAGGGGTGCTATTG-1FC_MI_GFPve_pos      2.6273021869  2.2272425018
## FC_Mpos_CTCCAACCATCCGAGC-1FC_MI_GFPve_pos      1.9660124950  1.9936056457
## FC_Mpos_CTCCCAATCCAACCAA-1FC_MI_GFPve_pos      0.8196785025  0.6908626877
## FC_Mpos_CTCCTTTGTGGAAATT-1FC_MI_GFPve_pos     -0.3924010076 -4.4927978791
## FC_Mpos_CTCTCGAGTAGAGACC-1FC_MI_GFPve_pos      3.2289431267  0.5579228721
## FC_Mpos_CTCTCGAGTCTCCTGT-1FC_MI_GFPve_pos      1.2483225040 -0.2943792023
## FC_Mpos_CTGCGAGCAGTCGCTG-1FC_MI_GFPve_pos      0.3286492996  1.3218202911
## FC_Mpos_CTGCTCAGTCTAGGTT-1FC_MI_GFPve_pos      0.8553115301  0.6658816658
## FC_Mpos_CTGGACGAGCTTCTAG-1FC_MI_GFPve_pos      3.0390501194  1.4036474548
## FC_Mpos_CTGTATTGTAGCTGAG-1FC_MI_GFPve_pos      1.4618058376  2.0647344909
## FC_Mpos_GAACACTTCGACATAC-1FC_MI_GFPve_pos      3.0700359516 -0.6850690521
## FC_Mpos_GAAGCCCTCCAGGACC-1FC_MI_GFPve_pos     -0.1948971689 -1.2693047203
## FC_Mpos_GAAGTAATCATGCCAA-1FC_MI_GFPve_pos      2.0188780002  2.2643566452
## FC_Mpos_GACGCTGGTTCGAGCC-1FC_MI_GFPve_pos      0.9995700292  1.0304928146
## FC_Mpos_GAGAAATTCAGCAGAG-1FC_MI_GFPve_pos      3.3287453823 -0.5572676338
## FC_Mpos_GAGCTGCAGCACCGAA-1FC_MI_GFPve_pos      0.2688609235  1.0387668930
## FC_Mpos_GAGGCAATCGACGTCG-1FC_MI_GFPve_pos      2.7030201130 -1.3187691845
## FC_Mpos_GAGTGTTGTCATCAGT-1FC_MI_GFPve_pos      0.4087408356 -4.4650954880
## FC_Mpos_GAGTTACTCGTTACCC-1FC_MI_GFPve_pos      1.6446219854  0.6324091278
## FC_Mpos_GATAGCTAGAGCTGCA-1FC_MI_GFPve_pos      2.2402768307  2.3934655510
## FC_Mpos_GATCATGAGCAACTCT-1FC_MI_GFPve_pos      2.9995289020 -0.6374792732
## FC_Mpos_GATCATGCAGCAGGAT-1FC_MI_GFPve_pos      0.8270322256  1.3887486778
## FC_Mpos_GATCATGTCCAAGGGA-1FC_MI_GFPve_pos      2.3673582248  0.2621269546
## FC_Mpos_GATGATCTCCAGCTCT-1FC_MI_GFPve_pos      1.6835096054  1.1309743248
## FC_Mpos_GATTCTTTCTAGATCG-1FC_MI_GFPve_pos      2.4036014252 -0.8474249520
## FC_Mpos_GCAACATCAACGGCCT-1FC_MI_GFPve_pos      2.9114031963  0.4844942413
```

```
## FC_Mpos_GCAACCGAGCAGTACG-1FC_MI_GFPve_pos      2.9896249943  1.0763717018
## FC_Mpos_GCACATACACAGAAGC-1FC_MI_GFPve_pos      2.7548113041  1.0741935097
## FC_Mpos_GCACTAATCCTACAAG-1FC_MI_GFPve_pos      3.3487134151 -0.5104903855
## FC_Mpos_GCAGCTGCAAACTAAG-1FC_MI_GFPve_pos      2.4995658569 -1.0825962700
## FC_Mpos_GCATCGGTCTCTGCCA-1FC_MI_GFPve_pos      0.9750666790 -0.7421798386
## FC_Mpos_GCATCTCTCGCCAGAC-1FC_MI_GFPve_pos      1.0440663271  2.0490789734
## FC_Mpos_GCCAACGAGGAGGGTG-1FC_MI_GFPve_pos      1.9790682964 -0.1791114487
## FC_Mpos_GCCAGCATCCCATAAG-1FC_MI_GFPve_pos      3.2900157146  1.4078002296
## FC_Mpos_GCCAGTGCAAGGGTCA-1FC_MI_GFPve_pos      0.9517119102  1.5220308624
## FC_Mpos_GCCAGTGTCATTCATC-1FC_MI_GFPve_pos      1.2553193741  1.7043752991
## FC_Mpos_GCCCGAATCAACGCTA-1FC_MI_GFPve_pos      1.3164524011  1.3630109154
## FC_Mpos_GCCGATGCAGTGTATC-1FC_MI_GFPve_pos      1.6354289226  1.0516376816
## FC_Mpos_GCCTGTTCAGCATGCC-1FC_MI_GFPve_pos      2.2178869419 -0.0458240189
## FC_Mpos_GCGATCGCATCCCGTT-1FC_MI_GFPve_pos      0.0305726402  0.8427892051
## FC_Mpos_GCGGATCTCTTCGACC-1FC_MI_GFPve_pos      2.8758685760 -1.0655257382
## FC_Mpos_GCTACCTCAGAACTTC-1FC_MI_GFPve_pos      2.7455780678  0.4945135437
## FC_Mpos_GCTTTCGCAGTGGCTC-1FC_MI_GFPve_pos      3.0848221950  1.9496727310
## FC_Mpos_GGAGGATAGGAAAGTG-1FC_MI_GFPve_pos      1.2919843368  2.4088135086
## FC_Mpos_GGATCTACACCAAAGG-1FC_MI_GFPve_pos      1.4388481550 -0.4221572556
## FC_Mpos_GGATGTTTCTTCACGC-1FC_MI_GFPve_pos      1.8937344723  1.7735219322
## FC_Mpos_GGGATGAAGGAATGTT-1FC_MI_GFPve_pos      1.2196048670  1.3705973945
## FC_Mpos_GGGCCATTCTGAGAGG-1FC_MI_GFPve_pos      2.4699881248 -0.1408614792
## FC_Mpos_GGGCTCATCTGTCAGA-1FC_MI_GFPve_pos      1.5875744991 -1.3517546334
## FC_Mpos_GGTCTGGAGCTGTTAC-1FC_MI_GFPve_pos      0.3169478648 -1.0971252598
## FC_Mpos_GGTGGCTAGCAAGTCG-1FC_MI_GFPve_pos      2.7108748131  2.0589185081
## FC_Mpos_GGTGGCTTCAACGAGG-1FC_MI_GFPve_pos      0.4111952953  2.4727158867
## FC_Mpos_GGTTAACTCTAGCATG-1FC_MI_GFPve_pos     -0.2217726126  1.3556895576
## FC_Mpos_GGTTGTATCCGCAAAT-1FC_MI_GFPve_pos      2.7308147125 -1.3474108853
## FC_Mpos_GTAACACCAACTGCCG-1FC_MI_GFPve_pos      3.0094399624  1.7828226410
## FC_Mpos_GTAACCACACTCAAGT-1FC_MI_GFPve_pos      2.4505701237  1.5679993950
## FC_Mpos_GTAACCATCCTAGCCT-1FC_MI_GFPve_pos      2.7154045276 -0.2568416275
## FC_Mpos_GTAATCGTCCTGATAG-1FC_MI_GFPve_pos      0.8417579107  0.0415263496
## FC_Mpos_GTATTGGGTCTTCCGT-1FC_MI_GFPve_pos      3.3691888027  0.4849715553
## FC_Mpos_GTCAGCGCACAGCTGC-1FC_MI_GFPve_pos      3.0362372570  0.1455326401
## FC_Mpos_GTCCCATCACGCGTCA-1FC_MI_GFPve_pos      1.2181506328  1.1431875549
## FC_Mpos_GTCCCATCACTTCCTG-1FC_MI_GFPve_pos      0.4203038387  1.4565720878
## FC_Mpos_GTCTAGAAGTGTTCAC-1FC_MI_GFPve_pos      2.4244308643  1.8346495949
## FC_Mpos_GTCTGTCCAGACGATG-1FC_MI_GFPve_pos      2.2883668117 -1.4719767250
## FC_Mpos_GTGACGCCACATACGT-1FC_MI_GFPve_pos      1.7877249889 -1.5203506626
## FC_Mpos_GTGAGCCAGACCTGGA-1FC_MI_GFPve_pos      2.7798237972 -1.1547619976
## FC_Mpos_GTGAGGATCTCATGGA-1FC_MI_GFPve_pos      1.4456192426  2.4416642509
## FC_Mpos_GTGAGTTTCAGCTGAT-1FC_MI_GFPve_pos      2.5691952877  0.4568696342
## FC_Mpos_GTGCGTGCACCGCTAG-1FC_MI_GFPve_pos      2.1410176925  0.2257523857
## FC_Mpos_GTGTAACCAGTCGCTG-1FC_MI_GFPve_pos      0.5506898932  0.6299438797
## FC_Mpos_GTGTGGCAGCCTCATA-1FC_MI_GFPve_pos      2.1183684044 -0.2313580193
## FC_Mpos_GTGTGGCAGGTACTGG-1FC_MI_GFPve_pos      2.3583681755 -1.3411640801
## FC_Mpos_GTTACCCCAAACCATC-1FC_MI_GFPve_pos      0.7500510983  1.6077394806
## FC_Mpos_GTTAGTGGTCCTTTGC-1FC_MI_GFPve_pos      1.9642183952  0.9802055679
## FC_Mpos_GTTCGCTTCAGAGTGG-1FC_MI_GFPve_pos      1.9461898975  1.9346638046
## FC_Mpos_TAAGCCACACGCGCTA-1FC_MI_GFPve_pos      2.7211513691 -1.4783947148
## FC_Mpos_TAAGCGTAGACATCAA-1FC_MI_GFPve_pos      0.3740605883 -2.8774491228
## FC_Mpos_TAAGTCGAGACATACA-1FC_MI_GFPve_pos      2.4437384777 -1.5089320816
## FC_Mpos_TAAGTCGCAATCTGCA-1FC_MI_GFPve_pos      1.1526653938  0.0168419205
## FC_Mpos_TAATTCCAGTTCGCAT-1FC_MI_GFPve_pos      0.7534690671 -0.2981981911
## FC_Mpos_TAATTCCCACCTGCGA-1FC_MI_GFPve_pos      2.6821048431 -1.1169924416
## FC_Mpos_TACACCCAGACTTGTC-1FC_MI_GFPve_pos      1.3204229049  1.9287300430
## FC_Mpos_TACCGAACAACCAGAG-1FC_MI_GFPve_pos      1.8758289985  1.8849153839
## FC_Mpos_TACGCTCTCAGCGCGT-1FC_MI_GFPve_pos      3.1961891823  1.2632732712
## FC_Mpos_TACGTCCAGGATTTAG-1FC_MI_GFPve_pos      1.0875882082 -0.4840573944
## FC_Mpos_TACTTACGTCCCTCAT-1FC_MI_GFPve_pos     -0.5280946918 -3.8471253790
## FC_Mpos_TAGACTGCACTACACA-1FC_MI_GFPve_pos      3.0309348278 -0.5491766609
## FC_Mpos_TAGACTGTCAGCCTTC-1FC_MI_GFPve_pos      2.4440493755 -1.8720061459
## FC_Mpos_TAGCACATCATGGTAC-1FC_MI_GFPve_pos      3.0152149372 -1.1399023213
## FC_Mpos_TAGCACATCCATCACC-1FC_MI_GFPve_pos      2.6881911926  1.3292942367
## FC_Mpos_TATATCCGTACGACAG-1FC_MI_GFPve_pos      2.6724021606 -1.4171154179
## FC_Mpos_TATTTCGGTGGTACAG-1FC_MI_GFPve_pos      2.0890057258  1.8365865074
## FC_Mpos_TCAAGACTCTACGCAA-1FC_MI_GFPve_pos      1.9887299709  0.5476584755
## FC_Mpos_TCAATCTTCTCTATGT-1FC_MI_GFPve_pos      1.7090295725  1.0314851127
## FC_Mpos_TCAATCTTCTTTCAGT-1FC_MI_GFPve_pos      2.5565416984 -0.6001066841
## FC_Mpos_TCAATTCTCTTTCCGG-1FC_MI_GFPve_pos      2.8092613392  1.3702855430
```

```
## FC_Mpos_TCACACCTCGAGCTGC-1FC_MI_GFPve_pos      2.1620545559  0.0332904182
## FC_Mpos_TCACATTAGGTAAACT-1FC_MI_GFPve_pos      2.7849633865  1.2505078636
## FC_Mpos_TCACGGGCATTCGGGC-1FC_MI_GFPve_pos      3.0253985100  1.5860753380
## FC_Mpos_TCACGGGGTTCTTAGG-1FC_MI_GFPve_pos      3.1391589813 -0.9028305687
## FC_Mpos_TCAGCCTGTATCGTGT-1FC_MI_GFPve_pos      2.2397039108  1.3524484955
## FC_Mpos_TCAGTTTAGAGACAAG-1FC_MI_GFPve_pos      1.2690800600  1.1110954605
## FC_Mpos_TCATACTGTACTAAGA-1FC_MI_GFPve_pos      2.2832036190 -1.9557659306
## FC_Mpos_TCATCCGAGATACCAA-1FC_MI_GFPve_pos      2.0249519520 -1.2657732643
## FC_Mpos_TCATGTTGTACTAAGA-1FC_MI_GFPve_pos      3.0725274257  0.9684248291
## FC_Mpos_TCATTCAAGCAGATAT-1FC_MI_GFPve_pos      2.7077741794  0.1094789825
## FC_Mpos_TCATTGTCAATGTCAC-1FC_MI_GFPve_pos      0.0478514157 -3.4646421351
## FC_Mpos_TCCAGAATCTCTCTAA-1FC_MI_GFPve_pos      1.3047937326  1.1102581344
## FC_Mpos_TCCCATGTCACGTCCT-1FC_MI_GFPve_pos      2.0301027469  2.0758800827
## FC_Mpos_TCCGAAACACTGGACC-1FC_MI_GFPve_pos      2.5240876846 -1.1809324898
## FC_Mpos_TCCGAAACATTGCTTT-1FC_MI_GFPve_pos      0.5713713341 -0.3993148483
## FC_Mpos_TCCTCCCCATGAGGGT-1FC_MI_GFPve_pos     -0.2680549301  1.6606746040
## FC_Mpos_TCCTCGACAAACTGCT-1FC_MI_GFPve_pos      2.7230186634 -0.2730955757
## FC_Mpos_TCCTCTTTCATTGCGA-1FC_MI_GFPve_pos     -0.3186712839 -1.4611172356
## FC_Mpos_TCGAAGTCACTCTGCT-1FC_MI_GFPve_pos      2.2250230484 -1.7549311794
## FC_Mpos_TCGCACTGTGTGGACA-1FC_MI_GFPve_pos      0.9056052141 -0.0939616837
## FC_Mpos_TCGCACTTCGCAACAT-1FC_MI_GFPve_pos     -0.8195912070 -4.3982485511
## FC_Mpos_TCGGTCTCAGTTCTAG-1FC_MI_GFPve_pos      2.9833095245 -0.2239675202
## FC_Mpos_TCGGTCTCATTGCCGG-1FC_MI_GFPve_pos      0.9205272369 -3.0535730995
## FC_Mpos_TCGTCCACACGGGCTT-1FC_MI_GFPve_pos      1.9435410671  1.9034414612
## FC_Mpos_TCGTCCACAGTAGGAC-1FC_MI_GFPve_pos      0.6610490255  0.1912155472
## FC_Mpos_TCTCAGCAGTTTCGAC-1FC_MI_GFPve_pos      0.4614954166 -0.5823025383
## FC_Mpos_TCTCAGCTCGGTGTAT-1FC_MI_GFPve_pos     -0.3305589534  0.6421490036
## FC_Mpos_TCTCCGAAGAAGATCT-1FC_MI_GFPve_pos      2.5757680111  2.2679667793
## FC_Mpos_TCTGGCTGTCTACATG-1FC_MI_GFPve_pos      2.0931849651 -0.6091274895
## FC_Mpos_TCTTAGTGTTAGAAAC-1FC_MI_GFPve_pos      2.4356630020 -1.6367215790
## FC_Mpos_TCTTTGACATGACTCA-1FC_MI_GFPve_pos      0.9423550539  2.0509372078
## FC_Mpos_TGAATGCTCCTCATAT-1FC_MI_GFPve_pos      0.5774020724 -0.2897638954
## FC_Mpos_TGACAGTAGTAGAGTT-1FC_MI_GFPve_pos      0.3759908848  1.2940345131
## FC_Mpos_TGACAGTGTGTGATGG-1FC_MI_GFPve_pos      2.9664662056  1.7112326942
## FC_Mpos_TGACCCTAGCACTCGC-1FC_MI_GFPve_pos      3.0171628170  0.1768665634
## FC_Mpos_TGACGCGCAGACCATT-1FC_MI_GFPve_pos      1.9798920326 -0.6594366707
## FC_Mpos_TGACTCCCATGTTACG-1FC_MI_GFPve_pos      0.9942958526  1.9252119385
## FC_Mpos_TGACTCCTCTGGACCG-1FC_MI_GFPve_pos      2.0318155460 -1.7840222992
## FC_Mpos_TGAGACTTCCCTCTAG-1FC_MI_GFPve_pos     -0.9477109738 -1.5387236752
## FC_Mpos_TGAGCATTCTTACCAT-1FC_MI_GFPve_pos      2.7846901588 -0.6257676758
## FC_Mpos_TGATGGTGTGGATCGA-1FC_MI_GFPve_pos      2.4639089279 -1.6107647099
## FC_Mpos_TGCAGATTCTGGGCCA-1FC_MI_GFPve_pos      0.8309459858 -0.8958954491
## FC_Mpos_TGGAGAGAGGTCGAGT-1FC_MI_GFPve_pos      3.1574356727  1.1207285248
## FC_Mpos_TGGGCGTAGATTGAGT-1FC_MI_GFPve_pos      2.1986935310 -0.7181024231
## FC_Mpos_TGTAACGAGGGCATGT-1FC_MI_GFPve_pos      1.5321965389  1.3243322693
## FC_Mpos_TGTGGCGGTGAAAGTT-1FC_MI_GFPve_pos      2.1519761257  1.2036395393
## FC_Mpos_TGTGTGACAAGACTGG-1FC_MI_GFPve_pos      1.7000315361  1.9646344505
## FC_Mpos_TGTTCTATCGCGGACT-1FC_MI_GFPve_pos      1.6407309942  1.5478668533
## FC_Mpos_TTAATCCCACGCTGCA-1FC_MI_GFPve_pos      2.7578003578 -0.7979755081
## FC_Mpos_TTACCGCTCGACATTG-1FC_MI_GFPve_pos     -0.7245061822 -3.6827212252
## FC_Mpos_TTACTGTCACACCAGC-1FC_MI_GFPve_pos      2.5526144676  0.2535600982
## FC_Mpos_TTACTGTTCACCATGA-1FC_MI_GFPve_pos      0.5079449587  1.2042160354
## FC_Mpos_TTACTGTTCTAGACCA-1FC_MI_GFPve_pos      0.9871406727 -1.1390037216
## FC_Mpos_TTAGGCATCCTAGCTC-1FC_MI_GFPve_pos      2.1201894455 -1.9037203469
## FC_Mpos_TTATTGCCAATCGCAT-1FC_MI_GFPve_pos      1.6280514173 -1.8129472412
## FC_Mpos_TTCCTTCTCCCGAGTG-1FC_MI_GFPve_pos     -0.8272215553 -3.6023455538
## FC_Mpos_TTCTAGTCAATCGAAA-1FC_MI_GFPve_pos      2.4336428814  1.3754878364
## FC_Mpos_TTCTCTCAGATACTGA-1FC_MI_GFPve_pos     -0.4843022056  1.1109404884
## FC_Mpos_TTCTGTAAGCCTTTGA-1FC_MI_GFPve_pos      2.8934877090 -0.9391152538
## FC_Mpos_TTCTTCCCACCGTGCA-1FC_MI_GFPve_pos      2.7982523613 -1.1794926800
## FC_Mpos_TTCTTCCGTGGTTTAC-1FC_MI_GFPve_pos      2.7515964680 -0.5107178368
## FC_Mpos_TTCTTCCTCGGCCAAC-1FC_MI_GFPve_pos      0.4931950145 -0.6856774964
## FC_Mpos_TTGAACGGTGATCATC-1FC_MI_GFPve_pos      1.5383385591  1.4557781540
## FC_Mpos_TTGGGATGTCACTGAT-1FC_MI_GFPve_pos      2.4798867874  0.9784641586
## FC_Mpos_TTGGTTTGTAAGGAGA-1FC_MI_GFPve_pos      2.5335650615  0.2326217018
## FC_Mpos_TTGTTCACAGGCCCTA-1FC_MI_GFPve_pos      2.3565223388 -0.0776304878
## FC_Mpos_TTGTTGTCAAATTGCC-1FC_MI_GFPve_pos      0.9072137050 -0.1710223831
## FC_Fpos_AACCCAAGTCAACATC-1FC_FI_GFPve_pos      1.6557761602  1.4274998031
## FC_Fpos_AACCTTTTCCGAACGC-1FC_FI_GFPve_pos     -1.4402711220  0.9454331718
## FC_Fpos_AAGACTCAGTATCTGC-1FC_FI_GFPve_pos      2.2922687702  0.9236455284
```

```
## FC_Fpos_AAGACTCCACGGTGAA-1FC_FI_GFPve_pos      2.9864149265 -1.0157451309
## FC_Fpos_AAGCATCCAAAGGCAC-1FC_FI_GFPve_pos     -1.1560637183  0.8698859535
## FC_Fpos_AATGCCAAGGGTTGCA-1FC_FI_GFPve_pos     -0.0706056408  0.8748979889
## FC_Fpos_AATGGAACAATTGAGA-1FC_FI_GFPve_pos      2.1569857769  1.8244781814
## FC_Fpos_AATGGCTGTTCTTGCC-1FC_FI_GFPve_pos      2.4516761474  1.2884159408
## FC_Fpos_AATTCCTGTAGCTTAC-1FC_FI_GFPve_pos      1.9800822906  1.1337738357
## FC_Fpos_ACAGAAAGTGGTCAAG-1FC_FI_GFPve_pos     -0.2232615068 -1.7140831627
## FC_Fpos_ACAGCCGGTCCTGGGT-1FC_FI_GFPve_pos     -0.4651184805 -3.8412641563
## FC_Fpos_ACATGCATCACCTTGC-1FC_FI_GFPve_pos     -2.0531140394  2.0821400009
## FC_Fpos_ACCAAACGTCTACATG-1FC_FI_GFPve_pos      2.5898575954  1.1968741737
## FC_Fpos_ACCAACATCTCGAACA-1FC_FI_GFPve_pos     -0.6011057325  0.7890029274
## FC_Fpos_ACCACAATCAACTTTC-1FC_FI_GFPve_pos      0.1447123163  2.5260725342
## FC_Fpos_ACCTACCAGGTCACTT-1FC_FI_GFPve_pos      0.6829369836 -3.7852955736
## FC_Fpos_ACCTGAAGTTTACACG-1FC_FI_GFPve_pos      1.9016018085  0.5987863861
## FC_Fpos_ACGTCCTAGGTTGTTC-1FC_FI_GFPve_pos      1.8203063183  1.2142615639
## FC_Fpos_ACTGTCCCATCCGCGA-1FC_FI_GFPve_pos      1.7422523670  1.3971829735
## FC_Fpos_AGAAATGCACTTGAGT-1FC_FI_GFPve_pos      2.1274490528  0.3655882202
## FC_Fpos_AGAAGTAAGGTACTGG-1FC_FI_GFPve_pos      2.0112207107  0.0012026153
## FC_Fpos_AGACACTCAAATTGGA-1FC_FI_GFPve_pos     -0.8669166990 -3.1739545740
## FC_Fpos_AGACAGGAGCACTAAA-1FC_FI_GFPve_pos      1.4651287966  1.4845648132
## FC_Fpos_AGAGAATTCTCAGGCG-1FC_FI_GFPve_pos      1.0554057531  1.9738350235
## FC_Fpos_AGAGAATTCTGTAAGC-1FC_FI_GFPve_pos      2.3070841007 -1.5489093937
## FC_Fpos_AGATCCATCCGCACGA-1FC_FI_GFPve_pos      2.1163311176  1.6487627350
## FC_Fpos_AGCCAGCGTTCCGGTG-1FC_FI_GFPve_pos      2.4235599212  1.7787075363
## FC_Fpos_AGGCATTCAGCGTATT-1FC_FI_GFPve_pos      0.9217875175  1.7068996750
## FC_Fpos_AGGTCATAGGATTCCT-1FC_FI_GFPve_pos      2.5854909591  2.1721349083
## FC_Fpos_AGTCACAGTAGTTACC-1FC_FI_GFPve_pos      2.5506923370  1.4478788696
## FC_Fpos_ATACCTTTCTTTCCGG-1FC_FI_GFPve_pos      3.1198158436 -0.5845708527
## FC_Fpos_ATCCACCGTATAGGAT-1FC_FI_GFPve_pos      0.4611579351  2.2294659935
## FC_Fpos_ATCCACCTCTCGACGG-1FC_FI_GFPve_pos      2.7232597046  2.0077433906
## FC_Fpos_ATCCCTGTCCCATACC-1FC_FI_GFPve_pos      2.2992465668  0.5092053734
## FC_Fpos_ATCCCTGTCGGCTATA-1FC_FI_GFPve_pos      2.8375282459 -1.0949453987
## FC_Fpos_ATTACCTTCGCCATAA-1FC_FI_GFPve_pos      2.5090351276  0.7049217544
## FC_Fpos_ATTACTCTCTACCAGA-1FC_FI_GFPve_pos     -1.3842928238  1.0897980057
## FC_Fpos_ATTCGTTTCAACCGAT-1FC_FI_GFPve_pos      2.2837073974  1.5374760948
## FC_Fpos_ATTCTACCAGAGTCAG-1FC_FI_GFPve_pos      1.5784616642  2.1126070343
## FC_Fpos_ATTCTACCAGCTGGTC-1FC_FI_GFPve_pos      1.5284772091  1.5276656471
## FC_Fpos_ATTTACCTCCTCATAT-1FC_FI_GFPve_pos      3.0353231602 -0.6861848511
## FC_Fpos_CAACGATTCTATTCGT-1FC_FI_GFPve_pos      0.8319544964  0.9942059837
## FC_Fpos_CAAGGGACATGACGGA-1FC_FI_GFPve_pos      1.8378436737  2.5298700653
## FC_Fpos_CACAACAGTTGACTAC-1FC_FI_GFPve_pos      3.1255626850 -0.7497644104
## FC_Fpos_CACATGACAATGTGGG-1FC_FI_GFPve_pos      1.4724209480 -0.8620497860
## FC_Fpos_CACCGTTCAACCAATC-1FC_FI_GFPve_pos      0.0165124707  0.6501427017
## FC_Fpos_CACCGTTGTTTGACAC-1FC_FI_GFPve_pos      2.8875012569 -0.7368368782
## FC_Fpos_CACGGGTTCATGGTAC-1FC_FI_GFPve_pos      1.7786386184  0.8065476738
## FC_Fpos_CAGCAATCAAGATGGC-1FC_FI_GFPve_pos      2.8447928600  1.8336487137
## FC_Fpos_CAGCACGAGAGCAAGA-1FC_FI_GFPve_pos      1.8476662807  2.1207404457
## FC_Fpos_CAGCACGAGGACTTCT-1FC_FI_GFPve_pos      2.2315986328  0.2886123977
## FC_Fpos_CATACAGCAGACAAGC-1FC_FI_GFPve_pos      1.6413284712  1.7302322708
## FC_Fpos_CATAGACTCTCCACTG-1FC_FI_GFPve_pos      1.9979183845 -0.1297216095
## FC_Fpos_CATTCATTCCTGCTAC-1FC_FI_GFPve_pos      2.0729131870  0.3312092148
## FC_Fpos_CATTGCCTCGGCTCTT-1FC_FI_GFPve_pos      1.0022280388  1.2281098686
## FC_Fpos_CCAATGATCTCTGCCA-1FC_FI_GFPve_pos      3.1869297199  1.6872153602
## FC_Fpos_CCACAAAGTTGCCGAC-1FC_FI_GFPve_pos     -0.0268246062  0.2326412521
## FC_Fpos_CCACCATCATACCACA-1FC_FI_GFPve_pos      1.9096944504  0.6722693764
## FC_Fpos_CCACGTTTCATATGGC-1FC_FI_GFPve_pos      2.4142208271  1.7075267158
## FC_Fpos_CCACTTGTCATACAGC-1FC_FI_GFPve_pos      0.4020337634  0.7118721329
## FC_Fpos_CCCAACTCACCGTGAC-1FC_FI_GFPve_pos      2.6712532215  1.5901222549
## FC_Fpos_CCCATTGTCGCCATAA-1FC_FI_GFPve_pos      3.1230220966 -0.5503978409
## FC_Fpos_CCCTCTCCAACTTGCA-1FC_FI_GFPve_pos      2.2360692196 -0.7041725792
## FC_Fpos_CCCTCTCGTAAGGCTG-1FC_FI_GFPve_pos      2.3293199711  0.6621704422
## FC_Fpos_CCGGACATCATATGGC-1FC_FI_GFPve_pos      2.1587755852  0.7004466377
## FC_Fpos_CCGGACATCCGTAATG-1FC_FI_GFPve_pos      2.3433914356 -0.9214727558
## FC_Fpos_CCTATCGTCAAGTAAG-1FC_FI_GFPve_pos      2.0384285621  1.7185807548
## FC_Fpos_CCTCAACCACAAGTTC-1FC_FI_GFPve_pos      1.9483304195  0.8227439247
## FC_Fpos_CCTCACACAGAACGCA-1FC_FI_GFPve_pos      1.7993499212  1.1046443306
## FC_Fpos_CCTGCATGTGCCTTCT-1FC_FI_GFPve_pos      1.0865782671  0.8241925560
## FC_Fpos_CCTGTTGAGCTTACGT-1FC_FI_GFPve_pos      2.0919895344  1.4297170959
## FC_Fpos_CCTTCAGTCGGCATTA-1FC_FI_GFPve_pos      0.8238587551 -1.1187708058
## FC_Fpos_CGCCATTTCGCATGAT-1FC_FI_GFPve_pos      2.7859156303  0.2682786308
```

```
## FC_Fpos_CGGAATTTCCTCATAT-1FC_FI_GFPve_pos     -0.0810043491 -1.2072701134
## FC_Fpos_CGGGACTAGACTGAGC-1FC_FI_GFPve_pos     -0.4336685367  0.0566859566
## FC_Fpos_CGGGTCAGTGCCAAGA-1FC_FI_GFPve_pos      1.5818363600  1.7236128174
## FC_Fpos_CGTAATGGTGTTAAAG-1FC_FI_GFPve_pos      1.0639898948  1.2445288025
## FC_Fpos_CTACAGACACATATCG-1FC_FI_GFPve_pos      0.9858485632  1.9594216667
## FC_Fpos_CTACCCACACGCACCA-1FC_FI_GFPve_pos      2.9486074619 -0.0050916352
## FC_Fpos_CTACTATAGTACTGTC-1FC_FI_GFPve_pos      1.8641603164 -1.0123772301
## FC_Fpos_CTACTATGTAGTAAGT-1FC_FI_GFPve_pos      2.0851960354 -0.1488952316
## FC_Fpos_CTATAGGAGAGCCCAA-1FC_FI_GFPve_pos     -0.2186837509 -0.7800123371
## FC_Fpos_CTATCCGCACTGAATC-1FC_FI_GFPve_pos      0.6394548588  0.8058038078
## FC_Fpos_CTATCCGGTTGATGTC-1FC_FI_GFPve_pos      2.6477439575  1.6175146423
## FC_Fpos_CTCTGGTCAGTTACCA-1FC_FI_GFPve_pos      1.9131546192  2.1602759682
## FC_Fpos_CTGCCATCAAGCCCAC-1FC_FI_GFPve_pos      2.5193162136 -0.0851282753
## FC_Fpos_CTTTCGGCATCACCAA-1FC_FI_GFPve_pos      2.5347216301  0.0090961777
## FC_Fpos_GAACGTTTCTTACCGC-1FC_FI_GFPve_pos      2.7797916107  0.5667581878
## FC_Fpos_GAACTGTAGTGTTGAA-1FC_FI_GFPve_pos      1.9664981537  1.7066707931
## FC_Fpos_GACATCAGTGAGCTCC-1FC_FI_GFPve_pos      3.1541967563  0.8351407371
## FC_Fpos_GACCAATTCTCGGCTT-1FC_FI_GFPve_pos      0.9617072515 -0.6148557343
## FC_Fpos_GACCAATTCTGACGCG-1FC_FI_GFPve_pos      2.2070574932  1.9596433960
## FC_Fpos_GACTGATTCAGGACAG-1FC_FI_GFPve_pos      2.5957677536  1.1624155365
## FC_Fpos_GACTTCCAGCTCTTCC-1FC_FI_GFPve_pos      0.7957654767  1.8583174072
## FC_Fpos_GAGATGGCAGGATTCT-1FC_FI_GFPve_pos      0.6647322349 -0.4016170181
## FC_Fpos_GAGTTACCAGCTCATA-1FC_FI_GFPve_pos      3.2338805370  1.6582546554
## FC_Fpos_GATCACACAGGCTATT-1FC_FI_GFPve_pos      2.3268196754  0.0456033073
## FC_Fpos_GATCCCTTCAACTTTC-1FC_FI_GFPve_pos      2.2911079101  0.9458079658
## FC_Fpos_GCACATACATTGAGCT-1FC_FI_GFPve_pos      1.9429266624  2.2137666069
## FC_Fpos_GCACGTGTCACGATCA-1FC_FI_GFPve_pos      2.5294077568  0.7528958641
## FC_Fpos_GCCAGTGCAATTGCCA-1FC_FI_GFPve_pos      2.3730905227 -0.4101414360
## FC_Fpos_GCGGATCCACATATGC-1FC_FI_GFPve_pos      1.9398389034  1.9551387153
## FC_Fpos_GCGTTTCTCGCCGAAC-1FC_FI_GFPve_pos      1.8120621614 -2.2066154160
## FC_Fpos_GCTACAAAGGGTCACA-1FC_FI_GFPve_pos      1.9327557258  1.9140801750
## FC_Fpos_GCTCAAACAGAACTAA-1FC_FI_GFPve_pos      2.7106731109  2.2539940201
## FC_Fpos_GGAAGTGGTCACCTTC-1FC_FI_GFPve_pos     -0.0717109792 -0.7546431698
## FC_Fpos_GGAGGTAGTCATGCAT-1FC_FI_GFPve_pos      0.8770457678 -1.6787850537
## FC_Fpos_GGGACTCTCAACACCA-1FC_FI_GFPve_pos      2.7323129349  0.4016366325
## FC_Fpos_GGGAGTATCTAGCAAC-1FC_FI_GFPve_pos      0.4890420608  1.7146010719
## FC_Fpos_GGGCGTTAGGTAGTAT-1FC_FI_GFPve_pos      1.0433267526 -2.8093628563
## FC_Fpos_GGTAACTAGAGTCAGC-1FC_FI_GFPve_pos      0.7502910070  0.2685637794
## FC_Fpos_GGTAACTTCCGTGGTG-1FC_FI_GFPve_pos      2.8698501758  2.4134846054
## FC_Fpos_GGTGAAGTCGAGAGAC-1FC_FI_GFPve_pos      2.2670784168  1.3597503029
## FC_Fpos_GTAGTACAGAGAGCAA-1FC_FI_GFPve_pos      1.1015989952  0.9307389580
## FC_Fpos_GTATTGGTCATAAGGA-1FC_FI_GFPve_pos      1.4837644272  2.2972159706
## FC_Fpos_GTCAAGTGTAGCGTAG-1FC_FI_GFPve_pos      1.9785716705 -1.0291585602
## FC_Fpos_GTCAGCGGTTCCGTTC-1FC_FI_GFPve_pos      0.9088016920  2.1105513893
## FC_Fpos_GTCATCCGTGGGAGAG-1FC_FI_GFPve_pos      2.6912319832 -0.8114189781
## FC_Fpos_GTCCACTAGCTTGTTG-1FC_FI_GFPve_pos      0.1491558425  1.4417486511
## FC_Fpos_GTCGAATGTATCTCTT-1FC_FI_GFPve_pos      0.6063521676  2.0380888305
## FC_Fpos_GTCGCGAAGCACCTGC-1FC_FI_GFPve_pos      1.5883746319  0.2753339134
## FC_Fpos_GTCTACCCACGTCGTG-1FC_FI_GFPve_pos      0.0075292014 -0.4268369354
## FC_Fpos_GTGAGGATCTATTTCG-1FC_FI_GFPve_pos      1.9458935432 -1.4476530232
## FC_Fpos_GTGCGTGGTCTCGGGT-1FC_FI_GFPve_pos      1.5391316585  0.4626512848
## FC_Fpos_GTGGAAGAGGAGTCTG-1FC_FI_GFPve_pos      2.4205110245  0.8017125450
## FC_Fpos_GTTACGATCCAGTACA-1FC_FI_GFPve_pos      3.1370935612  0.1127968155
## FC_Fpos_GTTGTCCCACTGCTTC-1FC_FI_GFPve_pos      2.4903454952  1.6547446571
## FC_Fpos_TAATCTCTCATAGCAC-1FC_FI_GFPve_pos     -0.0741905473  1.4057598434
## FC_Fpos_TACACCCTCTAGCAAC-1FC_FI_GFPve_pos      0.6532327704  0.1745453201
## FC_Fpos_TACATTCGTCAGACTT-1FC_FI_GFPve_pos      1.9882652931  0.4449100815
## FC_Fpos_TACCGAAAGGTAACTA-1FC_FI_GFPve_pos      1.0697287493  0.4222178779
## FC_Fpos_TACTGCCTCTATACGG-1FC_FI_GFPve_pos      1.7026314907  0.7558451019
## FC_Fpos_TAGGAGGAGGGATCAC-1FC_FI_GFPve_pos      2.7186780147  0.1276216827
## FC_Fpos_TAGGTTGAGTGCTACT-1FC_FI_GFPve_pos     -1.1304215617 -1.5838060059
## FC_Fpos_TAGGTTGCAACCGGAA-1FC_FI_GFPve_pos      0.0297813736  1.1577158294
## FC_Fpos_TAGGTTGGTACTCGAT-1FC_FI_GFPve_pos      1.6638034754  1.8370042167
## FC_Fpos_TATACCTCATACACCA-1FC_FI_GFPve_pos      0.4301890306  0.6974292122
## FC_Fpos_TATATCCCAGCTTTGA-1FC_FI_GFPve_pos     -0.4638295896  1.0055632911
## FC_Fpos_TATCGCCTCTAGCATG-1FC_FI_GFPve_pos      1.6966068916  1.9778724037
## FC_Fpos_TATCTGTCAACGGGTA-1FC_FI_GFPve_pos     -0.3203766026  1.0073485695
## FC_Fpos_TCAAGACTCGGTGTTA-1FC_FI_GFPve_pos      0.9830275945  0.9549966179
## FC_Fpos_TCACACCGTTCGTGCG-1FC_FI_GFPve_pos      0.1604500882 -3.7799222745
## FC_Fpos_TCACATTAGCGTCTCG-1FC_FI_GFPve_pos      0.5972997122 -0.3036327042
```

```
## FC_Fpos_TCACGCTCACTTCCTG-1FC_FI_GFPve_pos      1.8261175327  1.0721512161
## FC_Fpos_TCACTCGCACCCAATA-1FC_FI_GFPve_pos      0.8648475580  1.5152764641
## FC_Fpos_TCCCAGTGTGCTCTCT-1FC_FI_GFPve_pos      1.5152018242  2.1366496406
## FC_Fpos_TCCTCTTCATGTAACC-1FC_FI_GFPve_pos      1.0600509815  0.6819606148
## FC_Fpos_TCGGTCTTCTTTCCAA-1FC_FI_GFPve_pos      1.3029287033  1.2893424354
## FC_Fpos_TCTCACGCAACGGCTC-1FC_FI_GFPve_pos      1.2492107324  0.5069775902
## FC_Fpos_TCTGCCATCTACACAG-1FC_FI_GFPve_pos      1.1523150377  1.2470374428
## FC_Fpos_TGATGCAAGGATACAT-1FC_FI_GFPve_pos      1.0288934879  0.8072762810
## FC_Fpos_TGATTTCTCACAGAGG-1FC_FI_GFPve_pos      2.4364612274  1.5763516746
## FC_Fpos_TGCAGGCAGCAGTAAT-1FC_FI_GFPve_pos      2.2642479114  1.3695760093
## FC_Fpos_TGCAGTACAAGTGATA-1FC_FI_GFPve_pos     -0.1856061160 -1.8249840416
## FC_Fpos_TGCCGAGTCTCTGGTC-1FC_FI_GFPve_pos      0.9871667795  1.9378390632
## FC_Fpos_TGCGGGTAGGATGTTA-1FC_FI_GFPve_pos      1.6812975578  0.8543730102
## FC_Fpos_TGGAGAGTCTAGCAAC-1FC_FI_GFPve_pos      2.2452650242  0.8318300567
## FC_Fpos_TGGTAGTTCGCTACGG-1FC_FI_GFPve_pos      1.3381748371  2.2464771591
## FC_Fpos_TGTAAGCGTGGTTTAC-1FC_FI_GFPve_pos      2.7510707550 -0.1004261650
## FC_Fpos_TGTGATGCACCAGGTC-1FC_FI_GFPve_pos      1.7752342396  1.5611911140
## FC_Fpos_TGTTCATTCTTCGGAA-1FC_FI_GFPve_pos      2.6064348392  1.6616692863
## FC_Fpos_TGTTTGTTCCGCCTAT-1FC_FI_GFPve_pos      0.7775230579 -1.5498547234
## FC_Fpos_TTCCACGAGCTCGGCT-1FC_FI_GFPve_pos     -0.1192058191 -1.4327363648
## FC_Fpos_TTCCTCTGTTTCGCTC-1FC_FI_GFPve_pos      2.2693724804 -1.2720677532
## FC_Fpos_TTGCGTCCAGCTACAT-1FC_FI_GFPve_pos      1.7453690939  2.6776595436
## FC_Fpos_TTGGGATGTCAACCTA-1FC_FI_GFPve_pos      2.9384668045  2.2968297325
## FC_Fpos_TTGTTCATCCCTGTTG-1FC_FI_GFPve_pos      2.1455192737  1.7735109649
## FC_Fpos_TTGTTCATCCTGGGAC-1FC_FI_GFPve_pos      0.3030406408 -0.4791631378
## FC_Fpos_TTTCATGTCCAACACA-1FC_FI_GFPve_pos      0.4599463515  1.1048779808
## FC_Fpos_TTTCGATCACACCGCA-1FC_FI_GFPve_pos      0.3833291106  1.0904198013
## FC_Fpos_TTTGACTAGACGGTTG-1FC_FI_GFPve_pos     -0.0097693480  2.0522132240
## dCA2_Mpos_AACAAGATCCGCAACG-1dCA2_MI_GFPve_pos -4.1621639557 -1.8807840027
## dCA2_Mpos_AACAGGGCAGAGACTG-1dCA2_MI_GFPve_pos -3.4044837780 -2.2508279957
## dCA2_Mpos_AACCAACTCGAACTCA-1dCA2_MI_GFPve_pos -4.4428741760 -1.9786574520
## dCA2_Mpos_AACCTGAAGAGAGCAA-1dCA2_MI_GFPve_pos -1.9425376482 -0.4032420792
## dCA2_Mpos_AACCTGATCGAGTCTA-1dCA2_MI_GFPve_pos -3.4737524815 -2.5413765587
## dCA2_Mpos_AACCTTTAGCATCGAG-1dCA2_MI_GFPve_pos -3.9968173332 -0.8945040383
## dCA2_Mpos_AACGAAAAGCATTGAA-1dCA2_MI_GFPve_pos -4.4834453888 -1.7811829724
## dCA2_Mpos_AACGAAATCGGCTGTG-1dCA2_MI_GFPve_pos -3.0710179634 -0.6654224076
## dCA2_Mpos_AAGATAGCAATGAACA-1dCA2_MI_GFPve_pos -3.3031301327 -1.2956218399
## dCA2_Mpos_AAGCGAGAGCCAGTAG-1dCA2_MI_GFPve_pos -3.6780428715 -2.1889648117
## dCA2_Mpos_AAGGAATTCACCATCC-1dCA2_MI_GFPve_pos -4.4221908875 -1.0563258805
## dCA2_Mpos_AAGGAATTCCCTTCCC-1dCA2_MI_GFPve_pos -3.4417471714 -1.1390428223
## dCA2_Mpos_AAGTGAACAAGTGTCT-1dCA2_MI_GFPve_pos -4.1484033890 -1.7025017418
## dCA2_Mpos_AATCACGGTCACAGTT-1dCA2_MI_GFPve_pos -1.3608126469 -2.2871491589
## dCA2_Mpos_AATGAAGCATTCAGCA-1dCA2_MI_GFPve_pos -3.4006395169 -1.8791157879
## dCA2_Mpos_AATGAAGGTAGCGTAG-1dCA2_MI_GFPve_pos -2.5102934666 -0.2384876885
## dCA2_Mpos_AATGACCAGTCACAGG-1dCA2_MI_GFPve_pos -1.8228397198  0.3014211975
## dCA2_Mpos_AATGCCAAGCGACCCT-1dCA2_MI_GFPve_pos -1.8602815695 -1.0829860844
## dCA2_Mpos_AATGCCAAGTCAATCC-1dCA2_MI_GFPve_pos -1.7583615608 -1.9157933869
## dCA2_Mpos_AATGGAAGTCTGTAAC-1dCA2_MI_GFPve_pos -2.9496056862  0.7284622513
## dCA2_Mpos_AATGGCTCAATAACGA-1dCA2_MI_GFPve_pos -3.5069608517 -2.2266060986
## dCA2_Mpos_AATTCCTTCACGGGAA-1dCA2_MI_GFPve_pos -2.9832980461 -1.0670308747
## dCA2_Mpos_AATTTCCAGTTCACTG-1dCA2_MI_GFPve_pos -1.8826676436 -1.4844078697
## dCA2_Mpos_ACAAAGAAGGAACGTC-1dCA2_MI_GFPve_pos -1.5187777109  0.2603206955
## dCA2_Mpos_ACAAAGATCGTTCTGC-1dCA2_MI_GFPve_pos -3.4507398434 -1.3236651100
## dCA2_Mpos_ACAACCAAGCGTTAGG-1dCA2_MI_GFPve_pos -3.8427984543 -1.4421972908
## dCA2_Mpos_ACAAGCTGTTAGCGGA-1dCA2_MI_GFPve_pos -4.0403601952 -2.1614625134
## dCA2_Mpos_ACACAGTTCATAGCAC-1dCA2_MI_GFPve_pos -2.3391399212 -0.2460345902
## dCA2_Mpos_ACAGGGAAGTTCGGTT-1dCA2_MI_GFPve_pos -3.3174021073 -1.1236862816
## dCA2_Mpos_ACATTTCAGCTACAAA-1dCA2_MI_GFPve_pos -4.1565468140 -2.0822817959
## dCA2_Mpos_ACCAAACGTCATCGGC-1dCA2_MI_GFPve_pos -3.9486734696 -2.6589202561
## dCA2_Mpos_ACCAACAAGGTTGGAC-1dCA2_MI_GFPve_pos -2.3897428341  0.0400686584
## dCA2_Mpos_ACCCTCAAGAGTCCGA-1dCA2_MI_GFPve_pos -4.2757193871 -1.1315309681
## dCA2_Mpos_ACCCTCACAGGTACGA-1dCA2_MI_GFPve_pos -1.8455278702  0.1084652267
## dCA2_Mpos_ACCCTTGCAATGCAGG-1dCA2_MI_GFPve_pos -2.9937174149 -1.2013742604
## dCA2_Mpos_ACCTGTCAGCATCGAG-1dCA2_MI_GFPve_pos -4.5254919357 -1.9229452290
## dCA2_Mpos_ACGCACGGTTGGGATG-1dCA2_MI_GFPve_pos -3.7779428787 -0.3784861244
## dCA2_Mpos_ACGGTCGGTCCCTCAT-1dCA2_MI_GFPve_pos -4.1423475571 -2.5033676304
## dCA2_Mpos_ACGGTTATCAACACGT-1dCA2_MI_GFPve_pos -2.9042372532 -1.4240929760
## dCA2_Mpos_ACTATGGGTGCCCTTT-1dCA2_MI_GFPve_pos -1.7023234196 -0.6136669792
## dCA2_Mpos_ACTATTCTCTAGTGTG-1dCA2_MI_GFPve_pos -3.6069717236 -1.7098393120
## dCA2_Mpos_ACTGATGTCCTACGGG-1dCA2_MI_GFPve_pos -2.2314991780 -1.5364293732
```

```
## dCA2_Mpos_ACTGTGAGTAAGTAGT-1dCA2_MI_GFPve_pos -2.4312903710 -0.1418862023
## dCA2_Mpos_ACTTATCTCAATCAGC-1dCA2_MI_GFPve_pos -1.5014122553 -2.3027558006
## dCA2_Mpos_ACTTCGCCATGGCACC-1dCA2_MI_GFPve_pos -3.1695723362 -2.2920078911
## dCA2_Mpos_ACTTCGCGTGCAATAA-1dCA2_MI_GFPve_pos -3.4454815216 -2.0828363575
## dCA2_Mpos_ACTTCGCTCTAGTGTG-1dCA2_MI_GFPve_pos -3.4991547890 -0.2777127899
## dCA2_Mpos_AGAAATGGTAACATGA-1dCA2_MI_GFPve_pos -4.3152415581 -2.0264990010
## dCA2_Mpos_AGAACCTAGTCCGTCG-1dCA2_MI_GFPve_pos -2.0361995526  0.8155498825
## dCA2_Mpos_AGACCCGAGAGGCTGT-1dCA2_MI_GFPve_pos -2.9097246952 -0.7818467297
## dCA2_Mpos_AGAGAATGTACCGTCG-1dCA2_MI_GFPve_pos -2.1331951447  0.8343496643
## dCA2_Mpos_AGAGAGCTCAGCTGTA-1dCA2_MI_GFPve_pos -2.8519244023 -0.9685411133
## dCA2_Mpos_AGATCCACAATGGCAG-1dCA2_MI_GFPve_pos -2.6982459851 -1.4416861214
## dCA2_Mpos_AGATCCATCATTTGGG-1dCA2_MI_GFPve_pos -2.4010259934  0.6642222725
## dCA2_Mpos_AGCGATTCAATTGCAC-1dCA2_MI_GFPve_pos -3.3831090756 -2.3969919361
## dCA2_Mpos_AGCGATTTCCTGTAAG-1dCA2_MI_GFPve_pos -1.8972650595 -0.9383792557
## dCA2_Mpos_AGCGCCAGTAGGTACG-1dCA2_MI_GFPve_pos -3.0759579964 -0.3038835205
## dCA2_Mpos_AGCGCTGAGATGGCGT-1dCA2_MI_GFPve_pos -3.0288042851 -1.6469487824
## dCA2_Mpos_AGGACGATCTTGATTC-1dCA2_MI_GFPve_pos -4.2673027344 -1.9255673565
## dCA2_Mpos_AGGATCTGTAAGAACT-1dCA2_MI_GFPve_pos -2.5061106510  0.0886736236
## dCA2_Mpos_AGGCCACAGCGGACAT-1dCA2_MI_GFPve_pos -4.0420157738 -1.9209303536
## dCA2_Mpos_AGGGAGTGTTGCTAGT-1dCA2_MI_GFPve_pos -4.1764271088 -1.6070534863
## dCA2_Mpos_AGGGCTCCAAGATGGC-1dCA2_MI_GFPve_pos -3.3696801491 -1.4935352482
## dCA2_Mpos_AGGGCTCGTGGAGAAA-1dCA2_MI_GFPve_pos -2.1621187754  0.1643948875
## dCA2_Mpos_AGGTCATGTGCATACT-1dCA2_MI_GFPve_pos -2.6166047878 -1.1096784748
## dCA2_Mpos_AGGTTGTAGCAATTAG-1dCA2_MI_GFPve_pos -4.8135559387 -1.9716679730
## dCA2_Mpos_AGTAGTCTCGGTTGTA-1dCA2_MI_GFPve_pos -4.4668152161 -1.0053291001
## dCA2_Mpos_AGTCTCCAGGAAGAAC-1dCA2_MI_GFPve_pos -4.2529961891 -1.7298786320
## dCA2_Mpos_AGTCTCCAGGTACAAT-1dCA2_MI_GFPve_pos -3.3575248547 -2.0852212586
## dCA2_Mpos_AGTGATCCATGACTTG-1dCA2_MI_GFPve_pos -4.3226949997 -1.0618481316
## dCA2_Mpos_AGTGCCGTCTCATTGT-1dCA2_MI_GFPve_pos -4.3793470688 -2.1613657154
## dCA2_Mpos_AGTGTTGGTTGACTGT-1dCA2_MI_GFPve_pos -2.7840275593 -0.8825072922
## dCA2_Mpos_ATACTTCGTGTTCATG-1dCA2_MI_GFPve_pos -3.0429234333  0.1017551742
## dCA2_Mpos_ATAGACCGTTGTGGAG-1dCA2_MI_GFPve_pos -4.6283314057 -1.0055138745
## dCA2_Mpos_ATAGAGAGTACACGTT-1dCA2_MI_GFPve_pos -2.7706265278  0.0124931656
## dCA2_Mpos_ATCACGAAGTGAACAT-1dCA2_MI_GFPve_pos -4.1018743344 -2.5207979359
## dCA2_Mpos_ATCACGAGTTCTAACG-1dCA2_MI_GFPve_pos -2.6693844624 -0.5462526955
## dCA2_Mpos_ATCAGGTTCATTCATC-1dCA2_MI_GFPve_pos -3.5638422795 -1.4115349926
## dCA2_Mpos_ATCCACCAGACGGAAA-1dCA2_MI_GFPve_pos -3.5078670807 -2.0209750809
## dCA2_Mpos_ATCCATTGTGTTGATC-1dCA2_MI_GFPve_pos -3.1037859745 -0.5148272194
## dCA2_Mpos_ATCCCTGCAAGAGAGA-1dCA2_MI_GFPve_pos -1.8042335339 -0.5233540215
## dCA2_Mpos_ATCCTATCAACGGCCT-1dCA2_MI_GFPve_pos -3.1422893830 -2.2649991192
## dCA2_Mpos_ATCCTATCACTTGGGC-1dCA2_MI_GFPve_pos -3.8443274327 -2.1518649735
## dCA2_Mpos_ATCGTAGAGCTCGTGC-1dCA2_MI_GFPve_pos -4.2493645973 -1.0547089256
## dCA2_Mpos_ATCGTAGGTCTTGAGT-1dCA2_MI_GFPve_pos -3.7749035187 -2.7204107918
## dCA2_Mpos_ATCTCTAAGACGCCAA-1dCA2_MI_GFPve_pos -1.4847707577 -0.6554522194
## dCA2_Mpos_ATGAAAGTCATTTGGG-1dCA2_MI_GFPve_pos -2.6558155842 -0.7579927124
## dCA2_Mpos_ATGAGGGAGCTCCATA-1dCA2_MI_GFPve_pos -2.5185487099  0.4720216118
## dCA2_Mpos_ATGAGTCAGCCATTGT-1dCA2_MI_GFPve_pos -3.9034876652 -2.0624310650
## dCA2_Mpos_ATGCGATTCAAATGAG-1dCA2_MI_GFPve_pos  0.6930028968 -3.4893623270
## dCA2_Mpos_ATGGATCAGTCACTGT-1dCA2_MI_GFPve_pos -4.8239676781 -1.9200422444
## dCA2_Mpos_ATGGATCTCCCGAATA-1dCA2_MI_GFPve_pos -1.9239252634  0.4843960129
## dCA2_Mpos_ATGGGAGTCTCTGCTG-1dCA2_MI_GFPve_pos -3.3177556820 -2.0992970146
## dCA2_Mpos_ATGGGTTCAGTTTCGA-1dCA2_MI_GFPve_pos -2.4791011639  0.2171821914
## dCA2_Mpos_ATGTCTTTCTTCTGTA-1dCA2_MI_GFPve_pos -3.3465354271 -0.7337431587
## dCA2_Mpos_ATTCAGGAGCGTCTCG-1dCA2_MI_GFPve_pos -2.9703833885  0.0007868133
## dCA2_Mpos_ATTCCCGAGTCTTGGT-1dCA2_MI_GFPve_pos -1.9321702547  0.1815529190
## dCA2_Mpos_ATTCTACAGTAGACCG-1dCA2_MI_GFPve_pos -3.8009109326 -1.8695583023
## dCA2_Mpos_ATTCTACTCGACATAC-1dCA2_MI_GFPve_pos -2.2731795140 -1.2246873059
## dCA2_Mpos_ATTTACCAGATAGTCA-1dCA2_MI_GFPve_pos -3.3736774750 -2.0812873520
## dCA2_Mpos_ATTTCTGTCAGTCCGG-1dCA2_MI_GFPve_pos -4.4206073113 -1.4880859532
## dCA2_Mpos_CAACCAAAGTATGGCG-1dCA2_MI_GFPve_pos -1.5980009861 -0.4865688957
## dCA2_Mpos_CAACCTCGTGATATAG-1dCA2_MI_GFPve_pos -2.6743440457 -0.5435304321
## dCA2_Mpos_CAAGACTGTTTAGTCG-1dCA2_MI_GFPve_pos -3.8137669392 -0.9246987976
## dCA2_Mpos_CAAGAGGCAGGGTCTC-1dCA2_MI_GFPve_pos -4.0738758869 -2.4530098118
## dCA2_Mpos_CAAGGGAAGACTTCAC-1dCA2_MI_GFPve_pos -2.2146127053 -0.7668139614
## dCA2_Mpos_CAAGGGACACATGAAA-1dCA2_MI_GFPve_pos -2.2018229790 -0.8655013718
## dCA2_Mpos_CAATGACGTCTTGAAC-1dCA2_MI_GFPve_pos -3.5690197773 -2.0522448696
## dCA2_Mpos_CAATGACGTGACTCGC-1dCA2_MI_GFPve_pos -4.8732201882 -1.7554254212
## dCA2_Mpos_CACAGATAGAGGCGTT-1dCA2_MI_GFPve_pos -4.9035436936 -1.5502154507
## dCA2_Mpos_CACAGGCCAGTCGCTG-1dCA2_MI_GFPve_pos -1.1311504073 -1.3586406388
## dCA2_Mpos_CACATGAGTGAGATTA-1dCA2_MI_GFPve_pos -4.1985938378 -0.8617686905
```

```
## dCA2_Mpos_CACGGGTTCCCAGGAC-1dCA2_MI_GFPve_pos -4.6942260094 -1.2738659062
## dCA2_Mpos_CACTAAGTCCGTAGGC-1dCA2_MI_GFPve_pos -2.8086712189 -1.7915835060
## dCA2_Mpos_CACTAAGTCTCAACGA-1dCA2_MI_GFPve_pos -3.7951681442 -2.8271314778
## dCA2_Mpos_CACTGAATCAAGATAG-1dCA2_MI_GFPve_pos -3.6749832459 -2.8378925003
## dCA2_Mpos_CACTGGGAGACAACAT-1dCA2_MI_GFPve_pos -2.3186452217 -1.4425508656
## dCA2_Mpos_CACTGGGGTAGATCCT-1dCA2_MI_GFPve_pos -3.7870459385 -2.0188901104
## dCA2_Mpos_CAGATCAAGAGTGTTA-1dCA2_MI_GFPve_pos -2.5847256012  0.0501561485
## dCA2_Mpos_CAGATCATCACCATGA-1dCA2_MI_GFPve_pos -3.3853390045 -2.5015797295
## dCA2_Mpos_CAGCAATAGATGATTG-1dCA2_MI_GFPve_pos -3.9416408367 -1.2934517540
## dCA2_Mpos_CAGCAGCCACAACCGC-1dCA2_MI_GFPve_pos -2.5118954010 -1.4088816323
## dCA2_Mpos_CAGCAGCTCTGAGAAA-1dCA2_MI_GFPve_pos -4.3469688721 -1.4789385475
## dCA2_Mpos_CAGCGTGAGTGCTCAT-1dCA2_MI_GFPve_pos -3.8301932640 -1.4256920494
## dCA2_Mpos_CAGGCCATCCCGTGTT-1dCA2_MI_GFPve_pos -3.1976435013 -1.8424184002
## dCA2_Mpos_CAGTGCGCATCCTGTC-1dCA2_MI_GFPve_pos -3.4544486828 -1.3754927792
## dCA2_Mpos_CAGTTAGGTCGTTCAA-1dCA2_MI_GFPve_pos -4.9467556305 -1.8037736096
## dCA2_Mpos_CATCCCATCCATAGGT-1dCA2_MI_GFPve_pos -3.6892461605 -1.5121624150
## dCA2_Mpos_CATGCGGCACTTCAAG-1dCA2_MI_GFPve_pos -3.6514024563 -1.2220437206
## dCA2_Mpos_CATGGATGTACCGTGC-1dCA2_MI_GFPve_pos -4.7080600090 -1.8940002598
## dCA2_Mpos_CATGGTATCCATCTCG-1dCA2_MI_GFPve_pos -3.1440019436 -1.1380529083
## dCA2_Mpos_CATGGTATCCCGGTAG-1dCA2_MI_GFPve_pos -3.6834740467 -1.7908439316
## dCA2_Mpos_CCAAGCGAGGGAGGCA-1dCA2_MI_GFPve_pos -3.8777756520 -2.0338415779
## dCA2_Mpos_CCACCATGTCCTTGTC-1dCA2_MI_GFPve_pos -2.8153300114 -2.0629639305
## dCA2_Mpos_CCATCACAGTTCCTGA-1dCA2_MI_GFPve_pos -2.0418038197 -1.8817236103
## dCA2_Mpos_CCATCACTCAAACCTG-1dCA2_MI_GFPve_pos -1.9017942972 -1.9895546116
## dCA2_Mpos_CCCTGATCACCGTCGA-1dCA2_MI_GFPve_pos -3.2962040730 -1.9889917053
## dCA2_Mpos_CCGAACGTCACTCTTA-1dCA2_MI_GFPve_pos -2.8942467995 -1.0440704502
## dCA2_Mpos_CCGATGGCACCTTCCA-1dCA2_MI_GFPve_pos -2.1203455754  0.0345545135
## dCA2_Mpos_CCGGTAGTCTTCCTAA-1dCA2_MI_GFPve_pos -4.0572326012 -2.7605172075
## dCA2_Mpos_CCGTTCAAGCCGTTGC-1dCA2_MI_GFPve_pos -3.5134885140 -2.2282249607
## dCA2_Mpos_CCGTTCACAATCGCAT-1dCA2_MI_GFPve_pos -3.7402477093 -1.9395122208
## dCA2_Mpos_CCTACGTCATACCGTA-1dCA2_MI_GFPve_pos -1.5365209408  0.4623928390
## dCA2_Mpos_CCTCATGAGTAGTCAA-1dCA2_MI_GFPve_pos -1.5416227408  0.0212288223
## dCA2_Mpos_CCTCATGAGTTAGTAG-1dCA2_MI_GFPve_pos -4.3007972069 -1.4837660469
## dCA2_Mpos_CCTCCTCAGTCATCCA-1dCA2_MI_GFPve_pos -4.5901529618 -2.1409749665
## dCA2_Mpos_CCTCTAGCACTGGCCA-1dCA2_MI_GFPve_pos -2.9496393032 -0.7210797943
## dCA2_Mpos_CCTCTAGTCGTGCTCT-1dCA2_MI_GFPve_pos -2.3677208252 -0.5918903031
## dCA2_Mpos_CCTTCAGTCTAGGCCG-1dCA2_MI_GFPve_pos -3.1734351940 -2.2358212151
## dCA2_Mpos_CCTTGTGAGCGCATCC-1dCA2_MI_GFPve_pos -3.8804709740 -1.3055164494
## dCA2_Mpos_CGAAGGACAACACGTT-1dCA2_MI_GFPve_pos  0.4944220953 -1.8183018841
## dCA2_Mpos_CGAAGTTTCTCAGTCC-1dCA2_MI_GFPve_pos -2.5412385292 -0.5567073502
## dCA2_Mpos_CGAGAAGAGCATCCTA-1dCA2_MI_GFPve_pos -2.8039150067  0.3211732231
## dCA2_Mpos_CGAGAAGAGCGCCTTG-1dCA2_MI_GFPve_pos -4.4665791817 -1.4794606842
## dCA2_Mpos_CGAGGAAAGGTCGACA-1dCA2_MI_GFPve_pos -2.5429498978  0.5181322418
## dCA2_Mpos_CGAGGCTCAAAGACTA-1dCA2_MI_GFPve_pos -3.2074272461 -2.1719093003
## dCA2_Mpos_CGAGGCTTCCAATCCC-1dCA2_MI_GFPve_pos -2.0069403477 -1.3121921696
## dCA2_Mpos_CGAGTTAGTGCTGTCG-1dCA2_MI_GFPve_pos -2.1718997784  0.2197575889
## dCA2_Mpos_CGAGTTAGTGGGCTCT-1dCA2_MI_GFPve_pos -3.6927113362 -1.8790063538
## dCA2_Mpos_CGCATAAGTAGCTGCC-1dCA2_MI_GFPve_pos -2.6385121174 -0.6029653229
## dCA2_Mpos_CGCCAGACACTGCACG-1dCA2_MI_GFPve_pos -1.3108345099  0.7334709488
## dCA2_Mpos_CGCCAGATCCATACAG-1dCA2_MI_GFPve_pos -4.4539682694 -1.7900755085
## dCA2_Mpos_CGCCATTTCGCCTTTG-1dCA2_MI_GFPve_pos -1.9418786593 -0.0252117790
## dCA2_Mpos_CGGAACCCATGGTACT-1dCA2_MI_GFPve_pos -1.6236688919 -0.2487663902
## dCA2_Mpos_CGGAATTGTCGTTATG-1dCA2_MI_GFPve_pos -3.2299387284 -1.9842185654
## dCA2_Mpos_CGGAATTGTTGCATAC-1dCA2_MI_GFPve_pos -3.2437970467 -1.9258906521
## dCA2_Mpos_CGGGTGTAGATCCTAC-1dCA2_MI_GFPve_pos -3.5206551380 -1.6929399647
## dCA2_Mpos_CGTAATGGTGTTATCG-1dCA2_MI_GFPve_pos -3.2266442604 -1.0507545151
## dCA2_Mpos_CGTAGTACAACCACGC-1dCA2_MI_GFPve_pos -1.6475076504 -1.5407988705
## dCA2_Mpos_CGTCAAAAGGCTTAAA-1dCA2_MI_GFPve_pos -3.7241160698 -0.6572370209
## dCA2_Mpos_CGTCAAATCACAGAGG-1dCA2_MI_GFPve_pos -4.1165592499 -1.4362396874
## dCA2_Mpos_CGTCCATAGTCCGCGT-1dCA2_MI_GFPve_pos -2.4860679932  0.1975875221
## dCA2_Mpos_CGTCCATGTCTCGACG-1dCA2_MI_GFPve_pos -2.7469696827 -0.9570095219
## dCA2_Mpos_CGTGAATAGGACTTCT-1dCA2_MI_GFPve_pos -3.9019825287 -0.7371883072
## dCA2_Mpos_CGTGAATTCGCTAATG-1dCA2_MI_GFPve_pos -4.1424882240 -0.7422330059
## dCA2_Mpos_CGTGATATCTTGGATG-1dCA2_MI_GFPve_pos -4.9232785530 -1.4691631474
## dCA2_Mpos_CGTTAGATCATTCGGA-1dCA2_MI_GFPve_pos -3.0926439591 -1.1794094719
## dCA2_Mpos_CTAAGTGGTTTCGTTT-1dCA2_MI_GFPve_pos -3.8613633938 -1.0677024998
## dCA2_Mpos_CTACAGAGTGATTGGG-1dCA2_MI_GFPve_pos -2.4894864388 -0.8612749256
## dCA2_Mpos_CTACCCACATCCGAAT-1dCA2_MI_GFPve_pos -3.0404646225 -1.0283462681
## dCA2_Mpos_CTACGGGGTGACTCTA-1dCA2_MI_GFPve_pos -2.8212566204  0.7971988045
## dCA2_Mpos_CTATCCGCACATACTG-1dCA2_MI_GFPve_pos -3.0389349289 -0.9958510079
```

```
## dCA2_Mpos_CTCAACCGTGAAGCGT-1dCA2_MI_GFPve_pos -3.6827235050 -0.8458819069
## dCA2_Mpos_CTCAGTCCAGATCCTA-1dCA2_MI_GFPve_pos -3.4927384682 -2.7421114124
## dCA2_Mpos_CTCAGTCTCCGACATA-1dCA2_MI_GFPve_pos -3.9468955822 -2.6088070549
## dCA2_Mpos_CTCCACAAGGTTCACT-1dCA2_MI_GFPve_pos -2.9805521794 -0.5357880272
## dCA2_Mpos_CTCCATGCACGCCACA-1dCA2_MI_GFPve_pos -3.7928228207 -2.7900762238
## dCA2_Mpos_CTCCATGCAGCTTCGG-1dCA2_MI_GFPve_pos -0.8446500130  0.8549800239
## dCA2_Mpos_CTCCGATGTCAAGGCA-1dCA2_MI_GFPve_pos -3.3088450260 -2.1694831528
## dCA2_Mpos_CTCTGGTCACCGTCTT-1dCA2_MI_GFPve_pos -3.6781167813 -2.2034883179
## dCA2_Mpos_CTGCATCAGAAACTAC-1dCA2_MI_GFPve_pos -4.5736205406 -1.5410887875
## dCA2_Mpos_CTGCATCCAATAGAGT-1dCA2_MI_GFPve_pos -4.3892457314 -2.1103305496
## dCA2_Mpos_CTGCGAGCAATCTGCA-1dCA2_MI_GFPve_pos -2.2125046082 -0.6699332871
## dCA2_Mpos_CTGCTCAGTACCTATG-1dCA2_MI_GFPve_pos -3.7810971565 -2.3479709305
## dCA2_Mpos_CTGCTCATCGGAAACG-1dCA2_MI_GFPve_pos -4.5913083382 -1.4938065685
## dCA2_Mpos_CTGTACCGTCGTCGGT-1dCA2_MI_GFPve_pos -1.9889097042 -0.6524800934
## dCA2_Mpos_CTGTGAACAGTAGTGG-1dCA2_MI_GFPve_pos -0.3816005565 -0.4402665772
## dCA2_Mpos_CTTAGGAAGTATTGCC-1dCA2_MI_GFPve_pos -4.8968169518 -1.4426750817
## dCA2_Mpos_CTTCCTTTCTTACGGA-1dCA2_MI_GFPve_pos -4.4466607399 -2.1152083554
## dCA2_Mpos_CTTGAGACATAGCTGT-1dCA2_MI_GFPve_pos -4.5733587570 -1.6224882282
## dCA2_Mpos_CTTGATTTCGTGCGAC-1dCA2_MI_GFPve_pos -3.6996698208 -1.3056957401
## dCA2_Mpos_CTTTCAACAATTAGGA-1dCA2_MI_GFPve_pos -2.1335452624  0.5676226936
## dCA2_Mpos_GAACACTAGTTCAACC-1dCA2_MI_GFPve_pos -4.0409130879 -1.4550289787
## dCA2_Mpos_GAACACTCATCCTCAC-1dCA2_MI_GFPve_pos -4.0996274777 -2.6020941414
## dCA2_Mpos_GAACTGTGTCAAGCCC-1dCA2_MI_GFPve_pos -4.0108451672 -0.7065615334
## dCA2_Mpos_GAACTGTGTTTGACAC-1dCA2_MI_GFPve_pos -3.2160949536 -2.1231674828
## dCA2_Mpos_GAAGAATTCGGATTAC-1dCA2_MI_GFPve_pos -3.1648297138 -1.6410507836
## dCA2_Mpos_GAAGCCCTCCGATCGG-1dCA2_MI_GFPve_pos -4.6225740738 -1.5364100613
## dCA2_Mpos_GAAGGACAGTTACGAA-1dCA2_MI_GFPve_pos -3.5893893070 -2.2708837666
## dCA2_Mpos_GAAGGACTCTACGCGG-1dCA2_MI_GFPve_pos -3.0690450497 -1.5454392113
## dCA2_Mpos_GACAGCCAGCTGTACT-1dCA2_MI_GFPve_pos -4.0163035221 -2.5994474568
## dCA2_Mpos_GACATCATCGGCTTCT-1dCA2_MI_GFPve_pos -2.6865758724  0.0547471367
## dCA2_Mpos_GACCCAGGTCTCAAGT-1dCA2_MI_GFPve_pos -1.4569623299 -0.8547510781
## dCA2_Mpos_GACCGTGAGGACGCAT-1dCA2_MI_GFPve_pos -2.3475937672 -0.4469628014
## dCA2_Mpos_GACCGTGTCAGTGTCA-1dCA2_MI_GFPve_pos -1.3570501633 -2.2503833451
## dCA2_Mpos_GACGTTAAGCACTTTG-1dCA2_MI_GFPve_pos -3.5912086792 -2.1371812500
## dCA2_Mpos_GACGTTATCGCACGGT-1dCA2_MI_GFPve_pos -4.7188308068 -2.1528014816
## dCA2_Mpos_GACTTCCCAGCGAACA-1dCA2_MI_GFPve_pos -2.2871630020 -1.9937896408
## dCA2_Mpos_GACTTCCTCTGTGCGG-1dCA2_MI_GFPve_pos -4.8300125427 -1.9273256935
## dCA2_Mpos_GAGACTTCATATCTGG-1dCA2_MI_GFPve_pos -3.5405924149 -2.2507157006
## dCA2_Mpos_GAGCCTGAGCTAAACA-1dCA2_MI_GFPve_pos -3.6875002213 -1.1485156693
## dCA2_Mpos_GAGCCTGCATACACCA-1dCA2_MI_GFPve_pos -3.1993541546 -1.7068748154
## dCA2_Mpos_GAGCCTGGTCGGAACA-1dCA2_MI_GFPve_pos -4.3395120926 -1.0867187657
## dCA2_Mpos_GAGGCCTAGAGCATCG-1dCA2_MI_GFPve_pos -3.9415473767 -1.6590029873
## dCA2_Mpos_GAGGCCTTCATTTCGT-1dCA2_MI_GFPve_pos -1.4107562132 -0.4653234162
## dCA2_Mpos_GAGGGATCACAAGCCC-1dCA2_MI_GFPve_pos -4.4628684349 -1.1864578404
## dCA2_Mpos_GAGTCTACAATCGTCA-1dCA2_MI_GFPve_pos -4.2019703217 -1.5748421826
## dCA2_Mpos_GAGTCTAGTTTCTATC-1dCA2_MI_GFPve_pos -3.8072202034 -1.7669372238
## dCA2_Mpos_GAGTCTATCCACGAAT-1dCA2_MI_GFPve_pos -3.9188520737 -1.6965472378
## dCA2_Mpos_GAGTTACTCTCTATAC-1dCA2_MI_GFPve_pos -2.3464109726  0.5566535316
## dCA2_Mpos_GATCAGTTCTGCCTCA-1dCA2_MI_GFPve_pos -0.8638859816 -2.1468820252
## dCA2_Mpos_GATGCTAAGTCACGAG-1dCA2_MI_GFPve_pos -3.3504931755 -2.2058164753
## dCA2_Mpos_GATGGAGTCAATCTCT-1dCA2_MI_GFPve_pos -4.3136751480 -1.8301138558
## dCA2_Mpos_GCAACATTCACCTCTG-1dCA2_MI_GFPve_pos -1.4303848572 -2.4231133141
## dCA2_Mpos_GCACGGTAGGTACAAT-1dCA2_MI_GFPve_pos -3.3001391716 -0.6035012879
## dCA2_Mpos_GCACGTGTCATGGCCG-1dCA2_MI_GFPve_pos -3.7447488136 -1.6233570255
## dCA2_Mpos_GCACTAAGTACAGAAT-1dCA2_MI_GFPve_pos -4.2786719628 -1.8355965294
## dCA2_Mpos_GCAGCCACACATACTG-1dCA2_MI_GFPve_pos -4.0704545803 -2.1413910069
## dCA2_Mpos_GCATCTCCAGACGGAT-1dCA2_MI_GFPve_pos -3.0107841320  0.3611545883
## dCA2_Mpos_GCATGATCACCAGGTC-1dCA2_MI_GFPve_pos -2.2028915711 -0.6178593315
## dCA2_Mpos_GCCAACGAGTTGCTGT-1dCA2_MI_GFPve_pos -1.3556852169 -0.4053206124
## dCA2_Mpos_GCCAGTGTCAAAGGTA-1dCA2_MI_GFPve_pos -2.8431792088 -1.3257014431
## dCA2_Mpos_GCCATTCGTAATACCC-1dCA2_MI_GFPve_pos -2.7010195084 -1.8452546276
## dCA2_Mpos_GCCGATGGTCCGGACT-1dCA2_MI_GFPve_pos -4.1392190285 -1.9449090637
## dCA2_Mpos_GCGAGAACATATCTGG-1dCA2_MI_GFPve_pos -4.4201681442 -1.6455957569
## dCA2_Mpos_GCGATCGAGCCGATCC-1dCA2_MI_GFPve_pos -1.9771766491 -1.2616836704
## dCA2_Mpos_GCGATCGTCTTCCCAG-1dCA2_MI_GFPve_pos -2.0119912453  0.3339152656
## dCA2_Mpos_GCGGAAAAGAAACACT-1dCA2_MI_GFPve_pos -2.7714016266  0.1593904815
## dCA2_Mpos_GCTGCAGCAATGCTCA-1dCA2_MI_GFPve_pos -4.2144162483 -2.4703325905
## dCA2_Mpos_GCTGGGTTCATTCCTA-1dCA2_MI_GFPve_pos -4.8769271202 -1.6307084240
## dCA2_Mpos_GCTTCACAGGTTCCGC-1dCA2_MI_GFPve_pos -3.8864624329 -1.1227218785
## dCA2_Mpos_GCTTCACAGTTCCTGA-1dCA2_MI_GFPve_pos -3.6746296711 -2.0951120533
```

```
## dCA2_Mpos_GGAATCTTCGAAGGAC-1dCA2_MI_GFPve_pos -3.8714151211 -2.0285451092
## dCA2_Mpos_GGAGATGCAACCCGCA-1dCA2_MI_GFPve_pos -1.5105414219 -1.2220048584
## dCA2_Mpos_GGAGGATAGCAAGGAA-1dCA2_MI_GFPve_pos -4.6352441139 -1.7969622292
## dCA2_Mpos_GGATGTTAGAGAGCCT-1dCA2_MI_GFPve_pos -4.2450420685 -2.6742741741
## dCA2_Mpos_GGCAGTCCAAACACCT-1dCA2_MI_GFPve_pos -4.8983485527 -1.9031138100
## dCA2_Mpos_GGCAGTCGTAGTGGCA-1dCA2_MI_GFPve_pos -3.5574059315 -1.3179680981
## dCA2_Mpos_GGCGTCAAGAACAAGG-1dCA2_MI_GFPve_pos -3.5228724308 -0.4053053536
## dCA2_Mpos_GGCTTGGCACTTACAG-1dCA2_MI_GFPve_pos -3.5190379448 -2.5351569332
## dCA2_Mpos_GGGAAGTGTTAAAGTG-1dCA2_MI_GFPve_pos -2.3193795510  0.7793293320
## dCA2_Mpos_GGGACAATCTCCCTAG-1dCA2_MI_GFPve_pos -3.0199951954 -2.0559527554
## dCA2_Mpos_GGGACCTGTCGATTCA-1dCA2_MI_GFPve_pos -1.3476720877 -2.0033166088
## dCA2_Mpos_GGGAGATAGTCAACAA-1dCA2_MI_GFPve_pos -4.3006479569 -1.2330422081
## dCA2_Mpos_GGGAGTAAGTCTAGAA-1dCA2_MI_GFPve_pos  0.5519170933 -0.4036631264
## dCA2_Mpos_GGGATGATCTGAGAGG-1dCA2_MI_GFPve_pos -2.7768950291 -0.9308366455
## dCA2_Mpos_GGGCCATAGGGTTTCT-1dCA2_MI_GFPve_pos -2.9211118050 -0.8841187634
## dCA2_Mpos_GGGCCATCAGGTGTTT-1dCA2_MI_GFPve_pos -4.4202000923 -1.8527347721
## dCA2_Mpos_GGGCTCACATTCACAG-1dCA2_MI_GFPve_pos -4.1855838127 -2.2029435314
## dCA2_Mpos_GGGTCACGTCTCTCAC-1dCA2_MI_GFPve_pos -3.3537022896 -0.2209820427
## dCA2_Mpos_GGGTGTCAGTACGTCT-1dCA2_MI_GFPve_pos -3.0017683335 -0.4991459526
## dCA2_Mpos_GGTAATCAGCGATTCT-1dCA2_MI_GFPve_pos -4.0097169705 -2.0404174008
## dCA2_Mpos_GGTGATTAGAGGCTGT-1dCA2_MI_GFPve_pos -1.6230856009  0.7187075935
## dCA2_Mpos_GGTGTTAGTCCGAAAG-1dCA2_MI_GFPve_pos -4.5479519196 -1.5924133934
## dCA2_Mpos_GGTGTTAGTTTGTGGT-1dCA2_MI_GFPve_pos -4.5859191246 -1.3942260422
## dCA2_Mpos_GGTTAACGTATCCCTC-1dCA2_MI_GFPve_pos -1.5701192446 -0.4019145645
## dCA2_Mpos_GTAACACGTGTACAGG-1dCA2_MI_GFPve_pos -3.8898799248 -1.9413666405
## dCA2_Mpos_GTAACCAAGTATAACG-1dCA2_MI_GFPve_pos -2.9559230633 -0.8590786137
## dCA2_Mpos_GTAAGTCTCAAGCCAT-1dCA2_MI_GFPve_pos -3.2082478829 -1.2501740135
## dCA2_Mpos_GTACAGTAGGTGAGCT-1dCA2_MI_GFPve_pos -3.6447760887 -0.4966382660
## dCA2_Mpos_GTAGAAAAGACACACG-1dCA2_MI_GFPve_pos -3.7421033211 -1.9305293240
## dCA2_Mpos_GTAGAGGAGTTGCTCA-1dCA2_MI_GFPve_pos -4.5154683418 -1.8771357216
## dCA2_Mpos_GTATTGGTCGATTTCT-1dCA2_MI_GFPve_pos -4.2621586151 -1.3925983586
## dCA2_Mpos_GTATTTCTCCTGGGTG-1dCA2_MI_GFPve_pos  0.3807518058 -3.9624897517
## dCA2_Mpos_GTCAAACGTACATTGC-1dCA2_MI_GFPve_pos -4.2101561852 -2.1942057289
## dCA2_Mpos_GTCACGGAGCTAGTTC-1dCA2_MI_GFPve_pos -3.3017887898 -1.4761478581
## dCA2_Mpos_GTCACGGTCCGCACTT-1dCA2_MI_GFPve_pos -0.9112840123 -1.0039641537
## dCA2_Mpos_GTCAGCGAGAGCAGTC-1dCA2_MI_GFPve_pos -3.0667671986 -0.1232156433
## dCA2_Mpos_GTCATTTAGTCCGCCA-1dCA2_MI_GFPve_pos -3.2123887367 -0.7507760205
## dCA2_Mpos_GTCATTTCATAGACTC-1dCA2_MI_GFPve_pos -2.6263391800 -0.3697271027
## dCA2_Mpos_GTCCCATTCGTCCTCA-1dCA2_MI_GFPve_pos -4.3323352165 -1.5718562283
## dCA2_Mpos_GTCGCGAGTTGTACGT-1dCA2_MI_GFPve_pos -1.9607165880  1.2555499397
## dCA2_Mpos_GTCGTTCCACGTCATA-1dCA2_MI_GFPve_pos -3.5989506073 -0.2048496880
## dCA2_Mpos_GTCTACCAGAAGTATC-1dCA2_MI_GFPve_pos -2.7235483952 -1.9409970917
## dCA2_Mpos_GTCTAGATCGCCTTGT-1dCA2_MI_GFPve_pos -3.7373752423 -1.8450445809
## dCA2_Mpos_GTCTCACAGTAAGAGG-1dCA2_MI_GFPve_pos -4.8471839256 -1.8908664860
## dCA2_Mpos_GTCTGTCAGCAGGCTA-1dCA2_MI_GFPve_pos -4.0524744816 -1.5659348644
## dCA2_Mpos_GTCTGTCGTCCGACGT-1dCA2_MI_GFPve_pos -4.5805237122 -2.1692564167
## dCA2_Mpos_GTGAGCCTCAGACCGC-1dCA2_MI_GFPve_pos -3.1665892430 -2.2356071152
## dCA2_Mpos_GTGAGCCTCTTACTGT-1dCA2_MI_GFPve_pos -3.8448316879 -2.0062923111
## dCA2_Mpos_GTGAGTTTCGTGCACG-1dCA2_MI_GFPve_pos -2.4545738526  0.6419225059
## dCA2_Mpos_GTGCTGGGTGGTATGG-1dCA2_MI_GFPve_pos -1.7028271980  0.3410477958
## dCA2_Mpos_GTGCTGGTCCAGTGTA-1dCA2_MI_GFPve_pos -2.2900116272 -0.3327707924
## dCA2_Mpos_GTGCTTCAGACCTGGA-1dCA2_MI_GFPve_pos -3.4643201657 -0.4738731064
## dCA2_Mpos_GTGGAGAGTATGAAGT-1dCA2_MI_GFPve_pos -2.6052918263 -0.0444311775
## dCA2_Mpos_GTGGAGAGTTCTATCT-1dCA2_MI_GFPve_pos -4.6482307739 -1.3120791592
## dCA2_Mpos_GTGTAACCAGTTCCAA-1dCA2_MI_GFPve_pos -2.1048461027 -0.1878542580
## dCA2_Mpos_GTGTGGCGTAGCTTTG-1dCA2_MI_GFPve_pos -3.3377606697 -2.0497302689
## dCA2_Mpos_GTTACGAGTACTTGTG-1dCA2_MI_GFPve_pos -3.9350008793 -2.6429512181
## dCA2_Mpos_GTTAGACGTACTGTTG-1dCA2_MI_GFPve_pos -0.6647334821 -0.4370736756
## dCA2_Mpos_GTTAGTGCAGGTATGG-1dCA2_MI_GFPve_pos -3.8316981621 -2.1798536457
## dCA2_Mpos_GTTCCGTGTGTAAACA-1dCA2_MI_GFPve_pos -2.6744410820  0.3924136482
## dCA2_Mpos_GTTCGCTAGAGCCCAA-1dCA2_MI_GFPve_pos -1.9682025738 -0.5982656159
## dCA2_Mpos_GTTCGCTTCCAAACCA-1dCA2_MI_GFPve_pos -1.4873679705 -2.2298354782
## dCA2_Mpos_GTTGTCCGTAGACTGG-1dCA2_MI_GFPve_pos -4.8398325272 -1.4486884751
## dCA2_Mpos_TAACTTCGTGTTGAGG-1dCA2_MI_GFPve_pos -1.8191003628  0.2559719406
## dCA2_Mpos_TAAGCGTAGAATACAC-1dCA2_MI_GFPve_pos -4.6747982331 -1.4957267918
## dCA2_Mpos_TAAGTCGTCTGTGTGA-1dCA2_MI_GFPve_pos -3.5568277664 -0.4899701752
## dCA2_Mpos_TACAACGTCCTGGTCT-1dCA2_MI_GFPve_pos -1.1690860815 -2.0983462013
## dCA2_Mpos_TACACCCCAGGTTCAT-1dCA2_MI_GFPve_pos -3.8189163037 -1.4708854832
## dCA2_Mpos_TACAGGTCATTCTTCA-1dCA2_MI_GFPve_pos -3.4770803280 -1.5610215344
## dCA2_Mpos_TACATTCCACATGTTG-1dCA2_MI_GFPve_pos -4.9525763817 -1.8595561661
```

```
## dCA2_Mpos_TACCTCGCAGGCGAAT-1dCA2_MI_GFPve_pos -4.5656325646 -1.9453689732
## dCA2_Mpos_TACCTCGGTCCAGTTA-1dCA2_MI_GFPve_pos -3.3270942993 -1.3112804569
## dCA2_Mpos_TACGGTAAGATGCAGC-1dCA2_MI_GFPve_pos -4.4267985649 -1.8909353890
## dCA2_Mpos_TACGGTAAGTGCCGAA-1dCA2_MI_GFPve_pos -4.2311012573 -1.8937487282
## dCA2_Mpos_TACTTACTCTGACCCT-1dCA2_MI_GFPve_pos -3.3761379547 -2.5982207932
## dCA2_Mpos_TAGACTGAGTAGCCAG-1dCA2_MI_GFPve_pos -4.0606143303 -2.4876238980
## dCA2_Mpos_TAGACTGTCTGTAACG-1dCA2_MI_GFPve_pos -2.1450495548 -1.0156414189
## dCA2_Mpos_TAGATCGAGCCGATTT-1dCA2_MI_GFPve_pos -2.6432988472 -0.0359911598
## dCA2_Mpos_TAGCACAGTGAATTAG-1dCA2_MI_GFPve_pos -3.1524891682 -1.3961658158
## dCA2_Mpos_TAGTGCAAGTATAGAC-1dCA2_MI_GFPve_pos -1.5396691389  0.8463149391
## dCA2_Mpos_TATATCCGTTAAAGTG-1dCA2_MI_GFPve_pos -4.3829066582 -0.8148722328
## dCA2_Mpos_TATATCCTCGTCGATA-1dCA2_MI_GFPve_pos -3.2440686054 -0.1194286026
## dCA2_Mpos_TCAAGCAGTTTCGTTT-1dCA2_MI_GFPve_pos -3.7364311047 -1.9485654511
## dCA2_Mpos_TCAAGTGTCAGCCCAG-1dCA2_MI_GFPve_pos -2.6275889702  0.3220243774
## dCA2_Mpos_TCAATTCTCCTCATAT-1dCA2_MI_GFPve_pos -3.6948702164 -1.6920489945
## dCA2_Mpos_TCACATTCATGATGCT-1dCA2_MI_GFPve_pos -3.3615588970 -2.4286760964
## dCA2_Mpos_TCACTATAGGAAGTCC-1dCA2_MI_GFPve_pos -1.9635829754  0.3302107178
## dCA2_Mpos_TCACTATCAAAGGAGA-1dCA2_MI_GFPve_pos -2.4696276016  1.2976312958
## dCA2_Mpos_TCAGGTAAGCCTCAAT-1dCA2_MI_GFPve_pos -1.9662875958 -2.2859794773
## dCA2_Mpos_TCAGGTACACAACGTT-1dCA2_MI_GFPve_pos -4.3743908234 -2.6368791737
## dCA2_Mpos_TCAGGTAGTGCCTGCA-1dCA2_MI_GFPve_pos -4.0534863301 -1.1595546879
## dCA2_Mpos_TCATCATAGCGACTTT-1dCA2_MI_GFPve_pos -1.5344049759 -0.6250509896
## dCA2_Mpos_TCATGGAAGTCCCTAA-1dCA2_MI_GFPve_pos -4.3080150910 -1.7527351059
## dCA2_Mpos_TCCACGTCATCCGAAT-1dCA2_MI_GFPve_pos -1.7987201042 -2.3845193066
## dCA2_Mpos_TCCAGAACAGGCCCTA-1dCA2_MI_GFPve_pos -3.9326534100 -2.5909175553
## dCA2_Mpos_TCCAGAATCCTGGCTT-1dCA2_MI_GFPve_pos -2.7490200825 -1.4560165085
## dCA2_Mpos_TCCATGCAGTCTAGAA-1dCA2_MI_GFPve_pos -3.2118661232 -2.5264298596
## dCA2_Mpos_TCCCATGGTGGCCACT-1dCA2_MI_GFPve_pos -3.8315374680 -1.6343853153
## dCA2_Mpos_TCCGAAACAGAGGTTG-1dCA2_MI_GFPve_pos -4.5191914864 -1.3848533310
## dCA2_Mpos_TCCGAAACATGATCTG-1dCA2_MI_GFPve_pos -2.8188052006 -1.7050702252
## dCA2_Mpos_TCCGATCCATGCCATA-1dCA2_MI_GFPve_pos -3.4515414067 -1.9660679974
## dCA2_Mpos_TCCTAATAGCGTGCCT-1dCA2_MI_GFPve_pos -3.6457459755 -2.5159372963
## dCA2_Mpos_TCCTCTTGTGCCAAGA-1dCA2_MI_GFPve_pos -3.9143040009 -0.8560437836
## dCA2_Mpos_TCGAAGTTCTACTATC-1dCA2_MI_GFPve_pos -3.8492879696 -2.1943642773
## dCA2_Mpos_TCGACGGTCACTCTTA-1dCA2_MI_GFPve_pos -2.4429244824 -1.3026382603
## dCA2_Mpos_TCGGATAAGATTGGGC-1dCA2_MI_GFPve_pos -2.3262793846  0.6855354629
## dCA2_Mpos_TCGGATACACCTTCCA-1dCA2_MI_GFPve_pos -0.5550504155 -2.4082493462
## dCA2_Mpos_TCGGGCAGTTCAATCG-1dCA2_MI_GFPve_pos -4.4090831108 -2.5483734287
## dCA2_Mpos_TCGGGTGCATAGGAGC-1dCA2_MI_GFPve_pos -4.4380428620 -2.4284493603
## dCA2_Mpos_TCGGGTGGTAGGGTAC-1dCA2_MI_GFPve_pos -2.9209720917  0.1394810997
## dCA2_Mpos_TCGTGGGTCGCACGAC-1dCA2_MI_GFPve_pos -2.0984626837  0.9964652382
## dCA2_Mpos_TCTATCAGTGTCCACG-1dCA2_MI_GFPve_pos -2.1357035465  0.5370574318
## dCA2_Mpos_TCTCAGCCATTGAGCT-1dCA2_MI_GFPve_pos -3.8770944901 -1.6403491177
## dCA2_Mpos_TCTGTCGCATAAGCGG-1dCA2_MI_GFPve_pos -4.3097808190 -1.1057305016
## dCA2_Mpos_TCTTAGTGTGCAACAG-1dCA2_MI_GFPve_pos -1.8474689551  0.0780177437
## dCA2_Mpos_TCTTGCGAGAGTGTGC-1dCA2_MI_GFPve_pos -4.4163734741 -1.6335646786
## dCA2_Mpos_TCTTTGATCGTTAGTG-1dCA2_MI_GFPve_pos -2.6393451519  0.2550011001
## dCA2_Mpos_TGAATCGAGCTAAATG-1dCA2_MI_GFPve_pos -3.0464489289 -0.3297142662
## dCA2_Mpos_TGAATCGAGTATGTAG-1dCA2_MI_GFPve_pos -1.6099141665 -1.4555639900
## dCA2_Mpos_TGACTCCAGTCCCTAA-1dCA2_MI_GFPve_pos -3.9042727776 -2.8215540327
## dCA2_Mpos_TGAGACTGTGTTATCG-1dCA2_MI_GFPve_pos -4.0227308102 -2.5151059307
## dCA2_Mpos_TGAGCGCCAAGGCAAC-1dCA2_MI_GFPve_pos -3.9371094532 -0.6891398110
## dCA2_Mpos_TGAGGGAAGGTAGACC-1dCA2_MI_GFPve_pos -3.4770955868 -0.1829037346
## dCA2_Mpos_TGATCAGAGATGGCGT-1dCA2_MI_GFPve_pos -3.6671531029 -2.2779218830
## dCA2_Mpos_TGATCTTGTAAGGAGA-1dCA2_MI_GFPve_pos -2.5286118813  0.0405106865
## dCA2_Mpos_TGATGGTAGTCGGGAT-1dCA2_MI_GFPve_pos -2.6819281407  0.2971663795
## dCA2_Mpos_TGCACGGGTCCATAGT-1dCA2_MI_GFPve_pos -0.9470744558  0.2391395889
## dCA2_Mpos_TGCAGTACAAAGGAGA-1dCA2_MI_GFPve_pos -1.6359747477 -3.1547639050
## dCA2_Mpos_TGCGATAAGGCACCAA-1dCA2_MI_GFPve_pos -3.9978511162 -1.4534847416
## dCA2_Mpos_TGCGGGTAGGTCGAGT-1dCA2_MI_GFPve_pos -3.2324030228 -0.3834833779
## dCA2_Mpos_TGCTTCGAGTTCCATG-1dCA2_MI_GFPve_pos -2.8545944519 -0.4044923462
## dCA2_Mpos_TGGATCATCATCGTAG-1dCA2_MI_GFPve_pos -2.4165997334 -1.0727014221
## dCA2_Mpos_TGGATGTAGTGGAAGA-1dCA2_MI_GFPve_pos -4.7001874275 -1.4693791546
## dCA2_Mpos_TGGCGTGGTGCCTTTC-1dCA2_MI_GFPve_pos -2.6736783810 -1.6960599102
## dCA2_Mpos_TGGGAAGGTCTGCCTT-1dCA2_MI_GFPve_pos -2.5888407059 -1.1665014900
## dCA2_Mpos_TGGGAGAGTGGACAGT-1dCA2_MI_GFPve_pos -2.3865816422  0.5492916427
## dCA2_Mpos_TGGTACAAGAGAGCCT-1dCA2_MI_GFPve_pos -4.5022866554 -2.4732877888
## dCA2_Mpos_TGGTGATCATTGACAC-1dCA2_MI_GFPve_pos -3.1760349102 -1.9379186310
## dCA2_Mpos_TGGTGATGTTGCTTGA-1dCA2_MI_GFPve_pos -2.4240760632 -1.0775582470
## dCA2_Mpos_TGTACAGAGCTGGCTC-1dCA2_MI_GFPve_pos -3.9280247517 -0.9375607647
```

```
## dCA2_Mpos_TGTGATGCAAGACAAT-1dCA2_MI_GFPve_pos -1.2147332258  0.9623313270
## dCA2_Mpos_TGTGATGCACAGAGCA-1dCA2_MI_GFPve_pos -4.5042078324 -1.6036152519
## dCA2_Mpos_TGTGATGGTCGCATCG-1dCA2_MI_GFPve_pos -3.7228674717 -0.9421746411
## dCA2_Mpos_TGTGCGGAGACCATGG-1dCA2_MI_GFPve_pos -2.4576806851 -0.8164529480
## dCA2_Mpos_TGTGCGGTCTCTCAAT-1dCA2_MI_GFPve_pos -2.7734465428  0.6255369507
## dCA2_Mpos_TGTGCGGTCTGCGATA-1dCA2_MI_GFPve_pos -4.2669684715 -1.1840293087
## dCA2_Mpos_TGTTCATAGCGATTCT-1dCA2_MI_GFPve_pos -3.1703665085 -2.8966965355
## dCA2_Mpos_TGTTGAGAGCCTCATA-1dCA2_MI_GFPve_pos -3.9408557244 -1.2932133354
## dCA2_Mpos_TTACCATTCCTTCACG-1dCA2_MI_GFPve_pos -2.0590755768 -0.1639084496
## dCA2_Mpos_TTCCAATTCCAGTACA-1dCA2_MI_GFPve_pos -1.2808607645 -1.1519603409
## dCA2_Mpos_TTCCACGCAAACTAAG-1dCA2_MI_GFPve_pos -2.5098297425  0.2571544967
## dCA2_Mpos_TTCCTTCTCCCATACC-1dCA2_MI_GFPve_pos -2.0049398966  1.6638741813
## dCA2_Mpos_TTCGCTGCAATCCTAG-1dCA2_MI_GFPve_pos -4.6748492546 -1.2188715615
## dCA2_Mpos_TTCGCTGTCCACGGGT-1dCA2_MI_GFPve_pos -3.1339614220 -0.7659802117
## dCA2_Mpos_TTCTGTAAGATGTTCC-1dCA2_MI_GFPve_pos -4.2034203835 -2.1445493378
## dCA2_Mpos_TTGCATTGTTACTCAG-1dCA2_MI_GFPve_pos -3.7037536926 -1.9250068344
## dCA2_Mpos_TTGCGTCTCATTCGGA-1dCA2_MI_GFPve_pos -3.4690012760 -1.2652175106
## dCA2_Mpos_TTGCTGCCAACTTCTT-1dCA2_MI_GFPve_pos -4.1345009632 -2.3809880890
## dCA2_Mpos_TTGCTGCTCAAACCTG-1dCA2_MI_GFPve_pos -1.9065137930 -0.1911515869
## dCA2_Mpos_TTGGGTATCTACAGGT-1dCA2_MI_GFPve_pos -3.7869520016 -0.9742140450
## dCA2_Mpos_TTTACCAAGAGTCACG-1dCA2_MI_GFPve_pos -4.7211801834 -2.1012067475
## dCA2_Mpos_TTTACGTAGGTCACAG-1dCA2_MI_GFPve_pos -3.0979819126 -0.3689040817
## dCA2_Mpos_TTTCAGTCAGACACCC-1dCA2_MI_GFPve_pos -4.5041878052 -1.1475648560
## dCA2_Mpos_TTTGACTAGAGGATCC-1dCA2_MI_GFPve_pos -4.0864796467 -2.0197961010
## dCA2_Mpos_TTTGATCTCACAAGGG-1dCA2_MI_GFPve_pos -2.7683334179  0.5846495949
## dCA2_Mpos_TTTGTTGCAGTTGAAA-1dCA2_MI_GFPve_pos -2.6545026131  0.1593776069
## dCA2_Fpos_AAACGAAGTACCCACG-1dCA2_FI_GFPve_pos -3.5188429184 -1.6263565697
## dCA2_Fpos_AAACGCTGTTATCTTC-1dCA2_FI_GFPve_pos -1.6423381396 -2.1270599045
## dCA2_Fpos_AAAGTGAAGTAAACGT-1dCA2_FI_GFPve_pos -2.1669640370  1.1702714287
## dCA2_Fpos_AACAAGAAGCTTGTTG-1dCA2_FI_GFPve_pos -1.3156508036 -0.5945381798
## dCA2_Fpos_AACACACAGCAAATCA-1dCA2_FI_GFPve_pos -3.3289642163 -1.8375234284
## dCA2_Fpos_AACACACGTCTTGCTC-1dCA2_FI_GFPve_pos -4.6954438515 -2.1698152699
## dCA2_Fpos_AACCAACCATCGTGGC-1dCA2_FI_GFPve_pos -3.4433524437 -1.1192171253
## dCA2_Fpos_AACCACAAGATGCAGC-1dCA2_FI_GFPve_pos -1.8659740515  0.6733689628
## dCA2_Fpos_AACCTGACAGAGGTAC-1dCA2_FI_GFPve_pos -3.1949641533 -1.4444691815
## dCA2_Fpos_AACCTGAGTTTAAGGA-1dCA2_FI_GFPve_pos -4.8736393280 -2.0557582058
## dCA2_Fpos_AAGACAAAGCACCGTC-1dCA2_FI_GFPve_pos -2.7627723046 -0.7814984001
## dCA2_Fpos_AAGACTCAGTATGGAT-1dCA2_FI_GFPve_pos -4.5657365151 -1.9298081078
## dCA2_Fpos_AAGGAATTCACTGAAC-1dCA2_FI_GFPve_pos -4.7564661331 -1.8031396546
## dCA2_Fpos_AAGTACCAGACATCAA-1dCA2_FI_GFPve_pos -2.6020672150 -1.5847029366
## dCA2_Fpos_AAGTACCTCTCATTGT-1dCA2_FI_GFPve_pos -2.8028771706 -1.2720055260
## dCA2_Fpos_AAGTCGTTCTCCTGTG-1dCA2_FI_GFPve_pos -4.7243206330 -2.1647600807
## dCA2_Fpos_AATGACCTCCGTGGCA-1dCA2_FI_GFPve_pos -3.9930260010 -2.5328902878
## dCA2_Fpos_AATGGCTAGAATTCAG-1dCA2_FI_GFPve_pos -2.4789900608 -1.7578909077
## dCA2_Fpos_AATGGCTCACGCTTAA-1dCA2_FI_GFPve_pos -4.1887352295 -2.1769125141
## dCA2_Fpos_AATGGCTCAGGATGAC-1dCA2_FI_GFPve_pos -4.7257764168 -1.6066867985
## dCA2_Fpos_AATGGCTCATGGAAGC-1dCA2_FI_GFPve_pos -4.1667353935 -0.8520455040
## dCA2_Fpos_AATGGCTGTATTGACC-1dCA2_FI_GFPve_pos -1.2172227926  0.0026011787
## dCA2_Fpos_AATGGCTTCGTTACCC-1dCA2_FI_GFPve_pos -4.5976106949 -1.3200885929
## dCA2_Fpos_AATTCCTCACCGTCTT-1dCA2_FI_GFPve_pos -2.1531705685 -2.0832791008
## dCA2_Fpos_AATTCCTCATGACAGG-1dCA2_FI_GFPve_pos -4.7867290802 -2.0979411282
## dCA2_Fpos_ACAAAGACAGTTCCAA-1dCA2_FI_GFPve_pos -4.1231877632 -1.5809933819
## dCA2_Fpos_ACAGCCGGTTCAGTAC-1dCA2_FI_GFPve_pos -3.9437517948 -1.6667039074
## dCA2_Fpos_ACCAACAAGAGTCTGG-1dCA2_FI_GFPve_pos -1.9111383982  0.7803688370
## dCA2_Fpos_ACCACAAAGGTGATCG-1dCA2_FI_GFPve_pos -2.5289442368 -0.8313669838
## dCA2_Fpos_ACCATTTAGCTTTCTT-1dCA2_FI_GFPve_pos -4.0946881600 -2.1489672341
## dCA2_Fpos_ACCCAAAGTTCTCACC-1dCA2_FI_GFPve_pos -1.9105226822  0.3905682884
## dCA2_Fpos_ACGGGTCTCGAAGGAC-1dCA2_FI_GFPve_pos -4.5799376793 -2.1346332707
## dCA2_Fpos_ACGGTTAGTGACACAG-1dCA2_FI_GFPve_pos -4.7059867211 -1.8607053436
## dCA2_Fpos_ACTATTCCAATACAGA-1dCA2_FI_GFPve_pos -4.0106766052 -2.0500967182
## dCA2_Fpos_ACTATTCTCTTCTGTA-1dCA2_FI_GFPve_pos -3.9225649662 -2.5047509350
## dCA2_Fpos_ACTCTCGGTGCGCTCA-1dCA2_FI_GFPve_pos -3.4637203045 -0.5963086762
## dCA2_Fpos_AGACAGGTCACAAGAA-1dCA2_FI_GFPve_pos -4.3916184731 -1.8294186272
## dCA2_Fpos_AGAGCAGGTGTTATCG-1dCA2_FI_GFPve_pos -1.8124725647  1.0460796676
## dCA2_Fpos_AGAGCCCGTCTGCGCA-1dCA2_FI_GFPve_pos -4.8094208069 -1.5840842404
## dCA2_Fpos_AGATAGAGTGGATCAG-1dCA2_FI_GFPve_pos -4.3817226715 -2.5824844517
## dCA2_Fpos_AGCATCAAGTAGGGTC-1dCA2_FI_GFPve_pos -2.7003416844  0.7123465858
## dCA2_Fpos_AGCGATTCACTGGCCA-1dCA2_FI_GFPve_pos -3.6486685105 -1.9450633206
## dCA2_Fpos_AGGAATACACCGCTAG-1dCA2_FI_GFPve_pos -1.7685963936 -0.0257973351
## dCA2_Fpos_AGGAATACACGGTAGA-1dCA2_FI_GFPve_pos -1.4764737911 -0.3891210236
```

```
## dCA2_Fpos_AGGAGGTAGGATACAT-1dCA2_FI_GFPve_pos -4.6378924675 -1.6115898766
## dCA2_Fpos_AGGATCTTCCACGTGG-1dCA2_FI_GFPve_pos -4.1577439137 -2.2846052326
## dCA2_Fpos_AGGCATTAGAGAGCCT-1dCA2_FI_GFPve_pos -4.5944683380 -1.4542023815
## dCA2_Fpos_AGGGCCTGTCTAACGT-1dCA2_FI_GFPve_pos -4.4582950897 -2.3067309536
## dCA2_Fpos_AGGGCCTTCTGAGGCC-1dCA2_FI_GFPve_pos -3.3156835861 -0.2088427223
## dCA2_Fpos_AGGTAGGGTAGTCCTA-1dCA2_FI_GFPve_pos -2.9955591984  0.1591506325
## dCA2_Fpos_AGGTAGGGTTGTAGCT-1dCA2_FI_GFPve_pos -4.0441400833 -1.0285329498
## dCA2_Fpos_AGGTTGTCACTTCCTG-1dCA2_FI_GFPve_pos -3.1903400250 -0.6730241455
## dCA2_Fpos_AGTAACCAGGTGGTTG-1dCA2_FI_GFPve_pos -3.5447687931 -0.2311119713
## dCA2_Fpos_AGTCAACCACCAGTTA-1dCA2_FI_GFPve_pos -1.4760043450  0.5678110443
## dCA2_Fpos_AGTCATGTCTGTCGTC-1dCA2_FI_GFPve_pos -3.9462411232 -2.2337913193
## dCA2_Fpos_AGTGCCGGTGCTGTCG-1dCA2_FI_GFPve_pos -3.7479064293 -0.5664171852
## dCA2_Fpos_ATACTTCCACCTATCC-1dCA2_FI_GFPve_pos -4.1244361229 -2.3356423058
## dCA2_Fpos_ATATCCTCATACACCA-1dCA2_FI_GFPve_pos -4.4792611428 -1.2082810082
## dCA2_Fpos_ATCACTTGTCTAACGT-1dCA2_FI_GFPve_pos -3.9140438862 -2.1991629280
## dCA2_Fpos_ATCAGGTCATACTGAC-1dCA2_FI_GFPve_pos -4.5515520401 -2.1027071156
## dCA2_Fpos_ATCCACCGTACAAAGT-1dCA2_FI_GFPve_pos -3.9256026573 -1.0307559647
## dCA2_Fpos_ATCGCCTAGGTCATAA-1dCA2_FI_GFPve_pos -2.3849468060 -0.7528142609
## dCA2_Fpos_ATCGCCTGTACTTGTG-1dCA2_FI_GFPve_pos -3.8659076519 -2.1116351761
## dCA2_Fpos_ATCGTCCCACCGAATT-1dCA2_FI_GFPve_pos -3.5098986454 -1.9853303112
## dCA2_Fpos_ATCGTCCTCTTGGTGA-1dCA2_FI_GFPve_pos -3.6707594223 -0.2965869583
## dCA2_Fpos_ATCGTGATCATTCCTA-1dCA2_FI_GFPve_pos -2.3587519951  0.6588797890
## dCA2_Fpos_ATCTCTAAGAGGCTGT-1dCA2_FI_GFPve_pos -2.2593219109  0.7031741463
## dCA2_Fpos_ATGATCGGTGGAGAAA-1dCA2_FI_GFPve_pos -2.6202027626  0.5299911819
## dCA2_Fpos_ATGGGTTCAGGGACTA-1dCA2_FI_GFPve_pos -4.4233982392 -1.1008917965
## dCA2_Fpos_ATTACTCTCCTACCAC-1dCA2_FI_GFPve_pos -2.8568906613  0.2532348953
## dCA2_Fpos_ATTCCTAGTAAGCAAT-1dCA2_FI_GFPve_pos -3.3496825524 -0.9087700524
## dCA2_Fpos_ATTCGTTTCGTACACA-1dCA2_FI_GFPve_pos -2.3044738598  0.3896961532
## dCA2_Fpos_ATTCTACAGTCGTCTA-1dCA2_FI_GFPve_pos -1.9753590651  0.2940631233
## dCA2_Fpos_ATTGTTCCATCCAACA-1dCA2_FI_GFPve_pos -0.8646033831 -0.9852878727
## dCA2_Fpos_ATTTACCGTGTTCATG-1dCA2_FI_GFPve_pos -1.7285063095  0.1291523300
## dCA2_Fpos_ATTTCACAGCATCCTA-1dCA2_FI_GFPve_pos -1.6689058371 -0.4523400940
## dCA2_Fpos_CAAGCTATCGTTATCT-1dCA2_FI_GFPve_pos -3.2025012799 -0.5154647507
## dCA2_Fpos_CACAACAAGCACTGGA-1dCA2_FI_GFPve_pos -3.6811113186 -2.1338677086
## dCA2_Fpos_CACAACAGTAGCTGAG-1dCA2_FI_GFPve_pos -3.7163064308 -1.6752202191
## dCA2_Fpos_CACACAAGTGAGTTTC-1dCA2_FI_GFPve_pos -4.2234055824 -1.8753711857
## dCA2_Fpos_CACTGGGTCTGGGCGT-1dCA2_FI_GFPve_pos -1.8636668749  0.0060053192
## dCA2_Fpos_CAGCGTGAGCGCCATC-1dCA2_FI_GFPve_pos -4.0000202484 -1.7355394043
## dCA2_Fpos_CATAGACTCACCATAG-1dCA2_FI_GFPve_pos -0.7703748770 -2.3798534550
## dCA2_Fpos_CATCCACCACCCTATC-1dCA2_FI_GFPve_pos -2.3074698277 -0.8288311638
## dCA2_Fpos_CATCCGTAGCCTGTGC-1dCA2_FI_GFPve_pos -4.6941010781 -2.0475024857
## dCA2_Fpos_CATGCGGTCACTCTTA-1dCA2_FI_GFPve_pos -4.4753377266 -1.3931190647
## dCA2_Fpos_CATGCGGTCATTGGTG-1dCA2_FI_GFPve_pos -3.9524042435 -1.5702128090
## dCA2_Fpos_CATTCATCAAGAATAC-1dCA2_FI_GFPve_pos -3.7443251438 -2.0240194477
## dCA2_Fpos_CATTCATTCATTACGG-1dCA2_FI_GFPve_pos -4.3734662361 -2.1958374657
## dCA2_Fpos_CATTCTAGTTTCACAG-1dCA2_FI_GFPve_pos -2.3878381081 -1.8718483128
## dCA2_Fpos_CATTGAGCAGATTCGT-1dCA2_FI_GFPve_pos -3.4875757523 -0.5228695549
## dCA2_Fpos_CATTGTTGTGTGTGGA-1dCA2_FI_GFPve_pos -4.6625511475 -2.1001803555
## dCA2_Fpos_CCAAGCGCATCGAAGG-1dCA2_FI_GFPve_pos -3.5161213703 -0.5540914215
## dCA2_Fpos_CCACCATTCTGCAGCG-1dCA2_FI_GFPve_pos -4.2902877159 -1.1195432820
## dCA2_Fpos_CCCAACTCACCTCTGT-1dCA2_FI_GFPve_pos -1.9671077557  0.4296670280
## dCA2_Fpos_CCCATTGCAATTGTGC-1dCA2_FI_GFPve_pos -1.3842924662  0.1967244468
## dCA2_Fpos_CCCTTAGCACTGGACC-1dCA2_FI_GFPve_pos -1.5047741957  0.6723313652
## dCA2_Fpos_CCTAACCAGAGTGAAG-1dCA2_FI_GFPve_pos -2.2252962418  0.7734308563
## dCA2_Fpos_CCTACGTAGTGCACTT-1dCA2_FI_GFPve_pos -1.9879125185  0.8688278518
## dCA2_Fpos_CCTCAGTAGATAGCAT-1dCA2_FI_GFPve_pos -1.9794238634  1.0986576401
## dCA2_Fpos_CCTCAGTAGATGAATC-1dCA2_FI_GFPve_pos -3.8540162869 -2.6704625763
## dCA2_Fpos_CCTCTCCAGCCTCAGC-1dCA2_FI_GFPve_pos -1.5174115725 -0.6535787262
## dCA2_Fpos_CCTGCATAGTGTTGTC-1dCA2_FI_GFPve_pos -3.9787051506 -1.1581828274
## dCA2_Fpos_CCTGTTGGTCAGGCAA-1dCA2_FI_GFPve_pos -3.2721206970 -0.3747934975
## dCA2_Fpos_CCTTCAGAGTACAGCG-1dCA2_FI_GFPve_pos -4.3916833229 -1.1775505223
## dCA2_Fpos_CGAGAAGCAAGCGCAA-1dCA2_FI_GFPve_pos -3.4632544346 -0.6787728943
## dCA2_Fpos_CGCCAGAAGGCATCTT-1dCA2_FI_GFPve_pos -3.9246082134 -0.9869200863
## dCA2_Fpos_CGGGACTCATTAGGCT-1dCA2_FI_GFPve_pos -3.9737329312 -1.2786330857
## dCA2_Fpos_CGGGTCAGTGTCTTAG-1dCA2_FI_GFPve_pos -4.2495744057 -0.9737996735
## dCA2_Fpos_CGTAAGTGTGTTACTG-1dCA2_FI_GFPve_pos -4.2387311287 -1.8629176296
## dCA2_Fpos_CGTAATGCAATTCACG-1dCA2_FI_GFPve_pos -1.5869824715 -0.7118320145
## dCA2_Fpos_CTAAGTGAGACGCCAA-1dCA2_FI_GFPve_pos -1.6320469208 -0.5685519852
## dCA2_Fpos_CTACCCACAAAGACGC-1dCA2_FI_GFPve_pos -2.7425594158  0.2162165962
## dCA2_Fpos_CTACCCACATGGGATG-1dCA2_FI_GFPve_pos -3.4297582932 -0.1763801254
```

```
## dCA2_Fpos_CTACCTGTCACTGTTT-1dCA2_FI_GFPve_pos -4.7239253350 -1.6869120278
## dCA2_Fpos_CTACTATGTATCGCGC-1dCA2_FI_GFPve_pos -4.9428331680 -1.5760547795
## dCA2_Fpos_CTACTATTCGTTCCCA-1dCA2_FI_GFPve_pos -3.9582059212 -2.5236308254
## dCA2_Fpos_CTATAGGGTACGCGTC-1dCA2_FI_GFPve_pos -4.0617210694 -1.5889959015
## dCA2_Fpos_CTCAGAAGTGACCGAA-1dCA2_FI_GFPve_pos -4.0918254681 -2.0115010418
## dCA2_Fpos_CTCAGGGAGCTAAATG-1dCA2_FI_GFPve_pos -2.2197513409 -0.6501154579
## dCA2_Fpos_CTCATGCCAGAATTCC-1dCA2_FI_GFPve_pos -2.7620234318  0.0942125641
## dCA2_Fpos_CTCATTAGTCTACTGA-1dCA2_FI_GFPve_pos -0.8423853941 -2.8160354771
## dCA2_Fpos_CTCCATGTCATAGAGA-1dCA2_FI_GFPve_pos -2.2358341046  0.1031976066
## dCA2_Fpos_CTCCCTCTCTGTTGGA-1dCA2_FI_GFPve_pos -4.4027130432 -1.4867114701
## dCA2_Fpos_CTCCGATTCGCGAAGA-1dCA2_FI_GFPve_pos -4.1146919556 -0.7699904122
## dCA2_Fpos_CTCCTCCCAAGACAAT-1dCA2_FI_GFPve_pos -0.7244371004 -2.2053236641
## dCA2_Fpos_CTCCTCCGTTAGGACG-1dCA2_FI_GFPve_pos -3.5405003853 -1.2229940571
## dCA2_Fpos_CTCGAGGCACAACGAG-1dCA2_FI_GFPve_pos -3.9025301762 -2.2436611332
## dCA2_Fpos_CTCGAGGGTCTCAAGT-1dCA2_FI_GFPve_pos -3.3963103123 -1.5349809803
## dCA2_Fpos_CTGAATGAGGTAAAGG-1dCA2_FI_GFPve_pos -1.9717851706 -0.3524303116
## dCA2_Fpos_CTGAATGCAACCTATG-1dCA2_FI_GFPve_pos -2.8534262009 -1.3179807343
## dCA2_Fpos_CTGATCCTCTTCTCAA-1dCA2_FI_GFPve_pos -4.6321775742 -1.9776000656
## dCA2_Fpos_CTGCCATAGGGCAATC-1dCA2_FI_GFPve_pos -3.9461564846 -2.7930496611
## dCA2_Fpos_CTGCCATAGTACCCTA-1dCA2_FI_GFPve_pos -4.3140094109 -2.3326153435
## dCA2_Fpos_CTGCCATGTTGTAAAG-1dCA2_FI_GFPve_pos -3.7573735542 -2.3915450253
## dCA2_Fpos_CTGCGAGCAGGTCAGA-1dCA2_FI_GFPve_pos -2.2288250752  0.0228443466
## dCA2_Fpos_CTGGACGGTCAAGCCC-1dCA2_FI_GFPve_pos -1.7012481518 -0.5267691292
## dCA2_Fpos_CTGGCAGCAGCTCTGG-1dCA2_FI_GFPve_pos -2.1049123831 -0.0043754257
## dCA2_Fpos_CTGGCAGTCAAGCCGC-1dCA2_FI_GFPve_pos -4.5611769982 -2.3134326615
## dCA2_Fpos_CTGGTCTAGAGGTGCT-1dCA2_FI_GFPve_pos -2.6613802738  0.0784354530
## dCA2_Fpos_CTGTCGTGTACAACGG-1dCA2_FI_GFPve_pos -4.0303814240 -0.6777105011
## dCA2_Fpos_CTGTGAAAGCGGATCA-1dCA2_FI_GFPve_pos -2.3741609879  0.4221830688
## dCA2_Fpos_CTTCAATAGCGCCCAT-1dCA2_FI_GFPve_pos -0.9392187424 -1.7314376511
## dCA2_Fpos_CTTCGGTGTCGTGTTA-1dCA2_FI_GFPve_pos -3.1525988407 -0.2622327484
## dCA2_Fpos_GAAGGGTAGAAGCTCG-1dCA2_FI_GFPve_pos -4.3499753304 -2.3493232407
## dCA2_Fpos_GAATAGACAGGTCAGA-1dCA2_FI_GFPve_pos -3.3511771984 -0.1448616661
## dCA2_Fpos_GACATCAGTAGAATAC-1dCA2_FI_GFPve_pos -3.1796016522 -0.1933974899
## dCA2_Fpos_GACCCTTAGCTCCGAC-1dCA2_FI_GFPve_pos -1.2019697256  0.0399313293
## dCA2_Fpos_GAGAGGTCAGGTGACA-1dCA2_FI_GFPve_pos -4.0996880360 -1.6854295410
## dCA2_Fpos_GAGGCCTCAGATTTCG-1dCA2_FI_GFPve_pos -2.2011012860  1.1807098709
## dCA2_Fpos_GAGTCATTCCAAGGGA-1dCA2_FI_GFPve_pos -4.5640089340 -2.4049074330
## dCA2_Fpos_GAGTGAGAGATTGAGT-1dCA2_FI_GFPve_pos -1.2215096779  1.6742973648
## dCA2_Fpos_GAGTTTGGTATTCCTT-1dCA2_FI_GFPve_pos -1.7515562601 -0.6186976113
## dCA2_Fpos_GATCACAGTAGCCCTG-1dCA2_FI_GFPve_pos -3.0852899380 -0.3666467346
## dCA2_Fpos_GATTCGACAGTTTCGA-1dCA2_FI_GFPve_pos -3.9559686012 -1.4545831360
## dCA2_Fpos_GCATCTCTCACCTCTG-1dCA2_FI_GFPve_pos -4.1636986561 -1.9699580349
## dCA2_Fpos_GCATGATGTGAACGGT-1dCA2_FI_GFPve_pos -2.3011023827  0.8663368545
## dCA2_Fpos_GCCAACGAGCTGCCTG-1dCA2_FI_GFPve_pos -1.4031541176 -1.0571543850
## dCA2_Fpos_GCCAGGTAGATGACAT-1dCA2_FI_GFPve_pos -2.2240364380  0.0542507492
## dCA2_Fpos_GCCTGTTCACATATGC-1dCA2_FI_GFPve_pos -3.1757201977 -1.2721323647
## dCA2_Fpos_GCGTGCATCGTTAGTG-1dCA2_FI_GFPve_pos -2.1859555073  0.7479801498
## dCA2_Fpos_GGAGATGCACTTGTCC-1dCA2_FI_GFPve_pos -4.4195535011 -1.8565442242
## dCA2_Fpos_GGAGATGTCCCAGTGG-1dCA2_FI_GFPve_pos -4.6737158127 -2.0146853604
## dCA2_Fpos_GGCTGTGCACTAGGTT-1dCA2_FI_GFPve_pos -2.7531821556 -1.1625961937
## dCA2_Fpos_GGCTGTGCAGAACTTC-1dCA2_FI_GFPve_pos -2.1204581089  0.8942904793
## dCA2_Fpos_GGGCGTTGTGGTACAG-1dCA2_FI_GFPve_pos -2.5205848046  0.5795064293
## dCA2_Fpos_GGGTCACGTACTAGCT-1dCA2_FI_GFPve_pos -1.8315189905  0.7300262771
## dCA2_Fpos_GGGTCTGCAGAGTGTG-1dCA2_FI_GFPve_pos -3.2062635250 -0.3539824166
## dCA2_Fpos_GGTGTTAAGGGTTAGC-1dCA2_FI_GFPve_pos -4.2856490441 -1.2074317612
## dCA2_Fpos_GTAACACGTTCCATTT-1dCA2_FI_GFPve_pos -3.9382335968 -1.5060112156
## dCA2_Fpos_GTAAGTCAGGTGCCAA-1dCA2_FI_GFPve_pos -4.9241478272 -1.9343394913
## dCA2_Fpos_GTAATCGCAAGGTTGG-1dCA2_FI_GFPve_pos -4.0928783245 -1.9917835869
## dCA2_Fpos_GTACAACAGATGTAGT-1dCA2_FI_GFPve_pos -2.1421218939 -0.0926456131
## dCA2_Fpos_GTATTGGGTCCGGTCA-1dCA2_FI_GFPve_pos -4.5033457108 -1.6640250363
## dCA2_Fpos_GTATTTCGTAGCGCCT-1dCA2_FI_GFPve_pos -1.2628735132  1.3434272133
## dCA2_Fpos_GTATTTCGTTACAGCT-1dCA2_FI_GFPve_pos -3.6157762833 -1.4050728955
## dCA2_Fpos_GTCACGGAGGTGAGAA-1dCA2_FI_GFPve_pos -4.4174754448 -1.5361325421
## dCA2_Fpos_GTCATTTAGGCCCGTT-1dCA2_FI_GFPve_pos -1.5039934941 -3.1881967463
## dCA2_Fpos_GTCCTCATCTGAGTCA-1dCA2_FI_GFPve_pos -4.7226583786 -2.0367726959
## dCA2_Fpos_GTCTACCCACTCCACT-1dCA2_FI_GFPve_pos -1.6378391810 -1.7815963902
## dCA2_Fpos_GTCTTTAAGCGTGAGT-1dCA2_FI_GFPve_pos -2.4811942406  0.0086608253
## dCA2_Fpos_GTGAGTTCAAAGGGTC-1dCA2_FI_GFPve_pos -4.7011039086 -1.8602227844
## dCA2_Fpos_GTGATGTTCGACCTAA-1dCA2_FI_GFPve_pos -4.4583513565 -2.1693682350
## dCA2_Fpos_GTGCTTCAGACTTCGT-1dCA2_FI_GFPve_pos -1.7584438153  1.3524155937
```

```
## dCA2_Fpos_GTGGAGATCTTAATCC-1dCA2_FI_GFPve_pos -4.4478089638 -2.0462975182
## dCA2_Fpos_GTGGTTACACTTGGGC-1dCA2_FI_GFPve_pos -4.8258116074 -1.9485511460
## dCA2_Fpos_GTGTAACAGAGAGTGA-1dCA2_FI_GFPve_pos -1.8859263487  0.2091069542
## dCA2_Fpos_GTTACAGTCCAAATGC-1dCA2_FI_GFPve_pos -3.9575688667 -1.7303292431
## dCA2_Fpos_GTTGCTCCACGCGTCA-1dCA2_FI_GFPve_pos -4.4284951515 -1.4702319779
## dCA2_Fpos_GTTGCTCTCCTAGCTC-1dCA2_FI_GFPve_pos -1.0538002677  0.9365320526
## dCA2_Fpos_GTTGTAGCACAGACGA-1dCA2_FI_GFPve_pos -3.8879911728 -2.3015749134
## dCA2_Fpos_GTTGTGATCATTCTTG-1dCA2_FI_GFPve_pos -3.1365876026 -1.7373745121
## dCA2_Fpos_GTTTACTCAGCTGTTA-1dCA2_FI_GFPve_pos -4.0790653057 -1.5655524410
## dCA2_Fpos_GTTTGGAGTGCCTTCT-1dCA2_FI_GFPve_pos -3.9197819061 -1.1667737641
## dCA2_Fpos_TAACGACAGGTACTGG-1dCA2_FI_GFPve_pos -3.1474890537 -2.4813010372
## dCA2_Fpos_TAATTCCTCGGCCAAC-1dCA2_FI_GFPve_pos -3.4857513733 -0.5508474983
## dCA2_Fpos_TACACCCAGGATATGT-1dCA2_FI_GFPve_pos -4.9706151314 -2.0279988922
## dCA2_Fpos_TACACCCTCCAACCGG-1dCA2_FI_GFPve_pos -4.2780411072 -1.8826598801
## dCA2_Fpos_TACCCACGTTCGATTG-1dCA2_FI_GFPve_pos -4.5058624573 -2.1657437958
## dCA2_Fpos_TACCCGTGTCTGCGCA-1dCA2_FI_GFPve_pos -4.4974353142 -1.0884492077
## dCA2_Fpos_TACCGGGTCACTCTTA-1dCA2_FI_GFPve_pos -2.6491091080 -0.2038392700
## dCA2_Fpos_TACCTCGCACAACGAG-1dCA2_FI_GFPve_pos -3.8449134655 -2.3880035557
## dCA2_Fpos_TACGGGCCAGACCGCT-1dCA2_FI_GFPve_pos -4.0792212315 -1.3160147347
## dCA2_Fpos_TACGTCCTCCAGTTCC-1dCA2_FI_GFPve_pos -3.4946861096 -2.6725585140
## dCA2_Fpos_TATTGCTGTACATTGC-1dCA2_FI_GFPve_pos -3.8225052185 -1.2756542839
## dCA2_Fpos_TCAAGACGTTCGGGTC-1dCA2_FI_GFPve_pos -4.5844080277 -2.2068516888
## dCA2_Fpos_TCACACCTCCATACAG-1dCA2_FI_GFPve_pos -3.4648547001 -2.7118320145
## dCA2_Fpos_TCACGCTGTATCTTCT-1dCA2_FI_GFPve_pos -1.8460342713 -0.2174558319
## dCA2_Fpos_TCACGGGAGGGTTGCA-1dCA2_FI_GFPve_pos -1.7509233780  1.3127189003
## dCA2_Fpos_TCAGCCTCACGACGAA-1dCA2_FI_GFPve_pos -4.6788432427 -2.1741661705
## dCA2_Fpos_TCAGCCTCATACAGCT-1dCA2_FI_GFPve_pos -3.7559306450 -2.6571852841
## dCA2_Fpos_TCAGTTTTCACACCCT-1dCA2_FI_GFPve_pos -4.1122309990 -2.6802723088
## dCA2_Fpos_TCATACTCAAGGCAAC-1dCA2_FI_GFPve_pos -2.4357678719 -0.3822793640
## dCA2_Fpos_TCATATCCACACTTAG-1dCA2_FI_GFPve_pos -4.4555928536 -2.1594390549
## dCA2_Fpos_TCATGAGGTGTCATTG-1dCA2_FI_GFPve_pos -4.1771237679 -1.0255350746
## dCA2_Fpos_TCATGAGGTTGCGAAG-1dCA2_FI_GFPve_pos -2.2107872791  0.9107179962
## dCA2_Fpos_TCATGGATCCCGAGTG-1dCA2_FI_GFPve_pos -4.0880300827 -1.7942790665
## dCA2_Fpos_TCATTGTCAATAACGA-1dCA2_FI_GFPve_pos -2.1377224751  0.6657820068
## dCA2_Fpos_TCCCATGAGGGCCTCT-1dCA2_FI_GFPve_pos -2.4894988365 -0.9950027146
## dCA2_Fpos_TCGACCTCACTGGACC-1dCA2_FI_GFPve_pos -3.0532088108 -1.7301959671
## dCA2_Fpos_TCGACCTCAGAATCGG-1dCA2_FI_GFPve_pos -4.4988033600 -1.1744875588
## dCA2_Fpos_TCGACCTTCCGTCACT-1dCA2_FI_GFPve_pos -3.6323451824 -0.4445295014
## dCA2_Fpos_TCGATTTAGAGGACTC-1dCA2_FI_GFPve_pos -4.6635267563 -1.9513487496
## dCA2_Fpos_TCGATTTCAGTCGCAC-1dCA2_FI_GFPve_pos -4.1741573639 -1.2859429993
## dCA2_Fpos_TCGCAGGAGTAGGATT-1dCA2_FI_GFPve_pos -3.1007997818 -0.3451113381
## dCA2_Fpos_TCGTCCAGTCGAACGA-1dCA2_FI_GFPve_pos -2.5187020130 -0.7997312226
## dCA2_Fpos_TCTCACGCATAGGTAA-1dCA2_FI_GFPve_pos -1.4970263071 -2.2337758221
## dCA2_Fpos_TCTGCCATCAGGAGAC-1dCA2_FI_GFPve_pos -2.6080941982  0.1720533691
## dCA2_Fpos_TCTGGCTAGTAGTGCG-1dCA2_FI_GFPve_pos -1.9330873318  0.8416138016
## dCA2_Fpos_TGAACGTGTTGCGTAT-1dCA2_FI_GFPve_pos -4.6350376435 -2.1060523667
## dCA2_Fpos_TGAATCGAGAAGCTGC-1dCA2_FI_GFPve_pos -4.6251823731 -1.5344130673
## dCA2_Fpos_TGAATCGGTCAAAGTA-1dCA2_FI_GFPve_pos -4.5752680130 -1.6195575871
## dCA2_Fpos_TGACAGTTCATGTCTT-1dCA2_FI_GFPve_pos -3.8910856075 -1.7212433495
## dCA2_Fpos_TGACGCGAGGTTTACC-1dCA2_FI_GFPve_pos -3.6799197025 -1.8143064656
## dCA2_Fpos_TGACGCGGTGTTACAC-1dCA2_FI_GFPve_pos -2.4167666264  0.3288536392
## dCA2_Fpos_TGAGCGCTCTCGTGGG-1dCA2_FI_GFPve_pos -3.0855870075 -0.2001256623
## dCA2_Fpos_TGATGGTGTGGTTCTA-1dCA2_FI_GFPve_pos -4.2258236237 -1.8834526219
## dCA2_Fpos_TGCACGGGTCTTACAG-1dCA2_FI_GFPve_pos -3.8665027447 -2.0694157757
## dCA2_Fpos_TGCAGATTCCCTCAAC-1dCA2_FI_GFPve_pos -3.8575184174 -2.2877463974
## dCA2_Fpos_TGCCGAGGTCTGTCCT-1dCA2_FI_GFPve_pos -1.8130816050  0.5542192779
## dCA2_Fpos_TGCGATAAGGGTACAC-1dCA2_FI_GFPve_pos -2.9348826237 -0.4982018151
## dCA2_Fpos_TGCTCGTGTCAGACGA-1dCA2_FI_GFPve_pos -2.7312619515  0.1216097198
## dCA2_Fpos_TGCTCGTGTGGCATCC-1dCA2_FI_GFPve_pos -2.5822732277  0.6911306701
## dCA2_Fpos_TGGATGTTCGATACTG-1dCA2_FI_GFPve_pos -4.3328506775 -1.4144549050
## dCA2_Fpos_TGGGAAGAGTATAGAC-1dCA2_FI_GFPve_pos -4.3034999199 -1.5629858650
## dCA2_Fpos_TGGGTTATCGAGAAAT-1dCA2_FI_GFPve_pos -4.8997776337 -1.7303993382
## dCA2_Fpos_TGGTTAGCATCCCACT-1dCA2_FI_GFPve_pos -4.4430963822 -2.0815560497
## dCA2_Fpos_TGTAGACCAGGAACCA-1dCA2_FI_GFPve_pos -4.5468666382 -1.8370706715
## dCA2_Fpos_TGTCCACAGACTAAGT-1dCA2_FI_GFPve_pos -4.2561552353 -1.2864529766
## dCA2_Fpos_TGTCCACCAAGTATAG-1dCA2_FI_GFPve_pos -4.0498335190 -1.3619854130
## dCA2_Fpos_TGTCCTGGTCCACACG-1dCA2_FI_GFPve_pos -3.1910652943 -0.4973797478
## dCA2_Fpos_TGTTGAGTCATTACCT-1dCA2_FI_GFPve_pos -4.8121163674 -1.7333852925
## dCA2_Fpos_TTACCATCACGCTTAA-1dCA2_FI_GFPve_pos -1.2367471285  0.2847414337
## dCA2_Fpos_TTACTGTAGCGGTAGT-1dCA2_FI_GFPve_pos -2.5313625164  0.0522895180
```

```
## dCA2_Fpos_TTCAGGACACATGTTG-1dCA2_FI_GFPve_pos -1.4153057165 -0.9242789425
## dCA2_Fpos_TTCCACGCAGGTGACA-1dCA2_FI_GFPve_pos -3.5088383980 -1.4737329163
## dCA2_Fpos_TTCCGGTCAGACCATT-1dCA2_FI_GFPve_pos -1.7367520161 -0.3982796349
## dCA2_Fpos_TTCCTTCGTCACATTG-1dCA2_FI_GFPve_pos -0.7493131585  0.5796466194
## dCA2_Fpos_TTCTGTACAGACCCGT-1dCA2_FI_GFPve_pos -3.7110211678 -1.1743409313
## dCA2_Fpos_TTGAGTGCATCCTCAC-1dCA2_FI_GFPve_pos -4.4812986679 -1.8703281559
## dCA2_Fpos_TTGCGTCTCTCTAAGG-1dCA2_FI_GFPve_pos -1.9963418074 -0.0239295639
## dCA2_Fpos_TTGGGATAGCTGACTT-1dCA2_FI_GFPve_pos -4.7768966980 -1.3571030773
## dCA2_Fpos_TTGGGATTCTCACGAA-1dCA2_FI_GFPve_pos -3.5657706089 -0.5710491814
## dCA2_Fpos_TTGGGCGGTTAGAGAT-1dCA2_FI_GFPve_pos -2.8014902897  0.0126509987
## dCA2_Fpos_TTTAGTCAGGTACAAT-1dCA2_FI_GFPve_pos -4.0829694100 -1.7926606812
## dCA2_Fpos_TTTCCTCAGCGTGAAC-1dCA2_FI_GFPve_pos -1.3473485537  1.3139510475
## dCA2_Fpos_TTTGATCCATGGAATA-1dCA2_FI_GFPve_pos -2.8233301468 -0.3492512383
## dCA2_Fpos_TTTGGAGAGTAGAATC-1dCA2_FI_GFPve_pos -3.7878544159 -1.2302422203
## FC_Mpos_AAGACTCTCGACATCA-1FC_MI_GFPve_pos     -2.7244748898 -0.8936502613
## FC_Mpos_AAGCGAGAGCGTCAAG-1FC_MI_GFPve_pos     -1.2946382590 -2.3565048851
## FC_Mpos_ACATGCATCCAATCTT-1FC_MI_GFPve_pos     -3.5940222569 -0.4036202110
## FC_Mpos_ACCCTCATCAAGCCAT-1FC_MI_GFPve_pos     -3.5159292050 -2.5200113930
## FC_Mpos_ACCGTTCCACGGTGCT-1FC_MI_GFPve_pos     -2.3521182366 -4.1837972678
## FC_Mpos_ACCGTTCGTGCCCAGT-1FC_MI_GFPve_pos     -3.8482544251 -1.8034331478
## FC_Mpos_ACGGTTATCGATACAC-1FC_MI_GFPve_pos     -3.9980611630 -2.6563043274
## FC_Mpos_AGACTCAGTATGAGCG-1FC_MI_GFPve_pos     -4.4817774124 -2.0168642678
## FC_Mpos_AGCCAATGTTATAGAG-1FC_MI_GFPve_pos     -2.8846890755 -1.1054103054
## FC_Mpos_AGCGATTAGATGTTCC-1FC_MI_GFPve_pos     -0.0527463011  1.6494512878
## FC_Mpos_AGCGTCGCACATATCG-1FC_MI_GFPve_pos     -3.6298379727 -1.6361639179
## FC_Mpos_AGGACTTTCGTGTTCC-1FC_MI_GFPve_pos     -4.5316750832 -1.6091091313
## FC_Mpos_AGGCATTCACCTTCCA-1FC_MI_GFPve_pos     -4.5974352188 -2.0781254448
## FC_Mpos_AGGTTACGTAACACCT-1FC_MI_GFPve_pos     -2.9323484726 -1.6487750687
## FC_Mpos_ATAGAGACAGAGGGTT-1FC_MI_GFPve_pos     -4.6045481987 -1.1381272949
## FC_Mpos_ATCGTGAAGCATGCAG-1FC_MI_GFPve_pos     -2.1478416748 -3.8385424294
## FC_Mpos_ATGGAGGCACTATGTG-1FC_MI_GFPve_pos     -3.4881317444 -1.1822971977
## FC_Mpos_ATTATCCAGACCTCAT-1FC_MI_GFPve_pos     -1.9905683823  0.1845789276
## FC_Mpos_ATTCATCAGTGAGTTA-1FC_MI_GFPve_pos     -3.9908125229 -2.2427832760
## FC_Mpos_ATTCCATTCGTTCTCG-1FC_MI_GFPve_pos     -2.7287688084 -1.1800994553
## FC_Mpos_ATTCTTGCAAAGCTCT-1FC_MI_GFPve_pos     -3.2880928345 -1.5774123348
## FC_Mpos_CACATGAAGTTACGTC-1FC_MI_GFPve_pos     -4.7326028176 -1.2581824936
## FC_Mpos_CACGAATGTGTGCCTG-1FC_MI_GFPve_pos     -4.2799579926 -1.4283709206
## FC_Mpos_CACGGGTGTTCCGCTT-1FC_MI_GFPve_pos     -3.8925704785 -2.1380784191
## FC_Mpos_CACTGGGCACATAGCT-1FC_MI_GFPve_pos     -3.6223170586 -2.7588208594
## FC_Mpos_CATGCGGAGTAAGAGG-1FC_MI_GFPve_pos     -4.0586926766 -1.9531292595
## FC_Mpos_CCGATGGTCCCATGGG-1FC_MI_GFPve_pos     -4.3953702278 -1.0613181271
## FC_Mpos_CCTCTCCTCCCGGTAG-1FC_MI_GFPve_pos     -3.2060127087 -0.8920676388
## FC_Mpos_CGAGGAAGTTCCAGGC-1FC_MI_GFPve_pos     -4.7897312470 -1.7771806397
## FC_Mpos_CGATGGCAGGTCGCCT-1FC_MI_GFPve_pos     -3.4227475949 -0.6685280480
## FC_Mpos_CTCAGGGGTCGAATTC-1FC_MI_GFPve_pos     -3.9704360790 -0.8487646260
## FC_Mpos_GAACGTTTCTATTTCG-1FC_MI_GFPve_pos     -2.6587700672  0.3168454490
## FC_Mpos_GACCCAGCATAGACTC-1FC_MI_GFPve_pos     -4.5927474327 -2.2826244511
## FC_Mpos_GAGCCTGTCCAGGACC-1FC_MI_GFPve_pos     -4.6780364342 -1.1726681866
## FC_Mpos_GAGGGTAAGAAAGTCT-1FC_MI_GFPve_pos     -4.6602589913 -1.9518532433
## FC_Mpos_GAGTCATGTCCTGAAT-1FC_MI_GFPve_pos     -3.1018225975 -0.1832689919
## FC_Mpos_GATCCCTAGCGAAACC-1FC_MI_GFPve_pos     -4.2149903603 -1.3308894314
## FC_Mpos_GCAACATGTGAAGCGT-1FC_MI_GFPve_pos     -2.9873778649 -1.9620754399
## FC_Mpos_GCCATGGCAAACGTGG-1FC_MI_GFPve_pos     -3.0759846993 -1.1251806893
## FC_Mpos_GCGATCGAGATTGATG-1FC_MI_GFPve_pos     -2.3004464932 -0.0220193543
## FC_Mpos_GCTTCACCACGACGAA-1FC_MI_GFPve_pos     -2.9229941197 -1.1855966725
## FC_Mpos_GGAACCCAGTCATGCT-1FC_MI_GFPve_pos     -2.0964025088 -2.0002834477
## FC_Mpos_GGCACGTCAAGATCCT-1FC_MI_GFPve_pos     -3.8333768673 -2.0840315499
## FC_Mpos_GGGATCCTCTGAGTCA-1FC_MI_GFPve_pos     -3.2036247082 -2.3774869122
## FC_Mpos_GGGCTCAAGATCACTC-1FC_MI_GFPve_pos     -4.5932371445 -2.1452579178
## FC_Mpos_GGTTAACAGTATGGCG-1FC_MI_GFPve_pos     -0.3311285801 -4.0633881368
## FC_Mpos_GGTTCTCTCAACACGT-1FC_MI_GFPve_pos     -2.6287121601 -0.3645362534
## FC_Mpos_GTAACCACAGAGAGGG-1FC_MI_GFPve_pos     -4.0276880093 -2.1689600624
## FC_Mpos_GTAATGCTCCGTAGGC-1FC_MI_GFPve_pos     -4.2350632973 -2.1093675770
## FC_Mpos_GTAGAAACACAAATGA-1FC_MI_GFPve_pos     -2.2007326908  0.6865234695
## FC_Mpos_GTCATGATCTAAGAAG-1FC_MI_GFPve_pos     -4.0174074002 -1.2599706330
## FC_Mpos_GTGACGCGTTTCGGCG-1FC_MI_GFPve_pos     -4.0250861473 -0.6835312523
## FC_Mpos_GTGTTCCTCCCGAACG-1FC_MI_GFPve_pos     -3.5898015328 -0.9453308262
## FC_Mpos_TAAGTCGTCCGAGTGC-1FC_MI_GFPve_pos     -4.2292954750 -2.3300153889
## FC_Mpos_TAATTCCAGCCATTTG-1FC_MI_GFPve_pos     -3.6960463352 -2.7494051136
## FC_Mpos_TACTTGTAGTAAGAGG-1FC_MI_GFPve_pos     -0.7478627987  1.5236177765
```

```
## FC_Mpos_TAGTGCAAGGTAAGGA-1FC_MI_GFPve_pos     -2.3167631455  0.7213788353
## FC_Mpos_TCCATCGCAAGTATAG-1FC_MI_GFPve_pos     -4.1404299565 -2.0761601605
## FC_Mpos_TCGCTTGAGGACTTCT-1FC_MI_GFPve_pos     -4.1415927239 -2.3370752014
## FC_Mpos_TCGGGCAGTCTCCTGT-1FC_MI_GFPve_pos     -3.3429260082 -1.6622485794
## FC_Mpos_TCTATCAAGTCTAGCT-1FC_MI_GFPve_pos     -3.9640819855 -2.3964349903
## FC_Mpos_TCTCAGCAGTAAATGC-1FC_MI_GFPve_pos     -2.9639878102 -0.1266665138
## FC_Mpos_TCTTAGTGTGCGACAA-1FC_MI_GFPve_pos     -1.7093528338 -4.1027690448
## FC_Mpos_TGACAGTAGAGGGCGA-1FC_MI_GFPve_pos     -3.0560319252 -0.1271958031
## FC_Mpos_TGACTCCTCTGGCCAG-1FC_MI_GFPve_pos     -4.1017851658 -2.1660716213
## FC_Mpos_TGGGATTGTCTACGTA-1FC_MI_GFPve_pos     -0.4100295432 -1.9732150711
## FC_Mpos_TGTTGAGCACAAGCTT-1FC_MI_GFPve_pos     -2.6857831307 -0.6926903404
## FC_Mpos_TTAGGCAGTTCGAACT-1FC_MI_GFPve_pos     -3.5850190945 -2.7558012881
## FC_Mpos_TTCTAGTCATGACTGT-1FC_MI_GFPve_pos     -4.0514571495 -2.4715578236
## FC_Mpos_TTGTTGTCACTCCGGA-1FC_MI_GFPve_pos     -3.2358498402 -0.1707820572
## FC_Mpos_TTTGATCAGATGATTG-1FC_MI_GFPve_pos     -4.1160852738 -1.4475531258
## FC_Mpos_TTTGGAGCAAAGTATG-1FC_MI_GFPve_pos     -3.9965944119 -1.8978578724
## FC_Mpos_TTTGTTGGTGACGCCT-1FC_MI_GFPve_pos     -4.5762688942 -2.1979136147
## FC_Fpos_AACAACCCACCCTGTT-1FC_FI_GFPve_pos     -1.8212739058  0.6423106514
## FC_Fpos_AACGAAACAATCACGT-1FC_FI_GFPve_pos     -2.4895667858 -0.0732955612
## FC_Fpos_AACTTCTAGCCTCAGC-1FC_FI_GFPve_pos     -1.8448490925  1.7034435592
## FC_Fpos_AATTCCTAGCAAGTCG-1FC_FI_GFPve_pos     -2.8819954224 -1.3924920239
## FC_Fpos_ACACCAAGTCGCACAC-1FC_FI_GFPve_pos     -2.0010224409  0.9199791275
## FC_Fpos_ACATGCATCCCTCTTT-1FC_FI_GFPve_pos     -3.4419750996 -0.9886588730
## FC_Fpos_AGACACTAGGGTAGCT-1FC_FI_GFPve_pos     -3.9283499546 -1.4671384968
## FC_Fpos_AGCCAATGTTTCGTAG-1FC_FI_GFPve_pos     -2.1480190582  0.7073173843
## FC_Fpos_AGCGCTGGTAGACGTG-1FC_FI_GFPve_pos     -1.7630880900  0.3298063598
## FC_Fpos_AGGAAATTCGGCTGAC-1FC_FI_GFPve_pos     -3.3185775109 -2.4857453980
## FC_Fpos_AGGATAATCCGATAAC-1FC_FI_GFPve_pos     -3.3498861618 -1.1347384133
## FC_Fpos_AGGTAGGCACGTACAT-1FC_FI_GFPve_pos     -3.0585739441 -0.0742444672
## FC_Fpos_AGGTCATCACGTACAT-1FC_FI_GFPve_pos     -1.5288625784 -0.4746541657
## FC_Fpos_AGTAACCCAACGAGGT-1FC_FI_GFPve_pos     -3.2010552712 -1.0092096009
## FC_Fpos_AGTGTTGGTGGATGAC-1FC_FI_GFPve_pos     -3.2640690632  0.0308041893
## FC_Fpos_ATCCACCCAGGATGAC-1FC_FI_GFPve_pos     -2.0977879591  0.8106460892
## FC_Fpos_ATCCGTCGTTCTCTAT-1FC_FI_GFPve_pos     -3.0760919876  0.1773357712
## FC_Fpos_ATCCGTCTCACATACG-1FC_FI_GFPve_pos     -1.8866798706  0.9998803459
## FC_Fpos_ATCGTGAAGCACCAGA-1FC_FI_GFPve_pos     -0.6469956167 -2.0937773861
## FC_Fpos_ATCTTCAAGATGAATC-1FC_FI_GFPve_pos     -2.9551482029 -0.4116010346
## FC_Fpos_ATGTCCCGTTGGGAAC-1FC_FI_GFPve_pos     -2.0028503962 -1.1004335560
## FC_Fpos_CAAAGAAAGATTGCGG-1FC_FI_GFPve_pos     -2.9935927220 -0.3423323311
## FC_Fpos_CAAGAGGTCAACACGT-1FC_FI_GFPve_pos     -3.4883932896 -0.9821247734
## FC_Fpos_CACACAAAGTCTGTAC-1FC_FI_GFPve_pos     -3.8125729389 -0.5845799126
## FC_Fpos_CAGATACCAAGTGGTG-1FC_FI_GFPve_pos     -3.5797054596 -0.3420405068
## FC_Fpos_CAGATCATCTGCGGAC-1FC_FI_GFPve_pos     -3.5180508919 -0.6785788216
## FC_Fpos_CAGTTAGTCCTTCTTC-1FC_FI_GFPve_pos     -4.1128656693 -2.6149654068
## FC_Fpos_CATCAAGCAAGAGGCT-1FC_FI_GFPve_pos     -3.7388763256 -2.1806564011
## FC_Fpos_CATGCCTGTTGCGGCT-1FC_FI_GFPve_pos     -3.6699049301 -1.4221529640
## FC_Fpos_CATTGCCGTACTGTTG-1FC_FI_GFPve_pos     -1.0563778110  1.1050181709
## FC_Fpos_CATTGTTTCACCTTAT-1FC_FI_GFPve_pos     -4.2689707108 -1.3427631535
## FC_Fpos_CCAATGATCTCTGAGA-1FC_FI_GFPve_pos     -1.6147229500 -2.4335281529
## FC_Fpos_CCACACTAGCGGCTCT-1FC_FI_GFPve_pos     -2.4346499272  0.8598485313
## FC_Fpos_CCACGAGGTCCAATCA-1FC_FI_GFPve_pos     -4.3125951119 -2.1115555443
## FC_Fpos_CCCGAAGCAAGACCTT-1FC_FI_GFPve_pos     -1.4525968857 -1.3811435379
## FC_Fpos_CCCTAACGTACTGACT-1FC_FI_GFPve_pos     -4.0430931874 -1.8285927452
## FC_Fpos_CCCTGATCATAGGCGA-1FC_FI_GFPve_pos     -2.2645971604 -0.0618972458
## FC_Fpos_CCGTTCACACTAGAGG-1FC_FI_GFPve_pos     -1.4342275686  0.6972775780
## FC_Fpos_CCGTTCAGTACTAGCT-1FC_FI_GFPve_pos     -2.6420278378  0.4618382774
## FC_Fpos_CCTAACCCAGTCCCGA-1FC_FI_GFPve_pos     -4.6349198647 -1.6598078884
## FC_Fpos_CCTAACCTCATAGGCT-1FC_FI_GFPve_pos     -4.1895406075 -2.0868627705
## FC_Fpos_CCTAAGACAACGGCCT-1FC_FI_GFPve_pos     -1.0553020067 -1.6444668450
## FC_Fpos_CCTCACACAAATCCCA-1FC_FI_GFPve_pos     -4.5345060654 -1.8714849629
## FC_Fpos_CCTCCTCGTCCTCCTA-1FC_FI_GFPve_pos     -4.1194937058 -1.5901806034
## FC_Fpos_CCTCTCCCACGCGTGT-1FC_FI_GFPve_pos     -2.4375712700  0.2846327148
## FC_Fpos_CGAATTGGTTGTGTAC-1FC_FI_GFPve_pos     -3.2970323391 -1.0347487606
## FC_Fpos_CGAGGCTTCAACGAGG-1FC_FI_GFPve_pos     -3.9747094937 -0.6070284523
## FC_Fpos_CGGGCATTCACAACCA-1FC_FI_GFPve_pos     -3.5272571869 -0.3377413429
## FC_Fpos_CGTAGTAGTCACCCTT-1FC_FI_GFPve_pos     -2.8846516438 -0.6393150963
## FC_Fpos_CGTGCTTGTTGACTAC-1FC_FI_GFPve_pos     -3.9331829376 -1.1461689152
## FC_Fpos_CGTTCTGTCACACCCT-1FC_FI_GFPve_pos     -3.3846757240 -0.9523524918
## FC_Fpos_CTCATTATCTGGAGAG-1FC_FI_GFPve_pos     -0.8368028827 -2.0610203423
## FC_Fpos_CTCCCAACAGAGAGGG-1FC_FI_GFPve_pos     -2.2472474404  0.1859007202
```

```
## FC_Fpos_CTCCCAAGTCGGAACA-1FC_FI_GFPve_pos     -0.1984820474  0.2088585220
## FC_Fpos_CTCGAGGGTGGAGAAA-1FC_FI_GFPve_pos     -3.2254164047 -0.1448702492
## FC_Fpos_CTCTCGATCGTGGCTG-1FC_FI_GFPve_pos     -3.5639731713 -0.3830785431
## FC_Fpos_CTGCCATAGGAGCAAA-1FC_FI_GFPve_pos     -4.6850011177 -1.2497510590
## FC_Fpos_CTGGTCTCACTGGAAG-1FC_FI_GFPve_pos     -4.3789226837 -2.4619679131
## FC_Fpos_CTGTATTAGCGCTGCT-1FC_FI_GFPve_pos     -1.5279601641  0.0203500114
## FC_Fpos_CTGTGAACAATTAGGA-1FC_FI_GFPve_pos     -2.7931322880  0.6219926201
## FC_Fpos_CTTCCGAAGGACTAAT-1FC_FI_GFPve_pos     -3.7885899372 -1.5335383095
## FC_Fpos_CTTCGGTTCCCTCTTT-1FC_FI_GFPve_pos     -4.4154851265 -1.4432463326
## FC_Fpos_CTTCTAACAGAGTTGG-1FC_FI_GFPve_pos     -3.4976298638 -2.5519628205
## FC_Fpos_CTTTCAATCAAGTCTG-1FC_FI_GFPve_pos     -3.2950270004 -0.3530082382
## FC_Fpos_GAACGTTCAGAAGTGC-1FC_FI_GFPve_pos     -2.5837914772  0.6801967941
## FC_Fpos_GAAGTAATCTCCTGTG-1FC_FI_GFPve_pos     -0.5608018167 -0.6534042038
## FC_Fpos_GAGAAATCACGTTGGC-1FC_FI_GFPve_pos     -1.2335598297 -0.4036607422
## FC_Fpos_GAGCCTGTCTTTCCGG-1FC_FI_GFPve_pos     -3.5820295639 -0.8800158180
## FC_Fpos_GAGGGTAAGCGATGAC-1FC_FI_GFPve_pos     -1.2933736868 -0.2462820687
## FC_Fpos_GAGTGTTGTATCCCTC-1FC_FI_GFPve_pos     -4.3800298996 -1.3125388302
## FC_Fpos_GATAGAAAGAGGTCAC-1FC_FI_GFPve_pos     -2.2780775852  0.3685994468
## FC_Fpos_GATGACTTCCTGGTCT-1FC_FI_GFPve_pos     -2.7629661389  0.4662156425
## FC_Fpos_GATGAGGCAAAGGGCT-1FC_FI_GFPve_pos     -2.2367238827 -2.2765006699
## FC_Fpos_GCTGGGTCATGTGACT-1FC_FI_GFPve_pos     -2.4466745682 -1.5049941220
## FC_Fpos_GCTTTCGGTAGACACG-1FC_FI_GFPve_pos     -1.1471508570 -0.2077617325
## FC_Fpos_GCTTTCGGTTGAGTCT-1FC_FI_GFPve_pos     -3.4911904164 -0.6924557366
## FC_Fpos_GGAAGTGAGTCATCCA-1FC_FI_GFPve_pos      0.2930229180 -4.0621598043
## FC_Fpos_GGAATCTCATGGCTAT-1FC_FI_GFPve_pos      0.3473529987 -2.2210528530
## FC_Fpos_GGCTTGGCAAACTAAG-1FC_FI_GFPve_pos     -4.5107180901 -0.9394030251
## FC_Fpos_GGCTTTCTCAACACGT-1FC_FI_GFPve_pos     -3.3886782952 -1.2145821728
## FC_Fpos_GGGACTCAGCCGAACA-1FC_FI_GFPve_pos     -1.7252467699 -1.1001264729
## FC_Fpos_GGGTGTCCATGTTCGA-1FC_FI_GFPve_pos     -3.7383651562 -1.3107881226
## FC_Fpos_GGTTCTCTCCGATAGT-1FC_FI_GFPve_pos     -3.0166535206 -0.5302905716
## FC_Fpos_GTAACCATCAACTGAC-1FC_FI_GFPve_pos     -4.1716048546 -1.6462316193
## FC_Fpos_GTAGATCTCTCTGACC-1FC_FI_GFPve_pos     -3.2395813294 -0.2863726296
## FC_Fpos_GTCAGCGGTATGAGCG-1FC_FI_GFPve_pos     -2.7917482681  0.2065134369
## FC_Fpos_GTCATCCGTCTGCAAT-1FC_FI_GFPve_pos     -3.8428585358 -2.2859344162
## FC_Fpos_GTCATTTCAGGTGGAT-1FC_FI_GFPve_pos     -4.3417045899 -2.1248845734
## FC_Fpos_GTCCCATAGTGATTCC-1FC_FI_GFPve_pos     -1.9313586779 -1.2798514046
## FC_Fpos_GTCGCGAAGTTAACGA-1FC_FI_GFPve_pos     -3.5153143234 -0.5046200432
## FC_Fpos_GTTACAGCAAAGAACT-1FC_FI_GFPve_pos     -0.7124082871 -1.0541717686
## FC_Fpos_TAACCAGCAATACGCT-1FC_FI_GFPve_pos     -3.4734449215 -1.3181106724
## FC_Fpos_TAACCAGCACTTCAAG-1FC_FI_GFPve_pos     -4.8076059647 -1.4534539856
## FC_Fpos_TAATTCCTCCAGCCTT-1FC_FI_GFPve_pos     -3.3952255078 -2.6135694660
## FC_Fpos_TACCCACCACCTATCC-1FC_FI_GFPve_pos     -4.0328314133 -1.2594206013
## FC_Fpos_TACCGAAGTTTCCCAC-1FC_FI_GFPve_pos     -2.1209800072  0.8669777237
## FC_Fpos_TACCGGGGTCCTCCTA-1FC_FI_GFPve_pos     -3.0162086315 -1.0403468289
## FC_Fpos_TACTGCCCACGGTGTC-1FC_FI_GFPve_pos     -2.8945278950 -0.4211330094
## FC_Fpos_TACTGCCTCTGCGGGT-1FC_FI_GFPve_pos     -1.6190635987 -1.1068608441
## FC_Fpos_TACTTCATCACTAGCA-1FC_FI_GFPve_pos     -2.2984416313  0.8417902313
## FC_Fpos_TAGCACATCGCGCCAA-1FC_FI_GFPve_pos     -3.9794781037 -2.7737192787
## FC_Fpos_TATCCTATCCGCATAA-1FC_FI_GFPve_pos     -4.8332540817 -1.5390698590
## FC_Fpos_TCAATCTGTAGCTGTT-1FC_FI_GFPve_pos     -2.5022854634  0.2535553299
## FC_Fpos_TCAGGTACATCGGATT-1FC_FI_GFPve_pos     -1.4023325272 -0.6527561821
## FC_Fpos_TCCACCAGTAGCTGTT-1FC_FI_GFPve_pos     -3.7702846356 -2.5050475277
## FC_Fpos_TCCTGCAAGGTACTGG-1FC_FI_GFPve_pos     -0.7800477572 -0.3803548493
## FC_Fpos_TCCTTCTGTGATTCTG-1FC_FI_GFPve_pos     -3.4334246941 -0.5427069344
## FC_Fpos_TCGAACACACCTTCCA-1FC_FI_GFPve_pos     -0.8732142277  1.4711895309
## FC_Fpos_TCGGTCTGTCTACATG-1FC_FI_GFPve_pos     -2.0141143627 -0.9033465065
## FC_Fpos_TCTTAGTAGACACACG-1FC_FI_GFPve_pos     -1.0556030698  0.9106421791
## FC_Fpos_TCTTAGTCACCACTGG-1FC_FI_GFPve_pos     -0.9916629620 -0.3166851677
## FC_Fpos_TGAATGCAGACCAACG-1FC_FI_GFPve_pos     -2.3277966805 -3.5356865801
## FC_Fpos_TGACCCTGTATCAGGG-1FC_FI_GFPve_pos     -4.6134874649 -1.7123074211
## FC_Fpos_TGATGGTTCCATACTT-1FC_FI_GFPve_pos     -3.9635984726 -2.6997701802
## FC_Fpos_TGCATGAAGACAGCGT-1FC_FI_GFPve_pos     -2.7400960751 -0.7792765774
## FC_Fpos_TGCTGAAAGGTACATA-1FC_FI_GFPve_pos     -3.6208188362 -1.1797077336
## FC_Fpos_TGGGATTCATTCCTAT-1FC_FI_GFPve_pos     -1.2217547722  0.3070245109
## FC_Fpos_TGGTGATCATCACAGT-1FC_FI_GFPve_pos     -2.9770393200 -0.8008088745
## FC_Fpos_TGGTGATGTCTATGAC-1FC_FI_GFPve_pos     -1.2119123764  0.1635384880
## FC_Fpos_TGTTACTGTGTAGTGG-1FC_FI_GFPve_pos     -1.0581139155 -1.1509227432
## FC_Fpos_TTACAGGTCCTCACGT-1FC_FI_GFPve_pos     -3.0988139935 -1.2667810597
## FC_Fpos_TTACCATGTCGATTCA-1FC_FI_GFPve_pos     -1.5085053272 -4.5541755684
## FC_Fpos_TTACGTTCAAGAGTTA-1FC_FI_GFPve_pos     -1.7600180931 -1.3428530373
```

```
## FC_Fpos_TTCTAACCAAGGACAC-1FC_FI_GFPve_pos     -2.7130391426  0.3680587135
## FC_Fpos_TTGACCCGTATACGGG-1FC_FI_GFPve_pos     -2.2355298825  1.1069999061
## FC_Fpos_TTGTTGTTCTGCGGAC-1FC_FI_GFPve_pos     -3.4618942566 -0.3962907471
## FC_Fpos_TTTCCTCTCAAGATAG-1FC_FI_GFPve_pos     -1.7410237618  0.5015259109
## dCA2_Mpos_AAAGAACCATGTTCGA-1dCA2_MI_GFPve_pos -0.2590596877 -4.7168154173
## dCA2_Mpos_AAGACAATCGTCAACA-1dCA2_MI_GFPve_pos  0.2869616620 -4.2238039650
## dCA2_Mpos_AAGCCATAGGCTCACC-1dCA2_MI_GFPve_pos -1.1497490831 -5.0284150817
## dCA2_Mpos_AAGGTAAGTCGTATGT-1dCA2_MI_GFPve_pos -1.3848807640 -3.5546167769
## dCA2_Mpos_AATTTCCCATCAACCA-1dCA2_MI_GFPve_pos -1.6717626877 -3.8470527806
## dCA2_Mpos_ACAGAAAGTCAAAGCG-1dCA2_MI_GFPve_pos -0.6857719250 -5.0348396279
## dCA2_Mpos_ACATCGAGTCCGGATC-1dCA2_MI_GFPve_pos  1.1784610920 -3.1931308188
## dCA2_Mpos_ACCACAATCAAGGAGC-1dCA2_MI_GFPve_pos  1.2211251430 -1.8205017723
## dCA2_Mpos_ACCGTTCGTTAGGCCC-1dCA2_MI_GFPve_pos -0.9374853201 -5.0705381610
## dCA2_Mpos_ACCTGTCGTCGCAACC-1dCA2_MI_GFPve_pos -0.9054876991 -5.1920593955
## dCA2_Mpos_ACTGCAAGTCGGTAAG-1dCA2_MI_GFPve_pos -1.2694451637 -4.9388907083
## dCA2_Mpos_ACTGTGAAGTACCGGA-1dCA2_MI_GFPve_pos -1.0447028704 -5.2882635273
## dCA2_Mpos_ACTTTCAAGCCGAACA-1dCA2_MI_GFPve_pos -1.0162956543 -5.0457090713
## dCA2_Mpos_ACTTTGTCAATAACCC-1dCA2_MI_GFPve_pos -1.3239615984 -4.9728665181
## dCA2_Mpos_AGAAGCGAGTTGGAAT-1dCA2_MI_GFPve_pos -1.1839364596 -5.3207131304
## dCA2_Mpos_AGACAGGTCAATCAGC-1dCA2_MI_GFPve_pos -0.9277916379 -5.1478685118
## dCA2_Mpos_AGATCCATCCGACGGT-1dCA2_MI_GFPve_pos -1.2807808943 -3.3014642872
## dCA2_Mpos_AGGCATTCATCTATCT-1dCA2_MI_GFPve_pos -1.4586396046 -3.2189128079
## dCA2_Mpos_AGTAACCGTGACTGTT-1dCA2_MI_GFPve_pos -0.7980586119 -3.1663824000
## dCA2_Mpos_AGTAGTCCAAAGGCTG-1dCA2_MI_GFPve_pos  1.4634747677 -2.4741470494
## dCA2_Mpos_ATCACGACAGACCTAT-1dCA2_MI_GFPve_pos -1.0095368452 -5.2431676068
## dCA2_Mpos_ATCGTAGGTGTCCAAT-1dCA2_MI_GFPve_pos  0.7996873431 -3.6951864638
## dCA2_Mpos_ATTACTCCAGGTTACT-1dCA2_MI_GFPve_pos -0.9899313874 -5.1883020081
## dCA2_Mpos_ATTCCTAAGGACATCG-1dCA2_MI_GFPve_pos -1.3651702233 -3.0924893297
## dCA2_Mpos_ATTCGTTTCACCTCGT-1dCA2_MI_GFPve_pos -1.3244807549 -3.6648679651
## dCA2_Mpos_ATTTCACAGCGTGCCT-1dCA2_MI_GFPve_pos  0.8065646462 -3.0983136572
## dCA2_Mpos_CACGTTCTCTAAGCCA-1dCA2_MI_GFPve_pos -0.4620567687 -4.6215127684
## dCA2_Mpos_CAGTTAGCACGCGGTT-1dCA2_MI_GFPve_pos -1.0490141936 -5.2647001423
## dCA2_Mpos_CATCCGTTCACTGATG-1dCA2_MI_GFPve_pos -0.9219350047 -5.1384595610
## dCA2_Mpos_CCAATGAGTAGGGTAC-1dCA2_MI_GFPve_pos -1.2831431456 -3.2735848107
## dCA2_Mpos_CCAATGAGTTCATCGA-1dCA2_MI_GFPve_pos -1.4797720738 -3.7651275076
## dCA2_Mpos_CCTCATGCACAGCCTG-1dCA2_MI_GFPve_pos  1.1114720278 -3.1519707121
## dCA2_Mpos_CCTCTAGTCCGTTGGG-1dCA2_MI_GFPve_pos -1.2501454182 -5.0840405502
## dCA2_Mpos_CGAAGTTAGACGGTCA-1dCA2_MI_GFPve_pos -0.6317080982 -4.9703273304
## dCA2_Mpos_CGAAGTTTCCAAAGGG-1dCA2_MI_GFPve_pos -1.0606895514 -5.2540689983
## dCA2_Mpos_CGAATTGAGATTCGCT-1dCA2_MI_GFPve_pos  1.0179327898 -3.1307876982
## dCA2_Mpos_CGACAGCGTCCGGCAT-1dCA2_MI_GFPve_pos -1.4349851437 -3.2038715758
## dCA2_Mpos_CGAGAAGAGCATACTC-1dCA2_MI_GFPve_pos -0.8851903506 -5.1575807907
## dCA2_Mpos_CGGACACTCATCGCTC-1dCA2_MI_GFPve_pos -1.0101701446 -5.0702246703
## dCA2_Mpos_CGTAGTACATGGACAG-1dCA2_MI_GFPve_pos -0.7707227893 -3.2025216498
## dCA2_Mpos_CGTCAAAAGATACATG-1dCA2_MI_GFPve_pos -1.4663003512 -3.0290664114
## dCA2_Mpos_CGTGCTTAGAATGTTG-1dCA2_MI_GFPve_pos -0.7980483003 -4.6338922061
## dCA2_Mpos_CGTTAGATCCCATACC-1dCA2_MI_GFPve_pos -1.1022994943 -3.9598752536
## dCA2_Mpos_CTACCCAAGAGTTCGG-1dCA2_MI_GFPve_pos -0.9205815621 -5.2647292294
## dCA2_Mpos_CTAGGTAAGTTACTCG-1dCA2_MI_GFPve_pos -0.9773436613 -5.3440276660
## dCA2_Mpos_CTCAGTCCACTACGGC-1dCA2_MI_GFPve_pos  1.1220575743 -3.1237769999
## dCA2_Mpos_CTCATGCTCCTCGCAT-1dCA2_MI_GFPve_pos -1.2304750509 -3.7954770602
## dCA2_Mpos_CTCATTACAGAAGCTG-1dCA2_MI_GFPve_pos -1.1901454754 -4.9513108708
## dCA2_Mpos_CTCATTATCCTGATAG-1dCA2_MI_GFPve_pos -1.4130784102 -3.6453572191
## dCA2_Mpos_GAAGAATAGCCGTTAT-1dCA2_MI_GFPve_pos -0.8585402794 -5.0876308955
## dCA2_Mpos_GAAGGGTAGACGGTCA-1dCA2_MI_GFPve_pos  1.9786157779 -2.3112485089
## dCA2_Mpos_GAATAGAAGAGTCAGC-1dCA2_MI_GFPve_pos -0.9567757077 -5.0953854896
## dCA2_Mpos_GAATCGTCAAACTCGT-1dCA2_MI_GFPve_pos  1.3127457075 -1.7821325936
## dCA2_Mpos_GACCAATGTCAGATTC-1dCA2_MI_GFPve_pos -1.5583073921 -3.3565820135
## dCA2_Mpos_GACCGTGCATCGAAGG-1dCA2_MI_GFPve_pos -0.9128122754 -4.9379562117
## dCA2_Mpos_GAGAGGTAGAAGGTAG-1dCA2_MI_GFPve_pos  0.7750042252 -2.9880746283
## dCA2_Mpos_GAGATGGAGCCTATTG-1dCA2_MI_GFPve_pos -1.0392486878 -5.2152980246
## dCA2_Mpos_GATCAGTAGGCTCACC-1dCA2_MI_GFPve_pos -0.7797769137 -3.1803632654
## dCA2_Mpos_GATCAGTAGGGTGAAA-1dCA2_MI_GFPve_pos -1.4237152167 -3.8908317961
## dCA2_Mpos_GATTCGACATCTGCGG-1dCA2_MI_GFPve_pos -1.0790687628 -5.0916391589
## dCA2_Mpos_GCAGCCACAACGTTAC-1dCA2_MI_GFPve_pos -1.5490643807 -3.2238652386
## dCA2_Mpos_GCAGCTGAGGATTACA-1dCA2_MI_GFPve_pos -1.4263930149 -4.8552297183
## dCA2_Mpos_GCATCTCTCCGCCTAT-1dCA2_MI_GFPve_pos  1.0696822576 -3.1921195664
## dCA2_Mpos_GCGTTTCTCGTCTCAC-1dCA2_MI_GFPve_pos -1.1892269678 -3.5804479994
## dCA2_Mpos_GCTCAAAAGTCCTACA-1dCA2_MI_GFPve_pos -0.9614255257 -5.1071835734
## dCA2_Mpos_GCTGCAGGTCACCACG-1dCA2_MI_GFPve_pos -1.8832924195 -3.1279282250
```

```
## dCA2_Mpos_GGAACCCCATCCGGCA-1dCA2_MI_GFPve_pos -0.9564450808 -4.9353715755
## dCA2_Mpos_GGAGAACAGCACCAGA-1dCA2_MI_GFPve_pos -1.0805932112 -5.1914713062
## dCA2_Mpos_GGAGCAACAGAGGGTT-1dCA2_MI_GFPve_pos -1.3971696921 -3.2602123894
## dCA2_Mpos_GGAGGTAGTCCTACAA-1dCA2_MI_GFPve_pos  1.0956086092 -3.1929009833
## dCA2_Mpos_GGGCGTTAGCTTCTAG-1dCA2_MI_GFPve_pos -1.3296274967 -3.1808396258
## dCA2_Mpos_GTAGATCTCAGTGTTG-1dCA2_MI_GFPve_pos -0.9766934343 -3.5527949013
## dCA2_Mpos_GTAGCTATCCATGATG-1dCA2_MI_GFPve_pos -1.3476303644 -4.6992362715
## dCA2_Mpos_GTCCTCATCTTAGGAC-1dCA2_MI_GFPve_pos -0.8944611378 -5.2927719034
## dCA2_Mpos_GTCTCACAGCACGATG-1dCA2_MI_GFPve_pos -1.5465744562 -3.3009297528
## dCA2_Mpos_GTCTGTCTCCGTACGG-1dCA2_MI_GFPve_pos -1.4444420166 -3.2250750937
## dCA2_Mpos_GTGGAAGAGCGAGTCA-1dCA2_MI_GFPve_pos -1.4300607271 -4.4002540685
## dCA2_Mpos_GTTCCGTAGGTAGGCT-1dCA2_MI_GFPve_pos -1.1110710926 -3.5013432183
## dCA2_Mpos_TAAGCGTAGTGGAAGA-1dCA2_MI_GFPve_pos -1.2749284334 -3.3315720238
## dCA2_Mpos_TACCCGTCAGCGTAGA-1dCA2_MI_GFPve_pos -1.1597135492 -3.5792939343
## dCA2_Mpos_TACTTCAAGTTTGAGA-1dCA2_MI_GFPve_pos -1.0572242565 -5.1338255979
## dCA2_Mpos_TAGGTTGCAGTGGGTA-1dCA2_MI_GFPve_pos -1.0100609489 -5.2721018471
## dCA2_Mpos_TATCGCCAGTTAACGA-1dCA2_MI_GFPve_pos -0.7743471689 -3.2310328163
## dCA2_Mpos_TCACAAGCAACTAGAA-1dCA2_MI_GFPve_pos -1.2265324421 -3.6343265452
## dCA2_Mpos_TCACATTGTTGATCGT-1dCA2_MI_GFPve_pos -1.4977524109 -3.2279369511
## dCA2_Mpos_TCCACGTAGTGGGAAA-1dCA2_MI_GFPve_pos  1.1804950409 -3.0989963688
## dCA2_Mpos_TCCAGAATCTACTCAT-1dCA2_MI_GFPve_pos  0.5237326197 -2.6288692631
## dCA2_Mpos_TCTAACTTCCCTGTTG-1dCA2_MI_GFPve_pos -1.4966949053 -3.2086676039
## dCA2_Mpos_TGATGGTGTGACAACG-1dCA2_MI_GFPve_pos -0.7993963189 -3.2095489182
## dCA2_Mpos_TGCGGCATCTACTATC-1dCA2_MI_GFPve_pos -1.4627669878 -3.2200119175
## dCA2_Mpos_TGGTAGTCAGTTGGTT-1dCA2_MI_GFPve_pos  0.9339993171 -3.5580897011
## dCA2_Mpos_TGTTCATTCTAAGGAA-1dCA2_MI_GFPve_pos -1.5012713499 -3.2000995554
## dCA2_Mpos_TGTTCCGCATTCACCC-1dCA2_MI_GFPve_pos -1.0807552166 -5.2179013170
## dCA2_Mpos_TGTTGAGCAGTCGAGA-1dCA2_MI_GFPve_pos -1.4283959694 -3.4847161450
## dCA2_Mpos_TTACGCCTCGGCTGAC-1dCA2_MI_GFPve_pos -1.4661863871 -3.1867968716
## dCA2_Mpos_TTAGTCTCACTGTCCT-1dCA2_MI_GFPve_pos -1.4384537764 -3.8736309685
## dCA2_Mpos_TTCACGCTCGACCTAA-1dCA2_MI_GFPve_pos -0.1078276612 -3.9024714507
## dCA2_Mpos_TTGCGTCCAAGGAGTC-1dCA2_MI_GFPve_pos -1.1645055838 -3.4172662176
## dCA2_Mpos_TTGTTCAGTGCGAACA-1dCA2_MI_GFPve_pos -1.5127290316 -4.0912085451
## dCA2_Mpos_TTGTTGTAGAAACACT-1dCA2_MI_GFPve_pos -1.5121285744 -3.9188796558
## dCA2_Mpos_TTTCACAAGTTCTCTT-1dCA2_MI_GFPve_pos -0.8686604328 -5.1163562216
## dCA2_Fpos_AGGATCTTCACCTGGG-1dCA2_FI_GFPve_pos -0.7912481852 -4.4667462684
## dCA2_Fpos_AGGTCATTCCACAGGC-1dCA2_FI_GFPve_pos -1.6236722298 -3.6726208605
## dCA2_Fpos_AGTAGTCAGCTTACGT-1dCA2_FI_GFPve_pos -0.7875193782 -3.2352759518
## dCA2_Fpos_ATCATTCAGGTCATCT-1dCA2_FI_GFPve_pos -0.9644526072 -5.2251077570
## dCA2_Fpos_ATCTTCACAGAAGTGC-1dCA2_FI_GFPve_pos -1.1075981088 -5.2604849018
## dCA2_Fpos_ATGGGAGAGCATGGGT-1dCA2_FI_GFPve_pos  1.1060598068 -3.1809039988
## dCA2_Fpos_CCCTCAAGTTGTGGAG-1dCA2_FI_GFPve_pos -1.0616698094 -4.1536042966
## dCA2_Fpos_GCAACCGCAATAGTAG-1dCA2_FI_GFPve_pos -1.1566007562 -5.3008880891
## dCA2_Fpos_GCCCAGATCTTGTTAC-1dCA2_FI_GFPve_pos  1.0436357431 -2.9944451727
## dCA2_Fpos_GTAGAGGCAACAACAA-1dCA2_FI_GFPve_pos -1.3681042023 -3.7803595938
## dCA2_Fpos_TACGTCCTCCAATGCA-1dCA2_FI_GFPve_pos -1.5092131920 -3.2340093531
## dCA2_Fpos_TAGGAGGTCTGACCCT-1dCA2_FI_GFPve_pos  1.0347697906 -3.1641108669
## dCA2_Fpos_TCATGCCGTCCCGGTA-1dCA2_FI_GFPve_pos  1.1886050873 -3.1327811159
## dCA2_Fpos_TCGGGCAAGCGTTCCG-1dCA2_FI_GFPve_pos -1.5226931400 -4.0919318475
## dCA2_Fpos_TCTTTGAGTGTTTCTT-1dCA2_FI_GFPve_pos -1.5653711386 -3.3275302328
## dCA2_Fpos_TGAACGTGTCGACGCT-1dCA2_FI_GFPve_pos -0.7870209046 -3.2011372723
## dCA2_Fpos_TGCTGAAGTACCCAGC-1dCA2_FI_GFPve_pos -1.5331945248 -3.6662966885
## dCA2_Fpos_TGCTGAAGTGAATATG-1dCA2_FI_GFPve_pos  1.9203241043 -2.2327415623
## dCA2_Fpos_TGCTTCGCATAGAAAC-1dCA2_FI_GFPve_pos -1.3819751568 -3.2341037668
## dCA2_Fpos_TGTCCCACAGTCTACA-1dCA2_FI_GFPve_pos  1.1102023296 -3.0451393999
## dCA2_Fpos_TTGCATTGTGCAGTGA-1dCA2_FI_GFPve_pos -1.9552277155 -4.6053167708
## FC_Mpos_AAACCCACAGACTGCC-1FC_MI_GFPve_pos     -0.6766564079 -4.9406450965
## FC_Mpos_AAAGAACAGTGCAGCA-1FC_MI_GFPve_pos     -1.1464271970 -5.1758994140
## FC_Mpos_AAAGTCCGTGCGTTTA-1FC_MI_GFPve_pos     -2.0618181057 -3.6605784573
## FC_Mpos_AAATGGAAGCTCCGAC-1FC_MI_GFPve_pos     -1.7658060618 -4.1531373538
## FC_Mpos_AAATGGACAGTTCACA-1FC_MI_GFPve_pos     -0.6020818479 -4.3995878376
## FC_Mpos_AACCACAGTTTACCAG-1FC_MI_GFPve_pos     -0.7431363292 -4.9766336359
## FC_Mpos_AACGAAAAGGCAGGGA-1FC_MI_GFPve_pos     -2.2138862438 -3.6029385008
## FC_Mpos_AAGACAACAGTAGATA-1FC_MI_GFPve_pos      1.4350398950 -2.3337380566
## FC_Mpos_AAGACTCAGGTTCCGC-1FC_MI_GFPve_pos     -0.9151542611 -4.5796590872
## FC_Mpos_AAGATAGTCGCCTTTG-1FC_MI_GFPve_pos     -1.9199384279 -4.1717711963
## FC_Mpos_AAGCGAGCATCGGTTA-1FC_MI_GFPve_pos     -1.6043426819 -3.9874784626
## FC_Mpos_AAGCGAGTCGAAGCCC-1FC_MI_GFPve_pos     -1.0153761931 -4.9943110027
## FC_Mpos_AAGCGTTCAAAGCACG-1FC_MI_GFPve_pos     -1.9582810230 -4.5836085506
## FC_Mpos_AATGAAGAGTGAGCCA-1FC_MI_GFPve_pos     -1.5359341927 -3.6016626038
```

```
## FC_Mpos_AATGGCTAGATACAGT-1FC_MI_GFPve_pos     -1.8668544121 -4.3877625443
## FC_Mpos_ACAAAGAAGTTAGTGA-1FC_MI_GFPve_pos      1.4589369468 -2.4158944763
## FC_Mpos_ACAGCCGGTCGAGCAA-1FC_MI_GFPve_pos     -0.9145877786 -5.1915577926
## FC_Mpos_ACATCGAAGGGTACGT-1FC_MI_GFPve_pos     -1.6313426323 -4.2362888135
## FC_Mpos_ACCAACACAGCGGTCT-1FC_MI_GFPve_pos     -1.4069416352 -3.6456557192
## FC_Mpos_ACCATTTGTCATAAAG-1FC_MI_GFPve_pos     -1.8241475649 -4.1899870910
## FC_Mpos_ACCATTTGTGACTCGC-1FC_MI_GFPve_pos     -0.7991124220 -4.9660490729
## FC_Mpos_ACCATTTGTGTCATTG-1FC_MI_GFPve_pos     -1.0517005153 -5.0104300656
## FC_Mpos_ACGCACGCACAACGTT-1FC_MI_GFPve_pos     -1.9797630139 -3.7534224667
## FC_Mpos_ACGTAACGTTCTTAGG-1FC_MI_GFPve_pos     -0.6517624982 -5.0184921898
## FC_Mpos_ACTATCTGTGGGCTCT-1FC_MI_GFPve_pos     -1.4029623099 -4.5429903247
## FC_Mpos_ACTTCCGAGATACTGA-1FC_MI_GFPve_pos     -2.0516854353 -2.9389488377
## FC_Mpos_ACTTCGCAGGCTGAAC-1FC_MI_GFPve_pos      0.1621577911 -2.4689380802
## FC_Mpos_ACTTCGCAGGTGCTAG-1FC_MI_GFPve_pos     -0.8965923019 -4.9438714959
## FC_Mpos_ACTTTGTAGACCTGGA-1FC_MI_GFPve_pos     -2.1095521279 -2.9681582131
## FC_Mpos_AGAAATGCAACATCGT-1FC_MI_GFPve_pos     -0.3059510566 -4.7369239159
## FC_Mpos_AGACACTCACTACCCT-1FC_MI_GFPve_pos     -1.3173266478 -4.7376944319
## FC_Mpos_AGACCCGGTGGGCTCT-1FC_MI_GFPve_pos      1.5429254942 -2.6111814656
## FC_Mpos_AGAGAGCGTAGTACGG-1FC_MI_GFPve_pos     -1.3145919867 -4.9133427717
## FC_Mpos_AGAGAGCTCATGAAAG-1FC_MI_GFPve_pos      1.0328588657 -3.2812950291
## FC_Mpos_AGATAGAAGGTTAGTA-1FC_MI_GFPve_pos      0.6496547632 -3.9218405761
## FC_Mpos_AGCCAATAGTTCACTG-1FC_MI_GFPve_pos      1.0736447744 -3.5059357561
## FC_Mpos_AGCGTATCATACACCA-1FC_MI_GFPve_pos     -0.4262584276 -4.8078253769
## FC_Mpos_AGCTACACACGGTGCT-1FC_MI_GFPve_pos      0.0531824850 -4.2976029493
## FC_Mpos_AGGAATAAGGCAGGGA-1FC_MI_GFPve_pos     -0.1696211222 -4.6034516223
## FC_Mpos_AGGTCTACAAGGCAAC-1FC_MI_GFPve_pos      0.9551658802 -3.5347650923
## FC_Mpos_AGGTCTACACATGACT-1FC_MI_GFPve_pos     -1.3467663355 -4.8536365532
## FC_Mpos_AGGTTGTCACAAATCC-1FC_MI_GFPve_pos     -1.5605234928 -4.7504082821
## FC_Mpos_AGTAACCCAGAACTTC-1FC_MI_GFPve_pos      0.1575229935 -4.3842338897
## FC_Mpos_AGTTAGCGTCTGTGCG-1FC_MI_GFPve_pos     -1.7979632445 -3.7025018610
## FC_Mpos_AGTTCCCGTAAGCGGT-1FC_MI_GFPve_pos     -0.8368836470 -4.5573630311
## FC_Mpos_AGTTCCCGTACGAGCA-1FC_MI_GFPve_pos     -1.8019753523 -3.9572561659
## FC_Mpos_AGTTCCCGTACGCTTA-1FC_MI_GFPve_pos     -1.7111822195 -4.1487127222
## FC_Mpos_ATATCCTCAACGACAG-1FC_MI_GFPve_pos     -0.7762749620 -4.9568733849
## FC_Mpos_ATCACGATCCTCTTTC-1FC_MI_GFPve_pos     -2.4967317410 -3.3353864826
## FC_Mpos_ATCATTCGTCAGGTGA-1FC_MI_GFPve_pos     -1.9891055651 -3.0306044497
## FC_Mpos_ATCATTCTCGCACTCT-1FC_MI_GFPve_pos     -1.3364306517 -3.1479288019
## FC_Mpos_ATGAAAGAGATCGACG-1FC_MI_GFPve_pos      1.8593843155 -2.3743850865
## FC_Mpos_ATGAGTCGTCAAGTTC-1FC_MI_GFPve_pos     -1.4405697413 -4.8165080376
## FC_Mpos_ATGTCCCAGGAACATT-1FC_MI_GFPve_pos      1.2222747974 -3.1922212519
## FC_Mpos_ATTACTCGTCAACACT-1FC_MI_GFPve_pos     -1.3756038971 -4.4118834414
## FC_Mpos_ATTCTTGTCACTTTGT-1FC_MI_GFPve_pos     -1.6179096527 -4.1653167881
## FC_Mpos_ATTGGGTCAGGCTTGC-1FC_MI_GFPve_pos     -1.2145671673 -4.7909919411
## FC_Mpos_CAAAGAATCGGAATGG-1FC_MI_GFPve_pos     -0.6440957911 -4.8930802323
## FC_Mpos_CAACCTCGTACCTTCC-1FC_MI_GFPve_pos     -0.7514761515 -5.1859467603
## FC_Mpos_CAACGGCCATCAGCAT-1FC_MI_GFPve_pos     -1.8364329167 -3.3094623961
## FC_Mpos_CAAGCTAGTGACTATC-1FC_MI_GFPve_pos     -1.7641967364 -4.5999011375
## FC_Mpos_CAAGGGAGTTGCTCGG-1FC_MI_GFPve_pos      1.4822266273 -2.4982194580
## FC_Mpos_CAATACGAGGCGTTGA-1FC_MI_GFPve_pos     -1.1089811750 -4.0035825648
## FC_Mpos_CAATCGAGTTCTCCCA-1FC_MI_GFPve_pos     -2.1704094238 -3.1870057263
## FC_Mpos_CACAACATCAACCTTT-1FC_MI_GFPve_pos     -1.6620685883 -3.2174318709
## FC_Mpos_CACAGATCATTGTCGA-1FC_MI_GFPve_pos     -1.6941883393  2.6383467041
## FC_Mpos_CACGTGGAGAAGTATC-1FC_MI_GFPve_pos      0.9764348440 -3.5215509810
## FC_Mpos_CACGTTCAGGGAGGGT-1FC_MI_GFPve_pos      0.9814055137 -2.9577843823
## FC_Mpos_CAGATCAGTCCAAAGG-1FC_MI_GFPve_pos      0.5796766453 -3.5405906119
## FC_Mpos_CAGCGTGGTGATACCT-1FC_MI_GFPve_pos     -1.2824750967 -5.2120634295
## FC_Mpos_CAGTGCGGTCTTACAG-1FC_MI_GFPve_pos     -0.4062158741 -4.4572221019
## FC_Mpos_CAGTTCCGTCTTTCAT-1FC_MI_GFPve_pos     -0.7776343651 -4.9002339371
## FC_Mpos_CATGCCTAGTTGGCTT-1FC_MI_GFPve_pos     -1.5408081837 -4.1490308918
## FC_Mpos_CATGCGGTCCCTCTTT-1FC_MI_GFPve_pos     -0.0183966137 -4.3325940050
## FC_Mpos_CATTCATAGATTGATG-1FC_MI_GFPve_pos     -0.6965707965 -4.8780793376
## FC_Mpos_CATTTCACATCAACCA-1FC_MI_GFPve_pos     -1.6927631922 -4.8811942466
## FC_Mpos_CCAAGCGCACGCGTGT-1FC_MI_GFPve_pos     -2.2677302189 -3.3452924170
## FC_Mpos_CCAATGATCCTGTAAG-1FC_MI_GFPve_pos     -1.7117168732 -4.7274771143
## FC_Mpos_CCACACTAGAAGCGGG-1FC_MI_GFPve_pos     -1.4218776055 -4.5184279360
## FC_Mpos_CCCATTGAGTCATCCA-1FC_MI_GFPve_pos     -1.5694734879 -4.9178673305
## FC_Mpos_CCCATTGCATTGAAAG-1FC_MI_GFPve_pos     -1.2308673687 -4.9437871702
## FC_Mpos_CCCTAACGTCTCAGGC-1FC_MI_GFPve_pos     -1.7870045729 -4.2350652494
## FC_Mpos_CCGAACGCAGGTGAGT-1FC_MI_GFPve_pos     -1.2167285509 -5.2313235082
## FC_Mpos_CCGGACACACCTTCGT-1FC_MI_GFPve_pos     -2.9977845974 -1.3945402779
```

```
## FC_Mpos_CCGGACATCACTGGTA-1FC_MI_GFPve_pos     -0.8398137517 -4.8313936196
## FC_Mpos_CCGGTGACACAGTATC-1FC_MI_GFPve_pos     -1.1656820007 -4.7249221556
## FC_Mpos_CCGTAGGAGGTCGTGA-1FC_MI_GFPve_pos     -1.5502800770 -5.2168318011
## FC_Mpos_CCTGCATAGCGTTACT-1FC_MI_GFPve_pos     -1.8142974205 -4.1564184107
## FC_Mpos_CGAATTGTCCTGGGTG-1FC_MI_GFPve_pos     -1.4783159323 -4.2539994396
## FC_Mpos_CGAGGAAAGTGCTAGG-1FC_MI_GFPve_pos     -0.8531105347 -4.9507086553
## FC_Mpos_CGAGTTATCTGAGCAT-1FC_MI_GFPve_pos     -1.2456470556 -5.1015460469
## FC_Mpos_CGCATAATCTTAGTTC-1FC_MI_GFPve_pos     -1.1728864737 -5.0716566719
## FC_Mpos_CGGAACCAGGTTGGTG-1FC_MI_GFPve_pos      0.2001959257 -4.3101000764
## FC_Mpos_CGGGACTGTCCTCAGG-1FC_MI_GFPve_pos     -0.5199113615 -4.6355815359
## FC_Mpos_CGGGTGTCAGAACTCT-1FC_MI_GFPve_pos     -1.6696682997 -4.1420847453
## FC_Mpos_CGTCAAAGTTATAGCC-1FC_MI_GFPve_pos     -2.2830862828 -3.6245127596
## FC_Mpos_CGTCCATCAGCACCCA-1FC_MI_GFPve_pos     -1.0634774990 -5.0480074264
## FC_Mpos_CGTGATATCCACCTGT-1FC_MI_GFPve_pos     -1.4368889160 -3.7078590073
## FC_Mpos_CGTTAGAGTTGTGGAG-1FC_MI_GFPve_pos     -1.7978706188 -4.6175668515
## FC_Mpos_CTAACTTAGGGCCCTT-1FC_MI_GFPve_pos     -1.0944457479 -4.7639676478
## FC_Mpos_CTACAGAGTTTCGTGA-1FC_MI_GFPve_pos     -1.1130341716 -5.0745355465
## FC_Mpos_CTACCCACAGGCCCTA-1FC_MI_GFPve_pos     -2.3941380806 -3.1846301235
## FC_Mpos_CTCAACCTCGAACGCC-1FC_MI_GFPve_pos     -1.1364088483 -4.9297589190
## FC_Mpos_CTCCATGCACAGCCAC-1FC_MI_GFPve_pos     -0.5661165841 -4.5474793442
## FC_Mpos_CTGAGCGAGCTGAGTG-1FC_MI_GFPve_pos     -1.3484510012 -5.0461048164
## FC_Mpos_CTGCGAGTCAGAATAG-1FC_MI_GFPve_pos      0.1127864890 -4.4066527166
## FC_Mpos_CTTAGGAGTTGAAGTA-1FC_MI_GFPve_pos     -0.4780033953 -4.7590435404
## FC_Mpos_CTTCTCTAGCTGAAAT-1FC_MI_GFPve_pos     -1.7518706150 -4.4280226208
## FC_Mpos_GAACACTAGAGCAACC-1FC_MI_GFPve_pos     -0.5014723070 -4.8120704122
## FC_Mpos_GAAGAATAGCGGGTAT-1FC_MI_GFPve_pos      1.1961757116 -2.9310086645
## FC_Mpos_GAAGCCCCAATACGCT-1FC_MI_GFPve_pos     -0.7815666623 -4.9189819373
## FC_Mpos_GACTTCCAGGACAGCT-1FC_MI_GFPve_pos     -1.0873093434 -4.7191541843
## FC_Mpos_GAGACCCTCTCTTAAC-1FC_MI_GFPve_pos     -0.7681712337 -4.9324742831
## FC_Mpos_GAGCCTGTCATCACTT-1FC_MI_GFPve_pos      0.2632886343 -4.2934787251
## FC_Mpos_GAGGCCTCATACCACA-1FC_MI_GFPve_pos     -0.8415215440 -5.1647149600
## FC_Mpos_GAGGGTAAGCGATGCA-1FC_MI_GFPve_pos     -1.1132963128 -5.1227923013
## FC_Mpos_GAGTCATAGAAGTATC-1FC_MI_GFPve_pos      0.0985308759 -4.4420278348
## FC_Mpos_GAGTGAGAGAAGGTAG-1FC_MI_GFPve_pos     -1.3467037506 -4.7953954019
## FC_Mpos_GAGTGAGGTAGCTGAG-1FC_MI_GFPve_pos     -1.2507830687 -4.1434214987
## FC_Mpos_GAGTGTTCACGTACTA-1FC_MI_GFPve_pos     -0.8441915937 -4.9546465852
## FC_Mpos_GAGTTACCACAGCATT-1FC_MI_GFPve_pos     -1.5130611487 -4.8342849068
## FC_Mpos_GAGTTTGAGAATTCAG-1FC_MI_GFPve_pos     -1.6580688782 -4.7488020450
## FC_Mpos_GATGAGGCAAGGCTTT-1FC_MI_GFPve_pos     -1.3969724007 -3.6068097033
## FC_Mpos_GATGTTGAGCTACAAA-1FC_MI_GFPve_pos     -2.3479218312 -3.7127471842
## FC_Mpos_GCAACCGTCGTAGGAG-1FC_MI_GFPve_pos     -0.9426641889 -4.9666129328
## FC_Mpos_GCACGGTGTAGCGAGT-1FC_MI_GFPve_pos     -1.6733293362 -5.0240293421
## FC_Mpos_GCAGCCAGTAGATCCT-1FC_MI_GFPve_pos     -1.0149774380 -4.8828651883
## FC_Mpos_GCAGCTGTCAAGCCCG-1FC_MI_GFPve_pos     -0.5212112792 -4.4674051680
## FC_Mpos_GCAGGCTAGGTTGAGC-1FC_MI_GFPve_pos     -0.5665908582 -4.7967263773
## FC_Mpos_GCCATTCTCTTCCCAG-1FC_MI_GFPve_pos     -1.5644237824 -4.3723911621
## FC_Mpos_GCCTGTTAGAGCATTA-1FC_MI_GFPve_pos      0.2207952015 -4.2702763952
## FC_Mpos_GCCTGTTAGGGTACGT-1FC_MI_GFPve_pos     -1.9447779484 -3.1696270384
## FC_Mpos_GCGGAAACAGTCGTTA-1FC_MI_GFPve_pos      0.9072603159 -3.6172321715
## FC_Mpos_GCGGATCTCATTTGGG-1FC_MI_GFPve_pos     -0.0204762049 -4.7277475061
## FC_Mpos_GCGTGCATCCCTCAAC-1FC_MI_GFPve_pos     -1.5180700846 -4.4408012904
## FC_Mpos_GCTACCTCACAAAGTA-1FC_MI_GFPve_pos     -2.8817815609 -2.0813154854
## FC_Mpos_GCTTGGGTCGACCTAA-1FC_MI_GFPve_pos     -1.0428240128 -5.0461683251
## FC_Mpos_GGAATCTAGGCGTTGA-1FC_MI_GFPve_pos     -1.3901736565 -3.4338335671
## FC_Mpos_GGAGAACCAATACGCT-1FC_MI_GFPve_pos     -0.9445683189 -5.0781125702
## FC_Mpos_GGAGGATAGATCCCGC-1FC_MI_GFPve_pos     -2.0523949690 -2.6077246346
## FC_Mpos_GGGATCCAGTGGATAT-1FC_MI_GFPve_pos     -1.6646588869 -4.0275504269
## FC_Mpos_GGGATCCTCCCTATTA-1FC_MI_GFPve_pos     -1.4502836294 -3.7408575691
## FC_Mpos_GGGCCATCAACGGCCT-1FC_MI_GFPve_pos     -1.0494078822 -4.9416988917
## FC_Mpos_GGGTGAACAAGTTGGG-1FC_MI_GFPve_pos     -1.3313821383 -5.2525260724
## FC_Mpos_GGTAATCCACAAAGCG-1FC_MI_GFPve_pos     -1.4899701901 -4.5275664307
## FC_Mpos_GGTGAAGAGTAACAGT-1FC_MI_GFPve_pos     -1.0419961043 -3.6134512104
## FC_Mpos_GGTGAAGAGTAACCGG-1FC_MI_GFPve_pos     -1.3847631045 -3.8797264375
## FC_Mpos_GGTGGCTGTGTCTCCT-1FC_MI_GFPve_pos     -0.9261424370 -4.9931153335
## FC_Mpos_GGTGTCGCAAGTGGGT-1FC_MI_GFPve_pos     -0.2975292034 -4.4267252959
## FC_Mpos_GGTGTCGGTGTTAACC-1FC_MI_GFPve_pos      0.8144515924 -3.1332446017
## FC_Mpos_GGTGTCGTCCCTCGAT-1FC_MI_GFPve_pos     -1.4193483420 -4.7197506219
## FC_Mpos_GGTTGTAAGGCGATAC-1FC_MI_GFPve_pos     -1.5701953001 -4.8267933928
## FC_Mpos_GTAGATCCACTGTCCT-1FC_MI_GFPve_pos      1.3001917772 -2.8013250508
## FC_Mpos_GTAGATCTCTCTCGAC-1FC_MI_GFPve_pos     -1.9139607973 -4.0236470975
```

```
## FC_Mpos_GTAGCTAAGGGTCTTT-1FC_MI_GFPve_pos     -1.1885249443 -4.3572849311
## FC_Mpos_GTCAAACGTACCGTCG-1FC_MI_GFPve_pos     -1.3684732743 -3.9399707593
## FC_Mpos_GTCACTCCACATGTTG-1FC_MI_GFPve_pos     -1.2940400667 -4.6838311158
## FC_Mpos_GTCACTCCATGGTGGA-1FC_MI_GFPve_pos     -0.7864466138 -4.6189553268
## FC_Mpos_GTCATGAGTTGGATCT-1FC_MI_GFPve_pos     -1.5619319506  2.5866823517
## FC_Mpos_GTCGTTCCACGACAAG-1FC_MI_GFPve_pos     -1.3186151810 -3.7808076419
## FC_Mpos_GTGACGCAGCAACCAG-1FC_MI_GFPve_pos      0.2622512452 -4.0081705727
## FC_Mpos_GTGAGTTCATCAACCA-1FC_MI_GFPve_pos     -2.0401556320 -4.0651992835
## FC_Mpos_GTGGAAGAGCACCGTC-1FC_MI_GFPve_pos     -1.3015531130 -4.5488131471
## FC_Mpos_GTGGAGACAGTTGTCA-1FC_MI_GFPve_pos      1.8121385746 -2.0477730908
## FC_Mpos_GTTACCCAGCAAATGT-1FC_MI_GFPve_pos     -1.4055121966 -3.3369944967
## FC_Mpos_GTTAGACGTTCTCTAT-1FC_MI_GFPve_pos      1.3211857252 -3.1571578659
## FC_Mpos_GTTAGTGAGCATTGAA-1FC_MI_GFPve_pos      1.0883791618 -3.4432667174
## FC_Mpos_GTTGAACTCCGGACTG-1FC_MI_GFPve_pos     -2.0051187105 -3.1251699604
## FC_Mpos_GTTGCTCTCCCAGGAC-1FC_MI_GFPve_pos     -1.7808083124 -4.4410038270
## FC_Mpos_TAACACGCATGACGAG-1FC_MI_GFPve_pos      1.0461710863 -3.5647279896
## FC_Mpos_TAACGACGTTACCCTC-1FC_MI_GFPve_pos     -1.7105777092 -4.5728536226
## FC_Mpos_TAATTCCCAGGTCTCG-1FC_MI_GFPve_pos     -1.8264041968 -4.4315516926
## FC_Mpos_TAATTCCGTCACCGAC-1FC_MI_GFPve_pos     -0.6092081495 -4.7297626516
## FC_Mpos_TACAGGTCAAGCTGCC-1FC_MI_GFPve_pos      0.5910641722 -3.9545157828
## FC_Mpos_TAGACTGGTTTCCCAC-1FC_MI_GFPve_pos      1.0240740948 -3.6302534022
## FC_Mpos_TATTGCTGTAGCGTCC-1FC_MI_GFPve_pos     -1.5183843203 -5.0564040817
## FC_Mpos_TCAAGACTCTTGGAAC-1FC_MI_GFPve_pos     -1.6124840803 -4.9947292783
## FC_Mpos_TCAAGTGAGACGGATC-1FC_MI_GFPve_pos     -2.0702630110 -3.0037007012
## FC_Mpos_TCAATTCTCAGAGCAG-1FC_MI_GFPve_pos     -2.0136074848 -4.2516660966
## FC_Mpos_TCACACCAGTCTGGTT-1FC_MI_GFPve_pos     -1.7183971234 -4.3951502718
## FC_Mpos_TCACGGGTCGGTCACG-1FC_MI_GFPve_pos     -0.2140290193 -4.7015556038
## FC_Mpos_TCATCCGCAGGACATG-1FC_MI_GFPve_pos     -1.8020123072 -3.9840644874
## FC_Mpos_TCATGCCGTTGCCAAT-1FC_MI_GFPve_pos     -1.3782187529 -3.5004707255
## FC_Mpos_TCCACCATCCACAGGC-1FC_MI_GFPve_pos     -1.0857338734 -4.9430158861
## FC_Mpos_TCCACGTGTAATGCTC-1FC_MI_GFPve_pos     -1.7734908886 -4.1373756208
## FC_Mpos_TCCATGCGTACCTTCC-1FC_MI_GFPve_pos     -1.8650854655 -3.9613066591
## FC_Mpos_TCCGATCAGTAAACAC-1FC_MI_GFPve_pos     -1.4410065241 -4.7940719888
## FC_Mpos_TCGACCTAGCCATTGT-1FC_MI_GFPve_pos     -0.4106712170 -4.7114950489
## FC_Mpos_TCGGGCAAGCACTGGA-1FC_MI_GFPve_pos     -1.8813376255 -4.3443838634
## FC_Mpos_TCTAACTCACTGAGTT-1FC_MI_GFPve_pos     -1.1948036976 -5.1575957514
## FC_Mpos_TCTATACAGCTAATGA-1FC_MI_GFPve_pos     -0.7452474065 -4.1567749061
## FC_Mpos_TCTCACGTCTCCCAAC-1FC_MI_GFPve_pos     -1.7721998520 -4.8398127459
## FC_Mpos_TCTCTGGAGATGATTG-1FC_MI_GFPve_pos     -1.4890531130 -4.3487566687
## FC_Mpos_TCTGGCTCAGCTAACT-1FC_MI_GFPve_pos     -1.7147935457 -4.1321558036
## FC_Mpos_TCTGGCTTCCATAAGC-1FC_MI_GFPve_pos     -1.7657351322 -4.5803022213
## FC_Mpos_TCTTCCTTCTGGTGCG-1FC_MI_GFPve_pos     -1.0870189495 -4.2458921351
## FC_Mpos_TGACGCGGTCGGTGAA-1FC_MI_GFPve_pos     -2.0565033980 -4.5552423127
## FC_Mpos_TGACGCGTCTAGAGCT-1FC_MI_GFPve_pos     -0.5829371937 -4.8551911615
## FC_Mpos_TGACTCCAGACCAGCA-1FC_MI_GFPve_pos     -0.5305848248 -4.8495295994
## FC_Mpos_TGAGGTTAGCGTGAAC-1FC_MI_GFPve_pos      1.3958459072 -2.5895371117
## FC_Mpos_TGATCTTAGGTATAGT-1FC_MI_GFPve_pos      1.4865031414 -2.4806162991
## FC_Mpos_TGATGGTGTTGCCTAA-1FC_MI_GFPve_pos     -1.0709172316 -5.0177622713
## FC_Mpos_TGCGGGTGTGGACAGT-1FC_MI_GFPve_pos     -1.6507973499 -4.5681007184
## FC_Mpos_TGGATCATCGGTTCAA-1FC_MI_GFPve_pos     -4.1701567002 -2.7067815937
## FC_Mpos_TGGCGTGTCAGACTGT-1FC_MI_GFPve_pos     -1.7715288229 -3.1600964941
## FC_Mpos_TGGGAGAGTAAGGAGA-1FC_MI_GFPve_pos     -1.6941138334 -4.2504074849
## FC_Mpos_TGGGAGATCTTCCGTG-1FC_MI_GFPve_pos     -1.5956486292 -4.8392391779
## FC_Mpos_TGTAACGAGCAATAGT-1FC_MI_GFPve_pos     -0.7291995712 -5.0116547622
## FC_Mpos_TGTGTGATCGCCTCTA-1FC_MI_GFPve_pos     -1.0222126670 -5.0701532640
## FC_Mpos_TGTTCTAGTCATACCA-1FC_MI_GFPve_pos     -1.6017700262 -4.7549579224
## FC_Mpos_TGTTGAGAGCTGAAAT-1FC_MI_GFPve_pos     -1.4076374598 -5.1379443981
## FC_Mpos_TGTTGGAGTTAAGTCC-1FC_MI_GFPve_pos      1.4057493381 -1.9682006516
## FC_Mpos_TGTTTGTAGCAAATGT-1FC_MI_GFPve_pos     -2.0498999186 -3.1255930342
## FC_Mpos_TGTTTGTCAGCTTTGA-1FC_MI_GFPve_pos     -1.9404029675 -3.1161305584
## FC_Mpos_TTACCGCAGTCATACC-1FC_MI_GFPve_pos     -0.7106222935 -4.8273714982
## FC_Mpos_TTACCGCTCTATCGCC-1FC_MI_GFPve_pos     -1.7871217556 -4.1637807168
## FC_Mpos_TTACGCCAGGTTCATC-1FC_MI_GFPve_pos     -1.4605234690 -4.3841599204
## FC_Mpos_TTATTGCAGGTACATA-1FC_MI_GFPve_pos     -0.8871310778 -5.0710462012
## FC_Mpos_TTCACGCCAACAGCTT-1FC_MI_GFPve_pos     -1.9383671112 -4.0913914122
## FC_Mpos_TTCCTAAAGTTACGTC-1FC_MI_GFPve_pos     -1.2022676296 -3.9627207793
## FC_Mpos_TTCGCTGGTGAGTCAG-1FC_MI_GFPve_pos      1.4359702043 -1.8282622971
## FC_Mpos_TTCTGTATCCCATTTA-1FC_MI_GFPve_pos     -0.6394473441 -4.9555093773
## FC_Mpos_TTGGGCGAGAGCAACC-1FC_MI_GFPve_pos     -2.2354585953 -3.4720010437
## FC_Mpos_TTGGTTTTCACTTCTA-1FC_MI_GFPve_pos      0.0387775473 -4.2566224910
```

```
## FC_Mpos_TTTACCATCTTGGGCG-1FC_MI_GFPve_pos     -1.0802834459 -4.8581715502
## FC_Mpos_TTTAGTCCAGGAAGTC-1FC_MI_GFPve_pos     -1.8884473868 -3.6486385740
## FC_Mpos_TTTGATCTCGTTGTAG-1FC_MI_GFPve_pos     -1.6099943943 -5.1414533831
## FC_Mpos_TTTGGTTGTACGTGTT-1FC_MI_GFPve_pos      1.0810550623 -3.4515584387
## FC_Fpos_AAACGCTTCTAGATCG-1FC_FI_GFPve_pos      0.9473377638 -3.0985332407
## FC_Fpos_AAAGGATTCGGCCTTT-1FC_FI_GFPve_pos     -1.9533200093 -4.0363282837
## FC_Fpos_AAAGTGATCCGGTTCT-1FC_FI_GFPve_pos      0.9385373764 -3.5554489769
## FC_Fpos_AACAACCAGGTCCAGA-1FC_FI_GFPve_pos     -0.9453317352 -4.8759975113
## FC_Fpos_AACCAACAGACCATGG-1FC_FI_GFPve_pos     -1.4893671103 -4.4343121328
## FC_Fpos_AACCAACTCAGGTAAA-1FC_FI_GFPve_pos      1.6882785730 -1.8665907063
## FC_Fpos_AACCATGCACTTGAGT-1FC_FI_GFPve_pos     -2.4090621300 -3.7060478605
## FC_Fpos_AACGTCAGTGGTCCGT-1FC_FI_GFPve_pos      0.1851426534 -4.3872274138
## FC_Fpos_AACTTCTAGGGCAGTT-1FC_FI_GFPve_pos     -1.2362916298 -5.0222227253
## FC_Fpos_AACTTCTCAAATTGGA-1FC_FI_GFPve_pos     -1.6756397314 -4.9938107111
## FC_Fpos_AAGAACAAGTTAACAG-1FC_FI_GFPve_pos     -1.2644988127 -3.0787417568
## FC_Fpos_AAGAACAGTTAGGAGC-1FC_FI_GFPve_pos     -1.7030324764 -5.0050152220
## FC_Fpos_AAGACAAAGCATAGGC-1FC_FI_GFPve_pos      0.9783089332 -3.4742282309
## FC_Fpos_AAGACAAAGGCGATAC-1FC_FI_GFPve_pos     -1.6813245840 -4.4278048553
## FC_Fpos_AAGATAGGTGTTCCTC-1FC_FI_GFPve_pos     -1.8516641684 -3.7154324927
## FC_Fpos_AAGATAGTCTATCGTT-1FC_FI_GFPve_pos     -1.4815547295 -4.9497718193
## FC_Fpos_AAGCATCGTGCCGTTG-1FC_FI_GFPve_pos     -1.5901970692 -4.2616008498
## FC_Fpos_AAGCGTTTCATCCTGC-1FC_FI_GFPve_pos      1.1875928812 -3.2036550917
## FC_Fpos_AAGGTAACATATGGCT-1FC_FI_GFPve_pos     -0.5395007260 -4.6391173355
## FC_Fpos_AAGTACCGTATGTCCA-1FC_FI_GFPve_pos      0.9880100660 -3.3670688547
## FC_Fpos_AATCGTGAGTATAGAC-1FC_FI_GFPve_pos      1.0730484896 -3.6302271761
## FC_Fpos_AATGGCTGTTGGACTT-1FC_FI_GFPve_pos     -1.7509327955 -4.4276818015
## FC_Fpos_AATTCCTCAACGTTAC-1FC_FI_GFPve_pos     -1.0912053414 -3.2385741152
## FC_Fpos_ACAAAGAGTAGGTACG-1FC_FI_GFPve_pos      0.9366969042 -3.5293838658
## FC_Fpos_ACAGGGAAGCCTCATA-1FC_FI_GFPve_pos      1.0263554029 -3.6025300898
## FC_Fpos_ACCAAACTCTTCACGC-1FC_FI_GFPve_pos      0.6535769872 -3.9268609561
## FC_Fpos_ACCCAAACAAGACTGG-1FC_FI_GFPve_pos     -0.9768402405 -4.8574308880
## FC_Fpos_ACCTGAAAGGTCCTGC-1FC_FI_GFPve_pos     -0.4620103367 -4.6396838926
## FC_Fpos_ACCTGTCAGCTGAAAT-1FC_FI_GFPve_pos     -1.5903136559 -4.0962933578
## FC_Fpos_ACGCACGCACGGTGCT-1FC_FI_GFPve_pos     -1.4499839373 -5.1649952509
## FC_Fpos_ACGCACGCATATGCGT-1FC_FI_GFPve_pos     -1.7063884564 -4.6448591404
## FC_Fpos_ACGGAAGTCTTCGGTC-1FC_FI_GFPve_pos     -1.6705795355 -4.1779609598
## FC_Fpos_ACGTACACATACACCA-1FC_FI_GFPve_pos     -0.8953038283 -5.0769931652
## FC_Fpos_ACGTCCTCAGCTGGTC-1FC_FI_GFPve_pos      1.7531277113 -2.3311226048
## FC_Fpos_ACTGCAACACTGTGAT-1FC_FI_GFPve_pos     -1.8142012186 -4.7145740815
## FC_Fpos_AGACAAACAATGCAAA-1FC_FI_GFPve_pos     -1.3317050762 -4.7239646685
## FC_Fpos_AGATCGTTCGCCTTGT-1FC_FI_GFPve_pos     -1.3533074684 -3.4779662766
## FC_Fpos_AGCATCAGTACGCTAT-1FC_FI_GFPve_pos     -0.6198291309 -4.9649872311
## FC_Fpos_AGCCAATCACAGTCGC-1FC_FI_GFPve_pos     -0.8727163501 -3.8979260601
## FC_Fpos_AGCGATTCACTAGGCC-1FC_FI_GFPve_pos      0.7401859336 -2.9231377520
## FC_Fpos_AGCGCCAAGCTGCCTG-1FC_FI_GFPve_pos     -1.5847105808 -4.7116145190
## FC_Fpos_AGCGCCAAGGGCTGAT-1FC_FI_GFPve_pos      1.1715395622 -3.4573031344
## FC_Fpos_AGCGCTGCAGTCAGTT-1FC_FI_GFPve_pos      0.0694419764 -4.4480090119
## FC_Fpos_AGCTACACAGCAGTAG-1FC_FI_GFPve_pos     -1.3959292002 -2.9521858372
## FC_Fpos_AGCTTCCGTTCAAACC-1FC_FI_GFPve_pos     -1.8003257342 -3.5505984701
## FC_Fpos_AGGAAATCAGCTATAC-1FC_FI_GFPve_pos      0.7760669761 -3.7283776678
## FC_Fpos_AGGAATATCTGCGGCA-1FC_FI_GFPve_pos     -1.2457707949 -4.8374604784
## FC_Fpos_AGGCCACCACCCTTGT-1FC_FI_GFPve_pos     -1.2167513199 -5.1976506688
## FC_Fpos_AGGGCCTAGAGAGGTA-1FC_FI_GFPve_pos     -1.9228750296 -3.5741489805
## FC_Fpos_AGGGTGAGTAGCTCGC-1FC_FI_GFPve_pos     -1.5724042483 -4.1705104984
## FC_Fpos_AGGGTTTCAACCTATG-1FC_FI_GFPve_pos     -1.6621424980 -4.8638437517
## FC_Fpos_AGGTAGGTCTTCTGGC-1FC_FI_GFPve_pos      1.0153285436 -3.5982806362
## FC_Fpos_AGGTTGTCAACTTGCA-1FC_FI_GFPve_pos      1.1306129865 -3.2152986206
## FC_Fpos_AGTACCACAGGTGACA-1FC_FI_GFPve_pos     -1.5210132427 -4.1592285909
## FC_Fpos_AGTAGCTCAGCACGAA-1FC_FI_GFPve_pos     -0.5740338989 -4.8134287350
## FC_Fpos_AGTCACACACACGTGC-1FC_FI_GFPve_pos      1.7068133526 -2.3622340836
## FC_Fpos_AGTCATGAGGTGCGAT-1FC_FI_GFPve_pos     -2.1629785128  1.9744968735
## FC_Fpos_AGTCTCCCATGTAACC-1FC_FI_GFPve_pos     -0.8307010360 -4.6730038278
## FC_Fpos_AGTGTTGAGCAAATGT-1FC_FI_GFPve_pos     -1.2777223416 -4.7267616509
## FC_Fpos_ATACCTTCATCCGTGG-1FC_FI_GFPve_pos     -1.2414710350 -3.8198768772
## FC_Fpos_ATAGACCAGGTTGGTG-1FC_FI_GFPve_pos     -1.1809052057 -5.2439063467
## FC_Fpos_ATAGAGAGTCACGTGC-1FC_FI_GFPve_pos     -1.5793852635 -5.1888437845
## FC_Fpos_ATCACGACACCCTGTT-1FC_FI_GFPve_pos     -2.0418553181 -3.9449548401
## FC_Fpos_ATCCACCCAATTCGTG-1FC_FI_GFPve_pos      1.1172756128 -3.5168176569
## FC_Fpos_ATCCCTGAGCATCAGG-1FC_FI_GFPve_pos     -1.5468271799 -4.2001522459
## FC_Fpos_ATCGATGAGGGTACGT-1FC_FI_GFPve_pos      1.4478578739 -2.0014447846
```

```
## FC_Fpos_ATCGATGGTTGCTCAA-1FC_FI_GFPve_pos     -1.2122807331 -4.2284446634
## FC_Fpos_ATCGCCTAGCCGTTGC-1FC_FI_GFPve_pos     -1.4581128187 -4.9773820467
## FC_Fpos_ATCGTGATCTCCTGCA-1FC_FI_GFPve_pos     -1.5804122515 -4.0055938043
## FC_Fpos_ATCTCTAAGCGCGTTC-1FC_FI_GFPve_pos      1.4799394779 -2.1472673096
## FC_Fpos_ATCTCTAGTTAGTCGT-1FC_FI_GFPve_pos      1.5881159477 -2.6695015110
## FC_Fpos_ATGGGAGTCACTTCTA-1FC_FI_GFPve_pos     -1.9833779164 -3.8481942691
## FC_Fpos_ATGGGAGTCTTACGGA-1FC_FI_GFPve_pos     -1.2758745976 -4.2949512877
## FC_Fpos_ATTATCCTCTGGGCAC-1FC_FI_GFPve_pos     -2.1776876278 -3.5576231159
## FC_Fpos_ATTGTTCTCAACCTTT-1FC_FI_GFPve_pos     -1.7609381504 -4.3392912127
## FC_Fpos_CAAAGAAAGAATAACC-1FC_FI_GFPve_pos      0.9888326101 -3.6111080326
## FC_Fpos_CAAAGAAGTACTGCGC-1FC_FI_GFPve_pos     -1.6576579638 -4.5346276291
## FC_Fpos_CAAAGAAGTGCCGTTG-1FC_FI_GFPve_pos     -2.1176844664 -3.7447447457
## FC_Fpos_CAAAGAATCCGGTAAT-1FC_FI_GFPve_pos     -1.9930450745 -3.6640340962
## FC_Fpos_CAAGAGGAGGTTCTTG-1FC_FI_GFPve_pos      1.3401420288 -2.5345427670
## FC_Fpos_CAATGACTCAGTCACA-1FC_FI_GFPve_pos     -2.6406307049 -3.1520901598
## FC_Fpos_CACACAACAGCATTGT-1FC_FI_GFPve_pos     -1.6689369507 -4.1622908987
## FC_Fpos_CACAGGCAGAAGATCT-1FC_FI_GFPve_pos     -1.1296021767 -4.7348907605
## FC_Fpos_CAGAGCCGTCCTTAAG-1FC_FI_GFPve_pos      1.1949410610 -3.3562085308
## FC_Fpos_CAGATCATCTCACGAA-1FC_FI_GFPve_pos     -1.6033794708 -4.6262046002
## FC_Fpos_CAGCGTGCACATTCTT-1FC_FI_GFPve_pos     -2.1189561911 -3.9030151047
## FC_Fpos_CAGGTATGTCGTTGGC-1FC_FI_GFPve_pos      0.3999208979 -4.1863109745
## FC_Fpos_CATCAAGTCGGACCAC-1FC_FI_GFPve_pos     -2.1276894636 -2.7709229864
## FC_Fpos_CATCCCACAATACAGA-1FC_FI_GFPve_pos     -2.1026742287 -2.2679512181
## FC_Fpos_CATCGCTAGCATTGTC-1FC_FI_GFPve_pos     -1.9782992430 -4.1884160914
## FC_Fpos_CATCGGGTCTACGCAA-1FC_FI_GFPve_pos     -0.1867364157 -4.6892295905
## FC_Fpos_CATGCAACACTAGTAC-1FC_FI_GFPve_pos      0.8103794627 -3.7683015146
## FC_Fpos_CATGCCTCATAATGCC-1FC_FI_GFPve_pos     -1.3784215279 -4.2793478049
## FC_Fpos_CATGCGGAGATTAGCA-1FC_FI_GFPve_pos      0.6787227564 -3.5672671475
## FC_Fpos_CATGCTCTCTGAATGC-1FC_FI_GFPve_pos     -1.6660531588 -4.9825100429
## FC_Fpos_CATTCATGTGGTCCCA-1FC_FI_GFPve_pos      0.7676350050 -3.2270828165
## FC_Fpos_CATTCATTCACCACAA-1FC_FI_GFPve_pos     -1.7148896284 -4.4266292133
## FC_Fpos_CATTGTTTCAAGCCCG-1FC_FI_GFPve_pos     -1.1280023880 -4.8194686450
## FC_Fpos_CATTGTTTCTATTGTC-1FC_FI_GFPve_pos     -0.2765639402 -4.7834808320
## FC_Fpos_CCACAAAGTCTCCCTA-1FC_FI_GFPve_pos     -1.8619847126 -4.2808851577
## FC_Fpos_CCCTAACTCAAGCCGC-1FC_FI_GFPve_pos     -1.7297613449 -4.8826451726
## FC_Fpos_CCCTCTCGTGCCCGTA-1FC_FI_GFPve_pos     -1.3300373383 -5.0125063040
## FC_Fpos_CCGCAAGAGGTAGGCT-1FC_FI_GFPve_pos     -0.1457951512 -4.6515786179
## FC_Fpos_CCGGACACAACCGCCA-1FC_FI_GFPve_pos     -0.2916338123 -4.3054365792
## FC_Fpos_CCGTGAGCAGGACTTT-1FC_FI_GFPve_pos     -2.0368525810 -3.8452836551
## FC_Fpos_CCTATCGCACAACGTT-1FC_FI_GFPve_pos     -0.9953841396 -4.6023619600
## FC_Fpos_CCTCAACCACTCCGAG-1FC_FI_GFPve_pos     -1.6322980948 -5.0601378955
## FC_Fpos_CCTCAGTCACAATTCG-1FC_FI_GFPve_pos     -1.3134411402 -4.8264891632
## FC_Fpos_CCTGCATCACACGGTC-1FC_FI_GFPve_pos     -0.9783493109 -3.5781500019
## FC_Fpos_CCTTTGGTCAGACAAA-1FC_FI_GFPve_pos     -1.7124146052 -4.2349771537
## FC_Fpos_CGAGGCTGTGAGGAAA-1FC_FI_GFPve_pos      0.1028969757 -4.3511631765
## FC_Fpos_CGATCGGCAAACAGGC-1FC_FI_GFPve_pos     -0.4468159802 -4.8472355656
## FC_Fpos_CGATGGCAGAATAGTC-1FC_FI_GFPve_pos      1.3880884819 -2.6981513180
## FC_Fpos_CGCGTGAAGGCTAACG-1FC_FI_GFPve_pos     -1.8931441136 -4.1041531839
## FC_Fpos_CGGAATTAGGCGCTTC-1FC_FI_GFPve_pos     -1.6101381607 -3.0672133841
## FC_Fpos_CGGACACCACAGCTGC-1FC_FI_GFPve_pos     -1.3879954643 -4.4815664269
## FC_Fpos_CGGCAGTAGACATAAC-1FC_FI_GFPve_pos     -1.7723086901 -4.8048699252
## FC_Fpos_CGGGTGTGTTCGGGTC-1FC_FI_GFPve_pos     -0.9098599382 -5.0538798548
## FC_Fpos_CGTCCATCAGAGGAAA-1FC_FI_GFPve_pos      1.1503896885 -3.4661488213
## FC_Fpos_CGTGATAGTTGCGTAT-1FC_FI_GFPve_pos      1.3783556394 -2.8214025177
## FC_Fpos_CGTTAGACACCAGTTA-1FC_FI_GFPve_pos     -0.7717667051 -5.0770550646
## FC_Fpos_CTAACCCAGTGGATTA-1FC_FI_GFPve_pos     -1.5182113476 -4.8597406499
## FC_Fpos_CTAACTTAGGAGGTTC-1FC_FI_GFPve_pos     -1.1063298412 -5.1484166064
## FC_Fpos_CTAACTTCAACGCCCA-1FC_FI_GFPve_pos     -1.6910198755 -4.1789578475
## FC_Fpos_CTAAGTGAGCTCACTA-1FC_FI_GFPve_pos     -2.0520014591 -3.6954162993
## FC_Fpos_CTACCTGGTCTTACTT-1FC_FI_GFPve_pos     -1.0042352505 -5.0996629574
## FC_Fpos_CTAGACACATGTGCCG-1FC_FI_GFPve_pos     -0.3891297407 -4.7901692480
## FC_Fpos_CTCAACCGTCACGTGC-1FC_FI_GFPve_pos     -1.0985939927 -3.5922213473
## FC_Fpos_CTCCCTCCACAAGCCC-1FC_FI_GFPve_pos     -1.8738795347 -3.8926587381
## FC_Fpos_CTCCGATTCTGGAGAG-1FC_FI_GFPve_pos      0.9337461166 -3.6694867291
## FC_Fpos_CTCTCAGAGGGTGGGA-1FC_FI_GFPve_pos      1.2210815124 -3.1395890393
## FC_Fpos_CTGAATGCATAGTCAC-1FC_FI_GFPve_pos     -1.9948832817 -4.0789943017
## FC_Fpos_CTGAATGTCGGTAGAG-1FC_FI_GFPve_pos      1.1252137594 -3.1423619665
## FC_Fpos_CTGAGCGCAGACCTAT-1FC_FI_GFPve_pos     -1.3311995096 -4.8891800918
## FC_Fpos_CTGAGCGGTAGCCAGA-1FC_FI_GFPve_pos     -1.2671924658 -4.4370432474
## FC_Fpos_CTGATCCGTGACTGTT-1FC_FI_GFPve_pos     -0.8186129160 -4.8183599986
```

```
## FC_Fpos_CTGCAGGAGAGGCTGT-1FC_FI_GFPve_pos     -0.3506957419 -4.5188623049
## FC_Fpos_CTGCAGGTCTTAGCAG-1FC_FI_GFPve_pos     -1.0753182597 -4.9515137799
## FC_Fpos_CTGCATCAGGTTACAA-1FC_FI_GFPve_pos     -1.7468813486 -4.6097232528
## FC_Fpos_CTGCATCGTACGACAG-1FC_FI_GFPve_pos     -1.4728299208 -5.1349874117
## FC_Fpos_CTGCCTAAGCGTTGTT-1FC_FI_GFPve_pos     -1.6536961622 -5.0459850408
## FC_Fpos_CTGTGAACAAGTGCTT-1FC_FI_GFPve_pos     -1.4778030939 -4.3616181292
## FC_Fpos_CTTACCGTCCCGTAAA-1FC_FI_GFPve_pos     -1.8211554117 -3.5197083630
## FC_Fpos_CTTAGGATCATCACTT-1FC_FI_GFPve_pos     -1.7126734085 -4.7239776101
## FC_Fpos_CTTCGGTAGCGATGGT-1FC_FI_GFPve_pos     -1.7302454539 -4.3913151302
## FC_Fpos_CTTTCAACAGTTCTAG-1FC_FI_GFPve_pos     -1.7638015576 -4.5628736891
## FC_Fpos_GAAACCTTCCTATTGT-1FC_FI_GFPve_pos     -2.8043906517 -2.3618399777
## FC_Fpos_GAACGTTGTATTGACC-1FC_FI_GFPve_pos     -1.5618799753 -5.1464736141
## FC_Fpos_GAAGTAAGTTAAACCC-1FC_FI_GFPve_pos      0.1020556860 -4.1504845895
## FC_Fpos_GACACGCGTACCCGAC-1FC_FI_GFPve_pos      1.1283009224 -3.1907486595
## FC_Fpos_GACCCAGGTTCGGGTC-1FC_FI_GFPve_pos     -1.7494000025 -4.7518964788
## FC_Fpos_GACCCTTGTCAGTCCG-1FC_FI_GFPve_pos      1.4058437519 -2.0318853535
## FC_Fpos_GACGTTATCTGCTTAT-1FC_FI_GFPve_pos     -1.3923185892 -5.0994510628
## FC_Fpos_GACTTCCCACTCCGGA-1FC_FI_GFPve_pos     -1.0673950739 -4.8462998174
## FC_Fpos_GACTTCCTCTACCACC-1FC_FI_GFPve_pos     -0.6492063232 -4.8079315551
## FC_Fpos_GAGAAATCAAGCGCTC-1FC_FI_GFPve_pos     -0.4530850775 -4.5842818149
## FC_Fpos_GAGAAATGTACGAGTG-1FC_FI_GFPve_pos     -1.0579683609 -4.9887928046
## FC_Fpos_GAGACTTGTCGGTGAA-1FC_FI_GFPve_pos      1.4866644316 -2.2798702397
## FC_Fpos_GAGAGGTGTGAGCGAT-1FC_FI_GFPve_pos     -0.9225128122 -4.8744558074
## FC_Fpos_GAGGGTAGTAAGGAGA-1FC_FI_GFPve_pos     -1.7511598892 -4.5159155853
## FC_Fpos_GAGTCATTCTCGACGG-1FC_FI_GFPve_pos     -1.7849888630 -3.9788340606
## FC_Fpos_GAGTTGTTCATGCCCT-1FC_FI_GFPve_pos      1.1637172870 -3.3624833502
## FC_Fpos_GATCACACACCCTAGG-1FC_FI_GFPve_pos     -1.6572650500 -4.8704176076
## FC_Fpos_GATCAGTGTTCCTACC-1FC_FI_GFPve_pos     -2.0712245531 -4.5045082398
## FC_Fpos_GATCCCTAGGTATCTC-1FC_FI_GFPve_pos      1.1650779419 -3.3331529774
## FC_Fpos_GATGGAGCACGCAAAG-1FC_FI_GFPve_pos     -1.7704474755 -3.9198403634
## FC_Fpos_GCAACCGGTCGTGCCA-1FC_FI_GFPve_pos      1.4611975126 -2.3307547249
## FC_Fpos_GCACTAACAGATACTC-1FC_FI_GFPve_pos     -1.0604559012 -5.0157415249
## FC_Fpos_GCCAGGTAGAAGATCT-1FC_FI_GFPve_pos     -1.2276557512 -4.1207368054
## FC_Fpos_GCCATGGTCTACACTT-1FC_FI_GFPve_pos     -1.5673427410 -4.2929024376
## FC_Fpos_GCCCGAAAGCACTAGG-1FC_FI_GFPve_pos     -0.7762318678 -4.8288242124
## FC_Fpos_GCCGATGAGGAACGCT-1FC_FI_GFPve_pos     -1.1152400203 -4.9640077181
## FC_Fpos_GCCGTGATCATTACGG-1FC_FI_GFPve_pos     -0.7390948959 -5.1851421870
## FC_Fpos_GCTACAAGTCGTTATG-1FC_FI_GFPve_pos     -1.4359862633 -4.2169508614
## FC_Fpos_GCTGAATAGTCTTCGA-1FC_FI_GFPve_pos     -1.7038423844 -4.0143875756
## FC_Fpos_GCTGCAGAGCCTTTCC-1FC_FI_GFPve_pos     -1.7037663288 -4.2249424733
## FC_Fpos_GCTGCAGTCTCTCGCA-1FC_FI_GFPve_pos     -1.8419704266 -4.1253478803
## FC_Fpos_GCTTCACGTACGTGTT-1FC_FI_GFPve_pos     -1.5484971829 -5.0173095144
## FC_Fpos_GGACGTCAGAACCGCA-1FC_FI_GFPve_pos     -1.5515291519 -4.9734940209
## FC_Fpos_GGAGATGCATCCAACA-1FC_FI_GFPve_pos     -2.1268995829 -2.1717123665
## FC_Fpos_GGAGCAACAAGTGTCT-1FC_FI_GFPve_pos     -2.6510319538 -3.6014403977
## FC_Fpos_GGGAAGTGTGAGATCG-1FC_FI_GFPve_pos      1.3425406389 -2.7114393391
## FC_Fpos_GGGACCTGTGTGTGTT-1FC_FI_GFPve_pos     -1.7544953652 -4.6414582782
## FC_Fpos_GGGAGATGTTGTTGAC-1FC_FI_GFPve_pos     -1.2503784723 -4.7655216321
## FC_Fpos_GGGATCCTCGCAGTCG-1FC_FI_GFPve_pos     -1.0308117695 -5.0869283952
## FC_Fpos_GGGATGAGTGTCTCCT-1FC_FI_GFPve_pos     -0.8364644475 -5.3067018069
## FC_Fpos_GGGCTACAGGACAACC-1FC_FI_GFPve_pos     -1.7759688921 -3.8198741950
## FC_Fpos_GGGTAGACAAAGGCAC-1FC_FI_GFPve_pos     -1.7237627335 -4.0556835212
## FC_Fpos_GGGTCTGGTACGAGCA-1FC_FI_GFPve_pos     -1.2566425629 -4.9452800132
## FC_Fpos_GGTAATCAGTTGCTGT-1FC_FI_GFPve_pos      1.1186617784 -3.4391797461
## FC_Fpos_GGTAGAGTCGAGAGCA-1FC_FI_GFPve_pos     -2.0431015320 -4.4477324166
## FC_Fpos_GGTCTGGAGAAGTCCG-1FC_FI_GFPve_pos     -1.9906872339 -4.4582598187
## FC_Fpos_GGTGATTGTAAGTTGA-1FC_FI_GFPve_pos      1.0854153805 -1.6784839310
## FC_Fpos_GGTGTCGAGCACCCAC-1FC_FI_GFPve_pos     -0.3579342194 -4.7025379405
## FC_Fpos_GGTTGTATCGTAGCTA-1FC_FI_GFPve_pos     -1.8969877787 -3.5257299818
## FC_Fpos_GTAACCATCATGAGGG-1FC_FI_GFPve_pos     -1.9813706703 -3.8903496541
## FC_Fpos_GTAAGTCAGGTTCATC-1FC_FI_GFPve_pos      1.0180699997 -3.5041190066
## FC_Fpos_GTAATCGCAGCACGAA-1FC_FI_GFPve_pos     -1.7700321503 -4.0743735112
## FC_Fpos_GTACAACCACACCGAC-1FC_FI_GFPve_pos     -1.2120030947 -4.5546384283
## FC_Fpos_GTACAACTCAGTGGGA-1FC_FI_GFPve_pos      0.6394086652 -3.9110208430
## FC_Fpos_GTACAGTTCTTCGTGC-1FC_FI_GFPve_pos     -2.1821219750 -3.4977446474
## FC_Fpos_GTAGAAATCGTTTACT-1FC_FI_GFPve_pos     -1.0566422768 -4.6677808590
## FC_Fpos_GTAGGTTCATCTATCT-1FC_FI_GFPve_pos      0.9130243234 -3.5443727650
## FC_Fpos_GTATTGGGTGCTGATT-1FC_FI_GFPve_pos     -4.0618178673 -2.5107166447
## FC_Fpos_GTATTTCCAGTATGAA-1FC_FI_GFPve_pos     -0.3315666146 -4.7330321014
## FC_Fpos_GTCAAGTCAACTTCTT-1FC_FI_GFPve_pos     -1.9135268755 -3.2497269787
```

```
## FC_Fpos_GTCCACTCATCTCGTC-1FC_FI_GFPve_pos     -1.1126965709 -3.3383150973
## FC_Fpos_GTCCACTTCATTGAGC-1FC_FI_GFPve_pos      1.0061704092 -3.5831796803
## FC_Fpos_GTCCACTTCCCTTCCC-1FC_FI_GFPve_pos      0.9526406698 -3.6813672461
## FC_Fpos_GTCCCATCAATGAGCG-1FC_FI_GFPve_pos     -1.4558027811 -4.2673503734
## FC_Fpos_GTCCTCAGTCATACCA-1FC_FI_GFPve_pos     -1.8955377169 -3.9723695196
## FC_Fpos_GTCGCGACATCCTTGC-1FC_FI_GFPve_pos     -1.0219805069 -2.6220276036
## FC_Fpos_GTCTGTCTCTTGTGCC-1FC_FI_GFPve_pos     -1.6887043543 -4.1444390096
## FC_Fpos_GTGAGTTAGTGATTCC-1FC_FI_GFPve_pos     -1.4435437746 -5.0305931546
## FC_Fpos_GTGAGTTGTAATCAGA-1FC_FI_GFPve_pos      0.1454090946 -3.6142749466
## FC_Fpos_GTGCTTCAGTTCGGTT-1FC_FI_GFPve_pos     -1.4970402546 -4.4548971333
## FC_Fpos_GTGTGATTCTCAGTCC-1FC_FI_GFPve_pos      1.2746107750 -3.0331629194
## FC_Fpos_GTTCGCTAGACCAAGC-1FC_FI_GFPve_pos      1.1395995788 -3.4257981457
## FC_Fpos_GTTCGCTCACACAGCC-1FC_FI_GFPve_pos     -1.5901022978 -4.2706627824
## FC_Fpos_GTTCTATAGTGGGAAA-1FC_FI_GFPve_pos      0.8696656399 -3.9059800185
## FC_Fpos_GTTGCTCAGCATGCAG-1FC_FI_GFPve_pos     -1.5407455988 -5.0236217357
## FC_Fpos_TAATCTCAGCAGGTCA-1FC_FI_GFPve_pos      0.9765374832 -3.6784054913
## FC_Fpos_TACAACGGTCGCGGTT-1FC_FI_GFPve_pos     -1.6601817436  2.5679121338
## FC_Fpos_TACAGGTCAGATACCT-1FC_FI_GFPve_pos     -1.6045435496 -4.4922221728
## FC_Fpos_TACAGGTGTTGCAAGG-1FC_FI_GFPve_pos     -1.9356667824 -4.0488729157
## FC_Fpos_TACCTGCAGGGCAGAG-1FC_FI_GFPve_pos     -1.5746223755 -4.8309455939
## FC_Fpos_TACGGTAGTAACTGCT-1FC_FI_GFPve_pos     -1.6102874108  2.5697117172
## FC_Fpos_TATACCTCACAATTCG-1FC_FI_GFPve_pos     -1.7448323794 -4.1658177652
## FC_Fpos_TATCAGGGTAATTGGA-1FC_FI_GFPve_pos     -2.0085577793 -4.4763959326
## FC_Fpos_TATGTTCCAATAACCC-1FC_FI_GFPve_pos     -1.5797513552 -4.9990521647
## FC_Fpos_TATGTTCCAATGTCAC-1FC_FI_GFPve_pos      1.0877177887 -3.1643348612
## FC_Fpos_TATTGCTAGACGCAGT-1FC_FI_GFPve_pos     -1.7038528748 -4.2529800512
## FC_Fpos_TATTGCTCAAACTCTG-1FC_FI_GFPve_pos     -1.8834241457 -3.8930580296
## FC_Fpos_TCAAGACTCTTCCACG-1FC_FI_GFPve_pos     -0.8096663780 -3.3071382918
## FC_Fpos_TCAATCTCAATCGAAA-1FC_FI_GFPve_pos      0.8900301628 -3.6085160650
## FC_Fpos_TCACATTGTACTCGAT-1FC_FI_GFPve_pos     -1.6950616665 -4.0002134122
## FC_Fpos_TCACGGGGTTATGTGC-1FC_FI_GFPve_pos      0.1347982459 -4.3497988857
## FC_Fpos_TCACTCGGTCGAGTTT-1FC_FI_GFPve_pos      1.1937335901 -2.5682120003
## FC_Fpos_TCAGGGCTCTATTGTC-1FC_FI_GFPve_pos     -1.9224817581 -3.4966625847
## FC_Fpos_TCATGAGGTTTACTGG-1FC_FI_GFPve_pos     -1.1095425315 -4.8785208441
## FC_Fpos_TCATTGTGTCAGTCTA-1FC_FI_GFPve_pos     -0.6220044322 -4.8638151415
## FC_Fpos_TCCGTGTCAAATACAG-1FC_FI_GFPve_pos     -0.1249356992 -4.6949022241
## FC_Fpos_TCCTAATCATGAATCC-1FC_FI_GFPve_pos     -1.4725628920 -4.4090843476
## FC_Fpos_TCGAACAGTGAGATCG-1FC_FI_GFPve_pos      1.2619238071 -3.1985388674
## FC_Fpos_TCGAAGTGTCGCAGTC-1FC_FI_GFPve_pos     -1.7305009194 -4.5617853381
## FC_Fpos_TCGACCTAGAAACTGT-1FC_FI_GFPve_pos     -1.1375567742 -3.9544750728
## FC_Fpos_TCGCACTAGTCACAGG-1FC_FI_GFPve_pos     -1.6301066704 -4.7309759731
## FC_Fpos_TCGCTCAGTGGTCTCG-1FC_FI_GFPve_pos     -0.9788587518 -5.0373781301
## FC_Fpos_TCGGGCACAGCAGTGA-1FC_FI_GFPve_pos     -4.0432679482 -2.4795400776
## FC_Fpos_TCGGTCTCATACACCA-1FC_FI_GFPve_pos     -1.2054821081 -5.1218079604
## FC_Fpos_TCGTAGATCTCGTCAC-1FC_FI_GFPve_pos     -0.5711517759 -4.8594470151
## FC_Fpos_TCTACATGTACGTTCA-1FC_FI_GFPve_pos     -1.5125869341 -4.7388976732
## FC_Fpos_TCTACCGCATGGGCAA-1FC_FI_GFPve_pos     -1.0480260081 -4.1732892670
## FC_Fpos_TCTATCACAGCACGAA-1FC_FI_GFPve_pos     -0.6327325530 -3.9660551228
## FC_Fpos_TCTGTCGAGACATAGT-1FC_FI_GFPve_pos     -0.0019308157 -4.4925786384
## FC_Fpos_TCTTAGTCAAAGACGC-1FC_FI_GFPve_pos      0.1694409721 -4.3878499247
## FC_Fpos_TGAGACTGTGCATGTT-1FC_FI_GFPve_pos     -1.9615921803 -3.7715184964
## FC_Fpos_TGAGGTTGTCATTCCC-1FC_FI_GFPve_pos     -1.3914434738 -5.2240193762
## FC_Fpos_TGAGTCACAATACGCT-1FC_FI_GFPve_pos     -1.8502219744 -4.3433597066
## FC_Fpos_TGCGGCATCGGCTGTG-1FC_FI_GFPve_pos     -2.3913538285 -3.1148249783
## FC_Fpos_TGCTCGTCAAGAAATC-1FC_FI_GFPve_pos     -1.6089392729 -5.0157924272
## FC_Fpos_TGCTTCGAGTTGTACC-1FC_FI_GFPve_pos     -1.9454188175 -3.5955898442
## FC_Fpos_TGGCGTGAGGCACAAC-1FC_FI_GFPve_pos     -1.5007390804 -4.7005892225
## FC_Fpos_TGGTACATCCATGAGT-1FC_FI_GFPve_pos      1.0679218941 -3.4797857680
## FC_Fpos_TGTAACGTCTGAGAGG-1FC_FI_GFPve_pos     -1.0429471560 -4.9194650777
## FC_Fpos_TGTAGACGTCCAGTTA-1FC_FI_GFPve_pos     -2.1167005129 -3.6122171559
## FC_Fpos_TGTCAGAAGAGCCATG-1FC_FI_GFPve_pos     -1.3764158316 -3.1373092808
## FC_Fpos_TGTCCTGTCCATTGGA-1FC_FI_GFPve_pos     -1.5931497641 -4.6208590247
## FC_Fpos_TGTGTGACAAGCCTGC-1FC_FI_GFPve_pos     -0.9919020362 -4.6689991929
## FC_Fpos_TGTTGAGAGGACAAGA-1FC_FI_GFPve_pos     -1.6515496798 -4.1292689122
## FC_Fpos_TGTTTGTTCTGGGATT-1FC_FI_GFPve_pos      1.2488267593 -2.9869172253
## FC_Fpos_TTACTGTCAGAATCGG-1FC_FI_GFPve_pos      1.5550787620 -2.2418081440
## FC_Fpos_TTCAATCGTATTTCCT-1FC_FI_GFPve_pos     -1.2789350577 -2.8973471798
## FC_Fpos_TTCACGCCATAGATGA-1FC_FI_GFPve_pos      1.0887361936 -3.3861705937
## FC_Fpos_TTCAGGACAGGACTAG-1FC_FI_GFPve_pos     -1.0203749366 -5.1812607028
## FC_Fpos_TTCATTGTCTGTCGTC-1FC_FI_GFPve_pos     -2.0083800383 -3.5525675692
```

```
## FC_Fpos_TTCCAATCATGGAACG-1FC_FI_GFPve_pos     -2.1383583374 -3.9740335025
## FC_Fpos_TTCCTCTAGCACTCGC-1FC_FI_GFPve_pos     -1.6902700491 -4.1769958414
## FC_Fpos_TTCGCTGAGCTGGAGT-1FC_FI_GFPve_pos     -1.0987694688 -4.5748998948
## FC_Fpos_TTCTGTACAGAAGCGT-1FC_FI_GFPve_pos     -2.0449036188 -2.9278137364
## FC_Fpos_TTGCTGCGTGAAAGTT-1FC_FI_GFPve_pos     -1.3752161093 -4.4994023718
## FC_Fpos_TTTATGCTCTGGCTGG-1FC_FI_GFPve_pos     -1.6965750284 -3.7653038778
## FC_Fpos_TTTCATGGTGATATAG-1FC_FI_GFPve_pos     -0.4491770692 -4.8790801741
## FC_Fpos_TTTCCTCTCGAGCCTG-1FC_FI_GFPve_pos     -1.0790661998 -4.0778605022
## FC_Fpos_TTTGATCCATGCCATA-1FC_FI_GFPve_pos      0.7574896388 -3.7632413782
## dCA2_Mpos_AAAGAACTCCCGGTAG-1dCA2_MI_GFPve_pos -0.3154399373 -2.8151778855
## dCA2_Mpos_AACAACCAGGTCCGAA-1dCA2_MI_GFPve_pos  2.1693527870 -2.1822845616
## dCA2_Mpos_AACCACATCACTGCTC-1dCA2_MI_GFPve_pos  1.9092054539 -2.3406800904
## dCA2_Mpos_AACCATGCAAGCGCTC-1dCA2_MI_GFPve_pos  0.9115026169 -4.3645549692
## dCA2_Mpos_AAGACAATCAAATGCC-1dCA2_MI_GFPve_pos  2.3540644817 -1.5450176872
## dCA2_Mpos_ACATCCCTCACCTTGC-1dCA2_MI_GFPve_pos  2.3773851566 -1.6467346825
## dCA2_Mpos_ACATCCCTCCGTGACG-1dCA2_MI_GFPve_pos  2.5404658489 -1.5902027764
## dCA2_Mpos_ACCAAACCACGCTGCA-1dCA2_MI_GFPve_pos  2.2154939346 -1.9200234093
## dCA2_Mpos_ACCAACACAACCGATT-1dCA2_MI_GFPve_pos  0.5413430505 -3.1831559815
## dCA2_Mpos_AGACACTAGTCTAGCT-1dCA2_MI_GFPve_pos  1.4234844618 -2.6535916008
## dCA2_Mpos_AGACCATTCAAGCCGC-1dCA2_MI_GFPve_pos  1.2321462803 -2.2229153790
## dCA2_Mpos_AGACCATTCTACTGCC-1dCA2_MI_GFPve_pos  1.9472396545 -1.5320450939
## dCA2_Mpos_AGATCCATCAGTGTCA-1dCA2_MI_GFPve_pos  1.8510265522 -1.9911551155
## dCA2_Mpos_AGTCAACCAGCTTCGG-1dCA2_MI_GFPve_pos  0.6941825919 -3.3124712385
## dCA2_Mpos_AGTGATCCAGATGCGA-1dCA2_MI_GFPve_pos  2.0446248226 -2.1684636749
## dCA2_Mpos_ATCATTCAGCTGACTT-1dCA2_MI_GFPve_pos  1.2118983440 -2.4261319317
## dCA2_Mpos_ATCGCCTAGCCTCTCT-1dCA2_MI_GFPve_pos  1.8953261547 -2.2505991139
## dCA2_Mpos_ATGACCACATCCTTGC-1dCA2_MI_GFPve_pos  1.4355600052 -2.0678007282
## dCA2_Mpos_CACGTTCCAGACGCTC-1dCA2_MI_GFPve_pos  0.9627336435 -2.6123177685
## dCA2_Mpos_CCTCCAAAGCCATCCG-1dCA2_MI_GFPve_pos  1.8545999698 -2.2128383793
## dCA2_Mpos_CTAGACAGTGGTCTAT-1dCA2_MI_GFPve_pos  1.9986798935 -2.2667553105
## dCA2_Mpos_CTCACTGTCGGCTTCT-1dCA2_MI_GFPve_pos  1.3015241794 -2.2861771263
## dCA2_Mpos_CTCCACACAAGTTCCA-1dCA2_MI_GFPve_pos  1.9665184192 -1.8471793808
## dCA2_Mpos_GAACGTTAGATGAATC-1dCA2_MI_GFPve_pos  1.0984865598 -2.5443117298
## dCA2_Mpos_GAACTGTTCTACTCAT-1dCA2_MI_GFPve_pos  1.3742733173 -2.4514548458
## dCA2_Mpos_GAAGCGACACCCATAA-1dCA2_MI_GFPve_pos  1.8021968775 -1.7020792641
## dCA2_Mpos_GACCAATGTCGACTTA-1dCA2_MI_GFPve_pos  1.9297034912 -2.3082382359
## dCA2_Mpos_GACGTTAAGGAAACGA-1dCA2_MI_GFPve_pos  0.7410680823 -4.1715241946
## dCA2_Mpos_GACTCAACAACTGGTT-1dCA2_MI_GFPve_pos  2.1748421364 -1.9653477349
## dCA2_Mpos_GATGCTAAGCTCACTA-1dCA2_MI_GFPve_pos  1.8990638428 -2.2097668328
## dCA2_Mpos_GCACGTGCACAGTACT-1dCA2_MI_GFPve_pos  0.8944553308 -4.3026882686
## dCA2_Mpos_GCAGCCACATGACTAC-1dCA2_MI_GFPve_pos  2.4681413345 -1.7968339600
## dCA2_Mpos_GCAGCTGCACAACCGC-1dCA2_MI_GFPve_pos  2.5580454044 -1.7157075085
## dCA2_Mpos_GCCAGCATCCTCTCTT-1dCA2_MI_GFPve_pos  2.0880851917 -2.1313504852
## dCA2_Mpos_GGTTAACTCCATTCAT-1dCA2_MI_GFPve_pos  1.8650729828 -0.8627133049
## dCA2_Mpos_GTTTACTCACTGCGTG-1dCA2_MI_GFPve_pos  1.6131050758 -2.3100926556
## dCA2_Mpos_TACCCACAGTCACTGT-1dCA2_MI_GFPve_pos  0.8421074323  1.0798349701
## dCA2_Mpos_TACGCTCTCCATGCAA-1dCA2_MI_GFPve_pos  1.1863721781 -2.6928057350
## dCA2_Mpos_TCATACTCATCCTGTC-1dCA2_MI_GFPve_pos  2.1329798870 -2.0398397125
## dCA2_Mpos_TCATTACCAACTGGTT-1dCA2_MI_GFPve_pos  1.7043961458 -1.6678368725
## dCA2_Mpos_TCCGAAACAGTGTACT-1dCA2_MI_GFPve_pos  1.7017139368 -2.3114070572
## dCA2_Mpos_TCCGATCAGTGTCATC-1dCA2_MI_GFPve_pos  2.0652506523 -2.1860255875
## dCA2_Mpos_TCGCTCACACAGCCAC-1dCA2_MI_GFPve_pos -0.1791086435 -2.5207416691
## dCA2_Mpos_TCTATCATCATCACAG-1dCA2_MI_GFPve_pos  2.1740098171 -2.0245289482
## dCA2_Mpos_TGAACGTAGTGGTTAA-1dCA2_MI_GFPve_pos -0.1904738523 -3.0819135584
## dCA2_Mpos_TGATGCACATGACTGT-1dCA2_MI_GFPve_pos -0.0749552600 -3.4391382612
## dCA2_Mpos_TGGGAAGTCGTGGACC-1dCA2_MI_GFPve_pos  1.4025594168 -2.6114427723
## dCA2_Mpos_TGGTTAGAGAGCATTA-1dCA2_MI_GFPve_pos  2.3132052593 -1.8638262428
## dCA2_Mpos_TTTATGCTCATGCCCT-1dCA2_MI_GFPve_pos  2.1328766517 -1.4493772663
## dCA2_Fpos_CCGATGGAGCCTAACT-1dCA2_FI_GFPve_pos  1.8441641502 -2.3540911354
## dCA2_Fpos_CTGCGAGGTGTTGCCG-1dCA2_FI_GFPve_pos  1.3747354917 -2.6899923958
## dCA2_Fpos_GAAGAATAGCACTCTA-1dCA2_FI_GFPve_pos  1.3451245003 -2.2980725445
## dCA2_Fpos_TACATTCGTCGTGGAA-1dCA2_FI_GFPve_pos  0.7018528752 -2.2389626183
## FC_Mpos_AAACGCTCAATAGTGA-1FC_MI_GFPve_pos      0.6599128418 -4.2345547952
## FC_Mpos_AAAGAACGTCTCAAGT-1FC_MI_GFPve_pos      0.2977963559 -3.9640097298
## FC_Mpos_AAAGGGCTCTGCTCTG-1FC_MI_GFPve_pos      0.7714492850 -3.9584133305
## FC_Mpos_AACACACGTTCCTAAG-1FC_MI_GFPve_pos      0.6772573762 -4.3427252151
## FC_Mpos_AACCAACAGGTAGACC-1FC_MI_GFPve_pos      2.0238461666 -1.4642553009
## FC_Mpos_AACCAACTCTCAGAAC-1FC_MI_GFPve_pos     -0.1583439257 -3.8740622081
## FC_Mpos_AACCATGTCTTCTGGC-1FC_MI_GFPve_pos     -0.1970171198 -4.5217864789
## FC_Mpos_AACTTCTAGTCTTCCC-1FC_MI_GFPve_pos      0.5478012733 -4.3404266216
```

```
## FC_Mpos_AACTTCTGTGTCCTAA-1FC_MI_GFPve_pos      0.6736867003 -3.8793584980
## FC_Mpos_AAGTACCCATGCTGCG-1FC_MI_GFPve_pos      0.8408092432 -3.2473961987
## FC_Mpos_AATCACGAGTGGATTA-1FC_MI_GFPve_pos      0.4492067866 -4.2526960351
## FC_Mpos_AATGAAGCATCCGTGG-1FC_MI_GFPve_pos     -0.0544895060 -3.9990042724
## FC_Mpos_AATGAAGTCATCCTAT-1FC_MI_GFPve_pos      1.2830647402 -1.9745273270
## FC_Mpos_AATGACCAGGTAAAGG-1FC_MI_GFPve_pos      0.5530843906 -3.5701979078
## FC_Mpos_ACAAAGAGTGGCTGAA-1FC_MI_GFPve_pos     -0.3081048705 -4.3094089009
## FC_Mpos_ACAACCACATATCTGG-1FC_MI_GFPve_pos      2.0258088283 -1.7074012436
## FC_Mpos_ACACAGTCATAGGAGC-1FC_MI_GFPve_pos     -0.0022997536 -3.8537709393
## FC_Mpos_ACAGGGAGTTACCTTT-1FC_MI_GFPve_pos      1.9484765701 -2.1400417961
## FC_Mpos_ACAGGGAGTTCAAGTC-1FC_MI_GFPve_pos      0.4589786701 -3.9786901154
## FC_Mpos_ACATCGAGTATGAAGT-1FC_MI_GFPve_pos     -0.2203707598 -4.5240135469
## FC_Mpos_ACATTTCGTACAGAAT-1FC_MI_GFPve_pos      2.2222955398 -2.0275666393
## FC_Mpos_ACCACAACAACCGCTG-1FC_MI_GFPve_pos      1.3509650402 -2.4188720860
## FC_Mpos_ACCATTTTCCACACAA-1FC_MI_GFPve_pos      0.6571201258 -2.6548170723
## FC_Mpos_ACGGGTCCAATTCGTG-1FC_MI_GFPve_pos      0.4463928513 -4.2111383356
## FC_Mpos_ACGTACACATCACAGT-1FC_MI_GFPve_pos      0.5572043829 -4.3734062590
## FC_Mpos_ACGTAGTAGCGTATGG-1FC_MI_GFPve_pos      0.0928473346 -4.3952943958
## FC_Mpos_ACGTAGTAGGTACTGG-1FC_MI_GFPve_pos      0.3980183773 -3.7261410631
## FC_Mpos_ACGTAGTCACTCGATA-1FC_MI_GFPve_pos     -0.6812556811 -4.8600160606
## FC_Mpos_ACTCCCAGTGCGGATA-1FC_MI_GFPve_pos      0.5864802770 -3.7435716666
## FC_Mpos_ACTGTCCGTGAATTGA-1FC_MI_GFPve_pos      0.2983055286 -4.3071451165
## FC_Mpos_AGAAATGTCAGCTGAT-1FC_MI_GFPve_pos      1.3430267744 -2.7050609268
## FC_Mpos_AGAACCTAGTGGATAT-1FC_MI_GFPve_pos      1.6995154791 -1.9559290089
## FC_Mpos_AGAGCAGCAGGCGTTC-1FC_MI_GFPve_pos     -0.0314985193 -3.3598742165
## FC_Mpos_AGATAGACAACTCCAA-1FC_MI_GFPve_pos      0.3917102985 -3.8312692918
## FC_Mpos_AGATAGAGTAACCCTA-1FC_MI_GFPve_pos     -0.1514320277 -4.0504944004
## FC_Mpos_AGCGTATCATCGATCA-1FC_MI_GFPve_pos      0.2490425281 -4.6505226173
## FC_Mpos_AGCTACAAGTACCGGA-1FC_MI_GFPve_pos     -0.2492279090 -4.2737987615
## FC_Mpos_AGGAGGTCAAGCGATG-1FC_MI_GFPve_pos      0.5487567358 -3.9291082300
## FC_Mpos_AGGATCTGTGGACCTC-1FC_MI_GFPve_pos      0.5705887847 -3.7561188616
## FC_Mpos_AGGGCCTGTGCATTAC-1FC_MI_GFPve_pos      0.5165262990 -3.1986297049
## FC_Mpos_AGTCTCCAGAGGCCAT-1FC_MI_GFPve_pos     -0.7467685766 -3.9597502627
## FC_Mpos_AGTGATCGTCCTGTCT-1FC_MI_GFPve_pos      0.6690227561 -4.0323468723
## FC_Mpos_AGTGCCGAGCGATCGA-1FC_MI_GFPve_pos      0.7991718225 -4.1473449744
## FC_Mpos_AGTTCGATCATGCAGT-1FC_MI_GFPve_pos      0.5074690156 -3.7892524280
## FC_Mpos_ATCCGTCAGGGTATAT-1FC_MI_GFPve_pos      0.2168328337 -3.1667102255
## FC_Mpos_ATCGGCGAGACAGTCG-1FC_MI_GFPve_pos      0.8029635720 -3.7437629976
## FC_Mpos_ATCGTAGAGGACGCTA-1FC_MI_GFPve_pos      0.4163090758 -4.1882738151
## FC_Mpos_ATGATCGGTCCTCAGG-1FC_MI_GFPve_pos      0.2449394636 -3.8682090320
## FC_Mpos_ATGCATGGTCGCTTAA-1FC_MI_GFPve_pos      0.2319317632 -4.2916720785
## FC_Mpos_ATTACTCAGTCGCCCA-1FC_MI_GFPve_pos     -0.2167215250 -4.2879003264
## FC_Mpos_ATTCCATTCGGCTGAC-1FC_MI_GFPve_pos     -0.1880844325 -4.0819645800
## FC_Mpos_ATTCCTAAGTGAACAT-1FC_MI_GFPve_pos     -0.0732313447 -4.1925714888
## FC_Mpos_ATTCCTATCTAGACCA-1FC_MI_GFPve_pos      1.9636583500 -1.4010073818
## FC_Mpos_ATTGGGTAGTAGGGTC-1FC_MI_GFPve_pos      0.4314651661 -4.4373814441
## FC_Mpos_CAACAGTAGATGAATC-1FC_MI_GFPve_pos     -0.1672203456 -4.6688623034
## FC_Mpos_CAACCAAAGAGCCGTA-1FC_MI_GFPve_pos      0.8658719234 -3.5962218918
## FC_Mpos_CAACGGCTCTGTCCCA-1FC_MI_GFPve_pos      0.4851483874 -3.5869712509
## FC_Mpos_CAAGACTTCAGAGCGA-1FC_MI_GFPve_pos      0.8662440948 -4.0059168019
## FC_Mpos_CACACAAGTTGTGCCG-1FC_MI_GFPve_pos      0.0408625625 -4.1987495698
## FC_Mpos_CACGAATCATAAGCAA-1FC_MI_GFPve_pos     -2.6622085400 -2.7168116249
## FC_Mpos_CACTGAACAGGATTCT-1FC_MI_GFPve_pos     -0.1566654673 -4.5863543578
## FC_Mpos_CACTGAAGTGGTAATA-1FC_MI_GFPve_pos      0.2051948778 -3.5877670922
## FC_Mpos_CAGATTGAGGGAGGAC-1FC_MI_GFPve_pos      0.3127580755 -4.3989073016
## FC_Mpos_CAGCAATGTCTTTCTA-1FC_MI_GFPve_pos      2.1033818893 -2.0803341545
## FC_Mpos_CAGTTAGAGTCTTCGA-1FC_MI_GFPve_pos      0.4131588511 -4.2117775358
## FC_Mpos_CATCCACTCCGTGGGT-1FC_MI_GFPve_pos      0.4940266185 -4.2968682625
## FC_Mpos_CATCGCTTCGGTATGT-1FC_MI_GFPve_pos      0.7672686152 -3.9365198292
## FC_Mpos_CATCGGGCAATGCTCA-1FC_MI_GFPve_pos      0.0523000084 -4.4067290105
## FC_Mpos_CATGCCTCACAGCGCT-1FC_MI_GFPve_pos      1.5358195476 -2.0180251278
## FC_Mpos_CATTGAGGTTCTCCTG-1FC_MI_GFPve_pos     -0.0946487017 -4.5255456545
## FC_Mpos_CCAATTTGTAATCAAG-1FC_MI_GFPve_pos     -0.5729811914 -4.8160203003
## FC_Mpos_CCACGAGCATTGACAC-1FC_MI_GFPve_pos      0.0348217659 -4.0910258569
## FC_Mpos_CCATCACGTGCCCACA-1FC_MI_GFPve_pos      0.7058583073 -2.4934432186
## FC_Mpos_CCCATTGCATGAATCC-1FC_MI_GFPve_pos      1.9611637764 -1.3127188362
## FC_Mpos_CCCGGAATCTCCACTG-1FC_MI_GFPve_pos      0.5476981573 -2.8895140566
## FC_Mpos_CCGGGTAGTTAAACCC-1FC_MI_GFPve_pos     -0.0318586953 -4.3807504453
## FC_Mpos_CCGTTCACACTTCAGA-1FC_MI_GFPve_pos     -0.0724896319 -3.9590069331
## FC_Mpos_CCTATCGTCGAGCTGC-1FC_MI_GFPve_pos      0.5021123462 -3.7604285158
```

```
## FC_Mpos_CCTCTAGCAACCCGCA-1FC_MI_GFPve_pos      0.2116681807 -4.3684589841
## FC_Mpos_CCTCTCCAGGCCACCT-1FC_MI_GFPve_pos      0.7700320654 -4.3201803245
## FC_Mpos_CCTGTTGAGAGTTGAT-1FC_MI_GFPve_pos      0.3994583063 -4.1840765752
## FC_Mpos_CCTTCAGAGCGGTAGT-1FC_MI_GFPve_pos     -0.2096011691 -4.5893271394
## FC_Mpos_CGACAGCAGAGTCCGA-1FC_MI_GFPve_pos      0.6826740079 -3.7570125140
## FC_Mpos_CGCATAAAGTGCTAGG-1FC_MI_GFPve_pos      0.7443863325 -3.4418674387
## FC_Mpos_CGCATAAGTTGCGAAG-1FC_MI_GFPve_pos      0.6250569396 -4.2629040457
## FC_Mpos_CGTAAGTGTCCCTCAT-1FC_MI_GFPve_pos      0.7534335427 -4.0292181648
## FC_Mpos_CGTGTCTAGCGATTCT-1FC_MI_GFPve_pos      0.1894835107 -4.3279246964
## FC_Mpos_CGTGTCTCACGAGGTA-1FC_MI_GFPve_pos      0.4414634280 -4.2157858290
## FC_Mpos_CGTTCTGTCTTCTGGC-1FC_MI_GFPve_pos      0.4900264911 -3.3591869749
## FC_Mpos_CTAACTTAGATGCTTC-1FC_MI_GFPve_pos      0.5298835211 -4.1627776303
## FC_Mpos_CTACATTGTACTCGAT-1FC_MI_GFPve_pos      2.1597140007 -1.9490828194
## FC_Mpos_CTAGGTATCTCCGATC-1FC_MI_GFPve_pos      0.8257224731 -4.1504795827
## FC_Mpos_CTCATCGGTGTTACTG-1FC_MI_GFPve_pos     -0.2089670390 -4.5139142402
## FC_Mpos_CTCCACAAGCTACTAC-1FC_MI_GFPve_pos      1.9509368114 -2.2721273579
## FC_Mpos_CTCCTCCAGCAGGCAT-1FC_MI_GFPve_pos      1.6804237537 -2.4355351605
## FC_Mpos_CTGCAGGGTAGCGCCT-1FC_MI_GFPve_pos      0.6042806082 -4.3222981967
## FC_Mpos_CTGCGAGTCCATGCAA-1FC_MI_GFPve_pos      0.4090531640 -3.1297462858
## FC_Mpos_CTGGCAGGTTATTCTC-1FC_MI_GFPve_pos     -0.2096484914 -4.7201983087
## FC_Mpos_CTGTAGACAGCGGTCT-1FC_MI_GFPve_pos      0.4023849540 -4.2163392581
## FC_Mpos_CTGTAGATCTCGCAGG-1FC_MI_GFPve_pos      0.6424057655 -3.6783719935
## FC_Mpos_CTTCCGACAATCAAGA-1FC_MI_GFPve_pos      0.6168064647 -3.7316809572
## FC_Mpos_GAAGCGACAAGTGGCA-1FC_MI_GFPve_pos     -0.2858125992 -3.2345417418
## FC_Mpos_GAATAGAGTAATACCC-1FC_MI_GFPve_pos      0.7651867442 -4.0034818329
## FC_Mpos_GACACGCGTCCCGTGA-1FC_MI_GFPve_pos      0.8081848197 -2.9938647427
## FC_Mpos_GACCCTTTCTACAGGT-1FC_MI_GFPve_pos      0.6773746781 -3.5225888170
## FC_Mpos_GAGACTTGTTGACGGA-1FC_MI_GFPve_pos      0.5477332048 -4.4408576466
## FC_Mpos_GAGAGGTCATCATTTC-1FC_MI_GFPve_pos      0.8864731960 -4.1778421678
## FC_Mpos_GAGCCTGCAGGGATAC-1FC_MI_GFPve_pos      0.6884151869 -4.0334751286
## FC_Mpos_GAGTCTACATTAAAGG-1FC_MI_GFPve_pos      0.8059122138 -3.7063877262
## FC_Mpos_GAGTCTAGTCAGTCGC-1FC_MI_GFPve_pos      0.6078601055 -3.0540603079
## FC_Mpos_GAGTCTAGTCGAATGG-1FC_MI_GFPve_pos      0.4232371621 -3.3694314636
## FC_Mpos_GATCATGAGTCCCGGT-1FC_MI_GFPve_pos      0.3012611143 -4.1510566153
## FC_Mpos_GATGATCGTACTCCCT-1FC_MI_GFPve_pos      0.4013564758 -4.3013161995
## FC_Mpos_GATTCGATCATTGCCC-1FC_MI_GFPve_pos      0.2747672670 -4.1398673333
## FC_Mpos_GATTTCTAGCCTCATA-1FC_MI_GFPve_pos     -0.4646566041 -3.1984039225
## FC_Mpos_GCACTAAGTTCGGTTA-1FC_MI_GFPve_pos     -0.1206789404 -4.6350504510
## FC_Mpos_GCACTAAGTTCTGACA-1FC_MI_GFPve_pos      0.5351492934 -3.7708971776
## FC_Mpos_GCATCGGCAGCGTTTA-1FC_MI_GFPve_pos      0.7352073960 -3.5029396691
## FC_Mpos_GCATCGGCATAGATGA-1FC_MI_GFPve_pos     -0.0400068290 -4.7312298376
## FC_Mpos_GCCAGGTAGACTTCCA-1FC_MI_GFPve_pos      1.9142315559 -1.8424229302
## FC_Mpos_GCCGTGACATTATGCG-1FC_MI_GFPve_pos      0.4456753902 -4.0272997894
## FC_Mpos_GCCTGTTTCCTGCCAT-1FC_MI_GFPve_pos      2.2515082531 -2.0260221638
## FC_Mpos_GCGTTTCAGTGGCCTC-1FC_MI_GFPve_pos      0.3569660358 -4.3883231558
## FC_Mpos_GCTGGGTGTCAAAGAT-1FC_MI_GFPve_pos      0.4166565709 -4.3545258917
## FC_Mpos_GCTTGGGGTGCGGCTT-1FC_MI_GFPve_pos     -0.0994509585 -3.1056328692
## FC_Mpos_GGAGAACCAGCTGAAG-1FC_MI_GFPve_pos      0.5705510549 -3.8126401581
## FC_Mpos_GGAGCAACACTTCTCG-1FC_MI_GFPve_pos      0.4098659329 -3.3160378613
## FC_Mpos_GGAGCAAGTTATGTGC-1FC_MI_GFPve_pos      0.5589593224 -2.8945638813
## FC_Mpos_GGCACGTAGCTAGCCC-1FC_MI_GFPve_pos      0.2663938873 -4.5292704232
## FC_Mpos_GGCACGTCAACCGGAA-1FC_MI_GFPve_pos      0.6817142539 -3.7182162680
## FC_Mpos_GGCTGTGCAAGTCGTT-1FC_MI_GFPve_pos      0.2811554842 -4.2368773498
## FC_Mpos_GGGATCCGTAGCTTTG-1FC_MI_GFPve_pos      0.4599446826 -3.9241702832
## FC_Mpos_GGGCGTTAGCATGATA-1FC_MI_GFPve_pos     -0.1559559094 -3.9166258253
## FC_Mpos_GGGCGTTGTAGCGCCT-1FC_MI_GFPve_pos     -0.1502440363 -4.6409214713
## FC_Mpos_GGGTTTATCGCAGTTA-1FC_MI_GFPve_pos      0.8153116874 -3.8457900562
## FC_Mpos_GGTAATCAGAAATTCG-1FC_MI_GFPve_pos      0.1844779424 -4.3797172286
## FC_Mpos_GGTCTGGGTAGGAAAG-1FC_MI_GFPve_pos      0.8435734682 -3.9328185596
## FC_Mpos_GGTGAAGCACGAGGAT-1FC_MI_GFPve_pos      0.2365392022 -4.2696920909
## FC_Mpos_GGTGATTGTCTCCTGT-1FC_MI_GFPve_pos      0.6380684547 -4.3550957717
## FC_Mpos_GGTGTCGCACATGTTG-1FC_MI_GFPve_pos      0.1972650163 -4.4024552085
## FC_Mpos_GTAATCGCAGCTGTGC-1FC_MI_GFPve_pos     -0.0636591442 -4.5214626469
## FC_Mpos_GTACAACGTATATGGA-1FC_MI_GFPve_pos      1.7670740061 -2.1759311833
## FC_Mpos_GTCCACTTCGTTCATT-1FC_MI_GFPve_pos      2.4763195686 -1.7127632775
## FC_Mpos_GTCCCATTCGAAATCC-1FC_MI_GFPve_pos      0.5136182480 -4.2817822077
## FC_Mpos_GTCGCGAAGTTTGAGA-1FC_MI_GFPve_pos      0.7604130678 -3.8849128761
## FC_Mpos_GTCGCGACAAACCATC-1FC_MI_GFPve_pos      0.4630288653 -3.5599808373
## FC_Mpos_GTCGTTCCAGGACAGT-1FC_MI_GFPve_pos      0.5413850717 -3.2992490210
## FC_Mpos_GTCTAGATCGTTCATT-1FC_MI_GFPve_pos      0.3751825266 -3.4182456412
```

```
## FC_Mpos_GTGTCCTTCGCGCTGA-1FC_MI_GFPve_pos      0.6206734352 -4.0974803843
## FC_Mpos_GTTACGACAATAGTCC-1FC_MI_GFPve_pos      0.5358619265 -4.2399863280
## FC_Mpos_GTTCTATTCTCGTGGG-1FC_MI_GFPve_pos      0.5775745086 -4.0454238929
## FC_Mpos_GTTGAACTCCCGTTCA-1FC_MI_GFPve_pos      0.5533167891 -4.2922061063
## FC_Mpos_TAACACGTCAGGAAGC-1FC_MI_GFPve_pos      0.8905347757 -3.4639691986
## FC_Mpos_TAAGTCGGTCGCCTAG-1FC_MI_GFPve_pos      0.5389427118 -3.7838971891
## FC_Mpos_TAATCTCGTCGCAACC-1FC_MI_GFPve_pos      0.5475678615 -3.8729679264
## FC_Mpos_TACCGGGGTAACATGA-1FC_MI_GFPve_pos      0.1912481956 -4.1732628621
## FC_Mpos_TACCGGGTCTATCCAT-1FC_MI_GFPve_pos      0.5347345643 -4.1700733580
## FC_Mpos_TACCTCGGTGTCCACG-1FC_MI_GFPve_pos     -0.1115688585 -4.7022572696
## FC_Mpos_TACCTGCGTCGCTTAA-1FC_MI_GFPve_pos      0.7187193208 -2.7484066166
## FC_Mpos_TAGACCATCATCTACT-1FC_MI_GFPve_pos      0.3164206319 -4.4713265874
## FC_Mpos_TAGAGTCTCTCGGGAC-1FC_MI_GFPve_pos     -0.4310945995 -4.3057320990
## FC_Mpos_TAGGTTGAGACTAAGT-1FC_MI_GFPve_pos     -0.5623471565 -4.4059262552
## FC_Mpos_TATCGCCCAGAACTAA-1FC_MI_GFPve_pos      2.1004903488 -2.0277268566
## FC_Mpos_TATTCCAGTTGTGGCC-1FC_MI_GFPve_pos     -0.1741333358 -4.4721607246
## FC_Mpos_TATTGCTTCGTGCATA-1FC_MI_GFPve_pos      0.4851551823 -3.7571484722
## FC_Mpos_TATTGGGAGAGTTGTA-1FC_MI_GFPve_pos      0.6806231313 -2.9776521601
## FC_Mpos_TATTGGGGTATCAAGA-1FC_MI_GFPve_pos      0.5364664964 -3.0636957564
## FC_Mpos_TCACGCTCATCAGTCA-1FC_MI_GFPve_pos      0.6202179961 -3.9403646864
## FC_Mpos_TCACTATTCTGTTCAT-1FC_MI_GFPve_pos     -1.6070080824 -2.9124385990
## FC_Mpos_TCAGTGAGTCACCCTT-1FC_MI_GFPve_pos      0.1980336182 -4.7288091673
## FC_Mpos_TCATACTTCTTCCAGC-1FC_MI_GFPve_pos      2.0452292137 -2.1679908909
## FC_Mpos_TCATATCGTAGACACG-1FC_MI_GFPve_pos      0.3623866014 -3.1445690073
## FC_Mpos_TCATCCGGTGTTCCAA-1FC_MI_GFPve_pos      0.4705322556 -3.5889626421
## FC_Mpos_TCATGTTAGAGCTTTC-1FC_MI_GFPve_pos      0.5327227287 -4.4463765301
## FC_Mpos_TCATTACAGTCTACCA-1FC_MI_GFPve_pos      1.8476319485 -2.3314287342
## FC_Mpos_TCCACGTAGTCAGGGT-1FC_MI_GFPve_pos      0.3844791822 -4.3068730511
## FC_Mpos_TCCATGCAGGCACCAA-1FC_MI_GFPve_pos      0.6737849288 -3.7449919857
## FC_Mpos_TCCATGCCAACAGCCC-1FC_MI_GFPve_pos      0.3965828590 -4.3170982577
## FC_Mpos_TCCTAATGTCCAGCCA-1FC_MI_GFPve_pos      2.0171396904 -2.1701631226
## FC_Mpos_TCCTTTCAGGGCAGAG-1FC_MI_GFPve_pos      0.7237213425 -3.6512123026
## FC_Mpos_TCGACGGTCGGACGTC-1FC_MI_GFPve_pos      0.4427411132 -3.8164130129
## FC_Mpos_TCGTAGAAGCTAGTTC-1FC_MI_GFPve_pos     -0.2890555389 -3.9702208914
## FC_Mpos_TCTGCCATCAAGGAGC-1FC_MI_GFPve_pos      0.8500023060 -4.0137605943
## FC_Mpos_TCTTCCTGTTACACAC-1FC_MI_GFPve_pos      1.0714858942 -2.7130696453
## FC_Mpos_TCTTCCTGTTAGGAGC-1FC_MI_GFPve_pos      2.0508222751 -2.1758069672
## FC_Mpos_TGAATCGTCCTCAGAA-1FC_MI_GFPve_pos      0.2038675718 -4.2296338953
## FC_Mpos_TGACCCTAGTATTAGG-1FC_MI_GFPve_pos      0.5872082882 -3.4999174751
## FC_Mpos_TGATGCACAAGTGGGT-1FC_MI_GFPve_pos      0.8426233702 -4.0514212527
## FC_Mpos_TGCACGGAGAATGTTG-1FC_MI_GFPve_pos      0.3795556001 -4.3863915481
## FC_Mpos_TGCACGGGTACGAGCA-1FC_MI_GFPve_pos      0.0674670212 -4.5541143693
## FC_Mpos_TGCGACGGTAGGCAGT-1FC_MI_GFPve_pos      0.7659627371 -3.9803380050
## FC_Mpos_TGCGATACAAGGCAAC-1FC_MI_GFPve_pos      0.0929485433 -4.5301442422
## FC_Mpos_TGCTTGCTCACCCTCA-1FC_MI_GFPve_pos     -0.0527061127 -4.6308706157
## FC_Mpos_TGTAACGAGTGGCGAT-1FC_MI_GFPve_pos      0.5619331770 -3.6125132717
## FC_Mpos_TGTACAGCAGCTCTGG-1FC_MI_GFPve_pos      0.8229001932 -4.1855036892
## FC_Mpos_TGTGCGGGTCATAACC-1FC_MI_GFPve_pos      0.3119101100 -4.2921107389
## FC_Mpos_TTATTGCTCAATGCAC-1FC_MI_GFPve_pos      0.5962705188 -3.5343817152
## FC_Mpos_TTCACGCTCCATCACC-1FC_MI_GFPve_pos      0.3017289810 -4.5335625984
## FC_Mpos_TTCCACGCAGGTTACT-1FC_MI_GFPve_pos      0.6189266615 -4.0769496836
## FC_Mpos_TTCCTAACACGTCTCT-1FC_MI_GFPve_pos     -0.1221174985 -4.3957861044
## FC_Mpos_TTCGGTCCAGTTGCGC-1FC_MI_GFPve_pos     -0.0208075620 -4.5063364633
## FC_Mpos_TTCTGTAAGCGGATCA-1FC_MI_GFPve_pos      0.5018502049 -4.1363054790
## FC_Mpos_TTCTTGAGTTGATCGT-1FC_MI_GFPve_pos      0.4078664355 -4.0351703443
## FC_Mpos_TTGAACGAGGATAATC-1FC_MI_GFPve_pos      1.1938606672 -2.5537898220
## FC_Mpos_TTGACCCTCAGCTCTC-1FC_MI_GFPve_pos      0.1315495841 -4.5491638906
## FC_Mpos_TTGAGTGCATGTTTGG-1FC_MI_GFPve_pos      0.5735343270 -4.2368179836
## FC_Mpos_TTGCCTGCAGCCTTCT-1FC_MI_GFPve_pos      0.6572050624 -4.1316092290
## FC_Mpos_TTGTTGTGTATCACGT-1FC_MI_GFPve_pos      0.1129563622 -4.3222798087
## FC_Mpos_TTTAGTCGTCCGACGT-1FC_MI_GFPve_pos      0.6754141621 -4.0774024405
## FC_Mpos_TTTCAGTGTCAAATCC-1FC_MI_GFPve_pos      0.4790060692 -4.1799351253
## FC_Mpos_TTTGGTTGTGCGCTCA-1FC_MI_GFPve_pos      0.3803898864 -4.1770637907
## FC_Fpos_AAAGGATTCTAGCCTC-1FC_FI_GFPve_pos     -0.4963531621 -3.4893556513
## FC_Fpos_AAAGGTAAGTGTCATC-1FC_FI_GFPve_pos      0.2234825961 -3.8561005272
## FC_Fpos_AAAGTCCCAGAACTAA-1FC_FI_GFPve_pos      0.3157530598 -4.0413553395
## FC_Fpos_AACCCAACAAGCCTGC-1FC_FI_GFPve_pos      0.3492099814 -3.9600148477
## FC_Fpos_AACCTTTGTCTGTGGC-1FC_FI_GFPve_pos      0.2747193448 -4.2111205734
## FC_Fpos_AACGGGAAGCAGGTCA-1FC_FI_GFPve_pos      0.9081748895 -4.3760937430
## FC_Fpos_AAGGTAAAGGAACTAT-1FC_FI_GFPve_pos      0.8027959637 -4.2706538119
```

```
## FC_Fpos_AATAGAGCAACGCCCA-1FC_FI_GFPve_pos      0.6078919344 -3.9841494240
## FC_Fpos_AATGCCATCATCTATC-1FC_FI_GFPve_pos      2.2399597339 -2.0828871407
## FC_Fpos_ACACGCGTCGGTGTAT-1FC_FI_GFPve_pos      0.3819934301 -2.7257392086
## FC_Fpos_ACAGCCGCATGGTGGA-1FC_FI_GFPve_pos      0.6815807991 -3.9065550245
## FC_Fpos_ACCCTCATCTCAGGCG-1FC_FI_GFPve_pos      0.1133409910 -4.4088382401
## FC_Fpos_ACGGTTACAAGTGACG-1FC_FI_GFPve_pos      0.5826060347 -3.7982175626
## FC_Fpos_ACTGATGGTTTCGACA-1FC_FI_GFPve_pos      0.4072210960 -4.4353568234
## FC_Fpos_ACTGTGACACTGGACC-1FC_FI_GFPve_pos      0.3357703261 -4.1747684755
## FC_Fpos_ACTTTGTAGGTCGAGT-1FC_FI_GFPve_pos     -0.4166075416 -3.7831450381
## FC_Fpos_AGACCCGTCTCGGGAC-1FC_FI_GFPve_pos      0.2806899719 -3.8835166372
## FC_Fpos_AGAGCCCCAACAGCTT-1FC_FI_GFPve_pos      0.6487445407 -4.0518134512
## FC_Fpos_AGCCACGTCTTGGCTC-1FC_FI_GFPve_pos     -0.0981438436 -3.2812888302
## FC_Fpos_AGCGTATCACGTTCGG-1FC_FI_GFPve_pos      0.3288605683 -4.1583870925
## FC_Fpos_AGGATCTGTCATGGCC-1FC_FI_GFPve_pos     -0.0894846149 -4.4972812809
## FC_Fpos_AGGCCACAGTAACGAT-1FC_FI_GFPve_pos      0.6314106755 -3.6924591936
## FC_Fpos_AGGTCTAGTGACCGAA-1FC_FI_GFPve_pos      0.3522339039 -4.5486464031
## FC_Fpos_AGTCACAAGCCATTGT-1FC_FI_GFPve_pos      0.6399543457 -4.4232514180
## FC_Fpos_AGTCTCCGTCTTTCTA-1FC_FI_GFPve_pos      0.7657794528 -3.7814101853
## FC_Fpos_AGTGTTGCACGCGTCA-1FC_FI_GFPve_pos      0.3682924084 -3.8598767795
## FC_Fpos_ATAGAGATCGTCTCAC-1FC_FI_GFPve_pos      0.8485673837 -3.8853720702
## FC_Fpos_ATCAGGTAGAGTTGTA-1FC_FI_GFPve_pos      0.6360677652 -4.2556166627
## FC_Fpos_ATCGTGAGTAGGCTGA-1FC_FI_GFPve_pos      0.5185592942 -3.3213171639
## FC_Fpos_ATGTCCCCAATGTGGG-1FC_FI_GFPve_pos      1.4674403839 -2.5779037155
## FC_Fpos_ATTACCTGTACCAGAG-1FC_FI_GFPve_pos      1.9632024936 -2.2669219650
## FC_Fpos_ATTCGTTAGGACGGAG-1FC_FI_GFPve_pos      0.7522179656 -2.8779494442
## FC_Fpos_ATTCTTGGTGAGCCAA-1FC_FI_GFPve_pos     -0.0264799423 -3.7056257405
## FC_Fpos_ATTTCACTCGCAATTG-1FC_FI_GFPve_pos     -0.1558445977 -4.5969695427
## FC_Fpos_CAATACGAGCGCTTCG-1FC_FI_GFPve_pos      0.1902608149 -3.9479850806
## FC_Fpos_CAATGACTCGTGGACC-1FC_FI_GFPve_pos      1.3156287842 -2.5536350884
## FC_Fpos_CACAACAGTGACAGGT-1FC_FI_GFPve_pos      0.6153701715 -3.9549101271
## FC_Fpos_CACGTGGAGCTCACTA-1FC_FI_GFPve_pos      0.4293463998 -4.2934870698
## FC_Fpos_CATGCAACATGGCGCT-1FC_FI_GFPve_pos      1.5715884142 -2.5039834656
## FC_Fpos_CATGGATCAACTGAAA-1FC_FI_GFPve_pos      1.6326453857 -2.0013050713
## FC_Fpos_CATTGAGTCTGGGCCA-1FC_FI_GFPve_pos      0.4955175690 -4.3008761682
## FC_Fpos_CATTGTTCAGGACGAT-1FC_FI_GFPve_pos     -0.2477681316 -3.8152351059
## FC_Fpos_CATTTCAAGACTCCGC-1FC_FI_GFPve_pos      0.5140358977 -4.1062427439
## FC_Fpos_CCACAAACAATGCAAA-1FC_FI_GFPve_pos      0.6589471869 -3.9145151056
## FC_Fpos_CCACGAGGTATGGTAA-1FC_FI_GFPve_pos      0.4915837817 -4.4345777311
## FC_Fpos_CCACTTGAGATCGCCC-1FC_FI_GFPve_pos      0.1229949169 -4.4893940367
## FC_Fpos_CCCATTGTCTCACTCG-1FC_FI_GFPve_pos      0.6649968200 -3.9283531584
## FC_Fpos_CCGCAAGCAAGTATCC-1FC_FI_GFPve_pos     -0.0060904451 -4.6284659855
## FC_Fpos_CCGGACATCATTCCTA-1FC_FI_GFPve_pos     -0.0473168544 -3.8254309572
## FC_Fpos_CCTAACCGTCCGAAAG-1FC_FI_GFPve_pos     -0.1318044491 -4.4563099422
## FC_Fpos_CGATGGCAGGAGCTGT-1FC_FI_GFPve_pos      0.5758409672 -2.8485544600
## FC_Fpos_CGCAGGTCAAAGACGC-1FC_FI_GFPve_pos      0.3904264145 -3.8601149596
## FC_Fpos_CGGAGAAGTATGGTAA-1FC_FI_GFPve_pos     -0.3809133597 -3.4948995985
## FC_Fpos_CGTAGTAAGAGATTCA-1FC_FI_GFPve_pos      0.8452769689 -3.7980823793
## FC_Fpos_CGTCAAAAGGGATCTG-1FC_FI_GFPve_pos     -0.0422318257 -4.1560539283
## FC_Fpos_CGTCAAAGTGTCCACG-1FC_FI_GFPve_pos      0.2707431786 -3.7212502636
## FC_Fpos_CGTGAATTCGGTCACG-1FC_FI_GFPve_pos     -0.3088227786 -3.4321863331
## FC_Fpos_CGTGATAGTTGTAGCT-1FC_FI_GFPve_pos      0.8145405822 -3.9692262687
## FC_Fpos_CTACAGACACTGGATT-1FC_FI_GFPve_pos      0.4353689723 -3.6743935026
## FC_Fpos_CTACCTGAGTCTGCAT-1FC_FI_GFPve_pos      0.1152420513 -3.7958212175
## FC_Fpos_CTACCTGGTCCCTGTT-1FC_FI_GFPve_pos     -0.5097454913 -3.6678453364
## FC_Fpos_CTCCACATCAGCGCGT-1FC_FI_GFPve_pos     -0.1072283029 -4.2066842593
## FC_Fpos_CTCCTTTTCCCTCATG-1FC_FI_GFPve_pos      0.7543599777 -3.4811589398
## FC_Fpos_CTGCAGGGTTTCGTTT-1FC_FI_GFPve_pos      0.4810791783 -4.2108972348
## FC_Fpos_CTGCCATAGTAATACG-1FC_FI_GFPve_pos      0.4646538071 -3.2981750645
## FC_Fpos_CTTCCGACAGCTGTTA-1FC_FI_GFPve_pos      0.5957497530 -3.2483344950
## FC_Fpos_CTTCCTTTCGAAGCAG-1FC_FI_GFPve_pos      0.4494193964 -3.5600232758
## FC_Fpos_GAACACTAGATTGTGA-1FC_FI_GFPve_pos      0.3013932280 -3.2375684656
## FC_Fpos_GAAGTAACACAGTGTT-1FC_FI_GFPve_pos      0.3398229651 -3.4590285696
## FC_Fpos_GACACGCTCTACTGAG-1FC_FI_GFPve_pos      0.0098009132 -3.5573630013
## FC_Fpos_GACCAATTCACTTATC-1FC_FI_GFPve_pos      0.2167203896 -3.2689711727
## FC_Fpos_GACCTTCCATGTTACG-1FC_FI_GFPve_pos      0.1740895979 -3.5143673100
## FC_Fpos_GAGATGGAGCACGTCC-1FC_FI_GFPve_pos      0.2486871652 -3.8229045548
## FC_Fpos_GAGCCTGTCTAAGAAG-1FC_FI_GFPve_pos     -0.0804367416 -4.1543585338
## FC_Fpos_GAGTCATAGCACTGGA-1FC_FI_GFPve_pos      0.4737266474 -3.9227788724
## FC_Fpos_GAGTCTAGTTGTGCCG-1FC_FI_GFPve_pos      0.2593177132 -3.7070156016
## FC_Fpos_GAGTTGTGTAGGCAAC-1FC_FI_GFPve_pos      0.6152540617 -4.0818333306
```

```
## FC_Fpos_GATCCCTCAGTGAGCA-1FC_FI_GFPve_pos     -0.5890644617 -3.9819347538
## FC_Fpos_GATTCTTGTAAGGTCG-1FC_FI_GFPve_pos      0.7671485715 -4.1839418687
## FC_Fpos_GCACATACAGGTGAGT-1FC_FI_GFPve_pos     -0.5117122359 -3.0381558813
## FC_Fpos_GCACGTGAGAGTCACG-1FC_FI_GFPve_pos      0.0296435826 -4.7482490583
## FC_Fpos_GCAGTTAAGGAAGTCC-1FC_FI_GFPve_pos     -0.0884945296 -4.3600604810
## FC_Fpos_GCAGTTATCAAACCTG-1FC_FI_GFPve_pos     -0.2193368949 -4.0946052946
## FC_Fpos_GCATCGGTCAAGCGTT-1FC_FI_GFPve_pos      0.5044779353 -3.2397074379
## FC_Fpos_GCCTGTTCACGGTGTC-1FC_FI_GFPve_pos      0.1458022766 -4.6823721841
## FC_Fpos_GCTTGGGAGATCGACG-1FC_FI_GFPve_pos      0.5330653958 -3.3880262055
## FC_Fpos_GCTTTCGGTGTGTCCG-1FC_FI_GFPve_pos     -0.3937399514 -2.7329654373
## FC_Fpos_GGAGATGAGTCGGCAA-1FC_FI_GFPve_pos      1.1636309795 -2.6735710777
## FC_Fpos_GGAGGTATCGTTGTTT-1FC_FI_GFPve_pos      0.5385905080 -4.2850993849
## FC_Fpos_GGCAGTCCATCGCTGG-1FC_FI_GFPve_pos      1.0041432552 -2.5964235939
## FC_Fpos_GGGTATTTCCAACTGA-1FC_FI_GFPve_pos      0.0480425380 -4.6314662300
## FC_Fpos_GGTGTTATCAACACCA-1FC_FI_GFPve_pos      1.5246851616 -2.1263162770
## FC_Fpos_GGTTGTAGTCACATTG-1FC_FI_GFPve_pos     -0.2613409661 -4.4601021685
## FC_Fpos_GTAGGTTTCACATTGG-1FC_FI_GFPve_pos      0.9808038645 -4.3744927921
## FC_Fpos_GTAGTACTCGTGTCAA-1FC_FI_GFPve_pos      0.0543569349 -4.1745171227
## FC_Fpos_GTATTGGGTCGAACGA-1FC_FI_GFPve_pos     -0.0540699117 -3.8815631546
## FC_Fpos_GTCAAACCATCCGGCA-1FC_FI_GFPve_pos     -0.1013699822 -4.0936268248
## FC_Fpos_GTCACGGTCTTCTGTA-1FC_FI_GFPve_pos      0.6740293674 -2.6824011482
## FC_Fpos_GTCACTCAGAAGCGAA-1FC_FI_GFPve_pos     -0.2284693248 -2.8000116028
## FC_Fpos_GTCACTCAGTCGTCTA-1FC_FI_GFPve_pos      0.3087584548 -3.9991334953
## FC_Fpos_GTCCACTCAATTGCTG-1FC_FI_GFPve_pos     -0.6150943763 -4.1212735452
## FC_Fpos_GTCTCACAGTGCACTT-1FC_FI_GFPve_pos      1.7803137474 -2.3975977577
## FC_Fpos_GTGAGTTGTGCAACGA-1FC_FI_GFPve_pos      0.1760378353 -3.2463131823
## FC_Fpos_GTGAGTTTCGGTCTGG-1FC_FI_GFPve_pos     -0.2829935156 -3.3621854462
## FC_Fpos_GTGCTTCAGCGTGCTC-1FC_FI_GFPve_pos     -0.1596039767 -2.2753450550
## FC_Fpos_GTGGAAGTCGTCCTCA-1FC_FI_GFPve_pos      0.6218120627 -2.9485237278
## FC_Fpos_GTGGAGATCCATTGTT-1FC_FI_GFPve_pos      0.8436315231 -3.2190700449
## FC_Fpos_GTTCATTGTCTACAAC-1FC_FI_GFPve_pos      0.4027490191 -4.2789158799
## FC_Fpos_GTTGCTCCAAATGAGT-1FC_FI_GFPve_pos      0.8873610668 -4.1128138580
## FC_Fpos_TAAGCACAGTGCAACG-1FC_FI_GFPve_pos      1.6063640289 -1.8946265854
## FC_Fpos_TACGGTATCACTGCTC-1FC_FI_GFPve_pos      0.1113779120 -4.3007473029
## FC_Fpos_TACTTCAGTTAGGCTT-1FC_FI_GFPve_pos      0.4580790572 -3.8751632847
## FC_Fpos_TAGGGTTGTTACGGAG-1FC_FI_GFPve_pos      0.6838498287 -3.1056033053
## FC_Fpos_TAGTGCACACCAGTAT-1FC_FI_GFPve_pos     -0.1101170123 -4.3663541652
## FC_Fpos_TATACCTAGCATCTTG-1FC_FI_GFPve_pos      0.6984141044 -3.3122329392
## FC_Fpos_TATACCTCATGGCACC-1FC_FI_GFPve_pos      0.0009943716 -3.8061012902
## FC_Fpos_TATATCCTCGTGAGAG-1FC_FI_GFPve_pos      0.6837484412 -3.9948094763
## FC_Fpos_TATCTGTGTGCTTATG-1FC_FI_GFPve_pos      0.7095057540 -4.3432919957
## FC_Fpos_TATTGCTTCACAGTGT-1FC_FI_GFPve_pos     -0.3630996026 -3.5683009304
## FC_Fpos_TATTGGGCACCAGCCA-1FC_FI_GFPve_pos      0.4055902533 -3.8203789868
## FC_Fpos_TATTTCGGTCATCGCG-1FC_FI_GFPve_pos     -0.1241483591 -4.5410483010
## FC_Fpos_TCACAAGCAGGGAATC-1FC_FI_GFPve_pos      0.3077174597 -3.1071930803
## FC_Fpos_TCAGCCTCAACACAAA-1FC_FI_GFPve_pos     -0.4230080075 -4.0899790801
## FC_Fpos_TCATCATAGTGCACCC-1FC_FI_GFPve_pos      0.0309676848 -4.6154524960
## FC_Fpos_TCATGGAAGCTGTGCC-1FC_FI_GFPve_pos      0.5774431400 -3.3288889803
## FC_Fpos_TCCACGTAGGTGATCG-1FC_FI_GFPve_pos      0.2794377677 -3.0126794256
## FC_Fpos_TCCACGTGTTGGAGAC-1FC_FI_GFPve_pos      0.4506363444 -3.8576295652
## FC_Fpos_TCCATGCGTCCGGTGT-1FC_FI_GFPve_pos      0.6308439545 -2.8235381521
## FC_Fpos_TCCCAGTTCGTGTTCC-1FC_FI_GFPve_pos      0.1814520829 -4.0310537495
## FC_Fpos_TCCGAAATCTCGCGTT-1FC_FI_GFPve_pos     -0.3477227069 -3.7753716387
## FC_Fpos_TCCTTCTAGAGCATAT-1FC_FI_GFPve_pos      0.9491350822 -4.3034767188
## FC_Fpos_TCGACCTGTTAGGCCC-1FC_FI_GFPve_pos      0.6046053343 -4.3161967375
## FC_Fpos_TCGCAGGAGTTGGCGA-1FC_FI_GFPve_pos      0.0157992117 -4.3214552380
## FC_Fpos_TCGCAGGCAAACTCTG-1FC_FI_GFPve_pos     -0.5049215384 -3.8306667484
## FC_Fpos_TCGCTCAGTCAAACGG-1FC_FI_GFPve_pos      0.0784789197 -3.7593179025
## FC_Fpos_TCTAACTTCGTCTAAG-1FC_FI_GFPve_pos     -2.7593760319 -0.3912338890
## FC_Fpos_TCTGCCACACTGGCGT-1FC_FI_GFPve_pos      0.7958660297 -3.4830935873
## FC_Fpos_TCTGCCAGTACAGAGC-1FC_FI_GFPve_pos      0.4579060249 -2.8523233809
## FC_Fpos_TCTGGCTAGGCCTAAG-1FC_FI_GFPve_pos      0.3026721649 -3.8702036418
## FC_Fpos_TGACTCCAGACGATAT-1FC_FI_GFPve_pos      0.4417701535 -3.5469834484
## FC_Fpos_TGATTTCAGCGCACAA-1FC_FI_GFPve_pos     -0.3708162434 -4.1089839019
## FC_Fpos_TGCAGTATCTTGAGCA-1FC_FI_GFPve_pos      0.7681192331 -2.4956049599
## FC_Fpos_TGGCGTGAGCCGCTTG-1FC_FI_GFPve_pos      0.7619320326 -3.0651383080
## FC_Fpos_TGGGATTGTCGAAGCA-1FC_FI_GFPve_pos     -0.2158839643 -3.2694656529
## FC_Fpos_TGGTAGTAGTTTGGCT-1FC_FI_GFPve_pos      0.6036659055 -4.1559339441
## FC_Fpos_TGTCCACAGGACCCAA-1FC_FI_GFPve_pos     -0.2694271333 -2.7287039437
## FC_Fpos_TGTCCTGAGCAAATCA-1FC_FI_GFPve_pos     -0.3352147020 -3.2913614191
```

```
## FC_Fpos_TGTCCTGCAATTCACG-1FC_FI_GFPve_pos      0.4264419250 -3.6387341894
## FC_Fpos_TGTTGGAGTCGGCACT-1FC_FI_GFPve_pos      0.4262499385 -4.0408789195
## FC_Fpos_TTAATCCTCTTCGGAA-1FC_FI_GFPve_pos      0.4010102920 -3.8009277262
## FC_Fpos_TTCACGCTCATAGAGA-1FC_FI_GFPve_pos      0.6822180323 -3.1414588370
## FC_Fpos_TTCATGTGTTCAACGT-1FC_FI_GFPve_pos      0.5556181245 -3.4306814350
## FC_Fpos_TTCCAATGTCACGACC-1FC_FI_GFPve_pos      0.4213150315 -3.7605187573
## FC_Fpos_TTCCTTCAGATGTTCC-1FC_FI_GFPve_pos     -0.5760598011 -4.0209172048
## FC_Fpos_TTCCTTCCACAATGTC-1FC_FI_GFPve_pos     -0.1232406139 -4.5946360208
## FC_Fpos_TTCGGTCAGTTTCGGT-1FC_FI_GFPve_pos      0.4706884198 -3.4207454600
## FC_Fpos_TTCGGTCTCAACCGAT-1FC_FI_GFPve_pos     -0.2681258298 -4.7004138135
## FC_Fpos_TTCTGTACACCAAAGG-1FC_FI_GFPve_pos      0.5889665656 -3.4216273703
## FC_Fpos_TTCTTCCCAACGCATT-1FC_FI_GFPve_pos     -0.0431419887 -3.8976174869
## FC_Fpos_TTGAGTGTCGGACGTC-1FC_FI_GFPve_pos      0.5702091031 -4.0715981402
## FC_Fpos_TTGCCTGAGTATAGGT-1FC_FI_GFPve_pos     -0.3846977331 -4.7583439104
## FC_Fpos_TTGGATGAGGTCCAGA-1FC_FI_GFPve_pos      0.0289626144 -3.7734138049
## FC_Fpos_TTGTTGTCACCCTCTA-1FC_FI_GFPve_pos      0.9090221100 -4.3336645343
## FC_Fpos_TTGTTTGCAGACACCC-1FC_FI_GFPve_pos     -0.4445133634 -4.0190153398
## dCA2_Mpos_AAAGAACCAATCACGT-1dCA2_MI_GFPve_pos -1.4822323151 -2.9307139792
## dCA2_Mpos_AAAGTCCTCCGAGGCT-1dCA2_MI_GFPve_pos -2.5584809609 -1.7869174160
## dCA2_Mpos_AACAACCTCGACGAGA-1dCA2_MI_GFPve_pos  1.6942713432 -1.3766898789
## dCA2_Mpos_ACAACCAAGAGCCGAT-1dCA2_MI_GFPve_pos  1.5445943050 -1.1467883267
## dCA2_Mpos_ACACGCGAGCACCGAA-1dCA2_MI_GFPve_pos  0.6433290415 -0.1246514000
## dCA2_Mpos_ACGATGTGTCTACTGA-1dCA2_MI_GFPve_pos -2.2655057736 -1.4532615818
## dCA2_Mpos_ACGGAAGCAATGTCTG-1dCA2_MI_GFPve_pos -2.5293583698 -0.8695470967
## dCA2_Mpos_ACTGCAATCACGTCCT-1dCA2_MI_GFPve_pos  1.2858480387 -0.1881284394
## dCA2_Mpos_ACTGTCCCAGTCAGCC-1dCA2_MI_GFPve_pos  1.6140268020 -0.2505001702
## dCA2_Mpos_AGCTTCCAGTTGTCGT-1dCA2_MI_GFPve_pos -1.3249552078 -0.9059886612
## dCA2_Mpos_AGGATAAAGACGAAGA-1dCA2_MI_GFPve_pos  0.8592636757 -0.8831624665
## dCA2_Mpos_AGGCCACTCGATCCAA-1dCA2_MI_GFPve_pos -0.3034295268  0.3210163437
## dCA2_Mpos_AGGCTGCAGTCACTAC-1dCA2_MI_GFPve_pos  1.6671377592 -1.3218223728
## dCA2_Mpos_AGGCTGCCAAGACAAT-1dCA2_MI_GFPve_pos -0.4194666274  0.3782310806
## dCA2_Mpos_AGGGTTTCATTCTCTA-1dCA2_MI_GFPve_pos  0.4327299170 -0.4113874115
## dCA2_Mpos_ATGCCTCGTGTTGCCG-1dCA2_MI_GFPve_pos -2.3213891812 -1.0966789402
## dCA2_Mpos_ATGGGTTAGCAAGGAA-1dCA2_MI_GFPve_pos -2.2917883225 -1.2869634308
## dCA2_Mpos_ATTCTACTCAGGAAAT-1dCA2_MI_GFPve_pos -1.0565336175 -1.4003915467
## dCA2_Mpos_ATTTACCCACATGTTG-1dCA2_MI_GFPve_pos  1.1232068710 -1.0476526894
## dCA2_Mpos_CAAGACTTCAGCTAGT-1dCA2_MI_GFPve_pos -1.6338491268 -0.5944366135
## dCA2_Mpos_CACACAAAGCTTTCCC-1dCA2_MI_GFPve_pos  0.8322169356 -0.9619748272
## dCA2_Mpos_CACTAAGCACAAAGTA-1dCA2_MI_GFPve_pos  1.6663912706 -1.3661622681
## dCA2_Mpos_CAGCAATTCCGAGTGC-1dCA2_MI_GFPve_pos  0.2852307730 -0.1705579437
## dCA2_Mpos_CAGCCAGAGACAACTA-1dCA2_MI_GFPve_pos -1.0562338062 -1.2120761551
## dCA2_Mpos_CATCGTCAGTAATACG-1dCA2_MI_GFPve_pos  0.5138657265  0.1393347107
## dCA2_Mpos_CCAATTTCATTGCTTT-1dCA2_MI_GFPve_pos  0.6139903240 -0.4120249428
## dCA2_Mpos_CCATCACCATGACTCA-1dCA2_MI_GFPve_pos -1.4767011233 -0.9575357117
## dCA2_Mpos_CCGAACGCAGTTGTCA-1dCA2_MI_GFPve_pos  0.2875270717 -0.3296923317
## dCA2_Mpos_CCTCCTCCAGACCTAT-1dCA2_MI_GFPve_pos  1.6210340433 -1.2721922077
## dCA2_Mpos_CGAGAAGAGATAGCTA-1dCA2_MI_GFPve_pos -2.0789489575 -1.5323626675
## dCA2_Mpos_CGCCATTGTCACCGCA-1dCA2_MI_GFPve_pos  0.4176193647  0.5054135643
## dCA2_Mpos_CGGGTGTTCTCATGCC-1dCA2_MI_GFPve_pos -1.2148327656 -1.0568368115
## dCA2_Mpos_CGGTCAGGTAGCTGTT-1dCA2_MI_GFPve_pos  0.5272338562  0.4312482200
## dCA2_Mpos_CGTGATATCACGACTA-1dCA2_MI_GFPve_pos -2.7150833435 -1.5588109173
## dCA2_Mpos_CGTGCTTAGTAATCCC-1dCA2_MI_GFPve_pos -2.6242623158 -1.5590583958
## dCA2_Mpos_CTATAGGCAGGACATG-1dCA2_MI_GFPve_pos  0.9294910602  0.6368861519
## dCA2_Mpos_CTCAAGACATAGAGGC-1dCA2_MI_GFPve_pos  0.8922817879 -0.2137078919
## dCA2_Mpos_CTCATTAAGTCCTACA-1dCA2_MI_GFPve_pos  2.0797248058 -1.2188920654
## dCA2_Mpos_CTCCTTTCAGATACTC-1dCA2_MI_GFPve_pos  0.4877875023 -0.1034598030
## dCA2_Mpos_CTGCGAGGTTGCCGAC-1dCA2_MI_GFPve_pos -2.2872025795 -1.3939728417
## dCA2_Mpos_CTTCGGTCAGGCAATG-1dCA2_MI_GFPve_pos -2.2192532845 -1.4761213936
## dCA2_Mpos_GAACTGTTCACCACAA-1dCA2_MI_GFPve_pos -1.4264443941 -1.1067785897
## dCA2_Mpos_GACCCAGTCTGGCCTT-1dCA2_MI_GFPve_pos  0.0585069828 -0.0858072914
## dCA2_Mpos_GAGCCTGCAGGAGACT-1dCA2_MI_GFPve_pos -2.5298108883 -1.5056953110
## dCA2_Mpos_GAGGCAAAGTCTCCTC-1dCA2_MI_GFPve_pos  0.0756571762  0.4078507744
## dCA2_Mpos_GATTGGTTCTTCTTCC-1dCA2_MI_GFPve_pos  1.5089347534 -1.3645352997
## dCA2_Mpos_GCATTAGGTTGTCCCT-1dCA2_MI_GFPve_pos -2.6876203842 -1.6370455898
## dCA2_Mpos_GGAACCCTCTTCCTAA-1dCA2_MI_GFPve_pos  1.0692006521 -1.1636106648
## dCA2_Mpos_GGATCTATCTAAGCGT-1dCA2_MI_GFPve_pos -2.1728436775 -1.1458160557
## dCA2_Mpos_GGCTTTCCAAATCCCA-1dCA2_MI_GFPve_pos -1.4608917065 -1.4624409355
## dCA2_Mpos_GGGTGTCAGATACCAA-1dCA2_MI_GFPve_pos  0.5142617993 -0.0638427414
## dCA2_Mpos_GGTAACTAGTCACTCA-1dCA2_MI_GFPve_pos -1.3845323153 -1.4517254509
## dCA2_Mpos_GTCAAGTAGGCATTTC-1dCA2_MI_GFPve_pos  0.2261113100  0.4470944725
```

```
## dCA2_Mpos_GTCAGCGCAAGCCTGC-1dCA2_MI_GFPve_pos -1.4559877939 -0.8772661366
## dCA2_Mpos_GTCCCATTCCAAGCTA-1dCA2_MI_GFPve_pos -1.3209409542 -0.9922043957
## dCA2_Mpos_GTTAGACAGATAACAC-1dCA2_MI_GFPve_pos -1.6403168268 -1.4416615643
## dCA2_Mpos_TACATTCTCTAGCCAA-1dCA2_MI_GFPve_pos  0.4739067130 -0.6249203362
## dCA2_Mpos_TACCCACCACGGCACT-1dCA2_MI_GFPve_pos  0.8200075202 -0.9630465187
## dCA2_Mpos_TACCGAAAGCAGTAAT-1dCA2_MI_GFPve_pos -2.5693986244 -1.5740608849
## dCA2_Mpos_TACGCTCCACCTGCTT-1dCA2_MI_GFPve_pos -1.8252103157 -1.6163549103
## dCA2_Mpos_TACTTACAGTGGGAAA-1dCA2_MI_GFPve_pos -0.2705451973 -0.1630944885
## dCA2_Mpos_TACTTGTCAGGTGGAT-1dCA2_MI_GFPve_pos -1.6985580750 -0.6022972740
## dCA2_Mpos_TCAATCTTCGCGCTGA-1dCA2_MI_GFPve_pos -1.2360094614 -0.2944469132
## dCA2_Mpos_TCACATTCAGAGACTG-1dCA2_MI_GFPve_pos -1.8082378931 -1.2970451988
## dCA2_Mpos_TCACGCTAGCGTTAGG-1dCA2_MI_GFPve_pos  1.0916661196 -0.9599449314
## dCA2_Mpos_TCACTCGGTTGTGTAC-1dCA2_MI_GFPve_pos  0.0161586724 -0.3787779488
## dCA2_Mpos_TCAGCCTTCCTTTAGT-1dCA2_MI_GFPve_pos  0.6145162754  0.0701136909
## dCA2_Mpos_TCCTCTTAGTTAACGA-1dCA2_MI_GFPve_pos -1.7359955139 -2.9812837519
## dCA2_Mpos_TCGCAGGGTTTGGAGG-1dCA2_MI_GFPve_pos -1.7316996880 -1.2843129315
## dCA2_Mpos_TCGCTCAGTCTATGAC-1dCA2_MI_GFPve_pos -0.9409277864 -0.4307865776
## dCA2_Mpos_TCTTAGTGTGCAACGA-1dCA2_MI_GFPve_pos -1.1342053838 -0.5106444039
## dCA2_Mpos_TGGGAGACAGTAGATA-1dCA2_MI_GFPve_pos -0.7195964284 -0.4123248734
## dCA2_Mpos_TGGGCTGGTCTCGGAC-1dCA2_MI_GFPve_pos  1.0384974651 -0.4762863793
## dCA2_Mpos_TGTAAGCTCGATGGAG-1dCA2_MI_GFPve_pos -1.2029534407 -0.7006821312
## dCA2_Mpos_TGTCCCAAGCAGCCTC-1dCA2_MI_GFPve_pos  0.0206844680 -1.0847336926
## dCA2_Mpos_TGTGCGGGTACTGACT-1dCA2_MI_GFPve_pos  1.2538790874  0.0017891250
## dCA2_Mpos_TTGACCCGTTGCTTGA-1dCA2_MI_GFPve_pos  0.8663599663 -0.3289436974
## dCA2_Mpos_TTGGGCGTCTCGACCT-1dCA2_MI_GFPve_pos -1.5650088616 -0.8924450554
## dCA2_Mpos_TTGGGCGTCTGCGGGT-1dCA2_MI_GFPve_pos -0.9606267042 -0.0716151871
## dCA2_Mpos_TTGGGTATCCCTTGGT-1dCA2_MI_GFPve_pos -2.2645561524 -1.5344063915
## dCA2_Mpos_TTTACCACATCAGTCA-1dCA2_MI_GFPve_pos -1.4386339016 -1.1173607983
## dCA2_Mpos_TTTCCTCTCTATTCGT-1dCA2_MI_GFPve_pos  0.6118431263 -0.5586652436
## dCA2_Fpos_AAAGTCCGTGGACAGT-1dCA2_FI_GFPve_pos -1.4263534374 -1.0785131134
## dCA2_Fpos_ACACCAAAGAGGTATT-1dCA2_FI_GFPve_pos -1.1104798742 -0.2471336998
## dCA2_Fpos_ACAGGGAGTGTAAATG-1dCA2_FI_GFPve_pos  0.3783970051  0.3336487137
## dCA2_Fpos_ACTTCGCAGTAGGGTC-1dCA2_FI_GFPve_pos  1.3002154999 -0.1954197563
## dCA2_Fpos_ATGGATCGTATGCTTG-1dCA2_FI_GFPve_pos  1.1252623968 -0.2634267487
## dCA2_Fpos_CAACGATGTTTCCATT-1dCA2_FI_GFPve_pos -2.8938300438 -1.6038062252
## dCA2_Fpos_CACCAAAGTCTACTGA-1dCA2_FI_GFPve_pos  0.3069213204 -0.1062507309
## dCA2_Fpos_CAGAGCCGTATCGCAT-1dCA2_FI_GFPve_pos -1.3884029217 -0.6605338730
## dCA2_Fpos_CAGATTGTCATCGTAG-1dCA2_FI_GFPve_pos -2.1611685581 -1.4619114079
## dCA2_Fpos_CCACTTGGTCTTTCTA-1dCA2_FI_GFPve_pos -0.9201707072 -1.2358388581
## dCA2_Fpos_CCATCACGTTCTTCAT-1dCA2_FI_GFPve_pos -1.4161435194 -1.5861873306
## dCA2_Fpos_CCCTGATTCAAATGCC-1dCA2_FI_GFPve_pos -1.3477696009 -0.9674930252
## dCA2_Fpos_CCGATCTAGTGAGGCT-1dCA2_FI_GFPve_pos -2.7133734055 -1.5538110413
## dCA2_Fpos_CCGGGTATCTCCAAGA-1dCA2_FI_GFPve_pos -1.6391584702 -0.5723619141
## dCA2_Fpos_CGCGTGAAGCGCATCC-1dCA2_FI_GFPve_pos -2.1279983349 -1.1036417164
## dCA2_Fpos_CGTAAGTGTCCCTGAG-1dCA2_FI_GFPve_pos -2.5905120201 -1.4321441330
## dCA2_Fpos_CTATCCGGTGATGGCA-1dCA2_FI_GFPve_pos -0.1372966502 -0.0407833733
## dCA2_Fpos_CTCCACAAGATGCGAC-1dCA2_FI_GFPve_pos -2.4209947415 -1.3977493920
## dCA2_Fpos_CTCCCTCAGGTTGGAC-1dCA2_FI_GFPve_pos -2.2930552788 -1.6269261517
## dCA2_Fpos_CTGGACGTCATGGTAC-1dCA2_FI_GFPve_pos -1.5077752896 -0.9577366986
## dCA2_Fpos_GACACGCAGTCTTCGA-1dCA2_FI_GFPve_pos -1.4745256729 -0.7987546601
## dCA2_Fpos_GATGACTAGAGCAGTC-1dCA2_FI_GFPve_pos -0.2211182617  0.2014651619
## dCA2_Fpos_GGAGGATGTATTCCGA-1dCA2_FI_GFPve_pos -2.5050461121 -1.3916859307
## dCA2_Fpos_GGCTTTCGTAGATCGG-1dCA2_FI_GFPve_pos  0.7224315576  0.4683771454
## dCA2_Fpos_GGTGATTAGACGCATG-1dCA2_FI_GFPve_pos  0.3722528391  0.4608269058
## dCA2_Fpos_GTGGCGTCAGAAGCGT-1dCA2_FI_GFPve_pos -2.8524920769 -1.5949613728
## dCA2_Fpos_TAACCAGTCAACCTCC-1dCA2_FI_GFPve_pos -1.5977730580 -0.9758159794
## dCA2_Fpos_TATATCCCAACCACGC-1dCA2_FI_GFPve_pos -1.6752420492 -1.5319027580
## dCA2_Fpos_TATTGGGTCGCTCATC-1dCA2_FI_GFPve_pos -1.9144562311 -1.4265281834
## dCA2_Fpos_TCCATGCAGCTTTCCC-1dCA2_FI_GFPve_pos -1.7590379544 -1.4283926167
## dCA2_Fpos_TCCCAGTGTCGGAAAC-1dCA2_FI_GFPve_pos -2.1491607256 -0.6246738114
## dCA2_Fpos_TCCTGCACACCGGAAA-1dCA2_FI_GFPve_pos -1.4806555338 -1.0117890515
## dCA2_Fpos_TCTATACAGTCATACC-1dCA2_FI_GFPve_pos -1.5642524786 -1.0333232559
## dCA2_Fpos_TGACAGTTCTGGGTCG-1dCA2_FI_GFPve_pos -2.1019062824 -1.0048024334
## dCA2_Fpos_TGTTTGTGTCCTTAAG-1dCA2_FI_GFPve_pos  0.0988260083 -0.0131339707
## dCA2_Fpos_TTCCAATTCTCCACTG-1dCA2_FI_GFPve_pos -2.0626153774 -1.3769879021
## dCA2_Fpos_TTCTAACAGGGTGAAA-1dCA2_FI_GFPve_pos -1.6960418053 -1.2898761906
## dCA2_Fpos_TTGGATGGTCAGGTGA-1dCA2_FI_GFPve_pos  0.4854147010  0.1834898315
## FC_Mpos_AATGGCTCATGTGCCG-1FC_MI_GFPve_pos     -1.0929324455 -0.4949278511
## FC_Mpos_ACGATCAGTTGAATCC-1FC_MI_GFPve_pos     -0.0295990385  0.1511411987
## FC_Mpos_ACGTAGTTCCATAGAC-1FC_MI_GFPve_pos     -1.2696480580 -0.7102613129
```

```
## FC_Mpos_ACTCTCGTCACTCACC-1FC_MI_GFPve_pos     -0.8114307947 -0.3485397972
## FC_Mpos_AGAGAATAGGTCTGGA-1FC_MI_GFPve_pos     -1.3467365332 -0.9577507652
## FC_Mpos_ATGTCTTTCATACAGC-1FC_MI_GFPve_pos     -1.7355801888 -1.5684857048
## FC_Mpos_ATTCGTTTCATCAGTG-1FC_MI_GFPve_pos      0.7563331895 -0.0966825165
## FC_Mpos_CAAGACTAGAGCAGCT-1FC_MI_GFPve_pos      0.5436260872 -0.4936289467
## FC_Mpos_CACTGGGAGGACTTCT-1FC_MI_GFPve_pos     -2.0554110832 -0.8014588036
## FC_Mpos_CTACGGGAGGTCCCTG-1FC_MI_GFPve_pos     -0.3452861585  0.3997044884
## FC_Mpos_CTCCATGTCCTATTTG-1FC_MI_GFPve_pos     -1.3367514439 -1.4647094883
## FC_Mpos_GATCAGTGTTTCTTAC-1FC_MI_GFPve_pos     -2.4345922299 -1.1529199757
## FC_Mpos_GGGACAAGTTCGTGCG-1FC_MI_GFPve_pos     -2.3495592900 -1.1120102085
## FC_Mpos_GGTTCTCTCGCTAATG-1FC_MI_GFPve_pos      0.3968256883 -0.2251495995
## FC_Mpos_TCATTACCAAGCCCAC-1FC_MI_GFPve_pos     -1.9840005465 -1.7202629723
## FC_Mpos_TCATTTGTCCTTACCG-1FC_MI_GFPve_pos      0.3553803735  0.1230359398
## FC_Mpos_TCCATGCCACATACGT-1FC_MI_GFPve_pos      1.4835840635 -1.2925703205
## FC_Mpos_TCGACGGGTCGCAGTC-1FC_MI_GFPve_pos     -1.3510355778 -0.6666369118
## FC_Mpos_TGCTCGTCAGGTACGA-1FC_MI_GFPve_pos     -1.2030677624 -0.3637523331
## FC_Mpos_TGCTGAACAAGATGGC-1FC_MI_GFPve_pos     -1.0113861985 -0.4974383988
## FC_Mpos_TTACGCCCAGCCGTCA-1FC_MI_GFPve_pos     -2.1562678642 -1.6441867031
## FC_Mpos_TTCTAACTCTAGATCG-1FC_MI_GFPve_pos     -1.9561304875 -1.3684217610
## FC_Fpos_AAAGGGCCACTGCACG-1FC_FI_GFPve_pos     -1.9155835934 -0.7650978245
## FC_Fpos_AACCAACTCTATGTGG-1FC_FI_GFPve_pos      0.3407494001  0.3378982864
## FC_Fpos_AACCACAGTAGATTAG-1FC_FI_GFPve_pos      0.1157685988  0.5408439956
## FC_Fpos_AATCGTGTCCGGGACT-1FC_FI_GFPve_pos     -1.6892421074 -0.7918073811
## FC_Fpos_ACCCTTGAGAGCAGCT-1FC_FI_GFPve_pos     -1.9930124111 -0.9858357586
## FC_Fpos_AGGTCATAGGACACTG-1FC_FI_GFPve_pos     -2.5636277027 -1.1527547516
## FC_Fpos_ATCTCTAGTAGTGTGG-1FC_FI_GFPve_pos     -1.9475629158 -2.4786045231
## FC_Fpos_ATGATCGGTTCATCTT-1FC_FI_GFPve_pos     -1.1807023115 -0.5652260460
## FC_Fpos_ATTCATCCACCGTCGA-1FC_FI_GFPve_pos     -0.3575164951 -1.4166526474
## FC_Fpos_CACGAATTCGTCTAAG-1FC_FI_GFPve_pos      0.0787814133 -0.5550217308
## FC_Fpos_CAGATACGTCGGTGAA-1FC_FI_GFPve_pos     -2.0035840101 -2.7920727410
## FC_Fpos_CAGATACTCCGAGCTG-1FC_FI_GFPve_pos     -1.8362052269 -2.1062993683
## FC_Fpos_CAGGCCAAGTCATAGA-1FC_FI_GFPve_pos     -0.3851765670 -0.5684127487
## FC_Fpos_CCACGAGCATCGCTGG-1FC_FI_GFPve_pos      0.1503832810  0.3571396194
## FC_Fpos_CCCATTGGTATACCCA-1FC_FI_GFPve_pos      0.3867299132 -0.3480791725
## FC_Fpos_CGCGTGATCAACGAGG-1FC_FI_GFPve_pos      0.2802734844  0.4019713722
## FC_Fpos_CGTAATGTCTTGCGCT-1FC_FI_GFPve_pos     -2.4035394020 -1.2189340271
## FC_Fpos_CGTGCTTTCAACCCGG-1FC_FI_GFPve_pos     -2.1333685942 -1.5324093975
## FC_Fpos_CGTTAGACAGCTTCCT-1FC_FI_GFPve_pos     -0.5142004616 -0.2096438088
## FC_Fpos_CTAACCCCAGGAAGTC-1FC_FI_GFPve_pos     -1.2057349510  0.3649444900
## FC_Fpos_CTGTATTGTCTACTGA-1FC_FI_GFPve_pos     -1.3949974604 -0.9193956532
## FC_Fpos_GACGTTAAGACGTCGA-1FC_FI_GFPve_pos     -0.7006719537  0.2705331169
## FC_Fpos_GCAGCCATCTGCCTGT-1FC_FI_GFPve_pos     -2.0499374695 -1.6498253025
## FC_Fpos_GTAGAAAAGGACACTG-1FC_FI_GFPve_pos      0.6097431950 -0.2494892754
## FC_Fpos_GTCTACCGTTGACTGT-1FC_FI_GFPve_pos     -2.0434483118 -1.5311860718
## FC_Fpos_GTGATGTCAGGTACGA-1FC_FI_GFPve_pos     -1.3851215668 -0.8468482174
## FC_Fpos_GTTGCGGAGTTGGGAC-1FC_FI_GFPve_pos      0.4089702539  0.3956981025
## FC_Fpos_TACCTGCGTATCAAGA-1FC_FI_GFPve_pos      0.3009238415  0.0347037636
## FC_Fpos_TACTTACTCTCTTCAA-1FC_FI_GFPve_pos      0.5909127168 -0.8861934818
## FC_Fpos_TCCAGAATCCAAACCA-1FC_FI_GFPve_pos      0.0581141137  0.3481560074
## FC_Fpos_TCCGAAACATTGTACG-1FC_FI_GFPve_pos     -0.4564355738  0.5393715225
## FC_Fpos_TCCTCCCTCTCGCGTT-1FC_FI_GFPve_pos     -1.7737338371 -1.6334321179
## FC_Fpos_TTGACCCGTTCAAAGA-1FC_FI_GFPve_pos     -2.2317790814 -1.3958789982
## FC_Fpos_TTGTGTTAGTACGAGC-1FC_FI_GFPve_pos     -0.0505464725  0.4629278503
## dCA2_Mpos_AAATGGACACGCCAGT-1dCA2_MI_GFPve_pos -0.2177005201 -2.6605083622
## dCA2_Mpos_AATGGCTTCCATCTAT-1dCA2_MI_GFPve_pos -0.1260048650 -2.2410602249
## dCA2_Mpos_ACTCCCATCCCATTTA-1dCA2_MI_GFPve_pos  1.9849441223 -7.7949258961
## dCA2_Mpos_ACTTCGCAGTTGGAAT-1dCA2_MI_GFPve_pos -0.1587397946 -2.1708473839
## dCA2_Mpos_CAACGATGTCAGACTT-1dCA2_MI_GFPve_pos -0.4308345742 -3.3459340014
## dCA2_Mpos_CACCGTTAGGTCCCTG-1dCA2_MI_GFPve_pos -0.0384477802 -0.1983031906
## dCA2_Mpos_CATCCGTGTCAGTTTG-1dCA2_MI_GFPve_pos  0.1457081013 -3.2422359862
## dCA2_Mpos_CATCGGGCAGCAGATG-1dCA2_MI_GFPve_pos -0.1724721343 -2.1782588639
## dCA2_Mpos_CCCTAACTCGACTCCT-1dCA2_MI_GFPve_pos  0.0634517245 -2.3636564888
## dCA2_Mpos_CCTGTTGGTTGCTCCT-1dCA2_MI_GFPve_pos  2.0406937771 -7.8695315994
## dCA2_Mpos_CGGAATTAGCTTAGTC-1dCA2_MI_GFPve_pos -0.4642318912 -2.3160693325
## dCA2_Mpos_CGTGCTTCAGACCATT-1dCA2_MI_GFPve_pos -0.1921790648 -2.1882297672
## dCA2_Mpos_GGTGAAGTCTATCGGA-1dCA2_MI_GFPve_pos  2.4187417202  2.0436382614
## dCA2_Mpos_GTACAGTGTTCCTAAG-1dCA2_MI_GFPve_pos -0.1529384615 -2.1417970337
## dCA2_Mpos_TACCGAATCGGTCACG-1dCA2_MI_GFPve_pos -0.1516888138 -2.1464590706
## dCA2_Mpos_TAGGAGGCAAGTGCTT-1dCA2_MI_GFPve_pos -0.2878503330 -2.3748621620
## dCA2_Mpos_TATCGCCTCCTGCTAC-1dCA2_MI_GFPve_pos -0.1270522900 -2.2267093338
```

```
## dCA2_Mpos_TATCGCCTCGACATTG-1dCA2_MI_GFPve_pos  0.1187435262 -3.0408856549
## dCA2_Mpos_TATCTTGCACCTCTGT-1dCA2_MI_GFPve_pos  0.0905801229 -2.4119536556
## dCA2_Mpos_TCTGCCATCTCATTTG-1dCA2_MI_GFPve_pos -0.1421728000 -2.1936182656
## dCA2_Mpos_TGACCCTAGTAGTCAA-1dCA2_MI_GFPve_pos -0.2946822174 -2.3750824608
## dCA2_Mpos_TGTACAGGTACAGGTG-1dCA2_MI_GFPve_pos  2.0032868557 -7.8198263325
## dCA2_Mpos_TTGGGTACACACACTA-1dCA2_MI_GFPve_pos -0.4062091984 -2.3967814125
## dCA2_Mpos_TTTACGTGTCACAGAG-1dCA2_MI_GFPve_pos -0.2108488209 -2.2363688626
## dCA2_Mpos_TTTAGTCGTTATTCTC-1dCA2_MI_GFPve_pos -0.1308416255 -2.1736612000
## dCA2_Fpos_AAGACAACACTTACAG-1dCA2_FI_GFPve_pos -0.0703687869 -2.2840161003
## dCA2_Fpos_CCTCTAGAGATCGCTT-1dCA2_FI_GFPve_pos -0.4110237189 -2.1803180851
## dCA2_Fpos_CGGGCATAGTCATGAA-1dCA2_FI_GFPve_pos  1.9872675113 -7.7970573582
## dCA2_Fpos_CTGCTCAAGTCAATCC-1dCA2_FI_GFPve_pos -0.2797467016 -2.3012315907
## dCA2_Fpos_GCCATGGTCTGTCTCG-1dCA2_FI_GFPve_pos  0.1735826604 -3.1353449501
## dCA2_Fpos_GTTATGGGTTGCCGAC-1dCA2_FI_GFPve_pos -0.1525035990 -2.2156092800
## dCA2_Fpos_GTTGAACTCTACTATC-1dCA2_FI_GFPve_pos -0.2797750361 -2.4922043957
## dCA2_Fpos_TCAGTCCGTTGCCGCA-1dCA2_FI_GFPve_pos -0.2320599161 -2.2172751106
## FC_Mpos_AGGAATAAGTCTTCCC-1FC_MI_GFPve_pos     -0.2048849180 -0.8827755131
## FC_Mpos_ATCGCCTTCGCTCATC-1FC_MI_GFPve_pos      2.0017974548 -7.8224835076
## FC_Mpos_CATTGAGCAAATGGAT-1FC_MI_GFPve_pos     -1.7848802633 -3.3554929175
## FC_Mpos_CTCAATTAGAACAAGG-1FC_MI_GFPve_pos     -0.2226087242 -2.0141927876
## FC_Mpos_GATCCCTGTCGCAACC-1FC_MI_GFPve_pos      2.0013804607 -7.8219895043
## FC_Mpos_GATGGAGTCAGTGTTG-1FC_MI_GFPve_pos      0.2174217872 -1.7130896725
## FC_Mpos_GTAACACCAGTCGCTG-1FC_MI_GFPve_pos      0.9003278189 -2.8703338780
## FC_Mpos_TCCCAGTTCACACCGG-1FC_MI_GFPve_pos     -0.0378098614 -2.7275859989
## FC_Mpos_TGACGCGCACGTCGTG-1FC_MI_GFPve_pos     -0.1515975591 -1.5599162258
## FC_Mpos_TGTGTGAGTGTAGCAG-1FC_MI_GFPve_pos     -1.1450261302 -1.3370918907
## FC_Mpos_TTCCAATGTAGAGGAA-1FC_MI_GFPve_pos      2.0020704441 -7.8226675667
## FC_Fpos_AAAGGTAAGGATAATC-1FC_FI_GFPve_pos     -0.4666607804 -2.0155627407
## FC_Fpos_AAGCATCGTTCAAGGG-1FC_FI_GFPve_pos     -0.2550012298 -0.9614822544
## FC_Fpos_AAGTGAACAAGTGACG-1FC_FI_GFPve_pos     -0.8902263470 -2.0699443497
## FC_Fpos_AATAGAGCATGGCACC-1FC_FI_GFPve_pos     -0.5165745146  0.2084966026
## FC_Fpos_AATCACGGTTAAAGTG-1FC_FI_GFPve_pos     -0.1640294831 -0.9585892834
## FC_Fpos_ACAAAGAAGGTCACAG-1FC_FI_GFPve_pos     -0.4134414174 -1.9607228913
## FC_Fpos_ACATTTCGTCCAGGTC-1FC_FI_GFPve_pos     -0.5910031028 -1.4728037991
## FC_Fpos_ACCCAAATCGTGTGGC-1FC_FI_GFPve_pos      0.2038142555 -1.8418175854
## FC_Fpos_ACGGAAGCAGGATCTT-1FC_FI_GFPve_pos     -0.4608905144 -2.0288786568
## FC_Fpos_ACGTAGTAGCCAGAGT-1FC_FI_GFPve_pos     -0.7566057272 -1.3422045387
## FC_Fpos_ACTCTCGCAGTTGAAA-1FC_FI_GFPve_pos     -0.2399510048 -1.4330868401
## FC_Fpos_ACTGATGCAGCCTTCT-1FC_FI_GFPve_pos     -0.5442723758  0.0413742386
## FC_Fpos_AGATGAACAGGCATTT-1FC_FI_GFPve_pos     -0.1566730538 -1.5566272415
## FC_Fpos_AGCCAGCTCGGAATTC-1FC_FI_GFPve_pos      0.0449266903 -0.1857785858
## FC_Fpos_AGCGCTGTCCATTGCC-1FC_FI_GFPve_pos     -0.5850360222 -1.4609293617
## FC_Fpos_AGCGTCGAGACCTTTG-1FC_FI_GFPve_pos     -0.3937483556 -1.1190778889
## FC_Fpos_ATCACTTAGAGCATAT-1FC_FI_GFPve_pos     -0.9381763764 -1.7671250977
## FC_Fpos_ATCAGGTAGGAACGTC-1FC_FI_GFPve_pos     -0.1104802094 -1.7749778904
## FC_Fpos_ATGCGATGTTGCATCA-1FC_FI_GFPve_pos     -0.8250907488 -0.4101195015
## FC_Fpos_CAATACGAGCAAATGT-1FC_FI_GFPve_pos     -0.4249951966 -0.8693926014
## FC_Fpos_CAATCGAAGGATGCGT-1FC_FI_GFPve_pos     -0.6974150367 -2.0359050907
## FC_Fpos_CACTTCGTCGGTCGAC-1FC_FI_GFPve_pos     -0.5080829091 -0.1421675362
## FC_Fpos_CAGCGTGTCACATTGG-1FC_FI_GFPve_pos      0.0988583736 -1.9273693241
## FC_Fpos_CAGGTATGTTCCTAGA-1FC_FI_GFPve_pos     -0.3326657541 -0.1280336060
## FC_Fpos_CAGTTCCAGGAAGTAG-1FC_FI_GFPve_pos     -0.1635139665 -1.7468943276
## FC_Fpos_CATAAGCCAGTTCTAG-1FC_FI_GFPve_pos     -0.5217393167  0.4534740768
## FC_Fpos_CATCCCAAGCCAGAGT-1FC_FI_GFPve_pos     -1.5895891018 -2.0532931961
## FC_Fpos_CATCGCTAGTTGGCTT-1FC_FI_GFPve_pos     -0.9017830320 -2.0247502007
## FC_Fpos_CATCGGGAGGTTAAAC-1FC_FI_GFPve_pos     -0.2801353015 -1.8657037892
## FC_Fpos_CCAAGCGGTACAAAGT-1FC_FI_GFPve_pos     -0.4486524887 -0.8716263451
## FC_Fpos_CCACGAGTCGTTCCTG-1FC_FI_GFPve_pos     -1.4294574089 -3.1153863348
## FC_Fpos_CCTAAGAGTGACCGTC-1FC_FI_GFPve_pos     -0.2552856408 -1.7156028427
## FC_Fpos_CCTGTTGTCTGTGCTC-1FC_FI_GFPve_pos      0.0731179439  0.3260684334
## FC_Fpos_CGAATTGTCGTAACTG-1FC_FI_GFPve_pos     -1.9344432183 -3.1093912996
## FC_Fpos_CGAGTGCTCGTGGAAG-1FC_FI_GFPve_pos      2.0071282558 -7.8241572060
## FC_Fpos_CGATGCGAGTCCCAGC-1FC_FI_GFPve_pos     -0.2500773273 -1.6604678311
## FC_Fpos_CGCATAAGTTGTGTAC-1FC_FI_GFPve_pos     -0.2978516944 -1.4462413468
## FC_Fpos_CGCGTGAAGACGCCAA-1FC_FI_GFPve_pos     -0.3074117429 -1.6207110562
## FC_Fpos_CGTCAAACAGGTCAGA-1FC_FI_GFPve_pos     -0.1997161820 -1.6994375862
## FC_Fpos_CGTCCATAGTAAACGT-1FC_FI_GFPve_pos      0.2100974493 -0.0643605866
## FC_Fpos_CTACCCAGTTAAGGAT-1FC_FI_GFPve_pos     -0.3932492562 -1.8038828053
## FC_Fpos_CTACGGGAGAGGATGA-1FC_FI_GFPve_pos      0.2826446764 -0.1571769394
## FC_Fpos_CTAGGTAGTAGATTGA-1FC_FI_GFPve_pos     -0.5081834622  0.6790724121
```

```
## FC_Fpos_CTCAAGATCTCGCTTG-1FC_FI_GFPve_pos     -0.4212809332 -1.3709754624
## FC_Fpos_CTGAGGCAGGCGCTCT-1FC_FI_GFPve_pos     -0.5741573997 -1.9916948952
## FC_Fpos_CTGAGGCCAAGATTGA-1FC_FI_GFPve_pos     -0.3422167279 -2.1840371765
## FC_Fpos_CTGCCTAGTGTAGCAG-1FC_FI_GFPve_pos      2.0078609161 -7.8277454056
## FC_Fpos_CTGCGAGAGCCTGTGC-1FC_FI_GFPve_pos     -0.2640186451 -1.4703519024
## FC_Fpos_CTTGAGATCGTCGACG-1FC_FI_GFPve_pos     -0.7421392627 -1.8083858170
## FC_Fpos_CTTTCGGGTTGGCCGT-1FC_FI_GFPve_pos     -0.3128115959 -1.9282679238
## FC_Fpos_GAAGCCCGTTCTTCAT-1FC_FI_GFPve_pos     -0.9443358011 -1.8190366902
## FC_Fpos_GAAGTAACAGGCTCTG-1FC_FI_GFPve_pos      0.3512875609 -2.0591029801
## FC_Fpos_GACGCTGAGCATCTTG-1FC_FI_GFPve_pos     -0.5619798727 -1.6697616257
## FC_Fpos_GAGACTTAGCATGTTC-1FC_FI_GFPve_pos      0.5735117965  0.1306109749
## FC_Fpos_GAGGCCTCAGCGACAA-1FC_FI_GFPve_pos     -0.7377227969 -1.9018830933
## FC_Fpos_GAGGCCTCATGGTACT-1FC_FI_GFPve_pos      0.0759332202 -1.9785113014
## FC_Fpos_GAGGGTACAACTCGAT-1FC_FI_GFPve_pos     -0.5347439475 -0.1721324600
## FC_Fpos_GATGAGGGTCTGTAAC-1FC_FI_GFPve_pos     -0.7965454884 -2.2193004765
## FC_Fpos_GATGATCTCCGCATAA-1FC_FI_GFPve_pos     -1.0049123593 -1.7799455799
## FC_Fpos_GATGGAGGTGCCGGTT-1FC_FI_GFPve_pos     -0.2726026006 -1.9721781887
## FC_Fpos_GATGTTGCAAGTGGTG-1FC_FI_GFPve_pos     -0.1489020564 -1.1778759636
## FC_Fpos_GATTCGAAGAAAGCGA-1FC_FI_GFPve_pos     -0.5312958188 -1.1623143830
## FC_Fpos_GCACGTGGTGGCAACA-1FC_FI_GFPve_pos     -0.2477057107 -0.5901441254
## FC_Fpos_GCGATCGAGCAGTACG-1FC_FI_GFPve_pos     -0.5701201685  0.1942615829
## FC_Fpos_GCGATCGAGTGGACGT-1FC_FI_GFPve_pos     -0.0307360120 -1.5000223793
## FC_Fpos_GCGATCGTCTGAGATC-1FC_FI_GFPve_pos     -0.2330086879 -1.2056255020
## FC_Fpos_GCGGATCAGCGAGTAC-1FC_FI_GFPve_pos     -0.1844876144 -2.2190444149
## FC_Fpos_GCGGATCGTATTGAGA-1FC_FI_GFPve_pos      0.1706530027 -0.7521512188
## FC_Fpos_GCGTGCACAGGGCTTC-1FC_FI_GFPve_pos     -0.4111326910 -1.4496059097
## FC_Fpos_GGCACGTAGCGCCGTT-1FC_FI_GFPve_pos     -0.0821075044 -1.7010454811
## FC_Fpos_GGGCTACGTATCAGCT-1FC_FI_GFPve_pos     -0.5594945975  1.2327266060
## FC_Fpos_GGGTTTAAGGGCGAAG-1FC_FI_GFPve_pos     -0.6990687080 -1.7486703076
## FC_Fpos_GGTTAACCATGATGCT-1FC_FI_GFPve_pos     -0.2478747643 -1.5731749214
## FC_Fpos_GTAAGTCGTTTCGTTT-1FC_FI_GFPve_pos     -0.6588748224 -1.7153980412
## FC_Fpos_GTAGAGGAGGTCGAGT-1FC_FI_GFPve_pos     -0.3603933431 -1.5395233311
## FC_Fpos_GTAGGTTCACATGAAA-1FC_FI_GFPve_pos     -0.3976071305 -1.8407549538
## FC_Fpos_GTATTGGTCCCAATAG-1FC_FI_GFPve_pos      0.1291528814 -0.3744601883
## FC_Fpos_GTCAGCGTCGACATCA-1FC_FI_GFPve_pos      2.5998852424  0.5282326065
## FC_Fpos_GTCGCGAAGGGAGGAC-1FC_FI_GFPve_pos     -0.2649036400 -0.8895988144
## FC_Fpos_GTCGCGATCCTCTAAT-1FC_FI_GFPve_pos     -0.8231933541 -1.7601451554
## FC_Fpos_GTCGTTCCAGACATCT-1FC_FI_GFPve_pos     -0.6376699336 -0.5972294487
## FC_Fpos_GTGAGTTGTAATCAAG-1FC_FI_GFPve_pos     -0.2847889416 -1.0424375214
## FC_Fpos_GTGGAAGGTCGCACGT-1FC_FI_GFPve_pos     -0.6158884294 -2.2422403969
## FC_Fpos_GTGTTCCCAAACTGCT-1FC_FI_GFPve_pos      0.4755580001 -0.0463723816
## FC_Fpos_GTTCATTTCGGTAGAG-1FC_FI_GFPve_pos     -0.7663230725 -2.1087944187
## FC_Fpos_GTTGTCCCAATCGCAT-1FC_FI_GFPve_pos     -0.5012421735 -2.1059081234
## FC_Fpos_GTTGTCCGTTTACCAG-1FC_FI_GFPve_pos     -0.8126184769 -2.1123370804
## FC_Fpos_TAAGTCGCACGCCACA-1FC_FI_GFPve_pos     -1.0877781339  0.0341806732
## FC_Fpos_TACAACGTCGTTGTTT-1FC_FI_GFPve_pos     -0.5642037518 -1.6234838642
## FC_Fpos_TACGTCCCAAGCGGAT-1FC_FI_GFPve_pos      1.9278235607 -7.6753077187
## FC_Fpos_TAGACTGCAAGACCTT-1FC_FI_GFPve_pos     -0.2761082850 -1.8769859947
## FC_Fpos_TATCAGGTCAACGTGT-1FC_FI_GFPve_pos     -0.2859539367 -1.4408819355
## FC_Fpos_TATTGGGGTGTGTCGC-1FC_FI_GFPve_pos      0.2619156115 -0.1007694878
## FC_Fpos_TCAAGCAAGGGAGAAT-1FC_FI_GFPve_pos      0.7272701435  0.3766327225
## FC_Fpos_TCAAGTGGTTCTCCAC-1FC_FI_GFPve_pos     -0.5130916662 -3.3993891157
## FC_Fpos_TCACTATCACTCAGAT-1FC_FI_GFPve_pos      0.3442889027  0.0928216301
## FC_Fpos_TCATTGTAGCTAGCCC-1FC_FI_GFPve_pos     -0.4064566024  0.9617996536
## FC_Fpos_TCATTTGGTAGAGGAA-1FC_FI_GFPve_pos     -0.4724000521 -1.4636559166
## FC_Fpos_TCCAGAAAGAACTTCC-1FC_FI_GFPve_pos      0.2656231396 -0.0867123284
## FC_Fpos_TCGAACACAATGTGGG-1FC_FI_GFPve_pos     -0.8589519687  0.9837704025
## FC_Fpos_TCGTGCTAGCATGCGA-1FC_FI_GFPve_pos     -1.0306694932 -1.9818892159
## FC_Fpos_TCGTGGGCAAGCGCAA-1FC_FI_GFPve_pos     -0.5414895542 -1.9150359310
## FC_Fpos_TCTTGCGAGGAGTACC-1FC_FI_GFPve_pos     -0.2019639961 -1.3503343739
## FC_Fpos_TCTTTGAAGGGAGAAT-1FC_FI_GFPve_pos     -0.2045795672 -1.8325981774
## FC_Fpos_TGACGCGAGCTTGTTG-1FC_FI_GFPve_pos     -0.1492723074 -2.4491967835
## FC_Fpos_TGAGCATGTGGGATTG-1FC_FI_GFPve_pos     -0.4322796889 -1.2442288079
## FC_Fpos_TGAGGTTGTATCCCAA-1FC_FI_GFPve_pos     -0.3347119518 -1.4436733402
## FC_Fpos_TGATTCTGTGACATCT-1FC_FI_GFPve_pos     -0.6081213780 -1.0743050255
## FC_Fpos_TGCTGAAGTGCCTATA-1FC_FI_GFPve_pos     -0.1756317565 -0.7511081375
## FC_Fpos_TGGAGGATCTGCGGGT-1FC_FI_GFPve_pos     -0.2426484875 -1.5289876141
## FC_Fpos_TGGGATTTCTGGAAGG-1FC_FI_GFPve_pos      0.2551217549 -2.5651120819
## FC_Fpos_TGTAAGCTCTAGATCG-1FC_FI_GFPve_pos     -0.2585911728  1.4420085274
## FC_Fpos_TGTTCTATCGATTCCC-1FC_FI_GFPve_pos     -0.3881195761 -1.8968932308
```

```
## FC_Fpos_TGTTGAGTCATCACAG-1FC_FI_GFPve_pos     -0.4423374362 -1.3965463318
## FC_Fpos_TGTTTGTGTAATGCTC-1FC_FI_GFPve_pos     -0.3237668849 -1.9715058483
## FC_Fpos_TTACCGCGTGGCCACT-1FC_FI_GFPve_pos     -0.2137357384 -1.9668094792
## FC_Fpos_TTACGTTGTTGCATTG-1FC_FI_GFPve_pos      0.5966279082  0.1164412819
## FC_Fpos_TTAGGCAAGTCCCGGT-1FC_FI_GFPve_pos     -0.6506379671 -1.3442883171
## FC_Fpos_TTAGGGTGTAGTAAGT-1FC_FI_GFPve_pos     -0.6095750756 -1.6440851368
## FC_Fpos_TTCACGCGTCTTGGTA-1FC_FI_GFPve_pos      0.0723463170 -0.5940742172
## FC_Fpos_TTCCACGAGCTGTCCG-1FC_FI_GFPve_pos     -0.3062376059 -0.3871745743
## FC_Fpos_TTCTGTATCGGTTCAA-1FC_FI_GFPve_pos      0.2316509359 -1.8697852768
## FC_Fpos_TTGATGGGTTGGCCGT-1FC_FI_GFPve_pos     -0.0118759997 -1.8292510189
## FC_Fpos_TTGCATTAGAGGCGGA-1FC_FI_GFPve_pos     -0.6865251489 -0.4788374581
## FC_Fpos_TTGCGTCGTTGCCTAA-1FC_FI_GFPve_pos     -0.2081824705 -2.1534580864
## FC_Fpos_TTGCTGCGTAGTGATA-1FC_FI_GFPve_pos      0.0576588832 -2.3855271019
## FC_Fpos_TTGGATGAGAATCCCT-1FC_FI_GFPve_pos     -0.2468501754 -1.5706267037
## FC_Fpos_TTTAGTCGTATGAGCG-1FC_FI_GFPve_pos     -0.5836781390 -2.0692369618
## FC_Fpos_TTTATGCAGTCGTTAC-1FC_FI_GFPve_pos     -0.4383571871 -1.8640827812
##                                                       UMAP_3
## dCA2_Mpos_AAAGGTATCTAGAACC-1dCA2_MI_GFPve_pos  1.9179146213
## dCA2_Mpos_AACAGGGAGACCTCAT-1dCA2_MI_GFPve_pos -0.0323959666
## dCA2_Mpos_AACCATGGTACCAGAG-1dCA2_MI_GFPve_pos -0.9465831595
## dCA2_Mpos_AACCCAAAGCGTATAA-1dCA2_MI_GFPve_pos  0.4013845247
## dCA2_Mpos_AACGTCATCACGGACC-1dCA2_MI_GFPve_pos -0.0160203653
## dCA2_Mpos_AAGACAATCTGACGCG-1dCA2_MI_GFPve_pos -1.2275896149
## dCA2_Mpos_AAGGAATCATATCGGT-1dCA2_MI_GFPve_pos -0.3897465246
## dCA2_Mpos_AAGTTCGAGGATATAC-1dCA2_MI_GFPve_pos  0.3716487211
## dCA2_Mpos_AATCACGCAGAGTAAT-1dCA2_MI_GFPve_pos  1.4864736718
## dCA2_Mpos_AATGAAGAGAGTGAAG-1dCA2_MI_GFPve_pos -1.7817940789
## dCA2_Mpos_ACACTGAGTAATGATG-1dCA2_MI_GFPve_pos -1.5820694047
## dCA2_Mpos_ACAGCCGGTTATAGAG-1dCA2_MI_GFPve_pos -0.7655865627
## dCA2_Mpos_ACATGCATCGCTTGCT-1dCA2_MI_GFPve_pos  0.9771513266
## dCA2_Mpos_ACCAAACTCTCGTGGG-1dCA2_MI_GFPve_pos  1.2482510728
## dCA2_Mpos_ACCATTTTCAAGTCTG-1dCA2_MI_GFPve_pos -0.1584304350
## dCA2_Mpos_ACCCTTGTCGTGTCAA-1dCA2_MI_GFPve_pos -1.1722172576
## dCA2_Mpos_ACCGTTCAGGTACAAT-1dCA2_MI_GFPve_pos  0.3477910233
## dCA2_Mpos_ACCTGTCGTAGGTTTC-1dCA2_MI_GFPve_pos  0.9514440221
## dCA2_Mpos_ACGATGTCACCGGTCA-1dCA2_MI_GFPve_pos -0.3908714133
## dCA2_Mpos_ACGGTCGTCCCGTTGT-1dCA2_MI_GFPve_pos  0.1026393568
## dCA2_Mpos_ACGTAACCATGTTCAG-1dCA2_MI_GFPve_pos  0.4389481765
## dCA2_Mpos_ACGTACAAGGTCGTAG-1dCA2_MI_GFPve_pos  0.1545604167
## dCA2_Mpos_ACGTCCTCACGCGGTT-1dCA2_MI_GFPve_pos  0.9741306824
## dCA2_Mpos_ACGTCCTTCCGGACTG-1dCA2_MI_GFPve_pos -1.4630981761
## dCA2_Mpos_ACTACGATCGGCTATA-1dCA2_MI_GFPve_pos  1.4812070054
## dCA2_Mpos_ACTACGATCTTGAGCA-1dCA2_MI_GFPve_pos -0.2204056757
## dCA2_Mpos_ACTATGGGTTCGCGTG-1dCA2_MI_GFPve_pos  0.5906196458
## dCA2_Mpos_ACTCTCGAGCCTGAGA-1dCA2_MI_GFPve_pos  0.3351668191
## dCA2_Mpos_ACTTCCGCAGCTTTGA-1dCA2_MI_GFPve_pos -1.3698010522
## dCA2_Mpos_ACTTCGCTCGAGAACG-1dCA2_MI_GFPve_pos  1.3370190782
## dCA2_Mpos_ACTTCGCTCTGGTGGC-1dCA2_MI_GFPve_pos -0.1288045126
## dCA2_Mpos_AGAAATGGTCTTGCTC-1dCA2_MI_GFPve_pos  0.6638497633
## dCA2_Mpos_AGAACCTAGAAGGCTC-1dCA2_MI_GFPve_pos  0.7285500688
## dCA2_Mpos_AGACTCACACGGGTAA-1dCA2_MI_GFPve_pos  0.6014996094
## dCA2_Mpos_AGAGAGCCAAAGGAGA-1dCA2_MI_GFPve_pos  0.3344415945
## dCA2_Mpos_AGAGAGCCAGAATTCC-1dCA2_MI_GFPve_pos  1.0871879977
## dCA2_Mpos_AGATGAAGTCTTTCTA-1dCA2_MI_GFPve_pos -0.3427988368
## dCA2_Mpos_AGATGCTGTGCCTACG-1dCA2_MI_GFPve_pos  1.2634479922
## dCA2_Mpos_AGCGATTAGACTTCCA-1dCA2_MI_GFPve_pos  1.9804928226
## dCA2_Mpos_AGCGCCAAGGTAGCAC-1dCA2_MI_GFPve_pos  0.0894074689
## dCA2_Mpos_AGCGTCGCAGTTGGTT-1dCA2_MI_GFPve_pos  0.0892768155
## dCA2_Mpos_AGCGTCGCATCCTGTC-1dCA2_MI_GFPve_pos  0.2147177530
## dCA2_Mpos_AGCGTCGGTCAAATCC-1dCA2_MI_GFPve_pos -1.3135478574
## dCA2_Mpos_AGGACTTAGTTTCGAC-1dCA2_MI_GFPve_pos -0.6214537698
## dCA2_Mpos_AGGAGGTAGACCTGGA-1dCA2_MI_GFPve_pos -1.0800312835
## dCA2_Mpos_AGGATAACATCTCAAG-1dCA2_MI_GFPve_pos  0.0623914597
## dCA2_Mpos_AGGCATTAGGTGGCTA-1dCA2_MI_GFPve_pos  1.8877819699
## dCA2_Mpos_AGGTCATTCGATACTG-1dCA2_MI_GFPve_pos  0.5901804191
## dCA2_Mpos_AGTACCAAGCGCCGTT-1dCA2_MI_GFPve_pos -1.2599754411
## dCA2_Mpos_AGTAGCTTCCTCTTTC-1dCA2_MI_GFPve_pos -0.3135337907
## dCA2_Mpos_AGTAGTCAGCGTCGAA-1dCA2_MI_GFPve_pos -1.0105699378
## dCA2_Mpos_AGTCATGTCTGTAAGC-1dCA2_MI_GFPve_pos  0.7633744401
## dCA2_Mpos_AGTCTCCAGCTAGAAT-1dCA2_MI_GFPve_pos -0.8791519481
```

```
## dCA2_Mpos_AGTCTCCAGTTGCTCA-1dCA2_MI_GFPve_pos  2.1493205947
## dCA2_Mpos_AGTCTCCCAATGACCT-1dCA2_MI_GFPve_pos  0.7637119216
## dCA2_Mpos_AGTGCCGGTGTAAACA-1dCA2_MI_GFPve_pos -0.9852960187
## dCA2_Mpos_AGTGTTGGTACGCTAT-1dCA2_MI_GFPve_pos -0.7510580498
## dCA2_Mpos_AGTTAGCTCTGGACTA-1dCA2_MI_GFPve_pos -0.2525775808
## dCA2_Mpos_ATACCGAAGGCGACAT-1dCA2_MI_GFPve_pos -0.7907582002
## dCA2_Mpos_ATACCGATCATCGACA-1dCA2_MI_GFPve_pos  1.4729365033
## dCA2_Mpos_ATACCTTCACAAGGTG-1dCA2_MI_GFPve_pos  0.3822285158
## dCA2_Mpos_ATAGGCTAGAAACCCG-1dCA2_MI_GFPve_pos  1.8976435584
## dCA2_Mpos_ATCACGACACCAGCGT-1dCA2_MI_GFPve_pos -1.6572901088
## dCA2_Mpos_ATCAGGTAGCGTGAAC-1dCA2_MI_GFPve_pos -1.2779769975
## dCA2_Mpos_ATCCACCGTGGGACAT-1dCA2_MI_GFPve_pos -0.8530470806
## dCA2_Mpos_ATCCATTGTAGTACGG-1dCA2_MI_GFPve_pos  0.2466080261
## dCA2_Mpos_ATCCTATGTCGAATTC-1dCA2_MI_GFPve_pos  0.9122009796
## dCA2_Mpos_ATCGATGCACCCAATA-1dCA2_MI_GFPve_pos  0.1334051301
## dCA2_Mpos_ATCGGATGTCCACAGC-1dCA2_MI_GFPve_pos -1.3789390403
## dCA2_Mpos_ATCGTAGAGCCTGTCG-1dCA2_MI_GFPve_pos -0.4233552891
## dCA2_Mpos_ATGAAAGTCTACCCAC-1dCA2_MI_GFPve_pos -0.8369429189
## dCA2_Mpos_ATGACCACATGCCGCA-1dCA2_MI_GFPve_pos  1.7334587497
## dCA2_Mpos_ATGATCGCATGTGGCC-1dCA2_MI_GFPve_pos -1.1509926396
## dCA2_Mpos_ATGCATGTCACACGAT-1dCA2_MI_GFPve_pos  1.3467773122
## dCA2_Mpos_ATGTCCCTCTGTGCAA-1dCA2_MI_GFPve_pos  0.2323942107
## dCA2_Mpos_ATGTCTTTCGCTGTCT-1dCA2_MI_GFPve_pos  0.8694356603
## dCA2_Mpos_ATTATCCAGCTGTCCG-1dCA2_MI_GFPve_pos -1.1538391190
## dCA2_Mpos_ATTCACTTCACCGGTG-1dCA2_MI_GFPve_pos -0.8601887303
## dCA2_Mpos_ATTCCTATCCAAGCTA-1dCA2_MI_GFPve_pos -0.1454581695
## dCA2_Mpos_ATTCTTGGTCAGTCTA-1dCA2_MI_GFPve_pos  0.7750805539
## dCA2_Mpos_ATTCTTGTCTTTGCGC-1dCA2_MI_GFPve_pos  0.4775691551
## dCA2_Mpos_ATTGGGTTCTGACGCG-1dCA2_MI_GFPve_pos  1.2814544362
## dCA2_Mpos_ATTGTTCAGCCAAGCA-1dCA2_MI_GFPve_pos -0.1986987072
## dCA2_Mpos_ATTTACCAGAAGCGAA-1dCA2_MI_GFPve_pos -0.0350604283
## dCA2_Mpos_ATTTCACCAAGTCGTT-1dCA2_MI_GFPve_pos -1.5151879865
## dCA2_Mpos_ATTTCACTCGTCGCTT-1dCA2_MI_GFPve_pos  0.6236442846
## dCA2_Mpos_CAACAGTGTCCCGGTA-1dCA2_MI_GFPve_pos  0.1682162953
## dCA2_Mpos_CAACCTCTCTAGAGCT-1dCA2_MI_GFPve_pos  0.7731755895
## dCA2_Mpos_CAACGATCAGGCATTT-1dCA2_MI_GFPve_pos -1.7489154416
## dCA2_Mpos_CAAGCTAAGGCGTTAG-1dCA2_MI_GFPve_pos -0.3370188552
## dCA2_Mpos_CAAGGGAGTGCAACGA-1dCA2_MI_GFPve_pos  0.4493274612
## dCA2_Mpos_CAATCGAAGAGCCCAA-1dCA2_MI_GFPve_pos  1.2658218068
## dCA2_Mpos_CACCAAAAGACCAAGC-1dCA2_MI_GFPve_pos -1.2669639664
## dCA2_Mpos_CACGGGTAGCGCTTCG-1dCA2_MI_GFPve_pos  0.9517511052
## dCA2_Mpos_CACTGAAGTCAAGGCA-1dCA2_MI_GFPve_pos -0.9161710816
## dCA2_Mpos_CAGATACCACCGGAAA-1dCA2_MI_GFPve_pos  0.0034125296
## dCA2_Mpos_CAGATCAAGTCGAATA-1dCA2_MI_GFPve_pos -1.2247902232
## dCA2_Mpos_CAGATTGAGATGGTCG-1dCA2_MI_GFPve_pos -1.0470237809
## dCA2_Mpos_CAGCGTGAGGGATGTC-1dCA2_MI_GFPve_pos -0.2219995933
## dCA2_Mpos_CAGTGCGAGGAGCAAA-1dCA2_MI_GFPve_pos -1.7163014489
## dCA2_Mpos_CATACAGGTCTTGCGG-1dCA2_MI_GFPve_pos -0.6030743795
## dCA2_Mpos_CATCGCTTCTTTCGAT-1dCA2_MI_GFPve_pos  0.3302827102
## dCA2_Mpos_CATTCATGTAGGACTG-1dCA2_MI_GFPve_pos -0.1918603855
## dCA2_Mpos_CATTCATGTCTCGGAC-1dCA2_MI_GFPve_pos -1.0755031186
## dCA2_Mpos_CATTCCGTCGGACAAG-1dCA2_MI_GFPve_pos -0.7949751097
## dCA2_Mpos_CCAAGCGCACGTATAC-1dCA2_MI_GFPve_pos  2.3821463508
## dCA2_Mpos_CCAATTTTCGTGTTCC-1dCA2_MI_GFPve_pos  0.4592400414
## dCA2_Mpos_CCACGAGAGACTCCGC-1dCA2_MI_GFPve_pos  1.3666732234
## dCA2_Mpos_CCACTTGTCCTAGAGT-1dCA2_MI_GFPve_pos  0.2123220560
## dCA2_Mpos_CCCGAAGTCATGGAGG-1dCA2_MI_GFPve_pos  0.1815496606
## dCA2_Mpos_CCCGAAGTCCGGTAAT-1dCA2_MI_GFPve_pos  2.3558044357
## dCA2_Mpos_CCCTCTCAGTAAGAGG-1dCA2_MI_GFPve_pos  1.3656662625
## dCA2_Mpos_CCGATCTTCAGCAGAG-1dCA2_MI_GFPve_pos -0.1604444760
## dCA2_Mpos_CCGATGGGTATTTCCT-1dCA2_MI_GFPve_pos  1.8519797248
## dCA2_Mpos_CCGATGGTCCAAGCCG-1dCA2_MI_GFPve_pos -1.4588543254
## dCA2_Mpos_CCGCAAGTCGGCTGAC-1dCA2_MI_GFPve_pos  0.4175513906
## dCA2_Mpos_CCGGACAAGTGCAACG-1dCA2_MI_GFPve_pos  0.5233851296
## dCA2_Mpos_CCGTGAGCATGTGGTT-1dCA2_MI_GFPve_pos -0.5415443855
## dCA2_Mpos_CCTCACAAGTATAGGT-1dCA2_MI_GFPve_pos  0.1283645478
## dCA2_Mpos_CCTCAGTGTAGTCACT-1dCA2_MI_GFPve_pos  1.3375161809
## dCA2_Mpos_CCTCCTCCAGTAGTGG-1dCA2_MI_GFPve_pos  1.7267349881
## dCA2_Mpos_CCTCTAGTCTTCGTAT-1dCA2_MI_GFPve_pos -0.5220275002
## dCA2_Mpos_CCTCTCCAGGTCTTTG-1dCA2_MI_GFPve_pos -0.7892445999
```

```
## dCA2_Mpos_CCTGCATTCGGTCACG-1dCA2_MI_GFPve_pos -1.2298419553
## dCA2_Mpos_CCTTCAGAGTCACGCC-1dCA2_MI_GFPve_pos  1.9185779018
## dCA2_Mpos_CCTTGTGGTATAATGG-1dCA2_MI_GFPve_pos  0.4314331335
## dCA2_Mpos_CCTTGTGTCACGATAC-1dCA2_MI_GFPve_pos  0.6232614440
## dCA2_Mpos_CGACAGCTCGCTAGCG-1dCA2_MI_GFPve_pos  1.4848008079
## dCA2_Mpos_CGAGTGCGTAGGTCAG-1dCA2_MI_GFPve_pos  1.1509760541
## dCA2_Mpos_CGATCGGGTGTGCCTG-1dCA2_MI_GFPve_pos  1.5224035901
## dCA2_Mpos_CGATGCGAGCGATGGT-1dCA2_MI_GFPve_pos -1.1258783418
## dCA2_Mpos_CGATGGCCAACAAAGT-1dCA2_MI_GFPve_pos  0.2529478145
## dCA2_Mpos_CGCAGGTCAAGTCGTT-1dCA2_MI_GFPve_pos -1.2239562350
## dCA2_Mpos_CGCATGGTCGGTGTTA-1dCA2_MI_GFPve_pos  0.9168093723
## dCA2_Mpos_CGCCAGATCGCCGATG-1dCA2_MI_GFPve_pos  0.8647009653
## dCA2_Mpos_CGCGTGAAGAGTATAC-1dCA2_MI_GFPve_pos -0.3277553099
## dCA2_Mpos_CGGGACTGTAGTTAGA-1dCA2_MI_GFPve_pos -0.8202259022
## dCA2_Mpos_CGGGTGTTCTGATGGT-1dCA2_MI_GFPve_pos -0.8115184980
## dCA2_Mpos_CGGTCAGAGATTGCGG-1dCA2_MI_GFPve_pos  0.0930216831
## dCA2_Mpos_CGGTCAGGTTACGATC-1dCA2_MI_GFPve_pos  1.6744830531
## dCA2_Mpos_CGTAGTAGTGCTCTTC-1dCA2_MI_GFPve_pos  1.2578072471
## dCA2_Mpos_CGTCAAACACGACAAG-1dCA2_MI_GFPve_pos  2.2955145759
## dCA2_Mpos_CGTGTCTAGTGGCCTC-1dCA2_MI_GFPve_pos  0.1229788565
## dCA2_Mpos_CGTTGGGGTAGACGTG-1dCA2_MI_GFPve_pos -0.1584961193
## dCA2_Mpos_CTAAGTGGTGTAAATG-1dCA2_MI_GFPve_pos  1.0700517220
## dCA2_Mpos_CTACGGGTCTAGAGCT-1dCA2_MI_GFPve_pos -0.6672344881
## dCA2_Mpos_CTAGACAAGTGATAAC-1dCA2_MI_GFPve_pos  2.2883737010
## dCA2_Mpos_CTCAACCAGGGATCTG-1dCA2_MI_GFPve_pos  1.3391964359
## dCA2_Mpos_CTCAAGAGTCGTGATT-1dCA2_MI_GFPve_pos -0.5754008966
## dCA2_Mpos_CTCATCGGTACGAGCA-1dCA2_MI_GFPve_pos  1.0405185742
## dCA2_Mpos_CTCATGCTCTCATAGG-1dCA2_MI_GFPve_pos  0.5884715838
## dCA2_Mpos_CTCGAGGCATGAAAGT-1dCA2_MI_GFPve_pos -0.9394452649
## dCA2_Mpos_CTGAGCGAGGAGAATG-1dCA2_MI_GFPve_pos  0.2695232851
## dCA2_Mpos_CTGCAGGCAATTGCGT-1dCA2_MI_GFPve_pos  2.6430163306
## dCA2_Mpos_CTGCATCAGGACTAAT-1dCA2_MI_GFPve_pos -0.3946091312
## dCA2_Mpos_CTGCCATAGGAGATAG-1dCA2_MI_GFPve_pos  1.3588704986
## dCA2_Mpos_CTGCCATTCCTCGATC-1dCA2_MI_GFPve_pos -1.1447804051
## dCA2_Mpos_CTGCGAGAGCCTGCCA-1dCA2_MI_GFPve_pos -1.1087821799
## dCA2_Mpos_CTGGACGCACGTGAGA-1dCA2_MI_GFPve_pos -0.6206398087
## dCA2_Mpos_CTGGACGGTGACTAAA-1dCA2_MI_GFPve_pos  0.1614493911
## dCA2_Mpos_CTGGACGTCAGACCTA-1dCA2_MI_GFPve_pos  0.3095672948
## dCA2_Mpos_CTGGCAGCATGCGTGC-1dCA2_MI_GFPve_pos -1.3924183923
## dCA2_Mpos_CTGGTCTTCCAACCGG-1dCA2_MI_GFPve_pos  0.5377635879
## dCA2_Mpos_CTGGTCTTCGAGATAA-1dCA2_MI_GFPve_pos  0.8745667857
## dCA2_Mpos_CTGTAGACATGAGATA-1dCA2_MI_GFPve_pos  1.5760154647
## dCA2_Mpos_CTGTGAATCCGGACGT-1dCA2_MI_GFPve_pos  0.5598129851
## dCA2_Mpos_CTGTGGGCATGCAGGA-1dCA2_MI_GFPve_pos -0.3712288516
## dCA2_Mpos_CTGTGGGTCATGCCCT-1dCA2_MI_GFPve_pos -0.7247943359
## dCA2_Mpos_CTTAGGATCTCCGAGG-1dCA2_MI_GFPve_pos -0.3041482645
## dCA2_Mpos_CTTCGGTCAGGACTTT-1dCA2_MI_GFPve_pos -1.2255882102
## dCA2_Mpos_CTTGAGAGTTCAGCGC-1dCA2_MI_GFPve_pos -0.5360407310
## dCA2_Mpos_CTTTCAAAGGAGTCTG-1dCA2_MI_GFPve_pos -1.1798605996
## dCA2_Mpos_GAAACCTCACATGGTT-1dCA2_MI_GFPve_pos  0.3879645688
## dCA2_Mpos_GAAGAATTCGGCTATA-1dCA2_MI_GFPve_pos  1.1931388301
## dCA2_Mpos_GAAGCGAAGACTTAAG-1dCA2_MI_GFPve_pos  1.1991799993
## dCA2_Mpos_GAATAGACAGATACCT-1dCA2_MI_GFPve_pos -1.3028001862
## dCA2_Mpos_GAATAGAGTGCGACAA-1dCA2_MI_GFPve_pos -1.1037992316
## dCA2_Mpos_GAATCACAGGCATGCA-1dCA2_MI_GFPve_pos  1.9792570991
## dCA2_Mpos_GAATCGTTCGCGGACT-1dCA2_MI_GFPve_pos -0.9093172627
## dCA2_Mpos_GACCCAGCAAGAGGCT-1dCA2_MI_GFPve_pos -1.6502031165
## dCA2_Mpos_GACCCTTCACCTAAAC-1dCA2_MI_GFPve_pos  1.8127180261
## dCA2_Mpos_GACTCTCGTAAGGCTG-1dCA2_MI_GFPve_pos  2.1979806346
## dCA2_Mpos_GACTGATTCGCGTAGC-1dCA2_MI_GFPve_pos  2.3148610515
## dCA2_Mpos_GAGACCCCACTGGCGT-1dCA2_MI_GFPve_pos -0.2904493588
## dCA2_Mpos_GAGAGGTAGTTGCATC-1dCA2_MI_GFPve_pos  0.5759121103
## dCA2_Mpos_GAGAGGTCAAGATGTA-1dCA2_MI_GFPve_pos -0.3011052090
## dCA2_Mpos_GAGATGGTCTCCTGTG-1dCA2_MI_GFPve_pos -0.1799195486
## dCA2_Mpos_GAGCTGCCAGCTAACT-1dCA2_MI_GFPve_pos -0.4606370407
## dCA2_Mpos_GAGCTGCGTAGCTTAC-1dCA2_MI_GFPve_pos  1.1827529592
## dCA2_Mpos_GAGCTGCTCCTATTGT-1dCA2_MI_GFPve_pos -0.0819176960
## dCA2_Mpos_GAGGGTAAGTAGACCG-1dCA2_MI_GFPve_pos -1.5087838250
## dCA2_Mpos_GAGTCTAGTGATACCT-1dCA2_MI_GFPve_pos -0.5146918374
## dCA2_Mpos_GAGTGAGAGTCAGCGA-1dCA2_MI_GFPve_pos -0.3199307996
```

```
## dCA2_Mpos_GAGTGTTAGGGCGAGA-1dCA2_MI_GFPve_pos  1.0355715079
## dCA2_Mpos_GAGTTACAGAGCTGAC-1dCA2_MI_GFPve_pos  1.2436690253
## dCA2_Mpos_GAGTTTGGTTGGTACT-1dCA2_MI_GFPve_pos  0.1738923786
## dCA2_Mpos_GATCAGTAGCGAGAAA-1dCA2_MI_GFPve_pos -1.0475181418
## dCA2_Mpos_GATCAGTAGGCAGGTT-1dCA2_MI_GFPve_pos  0.0713049506
## dCA2_Mpos_GATCAGTAGTTCGGTT-1dCA2_MI_GFPve_pos -0.0239716696
## dCA2_Mpos_GATTCTTTCAGAATAG-1dCA2_MI_GFPve_pos -0.8422952371
## dCA2_Mpos_GCAGTTATCGGAATGG-1dCA2_MI_GFPve_pos  1.9376219434
## dCA2_Mpos_GCATCGGGTTGCTCCT-1dCA2_MI_GFPve_pos -0.1298820900
## dCA2_Mpos_GCCATGGCAAGGCGTA-1dCA2_MI_GFPve_pos -0.9244139272
## dCA2_Mpos_GCCATGGGTTCCTACC-1dCA2_MI_GFPve_pos -0.6392796117
## dCA2_Mpos_GCCGATGTCGCGGACT-1dCA2_MI_GFPve_pos  0.5657019836
## dCA2_Mpos_GCGATCGAGAGCTGCA-1dCA2_MI_GFPve_pos  0.8386508745
## dCA2_Mpos_GCGGATCAGAGAATCT-1dCA2_MI_GFPve_pos  1.1519374770
## dCA2_Mpos_GCGGATCTCAACGTGT-1dCA2_MI_GFPve_pos -0.6840555745
## dCA2_Mpos_GCGTGCAAGGTTCCAT-1dCA2_MI_GFPve_pos -0.7668773013
## dCA2_Mpos_GCGTGCAAGTTTGGCT-1dCA2_MI_GFPve_pos -1.5657810050
## dCA2_Mpos_GCTTTCGGTAGTGGCA-1dCA2_MI_GFPve_pos -1.6411179381
## dCA2_Mpos_GGAACCCCAACAGCCC-1dCA2_MI_GFPve_pos  1.2122495097
## dCA2_Mpos_GGAACCCGTAGTCTGT-1dCA2_MI_GFPve_pos -1.3385722714
## dCA2_Mpos_GGAACCCTCTCTGGTC-1dCA2_MI_GFPve_pos -1.0468144494
## dCA2_Mpos_GGAATCTCAGGGCTTC-1dCA2_MI_GFPve_pos -0.1350155997
## dCA2_Mpos_GGAATGGGTGTTAGCT-1dCA2_MI_GFPve_pos  0.1829076630
## dCA2_Mpos_GGAGGTATCATCGGGC-1dCA2_MI_GFPve_pos -1.0170441943
## dCA2_Mpos_GGATCTAGTCGTGTTA-1dCA2_MI_GFPve_pos  1.0410962624
## dCA2_Mpos_GGATGTTAGGAAGTCC-1dCA2_MI_GFPve_pos  2.2317950648
## dCA2_Mpos_GGCGTCAGTGACCTGC-1dCA2_MI_GFPve_pos  1.7480294627
## dCA2_Mpos_GGGACCTAGTTACGGG-1dCA2_MI_GFPve_pos  0.0694717054
## dCA2_Mpos_GGGAGTAGTTCGAGCC-1dCA2_MI_GFPve_pos -1.5457737523
## dCA2_Mpos_GGGAGTATCAGCGTCG-1dCA2_MI_GFPve_pos -0.8593935448
## dCA2_Mpos_GGGCGTTTCTTACCAT-1dCA2_MI_GFPve_pos  0.5707656962
## dCA2_Mpos_GGGCTCAGTTCGGTAT-1dCA2_MI_GFPve_pos -0.4836737591
## dCA2_Mpos_GGGTAGAAGGAGAATG-1dCA2_MI_GFPve_pos -0.6319637972
## dCA2_Mpos_GGGTCTGAGGGATCTG-1dCA2_MI_GFPve_pos -0.3890712934
## dCA2_Mpos_GGGTCTGAGTAGACCG-1dCA2_MI_GFPve_pos -1.0750522691
## dCA2_Mpos_GGGTCTGGTCTACATG-1dCA2_MI_GFPve_pos  0.4878134650
## dCA2_Mpos_GGTAGAGAGCCTCATA-1dCA2_MI_GFPve_pos  0.2664000613
## dCA2_Mpos_GGTGAAGGTAACGATA-1dCA2_MI_GFPve_pos -0.8842784839
## dCA2_Mpos_GGTGTCGGTTCTCCAC-1dCA2_MI_GFPve_pos -0.3235352712
## dCA2_Mpos_GGTGTCGGTTTACCTT-1dCA2_MI_GFPve_pos  1.3921812696
## dCA2_Mpos_GGTGTCGTCGCGTCGA-1dCA2_MI_GFPve_pos -0.9149787503
## dCA2_Mpos_GGTTAACAGGAAAGTG-1dCA2_MI_GFPve_pos  1.3341298026
## dCA2_Mpos_GGTTCTCTCGCCAGTG-1dCA2_MI_GFPve_pos -0.1109563487
## dCA2_Mpos_GTAACCACATAGCTGT-1dCA2_MI_GFPve_pos  0.5101624650
## dCA2_Mpos_GTAGAGGAGGAAAGAC-1dCA2_MI_GFPve_pos  0.1690495012
## dCA2_Mpos_GTAGATCAGTCATACC-1dCA2_MI_GFPve_pos -1.1460027772
## dCA2_Mpos_GTAGCTAGTAATTGGA-1dCA2_MI_GFPve_pos -0.6360199529
## dCA2_Mpos_GTAGGTTCATTGTAGC-1dCA2_MI_GFPve_pos -0.7885663586
## dCA2_Mpos_GTCACTCTCGGTAGGA-1dCA2_MI_GFPve_pos -0.9124004918
## dCA2_Mpos_GTCCCATGTAGGGAGG-1dCA2_MI_GFPve_pos -0.8971291858
## dCA2_Mpos_GTCGAATGTACAAACA-1dCA2_MI_GFPve_pos -1.4374238330
## dCA2_Mpos_GTCGTTCGTGTTCCTC-1dCA2_MI_GFPve_pos  1.6448414249
## dCA2_Mpos_GTCTAGAGTGATTCTG-1dCA2_MI_GFPve_pos -0.0299359786
## dCA2_Mpos_GTCTCACCATCGAAGG-1dCA2_MI_GFPve_pos -0.0863290506
## dCA2_Mpos_GTCTCACGTACCTAGT-1dCA2_MI_GFPve_pos  1.6313174886
## dCA2_Mpos_GTGACGCAGTACGAGC-1dCA2_MI_GFPve_pos -0.7630557495
## dCA2_Mpos_GTGAGGATCGTTCCTG-1dCA2_MI_GFPve_pos  1.3622243327
## dCA2_Mpos_GTGCTTCGTTAGGACG-1dCA2_MI_GFPve_pos  2.0838156861
## dCA2_Mpos_GTGGAGACAAACCACT-1dCA2_MI_GFPve_pos  0.0039152128
## dCA2_Mpos_GTGTAACCAGATTTCG-1dCA2_MI_GFPve_pos -1.5380364734
## dCA2_Mpos_GTTACCCGTTCGGTAT-1dCA2_MI_GFPve_pos  0.0463966389
## dCA2_Mpos_GTTACGACACCGTCGA-1dCA2_MI_GFPve_pos  0.7153487128
## dCA2_Mpos_GTTAGACGTCTTGAAC-1dCA2_MI_GFPve_pos -0.5108936506
## dCA2_Mpos_GTTGAACAGTCACGCC-1dCA2_MI_GFPve_pos -0.9661116677
## dCA2_Mpos_GTTTACTCATGTCTAG-1dCA2_MI_GFPve_pos  0.7099572343
## dCA2_Mpos_TAACACGTCGTTCTCG-1dCA2_MI_GFPve_pos -0.6652265864
## dCA2_Mpos_TAACGACAGGAAACGA-1dCA2_MI_GFPve_pos  1.0894095821
## dCA2_Mpos_TAAGTCGGTAACTTCG-1dCA2_MI_GFPve_pos  0.6825433892
## dCA2_Mpos_TACAACGGTGTTAACC-1dCA2_MI_GFPve_pos  0.9074137134
## dCA2_Mpos_TACCCACGTTCTCCCA-1dCA2_MI_GFPve_pos -0.9753047305
```

```
## dCA2_Mpos_TACCTGCGTGTCATGT-1dCA2_MI_GFPve_pos  0.8943505806
## dCA2_Mpos_TACTTACAGCTCTGTA-1dCA2_MI_GFPve_pos -1.1986906606
## dCA2_Mpos_TACTTACCAGAAATTG-1dCA2_MI_GFPve_pos -0.0908935147
## dCA2_Mpos_TAGAGTCCAATGGGTG-1dCA2_MI_GFPve_pos  2.1545638961
## dCA2_Mpos_TAGAGTCCAGTCCGTG-1dCA2_MI_GFPve_pos -0.8836058455
## dCA2_Mpos_TAGAGTCGTACGTGAG-1dCA2_MI_GFPve_pos  0.2816711885
## dCA2_Mpos_TAGCACACAACCTATG-1dCA2_MI_GFPve_pos -1.0993666726
## dCA2_Mpos_TAGGAGGAGATTGCGG-1dCA2_MI_GFPve_pos  0.5812875551
## dCA2_Mpos_TAGTGCAGTTAGGCCC-1dCA2_MI_GFPve_pos -0.8595408278
## dCA2_Mpos_TAGTGCAGTTTCACTT-1dCA2_MI_GFPve_pos  0.4328168554
## dCA2_Mpos_TATATCCAGGTGAGAA-1dCA2_MI_GFPve_pos  0.6791030688
## dCA2_Mpos_TATCCTAAGCTCTTCC-1dCA2_MI_GFPve_pos  0.8102722091
## dCA2_Mpos_TATCCTACAGAGAAAG-1dCA2_MI_GFPve_pos  1.4195705575
## dCA2_Mpos_TATCTGTCATAGCACT-1dCA2_MI_GFPve_pos  0.2988125754
## dCA2_Mpos_TATCTTGGTCGGATTT-1dCA2_MI_GFPve_pos  0.3247229678
## dCA2_Mpos_TATGTTCGTTAGGACG-1dCA2_MI_GFPve_pos -1.5386065322
## dCA2_Mpos_TATTGGGTCTTCCAGC-1dCA2_MI_GFPve_pos  0.1772795541
## dCA2_Mpos_TCAAGACAGTTTGAGA-1dCA2_MI_GFPve_pos -0.0365515875
## dCA2_Mpos_TCAAGCAGTTGGCCGT-1dCA2_MI_GFPve_pos  2.8204982204
## dCA2_Mpos_TCACATTGTTAGGGAC-1dCA2_MI_GFPve_pos  0.3238432628
## dCA2_Mpos_TCACGCTCAATAGGGC-1dCA2_MI_GFPve_pos  0.1666577530
## dCA2_Mpos_TCACTATTCCTTGAAG-1dCA2_MI_GFPve_pos  2.6278092784
## dCA2_Mpos_TCAGCCTGTGTTCCTC-1dCA2_MI_GFPve_pos -1.2949324923
## dCA2_Mpos_TCAGGGCAGGTGATAT-1dCA2_MI_GFPve_pos -0.9378995972
## dCA2_Mpos_TCAGTGAGTTAATGAG-1dCA2_MI_GFPve_pos  0.9508538169
## dCA2_Mpos_TCATTACCAATACGCT-1dCA2_MI_GFPve_pos  0.9996995253
## dCA2_Mpos_TCATTACTCTACGCGG-1dCA2_MI_GFPve_pos  2.3288235587
## dCA2_Mpos_TCATTCAGTCGTTGCG-1dCA2_MI_GFPve_pos  0.0507741531
## dCA2_Mpos_TCATTTGAGATGATTG-1dCA2_MI_GFPve_pos  2.0624651832
## dCA2_Mpos_TCATTTGTCGCCAACG-1dCA2_MI_GFPve_pos  1.0243233961
## dCA2_Mpos_TCCACGTCATATCTGG-1dCA2_MI_GFPve_pos -0.0310626405
## dCA2_Mpos_TCCAGAAAGTATTAGG-1dCA2_MI_GFPve_pos -1.0864499884
## dCA2_Mpos_TCCCAGTGTGGGCTCT-1dCA2_MI_GFPve_pos -0.1953255909
## dCA2_Mpos_TCCGATCAGTAGACAT-1dCA2_MI_GFPve_pos  0.4645197314
## dCA2_Mpos_TCCGTGTTCCCAGCGA-1dCA2_MI_GFPve_pos -0.1125151741
## dCA2_Mpos_TCCTGCAGTACAGCGA-1dCA2_MI_GFPve_pos -0.7681915837
## dCA2_Mpos_TCCTTCTGTACGGTTT-1dCA2_MI_GFPve_pos  0.7904581351
## dCA2_Mpos_TCCTTTCCAGCGTATT-1dCA2_MI_GFPve_pos  1.9631270093
## dCA2_Mpos_TCGACCTGTAAGTCAA-1dCA2_MI_GFPve_pos  1.3127078933
## dCA2_Mpos_TCGATTTAGGTGGTTG-1dCA2_MI_GFPve_pos -0.9455709534
## dCA2_Mpos_TCGCAGGAGCAGGGAG-1dCA2_MI_GFPve_pos  2.0524564904
## dCA2_Mpos_TCGCAGGTCGGCGATC-1dCA2_MI_GFPve_pos -0.0101859200
## dCA2_Mpos_TCGCTTGGTAGTCGGA-1dCA2_MI_GFPve_pos  2.2401356620
## dCA2_Mpos_TCGGGACCAGTAACCT-1dCA2_MI_GFPve_pos -0.2229725796
## dCA2_Mpos_TCGGGACTCGGAGATG-1dCA2_MI_GFPve_pos -0.9593898135
## dCA2_Mpos_TCGGTCTAGGTCGTGA-1dCA2_MI_GFPve_pos -1.0145967084
## dCA2_Mpos_TCGTAGATCATGAGAA-1dCA2_MI_GFPve_pos  2.1077911777
## dCA2_Mpos_TCGTGCTGTTCTAAGC-1dCA2_MI_GFPve_pos -1.5660102444
## dCA2_Mpos_TCGTGCTTCAAACGTC-1dCA2_MI_GFPve_pos -1.1926496106
## dCA2_Mpos_TCGTGGGTCAGTCAGT-1dCA2_MI_GFPve_pos  2.4021968764
## dCA2_Mpos_TCTATCACATGAATCC-1dCA2_MI_GFPve_pos -1.2982187348
## dCA2_Mpos_TCTCAGCAGAACGCGT-1dCA2_MI_GFPve_pos  1.2399185819
## dCA2_Mpos_TCTTCCTAGAGTCTGG-1dCA2_MI_GFPve_pos  0.3969753904
## dCA2_Mpos_TCTTCCTTCTCGAGTA-1dCA2_MI_GFPve_pos  0.6110570950
## dCA2_Mpos_TGAATCGCAAGACGAC-1dCA2_MI_GFPve_pos -0.9492043334
## dCA2_Mpos_TGAATGCTCGTCGACG-1dCA2_MI_GFPve_pos  2.0716563386
## dCA2_Mpos_TGAGCATCACTGCGTG-1dCA2_MI_GFPve_pos -0.9691726046
## dCA2_Mpos_TGAGCGCGTTTCGTGA-1dCA2_MI_GFPve_pos  0.3037878347
## dCA2_Mpos_TGAGGTTAGAGTCCGA-1dCA2_MI_GFPve_pos -1.5976108390
## dCA2_Mpos_TGAGGTTAGGTTCAGG-1dCA2_MI_GFPve_pos  1.6458859367
## dCA2_Mpos_TGAGGTTGTAAGATAC-1dCA2_MI_GFPve_pos -0.9311435300
## dCA2_Mpos_TGATTCTCACATTGTG-1dCA2_MI_GFPve_pos -1.2115738469
## dCA2_Mpos_TGATTTCAGCACCGTC-1dCA2_MI_GFPve_pos  0.5399704498
## dCA2_Mpos_TGCAGATCAAGTCATC-1dCA2_MI_GFPve_pos -1.2304668504
## dCA2_Mpos_TGCGACGCATAAGCGG-1dCA2_MI_GFPve_pos -0.6636405068
## dCA2_Mpos_TGCGGCAAGCTCCATA-1dCA2_MI_GFPve_pos -0.1239076632
## dCA2_Mpos_TGCGGCAGTCAGATTC-1dCA2_MI_GFPve_pos  2.0508426351
## dCA2_Mpos_TGCGGCAGTCTACAGT-1dCA2_MI_GFPve_pos -0.7489535290
## dCA2_Mpos_TGCGGGTGTACTAGCT-1dCA2_MI_GFPve_pos -0.1506736654
## dCA2_Mpos_TGCTCCAAGTGCCGAA-1dCA2_MI_GFPve_pos  0.6507587356
```

```
## dCA2_Mpos_TGCTTGCGTGATAGAT-1dCA2_MI_GFPve_pos -0.4463123756
## dCA2_Mpos_TGGAGGATCTAAGAAG-1dCA2_MI_GFPve_pos -0.7916201430
## dCA2_Mpos_TGGGAAGGTGTCCCTT-1dCA2_MI_GFPve_pos -0.8968277055
## dCA2_Mpos_TGGGAAGTCGTAGGAG-1dCA2_MI_GFPve_pos  1.9070370120
## dCA2_Mpos_TGGGAGAAGAAGTATC-1dCA2_MI_GFPve_pos -0.4213336783
## dCA2_Mpos_TGGGCGTCAAGCACAG-1dCA2_MI_GFPve_pos  1.9343842191
## dCA2_Mpos_TGGGCTGGTAAGCTCT-1dCA2_MI_GFPve_pos  0.3459438754
## dCA2_Mpos_TGGTGATAGTAAGCAT-1dCA2_MI_GFPve_pos -0.7047504264
## dCA2_Mpos_TGTACAGAGTCGGGAT-1dCA2_MI_GFPve_pos -1.2353695708
## dCA2_Mpos_TGTCCCATCCACGTCT-1dCA2_MI_GFPve_pos -1.5039722997
## dCA2_Mpos_TGTCCTGGTAACATGA-1dCA2_MI_GFPve_pos  2.3811561984
## dCA2_Mpos_TGTGAGTAGTTGCGAG-1dCA2_MI_GFPve_pos -0.1309468376
## dCA2_Mpos_TGTGTGATCTAGCAAC-1dCA2_MI_GFPve_pos -0.8724268156
## dCA2_Mpos_TGTTCCGCAGCCGGTT-1dCA2_MI_GFPve_pos -0.5471362549
## dCA2_Mpos_TGTTCTAGTACGCTAT-1dCA2_MI_GFPve_pos -0.2528226155
## dCA2_Mpos_TGTTCTAGTCAGTCGC-1dCA2_MI_GFPve_pos  1.9239958448
## dCA2_Mpos_TGTTCTAGTTAGGCTT-1dCA2_MI_GFPve_pos  0.5979713422
## dCA2_Mpos_TGTTTGTCACAACCGC-1dCA2_MI_GFPve_pos  0.4407403988
## dCA2_Mpos_TTACGTTAGGATACAT-1dCA2_MI_GFPve_pos  1.4364668054
## dCA2_Mpos_TTACTGTAGGCTCAAG-1dCA2_MI_GFPve_pos -1.1549829322
## dCA2_Mpos_TTAGTCTAGGTCATCT-1dCA2_MI_GFPve_pos -0.4363207298
## dCA2_Mpos_TTCAATCGTTGCCGCA-1dCA2_MI_GFPve_pos  1.7301388902
## dCA2_Mpos_TTCCGGTAGCCTATTG-1dCA2_MI_GFPve_pos  1.7344535512
## dCA2_Mpos_TTCCTAAGTTTCGATG-1dCA2_MI_GFPve_pos -0.0248381722
## dCA2_Mpos_TTCCTTCAGAGCAAGA-1dCA2_MI_GFPve_pos -1.2203598100
## dCA2_Mpos_TTCGATTGTTGGAGGT-1dCA2_MI_GFPve_pos  0.5802204830
## dCA2_Mpos_TTCGATTTCCGATGCG-1dCA2_MI_GFPve_pos -1.4769656735
## dCA2_Mpos_TTCTAACAGGAAGTGA-1dCA2_MI_GFPve_pos -1.2397986728
## dCA2_Mpos_TTCTGTATCGGACGTC-1dCA2_MI_GFPve_pos  0.6564028186
## dCA2_Mpos_TTCTTGAAGCAAGTGC-1dCA2_MI_GFPve_pos -0.2521886903
## dCA2_Mpos_TTCTTGACACATCATG-1dCA2_MI_GFPve_pos  0.0496054997
## dCA2_Mpos_TTGAGTGGTAAGGAGA-1dCA2_MI_GFPve_pos -1.5102170783
## dCA2_Mpos_TTGATGGAGTAACCTC-1dCA2_MI_GFPve_pos -0.2524972337
## dCA2_Mpos_TTGATGGAGTCGCTAT-1dCA2_MI_GFPve_pos  0.5780211908
## dCA2_Mpos_TTGCCTGCACGTCGGT-1dCA2_MI_GFPve_pos  1.7390994949
## dCA2_Mpos_TTGCGTCGTTGAAGTA-1dCA2_MI_GFPve_pos -0.3973870354
## dCA2_Mpos_TTGCTGCCATCTCATT-1dCA2_MI_GFPve_pos  0.3218653274
## dCA2_Mpos_TTGGGCGCACCTGAAT-1dCA2_MI_GFPve_pos  0.0686714982
## dCA2_Mpos_TTGTGGACACCTGCTT-1dCA2_MI_GFPve_pos  0.4757983309
## dCA2_Mpos_TTGTGGATCCATCGTC-1dCA2_MI_GFPve_pos -1.0034065324
## dCA2_Mpos_TTGTTTGAGGCTGAAC-1dCA2_MI_GFPve_pos -0.8918283659
## dCA2_Mpos_TTTCCTCAGCATGCGA-1dCA2_MI_GFPve_pos -0.8906712013
## dCA2_Mpos_TTTCGATCAGTGGGTA-1dCA2_MI_GFPve_pos  0.1147812844
## dCA2_Mpos_TTTGACTAGACCCTTA-1dCA2_MI_GFPve_pos  1.7062086982
## dCA2_Mpos_TTTGTTGCATGTGTCA-1dCA2_MI_GFPve_pos  0.7148829025
## dCA2_Fpos_AAACGAACAATTCTCT-1dCA2_FI_GFPve_pos  2.1172528190
## dCA2_Fpos_AAAGGATTCCGGTAAT-1dCA2_FI_GFPve_pos  0.9757254047
## dCA2_Fpos_AACAAGACAGAGCTAG-1dCA2_FI_GFPve_pos -1.1279851275
## dCA2_Fpos_AACCATGCAACTCATG-1dCA2_FI_GFPve_pos -0.9912277537
## dCA2_Fpos_AACCATGGTCCGATCG-1dCA2_FI_GFPve_pos  1.5706192178
## dCA2_Fpos_AACCTGACAATCCTTT-1dCA2_FI_GFPve_pos -0.4720100838
## dCA2_Fpos_AACGAAAAGGGCAGTT-1dCA2_FI_GFPve_pos -0.0825121182
## dCA2_Fpos_AACTTCTTCCAGCACG-1dCA2_FI_GFPve_pos  1.4305453223
## dCA2_Fpos_AAGCATCAGGGCCTCT-1dCA2_FI_GFPve_pos -0.2120419222
## dCA2_Fpos_AAGCATCTCCTATTTG-1dCA2_FI_GFPve_pos  1.6580991668
## dCA2_Fpos_AAGCGAGAGTCATTGC-1dCA2_FI_GFPve_pos  1.2684559745
## dCA2_Fpos_AAGCGAGGTAACCCGC-1dCA2_FI_GFPve_pos  0.8538247270
## dCA2_Fpos_AAGTGAAAGGAGCAAA-1dCA2_FI_GFPve_pos  0.2184634802
## dCA2_Fpos_AAGTTCGAGACAGTCG-1dCA2_FI_GFPve_pos  1.6434041185
## dCA2_Fpos_AAGTTCGGTCCTCCAT-1dCA2_FI_GFPve_pos  1.2081092519
## dCA2_Fpos_AATAGAGGTAACGGTG-1dCA2_FI_GFPve_pos  1.1456702871
## dCA2_Fpos_AATGAAGAGGACAACC-1dCA2_FI_GFPve_pos  0.3404241276
## dCA2_Fpos_AATGCCAGTTCATCTT-1dCA2_FI_GFPve_pos  0.6122625393
## dCA2_Fpos_AATGGAAAGGCAGGGA-1dCA2_FI_GFPve_pos  0.6783906144
## dCA2_Fpos_ACAAAGAGTTAAGGAT-1dCA2_FI_GFPve_pos  1.1429977340
## dCA2_Fpos_ACAACCATCACAATGC-1dCA2_FI_GFPve_pos  1.0415425224
## dCA2_Fpos_ACACGCGCATGAATAG-1dCA2_FI_GFPve_pos  0.9659901780
## dCA2_Fpos_ACACTGACATTACGGT-1dCA2_FI_GFPve_pos -1.2328695136
## dCA2_Fpos_ACCAACAGTTGTGTTG-1dCA2_FI_GFPve_pos  0.0779908669
## dCA2_Fpos_ACCCTTGAGGAACTCG-1dCA2_FI_GFPve_pos  0.9485000891
```

```
## dCA2_Fpos_ACCTGAAAGCTGAAGC-1dCA2_FI_GFPve_pos  0.3922023100
## dCA2_Fpos_ACGATGTAGGGAGGTG-1dCA2_FI_GFPve_pos  0.3712452573
## dCA2_Fpos_ACGATGTAGTAGTGCG-1dCA2_FI_GFPve_pos  1.5392594260
## dCA2_Fpos_ACGATGTGTCTGGTTA-1dCA2_FI_GFPve_pos  0.6646025104
## dCA2_Fpos_ACGCACGCAAAGTGTA-1dCA2_FI_GFPve_pos  0.6451008243
## dCA2_Fpos_ACGGAAGAGTTTCGAC-1dCA2_FI_GFPve_pos  2.4086878223
## dCA2_Fpos_ACGGGTCGTTGCTCGG-1dCA2_FI_GFPve_pos -0.8247034031
## dCA2_Fpos_ACGGTCGTCCTCACTG-1dCA2_FI_GFPve_pos -0.5660254436
## dCA2_Fpos_ACTATCTCAAACAGGC-1dCA2_FI_GFPve_pos -1.2984130460
## dCA2_Fpos_ACTCTCGAGCGCCATC-1dCA2_FI_GFPve_pos  1.4506056232
## dCA2_Fpos_ACTGCAAGTGACAGCA-1dCA2_FI_GFPve_pos  0.5150283498
## dCA2_Fpos_ACTTAGGTCGAAGCCC-1dCA2_FI_GFPve_pos -0.4520514088
## dCA2_Fpos_AGAAATGCACGCAAAG-1dCA2_FI_GFPve_pos -1.5879644233
## dCA2_Fpos_AGACAAATCGCGATCG-1dCA2_FI_GFPve_pos -0.1734537559
## dCA2_Fpos_AGACCATTCTGGGTCG-1dCA2_FI_GFPve_pos  0.4319764001
## dCA2_Fpos_AGACCCGCAACCACAT-1dCA2_FI_GFPve_pos  0.5388586444
## dCA2_Fpos_AGACCCGCAATCGTCA-1dCA2_FI_GFPve_pos -0.2505587416
## dCA2_Fpos_AGACCCGTCGTGCGAC-1dCA2_FI_GFPve_pos  0.1131324961
## dCA2_Fpos_AGAGAATCATCAGCTA-1dCA2_FI_GFPve_pos -1.3532168942
## dCA2_Fpos_AGAGAATGTAACGATA-1dCA2_FI_GFPve_pos  1.8854629916
## dCA2_Fpos_AGAGCCCCAGACGGAT-1dCA2_FI_GFPve_pos -1.2083951312
## dCA2_Fpos_AGAGCCCGTTAGAGTA-1dCA2_FI_GFPve_pos  1.1235095185
## dCA2_Fpos_AGATCCAGTCCGGATC-1dCA2_FI_GFPve_pos -0.6617876368
## dCA2_Fpos_AGATCGTTCCGCAAAT-1dCA2_FI_GFPve_pos  1.3126174134
## dCA2_Fpos_AGATGCTAGGTTTGAA-1dCA2_FI_GFPve_pos -0.0412334281
## dCA2_Fpos_AGATGCTCATTCGATG-1dCA2_FI_GFPve_pos  1.8280841035
## dCA2_Fpos_AGCCAATCAGAATCGG-1dCA2_FI_GFPve_pos  0.2510358167
## dCA2_Fpos_AGCGTATGTAGAGTTA-1dCA2_FI_GFPve_pos -0.7067790108
## dCA2_Fpos_AGCTCAAAGCGCATCC-1dCA2_FI_GFPve_pos  0.8489434642
## dCA2_Fpos_AGCTCAAGTACTGTTG-1dCA2_FI_GFPve_pos -0.9708507138
## dCA2_Fpos_AGCTCAATCCATTTCA-1dCA2_FI_GFPve_pos  1.0111975593
## dCA2_Fpos_AGGAATAAGGAATTAC-1dCA2_FI_GFPve_pos  0.5433975083
## dCA2_Fpos_AGGAGGTCACAAAGCG-1dCA2_FI_GFPve_pos -0.0304066318
## dCA2_Fpos_AGGGAGTCATCGGCCA-1dCA2_FI_GFPve_pos  0.8753666801
## dCA2_Fpos_AGGGCTCGTTCGAACT-1dCA2_FI_GFPve_pos -0.7461849528
## dCA2_Fpos_AGGGTCCGTCATAACC-1dCA2_FI_GFPve_pos  1.3686358852
## dCA2_Fpos_AGGTTACAGATCACTC-1dCA2_FI_GFPve_pos  0.1194290386
## dCA2_Fpos_AGGTTACGTCTATGAC-1dCA2_FI_GFPve_pos  0.3470695091
## dCA2_Fpos_AGGTTGTAGGCGTTGA-1dCA2_FI_GFPve_pos  2.5608584804
## dCA2_Fpos_AGTAACCTCCGCAACG-1dCA2_FI_GFPve_pos  1.3128161353
## dCA2_Fpos_AGTGACTAGAACCGCA-1dCA2_FI_GFPve_pos  0.2253044841
## dCA2_Fpos_AGTTAGCGTCGTAATC-1dCA2_FI_GFPve_pos -0.5735671001
## dCA2_Fpos_ATACCTTGTCCAGCCA-1dCA2_FI_GFPve_pos  1.1363758964
## dCA2_Fpos_ATACTTCAGTGGCGAT-1dCA2_FI_GFPve_pos  2.3330337924
## dCA2_Fpos_ATAGGCTGTACGTGAG-1dCA2_FI_GFPve_pos  0.6879238409
## dCA2_Fpos_ATCAGGTTCCGTGGTG-1dCA2_FI_GFPve_pos -0.8928163725
## dCA2_Fpos_ATCATTCAGGTTGAGC-1dCA2_FI_GFPve_pos -1.1495368558
## dCA2_Fpos_ATCCACCAGCAGAAAG-1dCA2_FI_GFPve_pos  0.1760186327
## dCA2_Fpos_ATCGATGCAACACGAG-1dCA2_FI_GFPve_pos -0.6694601851
## dCA2_Fpos_ATCGGATAGCATTTCG-1dCA2_FI_GFPve_pos -0.1637245792
## dCA2_Fpos_ATCGGATGTCAACCAT-1dCA2_FI_GFPve_pos  0.8325580281
## dCA2_Fpos_ATGAAAGTCCCGTGTT-1dCA2_FI_GFPve_pos  0.4469442887
## dCA2_Fpos_ATGAAAGTCTATACTC-1dCA2_FI_GFPve_pos -0.7502536255
## dCA2_Fpos_ATGAGGGTCAGAGCGA-1dCA2_FI_GFPve_pos  0.2939033431
## dCA2_Fpos_ATGATCGAGCGAGAAA-1dCA2_FI_GFPve_pos -0.4808126765
## dCA2_Fpos_ATGCATGCATGTTTGG-1dCA2_FI_GFPve_pos  1.4494420213
## dCA2_Fpos_ATGGAGGGTGGACCTC-1dCA2_FI_GFPve_pos  1.6283030433
## dCA2_Fpos_ATGGGTTCAAGTTCGT-1dCA2_FI_GFPve_pos  0.3979121966
## dCA2_Fpos_ATGGTTGGTAAGATTG-1dCA2_FI_GFPve_pos  0.7146558685
## dCA2_Fpos_ATTACCTCAGCAGATG-1dCA2_FI_GFPve_pos -1.3242577391
## dCA2_Fpos_ATTCCATAGGTTGACG-1dCA2_FI_GFPve_pos  1.4139031095
## dCA2_Fpos_ATTCCATCATAAGCAA-1dCA2_FI_GFPve_pos -0.8560726124
## dCA2_Fpos_ATTCTACAGGTTCCGC-1dCA2_FI_GFPve_pos  1.3706939143
## dCA2_Fpos_ATTCTTGCAAGAGTGC-1dCA2_FI_GFPve_pos  0.2561495823
## dCA2_Fpos_ATTGTTCAGCGTGAGT-1dCA2_FI_GFPve_pos  0.1385270094
## dCA2_Fpos_CAAAGAATCCGCACTT-1dCA2_FI_GFPve_pos  0.2961127532
## dCA2_Fpos_CAAAGAATCGATACAC-1dCA2_FI_GFPve_pos -0.6461163240
## dCA2_Fpos_CAAGACTTCGGTGAAG-1dCA2_FI_GFPve_pos  1.3812420291
## dCA2_Fpos_CAAGAGGTCTAGCCAA-1dCA2_FI_GFPve_pos -0.4986681419
## dCA2_Fpos_CAAGCTAAGGTTGACG-1dCA2_FI_GFPve_pos  0.7318528218
```

```
## dCA2_Fpos_CAAGCTAGTAGTTAGA-1dCA2_FI_GFPve_pos  1.5423389596
## dCA2_Fpos_CAAGCTATCCATGATG-1dCA2_FI_GFPve_pos  0.6337785644
## dCA2_Fpos_CAATACGGTACCTAAC-1dCA2_FI_GFPve_pos -0.8134249526
## dCA2_Fpos_CAATCGATCGCGTAGC-1dCA2_FI_GFPve_pos  1.9129973573
## dCA2_Fpos_CACAACAGTTCCTAGA-1dCA2_FI_GFPve_pos -0.6666477996
## dCA2_Fpos_CACAGATAGCGCACAA-1dCA2_FI_GFPve_pos  0.4139790458
## dCA2_Fpos_CACAGATAGGTTACAA-1dCA2_FI_GFPve_pos -0.0731332975
## dCA2_Fpos_CACATGACAAGCACCC-1dCA2_FI_GFPve_pos  1.3098699969
## dCA2_Fpos_CACGTGGTCGAGAAAT-1dCA2_FI_GFPve_pos  0.7073752207
## dCA2_Fpos_CACGTGGTCGTTCGCT-1dCA2_FI_GFPve_pos  1.4472012443
## dCA2_Fpos_CACTGGGAGATGGCAC-1dCA2_FI_GFPve_pos  0.4218099636
## dCA2_Fpos_CACTGTCAGCGTGAGT-1dCA2_FI_GFPve_pos  1.0413172168
## dCA2_Fpos_CATAAGCCATGTGCCG-1dCA2_FI_GFPve_pos  1.4045458955
## dCA2_Fpos_CATCGGGTCCTAGCTC-1dCA2_FI_GFPve_pos -0.3573214191
## dCA2_Fpos_CATCGTCTCGGAGATG-1dCA2_FI_GFPve_pos  0.1596883570
## dCA2_Fpos_CATGCGGAGGATCACG-1dCA2_FI_GFPve_pos  0.3988851232
## dCA2_Fpos_CATTCTACATGAAGCG-1dCA2_FI_GFPve_pos  0.4990677458
## dCA2_Fpos_CATTGTTTCCCGTGAG-1dCA2_FI_GFPve_pos -0.4834242540
## dCA2_Fpos_CATTTCATCGCTACAA-1dCA2_FI_GFPve_pos  0.4258713347
## dCA2_Fpos_CCAATGAGTGTCCATA-1dCA2_FI_GFPve_pos -0.0707423764
## dCA2_Fpos_CCCATTGGTGTGCTTA-1dCA2_FI_GFPve_pos  0.4274002833
## dCA2_Fpos_CCCTAACTCACCCTCA-1dCA2_FI_GFPve_pos  0.8626284522
## dCA2_Fpos_CCCTTAGAGACTTCCA-1dCA2_FI_GFPve_pos -0.2512039023
## dCA2_Fpos_CCCTTAGTCTGGCTGG-1dCA2_FI_GFPve_pos  1.1890361232
## dCA2_Fpos_CCGATCTCACTGGCGT-1dCA2_FI_GFPve_pos -1.3384310084
## dCA2_Fpos_CCGGTAGAGGCATCAG-1dCA2_FI_GFPve_pos -1.1603891927
## dCA2_Fpos_CCGGTAGAGTGATAGT-1dCA2_FI_GFPve_pos  0.0031887886
## dCA2_Fpos_CCGGTAGGTTGGAGAC-1dCA2_FI_GFPve_pos -1.6090760308
## dCA2_Fpos_CCGGTGAGTCCAATCA-1dCA2_FI_GFPve_pos  0.5340763551
## dCA2_Fpos_CCTAACCCATGACAAA-1dCA2_FI_GFPve_pos  0.9018872303
## dCA2_Fpos_CCTACGTAGTCACGAG-1dCA2_FI_GFPve_pos -1.3222407180
## dCA2_Fpos_CCTCCAATCCGTAATG-1dCA2_FI_GFPve_pos  0.7940379304
## dCA2_Fpos_CCTCCTCAGGTGCTAG-1dCA2_FI_GFPve_pos  2.3954725188
## dCA2_Fpos_CCTTGTGTCCTCAGAA-1dCA2_FI_GFPve_pos -0.7220444160
## dCA2_Fpos_CCTTTGGCAGCTAACT-1dCA2_FI_GFPve_pos  0.5077161116
## dCA2_Fpos_CCTTTGGTCGGTTCAA-1dCA2_FI_GFPve_pos -0.2959500926
## dCA2_Fpos_CGACAGCTCAGAGCAG-1dCA2_FI_GFPve_pos  0.5037425083
## dCA2_Fpos_CGATCGGAGACCAGAC-1dCA2_FI_GFPve_pos  0.3124209565
## dCA2_Fpos_CGCATGGGTAGGTGCA-1dCA2_FI_GFPve_pos  0.4400129241
## dCA2_Fpos_CGCCAGATCGGTCGAC-1dCA2_FI_GFPve_pos -1.7750530320
## dCA2_Fpos_CGCCATTAGCTGTCCG-1dCA2_FI_GFPve_pos  1.4339885635
## dCA2_Fpos_CGCGTGAGTTTCTATC-1dCA2_FI_GFPve_pos  0.0088423637
## dCA2_Fpos_CGGAACCGTTACCTTT-1dCA2_FI_GFPve_pos -0.9655025082
## dCA2_Fpos_CGGAACCTCCCTCTAG-1dCA2_FI_GFPve_pos -0.3236911970
## dCA2_Fpos_CGGAGAATCGAGATAA-1dCA2_FI_GFPve_pos  1.0457575244
## dCA2_Fpos_CGGCAGTAGGAGTACC-1dCA2_FI_GFPve_pos  1.5750218553
## dCA2_Fpos_CGGGTGTCACTACGGC-1dCA2_FI_GFPve_pos  0.9057239813
## dCA2_Fpos_CGTCCATTCAGTCTTT-1dCA2_FI_GFPve_pos  1.3075689000
## dCA2_Fpos_CGTGAATTCTCCACTG-1dCA2_FI_GFPve_pos -0.0368977862
## dCA2_Fpos_CGTGATAAGCTGCCTG-1dCA2_FI_GFPve_pos -0.3736925798
## dCA2_Fpos_CGTGCTTTCCGGTAAT-1dCA2_FI_GFPve_pos -1.0158047753
## dCA2_Fpos_CGTTCTGTCGAAGAAT-1dCA2_FI_GFPve_pos  1.6653572244
## dCA2_Fpos_CTAAGTGCAAGAGCTG-1dCA2_FI_GFPve_pos  2.4820318145
## dCA2_Fpos_CTACAGATCACTGATG-1dCA2_FI_GFPve_pos -0.5811052399
## dCA2_Fpos_CTACATTCACAAAGCG-1dCA2_FI_GFPve_pos  0.1162599379
## dCA2_Fpos_CTACTATCACGTCGGT-1dCA2_FI_GFPve_pos -0.1200504082
## dCA2_Fpos_CTATAGGAGTGACCTT-1dCA2_FI_GFPve_pos  2.1161487025
## dCA2_Fpos_CTCAAGACAGAGTGAC-1dCA2_FI_GFPve_pos -0.2701143401
## dCA2_Fpos_CTCACTGTCAGGAAGC-1dCA2_FI_GFPve_pos -1.1274613219
## dCA2_Fpos_CTCAGGGTCCCAGCGA-1dCA2_FI_GFPve_pos  0.4125026685
## dCA2_Fpos_CTCATCGGTAATGCTC-1dCA2_FI_GFPve_pos  1.2309596461
## dCA2_Fpos_CTCCAACTCACAACCA-1dCA2_FI_GFPve_pos  0.6077475471
## dCA2_Fpos_CTCCATGAGGCAGCTA-1dCA2_FI_GFPve_pos -0.8509233075
## dCA2_Fpos_CTCCCAATCGAAGGAC-1dCA2_FI_GFPve_pos  0.9695048851
## dCA2_Fpos_CTCCCTCCACTGAATC-1dCA2_FI_GFPve_pos -0.4779384213
## dCA2_Fpos_CTCTGGTAGCCGATAG-1dCA2_FI_GFPve_pos  0.7908317966
## dCA2_Fpos_CTGAGGCTCTAGGCCG-1dCA2_FI_GFPve_pos -0.8875925618
## dCA2_Fpos_CTGATCCAGAGCTTTC-1dCA2_FI_GFPve_pos -1.3763198930
## dCA2_Fpos_CTGATCCCAGACCCGT-1dCA2_FI_GFPve_pos -0.9787380772
## dCA2_Fpos_CTGCAGGGTAGCGCCT-1dCA2_FI_GFPve_pos  0.9311079902
```

```
## dCA2_Fpos_CTGCATCAGCGGGTTA-1dCA2_FI_GFPve_pos  0.3295459223
## dCA2_Fpos_CTGCATCAGGCTCAAG-1dCA2_FI_GFPve_pos  0.2628344220
## dCA2_Fpos_CTGCATCCAAGTGACG-1dCA2_FI_GFPve_pos -1.1759772378
## dCA2_Fpos_CTGCATCGTACGTGTT-1dCA2_FI_GFPve_pos -1.4273821193
## dCA2_Fpos_CTGCATCGTGACTCTA-1dCA2_FI_GFPve_pos -0.6278896409
## dCA2_Fpos_CTGCCTACAAGGTTGG-1dCA2_FI_GFPve_pos  0.1458548491
## dCA2_Fpos_CTGCGAGCAGCTGCCA-1dCA2_FI_GFPve_pos  2.3678932113
## dCA2_Fpos_CTGCTCAGTTAGGGAC-1dCA2_FI_GFPve_pos -1.2579604464
## dCA2_Fpos_CTGGTCTCACGGTAGA-1dCA2_FI_GFPve_pos -0.5629925209
## dCA2_Fpos_CTGGTCTTCTCGTCGT-1dCA2_FI_GFPve_pos  1.3543184919
## dCA2_Fpos_CTGTAGATCTCAGTCC-1dCA2_FI_GFPve_pos  0.1717412916
## dCA2_Fpos_CTGTCGTAGGATGAGA-1dCA2_FI_GFPve_pos  1.1114551944
## dCA2_Fpos_CTGTCGTAGTAAACGT-1dCA2_FI_GFPve_pos -0.5116226750
## dCA2_Fpos_CTGTCGTCATAAGATG-1dCA2_FI_GFPve_pos -0.9528635817
## dCA2_Fpos_CTTAGGATCTACCCAC-1dCA2_FI_GFPve_pos  1.0764118952
## dCA2_Fpos_CTTCCTTCATCATTGG-1dCA2_FI_GFPve_pos  0.0675966588
## dCA2_Fpos_CTTCTCTTCGATGGAG-1dCA2_FI_GFPve_pos -1.5711351710
## dCA2_Fpos_CTTGATTTCATGCATG-1dCA2_FI_GFPve_pos -0.7361161786
## dCA2_Fpos_GAACACTGTATGAGGC-1dCA2_FI_GFPve_pos -0.7070275026
## dCA2_Fpos_GAACTGTAGAGAAGGT-1dCA2_FI_GFPve_pos -1.0509902316
## dCA2_Fpos_GAATAGAGTACTGAGG-1dCA2_FI_GFPve_pos  1.2224073333
## dCA2_Fpos_GAATCACCAAGCGAGT-1dCA2_FI_GFPve_pos -0.9464051801
## dCA2_Fpos_GACACGCAGATAGTCA-1dCA2_FI_GFPve_pos  0.8525157494
## dCA2_Fpos_GACCCAGGTTCGTTCC-1dCA2_FI_GFPve_pos  1.5712861938
## dCA2_Fpos_GACCGTGAGCGCCGTT-1dCA2_FI_GFPve_pos  0.7567648215
## dCA2_Fpos_GACCGTGTCCAAGAGG-1dCA2_FI_GFPve_pos  1.3716921729
## dCA2_Fpos_GACGTTACATCGGAAG-1dCA2_FI_GFPve_pos  0.9529852790
## dCA2_Fpos_GAGACCCGTAGAGTTA-1dCA2_FI_GFPve_pos -1.2358857470
## dCA2_Fpos_GAGACTTGTTGTGCCG-1dCA2_FI_GFPve_pos  1.2023531121
## dCA2_Fpos_GAGGCAAAGATCGCTT-1dCA2_FI_GFPve_pos  0.7868335170
## dCA2_Fpos_GAGGCAACAGGAGGTT-1dCA2_FI_GFPve_pos  1.0489367766
## dCA2_Fpos_GAGGCCTGTTCCAGGC-1dCA2_FI_GFPve_pos -1.1131323653
## dCA2_Fpos_GAGGGTACAGTTAAAG-1dCA2_FI_GFPve_pos  0.4045688135
## dCA2_Fpos_GAGGGTATCCCGATCT-1dCA2_FI_GFPve_pos  1.3069586677
## dCA2_Fpos_GAGTGTTTCCTTCTTC-1dCA2_FI_GFPve_pos  0.5846168620
## dCA2_Fpos_GATCAGTTCATTGCGA-1dCA2_FI_GFPve_pos  0.4572276992
## dCA2_Fpos_GATCATGGTGGTTTAC-1dCA2_FI_GFPve_pos  0.5767265779
## dCA2_Fpos_GATGTTGGTGCATGTT-1dCA2_FI_GFPve_pos  1.2087181730
## dCA2_Fpos_GATTCTTGTTCGGGTC-1dCA2_FI_GFPve_pos  1.7604480905
## dCA2_Fpos_GCAACATAGGAAGTGA-1dCA2_FI_GFPve_pos  1.3404915256
## dCA2_Fpos_GCAACCGAGTCAGAGC-1dCA2_FI_GFPve_pos  2.1570193690
## dCA2_Fpos_GCACGGTGTAATCAGA-1dCA2_FI_GFPve_pos  1.5232911033
## dCA2_Fpos_GCACGTGTCTTGCGCT-1dCA2_FI_GFPve_pos  0.8566521925
## dCA2_Fpos_GCAGCCACACTTGGCG-1dCA2_FI_GFPve_pos  0.4619989914
## dCA2_Fpos_GCAGGCTGTCTGTCAA-1dCA2_FI_GFPve_pos -0.4173845726
## dCA2_Fpos_GCATCGGAGGGTTTCT-1dCA2_FI_GFPve_pos  2.4296460075
## dCA2_Fpos_GCCAGCAGTCTCACGG-1dCA2_FI_GFPve_pos -1.1942297297
## dCA2_Fpos_GCCAGGTGTTCTCTAT-1dCA2_FI_GFPve_pos  0.6339628023
## dCA2_Fpos_GCCAGGTTCAACACCA-1dCA2_FI_GFPve_pos  0.8223204536
## dCA2_Fpos_GCCAGTGAGCAAATGT-1dCA2_FI_GFPve_pos  1.9025560540
## dCA2_Fpos_GCCATGGTCAGCTCTC-1dCA2_FI_GFPve_pos  0.4398423356
## dCA2_Fpos_GCGAGAAGTATAGGGC-1dCA2_FI_GFPve_pos -1.1342134553
## dCA2_Fpos_GCGTGCAAGACTCATC-1dCA2_FI_GFPve_pos -0.8525368052
## dCA2_Fpos_GCGTGCACAGAATGTA-1dCA2_FI_GFPve_pos -0.6998061019
## dCA2_Fpos_GCGTGCAGTTGAGAGC-1dCA2_FI_GFPve_pos -0.5848000723
## dCA2_Fpos_GCTCAAAAGTCCTGCG-1dCA2_FI_GFPve_pos -1.0343831855
## dCA2_Fpos_GCTCAAATCGTAATGC-1dCA2_FI_GFPve_pos  1.2335729522
## dCA2_Fpos_GCTGCAGAGTCCGCGT-1dCA2_FI_GFPve_pos -0.7813454347
## dCA2_Fpos_GGAAGTGGTGTGTCGC-1dCA2_FI_GFPve_pos  0.6620104832
## dCA2_Fpos_GGAATGGCACATTCTT-1dCA2_FI_GFPve_pos  1.0190333647
## dCA2_Fpos_GGAGCAATCACTGTTT-1dCA2_FI_GFPve_pos  1.2608398122
## dCA2_Fpos_GGATCTACAATCAAGA-1dCA2_FI_GFPve_pos -0.7438091713
## dCA2_Fpos_GGATGTTAGACCTCAT-1dCA2_FI_GFPve_pos  0.1956108508
## dCA2_Fpos_GGCGTCATCTCGTGGG-1dCA2_FI_GFPve_pos  0.6700120491
## dCA2_Fpos_GGCTTGGGTCTCACGG-1dCA2_FI_GFPve_pos  1.1329759282
## dCA2_Fpos_GGCTTTCAGGTTAGTA-1dCA2_FI_GFPve_pos  1.7217820806
## dCA2_Fpos_GGGACAAAGTACTGGG-1dCA2_FI_GFPve_pos -0.3417658585
## dCA2_Fpos_GGGACCTAGCGACAGT-1dCA2_FI_GFPve_pos -0.8141350823
## dCA2_Fpos_GGGAGATCACTTGAAC-1dCA2_FI_GFPve_pos  0.8266653938
## dCA2_Fpos_GGGATGACAACTGAAA-1dCA2_FI_GFPve_pos -0.3195379453
```

```
## dCA2_Fpos_GGGCTACAGAAAGTCT-1dCA2_FI_GFPve_pos -1.1150337535
## dCA2_Fpos_GGGCTACGTCAGTTTG-1dCA2_FI_GFPve_pos  0.7334832472
## dCA2_Fpos_GGGCTCAGTAGCGTTT-1dCA2_FI_GFPve_pos  2.1893105430
## dCA2_Fpos_GGGTGTCCAAGTGCTT-1dCA2_FI_GFPve_pos  0.4695839209
## dCA2_Fpos_GGGTTATAGTTGCATC-1dCA2_FI_GFPve_pos  2.0704908294
## dCA2_Fpos_GGGTTTACAATCAAGA-1dCA2_FI_GFPve_pos -1.0945284444
## dCA2_Fpos_GGTAACTAGCTACGTT-1dCA2_FI_GFPve_pos  1.1448417825
## dCA2_Fpos_GGTGTCGGTGCCTTCT-1dCA2_FI_GFPve_pos -0.4312046247
## dCA2_Fpos_GTAACACAGTGGCAGT-1dCA2_FI_GFPve_pos  2.2721664828
## dCA2_Fpos_GTAATGCCATGCCGCA-1dCA2_FI_GFPve_pos -0.4733612614
## dCA2_Fpos_GTACAGTGTTCGGCGT-1dCA2_FI_GFPve_pos -0.4301867562
## dCA2_Fpos_GTAGAGGGTCAAGCGA-1dCA2_FI_GFPve_pos -0.5992025214
## dCA2_Fpos_GTAGATCCAGTCTTCC-1dCA2_FI_GFPve_pos -0.5882192331
## dCA2_Fpos_GTAGGTTGTGTGTGTT-1dCA2_FI_GFPve_pos  0.8537912292
## dCA2_Fpos_GTAGTACAGATACCAA-1dCA2_FI_GFPve_pos -0.0623206275
## dCA2_Fpos_GTATTGGAGTTATGGA-1dCA2_FI_GFPve_pos  1.3029996080
## dCA2_Fpos_GTATTGGTCGAACGGA-1dCA2_FI_GFPve_pos -1.1026218015
## dCA2_Fpos_GTATTTCTCCGTGCGA-1dCA2_FI_GFPve_pos  0.8316895885
## dCA2_Fpos_GTCAGCGGTATGGAAT-1dCA2_FI_GFPve_pos -0.2537818330
## dCA2_Fpos_GTCGTAAGTATGGAAT-1dCA2_FI_GFPve_pos -0.2317685860
## dCA2_Fpos_GTCTACCAGGGAGTTC-1dCA2_FI_GFPve_pos -0.9974973279
## dCA2_Fpos_GTCTTTATCGCTACGG-1dCA2_FI_GFPve_pos -1.0358741360
## dCA2_Fpos_GTGCGTGTCGCGTCGA-1dCA2_FI_GFPve_pos  1.8267871064
## dCA2_Fpos_GTGCTTCGTGAGTCAG-1dCA2_FI_GFPve_pos -0.1656872707
## dCA2_Fpos_GTGGAAGCAACTAGAA-1dCA2_FI_GFPve_pos  0.4860770625
## dCA2_Fpos_GTGGAAGGTCGTGGTC-1dCA2_FI_GFPve_pos -1.4626908379
## dCA2_Fpos_GTGTGATGTCTCAGGC-1dCA2_FI_GFPve_pos  2.3271703643
## dCA2_Fpos_GTGTGGCAGCGGTAGT-1dCA2_FI_GFPve_pos -0.2963653582
## dCA2_Fpos_GTGTTAGTCTTCACGC-1dCA2_FI_GFPve_pos -0.5686890560
## dCA2_Fpos_GTGTTCCGTTCTCAGA-1dCA2_FI_GFPve_pos -0.1130596774
## dCA2_Fpos_GTTAGTGGTCGCTCGA-1dCA2_FI_GFPve_pos  0.9946769399
## dCA2_Fpos_GTTCATTTCTTCGACC-1dCA2_FI_GFPve_pos -0.6855861026
## dCA2_Fpos_GTTGAACAGTCGAAGC-1dCA2_FI_GFPve_pos -0.0063646885
## dCA2_Fpos_GTTGCGGCAAAGCACG-1dCA2_FI_GFPve_pos  0.4355405194
## dCA2_Fpos_GTTGCGGCAGCTATTG-1dCA2_FI_GFPve_pos -0.7871996241
## dCA2_Fpos_GTTGTAGAGGAAAGGT-1dCA2_FI_GFPve_pos  1.0025620383
## dCA2_Fpos_TAACCAGCAGTTAGGG-1dCA2_FI_GFPve_pos  0.2846407664
## dCA2_Fpos_TAAGCGTCAATCGCAT-1dCA2_FI_GFPve_pos  0.2805869949
## dCA2_Fpos_TAATTCCGTCAAAGCG-1dCA2_FI_GFPve_pos -1.2027300673
## dCA2_Fpos_TACACCCTCACGGTCG-1dCA2_FI_GFPve_pos  0.7886972350
## dCA2_Fpos_TACACCCTCTGAGGCC-1dCA2_FI_GFPve_pos  0.6732658071
## dCA2_Fpos_TACACCCTCTGGTGGC-1dCA2_FI_GFPve_pos -0.7857143360
## dCA2_Fpos_TACAGGTGTACAATAG-1dCA2_FI_GFPve_pos  0.1256416419
## dCA2_Fpos_TACCCACCAACTCCAA-1dCA2_FI_GFPve_pos -0.6528903919
## dCA2_Fpos_TACCTCGAGGTTCTTG-1dCA2_FI_GFPve_pos  0.7746760172
## dCA2_Fpos_TACCTCGCATCCTGTC-1dCA2_FI_GFPve_pos  1.5884928626
## dCA2_Fpos_TACGCTCGTAGGTTTC-1dCA2_FI_GFPve_pos  0.4921590907
## dCA2_Fpos_TACGGTACACCTCAGG-1dCA2_FI_GFPve_pos  0.5541914028
## dCA2_Fpos_TACTTGTTCGTCTACC-1dCA2_FI_GFPve_pos  0.8277213496
## dCA2_Fpos_TAGACCAGTCTTTATC-1dCA2_FI_GFPve_pos -0.2576483565
## dCA2_Fpos_TAGATCGAGTTAGAAC-1dCA2_FI_GFPve_pos -1.2000143605
## dCA2_Fpos_TAGATCGTCAGGGATG-1dCA2_FI_GFPve_pos -1.3626613694
## dCA2_Fpos_TAGATCGTCCCTCTCC-1dCA2_FI_GFPve_pos  0.8744988364
## dCA2_Fpos_TAGCACAAGAAGCTCG-1dCA2_FI_GFPve_pos  0.3951666874
## dCA2_Fpos_TAGGTTGCATCTAACG-1dCA2_FI_GFPve_pos  0.0369647143
## dCA2_Fpos_TATCCTAGTAGACGGT-1dCA2_FI_GFPve_pos -0.4711046296
## dCA2_Fpos_TATCTGTTCCATCAGA-1dCA2_FI_GFPve_pos -0.0315650034
## dCA2_Fpos_TATTTCGGTTTCGGCG-1dCA2_FI_GFPve_pos -0.7631577330
## dCA2_Fpos_TCACACCAGAGCCGTA-1dCA2_FI_GFPve_pos -0.0141617882
## dCA2_Fpos_TCACACCAGGTTGGAC-1dCA2_FI_GFPve_pos -0.2500737088
## dCA2_Fpos_TCACGCTTCATCAGTG-1dCA2_FI_GFPve_pos -0.1965024846
## dCA2_Fpos_TCAGGGCAGTAAATGC-1dCA2_FI_GFPve_pos -0.8779389935
## dCA2_Fpos_TCAGGGCGTCCTGTTC-1dCA2_FI_GFPve_pos  0.6272377295
## dCA2_Fpos_TCAGTTTGTGAGCAGT-1dCA2_FI_GFPve_pos -1.0936199504
## dCA2_Fpos_TCATGCCAGTGTTCAC-1dCA2_FI_GFPve_pos  1.8845342321
## dCA2_Fpos_TCATTACCAGGAAGTC-1dCA2_FI_GFPve_pos  0.7145376724
## dCA2_Fpos_TCATTCAGTAGCGTCC-1dCA2_FI_GFPve_pos -0.6176740008
## dCA2_Fpos_TCATTGTCAGCACAGA-1dCA2_FI_GFPve_pos  0.7270165605
## dCA2_Fpos_TCATTTGCAGACCAGA-1dCA2_FI_GFPve_pos  0.2838155252
## dCA2_Fpos_TCCATCGGTGGGACAT-1dCA2_FI_GFPve_pos  0.6898752374
```

```
## dCA2_Fpos_TCCATCGTCTCGGTAA-1dCA2_FI_GFPve_pos  1.0953372640
## dCA2_Fpos_TCCCACAAGACAACAT-1dCA2_FI_GFPve_pos  0.1124558958
## dCA2_Fpos_TCCGAAAGTAACCAGG-1dCA2_FI_GFPve_pos -0.2913475710
## dCA2_Fpos_TCCTTCTAGTATGATG-1dCA2_FI_GFPve_pos  1.1692641896
## dCA2_Fpos_TCCTTCTCATCTCCCA-1dCA2_FI_GFPve_pos  0.0307408718
## dCA2_Fpos_TCGAACAAGCGCTGCT-1dCA2_FI_GFPve_pos -0.9060051518
## dCA2_Fpos_TCGACGGGTACATACC-1dCA2_FI_GFPve_pos -1.1851288157
## dCA2_Fpos_TCGATTTAGTTACGAA-1dCA2_FI_GFPve_pos  1.0002744717
## dCA2_Fpos_TCGCTCAAGCGGTATG-1dCA2_FI_GFPve_pos  1.4455842895
## dCA2_Fpos_TCGCTTGGTACGACAG-1dCA2_FI_GFPve_pos -0.3516646701
## dCA2_Fpos_TCGGATATCGAGAACG-1dCA2_FI_GFPve_pos  0.7722202462
## dCA2_Fpos_TCGGATATCTAGATCG-1dCA2_FI_GFPve_pos  0.6145390791
## dCA2_Fpos_TCGGTCTCATAACAGA-1dCA2_FI_GFPve_pos -0.4956548887
## dCA2_Fpos_TCGTAGAGTTGATGTC-1dCA2_FI_GFPve_pos -0.6762173849
## dCA2_Fpos_TCGTCCAGTAACGCGA-1dCA2_FI_GFPve_pos  1.2093011064
## dCA2_Fpos_TCGTGCTCACTCCTGT-1dCA2_FI_GFPve_pos  0.4225805623
## dCA2_Fpos_TCGTGCTGTATCTTCT-1dCA2_FI_GFPve_pos  2.3123044891
## dCA2_Fpos_TCTCAGCGTGTCCACG-1dCA2_FI_GFPve_pos  0.2230567334
## dCA2_Fpos_TCTCTGGAGAGTATAC-1dCA2_FI_GFPve_pos  0.7739731592
## dCA2_Fpos_TCTGCCAAGGCGAACT-1dCA2_FI_GFPve_pos  0.2976335359
## dCA2_Fpos_TCTGGCTTCGTGGACC-1dCA2_FI_GFPve_pos  0.7981153053
## dCA2_Fpos_TCTTCCTTCCGTACGG-1dCA2_FI_GFPve_pos -0.9781067448
## dCA2_Fpos_TCTTGCGCATCGAAGG-1dCA2_FI_GFPve_pos -0.3966333347
## dCA2_Fpos_TCTTGCGGTTCCTACC-1dCA2_FI_GFPve_pos  0.5187857074
## dCA2_Fpos_TGACCCTAGAGTCCGA-1dCA2_FI_GFPve_pos  0.6671067995
## dCA2_Fpos_TGACGCGTCGGCACTG-1dCA2_FI_GFPve_pos -0.5731790739
## dCA2_Fpos_TGAGACTGTCGCAGTC-1dCA2_FI_GFPve_pos -0.5948643761
## dCA2_Fpos_TGAGGAGAGTTGTCAC-1dCA2_FI_GFPve_pos  0.6393769306
## dCA2_Fpos_TGAGGAGTCCGGTAAT-1dCA2_FI_GFPve_pos  1.6937597913
## dCA2_Fpos_TGATCAGGTTAAGACA-1dCA2_FI_GFPve_pos  1.2054821176
## dCA2_Fpos_TGCACGGTCTGCTCTG-1dCA2_FI_GFPve_pos  1.5766788644
## dCA2_Fpos_TGCGACGAGCTTGTGT-1dCA2_FI_GFPve_pos  1.1466329021
## dCA2_Fpos_TGCGACGGTGTCCTAA-1dCA2_FI_GFPve_pos  1.0261904520
## dCA2_Fpos_TGCTCCACATCAGCTA-1dCA2_FI_GFPve_pos -0.5942882973
## dCA2_Fpos_TGCTCGTGTTCTAAGC-1dCA2_FI_GFPve_pos  1.6797221822
## dCA2_Fpos_TGCTCGTTCGTCAACA-1dCA2_FI_GFPve_pos  1.8778842611
## dCA2_Fpos_TGCTGAATCCGAGCTG-1dCA2_FI_GFPve_pos -0.1671684759
## dCA2_Fpos_TGGAGGAAGATGGCGT-1dCA2_FI_GFPve_pos -0.8582047897
## dCA2_Fpos_TGGGTTAGTAAGCGGT-1dCA2_FI_GFPve_pos -1.2871798354
## dCA2_Fpos_TGGTACATCAGACCGC-1dCA2_FI_GFPve_pos -0.8801999765
## dCA2_Fpos_TGGTGATGTAGGAGGG-1dCA2_FI_GFPve_pos  1.8678700847
## dCA2_Fpos_TGGTTAGTCGGTCAGC-1dCA2_FI_GFPve_pos  0.4479790551
## dCA2_Fpos_TGTCCCATCAAGCGTT-1dCA2_FI_GFPve_pos -0.8118701654
## dCA2_Fpos_TGTCCTGTCGCTAGCG-1dCA2_FI_GFPve_pos  1.6847164554
## dCA2_Fpos_TGTGGCGGTAGACGTG-1dCA2_FI_GFPve_pos -0.5605998116
## dCA2_Fpos_TGTTCATTCGACCACG-1dCA2_FI_GFPve_pos -0.2532118040
## dCA2_Fpos_TGTTTGTCAAACTCTG-1dCA2_FI_GFPve_pos  0.6319631857
## dCA2_Fpos_TTAATCCCAAGTACCT-1dCA2_FI_GFPve_pos  1.5125356836
## dCA2_Fpos_TTACAGGGTCCAAATC-1dCA2_FI_GFPve_pos  2.1629726810
## dCA2_Fpos_TTACAGGGTTAAAGTG-1dCA2_FI_GFPve_pos  1.1061611098
## dCA2_Fpos_TTAGGCAAGCACTCCG-1dCA2_FI_GFPve_pos -0.4033806401
## dCA2_Fpos_TTAGGCAGTATGGAAT-1dCA2_FI_GFPve_pos  0.4039215666
## dCA2_Fpos_TTAGGGTCAGAGTAAT-1dCA2_FI_GFPve_pos  2.0498336477
## dCA2_Fpos_TTCAATCCAAACCACT-1dCA2_FI_GFPve_pos  0.6711664719
## dCA2_Fpos_TTCAGGACAAATGGAT-1dCA2_FI_GFPve_pos  1.3882707280
## dCA2_Fpos_TTCCGTGAGCTGTGCC-1dCA2_FI_GFPve_pos  1.2197732848
## dCA2_Fpos_TTCCTCTAGGTAGTAT-1dCA2_FI_GFPve_pos  2.2115874213
## dCA2_Fpos_TTCTAACCAGGCAATG-1dCA2_FI_GFPve_pos -0.1268070864
## dCA2_Fpos_TTCTAGTAGGCTCTAT-1dCA2_FI_GFPve_pos  2.0250184459
## dCA2_Fpos_TTCTGTAAGTGATAAC-1dCA2_FI_GFPve_pos  0.3182049913
## dCA2_Fpos_TTCTGTAGTCTCTCCA-1dCA2_FI_GFPve_pos  0.1472071705
## dCA2_Fpos_TTGGATGAGACTTCAC-1dCA2_FI_GFPve_pos  2.2926118297
## dCA2_Fpos_TTGGGTACACTCCGGA-1dCA2_FI_GFPve_pos -0.7040785628
## dCA2_Fpos_TTGGGTATCAGAGTGG-1dCA2_FI_GFPve_pos -0.6094554740
## dCA2_Fpos_TTGGTTTAGGCTCTAT-1dCA2_FI_GFPve_pos  1.1247550172
## dCA2_Fpos_TTGTGGACATGCACTA-1dCA2_FI_GFPve_pos  0.1113741265
## dCA2_Fpos_TTGTTGTCAGTCCGTG-1dCA2_FI_GFPve_pos -0.2782917696
## dCA2_Fpos_TTTACGTAGCTCGGCT-1dCA2_FI_GFPve_pos  1.5662196798
## dCA2_Fpos_TTTACTGAGTCGAAGC-1dCA2_FI_GFPve_pos  1.0070127172
## dCA2_Fpos_TTTCATGCACAAATGA-1dCA2_FI_GFPve_pos  2.3621365947
```

```
## dCA2_Fpos_TTTCATGGTATCCCTC-1dCA2_FI_GFPve_pos  0.2217098621
## dCA2_Fpos_TTTGACTCATCGTGCG-1dCA2_FI_GFPve_pos  0.8483691735
## dCA2_Fpos_TTTGATCAGTGCCTCG-1dCA2_FI_GFPve_pos -1.1177264529
## FC_Mpos_AAAGGTATCTCTGCCA-1FC_MI_GFPve_pos      1.9710712356
## FC_Mpos_AAAGTGAAGCTTTCTT-1FC_MI_GFPve_pos     -0.2387873011
## FC_Mpos_AAAGTGAAGTTGGCTT-1FC_MI_GFPve_pos     -0.4935887533
## FC_Mpos_AAAGTGAGTCGAATTC-1FC_MI_GFPve_pos     -0.6085042435
## FC_Mpos_AACAAAGTCTGGTCAA-1FC_MI_GFPve_pos      0.1651500833
## FC_Mpos_AACCAACTCGGCCAAC-1FC_MI_GFPve_pos      1.4542106313
## FC_Mpos_AACCTGATCTCGACCT-1FC_MI_GFPve_pos     -0.0953783023
## FC_Mpos_AAGTACCAGTAGGAAG-1FC_MI_GFPve_pos     -0.0686523842
## FC_Mpos_AAGTACCGTCCTACGG-1FC_MI_GFPve_pos      0.1931772900
## FC_Mpos_AAGTACCTCCACCTGT-1FC_MI_GFPve_pos      1.9517990274
## FC_Mpos_AAGTGAAGTTCAAGGG-1FC_MI_GFPve_pos     -0.3884096521
## FC_Mpos_AATAGAGGTGTGTCGC-1FC_MI_GFPve_pos      1.5784010810
## FC_Mpos_AATCGTGAGGCATCAG-1FC_MI_GFPve_pos      0.7194663209
## FC_Mpos_AATCGTGCACCAAAGG-1FC_MI_GFPve_pos      2.5729253215
## FC_Mpos_AATGAAGGTACTAAGA-1FC_MI_GFPve_pos     -0.1817729252
## FC_Mpos_AATGCCACAGAAGCGT-1FC_MI_GFPve_pos      1.5677491349
## FC_Mpos_AATGGCTGTAGCACGA-1FC_MI_GFPve_pos      1.6234972400
## FC_Mpos_ACAAAGAGTTAGCGGA-1FC_MI_GFPve_pos      1.8228260202
## FC_Mpos_ACACAGTGTGCACGCT-1FC_MI_GFPve_pos     -1.0711929875
## FC_Mpos_ACACCAAGTCGATGCC-1FC_MI_GFPve_pos     -0.0388687479
## FC_Mpos_ACAGAAACAGCTATAC-1FC_MI_GFPve_pos     -1.5839250165
## FC_Mpos_ACCCTCAGTAGGTACG-1FC_MI_GFPve_pos      1.8354755563
## FC_Mpos_ACCCTTGCACTGTTCC-1FC_MI_GFPve_pos      0.6561128420
## FC_Mpos_ACCGTTCGTTCCGGTG-1FC_MI_GFPve_pos      1.5605819148
## FC_Mpos_ACCTGAACAGCGCGTT-1FC_MI_GFPve_pos     -0.7289379912
## FC_Mpos_ACGATCACAGTTCTAG-1FC_MI_GFPve_pos      0.5241844279
## FC_Mpos_ACGATCACATCGTTCC-1FC_MI_GFPve_pos      0.9442466420
## FC_Mpos_ACGGAAGGTATCACCA-1FC_MI_GFPve_pos      0.1639309374
## FC_Mpos_ACGGTCGTCGCATGAT-1FC_MI_GFPve_pos      0.3958154840
## FC_Mpos_ACGTACATCTTCTGTA-1FC_MI_GFPve_pos      0.6757162732
## FC_Mpos_ACTTAGGTCGTTCCTG-1FC_MI_GFPve_pos      2.5882718486
## FC_Mpos_ACTTATCGTAGGAGGG-1FC_MI_GFPve_pos      0.0795180590
## FC_Mpos_ACTTCCGCAATAGGAT-1FC_MI_GFPve_pos      0.8057310504
## FC_Mpos_AGAACAAGTACAAAGT-1FC_MI_GFPve_pos      0.8023289365
## FC_Mpos_AGAAGCGAGAGGATGA-1FC_MI_GFPve_pos      0.8917195124
## FC_Mpos_AGACCCGAGAGCCCAA-1FC_MI_GFPve_pos      2.2859580440
## FC_Mpos_AGACTCATCATTATCC-1FC_MI_GFPve_pos      0.8509384317
## FC_Mpos_AGAGCAGGTGCTTATG-1FC_MI_GFPve_pos      1.2797261399
## FC_Mpos_AGAGCCCTCTTGCAGA-1FC_MI_GFPve_pos      0.9085041804
## FC_Mpos_AGATGAAAGCGATTCT-1FC_MI_GFPve_pos     -0.2117608565
## FC_Mpos_AGATGAAGTTAAGGAT-1FC_MI_GFPve_pos      0.9507580918
## FC_Mpos_AGCATCATCACTCGAA-1FC_MI_GFPve_pos      1.0864966435
## FC_Mpos_AGCGTCGCAGGGTCTC-1FC_MI_GFPve_pos      0.3307367188
## FC_Mpos_AGCTTCCGTTAGAAGT-1FC_MI_GFPve_pos      0.8805413765
## FC_Mpos_AGGAATATCGATACGT-1FC_MI_GFPve_pos      1.0251815242
## FC_Mpos_AGGCATTAGGGTAATT-1FC_MI_GFPve_pos     -0.0706559497
## FC_Mpos_AGGCATTCATCAGCGC-1FC_MI_GFPve_pos      2.0984184665
## FC_Mpos_AGGCATTGTGGCGTAA-1FC_MI_GFPve_pos      0.8050708098
## FC_Mpos_AGGGCCTGTGTCACAT-1FC_MI_GFPve_pos     -0.4271599131
## FC_Mpos_AGGGTCCGTTTATGCG-1FC_MI_GFPve_pos     -0.4371579963
## FC_Mpos_AGGGTTTCATACAGAA-1FC_MI_GFPve_pos      2.5009004993
## FC_Mpos_AGGTCTATCCTCTAAT-1FC_MI_GFPve_pos      0.9548901242
## FC_Mpos_AGGTTGTCAATACCTG-1FC_MI_GFPve_pos      0.4472831113
## FC_Mpos_AGTACTGCAGCTACTA-1FC_MI_GFPve_pos      0.4574135763
## FC_Mpos_AGTAGCTAGTTCATCG-1FC_MI_GFPve_pos      2.6243643684
## FC_Mpos_AGTAGTCAGTGAGTTA-1FC_MI_GFPve_pos     -0.6059265214
## FC_Mpos_AGTAGTCCACTCTAGA-1FC_MI_GFPve_pos      1.4432936830
## FC_Mpos_AGTCATGAGTTAACGA-1FC_MI_GFPve_pos      1.5342695636
## FC_Mpos_AGTCTCCGTAGATGTA-1FC_MI_GFPve_pos      0.3987730664
## FC_Mpos_AGTGTTGAGTGTGTTC-1FC_MI_GFPve_pos      1.4024653358
## FC_Mpos_AGTTCGACAGCTGAGA-1FC_MI_GFPve_pos      0.8203097505
## FC_Mpos_ATACCGACACTCAGAT-1FC_MI_GFPve_pos      2.1584870738
## FC_Mpos_ATACCTTAGTTCATCG-1FC_MI_GFPve_pos      1.1504291219
## FC_Mpos_ATAGACCCATAGATCC-1FC_MI_GFPve_pos     -0.3850372809
## FC_Mpos_ATAGACCTCCATCCGT-1FC_MI_GFPve_pos      1.6106663865
## FC_Mpos_ATCCACCTCGGTAGGA-1FC_MI_GFPve_pos      1.9486817044
## FC_Mpos_ATCCCTGAGTGCTACT-1FC_MI_GFPve_pos      1.2650260848
```

```
## FC_Mpos_ATCGATGCACCTGCAG-1FC_MI_GFPve_pos     -1.4340186196
## FC_Mpos_ATCGATGGTGGAGGTT-1FC_MI_GFPve_pos      0.9671472234
## FC_Mpos_ATCGGATCACATTCGA-1FC_MI_GFPve_pos      1.2977909965
## FC_Mpos_ATCGGCGTCTTCCTAA-1FC_MI_GFPve_pos      0.7479666394
## FC_Mpos_ATCTCTATCGTGCAGC-1FC_MI_GFPve_pos      0.2341113013
## FC_Mpos_ATGAAAGGTGTCCCTT-1FC_MI_GFPve_pos      1.3685148877
## FC_Mpos_ATGACCATCCACTAGA-1FC_MI_GFPve_pos      2.0537362022
## FC_Mpos_ATGAGTCTCGTTCCCA-1FC_MI_GFPve_pos      0.0442655866
## FC_Mpos_ATGATCGGTCTACAAC-1FC_MI_GFPve_pos      0.1904880700
## FC_Mpos_ATGATCGTCGCTTACC-1FC_MI_GFPve_pos      0.9082208795
## FC_Mpos_ATGCCTCTCACCCTTG-1FC_MI_GFPve_pos      0.8517817182
## FC_Mpos_ATGCGATTCGGCATCG-1FC_MI_GFPve_pos      2.1150228900
## FC_Mpos_ATGGAGGAGATAGTCA-1FC_MI_GFPve_pos     -0.5700544196
## FC_Mpos_ATGGGAGTCGACATAC-1FC_MI_GFPve_pos     -0.9150893765
## FC_Mpos_ATGGGTTCACATAGCT-1FC_MI_GFPve_pos     -0.0001846703
## FC_Mpos_ATGGTTGAGCAGGTCA-1FC_MI_GFPve_pos     -0.9338183480
## FC_Mpos_ATGGTTGGTAGTACGG-1FC_MI_GFPve_pos      2.1207623405
## FC_Mpos_ATGGTTGTCTAAGAAG-1FC_MI_GFPve_pos      0.9345021171
## FC_Mpos_ATTACCTCACTCCTGT-1FC_MI_GFPve_pos      0.0320303810
## FC_Mpos_ATTACTCGTTGTGTTG-1FC_MI_GFPve_pos      0.4316984040
## FC_Mpos_ATTCACTGTGCTGTCG-1FC_MI_GFPve_pos      1.4499130172
## FC_Mpos_ATTCAGGCAATTCACG-1FC_MI_GFPve_pos      1.0098449511
## FC_Mpos_ATTCATCGTGTCTTGA-1FC_MI_GFPve_pos      0.2304792029
## FC_Mpos_ATTCCATCAGGTGTGA-1FC_MI_GFPve_pos     -0.1347684639
## FC_Mpos_ATTCTACTCCTCTTTC-1FC_MI_GFPve_pos      0.4518662912
## FC_Mpos_ATTCTTGTCGATAACC-1FC_MI_GFPve_pos      2.1578555030
## FC_Mpos_CAACAGTTCGCAGTGC-1FC_MI_GFPve_pos     -0.9278150874
## FC_Mpos_CAACCTCGTTGATGTC-1FC_MI_GFPve_pos      1.5890458745
## FC_Mpos_CAAGAGGAGTACAACA-1FC_MI_GFPve_pos      0.9089012069
## FC_Mpos_CAAGAGGCAGGTAGTG-1FC_MI_GFPve_pos     -0.3507267910
## FC_Mpos_CAAGGGACACAAATGA-1FC_MI_GFPve_pos      1.7249747438
## FC_Mpos_CAAGGGATCAATGTCG-1FC_MI_GFPve_pos      1.0314262432
## FC_Mpos_CAATACGTCTGGCCTT-1FC_MI_GFPve_pos      0.0862567888
## FC_Mpos_CAATCGACATGCCATA-1FC_MI_GFPve_pos      0.7418946547
## FC_Mpos_CAATTTCAGCCGTCGT-1FC_MI_GFPve_pos     -0.5540533739
## FC_Mpos_CAATTTCGTTGGGCCT-1FC_MI_GFPve_pos      0.5351691169
## FC_Mpos_CACAGGCCAGCCCAGT-1FC_MI_GFPve_pos      2.3994121474
## FC_Mpos_CACATGACAAGCAGGT-1FC_MI_GFPve_pos      1.2082370443
## FC_Mpos_CACCAAACATGACGTT-1FC_MI_GFPve_pos      0.0869793889
## FC_Mpos_CACGTGGTCTGTCGCT-1FC_MI_GFPve_pos     -0.7726165729
## FC_Mpos_CAGCACGTCCACTGGG-1FC_MI_GFPve_pos      2.5088772697
## FC_Mpos_CAGCAGCGTTGCTTGA-1FC_MI_GFPve_pos      2.0933775825
## FC_Mpos_CAGCCAGCAAGAGTTA-1FC_MI_GFPve_pos      1.3516676349
## FC_Mpos_CAGCGTGAGATAGTGT-1FC_MI_GFPve_pos      0.2241902200
## FC_Mpos_CAGGTATAGCATGAAT-1FC_MI_GFPve_pos      1.0798983497
## FC_Mpos_CAGTGCGCATAACTCG-1FC_MI_GFPve_pos      1.0605683250
## FC_Mpos_CATCGGGTCACATCAG-1FC_MI_GFPve_pos      0.0647085455
## FC_Mpos_CATGCGGAGGATGGCT-1FC_MI_GFPve_pos      0.5992322487
## FC_Mpos_CATTCATGTCATCTAG-1FC_MI_GFPve_pos     -0.6871976929
## FC_Mpos_CATTGAGCAGTGACCC-1FC_MI_GFPve_pos     -0.8065530139
## FC_Mpos_CCAATGACAATGAGCG-1FC_MI_GFPve_pos      0.2132587356
## FC_Mpos_CCAATGAGTTCTTAGG-1FC_MI_GFPve_pos      0.8428868574
## FC_Mpos_CCATAAGTCCAAAGGG-1FC_MI_GFPve_pos      0.6402383966
## FC_Mpos_CCCTAACTCAGCGTCG-1FC_MI_GFPve_pos     -0.9435920792
## FC_Mpos_CCCTCAAAGGTCTACT-1FC_MI_GFPve_pos      2.1535162849
## FC_Mpos_CCCTCAACAATCGAAA-1FC_MI_GFPve_pos      0.7348173184
## FC_Mpos_CCCTCTCTCTACCAGA-1FC_MI_GFPve_pos      0.6177086157
## FC_Mpos_CCGAACGAGGTGAGCT-1FC_MI_GFPve_pos     -0.8852669674
## FC_Mpos_CCGATGGAGTGTACCT-1FC_MI_GFPve_pos      0.6500254196
## FC_Mpos_CCGCAAGCATTCCTCG-1FC_MI_GFPve_pos      0.7657807392
## FC_Mpos_CCGGACAAGCAGCCCT-1FC_MI_GFPve_pos     -0.4560569602
## FC_Mpos_CCGGACAAGCCGAATG-1FC_MI_GFPve_pos      1.2251253051
## FC_Mpos_CCGGTAGCACCTGTCT-1FC_MI_GFPve_pos     -0.8387830334
## FC_Mpos_CCGTAGGGTCGGCTAC-1FC_MI_GFPve_pos      1.7321271819
## FC_Mpos_CCGTAGGGTTATCTGG-1FC_MI_GFPve_pos     -0.7776781398
## FC_Mpos_CCGTGAGCACTGGATT-1FC_MI_GFPve_pos     -0.8354277092
## FC_Mpos_CCTAACCAGTTGCCCG-1FC_MI_GFPve_pos      1.4285129232
## FC_Mpos_CCTACGTAGCAGGTCA-1FC_MI_GFPve_pos     -0.1031186211
## FC_Mpos_CCTCACAGTCAGCGTC-1FC_MI_GFPve_pos     -0.8423076349
## FC_Mpos_CCTCCAAAGAGCAACC-1FC_MI_GFPve_pos      0.4041021091
```

```
## FC_Mpos_CCTTGTGGTTGGACTT-1FC_MI_GFPve_pos      2.0448299569
## FC_Mpos_CGAAGGATCCATATGG-1FC_MI_GFPve_pos      0.2391337973
## FC_Mpos_CGACAGCCAGTTACCA-1FC_MI_GFPve_pos      0.1936984581
## FC_Mpos_CGAGGAACAAGTTTGC-1FC_MI_GFPve_pos     -0.2831761914
## FC_Mpos_CGATCGGTCGAACACT-1FC_MI_GFPve_pos     -0.3394234258
## FC_Mpos_CGCATAACAGAGGTTG-1FC_MI_GFPve_pos     -0.7016917544
## FC_Mpos_CGCCAGAAGCCATTTG-1FC_MI_GFPve_pos      2.2091989440
## FC_Mpos_CGGAACCAGAGTCAAT-1FC_MI_GFPve_pos      1.0463302058
## FC_Mpos_CGGGCATAGACGACGT-1FC_MI_GFPve_pos      1.9631670636
## FC_Mpos_CGGTCAGCAAACGAGC-1FC_MI_GFPve_pos      1.7440241498
## FC_Mpos_CGGTCAGTCGGAATGG-1FC_MI_GFPve_pos     -0.7832807380
## FC_Mpos_CGTAGTAAGCTAAATG-1FC_MI_GFPve_pos      0.9173711461
## FC_Mpos_CGTCCATCAGCAGACA-1FC_MI_GFPve_pos      1.0671425385
## FC_Mpos_CGTGAATAGCCGGATA-1FC_MI_GFPve_pos      1.8094862622
## FC_Mpos_CGTGATAGTGTGAATA-1FC_MI_GFPve_pos     -1.6391657668
## FC_Mpos_CGTTCTGCATCCTCAC-1FC_MI_GFPve_pos     -0.0483583587
## FC_Mpos_CGTTGGGTCCACTGGG-1FC_MI_GFPve_pos      1.4815829915
## FC_Mpos_CGTTGGGTCTACCAGA-1FC_MI_GFPve_pos      1.2408369703
## FC_Mpos_CTATCCGCAAAGGAGA-1FC_MI_GFPve_pos     -0.3827057260
## FC_Mpos_CTATCCGTCGCACGAC-1FC_MI_GFPve_pos      2.0503227634
## FC_Mpos_CTCAACCCATCCTTGC-1FC_MI_GFPve_pos      1.6702192945
## FC_Mpos_CTCCATGGTCATCGCG-1FC_MI_GFPve_pos      1.0778620166
## FC_Mpos_CTCCCAAAGACGGTTG-1FC_MI_GFPve_pos      0.8032672328
## FC_Mpos_CTCGAGGAGCCAGAGT-1FC_MI_GFPve_pos      1.6735757512
## FC_Mpos_CTCTGGTAGAACCCGA-1FC_MI_GFPve_pos      0.8594407362
## FC_Mpos_CTGAGGCAGACTTAAG-1FC_MI_GFPve_pos      0.0405237002
## FC_Mpos_CTGAGGCGTCGGATTT-1FC_MI_GFPve_pos      2.0314077062
## FC_Mpos_CTGCAGGAGCGCTTCG-1FC_MI_GFPve_pos      2.1519725246
## FC_Mpos_CTGCAGGTCAGCCTCT-1FC_MI_GFPve_pos      1.5655528230
## FC_Mpos_CTGCATCCAACGACTT-1FC_MI_GFPve_pos     -0.1421810317
## FC_Mpos_CTGCATCCACGCTGCA-1FC_MI_GFPve_pos     -0.4771948534
## FC_Mpos_CTGGCAGGTTTGGGAG-1FC_MI_GFPve_pos      1.2250064534
## FC_Mpos_CTGTAGATCCCGATCT-1FC_MI_GFPve_pos      0.0813629643
## FC_Mpos_CTGTCGTGTCACGACC-1FC_MI_GFPve_pos      0.3239927066
## FC_Mpos_CTGTGAACAGGATGAC-1FC_MI_GFPve_pos     -0.9447585421
## FC_Mpos_CTTACCGTCAGATGCT-1FC_MI_GFPve_pos     -1.3047788220
## FC_Mpos_CTTACCGTCTTCGATT-1FC_MI_GFPve_pos      1.3035937471
## FC_Mpos_CTTCAATAGATGTTAG-1FC_MI_GFPve_pos      1.2061536235
## FC_Mpos_CTTCAATTCAGCTTCC-1FC_MI_GFPve_pos      0.2567707819
## FC_Mpos_CTTCCGAGTTTGGCTA-1FC_MI_GFPve_pos      1.8523417634
## FC_Mpos_CTTCCGATCTGACGCG-1FC_MI_GFPve_pos      2.0521148366
## FC_Mpos_CTTTCAAGTAGCTCGC-1FC_MI_GFPve_pos     -0.0751148688
## FC_Mpos_GAAATGATCATGGGAG-1FC_MI_GFPve_pos     -1.4659676629
## FC_Mpos_GAACGTTAGGTAACTA-1FC_MI_GFPve_pos     -1.3035456019
## FC_Mpos_GAACGTTTCTCGTGGG-1FC_MI_GFPve_pos      1.2841027898
## FC_Mpos_GAAGAATTCTTGCGCT-1FC_MI_GFPve_pos      1.6130518836
## FC_Mpos_GAAGCGAGTCGTTGCG-1FC_MI_GFPve_pos      0.6649478001
## FC_Mpos_GAATCACAGTACAACA-1FC_MI_GFPve_pos      0.8886338991
## FC_Mpos_GACAGCCCAAGCAATA-1FC_MI_GFPve_pos      2.1599676532
## FC_Mpos_GACCCTTTCAGCCTCT-1FC_MI_GFPve_pos      1.6032401246
## FC_Mpos_GACCTTCAGAGTTGAT-1FC_MI_GFPve_pos     -0.2211549061
## FC_Mpos_GACTCTCGTCGCCTAG-1FC_MI_GFPve_pos      1.7769966048
## FC_Mpos_GACTTCCGTGACCGAA-1FC_MI_GFPve_pos      0.0012860594
## FC_Mpos_GAGAAATAGGTTGGAC-1FC_MI_GFPve_pos      0.7533814830
## FC_Mpos_GAGCCTGGTAACACGG-1FC_MI_GFPve_pos      0.9554770512
## FC_Mpos_GAGCTGCTCAGGAGAC-1FC_MI_GFPve_pos      1.3552876634
## FC_Mpos_GAGGCCTGTTCTCTAT-1FC_MI_GFPve_pos      0.2521531058
## FC_Mpos_GAGGGATCAGAAGTGC-1FC_MI_GFPve_pos     -0.0550400990
## FC_Mpos_GAGGGATGTTGCATTG-1FC_MI_GFPve_pos      0.0423178558
## FC_Mpos_GAGTCTAGTGGCAGAT-1FC_MI_GFPve_pos     -0.0432087409
## FC_Mpos_GAGTTACCAAAGTATG-1FC_MI_GFPve_pos      1.8536733312
## FC_Mpos_GAGTTACCACGCCACA-1FC_MI_GFPve_pos      0.5701876206
## FC_Mpos_GAGTTGTGTACGCTAT-1FC_MI_GFPve_pos      0.6338974756
## FC_Mpos_GAGTTGTGTTGAGTCT-1FC_MI_GFPve_pos      1.1445975227
## FC_Mpos_GATCGTAAGCTCCATA-1FC_MI_GFPve_pos      1.4801796598
## FC_Mpos_GATGATCGTCATAGTC-1FC_MI_GFPve_pos     -0.0676860737
## FC_Mpos_GATGTTGGTGACTATC-1FC_MI_GFPve_pos      0.0044262481
## FC_Mpos_GATTTCTCAAACTAGA-1FC_MI_GFPve_pos      1.0453635973
## FC_Mpos_GATTTCTTCACCTCGT-1FC_MI_GFPve_pos      2.1069590968
## FC_Mpos_GCAACATAGGGTAATT-1FC_MI_GFPve_pos     -0.5212630111
```

```
## FC_Mpos_GCAACATTCCGATCTC-1FC_MI_GFPve_pos     -0.0225393402
## FC_Mpos_GCAACCGTCATTGTTC-1FC_MI_GFPve_pos      1.4829122943
## FC_Mpos_GCACATATCATGAAAG-1FC_MI_GFPve_pos      2.8160409850
## FC_Mpos_GCACATATCTGTGCAA-1FC_MI_GFPve_pos      0.8752735180
## FC_Mpos_GCACTAATCCTACGAA-1FC_MI_GFPve_pos      0.5230491680
## FC_Mpos_GCAGTTAAGGAGCTGT-1FC_MI_GFPve_pos     -0.5477815944
## FC_Mpos_GCAGTTACAAGAGGTC-1FC_MI_GFPve_pos      0.9199400467
## FC_Mpos_GCATGATGTCTGTCAA-1FC_MI_GFPve_pos      1.3683849496
## FC_Mpos_GCATTAGAGCCACCGT-1FC_MI_GFPve_pos      0.3306382072
## FC_Mpos_GCCATGGCACTGCGAC-1FC_MI_GFPve_pos     -0.8948713618
## FC_Mpos_GCCCAGAAGAGTGTGC-1FC_MI_GFPve_pos      1.0318111104
## FC_Mpos_GCCCAGACATTGAGCT-1FC_MI_GFPve_pos      0.9145920796
## FC_Mpos_GCCCGAAGTTGGATCT-1FC_MI_GFPve_pos      1.4002041740
## FC_Mpos_GCCCGAATCAGGGATG-1FC_MI_GFPve_pos      1.2365335149
## FC_Mpos_GCGATCGAGTTGCCCG-1FC_MI_GFPve_pos      1.0152055544
## FC_Mpos_GCGTGCATCCAGGACC-1FC_MI_GFPve_pos     -0.4825245815
## FC_Mpos_GCTCAAAGTTAGGGTG-1FC_MI_GFPve_pos      1.7744605464
## FC_Mpos_GCTGAATAGGGCAGGA-1FC_MI_GFPve_pos      0.8579968733
## FC_Mpos_GCTTCACCAGTCTACA-1FC_MI_GFPve_pos      1.6332000417
## FC_Mpos_GCTTGGGGTGTTGCCG-1FC_MI_GFPve_pos      0.6050092501
## FC_Mpos_GGAACCCCATCACGGC-1FC_MI_GFPve_pos      0.2404763622
## FC_Mpos_GGAATCTGTTGCCGCA-1FC_MI_GFPve_pos     -0.3837018090
## FC_Mpos_GGACGTCCATCCTAAG-1FC_MI_GFPve_pos      0.0266010088
## FC_Mpos_GGAGAACAGAGCAGAA-1FC_MI_GFPve_pos      2.3371567649
## FC_Mpos_GGAGCAAGTATCCTCC-1FC_MI_GFPve_pos      0.6498339695
## FC_Mpos_GGCAGTCAGTTAACGA-1FC_MI_GFPve_pos      0.4350303632
## FC_Mpos_GGCAGTCCATGATGCT-1FC_MI_GFPve_pos      2.2575960082
## FC_Mpos_GGCGTCAGTCCGGCAT-1FC_MI_GFPve_pos     -0.8885991054
## FC_Mpos_GGGACAAAGGTTGTTC-1FC_MI_GFPve_pos      1.2292945308
## FC_Mpos_GGGACTCGTTATGTGC-1FC_MI_GFPve_pos     -0.1394812095
## FC_Mpos_GGGAGATGTGAGACCA-1FC_MI_GFPve_pos      2.1718480510
## FC_Mpos_GGGAGATTCGACATCA-1FC_MI_GFPve_pos      1.7178887052
## FC_Mpos_GGGATCCTCATTCATC-1FC_MI_GFPve_pos      2.4262046737
## FC_Mpos_GGGCCATAGATGGCGT-1FC_MI_GFPve_pos      0.3039571268
## FC_Mpos_GGGTATTAGTAGCATA-1FC_MI_GFPve_pos     -0.4439964968
## FC_Mpos_GGGTCACGTTTCCCAC-1FC_MI_GFPve_pos     -1.6257069188
## FC_Mpos_GGGTGTCAGTCGAAGC-1FC_MI_GFPve_pos      2.0296695155
## FC_Mpos_GGGTGTCAGTTTCTTC-1FC_MI_GFPve_pos      0.9061696452
## FC_Mpos_GGGTGTCCATGTTTGG-1FC_MI_GFPve_pos      1.8321231527
## FC_Mpos_GGTTGTAAGGTTCACT-1FC_MI_GFPve_pos     -0.6179977136
## FC_Mpos_GTAAGTCTCGCACGGT-1FC_MI_GFPve_pos      0.9600135011
## FC_Mpos_GTAGAGGCAGAGCCCT-1FC_MI_GFPve_pos     -1.3469652015
## FC_Mpos_GTAGGAGAGGAACTCG-1FC_MI_GFPve_pos     -1.4494004327
## FC_Mpos_GTAGGAGCAACGGTAG-1FC_MI_GFPve_pos      0.1374572774
## FC_Mpos_GTAGGTTCAATTCTTC-1FC_MI_GFPve_pos     -0.6667502003
## FC_Mpos_GTATTTCAGCAGGGAG-1FC_MI_GFPve_pos      0.1664424089
## FC_Mpos_GTATTTCGTCAACCTA-1FC_MI_GFPve_pos      1.2216395063
## FC_Mpos_GTCATGATCGGATACT-1FC_MI_GFPve_pos     -0.3303789573
## FC_Mpos_GTCCTCATCGTAACTG-1FC_MI_GFPve_pos      1.7714899701
## FC_Mpos_GTCTAGACATGACAGG-1FC_MI_GFPve_pos      1.7806179446
## FC_Mpos_GTCTCACGTAGGTCAG-1FC_MI_GFPve_pos      1.5590435189
## FC_Mpos_GTCTGTCCAAGGAGTC-1FC_MI_GFPve_pos      0.9824919624
## FC_Mpos_GTCTTTAAGGACTTCT-1FC_MI_GFPve_pos      1.7337126655
## FC_Mpos_GTGACGCGTAATTGGA-1FC_MI_GFPve_pos      1.5296304149
## FC_Mpos_GTGACGCGTTCTAAGC-1FC_MI_GFPve_pos      0.3565989656
## FC_Mpos_GTGACGCTCAGCGCGT-1FC_MI_GFPve_pos      2.2331719321
## FC_Mpos_GTGATGTTCACGAGGA-1FC_MI_GFPve_pos     -0.5119616466
## FC_Mpos_GTGCACGTCCACACCT-1FC_MI_GFPve_pos      1.1601295394
## FC_Mpos_GTGCAGCCAGTGTACT-1FC_MI_GFPve_pos      2.2859036845
## FC_Mpos_GTGCAGCTCCAGCAAT-1FC_MI_GFPve_pos      1.5927508992
## FC_Mpos_GTGCGTGAGAACTGAT-1FC_MI_GFPve_pos      0.4580148083
## FC_Mpos_GTGCGTGAGTACAGAT-1FC_MI_GFPve_pos     -0.2254848557
## FC_Mpos_GTGCGTGCACATTACG-1FC_MI_GFPve_pos      0.3541894776
## FC_Mpos_GTGCTGGAGTAAGAGG-1FC_MI_GFPve_pos     -0.0157756659
## FC_Mpos_GTGCTTCGTTGCACGC-1FC_MI_GFPve_pos      0.9428992194
## FC_Mpos_GTGGAAGAGAGGTTTA-1FC_MI_GFPve_pos      1.7340300006
## FC_Mpos_GTGGAAGAGGATGCGT-1FC_MI_GFPve_pos      0.3798316640
## FC_Mpos_GTGGCGTAGGTCCCTG-1FC_MI_GFPve_pos      1.3582429809
## FC_Mpos_GTGGCGTCAGAACGCA-1FC_MI_GFPve_pos      2.0449638290
## FC_Mpos_GTGGGAACAGCAAGAC-1FC_MI_GFPve_pos      0.1793039513
```

```
## FC_Mpos_GTGGGAATCAGGGTAG-1FC_MI_GFPve_pos      0.8171308560
## FC_Mpos_GTGTAACCACCAGCCA-1FC_MI_GFPve_pos      1.6482535524
## FC_Mpos_GTGTCCTAGAAGCTGC-1FC_MI_GFPve_pos      0.9141275210
## FC_Mpos_GTGTCCTCATCGGTTA-1FC_MI_GFPve_pos      1.8048303050
## FC_Mpos_GTGTTCCCAACTGTGT-1FC_MI_GFPve_pos      1.3329962415
## FC_Mpos_GTGTTCCCAGTGTGGA-1FC_MI_GFPve_pos      0.3536267800
## FC_Mpos_GTGTTCCGTTCTCGCT-1FC_MI_GFPve_pos      0.5935654563
## FC_Mpos_GTTACAGCAATGAAAC-1FC_MI_GFPve_pos      0.2417374176
## FC_Mpos_GTTATGGAGCAACTCT-1FC_MI_GFPve_pos      0.5467134041
## FC_Mpos_GTTCATTAGACATCCT-1FC_MI_GFPve_pos      0.6939914149
## FC_Mpos_GTTCTATAGAGCCGTA-1FC_MI_GFPve_pos     -0.1955110508
## FC_Mpos_GTTCTATTCCAGTTCC-1FC_MI_GFPve_pos     -1.2274226027
## FC_Mpos_GTTGTAGGTGGGATTG-1FC_MI_GFPve_pos      1.1534926814
## FC_Mpos_GTTGTGAGTCAACACT-1FC_MI_GFPve_pos      1.0229650659
## FC_Mpos_GTTTACTCAGAGTGAC-1FC_MI_GFPve_pos      0.1210342475
## FC_Mpos_TAAGCCATCGGAGTGA-1FC_MI_GFPve_pos     -0.2534287053
## FC_Mpos_TAAGTCGGTCGTGTTA-1FC_MI_GFPve_pos      0.1753328515
## FC_Mpos_TAAGTCGGTTAAAGTG-1FC_MI_GFPve_pos     -0.1256699788
## FC_Mpos_TACACCCCACTAGGTT-1FC_MI_GFPve_pos      0.1797860784
## FC_Mpos_TACCGGGTCTTCCAGC-1FC_MI_GFPve_pos      0.1344058418
## FC_Mpos_TACCTCGTCCGCGAGT-1FC_MI_GFPve_pos      1.7316826505
## FC_Mpos_TACGCTCGTTCTTGCC-1FC_MI_GFPve_pos      1.3770073575
## FC_Mpos_TACTTACTCGGTCGAC-1FC_MI_GFPve_pos     -0.2860158759
## FC_Mpos_TAGACCAGTTGCGGAA-1FC_MI_GFPve_pos      0.5865086240
## FC_Mpos_TAGAGTCGTAACTTCG-1FC_MI_GFPve_pos      2.4362275523
## FC_Mpos_TAGGAGGGTCATCACA-1FC_MI_GFPve_pos     -0.3402682322
## FC_Mpos_TAGGAGGGTCGAATTC-1FC_MI_GFPve_pos      1.8787906093
## FC_Mpos_TAGGTACAGTTCACTG-1FC_MI_GFPve_pos      0.6196466011
## FC_Mpos_TAGTGCACATCGAGCC-1FC_MI_GFPve_pos      1.2912716788
## FC_Mpos_TATATCCGTCGTCAGC-1FC_MI_GFPve_pos      0.9891597552
## FC_Mpos_TATATCCTCCCGTGTT-1FC_MI_GFPve_pos      0.9964253229
## FC_Mpos_TATCCTAGTTCCTTGC-1FC_MI_GFPve_pos     -0.7923392731
## FC_Mpos_TATCGCCGTATCCCTC-1FC_MI_GFPve_pos      1.3318161887
## FC_Mpos_TATCTGTAGTGGTGGT-1FC_MI_GFPve_pos      2.2114815635
## FC_Mpos_TCAAGCATCGCCTATC-1FC_MI_GFPve_pos      0.3861823899
## FC_Mpos_TCACACCCACTATGTG-1FC_MI_GFPve_pos      0.3043148292
## FC_Mpos_TCACGCTTCCTTGGAA-1FC_MI_GFPve_pos      1.9848514718
## FC_Mpos_TCACGGGTCCGGCAAC-1FC_MI_GFPve_pos     -0.0595633465
## FC_Mpos_TCACTATAGCGTCAAG-1FC_MI_GFPve_pos      0.7292686624
## FC_Mpos_TCACTCGCAGGACTAG-1FC_MI_GFPve_pos      0.9225178284
## FC_Mpos_TCATCATAGACCTCCG-1FC_MI_GFPve_pos      1.5368063373
## FC_Mpos_TCATCATCAGATTAAG-1FC_MI_GFPve_pos     -0.3122258561
## FC_Mpos_TCATCCGAGGCCTGAA-1FC_MI_GFPve_pos      2.1314566058
## FC_Mpos_TCATGAGAGTTAGTAG-1FC_MI_GFPve_pos      0.9575995726
## FC_Mpos_TCATGAGTCGATGCTA-1FC_MI_GFPve_pos     -0.3898506242
## FC_Mpos_TCATGGACAAATACAG-1FC_MI_GFPve_pos      0.9408537668
## FC_Mpos_TCCATGCAGTCTGGAG-1FC_MI_GFPve_pos      0.3819650633
## FC_Mpos_TCCATGCTCATACGGT-1FC_MI_GFPve_pos     -0.0798416513
## FC_Mpos_TCCCACAAGCGATTCT-1FC_MI_GFPve_pos      0.7731761259
## FC_Mpos_TCCCACAGTGGAGAAA-1FC_MI_GFPve_pos     -0.2522270459
## FC_Mpos_TCCCATGTCGTTCTCG-1FC_MI_GFPve_pos      0.7576414270
## FC_Mpos_TCCGATCAGTTGTAGA-1FC_MI_GFPve_pos     -0.3791961449
## FC_Mpos_TCCGATCCAAAGGTTA-1FC_MI_GFPve_pos      1.6024526281
## FC_Mpos_TCCGTGTCACTTCCTG-1FC_MI_GFPve_pos      0.1606814039
## FC_Mpos_TCCTCTTCAGATCCAT-1FC_MI_GFPve_pos      1.3246685189
## FC_Mpos_TCGACCTAGCTTACGT-1FC_MI_GFPve_pos      0.9348508639
## FC_Mpos_TCGACGGGTAACGCGA-1FC_MI_GFPve_pos     -0.9415971118
## FC_Mpos_TCGATTTTCCCGAATA-1FC_MI_GFPve_pos      1.7411416692
## FC_Mpos_TCGGATATCGCGTTTC-1FC_MI_GFPve_pos     -0.5144633728
## FC_Mpos_TCGGTCTAGACTTGTC-1FC_MI_GFPve_pos      1.8204400462
## FC_Mpos_TCGGTCTCATTGAGCT-1FC_MI_GFPve_pos     -0.1961661952
## FC_Mpos_TCGTGCTCAGTTAAAG-1FC_MI_GFPve_pos      1.3239046258
## FC_Mpos_TCGTGCTGTGATACAA-1FC_MI_GFPve_pos      0.1919649837
## FC_Mpos_TCTAACTCAGGTCAAG-1FC_MI_GFPve_pos      2.1535170001
## FC_Mpos_TCTACCGAGCGAGGAG-1FC_MI_GFPve_pos      0.7077407760
## FC_Mpos_TCTACCGGTACTGAGG-1FC_MI_GFPve_pos      0.2592162413
## FC_Mpos_TCTCAGCCACGTGTGC-1FC_MI_GFPve_pos      1.9106976909
## FC_Mpos_TCTCAGCGTATCGATC-1FC_MI_GFPve_pos      0.2240570781
## FC_Mpos_TCTCTGGTCTCGTGAA-1FC_MI_GFPve_pos     -0.7342581826
## FC_Mpos_TCTGCCAAGAGCAGCT-1FC_MI_GFPve_pos      1.7832324428
```

```
## FC_Mpos_TCTGGCTCATACCGTA-1FC_MI_GFPve_pos     -0.9510989266
## FC_Mpos_TCTGGCTGTCTGTAGT-1FC_MI_GFPve_pos      1.0423189921
## FC_Mpos_TCTTCCTTCAAAGAAC-1FC_MI_GFPve_pos     -0.1472322422
## FC_Mpos_TCTTGCGAGTTGAAGT-1FC_MI_GFPve_pos      0.9105837864
## FC_Mpos_TGAACGTTCGCTCATC-1FC_MI_GFPve_pos      0.8785384816
## FC_Mpos_TGACAGTCAGTCAACT-1FC_MI_GFPve_pos     -0.9141517955
## FC_Mpos_TGACAGTTCCCTCTTT-1FC_MI_GFPve_pos      1.6909978313
## FC_Mpos_TGACGCGCAGCTGTAT-1FC_MI_GFPve_pos      0.5671644730
## FC_Mpos_TGAGACTTCATTACGG-1FC_MI_GFPve_pos      1.0550947112
## FC_Mpos_TGAGGTTCAGTCGGAA-1FC_MI_GFPve_pos      0.4142871124
## FC_Mpos_TGAGGTTGTAACGATA-1FC_MI_GFPve_pos      0.1378276569
## FC_Mpos_TGATCAGTCGTAGCCG-1FC_MI_GFPve_pos      2.0556650085
## FC_Mpos_TGATGCACAGGTGTGA-1FC_MI_GFPve_pos     -1.1001269894
## FC_Mpos_TGATGCACATCTTCGC-1FC_MI_GFPve_pos     -0.0047908949
## FC_Mpos_TGATGCATCACTGTTT-1FC_MI_GFPve_pos      1.9303461236
## FC_Mpos_TGATGGTAGTTTGAGA-1FC_MI_GFPve_pos     -0.2553448158
## FC_Mpos_TGATTTCCATACCACA-1FC_MI_GFPve_pos     -0.1230918693
## FC_Mpos_TGCAGATGTACCGGAA-1FC_MI_GFPve_pos      0.5545052809
## FC_Mpos_TGCATGACAGCACGAA-1FC_MI_GFPve_pos      1.1794367952
## FC_Mpos_TGCCGAGCAGACGCTC-1FC_MI_GFPve_pos      0.4031857473
## FC_Mpos_TGCGGGTTCTCGACCT-1FC_MI_GFPve_pos      0.7229694051
## FC_Mpos_TGCTCGTCAGGTTTAC-1FC_MI_GFPve_pos     -0.0421231168
## FC_Mpos_TGGATCACATTGCCTC-1FC_MI_GFPve_pos      0.3971736950
## FC_Mpos_TGGATGTTCCAGCTCT-1FC_MI_GFPve_pos      1.7144156617
## FC_Mpos_TGGGCTGTCAGGTGTT-1FC_MI_GFPve_pos      1.6270276231
## FC_Mpos_TGGGTTATCACAATGC-1FC_MI_GFPve_pos      0.7226339502
## FC_Mpos_TGTAAGCTCCACGTGG-1FC_MI_GFPve_pos      0.5896758360
## FC_Mpos_TGTCAGAGTAATGCGG-1FC_MI_GFPve_pos      0.5882473809
## FC_Mpos_TGTCCACTCTGCCTCA-1FC_MI_GFPve_pos      1.9645277185
## FC_Mpos_TGTCCTGTCGTTATCT-1FC_MI_GFPve_pos     -0.9776628094
## FC_Mpos_TGTGATGGTGCCCTTT-1FC_MI_GFPve_pos      1.3820454997
## FC_Mpos_TGTGCGGCATGCCGGT-1FC_MI_GFPve_pos     -0.1454865712
## FC_Mpos_TGTGGCGTCGCAACAT-1FC_MI_GFPve_pos      0.5451394839
## FC_Mpos_TGTGTGACAAGCAGGT-1FC_MI_GFPve_pos      0.4916827304
## FC_Mpos_TGTTCCGTCATATGGC-1FC_MI_GFPve_pos      0.8298767728
## FC_Mpos_TGTTGAGGTCAGTCCG-1FC_MI_GFPve_pos      2.4762816352
## FC_Mpos_TGTTTGTCAATACGAA-1FC_MI_GFPve_pos     -0.3800990361
## FC_Mpos_TTAATCCTCCACTTTA-1FC_MI_GFPve_pos      1.5257204694
## FC_Mpos_TTACAGGGTGGAACAC-1FC_MI_GFPve_pos      1.6701439542
## FC_Mpos_TTACAGGTCTACTCAT-1FC_MI_GFPve_pos      2.3224675578
## FC_Mpos_TTACCATGTACGGATG-1FC_MI_GFPve_pos      2.3465402049
## FC_Mpos_TTACCGCTCCTCATAT-1FC_MI_GFPve_pos     -0.1444104242
## FC_Mpos_TTACGCCCATGGGCAA-1FC_MI_GFPve_pos      0.2328806293
## FC_Mpos_TTAGGCACACCAGGTC-1FC_MI_GFPve_pos      2.0617170257
## FC_Mpos_TTAGTCTTCAATCGGT-1FC_MI_GFPve_pos      2.3381664676
## FC_Mpos_TTATTGCAGGTTATAG-1FC_MI_GFPve_pos     -0.7327121573
## FC_Mpos_TTATTGCAGTCACACT-1FC_MI_GFPve_pos      0.5365538222
## FC_Mpos_TTATTGCCAGGACATG-1FC_MI_GFPve_pos      0.4371865017
## FC_Mpos_TTCAATCCACACGCCA-1FC_MI_GFPve_pos      0.4569779557
## FC_Mpos_TTCACGCAGACGACGT-1FC_MI_GFPve_pos      0.2565018756
## FC_Mpos_TTCACGCTCTGTAACG-1FC_MI_GFPve_pos      1.5111696643
## FC_Mpos_TTCCAATAGAGTTGAT-1FC_MI_GFPve_pos      1.4254552049
## FC_Mpos_TTCCGGTGTAGCGCTC-1FC_MI_GFPve_pos      0.6000510377
## FC_Mpos_TTCCGTGAGACGATAT-1FC_MI_GFPve_pos     -0.3674761432
## FC_Mpos_TTCCTAAAGGCCCACT-1FC_MI_GFPve_pos      1.0534995121
## FC_Mpos_TTCGCTGCAGGTCAGA-1FC_MI_GFPve_pos      1.2463247699
## FC_Mpos_TTCGGTCTCCTCTCGA-1FC_MI_GFPve_pos     -1.1273845511
## FC_Mpos_TTCTAACGTCCAGAAG-1FC_MI_GFPve_pos      1.0136789602
## FC_Mpos_TTCTTCCAGGCGTCCT-1FC_MI_GFPve_pos      0.1528687177
## FC_Mpos_TTCTTGACAACACGTT-1FC_MI_GFPve_pos      1.1861143035
## FC_Mpos_TTGAACGAGGCGAAGG-1FC_MI_GFPve_pos      2.4037105960
## FC_Mpos_TTGAGTGAGAAGCGGG-1FC_MI_GFPve_pos      1.4571520013
## FC_Mpos_TTGAGTGAGGTGCTTT-1FC_MI_GFPve_pos      0.3031605554
## FC_Mpos_TTGAGTGCAATAGGAT-1FC_MI_GFPve_pos      1.5587822122
## FC_Mpos_TTGAGTGTCGACGACC-1FC_MI_GFPve_pos      0.2939405364
## FC_Mpos_TTGCTGCAGCACTTTG-1FC_MI_GFPve_pos      1.0125630421
## FC_Mpos_TTGGGCGAGACCTCAT-1FC_MI_GFPve_pos      2.0877184791
## FC_Mpos_TTGTTCATCACTCACC-1FC_MI_GFPve_pos      0.8968736452
## FC_Mpos_TTTACGTTCAAGTGGG-1FC_MI_GFPve_pos     -0.4199158030
## FC_Mpos_TTTCACACATCGGAGA-1FC_MI_GFPve_pos      1.1330546064
```

```
## FC_Mpos_TTTCACATCGTCCATC-1FC_MI_GFPve_pos      0.4996142906
## FC_Mpos_TTTCATGTCGGAACTT-1FC_MI_GFPve_pos     -0.5198044854
## FC_Mpos_TTTCATGTCTACCACC-1FC_MI_GFPve_pos      0.7718349022
## FC_Mpos_TTTCGATGTATGGAAT-1FC_MI_GFPve_pos      2.2981800956
## FC_Mpos_TTTGTTGAGCATGATA-1FC_MI_GFPve_pos     -0.1100921261
## FC_Fpos_AAACGCTAGGTTCAGG-1FC_FI_GFPve_pos      1.1206008119
## FC_Fpos_AAAGGATCAGTTTCGA-1FC_FI_GFPve_pos      0.1184116031
## FC_Fpos_AAAGGGCCAGTCTGGC-1FC_FI_GFPve_pos      0.2162432743
## FC_Fpos_AAAGGTACAGAGTCTT-1FC_FI_GFPve_pos      1.6326546592
## FC_Fpos_AACACACTCGTTTACT-1FC_FI_GFPve_pos      1.0340643448
## FC_Fpos_AACAGGGAGGAACTCG-1FC_FI_GFPve_pos     -0.3282169181
## FC_Fpos_AACCACAAGTTCTCTT-1FC_FI_GFPve_pos     -0.3512541133
## FC_Fpos_AACCACAGTTGCTGAT-1FC_FI_GFPve_pos     -1.0076803046
## FC_Fpos_AACCTGATCATTTACC-1FC_FI_GFPve_pos      1.1832172794
## FC_Fpos_AACCTTTGTTCGGGTC-1FC_FI_GFPve_pos      0.3985155446
## FC_Fpos_AACGTCATCTAATTCC-1FC_FI_GFPve_pos      0.3612059576
## FC_Fpos_AACTTCTTCGCTGCGA-1FC_FI_GFPve_pos      0.5311930699
## FC_Fpos_AAGAACACACAGTCGC-1FC_FI_GFPve_pos     -0.3982057648
## FC_Fpos_AAGAACAGTCGGCTAC-1FC_FI_GFPve_pos     -0.5004814940
## FC_Fpos_AAGACTCTCAGCCCAG-1FC_FI_GFPve_pos      0.7435307426
## FC_Fpos_AAGCCATAGTATGTAG-1FC_FI_GFPve_pos      0.0752278024
## FC_Fpos_AAGCCATCACAGTGTT-1FC_FI_GFPve_pos      0.4915662927
## FC_Fpos_AAGCCATGTCGAGCAA-1FC_FI_GFPve_pos      1.1091887874
## FC_Fpos_AAGCGAGCACTTCAAG-1FC_FI_GFPve_pos     -1.0411081391
## FC_Fpos_AAGGAATCATTAAGCC-1FC_FI_GFPve_pos      1.9172667188
## FC_Fpos_AAGGTAACACAACATC-1FC_FI_GFPve_pos     -1.0794116336
## FC_Fpos_AAGGTAACACTACCGG-1FC_FI_GFPve_pos     -0.5223063308
## FC_Fpos_AAGTACCAGATGGGCT-1FC_FI_GFPve_pos     -0.3892370241
## FC_Fpos_AAGTACCGTTAAGACA-1FC_FI_GFPve_pos      2.0879249496
## FC_Fpos_AAGTACCTCCCGGTAG-1FC_FI_GFPve_pos     -0.6576764661
## FC_Fpos_AAGTCGTAGGAAAGAC-1FC_FI_GFPve_pos     -0.1559349078
## FC_Fpos_AAGTCGTTCCATCTCG-1FC_FI_GFPve_pos     -0.6855800825
## FC_Fpos_AAGTCGTTCTCTCGCA-1FC_FI_GFPve_pos      0.8533625526
## FC_Fpos_AAGTGAAGTTCCGCAG-1FC_FI_GFPve_pos      1.9573478622
## FC_Fpos_AAGTTCGCAACCCTAA-1FC_FI_GFPve_pos      1.7767136020
## FC_Fpos_AATCGTGCACGGAAGT-1FC_FI_GFPve_pos      0.0452163142
## FC_Fpos_AATGACCGTACCTAGT-1FC_FI_GFPve_pos     -0.3852061408
## FC_Fpos_AATGCCATCCATCTCG-1FC_FI_GFPve_pos     -0.9514730054
## FC_Fpos_ACAAAGAGTGCCAAGA-1FC_FI_GFPve_pos      2.4575657767
## FC_Fpos_ACAACCACAAGATCCT-1FC_FI_GFPve_pos      0.1674493921
## FC_Fpos_ACACAGTAGCTAGAGC-1FC_FI_GFPve_pos      1.1080117149
## FC_Fpos_ACACAGTTCGACATCA-1FC_FI_GFPve_pos      2.3775980396
## FC_Fpos_ACACCAAAGCACGATG-1FC_FI_GFPve_pos      1.6009870691
## FC_Fpos_ACACGCGAGTAACAGT-1FC_FI_GFPve_pos      1.5166096610
## FC_Fpos_ACACGCGGTACGGTTT-1FC_FI_GFPve_pos      0.1097884460
## FC_Fpos_ACACGCGTCGAATCCA-1FC_FI_GFPve_pos     -0.5899006205
## FC_Fpos_ACAGCCGCATTGACCA-1FC_FI_GFPve_pos      0.0083120090
## FC_Fpos_ACAGCCGTCGCCTTTG-1FC_FI_GFPve_pos      0.9740449709
## FC_Fpos_ACAGGGACAGGCTTGC-1FC_FI_GFPve_pos      1.0205072684
## FC_Fpos_ACAGGGATCGGTGTAT-1FC_FI_GFPve_pos      0.1439862696
## FC_Fpos_ACCAACAGTCGTCATA-1FC_FI_GFPve_pos      0.6357185764
## FC_Fpos_ACCCAAAGTCTTGCTC-1FC_FI_GFPve_pos      0.2253037763
## FC_Fpos_ACCCTCACAATTAGGA-1FC_FI_GFPve_pos      1.0291303915
## FC_Fpos_ACCGTTCGTAGAATAC-1FC_FI_GFPve_pos      2.4513974113
## FC_Fpos_ACCTACCAGCGGTATG-1FC_FI_GFPve_pos     -1.2139607745
## FC_Fpos_ACCTGAACAGACGGAT-1FC_FI_GFPve_pos      1.3343804998
## FC_Fpos_ACCTGAATCTGGGCCA-1FC_FI_GFPve_pos      0.7747361583
## FC_Fpos_ACCTGTCTCCGGACGT-1FC_FI_GFPve_pos      0.5800119561
## FC_Fpos_ACGCACGTCCAGTGCG-1FC_FI_GFPve_pos      0.5490085704
## FC_Fpos_ACGGAAGAGACTTCAC-1FC_FI_GFPve_pos     -0.0144249874
## FC_Fpos_ACGGGTCCAAATGCGG-1FC_FI_GFPve_pos      1.8899872226
## FC_Fpos_ACGGGTCCAGAACTTC-1FC_FI_GFPve_pos      0.4937941653
## FC_Fpos_ACGGGTCGTATGAAAC-1FC_FI_GFPve_pos     -0.6680135208
## FC_Fpos_ACGGTCGCATCTTTCA-1FC_FI_GFPve_pos     -0.1846265870
## FC_Fpos_ACGTACAAGTGCCCGT-1FC_FI_GFPve_pos      1.4970239324
## FC_Fpos_ACGTACAGTATGGTAA-1FC_FI_GFPve_pos     -0.6794170814
## FC_Fpos_ACGTAGTAGACTCTAC-1FC_FI_GFPve_pos      1.3274607581
## FC_Fpos_ACGTAGTGTAGCGCCT-1FC_FI_GFPve_pos      0.0481018608
## FC_Fpos_ACGTCCTAGCTGAGCA-1FC_FI_GFPve_pos     -0.4943912702
## FC_Fpos_ACGTCCTCAAGTCGTT-1FC_FI_GFPve_pos     -0.1367360878
```

```
## FC_Fpos_ACGTTCCCAGCACAAG-1FC_FI_GFPve_pos     -0.5139316397
## FC_Fpos_ACGTTCCGTCGCGTCA-1FC_FI_GFPve_pos      1.2455576581
## FC_Fpos_ACTATCTAGCATCAAA-1FC_FI_GFPve_pos     -0.5945959764
## FC_Fpos_ACTATGGTCAGTGGGA-1FC_FI_GFPve_pos      1.7509316129
## FC_Fpos_ACTGATGTCGCAGATT-1FC_FI_GFPve_pos      0.0074779612
## FC_Fpos_ACTGCAAAGATACGAT-1FC_FI_GFPve_pos     -0.4245997744
## FC_Fpos_ACTGCAAGTTTACTGG-1FC_FI_GFPve_pos      0.8769670648
## FC_Fpos_ACTGTCCAGCGGATCA-1FC_FI_GFPve_pos      0.5494764370
## FC_Fpos_ACTTAGGCAAAGGTTA-1FC_FI_GFPve_pos     -0.7323470789
## FC_Fpos_ACTTCGCTCGGCTGAC-1FC_FI_GFPve_pos      0.4218973440
## FC_Fpos_ACTTTCAAGGAGAGGC-1FC_FI_GFPve_pos      0.2182136772
## FC_Fpos_ACTTTCATCTAGTCAG-1FC_FI_GFPve_pos      1.4266744775
## FC_Fpos_ACTTTGTCAGAAGTGC-1FC_FI_GFPve_pos     -0.6572914797
## FC_Fpos_ACTTTGTGTTTGAACC-1FC_FI_GFPve_pos      0.2594892306
## FC_Fpos_AGAACAACAATAAGGT-1FC_FI_GFPve_pos      0.4542599541
## FC_Fpos_AGAACAAGTTAGAAAC-1FC_FI_GFPve_pos     -0.5516234117
## FC_Fpos_AGAACCTCAGTAACCT-1FC_FI_GFPve_pos      2.3293058795
## FC_Fpos_AGAAGCGGTAGGTCAG-1FC_FI_GFPve_pos     -0.6192635375
## FC_Fpos_AGACAAATCTATACGG-1FC_FI_GFPve_pos      0.6635345740
## FC_Fpos_AGACACTGTAACTGCT-1FC_FI_GFPve_pos     -0.9732158261
## FC_Fpos_AGACACTGTTATGTGC-1FC_FI_GFPve_pos      0.9883990211
## FC_Fpos_AGACAGGTCCTCAGGG-1FC_FI_GFPve_pos      0.8467994255
## FC_Fpos_AGACCATAGATTGAGT-1FC_FI_GFPve_pos     -0.8186274844
## FC_Fpos_AGACCATCACCCTGAG-1FC_FI_GFPve_pos      1.5131341142
## FC_Fpos_AGAGAATCATGACTCA-1FC_FI_GFPve_pos      0.2875476253
## FC_Fpos_AGAGAATGTACTGGGA-1FC_FI_GFPve_pos      1.6210201902
## FC_Fpos_AGAGCCCGTGAACGGT-1FC_FI_GFPve_pos      0.0551973765
## FC_Fpos_AGAGCCCGTGAGACCA-1FC_FI_GFPve_pos      1.1447750253
## FC_Fpos_AGATAGAGTCGTAATC-1FC_FI_GFPve_pos     -0.0596022385
## FC_Fpos_AGATGCTAGTAGCTCT-1FC_FI_GFPve_pos      1.6661384029
## FC_Fpos_AGCCAATGTCGATGCC-1FC_FI_GFPve_pos     -0.8803754526
## FC_Fpos_AGCCAATTCTATTCGT-1FC_FI_GFPve_pos      2.0767351312
## FC_Fpos_AGCCACGCAGATTAAG-1FC_FI_GFPve_pos     -0.8490657287
## FC_Fpos_AGCCAGCGTTTGTGGT-1FC_FI_GFPve_pos     -0.5790229517
## FC_Fpos_AGCGATTTCGTGAGAG-1FC_FI_GFPve_pos     -0.1069161075
## FC_Fpos_AGCGTATCAACGTATC-1FC_FI_GFPve_pos     -0.3994675356
## FC_Fpos_AGCTACAGTTCGTAAC-1FC_FI_GFPve_pos     -0.3531043011
## FC_Fpos_AGCTCAAAGGCTAGCA-1FC_FI_GFPve_pos     -0.1060253012
## FC_Fpos_AGGAATACACGTACAT-1FC_FI_GFPve_pos     -0.0894963222
## FC_Fpos_AGGAATAGTCTCAGGC-1FC_FI_GFPve_pos     -0.0253525006
## FC_Fpos_AGGACTTGTCATCAGT-1FC_FI_GFPve_pos      0.3470930827
## FC_Fpos_AGGAGGTAGGACCCAA-1FC_FI_GFPve_pos     -0.3929459828
## FC_Fpos_AGGAGGTGTCGGATTT-1FC_FI_GFPve_pos      0.9765303058
## FC_Fpos_AGGATAAGTTTCCAAG-1FC_FI_GFPve_pos      0.7959713859
## FC_Fpos_AGGCATTAGGCCACCT-1FC_FI_GFPve_pos      0.9619535131
## FC_Fpos_AGGCATTGTGGCTACC-1FC_FI_GFPve_pos      0.7565278930
## FC_Fpos_AGGCCACCACATACTG-1FC_FI_GFPve_pos      0.6099917216
## FC_Fpos_AGGCCACTCTGCGAGC-1FC_FI_GFPve_pos     -0.4085887152
## FC_Fpos_AGGGAGTCAACCCTCT-1FC_FI_GFPve_pos      0.0564940569
## FC_Fpos_AGGGCCTCAACCGTAT-1FC_FI_GFPve_pos      1.4040473623
## FC_Fpos_AGGGTTTCAGGAATAT-1FC_FI_GFPve_pos     -0.4066183048
## FC_Fpos_AGGGTTTGTCCCTAAA-1FC_FI_GFPve_pos      1.6947879714
## FC_Fpos_AGGTCATAGATTCGCT-1FC_FI_GFPve_pos      0.5948808414
## FC_Fpos_AGGTCATTCGACGCTG-1FC_FI_GFPve_pos      0.6216655297
## FC_Fpos_AGGTGTTTCCACAGGC-1FC_FI_GFPve_pos      0.6797329110
## FC_Fpos_AGGTTACAGCATCCCG-1FC_FI_GFPve_pos      2.0891694946
## FC_Fpos_AGGTTACGTCTTAGTG-1FC_FI_GFPve_pos      0.9945796651
## FC_Fpos_AGTAACCCACTCCCTA-1FC_FI_GFPve_pos      1.2999550027
## FC_Fpos_AGTACCAAGGCTCTAT-1FC_FI_GFPve_pos     -0.1784130710
## FC_Fpos_AGTACCAGTTTGGCTA-1FC_FI_GFPve_pos      1.1240385694
## FC_Fpos_AGTACTGAGTCCTACA-1FC_FI_GFPve_pos     -0.6353887397
## FC_Fpos_AGTACTGCACACACTA-1FC_FI_GFPve_pos     -0.8660283762
## FC_Fpos_AGTACTGTCGAACACT-1FC_FI_GFPve_pos     -0.9100505191
## FC_Fpos_AGTAGCTCACGGCTAC-1FC_FI_GFPve_pos      2.0485201997
## FC_Fpos_AGTAGTCGTTCCTTGC-1FC_FI_GFPve_pos     -1.1836414414
## FC_Fpos_AGTAGTCGTTGTGCCG-1FC_FI_GFPve_pos     -0.3573314923
## FC_Fpos_AGTCATGCAGCGTTGC-1FC_FI_GFPve_pos     -1.3959592658
## FC_Fpos_AGTCTCCCACGGTCTG-1FC_FI_GFPve_pos      0.1286940051
## FC_Fpos_AGTGACTAGCTAGAAT-1FC_FI_GFPve_pos     -1.2256286221
## FC_Fpos_AGTGTTGCAGAAGTGC-1FC_FI_GFPve_pos     -0.9630475121
```

```
## FC_Fpos_ATACCGAAGGAAAGGT-1FC_FI_GFPve_pos     -1.1263469535
## FC_Fpos_ATACCTTCAGCGCGTT-1FC_FI_GFPve_pos     -0.4197926598
## FC_Fpos_ATACTTCTCTCACTCG-1FC_FI_GFPve_pos      1.1489579601
## FC_Fpos_ATAGACCTCGGAGTGA-1FC_FI_GFPve_pos     -0.8705578523
## FC_Fpos_ATAGAGAGTATGCTAC-1FC_FI_GFPve_pos      0.2648348880
## FC_Fpos_ATAGGCTGTTGCATAC-1FC_FI_GFPve_pos      1.1562756223
## FC_Fpos_ATCACGATCGGTCAGC-1FC_FI_GFPve_pos     -1.0297087508
## FC_Fpos_ATCCCTGAGCTGCGAA-1FC_FI_GFPve_pos     -0.4296915608
## FC_Fpos_ATCCTATAGATGGGCT-1FC_FI_GFPve_pos      1.9069880170
## FC_Fpos_ATCGATGAGAGGTATT-1FC_FI_GFPve_pos      0.0381925103
## FC_Fpos_ATCGATGTCTAGTTCT-1FC_FI_GFPve_pos      0.5832368952
## FC_Fpos_ATCGCCTCACTAGTAC-1FC_FI_GFPve_pos      1.0207921189
## FC_Fpos_ATCGGATAGTACTGGG-1FC_FI_GFPve_pos     -0.3721913176
## FC_Fpos_ATCGGATGTATCACGT-1FC_FI_GFPve_pos     -0.8518043834
## FC_Fpos_ATCGGCGTCTGTGCGG-1FC_FI_GFPve_pos     -0.8068284469
## FC_Fpos_ATCTCTAAGTCAACAA-1FC_FI_GFPve_pos      0.9692943615
## FC_Fpos_ATGCGATAGTGATGGC-1FC_FI_GFPve_pos     -0.6289823132
## FC_Fpos_ATGGGAGTCGGCTGAC-1FC_FI_GFPve_pos      0.0238769275
## FC_Fpos_ATGGTTGCACCTGAAT-1FC_FI_GFPve_pos      0.5528839452
## FC_Fpos_ATTATCCTCGTTCGCT-1FC_FI_GFPve_pos      1.2989523334
## FC_Fpos_ATTCATCGTGAGGCAT-1FC_FI_GFPve_pos      0.5986126584
## FC_Fpos_ATTCCCGCACATAACC-1FC_FI_GFPve_pos      2.1877562923
## FC_Fpos_ATTCCTACATATCGGT-1FC_FI_GFPve_pos     -0.8071665841
## FC_Fpos_ATTCGTTAGAATCGTA-1FC_FI_GFPve_pos      1.7274628801
## FC_Fpos_ATTCTTGCAACCGCCA-1FC_FI_GFPve_pos     -0.2280412393
## FC_Fpos_ATTCTTGTCTTGGTGA-1FC_FI_GFPve_pos      0.6902769727
## FC_Fpos_ATTGGGTTCTGTAACG-1FC_FI_GFPve_pos      1.3832920713
## FC_Fpos_ATTGTTCTCACTACTT-1FC_FI_GFPve_pos      0.9092090053
## FC_Fpos_ATTTACCAGGATCATA-1FC_FI_GFPve_pos     -1.5922442752
## FC_Fpos_CAACAACCAGCGGTCT-1FC_FI_GFPve_pos      0.8254644198
## FC_Fpos_CAACAGTGTACGTGTT-1FC_FI_GFPve_pos      0.8728533906
## FC_Fpos_CAACAGTTCCTTTAGT-1FC_FI_GFPve_pos      0.7584053201
## FC_Fpos_CAACCTCGTAACTAAG-1FC_FI_GFPve_pos      2.0609623117
## FC_Fpos_CAACCTCTCAAACGAA-1FC_FI_GFPve_pos      1.3350818080
## FC_Fpos_CAAGACTTCAACGCTA-1FC_FI_GFPve_pos      2.1292924804
## FC_Fpos_CAAGAGGCAGCTCATA-1FC_FI_GFPve_pos      1.7653217239
## FC_Fpos_CAAGGGACACAGCATT-1FC_FI_GFPve_pos      0.4627318007
## FC_Fpos_CAAGGGACAGAGGGTT-1FC_FI_GFPve_pos     -0.9296350556
## FC_Fpos_CAATACGTCCAGTGCG-1FC_FI_GFPve_pos      0.2981906307
## FC_Fpos_CAATCGAGTAACTGCT-1FC_FI_GFPve_pos      1.3177453202
## FC_Fpos_CAATCGAGTCGCACGT-1FC_FI_GFPve_pos      0.7822193426
## FC_Fpos_CACAACATCGCTTAAG-1FC_FI_GFPve_pos      1.5597423238
## FC_Fpos_CACACAAGTAGAGGAA-1FC_FI_GFPve_pos     -0.0136662426
## FC_Fpos_CACATGAAGACGAAGA-1FC_FI_GFPve_pos      0.8465690536
## FC_Fpos_CACATGATCCTTATGT-1FC_FI_GFPve_pos      2.0978933496
## FC_Fpos_CACCAAAGTGTGACCC-1FC_FI_GFPve_pos      0.7148929161
## FC_Fpos_CACCGTTCATCCTCAC-1FC_FI_GFPve_pos      2.2489981574
## FC_Fpos_CACGAATAGCCTCAGC-1FC_FI_GFPve_pos     -0.1502964574
## FC_Fpos_CACGGGTGTACAGTAA-1FC_FI_GFPve_pos      0.3101801348
## FC_Fpos_CACGGGTGTCCCGTGA-1FC_FI_GFPve_pos      1.4979840440
## FC_Fpos_CAGAGCCCACATTACG-1FC_FI_GFPve_pos      0.2454421592
## FC_Fpos_CAGCAGCCATCATCTT-1FC_FI_GFPve_pos      2.2588725013
## FC_Fpos_CAGCCAGTCTAGGCCG-1FC_FI_GFPve_pos      0.1867501212
## FC_Fpos_CAGGCCAAGAAACCCG-1FC_FI_GFPve_pos      1.7159961385
## FC_Fpos_CAGGCCATCGTACCTC-1FC_FI_GFPve_pos     -0.0210090506
## FC_Fpos_CAGGGCTAGCGTCGAA-1FC_FI_GFPve_pos      0.2426549417
## FC_Fpos_CAGTTAGCATGATCTG-1FC_FI_GFPve_pos     -0.1975530165
## FC_Fpos_CAGTTCCCATTGTCGA-1FC_FI_GFPve_pos     -0.1959725993
## FC_Fpos_CATACAGGTGAATGAT-1FC_FI_GFPve_pos     -1.2611564475
## FC_Fpos_CATAGACGTGGCTAGA-1FC_FI_GFPve_pos      1.7382457179
## FC_Fpos_CATCAAGTCAGAGCGA-1FC_FI_GFPve_pos      1.5550464315
## FC_Fpos_CATCCACGTCACTGAT-1FC_FI_GFPve_pos     -0.5233321267
## FC_Fpos_CATCCACGTCTACACA-1FC_FI_GFPve_pos     -1.0201520997
## FC_Fpos_CATCCACGTTGAAGTA-1FC_FI_GFPve_pos      0.2005213198
## FC_Fpos_CATCCCATCTGGGTCG-1FC_FI_GFPve_pos     -0.7226024347
## FC_Fpos_CATCGCTAGGCATTTC-1FC_FI_GFPve_pos      0.7875482363
## FC_Fpos_CATGCGGTCGTAGGAG-1FC_FI_GFPve_pos      2.3182952327
## FC_Fpos_CATGGATAGTGGTGGT-1FC_FI_GFPve_pos     -0.7886843758
## FC_Fpos_CATTCATCAATCTAGC-1FC_FI_GFPve_pos      1.0659331006
## FC_Fpos_CATTGAGCACGACTAT-1FC_FI_GFPve_pos     -1.3527730780
```

```
## FC_Fpos_CATTGCCCAGGACTAG-1FC_FI_GFPve_pos     -0.4530615288
## FC_Fpos_CATTGCCGTGGACTGA-1FC_FI_GFPve_pos      2.2257635516
## FC_Fpos_CCACACTGTTCTCAGA-1FC_FI_GFPve_pos      1.3971882981
## FC_Fpos_CCACGAGGTGACAACG-1FC_FI_GFPve_pos      0.3411424083
## FC_Fpos_CCACTTGCAAGTGGCA-1FC_FI_GFPve_pos     -0.7722579914
## FC_Fpos_CCACTTGCAGCACAAG-1FC_FI_GFPve_pos     -0.4669970351
## FC_Fpos_CCATCACAGAGTGACC-1FC_FI_GFPve_pos     -0.2622070986
## FC_Fpos_CCCATTGGTCAAGCCC-1FC_FI_GFPve_pos     -0.2756384390
## FC_Fpos_CCCATTGGTCGTGATT-1FC_FI_GFPve_pos      0.0294659210
## FC_Fpos_CCCGGAACACGCGCAT-1FC_FI_GFPve_pos      1.8577743692
## FC_Fpos_CCCGGAAGTCAAGGCA-1FC_FI_GFPve_pos      2.6392357272
## FC_Fpos_CCCTAACTCCACTTTA-1FC_FI_GFPve_pos      1.3080145044
## FC_Fpos_CCCTCTCGTAACCAGG-1FC_FI_GFPve_pos     -0.4453093487
## FC_Fpos_CCGAACGCAAGGACAC-1FC_FI_GFPve_pos      0.5550650281
## FC_Fpos_CCGAACGGTCTTGCTC-1FC_FI_GFPve_pos      0.5850465519
## FC_Fpos_CCGATCTCAGAGGACT-1FC_FI_GFPve_pos      1.2804381770
## FC_Fpos_CCGGACAGTCAAGCCC-1FC_FI_GFPve_pos     -1.1349943953
## FC_Fpos_CCGGGTAAGATGGGCT-1FC_FI_GFPve_pos      0.1336449755
## FC_Fpos_CCGGTAGGTCTTACTT-1FC_FI_GFPve_pos     -0.3947861272
## FC_Fpos_CCGTTCATCGTGAGAG-1FC_FI_GFPve_pos     -1.0459094125
## FC_Fpos_CCGTTCATCTGACGCG-1FC_FI_GFPve_pos      2.0431126279
## FC_Fpos_CCTACGTGTCTAACTG-1FC_FI_GFPve_pos     -1.4221475201
## FC_Fpos_CCTATCGGTATGGGAC-1FC_FI_GFPve_pos      0.9033877176
## FC_Fpos_CCTCATGAGGCCTAGA-1FC_FI_GFPve_pos     -0.7264434772
## FC_Fpos_CCTCTAGCAAATGGCG-1FC_FI_GFPve_pos      1.0113050861
## FC_Fpos_CCTGTTGTCGTCCATC-1FC_FI_GFPve_pos      0.4349147600
## FC_Fpos_CCTTTGGAGTATGTAG-1FC_FI_GFPve_pos     -0.0487817901
## FC_Fpos_CGAAGGAAGATCCCGC-1FC_FI_GFPve_pos     -0.1054707396
## FC_Fpos_CGAAGGATCGGCTATA-1FC_FI_GFPve_pos     -0.4052294570
## FC_Fpos_CGAAGTTCATCGATCA-1FC_FI_GFPve_pos      0.1948277575
## FC_Fpos_CGACAGCCAGAGCCCT-1FC_FI_GFPve_pos     -0.2251051145
## FC_Fpos_CGAGGAACATGAGTAA-1FC_FI_GFPve_pos     -0.3755536753
## FC_Fpos_CGATCGGTCTGTCCGT-1FC_FI_GFPve_pos      0.7366998119
## FC_Fpos_CGATGCGCAAACTCGT-1FC_FI_GFPve_pos     -1.1617451983
## FC_Fpos_CGCAGGTCAACCGACC-1FC_FI_GFPve_pos     -0.3554189282
## FC_Fpos_CGCAGGTTCATTTACC-1FC_FI_GFPve_pos     -0.9387203532
## FC_Fpos_CGCATAACACGCCACA-1FC_FI_GFPve_pos      0.2333900255
## FC_Fpos_CGCATAATCTGTCAGA-1FC_FI_GFPve_pos      1.8679187221
## FC_Fpos_CGCATGGTCGAGTCCG-1FC_FI_GFPve_pos      0.0870727428
## FC_Fpos_CGCCAGAAGTCTTCCC-1FC_FI_GFPve_pos      0.2490443510
## FC_Fpos_CGCCAGAGTTACCGTA-1FC_FI_GFPve_pos     -0.1465120691
## FC_Fpos_CGGACACAGCATCCCG-1FC_FI_GFPve_pos     -0.7249194461
## FC_Fpos_CGGACACCAGTCGGTC-1FC_FI_GFPve_pos     -0.3209509629
## FC_Fpos_CGGACACTCGGTAAGG-1FC_FI_GFPve_pos     -0.2634597200
## FC_Fpos_CGGCAGTTCCATCGTC-1FC_FI_GFPve_pos      0.7337219638
## FC_Fpos_CGGGACTGTACTGCCG-1FC_FI_GFPve_pos      1.3917701167
## FC_Fpos_CGGGCATAGCTACAAA-1FC_FI_GFPve_pos      1.9680298490
## FC_Fpos_CGGGTGTAGCGGACAT-1FC_FI_GFPve_pos      2.0216785592
## FC_Fpos_CGTAATGAGGTATTGA-1FC_FI_GFPve_pos      0.0132314188
## FC_Fpos_CGTAATGTCGGTCTAA-1FC_FI_GFPve_pos     -0.2573339420
## FC_Fpos_CGTAGTATCGCAGTTA-1FC_FI_GFPve_pos     -0.3742998617
## FC_Fpos_CGTCAAACACCTCAGG-1FC_FI_GFPve_pos     -0.9370471316
## FC_Fpos_CGTCCATAGGTTCTAC-1FC_FI_GFPve_pos     -0.5756009894
## FC_Fpos_CGTCCATTCCCGTAAA-1FC_FI_GFPve_pos      0.0278531087
## FC_Fpos_CGTGAATAGTACAGAT-1FC_FI_GFPve_pos      1.6704661769
## FC_Fpos_CGTGAATCAACACGAG-1FC_FI_GFPve_pos      2.4455702228
## FC_Fpos_CGTGAATCATGGACAG-1FC_FI_GFPve_pos     -0.7479227262
## FC_Fpos_CGTGCTTGTCTTCTAT-1FC_FI_GFPve_pos      1.1017434520
## FC_Fpos_CGTGCTTTCTCGTCAC-1FC_FI_GFPve_pos     -0.9465402442
## FC_Fpos_CGTTCTGTCAGGACGA-1FC_FI_GFPve_pos      1.1434525174
## FC_Fpos_CGTTGGGCAATGGGTG-1FC_FI_GFPve_pos     -0.8435266095
## FC_Fpos_CGTTGGGTCCTTTAGT-1FC_FI_GFPve_pos      0.9987959785
## FC_Fpos_CTAACCCGTCGTATTG-1FC_FI_GFPve_pos     -1.0144898969
## FC_Fpos_CTAAGTGAGAGGCGTT-1FC_FI_GFPve_pos     -0.6499626714
## FC_Fpos_CTACATTAGGGTCAAC-1FC_FI_GFPve_pos      1.3383493346
## FC_Fpos_CTACCTGGTTAAGGGC-1FC_FI_GFPve_pos     -1.0570159035
## FC_Fpos_CTACTATGTGTCTTAG-1FC_FI_GFPve_pos     -1.6903603154
## FC_Fpos_CTATAGGGTTCAACGT-1FC_FI_GFPve_pos     -0.9373465854
## FC_Fpos_CTCAATTCAGGTACGA-1FC_FI_GFPve_pos     -1.0378329831
## FC_Fpos_CTCAGGGCAATAAGGT-1FC_FI_GFPve_pos      1.6890184802
```

```
## FC_Fpos_CTCAGGGGTGGAACAC-1FC_FI_GFPve_pos      2.3074166698
## FC_Fpos_CTCCCAAGTAGGACCA-1FC_FI_GFPve_pos     -0.9088985997
## FC_Fpos_CTCCCTCCATCCGTGG-1FC_FI_GFPve_pos     -0.6147569495
## FC_Fpos_CTCCCTCCATTGCTTT-1FC_FI_GFPve_pos     -0.9714618998
## FC_Fpos_CTCCCTCGTATACCTG-1FC_FI_GFPve_pos     -0.0656011182
## FC_Fpos_CTCCTCCGTGACTGAG-1FC_FI_GFPve_pos      0.0336977762
## FC_Fpos_CTCCTTTAGCGTCAAG-1FC_FI_GFPve_pos      0.7984134478
## FC_Fpos_CTCTCAGAGGATTCAA-1FC_FI_GFPve_pos      0.8570117873
## FC_Fpos_CTCTGGTGTCCTTAAG-1FC_FI_GFPve_pos      0.2152310235
## FC_Fpos_CTGAATGAGTCGAGGT-1FC_FI_GFPve_pos      2.0343446655
## FC_Fpos_CTGATCCTCTAAGCCA-1FC_FI_GFPve_pos     -0.4864483553
## FC_Fpos_CTGCAGGGTCACGTGC-1FC_FI_GFPve_pos     -0.7329943257
## FC_Fpos_CTGCATCAGTAACAGT-1FC_FI_GFPve_pos      1.3161742610
## FC_Fpos_CTGGCAGCATCATTTC-1FC_FI_GFPve_pos      2.2782583160
## FC_Fpos_CTGGCAGTCCCTCTAG-1FC_FI_GFPve_pos      1.8999982995
## FC_Fpos_CTGGTCTGTGGCTAGA-1FC_FI_GFPve_pos      0.1749336121
## FC_Fpos_CTGTAGACAGATAAAC-1FC_FI_GFPve_pos     -0.1233703422
## FC_Fpos_CTGTAGAGTACAAGTA-1FC_FI_GFPve_pos     -0.5675383883
## FC_Fpos_CTGTGGGTCCCGAGAC-1FC_FI_GFPve_pos      0.7245227617
## FC_Fpos_CTTAGGACAAATAAGC-1FC_FI_GFPve_pos     -0.1362658965
## FC_Fpos_CTTCAATGTCATCAGT-1FC_FI_GFPve_pos     -0.0003738108
## FC_Fpos_CTTCGGTTCGCTTGCT-1FC_FI_GFPve_pos      0.2199987245
## FC_Fpos_CTTCGGTTCGGTCGAC-1FC_FI_GFPve_pos      0.9546378178
## FC_Fpos_CTTCGGTTCTGAACGT-1FC_FI_GFPve_pos      0.8899434132
## FC_Fpos_CTTCTAATCTCCTGAC-1FC_FI_GFPve_pos      1.9619127435
## FC_Fpos_CTTGAGATCAGAGCAG-1FC_FI_GFPve_pos     -0.0153212252
## FC_Fpos_CTTTCGGCAAGCAGGT-1FC_FI_GFPve_pos      0.8740148467
## FC_Fpos_GAAACCTCATCGTTCC-1FC_FI_GFPve_pos     -0.4072089868
## FC_Fpos_GAAATGATCAATCTTC-1FC_FI_GFPve_pos     -0.7721557694
## FC_Fpos_GAACACTAGTAGCCAG-1FC_FI_GFPve_pos      1.8767411632
## FC_Fpos_GAACACTCAATGGCAG-1FC_FI_GFPve_pos      0.8710165543
## FC_Fpos_GAACGTTGTGTCTAAC-1FC_FI_GFPve_pos     -0.7688625532
## FC_Fpos_GAAGCCCTCTACTTCA-1FC_FI_GFPve_pos      1.2801232261
## FC_Fpos_GAAGGACGTACCCGAC-1FC_FI_GFPve_pos      2.2556102199
## FC_Fpos_GAAGGGTAGGTCCGAA-1FC_FI_GFPve_pos      0.8643150253
## FC_Fpos_GAAGTAACAACATACC-1FC_FI_GFPve_pos     -0.1573728996
## FC_Fpos_GAAGTAACAGGCACAA-1FC_FI_GFPve_pos     -0.8292260843
## FC_Fpos_GAATAGATCCTCACTG-1FC_FI_GFPve_pos      1.2817428035
## FC_Fpos_GAATCGTAGTCATGAA-1FC_FI_GFPve_pos      1.5936465186
## FC_Fpos_GACAGCCGTACCTGTA-1FC_FI_GFPve_pos     -1.0653864222
## FC_Fpos_GACATCACAGCGACAA-1FC_FI_GFPve_pos     -0.9062067347
## FC_Fpos_GACATCAGTCCTGAAT-1FC_FI_GFPve_pos      0.0840385051
## FC_Fpos_GACATCATCTTTACAC-1FC_FI_GFPve_pos      0.0597688516
## FC_Fpos_GACCAATGTCGGAAAC-1FC_FI_GFPve_pos      2.6494691295
## FC_Fpos_GACCCAGAGGTATAGT-1FC_FI_GFPve_pos      1.6916149778
## FC_Fpos_GACCCTTAGCGCCTAC-1FC_FI_GFPve_pos      1.1065120620
## FC_Fpos_GACCGTGAGTGACCTT-1FC_FI_GFPve_pos      0.9716091079
## FC_Fpos_GACGTTACACTCCTGT-1FC_FI_GFPve_pos      1.2325271291
## FC_Fpos_GACGTTATCAACACCA-1FC_FI_GFPve_pos      1.6950894517
## FC_Fpos_GACTCAAAGTCGCGAA-1FC_FI_GFPve_pos      1.0816786808
## FC_Fpos_GACTCAACATAATGAG-1FC_FI_GFPve_pos      2.0835381669
## FC_Fpos_GACTCTCAGTATGAAC-1FC_FI_GFPve_pos     -0.8690558153
## FC_Fpos_GAGACCCAGCGAGTAC-1FC_FI_GFPve_pos      0.6997768206
## FC_Fpos_GAGACTTAGCTGAAGC-1FC_FI_GFPve_pos      0.6080440802
## FC_Fpos_GAGACTTCACTACGGC-1FC_FI_GFPve_pos     -0.6025434213
## FC_Fpos_GAGCCTGTCGAAACAA-1FC_FI_GFPve_pos     -0.1406835842
## FC_Fpos_GAGCTGCAGTTGGCTT-1FC_FI_GFPve_pos      1.9725323838
## FC_Fpos_GAGGGTACAGCTGTTA-1FC_FI_GFPve_pos     -1.1254010278
## FC_Fpos_GAGTGTTAGTAGGAAG-1FC_FI_GFPve_pos      0.9523174686
## FC_Fpos_GAGTGTTCAGTTGTTG-1FC_FI_GFPve_pos      2.2256572169
## FC_Fpos_GAGTGTTTCGTAGAGG-1FC_FI_GFPve_pos      1.7089006824
## FC_Fpos_GAGTTACGTGGGAGAG-1FC_FI_GFPve_pos     -0.9901102858
## FC_Fpos_GATAGCTAGTCTAACC-1FC_FI_GFPve_pos      0.5881790740
## FC_Fpos_GATCATGTCACACCCT-1FC_FI_GFPve_pos     -0.0138755518
## FC_Fpos_GATCCCTTCTATCGTT-1FC_FI_GFPve_pos      0.8425910992
## FC_Fpos_GATGAGGCACGTCGTG-1FC_FI_GFPve_pos      0.5878967864
## FC_Fpos_GATGGAGCATAATCCG-1FC_FI_GFPve_pos     -0.6297913271
## FC_Fpos_GATGGAGCATCCCGTT-1FC_FI_GFPve_pos      1.7744805736
## FC_Fpos_GATGTTGAGAGAGCGG-1FC_FI_GFPve_pos      2.3649873656
## FC_Fpos_GATTCGATCATCGTAG-1FC_FI_GFPve_pos      0.6215268296
```

```
## FC_Fpos_GATTCGATCCCAATAG-1FC_FI_GFPve_pos     -0.0725528973
## FC_Fpos_GATTTCTTCCTGGCTT-1FC_FI_GFPve_pos      0.2557374013
## FC_Fpos_GCAACCGTCGAGAGCA-1FC_FI_GFPve_pos      0.2973418308
## FC_Fpos_GCAACCGTCTACACAG-1FC_FI_GFPve_pos     -0.1744275170
## FC_Fpos_GCACGGTCACCGGTCA-1FC_FI_GFPve_pos     -1.3271786052
## FC_Fpos_GCACGTGCATCACGGC-1FC_FI_GFPve_pos     -0.6915504413
## FC_Fpos_GCACTAAAGACTACCT-1FC_FI_GFPve_pos     -0.4873117286
## FC_Fpos_GCAGGCTGTATTGCCA-1FC_FI_GFPve_pos     -1.2284222918
## FC_Fpos_GCATCGGGTACCTATG-1FC_FI_GFPve_pos      0.5452532989
## FC_Fpos_GCATGATTCGACACCG-1FC_FI_GFPve_pos      2.4147143287
## FC_Fpos_GCCAGCACATGGAACG-1FC_FI_GFPve_pos     -0.7595756131
## FC_Fpos_GCCAGGTTCACGATAC-1FC_FI_GFPve_pos     -1.1545591431
## FC_Fpos_GCCGATGTCAAATAGG-1FC_FI_GFPve_pos      0.4491986555
## FC_Fpos_GCGGATCCATACGCAT-1FC_FI_GFPve_pos      1.5103191060
## FC_Fpos_GCGTGCACAAGGCTTT-1FC_FI_GFPve_pos     -0.0525130111
## FC_Fpos_GCGTTTCTCGTGCGAC-1FC_FI_GFPve_pos     -0.1362311470
## FC_Fpos_GCTACAACACGCTATA-1FC_FI_GFPve_pos      1.7422245664
## FC_Fpos_GCTACCTCATTGTCGA-1FC_FI_GFPve_pos     -0.3511512357
## FC_Fpos_GCTCAAACACGTCGGT-1FC_FI_GFPve_pos      1.0189590973
## FC_Fpos_GCTCAAAGTCTTGCTC-1FC_FI_GFPve_pos     -0.5742932039
## FC_Fpos_GCTGAATGTAGCTTAC-1FC_FI_GFPve_pos      0.3648925347
## FC_Fpos_GCTGGGTAGATAGCTA-1FC_FI_GFPve_pos      0.3251008612
## FC_Fpos_GCTGGGTCACGTTGGC-1FC_FI_GFPve_pos     -0.5381168204
## FC_Fpos_GCTGGGTGTTACGGAG-1FC_FI_GFPve_pos     -0.5513067919
## FC_Fpos_GCTGGGTTCTCGGTAA-1FC_FI_GFPve_pos     -0.2036398130
## FC_Fpos_GCTTTCGGTCCGGATC-1FC_FI_GFPve_pos      2.3021178168
## FC_Fpos_GGAAGTGAGTTCATGC-1FC_FI_GFPve_pos      1.0742712540
## FC_Fpos_GGAATCTAGATAGTGT-1FC_FI_GFPve_pos     -1.0168405848
## FC_Fpos_GGAGATGTCCGATAAC-1FC_FI_GFPve_pos      1.3956193847
## FC_Fpos_GGAGGATGTTCTCAGA-1FC_FI_GFPve_pos      0.8552585763
## FC_Fpos_GGCACGTTCCACTGAA-1FC_FI_GFPve_pos      1.0843245906
## FC_Fpos_GGCAGTCCATGTGGTT-1FC_FI_GFPve_pos     -0.7679168063
## FC_Fpos_GGCAGTCTCACCGCTT-1FC_FI_GFPve_pos      0.1867358458
## FC_Fpos_GGCTTTCAGAAGGCTC-1FC_FI_GFPve_pos      0.0756713839
## FC_Fpos_GGGACCTTCGTAATGC-1FC_FI_GFPve_pos     -0.2018019038
## FC_Fpos_GGGACTCCACAAGGTG-1FC_FI_GFPve_pos      1.2101768178
## FC_Fpos_GGGAGATTCATTTGTC-1FC_FI_GFPve_pos      1.5955493373
## FC_Fpos_GGGATGAAGTATGATG-1FC_FI_GFPve_pos     -1.2477737742
## FC_Fpos_GGGATGACAAGAATAC-1FC_FI_GFPve_pos      1.8243658466
## FC_Fpos_GGGCCATCATGTAACC-1FC_FI_GFPve_pos     -0.0155057835
## FC_Fpos_GGGCCATCATTCTCCG-1FC_FI_GFPve_pos      0.7725813312
## FC_Fpos_GGGCTCACATCCGTTC-1FC_FI_GFPve_pos      0.9487341565
## FC_Fpos_GGGTATTGTTAGTTCG-1FC_FI_GFPve_pos      1.5271654052
## FC_Fpos_GGGTCACGTAGTTACC-1FC_FI_GFPve_pos      0.7877475542
## FC_Fpos_GGGTGAACACATAGCT-1FC_FI_GFPve_pos      0.0882831251
## FC_Fpos_GGGTTATGTGTTTACG-1FC_FI_GFPve_pos     -0.4346919733
## FC_Fpos_GGGTTATTCCATTCGC-1FC_FI_GFPve_pos      1.6751545591
## FC_Fpos_GGGTTTAAGCAACTCT-1FC_FI_GFPve_pos      1.1066938562
## FC_Fpos_GGGTTTAGTCCTGAAT-1FC_FI_GFPve_pos      1.2160696906
## FC_Fpos_GGTAACTAGAGCCTGA-1FC_FI_GFPve_pos      1.0265226883
## FC_Fpos_GGTAATCCACCATATG-1FC_FI_GFPve_pos      0.6358399910
## FC_Fpos_GGTAATCTCGAGATGG-1FC_FI_GFPve_pos      0.4376895947
## FC_Fpos_GGTCACGAGTTTCGGT-1FC_FI_GFPve_pos      0.4897956175
## FC_Fpos_GGTGAAGTCATAGGCT-1FC_FI_GFPve_pos      0.5791360063
## FC_Fpos_GGTGGCTGTAGGACTG-1FC_FI_GFPve_pos      1.1097718400
## FC_Fpos_GGTGGCTTCATTGCTT-1FC_FI_GFPve_pos      0.1837464717
## FC_Fpos_GGTGGCTTCCACTAGA-1FC_FI_GFPve_pos     -0.6121438223
## FC_Fpos_GGTTAACCAGGTCAGA-1FC_FI_GFPve_pos     -1.2308981496
## FC_Fpos_GGTTAACGTAGCGTCC-1FC_FI_GFPve_pos      1.4197672528
## FC_Fpos_GGTTGTAAGATGCGAC-1FC_FI_GFPve_pos      1.4202437324
## FC_Fpos_GGTTGTAGTGAATGAT-1FC_FI_GFPve_pos      0.1072623387
## FC_Fpos_GTAACACAGCTAGAGC-1FC_FI_GFPve_pos      0.4576001984
## FC_Fpos_GTAACACCACTGAGTT-1FC_FI_GFPve_pos     -0.1266373622
## FC_Fpos_GTAAGTCTCATTACGG-1FC_FI_GFPve_pos      1.6775894088
## FC_Fpos_GTAAGTCTCTGAGGTT-1FC_FI_GFPve_pos     -0.6485900360
## FC_Fpos_GTAATGCAGTACAACA-1FC_FI_GFPve_pos      1.3721653146
## FC_Fpos_GTACAACAGTAACAGT-1FC_FI_GFPve_pos      0.0510948633
## FC_Fpos_GTACAACTCTCGCTCA-1FC_FI_GFPve_pos      0.0628659648
## FC_Fpos_GTACAGTGTGTCCTAA-1FC_FI_GFPve_pos     -0.4703556376
## FC_Fpos_GTAGAAACATGCAGCC-1FC_FI_GFPve_pos      0.0417791565
```

```
## FC_Fpos_GTAGAGGTCAACTCTT-1FC_FI_GFPve_pos      0.6663690132
## FC_Fpos_GTAGCTAAGCAAGTCG-1FC_FI_GFPve_pos     -0.5581756907
## FC_Fpos_GTAGGTTGTTATCTGG-1FC_FI_GFPve_pos      1.8049592895
## FC_Fpos_GTAGTACGTTCGTGCG-1FC_FI_GFPve_pos     -1.4722201901
## FC_Fpos_GTATTTCTCGTAGAGG-1FC_FI_GFPve_pos      0.2521067185
## FC_Fpos_GTCAAACTCATATGGC-1FC_FI_GFPve_pos     -0.4536750394
## FC_Fpos_GTCAAGTAGCCGAATG-1FC_FI_GFPve_pos      1.8780481738
## FC_Fpos_GTCACGGAGCAATTCC-1FC_FI_GFPve_pos     -0.9191979247
## FC_Fpos_GTCACGGCATGACTCA-1FC_FI_GFPve_pos      0.3635472876
## FC_Fpos_GTCACTCAGTCAGCCC-1FC_FI_GFPve_pos      1.9467513484
## FC_Fpos_GTCATCCAGCTGACAG-1FC_FI_GFPve_pos     -0.0280585515
## FC_Fpos_GTCATTTGTTACGGAG-1FC_FI_GFPve_pos     -0.5098258930
## FC_Fpos_GTCCTCACATGACTGT-1FC_FI_GFPve_pos      0.3799864871
## FC_Fpos_GTCGAATCATTAAAGG-1FC_FI_GFPve_pos      0.2722149265
## FC_Fpos_GTCGCGAGTATGATCC-1FC_FI_GFPve_pos     -0.9188206273
## FC_Fpos_GTCGTAATCCAGCCTT-1FC_FI_GFPve_pos     -0.6770812350
## FC_Fpos_GTCTACCAGGTCCCGT-1FC_FI_GFPve_pos      0.4106518549
## FC_Fpos_GTCTCACGTAGAATGT-1FC_FI_GFPve_pos     -0.3426068204
## FC_Fpos_GTGACGCCAACGGGTA-1FC_FI_GFPve_pos     -0.8418555337
## FC_Fpos_GTGACGCCACTGGATT-1FC_FI_GFPve_pos      0.9631574077
## FC_Fpos_GTGAGGATCTCAACGA-1FC_FI_GFPve_pos      0.1373127212
## FC_Fpos_GTGAGTTCAAATCGTC-1FC_FI_GFPve_pos      1.5559068841
## FC_Fpos_GTGCACGTCAAGCCGC-1FC_FI_GFPve_pos     -0.6031254607
## FC_Fpos_GTGCACGTCTGCGATA-1FC_FI_GFPve_pos      1.0394695920
## FC_Fpos_GTGCAGCAGGAAAGGT-1FC_FI_GFPve_pos     -1.4280787783
## FC_Fpos_GTGCGTGGTTCTTGTT-1FC_FI_GFPve_pos     -0.0036749917
## FC_Fpos_GTGCTGGCATTCAGCA-1FC_FI_GFPve_pos     -0.1618320721
## FC_Fpos_GTGCTGGGTAGCTTTG-1FC_FI_GFPve_pos     -1.1774792748
## FC_Fpos_GTGCTTCCACTAGTAC-1FC_FI_GFPve_pos      0.4084820074
## FC_Fpos_GTGCTTCTCTGTGCAA-1FC_FI_GFPve_pos      1.2130113763
## FC_Fpos_GTGGGAACACAGAGAC-1FC_FI_GFPve_pos      0.4918388647
## FC_Fpos_GTGGGAAGTGATACAA-1FC_FI_GFPve_pos      1.2430549783
## FC_Fpos_GTGGGAATCTAGCCAA-1FC_FI_GFPve_pos      0.9031577629
## FC_Fpos_GTGGTTAAGCAGCCTC-1FC_FI_GFPve_pos      2.2107839507
## FC_Fpos_GTGTTCCTCATCACAG-1FC_FI_GFPve_pos     -0.1288205313
## FC_Fpos_GTTACAGGTCGCTGCA-1FC_FI_GFPve_pos     -0.1916221457
## FC_Fpos_GTTACAGGTGTTACAC-1FC_FI_GFPve_pos     -0.4099973517
## FC_Fpos_GTTACGATCATAAGGA-1FC_FI_GFPve_pos     -1.4502424079
## FC_Fpos_GTTAGACGTATCGAGG-1FC_FI_GFPve_pos     -1.1420204716
## FC_Fpos_GTTAGTGAGTAGTCTC-1FC_FI_GFPve_pos     -0.4997334557
## FC_Fpos_GTTATGGAGGTACTGG-1FC_FI_GFPve_pos      0.0811423489
## FC_Fpos_GTTATGGTCAAATGAG-1FC_FI_GFPve_pos     -1.1883710700
## FC_Fpos_GTTCATTAGGTACAGC-1FC_FI_GFPve_pos     -0.0992518591
## FC_Fpos_GTTCATTGTCTCTCTG-1FC_FI_GFPve_pos      1.6924052161
## FC_Fpos_GTTCCGTCAGAGAATT-1FC_FI_GFPve_pos     -1.8440278846
## FC_Fpos_GTTCGCTGTTTCGTAG-1FC_FI_GFPve_pos      2.3037443084
## FC_Fpos_GTTCTATGTATGGGAC-1FC_FI_GFPve_pos      0.3209073317
## FC_Fpos_GTTCTATGTTACCCAA-1FC_FI_GFPve_pos      0.7260507864
## FC_Fpos_GTTGCGGTCAGTGATC-1FC_FI_GFPve_pos      1.5759326142
## FC_Fpos_GTTGCTCCATCAGCAT-1FC_FI_GFPve_pos     -1.0401703196
## FC_Fpos_GTTGTAGGTTGGGATG-1FC_FI_GFPve_pos     -0.5601888972
## FC_Fpos_GTTGTCCAGAACAGGA-1FC_FI_GFPve_pos     -0.7008001285
## FC_Fpos_GTTGTCCCATGGCGCT-1FC_FI_GFPve_pos     -1.1488448458
## FC_Fpos_GTTGTGAAGGTGATAT-1FC_FI_GFPve_pos      1.9751483125
## FC_Fpos_GTTGTGAAGTAGCAAT-1FC_FI_GFPve_pos     -0.8123357373
## FC_Fpos_GTTGTGACAAGAGTTA-1FC_FI_GFPve_pos     -0.6313610750
## FC_Fpos_GTTTACTGTTGCATGT-1FC_FI_GFPve_pos      1.3945559186
## FC_Fpos_TAACACGGTATGAAAC-1FC_FI_GFPve_pos     -1.0197801667
## FC_Fpos_TAACCAGCAGCGTACC-1FC_FI_GFPve_pos     -0.9516177254
## FC_Fpos_TAACGACTCGACGCTG-1FC_FI_GFPve_pos      1.0817239207
## FC_Fpos_TAACTTCAGCATGGGT-1FC_FI_GFPve_pos      0.6174919528
## FC_Fpos_TAAGCCAAGCGACTTT-1FC_FI_GFPve_pos      1.3146321696
## FC_Fpos_TAATTCCAGCACTCTA-1FC_FI_GFPve_pos      1.7754841966
## FC_Fpos_TAATTCCAGCTCCGAC-1FC_FI_GFPve_pos      2.0952318830
## FC_Fpos_TACAGGTAGTAATTGG-1FC_FI_GFPve_pos     -0.2557505446
## FC_Fpos_TACAGGTTCACTGGGC-1FC_FI_GFPve_pos      1.3960229081
## FC_Fpos_TACATTCGTAGTGATA-1FC_FI_GFPve_pos      1.2928762359
## FC_Fpos_TACCGAAGTGATATAG-1FC_FI_GFPve_pos     -0.2692503112
## FC_Fpos_TACCGGGAGACATAGT-1FC_FI_GFPve_pos      0.0227105317
## FC_Fpos_TACCGGGGTGTGTTTG-1FC_FI_GFPve_pos     -0.1145451444
```

```
## FC_Fpos_TACCTGCGTGTAGCAG-1FC_FI_GFPve_pos     -1.1112773495
## FC_Fpos_TACCTGCGTTTACTGG-1FC_FI_GFPve_pos     -0.0403903353
## FC_Fpos_TACGCTCTCATTCCTA-1FC_FI_GFPve_pos      0.6234677953
## FC_Fpos_TACGCTCTCCTGGCTT-1FC_FI_GFPve_pos      1.2179874105
## FC_Fpos_TACTGCCAGTCCCGGT-1FC_FI_GFPve_pos      0.1435787005
## FC_Fpos_TACTTACGTAGCCCTG-1FC_FI_GFPve_pos      2.4966945571
## FC_Fpos_TACTTCAAGGTGATCG-1FC_FI_GFPve_pos      1.5722801608
## FC_Fpos_TAGACTGTCTTAGTTC-1FC_FI_GFPve_pos     -0.0002337771
## FC_Fpos_TAGAGTCAGTTACGGG-1FC_FI_GFPve_pos     -0.4604832607
## FC_Fpos_TAGGTACGTCGCGGTT-1FC_FI_GFPve_pos      0.6207958979
## FC_Fpos_TAGTGCATCGCAATTG-1FC_FI_GFPve_pos     -1.6045785027
## FC_Fpos_TATACCTGTCAAGCCC-1FC_FI_GFPve_pos      2.1775689048
## FC_Fpos_TATCAGGCATAGCACT-1FC_FI_GFPve_pos     -0.5141674953
## FC_Fpos_TATCTTGTCCTAAGTG-1FC_FI_GFPve_pos      0.6191373986
## FC_Fpos_TATTCCAAGGTTTGAA-1FC_FI_GFPve_pos     -0.0440426695
## FC_Fpos_TATTCCACATGGAACG-1FC_FI_GFPve_pos      2.0258673352
## FC_Fpos_TATTGCTCAAAGCACG-1FC_FI_GFPve_pos      0.0325874460
## FC_Fpos_TATTTCGAGCAATTCC-1FC_FI_GFPve_pos     -1.0455218631
## FC_Fpos_TATTTCGGTCTTGAGT-1FC_FI_GFPve_pos     -0.2495051401
## FC_Fpos_TCAAGCATCGTCGCTT-1FC_FI_GFPve_pos      0.6127011699
## FC_Fpos_TCAAGTGCACGCGTCA-1FC_FI_GFPve_pos      0.7950852436
## FC_Fpos_TCAATCTGTGGATTTC-1FC_FI_GFPve_pos     -0.6710081178
## FC_Fpos_TCACGGGAGGTACTGG-1FC_FI_GFPve_pos      0.7552646918
## FC_Fpos_TCACGGGGTTGATGTC-1FC_FI_GFPve_pos     -0.1306758153
## FC_Fpos_TCAGGTAGTGCCGTAC-1FC_FI_GFPve_pos     -1.3345394212
## FC_Fpos_TCAGGTATCTATTCGT-1FC_FI_GFPve_pos     -0.5434562045
## FC_Fpos_TCAGTGAGTGCCGTAC-1FC_FI_GFPve_pos      0.4327639264
## FC_Fpos_TCAGTTTTCTGTCGCT-1FC_FI_GFPve_pos      1.1122983617
## FC_Fpos_TCATCATCACTTGGCG-1FC_FI_GFPve_pos      0.6974424643
## FC_Fpos_TCATGAGGTTTGCAGT-1FC_FI_GFPve_pos      0.0484981050
## FC_Fpos_TCATGCCAGCGTCTGC-1FC_FI_GFPve_pos      1.2625093383
## FC_Fpos_TCATGGAAGCCGATTT-1FC_FI_GFPve_pos     -1.1292096453
## FC_Fpos_TCATGTTCACATATCG-1FC_FI_GFPve_pos     -1.4223513680
## FC_Fpos_TCATGTTGTAAGCTCT-1FC_FI_GFPve_pos      1.6274058742
## FC_Fpos_TCATGTTGTACGATCT-1FC_FI_GFPve_pos      0.7711892647
## FC_Fpos_TCATTACAGGCATGCA-1FC_FI_GFPve_pos      1.4095035714
## FC_Fpos_TCATTACCATCTTCGC-1FC_FI_GFPve_pos     -0.1102771538
## FC_Fpos_TCATTGTTCTAGCAAC-1FC_FI_GFPve_pos      2.1633877677
## FC_Fpos_TCCACCACAGAAATTG-1FC_FI_GFPve_pos      2.4646453780
## FC_Fpos_TCCACCATCTTACTGT-1FC_FI_GFPve_pos      1.4511670989
## FC_Fpos_TCCAGAACAACGCCCA-1FC_FI_GFPve_pos      1.3754012508
## FC_Fpos_TCCATCGGTAAGATCA-1FC_FI_GFPve_pos      1.3497372789
## FC_Fpos_TCCCATGCACTTGTCC-1FC_FI_GFPve_pos      0.8186953587
## FC_Fpos_TCCGAAAGTGGAGGTT-1FC_FI_GFPve_pos     -0.1532833355
## FC_Fpos_TCCGATCAGGGTTAGC-1FC_FI_GFPve_pos     -0.7202540713
## FC_Fpos_TCCTAATAGACTCGAG-1FC_FI_GFPve_pos     -0.6901879984
## FC_Fpos_TCCTCCCAGTTGCGAG-1FC_FI_GFPve_pos      1.6934266013
## FC_Fpos_TCCTCGAAGATTGAGT-1FC_FI_GFPve_pos     -0.6567828732
## FC_Fpos_TCCTCTTAGCTCCACG-1FC_FI_GFPve_pos      0.8730912131
## FC_Fpos_TCCTCTTTCGAGTACT-1FC_FI_GFPve_pos      0.7229134363
## FC_Fpos_TCCTGCAAGCTCCACG-1FC_FI_GFPve_pos     -0.0335474091
## FC_Fpos_TCCTGCAGTGGCTAGA-1FC_FI_GFPve_pos      0.1137713368
## FC_Fpos_TCCTGCATCCATCCGT-1FC_FI_GFPve_pos     -0.1324083286
## FC_Fpos_TCCTTCTCAGGAATCG-1FC_FI_GFPve_pos     -0.5677990394
## FC_Fpos_TCCTTTCAGCTCATAC-1FC_FI_GFPve_pos      2.1006357593
## FC_Fpos_TCCTTTCCAGTTTGGT-1FC_FI_GFPve_pos      1.9747574252
## FC_Fpos_TCGAACAAGGTAAGTT-1FC_FI_GFPve_pos      0.5582185609
## FC_Fpos_TCGACCTTCAGGTAAA-1FC_FI_GFPve_pos      1.0196486038
## FC_Fpos_TCGCACTAGGCCGCTT-1FC_FI_GFPve_pos      1.0780618710
## FC_Fpos_TCGCACTAGGCGTTAG-1FC_FI_GFPve_pos      0.7135546130
## FC_Fpos_TCGCACTAGGGCTTCC-1FC_FI_GFPve_pos      0.6816305441
## FC_Fpos_TCGCACTGTACACGCC-1FC_FI_GFPve_pos     -0.2767283099
## FC_Fpos_TCGGATAAGTAGGGTC-1FC_FI_GFPve_pos     -0.5918659049
## FC_Fpos_TCGGATAGTTCCAGGC-1FC_FI_GFPve_pos      1.0140216273
## FC_Fpos_TCGGGCAAGAGCCTGA-1FC_FI_GFPve_pos      0.4706656856
## FC_Fpos_TCGGTCTCATGTGCTA-1FC_FI_GFPve_pos     -0.5830359536
## FC_Fpos_TCGTGCTAGACTCTTG-1FC_FI_GFPve_pos     -0.4136375266
## FC_Fpos_TCGTGCTCAAGGACAC-1FC_FI_GFPve_pos     -0.7849204617
## FC_Fpos_TCGTGCTCACGGTAGA-1FC_FI_GFPve_pos      2.1758868617
## FC_Fpos_TCGTGGGAGTAGCAAT-1FC_FI_GFPve_pos     -0.6139175969
```

```
## FC_Fpos_TCTAACTCATAGAGGC-1FC_FI_GFPve_pos      0.4041358751
## FC_Fpos_TCTAACTGTTATTCTC-1FC_FI_GFPve_pos      0.8257284683
## FC_Fpos_TCTATCACAGAGTTCT-1FC_FI_GFPve_pos     -0.6216002780
## FC_Fpos_TCTCACGGTGTTCCTC-1FC_FI_GFPve_pos      2.1076238078
## FC_Fpos_TCTCACGTCCTACTGC-1FC_FI_GFPve_pos      1.2457506580
## FC_Fpos_TCTCACGTCTCGTCAC-1FC_FI_GFPve_pos     -0.1560139733
## FC_Fpos_TCTCAGCTCCGCAACG-1FC_FI_GFPve_pos     -0.5044301229
## FC_Fpos_TCTCCGAAGTGCCCGT-1FC_FI_GFPve_pos      0.0065939424
## FC_Fpos_TCTTAGTGTACCAATC-1FC_FI_GFPve_pos      0.0001854894
## FC_Fpos_TCTTAGTGTAGACACG-1FC_FI_GFPve_pos      1.0691680831
## FC_Fpos_TCTTCCTCATCTCAAG-1FC_FI_GFPve_pos      0.8436669630
## FC_Fpos_TCTTGCGGTAGGAAAG-1FC_FI_GFPve_pos     -1.2649357396
## FC_Fpos_TCTTTGAAGTGCACAG-1FC_FI_GFPve_pos     -0.6370804983
## FC_Fpos_TGAACGTCAGAAGCTG-1FC_FI_GFPve_pos      1.4543732328
## FC_Fpos_TGAATCGCAATACGCT-1FC_FI_GFPve_pos      2.8807666225
## FC_Fpos_TGAATCGTCTAAGAAG-1FC_FI_GFPve_pos     -1.2644420939
## FC_Fpos_TGACAGTTCAACTCTT-1FC_FI_GFPve_pos      0.0461961364
## FC_Fpos_TGACTCCCAGCTTCGG-1FC_FI_GFPve_pos      0.9737412256
## FC_Fpos_TGAGCGCGTTTGGGAG-1FC_FI_GFPve_pos      1.6482970638
## FC_Fpos_TGAGGAGAGCCTGAGA-1FC_FI_GFPve_pos     -0.9472631293
## FC_Fpos_TGAGGAGTCAAGTAAG-1FC_FI_GFPve_pos      1.4933931750
## FC_Fpos_TGAGGTTCAGTAGAAT-1FC_FI_GFPve_pos      0.9507478995
## FC_Fpos_TGAGGTTGTCAGTTTG-1FC_FI_GFPve_pos      1.2766221685
## FC_Fpos_TGAGTCAAGAGCACTG-1FC_FI_GFPve_pos     -1.4840864020
## FC_Fpos_TGATCTTGTCTAACGT-1FC_FI_GFPve_pos      0.6060454768
## FC_Fpos_TGATGCAGTTTAGTCG-1FC_FI_GFPve_pos     -0.8736993748
## FC_Fpos_TGATTCTCAAAGTATG-1FC_FI_GFPve_pos      1.2113435191
## FC_Fpos_TGATTCTGTCATAACC-1FC_FI_GFPve_pos      1.2479864282
## FC_Fpos_TGATTTCCAGTAACGG-1FC_FI_GFPve_pos     -0.8351955491
## FC_Fpos_TGCAGATAGACCTCCG-1FC_FI_GFPve_pos     -0.0148676264
## FC_Fpos_TGCAGATTCTCTTCAA-1FC_FI_GFPve_pos      0.1163263067
## FC_Fpos_TGCGATAAGTGCTCGC-1FC_FI_GFPve_pos      0.0675949601
## FC_Fpos_TGCGGGTCAATTGAAG-1FC_FI_GFPve_pos     -0.0271869081
## FC_Fpos_TGCTCCAAGGTTCCGC-1FC_FI_GFPve_pos     -0.0205651524
## FC_Fpos_TGCTCCACAGTGACCC-1FC_FI_GFPve_pos      0.2324238939
## FC_Fpos_TGCTCGTGTCTCGACG-1FC_FI_GFPve_pos      0.6856875939
## FC_Fpos_TGCTTGCGTAGATTAG-1FC_FI_GFPve_pos     -0.5012034255
## FC_Fpos_TGGCGTGTCGAGTGGA-1FC_FI_GFPve_pos      1.4393366498
## FC_Fpos_TGGGAAGTCCGGCTTT-1FC_FI_GFPve_pos     -1.0931170064
## FC_Fpos_TGGGCGTGTACAGAGC-1FC_FI_GFPve_pos      1.3179852885
## FC_Fpos_TGGTGATCACGCCACA-1FC_FI_GFPve_pos     -0.8170995193
## FC_Fpos_TGGTGATTCACACCCT-1FC_FI_GFPve_pos      2.2069678230
## FC_Fpos_TGGTTAGCAAGCGAGT-1FC_FI_GFPve_pos     -0.8950140553
## FC_Fpos_TGTCAGACAGCAGTTT-1FC_FI_GFPve_pos     -0.5857971984
## FC_Fpos_TGTGCGGTCACATTGG-1FC_FI_GFPve_pos     -0.9839329797
## FC_Fpos_TGTGGCGAGATTGCGG-1FC_FI_GFPve_pos      0.5375479681
## FC_Fpos_TGTGTGACATTACTCT-1FC_FI_GFPve_pos     -0.4320447522
## FC_Fpos_TGTGTGAGTGACGTCC-1FC_FI_GFPve_pos      1.1213041467
## FC_Fpos_TGTTCCGTCCAATCTT-1FC_FI_GFPve_pos     -1.2960524636
## FC_Fpos_TGTTCTACAGTTACCA-1FC_FI_GFPve_pos     -1.2671032029
## FC_Fpos_TGTTCTATCTATACTC-1FC_FI_GFPve_pos     -0.1531758087
## FC_Fpos_TGTTGAGCAAGTGCTT-1FC_FI_GFPve_pos      2.1981027049
## FC_Fpos_TGTTGGACATGACGTT-1FC_FI_GFPve_pos     -0.2725066024
## FC_Fpos_TGTTTGTCACCTCAGG-1FC_FI_GFPve_pos     -0.1894824582
## FC_Fpos_TTACCATTCAATGCAC-1FC_FI_GFPve_pos      1.7387510461
## FC_Fpos_TTACCGCCATTCCTAT-1FC_FI_GFPve_pos      1.4249166173
## FC_Fpos_TTACGCCCATTGCAAC-1FC_FI_GFPve_pos     -0.5295406657
## FC_Fpos_TTACGTTTCCGAGTGC-1FC_FI_GFPve_pos      1.7691129369
## FC_Fpos_TTAGGCAGTCGTCAGC-1FC_FI_GFPve_pos      0.4389584583
## FC_Fpos_TTAGGGTCAAGTCGTT-1FC_FI_GFPve_pos     -0.8585185486
## FC_Fpos_TTAGTCTCAGTCCGTG-1FC_FI_GFPve_pos     -0.8682173568
## FC_Fpos_TTAGTCTGTCAAAGTA-1FC_FI_GFPve_pos      1.6849003953
## FC_Fpos_TTCACGCAGGCCTAGA-1FC_FI_GFPve_pos      0.4315068347
## FC_Fpos_TTCACGCTCCACGAAT-1FC_FI_GFPve_pos      0.5270675046
## FC_Fpos_TTCCAATAGTAGGCCA-1FC_FI_GFPve_pos      0.9602274222
## FC_Fpos_TTCCACGAGCCATTGT-1FC_FI_GFPve_pos     -0.2843916195
## FC_Fpos_TTCCACGAGGCGTTAG-1FC_FI_GFPve_pos      1.0274551434
## FC_Fpos_TTCCACGAGTGCGTCC-1FC_FI_GFPve_pos      1.0823191923
## FC_Fpos_TTCCACGCATCGGAAG-1FC_FI_GFPve_pos     -0.5652597028
## FC_Fpos_TTCCTAAGTTCACGAT-1FC_FI_GFPve_pos      2.8344631118
```

```
## FC_Fpos_TTCCTAAGTTGATGTC-1FC_FI_GFPve_pos      1.5413014812
## FC_Fpos_TTCCTCTTCAAGAATG-1FC_FI_GFPve_pos     -0.4960364776
## FC_Fpos_TTCCTTCAGACCTCCG-1FC_FI_GFPve_pos      0.7523026389
## FC_Fpos_TTCCTTCGTGAGTAGC-1FC_FI_GFPve_pos      1.2472680730
## FC_Fpos_TTCTCTCAGTACTGGG-1FC_FI_GFPve_pos      0.0215727014
## FC_Fpos_TTCTGTAAGCCTTTCC-1FC_FI_GFPve_pos      0.4718740803
## FC_Fpos_TTCTGTAGTCTTACAG-1FC_FI_GFPve_pos      0.7602602166
## FC_Fpos_TTGAGTGCATCCTCAC-1FC_FI_GFPve_pos      0.0764183176
## FC_Fpos_TTGATGGTCCCGTAAA-1FC_FI_GFPve_pos     -0.6613001304
## FC_Fpos_TTGCGTCCAGACGGAT-1FC_FI_GFPve_pos      2.6184461040
## FC_Fpos_TTGGATGAGATAGCAT-1FC_FI_GFPve_pos      1.0616542024
## FC_Fpos_TTGGATGGTCCAGAAG-1FC_FI_GFPve_pos      0.9857724232
## FC_Fpos_TTGGGTAGTCTGCCTT-1FC_FI_GFPve_pos      1.7936043662
## FC_Fpos_TTGGGTAGTTACAGCT-1FC_FI_GFPve_pos      1.1273593826
## FC_Fpos_TTGGTTTTCCACGTAA-1FC_FI_GFPve_pos      0.1185088313
## FC_Fpos_TTGTGTTAGAATCCCT-1FC_FI_GFPve_pos      1.4057335777
## FC_Fpos_TTGTTCAAGGATTTAG-1FC_FI_GFPve_pos      1.5451501531
## FC_Fpos_TTGTTGTAGAATTCAG-1FC_FI_GFPve_pos      2.0709659976
## FC_Fpos_TTGTTTGAGACATATG-1FC_FI_GFPve_pos      1.3113479537
## FC_Fpos_TTTACCACATTCGGGC-1FC_FI_GFPve_pos      1.8622955007
## FC_Fpos_TTTACTGGTGGAGGTT-1FC_FI_GFPve_pos      1.3972130937
## FC_Fpos_TTTATGCCACCGAATT-1FC_FI_GFPve_pos      0.2073612136
## FC_Fpos_TTTCACAAGTCATACC-1FC_FI_GFPve_pos      1.0141304058
## FC_Fpos_TTTCATGCAGGCAATG-1FC_FI_GFPve_pos      1.4572075528
## FC_Fpos_TTTGGAGCAAGAGGTC-1FC_FI_GFPve_pos     -1.5572716075
## FC_Fpos_TTTGGAGCACGCGCAT-1FC_FI_GFPve_pos      1.7860075158
## FC_Fpos_TTTGGTTCAGATCCAT-1FC_FI_GFPve_pos      1.2883347196
## dCA2_Mpos_AAACCCATCTGAGGCC-1dCA2_MI_GFPve_pos  0.8874563498
## dCA2_Mpos_AAACGAAGTCCAAAGG-1dCA2_MI_GFPve_pos -0.0823262560
## dCA2_Mpos_AAAGGATAGCAACTCT-1dCA2_MI_GFPve_pos  0.2670878840
## dCA2_Mpos_AAAGGATCAATGTCTG-1dCA2_MI_GFPve_pos  0.8166522903
## dCA2_Mpos_AAAGGGCCAGCTACTA-1dCA2_MI_GFPve_pos  0.2922207487
## dCA2_Mpos_AAAGGTAAGTTGAATG-1dCA2_MI_GFPve_pos  1.9517882985
## dCA2_Mpos_AAAGGTACAGAGTAAT-1dCA2_MI_GFPve_pos  0.1766596806
## dCA2_Mpos_AAAGGTATCTTGCGCT-1dCA2_MI_GFPve_pos  0.1958949489
## dCA2_Mpos_AAAGTCCTCGAGTCTA-1dCA2_MI_GFPve_pos  1.2208537978
## dCA2_Mpos_AAATGGAGTTAATCGC-1dCA2_MI_GFPve_pos  1.6747295779
## dCA2_Mpos_AACAAAGCACGCGCTA-1dCA2_MI_GFPve_pos  1.8034098071
## dCA2_Mpos_AACAAAGTCACTTGGA-1dCA2_MI_GFPve_pos  0.9936539454
## dCA2_Mpos_AACAAGACATATCTGG-1dCA2_MI_GFPve_pos  1.0539677066
## dCA2_Mpos_AACAGGGCAGCGATTT-1dCA2_MI_GFPve_pos -0.3524695712
## dCA2_Mpos_AACAGGGGTTTCCAAG-1dCA2_MI_GFPve_pos -0.5650903064
## dCA2_Mpos_AACCAACCACGACGCT-1dCA2_MI_GFPve_pos -0.7882428246
## dCA2_Mpos_AACCACACACATCCCT-1dCA2_MI_GFPve_pos  0.0066974533
## dCA2_Mpos_AACCTGACACCCTTGT-1dCA2_MI_GFPve_pos  1.3877392930
## dCA2_Mpos_AACCTGACACTTGTGA-1dCA2_MI_GFPve_pos -0.0303452986
## dCA2_Mpos_AACGAAACATCCGTGG-1dCA2_MI_GFPve_pos  2.0964941901
## dCA2_Mpos_AACGGGAAGTAGTCCT-1dCA2_MI_GFPve_pos  1.5441303176
## dCA2_Mpos_AACGTCAGTCCACTTC-1dCA2_MI_GFPve_pos  1.3050699157
## dCA2_Mpos_AACGTCAGTCCTGGGT-1dCA2_MI_GFPve_pos -0.9937795478
## dCA2_Mpos_AAGAACAGTCCAATCA-1dCA2_MI_GFPve_pos -1.5372413474
## dCA2_Mpos_AAGACAAAGACTGGGT-1dCA2_MI_GFPve_pos  0.1465742064
## dCA2_Mpos_AAGACAACAGTTTCAG-1dCA2_MI_GFPve_pos -0.6461761075
## dCA2_Mpos_AAGCGAGAGCGCCATC-1dCA2_MI_GFPve_pos  0.9628749413
## dCA2_Mpos_AAGCGAGCAGCAGACA-1dCA2_MI_GFPve_pos  0.0940974306
## dCA2_Mpos_AAGGAATAGTTAGTGA-1dCA2_MI_GFPve_pos  0.5904444677
## dCA2_Mpos_AAGTACCAGACTACCT-1dCA2_MI_GFPve_pos  0.5752351505
## dCA2_Mpos_AAGTACCCACACTTAG-1dCA2_MI_GFPve_pos -0.1229379552
## dCA2_Mpos_AAGTACCTCTCGCGTT-1dCA2_MI_GFPve_pos  0.1892074418
## dCA2_Mpos_AAGTCGTGTAACTTCG-1dCA2_MI_GFPve_pos  1.6857165021
## dCA2_Mpos_AAGTTCGGTCGTCAGC-1dCA2_MI_GFPve_pos  0.5887997967
## dCA2_Mpos_AATCACGAGTTGCGCC-1dCA2_MI_GFPve_pos  1.2147114200
## dCA2_Mpos_AATGACCTCATCGTAG-1dCA2_MI_GFPve_pos  1.2611725253
## dCA2_Mpos_AATGCCAAGTGGACGT-1dCA2_MI_GFPve_pos  0.9153659267
## dCA2_Mpos_AATGGCTAGTGCTCAT-1dCA2_MI_GFPve_pos  1.3394370002
## dCA2_Mpos_AATTTCCCAATTCTCT-1dCA2_MI_GFPve_pos  0.2358718884
## dCA2_Mpos_ACAACCAAGCGCAATG-1dCA2_MI_GFPve_pos  1.8837175292
## dCA2_Mpos_ACAACCAAGTGTCATC-1dCA2_MI_GFPve_pos  1.4158097429
## dCA2_Mpos_ACAACCAGTGACAGCA-1dCA2_MI_GFPve_pos  1.1327749414
## dCA2_Mpos_ACAAGCTAGTCACACT-1dCA2_MI_GFPve_pos -0.1886797326
```

```
## dCA2_Mpos_ACAAGCTCAGCATACT-1dCA2_MI_GFPve_pos -0.3238841968
## dCA2_Mpos_ACAAGCTTCTACGGGC-1dCA2_MI_GFPve_pos  1.0742371005
## dCA2_Mpos_ACACAGTGTCCACTCT-1dCA2_MI_GFPve_pos  2.4838981551
## dCA2_Mpos_ACACGCGAGCACACAG-1dCA2_MI_GFPve_pos -0.9683244305
## dCA2_Mpos_ACACGCGAGTATTGCC-1dCA2_MI_GFPve_pos -0.1238901841
## dCA2_Mpos_ACAGAAATCATTGCGA-1dCA2_MI_GFPve_pos  0.6287903113
## dCA2_Mpos_ACATCCCTCCGGACGT-1dCA2_MI_GFPve_pos  1.5769685430
## dCA2_Mpos_ACATGCAGTCCATACA-1dCA2_MI_GFPve_pos  1.9674584789
## dCA2_Mpos_ACCAAACGTTGCCGCA-1dCA2_MI_GFPve_pos -0.3699795085
## dCA2_Mpos_ACCAACACACCGTCTT-1dCA2_MI_GFPve_pos  0.7654777688
## dCA2_Mpos_ACCAACATCGGAAACG-1dCA2_MI_GFPve_pos  1.5935640258
## dCA2_Mpos_ACCACAAAGAGATCGC-1dCA2_MI_GFPve_pos  2.0007196588
## dCA2_Mpos_ACCCTCAAGTTGTCAC-1dCA2_MI_GFPve_pos -1.5227727967
## dCA2_Mpos_ACCCTCACAAGCTGCC-1dCA2_MI_GFPve_pos -0.3306550163
## dCA2_Mpos_ACCTACCGTATCGATC-1dCA2_MI_GFPve_pos -0.0722590166
## dCA2_Mpos_ACCTGAAAGATCCGAG-1dCA2_MI_GFPve_pos  1.3549147767
## dCA2_Mpos_ACCTGAACACAACCGC-1dCA2_MI_GFPve_pos -0.5926079231
## dCA2_Mpos_ACGATCAAGATGTTGA-1dCA2_MI_GFPve_pos  0.3094966037
## dCA2_Mpos_ACGATCATCCCATGGG-1dCA2_MI_GFPve_pos  0.6643538398
## dCA2_Mpos_ACGATGTAGCAGAAAG-1dCA2_MI_GFPve_pos  1.7294006271
## dCA2_Mpos_ACGATGTCAGGATTCT-1dCA2_MI_GFPve_pos  2.2988486213
## dCA2_Mpos_ACGCACGGTGAGTTTC-1dCA2_MI_GFPve_pos  1.5584213657
## dCA2_Mpos_ACGGGTCGTACGACTT-1dCA2_MI_GFPve_pos  1.7308801336
## dCA2_Mpos_ACGGTTAAGCCATCCG-1dCA2_MI_GFPve_pos -1.4249721843
## dCA2_Mpos_ACGGTTAAGTGTCATC-1dCA2_MI_GFPve_pos  1.1415760440
## dCA2_Mpos_ACGGTTACATAGAATG-1dCA2_MI_GFPve_pos  1.4461262149
## dCA2_Mpos_ACGTAACCACCTGTCT-1dCA2_MI_GFPve_pos  0.6373646778
## dCA2_Mpos_ACGTACATCATTGTTC-1dCA2_MI_GFPve_pos  1.3211084527
## dCA2_Mpos_ACGTTCCAGCGTATGG-1dCA2_MI_GFPve_pos  2.1566960735
## dCA2_Mpos_ACTATCTCAGAAATCA-1dCA2_MI_GFPve_pos -0.5714796263
## dCA2_Mpos_ACTATGGCATGCGTGC-1dCA2_MI_GFPve_pos  0.7844800872
## dCA2_Mpos_ACTATGGTCCAAGCTA-1dCA2_MI_GFPve_pos -0.1199289488
## dCA2_Mpos_ACTATTCAGAAACCCG-1dCA2_MI_GFPve_pos  1.2589445037
## dCA2_Mpos_ACTCCCAAGATCCGAG-1dCA2_MI_GFPve_pos  0.7122470659
## dCA2_Mpos_ACTCCCAAGGGTGGGA-1dCA2_MI_GFPve_pos  1.3914738816
## dCA2_Mpos_ACTGCAAGTACTGCGC-1dCA2_MI_GFPve_pos  1.3401145858
## dCA2_Mpos_ACTGTCCCAATTGAGA-1dCA2_MI_GFPve_pos  2.5251164359
## dCA2_Mpos_ACTGTGACAACCAATC-1dCA2_MI_GFPve_pos -0.5394381362
## dCA2_Mpos_ACTTATCTCCCATGGG-1dCA2_MI_GFPve_pos  1.1907000465
## dCA2_Mpos_ACTTTCACACCGTACG-1dCA2_MI_GFPve_pos  1.6075245065
## dCA2_Mpos_ACTTTCAGTAGGCTCC-1dCA2_MI_GFPve_pos  2.1548619193
## dCA2_Mpos_AGAAATGTCAGCGCAC-1dCA2_MI_GFPve_pos -0.4104450422
## dCA2_Mpos_AGAAGCGCAACGATCT-1dCA2_MI_GFPve_pos  2.2184820098
## dCA2_Mpos_AGACAAAAGCCATATC-1dCA2_MI_GFPve_pos  1.4203339738
## dCA2_Mpos_AGACAAAAGCGCCGTT-1dCA2_MI_GFPve_pos  1.7713124675
## dCA2_Mpos_AGACAAACAGCTGTTA-1dCA2_MI_GFPve_pos  0.5783565861
## dCA2_Mpos_AGACACTAGACGACTG-1dCA2_MI_GFPve_pos  2.2943570537
## dCA2_Mpos_AGACACTAGGGTATAT-1dCA2_MI_GFPve_pos  0.8018401188
## dCA2_Mpos_AGACAGGAGGTTTACC-1dCA2_MI_GFPve_pos -0.5022859054
## dCA2_Mpos_AGACCCGCATATCTCT-1dCA2_MI_GFPve_pos  0.4464962405
## dCA2_Mpos_AGACTCATCGGAGCAA-1dCA2_MI_GFPve_pos  0.9443452877
## dCA2_Mpos_AGAGAGCGTGATCGTT-1dCA2_MI_GFPve_pos  1.0761008782
## dCA2_Mpos_AGAGCAGCAAATCGGG-1dCA2_MI_GFPve_pos  2.1754691524
## dCA2_Mpos_AGAGCAGCACTCGATA-1dCA2_MI_GFPve_pos  1.8363925142
## dCA2_Mpos_AGAGCCCAGACCAAAT-1dCA2_MI_GFPve_pos  1.8820955676
## dCA2_Mpos_AGAGCCCTCTCACTCG-1dCA2_MI_GFPve_pos -0.1218822527
## dCA2_Mpos_AGATCCAAGGGTCACA-1dCA2_MI_GFPve_pos  0.0273429600
## dCA2_Mpos_AGATCGTCAACAGTGG-1dCA2_MI_GFPve_pos -0.5985564667
## dCA2_Mpos_AGATGAACAGCTGTAT-1dCA2_MI_GFPve_pos  1.7437469882
## dCA2_Mpos_AGATGCTCATCATTTC-1dCA2_MI_GFPve_pos  0.5780776066
## dCA2_Mpos_AGATGCTGTTCTTGTT-1dCA2_MI_GFPve_pos  0.3707669300
## dCA2_Mpos_AGCATCATCACGGACC-1dCA2_MI_GFPve_pos  2.1810302657
## dCA2_Mpos_AGCCAGCCATACCAGT-1dCA2_MI_GFPve_pos  0.0864282918
## dCA2_Mpos_AGCGATTGTGAGTAAT-1dCA2_MI_GFPve_pos  1.7577420396
## dCA2_Mpos_AGCGATTTCCGAGTGC-1dCA2_MI_GFPve_pos  1.3128843230
## dCA2_Mpos_AGCGCCATCACCATAG-1dCA2_MI_GFPve_pos  0.4505435449
## dCA2_Mpos_AGCGCTGAGTGAGTTA-1dCA2_MI_GFPve_pos  1.6454061193
## dCA2_Mpos_AGCGTATAGCCGTCGT-1dCA2_MI_GFPve_pos  1.9202006978
## dCA2_Mpos_AGCTACAGTCATCACA-1dCA2_MI_GFPve_pos  1.3340101165
## dCA2_Mpos_AGCTACAGTCGCAACC-1dCA2_MI_GFPve_pos -0.3463617044
```

```
## dCA2_Mpos_AGCTTCCCAAGCCTGC-1dCA2_MI_GFPve_pos -0.2355245727
## dCA2_Mpos_AGCTTCCCAGTAGGAC-1dCA2_MI_GFPve_pos  1.1963245792
## dCA2_Mpos_AGCTTCCGTGATTGGG-1dCA2_MI_GFPve_pos  1.3915164394
## dCA2_Mpos_AGGAGGTAGATGGTAT-1dCA2_MI_GFPve_pos  1.1387872619
## dCA2_Mpos_AGGATAAAGAGTTGAT-1dCA2_MI_GFPve_pos  0.2104480860
## dCA2_Mpos_AGGCTGCGTCACCCTT-1dCA2_MI_GFPve_pos  0.5055525524
## dCA2_Mpos_AGGCTGCGTTACGATC-1dCA2_MI_GFPve_pos -0.2345927137
## dCA2_Mpos_AGGGAGTGTTTAGAGA-1dCA2_MI_GFPve_pos -0.5714680033
## dCA2_Mpos_AGGGCCTAGATGCAGC-1dCA2_MI_GFPve_pos  0.7016434592
## dCA2_Mpos_AGGGTCCAGCATCAAA-1dCA2_MI_GFPve_pos  1.2709609193
## dCA2_Mpos_AGGGTCCAGGTAGCAC-1dCA2_MI_GFPve_pos  0.3459559155
## dCA2_Mpos_AGGGTGACATGGAACG-1dCA2_MI_GFPve_pos  2.1070070190
## dCA2_Mpos_AGGTCATCACTGGAAG-1dCA2_MI_GFPve_pos -0.5859448987
## dCA2_Mpos_AGGTCTAGTCGATTTG-1dCA2_MI_GFPve_pos  0.7287671489
## dCA2_Mpos_AGGTGTTTCCTCTGCA-1dCA2_MI_GFPve_pos  2.0550173444
## dCA2_Mpos_AGGTTACAGCGATTCT-1dCA2_MI_GFPve_pos  0.9093868655
## dCA2_Mpos_AGGTTGTCAAATTGCC-1dCA2_MI_GFPve_pos  0.1791532931
## dCA2_Mpos_AGGTTGTGTACGCTTA-1dCA2_MI_GFPve_pos  1.2488179130
## dCA2_Mpos_AGTAACCTCGCCTAGG-1dCA2_MI_GFPve_pos  2.2079460544
## dCA2_Mpos_AGTACCAGTGGTTTGT-1dCA2_MI_GFPve_pos -0.3730671185
## dCA2_Mpos_AGTACTGGTGTTTACG-1dCA2_MI_GFPve_pos  1.1425574941
## dCA2_Mpos_AGTCAACTCTCATAGG-1dCA2_MI_GFPve_pos  1.1167836112
## dCA2_Mpos_AGTCATGAGACGCCAA-1dCA2_MI_GFPve_pos  2.0731060428
## dCA2_Mpos_AGTCATGAGGGAGGTG-1dCA2_MI_GFPve_pos  2.1476225776
## dCA2_Mpos_AGTCATGGTGTGTCGC-1dCA2_MI_GFPve_pos  1.4093607587
## dCA2_Mpos_AGTCATGTCTAAGCCA-1dCA2_MI_GFPve_pos  2.2448654098
## dCA2_Mpos_AGTCTCCCACATGGTT-1dCA2_MI_GFPve_pos -0.6046475249
## dCA2_Mpos_AGTCTCCTCCGACATA-1dCA2_MI_GFPve_pos  0.2316320759
## dCA2_Mpos_AGTGACTGTGGATGAC-1dCA2_MI_GFPve_pos -0.4714701253
## dCA2_Mpos_AGTGACTGTTTCCATT-1dCA2_MI_GFPve_pos  0.0984281043
## dCA2_Mpos_AGTGATCAGCGGCTCT-1dCA2_MI_GFPve_pos  0.8924049658
## dCA2_Mpos_AGTGATCCACGGCCAT-1dCA2_MI_GFPve_pos  0.8720794839
## dCA2_Mpos_AGTTCCCGTAGCCCTG-1dCA2_MI_GFPve_pos  2.1583070678
## dCA2_Mpos_ATACCGAGTTCTTCAT-1dCA2_MI_GFPve_pos  1.6855280322
## dCA2_Mpos_ATACTTCCAAGTGGTG-1dCA2_MI_GFPve_pos  1.5498032493
## dCA2_Mpos_ATAGACCCATAAGCGG-1dCA2_MI_GFPve_pos -1.5713527280
## dCA2_Mpos_ATAGACCCATCGTTCC-1dCA2_MI_GFPve_pos  1.1303435487
## dCA2_Mpos_ATAGAGATCTCTGCTG-1dCA2_MI_GFPve_pos  1.0542623324
## dCA2_Mpos_ATAGGCTGTGACATCT-1dCA2_MI_GFPve_pos  0.0897697189
## dCA2_Mpos_ATAGGCTTCCTTATCA-1dCA2_MI_GFPve_pos  0.1460490112
## dCA2_Mpos_ATCACAGAGATGGGCT-1dCA2_MI_GFPve_pos -0.1626442152
## dCA2_Mpos_ATCACAGTCCCATTTA-1dCA2_MI_GFPve_pos  2.1667168063
## dCA2_Mpos_ATCACAGTCGCTGATA-1dCA2_MI_GFPve_pos -0.9834020215
## dCA2_Mpos_ATCACGAGTGTTCGTA-1dCA2_MI_GFPve_pos  1.9636881274
## dCA2_Mpos_ATCATTCGTCGTGATT-1dCA2_MI_GFPve_pos  1.4201931876
## dCA2_Mpos_ATCCACCAGAACAAGG-1dCA2_MI_GFPve_pos  0.6246988696
## dCA2_Mpos_ATCCACCCATCGAGCC-1dCA2_MI_GFPve_pos -0.8914393860
## dCA2_Mpos_ATCCACCGTACTGTTG-1dCA2_MI_GFPve_pos -0.5150641876
## dCA2_Mpos_ATCCCTGAGCAAGCCA-1dCA2_MI_GFPve_pos  2.0319150609
## dCA2_Mpos_ATCCCTGCAATCAAGA-1dCA2_MI_GFPve_pos  1.5228198690
## dCA2_Mpos_ATCCCTGCAGAGAAAG-1dCA2_MI_GFPve_pos  0.7219542188
## dCA2_Mpos_ATCCTATCATGGAATA-1dCA2_MI_GFPve_pos  1.9216487331
## dCA2_Mpos_ATCCTATTCCGACAGC-1dCA2_MI_GFPve_pos -0.0430517870
## dCA2_Mpos_ATCGATGTCCAACACA-1dCA2_MI_GFPve_pos -0.1755429583
## dCA2_Mpos_ATCGATGTCGCACTCT-1dCA2_MI_GFPve_pos  1.7772884292
## dCA2_Mpos_ATCGATGTCTAGACAC-1dCA2_MI_GFPve_pos  0.1340338548
## dCA2_Mpos_ATCGCCTCAGATGCGA-1dCA2_MI_GFPve_pos  0.9279001874
## dCA2_Mpos_ATCGGATTCGGCTGGT-1dCA2_MI_GFPve_pos  0.4736566168
## dCA2_Mpos_ATCGTAGCAGAGCCCT-1dCA2_MI_GFPve_pos  1.1265888137
## dCA2_Mpos_ATCGTAGGTTCGTACA-1dCA2_MI_GFPve_pos -0.7790404635
## dCA2_Mpos_ATCGTCCGTTGTATGC-1dCA2_MI_GFPve_pos  2.3827850742
## dCA2_Mpos_ATCGTGACACTTGAAC-1dCA2_MI_GFPve_pos  1.4982157869
## dCA2_Mpos_ATCTCTACATACTGTG-1dCA2_MI_GFPve_pos  1.5339544934
## dCA2_Mpos_ATCTTCAAGTCCCGAC-1dCA2_MI_GFPve_pos  0.9598204416
## dCA2_Mpos_ATCTTCACAGTTCTAG-1dCA2_MI_GFPve_pos  0.2947443200
## dCA2_Mpos_ATGAAAGTCCATAAGC-1dCA2_MI_GFPve_pos  1.2445640487
## dCA2_Mpos_ATGACCAAGTACTGGG-1dCA2_MI_GFPve_pos  1.8779655618
## dCA2_Mpos_ATGACCATCCCGAAAT-1dCA2_MI_GFPve_pos  1.6058750076
## dCA2_Mpos_ATGAGTCAGCGCTTCG-1dCA2_MI_GFPve_pos -1.1215533095
## dCA2_Mpos_ATGATCGGTCCTACAA-1dCA2_MI_GFPve_pos  2.2148444575
```

```
## dCA2_Mpos_ATGCATGCACAATTCG-1dCA2_MI_GFPve_pos  1.7488764447
## dCA2_Mpos_ATGCATGGTCTTACTT-1dCA2_MI_GFPve_pos  0.9711196346
## dCA2_Mpos_ATGCCTCGTCGCAACC-1dCA2_MI_GFPve_pos  1.6598128003
## dCA2_Mpos_ATGCGATAGGGTTGCA-1dCA2_MI_GFPve_pos  1.7529700917
## dCA2_Mpos_ATGCGATCAAGTGCAG-1dCA2_MI_GFPve_pos -0.5797236639
## dCA2_Mpos_ATGCGATGTGGAGAAA-1dCA2_MI_GFPve_pos  1.4508527440
## dCA2_Mpos_ATGGAGGCAAGACCTT-1dCA2_MI_GFPve_pos  1.7593310994
## dCA2_Mpos_ATGGGTTCATCCGCGA-1dCA2_MI_GFPve_pos  1.2784508390
## dCA2_Mpos_ATGTCCCCATAGAAAC-1dCA2_MI_GFPve_pos -0.1840963143
## dCA2_Mpos_ATGTCCCGTCACCGCA-1dCA2_MI_GFPve_pos -0.0479774850
## dCA2_Mpos_ATGTCTTAGCAACCAG-1dCA2_MI_GFPve_pos -0.5211589413
## dCA2_Mpos_ATTACTCCAGTAGAAT-1dCA2_MI_GFPve_pos  0.8155500812
## dCA2_Mpos_ATTATCCGTTTACCAG-1dCA2_MI_GFPve_pos -0.3872303742
## dCA2_Mpos_ATTATCCTCCACTTTA-1dCA2_MI_GFPve_pos  0.0191425530
## dCA2_Mpos_ATTCACTCACCTGCAG-1dCA2_MI_GFPve_pos  0.6182506007
## dCA2_Mpos_ATTCATCTCCGCGATG-1dCA2_MI_GFPve_pos  1.5004471463
## dCA2_Mpos_ATTCCTACATTCTCCG-1dCA2_MI_GFPve_pos  1.7803174180
## dCA2_Mpos_ATTCCTAGTTAGCTAC-1dCA2_MI_GFPve_pos  2.2387740535
## dCA2_Mpos_ATTCGTTAGCGTGCCT-1dCA2_MI_GFPve_pos  1.2534891290
## dCA2_Mpos_ATTCGTTGTATGACAA-1dCA2_MI_GFPve_pos -0.2370184737
## dCA2_Mpos_ATTCGTTTCTCGACGG-1dCA2_MI_GFPve_pos -0.7159885841
## dCA2_Mpos_ATTCTACCAAATGATG-1dCA2_MI_GFPve_pos -0.6175134259
## dCA2_Mpos_ATTGGGTCAACCGCCA-1dCA2_MI_GFPve_pos -0.8038949447
## dCA2_Mpos_ATTTACCTCAAACGAA-1dCA2_MI_GFPve_pos -0.4129228669
## dCA2_Mpos_ATTTCACCAGTGAGCA-1dCA2_MI_GFPve_pos  1.5052715463
## dCA2_Mpos_ATTTCTGGTGTTCAGT-1dCA2_MI_GFPve_pos  0.1962715608
## dCA2_Mpos_ATTTCTGTCTTGTGCC-1dCA2_MI_GFPve_pos  2.7814259452
## dCA2_Mpos_CAACAACCACAATGTC-1dCA2_MI_GFPve_pos  0.1324057297
## dCA2_Mpos_CAACCAACACTGCGTG-1dCA2_MI_GFPve_pos  1.9758528394
## dCA2_Mpos_CAACCTCCATTACGGT-1dCA2_MI_GFPve_pos  0.3872093183
## dCA2_Mpos_CAACGGCGTGGATCGA-1dCA2_MI_GFPve_pos  1.2896159810
## dCA2_Mpos_CAACGGCGTGTGAGCA-1dCA2_MI_GFPve_pos -0.8674052435
## dCA2_Mpos_CAAGCTAGTGCACATT-1dCA2_MI_GFPve_pos  2.0514923257
## dCA2_Mpos_CAATCGAAGTATGCAA-1dCA2_MI_GFPve_pos  0.0387741772
## dCA2_Mpos_CAATCGATCACGTAGT-1dCA2_MI_GFPve_pos  0.9751347226
## dCA2_Mpos_CAATGACAGGTCCTGC-1dCA2_MI_GFPve_pos  1.9685850066
## dCA2_Mpos_CAATTTCAGACTTCAC-1dCA2_MI_GFPve_pos  0.9268606228
## dCA2_Mpos_CAATTTCAGCATCTTG-1dCA2_MI_GFPve_pos  1.1860315723
## dCA2_Mpos_CAATTTCGTCTTTATC-1dCA2_MI_GFPve_pos  2.5244195384
## dCA2_Mpos_CACACAAAGTGGCGAT-1dCA2_MI_GFPve_pos  1.3737918061
## dCA2_Mpos_CACACAAAGTTTCGAC-1dCA2_MI_GFPve_pos  1.5564951820
## dCA2_Mpos_CACAGATCAAGTAGTA-1dCA2_MI_GFPve_pos  1.4958815498
## dCA2_Mpos_CACCAAACAAGTTCCA-1dCA2_MI_GFPve_pos -0.3514359074
## dCA2_Mpos_CACGAATCAAGAGTAT-1dCA2_MI_GFPve_pos  1.5707223338
## dCA2_Mpos_CACGAATCAGTTGGTT-1dCA2_MI_GFPve_pos -0.9085687476
## dCA2_Mpos_CACGAATGTGCTGATT-1dCA2_MI_GFPve_pos -0.5463144260
## dCA2_Mpos_CACGGGTTCTTTGCAT-1dCA2_MI_GFPve_pos  2.3047315997
## dCA2_Mpos_CACGTGGCACCGTGAC-1dCA2_MI_GFPve_pos  0.3041985405
## dCA2_Mpos_CACGTGGGTTCTCACC-1dCA2_MI_GFPve_pos  1.8540186805
## dCA2_Mpos_CACGTGGTCCATTGGA-1dCA2_MI_GFPve_pos  2.1223206443
## dCA2_Mpos_CACGTGGTCGATTTCT-1dCA2_MI_GFPve_pos  1.5170582456
## dCA2_Mpos_CACGTTCTCGCTTGAA-1dCA2_MI_GFPve_pos  2.4892125053
## dCA2_Mpos_CACTAAGTCAGCTAGT-1dCA2_MI_GFPve_pos -0.1738210398
## dCA2_Mpos_CACTAAGTCTCACGAA-1dCA2_MI_GFPve_pos  0.6051000280
## dCA2_Mpos_CACTGGGCAAGAGAGA-1dCA2_MI_GFPve_pos -0.4070762473
## dCA2_Mpos_CACTGTCGTATCAGGG-1dCA2_MI_GFPve_pos  0.2999314142
## dCA2_Mpos_CACTGTCGTCATAGTC-1dCA2_MI_GFPve_pos  0.3652864618
## dCA2_Mpos_CACTTCGAGACCTCCG-1dCA2_MI_GFPve_pos  0.2641637844
## dCA2_Mpos_CAGATCAGTATGAAAC-1dCA2_MI_GFPve_pos  0.7034594936
## dCA2_Mpos_CAGCACGAGTCTGCAT-1dCA2_MI_GFPve_pos  1.3939764423
## dCA2_Mpos_CAGCACGTCATTGGTG-1dCA2_MI_GFPve_pos  1.6353766841
## dCA2_Mpos_CAGCAGCCAAGCTGCC-1dCA2_MI_GFPve_pos -0.3017718273
## dCA2_Mpos_CAGCGTGAGTGAGGCT-1dCA2_MI_GFPve_pos  1.6247025651
## dCA2_Mpos_CAGCGTGCACATGTTG-1dCA2_MI_GFPve_pos  2.1999404353
## dCA2_Mpos_CAGGCCACATGGTACT-1dCA2_MI_GFPve_pos  0.8265269918
## dCA2_Mpos_CAGGGCTCACCCAACG-1dCA2_MI_GFPve_pos  0.1718053517
## dCA2_Mpos_CAGGGCTGTCACGTGC-1dCA2_MI_GFPve_pos  0.2363275510
## dCA2_Mpos_CAGGTATTCATCCCGT-1dCA2_MI_GFPve_pos -0.0074457678
## dCA2_Mpos_CAGGTATTCTCTATGT-1dCA2_MI_GFPve_pos  0.6380622310
## dCA2_Mpos_CAGTGCGAGTGATAGT-1dCA2_MI_GFPve_pos -0.8749156075
```

```
## dCA2_Mpos_CAGTGCGCAGAACGCA-1dCA2_MI_GFPve_pos  1.8818823022
## dCA2_Mpos_CAGTTAGAGAAGCGGG-1dCA2_MI_GFPve_pos  1.0118427200
## dCA2_Mpos_CAGTTAGCAACACAGG-1dCA2_MI_GFPve_pos -0.5798199254
## dCA2_Mpos_CAGTTAGGTAATACCC-1dCA2_MI_GFPve_pos  1.3877506179
## dCA2_Mpos_CAGTTCCAGAGAGTTT-1dCA2_MI_GFPve_pos  1.1843792123
## dCA2_Mpos_CAGTTCCTCATCCCGT-1dCA2_MI_GFPve_pos -0.6888990479
## dCA2_Mpos_CATACCCCATACAGAA-1dCA2_MI_GFPve_pos -0.3765404301
## dCA2_Mpos_CATACTTGTAGAGACC-1dCA2_MI_GFPve_pos  1.3532357139
## dCA2_Mpos_CATCCCAAGTGGTTCT-1dCA2_MI_GFPve_pos -0.2870055514
## dCA2_Mpos_CATCCCAGTCACCTTC-1dCA2_MI_GFPve_pos  0.3595969600
## dCA2_Mpos_CATCCCATCGTTAGAC-1dCA2_MI_GFPve_pos  2.0255824250
## dCA2_Mpos_CATCGGGGTGAAAGTT-1dCA2_MI_GFPve_pos  0.5087905747
## dCA2_Mpos_CATCGTCCACTCCTGT-1dCA2_MI_GFPve_pos  0.8114849848
## dCA2_Mpos_CATGCCTAGTATGTAG-1dCA2_MI_GFPve_pos  1.7504546565
## dCA2_Mpos_CATGCGGAGGGACTGT-1dCA2_MI_GFPve_pos  2.1466708106
## dCA2_Mpos_CATGCGGAGGTGCCAA-1dCA2_MI_GFPve_pos  0.7971171064
## dCA2_Mpos_CATGCGGCATAGGTAA-1dCA2_MI_GFPve_pos  1.4765758437
## dCA2_Mpos_CATGCTCCATCCGAAT-1dCA2_MI_GFPve_pos  0.6251510305
## dCA2_Mpos_CATGGTAGTAATTAGG-1dCA2_MI_GFPve_pos  0.6754350585
## dCA2_Mpos_CATGGTAGTCAAATCC-1dCA2_MI_GFPve_pos  1.7940099162
## dCA2_Mpos_CATTCCGAGGGTGAGG-1dCA2_MI_GFPve_pos  1.4591158551
## dCA2_Mpos_CATTGAGGTGATTCTG-1dCA2_MI_GFPve_pos  2.3971526546
## dCA2_Mpos_CATTGCCGTTAGCTAC-1dCA2_MI_GFPve_pos  1.8151509685
## dCA2_Mpos_CATTGTTGTTTGGGTT-1dCA2_MI_GFPve_pos  1.2688404245
## dCA2_Mpos_CATTTCATCTGTTCAT-1dCA2_MI_GFPve_pos -1.7399220544
## dCA2_Mpos_CCAATGACAGCTACCG-1dCA2_MI_GFPve_pos  0.9974509401
## dCA2_Mpos_CCACACTAGAGAGTGA-1dCA2_MI_GFPve_pos  0.6958630604
## dCA2_Mpos_CCACACTCAATGGCCC-1dCA2_MI_GFPve_pos -0.2079912501
## dCA2_Mpos_CCACCATTCGTAGTCA-1dCA2_MI_GFPve_pos -1.3944747525
## dCA2_Mpos_CCACGAGAGCTAAACA-1dCA2_MI_GFPve_pos  0.9480577630
## dCA2_Mpos_CCACGAGAGTATGCAA-1dCA2_MI_GFPve_pos -1.4629913646
## dCA2_Mpos_CCACGAGCATATGAAG-1dCA2_MI_GFPve_pos -1.3689096051
## dCA2_Mpos_CCACGAGGTATGGGAC-1dCA2_MI_GFPve_pos  0.0005751101
## dCA2_Mpos_CCACGTTCACCCTAGG-1dCA2_MI_GFPve_pos  1.4065030736
## dCA2_Mpos_CCACGTTCACGACTAT-1dCA2_MI_GFPve_pos  1.4066046399
## dCA2_Mpos_CCACGTTTCTTTCAGT-1dCA2_MI_GFPve_pos  1.2697607202
## dCA2_Mpos_CCATAAGGTCGACTGC-1dCA2_MI_GFPve_pos  1.0286174936
## dCA2_Mpos_CCCAACTAGGCTTAGG-1dCA2_MI_GFPve_pos  1.3584741277
## dCA2_Mpos_CCCATTGAGCTAATGA-1dCA2_MI_GFPve_pos  2.1265425605
## dCA2_Mpos_CCCATTGGTTCTCTCG-1dCA2_MI_GFPve_pos  2.2163193149
## dCA2_Mpos_CCCGAAGAGACCCGCT-1dCA2_MI_GFPve_pos -0.1395375061
## dCA2_Mpos_CCCGAAGAGTGTAGAT-1dCA2_MI_GFPve_pos  0.9636413974
## dCA2_Mpos_CCCGAAGGTGAAAGTT-1dCA2_MI_GFPve_pos  0.8467308802
## dCA2_Mpos_CCCGGAAGTGCACAAG-1dCA2_MI_GFPve_pos  0.7131499571
## dCA2_Mpos_CCCTGATCAATCTGCA-1dCA2_MI_GFPve_pos -0.4472611028
## dCA2_Mpos_CCCTGATTCTGTGTGA-1dCA2_MI_GFPve_pos  1.3454089088
## dCA2_Mpos_CCCTTAGAGACCATTC-1dCA2_MI_GFPve_pos -1.3185061293
## dCA2_Mpos_CCCTTAGAGCGACATG-1dCA2_MI_GFPve_pos  1.5401368064
## dCA2_Mpos_CCCTTAGAGGATTCCT-1dCA2_MI_GFPve_pos -0.3011090833
## dCA2_Mpos_CCCTTAGCACACACGC-1dCA2_MI_GFPve_pos  2.0282692832
## dCA2_Mpos_CCGATGGCACTGGCGT-1dCA2_MI_GFPve_pos  1.5561588926
## dCA2_Mpos_CCGCAAGCAAATCAGA-1dCA2_MI_GFPve_pos -0.2381124275
## dCA2_Mpos_CCGGACAGTGTAGGAC-1dCA2_MI_GFPve_pos  2.0463309211
## dCA2_Mpos_CCGGTAGAGTTACGGG-1dCA2_MI_GFPve_pos  1.0014898700
## dCA2_Mpos_CCGGTGATCCGGCAAC-1dCA2_MI_GFPve_pos  0.1262586766
## dCA2_Mpos_CCGTAGGTCTGGAGAG-1dCA2_MI_GFPve_pos  0.7326787037
## dCA2_Mpos_CCGTGAGCACGCGTGT-1dCA2_MI_GFPve_pos  2.1180410308
## dCA2_Mpos_CCGTGAGCATCGGCCA-1dCA2_MI_GFPve_pos  0.3585637433
## dCA2_Mpos_CCGTTCAGTGTAACGG-1dCA2_MI_GFPve_pos  2.1954798621
## dCA2_Mpos_CCTAACCCAACGTAAA-1dCA2_MI_GFPve_pos  0.6601285857
## dCA2_Mpos_CCTATCGTCTCTATGT-1dCA2_MI_GFPve_pos  0.3338756484
## dCA2_Mpos_CCTCATGCAACGATTC-1dCA2_MI_GFPve_pos -0.5376807528
## dCA2_Mpos_CCTCATGTCTCGAGTA-1dCA2_MI_GFPve_pos  0.8054442925
## dCA2_Mpos_CCTCTCCCAAGTGGAC-1dCA2_MI_GFPve_pos  1.1258963269
## dCA2_Mpos_CCTCTCCGTATGTCCA-1dCA2_MI_GFPve_pos  1.7779825849
## dCA2_Mpos_CCTGCATAGCATCAGG-1dCA2_MI_GFPve_pos -0.6292246657
## dCA2_Mpos_CCTGCATGTGTCCAAT-1dCA2_MI_GFPve_pos -1.3487861234
## dCA2_Mpos_CCTTCAGCAAGTATAG-1dCA2_MI_GFPve_pos -1.0528152066
## dCA2_Mpos_CCTTCAGCAAGTGCAG-1dCA2_MI_GFPve_pos  0.8476880712
## dCA2_Mpos_CCTTCAGCATGGCTGC-1dCA2_MI_GFPve_pos  0.4935006124
```

```
## dCA2_Mpos_CCTTGTGCAGACAAGC-1dCA2_MI_GFPve_pos  1.4707183761
## dCA2_Mpos_CGAAGGAAGAGCATAT-1dCA2_MI_GFPve_pos  1.0720640344
## dCA2_Mpos_CGAAGTTCAGCAGGAT-1dCA2_MI_GFPve_pos  0.3640919847
## dCA2_Mpos_CGAATTGTCAACCCGG-1dCA2_MI_GFPve_pos -0.6653954464
## dCA2_Mpos_CGAGAAGCAATGACCT-1dCA2_MI_GFPve_pos -0.4516909796
## dCA2_Mpos_CGAGGAAAGCTGCGAA-1dCA2_MI_GFPve_pos  0.3956683797
## dCA2_Mpos_CGAGGAATCGCTGACG-1dCA2_MI_GFPve_pos -0.0433532673
## dCA2_Mpos_CGAGGCTCACATCATG-1dCA2_MI_GFPve_pos -0.5001272636
## dCA2_Mpos_CGAGTTATCTTTGCAT-1dCA2_MI_GFPve_pos  2.2884831351
## dCA2_Mpos_CGATCGGTCTTCTGTA-1dCA2_MI_GFPve_pos  0.8293935579
## dCA2_Mpos_CGATGCGGTTAGGGAC-1dCA2_MI_GFPve_pos  0.2225916756
## dCA2_Mpos_CGCAGGTGTGTCATCA-1dCA2_MI_GFPve_pos  0.8663090986
## dCA2_Mpos_CGCATAACAGAGGCTA-1dCA2_MI_GFPve_pos  2.2583043498
## dCA2_Mpos_CGCATAATCCAAGCTA-1dCA2_MI_GFPve_pos  1.3182617349
## dCA2_Mpos_CGCATGGAGACCATAA-1dCA2_MI_GFPve_pos  1.5398380679
## dCA2_Mpos_CGCATGGCACAATGCT-1dCA2_MI_GFPve_pos -0.8040031272
## dCA2_Mpos_CGCATGGCATGTGCTA-1dCA2_MI_GFPve_pos  2.3285865707
## dCA2_Mpos_CGCCAGACAAACGGCA-1dCA2_MI_GFPve_pos -0.0521647709
## dCA2_Mpos_CGCCATTAGGCTGTAG-1dCA2_MI_GFPve_pos  1.1715446634
## dCA2_Mpos_CGCCATTAGGTAGACC-1dCA2_MI_GFPve_pos  2.4673070831
## dCA2_Mpos_CGCCATTGTGTAACGG-1dCA2_MI_GFPve_pos  1.7320781869
## dCA2_Mpos_CGCGTGAGTAGTATAG-1dCA2_MI_GFPve_pos  2.3448080939
## dCA2_Mpos_CGGAACCTCTAAGGAA-1dCA2_MI_GFPve_pos  1.4493037385
## dCA2_Mpos_CGGAGAAAGCACCAGA-1dCA2_MI_GFPve_pos -0.4932442385
## dCA2_Mpos_CGGCAGTAGATCCGAG-1dCA2_MI_GFPve_pos  1.8242365045
## dCA2_Mpos_CGGGACTAGAATAACC-1dCA2_MI_GFPve_pos  0.2996636105
## dCA2_Mpos_CGGGACTAGACTTCAC-1dCA2_MI_GFPve_pos  1.6239839715
## dCA2_Mpos_CGGTCAGCAATCCAGT-1dCA2_MI_GFPve_pos  1.3333475513
## dCA2_Mpos_CGTAAGTGTGGCTGCT-1dCA2_MI_GFPve_pos  0.1033799943
## dCA2_Mpos_CGTAAGTTCTCTGGTC-1dCA2_MI_GFPve_pos  0.7531899733
## dCA2_Mpos_CGTCAAAAGCCTCTCT-1dCA2_MI_GFPve_pos  1.3848326129
## dCA2_Mpos_CGTCAAACAATGTCTG-1dCA2_MI_GFPve_pos  2.2875053806
## dCA2_Mpos_CGTCAAACATCCGAGC-1dCA2_MI_GFPve_pos  1.0498704833
## dCA2_Mpos_CGTCAAACATCCTGTC-1dCA2_MI_GFPve_pos  0.6992005033
## dCA2_Mpos_CGTCAAAGTCGACTGC-1dCA2_MI_GFPve_pos  1.8914592189
## dCA2_Mpos_CGTCAAATCAACCCGG-1dCA2_MI_GFPve_pos  2.1703412456
## dCA2_Mpos_CGTGCTTAGACCTGGA-1dCA2_MI_GFPve_pos  1.8950394315
## dCA2_Mpos_CGTGCTTCAGGCCCTA-1dCA2_MI_GFPve_pos  0.7630415482
## dCA2_Mpos_CGTTAGAAGGTCGTCC-1dCA2_MI_GFPve_pos  1.0345524472
## dCA2_Mpos_CGTTGGGAGATACATG-1dCA2_MI_GFPve_pos  2.1918461246
## dCA2_Mpos_CGTTGGGCATTGTACG-1dCA2_MI_GFPve_pos  0.3195422066
## dCA2_Mpos_CGTTGGGGTATAGCTC-1dCA2_MI_GFPve_pos  2.4113361758
## dCA2_Mpos_CTAACCCTCAAGGACG-1dCA2_MI_GFPve_pos  0.5399841291
## dCA2_Mpos_CTAACTTCACGTACAT-1dCA2_MI_GFPve_pos  0.5360071880
## dCA2_Mpos_CTACATTTCAAGCCTA-1dCA2_MI_GFPve_pos  0.7825894875
## dCA2_Mpos_CTACCTGCATAAGCAA-1dCA2_MI_GFPve_pos  0.9225792212
## dCA2_Mpos_CTACCTGGTAGCGATG-1dCA2_MI_GFPve_pos  0.0574620364
## dCA2_Mpos_CTACCTGTCCGTATGA-1dCA2_MI_GFPve_pos  0.5981658620
## dCA2_Mpos_CTACTATAGAGATTCA-1dCA2_MI_GFPve_pos -1.4350493031
## dCA2_Mpos_CTAGACATCGTCAAAC-1dCA2_MI_GFPve_pos  1.7360401077
## dCA2_Mpos_CTAGGTAGTCCTCAGG-1dCA2_MI_GFPve_pos  0.0441838985
## dCA2_Mpos_CTATAGGCAACACACT-1dCA2_MI_GFPve_pos  0.1405406033
## dCA2_Mpos_CTCAACCAGACGCATG-1dCA2_MI_GFPve_pos  2.6608850879
## dCA2_Mpos_CTCAACCCAACTTCTT-1dCA2_MI_GFPve_pos -0.0442741441
## dCA2_Mpos_CTCAACCGTAATGTGA-1dCA2_MI_GFPve_pos -0.8175800520
## dCA2_Mpos_CTCAAGAGTGTTGATC-1dCA2_MI_GFPve_pos  0.7974418325
## dCA2_Mpos_CTCACTGAGATCACTC-1dCA2_MI_GFPve_pos  1.7466432971
## dCA2_Mpos_CTCAGAACAGGTTTAC-1dCA2_MI_GFPve_pos  0.8820474071
## dCA2_Mpos_CTCAGGGAGCTACTAC-1dCA2_MI_GFPve_pos -0.8312497216
## dCA2_Mpos_CTCAGTCAGTCACACT-1dCA2_MI_GFPve_pos  1.2768719119
## dCA2_Mpos_CTCAGTCTCCGCGATG-1dCA2_MI_GFPve_pos -1.2788113433
## dCA2_Mpos_CTCATCGCATGTGCCG-1dCA2_MI_GFPve_pos  0.5623993499
## dCA2_Mpos_CTCATGCTCATGCCAA-1dCA2_MI_GFPve_pos  1.2048654479
## dCA2_Mpos_CTCATTAAGTGACACG-1dCA2_MI_GFPve_pos  1.5435459537
## dCA2_Mpos_CTCCAACAGCTAATGA-1dCA2_MI_GFPve_pos -0.4741468507
## dCA2_Mpos_CTCCACACATCGATGT-1dCA2_MI_GFPve_pos  2.0170569343
## dCA2_Mpos_CTCCATGCATCCTATT-1dCA2_MI_GFPve_pos  2.1432928962
## dCA2_Mpos_CTCCCTCAGGAAAGAC-1dCA2_MI_GFPve_pos  2.0384900493
## dCA2_Mpos_CTCCTCCGTTACCTTT-1dCA2_MI_GFPve_pos  1.8254636449
## dCA2_Mpos_CTCTCAGAGGAGAATG-1dCA2_MI_GFPve_pos  1.3667822999
```

```
## dCA2_Mpos_CTCTCAGCATTGGCAT-1dCA2_MI_GFPve_pos  1.9062008781
## dCA2_Mpos_CTCTCGAGTTGTGTAC-1dCA2_MI_GFPve_pos -0.2225366610
## dCA2_Mpos_CTGAGGCTCGCCGAAC-1dCA2_MI_GFPve_pos  1.4577813071
## dCA2_Mpos_CTGCAGGGTATCTTCT-1dCA2_MI_GFPve_pos  2.2517933768
## dCA2_Mpos_CTGCAGGTCTAGTGTG-1dCA2_MI_GFPve_pos  0.3584011120
## dCA2_Mpos_CTGCATCGTGAATTGA-1dCA2_MI_GFPve_pos -0.0154891538
## dCA2_Mpos_CTGCATCTCCCTCATG-1dCA2_MI_GFPve_pos -1.1141693669
## dCA2_Mpos_CTGCTCAGTTGCCAAT-1dCA2_MI_GFPve_pos -1.1687380153
## dCA2_Mpos_CTGGCAGCAAGTGGTG-1dCA2_MI_GFPve_pos  2.0250927133
## dCA2_Mpos_CTGGCAGGTAGTGGCA-1dCA2_MI_GFPve_pos -0.6111228543
## dCA2_Mpos_CTGGTCTAGAAATTCG-1dCA2_MI_GFPve_pos -0.0484489727
## dCA2_Mpos_CTGTATTAGAGGTCAC-1dCA2_MI_GFPve_pos  0.9212362689
## dCA2_Mpos_CTTACCGAGGGATCGT-1dCA2_MI_GFPve_pos  1.8268489761
## dCA2_Mpos_CTTACCGGTGTCATGT-1dCA2_MI_GFPve_pos -0.1834729689
## dCA2_Mpos_CTTAGGAGTAGACAAT-1dCA2_MI_GFPve_pos -1.5639181214
## dCA2_Mpos_CTTCCTTTCTATTGTC-1dCA2_MI_GFPve_pos -0.6859508830
## dCA2_Mpos_CTTCTAAGTACTCGTA-1dCA2_MI_GFPve_pos  1.3118182344
## dCA2_Mpos_CTTGAGAAGGTAGCCA-1dCA2_MI_GFPve_pos  1.8998998326
## dCA2_Mpos_CTTTCGGAGGCGATAC-1dCA2_MI_GFPve_pos  0.0962664999
## dCA2_Mpos_GAAACCTCACAACCGC-1dCA2_MI_GFPve_pos -0.8565031963
## dCA2_Mpos_GAAACCTTCCGAGAAG-1dCA2_MI_GFPve_pos  0.1152040723
## dCA2_Mpos_GAAATGAAGATGAACT-1dCA2_MI_GFPve_pos  0.8783956689
## dCA2_Mpos_GAAATGAGTCGTGCCA-1dCA2_MI_GFPve_pos  1.0853904409
## dCA2_Mpos_GAAGAATGTTGGTGTT-1dCA2_MI_GFPve_pos  1.2054901046
## dCA2_Mpos_GAAGCCCCAAAGGCTG-1dCA2_MI_GFPve_pos  1.8471980018
## dCA2_Mpos_GAAGCGAGTTTCTATC-1dCA2_MI_GFPve_pos -1.3977043706
## dCA2_Mpos_GAAGGACTCTCCATAT-1dCA2_MI_GFPve_pos  0.3772579652
## dCA2_Mpos_GAAGGGTGTACCCGCA-1dCA2_MI_GFPve_pos  0.9569858832
## dCA2_Mpos_GAAGGGTTCTGGCTGG-1dCA2_MI_GFPve_pos  0.1867406142
## dCA2_Mpos_GAATAGATCCACAGGC-1dCA2_MI_GFPve_pos -0.8693887668
## dCA2_Mpos_GAATAGATCCTCTTTC-1dCA2_MI_GFPve_pos -0.1844324547
## dCA2_Mpos_GAATCACTCGAGAGCA-1dCA2_MI_GFPve_pos  0.7052965087
## dCA2_Mpos_GACAGCCGTACCTATG-1dCA2_MI_GFPve_pos  0.7944430036
## dCA2_Mpos_GACATCATCCGGTAAT-1dCA2_MI_GFPve_pos -0.4544150310
## dCA2_Mpos_GACCCAGGTGAAGCTG-1dCA2_MI_GFPve_pos  1.2516575975
## dCA2_Mpos_GACCCTTCAGGTGACA-1dCA2_MI_GFPve_pos -0.9998112994
## dCA2_Mpos_GACCGTGAGCAGAAAG-1dCA2_MI_GFPve_pos  1.7178829831
## dCA2_Mpos_GACTATGTCAGCCTTC-1dCA2_MI_GFPve_pos  2.0250352544
## dCA2_Mpos_GACTATGTCCGACAGC-1dCA2_MI_GFPve_pos  1.6185057086
## dCA2_Mpos_GACTCAAAGGCACCAA-1dCA2_MI_GFPve_pos  1.0330833120
## dCA2_Mpos_GACTCTCGTATCGATC-1dCA2_MI_GFPve_pos -0.0520911443
## dCA2_Mpos_GACTGATTCGAGCTGC-1dCA2_MI_GFPve_pos  1.1739489955
## dCA2_Mpos_GAGAAATAGATACCAA-1dCA2_MI_GFPve_pos  0.7808158321
## dCA2_Mpos_GAGACCCCATCCCGTT-1dCA2_MI_GFPve_pos -0.2470299977
## dCA2_Mpos_GAGAGGTCAGAGTAAT-1dCA2_MI_GFPve_pos  0.9301052612
## dCA2_Mpos_GAGAGGTCAGGTCAAG-1dCA2_MI_GFPve_pos  0.5754304153
## dCA2_Mpos_GAGAGGTGTGCCTATA-1dCA2_MI_GFPve_pos -1.3369412499
## dCA2_Mpos_GAGATGGTCAGTGATC-1dCA2_MI_GFPve_pos -0.9201335984
## dCA2_Mpos_GAGATGGTCTCGCGTT-1dCA2_MI_GFPve_pos -0.7535590010
## dCA2_Mpos_GAGCCTGGTGGGAGAG-1dCA2_MI_GFPve_pos  1.5131950301
## dCA2_Mpos_GAGCTGCTCAGACCCG-1dCA2_MI_GFPve_pos  0.5643098575
## dCA2_Mpos_GAGGCAAAGGAAGAAC-1dCA2_MI_GFPve_pos  2.0956193132
## dCA2_Mpos_GAGGCAATCAAACCTG-1dCA2_MI_GFPve_pos  1.2127598447
## dCA2_Mpos_GAGGCCTGTTCATCGA-1dCA2_MI_GFPve_pos  1.1815636081
## dCA2_Mpos_GAGGGTAAGGACAGTC-1dCA2_MI_GFPve_pos  0.9190795345
## dCA2_Mpos_GAGGGTACAAGAGTAT-1dCA2_MI_GFPve_pos  0.3768443388
## dCA2_Mpos_GAGGGTACAAGGCTTT-1dCA2_MI_GFPve_pos  1.0578446311
## dCA2_Mpos_GAGGGTAGTACTAGCT-1dCA2_MI_GFPve_pos  1.9965783281
## dCA2_Mpos_GAGTCATTCCATCTGC-1dCA2_MI_GFPve_pos  1.1912965697
## dCA2_Mpos_GAGTCTACATACGCAT-1dCA2_MI_GFPve_pos  0.1193859929
## dCA2_Mpos_GAGTGAGTCAACGCTA-1dCA2_MI_GFPve_pos  0.3181795550
## dCA2_Mpos_GAGTGAGTCTTCTCAA-1dCA2_MI_GFPve_pos  0.9180417580
## dCA2_Mpos_GAGTTACGTTAGAGAT-1dCA2_MI_GFPve_pos  0.2032386062
## dCA2_Mpos_GAGTTGTAGTGATAAC-1dCA2_MI_GFPve_pos -1.1077103692
## dCA2_Mpos_GAGTTGTCACTCTGCT-1dCA2_MI_GFPve_pos -0.6585612970
## dCA2_Mpos_GAGTTTGTCGCTTGAA-1dCA2_MI_GFPve_pos  1.4487196130
## dCA2_Mpos_GATAGAAGTGAAGCGT-1dCA2_MI_GFPve_pos  0.4018742663
## dCA2_Mpos_GATAGCTAGGTAACTA-1dCA2_MI_GFPve_pos  0.1898090464
## dCA2_Mpos_GATAGCTCAACGGTAG-1dCA2_MI_GFPve_pos -0.1631343740
## dCA2_Mpos_GATAGCTCAGCTTTGA-1dCA2_MI_GFPve_pos -0.3365534323
```

```
## dCA2_Mpos_GATCACAGTAGTGTGG-1dCA2_MI_GFPve_pos  1.1324926538
## dCA2_Mpos_GATCATGAGATTAGAC-1dCA2_MI_GFPve_pos -0.4893832880
## dCA2_Mpos_GATCATGCAGCGGATA-1dCA2_MI_GFPve_pos -0.8162340599
## dCA2_Mpos_GATCCCTAGCGGCTCT-1dCA2_MI_GFPve_pos -0.5534336644
## dCA2_Mpos_GATCGTATCATGCCAA-1dCA2_MI_GFPve_pos  1.9209434909
## dCA2_Mpos_GATCGTATCCTACGGG-1dCA2_MI_GFPve_pos  0.5577105505
## dCA2_Mpos_GATGATCCAAGAAACT-1dCA2_MI_GFPve_pos -1.3991348820
## dCA2_Mpos_GATGCTAAGCACTCTA-1dCA2_MI_GFPve_pos  0.9122100395
## dCA2_Mpos_GATGGAGAGTGCGCTC-1dCA2_MI_GFPve_pos  2.0108321828
## dCA2_Mpos_GATGGAGCAGGCCTGT-1dCA2_MI_GFPve_pos -0.9104192334
## dCA2_Mpos_GATGTTGGTCATGCAT-1dCA2_MI_GFPve_pos  0.2974285257
## dCA2_Mpos_GATTCGAAGCGTCGAA-1dCA2_MI_GFPve_pos  0.5754145903
## dCA2_Mpos_GATTCGACACCTGTCT-1dCA2_MI_GFPve_pos -0.4690275865
## dCA2_Mpos_GATTCTTGTGCCTGAC-1dCA2_MI_GFPve_pos  0.6892551107
## dCA2_Mpos_GATTCTTTCTTGGGCG-1dCA2_MI_GFPve_pos  0.3134482277
## dCA2_Mpos_GCAACCGCAAGGGTCA-1dCA2_MI_GFPve_pos  2.1998157424
## dCA2_Mpos_GCACATAGTACTCGCG-1dCA2_MI_GFPve_pos  0.4402721447
## dCA2_Mpos_GCACGTGCAAGCTGCC-1dCA2_MI_GFPve_pos  2.1400678081
## dCA2_Mpos_GCACGTGCAGATGCGA-1dCA2_MI_GFPve_pos  2.3944773597
## dCA2_Mpos_GCACGTGTCCCGAAAT-1dCA2_MI_GFPve_pos -1.5000780898
## dCA2_Mpos_GCACTAATCACGGACC-1dCA2_MI_GFPve_pos -0.4441329318
## dCA2_Mpos_GCAGCCAGTCGATTAC-1dCA2_MI_GFPve_pos  0.1773480472
## dCA2_Mpos_GCAGCCATCATCACTT-1dCA2_MI_GFPve_pos  2.0114275140
## dCA2_Mpos_GCAGCTGAGTCCCGGT-1dCA2_MI_GFPve_pos -1.6131562072
## dCA2_Mpos_GCAGGCTTCCATCAGA-1dCA2_MI_GFPve_pos -0.0870261567
## dCA2_Mpos_GCAGTTATCTGGGCAC-1dCA2_MI_GFPve_pos  0.5720578653
## dCA2_Mpos_GCATCGGAGGCACTAG-1dCA2_MI_GFPve_pos  2.5351633949
## dCA2_Mpos_GCATCGGCATGAGAAT-1dCA2_MI_GFPve_pos  1.4278317613
## dCA2_Mpos_GCATCGGGTAGGACTG-1dCA2_MI_GFPve_pos  0.8393734616
## dCA2_Mpos_GCATCGGGTCTAGGTT-1dCA2_MI_GFPve_pos -0.8712973076
## dCA2_Mpos_GCATCTCCAGAAACCG-1dCA2_MI_GFPve_pos  0.1473074702
## dCA2_Mpos_GCATGATTCAGTGTTG-1dCA2_MI_GFPve_pos -0.2019572037
## dCA2_Mpos_GCCAGGTAGGTTAAAC-1dCA2_MI_GFPve_pos  0.5354915780
## dCA2_Mpos_GCCAGGTGTGTCCACG-1dCA2_MI_GFPve_pos  2.5323047561
## dCA2_Mpos_GCCATGGGTCTACGAT-1dCA2_MI_GFPve_pos  0.4983858687
## dCA2_Mpos_GCCATTCAGCCTTTGA-1dCA2_MI_GFPve_pos  0.4778018219
## dCA2_Mpos_GCCATTCTCCTTGACC-1dCA2_MI_GFPve_pos  1.5553148908
## dCA2_Mpos_GCCCAGACACAACGAG-1dCA2_MI_GFPve_pos  1.4407491607
## dCA2_Mpos_GCCCAGACATTCTGTT-1dCA2_MI_GFPve_pos  1.3901046437
## dCA2_Mpos_GCCCAGAGTAGCGAGT-1dCA2_MI_GFPve_pos  0.4146650655
## dCA2_Mpos_GCCGATGCACGCGCAT-1dCA2_MI_GFPve_pos  1.3351008815
## dCA2_Mpos_GCCGTGATCAACTCTT-1dCA2_MI_GFPve_pos  1.8389977140
## dCA2_Mpos_GCCTGTTGTGGTCCCA-1dCA2_MI_GFPve_pos -0.7048247534
## dCA2_Mpos_GCGAGAAAGAGGGTGG-1dCA2_MI_GFPve_pos  1.2266155404
## dCA2_Mpos_GCGAGAATCCTTCACG-1dCA2_MI_GFPve_pos -1.0159396010
## dCA2_Mpos_GCTGAATAGACATCAA-1dCA2_MI_GFPve_pos -1.2886974889
## dCA2_Mpos_GCTGAATTCGTGGAAG-1dCA2_MI_GFPve_pos  1.7352325839
## dCA2_Mpos_GCTGCAGCAAATCGTC-1dCA2_MI_GFPve_pos -0.5836536961
## dCA2_Mpos_GCTGCAGTCACGACTA-1dCA2_MI_GFPve_pos  0.6164737863
## dCA2_Mpos_GCTGGGTCAAGCCATT-1dCA2_MI_GFPve_pos  0.3669710678
## dCA2_Mpos_GCTGGGTGTCACCCTT-1dCA2_MI_GFPve_pos  0.7467814011
## dCA2_Mpos_GGAACCCTCGCACGGT-1dCA2_MI_GFPve_pos -0.1734126883
## dCA2_Mpos_GGAAGTGAGTCCGTCG-1dCA2_MI_GFPve_pos  1.1085669917
## dCA2_Mpos_GGAATCTCAGGAAGTC-1dCA2_MI_GFPve_pos  2.0292235536
## dCA2_Mpos_GGAGAACGTCTGTGCG-1dCA2_MI_GFPve_pos -1.3979886847
## dCA2_Mpos_GGAGCAAAGCTCACTA-1dCA2_MI_GFPve_pos -0.0206096771
## dCA2_Mpos_GGAGGATAGTAGTCCT-1dCA2_MI_GFPve_pos  0.8499096555
## dCA2_Mpos_GGAGGTACAGTATACC-1dCA2_MI_GFPve_pos -0.0367737788
## dCA2_Mpos_GGATCTAAGCACCTGC-1dCA2_MI_GFPve_pos  1.0196766776
## dCA2_Mpos_GGCAGTCAGGGTTGCA-1dCA2_MI_GFPve_pos  1.7115641755
## dCA2_Mpos_GGCTGTGCAGTGGGTA-1dCA2_MI_GFPve_pos  1.2998615426
## dCA2_Mpos_GGCTGTGCATAGGCGA-1dCA2_MI_GFPve_pos  1.5777366084
## dCA2_Mpos_GGCTGTGGTACCCACG-1dCA2_MI_GFPve_pos -0.3541808503
## dCA2_Mpos_GGCTGTGGTCAAAGCG-1dCA2_MI_GFPve_pos  0.6241701764
## dCA2_Mpos_GGCTGTGGTCTGTGGC-1dCA2_MI_GFPve_pos  1.9539842529
## dCA2_Mpos_GGCTGTGTCGGCCCAA-1dCA2_MI_GFPve_pos  0.5558469814
## dCA2_Mpos_GGCTTGGAGTTCACTG-1dCA2_MI_GFPve_pos -0.1286359208
## dCA2_Mpos_GGCTTGGCAACAACAA-1dCA2_MI_GFPve_pos  1.2227473182
## dCA2_Mpos_GGCTTGGTCAGGAAAT-1dCA2_MI_GFPve_pos  0.8282002730
## dCA2_Mpos_GGGAAGTAGTCTAGCT-1dCA2_MI_GFPve_pos  2.1816611213
```

```
## dCA2_Mpos_GGGAAGTGTATTTCCT-1dCA2_MI_GFPve_pos -0.0280524271
## dCA2_Mpos_GGGACAATCAACACGT-1dCA2_MI_GFPve_pos  0.5425290686
## dCA2_Mpos_GGGACCTCATGCCATA-1dCA2_MI_GFPve_pos  1.8614615125
## dCA2_Mpos_GGGACCTGTACTGTTG-1dCA2_MI_GFPve_pos  0.0240854991
## dCA2_Mpos_GGGACTCTCCTCACGT-1dCA2_MI_GFPve_pos  0.4395546836
## dCA2_Mpos_GGGAGATTCGACGCGT-1dCA2_MI_GFPve_pos  0.9448147339
## dCA2_Mpos_GGGAGTACACCATAAC-1dCA2_MI_GFPve_pos  1.0278787536
## dCA2_Mpos_GGGAGTATCGACGTCG-1dCA2_MI_GFPve_pos -0.6256613212
## dCA2_Mpos_GGGCTACCACAGAGAC-1dCA2_MI_GFPve_pos  0.9184589309
## dCA2_Mpos_GGGCTCAGTACGAGCA-1dCA2_MI_GFPve_pos  1.2391060514
## dCA2_Mpos_GGGCTCATCCCAAGTA-1dCA2_MI_GFPve_pos  0.6741846127
## dCA2_Mpos_GGGTAGAGTCGAAACG-1dCA2_MI_GFPve_pos  0.9167013091
## dCA2_Mpos_GGGTCACTCGCTCATC-1dCA2_MI_GFPve_pos -0.2483836489
## dCA2_Mpos_GGGTCTGAGCCTGAAG-1dCA2_MI_GFPve_pos  1.5418821496
## dCA2_Mpos_GGGTGAATCACTTTGT-1dCA2_MI_GFPve_pos  0.0664380749
## dCA2_Mpos_GGGTTATGTCACGTGC-1dCA2_MI_GFPve_pos -1.5206005650
## dCA2_Mpos_GGGTTATTCATGGAGG-1dCA2_MI_GFPve_pos  0.4268048925
## dCA2_Mpos_GGTAACTTCCATCTGC-1dCA2_MI_GFPve_pos -0.3074906247
## dCA2_Mpos_GGTAATCAGCCGTAAG-1dCA2_MI_GFPve_pos  2.6228837890
## dCA2_Mpos_GGTAATCAGGCCCAAA-1dCA2_MI_GFPve_pos  0.6698554081
## dCA2_Mpos_GGTAATCCAAGCGGAT-1dCA2_MI_GFPve_pos -0.5728115516
## dCA2_Mpos_GGTAATCGTAGCTTTG-1dCA2_MI_GFPve_pos  2.0635692996
## dCA2_Mpos_GGTAATCGTTCGGCTG-1dCA2_MI_GFPve_pos  1.1872140092
## dCA2_Mpos_GGTAGAGAGTGCAACG-1dCA2_MI_GFPve_pos  0.6137375755
## dCA2_Mpos_GGTAGAGCATTCACAG-1dCA2_MI_GFPve_pos  0.0717931015
## dCA2_Mpos_GGTAGAGTCCATCAGA-1dCA2_MI_GFPve_pos  1.0583036465
## dCA2_Mpos_GGTGAAGTCCATCTCG-1dCA2_MI_GFPve_pos  0.5300809068
## dCA2_Mpos_GGTGATTGTAATCAGA-1dCA2_MI_GFPve_pos  1.2258627338
## dCA2_Mpos_GGTGGCTTCTCTAAGG-1dCA2_MI_GFPve_pos  1.1078504247
## dCA2_Mpos_GGTGTCGTCAACCCGG-1dCA2_MI_GFPve_pos -0.5499767738
## dCA2_Mpos_GGTGTTACAATGGCAG-1dCA2_MI_GFPve_pos  1.6332724017
## dCA2_Mpos_GGTTAACAGGTTAGTA-1dCA2_MI_GFPve_pos  0.7284974975
## dCA2_Mpos_GGTTAACGTCATCCCT-1dCA2_MI_GFPve_pos  0.3096136225
## dCA2_Mpos_GGTTAACTCATGGATC-1dCA2_MI_GFPve_pos  0.3804767055
## dCA2_Mpos_GGTTAACTCTTACCAT-1dCA2_MI_GFPve_pos  2.7586049957
## dCA2_Mpos_GGTTCTCAGAAGAGCA-1dCA2_MI_GFPve_pos  2.2031233234
## dCA2_Mpos_GGTTCTCAGCACTCCG-1dCA2_MI_GFPve_pos  1.2876148147
## dCA2_Mpos_GGTTCTCAGTGTACAA-1dCA2_MI_GFPve_pos  1.0923546476
## dCA2_Mpos_GGTTCTCGTTTAGAGA-1dCA2_MI_GFPve_pos  0.4909709317
## dCA2_Mpos_GTAACACGTATCCTCC-1dCA2_MI_GFPve_pos  2.3714127464
## dCA2_Mpos_GTAACACTCAAGAGTA-1dCA2_MI_GFPve_pos  1.1729717178
## dCA2_Mpos_GTAACCAAGATCGACG-1dCA2_MI_GFPve_pos  1.4356458110
## dCA2_Mpos_GTAACCAGTGGGCTCT-1dCA2_MI_GFPve_pos  0.8554248733
## dCA2_Mpos_GTAAGTCGTATACCTG-1dCA2_MI_GFPve_pos -0.7978274303
## dCA2_Mpos_GTAAGTCTCTACCACC-1dCA2_MI_GFPve_pos -1.2604881602
## dCA2_Mpos_GTAATCGAGCAAGCCA-1dCA2_MI_GFPve_pos -0.7064883190
## dCA2_Mpos_GTAATCGCAATCGCCG-1dCA2_MI_GFPve_pos  0.4454857928
## dCA2_Mpos_GTAATGCAGATGGTAT-1dCA2_MI_GFPve_pos  1.5415151042
## dCA2_Mpos_GTAATGCCACAGTCCG-1dCA2_MI_GFPve_pos -0.0792041349
## dCA2_Mpos_GTACAACGTATCGTAC-1dCA2_MI_GFPve_pos -0.7078796106
## dCA2_Mpos_GTACAGTGTGTTGATC-1dCA2_MI_GFPve_pos -0.7179685908
## dCA2_Mpos_GTAGAAAGTGGATTTC-1dCA2_MI_GFPve_pos -1.2488429623
## dCA2_Mpos_GTAGAGGGTCTCCCTA-1dCA2_MI_GFPve_pos  2.0735881252
## dCA2_Mpos_GTAGATCTCTGTACAG-1dCA2_MI_GFPve_pos  2.3719754142
## dCA2_Mpos_GTCAAACGTCCAGGTC-1dCA2_MI_GFPve_pos  0.7172851485
## dCA2_Mpos_GTCACTCAGACTGTTC-1dCA2_MI_GFPve_pos  1.7631052655
## dCA2_Mpos_GTCACTCTCTTGGTGA-1dCA2_MI_GFPve_pos  1.5132311506
## dCA2_Mpos_GTCAGCGGTTTAGACC-1dCA2_MI_GFPve_pos  1.0203369779
## dCA2_Mpos_GTCATCCCACTTGTCC-1dCA2_MI_GFPve_pos  1.3888095540
## dCA2_Mpos_GTCATTTAGCTGACTT-1dCA2_MI_GFPve_pos  1.8498076123
## dCA2_Mpos_GTCATTTGTAAGATTG-1dCA2_MI_GFPve_pos  0.3665504379
## dCA2_Mpos_GTCCACTCAGCCATTA-1dCA2_MI_GFPve_pos  1.5282833499
## dCA2_Mpos_GTCCACTGTACTCAAC-1dCA2_MI_GFPve_pos  1.5819051189
## dCA2_Mpos_GTCCACTTCGCTTGAA-1dCA2_MI_GFPve_pos -0.4895084577
## dCA2_Mpos_GTCCCATTCAAGTGTC-1dCA2_MI_GFPve_pos  1.6225212735
## dCA2_Mpos_GTCCTCAGTTTCGGCG-1dCA2_MI_GFPve_pos  2.7764892501
## dCA2_Mpos_GTCGCGACAGCCCAGT-1dCA2_MI_GFPve_pos  1.1824606580
## dCA2_Mpos_GTCGCGAGTTTACTGG-1dCA2_MI_GFPve_pos  0.5249078495
## dCA2_Mpos_GTCTACCAGCATCGAG-1dCA2_MI_GFPve_pos  0.5325373036
## dCA2_Mpos_GTCTAGACAGGCTATT-1dCA2_MI_GFPve_pos  0.9801201147
```

```
## dCA2_Mpos_GTCTAGAGTCTTGAAC-1dCA2_MI_GFPve_pos  2.0162328405
## dCA2_Mpos_GTCTCACGTCTCAGGC-1dCA2_MI_GFPve_pos  0.8228788895
## dCA2_Mpos_GTCTGTCCAGCTCGGT-1dCA2_MI_GFPve_pos  2.5905067844
## dCA2_Mpos_GTCTGTCGTCAGTCTA-1dCA2_MI_GFPve_pos  0.2514923138
## dCA2_Mpos_GTCTGTCTCCTGTAAG-1dCA2_MI_GFPve_pos  0.8840118570
## dCA2_Mpos_GTGAGCCGTATCGCGC-1dCA2_MI_GFPve_pos -3.2892973500
## dCA2_Mpos_GTGAGCCGTGTCCATA-1dCA2_MI_GFPve_pos  2.4952120704
## dCA2_Mpos_GTGAGTTAGTAACGAT-1dCA2_MI_GFPve_pos -0.6788604933
## dCA2_Mpos_GTGATGTTCGTTGTTT-1dCA2_MI_GFPve_pos  0.1417467778
## dCA2_Mpos_GTGCGTGAGCGGTAGT-1dCA2_MI_GFPve_pos  1.0953607482
## dCA2_Mpos_GTGCGTGCAACCGCCA-1dCA2_MI_GFPve_pos -1.5951466637
## dCA2_Mpos_GTGCGTGCAATTGGTC-1dCA2_MI_GFPve_pos  0.0029750166
## dCA2_Mpos_GTGCTGGTCACCTCAC-1dCA2_MI_GFPve_pos  1.3694647474
## dCA2_Mpos_GTGCTTCAGTGCACCC-1dCA2_MI_GFPve_pos  0.5462634486
## dCA2_Mpos_GTGCTTCCAAGAGTAT-1dCA2_MI_GFPve_pos -0.7010208445
## dCA2_Mpos_GTGCTTCCAGCTGAGA-1dCA2_MI_GFPve_pos  0.5770787818
## dCA2_Mpos_GTGGAAGCATGCCGCA-1dCA2_MI_GFPve_pos  0.6711848301
## dCA2_Mpos_GTGGCGTCACCAAATC-1dCA2_MI_GFPve_pos -0.0591263550
## dCA2_Mpos_GTGGCGTTCAGGACGA-1dCA2_MI_GFPve_pos  0.2810416741
## dCA2_Mpos_GTGGGAAAGACCTCCG-1dCA2_MI_GFPve_pos  0.4829208476
## dCA2_Mpos_GTGGGAAGTAAGGAGA-1dCA2_MI_GFPve_pos  1.6012371701
## dCA2_Mpos_GTGGTTACACACCTGG-1dCA2_MI_GFPve_pos  1.6009598893
## dCA2_Mpos_GTGGTTACATAAGCAA-1dCA2_MI_GFPve_pos -0.5632082897
## dCA2_Mpos_GTGTAACCAACGCATT-1dCA2_MI_GFPve_pos  1.3595485610
## dCA2_Mpos_GTGTCCTCAGCACGAA-1dCA2_MI_GFPve_pos  1.5826615018
## dCA2_Mpos_GTGTGATTCGCACGGT-1dCA2_MI_GFPve_pos  0.5420157236
## dCA2_Mpos_GTGTTAGCATGATAGA-1dCA2_MI_GFPve_pos  2.2281665725
## dCA2_Mpos_GTGTTAGTCTGAGTCA-1dCA2_MI_GFPve_pos  0.5418616754
## dCA2_Mpos_GTGTTCCCACTCCACT-1dCA2_MI_GFPve_pos  0.9968589467
## dCA2_Mpos_GTTACGAAGGAAACGA-1dCA2_MI_GFPve_pos  0.9486424846
## dCA2_Mpos_GTTACGATCCCGAAAT-1dCA2_MI_GFPve_pos -0.6263455230
## dCA2_Mpos_GTTAGACCAGATGCGA-1dCA2_MI_GFPve_pos  1.8898506087
## dCA2_Mpos_GTTCATTCAAGGACAC-1dCA2_MI_GFPve_pos -0.3130952197
## dCA2_Mpos_GTTCTATCATAGATCC-1dCA2_MI_GFPve_pos  0.2453761769
## dCA2_Mpos_GTTCTATGTGTCATCA-1dCA2_MI_GFPve_pos  0.0927783183
## dCA2_Mpos_GTTGAACTCAAATAGG-1dCA2_MI_GFPve_pos  1.4656771344
## dCA2_Mpos_GTTGCTCTCATCGCAA-1dCA2_MI_GFPve_pos  1.1837292833
## dCA2_Mpos_GTTGTCCAGTATCCTG-1dCA2_MI_GFPve_pos  1.5397498531
## dCA2_Mpos_GTTGTGACACAGCATT-1dCA2_MI_GFPve_pos  0.9348136110
## dCA2_Mpos_GTTGTGACACCATTCC-1dCA2_MI_GFPve_pos  2.6161894721
## dCA2_Mpos_GTTTACTCATAGTCGT-1dCA2_MI_GFPve_pos  0.4930221957
## dCA2_Mpos_GTTTACTGTGAATTGA-1dCA2_MI_GFPve_pos  1.6933349294
## dCA2_Mpos_TAACGACAGTTCTACG-1dCA2_MI_GFPve_pos -0.4381598311
## dCA2_Mpos_TAACGACCAGAGATGC-1dCA2_MI_GFPve_pos -1.6711200553
## dCA2_Mpos_TAACTTCCATATGGCT-1dCA2_MI_GFPve_pos -0.2652984339
## dCA2_Mpos_TAAGCACCAGCCGTTG-1dCA2_MI_GFPve_pos  1.2166640681
## dCA2_Mpos_TAAGCACGTACACGTT-1dCA2_MI_GFPve_pos -0.3049413639
## dCA2_Mpos_TAAGCGTAGGGACACT-1dCA2_MI_GFPve_pos  1.2117388171
## dCA2_Mpos_TAATCTCGTCCGAAGA-1dCA2_MI_GFPve_pos -0.2397792714
## dCA2_Mpos_TAATCTCGTGGGCTTC-1dCA2_MI_GFPve_pos -1.4963167983
## dCA2_Mpos_TAATTCCGTACCATAC-1dCA2_MI_GFPve_pos  2.5085809154
## dCA2_Mpos_TACATTCAGGAGAATG-1dCA2_MI_GFPve_pos  2.3783655090
## dCA2_Mpos_TACATTCCACAAGTGG-1dCA2_MI_GFPve_pos  0.8812572283
## dCA2_Mpos_TACATTCCACCCTCTA-1dCA2_MI_GFPve_pos  0.5687520129
## dCA2_Mpos_TACATTCCATCTCGTC-1dCA2_MI_GFPve_pos  0.2517097962
## dCA2_Mpos_TACATTCGTCGTTCAA-1dCA2_MI_GFPve_pos  0.4334037823
## dCA2_Mpos_TACCCACGTGGTACAG-1dCA2_MI_GFPve_pos  1.0414745731
## dCA2_Mpos_TACCGGGAGCGTCAGA-1dCA2_MI_GFPve_pos  2.0534827632
## dCA2_Mpos_TACCGGGGTTCTCCTG-1dCA2_MI_GFPve_pos  2.1743028087
## dCA2_Mpos_TACCTCGTCATTACGG-1dCA2_MI_GFPve_pos  1.1597396058
## dCA2_Mpos_TACCTGCTCATAGAGA-1dCA2_MI_GFPve_pos -1.0881645757
## dCA2_Mpos_TACCTGCTCTACTCAT-1dCA2_MI_GFPve_pos -0.4274437505
## dCA2_Mpos_TACCTGCTCTGGTGGC-1dCA2_MI_GFPve_pos  2.5953268928
## dCA2_Mpos_TACGGGCAGACTCATC-1dCA2_MI_GFPve_pos  1.3033350629
## dCA2_Mpos_TACGGTATCTGCGGGT-1dCA2_MI_GFPve_pos  2.0204987449
## dCA2_Mpos_TACTGCCTCCTCAGAA-1dCA2_MI_GFPve_pos -0.0668653923
## dCA2_Mpos_TACTTCAAGTCCCTAA-1dCA2_MI_GFPve_pos  0.6188583893
## dCA2_Mpos_TAGACCAGTTGCTGAT-1dCA2_MI_GFPve_pos  1.0234326643
## dCA2_Mpos_TAGACTGCAGGATTCT-1dCA2_MI_GFPve_pos  0.0809115765
## dCA2_Mpos_TAGACTGCAGTGCGCT-1dCA2_MI_GFPve_pos  1.7370148820
```

```
## dCA2_Mpos_TAGACTGGTGCTCGTG-1dCA2_MI_GFPve_pos -0.9383455592
## dCA2_Mpos_TAGAGTCAGCTCTGTA-1dCA2_MI_GFPve_pos -0.2604046839
## dCA2_Mpos_TAGCACATCGGAACTT-1dCA2_MI_GFPve_pos -1.0093290883
## dCA2_Mpos_TAGGAGGAGTTCGGTT-1dCA2_MI_GFPve_pos  0.5922946555
## dCA2_Mpos_TAGGGTTTCTATCGGA-1dCA2_MI_GFPve_pos  0.9875335616
## dCA2_Mpos_TAGGTTGGTCTTTCTA-1dCA2_MI_GFPve_pos  1.2070518655
## dCA2_Mpos_TAGTGCAAGCGAATGC-1dCA2_MI_GFPve_pos  0.3363635016
## dCA2_Mpos_TATACCTCAGCTGTAT-1dCA2_MI_GFPve_pos -0.0846448171
## dCA2_Mpos_TATCCTACAACAGATA-1dCA2_MI_GFPve_pos -0.2445485073
## dCA2_Mpos_TATCGCCGTAAGTTGA-1dCA2_MI_GFPve_pos -1.4834451752
## dCA2_Mpos_TATCTTGGTTCTTCAT-1dCA2_MI_GFPve_pos  1.4384440107
## dCA2_Mpos_TATGTTCAGTGTGTTC-1dCA2_MI_GFPve_pos -0.9932465630
## dCA2_Mpos_TATGTTCGTGAAAGTT-1dCA2_MI_GFPve_pos  2.0049047393
## dCA2_Mpos_TATTCCAGTTATGTCG-1dCA2_MI_GFPve_pos -0.0525928366
## dCA2_Mpos_TATTCCATCTGGCCAG-1dCA2_MI_GFPve_pos -0.0105387049
## dCA2_Mpos_TATTGGGAGTAAACTG-1dCA2_MI_GFPve_pos -0.3112425881
## dCA2_Mpos_TCAAGCACAAGTAGTA-1dCA2_MI_GFPve_pos  1.8572727365
## dCA2_Mpos_TCAAGTGCAGCGTACC-1dCA2_MI_GFPve_pos  0.4247090441
## dCA2_Mpos_TCAATCTTCGCAGTCG-1dCA2_MI_GFPve_pos  1.2647502345
## dCA2_Mpos_TCACAAGAGATAGGGA-1dCA2_MI_GFPve_pos -0.3397229689
## dCA2_Mpos_TCACATTCAAAGCAAT-1dCA2_MI_GFPve_pos  0.2893241865
## dCA2_Mpos_TCACATTCAATACGCT-1dCA2_MI_GFPve_pos  1.4409406108
## dCA2_Mpos_TCACATTTCAGCGCAC-1dCA2_MI_GFPve_pos  1.7696549815
## dCA2_Mpos_TCACATTTCATCGCAA-1dCA2_MI_GFPve_pos  0.1702553925
## dCA2_Mpos_TCACGCTTCAGGAGAC-1dCA2_MI_GFPve_pos  0.9618819279
## dCA2_Mpos_TCACTCGTCCTACTGC-1dCA2_MI_GFPve_pos  1.2566964549
## dCA2_Mpos_TCAGTCCCAACCGCTG-1dCA2_MI_GFPve_pos -0.4627780991
## dCA2_Mpos_TCAGTTGTCAGCGTC-1dCA2_MI_GFPve_pos -0.1788896399
## dCA2_Mpos_TCAGTTTTCCTTCAGC-1dCA2_MI_GFPve_pos -0.0979476946
## dCA2_Mpos_TCATACTGTCTTTCAT-1dCA2_MI_GFPve_pos  2.0785930080
## dCA2_Mpos_TCATACTTCCTTCTAA-1dCA2_MI_GFPve_pos  2.2156503123
## dCA2_Mpos_TCATATCAGGACCCAA-1dCA2_MI_GFPve_pos  1.5949057264
## dCA2_Mpos_TCATATCGTGGTACAG-1dCA2_MI_GFPve_pos  1.8342931194
## dCA2_Mpos_TCATATCTCACAAGGG-1dCA2_MI_GFPve_pos  1.4440092963
## dCA2_Mpos_TCATCATCACAGCTTA-1dCA2_MI_GFPve_pos  0.8190663380
## dCA2_Mpos_TCATGCCCATCCGGTG-1dCA2_MI_GFPve_pos  2.5884623450
## dCA2_Mpos_TCATGTTGTCAAAGAT-1dCA2_MI_GFPve_pos  0.4057484490
## dCA2_Mpos_TCATGTTGTTCGGTCG-1dCA2_MI_GFPve_pos  2.2296078128
## dCA2_Mpos_TCATTACCAGGGTCTC-1dCA2_MI_GFPve_pos -0.2429563182
## dCA2_Mpos_TCATTCATCATGGATC-1dCA2_MI_GFPve_pos  1.3452393932
## dCA2_Mpos_TCATTGTGTATCGTAC-1dCA2_MI_GFPve_pos  1.0738285107
## dCA2_Mpos_TCATTGTTCACTGTCC-1dCA2_MI_GFPve_pos  0.7385727090
## dCA2_Mpos_TCATTTGCAACTCGTA-1dCA2_MI_GFPve_pos  2.0914366168
## dCA2_Mpos_TCATTTGGTTGAGGAC-1dCA2_MI_GFPve_pos  2.1019576711
## dCA2_Mpos_TCCACGTGTCGACTGC-1dCA2_MI_GFPve_pos  2.5192913932
## dCA2_Mpos_TCCACGTTCGTAGCTA-1dCA2_MI_GFPve_pos  1.8986996335
## dCA2_Mpos_TCCAGAACAAAGGGCT-1dCA2_MI_GFPve_pos  1.6067787332
## dCA2_Mpos_TCCATCGCAAGGGCAT-1dCA2_MI_GFPve_pos  2.2237040919
## dCA2_Mpos_TCCATCGCAGGGCTTC-1dCA2_MI_GFPve_pos  1.7378356380
## dCA2_Mpos_TCCATGCCAGACCAGA-1dCA2_MI_GFPve_pos  0.1036308300
## dCA2_Mpos_TCCCACATCTTTGCAT-1dCA2_MI_GFPve_pos  1.4533476752
## dCA2_Mpos_TCCCAGTAGCACAAAT-1dCA2_MI_GFPve_pos  1.4545739812
## dCA2_Mpos_TCCCAGTGTAGGAAAG-1dCA2_MI_GFPve_pos  1.3125514907
## dCA2_Mpos_TCCCAGTGTGGCTTAT-1dCA2_MI_GFPve_pos  1.0026925725
## dCA2_Mpos_TCCCAGTTCTACTGAG-1dCA2_MI_GFPve_pos -0.3040665465
## dCA2_Mpos_TCCCATGAGGGAGTTC-1dCA2_MI_GFPve_pos -0.3258337158
## dCA2_Mpos_TCCCATGTCCTACGAA-1dCA2_MI_GFPve_pos -0.3026507872
## dCA2_Mpos_TCCGAAAAGAACGCGT-1dCA2_MI_GFPve_pos  1.5642312688
## dCA2_Mpos_TCCGATCCAGTTTGGT-1dCA2_MI_GFPve_pos  0.1613345889
## dCA2_Mpos_TCCGATCTCGGTGAAG-1dCA2_MI_GFPve_pos  0.5708496196
## dCA2_Mpos_TCCGTGTTCTAACGGT-1dCA2_MI_GFPve_pos  2.1187238616
## dCA2_Mpos_TCCTCGAGTCCCGGTA-1dCA2_MI_GFPve_pos  0.7958743495
## dCA2_Mpos_TCCTCTTCAGATCACT-1dCA2_MI_GFPve_pos  1.1572986764
## dCA2_Mpos_TCCTCTTTCACGATAC-1dCA2_MI_GFPve_pos  0.8106474799
## dCA2_Mpos_TCCTGCAAGCCAAGCA-1dCA2_MI_GFPve_pos  0.9317083878
## dCA2_Mpos_TCCTTTCCATGCCATA-1dCA2_MI_GFPve_pos  1.4444758815
## dCA2_Mpos_TCGAACAAGGCAGCTA-1dCA2_MI_GFPve_pos  1.2795708102
## dCA2_Mpos_TCGCACTCATCGGCCA-1dCA2_MI_GFPve_pos  0.9349818153
## dCA2_Mpos_TCGCACTGTGATTAGA-1dCA2_MI_GFPve_pos  0.7688205761
## dCA2_Mpos_TCGCAGGCAGATACTC-1dCA2_MI_GFPve_pos  2.5073001308
```

```
## dCA2_Mpos_TCGCAGGGTGACACAG-1dCA2_MI_GFPve_pos -0.1517849344
## dCA2_Mpos_TCGGATAAGAGTTCGG-1dCA2_MI_GFPve_pos  0.7363944573
## dCA2_Mpos_TCGGATATCGAACTCA-1dCA2_MI_GFPve_pos  0.9374381823
## dCA2_Mpos_TCGGGACCACAAAGTA-1dCA2_MI_GFPve_pos  1.8638152999
## dCA2_Mpos_TCGGGCAGTGCCCGTA-1dCA2_MI_GFPve_pos  1.6416912002
## dCA2_Mpos_TCGGGTGAGAAAGCGA-1dCA2_MI_GFPve_pos  1.9817960185
## dCA2_Mpos_TCGTCCAAGTTGTCAC-1dCA2_MI_GFPve_pos  0.4708581788
## dCA2_Mpos_TCGTGGGCAAGTCATC-1dCA2_MI_GFPve_pos  2.2213179988
## dCA2_Mpos_TCTAACTGTACTCGCG-1dCA2_MI_GFPve_pos  0.4692923946
## dCA2_Mpos_TCTACATTCACTCTTA-1dCA2_MI_GFPve_pos  1.8382141513
## dCA2_Mpos_TCTACCGCAATCGCGC-1dCA2_MI_GFPve_pos  1.3281575364
## dCA2_Mpos_TCTATACTCGAAGCAG-1dCA2_MI_GFPve_pos  2.1617512626
## dCA2_Mpos_TCTATCAAGAGCCTGA-1dCA2_MI_GFPve_pos  1.4307506007
## dCA2_Mpos_TCTCAGCAGAACAGGA-1dCA2_MI_GFPve_pos  0.5733863634
## dCA2_Mpos_TCTCAGCCAACTCCCT-1dCA2_MI_GFPve_pos  0.9742967409
## dCA2_Mpos_TCTCAGCCAGCAATTC-1dCA2_MI_GFPve_pos  1.8686907214
## dCA2_Mpos_TCTCCGATCTCGACCT-1dCA2_MI_GFPve_pos  1.5622492952
## dCA2_Mpos_TCTCCGATCTGGGCAC-1dCA2_MI_GFPve_pos  2.3114085120
## dCA2_Mpos_TCTGCCAAGAGTTGTA-1dCA2_MI_GFPve_pos  1.0815111322
## dCA2_Mpos_TCTGTCGCACGTTGGC-1dCA2_MI_GFPve_pos  2.0810816211
## dCA2_Mpos_TCTGTCGGTGGTCTCG-1dCA2_MI_GFPve_pos -0.1624735373
## dCA2_Mpos_TCTGTCGTCGGTTGTA-1dCA2_MI_GFPve_pos  2.0668990535
## dCA2_Mpos_TCTTAGTAGGAATTAC-1dCA2_MI_GFPve_pos  1.1668822688
## dCA2_Mpos_TCTTAGTAGGTTCCAT-1dCA2_MI_GFPve_pos  1.8787760657
## dCA2_Mpos_TCTTTGATCCCAAGCG-1dCA2_MI_GFPve_pos  2.0001785678
## dCA2_Mpos_TGAACGTGTCGCCTAG-1dCA2_MI_GFPve_pos  0.7946496529
## dCA2_Mpos_TGAACGTTCCGTCAAA-1dCA2_MI_GFPve_pos  0.9580280108
## dCA2_Mpos_TGACGCGGTACAAACA-1dCA2_MI_GFPve_pos  2.0568215770
## dCA2_Mpos_TGACTCCGTATGACAA-1dCA2_MI_GFPve_pos -0.3501719611
## dCA2_Mpos_TGAGCGCAGGTGGTTG-1dCA2_MI_GFPve_pos  2.2480218333
## dCA2_Mpos_TGAGCGCGTTGCCGAC-1dCA2_MI_GFPve_pos  1.1003899497
## dCA2_Mpos_TGAGGTTGTCACGACC-1dCA2_MI_GFPve_pos  0.1508580332
## dCA2_Mpos_TGAGTCATCTAGAACC-1dCA2_MI_GFPve_pos -0.3430383282
## dCA2_Mpos_TGATCAGAGACTGAGC-1dCA2_MI_GFPve_pos  1.7028688115
## dCA2_Mpos_TGATCAGGTCTGCCTT-1dCA2_MI_GFPve_pos  2.4440038127
## dCA2_Mpos_TGATCAGTCAGCACCG-1dCA2_MI_GFPve_pos  0.8634675664
## dCA2_Mpos_TGATCAGTCGCGGACT-1dCA2_MI_GFPve_pos -0.2879251915
## dCA2_Mpos_TGATCAGTCGTGAGAG-1dCA2_MI_GFPve_pos  1.9297195596
## dCA2_Mpos_TGATCTTCACCAGCCA-1dCA2_MI_GFPve_pos  2.0519739312
## dCA2_Mpos_TGATGGTCAGTCGTTA-1dCA2_MI_GFPve_pos  0.2989344967
## dCA2_Mpos_TGATGGTTCATCCCGT-1dCA2_MI_GFPve_pos  0.3655442816
## dCA2_Mpos_TGATTTCCAGTCGGAA-1dCA2_MI_GFPve_pos  1.2907756490
## dCA2_Mpos_TGCACGGTCTCCGATC-1dCA2_MI_GFPve_pos  2.2038581294
## dCA2_Mpos_TGCAGATTCCATCTCG-1dCA2_MI_GFPve_pos -0.2892080026
## dCA2_Mpos_TGCAGGCAGGAACATT-1dCA2_MI_GFPve_pos -0.1282060819
## dCA2_Mpos_TGCAGGCTCTCAACCC-1dCA2_MI_GFPve_pos  0.5757833940
## dCA2_Mpos_TGCAGTAAGACCACGA-1dCA2_MI_GFPve_pos  0.9241946263
## dCA2_Mpos_TGCAGTAAGAGCCATG-1dCA2_MI_GFPve_pos -1.3483733016
## dCA2_Mpos_TGCATCCCACAAATCC-1dCA2_MI_GFPve_pos  1.2345639390
## dCA2_Mpos_TGCGACGGTACGTTCA-1dCA2_MI_GFPve_pos  0.7132903856
## dCA2_Mpos_TGCGATATCTTGATTC-1dCA2_MI_GFPve_pos  0.0983115763
## dCA2_Mpos_TGCGGCATCACTAGCA-1dCA2_MI_GFPve_pos  1.4126639289
## dCA2_Mpos_TGCGGCATCAGAATAG-1dCA2_MI_GFPve_pos  0.3314609749
## dCA2_Mpos_TGCTGAACACACCAGC-1dCA2_MI_GFPve_pos -1.3319547253
## dCA2_Mpos_TGCTTCGTCCATATGG-1dCA2_MI_GFPve_pos -1.2728159505
## dCA2_Mpos_TGGAACTTCAAGCGTT-1dCA2_MI_GFPve_pos -0.9999030906
## dCA2_Mpos_TGGAGGACAATGAACA-1dCA2_MI_GFPve_pos  1.0362836045
## dCA2_Mpos_TGGATCACATGTCGTA-1dCA2_MI_GFPve_pos  1.8653788490
## dCA2_Mpos_TGGCGTGAGGCTAAAT-1dCA2_MI_GFPve_pos  0.6546686215
## dCA2_Mpos_TGGGATTTCAACGTGT-1dCA2_MI_GFPve_pos  0.0022854057
## dCA2_Mpos_TGGGCGTTCCTCTCGA-1dCA2_MI_GFPve_pos  1.3900397939
## dCA2_Mpos_TGGGCTGCACCGTCTT-1dCA2_MI_GFPve_pos  1.5201675815
## dCA2_Mpos_TGGGCTGGTGACCGAA-1dCA2_MI_GFPve_pos  1.2682291192
## dCA2_Mpos_TGGGCTGGTTCCTTGC-1dCA2_MI_GFPve_pos  0.9585030002
## dCA2_Mpos_TGGGTTAGTCCCTAAA-1dCA2_MI_GFPve_pos  0.7361759466
## dCA2_Mpos_TGGTAGTGTCCACAGC-1dCA2_MI_GFPve_pos  0.5915926022
## dCA2_Mpos_TGGTTAGCACGCGCAT-1dCA2_MI_GFPve_pos  1.8274698180
## dCA2_Mpos_TGGTTAGCAGGCGAAT-1dCA2_MI_GFPve_pos  0.7210652155
## dCA2_Mpos_TGGTTAGGTTAGAAAC-1dCA2_MI_GFPve_pos  0.4191633982
## dCA2_Mpos_TGGTTAGTCCTTGGAA-1dCA2_MI_GFPve_pos  2.1254122180
```

```
## dCA2_Mpos_TGGTTAGTCGTGGAAG-1dCA2_MI_GFPve_pos -0.2082914787
## dCA2_Mpos_TGTAACGCATATCTCT-1dCA2_MI_GFPve_pos  1.9985657853
## dCA2_Mpos_TGTAACGGTGGATCAG-1dCA2_MI_GFPve_pos -0.9310370761
## dCA2_Mpos_TGTAAGCAGTAGTCTC-1dCA2_MI_GFPve_pos  1.3535630626
## dCA2_Mpos_TGTAAGCGTAGCCCTG-1dCA2_MI_GFPve_pos  2.1479742450
## dCA2_Mpos_TGTACAGCATGGAGAC-1dCA2_MI_GFPve_pos  2.0170489473
## dCA2_Mpos_TGTAGACGTCGCATGC-1dCA2_MI_GFPve_pos  1.1921766919
## dCA2_Mpos_TGTCAGAGTTTACTTC-1dCA2_MI_GFPve_pos  1.4181713981
## dCA2_Mpos_TGTCCACGTCATATGC-1dCA2_MI_GFPve_pos  0.5018543822
## dCA2_Mpos_TGTCCACTCTTGGTCC-1dCA2_MI_GFPve_pos  1.2259516639
## dCA2_Mpos_TGTCCTGAGAATTCAG-1dCA2_MI_GFPve_pos  1.3196976108
## dCA2_Mpos_TGTCCTGCACCATTCC-1dCA2_MI_GFPve_pos  2.2778069896
## dCA2_Mpos_TGTCCTGGTACCCGCA-1dCA2_MI_GFPve_pos  1.2048354072
## dCA2_Mpos_TGTGAGTAGCATCTTG-1dCA2_MI_GFPve_pos  1.3054499549
## dCA2_Mpos_TGTGAGTCACATACGT-1dCA2_MI_GFPve_pos -0.9973014670
## dCA2_Mpos_TGTGCGGGTCATATGC-1dCA2_MI_GFPve_pos  2.0719885749
## dCA2_Mpos_TGTGGCGTCTGTGCGG-1dCA2_MI_GFPve_pos  0.1039889136
## dCA2_Mpos_TGTTACTTCTATTCGT-1dCA2_MI_GFPve_pos  1.3987767619
## dCA2_Mpos_TGTTCCGGTACTCGCG-1dCA2_MI_GFPve_pos  1.9261877460
## dCA2_Mpos_TGTTCCGTCCATTGGA-1dCA2_MI_GFPve_pos -1.0656263905
## dCA2_Mpos_TTACAGGTCTATTGTC-1dCA2_MI_GFPve_pos  0.7739150447
## dCA2_Mpos_TTACCGCCACACTTAG-1dCA2_MI_GFPve_pos  0.9046596808
## dCA2_Mpos_TTACCGCTCGGAATGG-1dCA2_MI_GFPve_pos  2.0734791679
## dCA2_Mpos_TTACGCCTCCCGAACG-1dCA2_MI_GFPve_pos -0.4404865580
## dCA2_Mpos_TTACGTTAGAAAGTCT-1dCA2_MI_GFPve_pos  2.0904681606
## dCA2_Mpos_TTAGGCAAGCATGCAG-1dCA2_MI_GFPve_pos  0.7910761160
## dCA2_Mpos_TTAGGCAGTGACAACG-1dCA2_MI_GFPve_pos  1.0992585940
## dCA2_Mpos_TTAGGGTGTAGTCTTG-1dCA2_MI_GFPve_pos  0.6339270992
## dCA2_Mpos_TTAGGGTTCTGAGGCC-1dCA2_MI_GFPve_pos  2.4416816158
## dCA2_Mpos_TTAGTCTAGTCAGCCC-1dCA2_MI_GFPve_pos  1.7210159225
## dCA2_Mpos_TTATTGCCAACACACT-1dCA2_MI_GFPve_pos -0.3452027219
## dCA2_Mpos_TTCAATCCAGGAAGTC-1dCA2_MI_GFPve_pos  1.4200522823
## dCA2_Mpos_TTCAATCGTGGCATCC-1dCA2_MI_GFPve_pos  2.0704885644
## dCA2_Mpos_TTCACGCAGACCAAGC-1dCA2_MI_GFPve_pos  1.9132174176
## dCA2_Mpos_TTCACGCCAACAGCTT-1dCA2_MI_GFPve_pos  2.4196476859
## dCA2_Mpos_TTCACGCGTTATGTGC-1dCA2_MI_GFPve_pos  1.3775325937
## dCA2_Mpos_TTCACGCTCCGATAGT-1dCA2_MI_GFPve_pos  0.8733330292
## dCA2_Mpos_TTCATGTCAATCACGT-1dCA2_MI_GFPve_pos  0.8388636631
## dCA2_Mpos_TTCATGTTCCGCTGTT-1dCA2_MI_GFPve_pos -0.1864215213
## dCA2_Mpos_TTCCAATCAGCGTTGC-1dCA2_MI_GFPve_pos  1.3972870034
## dCA2_Mpos_TTCCAATTCATGAGAA-1dCA2_MI_GFPve_pos  1.8028292579
## dCA2_Mpos_TTCCACGAGAGACAAG-1dCA2_MI_GFPve_pos  1.1426408214
## dCA2_Mpos_TTCCACGTCCAGGACC-1dCA2_MI_GFPve_pos -0.0854163247
## dCA2_Mpos_TTCCGGTTCACGATCA-1dCA2_MI_GFPve_pos  1.8532624168
## dCA2_Mpos_TTCCTTCAGAGAGCGG-1dCA2_MI_GFPve_pos  1.1496001405
## dCA2_Mpos_TTCGCTGAGCCATTCA-1dCA2_MI_GFPve_pos  0.7178153319
## dCA2_Mpos_TTCGGTCCAAAGACTA-1dCA2_MI_GFPve_pos  1.0174980683
## dCA2_Mpos_TTCGGTCCACGACCTG-1dCA2_MI_GFPve_pos  0.5107114834
## dCA2_Mpos_TTCTAACTCCATTCGC-1dCA2_MI_GFPve_pos  0.8709611816
## dCA2_Mpos_TTCTCTCTCGTAATGC-1dCA2_MI_GFPve_pos -0.3389531749
## dCA2_Mpos_TTCTTCCCAACCTAAC-1dCA2_MI_GFPve_pos  1.1251093072
## dCA2_Mpos_TTCTTCCTCGGCATCG-1dCA2_MI_GFPve_pos  2.3738753719
## dCA2_Mpos_TTGAACGGTTACGCCG-1dCA2_MI_GFPve_pos  0.9159024281
## dCA2_Mpos_TTGAGTGCACAGAGCA-1dCA2_MI_GFPve_pos -0.4902606684
## dCA2_Mpos_TTGATGGAGTTGCGCC-1dCA2_MI_GFPve_pos  2.1385843677
## dCA2_Mpos_TTGCATTAGAAGCCAC-1dCA2_MI_GFPve_pos  2.0954900903
## dCA2_Mpos_TTGCCTGAGCTCCACG-1dCA2_MI_GFPve_pos  1.6241480035
## dCA2_Mpos_TTGCCTGCACCGAATT-1dCA2_MI_GFPve_pos  0.3614172858
## dCA2_Mpos_TTGCGTCAGACCATGG-1dCA2_MI_GFPve_pos  2.3527686519
## dCA2_Mpos_TTGGATGAGTCGAAAT-1dCA2_MI_GFPve_pos  0.7312380594
## dCA2_Mpos_TTGGGATCAATCGAAA-1dCA2_MI_GFPve_pos  1.2001806421
## dCA2_Mpos_TTGGGATTCAAGTCGT-1dCA2_MI_GFPve_pos  1.4296443385
## dCA2_Mpos_TTGGGCGAGTCACGAG-1dCA2_MI_GFPve_pos  2.1958279533
## dCA2_Mpos_TTGGGTACAAGAGGCT-1dCA2_MI_GFPve_pos  0.7333375138
## dCA2_Mpos_TTGGGTAGTAACCCGC-1dCA2_MI_GFPve_pos  0.7118619007
## dCA2_Mpos_TTGGGTAGTCTACGTA-1dCA2_MI_GFPve_pos  0.6504807395
## dCA2_Mpos_TTGGTTTCATACCAGT-1dCA2_MI_GFPve_pos  2.5186810416
## dCA2_Mpos_TTGGTTTGTAGCGCCT-1dCA2_MI_GFPve_pos  2.0574372930
## dCA2_Mpos_TTGTGGATCGTGTCAA-1dCA2_MI_GFPve_pos  0.6061202807
## dCA2_Mpos_TTGTTGTAGACATCAA-1dCA2_MI_GFPve_pos  1.7781239671
```

```
## dCA2_Mpos_TTTACCAAGGTTAAAC-1dCA2_MI_GFPve_pos  0.0228985545
## dCA2_Mpos_TTTACCAGTCGATTAC-1dCA2_MI_GFPve_pos  0.4425894541
## dCA2_Mpos_TTTACGTTCGCACGAC-1dCA2_MI_GFPve_pos  0.1555025389
## dCA2_Mpos_TTTACTGTCCATAGGT-1dCA2_MI_GFPve_pos  0.7885292691
## dCA2_Mpos_TTTAGTCCACATATGC-1dCA2_MI_GFPve_pos  0.5920669657
## dCA2_Mpos_TTTCACAAGCTGAGCA-1dCA2_MI_GFPve_pos  0.3209119064
## dCA2_Mpos_TTTCACAAGTTAGTGA-1dCA2_MI_GFPve_pos  0.6997822446
## dCA2_Mpos_TTTCACACAAACCACT-1dCA2_MI_GFPve_pos  1.8904117269
## dCA2_Mpos_TTTCAGTGTATTGGCT-1dCA2_MI_GFPve_pos  2.0325357837
## dCA2_Mpos_TTTCAGTTCCCTCATG-1dCA2_MI_GFPve_pos  1.7229791803
## dCA2_Mpos_TTTCATGAGCACGTCC-1dCA2_MI_GFPve_pos -0.9824997264
## dCA2_Mpos_TTTCATGCAACGATTC-1dCA2_MI_GFPve_pos  0.4375721735
## dCA2_Mpos_TTTCATGCAGCATACT-1dCA2_MI_GFPve_pos -0.6364023166
## dCA2_Mpos_TTTCATGCATCGCCTT-1dCA2_MI_GFPve_pos -0.4333738762
## dCA2_Mpos_TTTCCTCCAGAGCGTA-1dCA2_MI_GFPve_pos  1.8215795678
## dCA2_Mpos_TTTCGATGTCTGTAAC-1dCA2_MI_GFPve_pos  1.8271423501
## dCA2_Mpos_TTTCGATTCTACAGGT-1dCA2_MI_GFPve_pos  0.8420590085
## dCA2_Fpos_AAAGGTATCCGATAAC-1dCA2_FI_GFPve_pos  2.1002299709
## dCA2_Fpos_AAAGTGACACAACGCC-1dCA2_FI_GFPve_pos  2.1173779887
## dCA2_Fpos_AAATGGACAATACGCT-1dCA2_FI_GFPve_pos  1.0192750615
## dCA2_Fpos_AAATGGACAGCTATAC-1dCA2_FI_GFPve_pos  1.7230101747
## dCA2_Fpos_AAATGGATCCTACTGC-1dCA2_FI_GFPve_pos  0.8407487792
## dCA2_Fpos_AACAAAGCATGCACTA-1dCA2_FI_GFPve_pos  1.3904060048
## dCA2_Fpos_AACAAAGTCGCGGTAC-1dCA2_FI_GFPve_pos  1.1216083688
## dCA2_Fpos_AACACACGTCAAGCGA-1dCA2_FI_GFPve_pos  1.6086403054
## dCA2_Fpos_AACACACGTCATAAAG-1dCA2_FI_GFPve_pos  1.1336857003
## dCA2_Fpos_AACCATGCAATAGAGT-1dCA2_FI_GFPve_pos  1.6872200889
## dCA2_Fpos_AACCATGGTGTTGACT-1dCA2_FI_GFPve_pos  0.7668971342
## dCA2_Fpos_AACCTTTTCTGCTTAT-1dCA2_FI_GFPve_pos  1.9332277698
## dCA2_Fpos_AACGAAAGTTCAGTAC-1dCA2_FI_GFPve_pos  1.8093968553
## dCA2_Fpos_AAGACAAGTATCGCTA-1dCA2_FI_GFPve_pos -1.8261866647
## dCA2_Fpos_AAGACTCGTGTCTTCC-1dCA2_FI_GFPve_pos -1.3197631913
## dCA2_Fpos_AAGCGTTGTTGCATGT-1dCA2_FI_GFPve_pos  2.3300537986
## dCA2_Fpos_AATCGACGTTAGTTCG-1dCA2_FI_GFPve_pos  1.8187379760
## dCA2_Fpos_AATTCCTGTCTCGACG-1dCA2_FI_GFPve_pos  2.0678979081
## dCA2_Fpos_AATTTCCGTACTAACC-1dCA2_FI_GFPve_pos  1.0506932658
## dCA2_Fpos_ACAAAGACAAGAATGT-1dCA2_FI_GFPve_pos  1.5118104142
## dCA2_Fpos_ACAAGCTCAATCTCTT-1dCA2_FI_GFPve_pos  0.7242484612
## dCA2_Fpos_ACACAGTAGACTCTAC-1dCA2_FI_GFPve_pos -0.3529756444
## dCA2_Fpos_ACACTGAGTCTGCGCA-1dCA2_FI_GFPve_pos -0.2088733095
## dCA2_Fpos_ACAGAAACAGAAATCA-1dCA2_FI_GFPve_pos -1.4650534468
## dCA2_Fpos_ACAGAAAGTTGCGAAG-1dCA2_FI_GFPve_pos  0.1038921079
## dCA2_Fpos_ACAGCCGGTCATTCCC-1dCA2_FI_GFPve_pos  1.6607929391
## dCA2_Fpos_ACATCCCTCGCAAGAG-1dCA2_FI_GFPve_pos  1.3426501674
## dCA2_Fpos_ACCACAAGTGTTCGAT-1dCA2_FI_GFPve_pos  1.6668783349
## dCA2_Fpos_ACCACAAGTTTCCAAG-1dCA2_FI_GFPve_pos -0.0868635702
## dCA2_Fpos_ACCCAAACAGTCTACA-1dCA2_FI_GFPve_pos  1.9027429742
## dCA2_Fpos_ACCCAAACATTGCTTT-1dCA2_FI_GFPve_pos  0.6283695621
## dCA2_Fpos_ACCCTTGGTCAAAGTA-1dCA2_FI_GFPve_pos -0.5223034697
## dCA2_Fpos_ACCTACCAGTAACAGT-1dCA2_FI_GFPve_pos  0.8296880645
## dCA2_Fpos_ACCTGAACACGCACCA-1dCA2_FI_GFPve_pos  2.3623251838
## dCA2_Fpos_ACGATGTCAGGTTTAC-1dCA2_FI_GFPve_pos  1.2146747035
## dCA2_Fpos_ACGTTCCTCTTCGGAA-1dCA2_FI_GFPve_pos  0.7008640689
## dCA2_Fpos_ACTATCTGTACTGCGC-1dCA2_FI_GFPve_pos  1.5044748706
## dCA2_Fpos_ACTCCCAAGAATCGTA-1dCA2_FI_GFPve_pos  1.3638458175
## dCA2_Fpos_ACTGATGCAACGATTC-1dCA2_FI_GFPve_pos -0.0798455852
## dCA2_Fpos_ACTGCAAGTTGAAGTA-1dCA2_FI_GFPve_pos -0.1101400631
## dCA2_Fpos_ACTGTGATCCGTTGAA-1dCA2_FI_GFPve_pos  2.7234759254
## dCA2_Fpos_ACTTATCAGGAGGTTC-1dCA2_FI_GFPve_pos  1.2728718442
## dCA2_Fpos_ACTTCCGGTTGCAACT-1dCA2_FI_GFPve_pos  1.3842520637
## dCA2_Fpos_ACTTTCAGTGTTGCCG-1dCA2_FI_GFPve_pos  0.8107230586
## dCA2_Fpos_ACTTTGTCAGGAGGTT-1dCA2_FI_GFPve_pos  1.6569996995
## dCA2_Fpos_AGAACAAAGTCGGCAA-1dCA2_FI_GFPve_pos  0.8428409619
## dCA2_Fpos_AGACAAAAGACATAAC-1dCA2_FI_GFPve_pos  1.1898411435
## dCA2_Fpos_AGACAGGCACAGCCTG-1dCA2_FI_GFPve_pos  2.3002149982
## dCA2_Fpos_AGACAGGGTCCCTGTT-1dCA2_FI_GFPve_pos  1.4151334686
## dCA2_Fpos_AGATCCAGTTTGGGTT-1dCA2_FI_GFPve_pos  2.3598492068
## dCA2_Fpos_AGATGAAAGTATGATG-1dCA2_FI_GFPve_pos  1.8648480100
## dCA2_Fpos_AGCATCACAGGAATCG-1dCA2_FI_GFPve_pos  1.6280671281
## dCA2_Fpos_AGCGCTGAGTTATGGA-1dCA2_FI_GFPve_pos  0.2578654063
```

```
## dCA2_Fpos_AGCTCAAAGGTTGGTG-1dCA2_FI_GFPve_pos  1.4779426974
## dCA2_Fpos_AGGACGAGTCCTACAA-1dCA2_FI_GFPve_pos  2.0017980260
## dCA2_Fpos_AGGCTGCTCCCATTCG-1dCA2_FI_GFPve_pos  1.0022942943
## dCA2_Fpos_AGGGAGTTCTTACCAT-1dCA2_FI_GFPve_pos  1.5415198726
## dCA2_Fpos_AGGGCCTTCCGCCTAT-1dCA2_FI_GFPve_pos -0.2242081242
## dCA2_Fpos_AGGTAGGTCAAGAGTA-1dCA2_FI_GFPve_pos  0.6197590751
## dCA2_Fpos_AGGTCTACAGCTACTA-1dCA2_FI_GFPve_pos  0.1230323592
## dCA2_Fpos_AGGTGTTGTAGACAGC-1dCA2_FI_GFPve_pos  1.8100953025
## dCA2_Fpos_AGGTTACAGTATTAGG-1dCA2_FI_GFPve_pos  2.2618927879
## dCA2_Fpos_AGTACCATCCTACCGT-1dCA2_FI_GFPve_pos  0.4477372092
## dCA2_Fpos_AGTACTGCATATACCG-1dCA2_FI_GFPve_pos  1.9239604396
## dCA2_Fpos_AGTACTGTCTTGGCTC-1dCA2_FI_GFPve_pos  1.8995997829
## dCA2_Fpos_AGTCTCCCAAGACCGA-1dCA2_FI_GFPve_pos -3.6070101338
## dCA2_Fpos_AGTGACTAGGAGATAG-1dCA2_FI_GFPve_pos  0.0551072468
## dCA2_Fpos_AGTGTTGCAACTCCCT-1dCA2_FI_GFPve_pos  2.3730695171
## dCA2_Fpos_AGTTAGCGTAGAGGAA-1dCA2_FI_GFPve_pos  1.2882192058
## dCA2_Fpos_AGTTCCCCACACGGAA-1dCA2_FI_GFPve_pos  2.3684194011
## dCA2_Fpos_AGTTCGAGTTTGGAGG-1dCA2_FI_GFPve_pos  2.1505052966
## dCA2_Fpos_ATACCTTAGGCACGAT-1dCA2_FI_GFPve_pos  1.9515578670
## dCA2_Fpos_ATACTTCGTACGTGTT-1dCA2_FI_GFPve_pos  2.2425923270
## dCA2_Fpos_ATCCTATAGGTTCTTG-1dCA2_FI_GFPve_pos  2.0462201757
## dCA2_Fpos_ATCGATGCATGCCGAC-1dCA2_FI_GFPve_pos  0.6266260070
## dCA2_Fpos_ATCGCCTCAATAGTCC-1dCA2_FI_GFPve_pos  1.3918073100
## dCA2_Fpos_ATCTCTAAGGACATCG-1dCA2_FI_GFPve_pos  1.9094101114
## dCA2_Fpos_ATGAAAGGTACTGGGA-1dCA2_FI_GFPve_pos  1.1186141891
## dCA2_Fpos_ATGACCACACCATAAC-1dCA2_FI_GFPve_pos  2.6904280109
## dCA2_Fpos_ATGAGTCTCCATTGTT-1dCA2_FI_GFPve_pos  1.6158664149
## dCA2_Fpos_ATGGATCAGTCCTGTA-1dCA2_FI_GFPve_pos  1.9408571643
## dCA2_Fpos_ATGTCTTTCATTACGG-1dCA2_FI_GFPve_pos  1.5685974282
## dCA2_Fpos_ATTACCTCACTTGTCC-1dCA2_FI_GFPve_pos  2.6724672240
## dCA2_Fpos_ATTCTACAGGGTTAGC-1dCA2_FI_GFPve_pos -1.7845233756
## dCA2_Fpos_ATTGTTCCACGCGCAT-1dCA2_FI_GFPve_pos  0.2936147374
## dCA2_Fpos_ATTTACCAGGTAGTCG-1dCA2_FI_GFPve_pos  0.9427834672
## dCA2_Fpos_ATTTCACGTCTTACTT-1dCA2_FI_GFPve_pos -1.3960508185
## dCA2_Fpos_ATTTCTGTCAACCTTT-1dCA2_FI_GFPve_pos -0.1438790190
## dCA2_Fpos_CAACAGTTCTCATAGG-1dCA2_FI_GFPve_pos  2.6050753516
## dCA2_Fpos_CAACCTCAGACTAAGT-1dCA2_FI_GFPve_pos -0.0583709853
## dCA2_Fpos_CAACCTCAGTAGACCG-1dCA2_FI_GFPve_pos  0.7547417206
## dCA2_Fpos_CAAGGGAGTCGTTCAA-1dCA2_FI_GFPve_pos  1.0052198691
## dCA2_Fpos_CAATGACTCTGGGCAC-1dCA2_FI_GFPve_pos  2.3609960002
## dCA2_Fpos_CAATTTCAGCCTGTGC-1dCA2_FI_GFPve_pos  1.5920761747
## dCA2_Fpos_CACACAATCCAACACA-1dCA2_FI_GFPve_pos  2.2977135104
## dCA2_Fpos_CACTAAGAGGCTCAAG-1dCA2_FI_GFPve_pos  1.7536978645
## dCA2_Fpos_CACTAAGCACTTGTGA-1dCA2_FI_GFPve_pos -0.3218642252
## dCA2_Fpos_CACTGGGGTGGATTTC-1dCA2_FI_GFPve_pos  1.6068745774
## dCA2_Fpos_CACTGTCGTATGGAAT-1dCA2_FI_GFPve_pos  1.3770205898
## dCA2_Fpos_CACTGTCTCTACCCAC-1dCA2_FI_GFPve_pos  1.2304174823
## dCA2_Fpos_CAGATACCATGGCGCT-1dCA2_FI_GFPve_pos  1.7995660228
## dCA2_Fpos_CAGATTGCACTCCCTA-1dCA2_FI_GFPve_pos -1.1441667157
## dCA2_Fpos_CAGCACGCATACAGCT-1dCA2_FI_GFPve_pos -0.2054525393
## dCA2_Fpos_CAGGCCAGTGTGCTTA-1dCA2_FI_GFPve_pos  2.5694813651
## dCA2_Fpos_CAGGCCATCGCGGTAC-1dCA2_FI_GFPve_pos  1.3736103696
## dCA2_Fpos_CAGGTATCACCCAATA-1dCA2_FI_GFPve_pos  0.6328414005
## dCA2_Fpos_CAGGTATTCTACACTT-1dCA2_FI_GFPve_pos  0.9697340054
## dCA2_Fpos_CAGTTAGAGGCCTGAA-1dCA2_FI_GFPve_pos  1.1781817598
## dCA2_Fpos_CAGTTCCAGCCAGTAG-1dCA2_FI_GFPve_pos  1.4957848710
## dCA2_Fpos_CAGTTCCGTGAAGCGT-1dCA2_FI_GFPve_pos  1.5966627521
## dCA2_Fpos_CATCAAGAGAGTTGCG-1dCA2_FI_GFPve_pos  2.3096408767
## dCA2_Fpos_CATCAAGAGATGGTAT-1dCA2_FI_GFPve_pos  1.9665715617
## dCA2_Fpos_CATCCACCAGTCTACA-1dCA2_FI_GFPve_pos  2.1923382205
## dCA2_Fpos_CATCCACCATAGTCAC-1dCA2_FI_GFPve_pos -1.3803529816
## dCA2_Fpos_CATCCGTTCAAGTCGT-1dCA2_FI_GFPve_pos  2.4224555415
## dCA2_Fpos_CATGAGTCACCATATG-1dCA2_FI_GFPve_pos  0.4603730065
## dCA2_Fpos_CATTCCGCATTGCCGG-1dCA2_FI_GFPve_pos  1.4045580548
## dCA2_Fpos_CATTGCCAGAACAAGG-1dCA2_FI_GFPve_pos  1.4816534442
## dCA2_Fpos_CATTGCCTCCGATGTA-1dCA2_FI_GFPve_pos  1.0867304725
## dCA2_Fpos_CATTGTTAGGCGACAT-1dCA2_FI_GFPve_pos  2.2355363292
## dCA2_Fpos_CCACCATAGTAGCATA-1dCA2_FI_GFPve_pos -0.8229097205
## dCA2_Fpos_CCACGAGAGACGAAGA-1dCA2_FI_GFPve_pos  2.3668115062
## dCA2_Fpos_CCCGGAAGTTAGAAGT-1dCA2_FI_GFPve_pos  0.1027545813
```

```
## dCA2_Fpos_CCCTAACCACCTTCGT-1dCA2_FI_GFPve_pos  1.0754375381
## dCA2_Fpos_CCCTTAGAGCAAACAT-1dCA2_FI_GFPve_pos  1.7689104003
## dCA2_Fpos_CCGAACGGTGGCAGAT-1dCA2_FI_GFPve_pos  2.4967191142
## dCA2_Fpos_CCGATCTTCTAGTGTG-1dCA2_FI_GFPve_pos -0.6477455574
## dCA2_Fpos_CCGGTGACACCACTGG-1dCA2_FI_GFPve_pos  2.0858688277
## dCA2_Fpos_CCGGTGATCGTCCATC-1dCA2_FI_GFPve_pos  2.3652591628
## dCA2_Fpos_CCGTAGGAGTTGCCCG-1dCA2_FI_GFPve_pos  1.0953717751
## dCA2_Fpos_CCTAACCAGACCATTC-1dCA2_FI_GFPve_pos -0.1560112315
## dCA2_Fpos_CCTATCGAGACTGAGC-1dCA2_FI_GFPve_pos  1.1997910661
## dCA2_Fpos_CCTATCGTCCTATTTG-1dCA2_FI_GFPve_pos  2.3042461795
## dCA2_Fpos_CCTCAACCATTATGCG-1dCA2_FI_GFPve_pos  1.5082902831
## dCA2_Fpos_CCTCATGTCAGAGTGG-1dCA2_FI_GFPve_pos  1.1766875905
## dCA2_Fpos_CCTTGTGGTAACATCC-1dCA2_FI_GFPve_pos  1.9037368221
## dCA2_Fpos_CCTTGTGTCTGTCCGT-1dCA2_FI_GFPve_pos  1.3082752151
## dCA2_Fpos_CCTTTGGTCAAAGACA-1dCA2_FI_GFPve_pos  1.1997253818
## dCA2_Fpos_CGAAGTTAGGGATCAC-1dCA2_FI_GFPve_pos  1.8630024118
## dCA2_Fpos_CGAATTGAGCACGTCC-1dCA2_FI_GFPve_pos  2.3413996619
## dCA2_Fpos_CGAGGCTAGCATGGGT-1dCA2_FI_GFPve_pos  2.7036483211
## dCA2_Fpos_CGAGTTAAGAGGTCAC-1dCA2_FI_GFPve_pos  0.0525621583
## dCA2_Fpos_CGATGGCCAGAGATGC-1dCA2_FI_GFPve_pos -0.3816732841
## dCA2_Fpos_CGATGGCTCTTCTGTA-1dCA2_FI_GFPve_pos  1.1143184823
## dCA2_Fpos_CGCATAAGTTGGCTAT-1dCA2_FI_GFPve_pos  0.9970949811
## dCA2_Fpos_CGCCAGATCTTGGATG-1dCA2_FI_GFPve_pos  1.3780877513
## dCA2_Fpos_CGGACACGTATCCTCC-1dCA2_FI_GFPve_pos  1.4905012769
## dCA2_Fpos_CGGACACTCTCTATGT-1dCA2_FI_GFPve_pos  0.0995282379
## dCA2_Fpos_CGGGTCAAGTCTAGCT-1dCA2_FI_GFPve_pos  1.6035381478
## dCA2_Fpos_CGTAATGAGCCAGACA-1dCA2_FI_GFPve_pos  2.0805654449
## dCA2_Fpos_CGTCCATAGGCTCCCA-1dCA2_FI_GFPve_pos  2.4532902164
## dCA2_Fpos_CGTCCATTCGAACTCA-1dCA2_FI_GFPve_pos  1.8413567466
## dCA2_Fpos_CGTGTCTTCAATCAGC-1dCA2_FI_GFPve_pos  1.0717985553
## dCA2_Fpos_CGTTGGGCACAGCCAC-1dCA2_FI_GFPve_pos  1.1137906236
## dCA2_Fpos_CTAACCCAGGGTTAGC-1dCA2_FI_GFPve_pos  0.7547399921
## dCA2_Fpos_CTAACCCGTCGGCTAC-1dCA2_FI_GFPve_pos -0.4809715825
## dCA2_Fpos_CTAACCCGTTTCTTAC-1dCA2_FI_GFPve_pos  1.8259857816
## dCA2_Fpos_CTAACTTTCAGCTTCC-1dCA2_FI_GFPve_pos  1.9378902835
## dCA2_Fpos_CTAAGTGCAAAGACGC-1dCA2_FI_GFPve_pos -0.6444114524
## dCA2_Fpos_CTACAGATCATTGCGA-1dCA2_FI_GFPve_pos  1.2872763795
## dCA2_Fpos_CTACATTGTTATGACC-1dCA2_FI_GFPve_pos  2.3154811782
## dCA2_Fpos_CTACCCACATCGCTGG-1dCA2_FI_GFPve_pos  0.9747047943
## dCA2_Fpos_CTAGGTAAGCATATGA-1dCA2_FI_GFPve_pos  0.1430426535
## dCA2_Fpos_CTATAGGAGTAGGGTC-1dCA2_FI_GFPve_pos  1.5779685897
## dCA2_Fpos_CTCAACCAGGCACAAC-1dCA2_FI_GFPve_pos  1.8415586871
## dCA2_Fpos_CTCAATTGTGAGTTGG-1dCA2_FI_GFPve_pos  2.4279317779
## dCA2_Fpos_CTCAGTCGTGCCGTAC-1dCA2_FI_GFPve_pos  1.6963035983
## dCA2_Fpos_CTCATCGTCTAGAACC-1dCA2_FI_GFPve_pos  1.5197606010
## dCA2_Fpos_CTCATGCTCCAATCCC-1dCA2_FI_GFPve_pos  0.5089057905
## dCA2_Fpos_CTCCAACAGGTTACAA-1dCA2_FI_GFPve_pos  1.9794909877
## dCA2_Fpos_CTCCCTCTCAGCCTTC-1dCA2_FI_GFPve_pos  1.3801070375
## dCA2_Fpos_CTCCGATAGGGACACT-1dCA2_FI_GFPve_pos  0.9704807920
## dCA2_Fpos_CTCGAGGGTCTTGCTC-1dCA2_FI_GFPve_pos  2.3936722202
## dCA2_Fpos_CTCGAGGTCCGTCACT-1dCA2_FI_GFPve_pos  0.7222575468
## dCA2_Fpos_CTCTCGATCATTGCTT-1dCA2_FI_GFPve_pos  1.4806971473
## dCA2_Fpos_CTCTGGTAGTCGTTAC-1dCA2_FI_GFPve_pos  0.6958199662
## dCA2_Fpos_CTGGCAGGTATCTCTT-1dCA2_FI_GFPve_pos  2.1996374053
## dCA2_Fpos_CTGTAGACAAGTTCCA-1dCA2_FI_GFPve_pos  2.5640714091
## dCA2_Fpos_CTGTAGACACGGCTAC-1dCA2_FI_GFPve_pos  1.0861594600
## dCA2_Fpos_CTGTATTGTCGTTGCG-1dCA2_FI_GFPve_pos  0.9732350630
## dCA2_Fpos_CTTACCGCAGCTCTGG-1dCA2_FI_GFPve_pos  1.0988244337
## dCA2_Fpos_CTTCAATGTATGATCC-1dCA2_FI_GFPve_pos  2.0925593299
## dCA2_Fpos_CTTCTCTGTGCCGGTT-1dCA2_FI_GFPve_pos  1.2977420015
## dCA2_Fpos_CTTGAGAGTGTTACAC-1dCA2_FI_GFPve_pos  1.4292833728
## dCA2_Fpos_CTTGATTCACACCTTC-1dCA2_FI_GFPve_pos  0.6240398807
## dCA2_Fpos_GAAATGAGTACTCAAC-1dCA2_FI_GFPve_pos  1.6917352599
## dCA2_Fpos_GAACTGTAGACTGTTC-1dCA2_FI_GFPve_pos  0.9210030956
## dCA2_Fpos_GAACTGTTCATTTCCA-1dCA2_FI_GFPve_pos  2.6375543994
## dCA2_Fpos_GAAGGACTCAGCGTCG-1dCA2_FI_GFPve_pos  1.7784882707
## dCA2_Fpos_GAAGGGTCAAAGCTAA-1dCA2_FI_GFPve_pos  0.9328067822
## dCA2_Fpos_GACAGCCTCTGCGGAC-1dCA2_FI_GFPve_pos  2.4709777755
## dCA2_Fpos_GACCCAGCATACCAGT-1dCA2_FI_GFPve_pos  1.7531965894
## dCA2_Fpos_GACCTTCAGCAGCCCT-1dCA2_FI_GFPve_pos  1.5423076076
```

```
## dCA2_Fpos_GACTATGCACGTCGGT-1dCA2_FI_GFPve_pos -0.0809382426
## dCA2_Fpos_GACTCTCGTCTTCATT-1dCA2_FI_GFPve_pos -0.2765898484
## dCA2_Fpos_GACTTCCTCAATGCAC-1dCA2_FI_GFPve_pos  2.0323270482
## dCA2_Fpos_GAGAAATGTTTGTGGT-1dCA2_FI_GFPve_pos -1.0286166745
## dCA2_Fpos_GAGACCCTCAGCTTGA-1dCA2_FI_GFPve_pos  1.1727859897
## dCA2_Fpos_GAGACTTGTCTATGAC-1dCA2_FI_GFPve_pos  0.6604384107
## dCA2_Fpos_GAGGCCTAGGCTATCT-1dCA2_FI_GFPve_pos  0.8778073711
## dCA2_Fpos_GAGTCATAGCCGTTAT-1dCA2_FI_GFPve_pos -0.7013284045
## dCA2_Fpos_GAGTCTAGTTAGAAAC-1dCA2_FI_GFPve_pos  0.6939041537
## dCA2_Fpos_GATAGAACACGGCACT-1dCA2_FI_GFPve_pos  1.6633477134
## dCA2_Fpos_GATAGCTCATTCATCT-1dCA2_FI_GFPve_pos  2.4079015178
## dCA2_Fpos_GATCACAGTTTCTTAC-1dCA2_FI_GFPve_pos  1.5746916455
## dCA2_Fpos_GATCATGCAGGCTACC-1dCA2_FI_GFPve_pos  2.1746749801
## dCA2_Fpos_GATCCCTAGCGGGTAT-1dCA2_FI_GFPve_pos  1.7056655807
## dCA2_Fpos_GATGACTCATGTCTAG-1dCA2_FI_GFPve_pos  1.7525091094
## dCA2_Fpos_GATGCTAAGAGGCGTT-1dCA2_FI_GFPve_pos  2.5872931403
## dCA2_Fpos_GATGCTAAGTGTGTTC-1dCA2_FI_GFPve_pos  0.9733067078
## dCA2_Fpos_GATGGAGGTCAACGCC-1dCA2_FI_GFPve_pos  1.0909212155
## dCA2_Fpos_GCACATAAGCCTCAAT-1dCA2_FI_GFPve_pos  1.4752824229
## dCA2_Fpos_GCACATACAATCCTAG-1dCA2_FI_GFPve_pos -0.4020521718
## dCA2_Fpos_GCCAGCAAGCCTAACT-1dCA2_FI_GFPve_pos -0.6454334336
## dCA2_Fpos_GCCAGGTTCGAGCCTG-1dCA2_FI_GFPve_pos  0.7417578620
## dCA2_Fpos_GCCAGGTTCGATCCAA-1dCA2_FI_GFPve_pos  0.6953282279
## dCA2_Fpos_GCCATGGTCCTGCCAT-1dCA2_FI_GFPve_pos  1.1600109262
## dCA2_Fpos_GCCCGAAGTAACGATA-1dCA2_FI_GFPve_pos -0.3799084800
## dCA2_Fpos_GCCGATGTCTGTCCCA-1dCA2_FI_GFPve_pos  2.2322842998
## dCA2_Fpos_GCCTGTTTCCGGACGT-1dCA2_FI_GFPve_pos  0.3129838926
## dCA2_Fpos_GCCTGTTTCGCTCTCA-1dCA2_FI_GFPve_pos  1.8162155074
## dCA2_Fpos_GCGAGAAAGTTGTCAC-1dCA2_FI_GFPve_pos  1.3812211675
## dCA2_Fpos_GCTACCTCAAATGCTC-1dCA2_FI_GFPve_pos  2.3008601588
## dCA2_Fpos_GCTCAAAAGAGTTGCG-1dCA2_FI_GFPve_pos  1.2134157342
## dCA2_Fpos_GCTGGGTCACTCCGAG-1dCA2_FI_GFPve_pos -0.6927267390
## dCA2_Fpos_GGAGATGAGTAACGTA-1dCA2_FI_GFPve_pos  1.7710055036
## dCA2_Fpos_GGATCTAAGGGCCTCT-1dCA2_FI_GFPve_pos  2.0729753894
## dCA2_Fpos_GGATCTAGTCACTTAG-1dCA2_FI_GFPve_pos  1.8338875694
## dCA2_Fpos_GGCGTCACAGTGTGCC-1dCA2_FI_GFPve_pos  1.0687050146
## dCA2_Fpos_GGCTGTGCATAACAGA-1dCA2_FI_GFPve_pos  1.1880906743
## dCA2_Fpos_GGGACCTTCAAACCTG-1dCA2_FI_GFPve_pos  1.4792035741
## dCA2_Fpos_GGGACTCTCCTCTTTC-1dCA2_FI_GFPve_pos  1.4258396548
## dCA2_Fpos_GGGAGATGTCTTGCTC-1dCA2_FI_GFPve_pos  0.6839112562
## dCA2_Fpos_GGGATCCGTCCGGATC-1dCA2_FI_GFPve_pos  0.7504500074
## dCA2_Fpos_GGGATGACAGTCAGCC-1dCA2_FI_GFPve_pos  1.2147099895
## dCA2_Fpos_GGGCGTTAGCGGTATG-1dCA2_FI_GFPve_pos  2.3517835063
## dCA2_Fpos_GGGTAGAAGCTGACTT-1dCA2_FI_GFPve_pos  1.9461207313
## dCA2_Fpos_GGGTCACCAGACAAAT-1dCA2_FI_GFPve_pos  2.0995953006
## dCA2_Fpos_GGTAGAGAGACTTAAG-1dCA2_FI_GFPve_pos  0.8307593984
## dCA2_Fpos_GGTCTGGTCGAGAGAC-1dCA2_FI_GFPve_pos  2.2055447025
## dCA2_Fpos_GGTGATTAGGGTTAAT-1dCA2_FI_GFPve_pos  1.4775825662
## dCA2_Fpos_GGTGATTGTCAGTCCG-1dCA2_FI_GFPve_pos  0.9366182012
## dCA2_Fpos_GGTGGCTTCTGACCCT-1dCA2_FI_GFPve_pos -0.3726055997
## dCA2_Fpos_GGTGTTAAGCTTTGTG-1dCA2_FI_GFPve_pos  2.5456108970
## dCA2_Fpos_GGTTAACTCGAACGCC-1dCA2_FI_GFPve_pos  1.8360674304
## dCA2_Fpos_GGTTCTCGTTTCGTGA-1dCA2_FI_GFPve_pos -0.9707648831
## dCA2_Fpos_GTAACACCAACAGTGG-1dCA2_FI_GFPve_pos  2.1008227987
## dCA2_Fpos_GTAATCGGTCTTGTCC-1dCA2_FI_GFPve_pos  2.2071885986
## dCA2_Fpos_GTAATCGTCGACGCTG-1dCA2_FI_GFPve_pos  1.5120330972
## dCA2_Fpos_GTAATGCAGTTTCGAC-1dCA2_FI_GFPve_pos  1.7231964988
## dCA2_Fpos_GTAGGAGGTATTGAGA-1dCA2_FI_GFPve_pos  1.3552955312
## dCA2_Fpos_GTAGGTTAGCATGATA-1dCA2_FI_GFPve_pos  0.4452747029
## dCA2_Fpos_GTAGTACAGCGAAACC-1dCA2_FI_GFPve_pos  1.0776933951
## dCA2_Fpos_GTAGTACGTCATCGCG-1dCA2_FI_GFPve_pos  1.6276239080
## dCA2_Fpos_GTAGTACGTTGGGAAC-1dCA2_FI_GFPve_pos  0.4836918635
## dCA2_Fpos_GTCAAACGTTGTCCCT-1dCA2_FI_GFPve_pos  1.9175157470
## dCA2_Fpos_GTCAAACTCTCTCGCA-1dCA2_FI_GFPve_pos -1.2225782948
## dCA2_Fpos_GTCAGCGAGTTCACTG-1dCA2_FI_GFPve_pos  1.4225796384
## dCA2_Fpos_GTCCTCATCTGAATGC-1dCA2_FI_GFPve_pos  1.7964888734
## dCA2_Fpos_GTCTAGAAGGGAGGCA-1dCA2_FI_GFPve_pos  0.0109066812
## dCA2_Fpos_GTCTAGATCCGAAGGA-1dCA2_FI_GFPve_pos  0.9406372828
## dCA2_Fpos_GTCTGTCTCATGAGTC-1dCA2_FI_GFPve_pos  1.4611607713
## dCA2_Fpos_GTGATGTAGTAATTGG-1dCA2_FI_GFPve_pos  0.7207819742
```

```
## dCA2_Fpos_GTGCTTCCAGAGTAAT-1dCA2_FI_GFPve_pos  1.5981041116
## dCA2_Fpos_GTGGAGATCAGTGGGA-1dCA2_FI_GFPve_pos  1.8999069852
## dCA2_Fpos_GTGTAACAGACCAAGC-1dCA2_FI_GFPve_pos  2.0923856420
## dCA2_Fpos_GTGTGGCAGGTCTTTG-1dCA2_FI_GFPve_pos  2.4715225620
## dCA2_Fpos_GTTCTATAGGACATCG-1dCA2_FI_GFPve_pos  2.2714257163
## dCA2_Fpos_GTTGCTCGTCCAGTTA-1dCA2_FI_GFPve_pos  2.3313581866
## dCA2_Fpos_GTTGTAGAGCATCCTA-1dCA2_FI_GFPve_pos  2.2221629543
## dCA2_Fpos_GTTGTAGCAGGACGAT-1dCA2_FI_GFPve_pos  1.1436577958
## dCA2_Fpos_GTTTACTAGTTGCTGT-1dCA2_FI_GFPve_pos  0.6118084711
## dCA2_Fpos_GTTTACTTCCATGCAA-1dCA2_FI_GFPve_pos  0.8726775569
## dCA2_Fpos_TAACCAGTCGTGGAAG-1dCA2_FI_GFPve_pos  2.3997459335
## dCA2_Fpos_TAAGCCAAGAAAGTCT-1dCA2_FI_GFPve_pos  1.0735040826
## dCA2_Fpos_TAAGCCACACATGGTT-1dCA2_FI_GFPve_pos  1.2679284734
## dCA2_Fpos_TAAGCCATCATTGCTT-1dCA2_FI_GFPve_pos -1.1174092370
## dCA2_Fpos_TAAGTCGTCTCGTTTA-1dCA2_FI_GFPve_pos  0.5137369258
## dCA2_Fpos_TAATTCCGTCCAAGAG-1dCA2_FI_GFPve_pos -0.3816698271
## dCA2_Fpos_TAATTCCTCTCAACGA-1dCA2_FI_GFPve_pos -0.3265280801
## dCA2_Fpos_TACAGGTAGGGTCACA-1dCA2_FI_GFPve_pos  1.1988991422
## dCA2_Fpos_TACCCGTAGCCTAGGA-1dCA2_FI_GFPve_pos  1.3408355636
## dCA2_Fpos_TACCCGTTCACTTCTA-1dCA2_FI_GFPve_pos  2.2138333244
## dCA2_Fpos_TACCGAAGTCCGGTGT-1dCA2_FI_GFPve_pos  0.7670462650
## dCA2_Fpos_TACCGGGAGCGATGGT-1dCA2_FI_GFPve_pos  1.2217712325
## dCA2_Fpos_TACCTGCAGCGGTATG-1dCA2_FI_GFPve_pos  2.1176519317
## dCA2_Fpos_TACCTGCCAAAGTGTA-1dCA2_FI_GFPve_pos  1.4067302865
## dCA2_Fpos_TACGGTACAGAGAATT-1dCA2_FI_GFPve_pos  1.6226905507
## dCA2_Fpos_TACTTACTCAACTGGT-1dCA2_FI_GFPve_pos  1.9903339071
## dCA2_Fpos_TACTTGTAGCCATCCG-1dCA2_FI_GFPve_pos  0.2435014767
## dCA2_Fpos_TAGACCATCATTTACC-1dCA2_FI_GFPve_pos  1.7023818416
## dCA2_Fpos_TAGGTACAGATTCGAA-1dCA2_FI_GFPve_pos  2.2969064635
## dCA2_Fpos_TAGGTTGTCGACGATT-1dCA2_FI_GFPve_pos  2.8338665885
## dCA2_Fpos_TAGTGCAAGTAGCATA-1dCA2_FI_GFPve_pos  0.4650306327
## dCA2_Fpos_TATACCTAGCAGCGAT-1dCA2_FI_GFPve_pos -1.4643623906
## dCA2_Fpos_TATACCTTCATCTGTT-1dCA2_FI_GFPve_pos -0.2420162636
## dCA2_Fpos_TATCTTGCAGCAGTCC-1dCA2_FI_GFPve_pos  1.9244403762
## dCA2_Fpos_TATCTTGGTTATTCCT-1dCA2_FI_GFPve_pos  1.6008808536
## dCA2_Fpos_TATGTTCCATCGTCCT-1dCA2_FI_GFPve_pos -0.3822757083
## dCA2_Fpos_TATTGCTCATCGCTGG-1dCA2_FI_GFPve_pos  0.7419617695
## dCA2_Fpos_TATTGCTTCTCCCATG-1dCA2_FI_GFPve_pos  1.7560440225
## dCA2_Fpos_TATTGGGCAGTTGTTG-1dCA2_FI_GFPve_pos  1.9870200080
## dCA2_Fpos_TATTTCGAGCAATAAC-1dCA2_FI_GFPve_pos -1.7502427178
## dCA2_Fpos_TATTTCGTCAGCTTGA-1dCA2_FI_GFPve_pos  2.1094784660
## dCA2_Fpos_TCACAAGCAAGCTGCC-1dCA2_FI_GFPve_pos  1.1333300990
## dCA2_Fpos_TCACACCCAACAGCTT-1dCA2_FI_GFPve_pos -0.6215638595
## dCA2_Fpos_TCACGGGCAACCAACT-1dCA2_FI_GFPve_pos -0.2101335305
## dCA2_Fpos_TCAGCCTAGCCGTAAG-1dCA2_FI_GFPve_pos  1.5462720317
## dCA2_Fpos_TCAGCCTAGGCACGAT-1dCA2_FI_GFPve_pos -0.0900999385
## dCA2_Fpos_TCAGCCTAGTCAGCGA-1dCA2_FI_GFPve_pos  1.0152949614
## dCA2_Fpos_TCAGGTAAGTGATTCC-1dCA2_FI_GFPve_pos  1.3149417562
## dCA2_Fpos_TCATCCGAGATTAGAC-1dCA2_FI_GFPve_pos -0.2583350318
## dCA2_Fpos_TCATCCGTCTCAATCT-1dCA2_FI_GFPve_pos  0.8860852045
## dCA2_Fpos_TCATGAGTCTGTGTGA-1dCA2_FI_GFPve_pos  1.9761921090
## dCA2_Fpos_TCATGCCAGGTCCCTG-1dCA2_FI_GFPve_pos  1.2963534517
## dCA2_Fpos_TCATTCACATGGTGGA-1dCA2_FI_GFPve_pos  2.1776137275
## dCA2_Fpos_TCCAGAAAGACTCTAC-1dCA2_FI_GFPve_pos  1.7760590238
## dCA2_Fpos_TCCAGAAAGTAGCAAT-1dCA2_FI_GFPve_pos  0.3497059864
## dCA2_Fpos_TCCAGAACACTGCTTC-1dCA2_FI_GFPve_pos  1.2303033990
## dCA2_Fpos_TCCATGCTCGGTAACT-1dCA2_FI_GFPve_pos  0.4055895430
## dCA2_Fpos_TCCGAAATCTCGACGG-1dCA2_FI_GFPve_pos  1.4622764510
## dCA2_Fpos_TCCTAATAGTCGGGAT-1dCA2_FI_GFPve_pos  1.9507347268
## dCA2_Fpos_TCCTCGAAGTCTAACC-1dCA2_FI_GFPve_pos -1.1291649419
## dCA2_Fpos_TCCTCGAGTGTTCCAA-1dCA2_FI_GFPve_pos  1.7381864709
## dCA2_Fpos_TCCTCGATCCTTGACC-1dCA2_FI_GFPve_pos  1.4697303695
## dCA2_Fpos_TCCTTCTAGAACTTCC-1dCA2_FI_GFPve_pos  1.1287766618
## dCA2_Fpos_TCGACCTAGATGACAT-1dCA2_FI_GFPve_pos  1.0251477284
## dCA2_Fpos_TCGACGGTCTCACCCA-1dCA2_FI_GFPve_pos  1.3350164813
## dCA2_Fpos_TCGCAGGGTGGTCTTA-1dCA2_FI_GFPve_pos  2.0395579261
## dCA2_Fpos_TCGGATACATGTCGTA-1dCA2_FI_GFPve_pos  0.1118830384
## dCA2_Fpos_TCGGATATCGATTGAC-1dCA2_FI_GFPve_pos -0.2693004387
## dCA2_Fpos_TCGGGTGGTGAGCTCC-1dCA2_FI_GFPve_pos  2.0999824924
## dCA2_Fpos_TCGGTCTCACTCCCTA-1dCA2_FI_GFPve_pos -1.0018414336
```

```
## dCA2_Fpos_TCTACCGCAATCTAGC-1dCA2_FI_GFPve_pos  1.7998480720
## dCA2_Fpos_TCTATACAGGAATGTT-1dCA2_FI_GFPve_pos  2.6908876819
## dCA2_Fpos_TCTATCAGTGAGTAAT-1dCA2_FI_GFPve_pos  2.0900806112
## dCA2_Fpos_TCTCAGCTCTAACGGT-1dCA2_FI_GFPve_pos  1.9505251569
## dCA2_Fpos_TCTCCGATCACGACTA-1dCA2_FI_GFPve_pos  1.4418822450
## dCA2_Fpos_TCTGCCAGTCAAAGAT-1dCA2_FI_GFPve_pos  0.3660570604
## dCA2_Fpos_TCTGGCTGTGGAGGTT-1dCA2_FI_GFPve_pos  0.2243796361
## dCA2_Fpos_TCTTAGTCACAATGCT-1dCA2_FI_GFPve_pos  2.0659443063
## dCA2_Fpos_TCTTAGTGTCGCTGCA-1dCA2_FI_GFPve_pos  2.7567670268
## dCA2_Fpos_TCTTTGAAGCCTCTGG-1dCA2_FI_GFPve_pos  1.8218061847
## dCA2_Fpos_TGAACGTAGTCCCGAC-1dCA2_FI_GFPve_pos  1.5629677695
## dCA2_Fpos_TGAATCGAGTGCACCC-1dCA2_FI_GFPve_pos  0.4719793421
## dCA2_Fpos_TGACAGTTCGGTCTGG-1dCA2_FI_GFPve_pos  1.7761709613
## dCA2_Fpos_TGACGCGGTTGGGAAC-1dCA2_FI_GFPve_pos  2.4597823543
## dCA2_Fpos_TGAGGAGTCAAGTGGG-1dCA2_FI_GFPve_pos  1.6206734103
## dCA2_Fpos_TGAGGGAGTATCGAAA-1dCA2_FI_GFPve_pos  2.3703169746
## dCA2_Fpos_TGATCAGCAGGTCTCG-1dCA2_FI_GFPve_pos  0.4751934928
## dCA2_Fpos_TGATGCAAGTACTGGG-1dCA2_FI_GFPve_pos  0.0587533561
## dCA2_Fpos_TGATGCACAAGCGCAA-1dCA2_FI_GFPve_pos  2.0772468967
## dCA2_Fpos_TGATGGTGTTTACGAC-1dCA2_FI_GFPve_pos  0.4379785878
## dCA2_Fpos_TGATTCTTCGAAACAA-1dCA2_FI_GFPve_pos  0.2547360880
## dCA2_Fpos_TGCACGGTCCACTAGA-1dCA2_FI_GFPve_pos -1.3643072921
## dCA2_Fpos_TGCAGATAGGTGCAGT-1dCA2_FI_GFPve_pos  0.1535065768
## dCA2_Fpos_TGCAGATTCAGGGTAG-1dCA2_FI_GFPve_pos  0.8275824708
## dCA2_Fpos_TGCAGATTCTCAACCC-1dCA2_FI_GFPve_pos  1.9280685109
## dCA2_Fpos_TGCAGGCCAGCGAGTA-1dCA2_FI_GFPve_pos -1.8052002269
## dCA2_Fpos_TGCGATACACTAGGTT-1dCA2_FI_GFPve_pos  1.5123606843
## dCA2_Fpos_TGCGGGTGTGTTTACG-1dCA2_FI_GFPve_pos  2.1854600829
## dCA2_Fpos_TGCTCCATCACGATAC-1dCA2_FI_GFPve_pos  0.7926190418
## dCA2_Fpos_TGCTGAAAGTTCCATG-1dCA2_FI_GFPve_pos  2.0502706689
## dCA2_Fpos_TGCTTCGAGGTGGCTA-1dCA2_FI_GFPve_pos  0.4115590972
## dCA2_Fpos_TGCTTGCTCGGATAAA-1dCA2_FI_GFPve_pos  1.9596855563
## dCA2_Fpos_TGGAACTGTATCGCGC-1dCA2_FI_GFPve_pos -0.0366902875
## dCA2_Fpos_TGGATCAAGAGCAGAA-1dCA2_FI_GFPve_pos  1.2370809240
## dCA2_Fpos_TGGATCATCAGAATAG-1dCA2_FI_GFPve_pos -0.4626090008
## dCA2_Fpos_TGGATGTCATGACGTT-1dCA2_FI_GFPve_pos  0.1660654244
## dCA2_Fpos_TGGGAGAGTCCACATA-1dCA2_FI_GFPve_pos  0.1526154650
## dCA2_Fpos_TGGGTTAGTGGAAATT-1dCA2_FI_GFPve_pos  1.0777324957
## dCA2_Fpos_TGTACAGGTTGTCCCT-1dCA2_FI_GFPve_pos  1.1851020974
## dCA2_Fpos_TGTAGACAGTATGTAG-1dCA2_FI_GFPve_pos  1.8504429979
## dCA2_Fpos_TGTCCTGGTATCACCA-1dCA2_FI_GFPve_pos -0.1809437352
## dCA2_Fpos_TGTGATGCACGTAGTT-1dCA2_FI_GFPve_pos  2.6057150287
## dCA2_Fpos_TGTGGCGAGCATCTTG-1dCA2_FI_GFPve_pos  0.1119686418
## dCA2_Fpos_TGTGGCGTCCATTGGA-1dCA2_FI_GFPve_pos  2.0732682866
## dCA2_Fpos_TGTGTGATCCTCAGGG-1dCA2_FI_GFPve_pos  2.0415385884
## dCA2_Fpos_TGTTCCGTCACAGAGG-1dCA2_FI_GFPve_pos  0.0159326566
## dCA2_Fpos_TGTTCTAAGCTCGTGC-1dCA2_FI_GFPve_pos  2.1235630435
## dCA2_Fpos_TTACGCCCATCCTAAG-1dCA2_FI_GFPve_pos  0.6456487698
## dCA2_Fpos_TTACGTTAGCGGCTCT-1dCA2_FI_GFPve_pos  2.5115160865
## dCA2_Fpos_TTACTGTTCTGCCTGT-1dCA2_FI_GFPve_pos  2.0312839669
## dCA2_Fpos_TTAGGCACAGGAACCA-1dCA2_FI_GFPve_pos -0.3299968320
## dCA2_Fpos_TTCAATCGTCACATTG-1dCA2_FI_GFPve_pos  2.4443409366
## dCA2_Fpos_TTCATTGCAACTTGGT-1dCA2_FI_GFPve_pos -0.0240719693
## dCA2_Fpos_TTCCAATCAACATACC-1dCA2_FI_GFPve_pos  2.4195344371
## dCA2_Fpos_TTCCTCTAGGCCCAAA-1dCA2_FI_GFPve_pos  1.3448717517
## dCA2_Fpos_TTCGATTTCTCTGCTG-1dCA2_FI_GFPve_pos  1.2281968517
## dCA2_Fpos_TTCGCTGGTTCTCCTG-1dCA2_FI_GFPve_pos  1.1172568721
## dCA2_Fpos_TTCTAACCACTCTCGT-1dCA2_FI_GFPve_pos  2.0088057441
## dCA2_Fpos_TTCTAGTTCAAGCTGT-1dCA2_FI_GFPve_pos  1.2364159745
## dCA2_Fpos_TTCTTCCCATCTAACG-1dCA2_FI_GFPve_pos  1.6481170577
## dCA2_Fpos_TTCTTCCTCATGAGGG-1dCA2_FI_GFPve_pos  1.0272203607
## dCA2_Fpos_TTGACCCAGGAACGTC-1dCA2_FI_GFPve_pos  2.0741887016
## dCA2_Fpos_TTGACCCGTCGCAGTC-1dCA2_FI_GFPve_pos -1.7485394555
## dCA2_Fpos_TTGCATTTCGTCTAAG-1dCA2_FI_GFPve_pos  1.5934306306
## dCA2_Fpos_TTGCTGCTCATTGCTT-1dCA2_FI_GFPve_pos  1.6975550574
## dCA2_Fpos_TTGCTGCTCCTACTGC-1dCA2_FI_GFPve_pos  1.5036231202
## dCA2_Fpos_TTTCCTCAGTCGCCAC-1dCA2_FI_GFPve_pos  1.5868909282
## dCA2_Fpos_TTTGACTAGGATTCAA-1dCA2_FI_GFPve_pos  1.2428350372
## dCA2_Fpos_TTTGGTTGTTCTCACC-1dCA2_FI_GFPve_pos  2.0245363635
## FC_Mpos_AAACGCTAGAAACTAC-1FC_MI_GFPve_pos      0.1549745945
```

```
## FC_Mpos_AAAGGATCACCTGCGA-1FC_MI_GFPve_pos     -0.1879874247
## FC_Mpos_AAAGGATTCATTGCTT-1FC_MI_GFPve_pos      1.4454595966
## FC_Mpos_AAAGGGCAGCTCTATG-1FC_MI_GFPve_pos      1.1377388162
## FC_Mpos_AAAGTCCTCAGTGATC-1FC_MI_GFPve_pos      2.5012888831
## FC_Mpos_AAATGGAAGAGGGTGG-1FC_MI_GFPve_pos      1.9990742130
## FC_Mpos_AAATGGACAAGCAATA-1FC_MI_GFPve_pos      0.0480667127
## FC_Mpos_AACAACCAGGAATGTT-1FC_MI_GFPve_pos      0.7536954803
## FC_Mpos_AACACACCACTTGGGC-1FC_MI_GFPve_pos     -0.2088814157
## FC_Mpos_AACGAAAGTCGACTTA-1FC_MI_GFPve_pos     -1.3347157317
## FC_Mpos_AACGTCATCGCCGAAC-1FC_MI_GFPve_pos      2.2809908313
## FC_Mpos_AAGAACACAATGTGGG-1FC_MI_GFPve_pos      1.8008435888
## FC_Mpos_AAGATAGAGCCATTTG-1FC_MI_GFPve_pos      2.1947860641
## FC_Mpos_AAGCGTTAGAGAGGGC-1FC_MI_GFPve_pos      1.3543515128
## FC_Mpos_AAGCGTTCACAGCGCT-1FC_MI_GFPve_pos      2.2933645171
## FC_Mpos_AATGCCACAGAGGTAC-1FC_MI_GFPve_pos      1.9482575578
## FC_Mpos_AATTCCTAGCTCTGTA-1FC_MI_GFPve_pos      2.2636363429
## FC_Mpos_ACACAGTAGTGCACCC-1FC_MI_GFPve_pos      2.0610389632
## FC_Mpos_ACACTGAAGGCCCAAA-1FC_MI_GFPve_pos      2.6995112819
## FC_Mpos_ACATTTCAGTCCCTAA-1FC_MI_GFPve_pos      0.8545890969
## FC_Mpos_ACCACAAAGCTCCCTT-1FC_MI_GFPve_pos      0.8141258282
## FC_Mpos_ACCACAACAGGTCTCG-1FC_MI_GFPve_pos      0.4247756821
## FC_Mpos_ACCATTTGTTCCAAAC-1FC_MI_GFPve_pos      0.6776195687
## FC_Mpos_ACCCAAAGTTGTGTAC-1FC_MI_GFPve_pos      0.6717680616
## FC_Mpos_ACCCTTGTCTCGGTCT-1FC_MI_GFPve_pos      1.2345166129
## FC_Mpos_ACCTGAAAGATGCTAA-1FC_MI_GFPve_pos      0.8535752815
## FC_Mpos_ACCTGAACACGCTGAC-1FC_MI_GFPve_pos      1.0459090990
## FC_Mpos_ACGCACGCATATAGCC-1FC_MI_GFPve_pos      0.2163781894
## FC_Mpos_ACGGTCGCACATATGC-1FC_MI_GFPve_pos      2.1946580333
## FC_Mpos_ACGTAGTTCCGTTTCG-1FC_MI_GFPve_pos      1.5922636909
## FC_Mpos_ACTTAGGAGGACATCG-1FC_MI_GFPve_pos     -0.5682439285
## FC_Mpos_ACTTATCTCACATACG-1FC_MI_GFPve_pos     -0.4205854612
## FC_Mpos_AGACCATAGAATAACC-1FC_MI_GFPve_pos      1.6196430844
## FC_Mpos_AGAGCAGCATAATCGC-1FC_MI_GFPve_pos      0.7660642189
## FC_Mpos_AGATCGTGTAATTAGG-1FC_MI_GFPve_pos     -1.0587664920
## FC_Mpos_AGCCAATGTAATGATG-1FC_MI_GFPve_pos     -1.3550667840
## FC_Mpos_AGCGCCAAGTATTGCC-1FC_MI_GFPve_pos      1.6035133523
## FC_Mpos_AGCTTCCGTCTGTTAG-1FC_MI_GFPve_pos     -0.4962936717
## FC_Mpos_AGGAATAGTCTTAGTG-1FC_MI_GFPve_pos     -0.5251360017
## FC_Mpos_AGGACGACAGAGGAAA-1FC_MI_GFPve_pos      0.3886155111
## FC_Mpos_AGGAGGTTCGTGGACC-1FC_MI_GFPve_pos      2.0992183608
## FC_Mpos_AGGCATTCACGCGCAT-1FC_MI_GFPve_pos      0.8585344476
## FC_Mpos_AGGCCACCAACCCTAA-1FC_MI_GFPve_pos      1.5768719835
## FC_Mpos_AGGTTACAGAGGTTTA-1FC_MI_GFPve_pos      0.3011112136
## FC_Mpos_AGTCTCCTCGACGTCG-1FC_MI_GFPve_pos      0.8409497661
## FC_Mpos_AGTGATCCAGCAAGAC-1FC_MI_GFPve_pos      0.6327096147
## FC_Mpos_AGTGATCGTGTGTACT-1FC_MI_GFPve_pos      0.5233638508
## FC_Mpos_ATACCGACACGCCACA-1FC_MI_GFPve_pos      1.6405956668
## FC_Mpos_ATACTTCTCGAACGGA-1FC_MI_GFPve_pos      1.9879650993
## FC_Mpos_ATATCCTCACGAAGAC-1FC_MI_GFPve_pos     -1.6501168090
## FC_Mpos_ATCACAGTCCACGGAC-1FC_MI_GFPve_pos     -1.2410786229
## FC_Mpos_ATCCACCTCCGTGTCT-1FC_MI_GFPve_pos      1.0458639187
## FC_Mpos_ATCTCTAGTATACGGG-1FC_MI_GFPve_pos      0.9267290754
## FC_Mpos_ATGAGTCAGGAACGAA-1FC_MI_GFPve_pos     -0.1053612458
## FC_Mpos_ATGATCGGTGCATCTA-1FC_MI_GFPve_pos     -0.5527816492
## FC_Mpos_ATGCATGCACCCTGTT-1FC_MI_GFPve_pos      1.4323458595
## FC_Mpos_ATGGGTTAGTTGCGCC-1FC_MI_GFPve_pos      1.4855865163
## FC_Mpos_ATGGGTTCAAATTGCC-1FC_MI_GFPve_pos      1.8067763967
## FC_Mpos_ATGTCCCCACAAAGCG-1FC_MI_GFPve_pos      0.5755648834
## FC_Mpos_ATGTCCCCACATTGTG-1FC_MI_GFPve_pos     -0.1425106901
## FC_Mpos_ATTACCTTCGCTTACC-1FC_MI_GFPve_pos      2.0582797450
## FC_Mpos_ATTCACTGTACTAAGA-1FC_MI_GFPve_pos     -0.5875064212
## FC_Mpos_ATTCCCGGTGGCAGAT-1FC_MI_GFPve_pos      1.8230531139
## FC_Mpos_ATTTCACGTCTCAAGT-1FC_MI_GFPve_pos      0.8277292771
## FC_Mpos_ATTTCACTCGACGAGA-1FC_MI_GFPve_pos     -0.0893423783
## FC_Mpos_CAACCTCAGATCCCAT-1FC_MI_GFPve_pos     -1.1805118399
## FC_Mpos_CAATTTCAGATGTTAG-1FC_MI_GFPve_pos      0.6537187618
## FC_Mpos_CACACAACATTGTAGC-1FC_MI_GFPve_pos      1.6954269332
## FC_Mpos_CACGTGGGTCCCTAAA-1FC_MI_GFPve_pos      1.1566479129
## FC_Mpos_CACTGGGCAGGTTCAT-1FC_MI_GFPve_pos      0.7135342879
## FC_Mpos_CAGATCACAAGTATCC-1FC_MI_GFPve_pos      0.4136833471
```

```
## FC_Mpos_CAGATCAGTTAAGTCC-1FC_MI_GFPve_pos     -0.2256800311
## FC_Mpos_CAGATCAGTTCACCGG-1FC_MI_GFPve_pos     -0.3578257936
## FC_Mpos_CAGCAATGTGATAGTA-1FC_MI_GFPve_pos      1.4501109046
## FC_Mpos_CAGCACGCAACATCGT-1FC_MI_GFPve_pos     -0.4530922848
## FC_Mpos_CAGTTCCCAGTCAGAG-1FC_MI_GFPve_pos      0.9874471349
## FC_Mpos_CATACCCGTGCCCACA-1FC_MI_GFPve_pos      0.8329372329
## FC_Mpos_CATACTTGTAGACACG-1FC_MI_GFPve_pos      2.1387340945
## FC_Mpos_CATGCGGAGATTGCGG-1FC_MI_GFPve_pos     -1.2296264249
## FC_Mpos_CATGCGGTCTCGAGTA-1FC_MI_GFPve_pos      1.1817586345
## FC_Mpos_CATTCCGCAACTCCAA-1FC_MI_GFPve_pos      0.0965598987
## FC_Mpos_CATTGAGGTTGTGGAG-1FC_MI_GFPve_pos      0.5005201918
## FC_Mpos_CATTGCCAGTCTTCGA-1FC_MI_GFPve_pos      0.9551064295
## FC_Mpos_CATTGTTTCTACTTCA-1FC_MI_GFPve_pos      1.8848490638
## FC_Mpos_CCAATGATCTGTAAGC-1FC_MI_GFPve_pos      1.0335853619
## FC_Mpos_CCACACTGTGACAACG-1FC_MI_GFPve_pos      0.3360135329
## FC_Mpos_CCACTTGCACCCATAA-1FC_MI_GFPve_pos     -1.0881010371
## FC_Mpos_CCACTTGCACGACGCT-1FC_MI_GFPve_pos     -0.0168240520
## FC_Mpos_CCCTAACGTGTTCAGT-1FC_MI_GFPve_pos      1.4810236615
## FC_Mpos_CCGAACGCAAGTGATA-1FC_MI_GFPve_pos     -0.7970866638
## FC_Mpos_CCGTAGGAGTGTCATC-1FC_MI_GFPve_pos     -1.4347618895
## FC_Mpos_CCGTAGGTCTATCCAT-1FC_MI_GFPve_pos      0.6218932790
## FC_Mpos_CCTAAGACAGTCTTCC-1FC_MI_GFPve_pos     -0.3826884108
## FC_Mpos_CCTAAGATCTCGGGAC-1FC_MI_GFPve_pos      2.0062059087
## FC_Mpos_CCTCACACAGCAGTGA-1FC_MI_GFPve_pos      2.1709854526
## FC_Mpos_CCTCAGTTCCATACTT-1FC_MI_GFPve_pos     -0.7123869139
## FC_Mpos_CCTCATGGTGTCTCCT-1FC_MI_GFPve_pos      0.3315879476
## FC_Mpos_CCTCCTCCACCAAATC-1FC_MI_GFPve_pos      0.9044485015
## FC_Mpos_CCTTCAGTCCTATGGA-1FC_MI_GFPve_pos      0.1240041663
## FC_Mpos_CGCATAAGTGTCCTAA-1FC_MI_GFPve_pos      0.5520273072
## FC_Mpos_CGCCAGAGTTGGGCCT-1FC_MI_GFPve_pos      0.6799150628
## FC_Mpos_CGGCAGTTCTGGGCCA-1FC_MI_GFPve_pos     -0.1993418413
## FC_Mpos_CGGGCATAGAATTTGG-1FC_MI_GFPve_pos      2.0512698812
## FC_Mpos_CGTCCATAGTATCCTG-1FC_MI_GFPve_pos      1.0247763318
## FC_Mpos_CGTGCTTCACGTCTCT-1FC_MI_GFPve_pos      0.1711684388
## FC_Mpos_CGTTCTGGTTCAAACC-1FC_MI_GFPve_pos      0.7982979340
## FC_Mpos_CTAACCCGTGGCCTCA-1FC_MI_GFPve_pos      0.7013658804
## FC_Mpos_CTAGGTACAGTGAGCA-1FC_MI_GFPve_pos     -0.9931069690
## FC_Mpos_CTCAACCTCCAGTGTA-1FC_MI_GFPve_pos      2.0502879543
## FC_Mpos_CTCACTGGTCTTGAGT-1FC_MI_GFPve_pos      0.2036815581
## FC_Mpos_CTCAGGGGTGCTATTG-1FC_MI_GFPve_pos      1.0018401665
## FC_Mpos_CTCCAACCATCCGAGC-1FC_MI_GFPve_pos     -0.2919522899
## FC_Mpos_CTCCCAATCCAACCAA-1FC_MI_GFPve_pos      1.1776606960
## FC_Mpos_CTCCTTTGTGGAAATT-1FC_MI_GFPve_pos      0.7417474908
## FC_Mpos_CTCTCGAGTAGAGACC-1FC_MI_GFPve_pos      1.5875524205
## FC_Mpos_CTCTCGAGTCTCCTGT-1FC_MI_GFPve_pos      1.0428348107
## FC_Mpos_CTGCGAGCAGTCGCTG-1FC_MI_GFPve_pos     -1.5684590417
## FC_Mpos_CTGCTCAGTCTAGGTT-1FC_MI_GFPve_pos      2.1163914127
## FC_Mpos_CTGGACGAGCTTCTAG-1FC_MI_GFPve_pos      0.4806899649
## FC_Mpos_CTGTATTGTAGCTGAG-1FC_MI_GFPve_pos     -0.6355077105
## FC_Mpos_GAACACTTCGACATAC-1FC_MI_GFPve_pos      0.3378524703
## FC_Mpos_GAAGCCCTCCAGGACC-1FC_MI_GFPve_pos     -1.1414471942
## FC_Mpos_GAAGTAATCATGCCAA-1FC_MI_GFPve_pos      0.1649152448
## FC_Mpos_GACGCTGGTTCGAGCC-1FC_MI_GFPve_pos      1.3238202256
## FC_Mpos_GAGAAATTCAGCAGAG-1FC_MI_GFPve_pos      0.0320803371
## FC_Mpos_GAGCTGCAGCACCGAA-1FC_MI_GFPve_pos      1.6579520625
## FC_Mpos_GAGGCAATCGACGTCG-1FC_MI_GFPve_pos      0.1954178286
## FC_Mpos_GAGTGTTGTCATCAGT-1FC_MI_GFPve_pos     -2.7274928170
## FC_Mpos_GAGTTACTCGTTACCC-1FC_MI_GFPve_pos     -1.0486986714
## FC_Mpos_GATAGCTAGAGCTGCA-1FC_MI_GFPve_pos      0.1126339899
## FC_Mpos_GATCATGAGCAACTCT-1FC_MI_GFPve_pos      0.8383894486
## FC_Mpos_GATCATGCAGCAGGAT-1FC_MI_GFPve_pos      0.4024668557
## FC_Mpos_GATCATGTCCAAGGGA-1FC_MI_GFPve_pos     -0.5612247067
## FC_Mpos_GATGATCTCCAGCTCT-1FC_MI_GFPve_pos      1.8932841939
## FC_Mpos_GATTCTTTCTAGATCG-1FC_MI_GFPve_pos      1.7834054155
## FC_Mpos_GCAACATCAACGGCCT-1FC_MI_GFPve_pos      0.5769198758
## FC_Mpos_GCAACCGAGCAGTACG-1FC_MI_GFPve_pos      0.0336178762
## FC_Mpos_GCACATACACAGAAGC-1FC_MI_GFPve_pos      1.3458007497
## FC_Mpos_GCACTAATCCTACAAG-1FC_MI_GFPve_pos      0.1444184591
## FC_Mpos_GCAGCTGCAAACTAAG-1FC_MI_GFPve_pos      0.8641945047
## FC_Mpos_GCATCGGTCTCTGCCA-1FC_MI_GFPve_pos      1.8680797738
```

```
## FC_Mpos_GCATCTCTCGCCAGAC-1FC_MI_GFPve_pos      2.6295611781
## FC_Mpos_GCCAACGAGGAGGGTG-1FC_MI_GFPve_pos      0.1401582879
## FC_Mpos_GCCAGCATCCCATAAG-1FC_MI_GFPve_pos      0.1905839888
## FC_Mpos_GCCAGTGCAAGGGTCA-1FC_MI_GFPve_pos      2.5585977954
## FC_Mpos_GCCAGTGTCATTCATC-1FC_MI_GFPve_pos      2.3824679774
## FC_Mpos_GCCCGAATCAACGCTA-1FC_MI_GFPve_pos      1.9467301292
## FC_Mpos_GCCGATGCAGTGTATC-1FC_MI_GFPve_pos      0.7112664503
## FC_Mpos_GCCTGTTCAGCATGCC-1FC_MI_GFPve_pos     -0.7151479202
## FC_Mpos_GCGATCGCATCCCGTT-1FC_MI_GFPve_pos      1.1289982719
## FC_Mpos_GCGGATCTCTTCGACC-1FC_MI_GFPve_pos     -0.4374486881
## FC_Mpos_GCTACCTCAGAACTTC-1FC_MI_GFPve_pos      0.8187039417
## FC_Mpos_GCTTTCGCAGTGGCTC-1FC_MI_GFPve_pos      1.0493240875
## FC_Mpos_GGAGGATAGGAAAGTG-1FC_MI_GFPve_pos      2.3562824649
## FC_Mpos_GGATCTACACCAAAGG-1FC_MI_GFPve_pos     -1.3721261102
## FC_Mpos_GGATGTTTCTTCACGC-1FC_MI_GFPve_pos      0.0396278274
## FC_Mpos_GGGATGAAGGAATGTT-1FC_MI_GFPve_pos      0.9070995373
## FC_Mpos_GGGCCATTCTGAGAGG-1FC_MI_GFPve_pos      0.0414495503
## FC_Mpos_GGGCTCATCTGTCAGA-1FC_MI_GFPve_pos     -1.3757728415
## FC_Mpos_GGTCTGGAGCTGTTAC-1FC_MI_GFPve_pos      1.4948012752
## FC_Mpos_GGTGGCTAGCAAGTCG-1FC_MI_GFPve_pos      1.1166832370
## FC_Mpos_GGTGGCTTCAACGAGG-1FC_MI_GFPve_pos      2.1365096492
## FC_Mpos_GGTTAACTCTAGCATG-1FC_MI_GFPve_pos     -0.4825097996
## FC_Mpos_GGTTGTATCCGCAAAT-1FC_MI_GFPve_pos      0.0926137098
## FC_Mpos_GTAACACCAACTGCCG-1FC_MI_GFPve_pos      0.3700580818
## FC_Mpos_GTAACCACACTCAAGT-1FC_MI_GFPve_pos      0.4193893594
## FC_Mpos_GTAACCATCCTAGCCT-1FC_MI_GFPve_pos      0.3622380716
## FC_Mpos_GTAATCGTCCTGATAG-1FC_MI_GFPve_pos      1.1967840118
## FC_Mpos_GTATTGGGTCTTCCGT-1FC_MI_GFPve_pos      0.4669631345
## FC_Mpos_GTCAGCGCACAGCTGC-1FC_MI_GFPve_pos      0.5619382483
## FC_Mpos_GTCCCATCACGCGTCA-1FC_MI_GFPve_pos     -0.2937975901
## FC_Mpos_GTCCCATCACTTCCTG-1FC_MI_GFPve_pos     -0.9736818152
## FC_Mpos_GTCTAGAAGTGTTCAC-1FC_MI_GFPve_pos      2.1107542438
## FC_Mpos_GTCTGTCCAGACGATG-1FC_MI_GFPve_pos      0.4074085576
## FC_Mpos_GTGACGCCACATACGT-1FC_MI_GFPve_pos     -1.1950528699
## FC_Mpos_GTGAGCCAGACCTGGA-1FC_MI_GFPve_pos      0.8191654009
## FC_Mpos_GTGAGGATCTCATGGA-1FC_MI_GFPve_pos      2.3596625251
## FC_Mpos_GTGAGTTTCAGCTGAT-1FC_MI_GFPve_pos     -0.3539086955
## FC_Mpos_GTGCGTGCACCGCTAG-1FC_MI_GFPve_pos      1.1030824107
## FC_Mpos_GTGTAACCAGTCGCTG-1FC_MI_GFPve_pos      2.1146376056
## FC_Mpos_GTGTGGCAGCCTCATA-1FC_MI_GFPve_pos     -0.0382364916
## FC_Mpos_GTGTGGCAGGTACTGG-1FC_MI_GFPve_pos      0.8277741594
## FC_Mpos_GTTACCCCAAACCATC-1FC_MI_GFPve_pos      1.8367406053
## FC_Mpos_GTTAGTGGTCCTTTGC-1FC_MI_GFPve_pos      1.1315348071
## FC_Mpos_GTTCGCTTCAGAGTGG-1FC_MI_GFPve_pos      1.7313333673
## FC_Mpos_TAAGCCACACGCGCTA-1FC_MI_GFPve_pos     -0.2974031049
## FC_Mpos_TAAGCGTAGACATCAA-1FC_MI_GFPve_pos     -3.3779394704
## FC_Mpos_TAAGTCGAGACATACA-1FC_MI_GFPve_pos      0.3700045568
## FC_Mpos_TAAGTCGCAATCTGCA-1FC_MI_GFPve_pos      1.3801780862
## FC_Mpos_TAATTCCAGTTCGCAT-1FC_MI_GFPve_pos      2.2046506328
## FC_Mpos_TAATTCCCACCTGCGA-1FC_MI_GFPve_pos      0.7857464475
## FC_Mpos_TACACCCAGACTTGTC-1FC_MI_GFPve_pos     -0.4319506961
## FC_Mpos_TACCGAACAACCAGAG-1FC_MI_GFPve_pos      2.3484797401
## FC_Mpos_TACGCTCTCAGCGCGT-1FC_MI_GFPve_pos      1.8311278743
## FC_Mpos_TACGTCCAGGATTTAG-1FC_MI_GFPve_pos      0.7745912594
## FC_Mpos_TACTTACGTCCCTCAT-1FC_MI_GFPve_pos      0.4992924256
## FC_Mpos_TAGACTGCACTACACA-1FC_MI_GFPve_pos      1.2376139087
## FC_Mpos_TAGACTGTCAGCCTTC-1FC_MI_GFPve_pos     -0.2309342402
## FC_Mpos_TAGCACATCATGGTAC-1FC_MI_GFPve_pos      0.3301301074
## FC_Mpos_TAGCACATCCATCACC-1FC_MI_GFPve_pos     -0.3688617128
## FC_Mpos_TATATCCGTACGACAG-1FC_MI_GFPve_pos      0.3584357125
## FC_Mpos_TATTTCGGTGGTACAG-1FC_MI_GFPve_pos      1.2128966970
## FC_Mpos_TCAAGACTCTACGCAA-1FC_MI_GFPve_pos      0.3514883590
## FC_Mpos_TCAATCTTCTCTATGT-1FC_MI_GFPve_pos      1.3033561629
## FC_Mpos_TCAATCTTCTTTCAGT-1FC_MI_GFPve_pos      1.2127206248
## FC_Mpos_TCAATTCTCTTTCCGG-1FC_MI_GFPve_pos      1.0492365283
## FC_Mpos_TCACACCTCGAGCTGC-1FC_MI_GFPve_pos      1.9309017581
## FC_Mpos_TCACATTAGGTAAACT-1FC_MI_GFPve_pos      1.7271977586
## FC_Mpos_TCACGGGCATTCGGGC-1FC_MI_GFPve_pos      1.1352417392
## FC_Mpos_TCACGGGGTTCTTAGG-1FC_MI_GFPve_pos     -0.0186210426
## FC_Mpos_TCAGCCTGTATCGTGT-1FC_MI_GFPve_pos     -0.0178396332
```

```
## FC_Mpos_TCAGTTTAGAGACAAG-1FC_MI_GFPve_pos     -0.7208828407
## FC_Mpos_TCATACTGTACTAAGA-1FC_MI_GFPve_pos     -0.0082207176
## FC_Mpos_TCATCCGAGATACCAA-1FC_MI_GFPve_pos     -0.8204150277
## FC_Mpos_TCATGTTGTACTAAGA-1FC_MI_GFPve_pos      0.5111511571
## FC_Mpos_TCATTCAAGCAGATAT-1FC_MI_GFPve_pos     -0.6066436249
## FC_Mpos_TCATTGTCAATGTCAC-1FC_MI_GFPve_pos      0.1437017841
## FC_Mpos_TCCAGAATCTCTCTAA-1FC_MI_GFPve_pos      1.1211464328
## FC_Mpos_TCCCATGTCACGTCCT-1FC_MI_GFPve_pos      0.9927750749
## FC_Mpos_TCCGAAACACTGGACC-1FC_MI_GFPve_pos      0.6875692529
## FC_Mpos_TCCGAAACATTGCTTT-1FC_MI_GFPve_pos      1.2197904510
## FC_Mpos_TCCTCCCCATGAGGGT-1FC_MI_GFPve_pos     -1.2228989678
## FC_Mpos_TCCTCGACAAACTGCT-1FC_MI_GFPve_pos     -0.5550706464
## FC_Mpos_TCCTCTTTCATTGCGA-1FC_MI_GFPve_pos      1.5185122413
## FC_Mpos_TCGAAGTCACTCTGCT-1FC_MI_GFPve_pos     -0.8570008951
## FC_Mpos_TCGCACTGTGTGGACA-1FC_MI_GFPve_pos      0.8762620610
## FC_Mpos_TCGCACTTCGCAACAT-1FC_MI_GFPve_pos      0.7967378420
## FC_Mpos_TCGGTCTCAGTTCTAG-1FC_MI_GFPve_pos     -0.3741649168
## FC_Mpos_TCGGTCTCATTGCCGG-1FC_MI_GFPve_pos     -3.4299085217
## FC_Mpos_TCGTCCACACGGGCTT-1FC_MI_GFPve_pos     -0.0922503995
## FC_Mpos_TCGTCCACAGTAGGAC-1FC_MI_GFPve_pos      1.1310732288
## FC_Mpos_TCTCAGCAGTTTCGAC-1FC_MI_GFPve_pos      1.8377346915
## FC_Mpos_TCTCAGCTCGGTGTAT-1FC_MI_GFPve_pos     -1.7632193642
## FC_Mpos_TCTCCGAAGAAGATCT-1FC_MI_GFPve_pos      1.5738832874
## FC_Mpos_TCTGGCTGTCTACATG-1FC_MI_GFPve_pos     -0.9510109502
## FC_Mpos_TCTTAGTGTTAGAAAC-1FC_MI_GFPve_pos     -0.2782346087
## FC_Mpos_TCTTTGACATGACTCA-1FC_MI_GFPve_pos      2.0160075349
## FC_Mpos_TGAATGCTCCTCATAT-1FC_MI_GFPve_pos      1.2567132634
## FC_Mpos_TGACAGTAGTAGAGTT-1FC_MI_GFPve_pos      1.1249633950
## FC_Mpos_TGACAGTGTGTGATGG-1FC_MI_GFPve_pos      2.1217198295
## FC_Mpos_TGACCCTAGCACTCGC-1FC_MI_GFPve_pos     -0.0674187469
## FC_Mpos_TGACGCGCAGACCATT-1FC_MI_GFPve_pos     -1.1614854413
## FC_Mpos_TGACTCCCATGTTACG-1FC_MI_GFPve_pos     -1.1542729216
## FC_Mpos_TGACTCCTCTGGACCG-1FC_MI_GFPve_pos     -1.0257526713
## FC_Mpos_TGAGACTTCCCTCTAG-1FC_MI_GFPve_pos      1.2646270913
## FC_Mpos_TGAGCATTCTTACCAT-1FC_MI_GFPve_pos      1.1225079221
## FC_Mpos_TGATGGTGTGGATCGA-1FC_MI_GFPve_pos      0.4953685922
## FC_Mpos_TGCAGATTCTGGGCCA-1FC_MI_GFPve_pos      1.6269071025
## FC_Mpos_TGGAGAGAGGTCGAGT-1FC_MI_GFPve_pos      0.5169918162
## FC_Mpos_TGGGCGTAGATTGAGT-1FC_MI_GFPve_pos      1.9445967597
## FC_Mpos_TGTAACGAGGGCATGT-1FC_MI_GFPve_pos     -0.3590332347
## FC_Mpos_TGTGGCGGTGAAAGTT-1FC_MI_GFPve_pos      0.9336389226
## FC_Mpos_TGTGTGACAAGACTGG-1FC_MI_GFPve_pos     -0.5138573128
## FC_Mpos_TGTTCTATCGCGGACT-1FC_MI_GFPve_pos     -0.6658899861
## FC_Mpos_TTAATCCCACGCTGCA-1FC_MI_GFPve_pos     -0.1050441640
## FC_Mpos_TTACCGCTCGACATTG-1FC_MI_GFPve_pos      0.1388817427
## FC_Mpos_TTACTGTCACACCAGC-1FC_MI_GFPve_pos     -0.2870929318
## FC_Mpos_TTACTGTTCACCATGA-1FC_MI_GFPve_pos     -0.6406489092
## FC_Mpos_TTACTGTTCTAGACCA-1FC_MI_GFPve_pos      0.3137655777
## FC_Mpos_TTAGGCATCCTAGCTC-1FC_MI_GFPve_pos     -1.0129634219
## FC_Mpos_TTATTGCCAATCGCAT-1FC_MI_GFPve_pos     -1.3207291442
## FC_Mpos_TTCCTTCTCCCGAGTG-1FC_MI_GFPve_pos      0.0460911912
## FC_Mpos_TTCTAGTCAATCGAAA-1FC_MI_GFPve_pos      1.7455894870
## FC_Mpos_TTCTCTCAGATACTGA-1FC_MI_GFPve_pos      0.3739222211
## FC_Mpos_TTCTGTAAGCCTTTGA-1FC_MI_GFPve_pos      0.0852532161
## FC_Mpos_TTCTTCCCACCGTGCA-1FC_MI_GFPve_pos      0.8347378296
## FC_Mpos_TTCTTCCGTGGTTTAC-1FC_MI_GFPve_pos      1.0358085555
## FC_Mpos_TTCTTCCTCGGCCAAC-1FC_MI_GFPve_pos      1.5070364398
## FC_Mpos_TTGAACGGTGATCATC-1FC_MI_GFPve_pos      1.3613731784
## FC_Mpos_TTGGGATGTCACTGAT-1FC_MI_GFPve_pos     -0.3292048949
## FC_Mpos_TTGGTTTGTAAGGAGA-1FC_MI_GFPve_pos      0.5140495223
## FC_Mpos_TTGTTCACAGGCCCTA-1FC_MI_GFPve_pos     -0.0194546583
## FC_Mpos_TTGTTGTCAAATTGCC-1FC_MI_GFPve_pos      1.1580674571
## FC_Fpos_AACCCAAGTCAACATC-1FC_FI_GFPve_pos     -1.0622513371
## FC_Fpos_AACCTTTTCCGAACGC-1FC_FI_GFPve_pos      0.3250464422
## FC_Fpos_AAGACTCAGTATCTGC-1FC_FI_GFPve_pos      0.5136951727
## FC_Fpos_AAGACTCCACGGTGAA-1FC_FI_GFPve_pos      1.3773620052
## FC_Fpos_AAGCATCCAAAGGCAC-1FC_FI_GFPve_pos     -1.1453151780
## FC_Fpos_AATGCCAAGGGTTGCA-1FC_FI_GFPve_pos     -0.9090735989
## FC_Fpos_AATGGAACAATTGAGA-1FC_FI_GFPve_pos      1.6698902768
## FC_Fpos_AATGGCTGTTCTTGCC-1FC_FI_GFPve_pos      1.0522980017
```

```
## FC_Fpos_AATTCCTGTAGCTTAC-1FC_FI_GFPve_pos      0.8876035732
## FC_Fpos_ACAGAAAGTGGTCAAG-1FC_FI_GFPve_pos     -1.1267969685
## FC_Fpos_ACAGCCGGTCCTGGGT-1FC_FI_GFPve_pos      1.2370093984
## FC_Fpos_ACATGCATCACCTTGC-1FC_FI_GFPve_pos      0.3305014295
## FC_Fpos_ACCAAACGTCTACATG-1FC_FI_GFPve_pos      1.6080290002
## FC_Fpos_ACCAACATCTCGAACA-1FC_FI_GFPve_pos      0.3714378101
## FC_Fpos_ACCACAATCAACTTTC-1FC_FI_GFPve_pos      0.0690702607
## FC_Fpos_ACCTACCAGGTCACTT-1FC_FI_GFPve_pos     -3.4938056546
## FC_Fpos_ACCTGAAGTTTACACG-1FC_FI_GFPve_pos      1.6117874307
## FC_Fpos_ACGTCCTAGGTTGTTC-1FC_FI_GFPve_pos     -0.6691885072
## FC_Fpos_ACTGTCCCATCCGCGA-1FC_FI_GFPve_pos      1.0354729218
## FC_Fpos_AGAAATGCACTTGAGT-1FC_FI_GFPve_pos      2.0524676961
## FC_Fpos_AGAAGTAAGGTACTGG-1FC_FI_GFPve_pos     -0.9924675303
## FC_Fpos_AGACACTCAAATTGGA-1FC_FI_GFPve_pos     -0.2250910180
## FC_Fpos_AGACAGGAGCACTAAA-1FC_FI_GFPve_pos     -0.9243961650
## FC_Fpos_AGAGAATTCTCAGGCG-1FC_FI_GFPve_pos      2.2508468551
## FC_Fpos_AGAGAATTCTGTAAGC-1FC_FI_GFPve_pos     -0.5606671649
## FC_Fpos_AGATCCATCCGCACGA-1FC_FI_GFPve_pos      0.0258598921
## FC_Fpos_AGCCAGCGTTCCGGTG-1FC_FI_GFPve_pos      2.4806749744
## FC_Fpos_AGGCATTCAGCGTATT-1FC_FI_GFPve_pos      2.5258705539
## FC_Fpos_AGGTCATAGGATTCCT-1FC_FI_GFPve_pos      0.3585541767
## FC_Fpos_AGTCACAGTAGTTACC-1FC_FI_GFPve_pos      1.5367159766
## FC_Fpos_ATACCTTTCTTTCCGG-1FC_FI_GFPve_pos      0.1505279166
## FC_Fpos_ATCCACCGTATAGGAT-1FC_FI_GFPve_pos      0.3077908081
## FC_Fpos_ATCCACCTCTCGACGG-1FC_FI_GFPve_pos      0.4319094044
## FC_Fpos_ATCCCTGTCCCATACC-1FC_FI_GFPve_pos      1.0497697515
## FC_Fpos_ATCCCTGTCGGCTATA-1FC_FI_GFPve_pos     -0.3151532787
## FC_Fpos_ATTACCTTCGCCATAA-1FC_FI_GFPve_pos      2.2764809055
## FC_Fpos_ATTACTCTCTACCAGA-1FC_FI_GFPve_pos      0.4992870015
## FC_Fpos_ATTCGTTTCAACCGAT-1FC_FI_GFPve_pos      0.3279014600
## FC_Fpos_ATTCTACCAGAGTCAG-1FC_FI_GFPve_pos     -0.5365862327
## FC_Fpos_ATTCTACCAGCTGGTC-1FC_FI_GFPve_pos      1.7869726342
## FC_Fpos_ATTTACCTCCTCATAT-1FC_FI_GFPve_pos      1.2569607419
## FC_Fpos_CAACGATTCTATTCGT-1FC_FI_GFPve_pos      1.5547996682
## FC_Fpos_CAAGGGACATGACGGA-1FC_FI_GFPve_pos      1.5324709338
## FC_Fpos_CACAACAGTTGACTAC-1FC_FI_GFPve_pos      1.1222095412
## FC_Fpos_CACATGACAATGTGGG-1FC_FI_GFPve_pos     -1.3853536921
## FC_Fpos_CACCGTTCAACCAATC-1FC_FI_GFPve_pos      1.1492935342
## FC_Fpos_CACCGTTGTTTGACAC-1FC_FI_GFPve_pos      1.5147228164
## FC_Fpos_CACGGGTTCATGGTAC-1FC_FI_GFPve_pos      1.7166606111
## FC_Fpos_CAGCAATCAAGATGGC-1FC_FI_GFPve_pos      1.2243579549
## FC_Fpos_CAGCACGAGAGCAAGA-1FC_FI_GFPve_pos     -0.1897667127
## FC_Fpos_CAGCACGAGGACTTCT-1FC_FI_GFPve_pos     -0.4589518386
## FC_Fpos_CATACAGCAGACAAGC-1FC_FI_GFPve_pos      2.6245825214
## FC_Fpos_CATAGACTCTCCACTG-1FC_FI_GFPve_pos      2.3020374698
## FC_Fpos_CATTCATTCCTGCTAC-1FC_FI_GFPve_pos      1.5020869893
## FC_Fpos_CATTGCCTCGGCTCTT-1FC_FI_GFPve_pos     -1.4316622096
## FC_Fpos_CCAATGATCTCTGCCA-1FC_FI_GFPve_pos      1.2980968875
## FC_Fpos_CCACAAAGTTGCCGAC-1FC_FI_GFPve_pos      0.8338037056
## FC_Fpos_CCACCATCATACCACA-1FC_FI_GFPve_pos     -0.1426439213
## FC_Fpos_CCACGTTTCATATGGC-1FC_FI_GFPve_pos      0.7358567042
## FC_Fpos_CCACTTGTCATACAGC-1FC_FI_GFPve_pos      1.9243254584
## FC_Fpos_CCCAACTCACCGTGAC-1FC_FI_GFPve_pos      0.5008827669
## FC_Fpos_CCCATTGTCGCCATAA-1FC_FI_GFPve_pos      0.4392448885
## FC_Fpos_CCCTCTCCAACTTGCA-1FC_FI_GFPve_pos      1.4630843324
## FC_Fpos_CCCTCTCGTAAGGCTG-1FC_FI_GFPve_pos     -0.0498128312
## FC_Fpos_CCGGACATCATATGGC-1FC_FI_GFPve_pos     -0.1831841248
## FC_Fpos_CCGGACATCCGTAATG-1FC_FI_GFPve_pos      1.4069243592
## FC_Fpos_CCTATCGTCAAGTAAG-1FC_FI_GFPve_pos      2.5632071418
## FC_Fpos_CCTCAACCACAAGTTC-1FC_FI_GFPve_pos      0.0586824862
## FC_Fpos_CCTCACACAGAACGCA-1FC_FI_GFPve_pos      1.4959502143
## FC_Fpos_CCTGCATGTGCCTTCT-1FC_FI_GFPve_pos      2.5764508170
## FC_Fpos_CCTGTTGAGCTTACGT-1FC_FI_GFPve_pos      2.1494343204
## FC_Fpos_CCTTCAGTCGGCATTA-1FC_FI_GFPve_pos     -1.9486439305
## FC_Fpos_CGCCATTTCGCATGAT-1FC_FI_GFPve_pos     -0.1320839303
## FC_Fpos_CGGAATTTCCTCATAT-1FC_FI_GFPve_pos     -1.2485138255
## FC_Fpos_CGGGACTAGACTGAGC-1FC_FI_GFPve_pos     -1.0325263816
## FC_Fpos_CGGGTCAGTGCCAAGA-1FC_FI_GFPve_pos     -0.9674059230
## FC_Fpos_CGTAATGGTGTTAAAG-1FC_FI_GFPve_pos     -1.3240809518
## FC_Fpos_CTACAGACACATATCG-1FC_FI_GFPve_pos      2.2011773509
```

```
## FC_Fpos_CTACCCACACGCACCA-1FC_FI_GFPve_pos      1.4190899057
## FC_Fpos_CTACTATAGTACTGTC-1FC_FI_GFPve_pos      1.8243210239
## FC_Fpos_CTACTATGTAGTAAGT-1FC_FI_GFPve_pos      1.2953596038
## FC_Fpos_CTATAGGAGAGCCCAA-1FC_FI_GFPve_pos     -2.0227012711
## FC_Fpos_CTATCCGCACTGAATC-1FC_FI_GFPve_pos      1.9482495708
## FC_Fpos_CTATCCGGTTGATGTC-1FC_FI_GFPve_pos      0.0251482723
## FC_Fpos_CTCTGGTCAGTTACCA-1FC_FI_GFPve_pos      0.8922253770
## FC_Fpos_CTGCCATCAAGCCCAC-1FC_FI_GFPve_pos      0.2151113522
## FC_Fpos_CTTTCGGCATCACCAA-1FC_FI_GFPve_pos     -0.8250628787
## FC_Fpos_GAACGTTTCTTACCGC-1FC_FI_GFPve_pos      1.5510536355
## FC_Fpos_GAACTGTAGTGTTGAA-1FC_FI_GFPve_pos      2.6153969688
## FC_Fpos_GACATCAGTGAGCTCC-1FC_FI_GFPve_pos      1.8233970327
## FC_Fpos_GACCAATTCTCGGCTT-1FC_FI_GFPve_pos      1.5395352763
## FC_Fpos_GACCAATTCTGACGCG-1FC_FI_GFPve_pos      1.4654393119
## FC_Fpos_GACTGATTCAGGACAG-1FC_FI_GFPve_pos      0.4460151118
## FC_Fpos_GACTTCCAGCTCTTCC-1FC_FI_GFPve_pos      2.0156643314
## FC_Fpos_GAGATGGCAGGATTCT-1FC_FI_GFPve_pos      1.5682696027
## FC_Fpos_GAGTTACCAGCTCATA-1FC_FI_GFPve_pos      0.5476456804
## FC_Fpos_GATCACACAGGCTATT-1FC_FI_GFPve_pos      0.7064871115
## FC_Fpos_GATCCCTTCAACTTTC-1FC_FI_GFPve_pos      0.2272719440
## FC_Fpos_GCACATACATTGAGCT-1FC_FI_GFPve_pos      1.3394073171
## FC_Fpos_GCACGTGTCACGATCA-1FC_FI_GFPve_pos      0.6076563520
## FC_Fpos_GCCAGTGCAATTGCCA-1FC_FI_GFPve_pos     -0.4185075837
## FC_Fpos_GCGGATCCACATATGC-1FC_FI_GFPve_pos      0.8081868810
## FC_Fpos_GCGTTTCTCGCCGAAC-1FC_FI_GFPve_pos     -0.4624801355
## FC_Fpos_GCTACAAAGGGTCACA-1FC_FI_GFPve_pos      2.5258753223
## FC_Fpos_GCTCAAACAGAACTAA-1FC_FI_GFPve_pos      0.2237322253
## FC_Fpos_GGAAGTGGTCACCTTC-1FC_FI_GFPve_pos     -0.8308254557
## FC_Fpos_GGAGGTAGTCATGCAT-1FC_FI_GFPve_pos     -0.9114836531
## FC_Fpos_GGGACTCTCAACACCA-1FC_FI_GFPve_pos      1.9327634496
## FC_Fpos_GGGAGTATCTAGCAAC-1FC_FI_GFPve_pos      2.2592456264
## FC_Fpos_GGGCGTTAGGTAGTAT-1FC_FI_GFPve_pos      0.6560907287
## FC_Fpos_GGTAACTAGAGTCAGC-1FC_FI_GFPve_pos      1.0170034093
## FC_Fpos_GGTAACTTCCGTGGTG-1FC_FI_GFPve_pos      0.9004003925
## FC_Fpos_GGTGAAGTCGAGAGAC-1FC_FI_GFPve_pos      0.2698403967
## FC_Fpos_GTAGTACAGAGAGCAA-1FC_FI_GFPve_pos      1.5414544267
## FC_Fpos_GTATTGGTCATAAGGA-1FC_FI_GFPve_pos      0.7749024553
## FC_Fpos_GTCAAGTGTAGCGTAG-1FC_FI_GFPve_pos      1.3855249805
## FC_Fpos_GTCAGCGGTTCCGTTC-1FC_FI_GFPve_pos      0.7136480135
## FC_Fpos_GTCATCCGTGGGAGAG-1FC_FI_GFPve_pos      0.4338078124
## FC_Fpos_GTCCACTAGCTTGTTG-1FC_FI_GFPve_pos      0.2519323898
## FC_Fpos_GTCGAATGTATCTCTT-1FC_FI_GFPve_pos      2.5439882201
## FC_Fpos_GTCGCGAAGCACCTGC-1FC_FI_GFPve_pos      0.7079775852
## FC_Fpos_GTCTACCCACGTCGTG-1FC_FI_GFPve_pos     -0.9132212477
## FC_Fpos_GTGAGGATCTATTTCG-1FC_FI_GFPve_pos      0.8976575655
## FC_Fpos_GTGCGTGGTCTCGGGT-1FC_FI_GFPve_pos      0.0314009858
## FC_Fpos_GTGGAAGAGGAGTCTG-1FC_FI_GFPve_pos      0.8194724840
## FC_Fpos_GTTACGATCCAGTACA-1FC_FI_GFPve_pos      1.2897611779
## FC_Fpos_GTTGTCCCACTGCTTC-1FC_FI_GFPve_pos      0.8767943305
## FC_Fpos_TAATCTCTCATAGCAC-1FC_FI_GFPve_pos      0.9210065526
## FC_Fpos_TACACCCTCTAGCAAC-1FC_FI_GFPve_pos      1.0828720850
## FC_Fpos_TACATTCGTCAGACTT-1FC_FI_GFPve_pos      2.1189992351
## FC_Fpos_TACCGAAAGGTAACTA-1FC_FI_GFPve_pos      1.1725783271
## FC_Fpos_TACTGCCTCTATACGG-1FC_FI_GFPve_pos      1.7974610252
## FC_Fpos_TAGGAGGAGGGATCAC-1FC_FI_GFPve_pos      1.7108009977
## FC_Fpos_TAGGTTGAGTGCTACT-1FC_FI_GFPve_pos      1.1203501147
## FC_Fpos_TAGGTTGCAACCGGAA-1FC_FI_GFPve_pos      0.3046248925
## FC_Fpos_TAGGTTGGTACTCGAT-1FC_FI_GFPve_pos      2.4597375316
## FC_Fpos_TATACCTCATACACCA-1FC_FI_GFPve_pos      0.9322830362
## FC_Fpos_TATATCCCAGCTTTGA-1FC_FI_GFPve_pos     -1.6581703502
## FC_Fpos_TATCGCCTCTAGCATG-1FC_FI_GFPve_pos     -0.1028222221
## FC_Fpos_TATCTGTCAACGGGTA-1FC_FI_GFPve_pos     -0.8729279714
## FC_Fpos_TCAAGACTCGGTGTTA-1FC_FI_GFPve_pos      2.4049684924
## FC_Fpos_TCACACCGTTCGTGCG-1FC_FI_GFPve_pos     -2.6597302037
## FC_Fpos_TCACATTAGCGTCTCG-1FC_FI_GFPve_pos      1.3960329217
## FC_Fpos_TCACGCTCACTTCCTG-1FC_FI_GFPve_pos      1.8473582191
## FC_Fpos_TCACTCGCACCCAATA-1FC_FI_GFPve_pos      2.2453701419
## FC_Fpos_TCCCAGTGTGCTCTCT-1FC_FI_GFPve_pos      2.3624524993
## FC_Fpos_TCCTCTTCATGTAACC-1FC_FI_GFPve_pos      1.6863541526
## FC_Fpos_TCGGTCTTCTTTCCAA-1FC_FI_GFPve_pos      2.1672780437
```

```
## FC_Fpos_TCTCACGCAACGGCTC-1FC_FI_GFPve_pos      1.2733855170
## FC_Fpos_TCTGCCATCTACACAG-1FC_FI_GFPve_pos      2.1426422519
## FC_Fpos_TGATGCAAGGATACAT-1FC_FI_GFPve_pos      1.8856997413
## FC_Fpos_TGATTTCTCACAGAGG-1FC_FI_GFPve_pos      0.4099019689
## FC_Fpos_TGCAGGCAGCAGTAAT-1FC_FI_GFPve_pos      2.6218671722
## FC_Fpos_TGCAGTACAAGTGATA-1FC_FI_GFPve_pos      0.7460240645
## FC_Fpos_TGCCGAGTCTCTGGTC-1FC_FI_GFPve_pos      2.6541368884
## FC_Fpos_TGCGGGTAGGATGTTA-1FC_FI_GFPve_pos      1.6185924930
## FC_Fpos_TGGAGAGTCTAGCAAC-1FC_FI_GFPve_pos      2.0918815059
## FC_Fpos_TGGTAGTTCGCTACGG-1FC_FI_GFPve_pos      2.1960658950
## FC_Fpos_TGTAAGCGTGGTTTAC-1FC_FI_GFPve_pos     -0.4919397312
## FC_Fpos_TGTGATGCACCAGGTC-1FC_FI_GFPve_pos      2.2989211005
## FC_Fpos_TGTTCATTCTTCGGAA-1FC_FI_GFPve_pos      0.9681515021
## FC_Fpos_TGTTTGTTCCGCCTAT-1FC_FI_GFPve_pos     -0.7400527674
## FC_Fpos_TTCCACGAGCTCGGCT-1FC_FI_GFPve_pos     -1.2101245004
## FC_Fpos_TTCCTCTGTTTCGCTC-1FC_FI_GFPve_pos      1.0858841461
## FC_Fpos_TTGCGTCCAGCTACAT-1FC_FI_GFPve_pos      1.9547099990
## FC_Fpos_TTGGGATGTCAACCTA-1FC_FI_GFPve_pos      0.5126831156
## FC_Fpos_TTGTTCATCCCTGTTG-1FC_FI_GFPve_pos      0.7463843745
## FC_Fpos_TTGTTCATCCTGGGAC-1FC_FI_GFPve_pos      1.1813626212
## FC_Fpos_TTTCATGTCCAACACA-1FC_FI_GFPve_pos      0.6301245016
## FC_Fpos_TTTCGATCACACCGCA-1FC_FI_GFPve_pos      2.0597929877
## FC_Fpos_TTTGACTAGACGGTTG-1FC_FI_GFPve_pos     -0.5541560727
## dCA2_Mpos_AACAAGATCCGCAACG-1dCA2_MI_GFPve_pos -2.5464537221
## dCA2_Mpos_AACAGGGCAGAGACTG-1dCA2_MI_GFPve_pos -2.4896154481
## dCA2_Mpos_AACCAACTCGAACTCA-1dCA2_MI_GFPve_pos -1.5905408936
## dCA2_Mpos_AACCTGAAGAGAGCAA-1dCA2_MI_GFPve_pos -2.2777772027
## dCA2_Mpos_AACCTGATCGAGTCTA-1dCA2_MI_GFPve_pos -2.0762982446
## dCA2_Mpos_AACCTTTAGCATCGAG-1dCA2_MI_GFPve_pos -1.8644549924
## dCA2_Mpos_AACGAAAAGCATTGAA-1dCA2_MI_GFPve_pos -1.9086494523
## dCA2_Mpos_AACGAAATCGGCTGTG-1dCA2_MI_GFPve_pos -2.1897244530
## dCA2_Mpos_AAGATAGCAATGAACA-1dCA2_MI_GFPve_pos -2.2311277467
## dCA2_Mpos_AAGCGAGAGCCAGTAG-1dCA2_MI_GFPve_pos -1.4807472306
## dCA2_Mpos_AAGGAATTCACCATCC-1dCA2_MI_GFPve_pos -1.6575166064
## dCA2_Mpos_AAGGAATTCCCTTCCC-1dCA2_MI_GFPve_pos -1.2577762681
## dCA2_Mpos_AAGTGAACAAGTGTCT-1dCA2_MI_GFPve_pos -1.6532253104
## dCA2_Mpos_AATCACGGTCACAGTT-1dCA2_MI_GFPve_pos -2.1606659966
## dCA2_Mpos_AATGAAGCATTCAGCA-1dCA2_MI_GFPve_pos -2.4189717847
## dCA2_Mpos_AATGAAGGTAGCGTAG-1dCA2_MI_GFPve_pos -0.3305213230
## dCA2_Mpos_AATGACCAGTCACAGG-1dCA2_MI_GFPve_pos -1.7389144975
## dCA2_Mpos_AATGCCAAGCGACCCT-1dCA2_MI_GFPve_pos -2.3211543637
## dCA2_Mpos_AATGCCAAGTCAATCC-1dCA2_MI_GFPve_pos -2.2787842828
## dCA2_Mpos_AATGGAAGTCTGTAAC-1dCA2_MI_GFPve_pos -1.4180619794
## dCA2_Mpos_AATGGCTCAATAACGA-1dCA2_MI_GFPve_pos -2.4492688256
## dCA2_Mpos_AATTCCTTCACGGGAA-1dCA2_MI_GFPve_pos -2.1686148721
## dCA2_Mpos_AATTTCCAGTTCACTG-1dCA2_MI_GFPve_pos -2.1476876813
## dCA2_Mpos_ACAAAGAAGGAACGTC-1dCA2_MI_GFPve_pos -1.9956517297
## dCA2_Mpos_ACAAAGATCGTTCTGC-1dCA2_MI_GFPve_pos -2.4809861260
## dCA2_Mpos_ACAACCAAGCGTTAGG-1dCA2_MI_GFPve_pos -2.4011676389
## dCA2_Mpos_ACAAGCTGTTAGCGGA-1dCA2_MI_GFPve_pos -2.2708113271
## dCA2_Mpos_ACACAGTTCATAGCAC-1dCA2_MI_GFPve_pos -2.0389840680
## dCA2_Mpos_ACAGGGAAGTTCGGTT-1dCA2_MI_GFPve_pos -2.2228803712
## dCA2_Mpos_ACATTTCAGCTACAAA-1dCA2_MI_GFPve_pos -1.6011998730
## dCA2_Mpos_ACCAAACGTCATCGGC-1dCA2_MI_GFPve_pos -2.2851646024
## dCA2_Mpos_ACCAACAAGGTTGGAC-1dCA2_MI_GFPve_pos -1.2278217154
## dCA2_Mpos_ACCCTCAAGAGTCCGA-1dCA2_MI_GFPve_pos -1.9770207482
## dCA2_Mpos_ACCCTCACAGGTACGA-1dCA2_MI_GFPve_pos -1.9165930825
## dCA2_Mpos_ACCCTTGCAATGCAGG-1dCA2_MI_GFPve_pos -2.3817520219
## dCA2_Mpos_ACCTGTCAGCATCGAG-1dCA2_MI_GFPve_pos -2.4211456853
## dCA2_Mpos_ACGCACGGTTGGGATG-1dCA2_MI_GFPve_pos -1.5552805739
## dCA2_Mpos_ACGGTCGGTCCCTCAT-1dCA2_MI_GFPve_pos -1.9103293496
## dCA2_Mpos_ACGGTTATCAACACGT-1dCA2_MI_GFPve_pos -2.4237961846
## dCA2_Mpos_ACTATGGGTGCCCTTT-1dCA2_MI_GFPve_pos -2.2145071107
## dCA2_Mpos_ACTATTCTCTAGTGTG-1dCA2_MI_GFPve_pos -1.1932603198
## dCA2_Mpos_ACTGATGTCCTACGGG-1dCA2_MI_GFPve_pos -2.3920934277
## dCA2_Mpos_ACTGTGAGTAAGTAGT-1dCA2_MI_GFPve_pos -2.1704611855
## dCA2_Mpos_ACTTATCTCAATCAGC-1dCA2_MI_GFPve_pos -2.1271660405
## dCA2_Mpos_ACTTCGCCATGGCACC-1dCA2_MI_GFPve_pos -2.1108388978
## dCA2_Mpos_ACTTCGCGTGCAATAA-1dCA2_MI_GFPve_pos -2.4035532552
## dCA2_Mpos_ACTTCGCTCTAGTGTG-1dCA2_MI_GFPve_pos -1.3657533007
```

```
## dCA2_Mpos_AGAAATGGTAACATGA-1dCA2_MI_GFPve_pos -1.9288547116
## dCA2_Mpos_AGAACCTAGTCCGTCG-1dCA2_MI_GFPve_pos -1.6923014003
## dCA2_Mpos_AGACCCGAGAGGCTGT-1dCA2_MI_GFPve_pos -0.9951326924
## dCA2_Mpos_AGAGAATGTACCGTCG-1dCA2_MI_GFPve_pos -1.5870952683
## dCA2_Mpos_AGAGAGCTCAGCTGTA-1dCA2_MI_GFPve_pos -2.2956645566
## dCA2_Mpos_AGATCCACAATGGCAG-1dCA2_MI_GFPve_pos -2.4809506017
## dCA2_Mpos_AGATCCATCATTTGGG-1dCA2_MI_GFPve_pos -1.5034762698
## dCA2_Mpos_AGCGATTCAATTGCAC-1dCA2_MI_GFPve_pos -2.1486189442
## dCA2_Mpos_AGCGATTTCCTGTAAG-1dCA2_MI_GFPve_pos  0.6316361946
## dCA2_Mpos_AGCGCCAGTAGGTACG-1dCA2_MI_GFPve_pos -1.8661819773
## dCA2_Mpos_AGCGCTGAGATGGCGT-1dCA2_MI_GFPve_pos -2.4579641896
## dCA2_Mpos_AGGACGATCTTGATTC-1dCA2_MI_GFPve_pos -2.4331655579
## dCA2_Mpos_AGGATCTGTAAGAACT-1dCA2_MI_GFPve_pos -1.9439017850
## dCA2_Mpos_AGGCCACAGCGGACAT-1dCA2_MI_GFPve_pos -2.3836295682
## dCA2_Mpos_AGGGAGTGTTGCTAGT-1dCA2_MI_GFPve_pos -2.4541103917
## dCA2_Mpos_AGGGCTCCAAGATGGC-1dCA2_MI_GFPve_pos -2.0182623940
## dCA2_Mpos_AGGGCTCGTGGAGAAA-1dCA2_MI_GFPve_pos -1.8667382079
## dCA2_Mpos_AGGTCATGTGCATACT-1dCA2_MI_GFPve_pos -2.3140823918
## dCA2_Mpos_AGGTTGTAGCAATTAG-1dCA2_MI_GFPve_pos -1.9729876595
## dCA2_Mpos_AGTAGTCTCGGTTGTA-1dCA2_MI_GFPve_pos -1.7714793759
## dCA2_Mpos_AGTCTCCAGGAAGAAC-1dCA2_MI_GFPve_pos -2.0507445412
## dCA2_Mpos_AGTCTCCAGGTACAAT-1dCA2_MI_GFPve_pos -2.1447269994
## dCA2_Mpos_AGTGATCCATGACTTG-1dCA2_MI_GFPve_pos -1.6754742938
## dCA2_Mpos_AGTGCCGTCTCATTGT-1dCA2_MI_GFPve_pos -1.3447538691
## dCA2_Mpos_AGTGTTGGTTGACTGT-1dCA2_MI_GFPve_pos -2.3562731820
## dCA2_Mpos_ATACTTCGTGTTCATG-1dCA2_MI_GFPve_pos -1.0888278561
## dCA2_Mpos_ATAGACCGTTGTGGAG-1dCA2_MI_GFPve_pos -1.9390192109
## dCA2_Mpos_ATAGAGAGTACACGTT-1dCA2_MI_GFPve_pos -1.8857122737
## dCA2_Mpos_ATCACGAAGTGAACAT-1dCA2_MI_GFPve_pos -2.3025047856
## dCA2_Mpos_ATCACGAGTTCTAACG-1dCA2_MI_GFPve_pos -2.3284323293
## dCA2_Mpos_ATCAGGTTCATTCATC-1dCA2_MI_GFPve_pos -2.2145440656
## dCA2_Mpos_ATCCACCAGACGGAAA-1dCA2_MI_GFPve_pos -0.5698069411
## dCA2_Mpos_ATCCATTGTGTTGATC-1dCA2_MI_GFPve_pos -1.8436000424
## dCA2_Mpos_ATCCCTGCAAGAGAGA-1dCA2_MI_GFPve_pos -2.2336089688
## dCA2_Mpos_ATCCTATCAACGGCCT-1dCA2_MI_GFPve_pos -2.1689486581
## dCA2_Mpos_ATCCTATCACTTGGGC-1dCA2_MI_GFPve_pos -2.3866958695
## dCA2_Mpos_ATCGTAGAGCTCGTGC-1dCA2_MI_GFPve_pos -2.1017901974
## dCA2_Mpos_ATCGTAGGTCTTGAGT-1dCA2_MI_GFPve_pos -1.9276654797
## dCA2_Mpos_ATCTCTAAGACGCCAA-1dCA2_MI_GFPve_pos -2.2387621480
## dCA2_Mpos_ATGAAAGTCATTTGGG-1dCA2_MI_GFPve_pos -2.3035800534
## dCA2_Mpos_ATGAGGGAGCTCCATA-1dCA2_MI_GFPve_pos -1.0068658667
## dCA2_Mpos_ATGAGTCAGCCATTGT-1dCA2_MI_GFPve_pos -1.5780491906
## dCA2_Mpos_ATGCGATTCAAATGAG-1dCA2_MI_GFPve_pos -0.0015892911
## dCA2_Mpos_ATGGATCAGTCACTGT-1dCA2_MI_GFPve_pos -1.2679377871
## dCA2_Mpos_ATGGATCTCCCGAATA-1dCA2_MI_GFPve_pos -0.3570170480
## dCA2_Mpos_ATGGGAGTCTCTGCTG-1dCA2_MI_GFPve_pos -0.5312321263
## dCA2_Mpos_ATGGGTTCAGTTTCGA-1dCA2_MI_GFPve_pos -1.6651144105
## dCA2_Mpos_ATGTCTTTCTTCTGTA-1dCA2_MI_GFPve_pos -1.7828556376
## dCA2_Mpos_ATTCAGGAGCGTCTCG-1dCA2_MI_GFPve_pos -1.7769860106
## dCA2_Mpos_ATTCCCGAGTCTTGGT-1dCA2_MI_GFPve_pos -1.1055502969
## dCA2_Mpos_ATTCTACAGTAGACCG-1dCA2_MI_GFPve_pos -2.4641611653
## dCA2_Mpos_ATTCTACTCGACATAC-1dCA2_MI_GFPve_pos -2.3568728047
## dCA2_Mpos_ATTTACCAGATAGTCA-1dCA2_MI_GFPve_pos -0.5064560252
## dCA2_Mpos_ATTTCTGTCAGTCCGG-1dCA2_MI_GFPve_pos -0.9798545914
## dCA2_Mpos_CAACCAAAGTATGGCG-1dCA2_MI_GFPve_pos -2.2747910100
## dCA2_Mpos_CAACCTCGTGATATAG-1dCA2_MI_GFPve_pos -2.2332320290
## dCA2_Mpos_CAAGACTGTTTAGTCG-1dCA2_MI_GFPve_pos -1.9216573315
## dCA2_Mpos_CAAGAGGCAGGGTCTC-1dCA2_MI_GFPve_pos -2.1768765527
## dCA2_Mpos_CAAGGGAAGACTTCAC-1dCA2_MI_GFPve_pos -2.0514266568
## dCA2_Mpos_CAAGGGACACATGAAA-1dCA2_MI_GFPve_pos -2.3030815202
## dCA2_Mpos_CAATGACGTCTTGAAC-1dCA2_MI_GFPve_pos -2.5870504456
## dCA2_Mpos_CAATGACGTGACTCGC-1dCA2_MI_GFPve_pos -1.7155582982
## dCA2_Mpos_CACAGATAGAGGCGTT-1dCA2_MI_GFPve_pos -1.3780305463
## dCA2_Mpos_CACAGGCCAGTCGCTG-1dCA2_MI_GFPve_pos -2.0837690907
## dCA2_Mpos_CACATGAGTGAGATTA-1dCA2_MI_GFPve_pos -1.6937044936
## dCA2_Mpos_CACGGGTTCCCAGGAC-1dCA2_MI_GFPve_pos -1.7095706540
## dCA2_Mpos_CACTAAGTCCGTAGGC-1dCA2_MI_GFPve_pos -2.3674814778
## dCA2_Mpos_CACTAAGTCTCAACGA-1dCA2_MI_GFPve_pos -2.0188913422
## dCA2_Mpos_CACTGAATCAAGATAG-1dCA2_MI_GFPve_pos -2.0879240113
## dCA2_Mpos_CACTGGGAGACAACAT-1dCA2_MI_GFPve_pos -2.3435876447
```

```
## dCA2_Mpos_CACTGGGGTAGATCCT-1dCA2_MI_GFPve_pos -2.5566859322
## dCA2_Mpos_CAGATCAAGAGTGTTA-1dCA2_MI_GFPve_pos -2.0414245206
## dCA2_Mpos_CAGATCATCACCATGA-1dCA2_MI_GFPve_pos -2.2934031564
## dCA2_Mpos_CAGCAATAGATGATTG-1dCA2_MI_GFPve_pos -1.3642768937
## dCA2_Mpos_CAGCAGCCACAACCGC-1dCA2_MI_GFPve_pos -2.4010453301
## dCA2_Mpos_CAGCAGCTCTGAGAAA-1dCA2_MI_GFPve_pos -2.0062925893
## dCA2_Mpos_CAGCGTGAGTGCTCAT-1dCA2_MI_GFPve_pos -2.2978184300
## dCA2_Mpos_CAGGCCATCCCGTGTT-1dCA2_MI_GFPve_pos -2.3175268250
## dCA2_Mpos_CAGTGCGCATCCTGTC-1dCA2_MI_GFPve_pos -2.3161947804
## dCA2_Mpos_CAGTTAGGTCGTTCAA-1dCA2_MI_GFPve_pos -1.4731148558
## dCA2_Mpos_CATCCCATCCATAGGT-1dCA2_MI_GFPve_pos -2.3775932866
## dCA2_Mpos_CATGCGGCACTTCAAG-1dCA2_MI_GFPve_pos -2.2043461877
## dCA2_Mpos_CATGGATGTACCGTGC-1dCA2_MI_GFPve_pos -2.0397851544
## dCA2_Mpos_CATGGTATCCATCTCG-1dCA2_MI_GFPve_pos -2.2636094170
## dCA2_Mpos_CATGGTATCCCGGTAG-1dCA2_MI_GFPve_pos -2.3055896836
## dCA2_Mpos_CCAAGCGAGGGAGGCA-1dCA2_MI_GFPve_pos -1.9251165467
## dCA2_Mpos_CCACCATGTCCTTGTC-1dCA2_MI_GFPve_pos -2.2162256318
## dCA2_Mpos_CCATCACAGTTCCTGA-1dCA2_MI_GFPve_pos -2.2574055272
## dCA2_Mpos_CCATCACTCAAACCTG-1dCA2_MI_GFPve_pos -2.2724509316
## dCA2_Mpos_CCCTGATCACCGTCGA-1dCA2_MI_GFPve_pos -2.5291702824
## dCA2_Mpos_CCGAACGTCACTCTTA-1dCA2_MI_GFPve_pos -0.9807812053
## dCA2_Mpos_CCGATGGCACCTTCCA-1dCA2_MI_GFPve_pos -1.7852536517
## dCA2_Mpos_CCGGTAGTCTTCCTAA-1dCA2_MI_GFPve_pos -2.0361011105
## dCA2_Mpos_CCGTTCAAGCCGTTGC-1dCA2_MI_GFPve_pos -2.1460807401
## dCA2_Mpos_CCGTTCACAATCGCAT-1dCA2_MI_GFPve_pos -0.8767203170
## dCA2_Mpos_CCTACGTCATACCGTA-1dCA2_MI_GFPve_pos -1.8979475575
## dCA2_Mpos_CCTCATGAGTAGTCAA-1dCA2_MI_GFPve_pos -1.9193012791
## dCA2_Mpos_CCTCATGAGTTAGTAG-1dCA2_MI_GFPve_pos -2.1123652535
## dCA2_Mpos_CCTCCTCAGTCATCCA-1dCA2_MI_GFPve_pos -1.4872711974
## dCA2_Mpos_CCTCTAGCACTGGCCA-1dCA2_MI_GFPve_pos -1.1763864833
## dCA2_Mpos_CCTCTAGTCGTGCTCT-1dCA2_MI_GFPve_pos -2.1458599644
## dCA2_Mpos_CCTTCAGTCTAGGCCG-1dCA2_MI_GFPve_pos -2.3024203854
## dCA2_Mpos_CCTTGTGAGCGCATCC-1dCA2_MI_GFPve_pos -2.2049405652
## dCA2_Mpos_CGAAGGACAACACGTT-1dCA2_MI_GFPve_pos -1.6718297082
## dCA2_Mpos_CGAAGTTTCTCAGTCC-1dCA2_MI_GFPve_pos -2.0627234059
## dCA2_Mpos_CGAGAAGAGCATCCTA-1dCA2_MI_GFPve_pos -1.6553299504
## dCA2_Mpos_CGAGAAGAGCGCCTTG-1dCA2_MI_GFPve_pos -1.9078600484
## dCA2_Mpos_CGAGGAAAGGTCGACA-1dCA2_MI_GFPve_pos -1.9233009892
## dCA2_Mpos_CGAGGCTCAAAGACTA-1dCA2_MI_GFPve_pos -1.9265396672
## dCA2_Mpos_CGAGGCTTCCAATCCC-1dCA2_MI_GFPve_pos -2.3309168893
## dCA2_Mpos_CGAGTTAGTGCTGTCG-1dCA2_MI_GFPve_pos -1.8952339965
## dCA2_Mpos_CGAGTTAGTGGGCTCT-1dCA2_MI_GFPve_pos -2.3391583043
## dCA2_Mpos_CGCATAAGTAGCTGCC-1dCA2_MI_GFPve_pos -2.0773303586
## dCA2_Mpos_CGCCAGACACTGCACG-1dCA2_MI_GFPve_pos -1.7403860169
## dCA2_Mpos_CGCCAGATCCATACAG-1dCA2_MI_GFPve_pos -2.3935098725
## dCA2_Mpos_CGCCATTTCGCCTTTG-1dCA2_MI_GFPve_pos -2.0098693448
## dCA2_Mpos_CGGAACCCATGGTACT-1dCA2_MI_GFPve_pos -2.1287217217
## dCA2_Mpos_CGGAATTGTCGTTATG-1dCA2_MI_GFPve_pos -2.3274765092
## dCA2_Mpos_CGGAATTGTTGCATAC-1dCA2_MI_GFPve_pos -2.5048577863
## dCA2_Mpos_CGGGTGTAGATCCTAC-1dCA2_MI_GFPve_pos -2.2700605470
## dCA2_Mpos_CGTAATGGTGTTATCG-1dCA2_MI_GFPve_pos -2.1774370747
## dCA2_Mpos_CGTAGTACAACCACGC-1dCA2_MI_GFPve_pos -2.3018214780
## dCA2_Mpos_CGTCAAAAGGCTTAAA-1dCA2_MI_GFPve_pos -2.0863394814
## dCA2_Mpos_CGTCAAATCACAGAGG-1dCA2_MI_GFPve_pos -1.9353201466
## dCA2_Mpos_CGTCCATAGTCCGCGT-1dCA2_MI_GFPve_pos -1.9121401387
## dCA2_Mpos_CGTCCATGTCTCGACG-1dCA2_MI_GFPve_pos -2.2351231652
## dCA2_Mpos_CGTGAATAGGACTTCT-1dCA2_MI_GFPve_pos -1.6328884440
## dCA2_Mpos_CGTGAATTCGCTAATG-1dCA2_MI_GFPve_pos -1.8592320758
## dCA2_Mpos_CGTGATATCTTGGATG-1dCA2_MI_GFPve_pos -1.4625468331
## dCA2_Mpos_CGTTAGATCATTCGGA-1dCA2_MI_GFPve_pos -2.4266505318
## dCA2_Mpos_CTAAGTGGTTTCGTTT-1dCA2_MI_GFPve_pos -2.0334029275
## dCA2_Mpos_CTACAGAGTGATTGGG-1dCA2_MI_GFPve_pos  0.2677289617
## dCA2_Mpos_CTACCCACATCCGAAT-1dCA2_MI_GFPve_pos -2.1016633588
## dCA2_Mpos_CTACGGGGTGACTCTA-1dCA2_MI_GFPve_pos -1.5177129584
## dCA2_Mpos_CTATCCGCACATACTG-1dCA2_MI_GFPve_pos -0.6946128088
## dCA2_Mpos_CTCAACCGTGAAGCGT-1dCA2_MI_GFPve_pos -1.4765062409
## dCA2_Mpos_CTCAGTCCAGATCCTA-1dCA2_MI_GFPve_pos -2.3021819669
## dCA2_Mpos_CTCAGTCTCCGACATA-1dCA2_MI_GFPve_pos -2.1815505105
## dCA2_Mpos_CTCCACAAGGTTCACT-1dCA2_MI_GFPve_pos -1.7478017884
## dCA2_Mpos_CTCCATGCACGCCACA-1dCA2_MI_GFPve_pos -2.1769957619
```

```
## dCA2_Mpos_CTCCATGCAGCTTCGG-1dCA2_MI_GFPve_pos -1.6872707682
## dCA2_Mpos_CTCCGATGTCAAGGCA-1dCA2_MI_GFPve_pos -2.4671151715
## dCA2_Mpos_CTCTGGTCACCGTCTT-1dCA2_MI_GFPve_pos -2.1658186990
## dCA2_Mpos_CTGCATCAGAAACTAC-1dCA2_MI_GFPve_pos -1.4267652112
## dCA2_Mpos_CTGCATCCAATAGAGT-1dCA2_MI_GFPve_pos -1.1873452740
## dCA2_Mpos_CTGCGAGCAATCTGCA-1dCA2_MI_GFPve_pos -2.1210889893
## dCA2_Mpos_CTGCTCAGTACCTATG-1dCA2_MI_GFPve_pos -2.4482309895
## dCA2_Mpos_CTGCTCATCGGAAACG-1dCA2_MI_GFPve_pos -1.1292904692
## dCA2_Mpos_CTGTACCGTCGTCGGT-1dCA2_MI_GFPve_pos -2.2053196507
## dCA2_Mpos_CTGTGAACAGTAGTGG-1dCA2_MI_GFPve_pos -1.1509966927
## dCA2_Mpos_CTTAGGAAGTATTGCC-1dCA2_MI_GFPve_pos -1.4873955327
## dCA2_Mpos_CTTCCTTTCTTACGGA-1dCA2_MI_GFPve_pos -1.8716076690
## dCA2_Mpos_CTTGAGACATAGCTGT-1dCA2_MI_GFPve_pos -1.5963691550
## dCA2_Mpos_CTTGATTTCGTGCGAC-1dCA2_MI_GFPve_pos -1.0953260737
## dCA2_Mpos_CTTTCAACAATTAGGA-1dCA2_MI_GFPve_pos -1.6859508830
## dCA2_Mpos_GAACACTAGTTCAACC-1dCA2_MI_GFPve_pos -2.1154654103
## dCA2_Mpos_GAACACTCATCCTCAC-1dCA2_MI_GFPve_pos -1.9561903554
## dCA2_Mpos_GAACTGTGTCAAGCCC-1dCA2_MI_GFPve_pos -1.9332237321
## dCA2_Mpos_GAACTGTGTTTGACAC-1dCA2_MI_GFPve_pos -2.4246349412
## dCA2_Mpos_GAAGAATTCGGATTAC-1dCA2_MI_GFPve_pos -2.5076806622
## dCA2_Mpos_GAAGCCCTCCGATCGG-1dCA2_MI_GFPve_pos -1.6623096543
## dCA2_Mpos_GAAGGACAGTTACGAA-1dCA2_MI_GFPve_pos -2.3791060525
## dCA2_Mpos_GAAGGACTCTACGCGG-1dCA2_MI_GFPve_pos -2.3276760655
## dCA2_Mpos_GACAGCCAGCTGTACT-1dCA2_MI_GFPve_pos -1.6408847647
## dCA2_Mpos_GACATCATCGGCTTCT-1dCA2_MI_GFPve_pos -2.0760083276
## dCA2_Mpos_GACCCAGGTCTCAAGT-1dCA2_MI_GFPve_pos -2.2342629510
## dCA2_Mpos_GACCGTGAGGACGCAT-1dCA2_MI_GFPve_pos -2.2908418256
## dCA2_Mpos_GACCGTGTCAGTGTCA-1dCA2_MI_GFPve_pos -2.2107904034
## dCA2_Mpos_GACGTTAAGCACTTTG-1dCA2_MI_GFPve_pos -1.9839429932
## dCA2_Mpos_GACGTTATCGCACGGT-1dCA2_MI_GFPve_pos -1.4570603448
## dCA2_Mpos_GACTTCCCAGCGAACA-1dCA2_MI_GFPve_pos -2.2179577428
## dCA2_Mpos_GACTTCCTCTGTGCGG-1dCA2_MI_GFPve_pos -1.5073856192
## dCA2_Mpos_GAGACTTCATATCTGG-1dCA2_MI_GFPve_pos -1.3718718367
## dCA2_Mpos_GAGCCTGAGCTAAACA-1dCA2_MI_GFPve_pos -2.0853960591
## dCA2_Mpos_GAGCCTGCATACACCA-1dCA2_MI_GFPve_pos -2.5553274232
## dCA2_Mpos_GAGCCTGGTCGGAACA-1dCA2_MI_GFPve_pos -1.9221856671
## dCA2_Mpos_GAGGCCTAGAGCATCG-1dCA2_MI_GFPve_pos -2.3996636945
## dCA2_Mpos_GAGGCCTTCATTTCGT-1dCA2_MI_GFPve_pos -2.0732853490
## dCA2_Mpos_GAGGGATCACAAGCCC-1dCA2_MI_GFPve_pos -1.9616038876
## dCA2_Mpos_GAGTCTACAATCGTCA-1dCA2_MI_GFPve_pos -1.9103610593
## dCA2_Mpos_GAGTCTAGTTTCTATC-1dCA2_MI_GFPve_pos -0.8420683223
## dCA2_Mpos_GAGTCTATCCACGAAT-1dCA2_MI_GFPve_pos -1.7651214677
## dCA2_Mpos_GAGTTACTCTCTATAC-1dCA2_MI_GFPve_pos -1.6241529065
## dCA2_Mpos_GATCAGTTCTGCCTCA-1dCA2_MI_GFPve_pos -2.1231947022
## dCA2_Mpos_GATGCTAAGTCACGAG-1dCA2_MI_GFPve_pos -0.6213874298
## dCA2_Mpos_GATGGAGTCAATCTCT-1dCA2_MI_GFPve_pos -2.2563757973
## dCA2_Mpos_GCAACATTCACCTCTG-1dCA2_MI_GFPve_pos -2.0584681111
## dCA2_Mpos_GCACGGTAGGTACAAT-1dCA2_MI_GFPve_pos -1.9984753209
## dCA2_Mpos_GCACGTGTCATGGCCG-1dCA2_MI_GFPve_pos -1.0943241196
## dCA2_Mpos_GCACTAAGTACAGAAT-1dCA2_MI_GFPve_pos -1.8961021977
## dCA2_Mpos_GCAGCCACACATACTG-1dCA2_MI_GFPve_pos -1.3575118857
## dCA2_Mpos_GCATCTCCAGACGGAT-1dCA2_MI_GFPve_pos -1.5922802764
## dCA2_Mpos_GCATGATCACCAGGTC-1dCA2_MI_GFPve_pos -2.1593906956
## dCA2_Mpos_GCCAACGAGTTGCTGT-1dCA2_MI_GFPve_pos -2.1378963071
## dCA2_Mpos_GCCAGTGTCAAAGGTA-1dCA2_MI_GFPve_pos -2.4639520722
## dCA2_Mpos_GCCATTCGTAATACCC-1dCA2_MI_GFPve_pos -2.3666267472
## dCA2_Mpos_GCCGATGGTCCGGACT-1dCA2_MI_GFPve_pos -1.1483031588
## dCA2_Mpos_GCGAGAACATATCTGG-1dCA2_MI_GFPve_pos -2.2468204575
## dCA2_Mpos_GCGATCGAGCCGATCC-1dCA2_MI_GFPve_pos -1.8636435347
## dCA2_Mpos_GCGATCGTCTTCCCAG-1dCA2_MI_GFPve_pos -1.5978781062
## dCA2_Mpos_GCGGAAAAGAAACACT-1dCA2_MI_GFPve_pos -1.6576232987
## dCA2_Mpos_GCTGCAGCAATGCTCA-1dCA2_MI_GFPve_pos -1.6815253573
## dCA2_Mpos_GCTGGGTTCATTCCTA-1dCA2_MI_GFPve_pos -1.3351386863
## dCA2_Mpos_GCTTCACAGGTTCCGC-1dCA2_MI_GFPve_pos -1.9618983346
## dCA2_Mpos_GCTTCACAGTTCCTGA-1dCA2_MI_GFPve_pos -1.5348643142
## dCA2_Mpos_GGAATCTTCGAAGGAC-1dCA2_MI_GFPve_pos -2.4796323853
## dCA2_Mpos_GGAGATGCAACCCGCA-1dCA2_MI_GFPve_pos -2.2578435021
## dCA2_Mpos_GGAGGATAGCAAGGAA-1dCA2_MI_GFPve_pos -1.1100876408
## dCA2_Mpos_GGATGTTAGAGAGCCT-1dCA2_MI_GFPve_pos -2.0686557370
## dCA2_Mpos_GGCAGTCCAAACACCT-1dCA2_MI_GFPve_pos -1.6268804150
```

```
## dCA2_Mpos_GGCAGTCGTAGTGGCA-1dCA2_MI_GFPve_pos -0.4328446465
## dCA2_Mpos_GGCGTCAAGAACAAGG-1dCA2_MI_GFPve_pos -1.6096931773
## dCA2_Mpos_GGCTTGGCACTTACAG-1dCA2_MI_GFPve_pos -2.3605313378
## dCA2_Mpos_GGGAAGTGTTAAAGTG-1dCA2_MI_GFPve_pos -1.4597319442
## dCA2_Mpos_GGGACAATCTCCCTAG-1dCA2_MI_GFPve_pos -2.4086804467
## dCA2_Mpos_GGGACCTGTCGATTCA-1dCA2_MI_GFPve_pos -2.1973636227
## dCA2_Mpos_GGGAGATAGTCAACAA-1dCA2_MI_GFPve_pos -1.8084367591
## dCA2_Mpos_GGGAGTAAGTCTAGAA-1dCA2_MI_GFPve_pos -1.7516795474
## dCA2_Mpos_GGGATGATCTGAGAGG-1dCA2_MI_GFPve_pos -2.3372316438
## dCA2_Mpos_GGGCCATAGGGTTTCT-1dCA2_MI_GFPve_pos -2.1935157853
## dCA2_Mpos_GGGCCATCAGGTGTTT-1dCA2_MI_GFPve_pos -2.3345294076
## dCA2_Mpos_GGGCTCACATTCACAG-1dCA2_MI_GFPve_pos -1.6321395713
## dCA2_Mpos_GGGTCACGTCTCTCAC-1dCA2_MI_GFPve_pos -1.3696781474
## dCA2_Mpos_GGGTGTCAGTACGTCT-1dCA2_MI_GFPve_pos -1.9344022351
## dCA2_Mpos_GGTAATCAGCGATTCT-1dCA2_MI_GFPve_pos -0.9325109797
## dCA2_Mpos_GGTGATTAGAGGCTGT-1dCA2_MI_GFPve_pos -1.3656355220
## dCA2_Mpos_GGTGTTAGTCCGAAAG-1dCA2_MI_GFPve_pos -1.6633287745
## dCA2_Mpos_GGTGTTAGTTTGTGGT-1dCA2_MI_GFPve_pos -1.9066472131
## dCA2_Mpos_GGTTAACGTATCCCTC-1dCA2_MI_GFPve_pos -1.9004478532
## dCA2_Mpos_GTAACACGTGTACAGG-1dCA2_MI_GFPve_pos -0.7736572104
## dCA2_Mpos_GTAACCAAGTATAACG-1dCA2_MI_GFPve_pos -2.3232333737
## dCA2_Mpos_GTAAGTCTCAAGCCAT-1dCA2_MI_GFPve_pos -2.1706256944
## dCA2_Mpos_GTACAGTAGGTGAGCT-1dCA2_MI_GFPve_pos -1.8610388118
## dCA2_Mpos_GTAGAAAAGACACACG-1dCA2_MI_GFPve_pos -2.3330748158
## dCA2_Mpos_GTAGAGGAGTTGCTCA-1dCA2_MI_GFPve_pos -2.3037035542
## dCA2_Mpos_GTATTGGTCGATTTCT-1dCA2_MI_GFPve_pos -1.2673806029
## dCA2_Mpos_GTATTTCTCCTGGGTG-1dCA2_MI_GFPve_pos -2.2680444794
## dCA2_Mpos_GTCAAACGTACATTGC-1dCA2_MI_GFPve_pos -2.0928549844
## dCA2_Mpos_GTCACGGAGCTAGTTC-1dCA2_MI_GFPve_pos -2.3635144311
## dCA2_Mpos_GTCACGGTCCGCACTT-1dCA2_MI_GFPve_pos -1.6500575620
## dCA2_Mpos_GTCAGCGAGAGCAGTC-1dCA2_MI_GFPve_pos -1.8321961242
## dCA2_Mpos_GTCATTTAGTCCGCCA-1dCA2_MI_GFPve_pos -1.5938159304
## dCA2_Mpos_GTCATTTCATAGACTC-1dCA2_MI_GFPve_pos -1.8833378630
## dCA2_Mpos_GTCCCATTCGTCCTCA-1dCA2_MI_GFPve_pos -1.8322194892
## dCA2_Mpos_GTCGCGAGTTGTACGT-1dCA2_MI_GFPve_pos -1.6304797011
## dCA2_Mpos_GTCGTTCCACGTCATA-1dCA2_MI_GFPve_pos -1.7369824725
## dCA2_Mpos_GTCTACCAGAAGTATC-1dCA2_MI_GFPve_pos -2.4089286404
## dCA2_Mpos_GTCTAGATCGCCTTGT-1dCA2_MI_GFPve_pos -1.8883084136
## dCA2_Mpos_GTCTCACAGTAAGAGG-1dCA2_MI_GFPve_pos -1.4448894339
## dCA2_Mpos_GTCTGTCAGCAGGCTA-1dCA2_MI_GFPve_pos -1.6637371856
## dCA2_Mpos_GTCTGTCGTCCGACGT-1dCA2_MI_GFPve_pos -1.9624435979
## dCA2_Mpos_GTGAGCCTCAGACCGC-1dCA2_MI_GFPve_pos -2.4367508965
## dCA2_Mpos_GTGAGCCTCTTACTGT-1dCA2_MI_GFPve_pos -2.4972648698
## dCA2_Mpos_GTGAGTTTCGTGCACG-1dCA2_MI_GFPve_pos -1.2049449759
## dCA2_Mpos_GTGCTGGGTGGTATGG-1dCA2_MI_GFPve_pos -1.1672501641
## dCA2_Mpos_GTGCTGGTCCAGTGTA-1dCA2_MI_GFPve_pos -0.6840469318
## dCA2_Mpos_GTGCTTCAGACCTGGA-1dCA2_MI_GFPve_pos -1.7642687636
## dCA2_Mpos_GTGGAGAGTATGAAGT-1dCA2_MI_GFPve_pos -2.0517966824
## dCA2_Mpos_GTGGAGAGTTCTATCT-1dCA2_MI_GFPve_pos -1.5138532000
## dCA2_Mpos_GTGTAACCAGTTCCAA-1dCA2_MI_GFPve_pos -2.1351060944
## dCA2_Mpos_GTGTGGCGTAGCTTTG-1dCA2_MI_GFPve_pos -0.4684237319
## dCA2_Mpos_GTTACGAGTACTTGTG-1dCA2_MI_GFPve_pos -2.0740902501
## dCA2_Mpos_GTTAGACGTACTGTTG-1dCA2_MI_GFPve_pos -1.0443321544
## dCA2_Mpos_GTTAGTGCAGGTATGG-1dCA2_MI_GFPve_pos -1.8944953757
## dCA2_Mpos_GTTCCGTGTGTAAACA-1dCA2_MI_GFPve_pos -1.7601705867
## dCA2_Mpos_GTTCGCTAGAGCCCAA-1dCA2_MI_GFPve_pos -2.2812564450
## dCA2_Mpos_GTTCGCTTCCAAACCA-1dCA2_MI_GFPve_pos -2.0948133546
## dCA2_Mpos_GTTGTCCGTAGACTGG-1dCA2_MI_GFPve_pos -1.4373200017
## dCA2_Mpos_TAACTTCGTGTTGAGG-1dCA2_MI_GFPve_pos -1.9406871873
## dCA2_Mpos_TAAGCGTAGAATACAC-1dCA2_MI_GFPve_pos -1.2048257666
## dCA2_Mpos_TAAGTCGTCTGTGTGA-1dCA2_MI_GFPve_pos -1.8909841853
## dCA2_Mpos_TACAACGTCCTGGTCT-1dCA2_MI_GFPve_pos -2.2308101731
## dCA2_Mpos_TACACCCCAGGTTCAT-1dCA2_MI_GFPve_pos -1.6127215701
## dCA2_Mpos_TACAGGTCATTCTTCA-1dCA2_MI_GFPve_pos -2.3975911218
## dCA2_Mpos_TACATTCCACATGTTG-1dCA2_MI_GFPve_pos -1.8496462184
## dCA2_Mpos_TACCTCGCAGGCGAAT-1dCA2_MI_GFPve_pos -2.3268020230
## dCA2_Mpos_TACCTCGGTCCAGTTA-1dCA2_MI_GFPve_pos -2.3913667279
## dCA2_Mpos_TACGGTAAGATGCAGC-1dCA2_MI_GFPve_pos -2.3642652112
## dCA2_Mpos_TACGGTAAGTGCCGAA-1dCA2_MI_GFPve_pos -1.8841544467
## dCA2_Mpos_TACTTACTCTGACCCT-1dCA2_MI_GFPve_pos -2.3960132676
```

```
## dCA2_Mpos_TAGACTGAGTAGCCAG-1dCA2_MI_GFPve_pos -1.8105816918
## dCA2_Mpos_TAGACTGTCTGTAACG-1dCA2_MI_GFPve_pos -1.8579850274
## dCA2_Mpos_TAGATCGAGCCGATTT-1dCA2_MI_GFPve_pos -1.9866180497
## dCA2_Mpos_TAGCACAGTGAATTAG-1dCA2_MI_GFPve_pos -2.3915228921
## dCA2_Mpos_TAGTGCAAGTATAGAC-1dCA2_MI_GFPve_pos -1.7319189387
## dCA2_Mpos_TATATCCGTTAAAGTG-1dCA2_MI_GFPve_pos -1.8457841950
## dCA2_Mpos_TATATCCTCGTCGATA-1dCA2_MI_GFPve_pos -1.5591046887
## dCA2_Mpos_TCAAGCAGTTTCGTTT-1dCA2_MI_GFPve_pos -2.3183822709
## dCA2_Mpos_TCAAGTGTCAGCCCAG-1dCA2_MI_GFPve_pos -1.6522910672
## dCA2_Mpos_TCAATTCTCCTCATAT-1dCA2_MI_GFPve_pos -0.5851279574
## dCA2_Mpos_TCACATTCATGATGCT-1dCA2_MI_GFPve_pos -2.3058860379
## dCA2_Mpos_TCACTATAGGAAGTCC-1dCA2_MI_GFPve_pos -1.6073962527
## dCA2_Mpos_TCACTATCAAAGGAGA-1dCA2_MI_GFPve_pos -1.0232184010
## dCA2_Mpos_TCAGGTAAGCCTCAAT-1dCA2_MI_GFPve_pos -2.1023633557
## dCA2_Mpos_TCAGGTACACAACGTT-1dCA2_MI_GFPve_pos -1.8851956206
## dCA2_Mpos_TCAGGTAGTGCCTGCA-1dCA2_MI_GFPve_pos -1.5353944379
## dCA2_Mpos_TCATCATAGCGACTTT-1dCA2_MI_GFPve_pos -2.2170963364
## dCA2_Mpos_TCATGGAAGTCCCTAA-1dCA2_MI_GFPve_pos -2.3557577210
## dCA2_Mpos_TCCACGTCATCCGAAT-1dCA2_MI_GFPve_pos -2.0453231412
## dCA2_Mpos_TCCAGAACAGGCCCTA-1dCA2_MI_GFPve_pos -2.3049194890
## dCA2_Mpos_TCCAGAATCCTGGCTT-1dCA2_MI_GFPve_pos -2.4691078740
## dCA2_Mpos_TCCATGCAGTCTAGAA-1dCA2_MI_GFPve_pos -2.4274523335
## dCA2_Mpos_TCCCATGGTGGCCACT-1dCA2_MI_GFPve_pos -0.9091578799
## dCA2_Mpos_TCCGAAACAGAGGTTG-1dCA2_MI_GFPve_pos -2.0547680932
## dCA2_Mpos_TCCGAAACATGATCTG-1dCA2_MI_GFPve_pos -2.4055779057
## dCA2_Mpos_TCCGATCCATGCCATA-1dCA2_MI_GFPve_pos -0.5864532071
## dCA2_Mpos_TCCTAATAGCGTGCCT-1dCA2_MI_GFPve_pos -2.4084549027
## dCA2_Mpos_TCCTCTTGTGCCAAGA-1dCA2_MI_GFPve_pos -1.9780457097
## dCA2_Mpos_TCGAAGTTCTACTATC-1dCA2_MI_GFPve_pos -1.5528091269
## dCA2_Mpos_TCGACGGTCACTCTTA-1dCA2_MI_GFPve_pos -2.3556544858
## dCA2_Mpos_TCGGATAAGATTGGGC-1dCA2_MI_GFPve_pos -1.6818287450
## dCA2_Mpos_TCGGATACACCTTCCA-1dCA2_MI_GFPve_pos -2.2484390813
## dCA2_Mpos_TCGGGCAGTTCAATCG-1dCA2_MI_GFPve_pos -1.6379520970
## dCA2_Mpos_TCGGGTGCATAGGAGC-1dCA2_MI_GFPve_pos -2.1081209260
## dCA2_Mpos_TCGGGTGGTAGGGTAC-1dCA2_MI_GFPve_pos -1.0265488702
## dCA2_Mpos_TCGTGGGTCGCACGAC-1dCA2_MI_GFPve_pos -1.5950884896
## dCA2_Mpos_TCTATCAGTGTCCACG-1dCA2_MI_GFPve_pos -1.7746543961
## dCA2_Mpos_TCTCAGCCATTGAGCT-1dCA2_MI_GFPve_pos -2.5142264443
## dCA2_Mpos_TCTGTCGCATAAGCGG-1dCA2_MI_GFPve_pos -1.7498056966
## dCA2_Mpos_TCTTAGTGTGCAACAG-1dCA2_MI_GFPve_pos -2.0671308118
## dCA2_Mpos_TCTTGCGAGAGTGTGC-1dCA2_MI_GFPve_pos -2.3296108323
## dCA2_Mpos_TCTTTGATCGTTAGTG-1dCA2_MI_GFPve_pos -1.9932739812
## dCA2_Mpos_TGAATCGAGCTAAATG-1dCA2_MI_GFPve_pos -1.9825368005
## dCA2_Mpos_TGAATCGAGTATGTAG-1dCA2_MI_GFPve_pos -2.2267005521
## dCA2_Mpos_TGACTCCAGTCCCTAA-1dCA2_MI_GFPve_pos -2.0106356221
## dCA2_Mpos_TGAGACTGTGTTATCG-1dCA2_MI_GFPve_pos -1.9290030080
## dCA2_Mpos_TGAGCGCCAAGGCAAC-1dCA2_MI_GFPve_pos -1.8422316390
## dCA2_Mpos_TGAGGGAAGGTAGACC-1dCA2_MI_GFPve_pos -1.7176963168
## dCA2_Mpos_TGATCAGAGATGGCGT-1dCA2_MI_GFPve_pos -1.4626026231
## dCA2_Mpos_TGATCTTGTAAGGAGA-1dCA2_MI_GFPve_pos -2.0262103158
## dCA2_Mpos_TGATGGTAGTCGGGAT-1dCA2_MI_GFPve_pos -1.8077123242
## dCA2_Mpos_TGCACGGGTCCATAGT-1dCA2_MI_GFPve_pos -1.9572405892
## dCA2_Mpos_TGCAGTACAAAGGAGA-1dCA2_MI_GFPve_pos -0.3832492011
## dCA2_Mpos_TGCGATAAGGCACCAA-1dCA2_MI_GFPve_pos -1.2578311044
## dCA2_Mpos_TGCGGGTAGGTCGAGT-1dCA2_MI_GFPve_pos -1.7760884839
## dCA2_Mpos_TGCTTCGAGTTCCATG-1dCA2_MI_GFPve_pos -1.9987518864
## dCA2_Mpos_TGGATCATCATCGTAG-1dCA2_MI_GFPve_pos  0.4092507583
## dCA2_Mpos_TGGATGTAGTGGAAGA-1dCA2_MI_GFPve_pos -1.3304814177
## dCA2_Mpos_TGGCGTGGTGCCTTTC-1dCA2_MI_GFPve_pos -2.4506263810
## dCA2_Mpos_TGGGAAGGTCTGCCTT-1dCA2_MI_GFPve_pos -2.4210851270
## dCA2_Mpos_TGGGAGAGTGGACAGT-1dCA2_MI_GFPve_pos -1.7519353705
## dCA2_Mpos_TGGTACAAGAGAGCCT-1dCA2_MI_GFPve_pos -2.0855002480
## dCA2_Mpos_TGGTGATCATTGACAC-1dCA2_MI_GFPve_pos -2.0855939465
## dCA2_Mpos_TGGTGATGTTGCTTGA-1dCA2_MI_GFPve_pos -2.2113552171
## dCA2_Mpos_TGTACAGAGCTGGCTC-1dCA2_MI_GFPve_pos -1.9313366490
## dCA2_Mpos_TGTGATGCAAGACAAT-1dCA2_MI_GFPve_pos  0.5996291262
## dCA2_Mpos_TGTGATGCACAGAGCA-1dCA2_MI_GFPve_pos -2.2881164628
## dCA2_Mpos_TGTGATGGTCGCATCG-1dCA2_MI_GFPve_pos -1.7467056590
## dCA2_Mpos_TGTGCGGAGACCATGG-1dCA2_MI_GFPve_pos -2.3419907170
## dCA2_Mpos_TGTGCGGTCTCTCAAT-1dCA2_MI_GFPve_pos -1.6456488448
```

```
## dCA2_Mpos_TGTGCGGTCTGCGATA-1dCA2_MI_GFPve_pos -1.4912402707
## dCA2_Mpos_TGTTCATAGCGATTCT-1dCA2_MI_GFPve_pos -0.9549965936
## dCA2_Mpos_TGTTGAGAGCCTCATA-1dCA2_MI_GFPve_pos -2.1857857781
## dCA2_Mpos_TTACCATTCCTTCACG-1dCA2_MI_GFPve_pos -1.8671882230
## dCA2_Mpos_TTCCAATTCCAGTACA-1dCA2_MI_GFPve_pos -2.2101085263
## dCA2_Mpos_TTCCACGCAAACTAAG-1dCA2_MI_GFPve_pos -1.9113285619
## dCA2_Mpos_TTCCTTCTCCCATACC-1dCA2_MI_GFPve_pos  0.4506524426
## dCA2_Mpos_TTCGCTGCAATCCTAG-1dCA2_MI_GFPve_pos -1.5614850598
## dCA2_Mpos_TTCGCTGTCCACGGGT-1dCA2_MI_GFPve_pos -2.1272888261
## dCA2_Mpos_TTCTGTAAGATGTTCC-1dCA2_MI_GFPve_pos -1.5659828263
## dCA2_Mpos_TTGCATTGTTACTCAG-1dCA2_MI_GFPve_pos -2.5659785348
## dCA2_Mpos_TTGCGTCTCATTCGGA-1dCA2_MI_GFPve_pos -2.2420821267
## dCA2_Mpos_TTGCTGCCAACTTCTT-1dCA2_MI_GFPve_pos -1.9543266373
## dCA2_Mpos_TTGCTGCTCAAACCTG-1dCA2_MI_GFPve_pos -1.9261300641
## dCA2_Mpos_TTGGGTATCTACAGGT-1dCA2_MI_GFPve_pos -1.5776134806
## dCA2_Mpos_TTTACCAAGAGTCACG-1dCA2_MI_GFPve_pos -1.4905617314
## dCA2_Mpos_TTTACGTAGGTCACAG-1dCA2_MI_GFPve_pos -1.4837226945
## dCA2_Mpos_TTTCAGTCAGACACCC-1dCA2_MI_GFPve_pos -1.6037634688
## dCA2_Mpos_TTTGACTAGAGGATCC-1dCA2_MI_GFPve_pos -1.4671911078
## dCA2_Mpos_TTTGATCTCACAAGGG-1dCA2_MI_GFPve_pos -1.6287797766
## dCA2_Mpos_TTTGTTGCAGTTGAAA-1dCA2_MI_GFPve_pos -1.0371891337
## dCA2_Fpos_AAACGAAGTACCCACG-1dCA2_FI_GFPve_pos -0.5369273501
## dCA2_Fpos_AAACGCTGTTATCTTC-1dCA2_FI_GFPve_pos -2.1388356763
## dCA2_Fpos_AAAGTGAAGTAAACGT-1dCA2_FI_GFPve_pos -1.4857939559
## dCA2_Fpos_AACAAGAAGCTTGTTG-1dCA2_FI_GFPve_pos -2.0511171895
## dCA2_Fpos_AACACACAGCAAATCA-1dCA2_FI_GFPve_pos -0.4086249548
## dCA2_Fpos_AACACACGTCTTGCTC-1dCA2_FI_GFPve_pos -1.9146738129
## dCA2_Fpos_AACCAACCATCGTGGC-1dCA2_FI_GFPve_pos -2.2752990800
## dCA2_Fpos_AACCACAAGATGCAGC-1dCA2_FI_GFPve_pos -1.6746425706
## dCA2_Fpos_AACCTGACAGAGGTAC-1dCA2_FI_GFPve_pos -2.2232580262
## dCA2_Fpos_AACCTGAGTTTAAGGA-1dCA2_FI_GFPve_pos -1.7343838292
## dCA2_Fpos_AAGACAAAGCACCGTC-1dCA2_FI_GFPve_pos -2.1515839177
## dCA2_Fpos_AAGACTCAGTATGGAT-1dCA2_FI_GFPve_pos -2.2378139573
## dCA2_Fpos_AAGGAATTCACTGAAC-1dCA2_FI_GFPve_pos -1.9633634167
## dCA2_Fpos_AAGTACCAGACATCAA-1dCA2_FI_GFPve_pos -2.4384346085
## dCA2_Fpos_AAGTACCTCTCATTGT-1dCA2_FI_GFPve_pos -2.4357030469
## dCA2_Fpos_AAGTCGTTCTCCTGTG-1dCA2_FI_GFPve_pos -1.5789337235
## dCA2_Fpos_AATGACCTCCGTGGCA-1dCA2_FI_GFPve_pos -1.8431320267
## dCA2_Fpos_AATGGCTAGAATTCAG-1dCA2_FI_GFPve_pos -2.1137292462
## dCA2_Fpos_AATGGCTCACGCTTAA-1dCA2_FI_GFPve_pos -1.7768006402
## dCA2_Fpos_AATGGCTCAGGATGAC-1dCA2_FI_GFPve_pos -1.4377251941
## dCA2_Fpos_AATGGCTCATGGAAGC-1dCA2_FI_GFPve_pos -1.9546470719
## dCA2_Fpos_AATGGCTGTATTGACC-1dCA2_FI_GFPve_pos -1.9913034516
## dCA2_Fpos_AATGGCTTCGTTACCC-1dCA2_FI_GFPve_pos -1.8430756408
## dCA2_Fpos_AATTCCTCACCGTCTT-1dCA2_FI_GFPve_pos -2.2760315018
## dCA2_Fpos_AATTCCTCATGACAGG-1dCA2_FI_GFPve_pos -1.6976285058
## dCA2_Fpos_ACAAAGACAGTTCCAA-1dCA2_FI_GFPve_pos -2.0983007031
## dCA2_Fpos_ACAGCCGGTTCAGTAC-1dCA2_FI_GFPve_pos -1.6293653327
## dCA2_Fpos_ACCAACAAGAGTCTGG-1dCA2_FI_GFPve_pos -1.6921438056
## dCA2_Fpos_ACCACAAAGGTGATCG-1dCA2_FI_GFPve_pos -2.1570475179
## dCA2_Fpos_ACCATTTAGCTTTCTT-1dCA2_FI_GFPve_pos -2.1240599232
## dCA2_Fpos_ACCCAAAGTTCTCACC-1dCA2_FI_GFPve_pos -1.5688883143
## dCA2_Fpos_ACGGGTCTCGAAGGAC-1dCA2_FI_GFPve_pos -1.6874398070
## dCA2_Fpos_ACGGTTAGTGACACAG-1dCA2_FI_GFPve_pos -1.7197189408
## dCA2_Fpos_ACTATTCCAATACAGA-1dCA2_FI_GFPve_pos -2.2253809052
## dCA2_Fpos_ACTATTCTCTTCTGTA-1dCA2_FI_GFPve_pos -2.0937838631
## dCA2_Fpos_ACTCTCGGTGCGCTCA-1dCA2_FI_GFPve_pos -1.6625746566
## dCA2_Fpos_AGACAGGTCACAAGAA-1dCA2_FI_GFPve_pos -2.3737094479
## dCA2_Fpos_AGAGCAGGTGTTATCG-1dCA2_FI_GFPve_pos -1.6035223084
## dCA2_Fpos_AGAGCCCGTCTGCGCA-1dCA2_FI_GFPve_pos -1.8362983542
## dCA2_Fpos_AGATAGAGTGGATCAG-1dCA2_FI_GFPve_pos -1.7259950715
## dCA2_Fpos_AGCATCAAGTAGGGTC-1dCA2_FI_GFPve_pos -1.4909216243
## dCA2_Fpos_AGCGATTCACTGGCCA-1dCA2_FI_GFPve_pos -1.2045697051
## dCA2_Fpos_AGGAATACACCGCTAG-1dCA2_FI_GFPve_pos -1.8997595387
## dCA2_Fpos_AGGAATACACGGTAGA-1dCA2_FI_GFPve_pos -2.1454992371
## dCA2_Fpos_AGGAGGTAGGATACAT-1dCA2_FI_GFPve_pos -1.3746537047
## dCA2_Fpos_AGGATCTTCCACGTGG-1dCA2_FI_GFPve_pos -1.4983350116
## dCA2_Fpos_AGGCATTAGAGAGCCT-1dCA2_FI_GFPve_pos -1.4640110808
## dCA2_Fpos_AGGGCCTGTCTAACGT-1dCA2_FI_GFPve_pos -2.0036766606
## dCA2_Fpos_AGGGCCTTCTGAGGCC-1dCA2_FI_GFPve_pos -1.8638092356
```

```
## dCA2_Fpos_AGGTAGGGTAGTCCTA-1dCA2_FI_GFPve_pos -1.7262665110
## dCA2_Fpos_AGGTAGGGTTGTAGCT-1dCA2_FI_GFPve_pos -1.6220774728
## dCA2_Fpos_AGGTTGTCACTTCCTG-1dCA2_FI_GFPve_pos -1.9920489865
## dCA2_Fpos_AGTAACCAGGTGGTTG-1dCA2_FI_GFPve_pos -1.6716350394
## dCA2_Fpos_AGTCAACCACCAGTTA-1dCA2_FI_GFPve_pos -1.9196839410
## dCA2_Fpos_AGTCATGTCTGTCGTC-1dCA2_FI_GFPve_pos -1.5471260625
## dCA2_Fpos_AGTGCCGGTGCTGTCG-1dCA2_FI_GFPve_pos -1.4863971549
## dCA2_Fpos_ATACTTCCACCTATCC-1dCA2_FI_GFPve_pos -2.0294351655
## dCA2_Fpos_ATATCCTCATACACCA-1dCA2_FI_GFPve_pos -1.8291435319
## dCA2_Fpos_ATCACTTGTCTAACGT-1dCA2_FI_GFPve_pos -1.2530114728
## dCA2_Fpos_ATCAGGTCATACTGAC-1dCA2_FI_GFPve_pos -1.8938201743
## dCA2_Fpos_ATCCACCGTACAAAGT-1dCA2_FI_GFPve_pos -1.8466659861
## dCA2_Fpos_ATCGCCTAGGTCATAA-1dCA2_FI_GFPve_pos -0.9367760497
## dCA2_Fpos_ATCGCCTGTACTTGTG-1dCA2_FI_GFPve_pos -1.3458662110
## dCA2_Fpos_ATCGTCCCACCGAATT-1dCA2_FI_GFPve_pos -2.4498260098
## dCA2_Fpos_ATCGTCCTCTTGGTGA-1dCA2_FI_GFPve_pos -1.3626110631
## dCA2_Fpos_ATCGTGATCATTCCTA-1dCA2_FI_GFPve_pos -1.4713836985
## dCA2_Fpos_ATCTCTAAGAGGCTGT-1dCA2_FI_GFPve_pos -1.5460282641
## dCA2_Fpos_ATGATCGGTGGAGAAA-1dCA2_FI_GFPve_pos -1.7806152182
## dCA2_Fpos_ATGGGTTCAGGGACTA-1dCA2_FI_GFPve_pos -1.7915593463
## dCA2_Fpos_ATTACTCTCCTACCAC-1dCA2_FI_GFPve_pos -1.6982514935
## dCA2_Fpos_ATTCCTAGTAAGCAAT-1dCA2_FI_GFPve_pos -1.5009937363
## dCA2_Fpos_ATTCGTTTCGTACACA-1dCA2_FI_GFPve_pos -0.7543426829
## dCA2_Fpos_ATTCTACAGTCGTCTA-1dCA2_FI_GFPve_pos -1.7832601147
## dCA2_Fpos_ATTGTTCCATCCAACA-1dCA2_FI_GFPve_pos -2.1676950532
## dCA2_Fpos_ATTTACCGTGTTCATG-1dCA2_FI_GFPve_pos -1.8144075948
## dCA2_Fpos_ATTTCACAGCATCCTA-1dCA2_FI_GFPve_pos -1.9888763505
## dCA2_Fpos_CAAGCTATCGTTATCT-1dCA2_FI_GFPve_pos -2.0036397057
## dCA2_Fpos_CACAACAAGCACTGGA-1dCA2_FI_GFPve_pos -1.1558442193
## dCA2_Fpos_CACAACAGTAGCTGAG-1dCA2_FI_GFPve_pos -2.4507799226
## dCA2_Fpos_CACACAAGTGAGTTTC-1dCA2_FI_GFPve_pos -1.4753120023
## dCA2_Fpos_CACTGGGTCTGGGCGT-1dCA2_FI_GFPve_pos -1.9700303155
## dCA2_Fpos_CAGCGTGAGCGCCATC-1dCA2_FI_GFPve_pos -2.5366087037
## dCA2_Fpos_CATAGACTCACCATAG-1dCA2_FI_GFPve_pos -2.1697082597
## dCA2_Fpos_CATCCACCACCCTATC-1dCA2_FI_GFPve_pos -2.3038234788
## dCA2_Fpos_CATCCGTAGCCTGTGC-1dCA2_FI_GFPve_pos -1.7967641431
## dCA2_Fpos_CATGCGGTCACTCTTA-1dCA2_FI_GFPve_pos -1.8787747699
## dCA2_Fpos_CATGCGGTCATTGGTG-1dCA2_FI_GFPve_pos -1.1124364215
## dCA2_Fpos_CATTCATCAAGAATAC-1dCA2_FI_GFPve_pos -1.2383241731
## dCA2_Fpos_CATTCATTCATTACGG-1dCA2_FI_GFPve_pos -1.2705098468
## dCA2_Fpos_CATTCTAGTTTCACAG-1dCA2_FI_GFPve_pos -2.3791096288
## dCA2_Fpos_CATTGAGCAGATTCGT-1dCA2_FI_GFPve_pos -1.6975731927
## dCA2_Fpos_CATTGTTGTGTGTGGA-1dCA2_FI_GFPve_pos -1.8805446702
## dCA2_Fpos_CCAAGCGCATCGAAGG-1dCA2_FI_GFPve_pos -1.9750027734
## dCA2_Fpos_CCACCATTCTGCAGCG-1dCA2_FI_GFPve_pos -1.5034569579
## dCA2_Fpos_CCCAACTCACCTCTGT-1dCA2_FI_GFPve_pos -1.5810843783
## dCA2_Fpos_CCCATTGCAATTGTGC-1dCA2_FI_GFPve_pos -1.9800424653
## dCA2_Fpos_CCCTTAGCACTGGACC-1dCA2_FI_GFPve_pos -1.6723972636
## dCA2_Fpos_CCTAACCAGAGTGAAG-1dCA2_FI_GFPve_pos -1.7281522828
## dCA2_Fpos_CCTACGTAGTGCACTT-1dCA2_FI_GFPve_pos -1.7670604068
## dCA2_Fpos_CCTCAGTAGATAGCAT-1dCA2_FI_GFPve_pos -1.1584004241
## dCA2_Fpos_CCTCAGTAGATGAATC-1dCA2_FI_GFPve_pos -2.0602009373
## dCA2_Fpos_CCTCTCCAGCCTCAGC-1dCA2_FI_GFPve_pos -1.9355173188
## dCA2_Fpos_CCTGCATAGTGTTGTC-1dCA2_FI_GFPve_pos -1.4661580401
## dCA2_Fpos_CCTGTTGGTCAGGCAA-1dCA2_FI_GFPve_pos -1.0175883847
## dCA2_Fpos_CCTTCAGAGTACAGCG-1dCA2_FI_GFPve_pos -1.4519221860
## dCA2_Fpos_CGAGAAGCAAGCGCAA-1dCA2_FI_GFPve_pos -1.8069789487
## dCA2_Fpos_CGCCAGAAGGCATCTT-1dCA2_FI_GFPve_pos -1.3624255734
## dCA2_Fpos_CGGGACTCATTAGGCT-1dCA2_FI_GFPve_pos -1.3241354304
## dCA2_Fpos_CGGGTCAGTGTCTTAG-1dCA2_FI_GFPve_pos -1.5005920010
## dCA2_Fpos_CGTAAGTGTGTTACTG-1dCA2_FI_GFPve_pos -1.8901382762
## dCA2_Fpos_CGTAATGCAATTCACG-1dCA2_FI_GFPve_pos -2.2374916154
## dCA2_Fpos_CTAAGTGAGACGCCAA-1dCA2_FI_GFPve_pos  0.8237254543
## dCA2_Fpos_CTACCCACAAAGACGC-1dCA2_FI_GFPve_pos -1.2695666629
## dCA2_Fpos_CTACCCACATGGGATG-1dCA2_FI_GFPve_pos -1.7767756062
## dCA2_Fpos_CTACCTGTCACTGTTT-1dCA2_FI_GFPve_pos -1.6073130446
## dCA2_Fpos_CTACTATGTATCGCGC-1dCA2_FI_GFPve_pos -1.4345289546
## dCA2_Fpos_CTACTATTCGTTCCCA-1dCA2_FI_GFPve_pos -1.7557321864
## dCA2_Fpos_CTATAGGGTACGCGTC-1dCA2_FI_GFPve_pos -1.0133417922
## dCA2_Fpos_CTCAGAAGTGACCGAA-1dCA2_FI_GFPve_pos -1.5389093238
```

```
## dCA2_Fpos_CTCAGGGAGCTAAATG-1dCA2_FI_GFPve_pos -2.2860643941
## dCA2_Fpos_CTCATGCCAGAATTCC-1dCA2_FI_GFPve_pos -1.6103180724
## dCA2_Fpos_CTCATTAGTCTACTGA-1dCA2_FI_GFPve_pos -2.2488346177
## dCA2_Fpos_CTCCATGTCATAGAGA-1dCA2_FI_GFPve_pos -1.8155058699
## dCA2_Fpos_CTCCCTCTCTGTTGGA-1dCA2_FI_GFPve_pos -1.7236301976
## dCA2_Fpos_CTCCGATTCGCGAAGA-1dCA2_FI_GFPve_pos -1.7373378354
## dCA2_Fpos_CTCCTCCCAAGACAAT-1dCA2_FI_GFPve_pos -1.6389291364
## dCA2_Fpos_CTCCTCCGTTAGGACG-1dCA2_FI_GFPve_pos -1.2362058240
## dCA2_Fpos_CTCGAGGCACAACGAG-1dCA2_FI_GFPve_pos -2.5449543076
## dCA2_Fpos_CTCGAGGGTCTCAAGT-1dCA2_FI_GFPve_pos -2.3268911916
## dCA2_Fpos_CTGAATGAGGTAAAGG-1dCA2_FI_GFPve_pos  0.3098735136
## dCA2_Fpos_CTGAATGCAACCTATG-1dCA2_FI_GFPve_pos -2.4811165410
## dCA2_Fpos_CTGATCCTCTTCTCAA-1dCA2_FI_GFPve_pos -1.2873609143
## dCA2_Fpos_CTGCCATAGGGCAATC-1dCA2_FI_GFPve_pos -2.0480434972
## dCA2_Fpos_CTGCCATAGTACCCTA-1dCA2_FI_GFPve_pos -1.4817096072
## dCA2_Fpos_CTGCCATGTTGTAAAG-1dCA2_FI_GFPve_pos -2.4523334580
## dCA2_Fpos_CTGCGAGCAGGTCAGA-1dCA2_FI_GFPve_pos -0.5268648940
## dCA2_Fpos_CTGGACGGTCAAGCCC-1dCA2_FI_GFPve_pos -2.1697299557
## dCA2_Fpos_CTGGCAGCAGCTCTGG-1dCA2_FI_GFPve_pos -1.8699472028
## dCA2_Fpos_CTGGCAGTCAAGCCGC-1dCA2_FI_GFPve_pos -1.6367691832
## dCA2_Fpos_CTGGTCTAGAGGTGCT-1dCA2_FI_GFPve_pos -1.7884157973
## dCA2_Fpos_CTGTCGTGTACAACGG-1dCA2_FI_GFPve_pos -1.8255221921
## dCA2_Fpos_CTGTGAAAGCGGATCA-1dCA2_FI_GFPve_pos -1.6757720786
## dCA2_Fpos_CTTCAATAGCGCCCAT-1dCA2_FI_GFPve_pos -2.0128903466
## dCA2_Fpos_CTTCGGTGTCGTGTTA-1dCA2_FI_GFPve_pos -1.6440724212
## dCA2_Fpos_GAAGGGTAGAAGCTCG-1dCA2_FI_GFPve_pos -1.4508311826
## dCA2_Fpos_GAATAGACAGGTCAGA-1dCA2_FI_GFPve_pos -1.6685240346
## dCA2_Fpos_GACATCAGTAGAATAC-1dCA2_FI_GFPve_pos -1.5755853730
## dCA2_Fpos_GACCCTTAGCTCCGAC-1dCA2_FI_GFPve_pos  0.0201096681
## dCA2_Fpos_GAGAGGTCAGGTGACA-1dCA2_FI_GFPve_pos -2.0863711911
## dCA2_Fpos_GAGGCCTCAGATTTCG-1dCA2_FI_GFPve_pos -1.6577452498
## dCA2_Fpos_GAGTCATTCCAAGGGA-1dCA2_FI_GFPve_pos -1.7730211097
## dCA2_Fpos_GAGTGAGAGATTGAGT-1dCA2_FI_GFPve_pos -1.7447953301
## dCA2_Fpos_GAGTTTGGTATTCCTT-1dCA2_FI_GFPve_pos -2.1805272179
## dCA2_Fpos_GATCACAGTAGCCCTG-1dCA2_FI_GFPve_pos -1.8256219703
## dCA2_Fpos_GATTCGACAGTTTCGA-1dCA2_FI_GFPve_pos -2.3298554498
## dCA2_Fpos_GCATCTCTCACCTCTG-1dCA2_FI_GFPve_pos -1.9718723374
## dCA2_Fpos_GCATGATGTGAACGGT-1dCA2_FI_GFPve_pos -1.5099873620
## dCA2_Fpos_GCCAACGAGCTGCCTG-1dCA2_FI_GFPve_pos -2.2522816735
## dCA2_Fpos_GCCAGGTAGATGACAT-1dCA2_FI_GFPve_pos -1.8159045058
## dCA2_Fpos_GCCTGTTCACATATGC-1dCA2_FI_GFPve_pos -2.2318842488
## dCA2_Fpos_GCGTGCATCGTTAGTG-1dCA2_FI_GFPve_pos -1.7512011605
## dCA2_Fpos_GGAGATGCACTTGTCC-1dCA2_FI_GFPve_pos -1.4420064765
## dCA2_Fpos_GGAGATGTCCCAGTGG-1dCA2_FI_GFPve_pos -2.2320921498
## dCA2_Fpos_GGCTGTGCACTAGGTT-1dCA2_FI_GFPve_pos  0.0032708642
## dCA2_Fpos_GGCTGTGCAGAACTTC-1dCA2_FI_GFPve_pos -1.6153993684
## dCA2_Fpos_GGGCGTTGTGGTACAG-1dCA2_FI_GFPve_pos -1.7565010863
## dCA2_Fpos_GGGTCACGTACTAGCT-1dCA2_FI_GFPve_pos -1.7105579453
## dCA2_Fpos_GGGTCTGCAGAGTGTG-1dCA2_FI_GFPve_pos -1.8162908631
## dCA2_Fpos_GGTGTTAAGGGTTAGC-1dCA2_FI_GFPve_pos -2.1614394265
## dCA2_Fpos_GTAACACGTTCCATTT-1dCA2_FI_GFPve_pos -1.4390391188
## dCA2_Fpos_GTAAGTCAGGTGCCAA-1dCA2_FI_GFPve_pos -1.8052960711
## dCA2_Fpos_GTAATCGCAAGGTTGG-1dCA2_FI_GFPve_pos -1.7201205569
## dCA2_Fpos_GTACAACAGATGTAGT-1dCA2_FI_GFPve_pos -2.1128342229
## dCA2_Fpos_GTATTGGGTCCGGTCA-1dCA2_FI_GFPve_pos -2.0346620160
## dCA2_Fpos_GTATTTCGTAGCGCCT-1dCA2_FI_GFPve_pos -1.5760650712
## dCA2_Fpos_GTATTTCGTTACAGCT-1dCA2_FI_GFPve_pos -2.4666657525
## dCA2_Fpos_GTCACGGAGGTGAGAA-1dCA2_FI_GFPve_pos -1.7928261834
## dCA2_Fpos_GTCATTTAGGCCCGTT-1dCA2_FI_GFPve_pos -0.1867594796
## dCA2_Fpos_GTCCTCATCTGAGTCA-1dCA2_FI_GFPve_pos -1.4444373923
## dCA2_Fpos_GTCTACCCACTCCACT-1dCA2_FI_GFPve_pos -2.2512292939
## dCA2_Fpos_GTCTTTAAGCGTGAGT-1dCA2_FI_GFPve_pos -1.9667611199
## dCA2_Fpos_GTGAGTTCAAAGGGTC-1dCA2_FI_GFPve_pos -2.1743724423
## dCA2_Fpos_GTGATGTTCGACCTAA-1dCA2_FI_GFPve_pos -1.9885261136
## dCA2_Fpos_GTGCTTCAGACTTCGT-1dCA2_FI_GFPve_pos -1.4406765776
## dCA2_Fpos_GTGGAGATCTTAATCC-1dCA2_FI_GFPve_pos -2.2855393964
## dCA2_Fpos_GTGGTTACACTTGGGC-1dCA2_FI_GFPve_pos -1.9289333897
## dCA2_Fpos_GTGTAACAGAGAGTGA-1dCA2_FI_GFPve_pos -1.9199335652
## dCA2_Fpos_GTTACAGTCCAAATGC-1dCA2_FI_GFPve_pos -2.1506929475
## dCA2_Fpos_GTTGCTCCACGCGTCA-1dCA2_FI_GFPve_pos -1.9595577794
```

```
## dCA2_Fpos_GTTGCTCTCCTAGCTC-1dCA2_FI_GFPve_pos -1.5893838483
## dCA2_Fpos_GTTGTAGCACAGACGA-1dCA2_FI_GFPve_pos -1.1146940070
## dCA2_Fpos_GTTGTGATCATTCTTG-1dCA2_FI_GFPve_pos -2.5340540486
## dCA2_Fpos_GTTTACTCAGCTGTTA-1dCA2_FI_GFPve_pos -1.5874199944
## dCA2_Fpos_GTTTGGAGTGCCTTCT-1dCA2_FI_GFPve_pos -2.2040124017
## dCA2_Fpos_TAACGACAGGTACTGG-1dCA2_FI_GFPve_pos -2.3510177213
## dCA2_Fpos_TAATTCCTCGGCCAAC-1dCA2_FI_GFPve_pos -1.3686831075
## dCA2_Fpos_TACACCCAGGATATGT-1dCA2_FI_GFPve_pos -1.8203097659
## dCA2_Fpos_TACACCCTCCAACCGG-1dCA2_FI_GFPve_pos -0.9904481250
## dCA2_Fpos_TACCCACGTTCGATTG-1dCA2_FI_GFPve_pos -1.4141392785
## dCA2_Fpos_TACCCGTGTCTGCGCA-1dCA2_FI_GFPve_pos -1.9716653901
## dCA2_Fpos_TACCGGGTCACTCTTA-1dCA2_FI_GFPve_pos -2.0167152959
## dCA2_Fpos_TACCTCGCACAACGAG-1dCA2_FI_GFPve_pos -2.4168403226
## dCA2_Fpos_TACGGGCCAGACCGCT-1dCA2_FI_GFPve_pos -1.0545971471
## dCA2_Fpos_TACGTCCTCCAGTTCC-1dCA2_FI_GFPve_pos -2.3463955003
## dCA2_Fpos_TATTGCTGTACATTGC-1dCA2_FI_GFPve_pos -2.1502468663
## dCA2_Fpos_TCAAGACGTTCGGGTC-1dCA2_FI_GFPve_pos -1.3119672614
## dCA2_Fpos_TCACACCTCCATACAG-1dCA2_FI_GFPve_pos -2.1465840417
## dCA2_Fpos_TCACGCTGTATCTTCT-1dCA2_FI_GFPve_pos -1.9730029183
## dCA2_Fpos_TCACGGGAGGGTTGCA-1dCA2_FI_GFPve_pos -1.2139630395
## dCA2_Fpos_TCAGCCTCACGACGAA-1dCA2_FI_GFPve_pos -2.0673806744
## dCA2_Fpos_TCAGCCTCATACAGCT-1dCA2_FI_GFPve_pos -2.1596834737
## dCA2_Fpos_TCAGTTTTCACACCCT-1dCA2_FI_GFPve_pos -1.9447124081
## dCA2_Fpos_TCATACTCAAGGCAAC-1dCA2_FI_GFPve_pos -2.0389778691
## dCA2_Fpos_TCATATCCACACTTAG-1dCA2_FI_GFPve_pos -1.5901342707
## dCA2_Fpos_TCATGAGGTGTCATTG-1dCA2_FI_GFPve_pos -1.9913919049
## dCA2_Fpos_TCATGAGGTTGCGAAG-1dCA2_FI_GFPve_pos -1.3681998330
## dCA2_Fpos_TCATGGATCCCGAGTG-1dCA2_FI_GFPve_pos -1.4994198161
## dCA2_Fpos_TCATTGTCAATAACGA-1dCA2_FI_GFPve_pos -1.7408124286
## dCA2_Fpos_TCCCATGAGGGCCTCT-1dCA2_FI_GFPve_pos -2.2993831712
## dCA2_Fpos_TCGACCTCACTGGACC-1dCA2_FI_GFPve_pos -2.3644907552
## dCA2_Fpos_TCGACCTCAGAATCGG-1dCA2_FI_GFPve_pos -1.8554189282
## dCA2_Fpos_TCGACCTTCCGTCACT-1dCA2_FI_GFPve_pos -1.8938114720
## dCA2_Fpos_TCGATTTAGAGGACTC-1dCA2_FI_GFPve_pos -2.1567912179
## dCA2_Fpos_TCGATTTCAGTCGCAC-1dCA2_FI_GFPve_pos -1.9346418458
## dCA2_Fpos_TCGCAGGAGTAGGATT-1dCA2_FI_GFPve_pos -1.1657030659
## dCA2_Fpos_TCGTCCAGTCGAACGA-1dCA2_FI_GFPve_pos -2.3474941331
## dCA2_Fpos_TCTCACGCATAGGTAA-1dCA2_FI_GFPve_pos -2.2599923688
## dCA2_Fpos_TCTGCCATCAGGAGAC-1dCA2_FI_GFPve_pos -1.8457564193
## dCA2_Fpos_TCTGGCTAGTAGTGCG-1dCA2_FI_GFPve_pos -1.3116885501
## dCA2_Fpos_TGAACGTGTTGCGTAT-1dCA2_FI_GFPve_pos -1.8189795094
## dCA2_Fpos_TGAATCGAGAAGCTGC-1dCA2_FI_GFPve_pos -2.1169028359
## dCA2_Fpos_TGAATCGGTCAAAGTA-1dCA2_FI_GFPve_pos -2.2054379063
## dCA2_Fpos_TGACAGTTCATGTCTT-1dCA2_FI_GFPve_pos -2.1674025136
## dCA2_Fpos_TGACGCGAGGTTTACC-1dCA2_FI_GFPve_pos -2.0442624169
## dCA2_Fpos_TGACGCGGTGTTACAC-1dCA2_FI_GFPve_pos -1.6963775235
## dCA2_Fpos_TGAGCGCTCTCGTGGG-1dCA2_FI_GFPve_pos -1.3392058688
## dCA2_Fpos_TGATGGTGTGGTTCTA-1dCA2_FI_GFPve_pos -1.0503823834
## dCA2_Fpos_TGCACGGGTCTTACAG-1dCA2_FI_GFPve_pos -2.5282600003
## dCA2_Fpos_TGCAGATTCCCTCAAC-1dCA2_FI_GFPve_pos -2.4700396138
## dCA2_Fpos_TGCCGAGGTCTGTCCT-1dCA2_FI_GFPve_pos -0.5601184445
## dCA2_Fpos_TGCGATAAGGGTACAC-1dCA2_FI_GFPve_pos -2.0455262738
## dCA2_Fpos_TGCTCGTGTCAGACGA-1dCA2_FI_GFPve_pos -1.8328005153
## dCA2_Fpos_TGCTCGTGTGGCATCC-1dCA2_FI_GFPve_pos -1.5808577615
## dCA2_Fpos_TGGATGTTCGATACTG-1dCA2_FI_GFPve_pos -1.3252192813
## dCA2_Fpos_TGGGAAGAGTATAGAC-1dCA2_FI_GFPve_pos -2.1243701058
## dCA2_Fpos_TGGGTTATCGAGAAAT-1dCA2_FI_GFPve_pos -1.3678779679
## dCA2_Fpos_TGGTTAGCATCCCACT-1dCA2_FI_GFPve_pos -1.9568369466
## dCA2_Fpos_TGTAGACCAGGAACCA-1dCA2_FI_GFPve_pos -2.2809565144
## dCA2_Fpos_TGTCCACAGACTAAGT-1dCA2_FI_GFPve_pos -1.6246504861
## dCA2_Fpos_TGTCCACCAAGTATAG-1dCA2_FI_GFPve_pos -2.2552135068
## dCA2_Fpos_TGTCCTGGTCCACACG-1dCA2_FI_GFPve_pos -0.6969369131
## dCA2_Fpos_TGTTGAGTCATTACCT-1dCA2_FI_GFPve_pos -1.8303583937
## dCA2_Fpos_TTACCATCACGCTTAA-1dCA2_FI_GFPve_pos -1.8383365947
## dCA2_Fpos_TTACTGTAGCGGTAGT-1dCA2_FI_GFPve_pos -1.8578764277
## dCA2_Fpos_TTCAGGACACATGTTG-1dCA2_FI_GFPve_pos -2.2380311566
## dCA2_Fpos_TTCCACGCAGGTGACA-1dCA2_FI_GFPve_pos -2.2372367459
## dCA2_Fpos_TTCCGGTCAGACCATT-1dCA2_FI_GFPve_pos -2.0840258675
## dCA2_Fpos_TTCCTTCGTCACATTG-1dCA2_FI_GFPve_pos -1.1224129277
## dCA2_Fpos_TTCTGTACAGACCCGT-1dCA2_FI_GFPve_pos -1.9611558991
```

```
## dCA2_Fpos_TTGAGTGCATCCTCAC-1dCA2_FI_GFPve_pos -2.1192524510
## dCA2_Fpos_TTGCGTCTCTCTAAGG-1dCA2_FI_GFPve_pos -2.1405544358
## dCA2_Fpos_TTGGGATAGCTGACTT-1dCA2_FI_GFPve_pos -1.5666884261
## dCA2_Fpos_TTGGGATTCTCACGAA-1dCA2_FI_GFPve_pos -1.9650511819
## dCA2_Fpos_TTGGGCGGTTAGAGAT-1dCA2_FI_GFPve_pos -1.8254212218
## dCA2_Fpos_TTTAGTCAGGTACAAT-1dCA2_FI_GFPve_pos -2.3093473988
## dCA2_Fpos_TTTCCTCAGCGTGAAC-1dCA2_FI_GFPve_pos -1.6751910525
## dCA2_Fpos_TTTGATCCATGGAATA-1dCA2_FI_GFPve_pos -1.7042470055
## dCA2_Fpos_TTTGGAGAGTAGAATC-1dCA2_FI_GFPve_pos -1.2185884791
## FC_Mpos_AAGACTCTCGACATCA-1FC_MI_GFPve_pos     -1.1166155415
## FC_Mpos_AAGCGAGAGCGTCAAG-1FC_MI_GFPve_pos     -1.9436900693
## FC_Mpos_ACATGCATCCAATCTT-1FC_MI_GFPve_pos     -1.4805296737
## FC_Mpos_ACCCTCATCAAGCCAT-1FC_MI_GFPve_pos     -1.4398233968
## FC_Mpos_ACCGTTCCACGGTGCT-1FC_MI_GFPve_pos      0.1127124836
## FC_Mpos_ACCGTTCGTGCCCAGT-1FC_MI_GFPve_pos     -1.1942776519
## FC_Mpos_ACGGTTATCGATACAC-1FC_MI_GFPve_pos     -2.1906547623
## FC_Mpos_AGACTCAGTATGAGCG-1FC_MI_GFPve_pos     -1.8231618481
## FC_Mpos_AGCCAATGTTATAGAG-1FC_MI_GFPve_pos     -0.1267822909
## FC_Mpos_AGCGATTAGATGTTCC-1FC_MI_GFPve_pos     -1.4018417674
## FC_Mpos_AGCGTCGCACATATCG-1FC_MI_GFPve_pos     -0.6183176714
## FC_Mpos_AGGACTTTCGTGTTCC-1FC_MI_GFPve_pos     -1.6580144244
## FC_Mpos_AGGCATTCACCTTCCA-1FC_MI_GFPve_pos     -2.1283192712
## FC_Mpos_AGGTTACGTAACACCT-1FC_MI_GFPve_pos     -2.4568302708
## FC_Mpos_ATAGAGACAGAGGGTT-1FC_MI_GFPve_pos     -1.8460580188
## FC_Mpos_ATCGTGAAGCATGCAG-1FC_MI_GFPve_pos      0.7889064473
## FC_Mpos_ATGGAGGCACTATGTG-1FC_MI_GFPve_pos     -1.3893272954
## FC_Mpos_ATTATCCAGACCTCAT-1FC_MI_GFPve_pos     -1.6642174798
## FC_Mpos_ATTCATCAGTGAGTTA-1FC_MI_GFPve_pos     -2.2939353066
## FC_Mpos_ATTCCATTCGTTCTCG-1FC_MI_GFPve_pos      0.1482251768
## FC_Mpos_ATTCTTGCAAAGCTCT-1FC_MI_GFPve_pos     -2.4540126401
## FC_Mpos_CACATGAAGTTACGTC-1FC_MI_GFPve_pos     -1.6994687396
## FC_Mpos_CACGAATGTGTGCCTG-1FC_MI_GFPve_pos     -2.0925347882
## FC_Mpos_CACGGGTGTTCCGCTT-1FC_MI_GFPve_pos     -1.7556825953
## FC_Mpos_CACTGGGCACATAGCT-1FC_MI_GFPve_pos     -2.2550063210
## FC_Mpos_CATGCGGAGTAAGAGG-1FC_MI_GFPve_pos     -0.8654971200
## FC_Mpos_CCGATGGTCCCATGGG-1FC_MI_GFPve_pos     -1.5002234059
## FC_Mpos_CCTCTCCTCCCGGTAG-1FC_MI_GFPve_pos     -0.3800408917
## FC_Mpos_CGAGGAAGTTCCAGGC-1FC_MI_GFPve_pos     -1.2628930884
## FC_Mpos_CGATGGCAGGTCGCCT-1FC_MI_GFPve_pos     -1.2598328667
## FC_Mpos_CTCAGGGGTCGAATTC-1FC_MI_GFPve_pos     -1.3819867450
## FC_Mpos_GAACGTTTCTATTTCG-1FC_MI_GFPve_pos     -1.2484184581
## FC_Mpos_GACCCAGCATAGACTC-1FC_MI_GFPve_pos     -1.9452445584
## FC_Mpos_GAGCCTGTCCAGGACC-1FC_MI_GFPve_pos     -1.6205935555
## FC_Mpos_GAGGGTAAGAAAGTCT-1FC_MI_GFPve_pos     -1.7601707059
## FC_Mpos_GAGTCATGTCCTGAAT-1FC_MI_GFPve_pos     -0.9487444239
## FC_Mpos_GATCCCTAGCGAAACC-1FC_MI_GFPve_pos     -1.9477186280
## FC_Mpos_GCAACATGTGAAGCGT-1FC_MI_GFPve_pos     -1.9289703446
## FC_Mpos_GCCATGGCAAACGTGG-1FC_MI_GFPve_pos     -0.3322706598
## FC_Mpos_GCGATCGAGATTGATG-1FC_MI_GFPve_pos     -2.0891332703
## FC_Mpos_GCTTCACCACGACGAA-1FC_MI_GFPve_pos     -0.1800240057
## FC_Mpos_GGAACCCAGTCATGCT-1FC_MI_GFPve_pos     -2.2393887120
## FC_Mpos_GGCACGTCAAGATCCT-1FC_MI_GFPve_pos     -0.8306863385
## FC_Mpos_GGGATCCTCTGAGTCA-1FC_MI_GFPve_pos     -2.3213396149
## FC_Mpos_GGGCTCAAGATCACTC-1FC_MI_GFPve_pos     -1.3772438842
## FC_Mpos_GGTTAACAGTATGGCG-1FC_MI_GFPve_pos     -1.9298727589
## FC_Mpos_GGTTCTCTCAACACGT-1FC_MI_GFPve_pos     -0.0902157056
## FC_Mpos_GTAACCACAGAGAGGG-1FC_MI_GFPve_pos     -1.4902279454
## FC_Mpos_GTAATGCTCCGTAGGC-1FC_MI_GFPve_pos     -1.8598147708
## FC_Mpos_GTAGAAACACAAATGA-1FC_MI_GFPve_pos     -0.9198041039
## FC_Mpos_GTCATGATCTAAGAAG-1FC_MI_GFPve_pos     -1.7269533950
## FC_Mpos_GTGACGCGTTTCGGCG-1FC_MI_GFPve_pos     -1.3627508956
## FC_Mpos_GTGTTCCTCCCGAACG-1FC_MI_GFPve_pos     -0.8180084902
## FC_Mpos_TAAGTCGTCCGAGTGC-1FC_MI_GFPve_pos     -1.2273017245
## FC_Mpos_TAATTCCAGCCATTTG-1FC_MI_GFPve_pos     -1.9944443780
## FC_Mpos_TACTTGTAGTAAGAGG-1FC_MI_GFPve_pos     -1.6870302039
## FC_Mpos_TAGTGCAAGGTAAGGA-1FC_MI_GFPve_pos     -0.6148969608
## FC_Mpos_TCCATCGCAAGTATAG-1FC_MI_GFPve_pos     -1.7925980168
## FC_Mpos_TCGCTTGAGGACTTCT-1FC_MI_GFPve_pos     -2.4627392369
## FC_Mpos_TCGGGCAGTCTCCTGT-1FC_MI_GFPve_pos     -2.3534352856
## FC_Mpos_TCTATCAAGTCTAGCT-1FC_MI_GFPve_pos     -1.8329155522
```

```
## FC_Mpos_TCTCAGCAGTAAATGC-1FC_MI_GFPve_pos     -1.1418078023
## FC_Mpos_TCTTAGTGTGCGACAA-1FC_MI_GFPve_pos     -0.5651400166
## FC_Mpos_TGACAGTAGAGGGCGA-1FC_MI_GFPve_pos     -1.6803840476
## FC_Mpos_TGACTCCTCTGGCCAG-1FC_MI_GFPve_pos     -0.9753599244
## FC_Mpos_TGGGATTGTCTACGTA-1FC_MI_GFPve_pos     -1.5871057587
## FC_Mpos_TGTTGAGCACAAGCTT-1FC_MI_GFPve_pos     -0.1751493591
## FC_Mpos_TTAGGCAGTTCGAACT-1FC_MI_GFPve_pos     -2.2008583623
## FC_Mpos_TTCTAGTCATGACTGT-1FC_MI_GFPve_pos     -1.5040308314
## FC_Mpos_TTGTTGTCACTCCGGA-1FC_MI_GFPve_pos     -0.7565785723
## FC_Mpos_TTTGATCAGATGATTG-1FC_MI_GFPve_pos     -1.4008575755
## FC_Mpos_TTTGGAGCAAAGTATG-1FC_MI_GFPve_pos     -1.1477468091
## FC_Mpos_TTTGTTGGTGACGCCT-1FC_MI_GFPve_pos     -1.4321457225
## FC_Fpos_AACAACCCACCCTGTT-1FC_FI_GFPve_pos     -1.2398105937
## FC_Fpos_AACGAAACAATCACGT-1FC_FI_GFPve_pos     -1.5461694079
## FC_Fpos_AACTTCTAGCCTCAGC-1FC_FI_GFPve_pos      0.6472839636
## FC_Fpos_AATTCCTAGCAAGTCG-1FC_FI_GFPve_pos      0.1641922851
## FC_Fpos_ACACCAAGTCGCACAC-1FC_FI_GFPve_pos     -0.6131775456
## FC_Fpos_ACATGCATCCCTCTTT-1FC_FI_GFPve_pos     -1.3284235078
## FC_Fpos_AGACACTAGGGTAGCT-1FC_FI_GFPve_pos     -1.5983117896
## FC_Fpos_AGCCAATGTTTCGTAG-1FC_FI_GFPve_pos     -1.4928577023
## FC_Fpos_AGCGCTGGTAGACGTG-1FC_FI_GFPve_pos     -1.0851632434
## FC_Fpos_AGGAAATTCGGCTGAC-1FC_FI_GFPve_pos     -1.8861134129
## FC_Fpos_AGGATAATCCGATAAC-1FC_FI_GFPve_pos     -0.8287945467
## FC_Fpos_AGGTAGGCACGTACAT-1FC_FI_GFPve_pos     -1.0456190186
## FC_Fpos_AGGTCATCACGTACAT-1FC_FI_GFPve_pos      0.4647171420
## FC_Fpos_AGTAACCCAACGAGGT-1FC_FI_GFPve_pos     -0.5220905619
## FC_Fpos_AGTGTTGGTGGATGAC-1FC_FI_GFPve_pos     -1.4965421038
## FC_Fpos_ATCCACCCAGGATGAC-1FC_FI_GFPve_pos     -1.6809202510
## FC_Fpos_ATCCGTCGTTCTCTAT-1FC_FI_GFPve_pos     -0.9680509644
## FC_Fpos_ATCCGTCTCACATACG-1FC_FI_GFPve_pos     -0.0473718273
## FC_Fpos_ATCGTGAAGCACCAGA-1FC_FI_GFPve_pos     -1.3935217934
## FC_Fpos_ATCTTCAAGATGAATC-1FC_FI_GFPve_pos     -0.7894266325
## FC_Fpos_ATGTCCCGTTGGGAAC-1FC_FI_GFPve_pos     -2.3087556439
## FC_Fpos_CAAAGAAAGATTGCGG-1FC_FI_GFPve_pos     -1.8290844041
## FC_Fpos_CAAGAGGTCAACACGT-1FC_FI_GFPve_pos     -1.1978373605
## FC_Fpos_CACACAAAGTCTGTAC-1FC_FI_GFPve_pos     -1.3262584287
## FC_Fpos_CAGATACCAAGTGGTG-1FC_FI_GFPve_pos     -1.3384344655
## FC_Fpos_CAGATCATCTGCGGAC-1FC_FI_GFPve_pos     -0.9643278199
## FC_Fpos_CAGTTAGTCCTTCTTC-1FC_FI_GFPve_pos     -1.9119675237
## FC_Fpos_CATCAAGCAAGAGGCT-1FC_FI_GFPve_pos     -1.9655516225
## FC_Fpos_CATGCCTGTTGCGGCT-1FC_FI_GFPve_pos     -0.9078760224
## FC_Fpos_CATTGCCGTACTGTTG-1FC_FI_GFPve_pos     -1.2000509578
## FC_Fpos_CATTGTTTCACCTTAT-1FC_FI_GFPve_pos     -1.8876267749
## FC_Fpos_CCAATGATCTCTGAGA-1FC_FI_GFPve_pos     -1.7962516623
## FC_Fpos_CCACACTAGCGGCTCT-1FC_FI_GFPve_pos     -1.1912028867
## FC_Fpos_CCACGAGGTCCAATCA-1FC_FI_GFPve_pos     -1.1274346190
## FC_Fpos_CCCGAAGCAAGACCTT-1FC_FI_GFPve_pos      1.0172590536
## FC_Fpos_CCCTAACGTACTGACT-1FC_FI_GFPve_pos     -1.9309589940
## FC_Fpos_CCCTGATCATAGGCGA-1FC_FI_GFPve_pos     -1.0313371497
## FC_Fpos_CCGTTCACACTAGAGG-1FC_FI_GFPve_pos      0.1462909987
## FC_Fpos_CCGTTCAGTACTAGCT-1FC_FI_GFPve_pos     -0.8752415853
## FC_Fpos_CCTAACCCAGTCCCGA-1FC_FI_GFPve_pos     -2.1750519353
## FC_Fpos_CCTAACCTCATAGGCT-1FC_FI_GFPve_pos     -1.5196307975
## FC_Fpos_CCTAAGACAACGGCCT-1FC_FI_GFPve_pos     -1.7908762770
## FC_Fpos_CCTCACACAAATCCCA-1FC_FI_GFPve_pos     -2.1692786294
## FC_Fpos_CCTCCTCGTCCTCCTA-1FC_FI_GFPve_pos     -1.8040067035
## FC_Fpos_CCTCTCCCACGCGTGT-1FC_FI_GFPve_pos     -1.7288289147
## FC_Fpos_CGAATTGGTTGTGTAC-1FC_FI_GFPve_pos     -0.9507209140
## FC_Fpos_CGAGGCTTCAACGAGG-1FC_FI_GFPve_pos     -1.7665782052
## FC_Fpos_CGGGCATTCACAACCA-1FC_FI_GFPve_pos     -1.5335377532
## FC_Fpos_CGTAGTAGTCACCCTT-1FC_FI_GFPve_pos     -0.6500095206
## FC_Fpos_CGTGCTTGTTGACTAC-1FC_FI_GFPve_pos     -1.5325013476
## FC_Fpos_CGTTCTGTCACACCCT-1FC_FI_GFPve_pos     -0.8128897029
## FC_Fpos_CTCATTATCTGGAGAG-1FC_FI_GFPve_pos     -2.0962691384
## FC_Fpos_CTCCCAACAGAGAGGG-1FC_FI_GFPve_pos     -0.3441680747
## FC_Fpos_CTCCCAAGTCGGAACA-1FC_FI_GFPve_pos     -1.0823187905
## FC_Fpos_CTCGAGGGTGGAGAAA-1FC_FI_GFPve_pos     -0.8599178868
## FC_Fpos_CTCTCGATCGTGGCTG-1FC_FI_GFPve_pos     -1.3813594657
## FC_Fpos_CTGCCATAGGAGCAAA-1FC_FI_GFPve_pos     -1.5278011638
## FC_Fpos_CTGGTCTCACTGGAAG-1FC_FI_GFPve_pos     -1.5714325982
```

```
## FC_Fpos_CTGTATTAGCGCTGCT-1FC_FI_GFPve_pos     -1.7801612693
## FC_Fpos_CTGTGAACAATTAGGA-1FC_FI_GFPve_pos     -1.6305139142
## FC_Fpos_CTTCCGAAGGACTAAT-1FC_FI_GFPve_pos     -1.0326182919
## FC_Fpos_CTTCGGTTCCCTCTTT-1FC_FI_GFPve_pos     -1.1279776173
## FC_Fpos_CTTCTAACAGAGTTGG-1FC_FI_GFPve_pos     -1.1040972548
## FC_Fpos_CTTTCAATCAAGTCTG-1FC_FI_GFPve_pos     -1.7305575686
## FC_Fpos_GAACGTTCAGAAGTGC-1FC_FI_GFPve_pos     -1.7086272317
## FC_Fpos_GAAGTAATCTCCTGTG-1FC_FI_GFPve_pos     -0.8429035621
## FC_Fpos_GAGAAATCACGTTGGC-1FC_FI_GFPve_pos      1.1947232408
## FC_Fpos_GAGCCTGTCTTTCCGG-1FC_FI_GFPve_pos     -1.2739040929
## FC_Fpos_GAGGGTAAGCGATGAC-1FC_FI_GFPve_pos     -0.6931039768
## FC_Fpos_GAGTGTTGTATCCCTC-1FC_FI_GFPve_pos     -1.2556940394
## FC_Fpos_GATAGAAAGAGGTCAC-1FC_FI_GFPve_pos     -1.7353335696
## FC_Fpos_GATGACTTCCTGGTCT-1FC_FI_GFPve_pos     -0.7139933186
## FC_Fpos_GATGAGGCAAAGGGCT-1FC_FI_GFPve_pos     -2.0361924249
## FC_Fpos_GCTGGGTCATGTGACT-1FC_FI_GFPve_pos     -2.3296034413
## FC_Fpos_GCTTTCGGTAGACACG-1FC_FI_GFPve_pos     -1.6160843449
## FC_Fpos_GCTTTCGGTTGAGTCT-1FC_FI_GFPve_pos     -1.1398111659
## FC_Fpos_GGAAGTGAGTCATCCA-1FC_FI_GFPve_pos     -1.2068864184
## FC_Fpos_GGAATCTCATGGCTAT-1FC_FI_GFPve_pos     -1.7105226594
## FC_Fpos_GGCTTGGCAAACTAAG-1FC_FI_GFPve_pos     -1.6674894172
## FC_Fpos_GGCTTTCTCAACACGT-1FC_FI_GFPve_pos     -1.2147454100
## FC_Fpos_GGGACTCAGCCGAACA-1FC_FI_GFPve_pos      0.0340179873
## FC_Fpos_GGGTGTCCATGTTCGA-1FC_FI_GFPve_pos     -0.7982089597
## FC_Fpos_GGTTCTCTCCGATAGT-1FC_FI_GFPve_pos     -1.9399020749
## FC_Fpos_GTAACCATCAACTGAC-1FC_FI_GFPve_pos     -0.9008783179
## FC_Fpos_GTAGATCTCTCTGACC-1FC_FI_GFPve_pos     -1.2399483996
## FC_Fpos_GTCAGCGGTATGAGCG-1FC_FI_GFPve_pos     -1.7996035891
## FC_Fpos_GTCATCCGTCTGCAAT-1FC_FI_GFPve_pos     -1.1619343835
## FC_Fpos_GTCATTTCAGGTGGAT-1FC_FI_GFPve_pos     -1.4720852452
## FC_Fpos_GTCCCATAGTGATTCC-1FC_FI_GFPve_pos     -0.0454238134
## FC_Fpos_GTCGCGAAGTTAACGA-1FC_FI_GFPve_pos     -1.6481226760
## FC_Fpos_GTTACAGCAAAGAACT-1FC_FI_GFPve_pos     -0.7875804382
## FC_Fpos_TAACCAGCAATACGCT-1FC_FI_GFPve_pos     -0.7196508723
## FC_Fpos_TAACCAGCACTTCAAG-1FC_FI_GFPve_pos     -1.5547972995
## FC_Fpos_TAATTCCTCCAGCCTT-1FC_FI_GFPve_pos     -2.1952955800
## FC_Fpos_TACCCACCACCTATCC-1FC_FI_GFPve_pos     -1.3203777152
## FC_Fpos_TACCGAAGTTTCCCAC-1FC_FI_GFPve_pos     -0.4988666254
## FC_Fpos_TACCGGGGTCCTCCTA-1FC_FI_GFPve_pos     -0.2313467937
## FC_Fpos_TACTGCCCACGGTGTC-1FC_FI_GFPve_pos     -0.4253168779
## FC_Fpos_TACTGCCTCTGCGGGT-1FC_FI_GFPve_pos     -0.2019438821
## FC_Fpos_TACTTCATCACTAGCA-1FC_FI_GFPve_pos     -1.5012043792
## FC_Fpos_TAGCACATCGCGCCAA-1FC_FI_GFPve_pos     -2.0027096349
## FC_Fpos_TATCCTATCCGCATAA-1FC_FI_GFPve_pos     -1.7190648394
## FC_Fpos_TCAATCTGTAGCTGTT-1FC_FI_GFPve_pos     -1.0222548323
## FC_Fpos_TCAGGTACATCGGATT-1FC_FI_GFPve_pos     -1.9674217778
## FC_Fpos_TCCACCAGTAGCTGTT-1FC_FI_GFPve_pos     -1.8183785754
## FC_Fpos_TCCTGCAAGGTACTGG-1FC_FI_GFPve_pos     -0.8519138771
## FC_Fpos_TCCTTCTGTGATTCTG-1FC_FI_GFPve_pos     -1.5322332459
## FC_Fpos_TCGAACACACCTTCCA-1FC_FI_GFPve_pos     -1.0502793866
## FC_Fpos_TCGGTCTGTCTACATG-1FC_FI_GFPve_pos      0.4207326991
## FC_Fpos_TCTTAGTAGACACACG-1FC_FI_GFPve_pos     -1.7238316613
## FC_Fpos_TCTTAGTCACCACTGG-1FC_FI_GFPve_pos     -0.4032729941
## FC_Fpos_TGAATGCAGACCAACG-1FC_FI_GFPve_pos     -0.6977017003
## FC_Fpos_TGACCCTGTATCAGGG-1FC_FI_GFPve_pos     -1.7344029027
## FC_Fpos_TGATGGTTCCATACTT-1FC_FI_GFPve_pos     -1.6382920819
## FC_Fpos_TGCATGAAGACAGCGT-1FC_FI_GFPve_pos     -0.4313335496
## FC_Fpos_TGCTGAAAGGTACATA-1FC_FI_GFPve_pos     -0.7285928803
## FC_Fpos_TGGGATTCATTCCTAT-1FC_FI_GFPve_pos     -1.9425275403
## FC_Fpos_TGGTGATCATCACAGT-1FC_FI_GFPve_pos     -0.3625015693
## FC_Fpos_TGGTGATGTCTATGAC-1FC_FI_GFPve_pos     -0.7548738199
## FC_Fpos_TGTTACTGTGTAGTGG-1FC_FI_GFPve_pos     -0.5799339491
## FC_Fpos_TTACAGGTCCTCACGT-1FC_FI_GFPve_pos     -0.2897813993
## FC_Fpos_TTACCATGTCGATTCA-1FC_FI_GFPve_pos     -0.5299995619
## FC_Fpos_TTACGTTCAAGAGTTA-1FC_FI_GFPve_pos     -0.5357911664
## FC_Fpos_TTCTAACCAAGGACAC-1FC_FI_GFPve_pos     -1.5641188699
## FC_Fpos_TTGACCCGTATACGGG-1FC_FI_GFPve_pos      0.0995917676
## FC_Fpos_TTGTTGTTCTGCGGAC-1FC_FI_GFPve_pos     -1.8469305116
## FC_Fpos_TTTCCTCTCAAGATAG-1FC_FI_GFPve_pos     -1.6880414563
## dCA2_Mpos_AAAGAACCATGTTCGA-1dCA2_MI_GFPve_pos  0.4992400629
```

```
## dCA2_Mpos_AAGACAATCGTCAACA-1dCA2_MI_GFPve_pos  0.3707835299
## dCA2_Mpos_AAGCCATAGGCTCACC-1dCA2_MI_GFPve_pos  0.8567228240
## dCA2_Mpos_AAGGTAAGTCGTATGT-1dCA2_MI_GFPve_pos -0.3391651886
## dCA2_Mpos_AATTTCCCATCAACCA-1dCA2_MI_GFPve_pos  0.1741310430
## dCA2_Mpos_ACAGAAAGTCAAAGCG-1dCA2_MI_GFPve_pos  0.7740122599
## dCA2_Mpos_ACATCGAGTCCGGATC-1dCA2_MI_GFPve_pos  0.2561810536
## dCA2_Mpos_ACCACAATCAAGGAGC-1dCA2_MI_GFPve_pos  0.8537536782
## dCA2_Mpos_ACCGTTCGTTAGGCCC-1dCA2_MI_GFPve_pos  1.1799021882
## dCA2_Mpos_ACCTGTCGTCGCAACC-1dCA2_MI_GFPve_pos  0.8342083019
## dCA2_Mpos_ACTGCAAGTCGGTAAG-1dCA2_MI_GFPve_pos  1.1653278989
## dCA2_Mpos_ACTGTGAAGTACCGGA-1dCA2_MI_GFPve_pos  0.9971051139
## dCA2_Mpos_ACTTTCAAGCCGAACA-1dCA2_MI_GFPve_pos  1.0190452856
## dCA2_Mpos_ACTTTGTCAATAACCC-1dCA2_MI_GFPve_pos  0.7368559760
## dCA2_Mpos_AGAAGCGAGTTGGAAT-1dCA2_MI_GFPve_pos  0.7047820014
## dCA2_Mpos_AGACAGGTCAATCAGC-1dCA2_MI_GFPve_pos  1.1376550121
## dCA2_Mpos_AGATCCATCCGACGGT-1dCA2_MI_GFPve_pos -0.4654708582
## dCA2_Mpos_AGGCATTCATCTATCT-1dCA2_MI_GFPve_pos -0.6723678189
## dCA2_Mpos_AGTAACCGTGACTGTT-1dCA2_MI_GFPve_pos -0.1698090213
## dCA2_Mpos_AGTAGTCCAAAGGCTG-1dCA2_MI_GFPve_pos  0.6579270286
## dCA2_Mpos_ATCACGACAGACCTAT-1dCA2_MI_GFPve_pos  0.8434774799
## dCA2_Mpos_ATCGTAGGTGTCCAAT-1dCA2_MI_GFPve_pos -0.0521916973
## dCA2_Mpos_ATTACTCCAGGTTACT-1dCA2_MI_GFPve_pos  1.0195705218
## dCA2_Mpos_ATTCCTAAGGACATCG-1dCA2_MI_GFPve_pos  1.0019310040
## dCA2_Mpos_ATTCGTTTCACCTCGT-1dCA2_MI_GFPve_pos  0.2205562962
## dCA2_Mpos_ATTTCACAGCGTGCCT-1dCA2_MI_GFPve_pos -1.0503430444
## dCA2_Mpos_CACGTTCTCTAAGCCA-1dCA2_MI_GFPve_pos  0.4228631181
## dCA2_Mpos_CAGTTAGCACGCGGTT-1dCA2_MI_GFPve_pos  1.1047302407
## dCA2_Mpos_CATCCGTTCACTGATG-1dCA2_MI_GFPve_pos  1.2521834296
## dCA2_Mpos_CCAATGAGTAGGGTAC-1dCA2_MI_GFPve_pos -0.5414728003
## dCA2_Mpos_CCAATGAGTTCATCGA-1dCA2_MI_GFPve_pos  0.2694549334
## dCA2_Mpos_CCTCATGCACAGCCTG-1dCA2_MI_GFPve_pos  0.2194168118
## dCA2_Mpos_CCTCTAGTCCGTTGGG-1dCA2_MI_GFPve_pos  0.7365834040
## dCA2_Mpos_CGAAGTTAGACGGTCA-1dCA2_MI_GFPve_pos  0.8257468862
## dCA2_Mpos_CGAAGTTTCCAAAGGG-1dCA2_MI_GFPve_pos  1.0980146450
## dCA2_Mpos_CGAATTGAGATTCGCT-1dCA2_MI_GFPve_pos  0.0602603776
## dCA2_Mpos_CGACAGCGTCCGGCAT-1dCA2_MI_GFPve_pos -0.6055438596
## dCA2_Mpos_CGAGAAGAGCATACTC-1dCA2_MI_GFPve_pos  1.0327570361
## dCA2_Mpos_CGGACACTCATCGCTC-1dCA2_MI_GFPve_pos  1.1655813379
## dCA2_Mpos_CGTAGTACATGGACAG-1dCA2_MI_GFPve_pos -0.1625694114
## dCA2_Mpos_CGTCAAAAGATACATG-1dCA2_MI_GFPve_pos  0.9999145788
## dCA2_Mpos_CGTGCTTAGAATGTTG-1dCA2_MI_GFPve_pos -0.0048445913
## dCA2_Mpos_CGTTAGATCCCATACC-1dCA2_MI_GFPve_pos  0.2988184911
## dCA2_Mpos_CTACCCAAGAGTTCGG-1dCA2_MI_GFPve_pos  1.1257514877
## dCA2_Mpos_CTAGGTAAGTTACTCG-1dCA2_MI_GFPve_pos  0.9499660653
## dCA2_Mpos_CTCAGTCCACTACGGC-1dCA2_MI_GFPve_pos  0.1411122826
## dCA2_Mpos_CTCATGCTCCTCGCAT-1dCA2_MI_GFPve_pos  0.1859700305
## dCA2_Mpos_CTCATTACAGAAGCTG-1dCA2_MI_GFPve_pos  1.1867520732
## dCA2_Mpos_CTCATTATCCTGATAG-1dCA2_MI_GFPve_pos -0.0259552526
## dCA2_Mpos_GAAGAATAGCCGTTAT-1dCA2_MI_GFPve_pos  1.0373356861
## dCA2_Mpos_GAAGGGTAGACGGTCA-1dCA2_MI_GFPve_pos -0.0755053687
## dCA2_Mpos_GAATAGAAGAGTCAGC-1dCA2_MI_GFPve_pos  1.0810773296
## dCA2_Mpos_GAATCGTCAAACTCGT-1dCA2_MI_GFPve_pos  0.9058853907
## dCA2_Mpos_GACCAATGTCAGATTC-1dCA2_MI_GFPve_pos -0.0978918600
## dCA2_Mpos_GACCGTGCATCGAAGG-1dCA2_MI_GFPve_pos  0.2766244930
## dCA2_Mpos_GAGAGGTAGAAGGTAG-1dCA2_MI_GFPve_pos -0.8204603868
## dCA2_Mpos_GAGATGGAGCCTATTG-1dCA2_MI_GFPve_pos  1.1210439128
## dCA2_Mpos_GATCAGTAGGCTCACC-1dCA2_MI_GFPve_pos -0.1954278725
## dCA2_Mpos_GATCAGTAGGGTGAAA-1dCA2_MI_GFPve_pos  0.2365333212
## dCA2_Mpos_GATTCGACATCTGCGG-1dCA2_MI_GFPve_pos  1.0136258525
## dCA2_Mpos_GCAGCCACAACGTTAC-1dCA2_MI_GFPve_pos -0.7752892452
## dCA2_Mpos_GCAGCTGAGGATTACA-1dCA2_MI_GFPve_pos  0.5004267317
## dCA2_Mpos_GCATCTCTCCGCCTAT-1dCA2_MI_GFPve_pos  0.2214852271
## dCA2_Mpos_GCGTTTCTCGTCTCAC-1dCA2_MI_GFPve_pos  0.0230811534
## dCA2_Mpos_GCTCAAAAGTCCTACA-1dCA2_MI_GFPve_pos  1.1245222015
## dCA2_Mpos_GCTGCAGGTCACCACG-1dCA2_MI_GFPve_pos  0.5669552607
## dCA2_Mpos_GGAACCCCATCCGGCA-1dCA2_MI_GFPve_pos  0.9269533676
## dCA2_Mpos_GGAGAACAGCACCAGA-1dCA2_MI_GFPve_pos  0.8004324955
## dCA2_Mpos_GGAGCAACAGAGGGTT-1dCA2_MI_GFPve_pos -0.7264063435
## dCA2_Mpos_GGAGGTAGTCCTACAA-1dCA2_MI_GFPve_pos  0.1282214445
## dCA2_Mpos_GGGCGTTAGCTTCTAG-1dCA2_MI_GFPve_pos -0.6765367466
```

```
## dCA2_Mpos_GTAGATCTCAGTGTTG-1dCA2_MI_GFPve_pos -0.0404955524
## dCA2_Mpos_GTAGCTATCCATGATG-1dCA2_MI_GFPve_pos  0.4219520611
## dCA2_Mpos_GTCCTCATCTTAGGAC-1dCA2_MI_GFPve_pos  0.7494260592
## dCA2_Mpos_GTCTCACAGCACGATG-1dCA2_MI_GFPve_pos -0.8191526013
## dCA2_Mpos_GTCTGTCTCCGTACGG-1dCA2_MI_GFPve_pos -0.6592064577
## dCA2_Mpos_GTGGAAGAGCGAGTCA-1dCA2_MI_GFPve_pos  0.9810448808
## dCA2_Mpos_GTTCCGTAGGTAGGCT-1dCA2_MI_GFPve_pos -0.1701978522
## dCA2_Mpos_TAAGCGTAGTGGAAGA-1dCA2_MI_GFPve_pos -0.5551783520
## dCA2_Mpos_TACCCGTCAGCGTAGA-1dCA2_MI_GFPve_pos -0.1254196542
## dCA2_Mpos_TACTTCAAGTTTGAGA-1dCA2_MI_GFPve_pos  0.9966648740
## dCA2_Mpos_TAGGTTGCAGTGGGTA-1dCA2_MI_GFPve_pos  1.0875835938
## dCA2_Mpos_TATCGCCAGTTAACGA-1dCA2_MI_GFPve_pos -0.1810797828
## dCA2_Mpos_TCACAAGCAACTAGAA-1dCA2_MI_GFPve_pos -0.0150913673
## dCA2_Mpos_TCACATTGTTGATCGT-1dCA2_MI_GFPve_pos -0.7186993438
## dCA2_Mpos_TCCACGTAGTGGGAAA-1dCA2_MI_GFPve_pos  0.3089964015
## dCA2_Mpos_TCCAGAATCTACTCAT-1dCA2_MI_GFPve_pos -0.7045399624
## dCA2_Mpos_TCTAACTTCCCTGTTG-1dCA2_MI_GFPve_pos -0.7650848704
## dCA2_Mpos_TGATGGTGTGACAACG-1dCA2_MI_GFPve_pos -0.1564613360
## dCA2_Mpos_TGCGGCATCTACTATC-1dCA2_MI_GFPve_pos -0.7343399602
## dCA2_Mpos_TGGTAGTCAGTTGGTT-1dCA2_MI_GFPve_pos -0.0838333892
## dCA2_Mpos_TGTTCATTCTAAGGAA-1dCA2_MI_GFPve_pos -0.7799186188
## dCA2_Mpos_TGTTCCGCATTCACCC-1dCA2_MI_GFPve_pos  1.0168178600
## dCA2_Mpos_TGTTGAGCAGTCGAGA-1dCA2_MI_GFPve_pos  0.4018089992
## dCA2_Mpos_TTACGCCTCGGCTGAC-1dCA2_MI_GFPve_pos -0.7274636703
## dCA2_Mpos_TTAGTCTCACTGTCCT-1dCA2_MI_GFPve_pos  0.1966180948
## dCA2_Mpos_TTCACGCTCGACCTAA-1dCA2_MI_GFPve_pos  0.0668693108
## dCA2_Mpos_TTGCGTCCAAGGAGTC-1dCA2_MI_GFPve_pos -0.3124402540
## dCA2_Mpos_TTGTTCAGTGCGAACA-1dCA2_MI_GFPve_pos  0.7303262276
## dCA2_Mpos_TTGTTGTAGAAACACT-1dCA2_MI_GFPve_pos  0.1083370938
## dCA2_Mpos_TTTCACAAGTTCTCTT-1dCA2_MI_GFPve_pos  0.9738432092
## dCA2_Fpos_AGGATCTTCACCTGGG-1dCA2_FI_GFPve_pos  1.2980915231
## dCA2_Fpos_AGGTCATTCCACAGGC-1dCA2_FI_GFPve_pos  0.3713799638
## dCA2_Fpos_AGTAGTCAGCTTACGT-1dCA2_FI_GFPve_pos -0.1588219183
## dCA2_Fpos_ATCATTCAGGTCATCT-1dCA2_FI_GFPve_pos  1.1750202102
## dCA2_Fpos_ATCTTCACAGAAGTGC-1dCA2_FI_GFPve_pos  0.9675407333
## dCA2_Fpos_ATGGGAGAGCATGGGT-1dCA2_FI_GFPve_pos  0.2254505900
## dCA2_Fpos_CCCTCAAGTTGTGGAG-1dCA2_FI_GFPve_pos  0.0257470456
## dCA2_Fpos_GCAACCGCAATAGTAG-1dCA2_FI_GFPve_pos  1.0438777131
## dCA2_Fpos_GCCCAGATCTTGTTAC-1dCA2_FI_GFPve_pos  0.2634191883
## dCA2_Fpos_GTAGAGGCAACAACAA-1dCA2_FI_GFPve_pos  0.2471747023
## dCA2_Fpos_TACGTCCTCCAATGCA-1dCA2_FI_GFPve_pos -0.7598447877
## dCA2_Fpos_TAGGAGGTCTGACCCT-1dCA2_FI_GFPve_pos  0.0035053340
## dCA2_Fpos_TCATGCCGTCCCGGTA-1dCA2_FI_GFPve_pos  0.3158058000
## dCA2_Fpos_TCGGGCAAGCGTTCCG-1dCA2_FI_GFPve_pos -0.5044767338
## dCA2_Fpos_TCTTTGAGTGTTTCTT-1dCA2_FI_GFPve_pos -0.7727921086
## dCA2_Fpos_TGAACGTGTCGACGCT-1dCA2_FI_GFPve_pos -0.1392633098
## dCA2_Fpos_TGCTGAAGTACCCAGC-1dCA2_FI_GFPve_pos -0.1110894906
## dCA2_Fpos_TGCTGAAGTGAATATG-1dCA2_FI_GFPve_pos -0.1439000892
## dCA2_Fpos_TGCTTCGCATAGAAAC-1dCA2_FI_GFPve_pos -0.6461597162
## dCA2_Fpos_TGTCCCACAGTCTACA-1dCA2_FI_GFPve_pos  0.2017807958
## dCA2_Fpos_TTGCATTGTGCAGTGA-1dCA2_FI_GFPve_pos  0.8132793826
## FC_Mpos_AAACCCACAGACTGCC-1FC_MI_GFPve_pos     -0.8375284749
## FC_Mpos_AAAGAACAGTGCAGCA-1FC_MI_GFPve_pos      0.5606837196
## FC_Mpos_AAAGTCCGTGCGTTTA-1FC_MI_GFPve_pos     -0.5858348089
## FC_Mpos_AAATGGAAGCTCCGAC-1FC_MI_GFPve_pos      0.5722452921
## FC_Mpos_AAATGGACAGTTCACA-1FC_MI_GFPve_pos      1.1000543756
## FC_Mpos_AACCACAGTTTACCAG-1FC_MI_GFPve_pos      0.6175432128
## FC_Mpos_AACGAAAAGGCAGGGA-1FC_MI_GFPve_pos     -0.0653237927
## FC_Mpos_AAGACAACAGTAGATA-1FC_MI_GFPve_pos      0.6780491990
## FC_Mpos_AAGACTCAGGTTCCGC-1FC_MI_GFPve_pos      0.3509832395
## FC_Mpos_AAGATAGTCGCCTTTG-1FC_MI_GFPve_pos      0.3423086924
## FC_Mpos_AAGCGAGCATCGGTTA-1FC_MI_GFPve_pos     -0.3391974049
## FC_Mpos_AAGCGAGTCGAAGCCC-1FC_MI_GFPve_pos     -0.2457024234
## FC_Mpos_AAGCGTTCAAAGCACG-1FC_MI_GFPve_pos      0.8226511997
## FC_Mpos_AATGAAGAGTGAGCCA-1FC_MI_GFPve_pos      1.4943013114
## FC_Mpos_AATGGCTAGATACAGT-1FC_MI_GFPve_pos     -0.0441651570
## FC_Mpos_ACAAAGAAGTTAGTGA-1FC_MI_GFPve_pos      0.5119715852
## FC_Mpos_ACAGCCGGTCGAGCAA-1FC_MI_GFPve_pos      0.5503522677
## FC_Mpos_ACATCGAAGGGTACGT-1FC_MI_GFPve_pos      1.1803288383
## FC_Mpos_ACCAACACAGCGGTCT-1FC_MI_GFPve_pos      0.8211776537
```

```
## FC_Mpos_ACCATTTGTCATAAAG-1FC_MI_GFPve_pos     -0.2223164814
## FC_Mpos_ACCATTTGTGACTCGC-1FC_MI_GFPve_pos      1.0930321736
## FC_Mpos_ACCATTTGTGTCATTG-1FC_MI_GFPve_pos      0.9747344178
## FC_Mpos_ACGCACGCACAACGTT-1FC_MI_GFPve_pos      0.4360956473
## FC_Mpos_ACGTAACGTTCTTAGG-1FC_MI_GFPve_pos      0.8253599924
## FC_Mpos_ACTATCTGTGGGCTCT-1FC_MI_GFPve_pos      0.5077490431
## FC_Mpos_ACTTCCGAGATACTGA-1FC_MI_GFPve_pos      0.7083833737
## FC_Mpos_ACTTCGCAGGCTGAAC-1FC_MI_GFPve_pos     -0.7397510486
## FC_Mpos_ACTTCGCAGGTGCTAG-1FC_MI_GFPve_pos     -0.8087428885
## FC_Mpos_ACTTTGTAGACCTGGA-1FC_MI_GFPve_pos      0.6443038506
## FC_Mpos_AGAAATGCAACATCGT-1FC_MI_GFPve_pos      0.2168274221
## FC_Mpos_AGACACTCACTACCCT-1FC_MI_GFPve_pos      0.0839888898
## FC_Mpos_AGACCCGGTGGGCTCT-1FC_MI_GFPve_pos      0.5223771555
## FC_Mpos_AGAGAGCGTAGTACGG-1FC_MI_GFPve_pos     -0.2992863434
## FC_Mpos_AGAGAGCTCATGAAAG-1FC_MI_GFPve_pos      0.0831405499
## FC_Mpos_AGATAGAAGGTTAGTA-1FC_MI_GFPve_pos      0.2493682874
## FC_Mpos_AGCCAATAGTTCACTG-1FC_MI_GFPve_pos      0.0106458810
## FC_Mpos_AGCGTATCATACACCA-1FC_MI_GFPve_pos     -1.1619364100
## FC_Mpos_AGCTACACACGGTGCT-1FC_MI_GFPve_pos      0.0441075784
## FC_Mpos_AGGAATAAGGCAGGGA-1FC_MI_GFPve_pos     -1.7536573487
## FC_Mpos_AGGTCTACAAGGCAAC-1FC_MI_GFPve_pos     -0.7542559582
## FC_Mpos_AGGTCTACACATGACT-1FC_MI_GFPve_pos     -0.0348590242
## FC_Mpos_AGGTTGTCACAAATCC-1FC_MI_GFPve_pos      0.0200523583
## FC_Mpos_AGTAACCCAGAACTTC-1FC_MI_GFPve_pos      0.3185252917
## FC_Mpos_AGTTAGCGTCTGTGCG-1FC_MI_GFPve_pos      0.3797650558
## FC_Mpos_AGTTCCCGTAAGCGGT-1FC_MI_GFPve_pos      1.2679693622
## FC_Mpos_AGTTCCCGTACGAGCA-1FC_MI_GFPve_pos      0.6590992731
## FC_Mpos_AGTTCCCGTACGCTTA-1FC_MI_GFPve_pos      0.5942659599
## FC_Mpos_ATATCCTCAACGACAG-1FC_MI_GFPve_pos      0.3323032183
## FC_Mpos_ATCACGATCCTCTTTC-1FC_MI_GFPve_pos     -0.1458333957
## FC_Mpos_ATCATTCGTCAGGTGA-1FC_MI_GFPve_pos      0.7847167176
## FC_Mpos_ATCATTCTCGCACTCT-1FC_MI_GFPve_pos     -0.9769647198
## FC_Mpos_ATGAAAGAGATCGACG-1FC_MI_GFPve_pos     -0.5741078335
## FC_Mpos_ATGAGTCGTCAAGTTC-1FC_MI_GFPve_pos      0.4368477685
## FC_Mpos_ATGTCCCAGGAACATT-1FC_MI_GFPve_pos      0.1512363290
## FC_Mpos_ATTACTCGTCAACACT-1FC_MI_GFPve_pos      0.0416649466
## FC_Mpos_ATTCTTGTCACTTTGT-1FC_MI_GFPve_pos      0.8044452590
## FC_Mpos_ATTGGGTCAGGCTTGC-1FC_MI_GFPve_pos      0.2142720518
## FC_Mpos_CAAAGAATCGGAATGG-1FC_MI_GFPve_pos     -1.0200493413
## FC_Mpos_CAACCTCGTACCTTCC-1FC_MI_GFPve_pos      0.8516233486
## FC_Mpos_CAACGGCCATCAGCAT-1FC_MI_GFPve_pos      1.2282268924
## FC_Mpos_CAAGCTAGTGACTATC-1FC_MI_GFPve_pos     -0.0315937477
## FC_Mpos_CAAGGGAGTTGCTCGG-1FC_MI_GFPve_pos      0.5867512447
## FC_Mpos_CAATACGAGGCGTTGA-1FC_MI_GFPve_pos     -1.0798688012
## FC_Mpos_CAATCGAGTTCTCCCA-1FC_MI_GFPve_pos     -0.4327932673
## FC_Mpos_CACAACATCAACCTTT-1FC_MI_GFPve_pos     -0.6668155270
## FC_Mpos_CACAGATCATTGTCGA-1FC_MI_GFPve_pos      1.6981650514
## FC_Mpos_CACGTGGAGAAGTATC-1FC_MI_GFPve_pos      0.2614590747
## FC_Mpos_CACGTTCAGGGAGGGT-1FC_MI_GFPve_pos      0.1632780654
## FC_Mpos_CAGATCAGTCCAAAGG-1FC_MI_GFPve_pos     -0.0170082527
## FC_Mpos_CAGCGTGGTGATACCT-1FC_MI_GFPve_pos      0.8830841703
## FC_Mpos_CAGTGCGGTCTTACAG-1FC_MI_GFPve_pos      1.0097171587
## FC_Mpos_CAGTTCCGTCTTTCAT-1FC_MI_GFPve_pos      0.4272444767
## FC_Mpos_CATGCCTAGTTGGCTT-1FC_MI_GFPve_pos      0.0534075608
## FC_Mpos_CATGCGGTCCCTCTTT-1FC_MI_GFPve_pos      0.6354833168
## FC_Mpos_CATTCATAGATTGATG-1FC_MI_GFPve_pos     -0.9373947459
## FC_Mpos_CATTTCACATCAACCA-1FC_MI_GFPve_pos      0.4010317547
## FC_Mpos_CCAAGCGCACGCGTGT-1FC_MI_GFPve_pos     -0.2942657250
## FC_Mpos_CCAATGATCCTGTAAG-1FC_MI_GFPve_pos     -0.0601788240
## FC_Mpos_CCACACTAGAAGCGGG-1FC_MI_GFPve_pos      1.2895965499
## FC_Mpos_CCCATTGAGTCATCCA-1FC_MI_GFPve_pos      0.8812234325
## FC_Mpos_CCCATTGCATTGAAAG-1FC_MI_GFPve_pos      0.3058354480
## FC_Mpos_CCCTAACGTCTCAGGC-1FC_MI_GFPve_pos      0.6596805376
## FC_Mpos_CCGAACGCAGGTGAGT-1FC_MI_GFPve_pos      1.0804905218
## FC_Mpos_CCGGACACACCTTCGT-1FC_MI_GFPve_pos     -0.1452773737
## FC_Mpos_CCGGACATCACTGGTA-1FC_MI_GFPve_pos     -0.9831098395
## FC_Mpos_CCGGTGACACAGTATC-1FC_MI_GFPve_pos     -0.6530521589
## FC_Mpos_CCGTAGGAGGTCGTGA-1FC_MI_GFPve_pos      0.4475077016
## FC_Mpos_CCTGCATAGCGTTACT-1FC_MI_GFPve_pos      0.9779728574
## FC_Mpos_CGAATTGTCCTGGGTG-1FC_MI_GFPve_pos      0.4007270557
```

```
## FC_Mpos_CGAGGAAAGTGCTAGG-1FC_MI_GFPve_pos     -0.7424519139
## FC_Mpos_CGAGTTATCTGAGCAT-1FC_MI_GFPve_pos      0.6535597367
## FC_Mpos_CGCATAATCTTAGTTC-1FC_MI_GFPve_pos      1.1658921165
## FC_Mpos_CGGAACCAGGTTGGTG-1FC_MI_GFPve_pos      0.2582930994
## FC_Mpos_CGGGACTGTCCTCAGG-1FC_MI_GFPve_pos      0.9680907649
## FC_Mpos_CGGGTGTCAGAACTCT-1FC_MI_GFPve_pos      0.2561918420
## FC_Mpos_CGTCAAAGTTATAGCC-1FC_MI_GFPve_pos     -0.3752521354
## FC_Mpos_CGTCCATCAGCACCCA-1FC_MI_GFPve_pos     -0.3464987057
## FC_Mpos_CGTGATATCCACCTGT-1FC_MI_GFPve_pos      1.4102091712
## FC_Mpos_CGTTAGAGTTGTGGAG-1FC_MI_GFPve_pos      0.8889045042
## FC_Mpos_CTAACTTAGGGCCCTT-1FC_MI_GFPve_pos      0.3684587700
## FC_Mpos_CTACAGAGTTTCGTGA-1FC_MI_GFPve_pos      0.9477336926
## FC_Mpos_CTACCCACAGGCCCTA-1FC_MI_GFPve_pos     -0.5682801681
## FC_Mpos_CTCAACCTCGAACGCC-1FC_MI_GFPve_pos      0.4576394779
## FC_Mpos_CTCCATGCACAGCCAC-1FC_MI_GFPve_pos      0.2123228160
## FC_Mpos_CTGAGCGAGCTGAGTG-1FC_MI_GFPve_pos      0.1948740405
## FC_Mpos_CTGCGAGTCAGAATAG-1FC_MI_GFPve_pos      0.3312815947
## FC_Mpos_CTTAGGAGTTGAAGTA-1FC_MI_GFPve_pos     -0.0348747450
## FC_Mpos_CTTCTCTAGCTGAAAT-1FC_MI_GFPve_pos      0.3211104346
## FC_Mpos_GAACACTAGAGCAACC-1FC_MI_GFPve_pos      0.6822314781
## FC_Mpos_GAAGAATAGCGGGTAT-1FC_MI_GFPve_pos      0.6516799254
## FC_Mpos_GAAGCCCCAATACGCT-1FC_MI_GFPve_pos      0.3031949175
## FC_Mpos_GACTTCCAGGACAGCT-1FC_MI_GFPve_pos      0.4339867455
## FC_Mpos_GAGACCCTCTCTTAAC-1FC_MI_GFPve_pos      0.6887742204
## FC_Mpos_GAGCCTGTCATCACTT-1FC_MI_GFPve_pos      0.3242295009
## FC_Mpos_GAGGCCTCATACCACA-1FC_MI_GFPve_pos      0.5675457043
## FC_Mpos_GAGGGTAAGCGATGCA-1FC_MI_GFPve_pos     -0.0270105975
## FC_Mpos_GAGTCATAGAAGTATC-1FC_MI_GFPve_pos      0.1892635298
## FC_Mpos_GAGTGAGAGAAGGTAG-1FC_MI_GFPve_pos     -0.4641408997
## FC_Mpos_GAGTGAGGTAGCTGAG-1FC_MI_GFPve_pos      1.0986433548
## FC_Mpos_GAGTGTTCACGTACTA-1FC_MI_GFPve_pos     -0.7327715832
## FC_Mpos_GAGTTACCACAGCATT-1FC_MI_GFPve_pos      0.1666637060
## FC_Mpos_GAGTTTGAGAATTCAG-1FC_MI_GFPve_pos      0.8655003232
## FC_Mpos_GATGAGGCAAGGCTTT-1FC_MI_GFPve_pos      1.5854282302
## FC_Mpos_GATGTTGAGCTACAAA-1FC_MI_GFPve_pos     -0.1575646179
## FC_Mpos_GCAACCGTCGTAGGAG-1FC_MI_GFPve_pos     -0.5460050183
## FC_Mpos_GCACGGTGTAGCGAGT-1FC_MI_GFPve_pos      0.2978990448
## FC_Mpos_GCAGCCAGTAGATCCT-1FC_MI_GFPve_pos     -0.6705716925
## FC_Mpos_GCAGCTGTCAAGCCCG-1FC_MI_GFPve_pos      1.1001412195
## FC_Mpos_GCAGGCTAGGTTGAGC-1FC_MI_GFPve_pos      0.3546634835
## FC_Mpos_GCCATTCTCTTCCCAG-1FC_MI_GFPve_pos     -0.0444741475
## FC_Mpos_GCCTGTTAGAGCATTA-1FC_MI_GFPve_pos      0.3599572999
## FC_Mpos_GCCTGTTAGGGTACGT-1FC_MI_GFPve_pos      0.7943479342
## FC_Mpos_GCGGAAACAGTCGTTA-1FC_MI_GFPve_pos     -0.4370828348
## FC_Mpos_GCGGATCTCATTTGGG-1FC_MI_GFPve_pos     -2.1818564015
## FC_Mpos_GCGTGCATCCCTCAAC-1FC_MI_GFPve_pos      1.3207194728
## FC_Mpos_GCTACCTCACAAAGTA-1FC_MI_GFPve_pos     -0.1148329603
## FC_Mpos_GCTTGGGTCGACCTAA-1FC_MI_GFPve_pos      0.1628227529
## FC_Mpos_GGAATCTAGGCGTTGA-1FC_MI_GFPve_pos      1.6223738116
## FC_Mpos_GGAGAACCAATACGCT-1FC_MI_GFPve_pos      1.1362318916
## FC_Mpos_GGAGGATAGATCCCGC-1FC_MI_GFPve_pos      0.7817121667
## FC_Mpos_GGGATCCAGTGGATAT-1FC_MI_GFPve_pos      0.5592664999
## FC_Mpos_GGGATCCTCCCTATTA-1FC_MI_GFPve_pos      1.5064042730
## FC_Mpos_GGGCCATCAACGGCCT-1FC_MI_GFPve_pos     -0.6955114680
## FC_Mpos_GGGTGAACAAGTTGGG-1FC_MI_GFPve_pos      0.5836542470
## FC_Mpos_GGTAATCCACAAAGCG-1FC_MI_GFPve_pos      0.2610934449
## FC_Mpos_GGTGAAGAGTAACAGT-1FC_MI_GFPve_pos     -0.1401603746
## FC_Mpos_GGTGAAGAGTAACCGG-1FC_MI_GFPve_pos      1.5317464990
## FC_Mpos_GGTGGCTGTGTCTCCT-1FC_MI_GFPve_pos      0.5387223286
## FC_Mpos_GGTGTCGCAAGTGGGT-1FC_MI_GFPve_pos     -1.4609077054
## FC_Mpos_GGTGTCGGTGTTAACC-1FC_MI_GFPve_pos     -1.3008639890
## FC_Mpos_GGTGTCGTCCCTCGAT-1FC_MI_GFPve_pos      1.0709079427
## FC_Mpos_GGTTGTAAGGCGATAC-1FC_MI_GFPve_pos      1.0354056877
## FC_Mpos_GTAGATCCACTGTCCT-1FC_MI_GFPve_pos      0.5378716213
## FC_Mpos_GTAGATCTCTCTCGAC-1FC_MI_GFPve_pos      0.0092701313
## FC_Mpos_GTAGCTAAGGGTCTTT-1FC_MI_GFPve_pos      1.3701175374
## FC_Mpos_GTCAAACGTACCGTCG-1FC_MI_GFPve_pos      1.5061058921
## FC_Mpos_GTCACTCCACATGTTG-1FC_MI_GFPve_pos     -0.6384666639
## FC_Mpos_GTCACTCCATGGTGGA-1FC_MI_GFPve_pos      1.2070701045
## FC_Mpos_GTCATGAGTTGGATCT-1FC_MI_GFPve_pos      1.8360407275
```

```
## FC_Mpos_GTCGTTCCACGACAAG-1FC_MI_GFPve_pos      1.5537854356
## FC_Mpos_GTGACGCAGCAACCAG-1FC_MI_GFPve_pos      0.6238272113
## FC_Mpos_GTGAGTTCATCAACCA-1FC_MI_GFPve_pos      0.8736914915
## FC_Mpos_GTGGAAGAGCACCGTC-1FC_MI_GFPve_pos     -0.7048044878
## FC_Mpos_GTGGAGACAGTTGTCA-1FC_MI_GFPve_pos      0.7798221034
## FC_Mpos_GTTACCCAGCAAATGT-1FC_MI_GFPve_pos      1.4224080962
## FC_Mpos_GTTAGACGTTCTCTAT-1FC_MI_GFPve_pos      0.1564284188
## FC_Mpos_GTTAGTGAGCATTGAA-1FC_MI_GFPve_pos      0.0969162134
## FC_Mpos_GTTGAACTCCGGACTG-1FC_MI_GFPve_pos      0.7166368884
## FC_Mpos_GTTGCTCTCCCAGGAC-1FC_MI_GFPve_pos      0.1831381765
## FC_Mpos_TAACACGCATGACGAG-1FC_MI_GFPve_pos     -0.0227247405
## FC_Mpos_TAACGACGTTACCCTC-1FC_MI_GFPve_pos      0.2212580887
## FC_Mpos_TAATTCCCAGGTCTCG-1FC_MI_GFPve_pos      1.0233910603
## FC_Mpos_TAATTCCGTCACCGAC-1FC_MI_GFPve_pos      0.4560598952
## FC_Mpos_TACAGGTCAAGCTGCC-1FC_MI_GFPve_pos      0.1378175539
## FC_Mpos_TAGACTGGTTTCCCAC-1FC_MI_GFPve_pos      0.0711224524
## FC_Mpos_TATTGCTGTAGCGTCC-1FC_MI_GFPve_pos      0.6027318758
## FC_Mpos_TCAAGACTCTTGGAAC-1FC_MI_GFPve_pos      0.7784895820
## FC_Mpos_TCAAGTGAGACGGATC-1FC_MI_GFPve_pos      0.6295375151
## FC_Mpos_TCAATTCTCAGAGCAG-1FC_MI_GFPve_pos      0.7067550940
## FC_Mpos_TCACACCAGTCTGGTT-1FC_MI_GFPve_pos      0.1443030459
## FC_Mpos_TCACGGGTCGGTCACG-1FC_MI_GFPve_pos      0.1774322239
## FC_Mpos_TCATCCGCAGGACATG-1FC_MI_GFPve_pos      0.5020351333
## FC_Mpos_TCATGCCGTTGCCAAT-1FC_MI_GFPve_pos      1.0140794439
## FC_Mpos_TCCACCATCCACAGGC-1FC_MI_GFPve_pos     -0.4777707535
## FC_Mpos_TCCACGTGTAATGCTC-1FC_MI_GFPve_pos      0.6400417012
## FC_Mpos_TCCATGCGTACCTTCC-1FC_MI_GFPve_pos      0.1599444335
## FC_Mpos_TCCGATCAGTAAACAC-1FC_MI_GFPve_pos      0.4242712182
## FC_Mpos_TCGACCTAGCCATTGT-1FC_MI_GFPve_pos      0.2848355097
## FC_Mpos_TCGGGCAAGCACTGGA-1FC_MI_GFPve_pos      0.3252432865
## FC_Mpos_TCTAACTCACTGAGTT-1FC_MI_GFPve_pos      0.5817077977
## FC_Mpos_TCTATACAGCTAATGA-1FC_MI_GFPve_pos      1.1204839868
## FC_Mpos_TCTCACGTCTCCCAAC-1FC_MI_GFPve_pos      0.8809299988
## FC_Mpos_TCTCTGGAGATGATTG-1FC_MI_GFPve_pos      0.9931011719
## FC_Mpos_TCTGGCTCAGCTAACT-1FC_MI_GFPve_pos      0.6565948052
## FC_Mpos_TCTGGCTTCCATAAGC-1FC_MI_GFPve_pos     -0.1893693286
## FC_Mpos_TCTTCCTTCTGGTGCG-1FC_MI_GFPve_pos      0.2240078223
## FC_Mpos_TGACGCGGTCGGTGAA-1FC_MI_GFPve_pos      0.3462736083
## FC_Mpos_TGACGCGTCTAGAGCT-1FC_MI_GFPve_pos     -0.7867644506
## FC_Mpos_TGACTCCAGACCAGCA-1FC_MI_GFPve_pos     -1.2882994490
## FC_Mpos_TGAGGTTAGCGTGAAC-1FC_MI_GFPve_pos      0.5959083361
## FC_Mpos_TGATCTTAGGTATAGT-1FC_MI_GFPve_pos      0.6534907145
## FC_Mpos_TGATGGTGTTGCCTAA-1FC_MI_GFPve_pos     -0.4549661952
## FC_Mpos_TGCGGGTGTGGACAGT-1FC_MI_GFPve_pos     -0.3957003432
## FC_Mpos_TGGATCATCGGTTCAA-1FC_MI_GFPve_pos     -1.5806487876
## FC_Mpos_TGGCGTGTCAGACTGT-1FC_MI_GFPve_pos      0.9866865796
## FC_Mpos_TGGGAGAGTAAGGAGA-1FC_MI_GFPve_pos      0.2607434464
## FC_Mpos_TGGGAGATCTTCCGTG-1FC_MI_GFPve_pos      0.1636382004
## FC_Mpos_TGTAACGAGCAATAGT-1FC_MI_GFPve_pos      0.4196410698
## FC_Mpos_TGTGTGATCGCCTCTA-1FC_MI_GFPve_pos      0.9602376146
## FC_Mpos_TGTTCTAGTCATACCA-1FC_MI_GFPve_pos     -0.2555410641
## FC_Mpos_TGTTGAGAGCTGAAAT-1FC_MI_GFPve_pos      0.2790690732
## FC_Mpos_TGTTGGAGTTAAGTCC-1FC_MI_GFPve_pos      0.8275720400
## FC_Mpos_TGTTTGTAGCAAATGT-1FC_MI_GFPve_pos      0.7040778321
## FC_Mpos_TGTTTGTCAGCTTTGA-1FC_MI_GFPve_pos      0.7756516856
## FC_Mpos_TTACCGCAGTCATACC-1FC_MI_GFPve_pos     -1.1019866544
## FC_Mpos_TTACCGCTCTATCGCC-1FC_MI_GFPve_pos      0.3045683873
## FC_Mpos_TTACGCCAGGTTCATC-1FC_MI_GFPve_pos      0.7893068713
## FC_Mpos_TTATTGCAGGTACATA-1FC_MI_GFPve_pos      0.4544728321
## FC_Mpos_TTCACGCCAACAGCTT-1FC_MI_GFPve_pos      0.2181938586
## FC_Mpos_TTCCTAAAGTTACGTC-1FC_MI_GFPve_pos      1.4906430167
## FC_Mpos_TTCGCTGGTGAGTCAG-1FC_MI_GFPve_pos      0.9873113555
## FC_Mpos_TTCTGTATCCCATTTA-1FC_MI_GFPve_pos     -0.8264650780
## FC_Mpos_TTGGGCGAGAGCAACC-1FC_MI_GFPve_pos     -0.2757293957
## FC_Mpos_TTGGTTTTCACTTCTA-1FC_MI_GFPve_pos      0.0368585733
## FC_Mpos_TTTACCATCTTGGGCG-1FC_MI_GFPve_pos      1.1562565488
## FC_Mpos_TTTAGTCCAGGAAGTC-1FC_MI_GFPve_pos      0.4997865183
## FC_Mpos_TTTGATCTCGTTGTAG-1FC_MI_GFPve_pos      0.6572889013
## FC_Mpos_TTTGGTTGTACGTGTT-1FC_MI_GFPve_pos      0.2836277557
## FC_Fpos_AAACGCTTCTAGATCG-1FC_FI_GFPve_pos      0.0280352903
```

```
## FC_Fpos_AAAGGATTCGGCCTTT-1FC_FI_GFPve_pos      0.3532380385
## FC_Fpos_AAAGTGATCCGGTTCT-1FC_FI_GFPve_pos      0.2838032467
## FC_Fpos_AACAACCAGGTCCAGA-1FC_FI_GFPve_pos     -0.7789385992
## FC_Fpos_AACCAACAGACCATGG-1FC_FI_GFPve_pos     -0.5515243488
## FC_Fpos_AACCAACTCAGGTAAA-1FC_FI_GFPve_pos      0.8091525358
## FC_Fpos_AACCATGCACTTGAGT-1FC_FI_GFPve_pos     -0.2922865229
## FC_Fpos_AACGTCAGTGGTCCGT-1FC_FI_GFPve_pos      0.2841965181
## FC_Fpos_AACTTCTAGGGCAGTT-1FC_FI_GFPve_pos      0.1005219286
## FC_Fpos_AACTTCTCAAATTGGA-1FC_FI_GFPve_pos      0.5426612419
## FC_Fpos_AAGAACAAGTTAACAG-1FC_FI_GFPve_pos      1.3569136781
## FC_Fpos_AAGAACAGTTAGGAGC-1FC_FI_GFPve_pos      0.4684592110
## FC_Fpos_AAGACAAAGCATAGGC-1FC_FI_GFPve_pos      0.1430610416
## FC_Fpos_AAGACAAAGGCGATAC-1FC_FI_GFPve_pos      0.1658498195
## FC_Fpos_AAGATAGGTGTTCCTC-1FC_FI_GFPve_pos      1.1784167213
## FC_Fpos_AAGATAGTCTATCGTT-1FC_FI_GFPve_pos     -0.0634870755
## FC_Fpos_AAGCATCGTGCCGTTG-1FC_FI_GFPve_pos     -0.5112755376
## FC_Fpos_AAGCGTTTCATCCTGC-1FC_FI_GFPve_pos      0.3138951582
## FC_Fpos_AAGGTAACATATGGCT-1FC_FI_GFPve_pos     -0.8692950683
## FC_Fpos_AAGTACCGTATGTCCA-1FC_FI_GFPve_pos      0.1481349205
## FC_Fpos_AATCGTGAGTATAGAC-1FC_FI_GFPve_pos      0.1396511642
## FC_Fpos_AATGGCTGTTGGACTT-1FC_FI_GFPve_pos      0.9014543218
## FC_Fpos_AATTCCTCAACGTTAC-1FC_FI_GFPve_pos     -0.3879704254
## FC_Fpos_ACAAAGAGTAGGTACG-1FC_FI_GFPve_pos      0.2363213224
## FC_Fpos_ACAGGGAAGCCTCATA-1FC_FI_GFPve_pos      0.1644352784
## FC_Fpos_ACCAAACTCTTCACGC-1FC_FI_GFPve_pos      0.1625629251
## FC_Fpos_ACCCAAACAAGACTGG-1FC_FI_GFPve_pos      1.1431612891
## FC_Fpos_ACCTGAAAGGTCCTGC-1FC_FI_GFPve_pos      0.2510007096
## FC_Fpos_ACCTGTCAGCTGAAAT-1FC_FI_GFPve_pos     -0.2504869479
## FC_Fpos_ACGCACGCACGGTGCT-1FC_FI_GFPve_pos      0.3812376482
## FC_Fpos_ACGCACGCATATGCGT-1FC_FI_GFPve_pos     -0.3383871692
## FC_Fpos_ACGGAAGTCTTCGGTC-1FC_FI_GFPve_pos     -0.5133685547
## FC_Fpos_ACGTACACATACACCA-1FC_FI_GFPve_pos      0.7615604920
## FC_Fpos_ACGTCCTCAGCTGGTC-1FC_FI_GFPve_pos      0.6750833911
## FC_Fpos_ACTGCAACACTGTGAT-1FC_FI_GFPve_pos      0.1443694418
## FC_Fpos_AGACAAACAATGCAAA-1FC_FI_GFPve_pos     -0.3710123378
## FC_Fpos_AGATCGTTCGCCTTGT-1FC_FI_GFPve_pos      1.5142331046
## FC_Fpos_AGCATCAGTACGCTAT-1FC_FI_GFPve_pos     -0.4454013782
## FC_Fpos_AGCCAATCACAGTCGC-1FC_FI_GFPve_pos      0.0563931351
## FC_Fpos_AGCGATTCACTAGGCC-1FC_FI_GFPve_pos     -0.5582764226
## FC_Fpos_AGCGCCAAGCTGCCTG-1FC_FI_GFPve_pos      1.1253007573
## FC_Fpos_AGCGCCAAGGGCTGAT-1FC_FI_GFPve_pos      0.2475676459
## FC_Fpos_AGCGCTGCAGTCAGTT-1FC_FI_GFPve_pos      0.3068629575
## FC_Fpos_AGCTACACAGCAGTAG-1FC_FI_GFPve_pos      1.3097797555
## FC_Fpos_AGCTTCCGTTCAAACC-1FC_FI_GFPve_pos      0.8380377812
## FC_Fpos_AGGAAATCAGCTATAC-1FC_FI_GFPve_pos      0.2381093425
## FC_Fpos_AGGAATATCTGCGGCA-1FC_FI_GFPve_pos     -0.2567718404
## FC_Fpos_AGGCCACCACCCTTGT-1FC_FI_GFPve_pos      0.3092271907
## FC_Fpos_AGGGCCTAGAGAGGTA-1FC_FI_GFPve_pos     -0.7360447126
## FC_Fpos_AGGGTGAGTAGCTCGC-1FC_FI_GFPve_pos     -0.4380940872
## FC_Fpos_AGGGTTTCAACCTATG-1FC_FI_GFPve_pos      0.8342412633
## FC_Fpos_AGGTAGGTCTTCTGGC-1FC_FI_GFPve_pos      0.1017226820
## FC_Fpos_AGGTTGTCAACTTGCA-1FC_FI_GFPve_pos      0.3828433675
## FC_Fpos_AGTACCACAGGTGACA-1FC_FI_GFPve_pos     -0.6122137385
## FC_Fpos_AGTAGCTCAGCACGAA-1FC_FI_GFPve_pos     -1.2218760329
## FC_Fpos_AGTCACACACACGTGC-1FC_FI_GFPve_pos      0.6274533195
## FC_Fpos_AGTCATGAGGTGCGAT-1FC_FI_GFPve_pos      0.8098004979
## FC_Fpos_AGTCTCCCATGTAACC-1FC_FI_GFPve_pos     -0.8463135439
## FC_Fpos_AGTGTTGAGCAAATGT-1FC_FI_GFPve_pos      1.1274450940
## FC_Fpos_ATACCTTCATCCGTGG-1FC_FI_GFPve_pos      1.4557800216
## FC_Fpos_ATAGACCAGGTTGGTG-1FC_FI_GFPve_pos      0.9151329321
## FC_Fpos_ATAGAGAGTCACGTGC-1FC_FI_GFPve_pos      0.3754423184
## FC_Fpos_ATCACGACACCCTGTT-1FC_FI_GFPve_pos     -0.3212814110
## FC_Fpos_ATCCACCCAATTCGTG-1FC_FI_GFPve_pos     -0.0547559934
## FC_Fpos_ATCCCTGAGCATCAGG-1FC_FI_GFPve_pos     -0.6098098832
## FC_Fpos_ATCGATGAGGGTACGT-1FC_FI_GFPve_pos      0.7091410083
## FC_Fpos_ATCGATGGTTGCTCAA-1FC_FI_GFPve_pos     -0.6897132474
## FC_Fpos_ATCGCCTAGCCGTTGC-1FC_FI_GFPve_pos      0.5070483607
## FC_Fpos_ATCGTGATCTCCTGCA-1FC_FI_GFPve_pos      0.5778803450
## FC_Fpos_ATCTCTAAGCGCGTTC-1FC_FI_GFPve_pos      0.6489858550
## FC_Fpos_ATCTCTAGTTAGTCGT-1FC_FI_GFPve_pos      0.4114202780
```

```
## FC_Fpos_ATGGGAGTCACTTCTA-1FC_FI_GFPve_pos      0.4831520540
## FC_Fpos_ATGGGAGTCTTACGGA-1FC_FI_GFPve_pos      1.3817506952
## FC_Fpos_ATTATCCTCTGGGCAC-1FC_FI_GFPve_pos     -0.3865100222
## FC_Fpos_ATTGTTCTCAACCTTT-1FC_FI_GFPve_pos     -0.2768853384
## FC_Fpos_CAAAGAAAGAATAACC-1FC_FI_GFPve_pos      0.1843511609
## FC_Fpos_CAAAGAAGTACTGCGC-1FC_FI_GFPve_pos      1.0734246296
## FC_Fpos_CAAAGAAGTGCCGTTG-1FC_FI_GFPve_pos     -0.6568277555
## FC_Fpos_CAAAGAATCCGGTAAT-1FC_FI_GFPve_pos      0.5006624681
## FC_Fpos_CAAGAGGAGGTTCTTG-1FC_FI_GFPve_pos      0.6252114696
## FC_Fpos_CAATGACTCAGTCACA-1FC_FI_GFPve_pos     -0.8552165705
## FC_Fpos_CACACAACAGCATTGT-1FC_FI_GFPve_pos      0.6146418375
## FC_Fpos_CACAGGCAGAAGATCT-1FC_FI_GFPve_pos      1.0466439051
## FC_Fpos_CAGAGCCGTCCTTAAG-1FC_FI_GFPve_pos      0.0073347387
## FC_Fpos_CAGATCATCTCACGAA-1FC_FI_GFPve_pos     -0.1118548888
## FC_Fpos_CAGCGTGCACATTCTT-1FC_FI_GFPve_pos      0.1348624786
## FC_Fpos_CAGGTATGTCGTTGGC-1FC_FI_GFPve_pos      0.2759599013
## FC_Fpos_CATCAAGTCGGACCAC-1FC_FI_GFPve_pos      0.6854854745
## FC_Fpos_CATCCCACAATACAGA-1FC_FI_GFPve_pos     -0.0337091761
## FC_Fpos_CATCGCTAGCATTGTC-1FC_FI_GFPve_pos     -0.1366095322
## FC_Fpos_CATCGGGTCTACGCAA-1FC_FI_GFPve_pos     -1.3529260236
## FC_Fpos_CATGCAACACTAGTAC-1FC_FI_GFPve_pos      0.1019635022
## FC_Fpos_CATGCCTCATAATGCC-1FC_FI_GFPve_pos     -0.7364491897
## FC_Fpos_CATGCGGAGATTAGCA-1FC_FI_GFPve_pos      0.5012770814
## FC_Fpos_CATGCTCTCTGAATGC-1FC_FI_GFPve_pos      0.4978425426
## FC_Fpos_CATTCATGTGGTCCCA-1FC_FI_GFPve_pos     -0.2080136913
## FC_Fpos_CATTCATTCACCACAA-1FC_FI_GFPve_pos      0.8187155647
## FC_Fpos_CATTGTTTCAAGCCCG-1FC_FI_GFPve_pos     -0.6660134273
## FC_Fpos_CATTGTTTCTATTGTC-1FC_FI_GFPve_pos     -1.6814549046
## FC_Fpos_CCACAAAGTCTCCCTA-1FC_FI_GFPve_pos      0.5707769913
## FC_Fpos_CCCTAACTCAAGCCGC-1FC_FI_GFPve_pos      0.2518683744
## FC_Fpos_CCCTCTCGTGCCCGTA-1FC_FI_GFPve_pos      0.5885289830
## FC_Fpos_CCGCAAGAGGTAGGCT-1FC_FI_GFPve_pos      0.2357714248
## FC_Fpos_CCGGACACAACCGCCA-1FC_FI_GFPve_pos      0.8769913239
## FC_Fpos_CCGTGAGCAGGACTTT-1FC_FI_GFPve_pos     -0.3563779968
## FC_Fpos_CCTATCGCACAACGTT-1FC_FI_GFPve_pos     -0.9203298169
## FC_Fpos_CCTCAACCACTCCGAG-1FC_FI_GFPve_pos      0.1711479795
## FC_Fpos_CCTCAGTCACAATTCG-1FC_FI_GFPve_pos     -0.3606545704
## FC_Fpos_CCTGCATCACACGGTC-1FC_FI_GFPve_pos     -0.0151812586
## FC_Fpos_CCTTTGGTCAGACAAA-1FC_FI_GFPve_pos      0.8537963552
## FC_Fpos_CGAGGCTGTGAGGAAA-1FC_FI_GFPve_pos      0.3559578580
## FC_Fpos_CGATCGGCAAACAGGC-1FC_FI_GFPve_pos     -1.1481591540
## FC_Fpos_CGATGGCAGAATAGTC-1FC_FI_GFPve_pos      0.5296583695
## FC_Fpos_CGCGTGAAGGCTAACG-1FC_FI_GFPve_pos      0.7735603971
## FC_Fpos_CGGAATTAGGCGCTTC-1FC_FI_GFPve_pos      1.0088325662
## FC_Fpos_CGGACACCACAGCTGC-1FC_FI_GFPve_pos      0.8656721038
## FC_Fpos_CGGCAGTAGACATAAC-1FC_FI_GFPve_pos      0.3183672500
## FC_Fpos_CGGGTGTGTTCGGGTC-1FC_FI_GFPve_pos      0.2205745054
## FC_Fpos_CGTCCATCAGAGGAAA-1FC_FI_GFPve_pos     -0.3437965887
## FC_Fpos_CGTGATAGTTGCGTAT-1FC_FI_GFPve_pos      0.3632717652
## FC_Fpos_CGTTAGACACCAGTTA-1FC_FI_GFPve_pos     -0.0455929118
## FC_Fpos_CTAACCCAGTGGATTA-1FC_FI_GFPve_pos      0.1742707786
## FC_Fpos_CTAACTTAGGAGGTTC-1FC_FI_GFPve_pos      0.1409129721
## FC_Fpos_CTAACTTCAACGCCCA-1FC_FI_GFPve_pos     -0.5147178250
## FC_Fpos_CTAAGTGAGCTCACTA-1FC_FI_GFPve_pos      0.8899627847
## FC_Fpos_CTACCTGGTCTTACTT-1FC_FI_GFPve_pos      0.3802326841
## FC_Fpos_CTAGACACATGTGCCG-1FC_FI_GFPve_pos     -1.5927751141
## FC_Fpos_CTCAACCGTCACGTGC-1FC_FI_GFPve_pos      0.0847660040
## FC_Fpos_CTCCCTCCACAAGCCC-1FC_FI_GFPve_pos      0.6696504873
## FC_Fpos_CTCCGATTCTGGAGAG-1FC_FI_GFPve_pos     -0.0356841015
## FC_Fpos_CTCTCAGAGGGTGGGA-1FC_FI_GFPve_pos      0.2242321742
## FC_Fpos_CTGAATGCATAGTCAC-1FC_FI_GFPve_pos     -0.1939049738
## FC_Fpos_CTGAATGTCGGTAGAG-1FC_FI_GFPve_pos      0.3526816589
## FC_Fpos_CTGAGCGCAGACCTAT-1FC_FI_GFPve_pos     -0.4082096892
## FC_Fpos_CTGAGCGGTAGCCAGA-1FC_FI_GFPve_pos     -0.7005441266
## FC_Fpos_CTGATCCGTGACTGTT-1FC_FI_GFPve_pos     -0.8955599147
## FC_Fpos_CTGCAGGAGAGGCTGT-1FC_FI_GFPve_pos     -1.0994856435
## FC_Fpos_CTGCAGGTCTTAGCAG-1FC_FI_GFPve_pos      0.3383177740
## FC_Fpos_CTGCATCAGGTTACAA-1FC_FI_GFPve_pos     -0.0741604793
## FC_Fpos_CTGCATCGTACGACAG-1FC_FI_GFPve_pos      0.1361803931
## FC_Fpos_CTGCCTAAGCGTTGTT-1FC_FI_GFPve_pos      0.2905341906
```

```
## FC_Fpos_CTGTGAACAAGTGCTT-1FC_FI_GFPve_pos     -0.6655074435
## FC_Fpos_CTTACCGTCCCGTAAA-1FC_FI_GFPve_pos      0.7225176019
## FC_Fpos_CTTAGGATCATCACTT-1FC_FI_GFPve_pos      0.9558454675
## FC_Fpos_CTTCGGTAGCGATGGT-1FC_FI_GFPve_pos      0.0090839562
## FC_Fpos_CTTTCAACAGTTCTAG-1FC_FI_GFPve_pos      0.0494493020
## FC_Fpos_GAAACCTTCCTATTGT-1FC_FI_GFPve_pos     -0.5164087492
## FC_Fpos_GAACGTTGTATTGACC-1FC_FI_GFPve_pos      0.4561648888
## FC_Fpos_GAAGTAAGTTAAACCC-1FC_FI_GFPve_pos      0.0558597517
## FC_Fpos_GACACGCGTACCCGAC-1FC_FI_GFPve_pos      0.2540951741
## FC_Fpos_GACCCAGGTTCGGGTC-1FC_FI_GFPve_pos      0.2832420540
## FC_Fpos_GACCCTTGTCAGTCCG-1FC_FI_GFPve_pos      0.8556575102
## FC_Fpos_GACGTTATCTGCTTAT-1FC_FI_GFPve_pos      0.7472699804
## FC_Fpos_GACTTCCCACTCCGGA-1FC_FI_GFPve_pos     -0.6500102359
## FC_Fpos_GACTTCCTCTACCACC-1FC_FI_GFPve_pos      0.5281486434
## FC_Fpos_GAGAAATCAAGCGCTC-1FC_FI_GFPve_pos      0.4572514815
## FC_Fpos_GAGAAATGTACGAGTG-1FC_FI_GFPve_pos     -0.2245325225
## FC_Fpos_GAGACTTGTCGGTGAA-1FC_FI_GFPve_pos      0.7026015443
## FC_Fpos_GAGAGGTGTGAGCGAT-1FC_FI_GFPve_pos     -0.6791267472
## FC_Fpos_GAGGGTAGTAAGGAGA-1FC_FI_GFPve_pos     -0.3315548974
## FC_Fpos_GAGTCATTCTCGACGG-1FC_FI_GFPve_pos      0.5247166080
## FC_Fpos_GAGTTGTTCATGCCCT-1FC_FI_GFPve_pos      0.2690922392
## FC_Fpos_GATCACACACCCTAGG-1FC_FI_GFPve_pos     -0.0906757044
## FC_Fpos_GATCAGTGTTCCTACC-1FC_FI_GFPve_pos      0.3581143541
## FC_Fpos_GATCCCTAGGTATCTC-1FC_FI_GFPve_pos      0.2230703903
## FC_Fpos_GATGGAGCACGCAAAG-1FC_FI_GFPve_pos      0.5705660803
## FC_Fpos_GCAACCGGTCGTGCCA-1FC_FI_GFPve_pos      0.6968696636
## FC_Fpos_GCACTAACAGATACTC-1FC_FI_GFPve_pos      0.4288625342
## FC_Fpos_GCCAGGTAGAAGATCT-1FC_FI_GFPve_pos      1.3745800180
## FC_Fpos_GCCATGGTCTACACTT-1FC_FI_GFPve_pos     -0.3388483005
## FC_Fpos_GCCCGAAAGCACTAGG-1FC_FI_GFPve_pos      0.6205190939
## FC_Fpos_GCCGATGAGGAACGCT-1FC_FI_GFPve_pos      0.2574986291
## FC_Fpos_GCCGTGATCATTACGG-1FC_FI_GFPve_pos      0.3408444745
## FC_Fpos_GCTACAAGTCGTTATG-1FC_FI_GFPve_pos     -0.7469338852
## FC_Fpos_GCTGAATAGTCTTCGA-1FC_FI_GFPve_pos      1.0974517984
## FC_Fpos_GCTGCAGAGCCTTTCC-1FC_FI_GFPve_pos      0.1103428558
## FC_Fpos_GCTGCAGTCTCTCGCA-1FC_FI_GFPve_pos      0.2866024298
## FC_Fpos_GCTTCACGTACGTGTT-1FC_FI_GFPve_pos     -0.1405050385
## FC_Fpos_GGACGTCAGAACCGCA-1FC_FI_GFPve_pos      0.8791560454
## FC_Fpos_GGAGATGCATCCAACA-1FC_FI_GFPve_pos      0.2814815563
## FC_Fpos_GGAGCAACAAGTGTCT-1FC_FI_GFPve_pos     -0.5486994463
## FC_Fpos_GGGAAGTGTGAGATCG-1FC_FI_GFPve_pos      0.2916894776
## FC_Fpos_GGGACCTGTGTGTGTT-1FC_FI_GFPve_pos      0.2341173661
## FC_Fpos_GGGAGATGTTGTTGAC-1FC_FI_GFPve_pos      1.0438486857
## FC_Fpos_GGGATCCTCGCAGTCG-1FC_FI_GFPve_pos     -0.2121814566
## FC_Fpos_GGGATGAGTGTCTCCT-1FC_FI_GFPve_pos      0.6802402062
## FC_Fpos_GGGCTACAGGACAACC-1FC_FI_GFPve_pos      0.7263962549
## FC_Fpos_GGGTAGACAAAGGCAC-1FC_FI_GFPve_pos     -0.5746018368
## FC_Fpos_GGGTCTGGTACGAGCA-1FC_FI_GFPve_pos     -0.0825591462
## FC_Fpos_GGTAATCAGTTGCTGT-1FC_FI_GFPve_pos      0.1392779310
## FC_Fpos_GGTAGAGTCGAGAGCA-1FC_FI_GFPve_pos      0.5940353793
## FC_Fpos_GGTCTGGAGAAGTCCG-1FC_FI_GFPve_pos      0.2890532834
## FC_Fpos_GGTGATTGTAAGTTGA-1FC_FI_GFPve_pos      0.6231662554
## FC_Fpos_GGTGTCGAGCACCCAC-1FC_FI_GFPve_pos      0.2676181567
## FC_Fpos_GGTTGTATCGTAGCTA-1FC_FI_GFPve_pos      0.6886048240
## FC_Fpos_GTAACCATCATGAGGG-1FC_FI_GFPve_pos      0.0160242987
## FC_Fpos_GTAAGTCAGGTTCATC-1FC_FI_GFPve_pos      0.1543832791
## FC_Fpos_GTAATCGCAGCACGAA-1FC_FI_GFPve_pos      0.4551811737
## FC_Fpos_GTACAACCACACCGAC-1FC_FI_GFPve_pos      1.0755349321
## FC_Fpos_GTACAACTCAGTGGGA-1FC_FI_GFPve_pos      0.2549113703
## FC_Fpos_GTACAGTTCTTCGTGC-1FC_FI_GFPve_pos     -0.7117106992
## FC_Fpos_GTAGAAATCGTTTACT-1FC_FI_GFPve_pos      1.2500284833
## FC_Fpos_GTAGGTTCATCTATCT-1FC_FI_GFPve_pos      0.0380556581
## FC_Fpos_GTATTGGGTGCTGATT-1FC_FI_GFPve_pos     -1.1871019679
## FC_Fpos_GTATTTCCAGTATGAA-1FC_FI_GFPve_pos      0.3179374558
## FC_Fpos_GTCAAGTCAACTTCTT-1FC_FI_GFPve_pos      0.6192067784
## FC_Fpos_GTCCACTCATCTCGTC-1FC_FI_GFPve_pos     -0.3172790485
## FC_Fpos_GTCCACTTCATTGAGC-1FC_FI_GFPve_pos      0.2378288281
## FC_Fpos_GTCCACTTCCCTTCCC-1FC_FI_GFPve_pos      0.0411941034
## FC_Fpos_GTCCCATCAATGAGCG-1FC_FI_GFPve_pos      0.0646655855
## FC_Fpos_GTCCTCAGTCATACCA-1FC_FI_GFPve_pos      0.0960302071
```

```
## FC_Fpos_GTCGCGACATCCTTGC-1FC_FI_GFPve_pos      1.3553174657
## FC_Fpos_GTCTGTCTCTTGTGCC-1FC_FI_GFPve_pos     -0.0537321943
## FC_Fpos_GTGAGTTAGTGATTCC-1FC_FI_GFPve_pos      0.2223189024
## FC_Fpos_GTGAGTTGTAATCAGA-1FC_FI_GFPve_pos     -0.4017180877
## FC_Fpos_GTGCTTCAGTTCGGTT-1FC_FI_GFPve_pos     -0.6021461564
## FC_Fpos_GTGTGATTCTCAGTCC-1FC_FI_GFPve_pos      0.3555198235
## FC_Fpos_GTTCGCTAGACCAAGC-1FC_FI_GFPve_pos      0.0830426586
## FC_Fpos_GTTCGCTCACACAGCC-1FC_FI_GFPve_pos      0.1233267986
## FC_Fpos_GTTCTATAGTGGGAAA-1FC_FI_GFPve_pos     -1.5347470122
## FC_Fpos_GTTGCTCAGCATGCAG-1FC_FI_GFPve_pos      0.4181907994
## FC_Fpos_TAATCTCAGCAGGTCA-1FC_FI_GFPve_pos      0.1244484854
## FC_Fpos_TACAACGGTCGCGGTT-1FC_FI_GFPve_pos      1.8701180143
## FC_Fpos_TACAGGTCAGATACCT-1FC_FI_GFPve_pos      1.2102329654
## FC_Fpos_TACAGGTGTTGCAAGG-1FC_FI_GFPve_pos      0.3118728769
## FC_Fpos_TACCTGCAGGGCAGAG-1FC_FI_GFPve_pos      0.1224126888
## FC_Fpos_TACGGTAGTAACTGCT-1FC_FI_GFPve_pos      1.8891171140
## FC_Fpos_TATACCTCACAATTCG-1FC_FI_GFPve_pos     -0.4078307229
## FC_Fpos_TATCAGGGTAATTGGA-1FC_FI_GFPve_pos      0.0839263347
## FC_Fpos_TATGTTCCAATAACCC-1FC_FI_GFPve_pos     -0.1342712420
## FC_Fpos_TATGTTCCAATGTCAC-1FC_FI_GFPve_pos      0.1234278192
## FC_Fpos_TATTGCTAGACGCAGT-1FC_FI_GFPve_pos      0.4252875072
## FC_Fpos_TATTGCTCAAACTCTG-1FC_FI_GFPve_pos      1.0072051806
## FC_Fpos_TCAAGACTCTTCCACG-1FC_FI_GFPve_pos     -0.1340544748
## FC_Fpos_TCAATCTCAATCGAAA-1FC_FI_GFPve_pos      0.1105205773
## FC_Fpos_TCACATTGTACTCGAT-1FC_FI_GFPve_pos      0.4777242762
## FC_Fpos_TCACGGGGTTATGTGC-1FC_FI_GFPve_pos     -0.0241191166
## FC_Fpos_TCACTCGGTCGAGTTT-1FC_FI_GFPve_pos      0.5807893497
## FC_Fpos_TCAGGGCTCTATTGTC-1FC_FI_GFPve_pos      0.5746862811
## FC_Fpos_TCATGAGGTTTACTGG-1FC_FI_GFPve_pos      1.1036807222
## FC_Fpos_TCATTGTGTCAGTCTA-1FC_FI_GFPve_pos     -1.0675641375
## FC_Fpos_TCCGTGTCAAATACAG-1FC_FI_GFPve_pos     -1.2906296330
## FC_Fpos_TCCTAATCATGAATCC-1FC_FI_GFPve_pos      0.6290422005
## FC_Fpos_TCGAACAGTGAGATCG-1FC_FI_GFPve_pos      0.0124711913
## FC_Fpos_TCGAAGTGTCGCAGTC-1FC_FI_GFPve_pos     -0.1979084688
## FC_Fpos_TCGACCTAGAAACTGT-1FC_FI_GFPve_pos      1.4973014516
## FC_Fpos_TCGCACTAGTCACAGG-1FC_FI_GFPve_pos     -0.2884097176
## FC_Fpos_TCGCTCAGTGGTCTCG-1FC_FI_GFPve_pos     -0.5599236565
## FC_Fpos_TCGGGCACAGCAGTGA-1FC_FI_GFPve_pos     -1.4993939477
## FC_Fpos_TCGGTCTCATACACCA-1FC_FI_GFPve_pos      0.2370379192
## FC_Fpos_TCGTAGATCTCGTCAC-1FC_FI_GFPve_pos     -1.1168003159
## FC_Fpos_TCTACATGTACGTTCA-1FC_FI_GFPve_pos      0.3338045103
## FC_Fpos_TCTACCGCATGGGCAA-1FC_FI_GFPve_pos     -0.8892825919
## FC_Fpos_TCTATCACAGCACGAA-1FC_FI_GFPve_pos     -0.6801385957
## FC_Fpos_TCTGTCGAGACATAGT-1FC_FI_GFPve_pos      0.2388643396
## FC_Fpos_TCTTAGTCAAAGACGC-1FC_FI_GFPve_pos      0.3138226045
## FC_Fpos_TGAGACTGTGCATGTT-1FC_FI_GFPve_pos      0.6060931606
## FC_Fpos_TGAGGTTGTCATTCCC-1FC_FI_GFPve_pos      0.6056151909
## FC_Fpos_TGAGTCACAATACGCT-1FC_FI_GFPve_pos     -0.2005083042
## FC_Fpos_TGCGGCATCGGCTGTG-1FC_FI_GFPve_pos     -0.7421104985
## FC_Fpos_TGCTCGTCAAGAAATC-1FC_FI_GFPve_pos      0.8191085977
## FC_Fpos_TGCTTCGAGTTGTACC-1FC_FI_GFPve_pos      0.6634843868
## FC_Fpos_TGGCGTGAGGCACAAC-1FC_FI_GFPve_pos      1.1603981179
## FC_Fpos_TGGTACATCCATGAGT-1FC_FI_GFPve_pos      0.1000497262
## FC_Fpos_TGTAACGTCTGAGAGG-1FC_FI_GFPve_pos      0.6091539783
## FC_Fpos_TGTAGACGTCCAGTTA-1FC_FI_GFPve_pos     -0.1374517011
## FC_Fpos_TGTCAGAAGAGCCATG-1FC_FI_GFPve_pos      1.1875586433
## FC_Fpos_TGTCCTGTCCATTGGA-1FC_FI_GFPve_pos      1.1649436874
## FC_Fpos_TGTGTGACAAGCCTGC-1FC_FI_GFPve_pos     -0.8138302046
## FC_Fpos_TGTTGAGAGGACAAGA-1FC_FI_GFPve_pos     -0.2497516053
## FC_Fpos_TGTTTGTTCTGGGATT-1FC_FI_GFPve_pos     -0.2338937002
## FC_Fpos_TTACTGTCAGAATCGG-1FC_FI_GFPve_pos      0.6641330046
## FC_Fpos_TTCAATCGTATTTCCT-1FC_FI_GFPve_pos      1.1744918746
## FC_Fpos_TTCACGCCATAGATGA-1FC_FI_GFPve_pos      0.2503384275
## FC_Fpos_TTCAGGACAGGACTAG-1FC_FI_GFPve_pos      0.2931837511
## FC_Fpos_TTCATTGTCTGTCGTC-1FC_FI_GFPve_pos      0.5556846780
## FC_Fpos_TTCCAATCATGGAACG-1FC_FI_GFPve_pos      0.8377287907
## FC_Fpos_TTCCTCTAGCACTCGC-1FC_FI_GFPve_pos      1.0102521104
## FC_Fpos_TTCGCTGAGCTGGAGT-1FC_FI_GFPve_pos     -0.8399928289
## FC_Fpos_TTCTGTACAGAAGCGT-1FC_FI_GFPve_pos      0.7228220028
## FC_Fpos_TTGCTGCGTGAAAGTT-1FC_FI_GFPve_pos      1.3157435579
```

```
## FC_Fpos_TTTATGCTCTGGCTGG-1FC_FI_GFPve_pos     -0.5990563112
## FC_Fpos_TTTCATGGTGATATAG-1FC_FI_GFPve_pos     -0.8843387442
## FC_Fpos_TTTCCTCTCGAGCCTG-1FC_FI_GFPve_pos     -1.0760463553
## FC_Fpos_TTTGATCCATGCCATA-1FC_FI_GFPve_pos      0.2154591379
## dCA2_Mpos_AAAGAACTCCCGGTAG-1dCA2_MI_GFPve_pos -1.6524047929
## dCA2_Mpos_AACAACCAGGTCCGAA-1dCA2_MI_GFPve_pos -0.7472382265
## dCA2_Mpos_AACCACATCACTGCTC-1dCA2_MI_GFPve_pos -0.8784763890
## dCA2_Mpos_AACCATGCAAGCGCTC-1dCA2_MI_GFPve_pos -2.8716473656
## dCA2_Mpos_AAGACAATCAAATGCC-1dCA2_MI_GFPve_pos -0.8766922432
## dCA2_Mpos_ACATCCCTCACCTTGC-1dCA2_MI_GFPve_pos -0.6791349130
## dCA2_Mpos_ACATCCCTCCGTGACG-1dCA2_MI_GFPve_pos -0.1983079689
## dCA2_Mpos_ACCAAACCACGCTGCA-1dCA2_MI_GFPve_pos -0.7364924031
## dCA2_Mpos_ACCAACACAACCGATT-1dCA2_MI_GFPve_pos -1.6318961459
## dCA2_Mpos_AGACACTAGTCTAGCT-1dCA2_MI_GFPve_pos -1.0528770762
## dCA2_Mpos_AGACCATTCAAGCCGC-1dCA2_MI_GFPve_pos -1.4822646457
## dCA2_Mpos_AGACCATTCTACTGCC-1dCA2_MI_GFPve_pos -1.1465287286
## dCA2_Mpos_AGATCCATCAGTGTCA-1dCA2_MI_GFPve_pos -1.1639773923
## dCA2_Mpos_AGTCAACCAGCTTCGG-1dCA2_MI_GFPve_pos -1.8449690419
## dCA2_Mpos_AGTGATCCAGATGCGA-1dCA2_MI_GFPve_pos -0.9289918023
## dCA2_Mpos_ATCATTCAGCTGACTT-1dCA2_MI_GFPve_pos -1.4870886880
## dCA2_Mpos_ATCGCCTAGCCTCTCT-1dCA2_MI_GFPve_pos -0.9973176795
## dCA2_Mpos_ATGACCACATCCTTGC-1dCA2_MI_GFPve_pos -1.4524644690
## dCA2_Mpos_CACGTTCCAGACGCTC-1dCA2_MI_GFPve_pos -1.5080598670
## dCA2_Mpos_CCTCCAAAGCCATCCG-1dCA2_MI_GFPve_pos -1.0883995372
## dCA2_Mpos_CTAGACAGTGGTCTAT-1dCA2_MI_GFPve_pos -0.9237396794
## dCA2_Mpos_CTCACTGTCGGCTTCT-1dCA2_MI_GFPve_pos -1.4527831155
## dCA2_Mpos_CTCCACACAAGTTCCA-1dCA2_MI_GFPve_pos -1.1072305518
## dCA2_Mpos_GAACGTTAGATGAATC-1dCA2_MI_GFPve_pos -1.4520063477
## dCA2_Mpos_GAACTGTTCTACTCAT-1dCA2_MI_GFPve_pos -1.3363723832
## dCA2_Mpos_GAAGCGACACCCATAA-1dCA2_MI_GFPve_pos -1.1404612141
## dCA2_Mpos_GACCAATGTCGACTTA-1dCA2_MI_GFPve_pos -0.9786340075
## dCA2_Mpos_GACGTTAAGGAAACGA-1dCA2_MI_GFPve_pos -2.1844148713
## dCA2_Mpos_GACTCAACAACTGGTT-1dCA2_MI_GFPve_pos -0.6985222774
## dCA2_Mpos_GATGCTAAGCTCACTA-1dCA2_MI_GFPve_pos -1.0609439688
## dCA2_Mpos_GCACGTGCACAGTACT-1dCA2_MI_GFPve_pos -2.7766718942
## dCA2_Mpos_GCAGCCACATGACTAC-1dCA2_MI_GFPve_pos -0.5868665653
## dCA2_Mpos_GCAGCTGCACAACCGC-1dCA2_MI_GFPve_pos -0.5042772966
## dCA2_Mpos_GCCAGCATCCTCTCTT-1dCA2_MI_GFPve_pos -0.9100070077
## dCA2_Mpos_GGTTAACTCCATTCAT-1dCA2_MI_GFPve_pos -1.2056179124
## dCA2_Mpos_GTTTACTCACTGCGTG-1dCA2_MI_GFPve_pos -1.0166801291
## dCA2_Mpos_TACCCACAGTCACTGT-1dCA2_MI_GFPve_pos -0.7418563443
## dCA2_Mpos_TACGCTCTCCATGCAA-1dCA2_MI_GFPve_pos -1.3024076300
## dCA2_Mpos_TCATACTCATCCTGTC-1dCA2_MI_GFPve_pos -0.8346867638
## dCA2_Mpos_TCATTACCAACTGGTT-1dCA2_MI_GFPve_pos -1.3610216456
## dCA2_Mpos_TCCGAAACAGTGTACT-1dCA2_MI_GFPve_pos -1.0520349818
## dCA2_Mpos_TCCGATCAGTGTCATC-1dCA2_MI_GFPve_pos -0.8408874350
## dCA2_Mpos_TCGCTCACACAGCCAC-1dCA2_MI_GFPve_pos -1.8836362439
## dCA2_Mpos_TCTATCATCATCACAG-1dCA2_MI_GFPve_pos -0.6629653650
## dCA2_Mpos_TGAACGTAGTGGTTAA-1dCA2_MI_GFPve_pos -1.5751434642
## dCA2_Mpos_TGATGCACATGACTGT-1dCA2_MI_GFPve_pos -1.5971435386
## dCA2_Mpos_TGGGAAGTCGTGGACC-1dCA2_MI_GFPve_pos -1.1675915795
## dCA2_Mpos_TGGTTAGAGAGCATTA-1dCA2_MI_GFPve_pos -0.1195332783
## dCA2_Mpos_TTTATGCTCATGCCCT-1dCA2_MI_GFPve_pos -1.0063589888
## dCA2_Fpos_CCGATGGAGCCTAACT-1dCA2_FI_GFPve_pos -1.1267625171
## dCA2_Fpos_CTGCGAGGTGTTGCCG-1dCA2_FI_GFPve_pos -0.7569624263
## dCA2_Fpos_GAAGAATAGCACTCTA-1dCA2_FI_GFPve_pos -1.5035114365
## dCA2_Fpos_TACATTCGTCGTGGAA-1dCA2_FI_GFPve_pos -1.6830409842
## FC_Mpos_AAACGCTCAATAGTGA-1FC_MI_GFPve_pos     -2.9221689778
## FC_Mpos_AAAGAACGTCTCAAGT-1FC_MI_GFPve_pos     -3.1027941781
## FC_Mpos_AAAGGGCTCTGCTCTG-1FC_MI_GFPve_pos     -2.6100564080
## FC_Mpos_AACACACGTTCCTAAG-1FC_MI_GFPve_pos     -3.1658015328
## FC_Mpos_AACCAACAGGTAGACC-1FC_MI_GFPve_pos     -1.1548628884
## FC_Mpos_AACCAACTCTCAGAAC-1FC_MI_GFPve_pos     -0.4745470362
## FC_Mpos_AACCATGTCTTCTGGC-1FC_MI_GFPve_pos     -1.9585447388
## FC_Mpos_AACTTCTAGTCTTCCC-1FC_MI_GFPve_pos     -2.9080307561
## FC_Mpos_AACTTCTGTGTCCTAA-1FC_MI_GFPve_pos     -3.5313093739
## FC_Mpos_AAGTACCCATGCTGCG-1FC_MI_GFPve_pos     -1.8835667449
## FC_Mpos_AATCACGAGTGGATTA-1FC_MI_GFPve_pos     -2.9551062661
## FC_Mpos_AATGAAGCATCCGTGG-1FC_MI_GFPve_pos     -2.3022570687
## FC_Mpos_AATGAAGTCATCCTAT-1FC_MI_GFPve_pos     -1.4818698245
```

```
## FC_Mpos_AATGACCAGGTAAAGG-1FC_MI_GFPve_pos     -3.5965313988
## FC_Mpos_ACAAAGAGTGGCTGAA-1FC_MI_GFPve_pos     -1.4724727946
## FC_Mpos_ACAACCACATATCTGG-1FC_MI_GFPve_pos     -1.1255862790
## FC_Mpos_ACACAGTCATAGGAGC-1FC_MI_GFPve_pos     -2.1654145795
## FC_Mpos_ACAGGGAGTTACCTTT-1FC_MI_GFPve_pos     -0.9981459456
## FC_Mpos_ACAGGGAGTTCAAGTC-1FC_MI_GFPve_pos     -3.4141385632
## FC_Mpos_ACATCGAGTATGAAGT-1FC_MI_GFPve_pos     -1.8085962611
## FC_Mpos_ACATTTCGTACAGAAT-1FC_MI_GFPve_pos     -0.5407782870
## FC_Mpos_ACCACAACAACCGCTG-1FC_MI_GFPve_pos     -1.3526386100
## FC_Mpos_ACCATTTTCCACACAA-1FC_MI_GFPve_pos     -1.4870247918
## FC_Mpos_ACGGGTCCAATTCGTG-1FC_MI_GFPve_pos     -3.0689385014
## FC_Mpos_ACGTACACATCACAGT-1FC_MI_GFPve_pos     -2.7759134847
## FC_Mpos_ACGTAGTAGCGTATGG-1FC_MI_GFPve_pos     -2.1067380982
## FC_Mpos_ACGTAGTAGGTACTGG-1FC_MI_GFPve_pos     -3.4318821507
## FC_Mpos_ACGTAGTCACTCGATA-1FC_MI_GFPve_pos     -1.2103574353
## FC_Mpos_ACTCCCAGTGCGGATA-1FC_MI_GFPve_pos     -3.4911630231
## FC_Mpos_ACTGTCCGTGAATTGA-1FC_MI_GFPve_pos     -2.6935966092
## FC_Mpos_AGAAATGTCAGCTGAT-1FC_MI_GFPve_pos     -0.9924420195
## FC_Mpos_AGAACCTAGTGGATAT-1FC_MI_GFPve_pos     -1.1895161944
## FC_Mpos_AGAGCAGCAGGCGTTC-1FC_MI_GFPve_pos     -1.8652637082
## FC_Mpos_AGATAGACAACTCCAA-1FC_MI_GFPve_pos     -3.3721454221
## FC_Mpos_AGATAGAGTAACCCTA-1FC_MI_GFPve_pos     -2.1272521096
## FC_Mpos_AGCGTATCATCGATCA-1FC_MI_GFPve_pos     -2.4395701962
## FC_Mpos_AGCTACAAGTACCGGA-1FC_MI_GFPve_pos     -1.6805750209
## FC_Mpos_AGGAGGTCAAGCGATG-1FC_MI_GFPve_pos     -3.4417440968
## FC_Mpos_AGGATCTGTGGACCTC-1FC_MI_GFPve_pos     -3.6362772065
## FC_Mpos_AGGGCCTGTGCATTAC-1FC_MI_GFPve_pos     -3.5862715321
## FC_Mpos_AGTCTCCAGAGGCCAT-1FC_MI_GFPve_pos     -1.5512595254
## FC_Mpos_AGTGATCGTCCTGTCT-1FC_MI_GFPve_pos     -3.4994049149
## FC_Mpos_AGTGCCGAGCGATCGA-1FC_MI_GFPve_pos     -3.3023350316
## FC_Mpos_AGTTCGATCATGCAGT-1FC_MI_GFPve_pos     -2.2524092274
## FC_Mpos_ATCCGTCAGGGTATAT-1FC_MI_GFPve_pos     -1.6811794120
## FC_Mpos_ATCGGCGAGACAGTCG-1FC_MI_GFPve_pos     -2.3683152276
## FC_Mpos_ATCGTAGAGGACGCTA-1FC_MI_GFPve_pos     -3.1030836182
## FC_Mpos_ATGATCGGTCCTCAGG-1FC_MI_GFPve_pos     -2.1706471520
## FC_Mpos_ATGCATGGTCGCTTAA-1FC_MI_GFPve_pos     -2.9268090802
## FC_Mpos_ATTACTCAGTCGCCCA-1FC_MI_GFPve_pos     -2.2222328263
## FC_Mpos_ATTCCATTCGGCTGAC-1FC_MI_GFPve_pos     -1.9585211354
## FC_Mpos_ATTCCTAAGTGAACAT-1FC_MI_GFPve_pos     -2.4963622170
## FC_Mpos_ATTCCTATCTAGACCA-1FC_MI_GFPve_pos     -1.1543159562
## FC_Mpos_ATTGGGTAGTAGGGTC-1FC_MI_GFPve_pos     -2.8940155583
## FC_Mpos_CAACAGTAGATGAATC-1FC_MI_GFPve_pos     -1.9439933377
## FC_Mpos_CAACCAAAGAGCCGTA-1FC_MI_GFPve_pos     -2.3656718808
## FC_Mpos_CAACGGCTCTGTCCCA-1FC_MI_GFPve_pos     -3.5899872857
## FC_Mpos_CAAGACTTCAGAGCGA-1FC_MI_GFPve_pos     -2.8435411530
## FC_Mpos_CACACAAGTTGTGCCG-1FC_MI_GFPve_pos     -2.3759284097
## FC_Mpos_CACGAATCATAAGCAA-1FC_MI_GFPve_pos     -1.3848487216
## FC_Mpos_CACTGAACAGGATTCT-1FC_MI_GFPve_pos     -1.9481344300
## FC_Mpos_CACTGAAGTGGTAATA-1FC_MI_GFPve_pos     -1.9117665368
## FC_Mpos_CAGATTGAGGGAGGAC-1FC_MI_GFPve_pos     -2.8671481686
## FC_Mpos_CAGCAATGTCTTTCTA-1FC_MI_GFPve_pos     -0.9050071317
## FC_Mpos_CAGTTAGAGTCTTCGA-1FC_MI_GFPve_pos     -2.3516318875
## FC_Mpos_CATCCACTCCGTGGGT-1FC_MI_GFPve_pos     -2.7240781861
## FC_Mpos_CATCGCTTCGGTATGT-1FC_MI_GFPve_pos     -3.4623928147
## FC_Mpos_CATCGGGCAATGCTCA-1FC_MI_GFPve_pos     -2.5105538445
## FC_Mpos_CATGCCTCACAGCGCT-1FC_MI_GFPve_pos     -1.3031132298
## FC_Mpos_CATTGAGGTTCTCCTG-1FC_MI_GFPve_pos     -1.3867969590
## FC_Mpos_CCAATTTGTAATCAAG-1FC_MI_GFPve_pos     -1.3640964108
## FC_Mpos_CCACGAGCATTGACAC-1FC_MI_GFPve_pos     -2.4225370961
## FC_Mpos_CCATCACGTGCCCACA-1FC_MI_GFPve_pos     -1.6101749020
## FC_Mpos_CCCATTGCATGAATCC-1FC_MI_GFPve_pos     -1.1310703831
## FC_Mpos_CCCGGAATCTCCACTG-1FC_MI_GFPve_pos     -3.4483265954
## FC_Mpos_CCGGGTAGTTAAACCC-1FC_MI_GFPve_pos     -1.9401965218
## FC_Mpos_CCGTTCACACTTCAGA-1FC_MI_GFPve_pos     -0.0668231177
## FC_Mpos_CCTATCGTCGAGCTGC-1FC_MI_GFPve_pos     -2.1250748711
## FC_Mpos_CCTCTAGCAACCCGCA-1FC_MI_GFPve_pos     -2.4653201180
## FC_Mpos_CCTCTCCAGGCCACCT-1FC_MI_GFPve_pos     -2.3511371690
## FC_Mpos_CCTGTTGAGAGTTGAT-1FC_MI_GFPve_pos     -2.3549995499
## FC_Mpos_CCTTCAGAGCGGTAGT-1FC_MI_GFPve_pos     -1.8588914948
## FC_Mpos_CGACAGCAGAGTCCGA-1FC_MI_GFPve_pos     -3.5795252400
```

```
## FC_Mpos_CGCATAAAGTGCTAGG-1FC_MI_GFPve_pos      -3.5413804131
## FC_Mpos_CGCATAAGTTGCGAAG-1FC_MI_GFPve_pos      -3.0777673798
## FC_Mpos_CGTAAGTGTCCCTCAT-1FC_MI_GFPve_pos      -1.9511222916
## FC_Mpos_CGTGTCTAGCGATTCT-1FC_MI_GFPve_pos      -2.0845589715
## FC_Mpos_CGTGTCTCACGAGGTA-1FC_MI_GFPve_pos      -2.4355108815
## FC_Mpos_CGTTCTGTCTTCTGGC-1FC_MI_GFPve_pos      -3.5437250214
## FC_Mpos_CTAACTTAGATGCTTC-1FC_MI_GFPve_pos      -3.2743086892
## FC_Mpos_CTACATTGTACTCGAT-1FC_MI_GFPve_pos      -0.7825027781
## FC_Mpos_CTAGGTATCTCCGATC-1FC_MI_GFPve_pos      -3.1846382695
## FC_Mpos_CTCATCGGTGTTACTG-1FC_MI_GFPve_pos      -1.9823422509
## FC_Mpos_CTCCACAAGCTACTAC-1FC_MI_GFPve_pos      -0.9072467166
## FC_Mpos_CTCCTCCAGCAGGCAT-1FC_MI_GFPve_pos      -0.9906023818
## FC_Mpos_CTGCAGGGTAGCGCCT-1FC_MI_GFPve_pos      -2.8174522000
## FC_Mpos_CTGCGAGTCCATGCAA-1FC_MI_GFPve_pos      -3.5514328557
## FC_Mpos_CTGGCAGGTTATTCTC-1FC_MI_GFPve_pos      -2.0165510255
## FC_Mpos_CTGTAGACAGCGGTCT-1FC_MI_GFPve_pos      -3.1025862771
## FC_Mpos_CTGTAGATCTCGCAGG-1FC_MI_GFPve_pos      -3.5519547540
## FC_Mpos_CTTCCGACAATCAAGA-1FC_MI_GFPve_pos      -3.5160365182
## FC_Mpos_GAAGCGACAAGTGGCA-1FC_MI_GFPve_pos      -1.6525082665
## FC_Mpos_GAATAGAGTAATACCC-1FC_MI_GFPve_pos      -2.8455951291
## FC_Mpos_GACACGCGTCCCGTGA-1FC_MI_GFPve_pos      -1.6571431237
## FC_Mpos_GACCCTTTCTACAGGT-1FC_MI_GFPve_pos      -3.4699692803
## FC_Mpos_GAGACTTGTTGACGGA-1FC_MI_GFPve_pos      -2.8767156678
## FC_Mpos_GAGAGGTCATCATTTC-1FC_MI_GFPve_pos      -3.1659317094
## FC_Mpos_GAGCCTGCAGGGATAC-1FC_MI_GFPve_pos      -3.3219695168
## FC_Mpos_GAGTCTACATTAAAGG-1FC_MI_GFPve_pos      -2.6165680962
## FC_Mpos_GAGTCTAGTCAGTCGC-1FC_MI_GFPve_pos      -3.5291047173
## FC_Mpos_GAGTCTAGTCGAATGG-1FC_MI_GFPve_pos      -3.5939338284
## FC_Mpos_GATCATGAGTCCCGGT-1FC_MI_GFPve_pos      -3.2784731465
## FC_Mpos_GATGATCGTACTCCCT-1FC_MI_GFPve_pos      -3.1586515981
## FC_Mpos_GATTCGATCATTGCCC-1FC_MI_GFPve_pos      -3.1416704732
## FC_Mpos_GATTTCTAGCCTCATA-1FC_MI_GFPve_pos      -1.5461080151
## FC_Mpos_GCACTAAGTTCGGTTA-1FC_MI_GFPve_pos      -2.0802307206
## FC_Mpos_GCACTAAGTTCTGACA-1FC_MI_GFPve_pos      -3.5269496518
## FC_Mpos_GCATCGGCAGCGTTTA-1FC_MI_GFPve_pos      -3.5588734227
## FC_Mpos_GCATCGGCATAGATGA-1FC_MI_GFPve_pos      -2.1930012780
## FC_Mpos_GCCAGGTAGACTTCCA-1FC_MI_GFPve_pos      -1.1619219857
## FC_Mpos_GCCGTGACATTATGCG-1FC_MI_GFPve_pos      -3.0426406937
## FC_Mpos_GCCTGTTTCCTGCCAT-1FC_MI_GFPve_pos      -0.6318827945
## FC_Mpos_GCGTTTCAGTGGCCTC-1FC_MI_GFPve_pos      -2.7995104867
## FC_Mpos_GCTGGGTGTCAAAGAT-1FC_MI_GFPve_pos      -2.9813127595
## FC_Mpos_GCTTGGGGTGCGGCTT-1FC_MI_GFPve_pos      -1.6911493617
## FC_Mpos_GGAGAACCAGCTGAAG-1FC_MI_GFPve_pos      -3.1389954167
## FC_Mpos_GGAGCAACACTTCTCG-1FC_MI_GFPve_pos      -3.5371260720
## FC_Mpos_GGAGCAAGTTATGTGC-1FC_MI_GFPve_pos      -3.4782548028
## FC_Mpos_GGCACGTAGCTAGCCC-1FC_MI_GFPve_pos      -2.8309762555
## FC_Mpos_GGCACGTCAACCGGAA-1FC_MI_GFPve_pos      -3.5484898167
## FC_Mpos_GGCTGTGCAAGTCGTT-1FC_MI_GFPve_pos      -3.0481069165
## FC_Mpos_GGGATCCGTAGCTTTG-1FC_MI_GFPve_pos      -3.2748432236
## FC_Mpos_GGGCGTTAGCATGATA-1FC_MI_GFPve_pos      -2.2610161382
## FC_Mpos_GGGCGTTGTAGCGCCT-1FC_MI_GFPve_pos      -1.9917023259
## FC_Mpos_GGGTTTATCGCAGTTA-1FC_MI_GFPve_pos      -2.4129564839
## FC_Mpos_GGTAATCAGAAATTCG-1FC_MI_GFPve_pos      -2.4721334058
## FC_Mpos_GGTCTGGGTAGGAAAG-1FC_MI_GFPve_pos      -3.4998958188
## FC_Mpos_GGTGAAGCACGAGGAT-1FC_MI_GFPve_pos      -3.0189585763
## FC_Mpos_GGTGATTGTCTCCTGT-1FC_MI_GFPve_pos      -3.0002808648
## FC_Mpos_GGTGTCGCACATGTTG-1FC_MI_GFPve_pos      -2.3205902654
## FC_Mpos_GTAATCGCAGCTGTGC-1FC_MI_GFPve_pos      -2.0790817815
## FC_Mpos_GTACAACGTATATGGA-1FC_MI_GFPve_pos      -1.1611974316
## FC_Mpos_GTCCACTTCGTTCATT-1FC_MI_GFPve_pos      -0.6810233074
## FC_Mpos_GTCCCATTCGAAATCC-1FC_MI_GFPve_pos      -2.5325591641
## FC_Mpos_GTCGCGAAGTTTGAGA-1FC_MI_GFPve_pos      -2.3939647752
## FC_Mpos_GTCGCGACAAACCATC-1FC_MI_GFPve_pos      -1.9318165856
## FC_Mpos_GTCGTTCCAGGACAGT-1FC_MI_GFPve_pos      -1.7521188336
## FC_Mpos_GTCTAGATCGTTCATT-1FC_MI_GFPve_pos      -1.8133784610
## FC_Mpos_GTGTCCTTCGCGCTGA-1FC_MI_GFPve_pos      -2.3473947125
## FC_Mpos_GTTACGACAATAGTCC-1FC_MI_GFPve_pos      -3.1644477921
## FC_Mpos_GTTCTATTCTCGTGGG-1FC_MI_GFPve_pos      -2.3461234647
## FC_Mpos_GTTGAACTCCCGTTCA-1FC_MI_GFPve_pos      -3.0593683797
## FC_Mpos_TAACACGTCAGGAAGC-1FC_MI_GFPve_pos      -2.0393807965
```

```
## FC_Mpos_TAAGTCGGTCGCCTAG-1FC_MI_GFPve_pos      -2.0436201173
## FC_Mpos_TAATCTCGTCGCAACC-1FC_MI_GFPve_pos      -2.2497828084
## FC_Mpos_TACCGGGGTAACATGA-1FC_MI_GFPve_pos      -2.2617023068
## FC_Mpos_TACCGGGTCTATCCAT-1FC_MI_GFPve_pos      -3.2679450589
## FC_Mpos_TACCTCGGTGTCCACG-1FC_MI_GFPve_pos      -2.1988582688
## FC_Mpos_TACCTGCGTCGCTTAA-1FC_MI_GFPve_pos      -1.1586582738
## FC_Mpos_TAGACCATCATCTACT-1FC_MI_GFPve_pos      -2.7222461777
## FC_Mpos_TAGAGTCTCTCGGGAC-1FC_MI_GFPve_pos      -1.7606127339
## FC_Mpos_TAGGTTGAGACTAAGT-1FC_MI_GFPve_pos      -1.2085149365
## FC_Mpos_TATCGCCCAGAACTAA-1FC_MI_GFPve_pos      -0.8193849402
## FC_Mpos_TATTCCAGTTGTGGCC-1FC_MI_GFPve_pos      -1.8834204751
## FC_Mpos_TATTGCTTCGTGCATA-1FC_MI_GFPve_pos      -3.5187542516
## FC_Mpos_TATTGGGAGAGTTGTA-1FC_MI_GFPve_pos      -1.4733686524
## FC_Mpos_TATTGGGGTATCAAGA-1FC_MI_GFPve_pos      -3.5597109872
## FC_Mpos_TCACGCTCATCAGTCA-1FC_MI_GFPve_pos      -3.5170154649
## FC_Mpos_TCACTATTCTGTTCAT-1FC_MI_GFPve_pos      -0.9634621220
## FC_Mpos_TCAGTGAGTCACCCTT-1FC_MI_GFPve_pos      -2.5130653458
## FC_Mpos_TCATACTTCTTCCAGC-1FC_MI_GFPve_pos      -0.8428113538
## FC_Mpos_TCATATCGTAGACACG-1FC_MI_GFPve_pos      -3.5209515172
## FC_Mpos_TCATCCGGTGTTCCAA-1FC_MI_GFPve_pos      -3.5806467610
## FC_Mpos_TCATGTTAGAGCTTTC-1FC_MI_GFPve_pos      -2.9913640099
## FC_Mpos_TCATTACAGTCTACCA-1FC_MI_GFPve_pos      -1.0386955815
## FC_Mpos_TCCACGTAGTCAGGGT-1FC_MI_GFPve_pos      -2.7573182660
## FC_Mpos_TCCATGCAGGCACCAA-1FC_MI_GFPve_pos      -3.3832075673
## FC_Mpos_TCCATGCCAACAGCCC-1FC_MI_GFPve_pos      -3.1357147771
## FC_Mpos_TCCTAATGTCCAGCCA-1FC_MI_GFPve_pos      -0.9315164166
## FC_Mpos_TCCTTTCAGGGCAGAG-1FC_MI_GFPve_pos      -2.1430883485
## FC_Mpos_TCGACGGTCGGACGTC-1FC_MI_GFPve_pos      -3.4332528191
## FC_Mpos_TCGTAGAAGCTAGTTC-1FC_MI_GFPve_pos      -1.6270792561
## FC_Mpos_TCTGCCATCAAGGAGC-1FC_MI_GFPve_pos      -3.2079844552
## FC_Mpos_TCTTCCTGTTACACAC-1FC_MI_GFPve_pos      -1.1945335942
## FC_Mpos_TCTTCCTGTTAGGAGC-1FC_MI_GFPve_pos      -0.6605603772
## FC_Mpos_TGAATCGTCCTCAGAA-1FC_MI_GFPve_pos      -2.9351654130
## FC_Mpos_TGACCCTAGTATTAGG-1FC_MI_GFPve_pos      -3.6560497361
## FC_Mpos_TGATGCACAAGTGGGT-1FC_MI_GFPve_pos      -1.9299881535
## FC_Mpos_TGCACGGAGAATGTTG-1FC_MI_GFPve_pos      -2.8352191525
## FC_Mpos_TGCACGGGTACGAGCA-1FC_MI_GFPve_pos      -2.1851694661
## FC_Mpos_TGCGACGGTAGGCAGT-1FC_MI_GFPve_pos      -2.7230327206
## FC_Mpos_TGCGATACAAGGCAAC-1FC_MI_GFPve_pos      -2.1915776807
## FC_Mpos_TGCTTGCTCACCCTCA-1FC_MI_GFPve_pos      -2.3427891808
## FC_Mpos_TGTAACGAGTGGCGAT-1FC_MI_GFPve_pos      -1.9505345898
## FC_Mpos_TGTACAGCAGCTCTGG-1FC_MI_GFPve_pos      -3.2188944894
## FC_Mpos_TGTGCGGGTCATAACC-1FC_MI_GFPve_pos      -2.8862192708
## FC_Mpos_TTATTGCTCAATGCAC-1FC_MI_GFPve_pos      -3.6127522069
## FC_Mpos_TTCACGCTCCATCACC-1FC_MI_GFPve_pos      -2.5527176934
## FC_Mpos_TTCCACGCAGGTTACT-1FC_MI_GFPve_pos      -3.3642275410
## FC_Mpos_TTCCTAACACGTCTCT-1FC_MI_GFPve_pos      -2.4076209145
## FC_Mpos_TTCGGTCCAGTTGCGC-1FC_MI_GFPve_pos      -2.2381260472
## FC_Mpos_TTCTGTAAGCGGATCA-1FC_MI_GFPve_pos      -2.5529704171
## FC_Mpos_TTCTTGAGTTGATCGT-1FC_MI_GFPve_pos      -2.3214027959
## FC_Mpos_TTGAACGAGGATAATC-1FC_MI_GFPve_pos      -1.3583308535
## FC_Mpos_TTGACCCTCAGCTCTC-1FC_MI_GFPve_pos      -2.6131787377
## FC_Mpos_TTGAGTGCATGTTTGG-1FC_MI_GFPve_pos      -3.1403007584
## FC_Mpos_TTGCCTGCAGCCTTCT-1FC_MI_GFPve_pos      -3.2489981728
## FC_Mpos_TTGTTGTGTATCACGT-1FC_MI_GFPve_pos      -2.2579426843
## FC_Mpos_TTTAGTCGTCCGACGT-1FC_MI_GFPve_pos      -3.3205599862
## FC_Mpos_TTTCAGTGTCAAATCC-1FC_MI_GFPve_pos      -2.8819413262
## FC_Mpos_TTTGGTTGTGCGCTCA-1FC_MI_GFPve_pos      -1.8912298756
## FC_Fpos_AAAGGATTCTAGCCTC-1FC_FI_GFPve_pos      -1.2687993127
## FC_Fpos_AAAGGTAAGTGTCATC-1FC_FI_GFPve_pos      -3.0391330796
## FC_Fpos_AAAGTCCCAGAACTAA-1FC_FI_GFPve_pos      -2.9777782040
## FC_Fpos_AACCCAACAAGCCTGC-1FC_FI_GFPve_pos      -2.4308812695
## FC_Fpos_AACCTTTGTCTGTGGC-1FC_FI_GFPve_pos      -3.0071356374
## FC_Fpos_AACGGGAAGCAGGTCA-1FC_FI_GFPve_pos      -2.5929639417
## FC_Fpos_AAGGTAAAGGAACTAT-1FC_FI_GFPve_pos      -2.2639629918
## FC_Fpos_AATAGAGCAACGCCCA-1FC_FI_GFPve_pos      -2.0803997594
## FC_Fpos_AATGCCATCATCTATC-1FC_FI_GFPve_pos      -0.4495761472
## FC_Fpos_ACACGCGTCGGTGTAT-1FC_FI_GFPve_pos      -1.5400755482
## FC_Fpos_ACAGCCGCATGGTGGA-1FC_FI_GFPve_pos      -2.7931349355
## FC_Fpos_ACCCTCATCTCAGGCG-1FC_FI_GFPve_pos      -2.3615195828
```

```
## FC_Fpos_ACGGTTACAAGTGACG-1FC_FI_GFPve_pos      -3.5421776849
## FC_Fpos_ACTGATGGTTTCGACA-1FC_FI_GFPve_pos      -2.8742985802
## FC_Fpos_ACTGTGACACTGGACC-1FC_FI_GFPve_pos      -3.1326892453
## FC_Fpos_ACTTTGTAGGTCGAGT-1FC_FI_GFPve_pos      -1.5559213239
## FC_Fpos_AGACCCGTCTCGGGAC-1FC_FI_GFPve_pos      -3.1871552544
## FC_Fpos_AGAGCCCCAACAGCTT-1FC_FI_GFPve_pos      -2.6793432313
## FC_Fpos_AGCCACGTCTTGGCTC-1FC_FI_GFPve_pos      -1.6920826512
## FC_Fpos_AGCGTATCACGTTCGG-1FC_FI_GFPve_pos      -3.0665533620
## FC_Fpos_AGGATCTGTCATGGCC-1FC_FI_GFPve_pos      -2.3759553510
## FC_Fpos_AGGCCACAGTAACGAT-1FC_FI_GFPve_pos      -2.1809897500
## FC_Fpos_AGGTCTAGTGACCGAA-1FC_FI_GFPve_pos      -2.6360325890
## FC_Fpos_AGTCACAAGCCATTGT-1FC_FI_GFPve_pos      -2.3969135361
## FC_Fpos_AGTCTCCGTCTTTCTA-1FC_FI_GFPve_pos      -2.2687149125
## FC_Fpos_AGTGTTGCACGCGTCA-1FC_FI_GFPve_pos      -3.3574435788
## FC_Fpos_ATAGAGATCGTCTCAC-1FC_FI_GFPve_pos      -3.5042765218
## FC_Fpos_ATCAGGTAGAGTTGTA-1FC_FI_GFPve_pos      -2.4137456494
## FC_Fpos_ATCGTGAGTAGGCTGA-1FC_FI_GFPve_pos      -1.8702133971
## FC_Fpos_ATGTCCCCAATGTGGG-1FC_FI_GFPve_pos      -1.0180535394
## FC_Fpos_ATTACCTGTACCAGAG-1FC_FI_GFPve_pos      -0.9454609233
## FC_Fpos_ATTCGTTAGGACGGAG-1FC_FI_GFPve_pos      -1.5269115048
## FC_Fpos_ATTCTTGGTGAGCCAA-1FC_FI_GFPve_pos      -2.1762096959
## FC_Fpos_ATTTCACTCGCAATTG-1FC_FI_GFPve_pos      -2.0907428342
## FC_Fpos_CAATACGAGCGCTTCG-1FC_FI_GFPve_pos      -2.9267242032
## FC_Fpos_CAATGACTCGTGGACC-1FC_FI_GFPve_pos      -1.2918815690
## FC_Fpos_CACAACAGTGACAGGT-1FC_FI_GFPve_pos      -3.3888902741
## FC_Fpos_CACGTGGAGCTCACTA-1FC_FI_GFPve_pos      -2.6956148225
## FC_Fpos_CATGCAACATGGCGCT-1FC_FI_GFPve_pos      -0.4436981159
## FC_Fpos_CATGGATCAACTGAAA-1FC_FI_GFPve_pos      -1.3668890076
## FC_Fpos_CATTGAGTCTGGGCCA-1FC_FI_GFPve_pos      -3.1082267838
## FC_Fpos_CATTGTTCAGGACGAT-1FC_FI_GFPve_pos      -1.7765951234
## FC_Fpos_CATTTCAAGACTCCGC-1FC_FI_GFPve_pos      -3.3622093278
## FC_Fpos_CCACAAACAATGCAAA-1FC_FI_GFPve_pos      -3.4310157376
## FC_Fpos_CCACGAGGTATGGTAA-1FC_FI_GFPve_pos      -3.0072641450
## FC_Fpos_CCACTTGAGATCGCCC-1FC_FI_GFPve_pos      -2.7704701501
## FC_Fpos_CCCATTGTCTCACTCG-1FC_FI_GFPve_pos      -3.4652891236
## FC_Fpos_CCGCAAGCAAGTATCC-1FC_FI_GFPve_pos      -2.2386357861
## FC_Fpos_CCGGACATCATTCCTA-1FC_FI_GFPve_pos      -1.8548547107
## FC_Fpos_CCTAACCGTCCGAAAG-1FC_FI_GFPve_pos      -2.2262437421
## FC_Fpos_CGATGGCAGGAGCTGT-1FC_FI_GFPve_pos      -1.6189667064
## FC_Fpos_CGCAGGTCAAAGACGC-1FC_FI_GFPve_pos      -3.3317992764
## FC_Fpos_CGGAGAAGTATGGTAA-1FC_FI_GFPve_pos      -1.4806761819
## FC_Fpos_CGTAGTAAGAGATTCA-1FC_FI_GFPve_pos      -1.7878962832
## FC_Fpos_CGTCAAAAGGGATCTG-1FC_FI_GFPve_pos      -2.2352368909
## FC_Fpos_CGTCAAAGTGTCCACG-1FC_FI_GFPve_pos      -1.9256041127
## FC_Fpos_CGTGAATTCGGTCACG-1FC_FI_GFPve_pos      -1.4766727763
## FC_Fpos_CGTGATAGTTGTAGCT-1FC_FI_GFPve_pos      -1.6678454953
## FC_Fpos_CTACAGACACTGGATT-1FC_FI_GFPve_pos      -3.5053424912
## FC_Fpos_CTACCTGAGTCTGCAT-1FC_FI_GFPve_pos      -2.0276792126
## FC_Fpos_CTACCTGGTCCCTGTT-1FC_FI_GFPve_pos      -2.1447618085
## FC_Fpos_CTCCACATCAGCGCGT-1FC_FI_GFPve_pos      -2.4163999635
## FC_Fpos_CTCCTTTTCCCTCATG-1FC_FI_GFPve_pos      -3.4470982629
## FC_Fpos_CTGCAGGGTTTCGTTT-1FC_FI_GFPve_pos      -3.1557247716
## FC_Fpos_CTGCCATAGTAATACG-1FC_FI_GFPve_pos      -3.5545730668
## FC_Fpos_CTTCCGACAGCTGTTA-1FC_FI_GFPve_pos      -3.5775551873
## FC_Fpos_CTTCCTTTCGAAGCAG-1FC_FI_GFPve_pos      -3.6258242207
## FC_Fpos_GAACACTAGATTGTGA-1FC_FI_GFPve_pos      -1.3893413621
## FC_Fpos_GAAGTAACACAGTGTT-1FC_FI_GFPve_pos      -1.3383859473
## FC_Fpos_GACACGCTCTACTGAG-1FC_FI_GFPve_pos      -1.9476337510
## FC_Fpos_GACCAATTCACTTATC-1FC_FI_GFPve_pos      -3.2614760476
## FC_Fpos_GACCTTCCATGTTACG-1FC_FI_GFPve_pos      -3.2633953172
## FC_Fpos_GAGATGGAGCACGTCC-1FC_FI_GFPve_pos      -2.6614110547
## FC_Fpos_GAGCCTGTCTAAGAAG-1FC_FI_GFPve_pos      -1.8476656752
## FC_Fpos_GAGTCATAGCACTGGA-1FC_FI_GFPve_pos      -3.3752744275
## FC_Fpos_GAGTCTAGTTGTGCCG-1FC_FI_GFPve_pos      -2.0365798550
## FC_Fpos_GAGTTGTGTAGGCAAC-1FC_FI_GFPve_pos      -2.3564255315
## FC_Fpos_GATCCCTCAGTGAGCA-1FC_FI_GFPve_pos      -1.6389666873
## FC_Fpos_GATTCTTGTAAGGTCG-1FC_FI_GFPve_pos      -1.9073917943
## FC_Fpos_GCACATACAGGTGAGT-1FC_FI_GFPve_pos      -1.5501357394
## FC_Fpos_GCACGTGAGAGTCACG-1FC_FI_GFPve_pos      -2.1333239156
## FC_Fpos_GCAGTTAAGGAAGTCC-1FC_FI_GFPve_pos      -2.2461066323
```

```
## FC_Fpos_GCAGTTATCAAACCTG-1FC_FI_GFPve_pos     -1.7833883839
## FC_Fpos_GCATCGGTCAAGCGTT-1FC_FI_GFPve_pos     -3.4785842973
## FC_Fpos_GCCTGTTCACGGTGTC-1FC_FI_GFPve_pos     -2.5650763589
## FC_Fpos_GCTTGGGAGATCGACG-1FC_FI_GFPve_pos     -1.7709878760
## FC_Fpos_GCTTTCGGTGTGTCCG-1FC_FI_GFPve_pos     -1.5513625222
## FC_Fpos_GGAGATGAGTCGGCAA-1FC_FI_GFPve_pos     -1.6559553223
## FC_Fpos_GGAGGTATCGTTGTTT-1FC_FI_GFPve_pos     -2.7091412621
## FC_Fpos_GGCAGTCCATCGCTGG-1FC_FI_GFPve_pos     -1.5265082198
## FC_Fpos_GGGTATTTCCAACTGA-1FC_FI_GFPve_pos     -2.1967842656
## FC_Fpos_GGTGTTATCAACACCA-1FC_FI_GFPve_pos     -1.3819662410
## FC_Fpos_GGTTGTAGTCACATTG-1FC_FI_GFPve_pos     -2.0458717423
## FC_Fpos_GTAGGTTTCACATTGG-1FC_FI_GFPve_pos     -2.7443280297
## FC_Fpos_GTAGTACTCGTGTCAA-1FC_FI_GFPve_pos     -2.5017428475
## FC_Fpos_GTATTGGGTCGAACGA-1FC_FI_GFPve_pos     -1.8962107974
## FC_Fpos_GTCAAACCATCCGGCA-1FC_FI_GFPve_pos     -2.3094809132
## FC_Fpos_GTCACGGTCTTCTGTA-1FC_FI_GFPve_pos     -0.9339861947
## FC_Fpos_GTCACTCAGAAGCGAA-1FC_FI_GFPve_pos     -1.4948128539
## FC_Fpos_GTCACTCAGTCGTCTA-1FC_FI_GFPve_pos     -3.2093689519
## FC_Fpos_GTCCACTCAATTGCTG-1FC_FI_GFPve_pos     -1.4304306584
## FC_Fpos_GTCTCACAGTGCACTT-1FC_FI_GFPve_pos     -1.0333743172
## FC_Fpos_GTGAGTTGTGCAACGA-1FC_FI_GFPve_pos     -1.7935322600
## FC_Fpos_GTGAGTTTCGGTCTGG-1FC_FI_GFPve_pos     -1.3097450810
## FC_Fpos_GTGCTTCAGCGTGCTC-1FC_FI_GFPve_pos     -1.4259065467
## FC_Fpos_GTGGAAGTCGTCCTCA-1FC_FI_GFPve_pos     -1.6054647046
## FC_Fpos_GTGGAGATCCATTGTT-1FC_FI_GFPve_pos     -1.5428117591
## FC_Fpos_GTTCATTGTCTACAAC-1FC_FI_GFPve_pos     -2.9471120911
## FC_Fpos_GTTGCTCCAAATGAGT-1FC_FI_GFPve_pos     -1.9584844189
## FC_Fpos_TAAGCACAGTGCAACG-1FC_FI_GFPve_pos     -1.4593131619
## FC_Fpos_TACGGTATCACTGCTC-1FC_FI_GFPve_pos     -2.7698891240
## FC_Fpos_TACTTCAGTTAGGCTT-1FC_FI_GFPve_pos     -3.4060175496
## FC_Fpos_TAGGGTTGTTACGGAG-1FC_FI_GFPve_pos     -1.6673050004
## FC_Fpos_TAGTGCACACCAGTAT-1FC_FI_GFPve_pos     -2.3201520520
## FC_Fpos_TATACCTAGCATCTTG-1FC_FI_GFPve_pos     -1.9269912320
## FC_Fpos_TATACCTCATGGCACC-1FC_FI_GFPve_pos     -2.2338006573
## FC_Fpos_TATATCCTCGTGAGAG-1FC_FI_GFPve_pos     -1.6741067248
## FC_Fpos_TATCTGTGTGCTTATG-1FC_FI_GFPve_pos     -2.1975364762
## FC_Fpos_TATTGCTTCACAGTGT-1FC_FI_GFPve_pos     -2.0674779492
## FC_Fpos_TATTGGGCACCAGCCA-1FC_FI_GFPve_pos     -3.3868801671
## FC_Fpos_TATTTCGGTCATCGCG-1FC_FI_GFPve_pos     -2.2405750829
## FC_Fpos_TCACAAGCAGGGAATC-1FC_FI_GFPve_pos     -3.3534326630
## FC_Fpos_TCAGCCTCAACACAAA-1FC_FI_GFPve_pos     -2.0430579263
## FC_Fpos_TCATCATAGTGCACCC-1FC_FI_GFPve_pos     -2.4735968190
## FC_Fpos_TCATGGAAGCTGTGCC-1FC_FI_GFPve_pos     -2.2060659009
## FC_Fpos_TCCACGTAGGTGATCG-1FC_FI_GFPve_pos     -1.8662579137
## FC_Fpos_TCCACGTGTTGGAGAC-1FC_FI_GFPve_pos     -3.4018981534
## FC_Fpos_TCCATGCGTCCGGTGT-1FC_FI_GFPve_pos     -1.7281214076
## FC_Fpos_TCCCAGTTCGTGTTCC-1FC_FI_GFPve_pos     -2.3706309872
## FC_Fpos_TCCGAAATCTCGCGTT-1FC_FI_GFPve_pos     -1.5917335826
## FC_Fpos_TCCTTCTAGAGCATAT-1FC_FI_GFPve_pos     -2.8831932622
## FC_Fpos_TCGACCTGTTAGGCCC-1FC_FI_GFPve_pos     -3.0354440289
## FC_Fpos_TCGCAGGAGTTGGCGA-1FC_FI_GFPve_pos     -2.5002343732
## FC_Fpos_TCGCAGGCAAACTCTG-1FC_FI_GFPve_pos     -1.7668579894
## FC_Fpos_TCGCTCAGTCAAACGG-1FC_FI_GFPve_pos     -3.0882995206
## FC_Fpos_TCTAACTTCGTCTAAG-1FC_FI_GFPve_pos     -1.8644968348
## FC_Fpos_TCTGCCACACTGGCGT-1FC_FI_GFPve_pos     -3.5380992966
## FC_Fpos_TCTGCCAGTACAGAGC-1FC_FI_GFPve_pos     -1.5615503865
## FC_Fpos_TCTGGCTAGGCCTAAG-1FC_FI_GFPve_pos     -2.2826013642
## FC_Fpos_TGACTCCAGACGATAT-1FC_FI_GFPve_pos     -3.5914778786
## FC_Fpos_TGATTTCAGCGCACAA-1FC_FI_GFPve_pos     -2.1344261246
## FC_Fpos_TGCAGTATCTTGAGCA-1FC_FI_GFPve_pos     -1.5004205781
## FC_Fpos_TGGCGTGAGCCGCTTG-1FC_FI_GFPve_pos     -3.4041066247
## FC_Fpos_TGGGATTGTCGAAGCA-1FC_FI_GFPve_pos     -1.5296430665
## FC_Fpos_TGGTAGTAGTTTGGCT-1FC_FI_GFPve_pos     -2.8747680264
## FC_Fpos_TGTCCACAGGACCCAA-1FC_FI_GFPve_pos     -1.6807206946
## FC_Fpos_TGTCCTGAGCAAATCA-1FC_FI_GFPve_pos     -1.5896611291
## FC_Fpos_TGTCCTGCAATTCACG-1FC_FI_GFPve_pos     -2.3339946347
## FC_Fpos_TGTTGGAGTCGGCACT-1FC_FI_GFPve_pos     -3.1956675130
## FC_Fpos_TTAATCCTCTTCGGAA-1FC_FI_GFPve_pos     -3.4197785931
## FC_Fpos_TTCACGCTCATAGAGA-1FC_FI_GFPve_pos     -1.3861892300
## FC_Fpos_TTCATGTGTTCAACGT-1FC_FI_GFPve_pos     -3.5351171571
```

```
## FC_Fpos_TTCCAATGTCACGACC-1FC_FI_GFPve_pos     -3.4451894837
## FC_Fpos_TTCCTTCAGATGTTCC-1FC_FI_GFPve_pos     -1.6901812630
## FC_Fpos_TTCCTTCCACAATGTC-1FC_FI_GFPve_pos     -2.1690156537
## FC_Fpos_TTCGGTCAGTTTCGGT-1FC_FI_GFPve_pos     -1.8014149743
## FC_Fpos_TTCGGTCTCAACCGAT-1FC_FI_GFPve_pos     -1.7945425587
## FC_Fpos_TTCTGTACACCAAAGG-1FC_FI_GFPve_pos     -1.7991482096
## FC_Fpos_TTCTTCCCAACGCATT-1FC_FI_GFPve_pos     -2.2654943543
## FC_Fpos_TTGAGTGTCGGACGTC-1FC_FI_GFPve_pos     -2.8209018784
## FC_Fpos_TTGCCTGAGTATAGGT-1FC_FI_GFPve_pos     -1.3081783133
## FC_Fpos_TTGGATGAGGTCCAGA-1FC_FI_GFPve_pos     -2.1267097073
## FC_Fpos_TTGTTGTCACCCTCTA-1FC_FI_GFPve_pos     -2.7219433862
## FC_Fpos_TTGTTTGCAGACACCC-1FC_FI_GFPve_pos     -1.9285025674
## dCA2_Mpos_AAAGAACCAATCACGT-1dCA2_MI_GFPve_pos  0.8765053672
## dCA2_Mpos_AAAGTCCTCCGAGGCT-1dCA2_MI_GFPve_pos  0.4660556835
## dCA2_Mpos_AACAACCTCGACGAGA-1dCA2_MI_GFPve_pos  1.3822704477
## dCA2_Mpos_ACAACCAAGAGCCGAT-1dCA2_MI_GFPve_pos  1.6679625434
## dCA2_Mpos_ACACGCGAGCACCGAA-1dCA2_MI_GFPve_pos  2.4308118743
## dCA2_Mpos_ACGATGTGTCTACTGA-1dCA2_MI_GFPve_pos  0.7772107047
## dCA2_Mpos_ACGGAAGCAATGTCTG-1dCA2_MI_GFPve_pos  0.2539464456
## dCA2_Mpos_ACTGCAATCACGTCCT-1dCA2_MI_GFPve_pos  2.3885254783
## dCA2_Mpos_ACTGTCCCAGTCAGCC-1dCA2_MI_GFPve_pos  2.1990313453
## dCA2_Mpos_AGCTTCCAGTTGTCGT-1dCA2_MI_GFPve_pos  1.4989464206
## dCA2_Mpos_AGGATAAAGACGAAGA-1dCA2_MI_GFPve_pos  2.0024058742
## dCA2_Mpos_AGGCCACTCGATCCAA-1dCA2_MI_GFPve_pos  2.3200967235
## dCA2_Mpos_AGGCTGCAGTCACTAC-1dCA2_MI_GFPve_pos  1.4496545715
## dCA2_Mpos_AGGCTGCCAAGACAAT-1dCA2_MI_GFPve_pos  1.9817635936
## dCA2_Mpos_AGGGTTTCATTCTCTA-1dCA2_MI_GFPve_pos  2.3185810966
## dCA2_Mpos_ATGCCTCGTGTTGCCG-1dCA2_MI_GFPve_pos  0.6658832950
## dCA2_Mpos_ATGGGTTAGCAAGGAA-1dCA2_MI_GFPve_pos  0.7339765948
## dCA2_Mpos_ATTCTACTCAGGAAAT-1dCA2_MI_GFPve_pos  1.3611758870
## dCA2_Mpos_ATTTACCCACATGTTG-1dCA2_MI_GFPve_pos  1.7008300943
## dCA2_Mpos_CAAGACTTCAGCTAGT-1dCA2_MI_GFPve_pos  1.1780718488
## dCA2_Mpos_CACACAAAGCTTTCCC-1dCA2_MI_GFPve_pos  1.8803765697
## dCA2_Mpos_CACTAAGCACAAAGTA-1dCA2_MI_GFPve_pos  1.3204522056
## dCA2_Mpos_CAGCAATTCCGAGTGC-1dCA2_MI_GFPve_pos  2.3333198947
## dCA2_Mpos_CAGCCAGAGACAACTA-1dCA2_MI_GFPve_pos  1.4772205276
## dCA2_Mpos_CATCGTCAGTAATACG-1dCA2_MI_GFPve_pos  2.5504760665
## dCA2_Mpos_CCAATTTCATTGCTTT-1dCA2_MI_GFPve_pos  2.2806348724
## dCA2_Mpos_CCATCACCATGACTCA-1dCA2_MI_GFPve_pos  1.3678700847
## dCA2_Mpos_CCGAACGCAGTTGTCA-1dCA2_MI_GFPve_pos  2.3141090793
## dCA2_Mpos_CCTCCTCCAGACCTAT-1dCA2_MI_GFPve_pos  1.5034912748
## dCA2_Mpos_CGAGAAGAGATAGCTA-1dCA2_MI_GFPve_pos  0.8892225546
## dCA2_Mpos_CGCCATTGTCACCGCA-1dCA2_MI_GFPve_pos  2.5501034183
## dCA2_Mpos_CGGGTGTTCTCATGCC-1dCA2_MI_GFPve_pos  1.6104745788
## dCA2_Mpos_CGGTCAGGTAGCTGTT-1dCA2_MI_GFPve_pos  2.4131553096
## dCA2_Mpos_CGTGATATCACGACTA-1dCA2_MI_GFPve_pos  0.3810659868
## dCA2_Mpos_CGTGCTTAGTAATCCC-1dCA2_MI_GFPve_pos  0.4849992019
## dCA2_Mpos_CTATAGGCAGGACATG-1dCA2_MI_GFPve_pos  2.6016871852
## dCA2_Mpos_CTCAAGACATAGAGGC-1dCA2_MI_GFPve_pos  2.4038836879
## dCA2_Mpos_CTCATTAAGTCCTACA-1dCA2_MI_GFPve_pos  1.4117412490
## dCA2_Mpos_CTCCTTTCAGATACTC-1dCA2_MI_GFPve_pos  2.4861915034
## dCA2_Mpos_CTGCGAGGTTGCCGAC-1dCA2_MI_GFPve_pos  0.7888714594
## dCA2_Mpos_CTTCGGTCAGGCAATG-1dCA2_MI_GFPve_pos  0.7949679417
## dCA2_Mpos_GAACTGTTCACCACAA-1dCA2_MI_GFPve_pos  1.3015946073
## dCA2_Mpos_GACCCAGTCTGGCCTT-1dCA2_MI_GFPve_pos  2.3063063544
## dCA2_Mpos_GAGCCTGCAGGAGACT-1dCA2_MI_GFPve_pos  0.5621817631
## dCA2_Mpos_GAGGCAAAGTCTCCTC-1dCA2_MI_GFPve_pos  2.2165489120
## dCA2_Mpos_GATTGGTTCTTCTTCC-1dCA2_MI_GFPve_pos  1.4593371076
## dCA2_Mpos_GCATTAGGTTGTCCCT-1dCA2_MI_GFPve_pos  0.4072417540
## dCA2_Mpos_GGAACCCTCTTCCTAA-1dCA2_MI_GFPve_pos  1.7201685828
## dCA2_Mpos_GGATCTATCTAAGCGT-1dCA2_MI_GFPve_pos  0.6142855806
## dCA2_Mpos_GGCTTTCCAAATCCCA-1dCA2_MI_GFPve_pos  1.1714962644
## dCA2_Mpos_GGGTGTCAGATACCAA-1dCA2_MI_GFPve_pos  2.4563703460
## dCA2_Mpos_GGTAACTAGTCACTCA-1dCA2_MI_GFPve_pos  1.2417956514
## dCA2_Mpos_GTCAAGTAGGCATTTC-1dCA2_MI_GFPve_pos  2.5358424110
## dCA2_Mpos_GTCAGCGCAAGCCTGC-1dCA2_MI_GFPve_pos  1.3838872832
## dCA2_Mpos_GTCCCATTCCAAGCTA-1dCA2_MI_GFPve_pos  1.5288358850
## dCA2_Mpos_GTTAGACAGATAACAC-1dCA2_MI_GFPve_pos  1.1565318031
## dCA2_Mpos_TACATTCTCTAGCCAA-1dCA2_MI_GFPve_pos  2.1559395713
## dCA2_Mpos_TACCCACCACGGCACT-1dCA2_MI_GFPve_pos  1.8408726377
```

```
## dCA2_Mpos_TACCGAAAGCAGTAAT-1dCA2_MI_GFPve_pos  0.5452070457
## dCA2_Mpos_TACGCTCCACCTGCTT-1dCA2_MI_GFPve_pos  1.0580404920
## dCA2_Mpos_TACTTACAGTGGGAAA-1dCA2_MI_GFPve_pos  2.1570587081
## dCA2_Mpos_TACTTGTCAGGTGGAT-1dCA2_MI_GFPve_pos  1.0728660745
## dCA2_Mpos_TCAATCTTCGCGCTGA-1dCA2_MI_GFPve_pos  1.1897514981
## dCA2_Mpos_TCACATTCAGAGACTG-1dCA2_MI_GFPve_pos  1.0102271957
## dCA2_Mpos_TCACGCTAGCGTTAGG-1dCA2_MI_GFPve_pos  1.9159120244
## dCA2_Mpos_TCACTCGGTTGTGTAC-1dCA2_MI_GFPve_pos  2.1787421626
## dCA2_Mpos_TCAGCCTTCCTTTAGT-1dCA2_MI_GFPve_pos  2.4869522971
## dCA2_Mpos_TCCTCTTAGTTAACGA-1dCA2_MI_GFPve_pos  0.7529463095
## dCA2_Mpos_TCGCAGGGTTTGGAGG-1dCA2_MI_GFPve_pos  1.1341535968
## dCA2_Mpos_TCGCTCAGTCTATGAC-1dCA2_MI_GFPve_pos  1.8306900186
## dCA2_Mpos_TCTTAGTGTGCAACGA-1dCA2_MI_GFPve_pos  1.5786112470
## dCA2_Mpos_TGGGAGACAGTAGATA-1dCA2_MI_GFPve_pos  1.4277758521
## dCA2_Mpos_TGGGCTGGTCTCGGAC-1dCA2_MI_GFPve_pos  2.2654263896
## dCA2_Mpos_TGTAAGCTCGATGGAG-1dCA2_MI_GFPve_pos  1.6680408639
## dCA2_Mpos_TGTCCCAAGCAGCCTC-1dCA2_MI_GFPve_pos  1.7904201669
## dCA2_Mpos_TGTGCGGGTACTGACT-1dCA2_MI_GFPve_pos  2.4171600265
## dCA2_Mpos_TTGACCCGTTGCTTGA-1dCA2_MI_GFPve_pos  2.3625435752
## dCA2_Mpos_TTGGGCGTCTCGACCT-1dCA2_MI_GFPve_pos  1.3356033487
## dCA2_Mpos_TTGGGCGTCTGCGGGT-1dCA2_MI_GFPve_pos  1.3439402503
## dCA2_Mpos_TTGGGTATCCCTTGGT-1dCA2_MI_GFPve_pos  0.7435964865
## dCA2_Mpos_TTTACCACATCAGTCA-1dCA2_MI_GFPve_pos  1.4340819043
## dCA2_Mpos_TTTCCTCTCTATTCGT-1dCA2_MI_GFPve_pos  2.1642434520
## dCA2_Fpos_AAAGTCCGTGGACAGT-1dCA2_FI_GFPve_pos  1.3053517265
## dCA2_Fpos_ACACCAAAGAGGTATT-1dCA2_FI_GFPve_pos  1.3204278869
## dCA2_Fpos_ACAGGGAGTGTAAATG-1dCA2_FI_GFPve_pos  2.5815544051
## dCA2_Fpos_ACTTCGCAGTAGGGTC-1dCA2_FI_GFPve_pos  2.3264002723
## dCA2_Fpos_ATGGATCGTATGCTTG-1dCA2_FI_GFPve_pos  2.3462605399
## dCA2_Fpos_CAACGATGTTTCCATT-1dCA2_FI_GFPve_pos  0.1352641617
## dCA2_Fpos_CACCAAAGTCTACTGA-1dCA2_FI_GFPve_pos  2.3741140289
## dCA2_Fpos_CAGAGCCGTATCGCAT-1dCA2_FI_GFPve_pos  1.3780213517
## dCA2_Fpos_CAGATTGTCATCGTAG-1dCA2_FI_GFPve_pos  0.8409592432
## dCA2_Fpos_CCACTTGGTCTTTCTA-1dCA2_FI_GFPve_pos  1.3399261159
## dCA2_Fpos_CCATCACGTTCTTCAT-1dCA2_FI_GFPve_pos  1.1449887675
## dCA2_Fpos_CCCTGATTCAAATGCC-1dCA2_FI_GFPve_pos  1.5225293559
## dCA2_Fpos_CCGATCTAGTGAGGCT-1dCA2_FI_GFPve_pos  0.3981521351
## dCA2_Fpos_CCGGGTATCTCCAAGA-1dCA2_FI_GFPve_pos  1.1464068813
## dCA2_Fpos_CGCGTGAAGCGCATCC-1dCA2_FI_GFPve_pos  0.7865990323
## dCA2_Fpos_CGTAAGTGTCCCTGAG-1dCA2_FI_GFPve_pos  0.5021098179
## dCA2_Fpos_CTATCCGGTGATGGCA-1dCA2_FI_GFPve_pos  2.2820711059
## dCA2_Fpos_CTCCACAAGATGCGAC-1dCA2_FI_GFPve_pos  0.5962531192
## dCA2_Fpos_CTCCCTCAGGTTGGAC-1dCA2_FI_GFPve_pos  0.7124493641
## dCA2_Fpos_CTGGACGTCATGGTAC-1dCA2_FI_GFPve_pos  1.4094729346
## dCA2_Fpos_GACACGCAGTCTTCGA-1dCA2_FI_GFPve_pos  1.3470218105
## dCA2_Fpos_GATGACTAGAGCAGTC-1dCA2_FI_GFPve_pos  2.1414794845
## dCA2_Fpos_GGAGGATGTATTCCGA-1dCA2_FI_GFPve_pos  0.5390695852
## dCA2_Fpos_GGCTTTCGTAGATCGG-1dCA2_FI_GFPve_pos  2.6068546695
## dCA2_Fpos_GGTGATTAGACGCATG-1dCA2_FI_GFPve_pos  2.5011713428
## dCA2_Fpos_GTGGCGTCAGAAGCGT-1dCA2_FI_GFPve_pos  0.2169704360
## dCA2_Fpos_TAACCAGTCAACCTCC-1dCA2_FI_GFPve_pos  1.3450890702
## dCA2_Fpos_TATATCCCAACCACGC-1dCA2_FI_GFPve_pos  1.1370992583
## dCA2_Fpos_TATTGGGTCGCTCATC-1dCA2_FI_GFPve_pos  1.0253823919
## dCA2_Fpos_TCCATGCAGCTTTCCC-1dCA2_FI_GFPve_pos  1.1468470020
## dCA2_Fpos_TCCCAGTGTCGGAAAC-1dCA2_FI_GFPve_pos  0.5438555998
## dCA2_Fpos_TCCTGCACACCGGAAA-1dCA2_FI_GFPve_pos  1.4101034326
## dCA2_Fpos_TCTATACAGTCATACC-1dCA2_FI_GFPve_pos  1.4004967136
## dCA2_Fpos_TGACAGTTCTGGGTCG-1dCA2_FI_GFPve_pos  0.8154700917
## dCA2_Fpos_TGTTTGTGTCCTTAAG-1dCA2_FI_GFPve_pos  2.3926782531
## dCA2_Fpos_TTCCAATTCTCCACTG-1dCA2_FI_GFPve_pos  0.8743236584
## dCA2_Fpos_TTCTAACAGGGTGAAA-1dCA2_FI_GFPve_pos  1.2023066205
## dCA2_Fpos_TTGGATGGTCAGGTGA-1dCA2_FI_GFPve_pos  2.5333020610
## FC_Mpos_AATGGCTCATGTGCCG-1FC_MI_GFPve_pos      1.7391086740
## FC_Mpos_ACGATCAGTTGAATCC-1FC_MI_GFPve_pos      2.3636760635
## FC_Mpos_ACGTAGTTCCATAGAC-1FC_MI_GFPve_pos      1.6163681668
## FC_Mpos_ACTCTCGTCACTCACC-1FC_MI_GFPve_pos      1.8883964938
## FC_Mpos_AGAGAATAGGTCTGGA-1FC_MI_GFPve_pos      1.4944714231
## FC_Mpos_ATGTCTTTCATACAGC-1FC_MI_GFPve_pos      1.0905790848
## FC_Mpos_ATTCGTTTCATCAGTG-1FC_MI_GFPve_pos      2.4363083762
## FC_Mpos_CAAGACTAGAGCAGCT-1FC_MI_GFPve_pos      2.1937723083
```

```
## FC_Mpos_CACTGGGAGGACTTCT-1FC_MI_GFPve_pos      0.6523128671
## FC_Mpos_CTACGGGAGGTCCCTG-1FC_MI_GFPve_pos      2.0380310935
## FC_Mpos_CTCCATGTCCTATTTG-1FC_MI_GFPve_pos      1.2306109590
## FC_Mpos_GATCAGTGTTTCTTAC-1FC_MI_GFPve_pos      0.5737259907
## FC_Mpos_GGGACAAGTTCGTGCG-1FC_MI_GFPve_pos      0.5557474715
## FC_Mpos_GGTTCTCTCGCTAATG-1FC_MI_GFPve_pos      2.3387448711
## FC_Mpos_TCATTACCAAGCCCAC-1FC_MI_GFPve_pos      0.8745663685
## FC_Mpos_TCATTTGTCCTTACCG-1FC_MI_GFPve_pos      2.3445553703
## FC_Mpos_TCCATGCCACATACGT-1FC_MI_GFPve_pos      1.5468351764
## FC_Mpos_TCGACGGGTCGCAGTC-1FC_MI_GFPve_pos      1.5972206515
## FC_Mpos_TGCTCGTCAGGTACGA-1FC_MI_GFPve_pos      1.6498579902
## FC_Mpos_TGCTGAACAAGATGGC-1FC_MI_GFPve_pos      1.7271484060
## FC_Mpos_TTACGCCCAGCCGTCA-1FC_MI_GFPve_pos      0.6210827154
## FC_Mpos_TTCTAACTCTAGATCG-1FC_MI_GFPve_pos      0.9975063128
## FC_Fpos_AAAGGGCCACTGCACG-1FC_FI_GFPve_pos      0.9286874456
## FC_Fpos_AACCAACTCTATGTGG-1FC_FI_GFPve_pos      2.5064420623
## FC_Fpos_AACCACAGTAGATTAG-1FC_FI_GFPve_pos      2.4458105487
## FC_Fpos_AATCGTGTCCGGGACT-1FC_FI_GFPve_pos      1.1677814645
## FC_Fpos_ACCCTTGAGAGCAGCT-1FC_FI_GFPve_pos      0.8020567817
## FC_Fpos_AGGTCATAGGACACTG-1FC_FI_GFPve_pos      0.3246623797
## FC_Fpos_ATCTCTAGTAGTGTGG-1FC_FI_GFPve_pos      0.8489615244
## FC_Fpos_ATGATCGGTTCATCTT-1FC_FI_GFPve_pos      1.6957700175
## FC_Fpos_ATTCATCCACCGTCGA-1FC_FI_GFPve_pos      1.5905706805
## FC_Fpos_CACGAATTCGTCTAAG-1FC_FI_GFPve_pos      2.1311571521
## FC_Fpos_CAGATACGTCGGTGAA-1FC_FI_GFPve_pos      0.7820185942
## FC_Fpos_CAGATACTCCGAGCTG-1FC_FI_GFPve_pos      0.9484221858
## FC_Fpos_CAGGCCAAGTCATAGA-1FC_FI_GFPve_pos      1.9967595262
## FC_Fpos_CCACGAGCATCGCTGG-1FC_FI_GFPve_pos      2.4753510875
## FC_Fpos_CCCATTGGTATACCCA-1FC_FI_GFPve_pos      2.3193130416
## FC_Fpos_CGCGTGATCAACGAGG-1FC_FI_GFPve_pos      2.4077854079
## FC_Fpos_CGTAATGTCTTGCGCT-1FC_FI_GFPve_pos      0.5572552306
## FC_Fpos_CGTGCTTTCAACCCGG-1FC_FI_GFPve_pos      0.7813376111
## FC_Fpos_CGTTAGACAGCTTCCT-1FC_FI_GFPve_pos      2.0613949222
## FC_Fpos_CTAACCCCAGGAAGTC-1FC_FI_GFPve_pos      1.5927847547
## FC_Fpos_CTGTATTGTCTACTGA-1FC_FI_GFPve_pos      1.4558237714
## FC_Fpos_GACGTTAAGACGTCGA-1FC_FI_GFPve_pos      1.9409457368
## FC_Fpos_GCAGCCATCTGCCTGT-1FC_FI_GFPve_pos      0.7264400644
## FC_Fpos_GTAGAAAAGGACACTG-1FC_FI_GFPve_pos      2.3399169368
## FC_Fpos_GTCTACCGTTGACTGT-1FC_FI_GFPve_pos      0.8627938551
## FC_Fpos_GTGATGTCAGGTACGA-1FC_FI_GFPve_pos      1.5022550744
## FC_Fpos_GTTGCGGAGTTGGGAC-1FC_FI_GFPve_pos      2.5706496162
## FC_Fpos_TACCTGCGTATCAAGA-1FC_FI_GFPve_pos      2.4909121913
## FC_Fpos_TACTTACTCTCTTCAA-1FC_FI_GFPve_pos      1.9088612718
## FC_Fpos_TCCAGAATCCAAACCA-1FC_FI_GFPve_pos      2.5052905006
## FC_Fpos_TCCGAAACATTGTACG-1FC_FI_GFPve_pos      2.1041884345
## FC_Fpos_TCCTCCCTCTCGCGTT-1FC_FI_GFPve_pos      1.0524730605
## FC_Fpos_TTGACCCGTTCAAAGA-1FC_FI_GFPve_pos      0.6632493061
## FC_Fpos_TTGTGTTAGTACGAGC-1FC_FI_GFPve_pos      2.4304885787
## dCA2_Mpos_AAATGGACACGCCAGT-1dCA2_MI_GFPve_pos -1.3912718373
## dCA2_Mpos_AATGGCTTCCATCTAT-1dCA2_MI_GFPve_pos -0.7465659457
## dCA2_Mpos_ACTCCCATCCCATTTA-1dCA2_MI_GFPve_pos -1.3009190636
## dCA2_Mpos_ACTTCGCAGTTGGAAT-1dCA2_MI_GFPve_pos -0.7686762291
## dCA2_Mpos_CAACGATGTCAGACTT-1dCA2_MI_GFPve_pos -0.9473844844
## dCA2_Mpos_CACCGTTAGGTCCCTG-1dCA2_MI_GFPve_pos -0.8542804795
## dCA2_Mpos_CATCCGTGTCAGTTTG-1dCA2_MI_GFPve_pos -1.8290441113
## dCA2_Mpos_CATCGGGCAGCAGATG-1dCA2_MI_GFPve_pos -0.7740945893
## dCA2_Mpos_CCCTAACTCGACTCCT-1dCA2_MI_GFPve_pos -0.7471647340
## dCA2_Mpos_CCTGTTGGTTGCTCCT-1dCA2_MI_GFPve_pos -1.3430380898
## dCA2_Mpos_CGGAATTAGCTTAGTC-1dCA2_MI_GFPve_pos -1.1933779794
## dCA2_Mpos_CGTGCTTCAGACCATT-1dCA2_MI_GFPve_pos -0.7913349348
## dCA2_Mpos_GGTGAAGTCTATCGGA-1dCA2_MI_GFPve_pos -0.0365717339
## dCA2_Mpos_GTACAGTGTTCCTAAG-1dCA2_MI_GFPve_pos -0.7766184884
## dCA2_Mpos_TACCGAATCGGTCACG-1dCA2_MI_GFPve_pos -0.7941135841
## dCA2_Mpos_TAGGAGGCAAGTGCTT-1dCA2_MI_GFPve_pos -1.2286735850
## dCA2_Mpos_TATCGCCTCCTGCTAC-1dCA2_MI_GFPve_pos -0.7667045670
## dCA2_Mpos_TATCGCCTCGACATTG-1dCA2_MI_GFPve_pos -1.7053048688
## dCA2_Mpos_TATCTTGCACCTCTGT-1dCA2_MI_GFPve_pos -0.7864257770
## dCA2_Mpos_TCTGCCATCTCATTTG-1dCA2_MI_GFPve_pos -0.7783391553
## dCA2_Mpos_TGACCCTAGTAGTCAA-1dCA2_MI_GFPve_pos -1.2310270148
## dCA2_Mpos_TGTACAGGTACAGGTG-1dCA2_MI_GFPve_pos -1.3007968741
```

```
## dCA2_Mpos_TTGGGTACACACACTA-1dCA2_MI_GFPve_pos -1.2623931247
## dCA2_Mpos_TTTACGTGTCACAGAG-1dCA2_MI_GFPve_pos -0.8611208874
## dCA2_Mpos_TTTAGTCGTTATTCTC-1dCA2_MI_GFPve_pos -0.7784794646
## dCA2_Fpos_AAGACAACACTTACAG-1dCA2_FI_GFPve_pos -0.7637521702
## dCA2_Fpos_CCTCTAGAGATCGCTT-1dCA2_FI_GFPve_pos -1.1126123744
## dCA2_Fpos_CGGGCATAGTCATGAA-1dCA2_FI_GFPve_pos -1.3025536614
## dCA2_Fpos_CTGCTCAAGTCAATCC-1dCA2_FI_GFPve_pos -1.0093442279
## dCA2_Fpos_GCCATGGTCTGTCTCG-1dCA2_FI_GFPve_pos -1.7595881301
## dCA2_Fpos_GTTATGGGTTGCCGAC-1dCA2_FI_GFPve_pos -0.8242191153
## dCA2_Fpos_GTTGAACTCTACTATC-1dCA2_FI_GFPve_pos -1.3153083401
## dCA2_Fpos_TCAGTCCGTTGCCGCA-1dCA2_FI_GFPve_pos -0.8905757743
## FC_Mpos_AGGAATAAGTCTTCCC-1FC_MI_GFPve_pos     -1.0053482133
## FC_Mpos_ATCGCCTTCGCTCATC-1FC_MI_GFPve_pos     -1.3090763169
## FC_Mpos_CATTGAGCAAATGGAT-1FC_MI_GFPve_pos     -0.6508786159
## FC_Mpos_CTCAATTAGAACAAGG-1FC_MI_GFPve_pos     -0.8690036016
## FC_Mpos_GATCCCTGTCGCAACC-1FC_MI_GFPve_pos     -1.3086379844
## FC_Mpos_GATGGAGTCAGTGTTG-1FC_MI_GFPve_pos     -0.8983781415
## FC_Mpos_GTAACACCAGTCGCTG-1FC_MI_GFPve_pos     -0.2151419895
## FC_Mpos_TCCCAGTTCACACCGG-1FC_MI_GFPve_pos     -0.7752744036
## FC_Mpos_TGACGCGCACGTCGTG-1FC_MI_GFPve_pos     -1.1410257893
## FC_Mpos_TGTGTGAGTGTAGCAG-1FC_MI_GFPve_pos     -1.1472604352
## FC_Mpos_TTCCAATGTAGAGGAA-1FC_MI_GFPve_pos     -1.3099845725
## FC_Fpos_AAAGGTAAGGATAATC-1FC_FI_GFPve_pos     -1.4084744530
## FC_Fpos_AAGCATCGTTCAAGGG-1FC_FI_GFPve_pos     -0.9875953275
## FC_Fpos_AAGTGAACAAGTGACG-1FC_FI_GFPve_pos     -1.2393214780
## FC_Fpos_AATAGAGCATGGCACC-1FC_FI_GFPve_pos     -0.7713049727
## FC_Fpos_AATCACGGTTAAAGTG-1FC_FI_GFPve_pos     -1.0269832688
## FC_Fpos_ACAAAGAAGGTCACAG-1FC_FI_GFPve_pos     -1.4628329354
## FC_Fpos_ACATTTCGTCCAGGTC-1FC_FI_GFPve_pos     -1.0876892882
## FC_Fpos_ACCCAAATCGTGTGGC-1FC_FI_GFPve_pos     -1.0600929337
## FC_Fpos_ACGGAAGCAGGATCTT-1FC_FI_GFPve_pos     -1.2953859645
## FC_Fpos_ACGTAGTAGCCAGAGT-1FC_FI_GFPve_pos     -1.0093845206
## FC_Fpos_ACTCTCGCAGTTGAAA-1FC_FI_GFPve_pos     -1.2153039056
## FC_Fpos_ACTGATGCAGCCTTCT-1FC_FI_GFPve_pos     -0.9803692179
## FC_Fpos_AGATGAACAGGCATTT-1FC_FI_GFPve_pos     -1.4098793345
## FC_Fpos_AGCCAGCTCGGAATTC-1FC_FI_GFPve_pos     -0.4237598258
## FC_Fpos_AGCGCTGTCCATTGCC-1FC_FI_GFPve_pos     -1.0387333709
## FC_Fpos_AGCGTCGAGACCTTTG-1FC_FI_GFPve_pos     -1.1546679812
## FC_Fpos_ATCACTTAGAGCATAT-1FC_FI_GFPve_pos     -1.1315345841
## FC_Fpos_ATCAGGTAGGAACGTC-1FC_FI_GFPve_pos     -1.3552103120
## FC_Fpos_ATGCGATGTTGCATCA-1FC_FI_GFPve_pos     -0.7088902074
## FC_Fpos_CAATACGAGCAAATGT-1FC_FI_GFPve_pos     -0.9793529588
## FC_Fpos_CAATCGAAGGATGCGT-1FC_FI_GFPve_pos     -1.1741411763
## FC_Fpos_CACTTCGTCGGTCGAC-1FC_FI_GFPve_pos     -0.9576160985
## FC_Fpos_CAGCGTGTCACATTGG-1FC_FI_GFPve_pos     -1.0892453271
## FC_Fpos_CAGGTATGTTCCTAGA-1FC_FI_GFPve_pos     -0.6997405368
## FC_Fpos_CAGTTCCAGGAAGTAG-1FC_FI_GFPve_pos     -0.9049104529
## FC_Fpos_CATAAGCCAGTTCTAG-1FC_FI_GFPve_pos     -0.3819179910
## FC_Fpos_CATCCCAAGCCAGAGT-1FC_FI_GFPve_pos     -1.2027559357
## FC_Fpos_CATCGCTAGTTGGCTT-1FC_FI_GFPve_pos     -1.1965210515
## FC_Fpos_CATCGGGAGGTTAAAC-1FC_FI_GFPve_pos     -1.4210037070
## FC_Fpos_CCAAGCGGTACAAAGT-1FC_FI_GFPve_pos     -1.1693252402
## FC_Fpos_CCACGAGTCGTTCCTG-1FC_FI_GFPve_pos     -0.8018757182
## FC_Fpos_CCTAAGAGTGACCGTC-1FC_FI_GFPve_pos     -1.1714377480
## FC_Fpos_CCTGTTGTCTGTGCTC-1FC_FI_GFPve_pos     -0.2231676358
## FC_Fpos_CGAATTGTCGTAACTG-1FC_FI_GFPve_pos     -0.8481453734
## FC_Fpos_CGAGTGCTCGTGGAAG-1FC_FI_GFPve_pos     -1.3101540881
## FC_Fpos_CGATGCGAGTCCCAGC-1FC_FI_GFPve_pos     -1.2626121121
## FC_Fpos_CGCATAAGTTGTGTAC-1FC_FI_GFPve_pos     -1.0564065057
## FC_Fpos_CGCGTGAAGACGCCAA-1FC_FI_GFPve_pos     -1.2512672025
## FC_Fpos_CGTCAAACAGGTCAGA-1FC_FI_GFPve_pos     -1.1326711255
## FC_Fpos_CGTCCATAGTAAACGT-1FC_FI_GFPve_pos     -0.8333579379
## FC_Fpos_CTACCCAGTTAAGGAT-1FC_FI_GFPve_pos     -1.3795542794
## FC_Fpos_CTACGGGAGAGGATGA-1FC_FI_GFPve_pos     -0.6879375654
## FC_Fpos_CTAGGTAGTAGATTGA-1FC_FI_GFPve_pos     -0.2554900723
## FC_Fpos_CTCAAGATCTCGCTTG-1FC_FI_GFPve_pos     -1.2241333800
## FC_Fpos_CTGAGGCAGGCGCTCT-1FC_FI_GFPve_pos     -1.2294417697
## FC_Fpos_CTGAGGCCAAGATTGA-1FC_FI_GFPve_pos     -1.1750426369
## FC_Fpos_CTGCCTAGTGTAGCAG-1FC_FI_GFPve_pos     -1.3061894255
## FC_Fpos_CTGCGAGAGCCTGTGC-1FC_FI_GFPve_pos     -1.1439712125
```

```
## FC_Fpos_CTTGAGATCGTCGACG-1FC_FI_GFPve_pos     -1.1038135367
## FC_Fpos_CTTTCGGGTTGGCCGT-1FC_FI_GFPve_pos     -1.4686812239
## FC_Fpos_GAAGCCCGTTCTTCAT-1FC_FI_GFPve_pos     -1.1427685099
## FC_Fpos_GAAGTAACAGGCTCTG-1FC_FI_GFPve_pos     -0.7796795445
## FC_Fpos_GACGCTGAGCATCTTG-1FC_FI_GFPve_pos     -1.3082103806
## FC_Fpos_GAGACTTAGCATGTTC-1FC_FI_GFPve_pos     -0.4554139453
## FC_Fpos_GAGGCCTCAGCGACAA-1FC_FI_GFPve_pos     -1.1243481713
## FC_Fpos_GAGGCCTCATGGTACT-1FC_FI_GFPve_pos     -1.4362911063
## FC_Fpos_GAGGGTACAACTCGAT-1FC_FI_GFPve_pos     -0.8338794785
## FC_Fpos_GATGAGGGTCTGTAAC-1FC_FI_GFPve_pos     -1.1655861216
## FC_Fpos_GATGATCTCCGCATAA-1FC_FI_GFPve_pos     -1.1633158999
## FC_Fpos_GATGGAGGTGCCGGTT-1FC_FI_GFPve_pos     -1.4701011258
## FC_Fpos_GATGTTGCAAGTGGTG-1FC_FI_GFPve_pos     -1.0018327313
## FC_Fpos_GATTCGAAGAAAGCGA-1FC_FI_GFPve_pos     -1.2922494489
## FC_Fpos_GCACGTGGTGGCAACA-1FC_FI_GFPve_pos     -1.0206899720
## FC_Fpos_GCGATCGAGCAGTACG-1FC_FI_GFPve_pos     -0.3787221390
## FC_Fpos_GCGATCGAGTGGACGT-1FC_FI_GFPve_pos     -1.1941698867
## FC_Fpos_GCGATCGTCTGAGATC-1FC_FI_GFPve_pos     -1.0970771390
## FC_Fpos_GCGGATCAGCGAGTAC-1FC_FI_GFPve_pos     -1.4303902465
## FC_Fpos_GCGGATCGTATTGAGA-1FC_FI_GFPve_pos     -0.8088253814
## FC_Fpos_GCGTGCACAGGGCTTC-1FC_FI_GFPve_pos     -0.9509024697
## FC_Fpos_GGCACGTAGCGCCGTT-1FC_FI_GFPve_pos     -1.3045876103
## FC_Fpos_GGGCTACGTATCAGCT-1FC_FI_GFPve_pos     -1.3855266648
## FC_Fpos_GGGTTTAAGGGCGAAG-1FC_FI_GFPve_pos     -1.1404517966
## FC_Fpos_GGTTAACCATGATGCT-1FC_FI_GFPve_pos     -1.2265894490
## FC_Fpos_GTAAGTCGTTTCGTTT-1FC_FI_GFPve_pos     -1.2869201976
## FC_Fpos_GTAGAGGAGGTCGAGT-1FC_FI_GFPve_pos     -1.2241821366
## FC_Fpos_GTAGGTTCACATGAAA-1FC_FI_GFPve_pos     -1.0254390317
## FC_Fpos_GTATTGGTCCCAATAG-1FC_FI_GFPve_pos     -0.6954982954
## FC_Fpos_GTCAGCGTCGACATCA-1FC_FI_GFPve_pos      0.5493079943
## FC_Fpos_GTCGCGAAGGGAGGAC-1FC_FI_GFPve_pos     -1.0507116395
## FC_Fpos_GTCGCGATCCTCTAAT-1FC_FI_GFPve_pos     -1.2506337243
## FC_Fpos_GTCGTTCCAGACATCT-1FC_FI_GFPve_pos     -1.2707525569
## FC_Fpos_GTGAGTTGTAATCAAG-1FC_FI_GFPve_pos     -1.3123945075
## FC_Fpos_GTGGAAGGTCGCACGT-1FC_FI_GFPve_pos     -1.2922620850
## FC_Fpos_GTGTTCCCAAACTGCT-1FC_FI_GFPve_pos     -0.4680055457
## FC_Fpos_GTTCATTTCGGTAGAG-1FC_FI_GFPve_pos     -1.0971872884
## FC_Fpos_GTTGTCCCAATCGCAT-1FC_FI_GFPve_pos     -1.2190517264
## FC_Fpos_GTTGTCCGTTTACCAG-1FC_FI_GFPve_pos     -1.0188974219
## FC_Fpos_TAAGTCGCACGCCACA-1FC_FI_GFPve_pos     -0.8307642418
## FC_Fpos_TACAACGTCGTTGTTT-1FC_FI_GFPve_pos     -0.9681176024
## FC_Fpos_TACGTCCCAAGCGGAT-1FC_FI_GFPve_pos     -1.2819181758
## FC_Fpos_TAGACTGCAAGACCTT-1FC_FI_GFPve_pos     -1.4639750796
## FC_Fpos_TATCAGGTCAACGTGT-1FC_FI_GFPve_pos     -1.0907001572
## FC_Fpos_TATTGGGGTGTGTCGC-1FC_FI_GFPve_pos     -0.2468713897
## FC_Fpos_TCAAGCAAGGGAGAAT-1FC_FI_GFPve_pos     -0.6703909117
## FC_Fpos_TCAAGTGGTTCTCCAC-1FC_FI_GFPve_pos     -1.0597045499
## FC_Fpos_TCACTATCACTCAGAT-1FC_FI_GFPve_pos     -0.3092040199
## FC_Fpos_TCATTGTAGCTAGCCC-1FC_FI_GFPve_pos     -0.4370041567
## FC_Fpos_TCATTTGGTAGAGGAA-1FC_FI_GFPve_pos     -1.0991947728
## FC_Fpos_TCCAGAAAGAACTTCC-1FC_FI_GFPve_pos     -0.4391785937
## FC_Fpos_TCGAACACAATGTGGG-1FC_FI_GFPve_pos     -0.5462142306
## FC_Fpos_TCGTGCTAGCATGCGA-1FC_FI_GFPve_pos     -1.3072366792
## FC_Fpos_TCGTGGGCAAGCGCAA-1FC_FI_GFPve_pos     -1.0989884215
## FC_Fpos_TCTTGCGAGGAGTACC-1FC_FI_GFPve_pos     -1.3597649413
## FC_Fpos_TCTTTGAAGGGAGAAT-1FC_FI_GFPve_pos     -1.4344903308
## FC_Fpos_TGACGCGAGCTTGTTG-1FC_FI_GFPve_pos     -1.5721802789
## FC_Fpos_TGAGCATGTGGGATTG-1FC_FI_GFPve_pos     -1.1442596989
## FC_Fpos_TGAGGTTGTATCCCAA-1FC_FI_GFPve_pos     -1.1790465194
## FC_Fpos_TGATTCTGTGACATCT-1FC_FI_GFPve_pos     -0.9595310765
## FC_Fpos_TGCTGAAGTGCCTATA-1FC_FI_GFPve_pos     -1.0336450415
## FC_Fpos_TGGAGGATCTGCGGGT-1FC_FI_GFPve_pos     -1.2870949584
## FC_Fpos_TGGGATTTCTGGAAGG-1FC_FI_GFPve_pos     -0.7718409973
## FC_Fpos_TGTAAGCTCTAGATCG-1FC_FI_GFPve_pos     -0.3618246394
## FC_Fpos_TGTTCTATCGATTCCC-1FC_FI_GFPve_pos     -1.3488189059
## FC_Fpos_TGTTGAGTCATCACAG-1FC_FI_GFPve_pos     -1.1344223100
## FC_Fpos_TGTTTGTGTAATGCTC-1FC_FI_GFPve_pos     -1.3367391902
## FC_Fpos_TTACCGCGTGGCCACT-1FC_FI_GFPve_pos     -1.3154302913
## FC_Fpos_TTACGTTGTTGCATTG-1FC_FI_GFPve_pos     -0.1178673881
## FC_Fpos_TTAGGCAAGTCCCGGT-1FC_FI_GFPve_pos     -1.0298886376
```

```
## FC_Fpos_TTAGGGTGTAGTAAGT-1FC_FI_GFPve_pos     -1.6221237260
## FC_Fpos_TTCACGCGTCTTGGTA-1FC_FI_GFPve_pos     -0.8767808753
## FC_Fpos_TTCCACGAGCTGTCCG-1FC_FI_GFPve_pos     -0.9499802666
## FC_Fpos_TTCTGTATCGGTTCAA-1FC_FI_GFPve_pos     -1.0930690842
## FC_Fpos_TTGATGGGTTGGCCGT-1FC_FI_GFPve_pos     -1.2452637034
## FC_Fpos_TTGCATTAGAGGCGGA-1FC_FI_GFPve_pos     -0.9310166913
## FC_Fpos_TTGCGTCGTTGCCTAA-1FC_FI_GFPve_pos     -1.4548837024
## FC_Fpos_TTGCTGCGTAGTGATA-1FC_FI_GFPve_pos     -1.4732652980
## FC_Fpos_TTGGATGAGAATCCCT-1FC_FI_GFPve_pos     -1.2106865722
## FC_Fpos_TTTAGTCGTATGAGCG-1FC_FI_GFPve_pos     -1.3097840625
## FC_Fpos_TTTATGCAGTCGTTAC-1FC_FI_GFPve_pos     -0.9630615788
## attr(,"scaled:center")
## [1]  0.1673992  4.7414837 -0.1033623
```

```
# Prepare a dataframe for cell plotting
plot.data <- FetchData(object = yourseuratobject, vars = c("UMAP_1", "UMAP_2", "UMAP_3", "seurat_clusters"))

# Make a column of row name identities (these will be your cell/barcode names)
plot.data$label <- paste(rownames(plot.data))

# Plot your data, in this example my Seurat object had 21 clusters (0-20)
fig <- plot_ly(data = plot.data,
        x = ~UMAP_1, y = ~UMAP_2, z = ~UMAP_3,
        color = ~seurat_clusters,
        colors = c("lightseagreen",
                   "gray50",
                   "darkgreen",
                   "red4",
                   "red",
                   "turquoise4",
                   "black",
                   "yellow4",
                   "royalblue1",
                   "lightcyan3",
                   "peachpuff3",
                   "khaki3",
                   "gray20",
                   "orange2",
                   "royalblue4",
                   "yellow3",
                   "gray80",
                   "darkorchid1",
                   "lawngreen",
                   "plum2",
                   "darkmagenta"),
        type = "scatter3d",
        mode = "markers",
        marker = list(size = 5, width=2), # controls size of points
        text=~label, #This is that extra column we made earlier for which we will use for cell ID
        hoverinfo="text") #When you visualize your plotly object, hovering your mouse pointer over a point shows
cell names
```

```
## Error in plot_ly(data = plot.data, x = ~UMAP_1, y = ~UMAP_2, z = ~UMAP_3, : could not find function "plot_ly"
```

```
# xaxis
axx <- list(
  nticks = 4,
  range = c(-10,10) #select range of xaxis
)

# yaxis
axy <- list(
  nticks = 4,
  range = c(-10,10) #select range of yaxis
)

#zaxis
axz <- list(
  nticks = 4,
  range = c(-10,10) #select range of zaxis
)

fig <- fig %>% layout(scene = list(xaxis=axx,yaxis=axy,zaxis=axz))
```

```
## Error in layout(., scene = list(xaxis = axx, yaxis = axy, zaxis = axz)): unused argument (scene = list(xaxis =
axx, yaxis = axy, zaxis = axz))
```

```
fig_cube <- fig %>% layout(scene = list(xaxis=axx,yaxis=axy,zaxis=axz, aspectmode='cube')) # To maintain cubic as
pect
```

```
## Error in layout(., scene = list(xaxis = axx, yaxis = axy, zaxis = axz, : unused argument (scene = list(xaxis =
axx, yaxis = axy, zaxis = axz, aspectmode = "cube"))
```

```
fig
```

```
## Error in eval(expr, envir, enclos): object 'fig' not found
```

```
fig_cube
```

```
## Error in eval(expr, envir, enclos): object 'fig_cube' not found
```

object = dCA2.integrated_clean_7celltype

Cluster tree

```
#cluster tree to go with the violin plot based on cell type
pdf(file="/Users/lees44/R_project/FC_vs_dCA2_comparison_vector_integration_042821/cluster_tree_clean_7celltype.pd
f",width=20,height=8,paper='special')
#https://www.rdocumentation.org/packages/ape/versions/5.2/topics/plot.phylo
dCA2.integrated_clean_7celltype <- BuildClusterTree(dCA2.integrated_clean_7celltype, verbose = FALSE, reorder = F
ALSE)
plot(dCA2.integrated_clean_7celltype@tools$BuildClusterTree, type = "phylogram", edge.color = "black", edge.width
= 4,edge.lty = 1, srt = 0, label.offset = 10, direction = "downwards", tip.color = "black")
dev.off()
```

```
## quartz_off_screen
##                 2
```

Find markers for clean subsets 7 cell types

###########switch back to RNA for gene anysis ################################

see https://satijalab.org/seurat/archive/v3.0/immune_alignment.html (https://satijalab.org/seurat/archive/v3.0/immune_alignment.html)

switched back to RNA from integrated assay for gene analysis 08.13.21

```
DefaultAssay(dCA2.integrated_clean_7celltype) <- "integrated"
```

```
# Find markers for every cluster compared to all remaining cells, report only the positive ones
combined_markers_clean_7celltype <- FindAllMarkers(object = dCA2.integrated_clean_7celltype,
                          only.pos = TRUE,
                          logfc.threshold = 0.25)
```

```
## Calculating cluster Type1
```

```
## Calculating cluster Type2
```

```
## Calculating cluster Type3
```

```
## Calculating cluster Type4
```

```
## Calculating cluster Type5
```

```
## Calculating cluster Type6
```

```
## Calculating cluster Type7
```

```
View(combined_markers_clean_7celltype)

# Save markers to file
write.csv(combined_markers_clean_7celltype,
          file = "/Users/lees44/R_project/FC_vs_dCA2_comparison_vector_integration_042821/combined_all_markers_cl
ean_7celltype.csv",
          quote = FALSE,
          row.names = FALSE)
```

```
# Extract top 50 markers per cluster
top50_comb <- combined_markers_clean_7celltype %>%
        group_by(cluster) %>%
        top_n(n = 50,
              wt = avg_log2FC)

# Visualize top 50 markers per cluster
View(top50_comb)

# Save markers to file
write.csv(combined_markers_clean_7celltype,
          file = "/Users/lees44/R_project/FC_vs_dCA2_comparison_vector_integration_042821/top50_markers_clean_7ce
lltypes.csv",
          quote = FALSE,
          row.names = FALSE)
```

Heatmap for different cell types from combined markers for clean 7 cell types

DoHeatmap generates an expression heatmap for given cells and features. In this case, we are plotting the top 20 markers (or all markers if less than 20) for each cluster. https://satijalab.org/seurat/archive/v3.2/pbmc3k_tutorial.html (https://satijalab.org/seurat/archive/v3.2/pbmc3k_tutorial.html)

###################scale data after subsetting 08.13.21#########################

```
combined_markers_clean_7celltype %>%
    group_by(cluster) %>%
    top_n(n = 20, wt = avg_log2FC) -> top20
DoHeatmap(dCA2.integrated_clean_7celltype, features = top20$gene) + NoLegend()
```

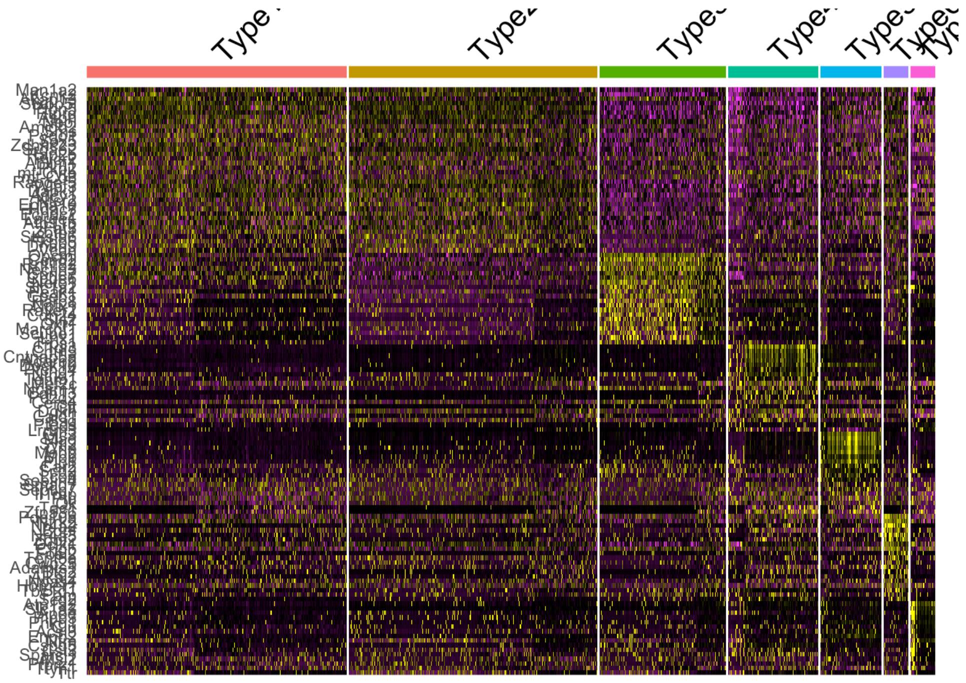

```
ggsave(file="/Users/lees44/R_project/FC_vs_dCA2_comparison_vector_integration_042821/heatmap_top20_celltype_clean
_7celltypes.pdf",width=50,height=25, limitsize = FALSE)
```

Dot plot for Top 10 markers from combined markers subset clean 7 cell types

https://blog.dolomite-bio.com/visualization-of-single-cell-data-from-seurat-object-to-umap/ (https://blog.dolomite-bio.com/visualization-of-single-cell-data-from-seurat-object-to-umap/)

Change font and dot size

https://satijalab.org/seurat/reference/seurattheme (https://satijalab.org/seurat/reference/seurattheme)

```
#subset clean markers 7 cell type
features <- c("Man1a2", "Kcnk2", "Akap13", "Asph", "Ptpn5", "Nebl", "Amigo2", "Sms", "Pygo1", "Sox5", "Stxbp5l",
"Rapgef5", "Dapk1", "Ntsr2", "Epha10", "Hdac7", "St8sia6", "Vcan", "Opcml", "Ccnd2", "Rnf182", "Nectin3", "Neto
1", "Cpeb1", "Parp8", "Nptx1", "Cdh24", "Slit2", "C1ql3", "Adarb2", "Ptchd4", "Ntng1", "Tenm1", "Mef2c", "Cdh13",
"Cit", "Cald1", "Ptgds","Gjc3","Mag", "Mbp", "Plp1", "Car2", "Septin7","Il1rap", "Ptn", "Tac1", "Pde10a", "Ntrk
2", "Nr4a2", "Zdbf2", "Chgb", "Tent5a", "Adamts1", "Scg2", "Atp1a2", "Apoe", "Clu", "Enpp2", "Cspg5", "Slc1a2", "
Cst3", "Ptprz1", "Ttr", "Qk")
```

```
DotPlot(dCA2.integrated_clean_7celltype, features = features) + RotatedAxis() + scale_size(range = c(1, 20))  + F
ontSize(x.text = 30, y.text = 30)
```

```
## Scale for 'size' is already present. Adding another scale for 'size', which
## will replace the existing scale.
```

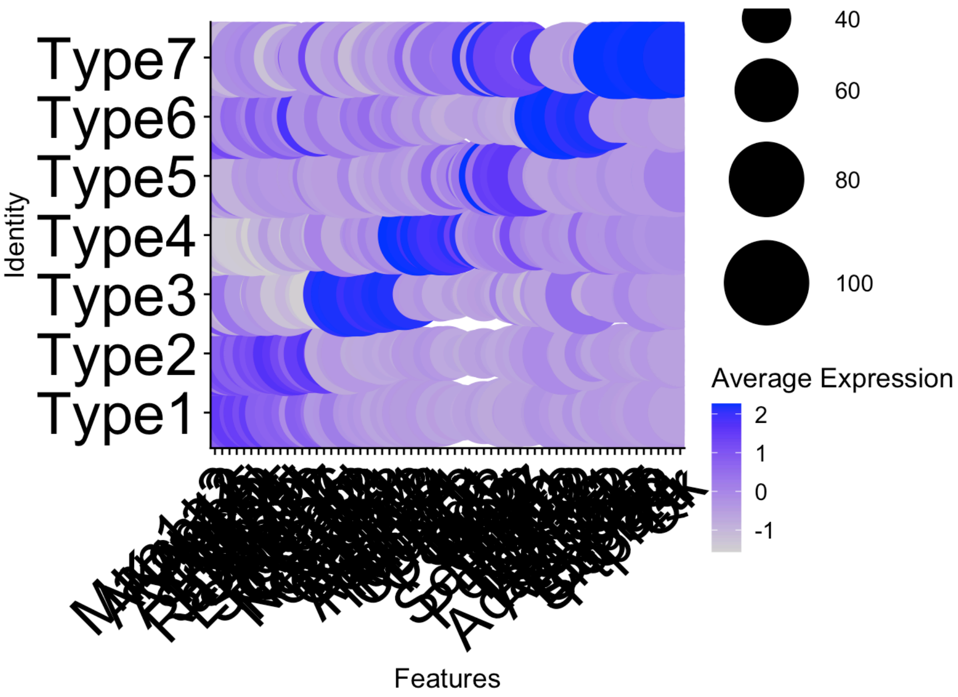


```
ggsave(file="/Users/lees44/R_project/FC_vs_dCA2_comparison_vector_integration_042821/dotplot_top10_celltype_clean
_7celltypes.pdf", width=50, height=25, limitsize = FALSE)
```

Dot plot for Top 10 markers from combined markers subset clean 7 cell types

https://blog.dolomite-bio.com/visualization-of-single-cell-data-from-seurat-object-to-umap/ (https://blog.dolomite-bio.com/visualization-of-single-cell-data-from-seurat-object-to-umap/)

Change font and dot size

https://satijalab.org/seurat/reference/seurattheme (https://satijalab.org/seurat/reference/seurattheme)

```
#subset clean markers 7 cell type
features <- c("Man1a2", "Kcnk2", "Akap13", "Stxbp5l", "Ptpn5", "Ptprd", "Asph", "Nebl", "Amigo2", "Sms", "Pygo1",
"Sox5", "Zdhhc23", "Srgap2", "Rap2b", "Pycr2", "Pkp2", "mt-Cytb", "mt-Co3", "Rapgef5", "Wipf3", "Dapk1", "Adcy1",
"Ntsr2", "Epha10", "Hdac7", "Echdc2", "Large1", "Atp11b", "Fat3", "Zbtb1", "St8sia6", "Fkbp5", "Dnah9","Vcan", "O
pcml", "Ccnd2", "Rnf182", "Nectin3", "Cpne7", "Cpne6",  "Neto1", "Slc1a2", "Peak1", "Cpeb1", "Parp8", "Nptx1", "R
aver2", "Cdh24","Slit2", "Oxr1", "Marchf1", "Scube1", "Lnx1", "Trps1", "C1ql3", "Cntn5", "Cntnap5a", "Adarb2", "D
ock10","Ptchd4", "Ntng1", "Tenm1", "Mef2c", "Ncam1","Cdh13", "Pdzd2", "Cers4","Cit", "Dgkh", "Cald1", "Bsg", "Ptg
ds", "Lrrtm3", "Gjc3","Mag", "Sytl2", "Cnp", "Mobp", "Mbp", "Plp1", "Car2", "Scd2", "Lcorl", "Septin4", "Smad7",
"Septin7","Il1rap", "Ptn", "Qk", "Tac1", "Zfp536", "Pde10a", "Ntrk2", "Nr4a2", "Bdnf", "Nr4a3",  "Zdbf2", "Dclk
1", "Chgb", "Fosl2", "Tent5a", "Cwc25", "Adamts1", "Btg2", "Alkal2", "Npas4", "Homer1", "Tbc1d1", "Pam","Scg2", "
Atp1a2", "Slc1a3", "Apoe", "Plpp3", "Phkg1", "Clu", "Acsl3","Enpp2", "Ntm", "Cspg5", "Cst3", "Sparcl1", "Msi", "P
tprz1", "Ttyh1", "Ttr")
```

```
DotPlot(dCA2.integrated_clean_7celltype, features = features) + RotatedAxis() + scale_size(range = c(1, 10))  + F
ontSize(x.text = 20, y.text = 20)
```

```
## Warning in FetchData(object = object, vars = features, cells = cells): The
## following requested variables were not found: mt-Cytb, mt-Co3, Msi
```

```
## Scale for 'size' is already present. Adding another scale for 'size', which
## will replace the existing scale.
```

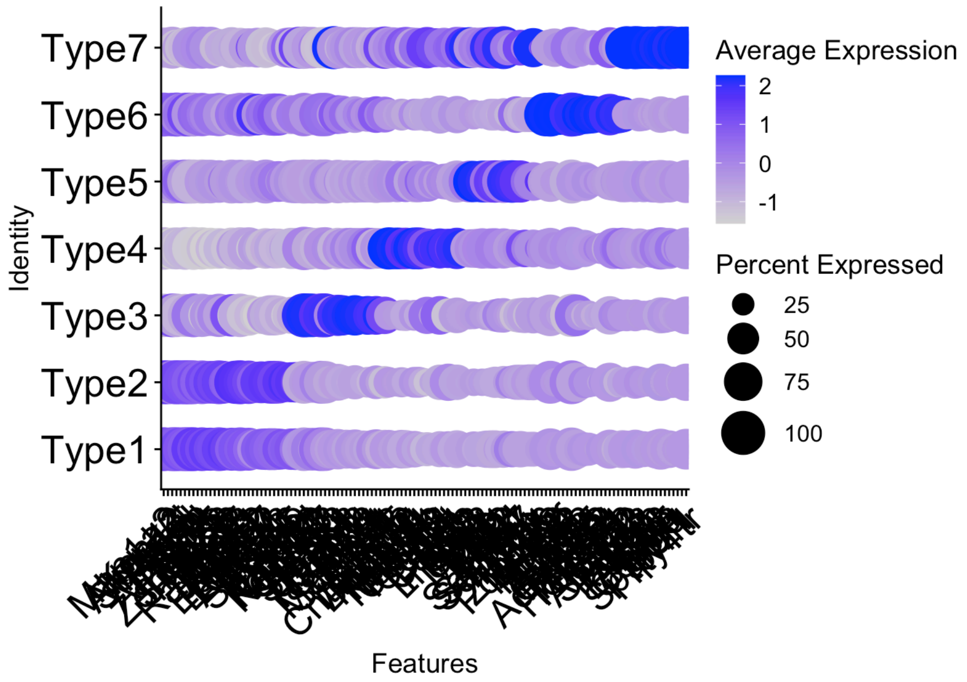


```
ggsave(file="/Users/lees44/R_project/FC_vs_dCA2_comparison_vector_integration_042821/dotplot_top10_celltype_clean_7celltypes_Top20 genes.pdf", width=50, height=25, limitsize = FALSE)
```

###########subset clean markers 7 cell type dotplot as list top#20 markers###################

generating excel table for the 7 cell types aligned to each other

see: https://github.com/satijalab/seurat/issues/1888 (https://github.com/satijalab/seurat/issues/1888)

```
#subset clean markers 7 cell type dotplot as list top#20 markers
 my_genes <- c("Man1a2", "Kcnk2", "Akap13", "Stxbp5l", "Ptpn5", "Ptprd", "Asph", "Nebl", "Amigo2", "Sms", "Pygo1", "Sox5", "Zdhhc23", "Srgap2", "Rap2b", "Pycr2", "Pkp2", "mt-Cytb", "mt-Co3", "Rapgef5", "Wipf3", "Dapk1", "Adcy1", "Ntsr2", "Epha10", "Hdac7", "Echdc2", "Large1", "Atp11b", "Fat3", "Zbtb1", "St8sia6", "Fkbp5", "Dnah9","Vcan", "Opcml", "Ccnd2", "Rnf182", "Nectin3", "Cpne7", "Cpne6",  "Neto1", "Slc1a2", "Peak1", "Cpeb1", "Parp8", "Nptx1", "Raver2", "Cdh24","Slit2", "Oxr1", "Marchf1", "Scube1", "Lnx1", "Trps1", "C1ql3", "Cntn5", "Cntnap5a", "Adarb2", "Dock10","Ptchd4", "Ntng1", "Tenm1", "Mef2c", "Ncam1","Cdh13", "Pdzd2", "Cers4","Cit", "Dgkh", "Cald1", "Bsg", "Ptgds", "Lrrtm3", "Gjc3","Mag", "Sytl2", "Cnp", "Mobp", "Mbp", "Plp1", "Car2", "Scd2", "Lcorl", "Septin4", "Smad7", "Septin7","Il1rap", "Ptn", "Qk", "Tac1", "Zfp536", "Pde10a", "Ntrk2", "Nr4a2", "Bdnf", "Nr4a3",  "Zdbf2", "Dclk1", "Chgb", "Fosl2", "Tent5a", "Cwc25", "Adamts1", "Btg2", "Alkal2", "Npas4", "Homer1", "Tbc1d1", "Pam","Scg2", "Atp1a2", "Slc1a3", "Apoe", "Plpp3", "Phkg1", "Clu", "Acsl3","Enpp2", "Ntm", "Cspg5", "Cst3", "Sparcl1", "Msi", "Ptprz1", "Ttyh1", "Ttr")
a <- DotPlot(dCA2.integrated_clean_7celltype, features = my_genes)
```

```
## Warning in FetchData(object = object, vars = features, cells = cells): The
## following requested variables were not found: mt-Cytb, mt-Co3, Msi
```

```
a$data
```

```
##                avg.exp    pct.exp features.plot    id avg.exp.scaled
## Man1a2    12.8755332891  99.016393        Man1a2 Type1    1.252079205
## Kcnk2      8.3232363505  94.590164         Kcnk2 Type1    1.049185415
## Akap13    12.9073050464  99.344262        Akap13 Type1    0.911432191
## Stxbp5l   22.5737134426  99.508197       Stxbp5l Type1    0.822136096
## Ptpn5      8.5865058452  96.666667         Ptpn5 Type1    1.193270264
## Ptprd     17.6716324146  99.672131         Ptprd Type1    0.984760637
## Asph      14.8376162122  99.398907          Asph Type1    0.800172077
## Nebl      11.8541061362  99.071038          Nebl Type1    1.544543398
## Amigo2     3.0961754525  85.573770        Amigo2 Type1    1.172845558
## Sms       10.0613386336  98.251366           Sms Type1    1.449068775
## Pygo1      3.9648340033  92.349727         Pygo1 Type1    1.167931062
```

```
## Sox5       5.3777527742  94.972678         Sox5 Type1    1.482359550
## Zdhhc23    4.7499467154  90.437158      Zdhhc23 Type1    1.205933185
## Srgap2    10.5856262394  98.633880       Srgap2 Type1    1.094564132
## Rap2b      5.3483032979  93.224044        Rap2b Type1    1.408739764
## Pycr2      1.7651775513  76.885246        Pycr2 Type1    1.551371465
## Pkp2       4.3398657327  94.426230         Pkp2 Type1    1.302390911
## Rapgef5    5.4991974556  89.453552      Rapgef5 Type1    0.704565331
## Wipf3     11.1249843472  97.923497        Wipf3 Type1    0.986788208
## Dapk1      3.4166011456  83.770492        Dapk1 Type1    0.829986456
## Adcy1      5.8792236426  94.153005        Adcy1 Type1    0.981283634
## Ntsr2      3.3176574709  81.256831        Ntsr2 Type1    0.602642513
## Epha10     2.6215177789  80.874317       Epha10 Type1   -0.085229290
## Hdac7      3.6111623467  85.573770        Hdac7 Type1    0.806261916
## Echdc2     2.8215386628  87.923497       Echdc2 Type1    0.926717590
## Large1     4.8231498310  89.562842       Large1 Type1    0.828909712
## Atp11b     4.8692087115  89.508197       Atp11b Type1    0.623560425
## Fat3       4.0580921268  87.103825         Fat3 Type1    0.587571350
## Zbtb1      2.8742780731  82.622951        Zbtb1 Type1    1.059588617
## St8sia6    0.9893329524  74.262295      St8sia6 Type1    0.692775820
## Fkbp5      2.0577977386  87.650273        Fkbp5 Type1    0.849313940
## Dnah9      1.9989624445  88.633880        Dnah9 Type1    0.962194399
## Vcan       1.0332192171  80.983607         Vcan Type1    0.316709767
## Opcml      7.7668480948  97.377049        Opcml Type1   -0.351050440
## Ccnd2      2.2504185878  89.562842        Ccnd2 Type1   -0.347108455
## Rnf182     4.3497645889  93.333333       Rnf182 Type1    0.201502934
## Nectin3    0.9866700362  68.579235      Nectin3 Type1   -0.341382969
## Cpne7      9.4324422181  97.978142        Cpne7 Type1    0.513750284
## Cpne6      6.8259824415  97.103825        Cpne6 Type1    0.045629198
## Neto1      1.0382197965  75.519126        Neto1 Type1   -0.391942721
## Slc1a2     0.5282891607  73.114754       Slc1a2 Type1   -0.561181571
## Peak1      1.3173173875  82.950820        Peak1 Type1   -0.387398265
## Cpeb1      1.1658333370  69.672131        Cpeb1 Type1   -0.320301275
## Parp8      0.4147206879  76.229508        Parp8 Type1   -0.427028424
## Nptx1      0.1147676961  74.098361        Nptx1 Type1   -0.607392618
## Raver2     0.9256527691  78.907104       Raver2 Type1   -0.248200741
## Cdh24      0.3042444215  66.612022        Cdh24 Type1   -0.451627987
## Slit2      0.1455582304  70.163934        Slit2 Type1   -0.574601561
## Oxr1       0.7161478563  66.448087         Oxr1 Type1   -0.457071444
## Marchf1    0.5255629014  72.732240      Marchf1 Type1   -0.625821462
## Scube1     0.6486722410  80.327869       Scube1 Type1   -0.147142713
## Lnx1       0.1646190251  69.071038         Lnx1 Type1   -0.453830113
## Trps1      0.6218077910  75.464481        Trps1 Type1   -0.832076368
## C1ql3      0.0381486566  74.918033        C1ql3 Type1   -0.399984027
## Cntn5      0.0096197588  47.540984        Cntn5 Type1   -0.505892060
## Cntnap5a   0.0401261611  69.945355     Cntnap5a Type1   -0.542494053
## Adarb2     0.0668014962  75.081967       Adarb2 Type1   -0.439162073
## Dock10     0.0274568335  73.060109       Dock10 Type1   -0.745190996
## Ptchd4     0.0555420920  33.224044       Ptchd4 Type1   -0.578910593
## Ntng1      0.4583978519  66.721311        Ntng1 Type1   -0.382216820
## Tenm1      0.3179875376  59.617486        Tenm1 Type1   -0.652020391
## Mef2c      0.6838160902  49.453552        Mef2c Type1   -0.511722087
## Ncam1      0.5095425537  57.704918        Ncam1 Type1   -0.979633884
## Cdh13      0.0218688358  48.087432        Cdh13 Type1   -0.629812786
## Pdzd2      0.0395751063  37.377049        Pdzd2 Type1   -0.776763088
## Cers4      1.3177646419  64.644809        Cers4 Type1   -0.715500429
## Cit        0.0871250435  44.808743          Cit Type1   -0.649303547
## Dgkh       1.5516111421  64.371585         Dgkh Type1   -0.397403477
## Cald1      0.4155922588  45.027322        Cald1 Type1   -0.698115038
## Bsg        0.9723915974  72.021858          Bsg Type1   -0.735345858
## Ptgds      0.0986250959  58.360656        Ptgds Type1   -0.551679909
## Lrrtm3     0.0637831972  31.147541       Lrrtm3 Type1   -0.567899584
## Gjc3       0.0129095900  54.262295         Gjc3 Type1   -0.519944955
## Mag        0.0519853371  63.005464          Mag Type1   -0.470195403
## Sytl2      0.0219688412  51.639344        Sytl2 Type1   -0.706900278
## Cnp        0.1085391957  63.934426          Cnp Type1   -0.636167319
## Mobp       0.0658924638  69.836066         Mobp Type1   -0.664545813
## Mbp        0.1480934133  71.311475          Mbp Type1   -0.649893502
## Plp1       0.3311863239  71.693989         Plp1 Type1   -0.785701316
## Car2       0.4967391657  49.071038         Car2 Type1   -0.472998034
## Scd2       0.5136280507  69.234973         Scd2 Type1   -0.736213314
```

```
## Lcorl      0.3933260552  58.087432        Lcorl Type1   -0.476956004
## Septin4    0.8712442289  65.737705      Septin4 Type1    0.200034239
## Smad7      1.1585974539  63.606557        Smad7 Type1   -0.201614892
## Septin7    1.2770728401  66.284153      Septin7 Type1   -0.539770057
## Il1rap     1.2350252772  60.000000       Il1rap Type1   -0.581282974
## Ptn        1.2821958203  59.945355          Ptn Type1   -0.523314500
## Qk         1.3656704931  60.710383           Qk Type1   -0.546500349
## Tac1       0.1583074601  31.366120         Tac1 Type1   -0.499836654
## Zfp536     0.1598084988  14.699454       Zfp536 Type1   -0.657139838
## Pde10a     2.5511635798  70.983607       Pde10a Type1   -0.437386924
## Ntrk2      5.0529479338  88.743169        Ntrk2 Type1   -0.299990942
## Nr4a2      0.1425437431  61.366120        Nr4a2 Type1   -0.414070381
## Bdnf       0.6992519550  49.016393         Bdnf Type1   -0.386783444
## Nr4a3      0.2416963275  56.612022        Nr4a3 Type1   -0.489591092
## Zdbf2      0.1482404522  35.519126        Zdbf2 Type1   -0.566746763
## Dclk1      8.9388895281  96.721311        Dclk1 Type1   -0.337001273
## Chgb       2.5369625011  78.907104         Chgb Type1   -0.591972235
## Fosl2      0.0933975992  50.601093        Fosl2 Type1   -0.482813557
## Tent5a     0.6180400013  56.284153       Tent5a Type1   -0.313317889
## Cwc25      0.4792346039  54.098361        Cwc25 Type1   -0.365979021
## Adamts1    0.6356544794  45.792350      Adamts1 Type1   -0.366494841
## Btg2       0.1362110251  30.218579         Btg2 Type1   -0.532797958
## Alkal2     0.1630222576  47.650273       Alkal2 Type1   -0.338111217
## Npas4      0.6302917636  49.726776        Npas4 Type1   -0.387237890
## Homer1     2.3015515762  77.049180       Homer1 Type1   -0.423239794
## Tbc1d1     0.2454583509  58.087432       Tbc1d1 Type1   -0.456131407
## Pam        0.4750763904  56.010929          Pam Type1   -0.860745023
## Scg2       0.1537447948  72.677596         Scg2 Type1   -0.620096957
## Atp1a2     0.0719436503  69.617486       Atp1a2 Type1   -0.421113666
## Slc1a3     0.0184928106  66.065574       Slc1a3 Type1   -0.404394663
## Apoe       0.2604232075  76.830601         Apoe Type1   -0.400849857
## Plpp3      0.1886139444  66.120219        Plpp3 Type1   -0.352476304
## Phkg1      0.1539210769  64.262295        Phkg1 Type1   -0.382679868
## Clu        0.2276265077  58.579235          Clu Type1   -0.431752214
## Acsl3      0.3390669782  60.327869        Acsl3 Type1   -0.494622032
## Enpp2      0.1150589870  75.300546        Enpp2 Type1   -0.488584884
## Ntm        1.1981208068  85.355191          Ntm Type1   -0.382890567
## Cspg5      0.1395973014  67.103825        Cspg5 Type1   -0.505506056
## Cst3       0.5901521302  71.147541         Cst3 Type1   -0.419883119
## Sparcl1    0.3591868358  56.612022      Sparcl1 Type1   -0.521198914
## Ptprz1     0.5111851323  45.846995       Ptprz1 Type1   -0.414409587
## Ttyh1      0.7601830547  58.251366        Ttyh1 Type1   -0.514795433
## Ttr        0.9299610805  84.590164          Ttr Type1   -0.398842668
## Man1a21   12.5415829129  99.656357       Man1a2 Type2    1.123691475
## Kcnk21     9.4810256072  98.281787        Kcnk2 Type2    1.501327774
## Akap131   10.3357134872  97.365407       Akap13 Type2    0.178086143
## Stxbp5l1  25.8877930525 100.000000      Stxbp5l Type2    1.275913618
## Ptpn51     8.6983168799  98.167239        Ptpn5 Type2    1.249417504
## Ptprd1    18.0906728824 100.000000        Ptprd Type2    1.092157838
## Asph1     16.3103200822  99.656357         Asph Type2    1.085064263
## Nebl1     10.5557397441  99.140893         Nebl Type2    0.793275106
## Amigo21    3.0587533941  83.619702       Amigo2 Type2    1.125632874
## Sms1       8.9499876283  98.109966          Sms Type2    0.765490671
## Pygo11     4.0169764068  90.435281        Pygo1 Type2    1.219902748
## Sox51      4.8780729231  93.470790         Sox5 Type2    1.036893828
## Zdhhc231   4.9497529035  91.867125      Zdhhc23 Type2    1.412960384
## Srgap21   10.9063485191  98.739977       Srgap2 Type2    1.236786750
## Rap2b1     5.3731154976  93.986254        Rap2b Type2    1.434840984
## Pycr21     1.5236525329  69.702176        Pycr2 Type2    0.854861738
## Pkp21      3.8362561737  89.690722         Pkp2 Type2    0.689456787
## Rapgef51   7.3284562604  96.048110      Rapgef5 Type2    1.721540331
## Wipf31    12.6822755280  99.255441        Wipf3 Type2    1.669335753
## Dapk11     4.1413136665  89.175258        Dapk1 Type2    1.582537389
## Adcy11     6.4994927835  96.678121        Adcy1 Type2    1.452677355
## Ntsr21     3.9190575170  87.457045        Ntsr2 Type2    1.216899389
## Epha101    3.7139588779  84.249714       Epha10 Type2    0.720102989
## Hdac71     4.2227890759  89.690722        Hdac7 Type2    1.478355711
## Echdc21    3.2301487276  85.910653       Echdc2 Type2    1.483667800
## Large11    5.6383152241  93.241695       Large1 Type2    1.596429276
## Atp11b1    5.8113546791  93.757159       Atp11b Type2    1.502576435
```

```
## Fat31       4.7419423651  92.497136          Fat3 Type2    1.242463838
## Zbtb11      3.2490096118  82.416953         Zbtb1 Type2    1.570031053
## St8sia61    1.3576617340  68.671249       St8sia6 Type2    1.766936419
## Fkbp51      2.3928409051  81.557847         Fkbp5 Type2    1.502607591
## Dnah91      2.3534112042  83.104238         Dnah9 Type2    1.626231927
## Vcan1       1.4361173023  70.160367          Vcan Type2    1.222794355
## Opcml1      5.9975853451  94.100802         Opcml Type2   -0.749596352
## Ccnd21      1.9250851452  74.513173         Ccnd2 Type2   -0.468130814
## Rnf1821     3.2541795356  85.452463        Rnf182 Type2   -0.373443120
## Nectin31    0.5921874209  46.563574       Nectin3 Type2   -0.830184020
## Cpne71      7.6735923714  94.100802         Cpne7 Type2    0.054037695
## Cpne61      5.5140411618  92.611684         Cpne6 Type2   -0.499264634
## Neto11      0.7453509959  52.806415         Neto1 Type2   -0.755459966
## Slc1a21     0.3513947160  47.250859        Slc1a2 Type2   -0.598823245
## Peak11      1.2181318029  67.926690         Peak1 Type2   -0.514591769
## Cpeb11      0.8973044289  58.361970         Cpeb1 Type2   -0.696246227
## Parp81      0.3171309055  45.475372         Parp8 Type2   -0.628910408
## Nptx11      0.0807819372  41.008018         Nptx1 Type2   -0.689221224
## Raver21     0.7751192609  57.903780        Raver2 Type2   -0.519320718
## Cdh241      0.2446855413  42.153494         Cdh24 Type2   -0.584128267
## Slit21      0.0779848519  34.421535         Slit2 Type2   -0.738833346
## Oxr11       0.5906850450  45.933562          Oxr1 Type2   -0.714520625
## Marchf11    0.3574472910  46.792669       Marchf1 Type2   -0.998310802
## Scube11     0.5734356569  56.242841        Scube1 Type2   -0.310154167
## Lnx11       0.1252302246  43.069874          Lnx1 Type2   -0.561904680
## Trps11      0.4656097272  51.660939         Trps1 Type2   -1.134162255
## C1ql31      0.0339182783  45.589920         C1ql3 Type2   -0.404677645
## Cntn51      0.0138505843  25.085911         Cntn5 Type2   -0.487898648
## Cntnap5a1   0.0494040156  40.091638      Cntnap5a Type2   -0.510267407
## Adarb21     0.0600183714  44.272623        Adarb2 Type2   -0.460058857
## Dock101     0.0473887193  39.747995        Dock10 Type2   -0.688357002
## Ptchd41     0.0483126277  20.274914        Ptchd4 Type2   -0.599829680
## Ntng11      0.5050691019  46.391753         Ntng1 Type2   -0.314976646
## Tenm11      0.4286490908  46.964490         Tenm1 Type2   -0.412992429
## Mef2c1      0.5434386939  40.950745         Mef2c Type2   -0.692366286
## Ncam11      0.3222612889  36.139748         Ncam1 Type2   -1.209312243
## Cdh131      0.0119281372  27.434135         Cdh13 Type2   -0.660263121
## Pdzd21      0.0435130885  25.658648         Pdzd2 Type2   -0.764252442
## Cers41      1.2563752756  62.428408         Cers4 Type2   -0.810742393
## Cit1        0.0671798360  25.773196           Cit Type2   -0.694895571
## Dgkh1       1.7832341936  69.702176          Dgkh Type2   -0.102093189
## Cald11      0.3352731237  37.457045         Cald1 Type2   -0.803333256
## Bsg1        1.0231019954  62.485682           Bsg Type2   -0.646315750
## Ptgds1      0.1626216812  41.294387         Ptgds Type2   -0.515916176
## Lrrtm31     0.0458124886  21.649485        Lrrtm3 Type2   -0.609055484
## Gjc31       0.0088874516  26.231386          Gjc3 Type2   -0.526092033
## Mag1        0.0589015654  36.941581           Mag Type2   -0.438598993
## Sytl21      0.0433222836  30.870561         Sytl2 Type2   -0.564287720
## Cnp1        0.1480792809  36.597938           Cnp Type2   -0.433675263
## Mobp1       0.0748088271  36.254296          Mobp Type2   -0.617253683
## Mbp1        0.1866507424  41.523482           Mbp Type2   -0.602035784
## Plp11       0.3629151664  48.052692          Plp1 Type2   -0.737099611
## Car21       0.3611981395  35.452463          Car2 Type2   -0.742925217
## Scd21       0.5202333341  52.462772          Scd2 Type2   -0.728267066
## Lcorl1      0.3881225848  42.325315         Lcorl Type2   -0.497730968
## Septin41    0.8337879189  54.581901       Septin4 Type2    0.098667719
## Smad71      1.0620504109  61.168385         Smad7 Type2   -0.429707796
## Septin71    1.2632454965  61.569301       Septin7 Type2   -0.562853950
## Il1rap1     1.1720047264  60.996564        Il1rap Type2   -0.679540205
## Ptn1        1.4083506582  64.032073           Ptn Type2   -0.426746157
## Qk1         1.4069704475  64.604811            Qk Type2   -0.527162195
## Tac11       0.0608484568  20.045819          Tac1 Type2   -0.856995707
## Zfp5361     0.0821604384  11.282932        Zfp536 Type2   -0.874643126
## Pde10a1     2.6305921815  73.711340        Pde10a Type2   -0.415543416
## Ntrk21      5.9227453474  94.444444         Ntrk2 Type2   -0.081951507
## Nr4a21      0.1419866000  39.805269         Nr4a2 Type2   -0.414482401
## Bdnf1       0.6331755767  43.585338          Bdnf Type2   -0.440965915
## Nr4a31      0.2157771016  39.175258         Nr4a3 Type2   -0.530976198
## Zdbf21      0.1602532109  28.350515         Zdbf2 Type2   -0.544601214
## Dclk11     10.7105111285  98.797251         Dclk1 Type2    0.192543349
```

```
## Chgb1       2.5409438297  78.407789          Chgb Type2   -0.589991456
## Fosl21      0.0631938195  32.302405         Fosl2 Type2   -0.578855396
## Tent5a1     0.8234781610  49.369989        Tent5a Type2   -0.048786800
## Cwc251      0.4951195968  44.845361         Cwc25 Type2   -0.337881013
## Adamts11    0.6650689620  40.148912       Adamts1 Type2   -0.323459281
## Btg21       0.0961330011  20.045819          Btg2 Type2   -0.639205070
## Alkal21     0.2017296023  31.271478        Alkal2 Type2   -0.251012303
## Npas41      0.8085736364  51.260023         Npas4 Type2   -0.101200930
## Homer11     2.2993479318  77.262314        Homer1 Type2   -0.425097926
## Tbc1d11     0.2888840163  41.179840        Tbc1d1 Type2   -0.359475068
## Pam1        0.4426733520  44.387171           Pam Type2   -0.933338073
## Scg21       0.1584564046  45.532646          Scg2 Type2   -0.611520272
## Atp1a21     0.0924258917  39.747995        Atp1a2 Type2   -0.411443464
## Slc1a31     0.0202768787  39.747995        Slc1a3 Type2   -0.402137616
## Apoe1       0.3084716809  53.436426          Apoe Type2   -0.388893800
## Plpp31      0.2858480430  47.021764         Plpp3 Type2   -0.237015620
## Phkg11      0.1674966210  40.950745         Phkg1 Type2   -0.365594615
## Clu1        0.2472171769  39.919817           Clu Type2   -0.414112880
## Acsl31      0.3339923769  41.008018         Acsl3 Type2   -0.499979310
## Enpp21      0.1370330607  48.109966         Enpp2 Type2   -0.465651672
## Ntm1        1.2389114423  69.988545           Ntm Type2   -0.359897137
## Cspg51      0.1226642747  43.585338         Cspg5 Type2   -0.516846297
## Cst31       0.5950463650  49.541810          Cst3 Type2   -0.418512931
## Sparcl11    0.4735014745  47.537228       Sparcl1 Type2   -0.411377152
## Ptprz11     0.5670549258  43.814433        Ptprz1 Type2   -0.380983799
## Ttyh11      0.7277593633  53.150057         Ttyh1 Type2   -0.550800148
## Ttr1        1.0790125385  69.759450           Ttr Type2   -0.373996771
## Man1a22     6.9902001674  96.501129        Man1a2 Type3   -1.010546065
## Kcnk22      2.6057338450  67.155756         Kcnk2 Type3   -1.183626002
## Akap132    13.3993560619  98.081264        Akap13 Type3    1.051751390
## Stxbp5l2   12.4948419612  96.952596       Stxbp5l Type3   -0.557904780
## Ptpn52      4.1756059504  85.553047         Ptpn5 Type3   -1.021715527
## Ptprd2      9.9651313717  94.469526         Ptprd Type3   -0.990362790
## Asph2       6.7959532207  92.889391          Asph Type3   -0.755474650
## Nebl2       9.8531062345  98.645598          Nebl Type3    0.386713229
## Amigo22     1.6308254365  61.963883        Amigo2 Type3   -0.675879781
## Sms2        7.1698753516  95.823928           Sms Type3   -0.329434128
## Pygo12      1.4867640938  63.769752         Pygo1 Type3   -1.302025476
## Sox52       3.3593999857  86.907449          Sox5 Type3   -0.317006544
## Zdhhc232    2.9692522723  80.022573       Zdhhc23 Type3   -0.639115688
## Srgap22     8.0677843825  96.049661        Srgap2 Type3   -0.021959673
## Rap2b2      3.4772198330  83.521445         Rap2b Type3   -0.559548527
## Pycr22      1.2453109939  63.205418         Pycr2 Type3    0.052180549
## Pkp22       4.1530293122  92.663657          Pkp2 Type3    1.074995669
## Rapgef52    1.9512661392  68.961625       Rapgef5 Type3   -1.267904307
## Wipf32      7.2478566481  96.726862         Wipf3 Type3   -0.712524049
## Dapk12      1.4374901622  62.979684         Dapk1 Type3   -1.225148199
## Adcy12      2.9236206663  84.085779         Adcy1 Type3   -1.264923042
## Ntsr22      1.7446592980  69.187359         Ntsr2 Type3   -1.003983474
## Epha102     1.0157308628  59.029345        Epha10 Type3   -1.268992916
## Hdac72      1.5147637775  69.751693         Hdac7 Type3   -1.497392264
## Echdc22     1.8279213659  69.751693        Echdc2 Type3   -0.427618511
## Large12     3.0899384346  82.167043        Large1 Type3   -0.802996730
## Atp11b2     3.4342084900  82.054176        Atp11b Type3   -0.715285372
## Fat32       2.4139920704  77.200903          Fat3 Type3   -0.986909022
## Zbtb12      1.5398097955  62.076749         Zbtb1 Type3   -0.758164008
## St8sia62    0.2911477314  48.645598       St8sia6 Type3   -1.343348135
## Fkbp52      1.0583674134  68.397291         Fkbp5 Type3   -1.099454312
## Dnah92      1.0766322395  69.074492         Dnah9 Type3   -0.765733413
## Vcan2       0.3545224761  52.031603          Vcan Type3   -1.209623267
## Opcml2     18.6918576538  99.887133         Opcml Type3    2.109928241
## Ccnd22      9.1378499945  97.855530         Ccnd2 Type3    2.214980421
## Rnf1822     8.1061249107  97.629797        Rnf182 Type3    2.172782605
## Nectin32    3.0174188646  83.521445       Nectin3 Type3    2.174905770
## Cpne72     13.8253519137  98.645598         Cpne7 Type3    1.661930100
## Cpne62     11.3549740232  99.209932         Cpne6 Type3    1.926673432
## Neto12      2.9646569129  83.634312         Neto1 Type3    1.999206869
## Slc1a22     2.9734420163  79.796840        Slc1a2 Type3   -0.040873289
## Peak12      3.0734685313  86.455982         Peak1 Type3    1.864652957
## Cpeb12      2.9168229732  82.731377         Cpeb1 Type3    2.131113432
```

```
## Parp82      1.6268427156  75.282167          Parp8 Type3    2.080463587
## Nptx12      1.2740106753  71.331828          Nptx1 Type3    2.183754038
## Raver22     2.2967971704  82.844244         Raver2 Type3    2.221313477
## Cdh242      1.5006927058  72.799097          Cdh24 Type3    2.210103259
## Slit22      1.2649051162  67.607223          Slit2 Type3    2.145883054
## Oxr12       1.9900747336  75.959368           Oxr1 Type3    2.157021348
## Marchf12    1.6674764728  70.654628        Marchf1 Type3    1.904286427
## Scube12     1.5666666774  75.733634         Scube1 Type3    1.841831264
## Lnx12       1.1363186185  61.399549           Lnx1 Type3    2.212308778
## Trps12      1.6965769137  70.541761          Trps1 Type3    1.246519077
## C1ql32      0.0429927908  42.663657          C1ql3 Type3   -0.394609445
## Cntn52      0.0146218957  21.218962          Cntn5 Type3   -0.484618313
## Cntnap5a2   0.0570785302  37.471783       Cntnap5a Type3   -0.483609965
## Adarb22     0.0559685202  46.049661         Adarb2 Type3   -0.472535241
## Dock102     0.0320252986  47.855530         Dock10 Type3   -0.732164425
## Ptchd42     0.0295441598  17.607223         Ptchd4 Type3   -0.654137882
## Ntng12      0.5146156637  46.388262          Ntng1 Type3   -0.301222730
## Tenm12      0.2868334916  42.776524          Tenm1 Type3   -0.719312851
## Mef2c2      0.5348567284  40.067720          Mef2c Type3   -0.703409961
## Ncam12      1.5763215508  69.638826          Ncam1 Type3    0.328644600
## Cdh132      0.0728009958  31.151242          Cdh13 Type3   -0.473797461
## Pdzd22      0.0789890321  30.022573          Pdzd2 Type3   -0.651548286
## Cers42      2.2733000818  75.282167          Cers4 Type3    0.766956192
## Cit2        0.1389356297  29.345372            Cit Type3   -0.530871613
## Dgkh2       0.8889354545  57.449210           Dgkh Type3   -1.242288935
## Cald12      0.2928665448  38.713318          Cald1 Type3   -0.858885954
## Bsg2        1.1148901589  63.431151            Bsg Type3   -0.485167139
## Ptgds2      0.1536576594  45.936795          Ptgds Type3   -0.520925613
## Lrrtm32     0.0369989152  10.496614         Lrrtm3 Type3   -0.629240027
## Gjc32       0.0228725033  38.261851           Gjc3 Type3   -0.504718525
## Mag2        0.0490263510  37.246050            Mag Type3   -0.483713368
## Sytl22      0.0890129790  32.844244          Sytl2 Type3   -0.259134721
## Cnp2        0.1358379835  32.167043            Cnp Type3   -0.496365201
## Mobp2       0.1085118555  42.776524           Mobp Type3   -0.438493806
## Mbp2        0.2316020785  42.212190            Mbp Type3   -0.546241764
## Plp12       0.3977770009  51.805869           Plp1 Type3   -0.683698842
## Car22       0.7823656789  56.884876           Car2 Type3    0.095821328
## Scd22       0.8720587926  59.142212           Scd2 Type3   -0.305016099
## Lcorl2      0.4062040209  40.970655          Lcorl Type3   -0.425540459
## Septin42    0.5771112434  46.275395        Septin4 Type3   -0.595966144
## Smad72      0.7394637884  55.417607          Smad7 Type3   -1.191820410
## Septin72    1.1099734952  59.819413        Septin7 Type3   -0.818732059
## Il1rap2     1.2406071972  62.528217         Il1rap Type3   -0.572580035
## Ptn2        0.5934798165  51.128668            Ptn Type3   -1.050509210
## Qk2         1.4995814978  65.237020             Qk Type3   -0.483798304
## Tac12       0.0469939967  23.250564           Tac1 Type3   -0.907768296
## Zfp5362     0.0823362132   8.803612         Zfp536 Type3   -0.874150756
## Pde10a2     2.8866523056  75.733634         Pde10a Type3   -0.345124809
## Ntrk22      4.0461555963  88.148984          Ntrk2 Type3   -0.552372046
## Nr4a22      0.1689218429  44.018059          Nr4a2 Type3   -0.394563143
## Bdnf2       0.4779660444  30.812641           Bdnf Type3   -0.568237357
## Nr4a32      0.3304835608  37.133183          Nr4a3 Type3   -0.347824933
## Zdbf22      0.2624539447  37.358916          Zdbf2 Type3   -0.356193915
## Dclk12      6.1024869001  95.259594          Dclk1 Type3   -1.184813106
## Chgb2       4.6373405947  89.729120           Chgb Type3    0.453001900
## Fosl22      0.1458485289  44.356659          Fosl2 Type3   -0.316030333
## Tent5a2     0.1970157826  33.634312         Tent5a Type3   -0.855446944
## Cwc252      0.4172523690  43.340858          Cwc25 Type3   -0.475615663
## Adamts12    0.3038182657  28.103837        Adamts1 Type3   -0.851995724
## Btg22       0.0864535242  30.135440           Btg2 Type3   -0.664904070
## Alkal22     0.0789284849  31.715576         Alkal2 Type3   -0.527338260
## Npas42      0.4390300205  41.196388          Npas4 Type3   -0.694099865
## Homer12     1.9342437534  74.266366         Homer1 Type3   -0.732956953
## Tbc1d12     0.4477486772  47.629797         Tbc1d1 Type3   -0.005875959
## Pam2        0.7963389463  52.257336            Pam Type3   -0.141015338
## Scg22       0.2588596765  45.033860           Scg2 Type3   -0.428753176
## Atp1a22     0.1184283973  49.548533         Atp1a2 Type3   -0.399167001
## Slc1a32     0.0435900369  44.582393         Slc1a3 Type3   -0.372643847
## Apoe2       0.2275017224  49.661400           Apoe Type3   -0.409041816
## Plpp32      0.0807506075  33.069977          Plpp3 Type3   -0.480558682
```

```
## Phkg12     0.1450928157  42.099323         Phkg1 Type3   -0.393790514
## Clu2       0.2471547480  31.602709           Clu Type3   -0.414169090
## Acsl32     0.4064527752  47.291196         Acsl3 Type3   -0.423482562
## Enpp22     0.1908461634  46.613995         Enpp2 Type3   -0.409489700
## Ntm2       0.7606745518  65.011287           Ntm Type3   -0.629476332
## Cspg52     0.4989755030  60.835214         Cspg5 Type3   -0.264826374
## Cst32      0.6336497622  54.176072          Cst3 Type3   -0.407705536
## Sparcl12   0.3120951107  42.550790       Sparcl1 Type3   -0.566439806
## Ptprz12    0.3928170719  43.792325        Ptprz1 Type3   -0.485226849
## Ttyh12     0.8610702581  57.674944         Ttyh1 Type3   -0.402765777
## Ttr2       1.0213642215  71.444695           Ttr Type3   -0.383606366
## Man1a23    6.9420980782  92.598425        Man1a2 Type4   -1.029038986
## Kcnk23     3.4666625368  83.779528         Kcnk2 Type4   -0.847414271
## Akap133    4.2224374796  88.031496        Akap13 Type4   -1.565249337
## Stxbp5l3   9.3986777964  90.078740       Stxbp5l Type4   -0.981844417
## Ptpn53     3.7403958628  80.787402         Ptpn5 Type4   -1.240261444
## Ptprd3    10.0249668647  93.228346         Ptprd Type4   -0.975027363
## Asph3      5.8707549386  89.133858          Asph Type4   -0.934452765
## Nebl3      6.8131396873  94.015748          Nebl Type4   -1.372289845
## Amigo23    0.9675128565  46.929134        Amigo2 Type4   -1.512732929
## Sms3       5.3570555285  91.338583           Sms Type4   -1.444476929
## Pygo13     1.6843273335  77.637795         Pygo1 Type4   -1.105109072
## Sox53      1.9992644940  86.614173          Sox5 Type4   -1.529570424
## Zdhhc233   2.5951575559  73.228346       Zdhhc23 Type4   -1.026730215
## Srgap23    5.0243693097  83.779528        Srgap2 Type4   -1.371546172
## Rap2b3     3.0987860899  78.267717         Rap2b Type4   -0.957642316
## Pycr23     0.7761546376  52.913386         Pycr2 Type4   -1.300772315
## Pkp23      2.2759338379  88.346457          Pkp2 Type4   -1.209583426
## Rapgef53   3.3552501030  76.220472       Rapgef5 Type4   -0.487360522
## Wipf33     6.7373711547  91.496063         Wipf3 Type4   -0.936265520
## Dapk13     1.7764420658  69.921260         Dapk1 Type4   -0.873176122
## Adcy13     3.8144891282  91.338583         Adcy1 Type4   -0.587878560
## Ntsr23     1.4462017862  55.748031         Ntsr2 Type4   -1.308821460
## Epha103    2.0724738415  74.803150        Epha10 Type4   -0.489976792
## Hdac73     2.2915648529  67.874016         Hdac7 Type4   -0.643794541
## Echdc23    1.3553872411  83.149606        Echdc2 Type4   -1.071699513
## Large13    3.0252871854  77.007874        Large1 Type4   -0.863869160
## Atp11b3    3.4942832861  80.629921        Atp11b Type4   -0.659235981
## Fat33      2.3705420817  79.685039          Fat3 Type4   -1.028519114
## Zbtb13     1.4413924114  57.637795         Zbtb1 Type4   -0.892223740
## St8sia63   0.5492452897  61.259843       St8sia6 Type4   -0.590655862
## Fkbp53     1.1219641370  85.826772         Fkbp5 Type4   -0.975448393
## Dnah93     0.9727322055  84.724409         Dnah9 Type4   -0.960383613
## Vcan3      0.4590331969  77.637795          Vcan Type4   -0.974587270
## Opcml3     9.5012234567  96.535433         Opcml Type4    0.039636705
## Ccnd23     3.2929060582  91.181102         Ccnd2 Type4    0.040691495
## Rnf1823    2.5368014448  86.614173        Rnf182 Type4   -0.749912022
## Nectin33   0.9885964192  47.716535       Nectin3 Type4   -0.338995999
## Cpne73     3.9518860463  89.448819         Cpne7 Type4   -0.918708930
## Cpne63     6.1383522961  92.440945         Cpne6 Type4   -0.239966987
## Neto13     1.2809929519  61.574803         Neto1 Type4   -0.090605643
## Slc1a23    2.1975480848  69.606299        Slc1a2 Type4   -0.205977093
## Peak13     0.8923723279  86.299213         Peak1 Type4   -0.932338856
## Cpeb13     1.2265774497  77.322835         Cpeb1 Type4   -0.235258504
## Parp83     0.4346866878  85.511811         Parp8 Type4   -0.385725169
## Nptx13     0.4034726345  85.984252         Nptx1 Type4    0.087731561
## Raver23    0.8399808601  84.251969        Raver2 Type4   -0.402501044
## Cdh243     0.4328561396  83.937008         Cdh24 Type4   -0.165506277
## Slit23     0.3140308308  84.251969         Slit2 Type4   -0.165142149
## Oxr13      1.0353192327  87.401575          Oxr1 Type4    0.197866927
## Marchf13   1.1339001107  85.669291       Marchf1 Type4    0.722055305
## Scube13    0.3294556568  83.779528        Scube1 Type4   -0.838773788
## Lnx13      0.2375267060  82.204724          Lnx1 Type4   -0.253786802
## Trps13     0.7919044670  72.125984         Trps1 Type4   -0.503110669
## C1ql33     2.4412830962  91.181102         C1ql3 Type4    2.266301227
## Cntn53     0.6558257202  87.244094         Cntn5 Type4    2.242377735
## Cntnap5a3  0.8280049499  88.976378      Cntnap5a Type4    2.194204499
## Adarb23    0.9332102562  86.929134        Adarb2 Type4    2.229984834
## Dock103    0.8741063872  87.716535        Dock10 Type4    1.668954673
## Ptchd43    0.9431966747  75.590551        Ptchd4 Type4    1.989595520
```

```
## Ntng13     2.2755585573  89.448819         Ntng1 Type4    2.235801809
## Tenm13     1.5794370036  90.078740         Tenm1 Type4    2.072699137
## Mef2c3     2.5349578363  82.834646         Mef2c Type4    1.870413691
## Ncam13     2.0957156364  80.629921         Ncam1 Type4    0.965620120
## Cdh133     0.8584462348  62.677165         Cdh13 Type4    1.932790000
## Pdzd23     0.7907308558  59.055118         Pdzd2 Type4    1.609596976
## Cers43     3.0252498860  81.259843         Cers4 Type4    1.933561761
## Cit3       1.2930795034  60.787402           Cit Type4    2.107343876
## Dgkh3      3.3136333417  78.425197          Dgkh Type4    1.849105579
## Cald13     2.0425081848  63.307087         Cald1 Type4    1.433147872
## Bsg3       2.5081305750  87.086614           Bsg Type4    1.960886341
## Ptgds3     4.9184004340  68.031496         Ptgds Type4    2.141794395
## Lrrtm33    0.7781311577  36.062992        Lrrtm3 Type4    1.068075599
## Gjc33      0.0730739433  74.015748          Gjc3 Type4   -0.427995114
## Mag3       0.1279639889  74.960630           Mag Type4   -0.123091099
## Sytl23     0.0851601892  77.322835         Sytl2 Type4   -0.284866227
## Cnp3       0.2180604482  82.677165           Cnp Type4   -0.075288821
## Mobp3      0.1933721998  78.582677          Mobp Type4    0.011602977
## Mbp3       0.6411337970  81.889764           Mbp Type4   -0.037927155
## Plp13      0.9300363568  82.834646          Plp1 Type4    0.131607053
## Car23      0.4034718553  47.401575          Car2 Type4   -0.658737976
## Scd23      1.0383407350  82.047244          Scd2 Type4   -0.104976588
## Lcorl3     0.5138850206  42.677165         Lcorl Type4    0.004378173
## Septin43   0.6210966220  73.858268       Septin4 Type4   -0.476930266
## Smad73     1.1397276035  69.291339         Smad7 Type4   -0.246195014
## Septin73   1.5081694087  61.889764       Septin7 Type4   -0.153968655
## Il1rap3    2.3673901338  71.023622        Il1rap Type4    1.184221037
## Ptn3       2.1043231182  62.677165           Ptn Type4    0.106003187
## Qk3        2.1389332916  71.496063            Qk Type4   -0.184430337
## Tac13      0.4730666311  30.866142          Tac1 Type4    0.653664653
## Zfp5363    0.7250763539  46.456693        Zfp536 Type4    0.926256018
## Pde10a3    3.7146335598  76.062992        Pde10a Type4   -0.117423264
## Ntrk23     5.0927420244  85.984252         Ntrk2 Type4   -0.290015422
## Nr4a23     0.4879024646  52.125984         Nr4a2 Type4   -0.158669322
## Bdnf3      0.9832178114  47.244094          Bdnf Type4   -0.153932125
## Nr4a33     0.3624086989  54.015748         Nr4a3 Type4   -0.296850215
## Zdbf23     0.3532266259  28.031496         Zdbf2 Type4   -0.188854259
## Dclk13     9.7278324545  95.905512         Dclk1 Type4   -0.101183148
## Chgb3      3.2035305544  76.850394          Chgb Type4   -0.260343193
## Fosl23     0.1726034004  42.519685         Fosl2 Type4   -0.230955316
## Tent5a3    0.5812213008  69.763780        Tent5a Type4   -0.360727246
## Cwc253     0.4694448321  56.220472         Cwc25 Type4   -0.383295558
## Adamts13   0.7240799606  38.740157       Adamts1 Type4   -0.237121833
## Btg23      0.4008580681  35.275591          Btg2 Type4    0.169839659
## Alkal23    0.2614500087  47.559055        Alkal2 Type4   -0.116629987
## Npas43     0.7454920300  36.692913         Npas4 Type4   -0.202409598
## Homer13    1.9159618631  65.039370        Homer1 Type4   -0.748372402
## Tbc1d13    0.4297437985  85.984252        Tbc1d1 Type4   -0.045951008
## Pam3       0.9928656962  68.661417           Pam Type4    0.299266710
## Scg23      0.7872429579  87.086614          Scg2 Type4    0.533078808
## Atp1a23    0.4716261334  84.409449        Atp1a2 Type4   -0.232413106
## Slc1a33    0.0932218511  70.393701        Slc1a3 Type4   -0.309854017
## Apoe3      0.6834685341  74.645669          Apoe Type4   -0.295582123
## Plpp33     0.1338569874  81.259843         Plpp3 Type4   -0.417497482
## Phkg13     0.0809360392  68.346457         Phkg1 Type4   -0.474533852
## Clu3       0.4752996161  57.165354           Clu Type4   -0.208748665
## Acsl33     0.6642822754  74.803150         Acsl3 Type4   -0.151290871
## Enpp23     0.3168812523  81.102362         Enpp2 Type4   -0.277953350
## Ntm3       2.2877863691  91.338583           Ntm Type4    0.231347211
## Cspg53     0.8131330717  85.826772         Cspg5 Type4   -0.054431466
## Cst33      1.3017968065  84.881890          Cst3 Type4   -0.220651301
## Sparcl13   1.2605493537  70.078740       Sparcl1 Type4    0.344737599
## Ptprz13    0.7658831101  54.488189        Ptprz1 Type4   -0.262028842
## Ttyh13     1.3398012343  66.771654         Ttyh1 Type4    0.128838479
## Ttr3       1.5324772394  88.818898           Ttr Type4   -0.298407191
## Man1a24    8.8705825168  95.093458        Man1a2 Type5   -0.287630211
## Kcnk24     5.3932405642  96.495327         Kcnk2 Type5   -0.095042862
## Akap134    6.8448194427 100.000000        Akap13 Type5   -0.817419284
## Stxbp5l4  14.6614559685  96.028037       Stxbp5l Type5   -0.261243006
## Ptpn54     6.9411701657  92.289720         Ptpn5 Type5    0.367045403
```

```
## Ptprd4    16.5626570405  99.299065        Ptprd Type5    0.700537845
## Asph4      8.3671434220 100.000000         Asph Type5   -0.451530442
## Nebl4      7.5863350685  99.532710         Nebl Type5   -0.924899049
## Amigo24    1.9130120903  61.214953       Amigo2 Type5   -0.319865442
## Sms4       6.2365684220 100.000000          Sms Type5   -0.903499548
## Pygo14     2.4185065070  95.327103        Pygo1 Type5   -0.373333652
## Sox54      3.1523112786  97.897196         Sox5 Type5   -0.501626597
## Zdhhc234   3.2420616576  75.934579      Zdhhc23 Type5   -0.356446951
## Srgap24    7.5604429994  92.757009       Srgap2 Type5   -0.246937556
## Rap2b4     3.5946480114  74.532710        Rap2b Type5   -0.436019827
## Pycr24     1.3220931351  75.934579        Pycr2 Type5    0.273604849
## Pkp24      2.8709734138  99.065421         Pkp2 Type5   -0.485371475
## Rapgef54   3.5004281176  78.971963      Rapgef5 Type5   -0.406648918
## Wipf34     8.2711195859  97.663551        Wipf3 Type5   -0.264036563
## Dapk14     2.2632995911  77.803738        Dapk1 Type5   -0.367616919
## Adcy14     3.7237312437  96.261682        Adcy1 Type5   -0.656852968
## Ntsr24     2.2497447592  62.616822        Ntsr2 Type5   -0.488100213
## Epha104    2.1989037888  84.813084       Epha10 Type5   -0.396774405
## Hdac74     2.4665284032  73.130841        Hdac7 Type5   -0.451533623
## Echdc24    1.9605716434  99.299065       Echdc2 Type5   -0.246811413
## Large14    3.9138226192  78.971963       Large1 Type5   -0.027267981
## Atp11b4    4.0814514929  80.140187       Atp11b Type5   -0.111411894
## Fat34      3.7278441643  85.046729         Fat3 Type5    0.271307791
## Zbtb14     1.9466336188  60.747664        Zbtb1 Type5   -0.204006907
## St8sia64   0.7991685412  83.411215      St8sia6 Type5    0.138197606
## Fkbp54     1.4070335189  93.925234        Fkbp5 Type5   -0.419597578
## Dnah94     1.3761132255  96.728972        Dnah9 Type5   -0.204674562
## Vcan4      0.6908472638  95.093458         Vcan Type5   -0.453256538
## Opcml4     6.6809526259 100.000000        Opcml Type5   -0.595660349
## Ccnd24     1.8077705735  98.364486        Ccnd2 Type5   -0.511771228
## Rnf1824    2.9938768677  99.299065       Rnf182 Type5   -0.510045925
## Nectin34   0.9722135231  44.158879      Nectin3 Type5   -0.359295948
## Cpne74     4.9602744613 100.000000        Cpne7 Type5   -0.655145301
## Cpne64     5.6589920962 100.000000        Cpne6 Type5   -0.439061581
## Neto14     1.3225847986  80.140187        Neto1 Type5   -0.038980638
## Slc1a24    1.2786407204  76.401869       Slc1a2 Type5   -0.401512972
## Peak14     1.1657258241  97.429907        Peak1 Type5   -0.581796092
## Cpeb14     1.2988151135  67.523364        Cpeb1 Type5   -0.134124569
## Parp84     0.2850950508  92.990654        Parp8 Type5   -0.695182324
## Nptx14     0.1481781420  89.485981        Nptx1 Type5   -0.526949211
## Raver24    0.6930640782  92.056075       Raver2 Type5   -0.667107078
## Cdh244     0.3301317238  89.018692        Cdh24 Type5   -0.394036663
## Slit24     0.1578286146  89.252336        Slit2 Type5   -0.544779357
## Oxr14      0.8100551496  88.317757         Oxr1 Type5   -0.264374059
## Marchf14   0.5280152865  89.252336      Marchf1 Type5   -0.620387777
## Scube14    0.4588884266  94.158879       Scube1 Type5   -0.558338089
## Lnx14      0.1595247047  88.785047         Lnx1 Type5   -0.467807855
## Trps14     0.7757429301  88.785047        Trps1 Type5   -0.534366960
## C1ql34     0.1291718277  88.551402        C1ql3 Type5   -0.298993531
## Cntn54     0.0717971209  83.177570        Cntn5 Type5   -0.241455988
## Cntnap5a4  0.1380033484  87.616822     Cntnap5a Type5   -0.202517447
## Adarb24    0.0840615533  71.962617       Adarb2 Type5   -0.385988987
## Dock104    0.3973398987  86.448598       Dock10 Type5    0.309497565
## Ptchd44    0.1639593769  54.672897       Ptchd4 Type5   -0.265195704
## Ntng14     0.2967602959  87.383178        Ntng1 Type5   -0.615091178
## Tenm14     0.5673194909  87.850467        Tenm1 Type5   -0.113465609
## Mef2c4     1.6415102278  65.654206        Mef2c Type5    0.720683521
## Ncam14     1.3611688435  83.411215        Ncam1 Type5    0.064785207
## Cdh134     0.0901625056  58.644860        Cdh13 Type5   -0.420615707
## Pdzd24     0.2750054083  58.644860        Pdzd2 Type5   -0.028820373
## Cers44     1.4472285297  55.140187        Cers4 Type5   -0.514644877
## Cit4       0.3546060457  62.850467          Cit Type5   -0.037878456
## Dgkh4      1.7632155493  65.420561         Dgkh Type5   -0.127616174
## Cald14     1.5517597740  56.308411        Cald1 Type5    0.790266534
## Bsg4       1.0626230781  89.953271          Bsg Type5   -0.576930250
## Ptgds4     0.4337215613  72.196262        Ptgds Type5   -0.364415222
## Lrrtm34    0.1434229092  22.429907       Lrrtm3 Type5   -0.385511449
## Gjc34      0.6016795324  89.018692         Gjc3 Type5    0.379878602
## Mag4       0.6474663673  86.915888          Mag Type5    2.250227025
## Sytl24     0.4543653372  85.280374        Sytl2 Type5    2.180932488
```

```
## Cnp4        0.6605585550  89.252336          Cnp Type5    2.190825454
## Mobp4       0.5846810911  87.850467         Mobp Type5    2.087093887
## Mbp4        1.0261716724  90.887850          Mbp Type5    0.439985467
## Plp14       1.8079129712  91.588785         Plp1 Type5    1.476323630
## Car24       1.4447908022  89.953271         Car2 Type5    1.415027418
## Scd24       1.3714735901  93.691589         Scd2 Type5    0.295786915
## Lcorl4      1.0332570203  87.149533        Lcorl Type5    2.077981689
## Septin44    1.5527023588  92.757009      Septin4 Type5    2.044237231
## Smad74      2.0644270166  89.485981        Smad7 Type5    1.938412275
## Septin74    2.5423160005  90.887850      Septin7 Type5    1.572475002
## Il1rap4     2.6346359070  72.196262       Il1rap Type5    1.600891938
## Ptn4        2.7308163359  67.757009          Ptn Type5    0.585567926
## Qk4         2.7349859575  75.000000           Qk Type5    0.094663389
## Tac14       0.7507584329  31.308411         Tac1 Type5    1.671324774
## Zfp5364     0.7149160513  36.682243       Zfp536 Type5    0.897795562
## Pde10a4     2.4960258176  67.289720       Pde10a Type5   -0.452550255
## Ntrk24      4.1310240351  84.345794        Ntrk2 Type5   -0.531097361
## Nr4a24      0.1956357440  37.383178        Nr4a2 Type5   -0.374807572
## Bdnf4       0.5824700698  30.841121         Bdnf Type5   -0.482544308
## Nr4a34      0.3081761452  58.411215        Nr4a3 Type5   -0.383443079
## Zdbf24      0.2747304050  31.308411        Zdbf2 Type5   -0.333562231
## Dclk14      7.3311055274  99.532710        Dclk1 Type5   -0.817574196
## Chgb4       3.0838243669  76.635514         Chgb Type5   -0.319899075
## Fosl24      0.1291005895  42.056075        Fosl2 Type5   -0.369285353
## Tent5a4     0.5304046122  76.168224       Tent5a Type5   -0.426161019
## Cwc254      0.5780813052  57.710280        Cwc25 Type5   -0.191135043
## Adamts14    0.6352625175  34.813084      Adamts1 Type5   -0.367068310
## Btg24       0.2778962246  33.878505         Btg2 Type5   -0.156623905
## Alkal24     0.1361680702  40.887850       Alkal2 Type5   -0.398538266
## Npas44      0.7404415037  38.084112        Npas4 Type5   -0.210512706
## Homer14     2.2820407834  68.457944       Homer1 Type5   -0.439691464
## Tbc1d14     0.1833540972  75.467290       Tbc1d1 Type5   -0.594362331
## Pam4        0.7335269208  62.616822          Pam Type5   -0.281734131
## Scg24       0.2230254665  87.850467         Scg2 Type5   -0.493983266
## Atp1a24     0.1800548157  86.915888       Atp1a2 Type5   -0.370071556
## Slc1a34     0.0495794032  84.579439       Slc1a3 Type5   -0.365066625
## Apoe4       0.3605528582  87.616822         Apoe Type5   -0.375934273
## Plpp34      0.1269897277  82.943925        Plpp3 Type5   -0.425652014
## Phkg14      0.1972273247  87.383178        Phkg1 Type5   -0.328177581
## Clu4        0.2335612855  70.093458          Clu Type5   -0.426408572
## Acsl34      0.7138836186  89.018692        Acsl3 Type5   -0.098926524
## Enpp24      0.3908142446  89.485981        Enpp2 Type5   -0.200793282
## Ntm4        0.9731485863  95.327103          Ntm Type5   -0.509706025
## Cspg54      0.2560445440  85.514019        Cspg5 Type5   -0.427520009
## Cst34       0.5930805094  89.953271         Cst3 Type5   -0.419063291
## Sparcl14    0.4242240222  65.420561      Sparcl1 Type5   -0.458717866
## Ptprz14     0.8126013029  68.457944       Ptprz1 Type5   -0.234078274
## Ttyh14      0.8822258498  43.457944        Ttyh1 Type5   -0.379273663
## Ttr4        0.8050141591  94.626168          Ttr Type5   -0.419670497
## Man1a25    11.3999770547  99.431818       Man1a2 Type6    0.684799366
## Kcnk25      3.9099589876  78.977273        Kcnk2 Type6   -0.674297185
## Akap135    12.4659148473  97.727273       Akap13 Type6    0.785560041
## Stxbp5l5   23.3310183285 100.000000      Stxbp5l Type6    0.925829421
## Ptpn55      6.2068695840  90.909091        Ptpn5 Type6   -0.001692359
## Ptprd5     15.1195120971  98.863636        Ptprd Type6    0.330669682
## Asph5      17.1449778019  98.863636         Asph Type6    1.246527452
## Nebl5       8.7238521235  98.863636         Nebl Type6   -0.266702334
## Amigo25     2.6412389485  81.818182       Amigo2 Type6    0.598885321
## Sms5        8.5903472506  96.590909          Sms Type6    0.544280372
## Pygo15      3.0762731870  78.977273        Pygo1 Type6    0.282279455
## Sox55       3.7343332589  89.204545         Sox5 Type6    0.017247320
## Zdhhc235    3.9015169413  89.204545      Zdhhc23 Type6    0.326841096
## Srgap25     9.0038417052  96.022727       Srgap2 Type6    0.393130052
## Rap2b5      3.8374601497  84.659091        Rap2b Type6   -0.180593333
## Pycr25      1.0433013769  58.522727        Pycr2 Type6   -0.530374683
## Pkp25       2.8118684646  85.227273         Pkp2 Type6   -0.557307044
## Rapgef55    4.8886968110  89.204545      Rapgef5 Type6    0.365157975
## Wipf35      9.1769717111  97.159091        Wipf3 Type6    0.132990756
## Dapk15      3.0773027311  82.386364        Dapk1 Type6    0.477654558
## Adcy15      5.3258792897  97.159091        Adcy1 Type6    0.560751584
```

```
## Ntsr25      2.6225587225  80.113636         Ntsr2 Type6   -0.107316171
## Epha105     5.2264648520  88.068182        Epha10 Type6    1.835101211
## Hdac75      2.6917898267  80.113636         Hdac7 Type6   -0.204002248
## Echdc25     2.4955805708  80.681818        Echdc2 Type6    0.482424991
## Large15     4.3439933618  89.772727        Large1 Type6    0.377759592
## Atp11b5     4.9793153501  92.045455        Atp11b Type6    0.726289197
## Fat35       4.4443973857  90.340909          Fat3 Type6    0.957518449
## Zbtb15      2.2459413227  73.863636         Zbtb1 Type6    0.203696576
## St8sia65    0.6207157745  49.431818       St8sia6 Type6   -0.382225832
## Fkbp55      1.3609979819  76.704545         Fkbp5 Type6   -0.509361307
## Dnah95      1.6171151586  73.863636         Dnah9 Type6    0.246827449
## Vcan5       0.7944811266  56.818182          Vcan Type6   -0.220192522
## Opcml5     10.3143279436  97.159091         Opcml Type6    0.222797442
## Ccnd25      2.3988097899  73.863636         Ccnd2 Type6   -0.291907695
## Rnf1825     3.3198963262  85.795455        Rnf182 Type6   -0.338955967
## Nectin35    0.9013878697  52.840909       Nectin3 Type6   -0.447055592
## Cpne75      9.5065368990  96.590909         Cpne7 Type6    0.533116496
## Cpne65      7.8251261106  96.590909         Cpne6 Type6    0.460607545
## Neto15      1.6351045664  70.454545         Neto1 Type6    0.348927957
## Slc1a25     1.2102873125  56.818182        Slc1a2 Type6   -0.416058012
## Peak15      2.3114437305  75.568182         Peak1 Type6    0.887448409
## Cpeb15      1.4927017061  72.727273         Cpeb1 Type6    0.137319896
## Parp85      0.8666393019  60.227273         Parp8 Type6    0.507846356
## Nptx15      0.2856802718  47.159091         Nptx1 Type6   -0.195880905
## Raver25     0.8658846703  67.045455        Raver2 Type6   -0.355846711
## Cdh245      0.5107273787  53.409091         Cdh24 Type6    0.007733396
## Slit25      0.4827799580  53.409091         Slit2 Type6    0.244989339
## Oxr15       0.8220579987  58.522727          Oxr1 Type6   -0.239744261
## Marchf15    0.7471366406  55.681818       Marchf1 Type6   -0.134886351
## Scube15     1.1157751755  67.045455        Scube1 Type6    0.864906538
## Lnx15       0.3551886753  51.704545          Lnx1 Type6    0.069052843
## Trps15      1.2794838408  64.204545         Trps1 Type6    0.439864194
## C1ql35      0.0470676701  47.727273         C1ql3 Type6   -0.390088353
## Cntn55      0.0306752907  31.250000         Cntn5 Type6   -0.416344321
## Cntnap5a5   0.0285235471  41.477273      Cntnap5a Type6   -0.582795756
## Adarb25     0.0471757588  50.000000        Adarb2 Type6   -0.499623115
## Dock105     0.0142873062  42.613636        Dock10 Type6   -0.782742728
## Ptchd45     0.0500556425  18.750000        Ptchd4 Type6   -0.594786114
## Ntng15      0.6575564221  49.431818         Ntng1 Type6   -0.095285210
## Tenm15      0.3338065389  48.863636         Tenm1 Type6   -0.617851491
## Mef2c5      0.4224890963  38.636364          Mef2c Type6  -0.848009885
## Ncam15      0.7936502285  54.545455         Ncam1 Type6   -0.631209366
## Cdh135      0.0378778698  26.704545         Cdh13 Type6   -0.580773934
## Pdzd25      0.0980100313  28.409091         Pdzd2 Type6   -0.591120135
## Cers45      1.4995990361  72.727273         Cers4 Type6   -0.433395140
## Cit5        0.1450029682  33.522727           Cit Type6   -0.517002505
## Dgkh5       1.3591373710  57.386364          Dgkh Type6   -0.642799971
## Cald15      0.3526283128  28.409091         Cald1 Type6   -0.780597926
## Bsg5        1.2406622412  69.886364           Bsg Type6   -0.264354395
## Ptgds5      0.0966279989  48.295455         Ptgds Type6   -0.552795963
## Lrrtm35     0.0231875990  17.045455        Lrrtm3 Type6   -0.660870230
## Gjc35      -0.0007356181  28.409091          Gjc3 Type6   -0.540799076
## Mag5        0.0646475694  43.181818           Mag Type6   -0.412348689
## Sytl25      0.0518881185  34.090909         Sytl2 Type6   -0.507079350
## Cnp5        0.1166780212  38.636364           Cnp Type6   -0.594486896
## Mobp5       0.0559789379  39.204545          Mobp Type6   -0.717126864
## Mbp5        0.1312961563  49.431818           Mbp Type6   -0.670742415
## Plp15       0.3873343469  49.431818          Plp1 Type6   -0.699694723
## Car25       0.2407849040  33.522727          Car2 Type6   -0.982725714
## Scd25       0.6836858267  52.840909          Scd2 Type6   -0.531631422
## Lcorl5      0.2696453614  36.363636         Lcorl Type6   -0.970753748
## Septin45    0.4011464042  49.431818       Septin4 Type6   -1.072172779
## Smad75      1.4890571599  67.045455         Smad7 Type6    0.579097924
## Septin75    1.1373940710  57.954545       Septin7 Type6   -0.772955107
## Il1rap5     0.9969342130  60.795455        Il1rap Type6   -0.952497893
## Ptn5        1.1607902547  60.227273           Ptn Type6   -0.616247395
## Qk5         1.3519485735  61.363636            Qk Type6   -0.552925455
## Tac15       0.0964995186  23.295455          Tac1 Type6   -0.726344879
## Zfp5365     0.1233602156  11.931818        Zfp536 Type6   -0.759236676
## Pde10a5    12.3255333788  97.727273        Pde10a Type6    2.250643869
```

```
## Ntrk25    15.1704538427 100.000000         Ntrk2 Type6    2.236249375
## Nr4a25     3.7522030330  86.931818         Nr4a2 Type6    2.255359253
## Bdnf5      3.9065964147  84.659091          Bdnf Type6    2.243231289
## Nr4a35     1.9599342596  78.977273         Nr4a3 Type6    2.253911349
## Zdbf25     1.6738219137  66.477273         Zdbf2 Type6    2.245666331
## Dclk15    16.4611933012 100.000000         Dclk1 Type6    1.911444731
## Chgb5      7.9095590661  93.750000          Chgb Type6    2.080986726
## Fosl25     0.9531843005  64.204545         Fosl2 Type6    2.251132223
## Tent5a5    2.5662893599  76.704545        Tent5a Type6    2.195332530
## Cwc255     1.9616121617  73.863636         Cwc25 Type6    2.256109373
## Adamts15   2.3900303548  70.454545       Adamts1 Type6    2.200286606
## Btg25      1.1529583907  55.681818          Btg2 Type6    2.166665216
## Alkal25    1.3075371380  61.363636        Alkal2 Type6    2.237265755
## Npas45     2.2503164769  72.159091         Npas4 Type6    2.211943604
## Homer15    5.0473722773  88.068182        Homer1 Type6    1.892060163
## Tbc1d15    1.4328398746  67.613636        Tbc1d1 Type6    2.186728538
## Pam5       1.7721947881  70.454545           Pam Type6    2.045210245
## Scg25      1.6362522559  63.636364          Scg2 Type6    2.078555974
## Atp1a25    0.0564518535  43.181818        Atp1a2 Type6   -0.428427749
## Slc1a35    0.0122355159  42.613636        Slc1a3 Type6   -0.412310845
## Apoe5      0.2805700671  43.750000          Apoe Type6   -0.395836649
## Plpp35     0.1925395688  56.250000         Plpp3 Type6   -0.347814819
## Phkg15     0.2036742351  51.704545         Phkg1 Type6   -0.320063940
## Clu5       0.3010284499  42.045455           Clu Type6   -0.365661496
## Acsl35     0.2746898549  41.477273         Acsl3 Type6   -0.562585233
## Enpp25     0.1872190429  48.863636         Enpp2 Type6   -0.413275140
## Ntm5       0.9442613727  66.477273           Ntm Type6   -0.525989569
## Cspg55     0.1962385699  56.250000         Cspg5 Type6   -0.467572752
## Cst35      0.7463369561  55.113636          Cst3 Type6   -0.376157665
## Sparcl15   0.3430980865  46.022727       Sparcl1 Type6   -0.536655331
## Ptprz15    0.4012479134  36.363636        Ptprz1 Type6   -0.480182844
## Ttyh15     0.7841727797  53.977273         Ttyh1 Type6   -0.488156171
## Ttr5       0.9733416374  73.863636           Ttr Type6   -0.391611415
## Man1a26    7.7112046800  82.080925        Man1a2 Type7   -0.733354784
## Kcnk26     6.2764426746  73.988439         Kcnk2 Type7    0.249867132
## Akap136    7.8030414568  90.173410        Akap13 Type7   -0.544161144
## Stxbp5l6   7.6382686801  65.895954       Stxbp5l Type7   -1.222886932
## Ptpn56     5.1228138180  66.473988         Ptpn5 Type7   -0.546063842
## Ptprd6     9.3706049112  81.502890         Ptprd Type7   -1.142735849
## Asph6      5.5820310503  86.127168          Asph Type7   -0.990305935
## Nebl6      8.9071516405  91.329480          Nebl Type7   -0.160640504
## Amigo26    1.8583048351  41.618497        Amigo2 Type7   -0.388885600
## Sms6       7.5730780121  90.751445           Sms Type7   -0.081429214
## Pygo16     2.9037839264  83.815029         Pygo1 Type7    0.110354936
## Sox56      3.5037736064  89.595376          Sox5 Type7   -0.188297134
## Zdhhc236   2.6948433091  45.664740       Zdhhc23 Type7   -0.923441811
## Srgap26    5.6727220679  69.942197        Srgap2 Type7   -1.084037532
## Rap2b6     3.3344107065  67.052023         Rap2b Type7   -0.709776744
## Pycr26     0.9148261062  52.601156         Pycr2 Type7   -0.900871603
## Pkp26      2.6004822313  85.549133          Pkp2 Type7   -0.814581422
## Rapgef56   3.0998502149  58.959538       Rapgef5 Type7   -0.629349890
## Wipf36     6.8742137128  79.768786         Wipf3 Type7   -0.876288585
## Dapk16     2.2087738465  47.976879         Dapk1 Type7   -0.424237163
## Adcy16     3.9497824420  76.300578         Adcy1 Type7   -0.485058003
## Ntsr26     3.7935212779  58.381503         Ntsr2 Type7    1.088679415
## Epha106    2.3108750011  80.346821        Epha10 Type7   -0.314230796
## Hdac76     3.3434702423  52.023121         Hdac7 Type7    0.512105049
## Echdc26    1.3003766372  83.815029        Echdc2 Type7   -1.146680944
## Large16    2.7649766737  71.676301        Large1 Type7   -1.108964710
## Atp11b6    2.7362320536  46.820809        Atp11b Type7   -1.366492811
## Fat36      2.3549684360  50.289017          Fat3 Type7   -1.043433291
## Zbtb16     1.3777448448  42.774566         Zbtb1 Type7   -0.978921590
## St8sia66   0.6551928517  80.346821       St8sia6 Type7   -0.281680016
## Fkbp56     1.9565740854  82.080925         Fkbp5 Type7    0.651940059
## Dnah96     1.0025818434  84.393064         Dnah9 Type7   -0.904462187
## Vcan6      1.4785204212  90.173410          Vcan Type7    1.318155475
## Opcml6     6.3240559838  91.907514         Opcml Type7   -0.676055247
## Ccnd26     1.4717914502  87.283237         Ccnd2 Type7   -0.636753725
## Rnf1826    3.1998993771  87.861272        Rnf182 Type7   -0.401928505
## Nectin36   1.3767871028  65.895954       Nectin3 Type7    0.142008757
```

```
## Cpne76     2.9178338080  85.549133         Cpne7 Type7   -1.188980343
## Cpne66     3.6953787790  87.283237         Cpne6 Type7   -1.254616974
## Neto16     0.4910176737  77.456647         Neto1 Type7   -1.071145858
## Slc1a26   13.6190605062  95.375723        Slc1a2 Type7    2.224426182
## Peak16     1.3574162069  90.173410         Peak1 Type7   -0.335976384
## Cpeb16     0.7642656391  61.849711         Cpeb1 Type7   -0.882502752
## Parp86     0.4029087117  81.502890         Parp8 Type7   -0.451463617
## Nptx16     0.2623551129  85.549133         Nptx1 Type7   -0.252041641
## Raver26    1.0477272572  86.127168        Raver2 Type7   -0.028337185
## Cdh246     0.2274206070  85.549133         Cdh24 Type7   -0.622537461
## Slit26     0.2307638577  83.815029         Slit2 Type7   -0.367515982
## Oxr16      0.6079086354  60.115607          Oxr1 Type7   -0.679177887
## Marchf16   0.6965655721  84.393064       Marchf1 Type7   -0.246935339
## Scube16    0.3231993404  85.549133        Scube1 Type7   -0.852329045
## Lnx16      0.1317440297  84.393064          Lnx1 Type7   -0.544032172
## Trps16     1.7331923107  91.907514         Trps1 Type7    1.317332981
## C1ql36     0.0580096195  86.127168         C1ql3 Type7   -0.377948226
## Cntn56     0.1036075504  84.393064         Cntn5 Type7   -0.106168405
## Cntnap5a6  0.2330076342  76.300578      Cntnap5a Type7    0.127480130
## Adarb26    0.2182428115  82.658960        Adarb2 Type7    0.027383439
## Dock106    0.6289814954  87.861272        Dock10 Type7    0.970002913
## Ptchd46    0.4986512405  83.236994        Ptchd4 Type7    0.703264454
## Ntng16     0.3578977768  80.924855         Ntng1 Type7   -0.527009225
## Tenm16     0.8249174333  78.612717         Tenm1 Type7    0.442943635
## Mef2c6     1.2092345861  42.196532         Mef2c Type7    0.164411008
## Ncam16     2.4997377726  87.283237         Ncam1 Type7    1.461105566
## Cdh136     0.4992410036  38.150289         Cdh13 Type7    0.832473009
## Pdzd26     0.6627169586  76.878613         Pdzd2 Type7    1.202907348
## Cers46     1.6331265499  38.150289         Cers4 Type7   -0.226235113
## Cit6       0.5123084862  80.346821           Cit Type7    0.322607815
## Dgkh6      2.3834012085  46.242775          Dgkh Type7    0.663096168
## Cald16     1.6488979810  39.884393         Cald1 Type7    0.917517767
## Bsg6       1.8168459461  88.439306           Bsg Type7    0.747227052
## Ptgds6     1.7370572706  85.549133         Ptgds Type7    0.363938487
## Lrrtm36    1.0909581218  83.236994        Lrrtm3 Type7    1.784501176
## Gjc36      1.7531418620  85.549133          Gjc3 Type7    2.139671100
## Mag6       0.0843630772  83.236994           Mag Type7   -0.322279473
## Sytl26     0.1489756112  77.456647         Sytl2 Type7    0.141335809
## Cnp6       0.2415797873  83.236994           Cnp Type7    0.045158045
## Mobp6      0.2550468094  84.971098          Mobp Type7    0.338723302
## Mbp6       2.3368850104  87.861272           Mbp Type7    2.066855152
## Plp16      1.6916694776  89.017341          Plp1 Type7    1.298263809
## Car26      1.4103996771  81.502890          Car2 Type7    1.346538195
## Scd26      2.8797940233  94.219653          Scd2 Type7    2.110317574
## Lcorl6     0.5850789246  63.005780         Lcorl Type7    0.288621317
## Septin46   0.7242131919  43.352601       Septin4 Type7   -0.197870000
## Smad76     1.0542348556  40.462428         Smad7 Type7   -0.448172087
## Septin76   2.3646093902  64.161850       Septin7 Type7    1.275804827
## Il1rap6    1.6083559856  37.572254        Il1rap Type7    0.000788131
## Ptn6       4.4809435929  50.289017           Ptn Type7    1.925246148
## Qk6        7.2316217684  78.034682            Qk Type7    2.200153250
## Tac16      0.4764206357  16.184971          Tac1 Type7    0.665956109
## Zfp5366    0.8731812029  27.745665        Zfp536 Type7    1.341118816
## Pde10a6    2.3867019559  44.508671        Pde10a Type7   -0.482615201
## Ntrk26     4.3315808494  67.052023         Ntrk2 Type7   -0.480822097
## Nr4a26     0.0280159467  74.566474         Nr4a2 Type7   -0.498766433
## Bdnf6      0.9139053948  46.820809          Bdnf Type7   -0.210768140
## Nr4a36     0.4197924615  53.179191         Nr4a3 Type7   -0.205225831
## Zdbf26     0.3169621279  15.028902         Zdbf2 Type7   -0.255707948
## Dclk16    11.1924061088  89.017341         Dclk1 Type7    0.336583643
## Chgb6      2.1755469616  42.196532          Chgb Type7   -0.771782666
## Fosl26     0.1593204852  72.832370         Fosl2 Type7   -0.273192268
## Tent5a6    0.7131169901  56.069364        Tent5a Type7   -0.190892633
## Cwc256     0.4022213805  30.057803         Cwc25 Type7   -0.502203074
## Adamts16   0.8491421479  60.693642       Adamts1 Type7   -0.054146618
## Btg26      0.2077078847  34.682081          Btg2 Type7   -0.342973872
## Alkal26    0.0441325652  46.242775        Alkal2 Type7   -0.605635722
## Npas46     0.4874075062  16.184971         Npas4 Type7   -0.616482614
## Homer16    3.8439196029  84.971098        Homer1 Type7    0.877298375
## Tbc1d16    0.1246915408  45.086705        Tbc1d1 Type7   -0.724932765
```

```
## Pam6       0.8023072770  72.254335           Pam Type7   -0.127644390
## Scg26      0.2431878287  84.393064          Scg2 Type7   -0.457281112
## Atp1a26    5.7563356060  94.797688        Atp1a2 Type7    2.262636541
## Slc1a36    2.1296111437  90.751445        Slc1a3 Type7    2.266407614
## Apoe6     10.9783984979  95.375723          Apoe Type7    2.266138517
## Plpp36     2.3895403120  93.063584         Plpp3 Type7    2.261014921
## Phkg16     2.2575790196  90.173410         Phkg1 Type7    2.264840370
## Clu6       3.2180985605  91.329480           Clu Type7    2.260852916
## Acsl36     2.9207641230  91.329480         Acsl3 Type7    2.230886533
## Enpp26     2.7446153193  88.439306         Enpp2 Type7    2.255748029
## Ntm6       5.7387113713  95.375723           Ntm Type7    2.176612419
## Cspg56     4.2342102549  93.063584         Cspg5 Type7    2.236702954
## Cst36     10.1695920452  94.797688          Cst3 Type7    2.261973842
## Sparcl16   3.1393033799  92.485549       Sparcl1 Type7    2.149651469
## Ptprz16    4.9761832805  82.080925        Ptprz1 Type7    2.256910195
## Ttyh16     3.2112265356  90.751445         Ttyh1 Type7    2.206952713
## Ttr6      16.9172605666  91.329480           Ttr Type7    2.266134908
```

```
# Save markers to file
write.csv(a$data,
          file = "/Users/lees44/R_project/FC_vs_dCA2_comparison_vector_integration_042821/combined_all_markers_cl
ean_7celltype_avg_exp.csv",
          quote = TRUE,
          row.names = TRUE)
```

Find markers for clean subsets 7 cell types with min.diff.pct = 0.4 and logfc.threshold = 0.4

```
# Find markers for every cluster compared to all remaining cells, report only the positive ones
combined_markers_clean_7celltype_pctdiff0.5 <- FindAllMarkers(object = dCA2.integrated_clean_7celltype,
                          only.pos = TRUE,
                          logfc.threshold = 0.4,
                          min.diff.pct = 0.4)
```

```
## Calculating cluster Type1
```

```
## Warning in FindMarkers.default(object = data.use, slot = data.slot, counts =
## counts, : No features pass min.diff.pct threshold; returning empty data.frame
```

```
## Calculating cluster Type2
```

```
## Warning in FindMarkers.default(object = data.use, slot = data.slot, counts =
## counts, : No features pass min.diff.pct threshold; returning empty data.frame
```

```
## Calculating cluster Type3
```

```
## Warning in FindMarkers.default(object = data.use, slot = data.slot, counts =
## counts, : No features pass min.diff.pct threshold; returning empty data.frame
```

```
## Calculating cluster Type4
```

```
## Calculating cluster Type5
```

```
## Calculating cluster Type6
```

```
## Calculating cluster Type7
```

```
View(combined_markers_clean_7celltype_pctdiff0.5)

# Save markers to file
write.csv(combined_markers_clean_7celltype_pctdiff0.5,
          file = "/Users/lees44/R_project/FC_vs_dCA2_comparison_vector_integration_042821/combined_all_markers_cl
ean_7celltype_pctdiff0.5.csv",
          quote = FALSE,
          row.names = FALSE)
```

```
combined_markers_clean_7celltype_pctdiff0.5 %>%
    group_by(cluster) %>%
    top_n(n = 10, wt = avg_log2FC) -> top10
DoHeatmap(dCA2.integrated_clean_7celltype, features = top10$gene) + NoLegend()
```

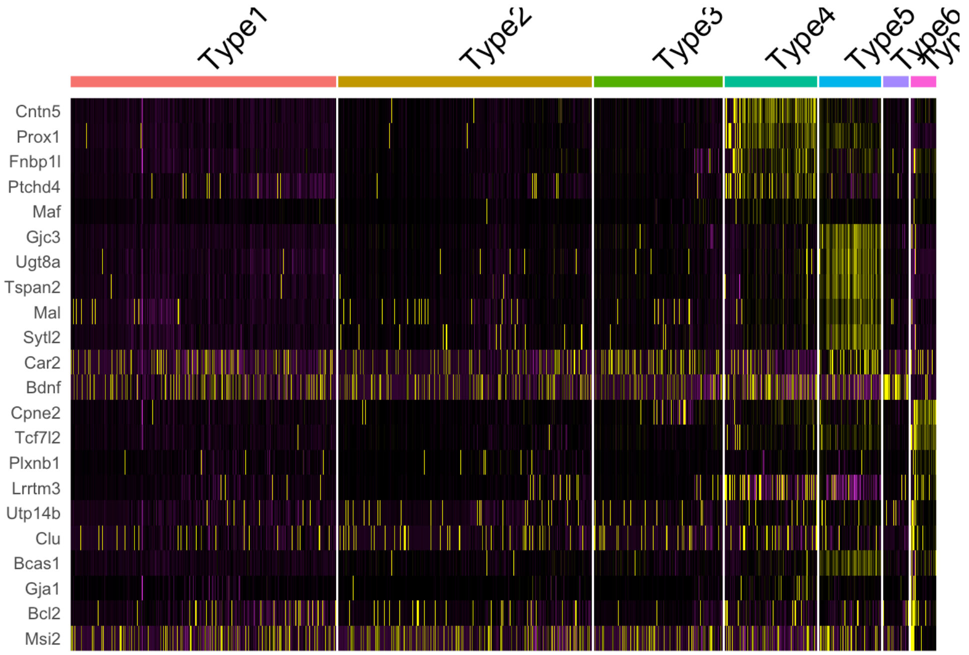


```
ggsave(file="/Users/lees44/R_project/FC_vs_dCA2_comparison_vector_integration_042821/heatmap_top10_celltype_clean
_7celltypes_pctdiff0.5.pdf",width=50,height=25, limitsize = FALSE)
```

```
#subset clean markers 7 cell type
features <- c("Cntn5", "Prox1", "Fnbp1l1", "Ptchd4", "Maf", "Gjc3", "Ugt8a", "Tspan2", "Mal", "Sytl2", "Car2", "B
dnf", "Cpne2", "Tcf7l2", "Plxnb1   ", "Lrrtm3", "Utp14b", "Clu", "Bcas1", "Gja1", "Bcl2", "Msi2")
```

```
DotPlot(dCA2.integrated_clean_7celltype, features = features) + RotatedAxis() + scale_size(range = c(1, 20))  + F
ontSize(x.text = 30, y.text = 30)
```

```
## Warning in FetchData(object = object, vars = features, cells = cells): The
## following requested variables were not found: Fnbp1l1, Plxnb1
```

```
## Scale for 'size' is already present. Adding another scale for 'size', which
## will replace the existing scale.
```

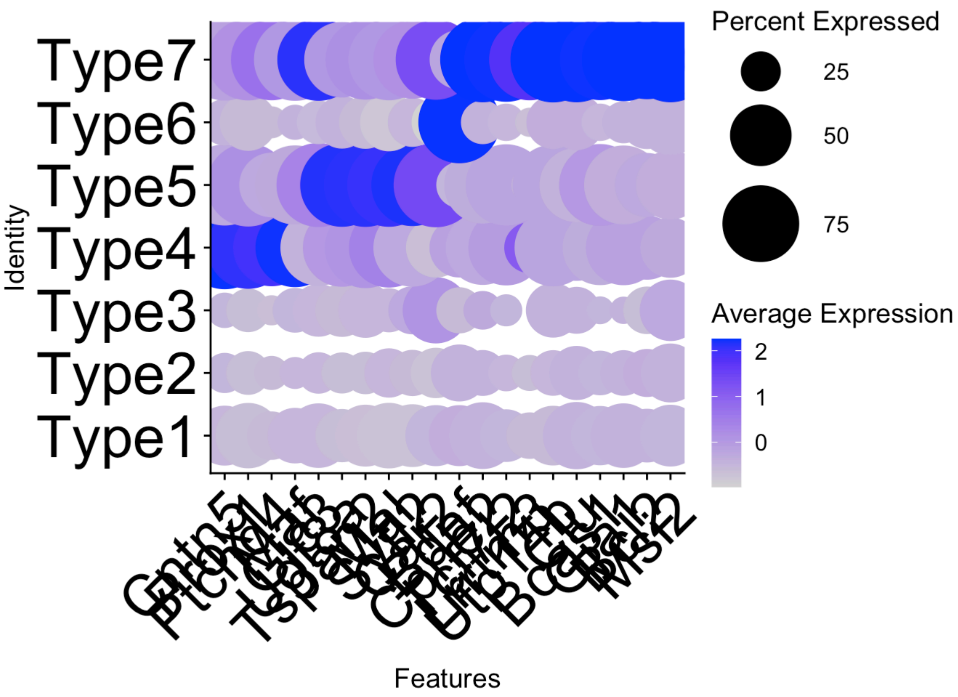


```
ggsave(file="/Users/lees44/R_project/FC_vs_dCA2_comparison_vector_integration_042821/dotplot_top10_celltype_clean_7celltypes_pctdiff0.5.pdf", width=50, height=25, limitsize = FALSE)
```

Cell type marker distribution

```
#Tac1
F<-FeaturePlot(object =dCA2.integrated_clean_7celltype, features =  c("Tac1"))
ggsave(file="/Users/lees44/R_project/FC_vs_dCA2_comparison_vector_integration_042821/clean_7celltype_Tac1.pdf",width=10,height=10)
plot(F)
```

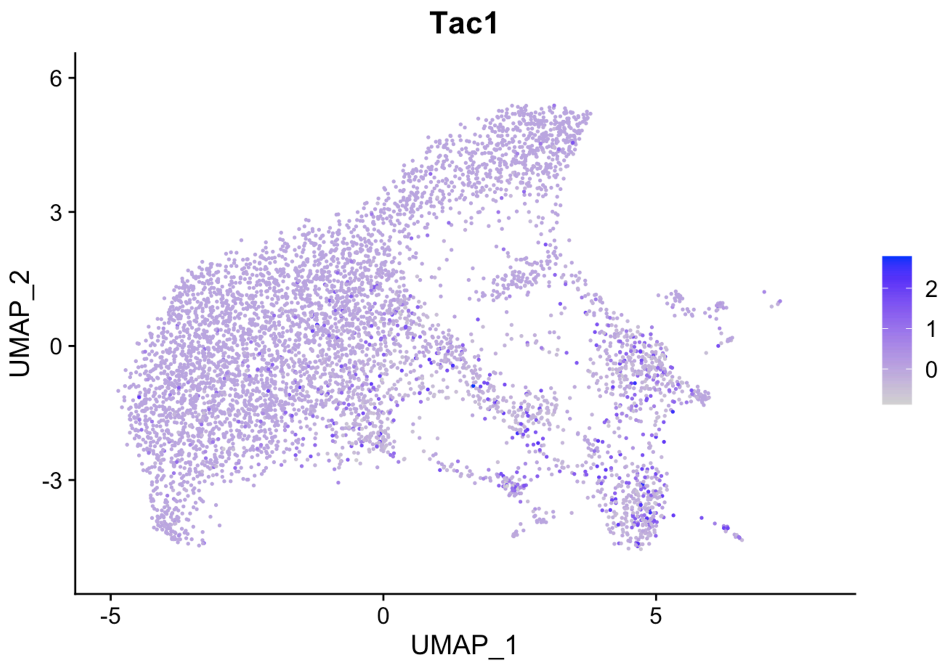

```
#Clu
F<-FeaturePlot(object =dCA2.integrated_clean_7celltype, features =  c("Clu"))
ggsave(file="/Users/lees44/R_project/FC_vs_dCA2_comparison_vector_integration_042821/clean_7celltype_Clu.pdf",wid
th=10,height=10)
plot(F)
```

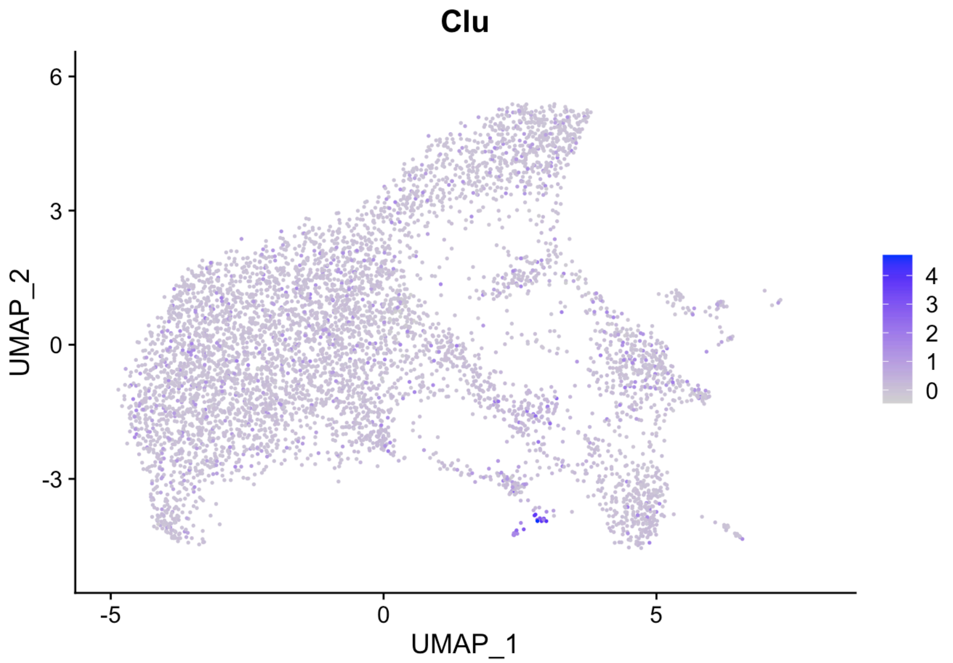


```
#Mef2c
F<-FeaturePlot(object =dCA2.integrated_clean_7celltype, features =  c("Mef2c"))
ggsave(file="/Users/lees44/R_project/FC_vs_dCA2_comparison_vector_integration_042821/clean_7celltype_Mef2c.pdf",w
idth=10,height=10)
plot(F)
```

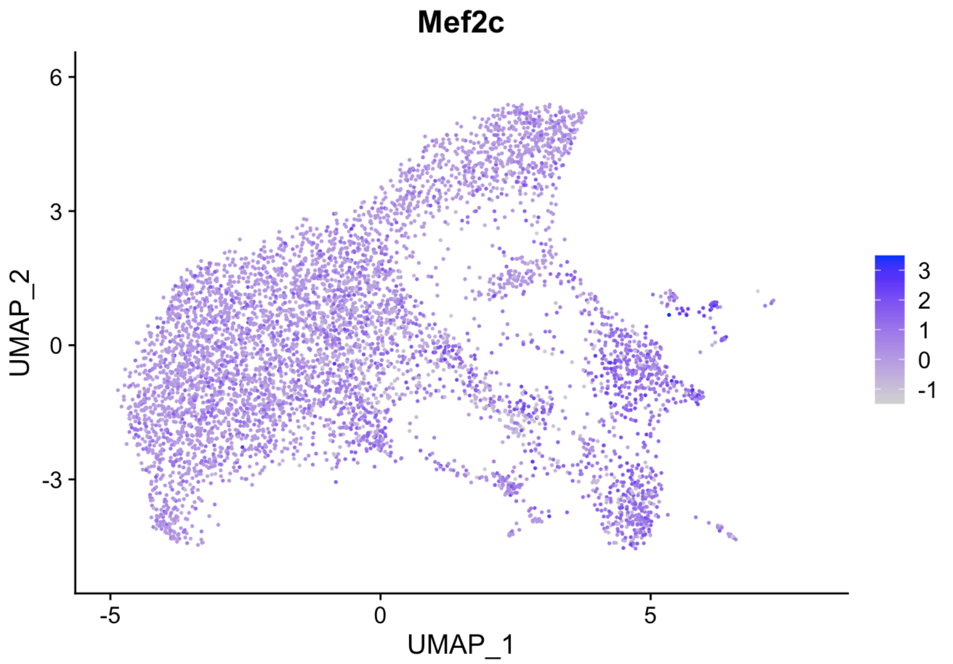

```
#Kcnk2
F<-FeaturePlot(object =dCA2.integrated_clean_7celltype, features =  c("Kcnk2"))
ggsave(file="/Users/lees44/R_project/FC_vs_dCA2_comparison_vector_integration_042821/clean_7celltype_Kcnk2.pdf",w
idth=10,height=10)
plot(F)
```

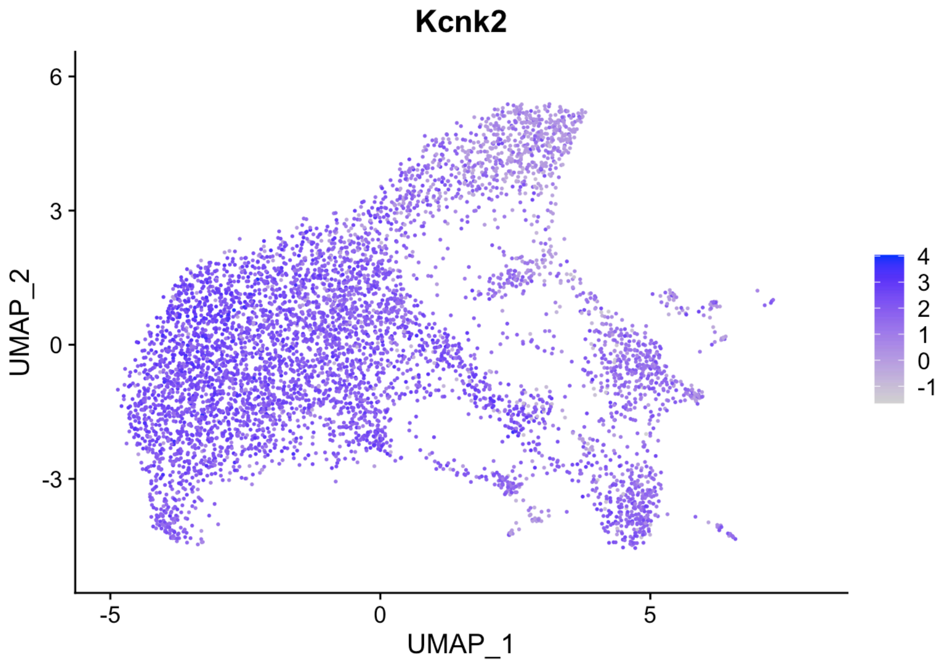


```
#BDNF
F<-FeaturePlot(object =dCA2.integrated_clean_7celltype, features =  c("Bdnf"))
ggsave(file="/Users/lees44/R_project/FC_vs_dCA2_comparison_vector_integration_042821/clean_7celltype_Bdnf.pdf",wi
dth=10,height=10)
plot(F)
```

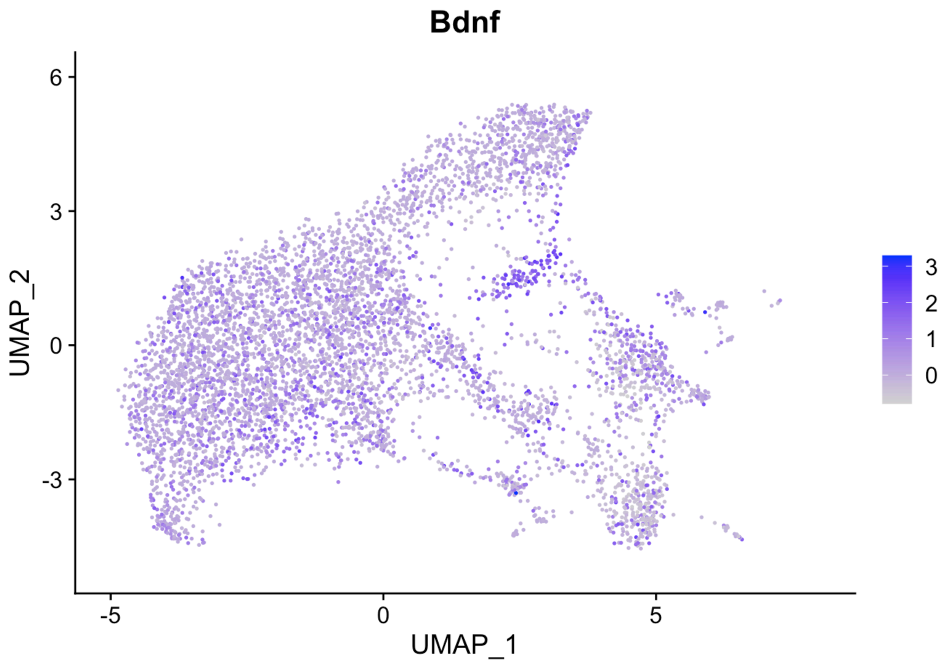

```
#Amigo2
F<-FeaturePlot(object =dCA2.integrated_clean_7celltype, features =  c("Amigo2"))
ggsave(file="/Users/lees44/R_project/FC_vs_dCA2_comparison_vector_integration_042821/clean_7celltype_Amigo2.pdf",
width=10,height=10)
plot(F)
```

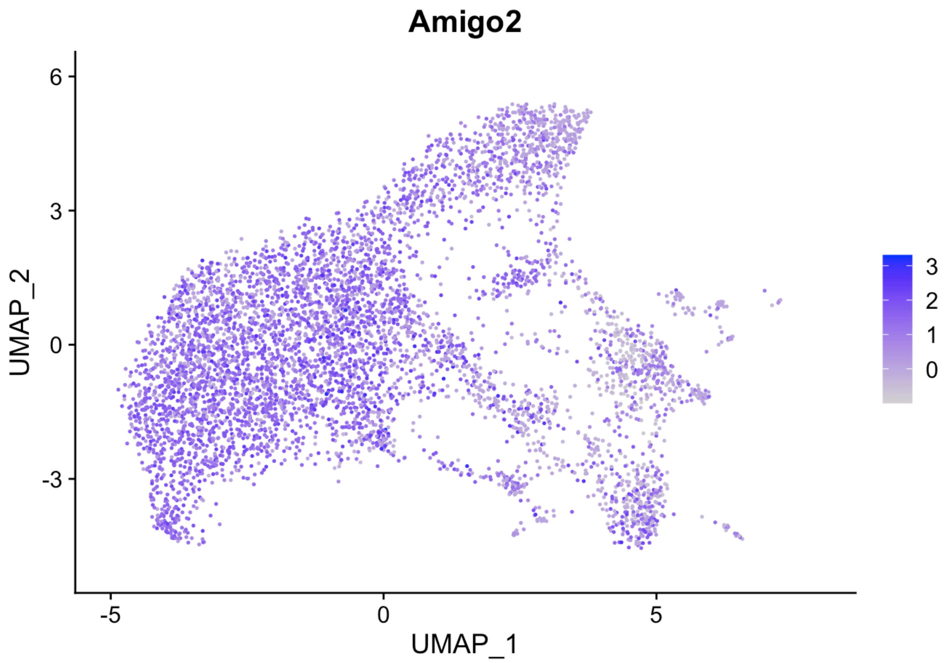


```
#Cpne7
F<-FeaturePlot(object =dCA2.integrated_clean_7celltype, features =  c("Cpne7"))
ggsave(file="/Users/lees44/R_project/FC_vs_dCA2_comparison_vector_integration_042821/clean_7celltype_Cpne7.pdf",w
idth=10,height=10)
plot(F)
```

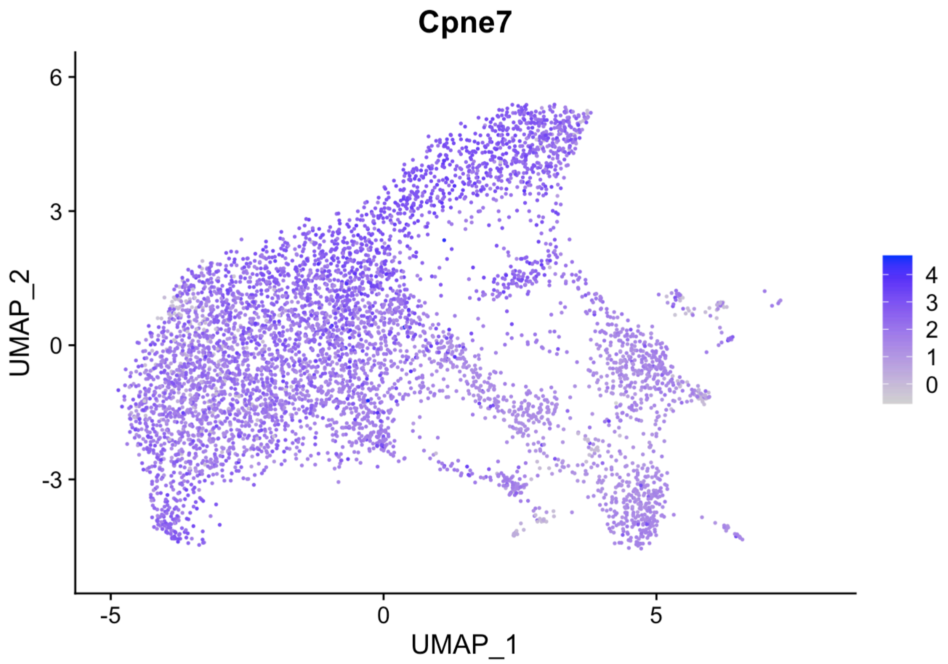

```
#Ccnd2
F<-FeaturePlot(object =dCA2.integrated_clean_7celltype, features =  c("Ccnd2"))
ggsave(file="/Users/lees44/R_project/FC_vs_dCA2_comparison_vector_integration_042821/clean_7celltype_Ccnd2.pdf",width=10,height=10)
plot(F)
```

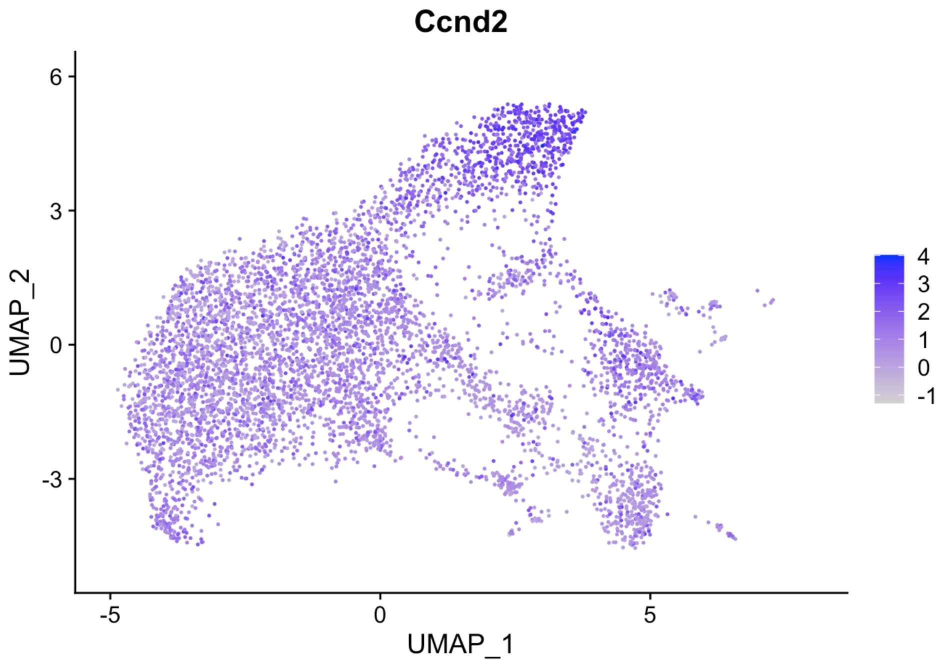


############################################08.05.2021##############################################################################
Visualize co-expression of two features simultaneously https://satijalab.org/seurat/articles/visualization_vignette.html (https://satijalab.org/seurat/articles/visualization_vignette.html)

```
# Visualize co-expression of two features simultaneously
FeaturePlot(dCA2.integrated_clean_7celltype, features = c("Mef2c", "Kcnk2"), blend = TRUE)
```

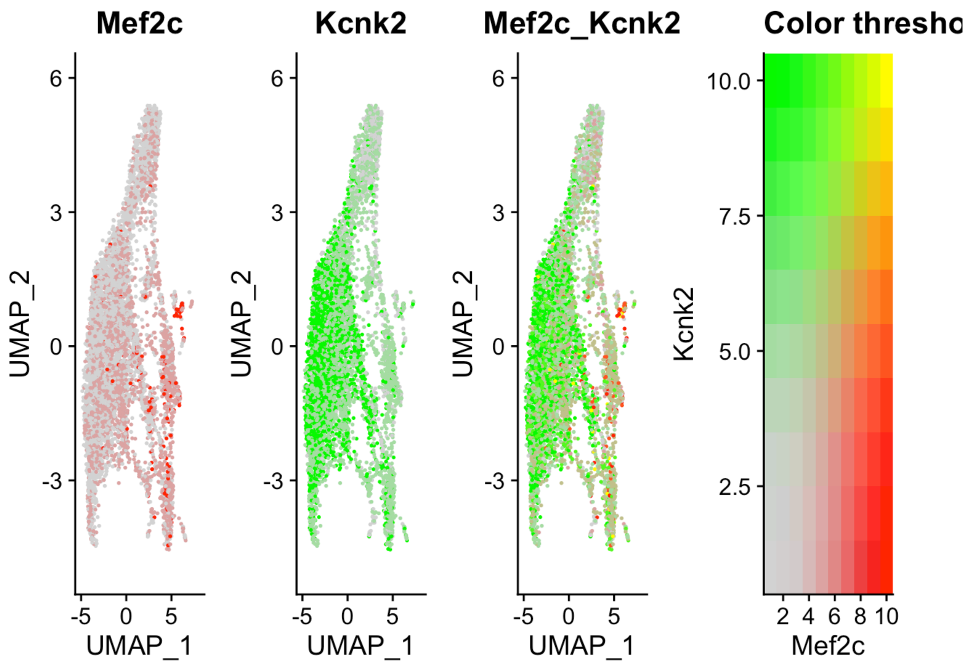

```
ggsave(file="/Users/lees44/R_project/FC_vs_dCA2_comparison_vector_integration_042821/clean_7celltype_merged_Mef2c
_Kcnk2.pdf",width=10,height=10)
```

```
# Visualize co-expression of two features simultaneously
FeaturePlot(dCA2.integrated_clean_7celltype, features = c("Kcnk2", "Amigo2"), blend = TRUE)
```

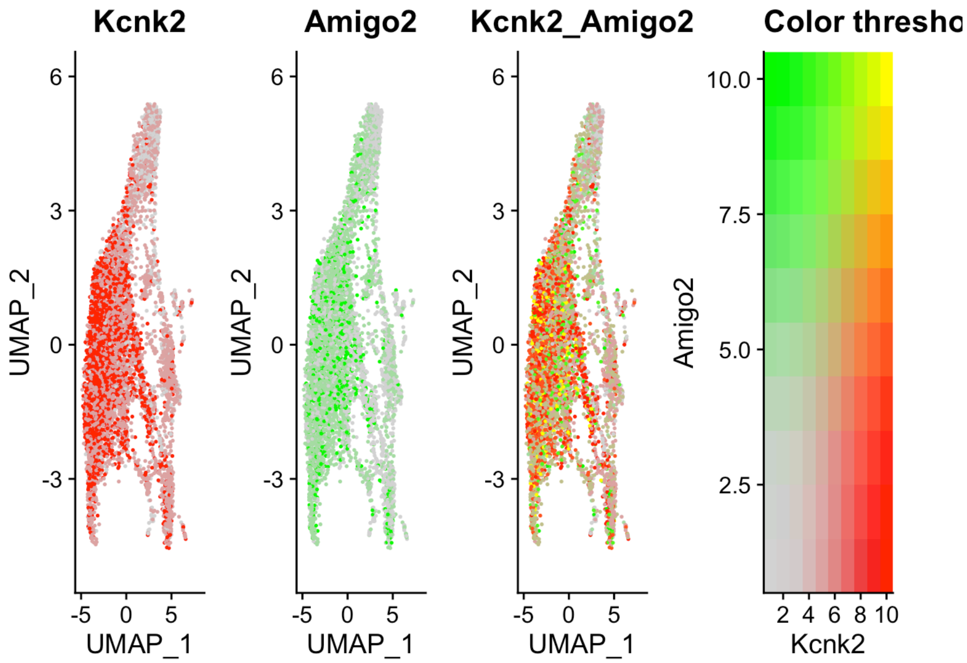


```
ggsave(file="/Users/lees44/R_project/FC_vs_dCA2_comparison_vector_integration_042821/clean_7celltype_merged_Kcnk2
_Amigo2.pdf",width=10,height=10)
```

```
# Visualize co-expression of two features simultaneously
FeaturePlot(dCA2.integrated_clean_7celltype, features = c("Amigo2", "Ccnd2"), blend = TRUE)
```

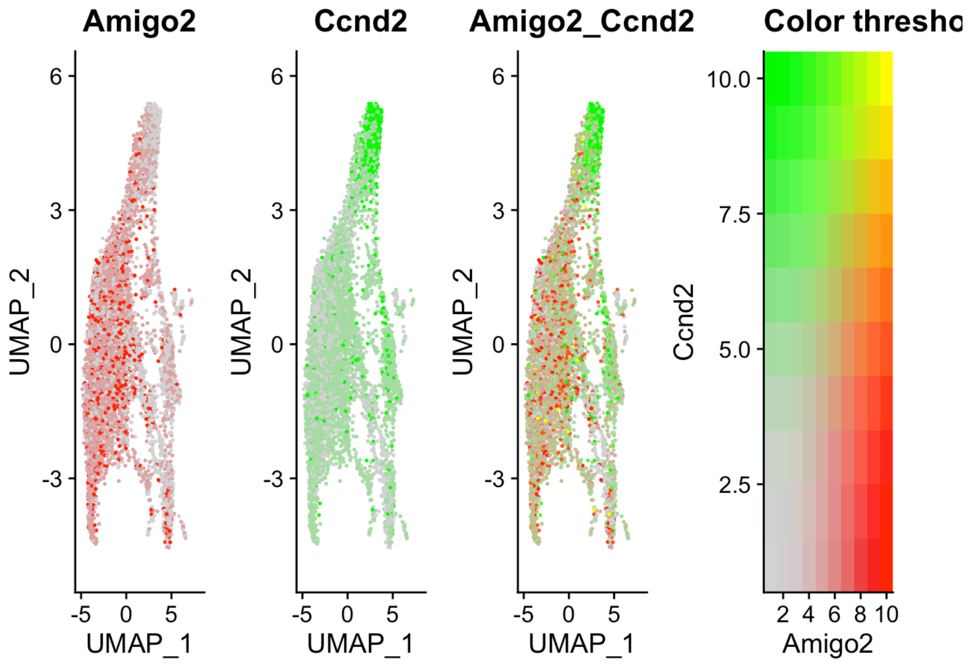

```
ggsave(file="/Users/lees44/R_project/FC_vs_dCA2_comparison_vector_integration_042821/clean_7celltype_merged_Amigo2_Ccnd2.pdf",width=10,height=10)
```

```
# Visualize co-expression of two features simultaneously
FeaturePlot(dCA2.integrated_clean_7celltype, features = c("Nptx1", "C1ql3"), blend = TRUE, max.cutoff = 2)
```

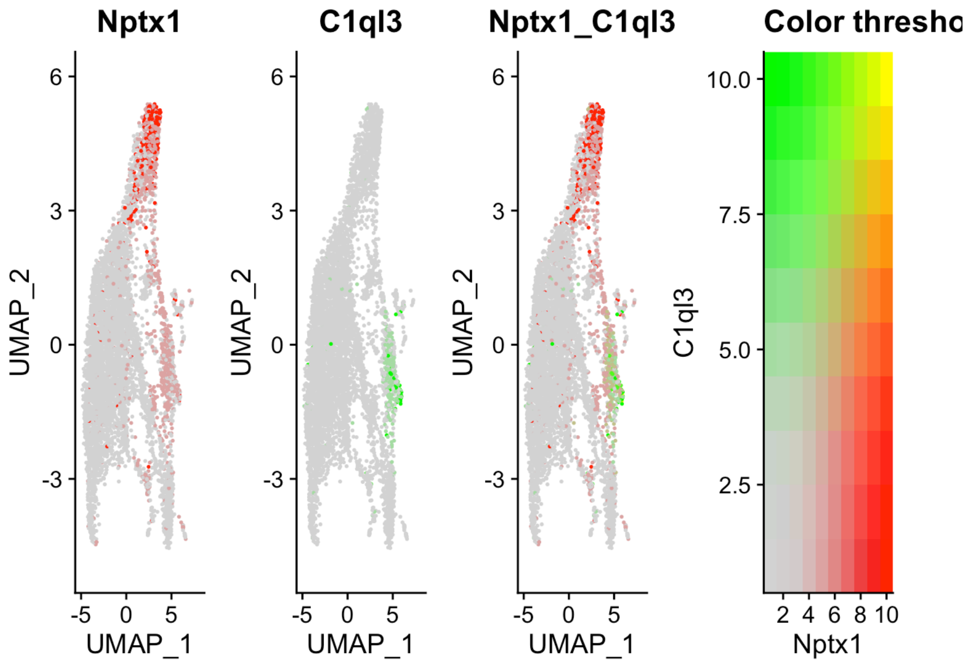


```
ggsave(file="/Users/lees44/R_project/FC_vs_dCA2_comparison_vector_integration_042821/clean_7celltype_merged_Nptx1_C1ql3.pdf",width=10,height=10)
```

```
# Visualize co-expression of two features simultaneously
FeaturePlot(dCA2.integrated_clean_7celltype, features = c("Ptgds", "Atp1a2"), blend = TRUE, blend.threshold = 0.2, max.cutoff = 2)
```

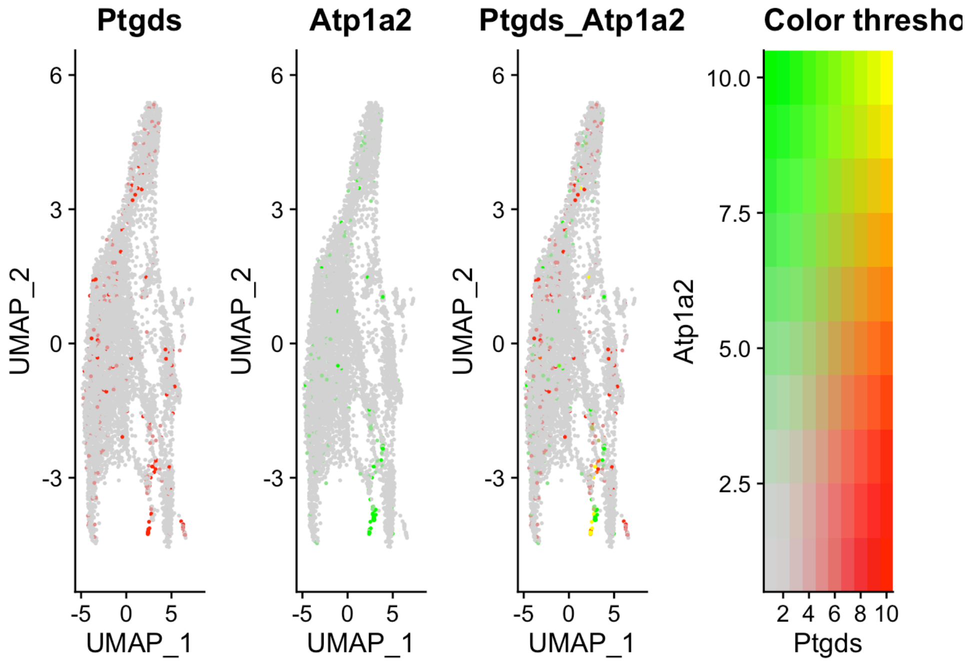

```
ggsave(file="/Users/lees44/R_project/FC_vs_dCA2_comparison_vector_integration_042821/clean_7celltype_merged_Ptgds_Aptla2.pdf",width=10,height=10)
```

```
# Visualize co-expression of two features simultaneously
FeaturePlot(dCA2.integrated_clean_7celltype, features = c("Ptgds", "Atp1a2"), blend = TRUE,blend.threshold = 0.1,
max.cutoff = 2)
```

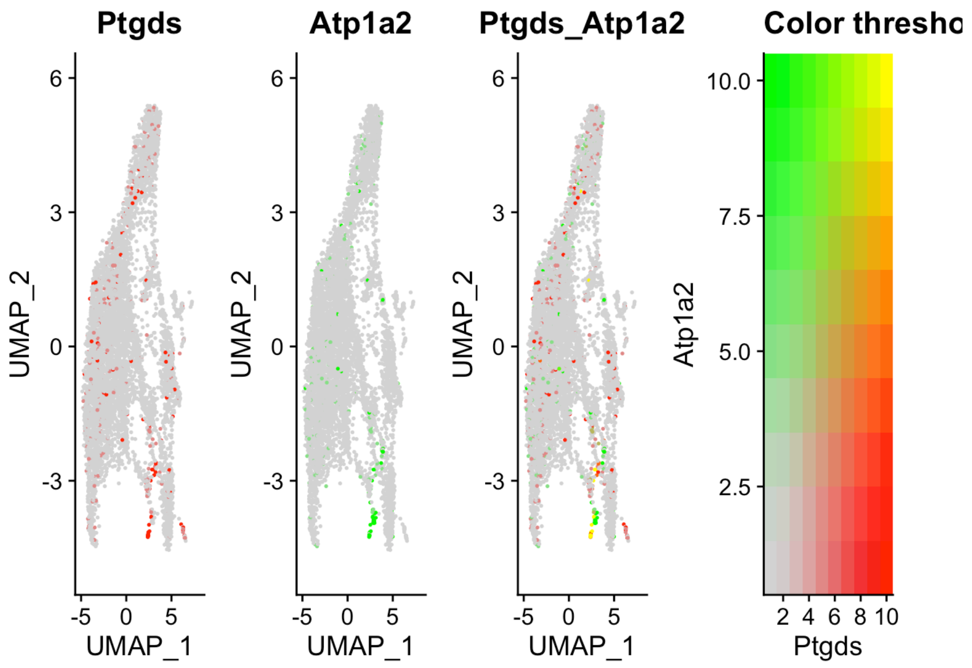


```
ggsave(file="/Users/lees44/R_project/FC_vs_dCA2_comparison_vector_integration_042821/clean_7celltype_merged_Ptgds_Apoe.pdf",width=10,height=10)
```

```
# Visualize co-expression of two features simultaneously
FeaturePlot(dCA2.integrated_clean_7celltype, features = c("Gjc3", "Atp1a2"), blend = TRUE)
```

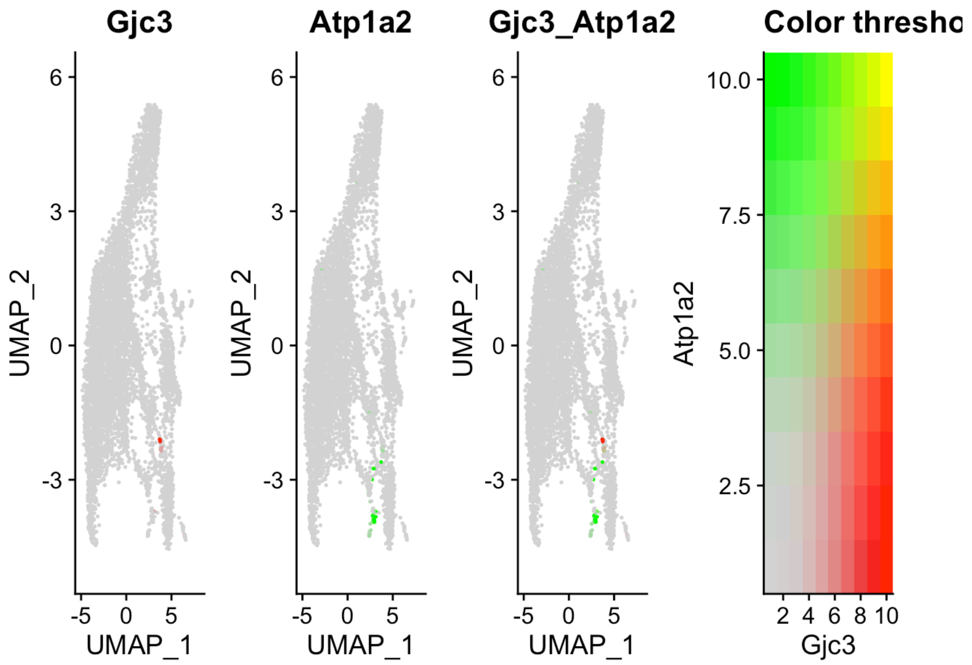

```
ggsave(file="/Users/lees44/R_project/FC_vs_dCA2_comparison_vector_integration_042821/clean_7celltype_merged_Gjc3_Apt1a2.pdf",width=10,height=10)
```

rcolor brewer for better color choices. https://www.r-graph-gallery.com/38-rcolorbrewers-palettes.html (https://www.r-graph-gallery.com/38-rcolorbrewers-palettes.html)

```
install.packages("RColorBrewer")
```

```
## Installing package into '/Users/lees44/Library/R/4.0/library'
## (as 'lib' is unspecified)
```

```
## Error in contrib.url(repos, "source"): trying to use CRAN without setting a mirror
```

```
library("RColorBrewer")
```

change point size: see https://www.rdocumentation.org/packages/Seurat/versions/4.0.3/topics/FeaturePlot (https://www.rdocumentation.org/packages/Seurat/versions/4.0.3/topics/FeaturePlot)

```
#Ccnd2
FeaturePlot(object =dCA2.integrated_clean_7celltype, features =  c("Ccnd2"), blend.threshold = 0.5, min.cutoff = 0, max.cutoff = 2, pt.size = 1.5) + scale_colour_gradientn(colours = rev(brewer.pal(n = 11, name = "Spectral")))
```

```
## Scale for 'colour' is already present. Adding another scale for 'colour',
## which will replace the existing scale.
```

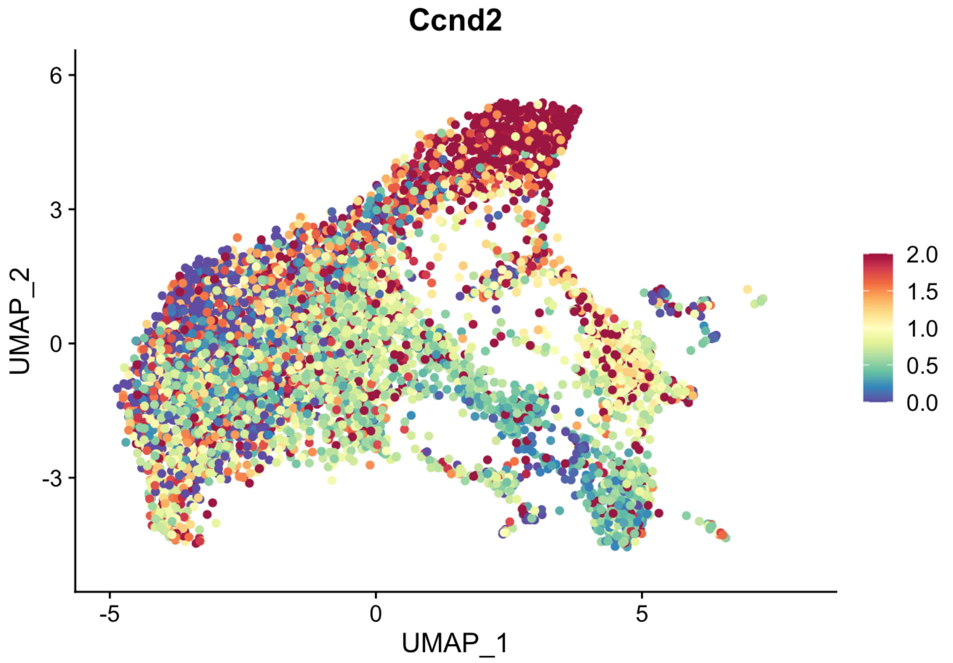


```
ggsave(file="/Users/lees44/R_project/FC_vs_dCA2_comparison_vector_integration_042821/clean_7celltype_Ccnd2_color.pdf",width=10,height=10)
```

Trying different color palattes fo TAC1 expression.

```
#Tac1
FeaturePlot(object =dCA2.integrated_clean_7celltype, features =  c("Tac1"), blend.threshold = 0.5, min.cutoff = 0, max.cutoff = 2, pt.size = 1.5, cols = c("pink", "red"))
```

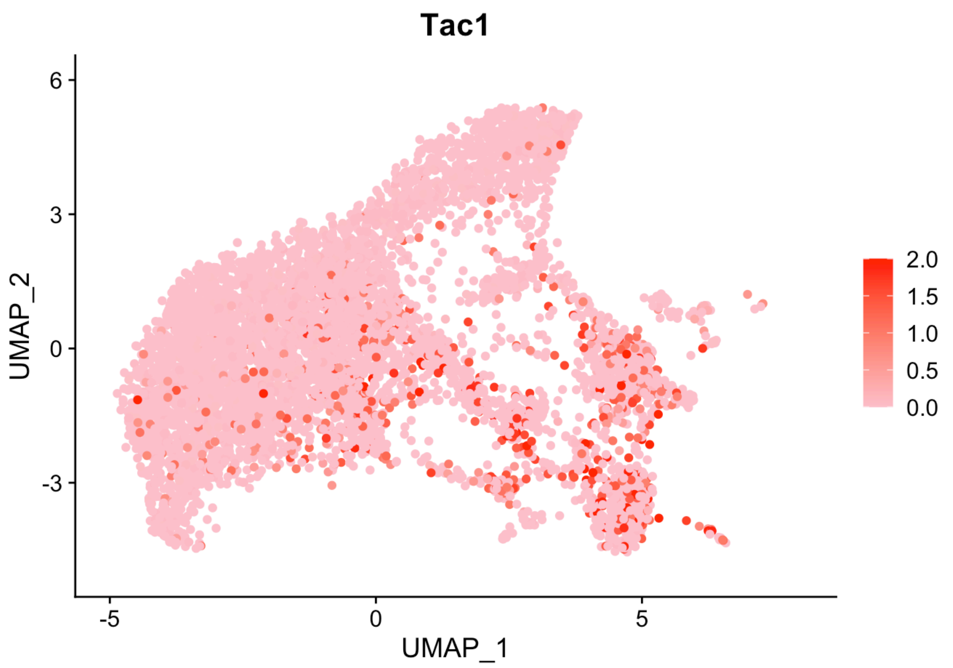


```
ggsave(file="/Users/lees44/R_project/FC_vs_dCA2_comparison_vector_integration_042821/clean_7celltype_Tac1_color.pdf",width=10,height=10)
```

```
#Tac1
FeaturePlot(object =dCA2.integrated_clean_7celltype, features =  c("Tac1"), blend.threshold = 0.5, min.cutoff = 0, max.cutoff = 2, pt.size = 1.5, cols = c("lightblue", "red"))
```

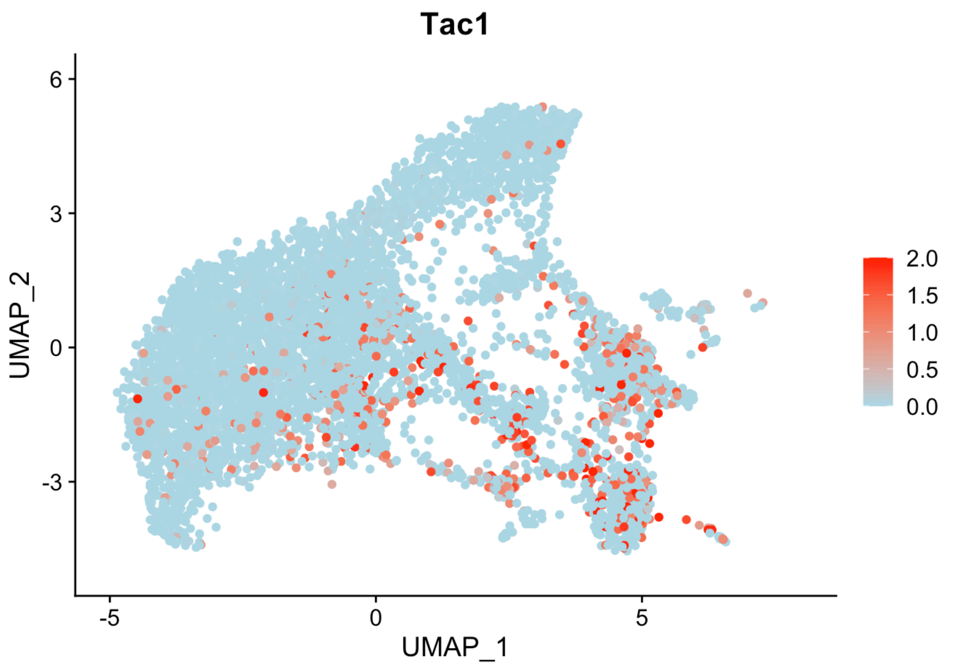


```
ggsave(file="/Users/lees44/R_project/FC_vs_dCA2_comparison_vector_integration_042821/clean_7celltype_Tac1_blue_red_color.pdf",width=10,height=10)
```

```
#Tac1
FeaturePlot(object =dCA2.integrated_clean_7celltype, features =  c("Tac1"), blend.threshold = 0.5, min.cutoff =
0, max.cutoff = 2, pt.size = 1.5) + scale_colour_gradientn(colours = rev(brewer.pal(n = 11, name = "Spectral")))
```

```
## Scale for 'colour' is already present. Adding another scale for 'colour',
## which will replace the existing scale.
```

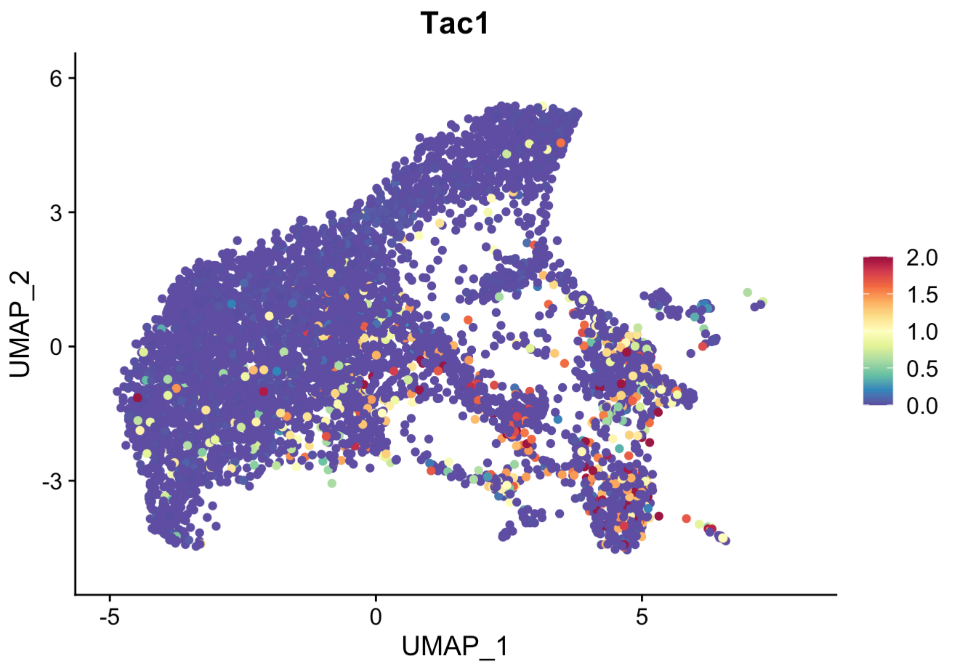


```
ggsave(file="/Users/lees44/R_project/FC_vs_dCA2_comparison_vector_integration_042821/clean_7celltype_Tac1_color_s
pectral.pdf",width=10,height=10)
```

```
#Tac1
FeaturePlot(object =dCA2.integrated_clean_7celltype, features =  c("Cdh13", "C1ql13", "Ptgds"), blend.threshold =
0.5, min.cutoff = 0, max.cutoff = 2, pt=1) + scale_fill_continuous(type= "viridis")
```

```
## Warning in FetchData(object = object, vars = c(dims, "ident", features), : The
## following requested variables were not found: C1ql13
```

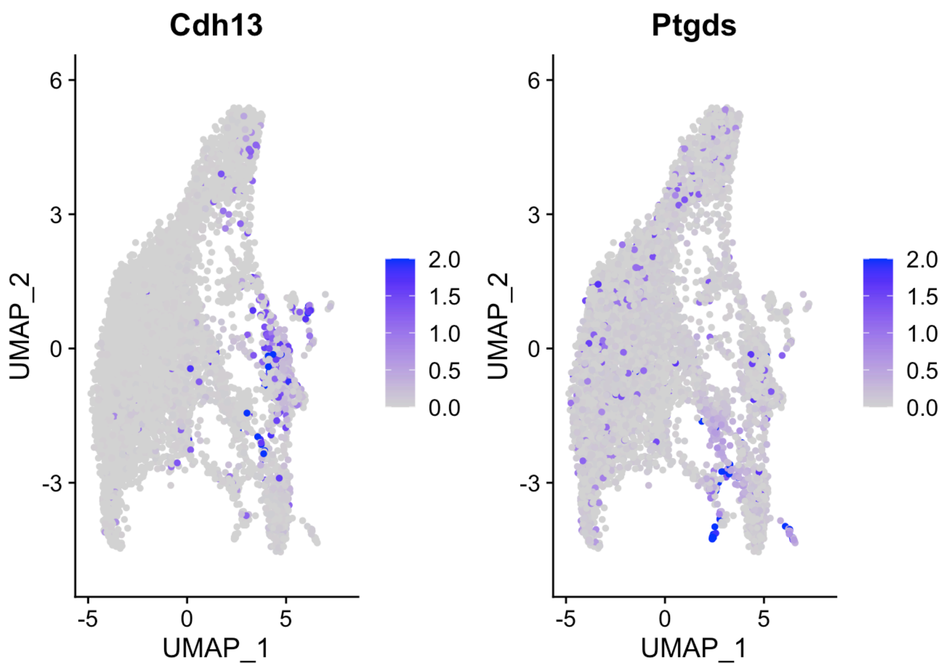


```
ggsave(file="/Users/lees44/R_project/FC_vs_dCA2_comparison_vector_integration_042821/clean_7celltype_Cdh13_C1ql13
_Ptgds_color_viridis.pdf",width=10,height=10)
```

```
#Tac1
FeaturePlot(object =dCA2.integrated_clean_7celltype, features =  c("Cdh13", "C1ql3", "Ptgds"), blend.threshold =
0.5, min.cutoff = 0, max.cutoff = 2, pt=1) + scale_fill_continuous(type= "viridis")
```

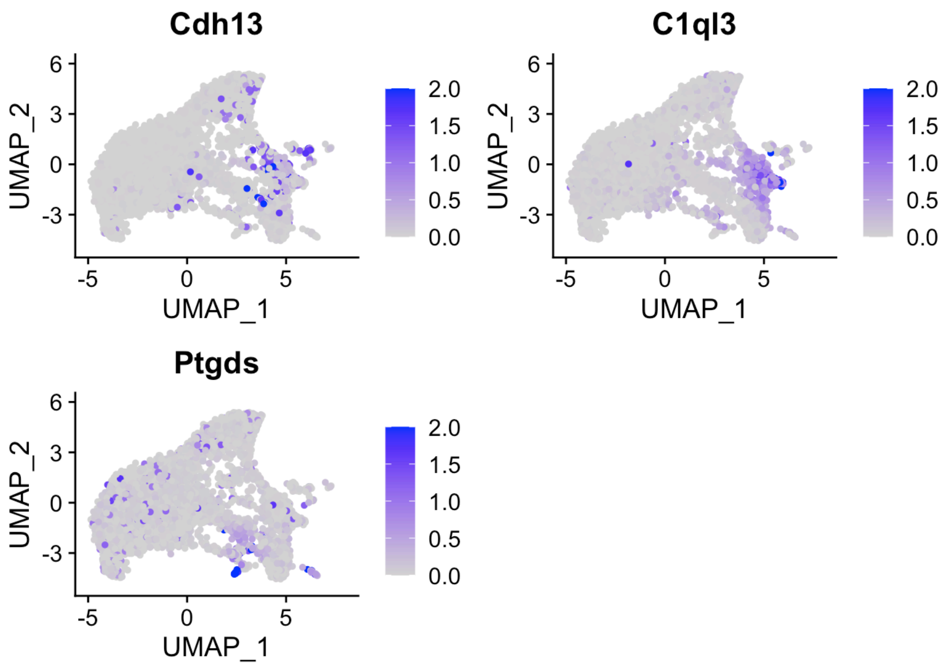


```
ggsave(file="/Users/lees44/R_project/FC_vs_dCA2_comparison_vector_integration_042821/clean_7celltype_Cdh13_C1ql13
_Ptgds_color_viridis.pdf",width=10,height=10)
```

```
#Tac1
FeaturePlot(object =dCA2.integrated_clean_7celltype, features =  c("Slit2", "Lnx1", "Nptx1"), blend.threshold =
0.5, min.cutoff = 0, max.cutoff = 2, pt=1) + scale_fill_continuous(type= "viridis")
```

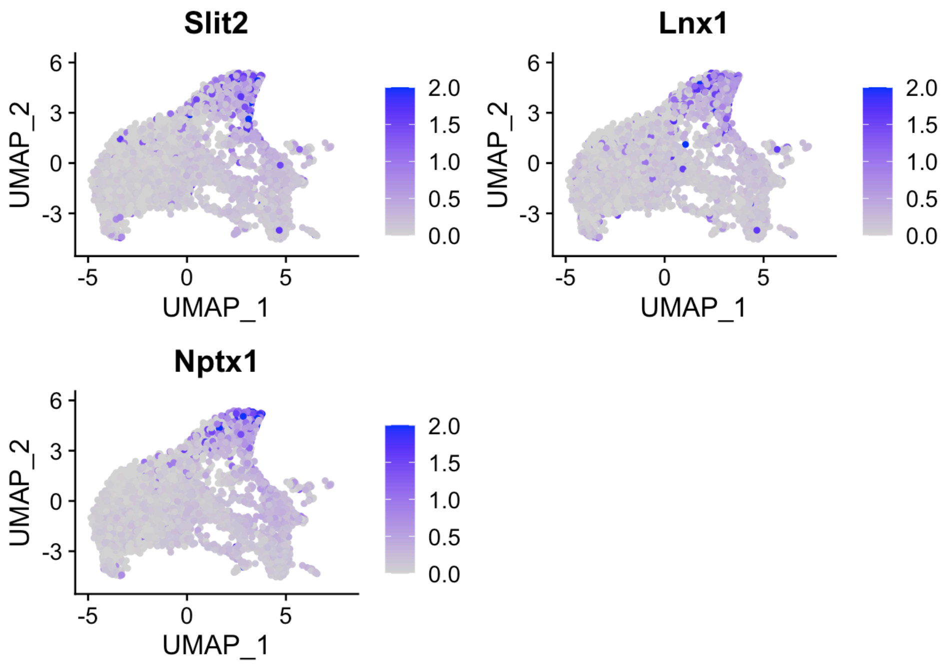


```
ggsave(file="/Users/lees44/R_project/FC_vs_dCA2_comparison_vector_integration_042821/clean_7celltype_Slit2_Lnx1_N
ptx1_color_viridis.pdf",width=10,height=10)
```

For all the gene list combined markers for 7 cell types 08.23.2021 All of 900 genes to vizualize best cell type markers. The genes were chosen based on their RNA transcript number. Refer to the extract raw count code

```
dCA2.integrated = readRDS(file="/Users/lees44/R_project/dCA2.integrated.rds")
# Switch to the desired identity
Idents(object = dCA2.integrated) <- "type"
Cell_type<-c("Type1", "Type2", "Type3", "Type4", "Type5", "Type6", "Type7")


# genes of interest
gene_df = read.csv("/Users/lees44/R_project/FC_vs_dCA2_comparison_vector_integration_042821/combined_all_markers_
clean_7celltype.csv", header = T)
gene_list <- gene_df$gene


# generate the workbook

for (i in 1:length(Cell_type)){
  tryCatch({
    subset = subset(x = dCA2.integrated, idents = Cell_type[i])
    raw_counts = as.data.frame(subset@assays$RNA@counts)
    filtered_raw_counts = raw_counts[row.names(raw_counts) %in% gene_list,]
    filtered_raw_counts <- filtered_raw_counts[order(match(rownames(filtered_raw_counts), gene_list)),]
    write.csv(t(filtered_raw_counts), paste0(Cell_type[i],".csv"))
    gc()
  }, error=function(e){})

}
```

```
# genes of interest
gene_df = read.csv("/Users/lees44/R_project/FC_vs_dCA2_comparison_vector_integration_042821/FEATUREPLOT_GENELIST/Gene_64.csv", header = T)
gene_list <- gene_df$gene
```

```
#Gene list
FeaturePlot(object =dCA2.integrated_clean_7celltype, features = gene_list, blend.threshold = 0.5, min.cutoff = 0, max.cutoff = 2, pt=0.2) + scale_fill_continuousex(type= "viridis")
```

```
## Error in scale_fill_continuousex(type = "viridis"): could not find function "scale_fill_continuousex"
```

```
ggsave(file="/Users/lees44/R_project/FC_vs_dCA2_comparison_vector_integration_042821/FEATUREPLOT_GENELIST/gene_64.pdf", width = 10, height = 10)
```

```
#Tac1
FeaturePlot(object =dCA2.integrated_clean_7celltype, features =  c("Tac1"), blend.threshold = 0.5, min.cutoff = 0, max.cutoff = 2, pt.size = 1.5) + scale_colour_gradientn(colours = rev(brewer.pal(n = 11, name = "RdYlGn")))
```

```
## Scale for 'colour' is already present. Adding another scale for 'colour',
## which will replace the existing scale.
```

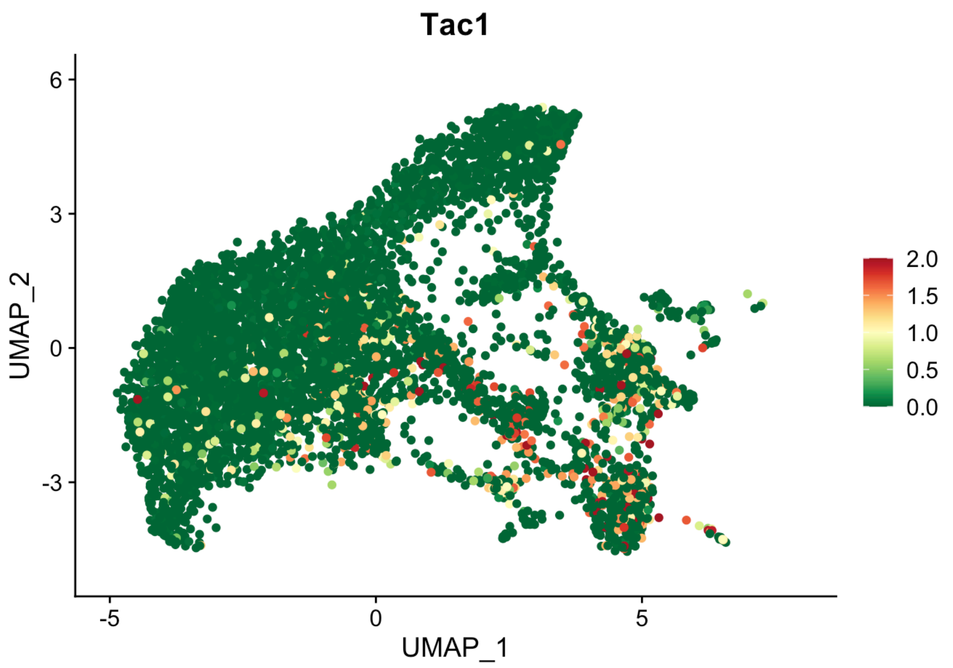


```
ggsave(file="/Users/lees44/R_project/FC_vs_dCA2_comparison_vector_integration_042821/clean_7celltype_Tac1_color_set1.pdf",width=10,height=10)
```

```
#Tac1
FeaturePlot(object =dCA2.integrated_clean_7celltype, features =  c("Tac1"), blend.threshold = 0.5, min.cutoff = 0, max.cutoff = 2, pt.size = 1.5) + scale_colour_gradientn(colours = rev(brewer.pal(n = 11, name = "RdBu")))
```

```
## Scale for 'colour' is already present. Adding another scale for 'colour',
## which will replace the existing scale.
```

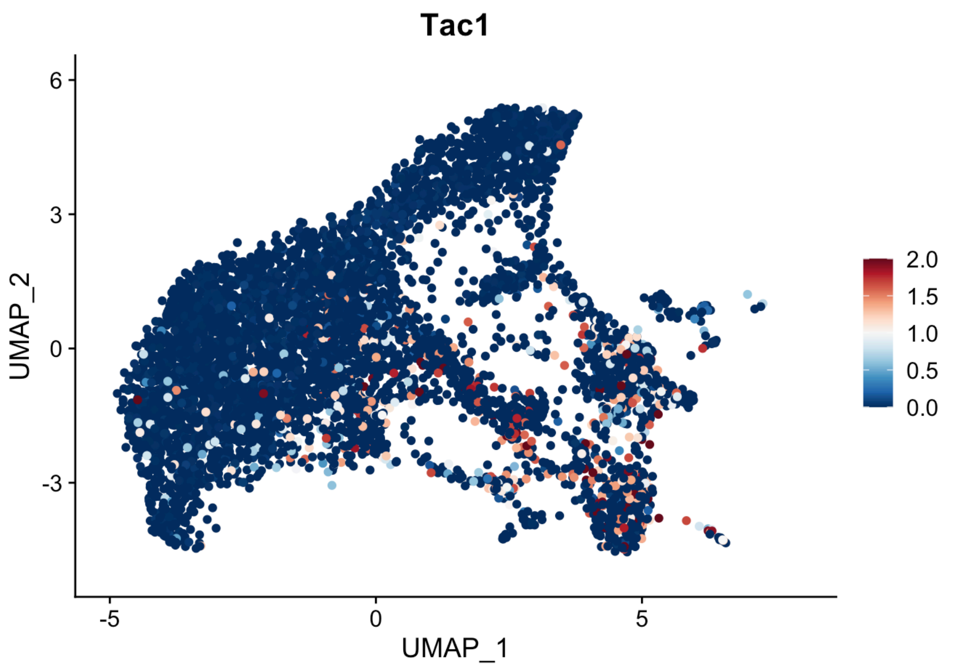


```
ggsave(file="/Users/lees44/R_project/FC_vs_dCA2_comparison_vector_integration_042821/clean_7celltype_Tac1_color_RdBu.pdf",width=10,height=10)
```

```
#Mef2c
FeaturePlot(object =dCA2.integrated_clean_7celltype, features =  c("Mef2c"), blend.threshold = 0.5, min.cutoff = 0, max.cutoff = 2, pt.size = 1.5) + scale_colour_gradientn(colours = rev(brewer.pal(n = 11, name = "RdBu")))
```

```
## Scale for 'colour' is already present. Adding another scale for 'colour',
## which will replace the existing scale.
```

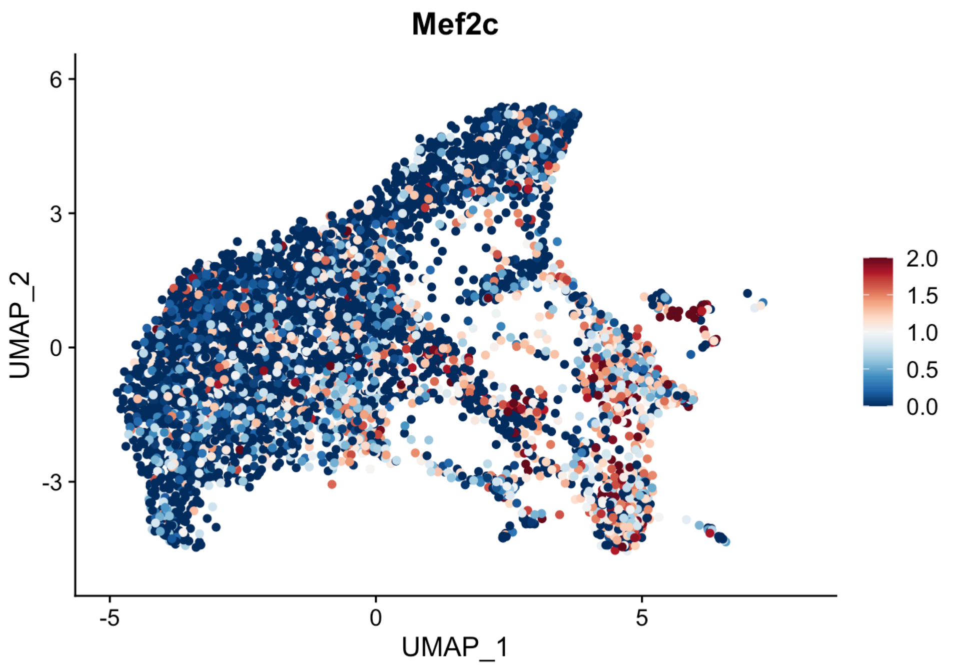


```
ggsave(file="/Users/lees44/R_project/FC_vs_dCA2_comparison_vector_integration_042821/clean_7celltype_Mef2c_color.pdf",width=10,height=10)
```

```
#Mef2c
FeaturePlot(object =dCA2.integrated_clean_7celltype, features =  c("Mef2c"), blend.threshold = 0.5, min.cutoff =
0, max.cutoff = 2, pt.size = 1.5, cols = c("lightblue", "red"))
```

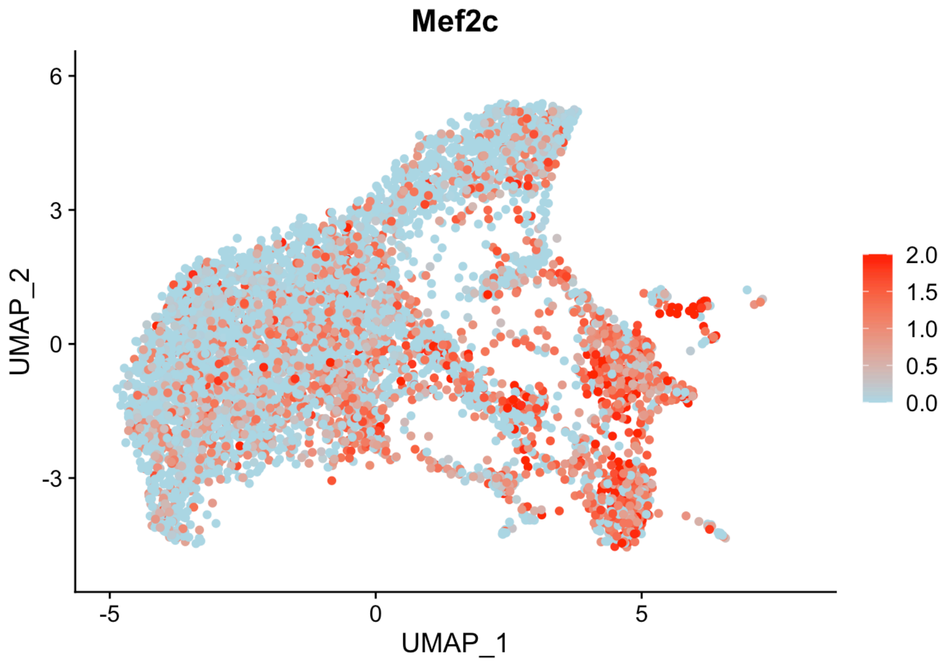


```
ggsave(file="/Users/lees44/R_project/FC_vs_dCA2_comparison_vector_integration_042821/clean_7celltype_Mef2c_blue_r
ed_color.pdf",width=10,height=10)
```

```
#Clu
FeaturePlot(object =dCA2.integrated_clean_7celltype, features =  c("Clu"), blend.threshold = 0.5, min.cutoff = 0,
max.cutoff = 2, pt.size = 1.5) + scale_colour_gradientn(colours = rev(brewer.pal(n = 11, name = "RdBu")))
```

```
## Scale for 'colour' is already present. Adding another scale for 'colour',
## which will replace the existing scale.
```

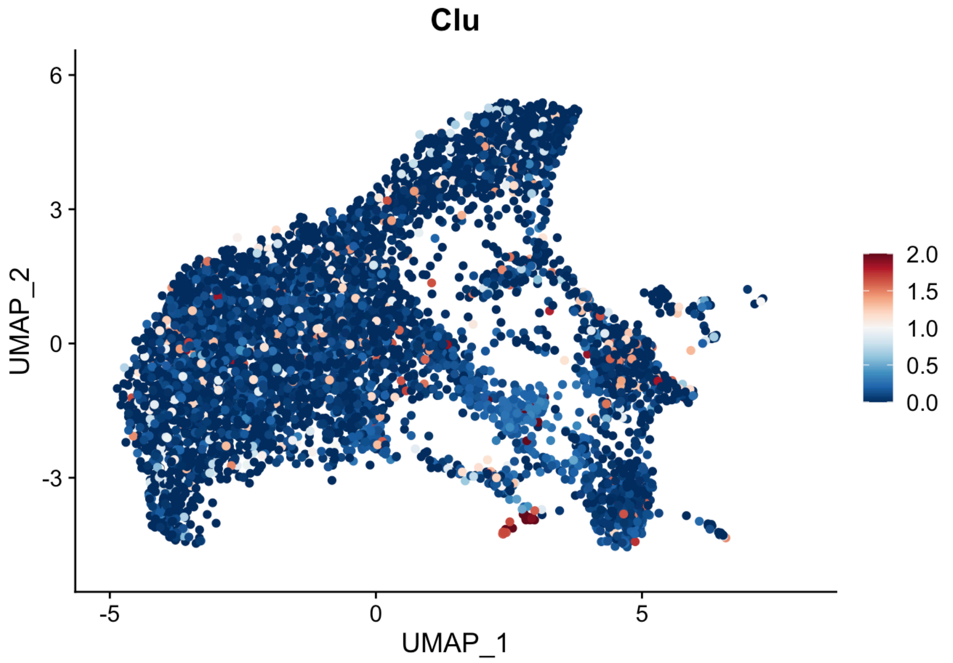


```
ggsave(file="/Users/lees44/R_project/FC_vs_dCA2_comparison_vector_integration_042821/clean_7celltype_Clu_color.pdf",width=10,height=10)
```

```
#Kcnk2
FeaturePlot(object =dCA2.integrated_clean_7celltype, features =  c("Kcnk2"), blend.threshold = 0.5, min.cutoff = 0, max.cutoff = 2, pt.size = 1.5) + scale_colour_gradientn(colours = rev(brewer.pal(n = 11, name = "RdBu")))
```

```
## Scale for 'colour' is already present. Adding another scale for 'colour',
## which will replace the existing scale.
```

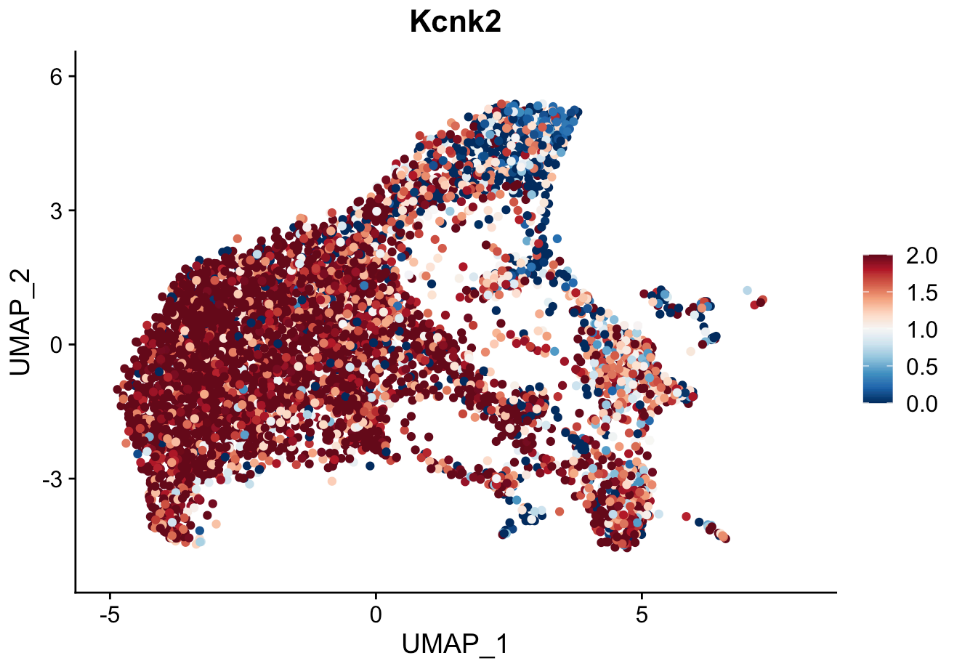


```
ggsave(file="/Users/lees44/R_project/FC_vs_dCA2_comparison_vector_integration_042821/clean_7celltype_Kcnk2_color.pdf",width=10,height=10)
```

```
#Bdnf
FeaturePlot(object =dCA2.integrated_clean_7celltype, features =  c("Bdnf"), blend.threshold = 0.5, min.cutoff =
0, max.cutoff = 2, pt.size = 1.5) + scale_colour_gradientn(colours = rev(brewer.pal(n = 11, name = "RdBu")))
```

```
## Scale for 'colour' is already present. Adding another scale for 'colour',
## which will replace the existing scale.
```

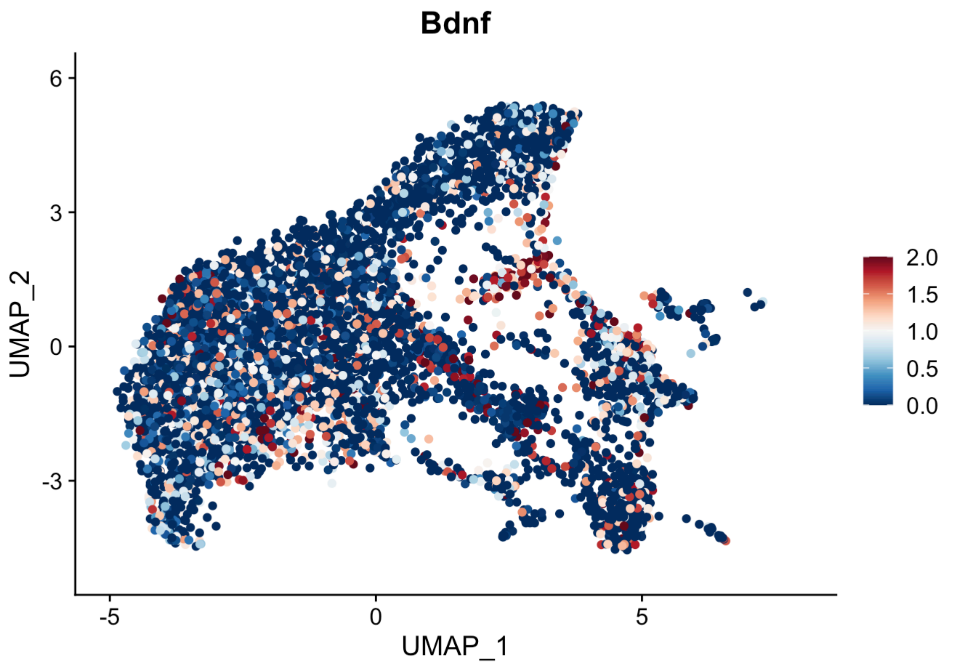


```
ggsave(file="/Users/lees44/R_project/FC_vs_dCA2_comparison_vector_integration_042821/clean_7celltype_Bdnf_color.p
df",width=10,height=10)
```

```
#Amigo2
FeaturePlot(object =dCA2.integrated_clean_7celltype, features =  c("Amigo2"), blend.threshold = 0.5, min.cutoff =
0, max.cutoff = 2, pt.size = 1.5) + scale_colour_gradientn(colours = rev(brewer.pal(n = 11, name = "RdBu")))
```

```
## Scale for 'colour' is already present. Adding another scale for 'colour',
## which will replace the existing scale.
```

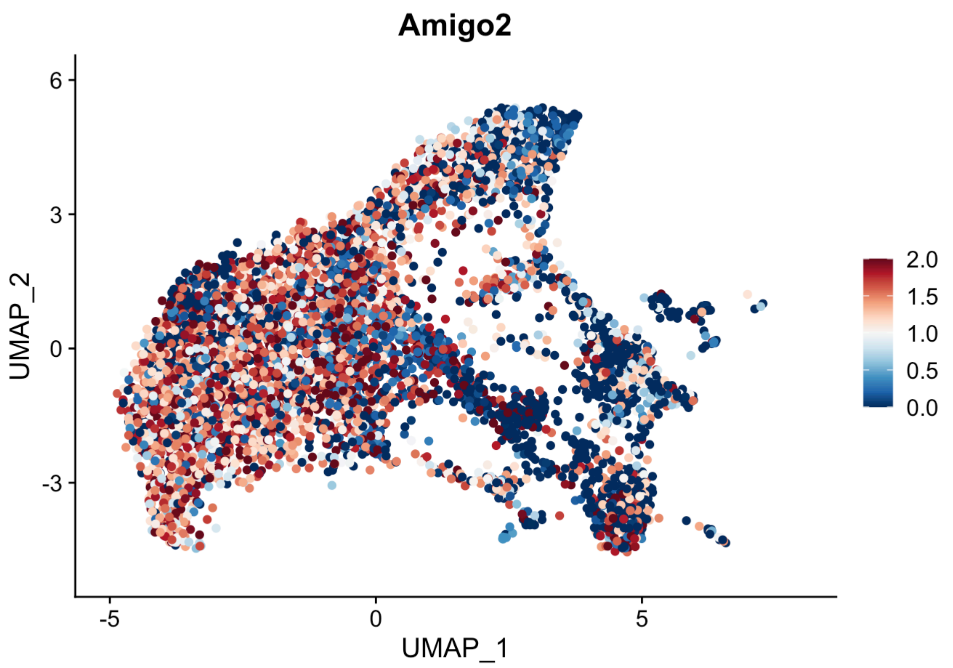


```
ggsave(file="/Users/lees44/R_project/FC_vs_dCA2_comparison_vector_integration_042821/clean_7celltype_Amigo2_colo
r.pdf",width=10,height=10)
```

```
#Ccnd2
FeaturePlot(object =dCA2.integrated_clean_7celltype, features =  c("Ccnd2"), blend.threshold = 0.5, min.cutoff =
0, max.cutoff = 2) + scale_colour_gradientn(colours = rev(brewer.pal(n = 11, name = "RdBu")))
```

```
## Scale for 'colour' is already present. Adding another scale for 'colour',
## which will replace the existing scale.
```

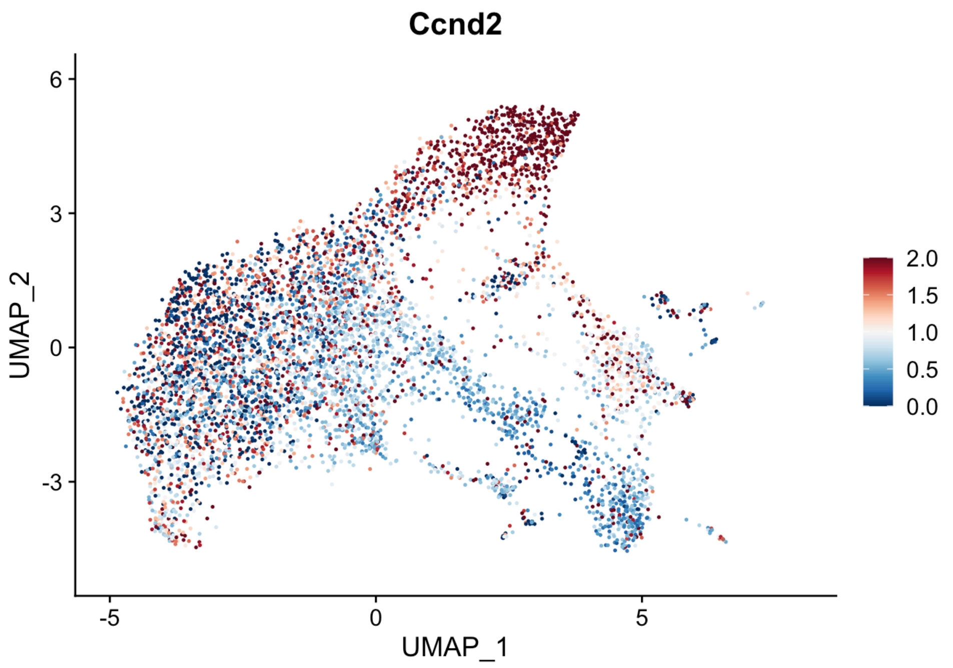


```
ggsave(file="/Users/lees44/R_project/FC_vs_dCA2_comparison_vector_integration_042821/clean_7celltype_Ccnd2_color.
pdf",width=10,height=10)
```

######################################081921########################

Cell type numbers for 7 cell types

```
# How many cells are in each cluster
table(Idents(dCA2.integrated_clean_7celltype))
```

```
##
## Type1 Type2 Type3 Type4 Type5 Type6 Type7
##  1830  1746   886   635   428   176   173
```

```
df <-data.frame(
  group = c("Type1", "Type2", "Type3", "Type4", "Type5", "Type6","Type7"),
  value = c(1830, 1746, 886, 635, 428, 176, 173)
)
head(df)
```

```
##   group value
## 1 Type1  1830
## 2 Type2  1746
## 3 Type3   886
## 4 Type4   635
## 5 Type5   428
## 6 Type6   176
```

```
bp<- ggplot(df, aes(x="", y=value, fill=group))+
geom_bar(width = 1, stat = "identity")
bp
```

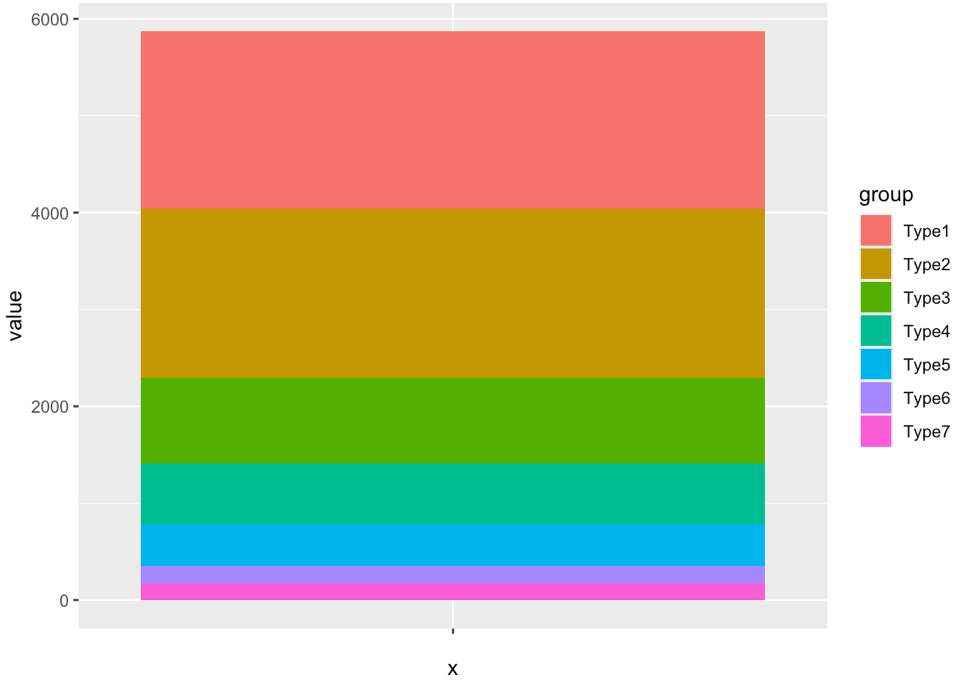


```
pie <- bp + coord_polar("y", start=0)
pie
```

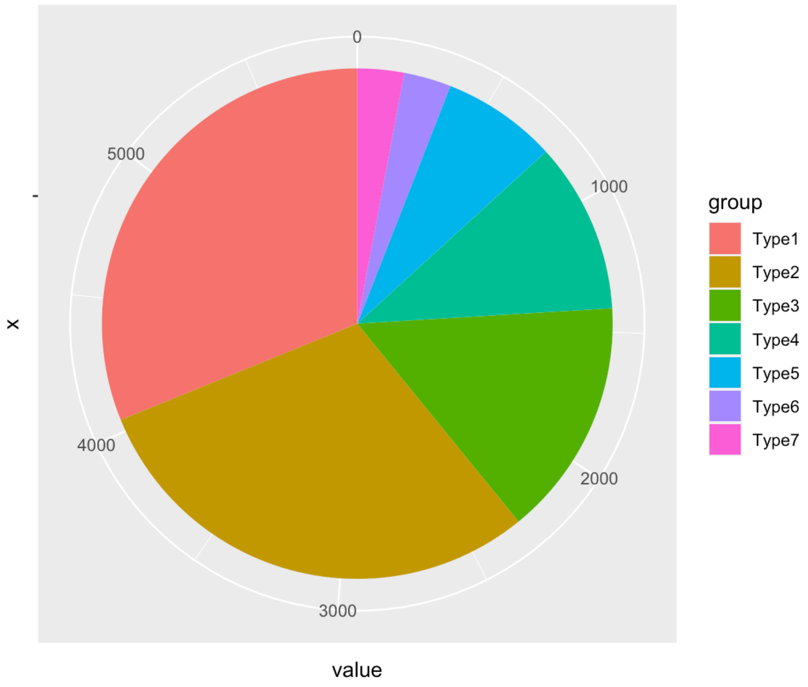


```
ggsave(file="/Users/lees44/R_project/FC_vs_dCA2_comparison_vector_integration_042821/cell_number_cluster_pie_7cel
ltype.pdf",width=10,height=7)
```

```
# How many cells are in each group?
table(dCA2.integrated_clean_7celltype$protocol)
```

```
##
## dCA2_Fpos dCA2_Mpos   FC_Fpos   FC_Mpos
##      1119      1953      1562      1240
```

```
df <-data.frame(
  group = c("dCA2_Fpos", "dCA2_Mpos", "FC_Fpos", "FC_Mpos"),
  value = c(1119, 1953, 1562, 1240)
)
head(df)
```

```
##       group value
## 1 dCA2_Fpos  1119
## 2 dCA2_Mpos  1953
## 3   FC_Fpos  1562
## 4   FC_Mpos  1240
```

```
bp<- ggplot(df, aes(x="", y=value, fill=group))+
geom_bar(width = 1, stat = "identity")
bp
```

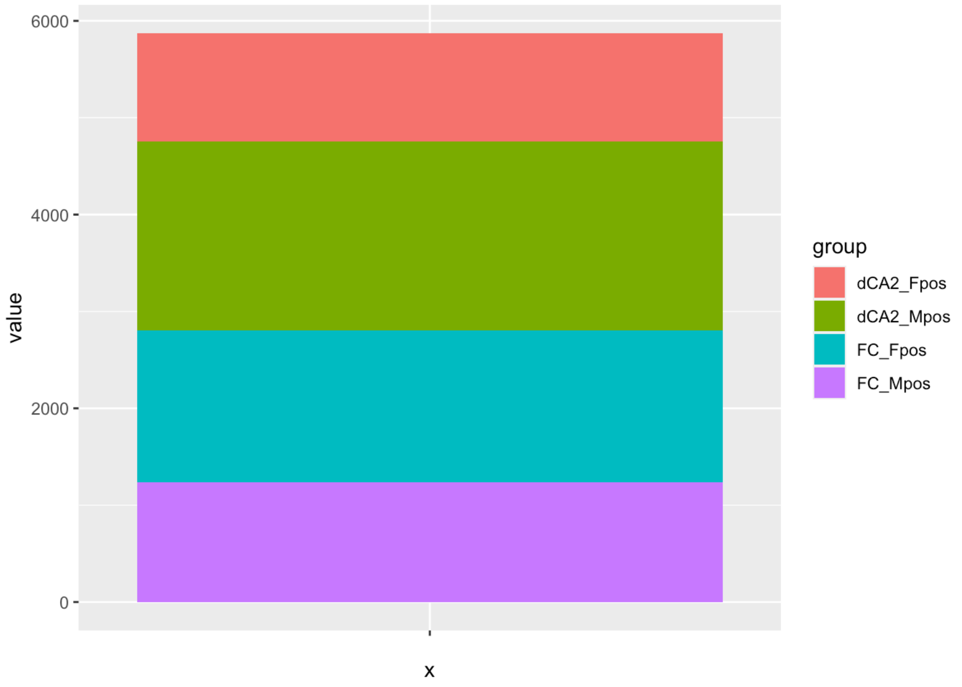


```
pie <- bp + coord_polar("y", start=0)
pie
```

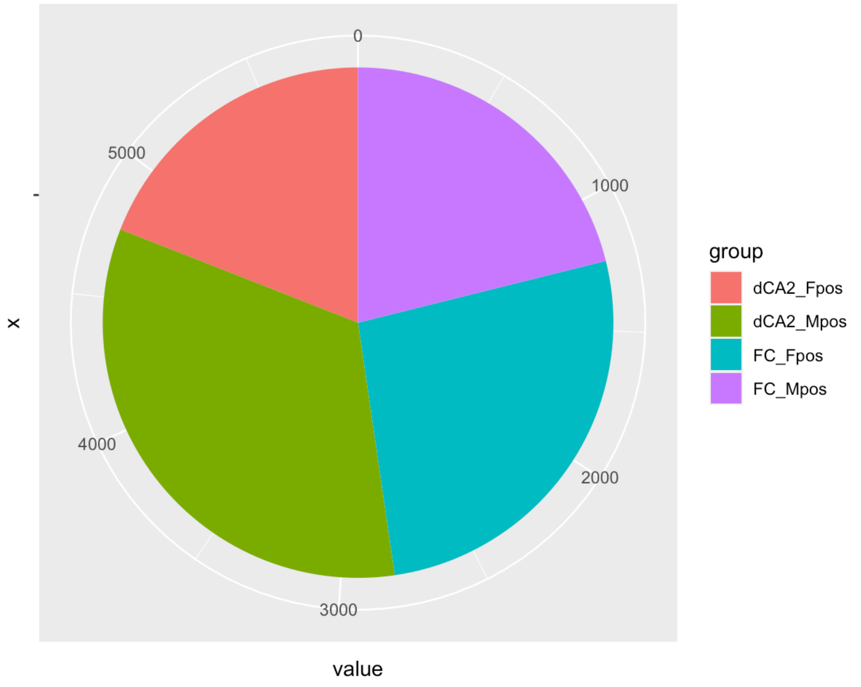


```
ggsave(file="/Users/lees44/R_project/FC_vs_dCA2_comparison_vector_integration_042821/cell_number_group_pie_7cellt
ype.pdf",width=10,height=7)
```

```
# What proportion of cells are in each cluster?
prop.table(table(Idents(dCA2.integrated_clean_7celltype)))
```

```
##
##      Type1      Type2      Type3      Type4      Type5      Type6      Type7
## 0.31154239 0.29724208 0.15083418 0.10810351 0.07286347 0.02996255 0.02945182
```

```
# How does cluster membership vary by group?
table(Idents(dCA2.integrated_clean_7celltype), dCA2.integrated_clean_7celltype$protocol)
```

```
##
##          dCA2_Fpos dCA2_Mpos FC_Fpos FC_Mpos
##   Type1       378       378     652     422
##   Type2       402       900     171     273
##   Type3       268       418     127      73
##   Type4        21       101     285     228
##   Type5         4        49     164     211
##   Type6        38        82      34      22
##   Type7         8        25     129      11
```

pie graph for different cell types

```
#type1
df <-data.frame(
  group = c("dCA2_Fpos", "dCA2_Mpos", "FC_Fpos", "FC_Mpos"),
  value = c(378, 378, 652, 422)
)
head(df)
```

```
##       group value
## 1 dCA2_Fpos   378
## 2 dCA2_Mpos   378
## 3   FC_Fpos   652
## 4   FC_Mpos   422
```

```
bp<- ggplot(df, aes(x="", y=value, fill=group))+
geom_bar(width = 1, stat = "identity")
bp
```

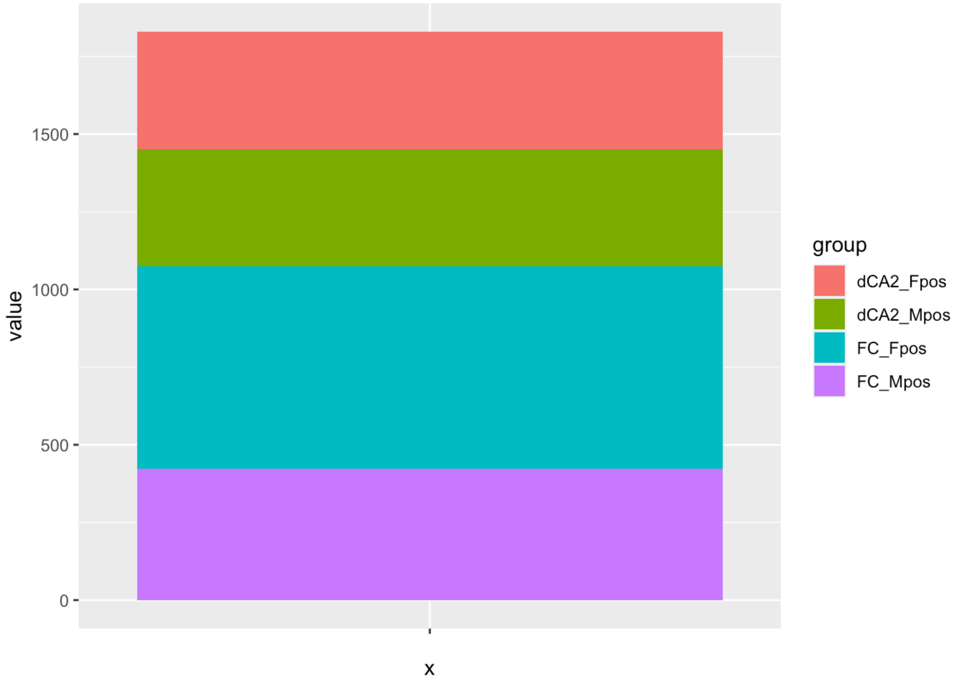


```
pie <- bp + coord_polar("y", start=0)
pie
```

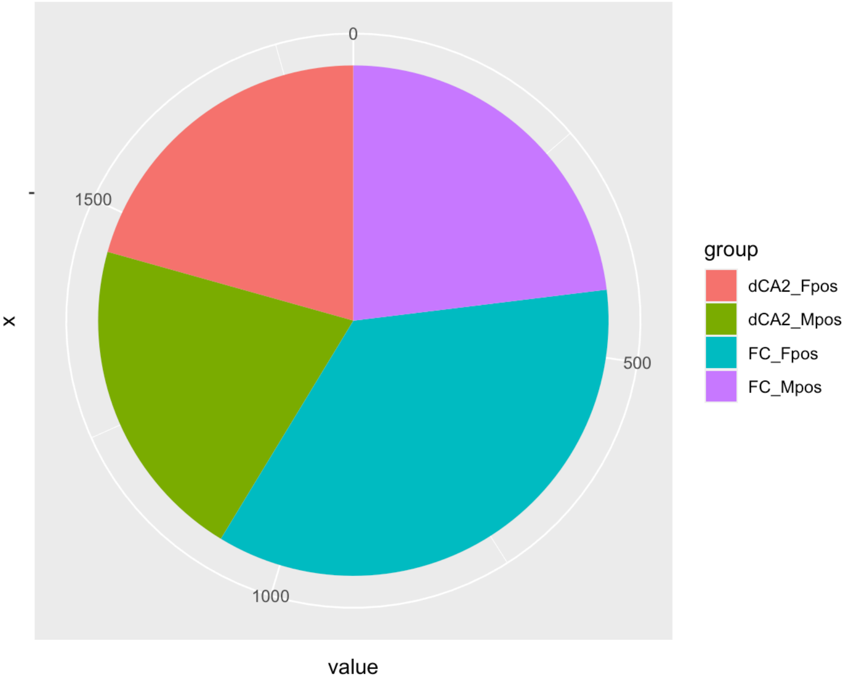


```
ggsave(file="/Users/lees44/R_project/FC_vs_dCA2_comparison_vector_integration_042821/cell_number_group_type1_pie_
7celltype.pdf",width=10,height=7)
```

```
#type2
df <-data.frame(
  group = c("dCA2_Fpos", "dCA2_Mpos", "FC_Fpos", "FC_Mpos"),
  value = c(402, 900, 171, 273)
)
head(df)
```

```
##       group value
## 1 dCA2_Fpos   402
## 2 dCA2_Mpos   900
## 3   FC_Fpos   171
## 4   FC_Mpos   273
```

```
bp<- ggplot(df, aes(x="", y=value, fill=group))+
geom_bar(width = 1, stat = "identity")
bp
```

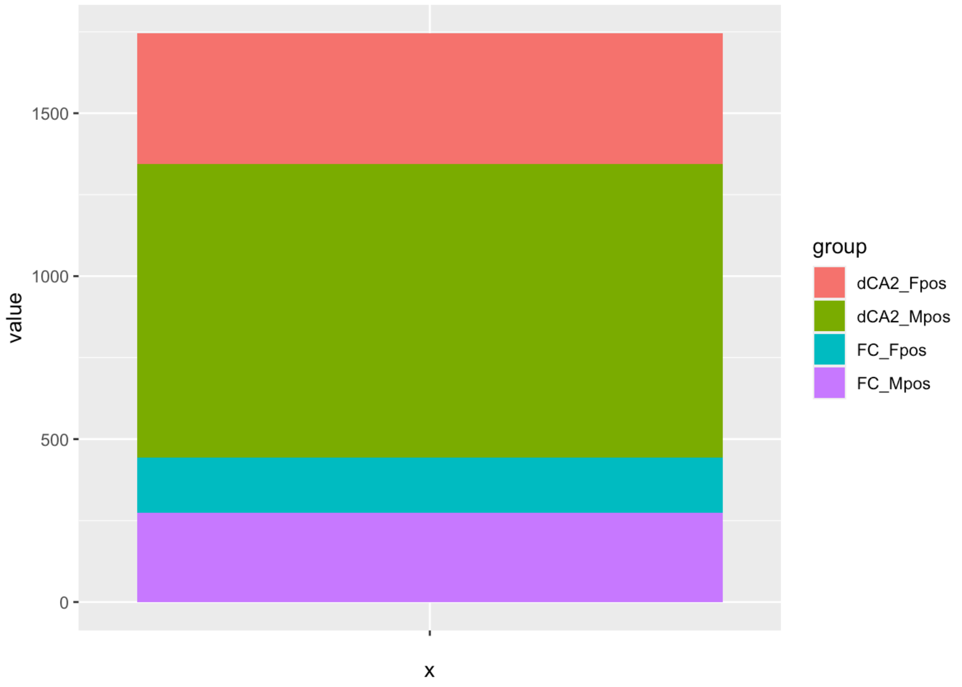


```
pie <- bp + coord_polar("y", start=0)
pie
```

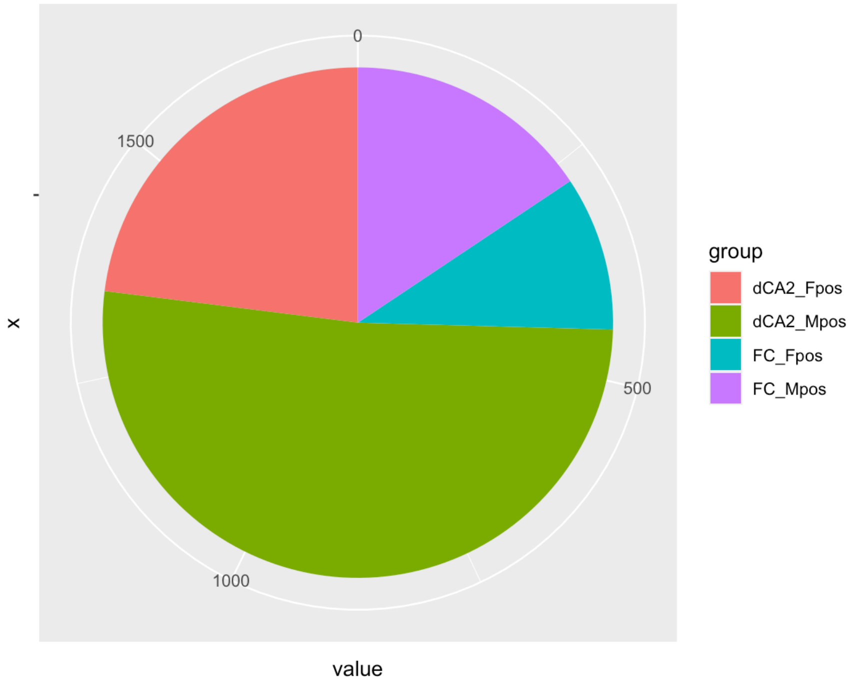


```
ggsave(file="/Users/lees44/R_project/FC_vs_dCA2_comparison_vector_integration_042821/cell_number_group_type2_pie_
7celltype.pdf",width=10,height=7)
```

```
#type3
df <-data.frame(
  group = c("dCA2_Fpos", "dCA2_Mpos", "FC_Fpos", "FC_Mpos"),
  value = c(268, 418, 127, 73)
)
head(df)
```

```
##         group value
## 1 dCA2_Fpos   268
## 2 dCA2_Mpos   418
## 3   FC_Fpos   127
## 4   FC_Mpos    73
```

```
bp<- ggplot(df, aes(x="", y=value, fill=group))+
geom_bar(width = 1, stat = "identity")
bp
```

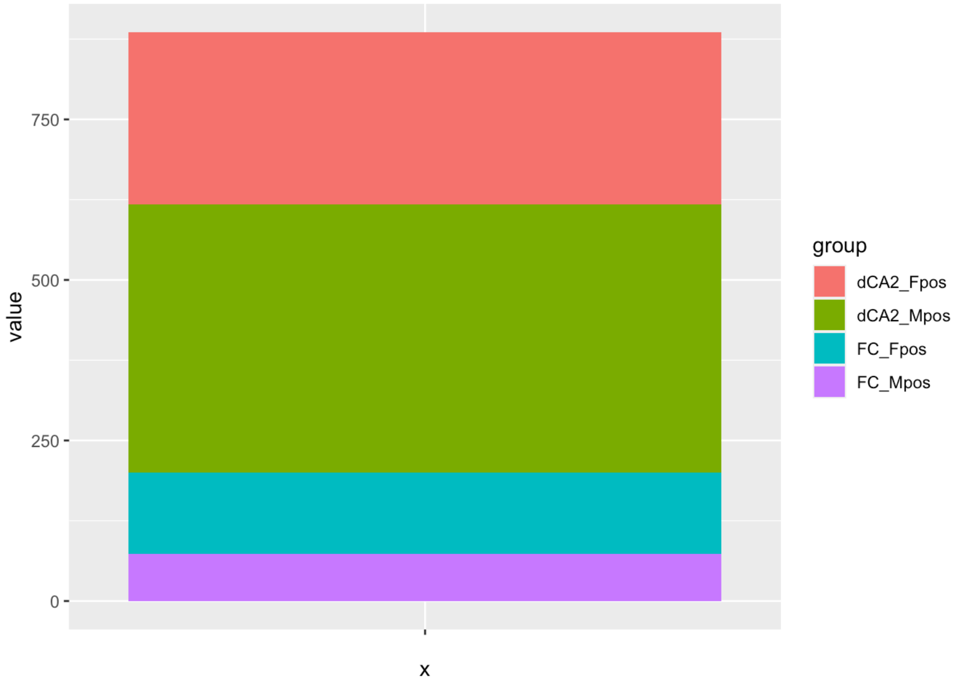


```
pie <- bp + coord_polar("y", start=0)
pie
```

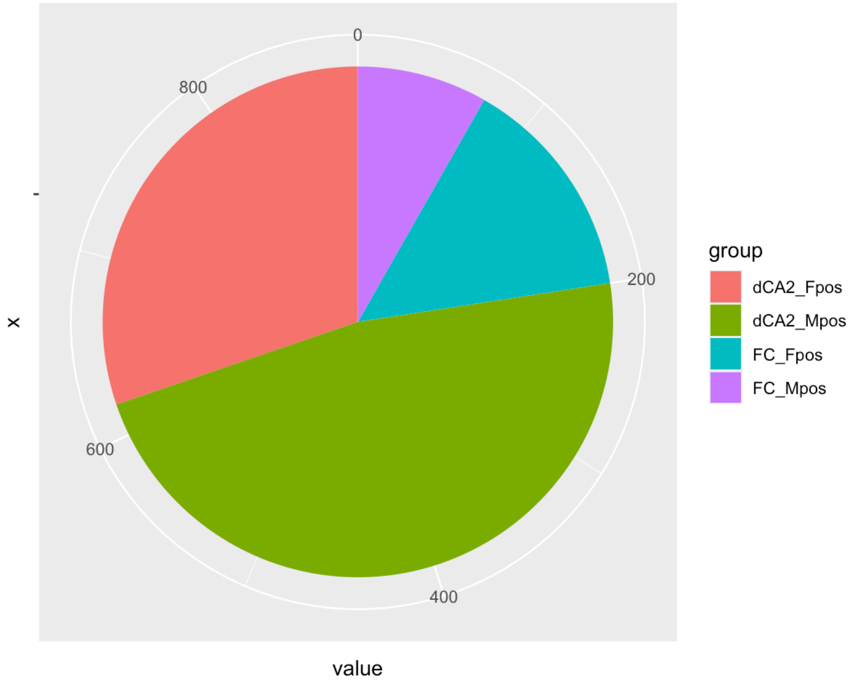

```
ggsave(file="/Users/lees44/R_project/FC_vs_dCA2_comparison_vector_integration_042821/cell_number_group_type3_pie_
7celltype.pdf",width=10,height=7)
```

```
#type4
df <-data.frame(
  group = c("dCA2_Fpos", "dCA2_Mpos", "FC_Fpos", "FC_Mpos"),
  value = c(21, 101, 285, 228)
)
head(df)
```

```
##       group value
## 1 dCA2_Fpos    21
## 2 dCA2_Mpos   101
## 3   FC_Fpos   285
## 4   FC_Mpos   228
```

```
bp<- ggplot(df, aes(x="", y=value, fill=group))+
geom_bar(width = 1, stat = "identity")
bp
```

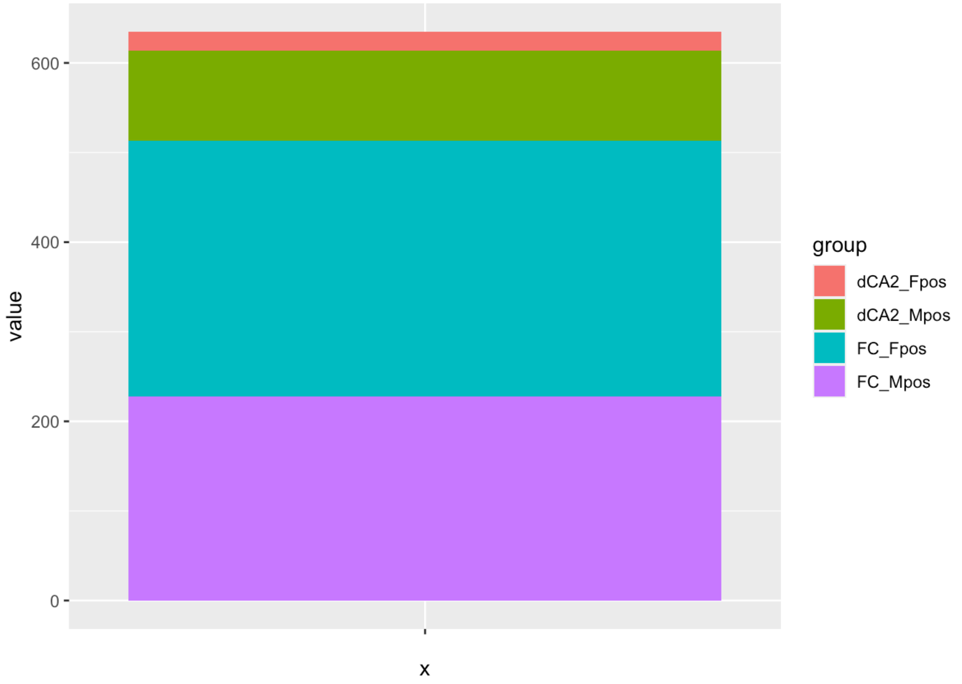


```
pie <- bp + coord_polar("y", start=0)
pie
```

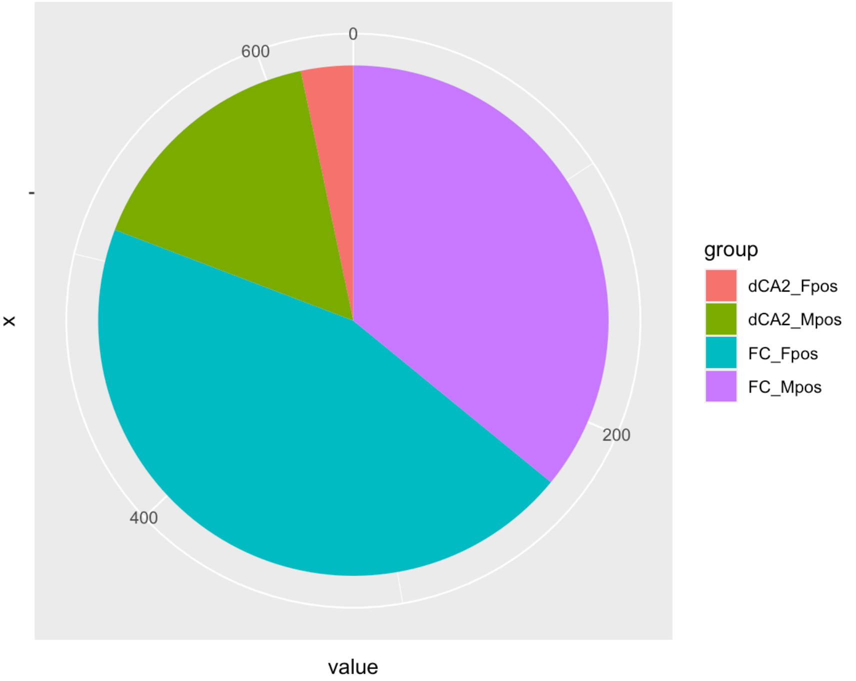


```
ggsave(file="/Users/lees44/R_project/FC_vs_dCA2_comparison_vector_integration_042821/cell_number_group_type4_pie_
7celltype.pdf",width=10,height=7)
```

```
#type5
df <-data.frame(
  group = c("dCA2_Fpos", "dCA2_Mpos", "FC_Fpos", "FC_Mpos"),
  value = c(4, 49, 164, 211)
)
head(df)
```

```
##       group value
## 1 dCA2_Fpos     4
## 2 dCA2_Mpos    49
## 3   FC_Fpos   164
## 4   FC_Mpos   211
```

```
bp<- ggplot(df, aes(x="", y=value, fill=group))+
geom_bar(width = 1, stat = "identity")
bp
```

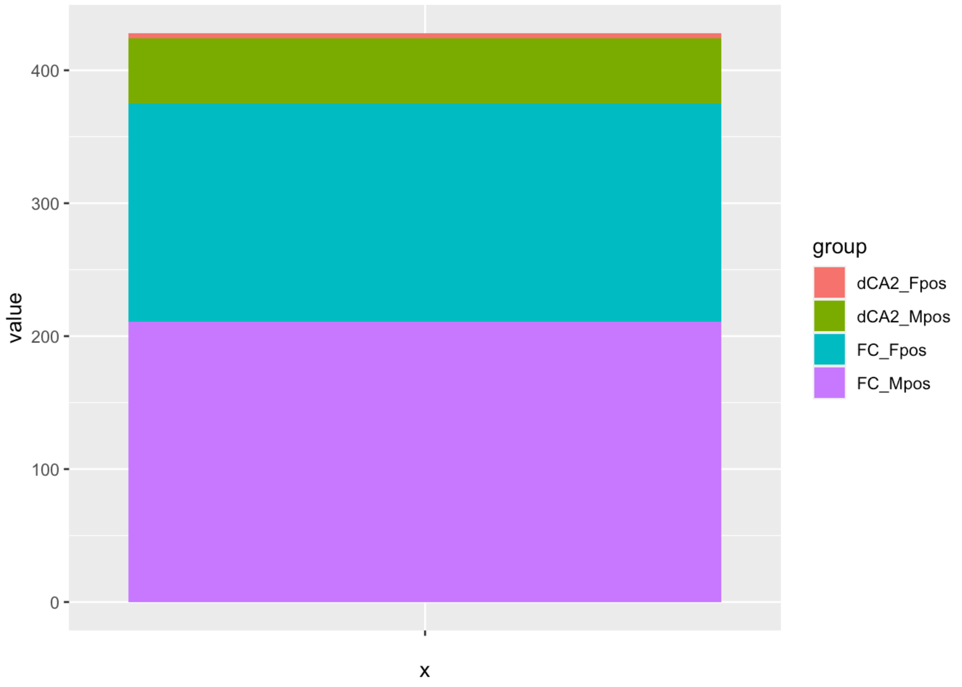


```
pie <- bp + coord_polar("y", start=0)
pie
```

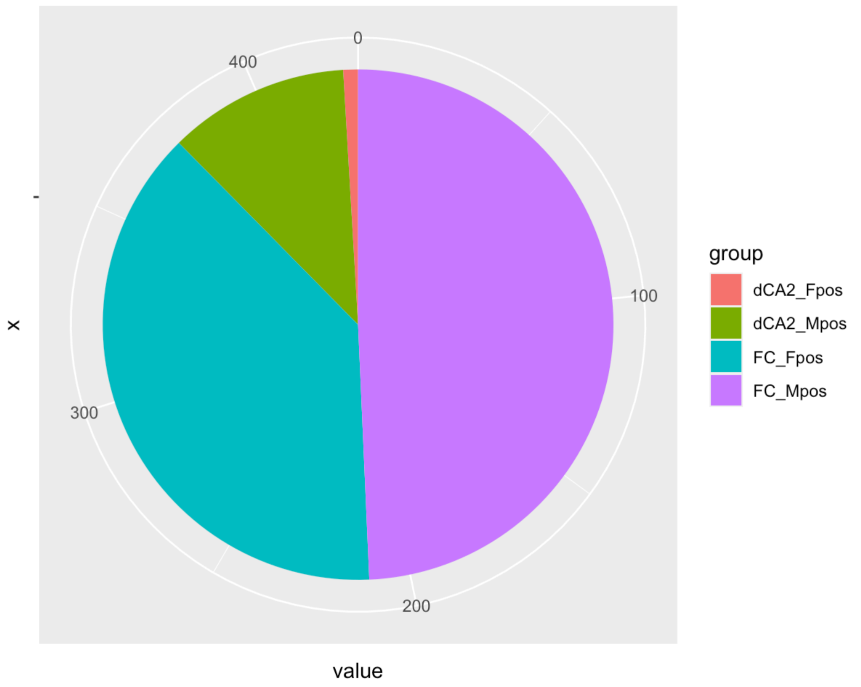


```
ggsave(file="/Users/lees44/R_project/FC_vs_dCA2_comparison_vector_integration_042821/cell_number_group_type3_pie_
7celltype.pdf",width=10,height=7)
```

```
#type6
df <-data.frame(
  group = c("dCA2_Fpos", "dCA2_Mpos", "FC_Fpos", "FC_Mpos"),
  value = c(38, 82, 34, 22)
)
head(df)
```

```
##       group value
## 1 dCA2_Fpos    38
## 2 dCA2_Mpos    82
## 3   FC_Fpos    34
## 4   FC_Mpos    22
```

```
bp<- ggplot(df, aes(x="", y=value, fill=group))+
geom_bar(width = 1, stat = "identity")
bp
```

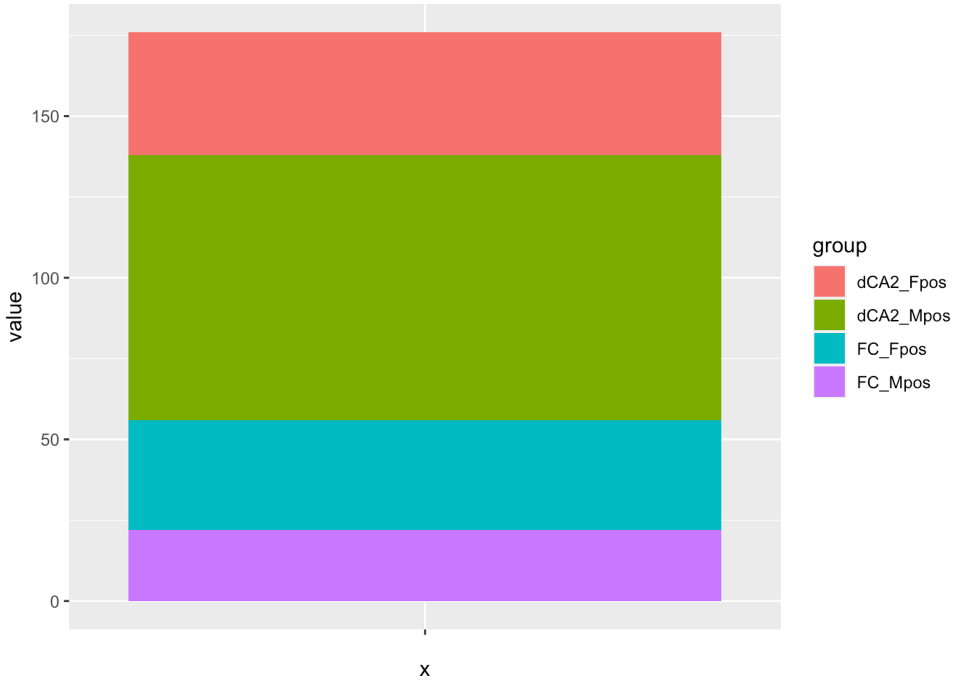


```
pie <- bp + coord_polar("y", start=0)
pie
```

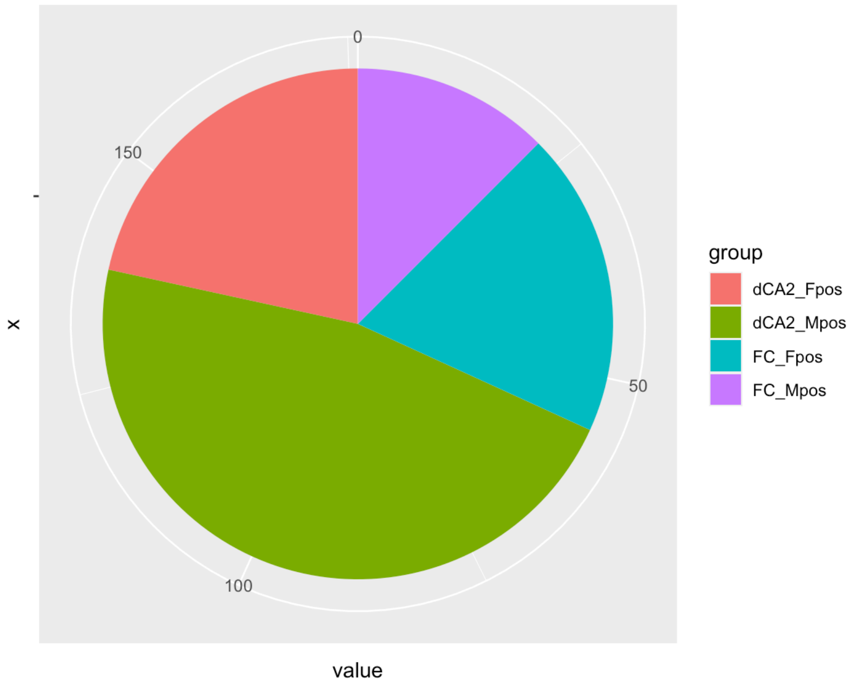

```
ggsave(file="/Users/lees44/R_project/FC_vs_dCA2_comparison_vector_integration_042821/cell_number_group_type6_pie_
7celltype.pdf",width=10,height=7)
```

```
#type7
df <-data.frame(
  group = c("dCA2_Fpos", "dCA2_Mpos", "FC_Fpos", "FC_Mpos"),
  value = c(8, 25, 129, 11)
)
head(df)
```

```
##       group value
## 1 dCA2_Fpos     8
## 2 dCA2_Mpos    25
## 3   FC_Fpos   129
## 4   FC_Mpos    11
```

```
bp<- ggplot(df, aes(x="", y=value, fill=group))+
geom_bar(width = 1, stat = "identity")
bp
```

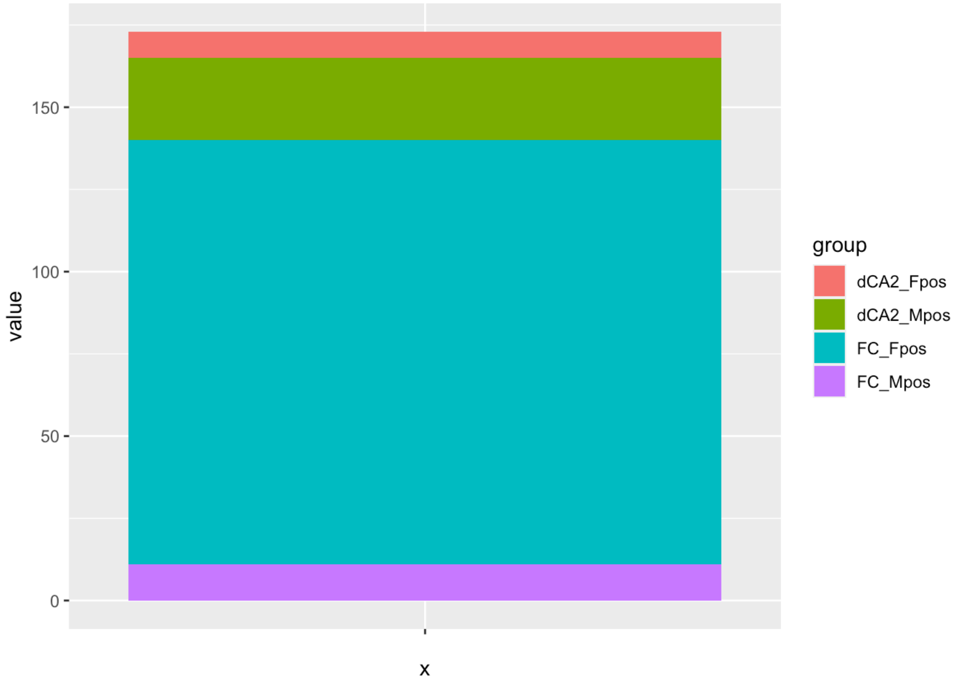


```
pie <- bp + coord_polar("y", start=0)
pie
```

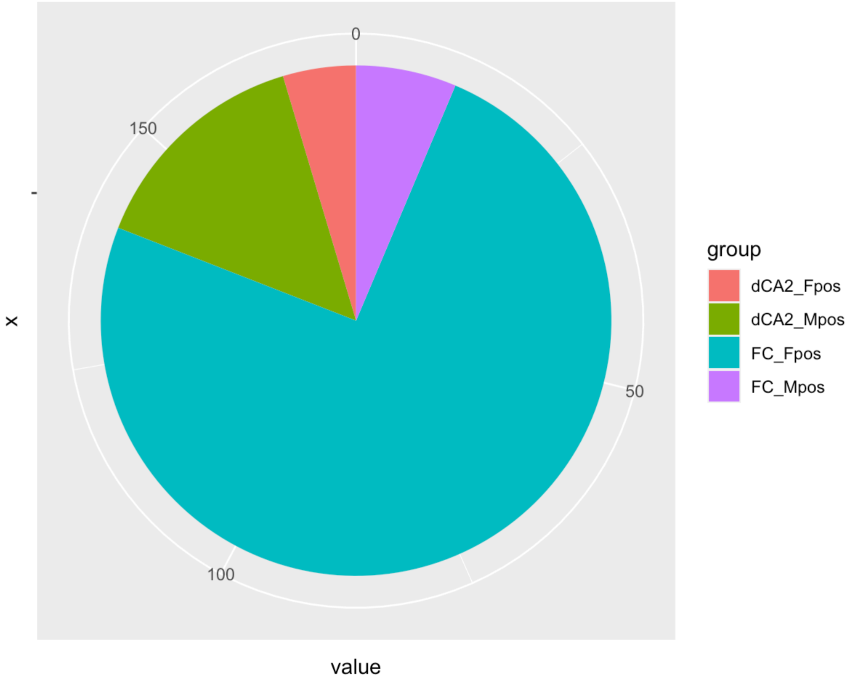


```
ggsave(file="/Users/lees44/R_project/FC_vs_dCA2_comparison_vector_integration_042821/cell_number_group_type7_pie_
7celltype.pdf",width=10,height=7)
```

########percentage within the group 7 cell types ####################

```
#percentage of nuclei dCA2_Fpos
df <-data.frame(
  group = c("Type1", "Type2", "Type3", "Type4", "Type5", "Type6", "Type7"),
  value = c(33.8, 35.9, 23.9, 1.9, 0.4, 3.4, 0.7)
)
head(df)
```

```
##   group value
## 1 Type1  33.8
## 2 Type2  35.9
## 3 Type3  23.9
## 4 Type4   1.9
## 5 Type5   0.4
## 6 Type6   3.4
```

```
bp<- ggplot(df, aes(x="", y=value, fill=group))+
geom_bar(width = 1, stat = "identity")+ ggtitle("dCA2_Fpos")
bp
```

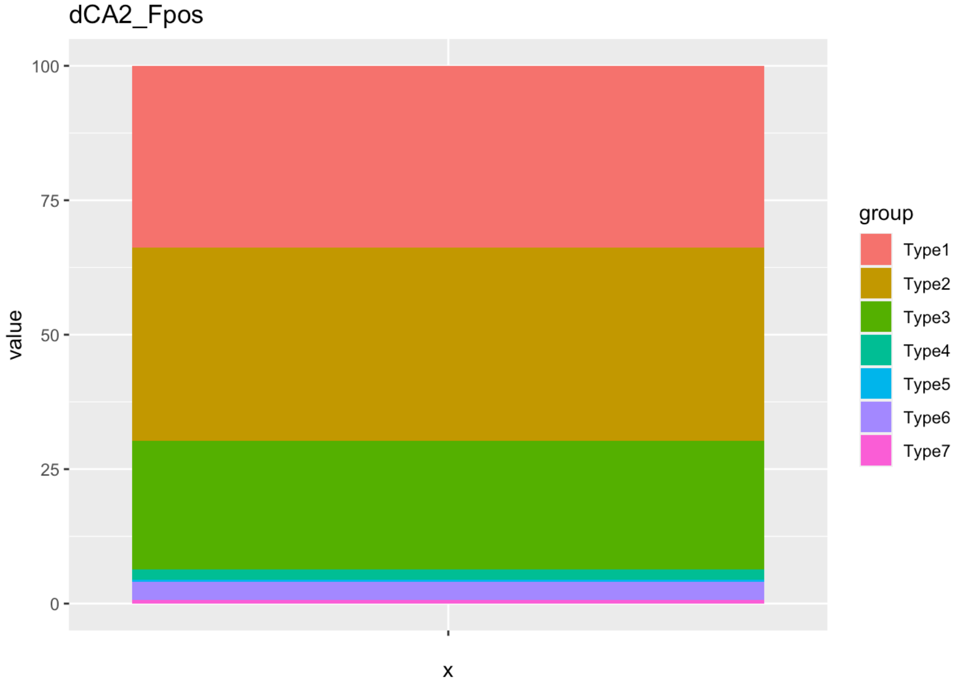


```
pie <- bp + coord_polar("y", start=0)
pie
```

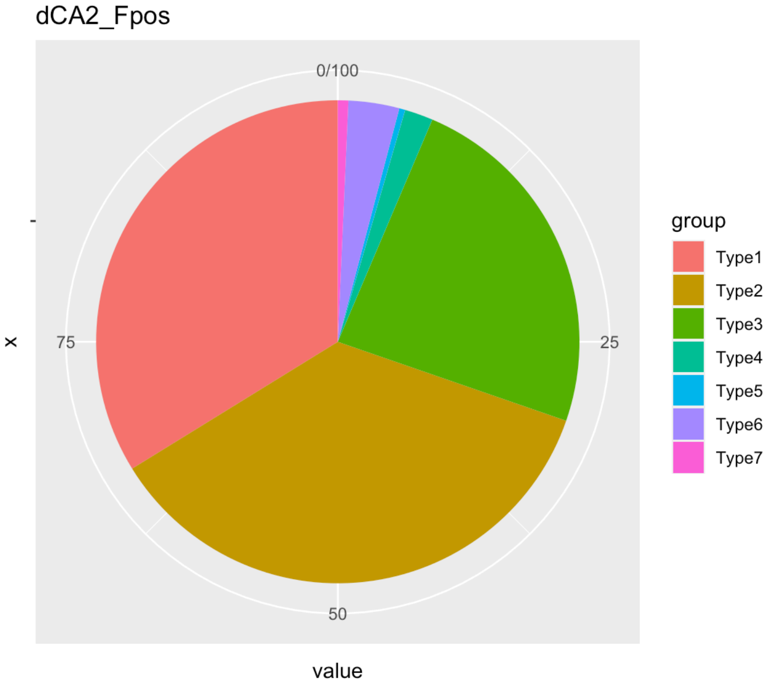


```
ggsave(file="/Users/lees44/R_project/FC_vs_dCA2_comparison_vector_integration_042821/cell_type_percentage_dCA2_Fp
os_7celltype.pdf",width=10,height=7)
```

```
#percentage of nuclei dCA2_Mpos
df <-data.frame(
  group = c("Type1", "Type2", "Type3", "Type4", "Type5", "Type6", "Type7"),
  value = c(19.4, 46.1, 21.1, 5.1, 2.5, 4.1, 1.3)
)
head(df)
```

```
##   group value
## 1 Type1  19.4
## 2 Type2  46.1
## 3 Type3  21.1
## 4 Type4   5.1
## 5 Type5   2.5
## 6 Type6   4.1
```

```
bp<- ggplot(df, aes(x="", y=value, fill=group))+
geom_bar(width = 1, stat = "identity")+ ggtitle("dCA2_Mpos")
bp
```

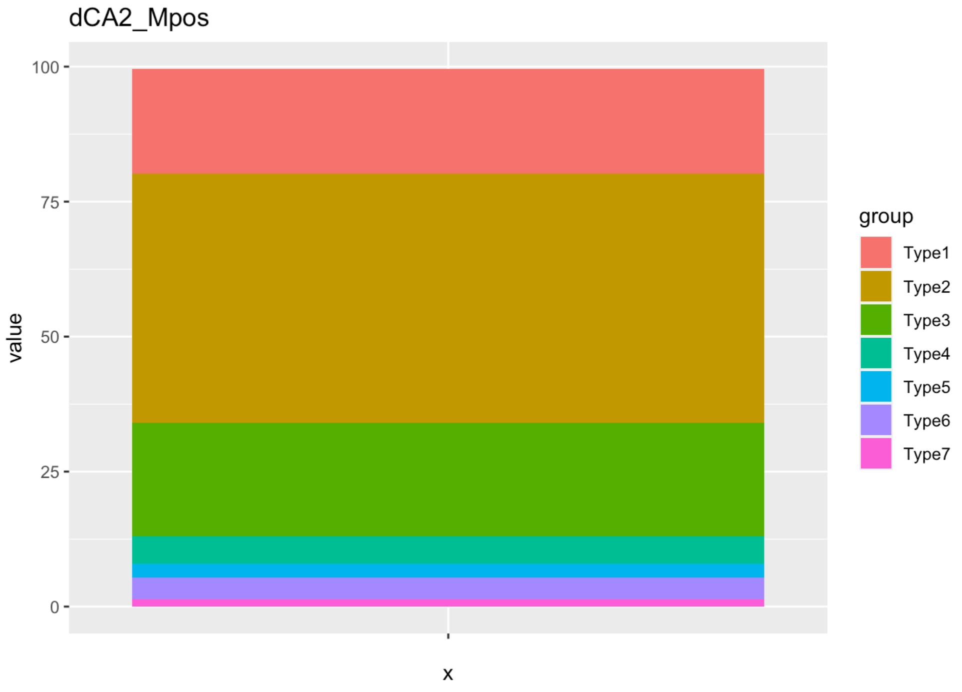


```
pie <- bp + coord_polar("y", start=0)
pie
```

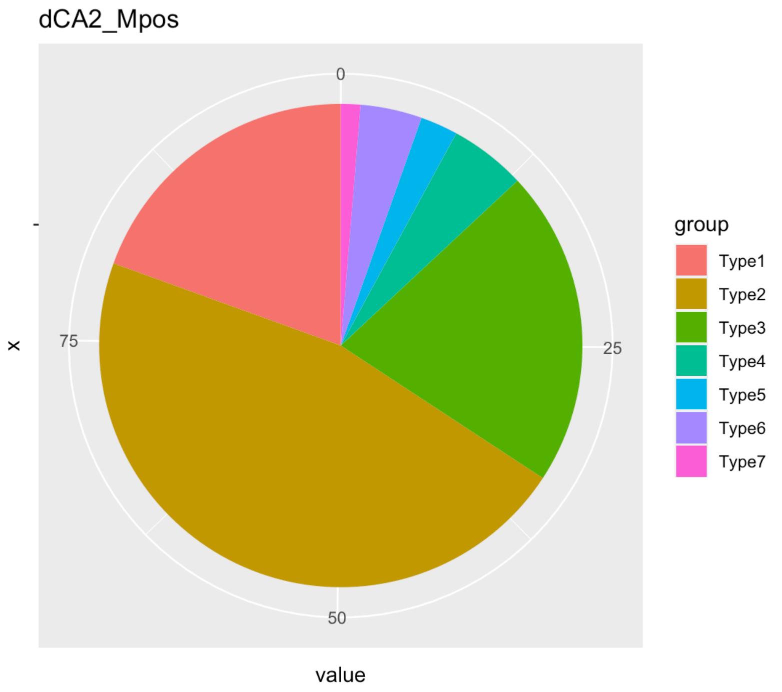


```
ggsave(file="/Users/lees44/R_project/FC_vs_dCA2_comparison_vector_integration_042821/cell_type_percentage_dCA2_Mpos_7celltype.pdf",width=10,height=7)
```

```
#percentage of nuclei FC_Fpos
df <-data.frame(
  group = c("Type1", "Type2", "Type3", "Type4", "Type5", "Type6", "Type7"),
  value = c(41.7, 10.9, 8.1, 18.2, 10.5, 2.2, 8.3)
)
head(df)
```

```
##   group value
## 1 Type1  41.7
## 2 Type2  10.9
## 3 Type3   8.1
## 4 Type4  18.2
## 5 Type5  10.5
## 6 Type6   2.2
```

```
bp<- ggplot(df, aes(x="", y=value, fill=group))+
geom_bar(width = 1, stat = "identity")+ ggtitle("FC_Fpos")
bp
```

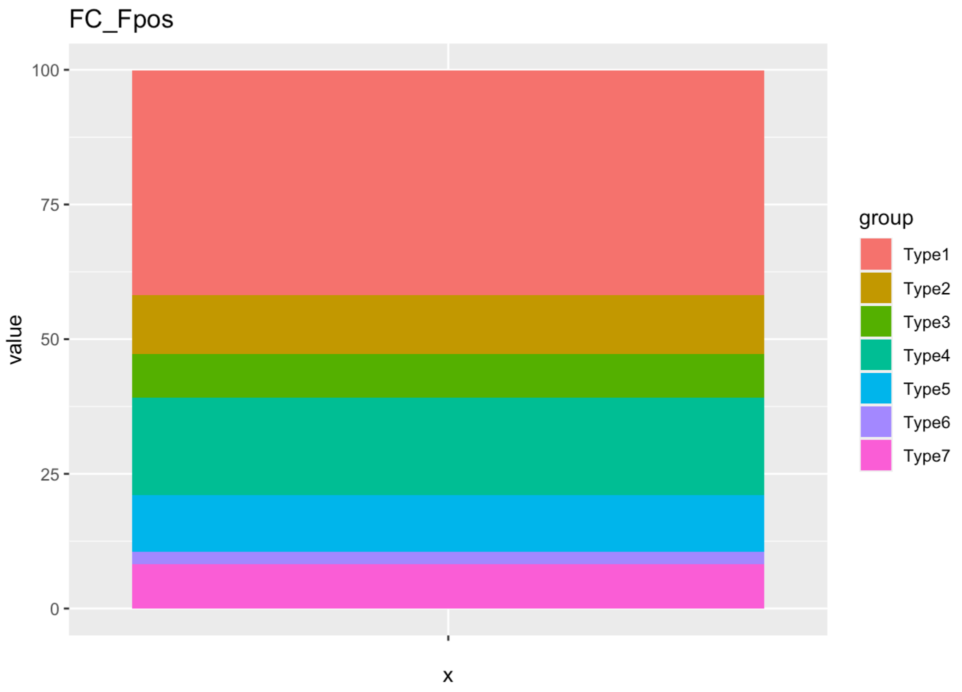


```
pie <- bp + coord_polar("y", start=0)
pie
```

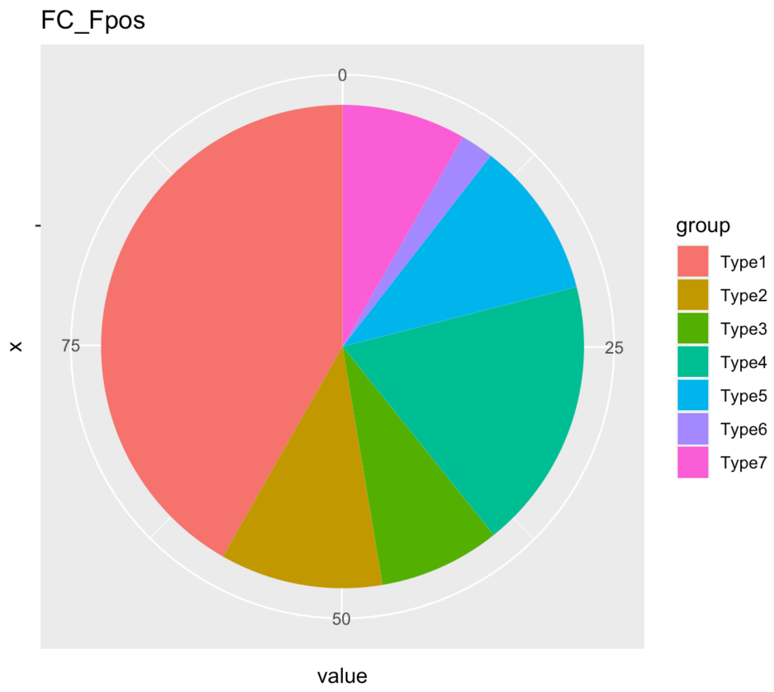


```
ggsave(file="/Users/lees44/R_project/FC_vs_dCA2_comparison_vector_integration_042821/cell_type_percentage_FC_Fpos
_7celltype.pdf",width=10,height=7)
```

```
#percentage of nuclei FC_Mpos
df <-data.frame(
  group = c("Type1", "Type2", "Type3", "Type4", "Type5", "Type6", "Type7"),
  value = c(41.7,10.9, 8.1, 18.2, 10.5, 2.2, 8.3)
)
head(df)
```

```
##   group value
## 1 Type1  41.7
## 2 Type2  10.9
## 3 Type3   8.1
## 4 Type4  18.2
## 5 Type5  10.5
## 6 Type6   2.2
```

```
bp<- ggplot(df, aes(x="", y=value, fill=group))+
geom_bar(width = 1, stat = "identity")+ ggtitle("FC_Mpos")
bp
```

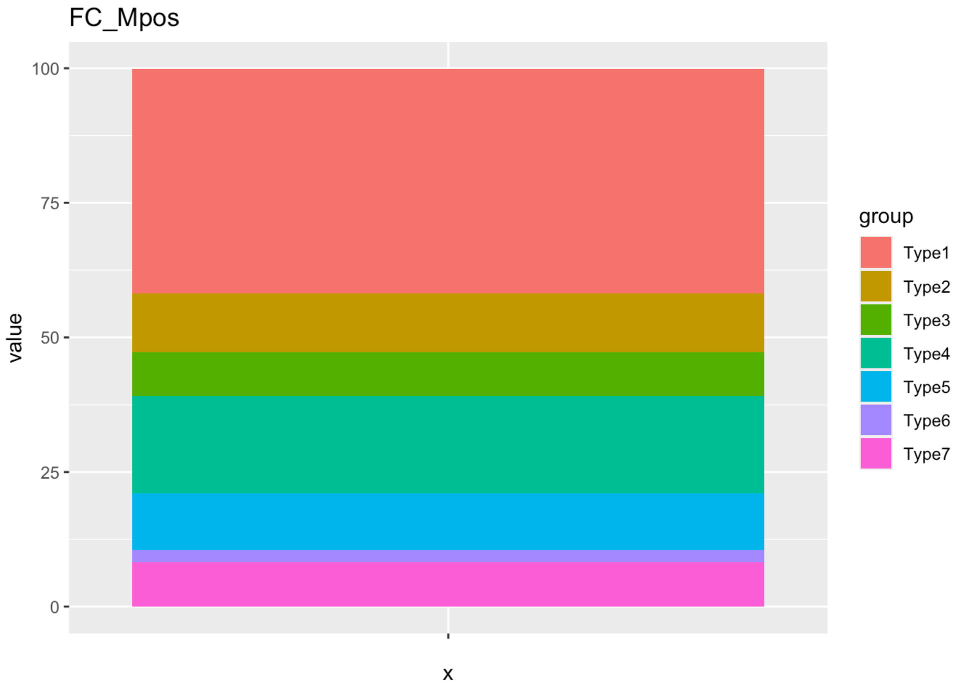


```
pie <- bp + coord_polar("y", start=0)
pie
```

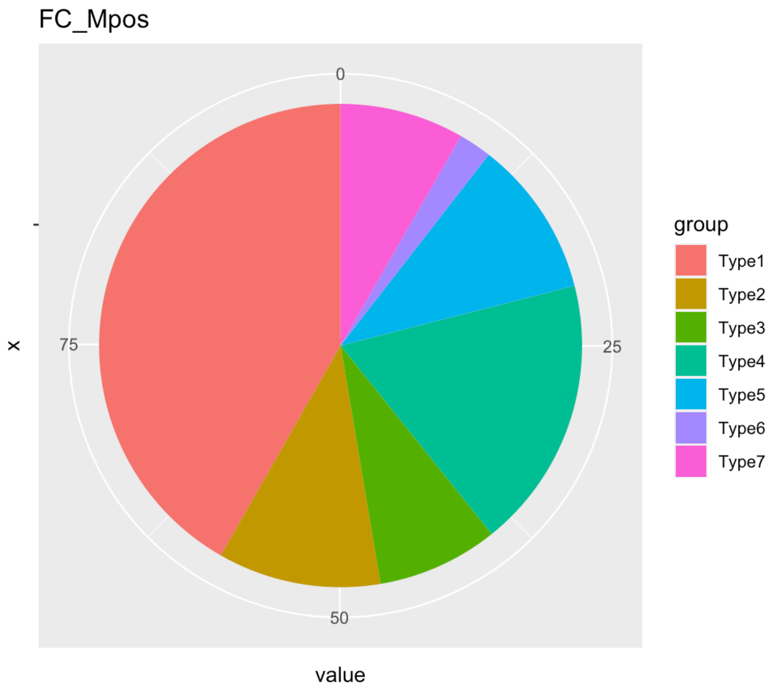


```
ggsave(file="/Users/lees44/R_project/FC_vs_dCA2_comparison_vector_integration_042821/cell_type_percentage_FC_Mpos
_7celltype.pdf",width=10,height=7)
```

```
#percentage of nuclei dCA2
df <-data.frame(
  group = c("Type1", "Type2", "Type3", "Type4", "Type5", "Type6", "Type7"),
  value = c(24.6, 42.4, 22.3, 4, 1.7, 3.9, 1.1)
)
head(df)
```

```
##   group value
## 1 Type1  24.6
## 2 Type2  42.4
## 3 Type3  22.3
## 4 Type4   4.0
## 5 Type5   1.7
## 6 Type6   3.9
```

```
bp<- ggplot(df, aes(x="", y=value, fill=group))+
geom_bar(width = 1, stat = "identity")+ ggtitle("dCA2")
bp
```

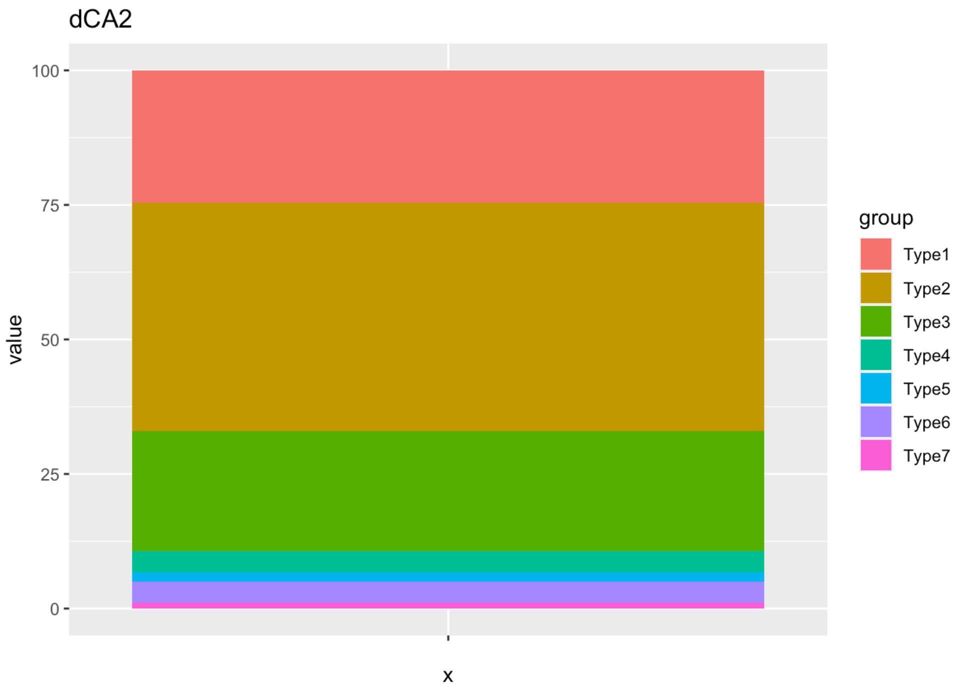


```
pie <- bp + coord_polar("y", start=0)
pie
```

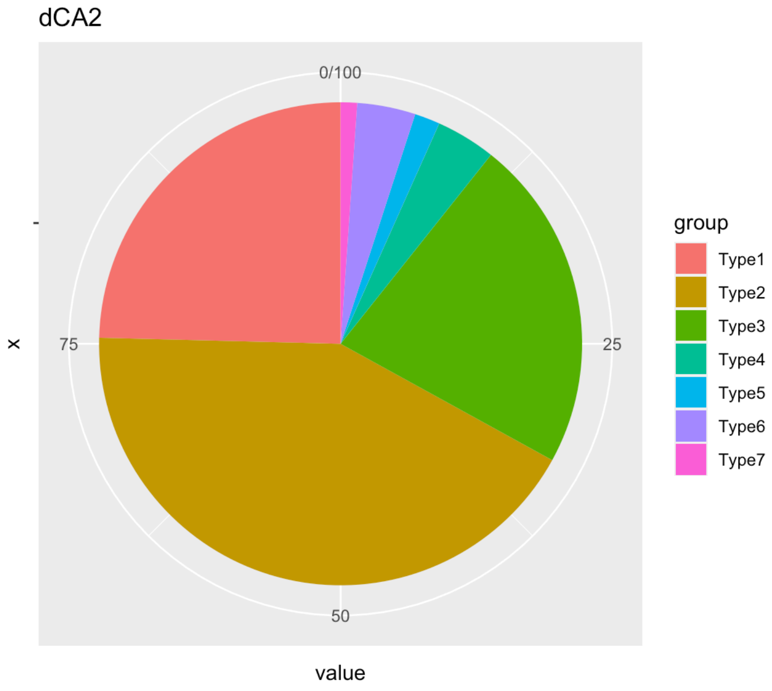


```
ggsave(file="/Users/lees44/R_project/FC_vs_dCA2_comparison_vector_integration_042821/cell_type_percentage_dCA2_7c
elltype.pdf",width=10,height=7)
```

```
#percentage of nuclei FC
df <-data.frame(
  group = c("Type1", "Type2", "Type3", "Type4", "Type5", "Type6", "Type7"),
  value = c(38.3, 16, 7.1, 18, 13.4, 2, 5)
)
head(df)
```

```
##   group value
## 1 Type1  38.3
## 2 Type2  16.0
## 3 Type3   7.1
## 4 Type4  18.0
## 5 Type5  13.4
## 6 Type6   2.0
```

```
bp<- ggplot(df, aes(x="", y=value, fill=group))+
geom_bar(width = 1, stat = "identity")+ ggtitle("FC")
bp
```

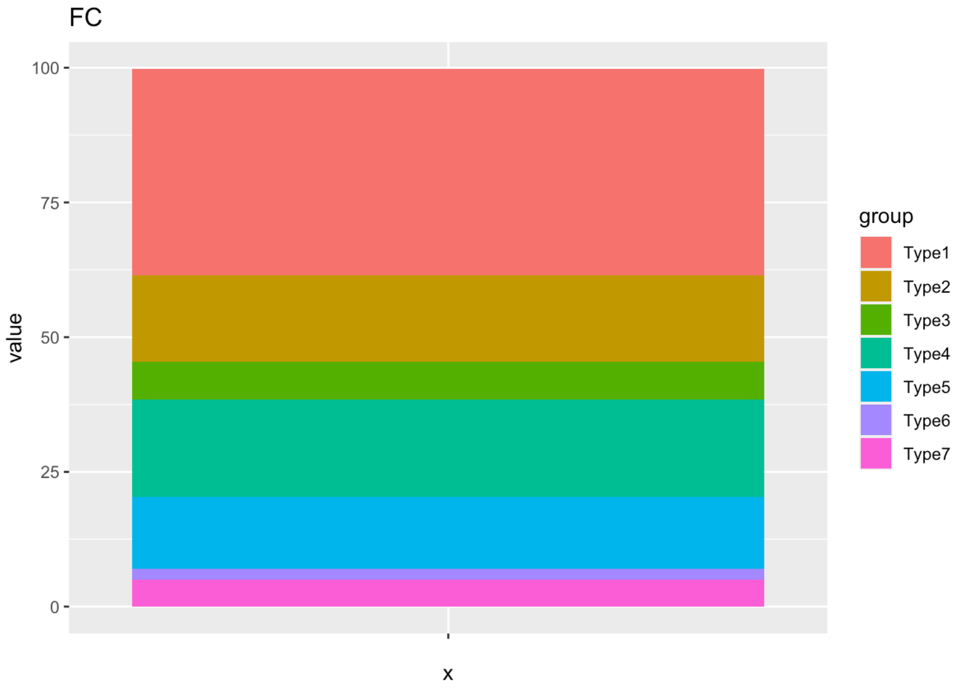


```
pie <- bp + coord_polar("y", start=0)
pie
```

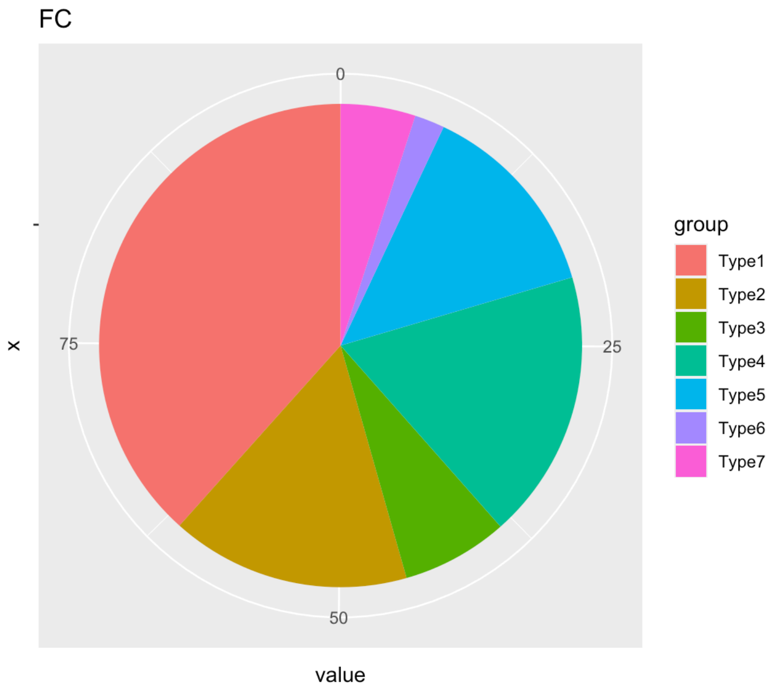


```
ggsave(file="/Users/lees44/R_project/FC_vs_dCA2_comparison_vector_integration_042821/cell_type_percentage_FC_7cel
ltpe.pdf",width=10,height=7)
```

###########################################UMI plot for 7 cell types 08.24.21##################################

UMI plot for each cluster Identifying conserved markers for each clusters for 7 cell types

```
library("rJava")
```

```
library("xlsx")
```

```
library("metap")
```

```
#divide by avarage read number
length(Type1_barcode)/10000
```

```
## [1] 0.183
```

```
length(Type2_barcode)/10000
```

```
## [1] 0.1746
```

```
length(Type3_barcode)/10000
```

```
## [1] 0.0886
```

```
length(Type4_barcode)/10000
```

```
## [1] 0.0635
```

```
length(Type5_barcode)/10000
```

```
## [1] 0.0428
```

```
length(Type6_barcode)/10000
```

```
## [1] 0.0176
```

```
length(Type7_barcode)/10000
```

```
## [1] 0.0173
```

```
type<-numeric()
for (i in 1:dim(dCA2.integrated_clean_7celltype@meta.data)[1]){
if(rownames(dCA2.integrated_clean_7celltype@meta.data)[i] %in% Type1_barcode){type[i]<-"Type1"}
else if(rownames(dCA2.integrated_clean_7celltype@meta.data)[i] %in% Type2_barcode){type[i]<-"Type2"}
else if(rownames(dCA2.integrated_clean_7celltype@meta.data)[i] %in% Type3_barcode){type[i]<-"Type3"}
else if(rownames(dCA2.integrated_clean_7celltype@meta.data)[i] %in% Type4_barcode){type[i]<-"Type4"}
else if(rownames(dCA2.integrated_clean_7celltype@meta.data)[i] %in% Type5_barcode){type[i]<-"Type5"}
else if(rownames(dCA2.integrated_clean_7celltype@meta.data)[i] %in% Type6_barcode){type[i]<-"Type6"}
else if(rownames(dCA2.integrated_clean_7celltype@meta.data)[i] %in% Type7_barcode){type[i]<-"Type7"}
else{type[i]<-"NA"}
}
dCA2.integrated_clean_7celltype@meta.data$type<-type
```

```
colors<-c("Type1"="#19647e","Type2"="#ffc857","Type3"="#9B0C1E","Type4"="#4b3f72","Type5"="#ffc857","Type6"="#676
833", "Type7"="#19647e")
```

```
subset_meta<-subset(dCA2.integrated@meta.data,type=="Type1"|type=="Type2"|type=="Type3"|type=="Type4"|type=="Type
5"|type=="Type6"|type=="Type7")
ggplot(subset_meta,aes_string(x="type",y="nCount_RNA",fill="type"))+geom_violin(scale = "width")+scale_fill_manua
l(values=colors)+
stat_summary(fun=median, geom="point", size=0.6, color="red")+ylab("UMI")+
theme(axis.title.x=element_blank(),axis.text.y=element_text(size=10),axis.title.y=element_text(size=10,angle=0,fa
ce="bold",margin = margin(t = 10, r = 8, b = 0, l = 10),vjust=0.5),axis.text.x=element_text(size=10,face="bold"),
axis.title=element_text(size=15,face="bold"),panel.grid.major = element_blank(), panel.grid.minor = element_blan
k(),panel.background = element_blank(), axis.line = element_line(colour = "black"),legend.position="none",plot.ma
rgin = unit(c(0,
0,0, 0), "cm"))
```

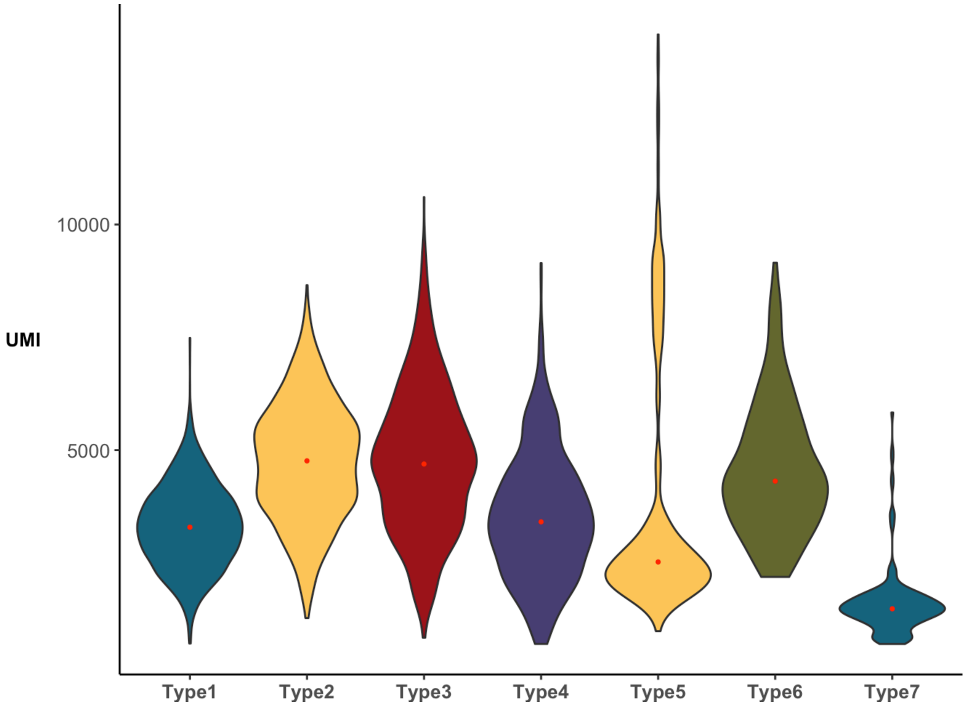


```
ggsave(file="/Users/lees44/R_project/FC_vs_dCA2_comparison_vector_integration_042821/cell_type_UMI_count_clean_7c
elltype.pdf",width=10,height=7)
```

#########violin plot for 7 cell types with CA2 markers 08.24.21 ####################

```
Cell_type<-rev(c("Type7", "Type4", "Type5", "Type3", "Type6", "Type1", "Type2"))
gene_list<-c("Avpr1b" ,"Amigo2","Rgs14", "Map3k15", "Pcp4")
```

```
Cell_number <- NULL
for (i in 1:length(Cell_type)){
                temp1 <- data.frame(type=factor(rep(Cell_type[i],sum(Cell_type[i]==Idents(dCA2.integrated)))))
                temp2 <- as.data.frame(dCA2.integrated_clean_7celltype@assays$RNA@data[gene_list,Idents(dCA2.inte
grated_clean_7celltype)==Cell_type[i]])
                temp3 <- cbind(temp1,t(temp2))
                Cell_number <- rbind(temp3,Cell_number)
}
rownames(Cell_number) <- NULL
```

```
colors<-c("#ffc857","#ffc857","#ffc857","#9B0C1E","#9B0C1E","#9B0C1E","#9B0C1E")
```

```
for (k in 1:length(gene_list))
    {
if (k==length(gene_list)){
assign(paste("P",k,sep=""),ggplot(Cell_number,aes_string(x="type",y=gene_list[k],fill="type"))+geom_violin(scale
= "width")+scale_fill_manual(values=colors)
+stat_summary(fun=median, geom="point", size=0.6, color="red")+ylab(gene_list[k])+ theme(axis.title.x=element_bla
nk(),
axis.text.y=element_text(size=5),axis.title.y=element_text(size=6,angle=0,face="bold",margin = margin(t = 10, r =
8, b = 0, l = 10),vjust=0.5),axis.text.x=element_text(size=5,face="bold")
,axis.title=element_text(size=4,face="bold"),panel.grid.major = element_blank(), panel.grid.minor = element_blan
k(),
panel.background = element_blank(), axis.line = element_line(colour = "black"),legend.position="none",plot.margin
= unit(c(0, 0,0, 0), "cm")))}
else{
assign(paste("P",k,sep=""),ggplot(Cell_number,aes_string(x="type",y=gene_list[k],fill="type"))+geom_violin(scale
= "width")+scale_fill_manual(values=colors)
+stat_summary(fun=median, geom="point", size=0.6, color="red")+ylab(gene_list[k])+ theme(axis.title.x=element_bla
nk(),
axis.text.x=element_blank(),axis.text.y=element_text(size=5),axis.title.y=element_text(size=6,angle=0,face="bold",
margin = margin(t = 10, r = 8, b = 0, l = 10),vjust=0.5),axis.ticks.x=element_blank(),axis.text=element_text(size
=5)
,axis.title=element_text(size=4,face="bold"),panel.grid.major = element_blank(), panel.grid.minor = element_blan
k(),
panel.background = element_blank(), axis.line = element_line(colour = "black"),legend.position="none",plot.margin
= unit(c(0, 0,0, 0), "cm")))}

}
```

```
P1
```

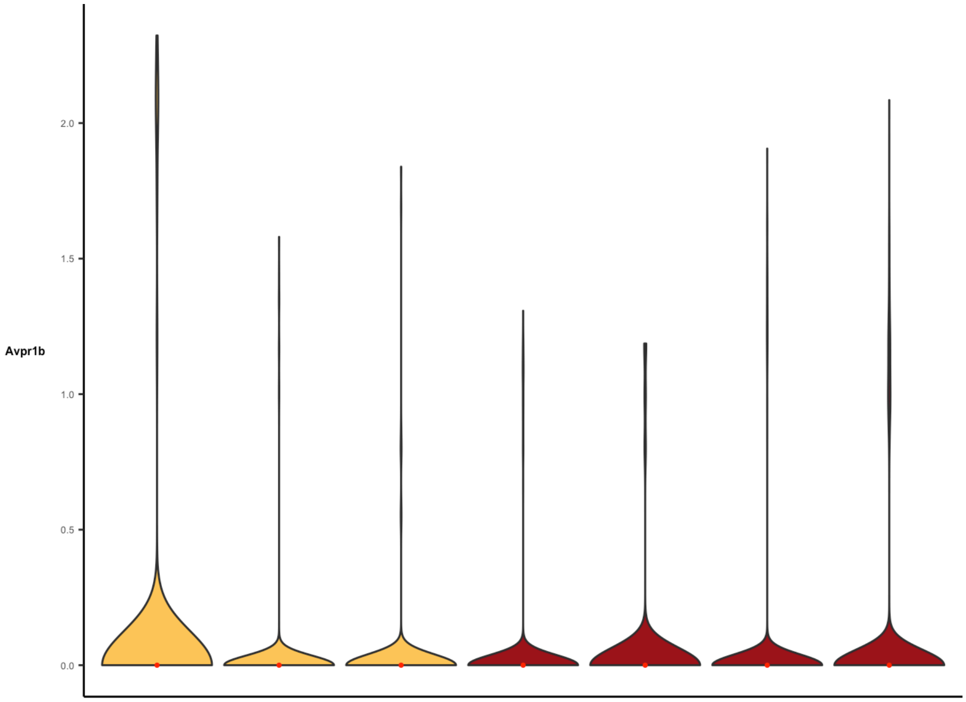


```
P2
```

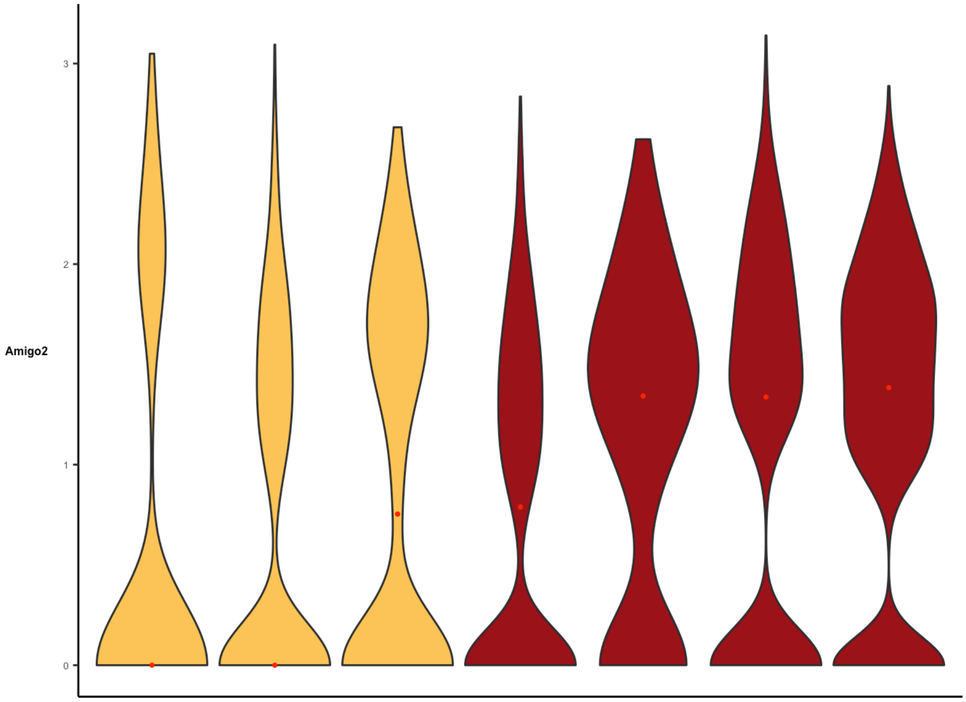


```
P3
```

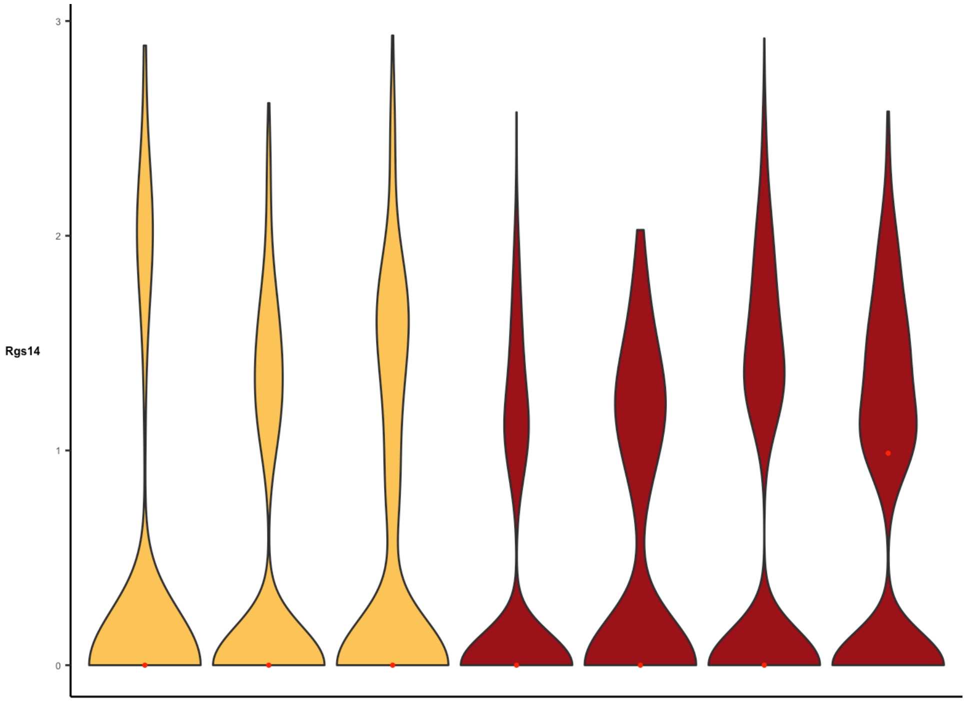


```
P4
```

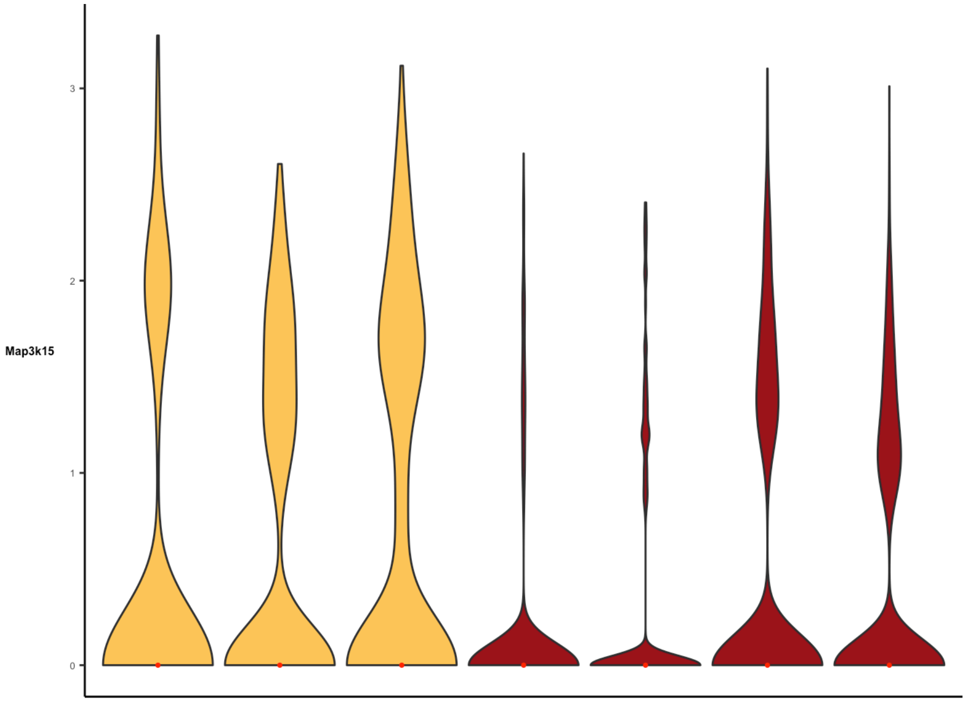
Map3k15
0
1
2
3

```
P5
```

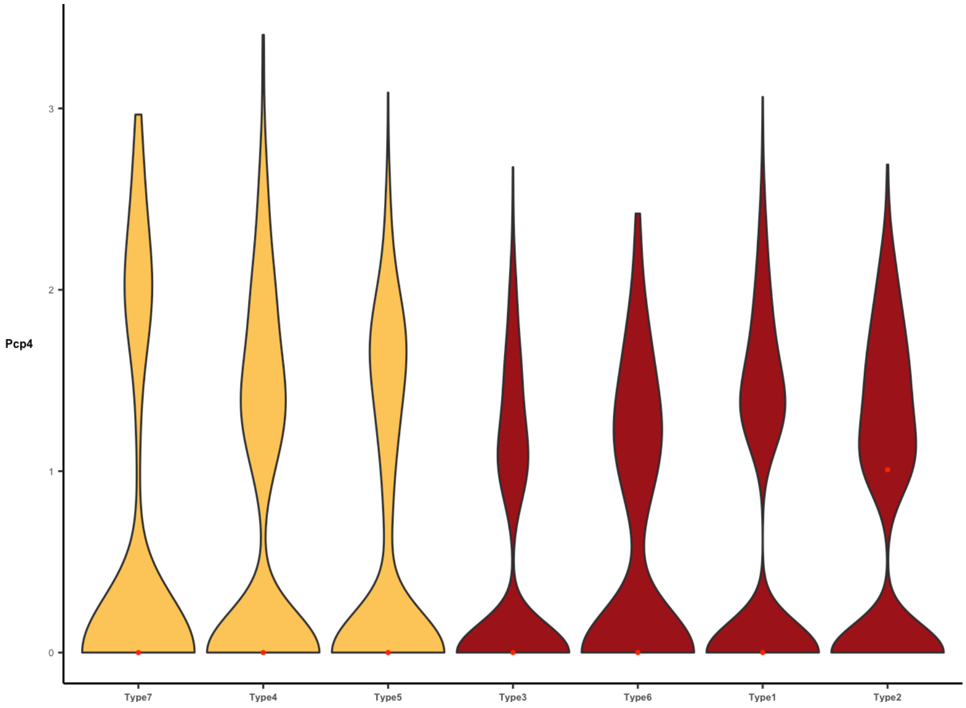
Pcp4
0
1
2
3
Type7
Type4
Type5
Type3
Type6
Type1
Type2

```
merge<-list()
for (i in length(gene_list):1){
       if (length(merge)==0){
        merge<-ggplotGrob(eval(parse(text=paste("P",i,sep = ""))))}else{merge<-rbind(ggplotGrob(eval(parse(text=p
aste("P",i,sep = "")))),merge,size = "last")}}

 #all genes
pdf(paste("/Users/lees44/R_project/FC_vs_dCA2_comparison_vector_integration_042821/Violin_plot_dCA2_markers_7cell
type.pdf",sep=""),height=6, width=50 , paper = "letter")


grid.newpage()
grid.draw(merge)
```

#########violin plot for 7 cell types with FC vs dCA2 markers 08.24.21 #####################

```
Cell_type<-rev(c("Type7", "Type4", "Type5", "Type3", "Type6", "Type1", "Type2"))
gene_list<-c("Mef2c","Zfp536", "Tac1", "Scn1a", "Cpne7", "Apoe", "Ptgds", "Ttr")
```

```
Cell_number <- NULL
for (i in 1:length(Cell_type)){
                temp1 <- data.frame(type=factor(rep(Cell_type[i],sum(Cell_type[i]==Idents(dCA2.integrated)))))
                temp2 <- as.data.frame(dCA2.integrated_clean_7celltype@assays$RNA@data[gene_list,Idents(dCA2.inte
grated_clean_7celltype)==Cell_type[i]])
                temp3 <- cbind(temp1,t(temp2))
                Cell_number <- rbind(temp3,Cell_number)
}
rownames(Cell_number) <- NULL
```

```
colors<-c("#ffc857","#ffc857","#ffc857","#9B0C1E","#9B0C1E","#9B0C1E","#9B0C1E")
```

```
for (k in 1:length(gene_list))
    {
if (k==length(gene_list)){
assign(paste("P",k,sep=""),ggplot(Cell_number,aes_string(x="type",y=gene_list[k],fill="type"))+geom_violin(scale
= "width")+scale_fill_manual(values=colors)
+stat_summary(fun=median, geom="point", size=0.6, color="red")+ylab(gene_list[k])+ theme(axis.title.x=element_bla
nk(),
axis.text.y=element_text(size=5),axis.title.y=element_text(size=6,angle=0,face="bold",margin = margin(t = 10, r =
8, b = 0, l = 10),vjust=0.5),axis.text.x=element_text(size=5,face="bold")
,axis.title=element_text(size=4,face="bold"),panel.grid.major = element_blank(), panel.grid.minor = element_blan
k(),
panel.background = element_blank(), axis.line = element_line(colour = "black"),legend.position="none",plot.margin
= unit(c(0, 0,0, 0), "cm")))}
else{
assign(paste("P",k,sep=""),ggplot(Cell_number,aes_string(x="type",y=gene_list[k],fill="type"))+geom_violin(scale
= "width")+scale_fill_manual(values=colors)
+stat_summary(fun=median, geom="point", size=0.6, color="red")+ylab(gene_list[k])+ theme(axis.title.x=element_bla
nk(),
axis.text.x=element_blank(),axis.text.y=element_text(size=5),axis.title.y=element_text(size=6,angle=0,face="bold",
margin = margin(t = 10, r = 8, b = 0, l = 10),vjust=0.5),axis.ticks.x=element_blank(),axis.text=element_text(size
=5)
,axis.title=element_text(size=4,face="bold"),panel.grid.major = element_blank(), panel.grid.minor = element_blan
k(),
panel.background = element_blank(), axis.line = element_line(colour = "black"),legend.position="none",plot.margin
= unit(c(0, 0,0, 0), "cm")))}

}
```

```
P1
```

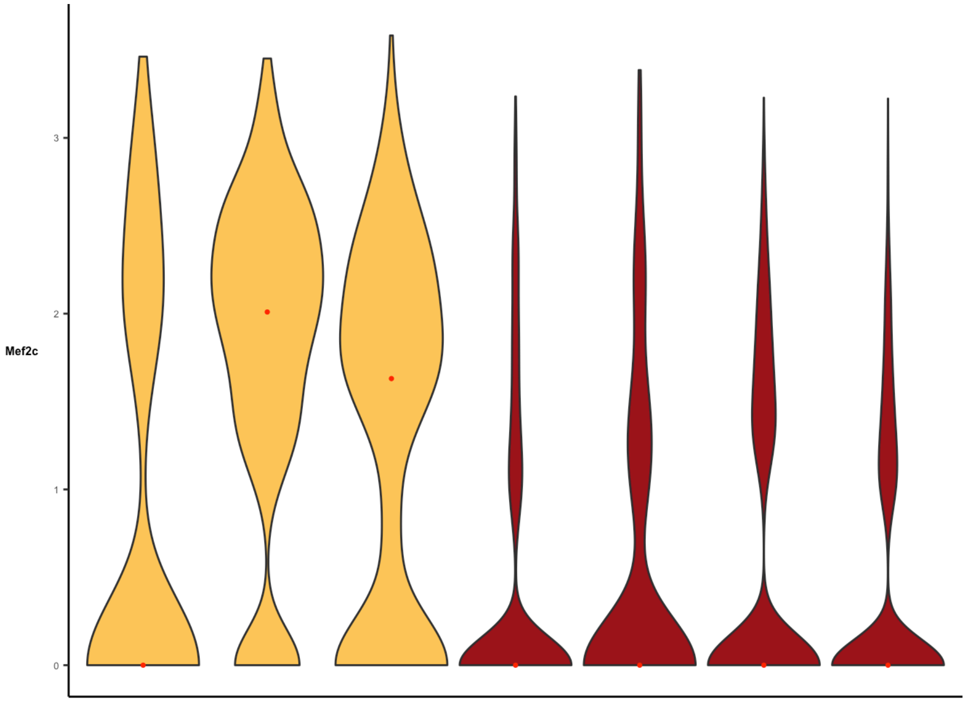


P2

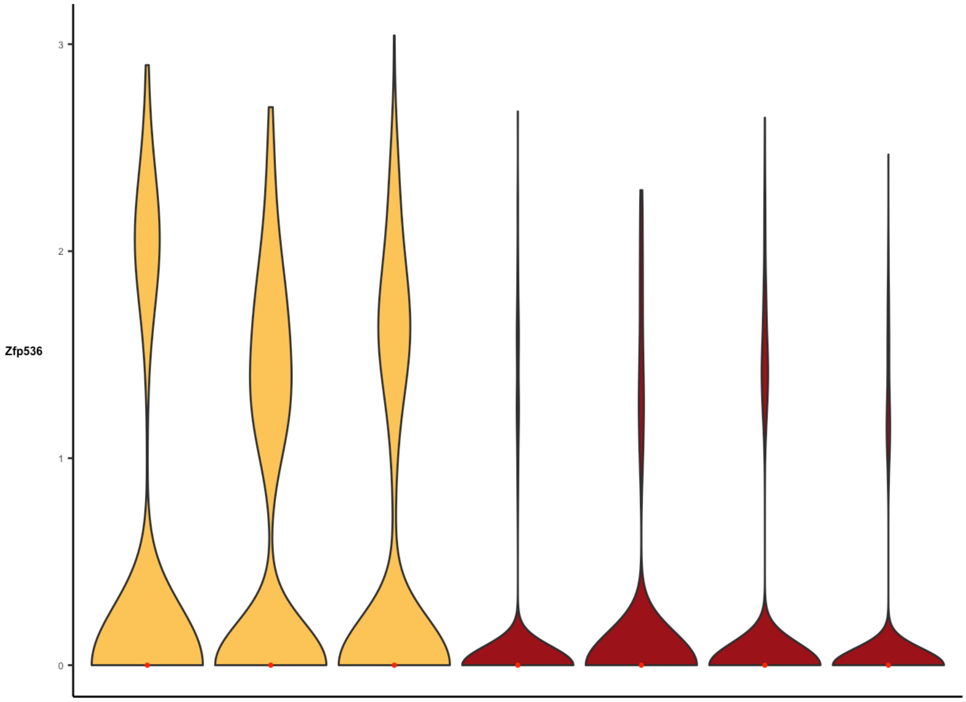


P3

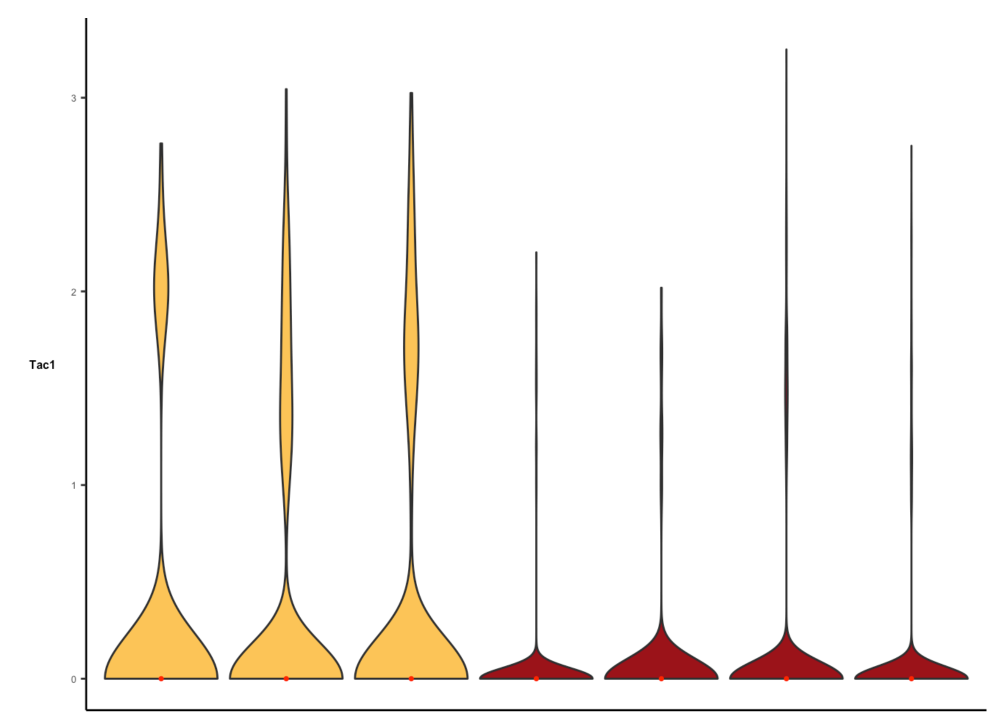


```
P4
```

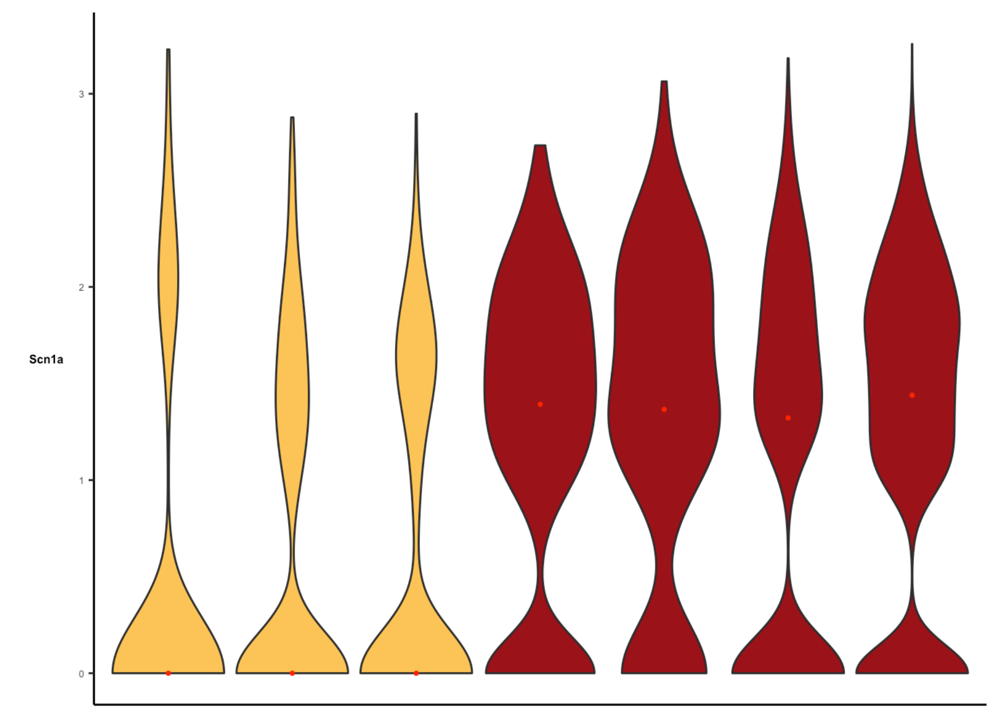


```
P5
```

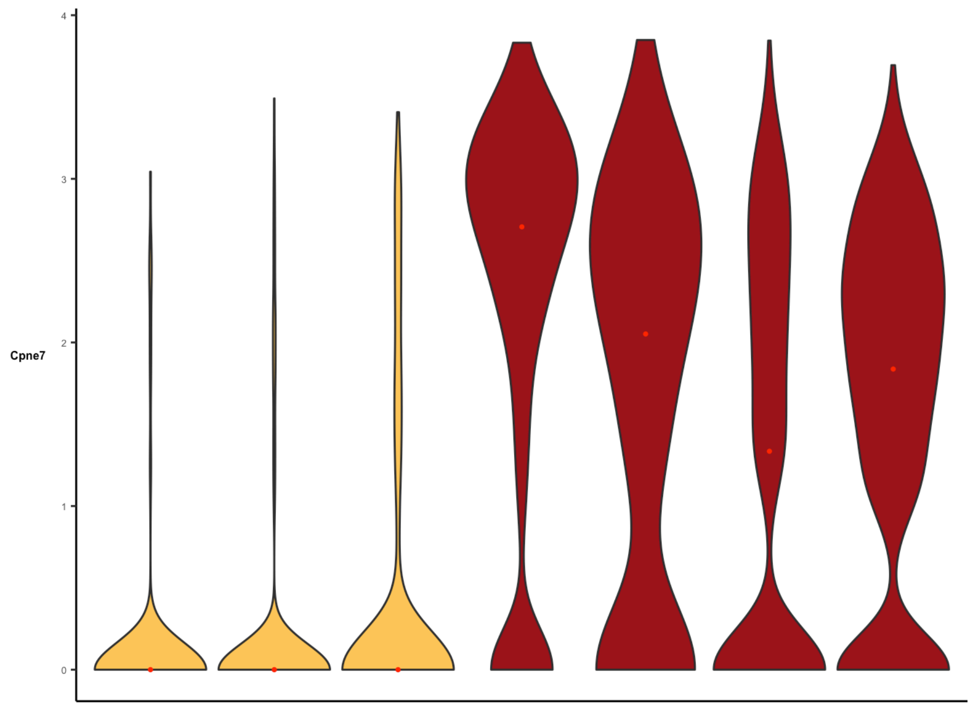


```
P6
```

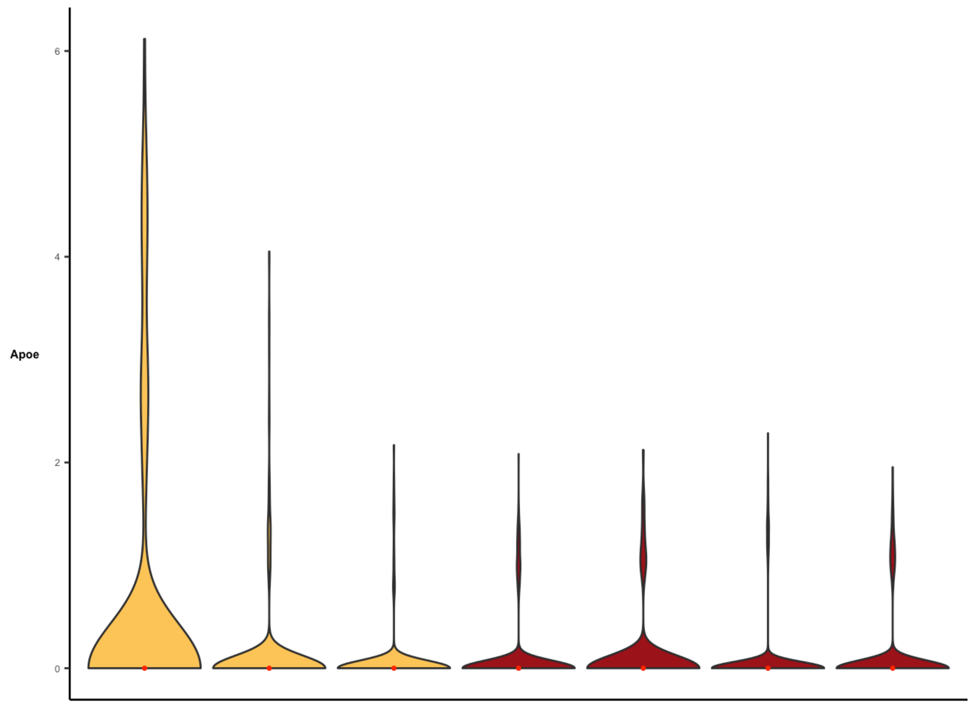


```
P7
```

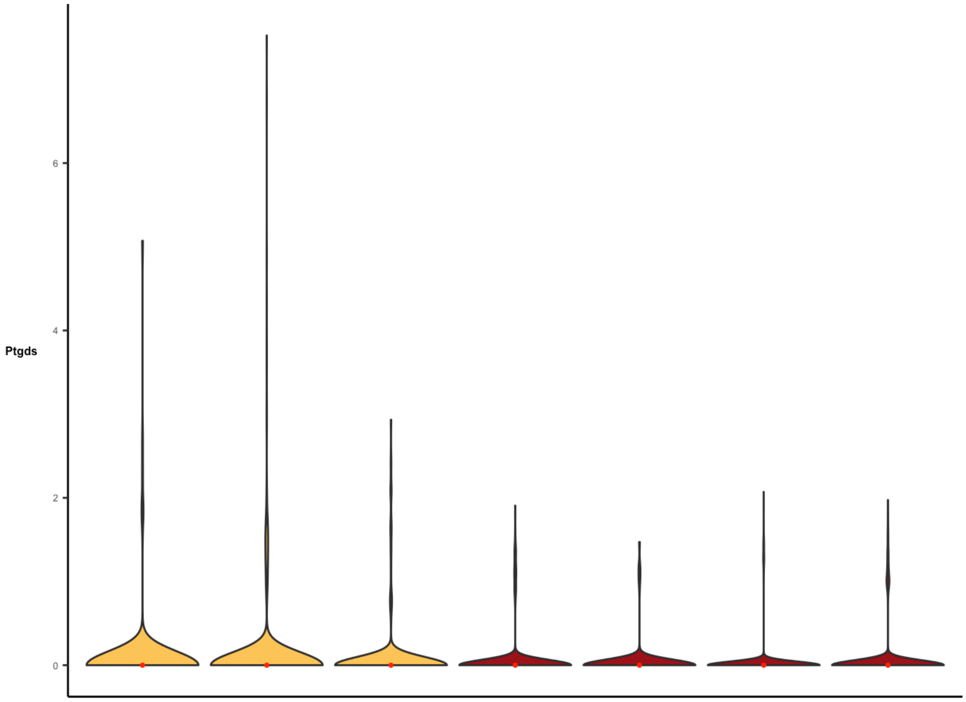

Ptgds
0
2
4
6


```
P8
```

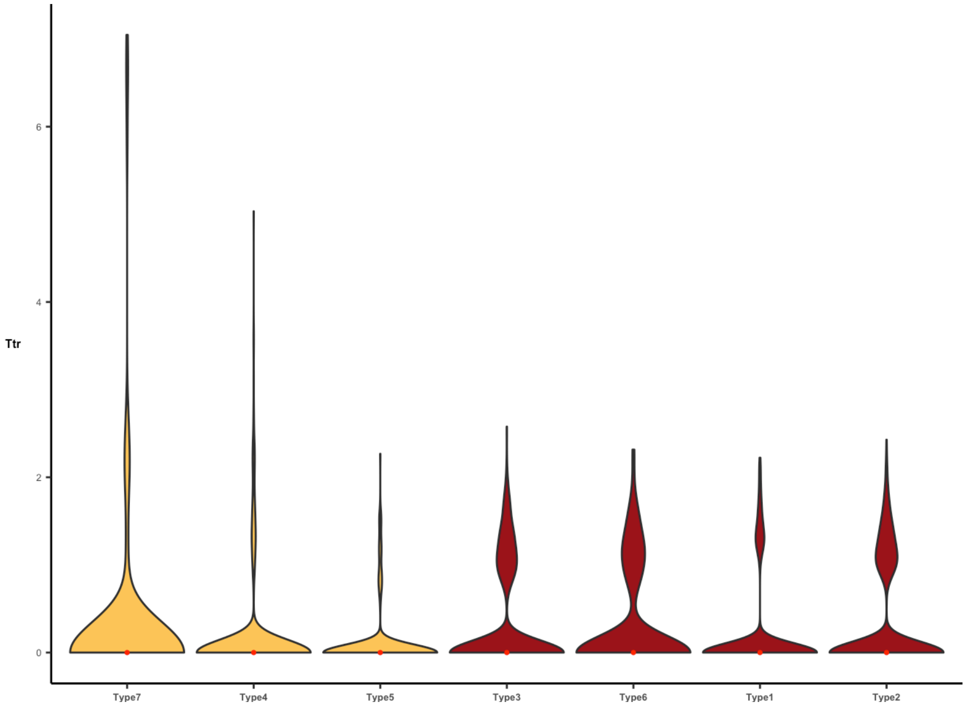

Ttr
0
2
4
6
Type7
Type4
Type5
Type3
Type6
Type1
Type2

```
merge<-list()
for (i in length(gene_list):1){
       if (length(merge)==0){
        merge<-ggplotGrob(eval(parse(text=paste("P",i,sep = ""))))}else{merge<-rbind(ggplotGrob(eval(parse(text=p
aste("P",i,sep = "")))),merge,size = "last")}}

 #all genes
pdf(paste("/Users/lees44/R_project/FC_vs_dCA2_comparison_vector_integration_042821/Violin_plot_FcvsdCA2_markers_7
celltype.pdf",sep=""),height=6, width=50 , paper = "letter")


grid.newpage()
grid.draw(merge)
```

#########violin plot for 7 cell types with neurotransmitters 08.24.21 ####################

```
Cell_type<-rev(c("Type7", "Type4", "Type5", "Type3", "Type6", "Type1", "Type2"))
gene_list<-c("Gad1", "Sst", "Cck", "Pvalb","Htr3a", "Nos1","Slc17a7","Calb2")
```

```
Cell_number <- NULL
for (i in 1:length(Cell_type)){
                temp1 <- data.frame(type=factor(rep(Cell_type[i],sum(Cell_type[i]==Idents(dCA2.integrated)))))
                temp2 <- as.data.frame(dCA2.integrated_clean_7celltype@assays$RNA@data[gene_list,Idents(dCA2.inte
grated_clean_7celltype)==Cell_type[i]])
                temp3 <- cbind(temp1,t(temp2))
                Cell_number <- rbind(temp3,Cell_number)
}
rownames(Cell_number) <- NULL
```

```
colors<-c("#ffc857","#ffc857","#ffc857","#9B0C1E","#9B0C1E","#9B0C1E","#9B0C1E")
```

```
for (k in 1:length(gene_list))
    {
if (k==length(gene_list)){
assign(paste("P",k,sep=""),ggplot(Cell_number,aes_string(x="type",y=gene_list[k],fill="type"))+geom_violin(scale
= "width")+scale_fill_manual(values=colors)
+stat_summary(fun=median, geom="point", size=0.6, color="red")+ylab(gene_list[k])+ theme(axis.title.x=element_bla
nk(),
axis.text.y=element_text(size=5),axis.title.y=element_text(size=6,angle=0,face="bold",margin = margin(t = 10, r =
8, b = 0, l = 10),vjust=0.5),axis.text.x=element_text(size=5,face="bold")
,axis.title=element_text(size=4,face="bold"),panel.grid.major = element_blank(), panel.grid.minor = element_blan
k(),
panel.background = element_blank(), axis.line = element_line(colour = "black"),legend.position="none",plot.margin
= unit(c(0, 0,0, 0), "cm")))}
else{
assign(paste("P",k,sep=""),ggplot(Cell_number,aes_string(x="type",y=gene_list[k],fill="type"))+geom_violin(scale
= "width")+scale_fill_manual(values=colors)
+stat_summary(fun=median, geom="point", size=0.6, color="red")+ylab(gene_list[k])+ theme(axis.title.x=element_bla
nk(),
axis.text.x=element_blank(),axis.text.y=element_text(size=5),axis.title.y=element_text(size=6,angle=0,face="bold",
margin = margin(t = 10, r = 8, b = 0, l = 10),vjust=0.5),axis.ticks.x=element_blank(),axis.text=element_text(size
=5)
,axis.title=element_text(size=4,face="bold"),panel.grid.major = element_blank(), panel.grid.minor = element_blan
k(),
panel.background = element_blank(), axis.line = element_line(colour = "black"),legend.position="none",plot.margin
= unit(c(0, 0,0, 0), "cm")))}

}
```

```
P1
```

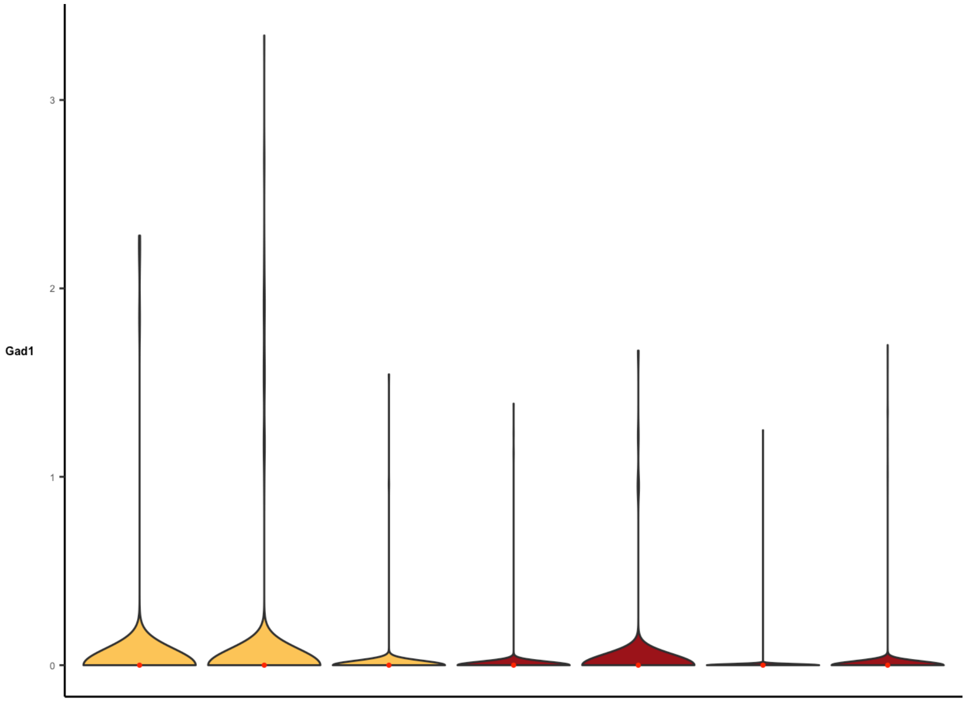


```
P2
```

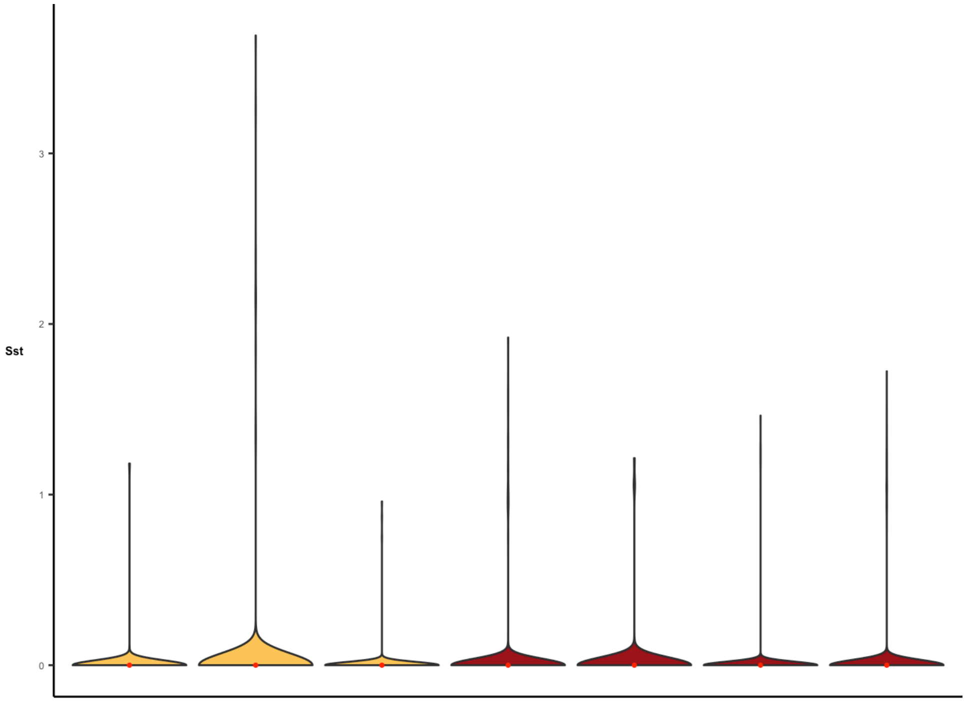


```
P3
```

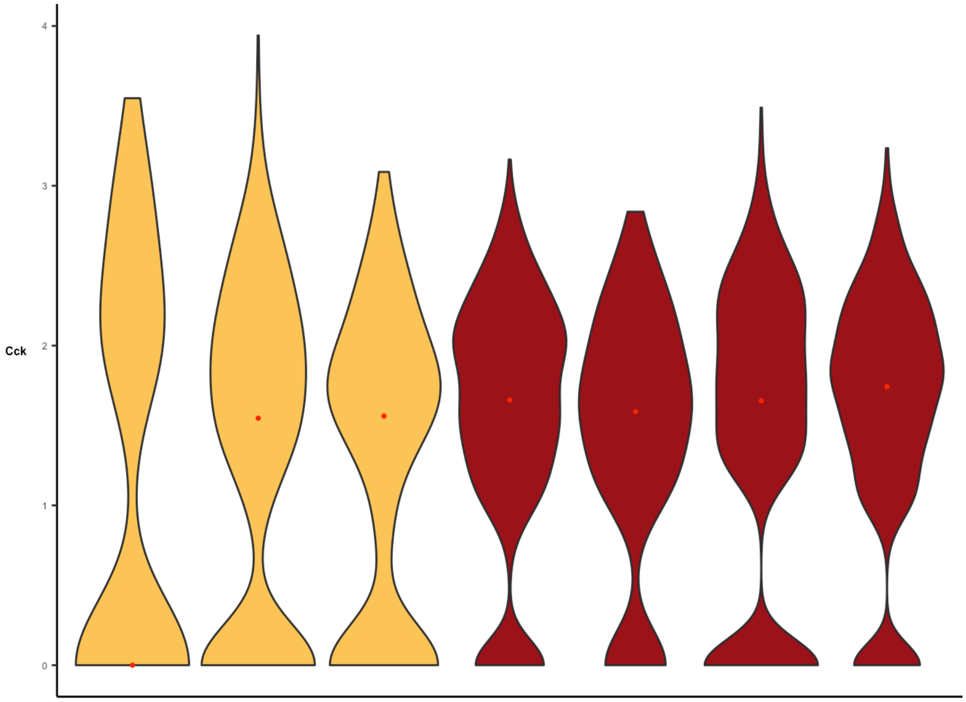


```
P4
```

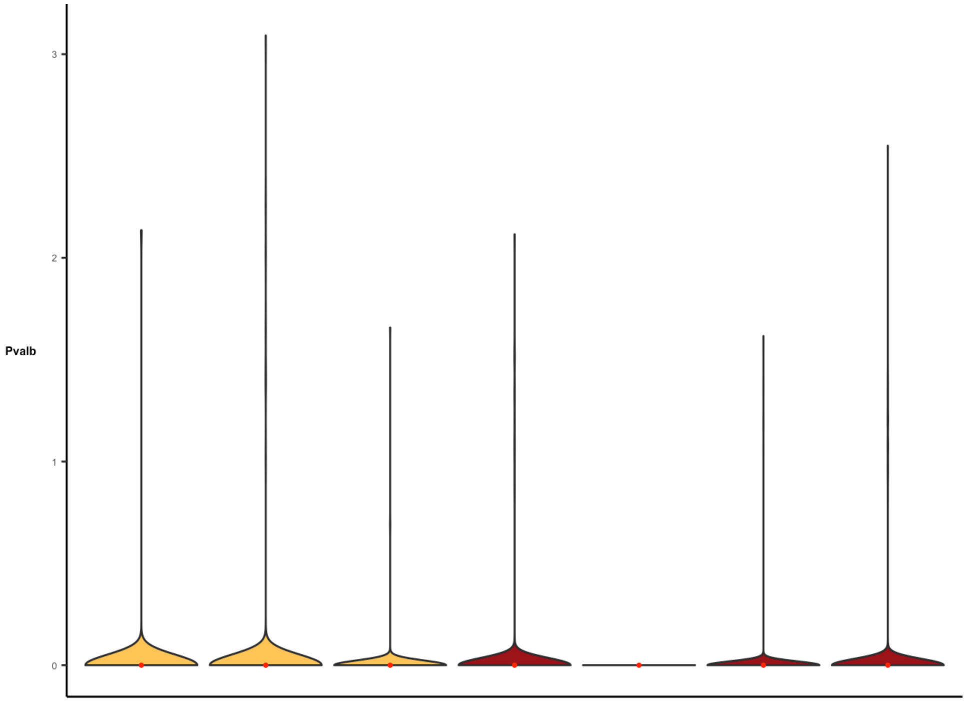


```
P5
```

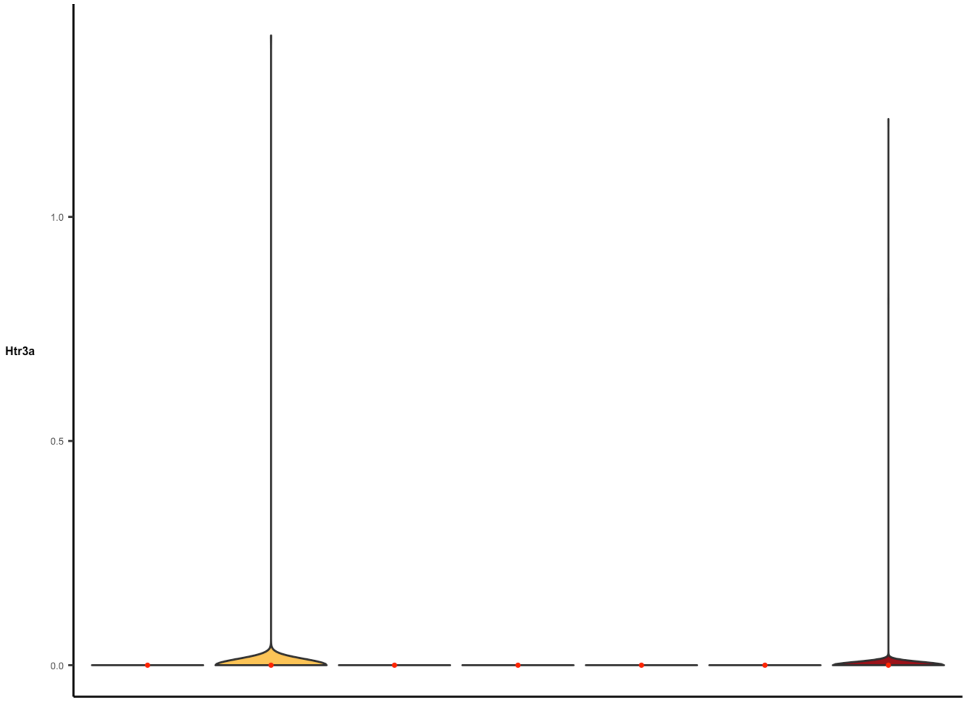


P6

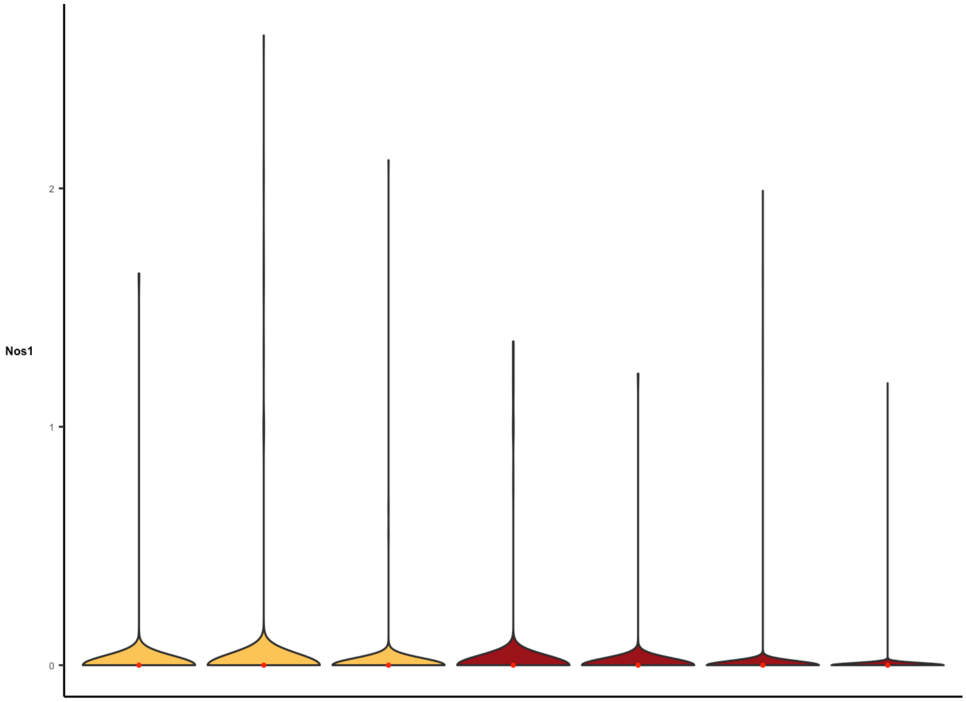


P7

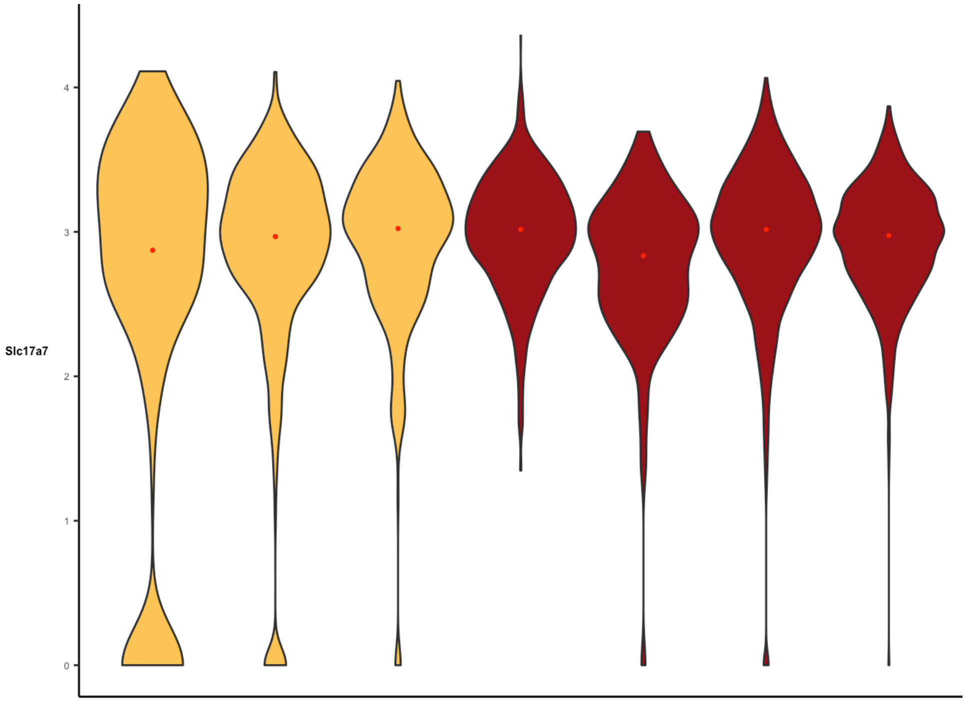
Slc17a7
0
1
2
3
4

```
P8
```

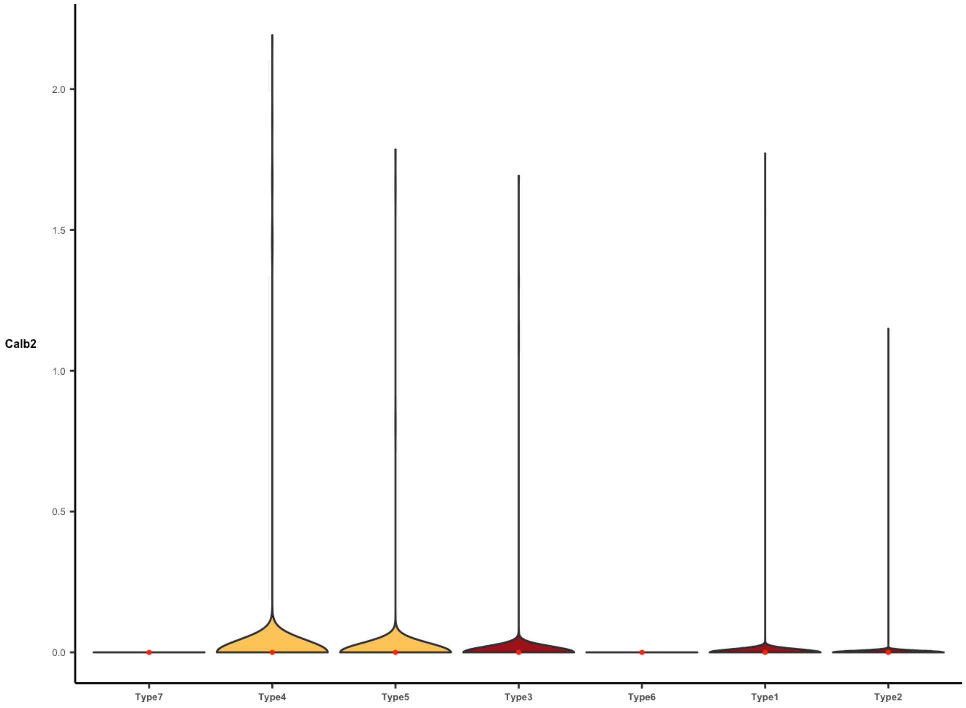
Calb2
0.0
0.5
1.0
1.5
2.0
Type7
Type4
Type5
Type3
Type6
Type1
Type2

```
merge<-list()
for (i in length(gene_list):1){
       if (length(merge)==0){
        merge<-ggplotGrob(eval(parse(text=paste("P",i,sep = ""))))}else{merge<-rbind(ggplotGrob(eval(parse(text=p
aste("P",i,sep = "")))),merge,size = "last")}}

 #all genes
pdf(paste("/Users/lees44/R_project/FC_vs_dCA2_comparison_vector_integration_042821/Violin_plot_neuronalmarker_7ce
lltype.pdf",sep=""),height=6, width=50 , paper = "letter")


grid.newpage()
grid.draw(merge)
```

##########################violin plot with each cell type markers################################

Violinplot for potassium channels, figure goes with cluster tree

```
Cell_type<-rev(c("Type5", "Type7", "Type4", "Type1", "Type6", "Type2", "Type3"))
gene_list<-c("Zfp536", "Tac1", "Clu", "Mef2c", "Kcnk2", "Bdnf", "Amigo2", "Cpne7")
```

```
Cell_number <- NULL
for (i in 1:length(Cell_type)){
                temp1 <- data.frame(type=factor(rep(Cell_type[i],sum(Cell_type[i]==Idents(dCA2.integrated_clean_7
celltype)))))
                temp2 <- as.data.frame(dCA2.integrated_clean_7celltype@assays$RNA@data[gene_list,Idents(dCA2.inte
grated_clean_7celltype)==Cell_type[i]])
                temp3 <- cbind(temp1,t(temp2))
                Cell_number <- rbind(temp3,Cell_number)
}
rownames(Cell_number) <- NULL
```

```
colors<-c("#ffc857","#ffc857","#ffc857","#9B0C1E", "#9B0C1E", "#9B0C1E", "#E8C6C7")
```

```
for (k in 1:length(gene_list))
    {
if (k==length(gene_list)){
assign(paste("P",k,sep=""),ggplot(Cell_number,aes_string(x="type",y=gene_list[k],fill="type"))+geom_violin(scale
= "width")+scale_fill_manual(values=colors)
+stat_summary(fun=median, geom="point", size=0.6, color="red")+ylab(gene_list[k])+ theme(axis.title.x=element_bla
nk(),
axis.text.y=element_text(size=5),axis.title.y=element_text(size=6,angle=0,face="bold",margin = margin(t = 10, r =
8, b = 0, l = 10),vjust=0.5),axis.text.x=element_text(size=5,face="bold")
,axis.title=element_text(size=4,face="bold"),panel.grid.major = element_blank(), panel.grid.minor = element_blan
k(),
panel.background = element_blank(), axis.line = element_line(colour = "black"),legend.position="none",plot.margin
= unit(c(0, 0,0, 0), "cm")))}
else{
assign(paste("P",k,sep=""),ggplot(Cell_number,aes_string(x="type",y=gene_list[k],fill="type"))+geom_violin(scale
= "width")+scale_fill_manual(values=colors)
+stat_summary(fun=median, geom="point", size=0.6, color="red")+ylab(gene_list[k])+ theme(axis.title.x=element_bla
nk(),
axis.text.x=element_blank(),axis.text.y=element_text(size=5),axis.title.y=element_text(size=6,angle=0,face="bold",
margin = margin(t = 10, r = 8, b = 0, l = 10),vjust=0.5),axis.ticks.x=element_blank(),axis.text=element_text(size
=5)
,axis.title=element_text(size=4,face="bold"),panel.grid.major = element_blank(), panel.grid.minor = element_blan
k(),
panel.background = element_blank(), axis.line = element_line(colour = "black"),legend.position="none",plot.margin
= unit(c(0, 0,0, 0), "cm")))}

}
```

```
P1
```

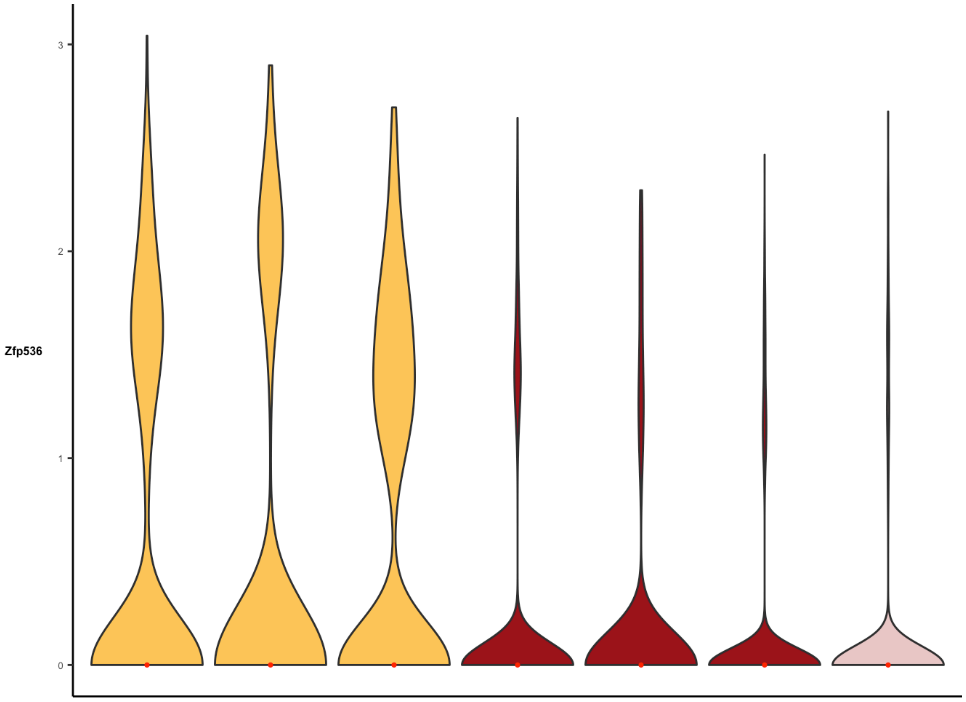


```
P2
```

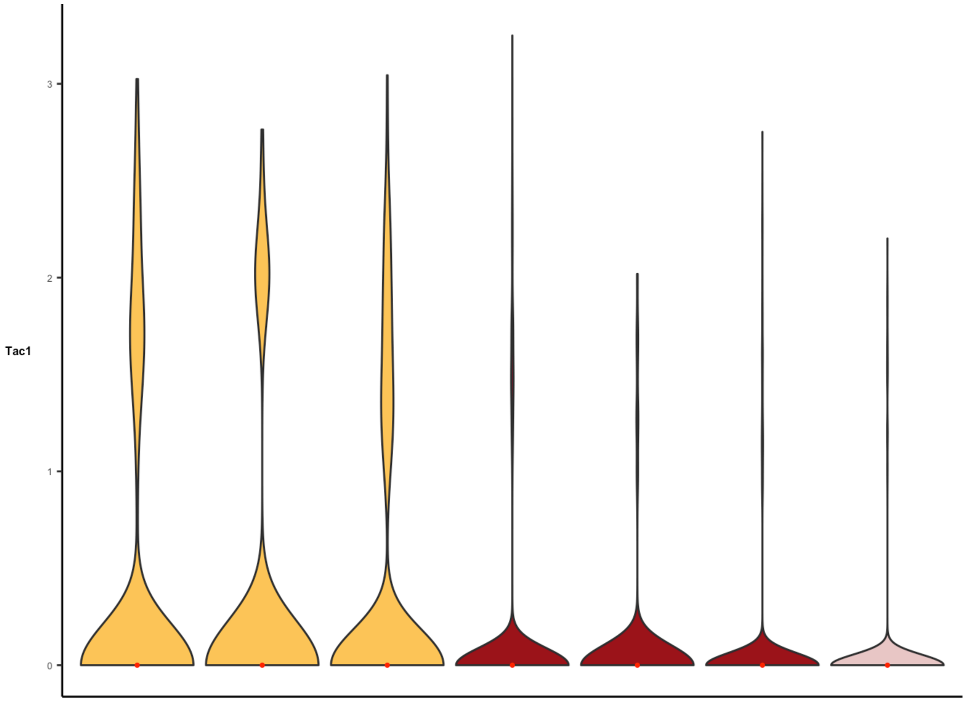


```
P3
```

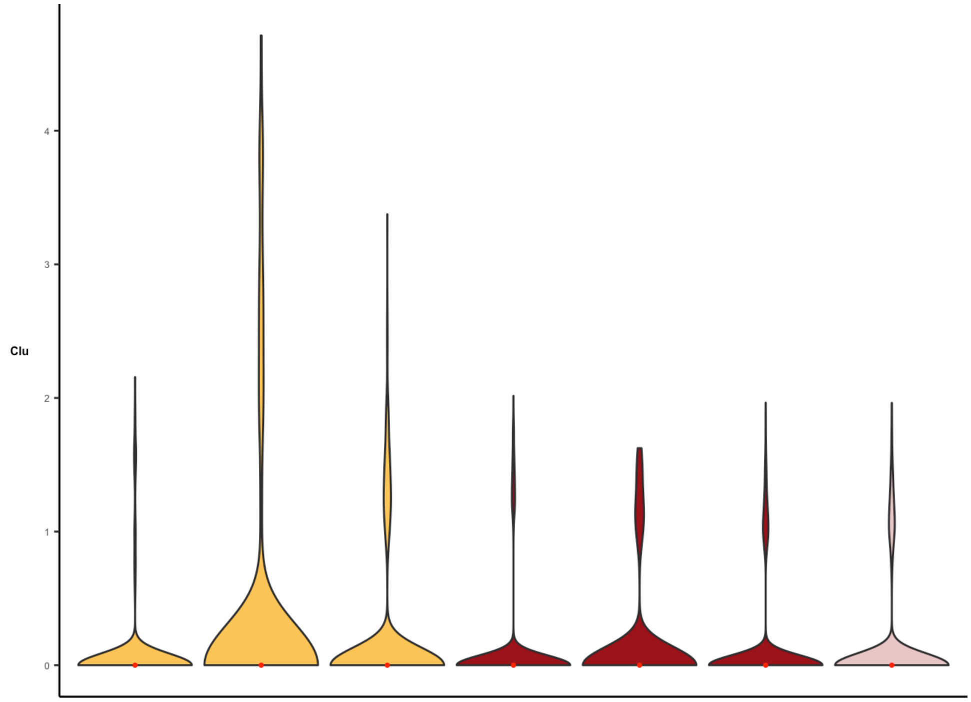


```
P4
```

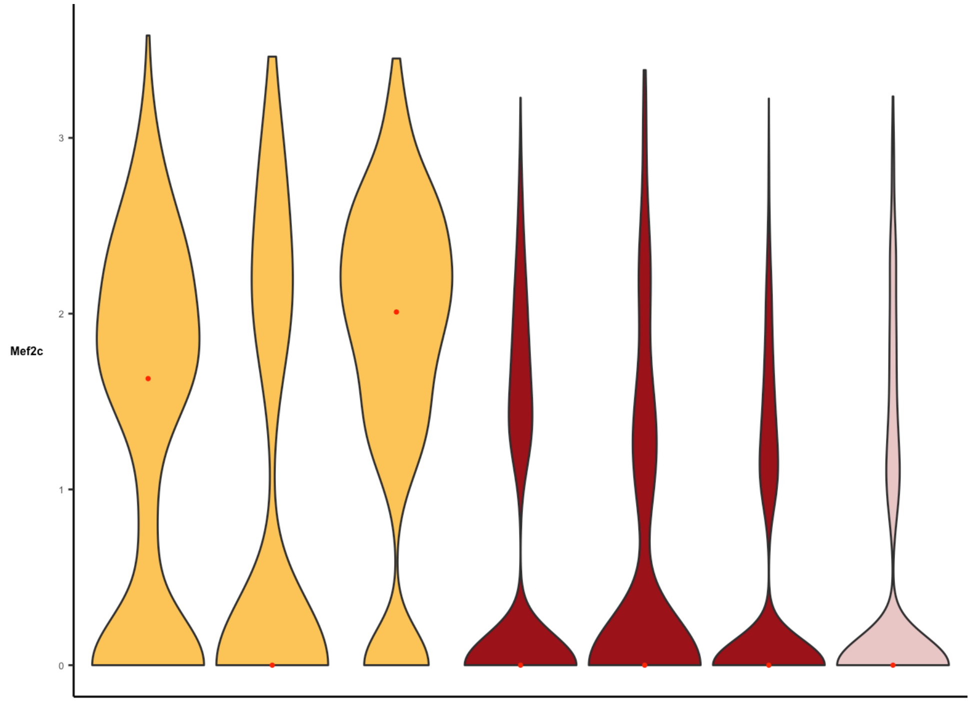


```
P5
```

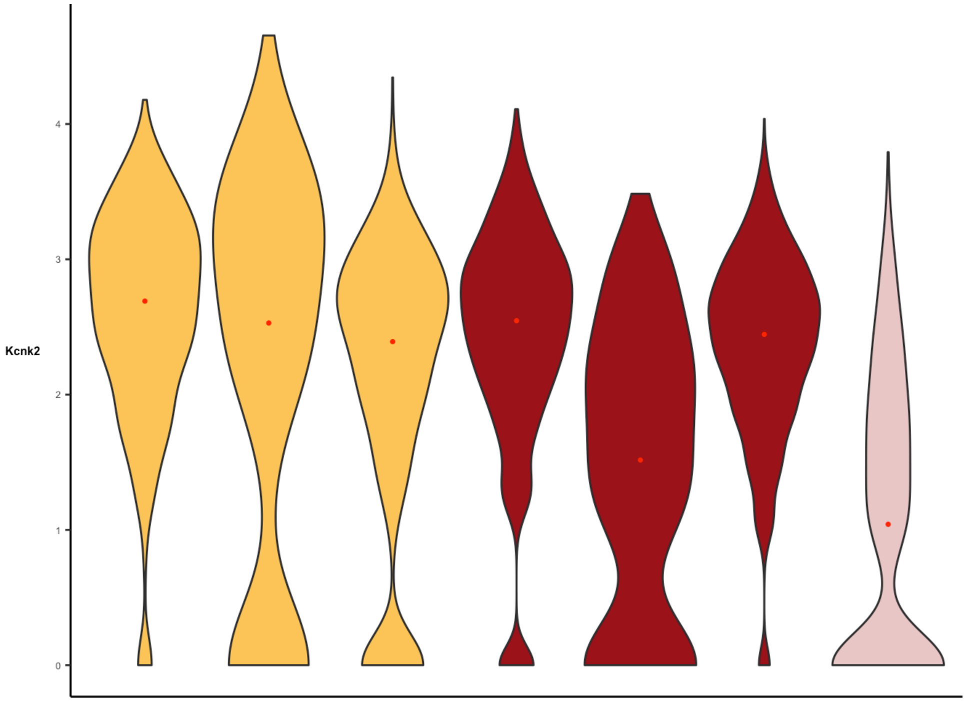


```
P6
```

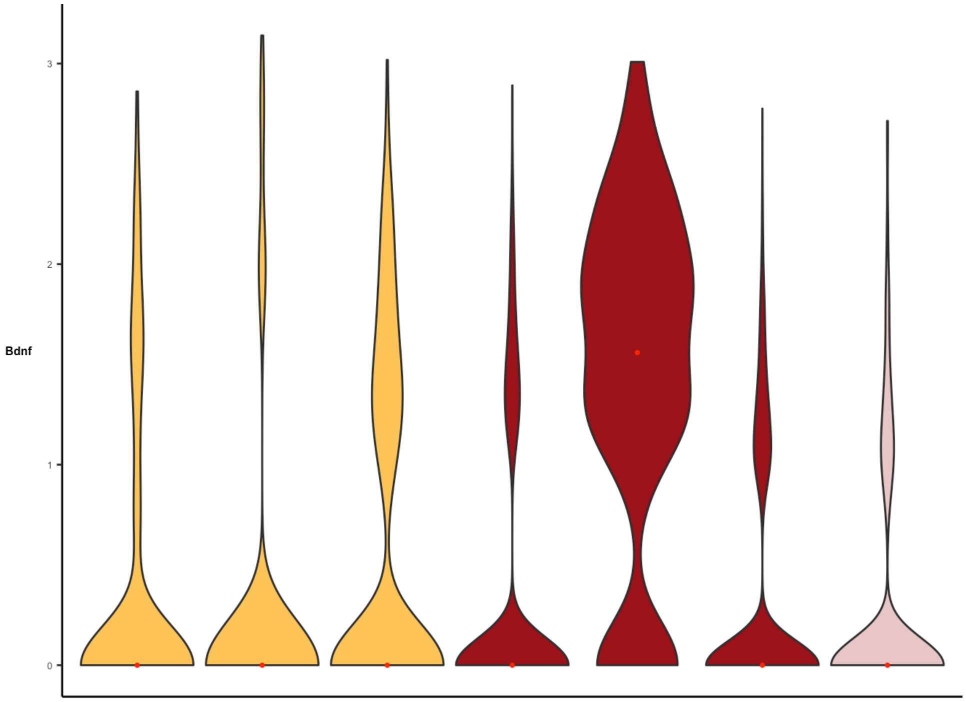


```
P7
```

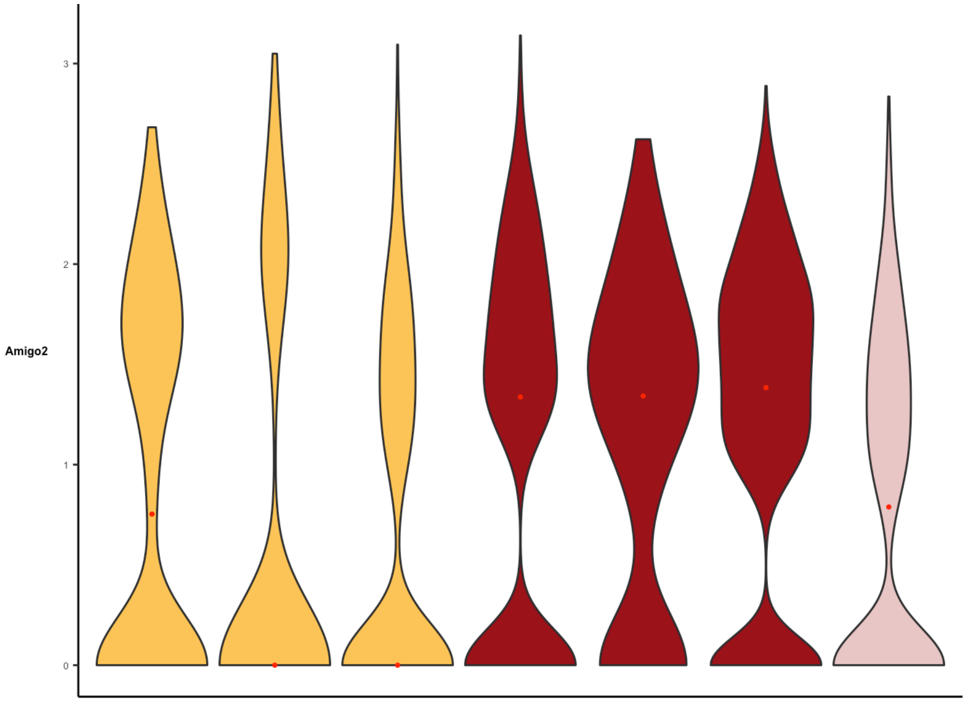


```
merge<-list()
for (i in length(gene_list):1){
       if (length(merge)==0){
        merge<-ggplotGrob(eval(parse(text=paste("P",i,sep = ""))))}else{merge<-rbind(ggplotGrob(eval(parse(text=p
aste("P",i,sep = "")))),merge,size = "last")}}

 #all genes
pdf(paste("/Users/lees44/R_project/FC_vs_dCA2_comparison_vector_integration_042821/cellytype7.pdf",".pdf",sep=""),
height=6, width=50 , paper = "letter")


grid.newpage()
grid.draw(merge)
```

```
DefaultAssay(dCA2.integrated) <- "RNA" #switch back and forth with integrated
```

Generalized dCA2 vs FC

Differential expression testing: see https://satijalab.org/seurat/archive/v3.1/de_vignette.html (https://satijalab.org/seurat/archive/v3.1/de_vignette.html) https://satijalab.org/seurat/reference/findmarkers (https://satijalab.org/seurat/reference/findmarkers)

```
dCA2.integrated@meta.data$Group = "no"
dCA2.integrated@meta.data$Group[dCA2.integrated@meta.data$protocol == "dCA2_Fpos" | dCA2.integrated@meta.data$pro
tocol == "dCA2_Mpos"] = "dCA2"
dCA2.integrated@meta.data$Group[dCA2.integrated@meta.data$protocol == "FC_Fpos" | dCA2.integrated@meta.data$proto
col == "FC_Mpos"] = "FC"

table(dCA2.integrated@meta.data$Group)
```

```
##
## dCA2   FC
## 3119 2809
```

```
Idents(object = dCA2.integrated) <- "Group"

dCA2vsFC.markers <- FindMarkers(dCA2.integrated, ident.1 = "FC" , ident.2 = "dCA2", min.pct = 0.25)
head(dCA2vsFC.markers, n = 10)
```

```
##          p_val avg_log2FC pct.1 pct.2 p_val_adj
## Sphkap      0 -1.2696400 0.261 0.745         0
## Cacnb2      0 -1.1937076 0.412 0.840         0
## Nebl        0 -1.1155057 0.719 0.956         0
## Snhg11      0 -1.2958831 0.989 0.995         0
## Sms         0 -1.1882048 0.600 0.924         0
## Trim2       0 -1.1185150 0.611 0.933         0
## Asph        0 -1.3768794 0.677 0.931         0
## Epha5       0  0.8209458 0.999 0.975         0
## Akap13      0 -2.7928265 0.280 0.924         0
## Socs2       0 -1.2298497 0.191 0.703         0
```

```
write.csv(dCA2vsFC.markers, "/Users/lees44/R_project/FC_vs_dCA2_comparison_vector_integration_042821/FCvsdCA2top_
10markers_integrated.csv")
```

```
write.csv(dCA2vsFC.markers, "/Users/lees44/R_project/FC_vs_dCA2_comparison_vector_integration_042821/top_10marker
s_RNAassay.csv", quote = F)
```

```
## Error in write.table(dCA2vsFC.markers, "/Users/lees44/R_project/FC_vs_dCA2_comparison_vector_integration_04282
1/top_10markers_RNAassay.csv", : invalid 'quote' specification
```

```
p3<-DimPlot(dCA2.integrated, reduction = "umap", group.by = "Group", pt=1)
plot_grid(p3)
```

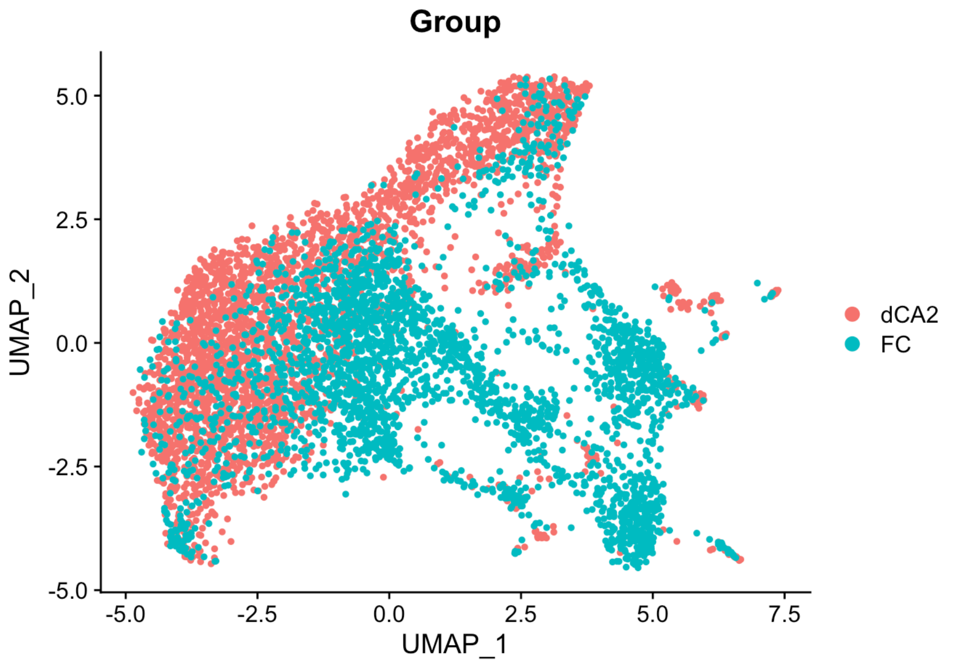


```
ggsave(file="/Users/lees44/R_project/FC_vs_dCA2_comparison_vector_integration_042821/FC_dCA2_UMAP_Group.pdf",widt
h=10,height=10)
```

```
p4<-DimPlot(dCA2.integrated, reduction = "umap", split.by = "Group",  pt=1)
plot_grid(p4)
```

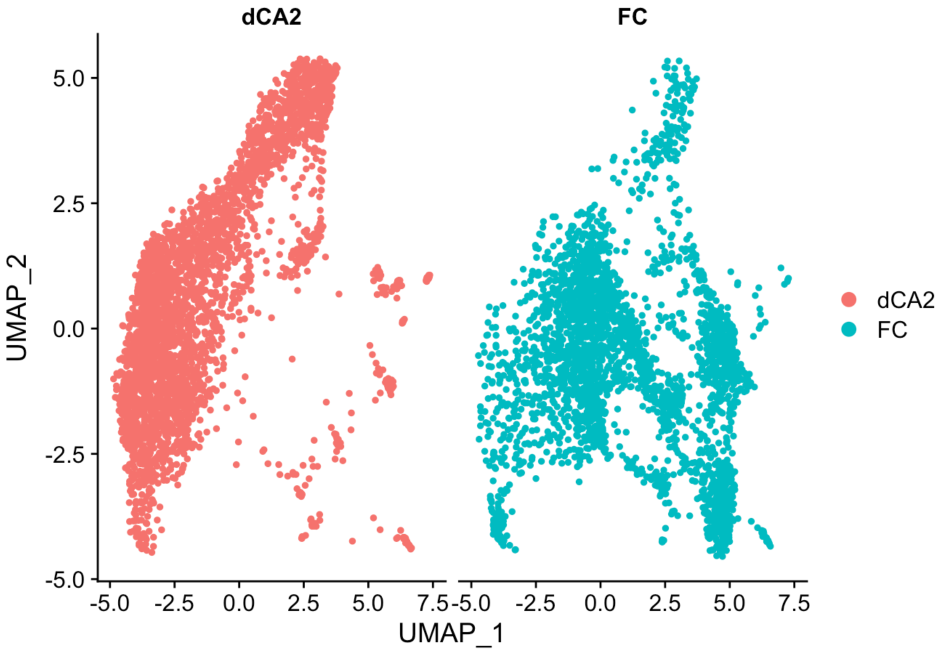


```
ggsave(file="/Users/lees44/R_project/FC_vs_dCA2_comparison_vector_integration_042821/FC_dCA2_UMAP_Group_split.pd
f",width=10,height=10)
```

Violin plot for genes in dCA2 vs FC see https://ggplot2.tidyverse.org/reference/theme.html (https://ggplot2.tidyverse.org/reference/theme.html) changing graph using ggplot theme

```
VlnPlot(object = dCA2.integrated, features = "Scn1a", pt.size = 0.5) +theme(title = element_text(size = 30), axi
s.title.x = element_text(size =20), axis.title.y = element_text(size = 20), axis.text.x = element_text(size = 2
0), legend.text = element_text(size =20))
```

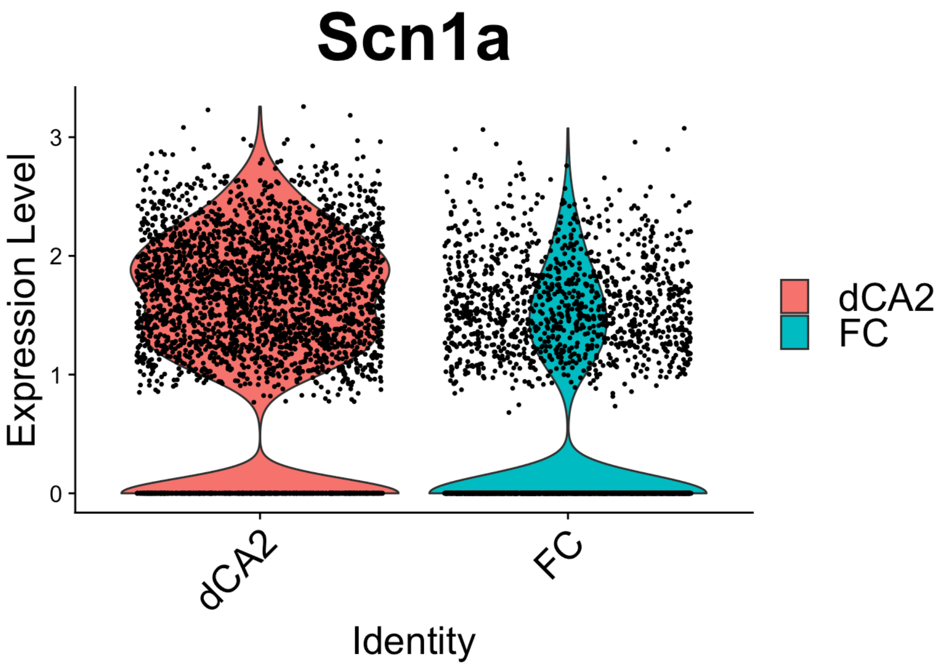


```
ggsave(file="/Users/lees44/R_project/FC_vs_dCA2_comparison_vector_integration_042821/Scn1a_FCvsCA2.pdf", width=7,
height=10)
```

```
VlnPlot(object = dCA2.integrated, features = "Cpne7", pt.size = 0.5)+theme(title = element_text(size = 30), axis.
title.x = element_text(size =20), axis.title.y = element_text(size = 20), axis.text.x = element_text(size = 20),
legend.text = element_text(size =20))
```

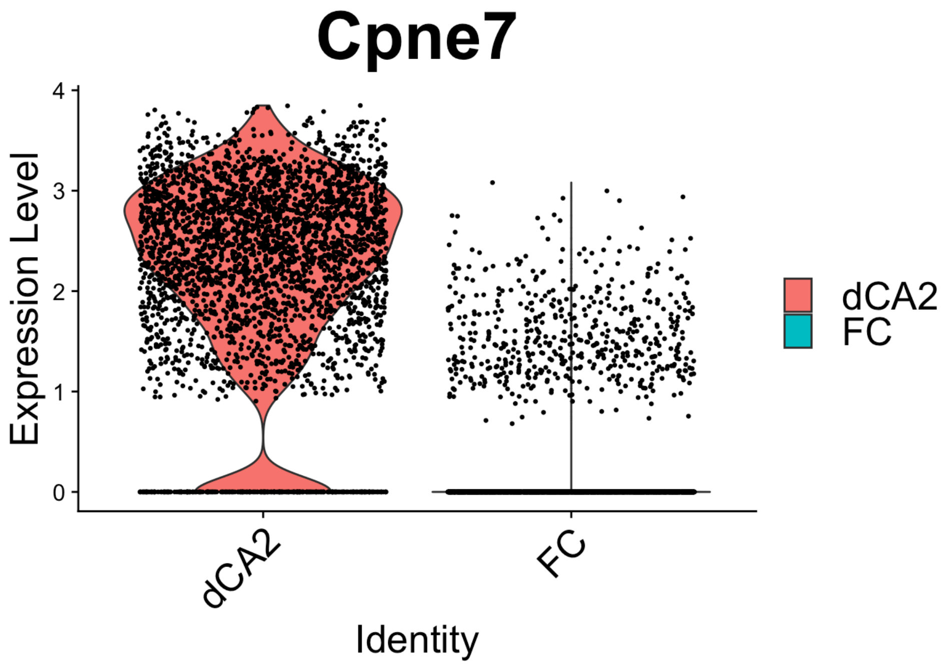


```
ggsave(file="/Users/lees44/R_project/FC_vs_dCA2_comparison_vector_integration_042821/Cpne7_FCvsCA2.pdf", width=7,
height=10)
```

```
VlnPlot(object = dCA2.integrated, features = "Ptgds", pt.size = 0.5)+theme(title = element_text(size = 30), axis.
title.x = element_text(size =20), axis.title.y = element_text(size = 20), axis.text.x = element_text(size = 20),
legend.text = element_text(size =20))
```

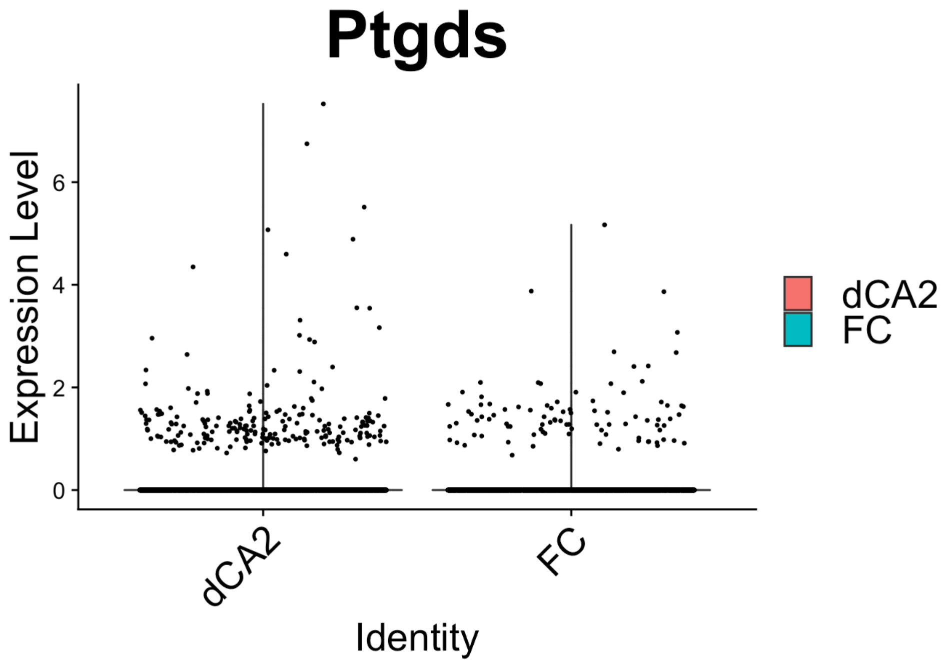

```
ggsave(file="/Users/lees44/R_project/FC_vs_dCA2_comparison_vector_integration_042821/Ptgds_FCvsCA2.pdf", width=7,
height=10)
```

```
VlnPlot(object = dCA2.integrated, features = "Ttr", pt.size = 0.5) + theme(title = element_text(size = 30), axis.
title.x = element_text(size =20), axis.title.y = element_text(size = 20), axis.text.x = element_text(size = 20),
legend.text = element_text(size =20))
```

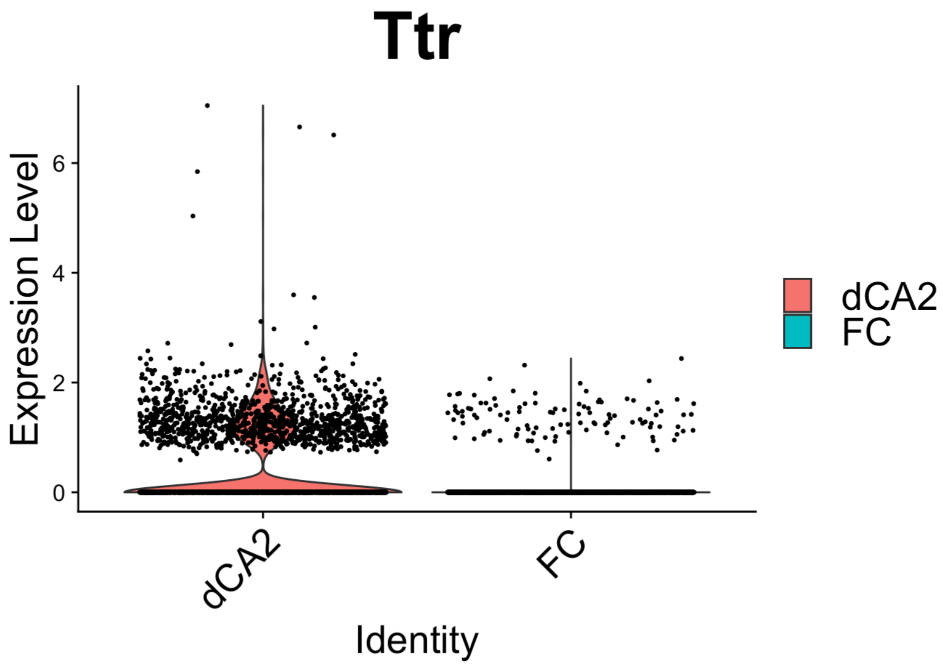


```
ggsave(file="/Users/lees44/R_project/FC_vs_dCA2_comparison_vector_integration_042821/Ttr_FCvsCA2.pdf", width=7,he
ight=10)
```

```
VlnPlot(object = dCA2.integrated, features = "Apoe", pt.size = 0.5) + theme(title = element_text(size = 30), axi
s.title.x = element_text(size =20), axis.title.y = element_text(size = 20), axis.text.x = element_text(size = 2
0), legend.text = element_text(size =20))
```

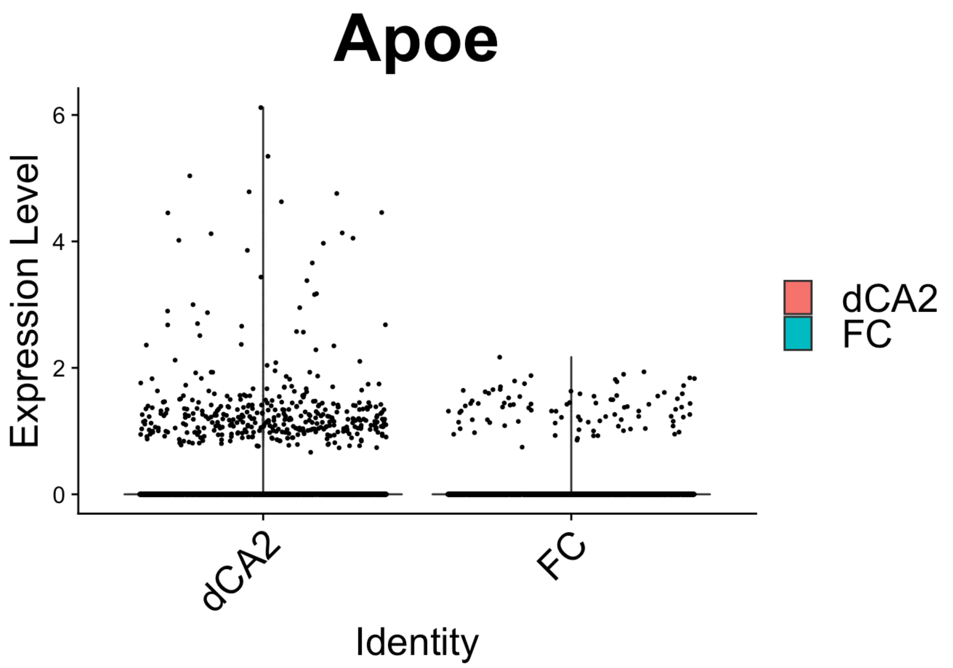


```
ggsave(file="/Users/lees44/R_project/FC_vs_dCA2_comparison_vector_integration_042821/Apoe_FCvsCA2.pdf", width=7,h
eight=10)
```

```
VlnPlot(object = dCA2.integrated, features = "Mef2c", pt.size = 0.5) + theme(title = element_text(size = 30), axi
s.title.x = element_text(size =20), axis.title.y = element_text(size = 20), axis.text.x = element_text(size = 2
0), legend.text = element_text(size =20))
```

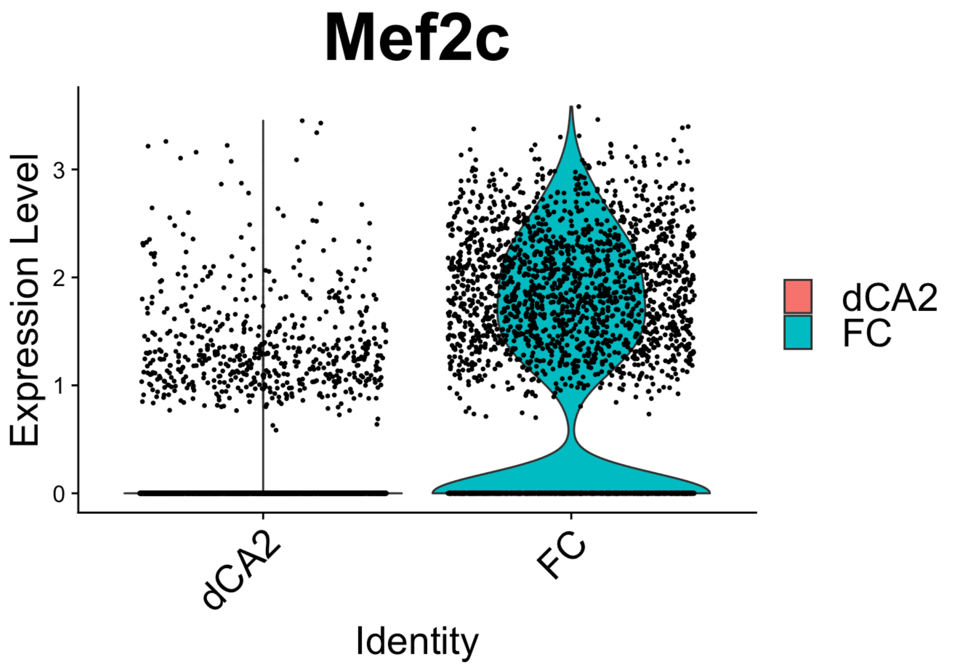


```
ggsave(file="/Users/lees44/R_project/FC_vs_dCA2_comparison_vector_integration_042821/Mef2c_FCvsCA2.pdf", width=7,
height=10)
```

```
VlnPlot(object = dCA2.integrated, features = "Tac1", pt.size = 0.5) + theme(title = element_text(size = 30), axi
s.title.x = element_text(size =20), axis.title.y = element_text(size = 20), axis.text.x = element_text(size = 2
0), legend.text = element_text(size =20))
```

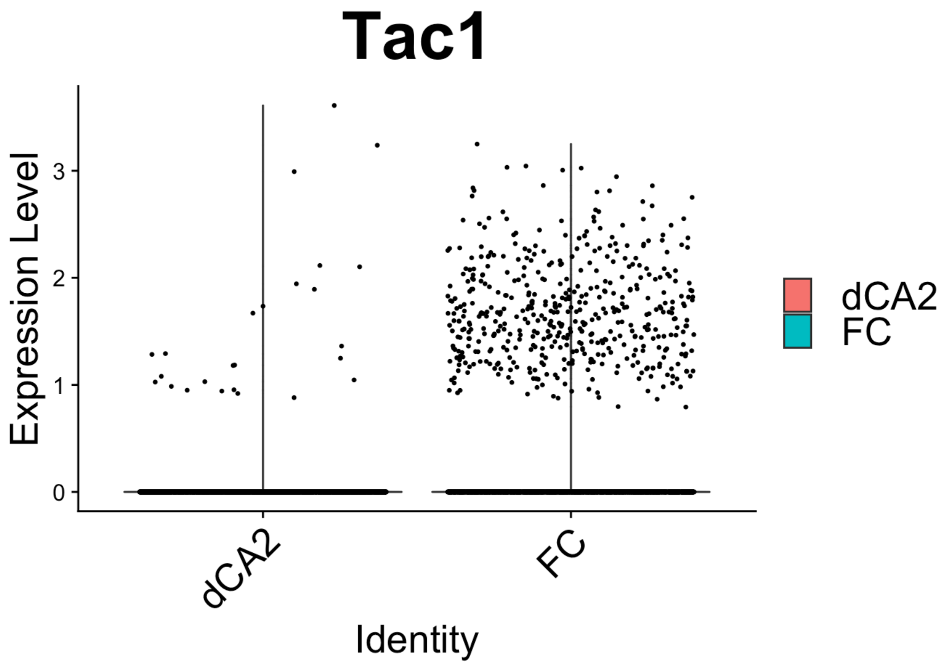


```
ggsave(file="/Users/lees44/R_project/FC_vs_dCA2_comparison_vector_integration_042821/Tac1_FCvsCA2.pdf", width=7,h
eight=10)
```

```
VlnPlot(object = dCA2.integrated, features = "Zfp536", pt.size = 0.5) + theme(title = element_text(size = 30), ax
is.title.x = element_text(size =20), axis.title.y = element_text(size = 20), axis.text.x = element_text(size = 2
0), legend.text = element_text(size =20))
```

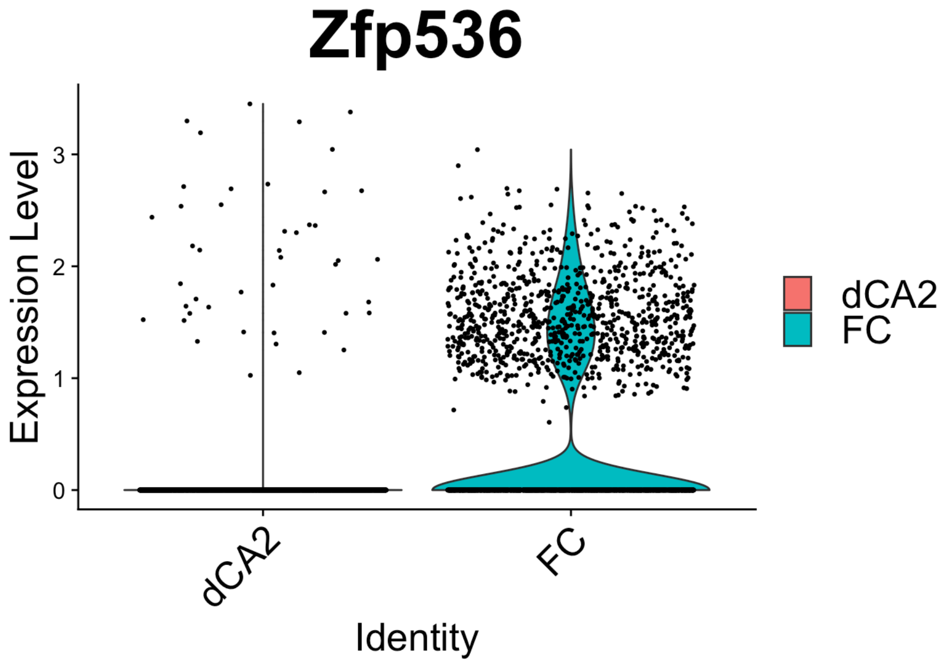

```
ggsave(file="/Users/lees44/R_project/FC_vs_dCA2_comparison_vector_integration_042821/Zfp536_FCvsCA2.pdf", width=
7,height=10)
```

```
VlnPlot(object = dCA2.integrated, features = "Igfbp4", pt.size = 0.5) + theme(title = element_text(size = 30), ax
is.title.x = element_text(size =20), axis.title.y = element_text(size = 20), axis.text.x = element_text(size = 2
0), legend.text = element_text(size =20))
```

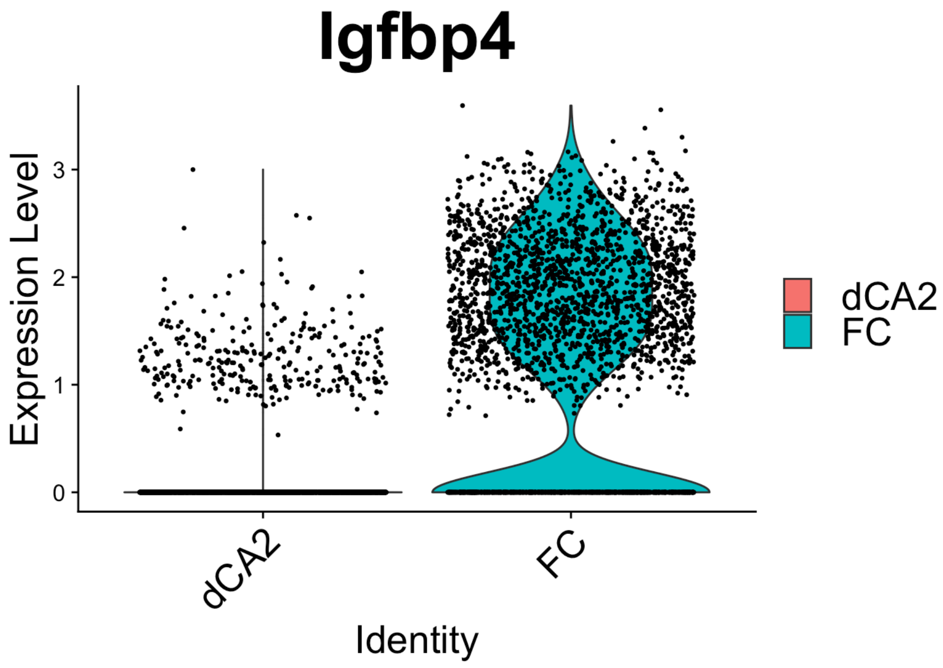


```
ggsave(file="/Users/lees44/R_project/FC_vs_dCA2_comparison_vector_integration_042821/Igfbp4_FCvsCA2.pdf", width=
7,height=10)
```

```
VlnPlot(object = dCA2.integrated, features = "Ntng1", pt.size = 0.5) + theme(title = element_text(size = 30), axi
s.title.x = element_text(size =20), axis.title.y = element_text(size = 20), axis.text.x = element_text(size = 2
0), legend.text = element_text(size =20))
```

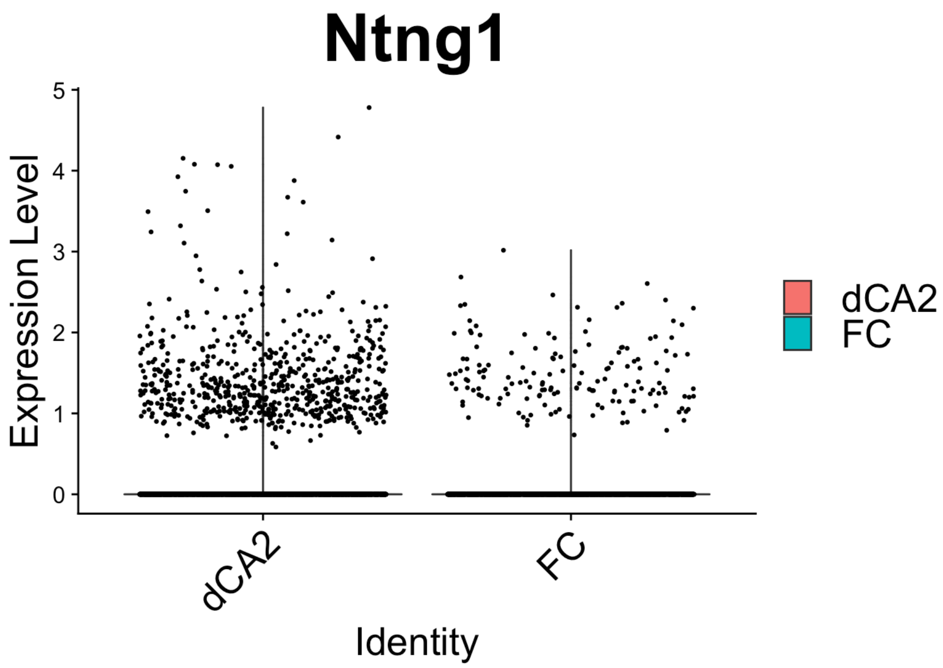


```
ggsave(file="/Users/lees44/R_project/FC_vs_dCA2_comparison_vector_integration_042821/Ntng1_FCvsCA2.pdf", width=7,
height=10)
```

```
average<-(AverageExpression(object = dCA2.integrated, features = "Scn1a", group.by = "Group"))
average
```

```
## $RNA
##          dCA2       FC
## [1,] 4.247062 1.919833
##
## $integrated
##          dCA2       FC
## [1,] 4.210497 3.825983
```

```
AverageExpression(dCA2.integrated, ident.1 = "dCA2", ident.2 = "FC", features = "Scn1a")
```

```
## The following functions and any applicable methods accept the dots: CreateSeuratObject
```

```
## $RNA
##          dCA2       FC
## [1,] 4.247062 1.919833
##
## $integrated
##          dCA2       FC
## [1,] 4.210497 3.825983
```

plot by subsetting by sex within the region 08122021

```
dCA2.integrated@meta.data$Sex = "no"
dCA2.integrated@meta.data$Sex[dCA2.integrated@meta.data$protocol == "dCA2_Fpos"] = "dCA2 female"
dCA2.integrated@meta.data$Sex[dCA2.integrated@meta.data$protocol == "dCA2_Mpos"] = "dCA2 male"
dCA2.integrated@meta.data$Sex[dCA2.integrated@meta.data$protocol == "FC_Fpos"] = "FC female"
dCA2.integrated@meta.data$Sex[dCA2.integrated@meta.data$protocol == "FC_Mpos"] = "FC male"
table(dCA2.integrated@meta.data$Sex)
```

```
##
## dCA2 female   dCA2 male   FC female     FC male
##        1138        1981        1566        1243
```

```
Idents(object = dCA2.integrated) <- "Sex"
```

```
dCA2_subsetbysex <- subset(x = dCA2.integrated, idents = c("FC female", "FC male"), invert = TRUE)
p4<-DimPlot(dCA2_subsetbysex, reduction = "umap", group.by = "Sex", pt=1)
plot_grid(p4)
```

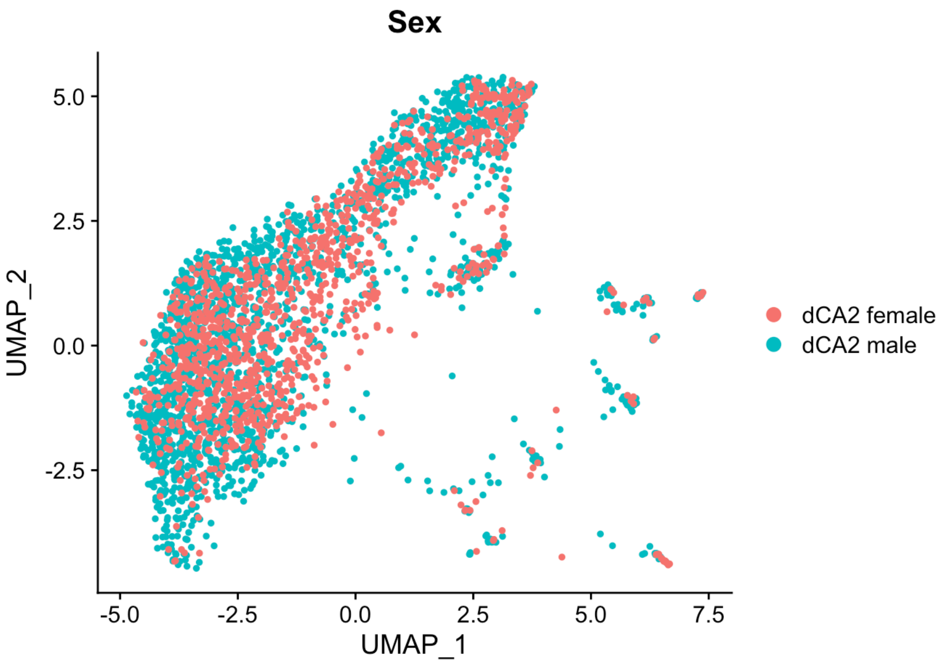


```
ggsave(file="/Users/lees44/R_project/FC_vs_dCA2_comparison_vector_integration_042821/dCA2_UMAP_by_sex.pdf",width=
10,height=10)
```

```
p5<-DimPlot(dCA2_subsetbysex, reduction = "umap", split.by = "Sex", pt=1)
plot_grid(p5)
```

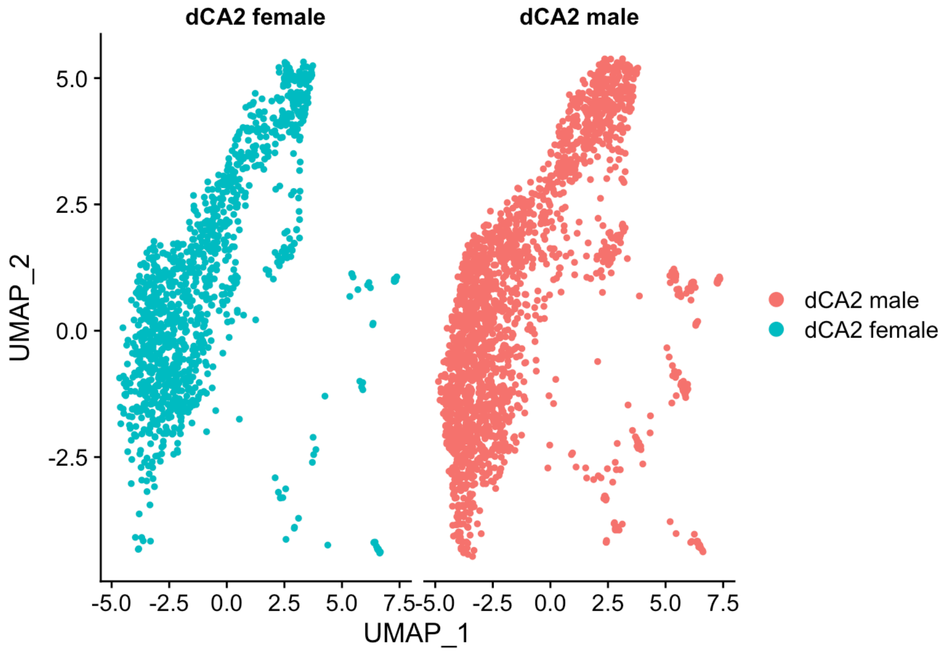


```
ggsave(file="/Users/lees44/R_project/FC_vs_dCA2_comparison_vector_integration_042821/dCA2_UMAP_splitby_sex.pdf",w
idth=10,height=10)
```

```
FC_subsetbysex <- subset(x = dCA2.integrated, idents = c("dCA2 female", "dCA2 male"), invert = TRUE)
p4<-DimPlot(FC_subsetbysex, reduction = "umap", group.by = "Sex", pt=1)
plot_grid(p4)
```

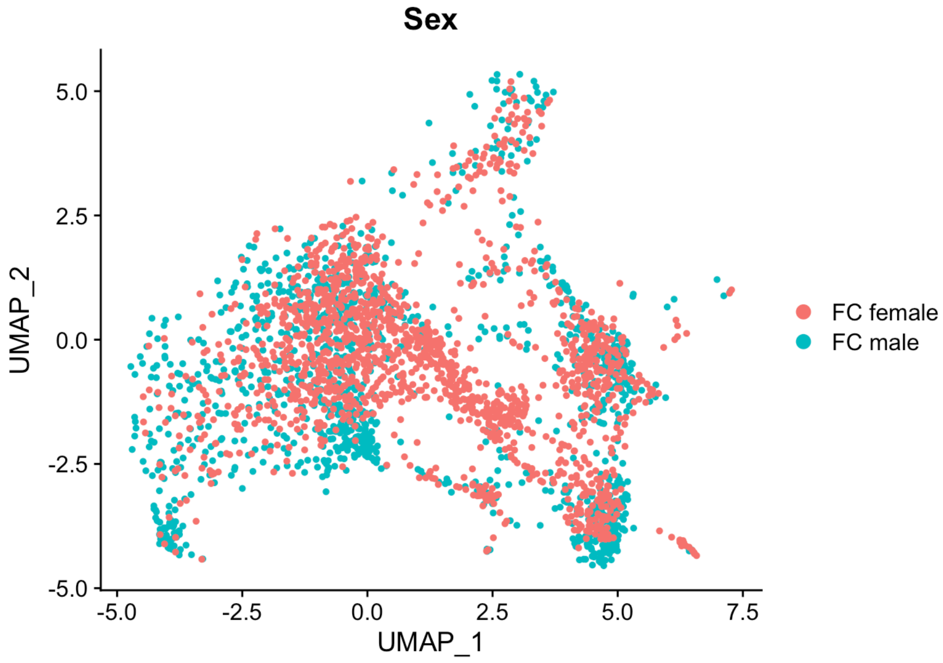


```
ggsave(file="/Users/lees44/R_project/FC_vs_dCA2_comparison_vector_integration_042821/FC2_UMAP_by_sex.pdf",width=1
0,height=10)
```

```
p5<-DimPlot(FC_subsetbysex, reduction = "umap", split.by = "Sex", pt=1)
plot_grid(p5)
```

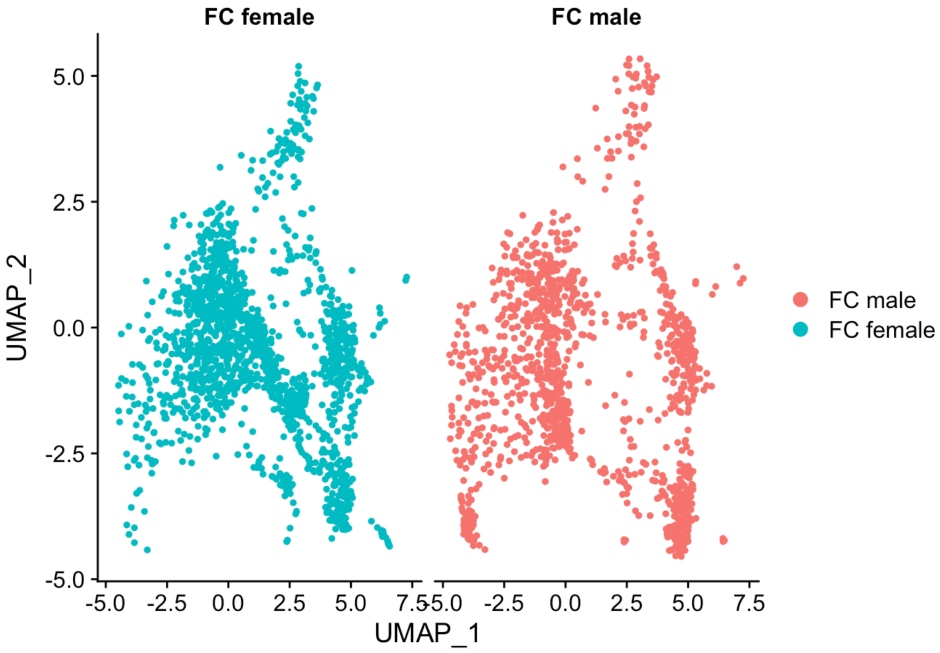


```
ggsave(file="/Users/lees44/R_project/FC_vs_dCA2_comparison_vector_integration_042821/FC2_UMAP_splitby_sex.pdf",wi
dth=10,height=10)
```

```
Idents(object = dCA2.integrated) <- "Type"
```
