## Supplemental table1 for "Investigation of the Fasciola Cinereum, Absent in BTBR mice, and Comparison with the Hippocampal Area CA2"

| | p_val | avg_log2FC | pct.1 | pct.2 | p_val_adj |
|---|---|---|---|---|---|
| Sphkap | 0 | -1.26964 | 0.261 | 0.745 | 0 |
| Cacnb2 | 0 | -1.1937076 | 0.412 | 0.84 | 0 |
| Nebl | 0 | -1.1155057 | 0.719 | 0.956 | 0 |
| Snhg11 | 0 | -1.2958831 | 0.989 | 0.995 | 0 |
| Sms | 0 | -1.1882048 | 0.6 | 0.924 | 0 |
| Trim2 | 0 | -1.118515 | 0.611 | 0.933 | 0 |
| Asph | 0 | -1.3768794 | 0.677 | 0.931 | 0 |
| Epha5 | 0 | 0.8209458 | 0.999 | 0.975 | 0 |
| Akap13 | 0 | -2.7928265 | 0.28 | 0.924 | 0 |
| Socs2 | 0 | -1.2298497 | 0.191 | 0.703 | 0 |
| Cpne7 | 0 | -2.7864114 | 0.189 | 0.901 | 0 |
| Cpne6 | 0 | -1.9279874 | 0.302 | 0.882 | 0 |
| Opcml | 0 | -1.572709 | 0.557 | 0.923 | 0 |
| Adcy1 | 0 | 1.20537206 | 0.963 | 0.822 | 0 |
| Rpl26 | 0 | -1.3029142 | 0.27 | 0.758 | 0 |
| Igfbp4 | 0 | 2.13898758 | 0.67 | 0.094 | 0 |
| Brd9 | 0 | -0.9816338 | 0.879 | 0.974 | 0 |
| Mef2c | 0 | 1.67400172 | 0.642 | 0.183 | 0 |
| Ddx17 | 0 | 0.8792549 | 0.978 | 0.93 | 0 |
| Kalrn | 0 | 0.62120767 | 0.996 | 0.974 | 0 |
| Cadm2 | 0 | 1.42340571 | 0.953 | 0.798 | 0 |
| Trpm3 | 0 | 1.56528263 | 0.78 | 0.352 | 0 |
| Spaar | 3.61E-304 | 1.42455776 | 0.41 | 0.02 | 5.32E-300 |
| Elavl2 | 2.30E-301 | -1.1393007 | 0.371 | 0.843 | 3.40E-297 |
| Grik4 | 2.37E-297 | -1.1911354 | 0.164 | 0.648 | 3.49E-293 |
| Celf2 | 1.85E-290 | 0.61922202 | 0.995 | 0.978 | 2.73E-286 |
| Notch2 | 1.07E-289 | 1.54921208 | 0.473 | 0.071 | 1.58E-285 |
| Tenm2 | 3.68E-282 | -1.2021324 | 0.027 | 0.427 | 5.43E-278 |
| Rnf182 | 9.32E-281 | -1.2903942 | 0.365 | 0.785 | 1.37E-276 |
| Gpi1 | 6.17E-280 | -1.1014338 | 0.568 | 0.887 | 9.10E-276 |
| Dock4 | 1.13E-277 | -0.7253951 | 0.905 | 0.968 | 1.66E-273 |
| Cald1 | 2.36E-275 | 1.45616929 | 0.503 | 0.096 | 3.48E-271 |
| Nos1ap | 3.99E-272 | -0.9455727 | 0.712 | 0.934 | 5.89E-268 |
| Echdc2 | 1.46E-266 | -1.2409369 | 0.244 | 0.691 | 2.15E-262 |
| Gm2115 | 2.49E-264 | -1.1501792 | 0.074 | 0.49 | 3.67E-260 |
| Tafa1 | 4.43E-260 | -0.7684315 | 0.949 | 0.961 | 6.53E-256 |
| Col4a2 | 2.13E-259 | -1.1877184 | 0.332 | 0.743 | 3.14E-255 |
| Peak1 | 3.72E-255 | -1.1861423 | 0.088 | 0.497 | 5.48E-251 |
| Adgrl2 | 3.79E-255 | 1.36919249 | 0.358 | 0.017 | 5.59E-251 |
| Col11a1 | 1.73E-253 | -1.3306561 | 0.267 | 0.703 | 2.56E-249 |
| Slc4a3 | 3.55E-248 | -1.0073883 | 0.457 | 0.834 | 5.24E-244 |
| Grin2a | 9.67E-245 | -0.6462641 | 0.945 | 0.973 | 1.43E-240 |

| | | | | | |
|---|---|---|---|---|---|
| Samd5 | 3.52E-243 | -1.086162 | 0.125 | 0.543 | 5.19E-239 |
| Chka | 1.36E-239 | -0.747464 | 0.878 | 0.965 | 2.00E-235 |
| Arhgap6 | 7.87E-238 | 1.35822205 | 0.312 | 0.005 | 1.16E-233 |
| Fat3 | 9.36E-236 | 1.03194588 | 0.853 | 0.702 | 1.38E-231 |
| Cacna1d | 1.58E-234 | -0.8177982 | 0.755 | 0.935 | 2.32E-230 |
| Zfp536 | 1.39E-227 | 1.06070555 | 0.328 | 0.014 | 2.05E-223 |
| Efr3b | 5.14E-227 | -1.0250725 | 0.344 | 0.736 | 7.58E-223 |
| Plcb1 | 1.07E-225 | -0.9516355 | 0.471 | 0.836 | 1.57E-221 |
| R3hdm1 | 1.44E-220 | 0.54256469 | 0.992 | 0.981 | 2.12E-216 |
| Ube3a | 1.33E-218 | -0.5764924 | 0.975 | 0.981 | 1.96E-214 |
| Dclk1 | 2.17E-218 | 0.72521969 | 0.977 | 0.939 | 3.20E-214 |
| Ints6l | 1.46E-217 | 1.04488752 | 0.78 | 0.56 | 2.16E-213 |
| Onecut2 | 1.24E-216 | -1.0085272 | 0.269 | 0.687 | 1.83E-212 |
| Atp6v0b | 7.81E-209 | -0.4947375 | 0.986 | 0.991 | 1.15E-204 |
| Ccnd2 | 1.53E-207 | -1.3345454 | 0.195 | 0.583 | 2.25E-203 |
| Camkv | 5.37E-207 | -0.7571621 | 0.774 | 0.937 | 7.93E-203 |
| Auts2 | 1.05E-206 | -0.5012744 | 0.985 | 0.988 | 1.54E-202 |
| Syn2 | 7.00E-205 | -0.9661031 | 0.364 | 0.743 | 1.03E-200 |
| Vxn | 3.62E-204 | -0.993597 | 0.088 | 0.448 | 5.35E-200 |
| Unc80 | 1.47E-203 | 0.67937267 | 0.958 | 0.923 | 2.16E-199 |
| Ttr | 1.97E-203 | -1.4802916 | 0.039 | 0.367 | 2.91E-199 |
| Cpne8 | 1.58E-200 | -0.8786676 | 0.014 | 0.31 | 2.33E-196 |
| Igsf3 | 2.78E-200 | 1.12095532 | 0.358 | 0.049 | 4.10E-196 |
| Prdx6 | 2.07E-196 | -0.9930064 | 0.274 | 0.668 | 3.05E-192 |
| Map3k15 | 6.57E-196 | 1.22784888 | 0.478 | 0.154 | 9.70E-192 |
| 2010300C02I | 1.52E-191 | 0.85095994 | 0.863 | 0.748 | 2.24E-187 |
| Pfkl | 1.23E-189 | -0.89584 | 0.218 | 0.605 | 1.82E-185 |
| Pkp2 | 5.85E-188 | -0.8853375 | 0.365 | 0.737 | 8.63E-184 |
| Igf1r | 6.30E-188 | -0.9457996 | 0.39 | 0.755 | 9.29E-184 |
| Kcna1 | 2.20E-187 | -0.8524419 | 0.552 | 0.857 | 3.25E-183 |
| Scn1a | 3.40E-187 | -0.8456242 | 0.436 | 0.801 | 5.02E-183 |
| Sema6d | 6.14E-187 | 0.98680818 | 0.312 | 0.031 | 9.06E-183 |
| Syne1 | 1.00E-185 | 0.99707753 | 0.766 | 0.556 | 1.48E-181 |
| Entpd6 | 1.31E-185 | -0.9343211 | 0.349 | 0.71 | 1.94E-181 |
| Traip | 3.27E-184 | -0.9324236 | 0.162 | 0.529 | 4.82E-180 |
| Atp8a1 | 7.53E-183 | 0.84532049 | 0.841 | 0.706 | 1.11E-178 |
| Nmrk1 | 8.43E-183 | -0.9227872 | 0.186 | 0.559 | 1.24E-178 |
| Cacna1e | 2.09E-181 | 0.71909132 | 0.913 | 0.844 | 3.08E-177 |
| Ptn | 4.83E-181 | 1.0290888 | 0.615 | 0.318 | 7.13E-177 |
| Cul4a | 3.77E-180 | -0.9011293 | 0.169 | 0.536 | 5.57E-176 |
| Rph3a | 2.57E-179 | 1.00057287 | 0.342 | 0.053 | 3.79E-175 |
| 5031439G07 | 2.58E-178 | -0.9121324 | 0.123 | 0.472 | 3.81E-174 |
| Scube1 | 4.49E-177 | -0.8983201 | 0.013 | 0.28 | 6.62E-173 |

| | | | | | |
|---|---|---|---|---|---|
| Sash1 | 5.75E-177 | 1.13290457 | 0.627 | 0.348 | 8.49E-173 |
| Trpc5 | 1.40E-173 | -0.882105 | 0.292 | 0.665 | 2.06E-169 |
| Parp8 | 1.41E-172 | -0.807426 | 0.011 | 0.27 | 2.08E-168 |
| Cntn3 | 1.17E-171 | 0.97550541 | 0.272 | 0.021 | 1.72E-167 |
| Osbpl3 | 2.45E-170 | -0.9123316 | 0.098 | 0.423 | 3.61E-166 |
| Ntm | 4.41E-170 | -1.123169 | 0.062 | 0.365 | 6.51E-166 |
| Tmem108 | 3.25E-169 | -0.8355625 | 0.591 | 0.869 | 4.80E-165 |
| Timp4 | 2.01E-168 | -0.8863593 | 0.2 | 0.566 | 2.97E-164 |
| Atp2a2 | 4.77E-167 | 0.73242307 | 0.889 | 0.819 | 7.04E-163 |
| Dpyd | 1.64E-166 | -0.8518723 | 0.121 | 0.453 | 2.41E-162 |
| Nrxn3 | 7.91E-166 | -0.4912186 | 0.971 | 0.969 | 1.17E-161 |
| Gria2 | 6.41E-164 | -0.2541448 | 0.999 | 0.99 | 9.45E-160 |
| Dync1i2 | 1.92E-163 | -0.6938996 | 0.759 | 0.919 | 2.83E-159 |
| Nrgn | 2.14E-161 | -0.836348 | 0.394 | 0.75 | 3.15E-157 |
| Fgfr1 | 1.84E-159 | -0.8022134 | 0.466 | 0.782 | 2.72E-155 |
| Syn3 | 2.46E-159 | 0.91131519 | 0.277 | 0.03 | 3.62E-155 |
| Il1rap | 7.49E-159 | 1.03856819 | 0.621 | 0.367 | 1.10E-154 |
| Rnf165 | 8.75E-159 | -0.7151088 | 0.649 | 0.888 | 1.29E-154 |
| Slc24a2 | 8.78E-159 | 0.60370648 | 0.941 | 0.913 | 1.30E-154 |
| Grb14 | 5.05E-156 | -0.8133558 | 0.275 | 0.634 | 7.45E-152 |
| Wipf3 | 2.56E-155 | 0.57435426 | 0.969 | 0.933 | 3.77E-151 |
| Sox5 | 5.64E-154 | -0.81802 | 0.405 | 0.744 | 8.33E-150 |
| Arhgef26 | 8.40E-154 | -0.73549 | 0.6 | 0.857 | 1.24E-149 |
| Ly6e | 2.43E-151 | -0.7432649 | 0.542 | 0.831 | 3.58E-147 |
| Rapgef4 | 1.82E-147 | 0.7986043 | 0.806 | 0.657 | 2.68E-143 |
| Pabpn1 | 6.59E-144 | -0.6333387 | 0.705 | 0.922 | 9.72E-140 |
| Gria1 | 1.23E-141 | 0.50846961 | 0.972 | 0.956 | 1.82E-137 |
| Rasl11b | 9.76E-141 | -0.7799078 | 0.223 | 0.563 | 1.44E-136 |
| Stard5 | 1.90E-140 | -0.8161957 | 0.314 | 0.65 | 2.81E-136 |
| Trank1 | 4.68E-140 | 0.7934554 | 0.795 | 0.66 | 6.91E-136 |
| Mtdh | 2.76E-139 | 0.73470065 | 0.838 | 0.736 | 4.07E-135 |
| Gabra2 | 9.33E-139 | -0.7296916 | 0.466 | 0.782 | 1.38E-134 |
| Dapk1 | 1.41E-137 | 0.75920747 | 0.802 | 0.66 | 2.08E-133 |
| Hnrnpu | 4.40E-136 | 0.59701838 | 0.915 | 0.887 | 6.49E-132 |
| Tenm4 | 7.67E-136 | -0.7493618 | 0.109 | 0.403 | 1.13E-131 |
| Dpp6 | 1.02E-135 | 0.62714412 | 0.906 | 0.84 | 1.51E-131 |
| Mas1 | 1.88E-134 | -0.7457865 | 0.159 | 0.469 | 2.78E-130 |
| Col4a1 | 3.33E-133 | -0.7854815 | 0.343 | 0.66 | 4.91E-129 |
| Rb1 | 5.85E-133 | 0.99500897 | 0.516 | 0.272 | 8.63E-129 |
| Nrp1 | 2.14E-132 | -0.6131292 | 0.731 | 0.899 | 3.16E-128 |
| Zbtb20 | 2.25E-132 | 0.32014281 | 0.999 | 0.99 | 3.32E-128 |
| Snrnp70 | 2.91E-132 | -0.3562658 | 0.984 | 0.992 | 4.29E-128 |
| Nr2f1 | 3.15E-132 | -0.6834061 | 0.03 | 0.259 | 4.64E-128 |

| | | | | | |
|---|---|---|---|---|---|
| Slc6a7 | 1.30E-131 | -0.7029869 | 0.105 | 0.391 | 1.92E-127 |
| Dgkh | 5.39E-129 | 0.86287227 | 0.641 | 0.417 | 7.95E-125 |
| Mast4 | 1.58E-128 | -0.7014694 | 0.578 | 0.842 | 2.33E-124 |
| Robo2 | 1.76E-127 | -0.7038522 | 0.143 | 0.437 | 2.60E-123 |
| Tacc1 | 2.28E-127 | 0.75360924 | 0.807 | 0.656 | 3.37E-123 |
| Kcnk2 | 3.66E-127 | 0.66676801 | 0.935 | 0.804 | 5.40E-123 |
| Slc24a5 | 4.20E-127 | 0.80971981 | 0.847 | 0.769 | 6.20E-123 |
| Cdh10 | 1.05E-126 | -0.7743114 | 0.101 | 0.375 | 1.55E-122 |
| Tigd2 | 1.29E-126 | -0.7146982 | 0.361 | 0.685 | 1.91E-122 |
| Ndst1 | 2.48E-126 | 0.89822918 | 0.798 | 0.676 | 3.66E-122 |
| Dmd | 7.52E-126 | -0.6946536 | 0.455 | 0.76 | 1.11E-121 |
| Tspan33 | 1.18E-125 | -0.7066146 | 0.062 | 0.31 | 1.74E-121 |
| Rock2 | 4.48E-125 | 0.68500679 | 0.845 | 0.774 | 6.61E-121 |
| Bin1 | 9.23E-125 | -0.7021503 | 0.441 | 0.745 | 1.36E-120 |
| Kcnn2 | 1.53E-124 | 0.75320991 | 0.786 | 0.661 | 2.26E-120 |
| Map1b | 2.45E-123 | 0.43336462 | 0.992 | 0.979 | 3.62E-119 |
| Homer3 | 2.68E-123 | -0.6992363 | 0.107 | 0.379 | 3.95E-119 |
| Gapdh | 1.63E-122 | -0.7461126 | 0.335 | 0.657 | 2.41E-118 |
| Srsf7 | 2.33E-121 | -0.5130644 | 0.832 | 0.955 | 3.44E-117 |
| Fez2 | 2.68E-120 | -0.6887756 | 0.063 | 0.304 | 3.95E-116 |
| Aldoa | 2.71E-120 | -0.6792718 | 0.389 | 0.714 | 3.99E-116 |
| Palmd | 1.16E-118 | -0.7052288 | 0.119 | 0.388 | 1.71E-114 |
| Rbfox1 | 7.27E-118 | -0.6480491 | 0.426 | 0.736 | 1.07E-113 |
| Gins2 | 2.81E-117 | -0.761724 | 0.206 | 0.502 | 4.14E-113 |
| Tcf12 | 1.52E-116 | -0.6918141 | 0.325 | 0.638 | 2.25E-112 |
| Lrp1b | 5.57E-116 | -0.3941201 | 0.984 | 0.983 | 8.21E-112 |
| Rims1 | 5.77E-115 | 0.61274992 | 0.889 | 0.849 | 8.52E-111 |
| Tcf4 | 7.91E-114 | 0.34452825 | 0.996 | 0.985 | 1.17E-109 |
| Tmed9 | 1.99E-113 | -0.6393072 | 0.48 | 0.776 | 2.93E-109 |
| Prkca | 5.47E-113 | 0.51575271 | 0.965 | 0.928 | 8.07E-109 |
| Efhd2 | 8.00E-113 | 0.7900629 | 0.27 | 0.061 | 1.18E-108 |
| Kif5c | 1.13E-112 | 0.71485592 | 0.827 | 0.748 | 1.67E-108 |
| Cnrip1 | 1.63E-112 | -0.6253624 | 0.566 | 0.823 | 2.41E-108 |
| Sat1 | 1.49E-111 | 1.05083877 | 0.431 | 0.202 | 2.21E-107 |
| Dgkg | 2.06E-111 | -0.5562677 | 0.718 | 0.895 | 3.03E-107 |
| Alcam | 8.37E-110 | 0.81954159 | 0.295 | 0.08 | 1.23E-105 |
| Plxnc1 | 2.13E-109 | 0.87533366 | 0.387 | 0.16 | 3.14E-105 |
| Tiam2 | 1.60E-108 | 0.8726684 | 0.463 | 0.228 | 2.36E-104 |
| Pebp1 | 1.74E-108 | -0.6126066 | 0.563 | 0.835 | 2.57E-104 |
| Kcnk9 | 2.67E-108 | -0.6673034 | 0.134 | 0.396 | 3.94E-104 |
| Hpca | 5.79E-108 | -0.5302287 | 0.745 | 0.919 | 8.54E-104 |
| Crybb3 | 6.62E-108 | -0.6800733 | 0.077 | 0.31 | 9.77E-104 |
| Cpt1c | 8.45E-108 | -0.6156691 | 0.487 | 0.757 | 1.25E-103 |

| | | | | | |
|---|---|---|---|---|---|
| Lipe | 1.52E-107 | -0.6245115 | 0.106 | 0.359 | 2.24E-103 |
| Kcnma1 | 9.25E-106 | 0.57265311 | 0.901 | 0.866 | 1.36E-101 |
| Stmn2 | 2.94E-105 | -0.6734121 | 0.329 | 0.629 | 4.33E-101 |
| Kcnj4 | 3.25E-105 | 0.84281586 | 0.316 | 0.102 | 4.80E-101 |
| Ywhah | 4.19E-105 | -0.6563029 | 0.392 | 0.682 | 6.18E-101 |
| Atpif1 | 8.77E-105 | -0.645431 | 0.404 | 0.704 | 1.29E-100 |
| Ryr2 | 8.96E-105 | 0.51950086 | 0.919 | 0.883 | 1.32E-100 |
| Xkr6 | 9.53E-105 | -0.680072 | 0.177 | 0.449 | 1.41E-100 |
| Cnksr2 | 6.24E-104 | 0.6892102 | 0.776 | 0.69 | 9.20E-100 |
| Nr3c2 | 1.06E-103 | 0.73625836 | 0.692 | 0.557 | 1.57E-99 |
| Caskin1 | 7.31E-103 | -0.6387678 | 0.319 | 0.621 | 1.08E-98 |
| Zranb2 | 1.21E-102 | -0.3201023 | 0.985 | 0.983 | 1.78E-98 |
| Htr1f | 1.39E-102 | -0.5999165 | 0.086 | 0.318 | 2.06E-98 |
| Luzp2 | 1.72E-102 | -0.6512628 | 0.188 | 0.464 | 2.54E-98 |
| Inhba | 1.74E-102 | 0.79799167 | 0.258 | 0.063 | 2.57E-98 |
| Nrxn2 | 1.04E-101 | -0.5923063 | 0.587 | 0.827 | 1.54E-97 |
| Hnrnph1 | 1.33E-100 | 0.70213747 | 0.708 | 0.586 | 1.97E-96 |
| Dock11 | 4.33E-100 | 0.85146195 | 0.471 | 0.258 | 6.40E-96 |
| Selenow | 5.51E-100 | -0.6294394 | 0.366 | 0.673 | 8.14E-96 |
| Kif1b | 2.34E-99 | 0.58157043 | 0.888 | 0.848 | 3.45E-95 |
| Stxbp5l | 1.30E-98 | -0.4197302 | 0.921 | 0.961 | 1.92E-94 |
| Cacna1a | 2.92E-98 | -0.403285 | 0.927 | 0.963 | 4.31E-94 |
| Mrps5 | 4.15E-98 | -0.5656866 | 0.528 | 0.796 | 6.12E-94 |
| Ppib | 8.05E-98 | -0.5698468 | 0.586 | 0.82 | 1.19E-93 |
| Ackr1 | 1.30E-97 | -0.5492499 | 0.072 | 0.288 | 1.92E-93 |
| Unc13b | 2.46E-97 | -0.5861156 | 0.428 | 0.711 | 3.63E-93 |
| Tgfb2 | 5.58E-97 | -0.6016448 | 0.067 | 0.277 | 8.24E-93 |
| Sorl1 | 9.08E-97 | 0.68155061 | 0.741 | 0.633 | 1.34E-92 |
| Cadm3 | 1.82E-96 | -0.6189082 | 0.232 | 0.508 | 2.69E-92 |
| Ndufa10 | 2.45E-96 | -0.5760281 | 0.37 | 0.662 | 3.61E-92 |
| Dmxl1 | 4.48E-96 | 0.64582033 | 0.793 | 0.718 | 6.61E-92 |
| Cdc37l1 | 7.72E-96 | 0.49022282 | 0.914 | 0.884 | 1.14E-91 |
| Epha7 | 4.73E-95 | 0.68719233 | 0.728 | 0.615 | 6.99E-91 |
| Lsamp | 1.40E-94 | -0.4489273 | 0.823 | 0.946 | 2.06E-90 |
| Thoc1 | 7.56E-94 | -0.5304414 | 0.593 | 0.828 | 1.12E-89 |
| Net1 | 1.13E-93 | 0.8085282 | 0.298 | 0.098 | 1.67E-89 |
| Bcl9 | 1.18E-93 | -0.5954655 | 0.289 | 0.576 | 1.74E-89 |
| Trhde | 4.36E-93 | -0.5828564 | 0.5 | 0.772 | 6.43E-89 |
| Pid1 | 2.26E-92 | -0.5950891 | 0.237 | 0.518 | 3.34E-88 |
| B3gat1 | 8.39E-91 | -0.5722978 | 0.161 | 0.412 | 1.24E-86 |
| Fkbp1a | 5.71E-90 | -0.5621288 | 0.479 | 0.747 | 8.43E-86 |
| Myo18a | 1.16E-89 | -0.5800301 | 0.475 | 0.731 | 1.71E-85 |
| Kcnb2 | 1.67E-89 | 0.80862599 | 0.38 | 0.176 | 2.47E-85 |

| | | | | | |
|---|---|---|---|---|---|
| Ly6h | 2.60E-89 | -0.4645651 | 0.818 | 0.937 | 3.83E-85 |
| Arhgap20 | 3.33E-89 | -0.5586633 | 0.073 | 0.277 | 4.92E-85 |
| Nefm | 1.85E-88 | -0.608857 | 0.076 | 0.28 | 2.73E-84 |
| Zfhx4 | 3.07E-88 | 0.92544657 | 0.363 | 0.163 | 4.52E-84 |
| Prpf4b | 4.56E-88 | -0.3420982 | 0.974 | 0.988 | 6.73E-84 |
| Mcf2l | 7.73E-88 | -0.5801475 | 0.284 | 0.562 | 1.14E-83 |
| Lyst | 9.22E-88 | 0.72682009 | 0.639 | 0.497 | 1.36E-83 |
| Mctp1 | 1.01E-87 | 0.56101755 | 0.849 | 0.789 | 1.48E-83 |
| Sorcs2 | 1.16E-86 | 0.74122797 | 0.588 | 0.411 | 1.71E-82 |
| Igsf9b | 1.69E-86 | 0.71540071 | 0.549 | 0.356 | 2.49E-82 |
| Asap2 | 1.76E-86 | 0.78604993 | 0.326 | 0.128 | 2.59E-82 |
| Flna | 8.57E-86 | 0.75975399 | 0.639 | 0.486 | 1.26E-81 |
| Lynx1 | 2.48E-85 | -0.5225814 | 0.661 | 0.857 | 3.66E-81 |
| Nfia | 2.90E-85 | 0.84446008 | 0.552 | 0.375 | 4.28E-81 |
| Dgkb | 8.87E-85 | -0.5395469 | 0.08 | 0.282 | 1.31E-80 |
| Kcnc1 | 1.12E-84 | -0.6024487 | 0.352 | 0.623 | 1.66E-80 |
| Rnps1 | 2.51E-84 | -0.4884933 | 0.631 | 0.846 | 3.71E-80 |
| Cdk14 | 7.88E-84 | -0.4584446 | 0.75 | 0.903 | 1.16E-79 |
| Tlk1 | 1.56E-83 | 0.79075402 | 0.485 | 0.302 | 2.30E-79 |
| Crym | 1.64E-83 | -0.5630124 | 0.407 | 0.68 | 2.41E-79 |
| Scn2a | 2.67E-83 | 0.48408174 | 0.905 | 0.883 | 3.94E-79 |
| Tmem181a | 3.45E-83 | -0.5548813 | 0.299 | 0.567 | 5.09E-79 |
| Nefl | 3.45E-83 | -0.5786852 | 0.21 | 0.463 | 5.10E-79 |
| Slc25a23 | 6.89E-83 | -0.558903 | 0.456 | 0.715 | 1.02E-78 |
| Limch1 | 9.20E-83 | 0.70482408 | 0.539 | 0.349 | 1.36E-78 |
| Map2k2 | 1.18E-82 | -0.5217753 | 0.289 | 0.567 | 1.75E-78 |
| Hdac4 | 1.40E-82 | -0.5722741 | 0.317 | 0.585 | 2.07E-78 |
| Slc4a4 | 2.74E-82 | 0.80142626 | 0.428 | 0.233 | 4.04E-78 |
| Nrcam | 3.29E-82 | -0.3956485 | 0.874 | 0.947 | 4.86E-78 |
| Camkk2 | 5.41E-82 | 0.79952297 | 0.553 | 0.38 | 7.98E-78 |
| Gabarapl2 | 6.86E-82 | -0.5914228 | 0.384 | 0.664 | 1.01E-77 |
| Prox2 | 7.50E-82 | -0.5129954 | 0.104 | 0.317 | 1.11E-77 |
| Map4k4 | 1.23E-81 | -0.5429234 | 0.553 | 0.799 | 1.81E-77 |
| Macf1 | 2.19E-81 | 0.43488812 | 0.937 | 0.925 | 3.23E-77 |
| mt-Co1 | 5.12E-81 | 0.73901822 | 0.679 | 0.541 | 7.55E-77 |
| Osbp2 | 1.23E-80 | -0.553671 | 0.305 | 0.569 | 1.82E-76 |
| mt-Nd2 | 1.46E-80 | 0.80495537 | 0.443 | 0.243 | 2.16E-76 |
| Pcdh7 | 4.11E-80 | -0.5665374 | 0.15 | 0.38 | 6.06E-76 |
| Atp5g1 | 4.29E-80 | -0.5106375 | 0.231 | 0.493 | 6.33E-76 |
| Eno1 | 7.95E-80 | -0.5675138 | 0.24 | 0.492 | 1.17E-75 |
| Zdhhc14 | 9.20E-80 | -0.5019497 | 0.087 | 0.286 | 1.36E-75 |
| Mmp16 | 1.14E-79 | 0.70837477 | 0.634 | 0.508 | 1.68E-75 |
| Whamm | 1.18E-79 | -0.5307382 | 0.254 | 0.512 | 1.74E-75 |

| | | | | | |
|---|---|---|---|---|---|
| Dnm3 | 3.80E-79 | -0.5771456 | 0.294 | 0.562 | 5.60E-75 |
| Armc9 | 4.04E-79 | -0.5251448 | 0.449 | 0.713 | 5.97E-75 |
| Ccdc88c | 1.44E-78 | -0.5258912 | 0.267 | 0.534 | 2.12E-74 |
| Zfp30 | 2.58E-78 | -0.5489058 | 0.257 | 0.514 | 3.80E-74 |
| Bcl11b | 2.83E-78 | 0.64001415 | 0.699 | 0.571 | 4.17E-74 |
| Tbata | 6.25E-78 | -0.4808195 | 0.075 | 0.265 | 9.22E-74 |
| Rab3a | 1.07E-77 | -0.5348689 | 0.264 | 0.521 | 1.58E-73 |
| Birc6 | 1.87E-77 | 0.61250757 | 0.708 | 0.61 | 2.76E-73 |
| Taf1d | 1.90E-77 | -0.5057695 | 0.583 | 0.815 | 2.80E-73 |
| Ttc19 | 2.31E-77 | -0.408263 | 0.835 | 0.944 | 3.41E-73 |
| Ryr3 | 2.62E-77 | 0.5629215 | 0.789 | 0.71 | 3.87E-73 |
| Pde4d | 1.61E-76 | -0.4757317 | 0.701 | 0.87 | 2.38E-72 |
| Chst2 | 2.68E-76 | -0.5116303 | 0.126 | 0.34 | 3.95E-72 |
| Lrtm1 | 2.92E-76 | -0.5309532 | 0.54 | 0.775 | 4.31E-72 |
| Tango2 | 3.51E-76 | -0.5331224 | 0.388 | 0.652 | 5.17E-72 |
| Kmt2a | 4.09E-76 | 0.50232074 | 0.86 | 0.825 | 6.03E-72 |
| Ric1 | 4.66E-76 | 0.72419265 | 0.526 | 0.359 | 6.88E-72 |
| Cog5 | 6.29E-76 | 0.78283926 | 0.514 | 0.351 | 9.28E-72 |
| Nwd2 | 7.19E-76 | -0.4959576 | 0.087 | 0.281 | 1.06E-71 |
| Etnk1 | 9.46E-76 | 0.46523488 | 0.896 | 0.878 | 1.40E-71 |
| Shisa4 | 1.10E-75 | -0.6512599 | 0.109 | 0.311 | 1.62E-71 |
| Snx32 | 2.32E-75 | -0.5180896 | 0.314 | 0.573 | 3.43E-71 |
| Grip1 | 4.21E-75 | -0.5306739 | 0.104 | 0.304 | 6.21E-71 |
| Il1rapl1 | 1.72E-74 | 0.53229459 | 0.832 | 0.746 | 2.54E-70 |
| Nell2 | 1.82E-74 | 0.60832917 | 0.729 | 0.665 | 2.69E-70 |
| Raver2 | 2.20E-74 | -0.5617525 | 0.161 | 0.38 | 3.25E-70 |
| Dab1 | 9.80E-74 | 0.59375804 | 0.718 | 0.635 | 1.45E-69 |
| mt-Nd4 | 1.05E-73 | 0.79720971 | 0.515 | 0.336 | 1.56E-69 |
| Mvb12a | 2.00E-73 | -0.4913812 | 0.171 | 0.402 | 2.96E-69 |
| Fbxw7 | 3.93E-73 | 0.65877601 | 0.71 | 0.633 | 5.81E-69 |
| Nrbp2 | 2.21E-72 | -0.49404 | 0.27 | 0.522 | 3.27E-68 |
| Pex5l | 3.86E-72 | -0.5260741 | 0.447 | 0.698 | 5.69E-68 |
| Kcnj3 | 4.37E-72 | 0.71698825 | 0.459 | 0.287 | 6.45E-68 |
| Dgkz | 6.94E-72 | 0.52875396 | 0.805 | 0.744 | 1.02E-67 |
| Jakmip1 | 9.25E-72 | -0.4890856 | 0.138 | 0.35 | 1.36E-67 |
| Cabp7 | 3.14E-71 | 0.73150102 | 0.54 | 0.385 | 4.63E-67 |
| Dnajb4 | 4.28E-71 | -0.4987251 | 0.364 | 0.618 | 6.31E-67 |
| Rgs11 | 3.54E-70 | -0.4691104 | 0.594 | 0.81 | 5.22E-66 |
| Cnot3 | 9.87E-70 | -0.4894763 | 0.461 | 0.708 | 1.46E-65 |
| Elmo1 | 1.34E-69 | 0.71635743 | 0.326 | 0.15 | 1.97E-65 |
| Celf3 | 1.37E-69 | 0.51671452 | 0.835 | 0.778 | 2.02E-65 |
| Spink8 | 1.44E-69 | -0.4890569 | 0.122 | 0.324 | 2.12E-65 |
| Thoc2l | 3.05E-69 | -0.3712016 | 0.86 | 0.954 | 4.50E-65 |

| | | | | | |
|---|---|---|---|---|---|
| Sugp1 | 3.55E-69 | -0.4915695 | 0.365 | 0.613 | 5.24E-65 |
| Ralgapa1 | 3.92E-69 | 0.45745245 | 0.855 | 0.808 | 5.78E-65 |
| Lmo4 | 7.61E-69 | -0.5887208 | 0.187 | 0.405 | 1.12E-64 |
| Spon1 | 8.37E-69 | 0.6675704 | 0.324 | 0.146 | 1.23E-64 |
| Ccbe1 | 8.65E-69 | -0.4687293 | 0.165 | 0.381 | 1.28E-64 |
| Lrp12 | 1.27E-68 | 0.67056039 | 0.279 | 0.114 | 1.88E-64 |
| Rbm3 | 2.02E-68 | 0.62365893 | 0.634 | 0.518 | 2.98E-64 |
| Rab3c | 3.20E-68 | -0.5378215 | 0.305 | 0.541 | 4.72E-64 |
| Ubb | 3.51E-68 | -0.5196997 | 0.486 | 0.733 | 5.18E-64 |
| mt-Atp6 | 5.49E-68 | 0.63389264 | 0.557 | 0.385 | 8.11E-64 |
| Csmd1 | 3.05E-67 | 0.41268274 | 0.935 | 0.93 | 4.51E-63 |
| Cfap100 | 3.33E-67 | -0.4932739 | 0.12 | 0.311 | 4.92E-63 |
| Sybu | 5.47E-67 | -0.4948689 | 0.279 | 0.529 | 8.07E-63 |
| Selenom | 8.88E-67 | -0.5060022 | 0.46 | 0.707 | 1.31E-62 |
| 2300009A05I | 1.67E-66 | -0.47683 | 0.216 | 0.445 | 2.47E-62 |
| Kcnj6 | 2.51E-66 | 0.49672683 | 0.809 | 0.766 | 3.70E-62 |
| Armh4 | 3.06E-66 | -0.4451646 | 0.081 | 0.253 | 4.51E-62 |
| Gas2l1 | 3.22E-66 | -0.4446779 | 0.106 | 0.296 | 4.75E-62 |
| Cck | 3.63E-66 | -0.3891635 | 0.67 | 0.885 | 5.36E-62 |
| Slc38a1 | 4.56E-66 | 0.68981093 | 0.558 | 0.425 | 6.72E-62 |
| Ttc21b | 5.38E-66 | -0.4847367 | 0.265 | 0.499 | 7.95E-62 |
| Osbpl1a | 2.43E-65 | -0.4699868 | 0.11 | 0.298 | 3.58E-61 |
| Nr1d2 | 2.95E-65 | 0.62900719 | 0.323 | 0.153 | 4.35E-61 |
| Pisd | 3.79E-65 | -0.5005525 | 0.529 | 0.748 | 5.60E-61 |
| Ccdc88a | 4.88E-65 | 0.51332396 | 0.765 | 0.705 | 7.20E-61 |
| Ank2 | 1.09E-64 | 0.2729278 | 0.994 | 0.987 | 1.60E-60 |
| Sez6l2 | 6.68E-64 | -0.4836895 | 0.346 | 0.59 | 9.85E-60 |
| Pex3 | 7.15E-64 | -0.4605875 | 0.257 | 0.494 | 1.06E-59 |
| Marchf1 | 1.62E-63 | -0.4987475 | 0.093 | 0.268 | 2.38E-59 |
| Marchf2 | 1.72E-63 | -0.4630995 | 0.263 | 0.501 | 2.53E-59 |
| Man1a2 | 2.64E-63 | -0.4049101 | 0.864 | 0.944 | 3.89E-59 |
| Dennd4a | 3.08E-63 | 0.6703568 | 0.576 | 0.45 | 4.55E-59 |
| Nav3 | 5.85E-63 | -0.4609957 | 0.565 | 0.779 | 8.64E-59 |
| Dock3 | 6.26E-63 | 0.65695635 | 0.455 | 0.291 | 9.24E-59 |
| Hunk | 1.04E-62 | 0.68263684 | 0.448 | 0.285 | 1.53E-58 |
| Grm5 | 1.11E-62 | -0.3022564 | 0.939 | 0.964 | 1.64E-58 |
| Lrrtm1 | 1.26E-62 | -0.4765593 | 0.107 | 0.283 | 1.86E-58 |
| Pycard | 2.17E-62 | -0.4689929 | 0.096 | 0.273 | 3.20E-58 |
| Foxp1 | 3.74E-62 | -0.4926909 | 0.413 | 0.648 | 5.52E-58 |
| Srgap3 | 3.82E-62 | -0.4503228 | 0.669 | 0.837 | 5.64E-58 |
| Hsp90b1 | 4.22E-62 | 0.67898494 | 0.521 | 0.383 | 6.22E-58 |
| Gpm6b | 4.68E-62 | -0.3977938 | 0.811 | 0.929 | 6.91E-58 |
| Fzd3 | 1.16E-61 | 0.63785558 | 0.296 | 0.135 | 1.71E-57 |

| | | | | | |
|---|---|---|---|---|---|
| Eid1 | 1.55E-61 | -0.4268354 | 0.523 | 0.762 | 2.29E-57 |
| Rbm33 | 2.81E-61 | 0.64381534 | 0.548 | 0.419 | 4.15E-57 |
| Susd4 | 3.87E-61 | -0.434632 | 0.367 | 0.619 | 5.71E-57 |
| Ptprd | 4.74E-61 | -0.3281223 | 0.944 | 0.966 | 6.99E-57 |
| Dnah9 | 5.53E-61 | -0.4557383 | 0.295 | 0.529 | 8.15E-57 |
| Knop1 | 9.86E-61 | -0.4353737 | 0.361 | 0.614 | 1.46E-56 |
| Calm2 | 1.08E-60 | -0.3339194 | 0.832 | 0.941 | 1.60E-56 |
| Basp1 | 1.24E-60 | -0.4726694 | 0.497 | 0.723 | 1.83E-56 |
| Arhgdig | 2.92E-60 | -0.432138 | 0.136 | 0.325 | 4.31E-56 |
| Polb | 3.70E-60 | -0.417242 | 0.583 | 0.799 | 5.45E-56 |
| Mpp3 | 5.42E-60 | -0.4517805 | 0.388 | 0.635 | 8.00E-56 |
| Shisa6 | 6.04E-60 | -0.4671921 | 0.24 | 0.458 | 8.91E-56 |
| Rnpepl1 | 8.51E-60 | -0.4352403 | 0.18 | 0.39 | 1.26E-55 |
| Wbp11 | 1.10E-59 | -0.4291743 | 0.237 | 0.458 | 1.63E-55 |
| Taok3 | 1.15E-59 | -0.4524055 | 0.278 | 0.507 | 1.70E-55 |
| Lsm11 | 1.30E-59 | -0.4449847 | 0.209 | 0.423 | 1.92E-55 |
| Tet2 | 1.38E-59 | 0.54992841 | 0.673 | 0.595 | 2.04E-55 |
| Timm9 | 1.63E-59 | -0.4433023 | 0.541 | 0.755 | 2.41E-55 |
| Rnf112 | 1.76E-59 | -0.370335 | 0.857 | 0.935 | 2.60E-55 |
| Hook2 | 2.03E-59 | -0.4170276 | 0.141 | 0.333 | 3.00E-55 |
| Vcan | 2.08E-59 | -0.5063614 | 0.129 | 0.314 | 3.06E-55 |
| Acan | 2.77E-59 | 0.63849089 | 0.314 | 0.152 | 4.09E-55 |
| Fam220a.1 | 7.17E-59 | -0.4655053 | 0.325 | 0.554 | 1.06E-54 |
| Cend1 | 1.28E-58 | -0.4358514 | 0.138 | 0.326 | 1.89E-54 |
| Hras | 2.01E-58 | -0.4533599 | 0.198 | 0.406 | 2.96E-54 |
| Cst3 | 2.93E-58 | -0.7759614 | 0.099 | 0.269 | 4.32E-54 |
| Pdlim7 | 1.28E-57 | -0.4135404 | 0.529 | 0.754 | 1.90E-53 |
| Gm38394 | 1.42E-57 | -0.426854 | 0.432 | 0.671 | 2.10E-53 |
| Qk | 2.19E-57 | 0.33151229 | 0.578 | 0.461 | 3.23E-53 |
| Dync1i1 | 2.41E-57 | -0.4176841 | 0.235 | 0.45 | 3.56E-53 |
| Abcd4 | 3.01E-57 | -0.4657495 | 0.258 | 0.478 | 4.45E-53 |
| Pxn | 3.40E-57 | -0.4078255 | 0.164 | 0.361 | 5.02E-53 |
| Fyco1 | 3.70E-57 | 0.61777472 | 0.581 | 0.469 | 5.46E-53 |
| Uvrag | 4.40E-57 | 0.58356234 | 0.695 | 0.621 | 6.49E-53 |
| Igf2bp3 | 4.40E-57 | -0.3936152 | 0.498 | 0.735 | 6.49E-53 |
| Eml5 | 1.36E-56 | 0.50868659 | 0.765 | 0.724 | 2.01E-52 |
| Mmp17 | 2.22E-56 | 0.62966915 | 0.523 | 0.395 | 3.27E-52 |
| Slc4a10 | 2.36E-56 | 0.61985356 | 0.438 | 0.29 | 3.49E-52 |
| Atp5j2 | 2.41E-56 | -0.4599773 | 0.248 | 0.466 | 3.56E-52 |
| Srgap2 | 2.48E-56 | -0.3776062 | 0.843 | 0.921 | 3.66E-52 |
| Adgrl3 | 3.79E-56 | -0.4760947 | 0.572 | 0.769 | 5.59E-52 |
| Pcbp2 | 7.89E-56 | 0.60529164 | 0.511 | 0.375 | 1.16E-51 |
| Oxr1 | 9.41E-56 | -0.4695186 | 0.162 | 0.35 | 1.39E-51 |

| | | | | | |
|---|---|---|---|---|---|
| Meis2 | 1.02E-55 | -0.4526448 | 0.195 | 0.409 | 1.51E-51 |
| Pianp | 1.58E-55 | 0.64680848 | 0.413 | 0.264 | 2.34E-51 |
| Ripor2 | 1.69E-55 | -0.3999969 | 0.114 | 0.286 | 2.49E-51 |
| Akap9 | 1.73E-55 | 0.54076227 | 0.667 | 0.585 | 2.55E-51 |
| Atp6v0c | 3.31E-55 | -0.3932334 | 0.648 | 0.858 | 4.88E-51 |
| Flrt3 | 3.74E-55 | -0.4444961 | 0.207 | 0.414 | 5.52E-51 |
| Prkx | 4.39E-55 | -0.3959883 | 0.136 | 0.317 | 6.47E-51 |
| Ubr3 | 5.07E-55 | 0.54449602 | 0.667 | 0.6 | 7.48E-51 |
| Ntrk2 | 5.82E-55 | 0.47050798 | 0.841 | 0.812 | 8.58E-51 |
| Rbm39 | 6.35E-55 | -0.2963136 | 0.958 | 0.981 | 9.38E-51 |
| Rcor3 | 1.19E-54 | -0.4150756 | 0.506 | 0.728 | 1.76E-50 |
| Gria3 | 1.21E-54 | 0.36650113 | 0.929 | 0.911 | 1.79E-50 |
| Tafa2 | 2.07E-54 | 0.59892485 | 0.27 | 0.125 | 3.05E-50 |
| Caln1 | 2.16E-54 | -0.3956274 | 0.131 | 0.308 | 3.19E-50 |
| Arhgef4 | 2.23E-54 | -0.4076855 | 0.318 | 0.547 | 3.29E-50 |
| Mical2 | 4.29E-54 | -0.3110742 | 0.885 | 0.943 | 6.33E-50 |
| Sncb | 7.67E-54 | -0.4372115 | 0.162 | 0.347 | 1.13E-49 |
| Fgf12 | 8.64E-54 | 0.61368624 | 0.343 | 0.192 | 1.28E-49 |
| Zc3h6 | 1.96E-53 | -0.4319711 | 0.317 | 0.539 | 2.89E-49 |
| Ppfia2 | 2.35E-53 | 0.40066454 | 0.878 | 0.847 | 3.46E-49 |
| Arhgef28 | 3.73E-53 | 0.55451186 | 0.665 | 0.577 | 5.51E-49 |
| Acap2 | 5.25E-53 | 0.41019466 | 0.879 | 0.867 | 7.75E-49 |
| Nav2 | 6.66E-53 | -0.3812984 | 0.711 | 0.861 | 9.83E-49 |
| Actr1b | 6.68E-53 | -0.4068091 | 0.491 | 0.706 | 9.86E-49 |
| Osbpl6 | 9.01E-53 | -0.4006041 | 0.254 | 0.472 | 1.33E-48 |
| Kctd4 | 9.99E-53 | -0.4716419 | 0.258 | 0.46 | 1.47E-48 |
| Tubb4b | 1.03E-52 | -0.3895663 | 0.14 | 0.318 | 1.52E-48 |
| Rgs7bp | 1.37E-52 | -0.4548123 | 0.269 | 0.481 | 2.02E-48 |
| Ndrg3 | 1.57E-52 | 0.39439239 | 0.884 | 0.862 | 2.32E-48 |
| Trpc4 | 1.64E-52 | -0.435303 | 0.311 | 0.529 | 2.42E-48 |
| Sel1l3 | 2.26E-52 | -0.4435202 | 0.201 | 0.399 | 3.34E-48 |
| Fkbp5 | 3.21E-52 | -0.4278667 | 0.289 | 0.499 | 4.73E-48 |
| Ppp1r12a | 6.19E-52 | 0.51832128 | 0.686 | 0.634 | 9.14E-48 |
| Rap1a | 1.26E-51 | -0.4147259 | 0.189 | 0.382 | 1.86E-47 |
| Tnrc6c | 2.05E-51 | -0.2704966 | 0.951 | 0.98 | 3.02E-47 |
| Stmn3 | 2.44E-51 | -0.3732484 | 0.367 | 0.612 | 3.60E-47 |
| Pak1 | 2.54E-51 | -0.4159568 | 0.308 | 0.53 | 3.75E-47 |
| Ahcyl2 | 2.67E-51 | 0.38669165 | 0.901 | 0.874 | 3.94E-47 |
| Rtn3 | 2.94E-51 | 0.51514473 | 0.669 | 0.616 | 4.34E-47 |
| Pdcd4 | 4.91E-51 | -0.4445077 | 0.172 | 0.357 | 7.24E-47 |
| Npdc1 | 5.49E-51 | -0.4032919 | 0.417 | 0.642 | 8.09E-47 |
| Rps13 | 6.98E-51 | -0.3904765 | 0.393 | 0.626 | 1.03E-46 |
| Negr1 | 7.10E-51 | 0.36922959 | 0.93 | 0.9 | 1.05E-46 |

| | | | | | |
|---|---|---|---|---|---|
| Phactr2 | 2.04E-50 | -0.3444675 | 0.109 | 0.273 | 3.01E-46 |
| Prdm8 | 2.62E-50 | 0.56994353 | 0.627 | 0.544 | 3.86E-46 |
| Wscd2 | 3.13E-50 | -0.3927248 | 0.475 | 0.693 | 4.62E-46 |
| Anapc5 | 4.08E-50 | -0.3822075 | 0.579 | 0.784 | 6.02E-46 |
| Ppwd1 | 4.94E-50 | -0.4015593 | 0.419 | 0.651 | 7.29E-46 |
| Nrg3 | 5.45E-50 | 0.55597714 | 0.582 | 0.497 | 8.04E-46 |
| Nbea | 6.04E-50 | 0.3608798 | 0.908 | 0.906 | 8.92E-46 |
| Arfgef3 | 6.27E-50 | 0.58065855 | 0.563 | 0.458 | 9.25E-46 |
| Nagk | 1.00E-49 | -0.3539968 | 0.153 | 0.335 | 1.48E-45 |
| Csmd2 | 1.57E-49 | -0.4119427 | 0.363 | 0.581 | 2.32E-45 |
| Lage3 | 1.85E-49 | -0.4062773 | 0.315 | 0.534 | 2.72E-45 |
| Trio | 2.83E-49 | -0.3761105 | 0.533 | 0.738 | 4.18E-45 |
| Mcur1 | 4.06E-49 | -0.3926825 | 0.201 | 0.391 | 5.99E-45 |
| Pabpc1 | 5.35E-49 | -0.411309 | 0.351 | 0.562 | 7.90E-45 |
| Hacd2 | 8.62E-49 | 0.63205306 | 0.354 | 0.213 | 1.27E-44 |
| Agrn | 9.05E-49 | -0.428387 | 0.269 | 0.471 | 1.33E-44 |
| Psmd9 | 9.75E-49 | -0.3773955 | 0.271 | 0.489 | 1.44E-44 |
| Polr3d | 1.07E-48 | -0.3688022 | 0.127 | 0.292 | 1.57E-44 |
| Tmsb4x | 1.25E-48 | 0.43513076 | 0.849 | 0.852 | 1.85E-44 |
| Pet100 | 2.18E-48 | -0.3891407 | 0.287 | 0.5 | 3.21E-44 |
| Mpg | 2.35E-48 | -0.3538901 | 0.097 | 0.25 | 3.47E-44 |
| Vps13c | 2.65E-48 | -0.3667102 | 0.731 | 0.867 | 3.91E-44 |
| Zbtb16 | 2.87E-48 | 0.60271355 | 0.536 | 0.431 | 4.23E-44 |
| Chd3 | 6.33E-48 | 0.41355549 | 0.822 | 0.806 | 9.34E-44 |
| Hnrnpa2b1 | 6.88E-48 | 0.34462915 | 0.866 | 0.839 | 1.02E-43 |
| Kif21a | 7.50E-48 | -0.3658227 | 0.686 | 0.862 | 1.11E-43 |
| Las1l | 9.04E-48 | -0.370565 | 0.414 | 0.645 | 1.33E-43 |
| Jph1 | 1.02E-47 | 0.54022449 | 0.611 | 0.522 | 1.50E-43 |
| Dstn | 1.17E-47 | -0.3901782 | 0.354 | 0.57 | 1.72E-43 |
| Gm10033 | 1.37E-47 | -0.369582 | 0.151 | 0.325 | 2.03E-43 |
| Manba | 1.53E-47 | -0.3657999 | 0.108 | 0.261 | 2.26E-43 |
| Shisal1 | 1.71E-47 | 0.59363395 | 0.284 | 0.146 | 2.52E-43 |
| Cox4i1 | 1.79E-47 | -0.420877 | 0.265 | 0.474 | 2.64E-43 |
| Ano6 | 1.81E-47 | -0.3599201 | 0.131 | 0.297 | 2.66E-43 |
| Ak2 | 1.90E-47 | -0.3812667 | 0.251 | 0.455 | 2.80E-43 |
| Psma1 | 2.49E-47 | -0.3711902 | 0.551 | 0.76 | 3.67E-43 |
| Rock1 | 2.78E-47 | 0.48555001 | 0.715 | 0.665 | 4.10E-43 |
| Mfsd4a | 3.01E-47 | 0.5323746 | 0.604 | 0.52 | 4.44E-43 |
| Usp20 | 4.98E-47 | -0.3780092 | 0.22 | 0.413 | 7.35E-43 |
| Fam214b | 6.34E-47 | -0.3669478 | 0.253 | 0.454 | 9.35E-43 |
| Pgp | 7.11E-47 | -0.3778606 | 0.2 | 0.388 | 1.05E-42 |
| R3hdm2 | 1.10E-46 | 0.43888896 | 0.778 | 0.735 | 1.62E-42 |
| Ncor2 | 1.29E-46 | 0.501664 | 0.652 | 0.587 | 1.91E-42 |

| | | | | | |
|---|---|---|---|---|---|
| Cdc14b | 1.35E-46 | 0.60884267 | 0.294 | 0.159 | 1.99E-42 |
| L1cam | 1.43E-46 | -0.3686181 | 0.201 | 0.391 | 2.11E-42 |
| Slc12a5 | 2.34E-46 | 0.55426794 | 0.561 | 0.465 | 3.46E-42 |
| Serinc1 | 2.60E-46 | 0.55411749 | 0.539 | 0.437 | 3.83E-42 |
| Zer1 | 3.15E-46 | -0.3366894 | 0.713 | 0.861 | 4.64E-42 |
| Arhgef9 | 4.28E-46 | 0.48480759 | 0.681 | 0.627 | 6.31E-42 |
| Zfp207 | 6.66E-46 | 0.45213177 | 0.714 | 0.675 | 9.83E-42 |
| Prrt1 | 8.63E-46 | -0.3626704 | 0.511 | 0.722 | 1.27E-41 |
| Xpr1 | 1.47E-45 | 0.6000341 | 0.456 | 0.331 | 2.17E-41 |
| Taok1 | 1.60E-45 | 0.53167144 | 0.58 | 0.492 | 2.37E-41 |
| Nlgn1 | 1.68E-45 | 0.4877388 | 0.693 | 0.648 | 2.47E-41 |
| Slc38a2 | 2.86E-45 | 0.45954869 | 0.754 | 0.7 | 4.22E-41 |
| Golim4 | 3.33E-45 | -0.3627298 | 0.124 | 0.279 | 4.92E-41 |
| Olfm1 | 3.46E-45 | -0.4375908 | 0.246 | 0.441 | 5.11E-41 |
| Dsel | 3.77E-45 | -0.3441817 | 0.132 | 0.292 | 5.56E-41 |
| Elavl3 | 4.82E-45 | -0.3687723 | 0.512 | 0.722 | 7.11E-41 |
| Safb | 5.05E-45 | -0.3981751 | 0.513 | 0.711 | 7.46E-41 |
| Nsg2 | 5.59E-45 | -0.3976419 | 0.265 | 0.462 | 8.25E-41 |
| Tuba1a | 6.33E-45 | -0.4161724 | 0.203 | 0.385 | 9.35E-41 |
| Rchy1 | 6.75E-45 | -0.3850474 | 0.512 | 0.727 | 9.96E-41 |
| P2ry14 | 7.02E-45 | -0.3788878 | 0.273 | 0.475 | 1.04E-40 |
| Begain | 8.12E-45 | -0.3236203 | 0.206 | 0.395 | 1.20E-40 |
| Hecw2 | 1.41E-44 | 0.58461211 | 0.381 | 0.247 | 2.07E-40 |
| Sirt3 | 1.58E-44 | -0.3851366 | 0.29 | 0.496 | 2.33E-40 |
| Diaph2 | 2.85E-44 | 0.56848517 | 0.352 | 0.217 | 4.20E-40 |
| Mpped2 | 3.02E-44 | 0.41508376 | 0.821 | 0.786 | 4.45E-40 |
| Fam229b | 3.05E-44 | -0.3376109 | 0.113 | 0.262 | 4.50E-40 |
| Kndc1 | 3.20E-44 | -0.3910688 | 0.185 | 0.363 | 4.72E-40 |
| Ablim2 | 3.24E-44 | -0.3941733 | 0.217 | 0.403 | 4.78E-40 |
| Ak5 | 6.89E-44 | 0.49602656 | 0.631 | 0.559 | 1.02E-39 |
| Scai | 8.88E-44 | -0.3736257 | 0.52 | 0.729 | 1.31E-39 |
| Nmt1 | 1.08E-43 | -0.3869228 | 0.399 | 0.611 | 1.59E-39 |
| Atxn1 | 1.10E-43 | 0.48178162 | 0.669 | 0.613 | 1.62E-39 |
| Clk4 | 1.18E-43 | 0.43549426 | 0.761 | 0.738 | 1.75E-39 |
| Hook1 | 1.30E-43 | 0.47538948 | 0.677 | 0.635 | 1.92E-39 |
| Cdc42se2 | 1.56E-43 | -0.3878032 | 0.323 | 0.528 | 2.30E-39 |
| Gpr158 | 1.80E-43 | 0.56359531 | 0.478 | 0.368 | 2.65E-39 |
| Tle4 | 2.04E-43 | -0.3942168 | 0.28 | 0.477 | 3.01E-39 |
| Tmem214 | 2.16E-43 | -0.3617812 | 0.295 | 0.495 | 3.19E-39 |
| Ralyl | 2.65E-43 | -0.3346506 | 0.642 | 0.819 | 3.91E-39 |
| Clip1 | 2.90E-43 | 0.39698159 | 0.818 | 0.804 | 4.27E-39 |
| Saa3 | 3.04E-43 | -0.3202807 | 0.126 | 0.281 | 4.49E-39 |
| Ncdn | 4.32E-43 | -0.3847684 | 0.382 | 0.585 | 6.37E-39 |

| | | | | | |
|---|---|---|---|---|---|
| Sf3b1 | 7.53E-43 | 0.29267747 | 0.964 | 0.959 | 1.11E-38 |
| Mdh1 | 1.20E-42 | -0.3632163 | 0.252 | 0.448 | 1.77E-38 |
| Tasor | 1.41E-42 | 0.56413971 | 0.514 | 0.416 | 2.08E-38 |
| Iqsec3 | 1.88E-42 | -0.339006 | 0.226 | 0.414 | 2.78E-38 |
| Ankrd12 | 2.62E-42 | 0.41422681 | 0.819 | 0.816 | 3.87E-38 |
| Agk | 2.63E-42 | -0.3510016 | 0.274 | 0.472 | 3.88E-38 |
| Ptk2b | 3.21E-42 | 0.44765625 | 0.703 | 0.655 | 4.73E-38 |
| Fam126b | 3.30E-42 | 0.55978741 | 0.596 | 0.519 | 4.87E-38 |
| Mprip | 4.09E-42 | 0.44862209 | 0.711 | 0.67 | 6.03E-38 |
| Nckap1 | 4.65E-42 | 0.48364292 | 0.604 | 0.53 | 6.86E-38 |
| Map1lc3a | 4.90E-42 | -0.3646389 | 0.176 | 0.348 | 7.23E-38 |
| Mef2a | 6.86E-42 | -0.3480683 | 0.621 | 0.798 | 1.01E-37 |
| Tiam1 | 7.03E-42 | 0.54361455 | 0.306 | 0.177 | 1.04E-37 |
| Ext1 | 1.03E-41 | 0.55132688 | 0.393 | 0.268 | 1.52E-37 |
| Kat6b | 1.48E-41 | 0.52852541 | 0.569 | 0.487 | 2.18E-37 |
| Mt3 | 1.89E-41 | -0.3760429 | 0.523 | 0.731 | 2.78E-37 |
| Zfp955b | 2.14E-41 | -0.3395582 | 0.211 | 0.391 | 3.16E-37 |
| Yeats2 | 5.39E-41 | -0.3524125 | 0.503 | 0.706 | 7.95E-37 |
| Nudcd3 | 2.40E-40 | -0.3400397 | 0.6 | 0.784 | 3.54E-36 |
| D3Ertd751e | 3.01E-40 | -0.3092393 | 0.126 | 0.274 | 4.44E-36 |
| Gk | 3.05E-40 | 0.55938715 | 0.35 | 0.225 | 4.50E-36 |
| Usp46 | 3.07E-40 | -0.3563826 | 0.347 | 0.548 | 4.53E-36 |
| Grm7 | 3.27E-40 | 0.54455203 | 0.528 | 0.435 | 4.83E-36 |
| Cdkl5 | 3.31E-40 | 0.55686635 | 0.502 | 0.405 | 4.88E-36 |
| Polr1a | 5.54E-40 | -0.2841637 | 0.144 | 0.302 | 8.17E-36 |
| Atp5d | 5.63E-40 | -0.3318347 | 0.208 | 0.385 | 8.31E-36 |
| Lyrm9 | 7.12E-40 | -0.327485 | 0.127 | 0.274 | 1.05E-35 |
| Maml3 | 7.44E-40 | 0.55185969 | 0.429 | 0.306 | 1.10E-35 |
| Rbm4b | 1.21E-39 | -0.321509 | 0.599 | 0.797 | 1.79E-35 |
| Puf60 | 1.48E-39 | -0.3406313 | 0.318 | 0.521 | 2.18E-35 |
| Psmc3 | 1.63E-39 | -0.3563741 | 0.313 | 0.508 | 2.40E-35 |
| Atp2b2 | 2.59E-39 | 0.41555206 | 0.722 | 0.692 | 3.82E-35 |
| Gsdme | 2.72E-39 | -0.3853968 | 0.284 | 0.476 | 4.01E-35 |
| Cpsf6 | 2.72E-39 | 0.42739684 | 0.719 | 0.681 | 4.01E-35 |
| Lypla2 | 4.11E-39 | -0.3510932 | 0.271 | 0.457 | 6.07E-35 |
| Sag | 4.83E-39 | -0.3105483 | 0.112 | 0.252 | 7.13E-35 |
| Pfkfb4 | 5.23E-39 | -0.3559614 | 0.19 | 0.356 | 7.71E-35 |
| Sh3bp1 | 5.39E-39 | -0.3225469 | 0.114 | 0.254 | 7.96E-35 |
| Atp1b1 | 6.74E-39 | 0.34294213 | 0.899 | 0.929 | 9.94E-35 |
| mt-Co2 | 7.32E-39 | 0.46874091 | 0.566 | 0.465 | 1.08E-34 |
| Ubl5 | 8.28E-39 | -0.329749 | 0.27 | 0.458 | 1.22E-34 |
| Chchd10 | 1.46E-38 | -0.3525022 | 0.134 | 0.278 | 2.16E-34 |
| Dbndd2 | 1.69E-38 | -0.3351625 | 0.217 | 0.394 | 2.49E-34 |

| | | | | | |
|---|---|---|---|---|---|
| Top2b | 1.76E-38 | 0.52433626 | 0.436 | 0.324 | 2.60E-34 |
| Zmynd8 | 2.56E-38 | 0.36738622 | 0.881 | 0.862 | 3.78E-34 |
| Tubb2a | 3.02E-38 | -0.3234909 | 0.359 | 0.563 | 4.45E-34 |
| Dalrd3 | 3.28E-38 | -0.2904111 | 0.637 | 0.814 | 4.85E-34 |
| Sptbn1 | 3.49E-38 | -0.2510949 | 0.899 | 0.965 | 5.15E-34 |
| Kcnh1 | 3.97E-38 | -0.3364723 | 0.21 | 0.382 | 5.86E-34 |
| Zfp365 | 4.57E-38 | 0.55406672 | 0.323 | 0.203 | 6.74E-34 |
| Atp6v1g2 | 4.65E-38 | -0.3326067 | 0.26 | 0.437 | 6.86E-34 |
| Chn1 | 8.30E-38 | 0.33026645 | 0.896 | 0.892 | 1.22E-33 |
| Hnrnpm | 1.39E-37 | -0.3646715 | 0.508 | 0.688 | 2.05E-33 |
| Zfr2 | 1.53E-37 | -0.3306797 | 0.364 | 0.562 | 2.25E-33 |
| Man2b1 | 1.97E-37 | -0.3299084 | 0.272 | 0.459 | 2.90E-33 |
| Faim | 2.01E-37 | -0.3269907 | 0.143 | 0.29 | 2.97E-33 |
| Mapk8ip1 | 2.88E-37 | -0.327866 | 0.478 | 0.677 | 4.25E-33 |
| Pea15a | 3.14E-37 | -0.3021595 | 0.12 | 0.259 | 4.63E-33 |
| Reps2 | 3.42E-37 | -0.3306149 | 0.463 | 0.675 | 5.04E-33 |
| Hint1 | 3.69E-37 | -0.321815 | 0.152 | 0.303 | 5.44E-33 |
| Btbd3 | 4.17E-37 | 0.48105059 | 0.613 | 0.525 | 6.15E-33 |
| Huwe1 | 4.53E-37 | 0.4194633 | 0.672 | 0.621 | 6.69E-33 |
| Arhgef25 | 4.76E-37 | 0.31443175 | 0.936 | 0.921 | 7.03E-33 |
| Tecr | 5.34E-37 | -0.3603631 | 0.347 | 0.542 | 7.89E-33 |
| Tspoap1 | 6.19E-37 | 0.35549869 | 0.853 | 0.836 | 9.14E-33 |
| Tdrp | 8.16E-37 | -0.3024369 | 0.131 | 0.275 | 1.20E-32 |
| Grin1 | 8.17E-37 | -0.2675105 | 0.794 | 0.907 | 1.21E-32 |
| Syt14 | 8.69E-37 | 0.53074949 | 0.428 | 0.319 | 1.28E-32 |
| Uqcrc1 | 9.27E-37 | -0.2879486 | 0.161 | 0.317 | 1.37E-32 |
| Mdga1 | 1.20E-36 | -0.334279 | 0.205 | 0.371 | 1.76E-32 |
| Phactr1 | 1.26E-36 | -0.3198104 | 0.703 | 0.856 | 1.85E-32 |
| Mrtfb | 1.27E-36 | 0.48692249 | 0.544 | 0.463 | 1.87E-32 |
| Rps6kb2 | 1.37E-36 | -0.3218736 | 0.54 | 0.726 | 2.02E-32 |
| Gng3 | 1.43E-36 | -0.3264786 | 0.155 | 0.305 | 2.10E-32 |
| Map4k3 | 2.17E-36 | 0.44197403 | 0.651 | 0.605 | 3.20E-32 |
| Flnb | 2.38E-36 | -0.3000103 | 0.244 | 0.421 | 3.51E-32 |
| Kcnd2 | 2.70E-36 | 0.52593488 | 0.427 | 0.318 | 3.99E-32 |
| Nsg1 | 4.05E-36 | -0.2998056 | 0.119 | 0.256 | 5.98E-32 |
| Prkce | 6.37E-36 | 0.44582861 | 0.674 | 0.644 | 9.40E-32 |
| Hpgds | 6.85E-36 | -0.3329822 | 0.242 | 0.413 | 1.01E-31 |
| Mif | 8.14E-36 | -0.2964417 | 0.148 | 0.292 | 1.20E-31 |
| Unc5a | 1.04E-35 | -0.2745079 | 0.13 | 0.273 | 1.54E-31 |
| Tnfrsf25 | 1.18E-35 | -0.290236 | 0.217 | 0.391 | 1.74E-31 |
| Apbb2 | 1.71E-35 | 0.3932654 | 0.735 | 0.699 | 2.52E-31 |
| Ppp1r9a | 2.04E-35 | 0.48908492 | 0.555 | 0.476 | 3.01E-31 |
| Abraxas2 | 2.19E-35 | -0.3008282 | 0.189 | 0.349 | 3.23E-31 |

| | | | | | |
|---|---|---|---|---|---|
| Kcnq5 | 2.32E-35 | -0.3231502 | 0.227 | 0.395 | 3.43E-31 |
| Eps15 | 2.37E-35 | -0.3117778 | 0.446 | 0.648 | 3.50E-31 |
| Arhgef11 | 2.58E-35 | -0.3064151 | 0.193 | 0.354 | 3.80E-31 |
| Tfcp2 | 2.74E-35 | -0.2935068 | 0.163 | 0.314 | 4.04E-31 |
| Sppl2b | 2.98E-35 | -0.2993155 | 0.344 | 0.544 | 4.40E-31 |
| Mink1 | 2.99E-35 | -0.3435506 | 0.417 | 0.609 | 4.41E-31 |
| Rbm28 | 3.32E-35 | -0.3214167 | 0.543 | 0.724 | 4.90E-31 |
| Mttp | 3.73E-35 | -0.3073045 | 0.159 | 0.309 | 5.50E-31 |
| Rab11fip2 | 5.08E-35 | -0.3198559 | 0.614 | 0.791 | 7.50E-31 |
| Vti1b | 5.73E-35 | -0.3152914 | 0.182 | 0.336 | 8.46E-31 |
| Sec61a2 | 7.07E-35 | -0.3256187 | 0.55 | 0.732 | 1.04E-30 |
| Lars2 | 8.50E-35 | 0.49323812 | 0.361 | 0.245 | 1.25E-30 |
| Msh3 | 1.24E-34 | -0.3218156 | 0.286 | 0.471 | 1.83E-30 |
| Mrpl38 | 1.88E-34 | -0.3316568 | 0.285 | 0.465 | 2.77E-30 |
| Peg3 | 1.92E-34 | 0.39586326 | 0.788 | 0.772 | 2.84E-30 |
| Adgrb1 | 2.01E-34 | -0.3342698 | 0.571 | 0.744 | 2.97E-30 |
| Uqcr11 | 2.05E-34 | -0.3178566 | 0.153 | 0.296 | 3.03E-30 |
| Cep70 | 2.63E-34 | -0.3028353 | 0.347 | 0.536 | 3.88E-30 |
| Pkd1 | 2.64E-34 | -0.3138724 | 0.619 | 0.789 | 3.89E-30 |
| Runx1t1 | 3.19E-34 | -0.3052428 | 0.243 | 0.412 | 4.71E-30 |
| Dip2b | 3.32E-34 | -0.2920841 | 0.4 | 0.606 | 4.90E-30 |
| Pias1 | 3.49E-34 | -0.31208 | 0.23 | 0.397 | 5.15E-30 |
| Rps12 | 3.67E-34 | -0.3003892 | 0.308 | 0.502 | 5.41E-30 |
| Pmm2 | 3.95E-34 | -0.288577 | 0.197 | 0.357 | 5.82E-30 |
| Mllt11 | 4.28E-34 | -0.3276128 | 0.303 | 0.485 | 6.31E-30 |
| Ncoa7 | 5.58E-34 | -0.3056398 | 0.578 | 0.762 | 8.23E-30 |
| Atp5b | 6.53E-34 | -0.3096973 | 0.395 | 0.594 | 9.64E-30 |
| Telo2 | 7.01E-34 | -0.2956924 | 0.225 | 0.392 | 1.03E-29 |
| Gpatch11 | 7.41E-34 | -0.2674903 | 0.156 | 0.305 | 1.09E-29 |
| Akt3 | 7.97E-34 | 0.50908114 | 0.391 | 0.284 | 1.18E-29 |
| Hdac7 | 8.38E-34 | 0.38547395 | 0.703 | 0.625 | 1.24E-29 |
| Tnfrsf21 | 8.97E-34 | -0.3033315 | 0.151 | 0.29 | 1.32E-29 |
| Jak1 | 9.81E-34 | -0.3346538 | 0.35 | 0.525 | 1.45E-29 |
| Ephb2 | 1.16E-33 | -0.2904441 | 0.14 | 0.278 | 1.71E-29 |
| Ptdss2 | 2.01E-33 | -0.2796073 | 0.279 | 0.46 | 2.96E-29 |
| Hivep2 | 2.02E-33 | 0.26764232 | 0.953 | 0.942 | 2.98E-29 |
| Mical3 | 2.15E-33 | -0.3195385 | 0.412 | 0.597 | 3.17E-29 |
| Braf | 2.39E-33 | 0.43507986 | 0.682 | 0.642 | 3.53E-29 |
| Dipk1b | 2.55E-33 | -0.3048881 | 0.345 | 0.532 | 3.77E-29 |
| Tln2 | 2.65E-33 | -0.3097268 | 0.28 | 0.457 | 3.91E-29 |
| Cep192 | 2.69E-33 | -0.2874576 | 0.147 | 0.288 | 3.97E-29 |
| Slc25a37 | 2.87E-33 | 0.54599841 | 0.367 | 0.258 | 4.24E-29 |
| Usp3 | 3.16E-33 | 0.5114734 | 0.51 | 0.434 | 4.66E-29 |

| | | | | | |
|---|---|---|---|---|---|
| Ndufa11 | 3.68E-33 | -0.3065528 | 0.122 | 0.254 | 5.43E-29 |
| Glrx3 | 3.82E-33 | -0.2857684 | 0.204 | 0.366 | 5.64E-29 |
| Klhl29 | 4.75E-33 | 0.51834717 | 0.492 | 0.408 | 7.01E-29 |
| Rpl38 | 6.42E-33 | -0.3311608 | 0.168 | 0.316 | 9.48E-29 |
| Ndufa4 | 6.74E-33 | -0.2942425 | 0.203 | 0.363 | 9.95E-29 |
| Slc25a27 | 9.16E-33 | -0.3037811 | 0.464 | 0.661 | 1.35E-28 |
| Gprasp2 | 9.60E-33 | -0.2765186 | 0.126 | 0.257 | 1.42E-28 |
| Rapgef6 | 9.78E-33 | -0.2918848 | 0.581 | 0.764 | 1.44E-28 |
| Smpd3 | 1.24E-32 | -0.3225388 | 0.538 | 0.712 | 1.83E-28 |
| Pdcd11 | 1.26E-32 | -0.2731362 | 0.179 | 0.332 | 1.86E-28 |
| Srrm4 | 1.31E-32 | -0.2706023 | 0.586 | 0.768 | 1.94E-28 |
| Ppp1r12c | 1.59E-32 | 0.47939484 | 0.593 | 0.541 | 2.35E-28 |
| Ndfip1 | 1.61E-32 | -0.3295324 | 0.472 | 0.659 | 2.37E-28 |
| Eif3m | 1.68E-32 | -0.2808258 | 0.154 | 0.295 | 2.49E-28 |
| Trmt1l | 1.96E-32 | -0.3117539 | 0.38 | 0.568 | 2.89E-28 |
| Ptpru | 2.17E-32 | 0.53632075 | 0.41 | 0.311 | 3.20E-28 |
| Iffo1 | 2.34E-32 | -0.2905938 | 0.382 | 0.57 | 3.46E-28 |
| Atp5md | 2.84E-32 | -0.323508 | 0.171 | 0.315 | 4.18E-28 |
| Dnajc21 | 2.92E-32 | -0.297453 | 0.235 | 0.399 | 4.30E-28 |
| Ankrd27 | 3.86E-32 | -0.2837243 | 0.145 | 0.281 | 5.70E-28 |
| Podxl2 | 3.87E-32 | -0.3347864 | 0.37 | 0.545 | 5.71E-28 |
| Nptxr | 4.18E-32 | 0.56173582 | 0.374 | 0.271 | 6.17E-28 |
| Tacc2 | 4.50E-32 | 0.42730081 | 0.643 | 0.589 | 6.64E-28 |
| Nisch | 4.58E-32 | -0.2716245 | 0.707 | 0.844 | 6.76E-28 |
| Lman2l | 5.90E-32 | -0.2942298 | 0.505 | 0.695 | 8.71E-28 |
| 1110008P14l | 6.80E-32 | -0.2967355 | 0.138 | 0.271 | 1.00E-27 |
| Atp6v1e1 | 7.09E-32 | -0.3027581 | 0.196 | 0.346 | 1.05E-27 |
| Slc50a1 | 7.19E-32 | -0.2825888 | 0.279 | 0.452 | 1.06E-27 |
| Rasal1 | 7.78E-32 | 0.52321534 | 0.394 | 0.293 | 1.15E-27 |
| Nptn | 8.38E-32 | 0.40766142 | 0.601 | 0.544 | 1.24E-27 |
| Cabin1 | 8.45E-32 | -0.2903418 | 0.595 | 0.768 | 1.25E-27 |
| Ttc7b | 9.05E-32 | 0.51353843 | 0.392 | 0.291 | 1.33E-27 |
| Prdx5 | 9.12E-32 | -0.2920943 | 0.173 | 0.318 | 1.35E-27 |
| Exoc6b | 1.38E-31 | 0.37184821 | 0.73 | 0.71 | 2.03E-27 |
| Smpd4 | 1.50E-31 | -0.3099466 | 0.359 | 0.534 | 2.21E-27 |
| Sidt1 | 1.74E-31 | 0.49498889 | 0.552 | 0.479 | 2.56E-27 |
| Wdr18 | 1.94E-31 | -0.2721934 | 0.194 | 0.346 | 2.87E-27 |
| Polg | 2.06E-31 | 0.45371502 | 0.559 | 0.49 | 3.04E-27 |
| Fam126a | 2.07E-31 | -0.30126 | 0.183 | 0.327 | 3.05E-27 |
| Rrbp1 | 2.22E-31 | -0.3082932 | 0.151 | 0.291 | 3.27E-27 |
| Phka2 | 2.29E-31 | -0.3023355 | 0.44 | 0.63 | 3.37E-27 |
| Grk3 | 2.47E-31 | -0.3363085 | 0.541 | 0.713 | 3.65E-27 |
| Sptan1 | 2.55E-31 | -0.2697209 | 0.673 | 0.832 | 3.76E-27 |

| | | | | | |
|---|---|---|---|---|---|
| Ccdc28a | 3.25E-31 | -0.2569048 | 0.138 | 0.273 | 4.80E-27 |
| Pygo1 | 3.34E-31 | -0.3077683 | 0.473 | 0.655 | 4.93E-27 |
| D10Wsu102e | 4.22E-31 | 0.51175554 | 0.574 | 0.522 | 6.23E-27 |
| Degs1 | 4.52E-31 | -0.2814281 | 0.16 | 0.3 | 6.67E-27 |
| Thap3 | 4.57E-31 | -0.286649 | 0.274 | 0.447 | 6.74E-27 |
| Nsun6 | 4.99E-31 | -0.2756321 | 0.231 | 0.395 | 7.36E-27 |
| Srpk1 | 5.15E-31 | -0.2761048 | 0.559 | 0.738 | 7.61E-27 |
| Izumo4 | 8.17E-31 | -0.2754108 | 0.283 | 0.456 | 1.21E-26 |
| Usp2 | 9.76E-31 | 0.51083295 | 0.334 | 0.225 | 1.44E-26 |
| Pgam1 | 1.19E-30 | -0.2777934 | 0.19 | 0.339 | 1.75E-26 |
| Ddn | 1.23E-30 | 0.49981345 | 0.426 | 0.333 | 1.81E-26 |
| Rnf170 | 1.32E-30 | -0.2778967 | 0.203 | 0.356 | 1.95E-26 |
| Lekr1 | 1.77E-30 | -0.2798502 | 0.216 | 0.369 | 2.62E-26 |
| Hspa12a | 1.82E-30 | -0.2763039 | 0.124 | 0.25 | 2.69E-26 |
| Ccdc85b | 1.92E-30 | 0.47331943 | 0.481 | 0.394 | 2.83E-26 |
| Uqcrq | 2.34E-30 | -0.2810637 | 0.161 | 0.299 | 3.46E-26 |
| Dkk3 | 2.38E-30 | -0.2853511 | 0.173 | 0.314 | 3.51E-26 |
| Tnrc6a | 2.86E-30 | -0.303823 | 0.52 | 0.699 | 4.22E-26 |
| Eif2s2 | 3.31E-30 | -0.269433 | 0.397 | 0.579 | 4.89E-26 |
| Kif13b | 3.61E-30 | -0.2997279 | 0.218 | 0.368 | 5.32E-26 |
| Suclg1 | 3.84E-30 | -0.2566149 | 0.167 | 0.307 | 5.67E-26 |
| Coro2b | 4.35E-30 | -0.2630099 | 0.154 | 0.288 | 6.42E-26 |
| Tmed4 | 4.35E-30 | -0.2895025 | 0.215 | 0.367 | 6.43E-26 |
| Got2 | 4.39E-30 | -0.2868137 | 0.25 | 0.415 | 6.48E-26 |
| Abr | 4.62E-30 | 0.35130236 | 0.758 | 0.751 | 6.82E-26 |
| Dmac2 | 4.99E-30 | -0.273641 | 0.197 | 0.346 | 7.36E-26 |
| Arel1 | 5.36E-30 | -0.2755567 | 0.189 | 0.333 | 7.90E-26 |
| Rapgef5 | 5.89E-30 | 0.32791111 | 0.84 | 0.794 | 8.69E-26 |
| Wasl | 6.16E-30 | 0.41263572 | 0.649 | 0.609 | 9.09E-26 |
| Stag1 | 6.46E-30 | 0.4852359 | 0.489 | 0.413 | 9.53E-26 |
| Plxna4 | 7.23E-30 | 0.46149976 | 0.565 | 0.505 | 1.07E-25 |
| Arhgef3 | 7.33E-30 | 0.46576574 | 0.262 | 0.159 | 1.08E-25 |
| Tmem234 | 8.98E-30 | -0.265258 | 0.54 | 0.735 | 1.33E-25 |
| Atrx | 8.99E-30 | 0.29590791 | 0.868 | 0.878 | 1.33E-25 |
| 4932438A13I | 9.98E-30 | 0.41063622 | 0.593 | 0.541 | 1.47E-25 |
| Zbtb18 | 1.03E-29 | 0.43238122 | 0.583 | 0.54 | 1.52E-25 |
| Unc5c | 1.05E-29 | -0.2911653 | 0.542 | 0.713 | 1.55E-25 |
| Emc9 | 1.40E-29 | -0.2655716 | 0.184 | 0.329 | 2.06E-25 |
| Adam15 | 1.71E-29 | -0.2627324 | 0.297 | 0.474 | 2.52E-25 |
| Sumo2 | 1.84E-29 | -0.2750161 | 0.425 | 0.615 | 2.71E-25 |
| Macrod2 | 2.02E-29 | -0.274249 | 0.26 | 0.421 | 2.99E-25 |
| Wdr43 | 2.07E-29 | -0.2622181 | 0.326 | 0.503 | 3.06E-25 |
| Aimp1 | 2.16E-29 | -0.2960519 | 0.364 | 0.545 | 3.19E-25 |

| | | | | | |
|---|---|---|---|---|---|
| Rab34 | 2.65E-29 | -0.2940793 | 0.535 | 0.719 | 3.91E-25 |
| Tsnax | 2.83E-29 | -0.2737622 | 0.266 | 0.428 | 4.18E-25 |
| Elfn2 | 3.03E-29 | -0.2605816 | 0.171 | 0.308 | 4.47E-25 |
| Tia1 | 3.17E-29 | 0.25241423 | 0.925 | 0.937 | 4.68E-25 |
| Clock | 3.29E-29 | -0.255832 | 0.622 | 0.789 | 4.85E-25 |
| Ssh2 | 4.60E-29 | 0.4304235 | 0.617 | 0.573 | 6.78E-25 |
| Aifm3 | 4.91E-29 | 0.50154196 | 0.36 | 0.261 | 7.25E-25 |
| Tpi1 | 5.82E-29 | -0.2936314 | 0.311 | 0.483 | 8.58E-25 |
| Gpr162 | 5.94E-29 | 0.46508025 | 0.521 | 0.454 | 8.76E-25 |
| Bcl7a | 6.82E-29 | 0.50228514 | 0.429 | 0.339 | 1.01E-24 |
| Stk38 | 7.34E-29 | -0.2849681 | 0.541 | 0.725 | 1.08E-24 |
| Rpp21 | 7.41E-29 | -0.2732027 | 0.232 | 0.387 | 1.09E-24 |
| BC005537 | 8.14E-29 | 0.42551915 | 0.54 | 0.479 | 1.20E-24 |
| Arap2 | 8.56E-29 | -0.2684208 | 0.699 | 0.835 | 1.26E-24 |
| Rint1 | 9.44E-29 | -0.2784696 | 0.146 | 0.273 | 1.39E-24 |
| Ckmt1 | 1.00E-28 | -0.2743015 | 0.189 | 0.331 | 1.48E-24 |
| Esrra | 1.11E-28 | -0.2775736 | 0.288 | 0.457 | 1.64E-24 |
| Larp1b | 1.48E-28 | -0.2571689 | 0.244 | 0.4 | 2.18E-24 |
| Ralgps2 | 1.70E-28 | -0.2620533 | 0.514 | 0.697 | 2.50E-24 |
| Gkap1 | 1.97E-28 | -0.287252 | 0.258 | 0.417 | 2.91E-24 |
| Josd2 | 2.03E-28 | -0.2775402 | 0.158 | 0.29 | 2.99E-24 |
| Sbf2 | 2.11E-28 | -0.2835099 | 0.266 | 0.426 | 3.11E-24 |
| Hint3 | 2.51E-28 | -0.2562988 | 0.131 | 0.253 | 3.70E-24 |
| Chgb | 2.52E-28 | 0.39768048 | 0.66 | 0.634 | 3.71E-24 |
| Mia2 | 2.53E-28 | 0.4156393 | 0.65 | 0.61 | 3.73E-24 |
| Far1 | 2.92E-28 | 0.41302829 | 0.621 | 0.588 | 4.31E-24 |
| Cs | 2.94E-28 | -0.2514692 | 0.198 | 0.344 | 4.34E-24 |
| Rpl36 | 3.09E-28 | -0.2903283 | 0.148 | 0.275 | 4.56E-24 |
| Pde10a | 4.81E-28 | 0.28521331 | 0.656 | 0.575 | 7.10E-24 |
| Eif4g2 | 6.10E-28 | 0.44132282 | 0.475 | 0.397 | 9.01E-24 |
| Rnf187 | 6.81E-28 | -0.2896314 | 0.188 | 0.326 | 1.00E-23 |
| Fam184a | 7.80E-28 | 0.46054303 | 0.29 | 0.188 | 1.15E-23 |
| Cox6c | 9.56E-28 | -0.2761465 | 0.291 | 0.452 | 1.41E-23 |
| Hivep3 | 1.01E-27 | -0.2699063 | 0.545 | 0.726 | 1.50E-23 |
| Rpl21 | 1.02E-27 | -0.281664 | 0.301 | 0.467 | 1.50E-23 |
| Napg | 1.05E-27 | -0.2528803 | 0.645 | 0.817 | 1.55E-23 |
| Rpl28 | 1.09E-27 | -0.2645404 | 0.24 | 0.393 | 1.61E-23 |
| Clasrp | 1.10E-27 | -0.261207 | 0.297 | 0.464 | 1.63E-23 |
| Dst | 1.11E-27 | 0.30823897 | 0.789 | 0.805 | 1.64E-23 |
| Kcnq2 | 1.16E-27 | -0.2527583 | 0.637 | 0.794 | 1.71E-23 |
| Fopnl | 1.21E-27 | -0.2553727 | 0.251 | 0.409 | 1.79E-23 |
| Thsd4 | 1.24E-27 | -0.2679639 | 0.281 | 0.44 | 1.83E-23 |
| Dlg1 | 1.41E-27 | 0.45950942 | 0.273 | 0.177 | 2.09E-23 |

| | | | | | |
|---|---|---|---|---|---|
| Lrp1 | 1.47E-27 | -0.2788127 | 0.458 | 0.637 | 2.17E-23 |
| Ap2a2 | 1.48E-27 | -0.2718482 | 0.518 | 0.688 | 2.19E-23 |
| Bsg | 1.71E-27 | -0.3888225 | 0.234 | 0.385 | 2.53E-23 |
| Mon2 | 1.96E-27 | 0.46100841 | 0.472 | 0.402 | 2.89E-23 |
| Kansl3 | 1.98E-27 | -0.2590517 | 0.312 | 0.483 | 2.92E-23 |
| Lrrtm4 | 2.06E-27 | 0.40719131 | 0.566 | 0.507 | 3.05E-23 |
| Herc2 | 2.20E-27 | 0.31941059 | 0.8 | 0.789 | 3.25E-23 |
| Elmod1 | 2.55E-27 | 0.39277066 | 0.641 | 0.598 | 3.76E-23 |
| Plppr4 | 3.16E-27 | -0.2861799 | 0.661 | 0.799 | 4.67E-23 |
| Ssbp4 | 3.56E-27 | -0.2734914 | 0.615 | 0.775 | 5.26E-23 |
| Dock9 | 4.40E-27 | -0.2559767 | 0.594 | 0.757 | 6.49E-23 |
| Rims2 | 5.12E-27 | -0.2897043 | 0.604 | 0.761 | 7.56E-23 |
| 4930430F08I | 5.49E-27 | -0.2543768 | 0.155 | 0.282 | 8.10E-23 |
| Adgrl1 | 5.49E-27 | 0.35726078 | 0.687 | 0.659 | 8.10E-23 |
| Ube2i | 5.58E-27 | -0.2650491 | 0.214 | 0.357 | 8.23E-23 |
| Kifc2 | 5.99E-27 | -0.2666928 | 0.539 | 0.718 | 8.84E-23 |
| Trip11 | 6.11E-27 | 0.36249032 | 0.675 | 0.644 | 9.02E-23 |
| Ap4m1 | 6.13E-27 | -0.2630999 | 0.234 | 0.385 | 9.04E-23 |
| Chchd2 | 6.26E-27 | -0.2598373 | 0.222 | 0.371 | 9.23E-23 |
| Bclaf1 | 7.33E-27 | 0.31655655 | 0.833 | 0.838 | 1.08E-22 |
| Tmeff2 | 7.89E-27 | -0.3267047 | 0.134 | 0.253 | 1.16E-22 |
| Zeb1 | 9.48E-27 | 0.42039667 | 0.343 | 0.246 | 1.40E-22 |
| Ogfrl1 | 9.85E-27 | -0.2606048 | 0.32 | 0.487 | 1.45E-22 |
| Rlf | 1.02E-26 | -0.265228 | 0.48 | 0.654 | 1.50E-22 |
| Myl6 | 1.05E-26 | -0.2592974 | 0.459 | 0.646 | 1.54E-22 |
| Cstf2 | 1.08E-26 | 0.43741964 | 0.508 | 0.447 | 1.59E-22 |
| Shisa7 | 1.41E-26 | 0.47097592 | 0.452 | 0.374 | 2.08E-22 |
| Gpm6a | 1.49E-26 | 0.37483405 | 0.594 | 0.55 | 2.20E-22 |
| Clmn | 1.61E-26 | 0.32251835 | 0.838 | 0.833 | 2.37E-22 |
| Htt | 2.20E-26 | 0.39281765 | 0.637 | 0.594 | 3.25E-22 |
| Atp5l | 2.29E-26 | -0.256296 | 0.385 | 0.568 | 3.38E-22 |
| Gtpbp2 | 3.69E-26 | -0.2551754 | 0.494 | 0.673 | 5.44E-22 |
| Nufip2 | 3.99E-26 | 0.42448854 | 0.579 | 0.545 | 5.89E-22 |
| Kif5a | 4.28E-26 | -0.2513105 | 0.369 | 0.536 | 6.32E-22 |
| Arhgap44 | 4.69E-26 | 0.47523224 | 0.375 | 0.284 | 6.91E-22 |
| Slc24a4 | 5.74E-26 | 0.4412689 | 0.293 | 0.197 | 8.46E-22 |
| Rbm26 | 6.00E-26 | 0.29199511 | 0.82 | 0.824 | 8.85E-22 |
| Bdnf | 6.85E-26 | 0.47853917 | 0.322 | 0.22 | 1.01E-21 |
| Haghl | 8.39E-26 | -0.2549313 | 0.29 | 0.45 | 1.24E-21 |
| Celf1 | 8.60E-26 | 0.35148463 | 0.743 | 0.733 | 1.27E-21 |
| Cpne4 | 9.88E-26 | 0.3025227 | 0.896 | 0.864 | 1.46E-21 |
| Gba2 | 1.58E-25 | 0.48979491 | 0.491 | 0.431 | 2.32E-21 |
| Rprd1a | 1.76E-25 | 0.43401981 | 0.519 | 0.455 | 2.60E-21 |

| | | | | | |
|---|---|---|---|---|---|
| Pdhb | 2.00E-25 | -0.2598644 | 0.397 | 0.564 | 2.96E-21 |
| Pwwp2a | 2.06E-25 | 0.40965908 | 0.61 | 0.596 | 3.04E-21 |
| Kmt2c | 2.37E-25 | 0.37342903 | 0.632 | 0.6 | 3.50E-21 |
| Ash1l | 2.40E-25 | 0.31889398 | 0.753 | 0.756 | 3.55E-21 |
| Cox5a | 2.77E-25 | -0.2590354 | 0.18 | 0.309 | 4.08E-21 |
| Stxbp5 | 2.77E-25 | 0.40811103 | 0.535 | 0.483 | 4.08E-21 |
| Pcdh1 | 3.89E-25 | 0.40211641 | 0.57 | 0.526 | 5.74E-21 |
| Hdgfl2 | 4.02E-25 | -0.2546744 | 0.195 | 0.33 | 5.93E-21 |
| Pgap1 | 4.29E-25 | 0.4635582 | 0.306 | 0.214 | 6.33E-21 |
| Epb41l3 | 4.36E-25 | -0.2847294 | 0.262 | 0.411 | 6.43E-21 |
| Idh3b | 4.42E-25 | -0.2596742 | 0.466 | 0.633 | 6.53E-21 |
| Fry | 5.00E-25 | 0.26919238 | 0.865 | 0.875 | 7.37E-21 |
| Brinp2 | 5.46E-25 | 0.26547554 | 0.897 | 0.894 | 8.06E-21 |
| Tial1 | 6.68E-25 | 0.3732904 | 0.601 | 0.561 | 9.86E-21 |
| Lix1 | 6.70E-25 | -0.2538069 | 0.169 | 0.293 | 9.89E-21 |
| Lgi1 | 1.04E-24 | -0.2511386 | 0.729 | 0.851 | 1.54E-20 |
| Tardbp | 1.07E-24 | 0.32338078 | 0.716 | 0.704 | 1.58E-20 |
| Strn4 | 1.55E-24 | -0.2531484 | 0.399 | 0.572 | 2.28E-20 |
| Atp11a | 1.64E-24 | 0.4407262 | 0.308 | 0.216 | 2.42E-20 |
| Strbp | 2.08E-24 | 0.30198328 | 0.778 | 0.762 | 3.07E-20 |
| Calm1 | 2.24E-24 | 0.30545807 | 0.862 | 0.898 | 3.31E-20 |
| Drap1 | 2.49E-24 | 0.43976731 | 0.528 | 0.479 | 3.68E-20 |
| Crebbp | 2.50E-24 | 0.41305744 | 0.522 | 0.466 | 3.70E-20 |
| Rps27 | 3.83E-24 | -0.2900413 | 0.255 | 0.396 | 5.66E-20 |
| Ppia | 4.91E-24 | -0.279816 | 0.527 | 0.702 | 7.25E-20 |
| Chd2 | 4.95E-24 | 0.43834345 | 0.547 | 0.507 | 7.30E-20 |
| Senp7 | 4.95E-24 | 0.38278062 | 0.564 | 0.517 | 7.31E-20 |
| Mast3 | 5.52E-24 | -0.2796557 | 0.497 | 0.645 | 8.14E-20 |
| Kcnv1 | 5.98E-24 | 0.44082676 | 0.314 | 0.223 | 8.83E-20 |
| Plxna2 | 7.24E-24 | 0.4389163 | 0.48 | 0.416 | 1.07E-19 |
| Kcnmb4 | 7.40E-24 | 0.45299365 | 0.362 | 0.277 | 1.09E-19 |
| Tsc22d4 | 7.42E-24 | -0.2699079 | 0.145 | 0.259 | 1.09E-19 |
| Cdh11 | 7.90E-24 | 0.41181355 | 0.605 | 0.58 | 1.17E-19 |
| Myh10 | 1.05E-23 | 0.44759253 | 0.413 | 0.336 | 1.54E-19 |
| Aplp2 | 1.05E-23 | 0.33410698 | 0.665 | 0.645 | 1.55E-19 |
| Usp34 | 1.16E-23 | 0.29335205 | 0.816 | 0.829 | 1.71E-19 |
| Celsr2 | 1.17E-23 | 0.43483723 | 0.427 | 0.351 | 1.73E-19 |
| Smg1 | 1.27E-23 | 0.37243958 | 0.65 | 0.643 | 1.87E-19 |
| Lingo3 | 1.61E-23 | 0.45190532 | 0.352 | 0.268 | 2.38E-19 |
| Dhx36 | 1.75E-23 | 0.32634659 | 0.714 | 0.706 | 2.58E-19 |
| Pde1a | 2.34E-23 | -0.26033 | 0.343 | 0.504 | 3.46E-19 |
| Setd5 | 2.76E-23 | 0.36295261 | 0.592 | 0.56 | 4.08E-19 |
| Ablim1 | 3.19E-23 | -0.2667117 | 0.576 | 0.725 | 4.71E-19 |

| | | | | | |
|---|---|---|---|---|---|
| Hmgn3 | 5.34E-23 | 0.35083847 | 0.698 | 0.681 | 7.87E-19 |
| Srsf1 | 5.55E-23 | 0.3690501 | 0.557 | 0.506 | 8.18E-19 |
| Fam135b | 5.65E-23 | 0.45306629 | 0.422 | 0.352 | 8.34E-19 |
| Neurod6 | 6.84E-23 | -0.253897 | 0.599 | 0.757 | 1.01E-18 |
| Slc25a4 | 8.30E-23 | -0.25801 | 0.471 | 0.646 | 1.22E-18 |
| Cacna1h | 8.79E-23 | 0.4108684 | 0.577 | 0.545 | 1.30E-18 |
| Ncoa1 | 9.61E-23 | 0.41506389 | 0.48 | 0.415 | 1.42E-18 |
| Hnrnpa3 | 1.14E-22 | 0.37813165 | 0.563 | 0.529 | 1.67E-18 |
| Med13 | 1.28E-22 | 0.40758152 | 0.452 | 0.384 | 1.88E-18 |
| Phf20 | 1.35E-22 | -0.2596007 | 0.487 | 0.662 | 1.99E-18 |
| Spock2 | 2.76E-22 | -0.2733316 | 0.333 | 0.487 | 4.07E-18 |
| Kmt2e | 2.77E-22 | 0.33381776 | 0.728 | 0.719 | 4.09E-18 |
| Ttyh1 | 3.68E-22 | 0.32523133 | 0.323 | 0.231 | 5.43E-18 |
| Frmd4a | 3.88E-22 | -0.2617593 | 0.614 | 0.766 | 5.72E-18 |
| Ccdc30 | 4.00E-22 | 0.35729166 | 0.609 | 0.58 | 5.91E-18 |
| Prrc2c | 4.52E-22 | 0.31393414 | 0.736 | 0.747 | 6.67E-18 |
| Zmynd11 | 4.60E-22 | 0.33948009 | 0.687 | 0.68 | 6.78E-18 |
| Nsf | 5.32E-22 | 0.28521955 | 0.831 | 0.845 | 7.85E-18 |
| Kmt2d | 6.69E-22 | 0.42475578 | 0.382 | 0.304 | 9.87E-18 |
| Ptma | 8.61E-22 | 0.42103834 | 0.387 | 0.307 | 1.27E-17 |
| Dscam | 9.03E-22 | 0.44281218 | 0.396 | 0.324 | 1.33E-17 |
| Zfp266 | 1.91E-21 | -0.2570284 | 0.556 | 0.716 | 2.81E-17 |
| Syt4 | 2.65E-21 | 0.43021157 | 0.313 | 0.229 | 3.91E-17 |
| Herc1 | 2.85E-21 | 0.27060049 | 0.819 | 0.832 | 4.21E-17 |
| Cdk5r1 | 5.96E-21 | 0.42597228 | 0.362 | 0.284 | 8.79E-17 |
| Olfml2b | 8.31E-21 | 0.41535837 | 0.39 | 0.316 | 1.23E-16 |
| Csde1 | 8.48E-21 | 0.39494937 | 0.522 | 0.483 | 1.25E-16 |
| Xpo1 | 8.52E-21 | 0.42676678 | 0.311 | 0.231 | 1.26E-16 |
| Tsga10 | 9.38E-21 | 0.45740662 | 0.418 | 0.359 | 1.38E-16 |
| Usp9x | 1.12E-20 | 0.38570715 | 0.464 | 0.404 | 1.66E-16 |
| Atp11b | 1.50E-20 | 0.26529509 | 0.811 | 0.818 | 2.22E-16 |
| Ncl | 2.42E-20 | 0.37167153 | 0.553 | 0.513 | 3.57E-16 |
| Ubr2 | 2.87E-20 | 0.34600557 | 0.554 | 0.514 | 4.24E-16 |
| Plbd2 | 3.58E-20 | 0.42793416 | 0.354 | 0.279 | 5.28E-16 |
| Rasa1 | 4.90E-20 | 0.41606542 | 0.316 | 0.242 | 7.23E-16 |
| Rev3l | 5.32E-20 | 0.3317754 | 0.644 | 0.632 | 7.85E-16 |
| Hmbox1 | 8.02E-20 | 0.38752143 | 0.508 | 0.463 | 1.18E-15 |
| Lrrn2 | 9.60E-20 | 0.40908209 | 0.491 | 0.449 | 1.42E-15 |
| Pde7a | 9.81E-20 | 0.40478233 | 0.339 | 0.26 | 1.45E-15 |
| Slc25a3 | 1.68E-19 | 0.28671414 | 0.708 | 0.698 | 2.48E-15 |
| mt-Nd1 | 1.68E-19 | 0.38835066 | 0.378 | 0.297 | 2.48E-15 |
| Matr3 | 1.74E-19 | 0.36685399 | 0.588 | 0.573 | 2.56E-15 |
| Rabgap1l | 1.74E-19 | -0.2590562 | 0.624 | 0.754 | 2.57E-15 |

| | | | | | |
|---|---|---|---|---|---|
| Hnrnph3 | 1.78E-19 | 0.40469069 | 0.334 | 0.259 | 2.63E-15 |
| S100pbp | 1.81E-19 | 0.40885285 | 0.308 | 0.231 | 2.68E-15 |
| Calr | 2.47E-19 | 0.41987159 | 0.454 | 0.405 | 3.65E-15 |
| Abl2 | 3.66E-19 | 0.37501013 | 0.528 | 0.481 | 5.41E-15 |
| Phip | 5.03E-19 | 0.35871233 | 0.65 | 0.64 | 7.43E-15 |
| Rev1 | 6.88E-19 | 0.33672863 | 0.627 | 0.613 | 1.01E-14 |
| Sntb2 | 8.52E-19 | 0.40998844 | 0.325 | 0.246 | 1.26E-14 |
| Rnf150 | 8.63E-19 | 0.40976059 | 0.42 | 0.358 | 1.27E-14 |
| Nfat5 | 9.29E-19 | 0.25589718 | 0.845 | 0.866 | 1.37E-14 |
| Shank2 | 1.03E-18 | 0.35021359 | 0.518 | 0.472 | 1.53E-14 |
| Psap | 1.24E-18 | 0.35604542 | 0.535 | 0.505 | 1.83E-14 |
| Sec62 | 1.34E-18 | 0.37918098 | 0.568 | 0.535 | 1.98E-14 |
| Gabrg2 | 1.51E-18 | 0.34813342 | 0.593 | 0.572 | 2.22E-14 |
| Robo1 | 1.61E-18 | 0.39962492 | 0.445 | 0.391 | 2.38E-14 |
| Ttll7 | 1.65E-18 | -0.2609958 | 0.469 | 0.626 | 2.43E-14 |
| Cep170 | 1.70E-18 | 0.40095278 | 0.426 | 0.37 | 2.50E-14 |
| Galnt11 | 1.84E-18 | 0.31906737 | 0.687 | 0.695 | 2.72E-14 |
| Ncald | 2.07E-18 | 0.34009139 | 0.473 | 0.407 | 3.06E-14 |
| Tspan7 | 2.29E-18 | 0.38899072 | 0.363 | 0.293 | 3.38E-14 |
| Usp15 | 2.54E-18 | 0.33298754 | 0.583 | 0.566 | 3.74E-14 |
| Ppp2r5a | 2.90E-18 | 0.39577585 | 0.267 | 0.193 | 4.28E-14 |
| Scd2 | 3.05E-18 | -0.3209434 | 0.152 | 0.255 | 4.50E-14 |
| Dzip1 | 3.27E-18 | 0.37632459 | 0.436 | 0.378 | 4.82E-14 |
| Mef2d | 3.71E-18 | 0.41797559 | 0.336 | 0.265 | 5.47E-14 |
| Map3k4 | 4.45E-18 | 0.4143704 | 0.393 | 0.33 | 6.56E-14 |
| Eif4a2 | 6.25E-18 | 0.34802772 | 0.48 | 0.431 | 9.22E-14 |
| Fam49a | 6.79E-18 | 0.36344143 | 0.435 | 0.376 | 1.00E-13 |
| Skil | 8.28E-18 | 0.38837202 | 0.482 | 0.435 | 1.22E-13 |
| Lats2 | 9.18E-18 | 0.38030596 | 0.25 | 0.174 | 1.35E-13 |
| Neto1 | 1.02E-17 | -0.2819301 | 0.279 | 0.397 | 1.50E-13 |
| Purb | 1.07E-17 | 0.33586795 | 0.597 | 0.578 | 1.59E-13 |
| Zfp638 | 1.23E-17 | 0.27850825 | 0.681 | 0.677 | 1.82E-13 |
| Birc2 | 1.36E-17 | 0.38241734 | 0.422 | 0.372 | 2.01E-13 |
| Agbl4 | 2.09E-17 | 0.39727225 | 0.323 | 0.253 | 3.09E-13 |
| Crtc3 | 2.17E-17 | 0.33214134 | 0.615 | 0.602 | 3.20E-13 |
| Cers4 | 2.32E-17 | 0.36162623 | 0.491 | 0.448 | 3.42E-13 |
| Prkar1a | 2.39E-17 | 0.35481873 | 0.529 | 0.494 | 3.53E-13 |
| Adamts1 | 3.19E-17 | 0.37460862 | 0.264 | 0.186 | 4.71E-13 |
| Srsf2 | 3.64E-17 | 0.30311159 | 0.649 | 0.641 | 5.38E-13 |
| Kif1a | 4.32E-17 | 0.31299973 | 0.688 | 0.691 | 6.38E-13 |
| Eif1b | 1.35E-16 | 0.44144146 | 0.444 | 0.399 | 1.99E-12 |
| Ccnl1 | 3.02E-16 | 0.28166038 | 0.719 | 0.742 | 4.45E-12 |
| Sacs | 3.14E-16 | 0.39112193 | 0.452 | 0.416 | 4.64E-12 |

| | | | | | |
|---|---|---|---|---|---|
| Sv2b | 4.69E-16 | 0.2659546 | 0.744 | 0.764 | 6.92E-12 |
| Hspa5 | 5.41E-16 | 0.39138394 | 0.347 | 0.284 | 7.99E-12 |
| Tafa5 | 6.33E-16 | 0.36538062 | 0.441 | 0.396 | 9.34E-12 |
| Atp1a1 | 6.35E-16 | 0.38003988 | 0.355 | 0.292 | 9.37E-12 |
| Cfap69 | 7.13E-16 | 0.40317492 | 0.294 | 0.23 | 1.05E-11 |
| Stk39 | 7.81E-16 | 0.39337159 | 0.298 | 0.234 | 1.15E-11 |
| Gphn | 9.81E-16 | 0.3511656 | 0.465 | 0.421 | 1.45E-11 |
| Prpf38b | 1.55E-15 | 0.32413967 | 0.613 | 0.612 | 2.29E-11 |
| Pigk | 1.74E-15 | 0.41095199 | 0.414 | 0.366 | 2.57E-11 |
| Homer1 | 1.86E-15 | 0.31109745 | 0.581 | 0.558 | 2.75E-11 |
| Gpr85 | 1.95E-15 | 0.38077315 | 0.457 | 0.416 | 2.88E-11 |
| Lrp8 | 2.03E-15 | 0.35133794 | 0.52 | 0.492 | 3.00E-11 |
| Dusp3 | 2.31E-15 | 0.36501386 | 0.402 | 0.347 | 3.42E-11 |
| Laptm4a | 2.58E-15 | 0.35152603 | 0.35 | 0.288 | 3.80E-11 |
| Chrd | 2.88E-15 | 0.36983664 | 0.503 | 0.488 | 4.26E-11 |
| Bptf | 3.62E-15 | 0.2629449 | 0.681 | 0.687 | 5.34E-11 |
| Pkd2 | 3.88E-15 | 0.38593824 | 0.402 | 0.351 | 5.72E-11 |
| Tnrc18 | 4.68E-15 | 0.36053691 | 0.291 | 0.224 | 6.91E-11 |
| Enox1 | 5.85E-15 | 0.31221554 | 0.623 | 0.618 | 8.64E-11 |
| Tmem161b | 6.87E-15 | 0.35776583 | 0.341 | 0.279 | 1.01E-10 |
| Ptk2 | 6.91E-15 | 0.31981396 | 0.552 | 0.539 | 1.02E-10 |
| Prrc2b | 9.03E-15 | 0.34900239 | 0.413 | 0.358 | 1.33E-10 |
| Slc16a7 | 1.03E-14 | 0.36026674 | 0.26 | 0.192 | 1.53E-10 |
| Vmp1 | 1.15E-14 | 0.34543521 | 0.367 | 0.309 | 1.70E-10 |
| App | 1.56E-14 | 0.27399071 | 0.634 | 0.646 | 2.31E-10 |
| Ranbp2 | 1.59E-14 | 0.32826798 | 0.471 | 0.446 | 2.34E-10 |
| Mib1 | 1.99E-14 | 0.33152169 | 0.483 | 0.448 | 2.93E-10 |
| Relch | 2.54E-14 | 0.32962284 | 0.498 | 0.472 | 3.75E-10 |
| Flrt2 | 2.88E-14 | 0.34074425 | 0.455 | 0.412 | 4.25E-10 |
| Smg7 | 3.22E-14 | 0.36724307 | 0.282 | 0.221 | 4.76E-10 |
| Cop1 | 3.55E-14 | 0.33267934 | 0.456 | 0.413 | 5.24E-10 |
| Fam76b | 3.77E-14 | 0.37661772 | 0.422 | 0.385 | 5.56E-10 |
| Otud4 | 4.17E-14 | 0.37690382 | 0.356 | 0.306 | 6.16E-10 |
| Araf | 4.96E-14 | 0.35365388 | 0.341 | 0.281 | 7.33E-10 |
| Cacna2d1 | 5.10E-14 | 0.34322805 | 0.443 | 0.401 | 7.53E-10 |
| Sptbn2 | 5.17E-14 | 0.29083591 | 0.577 | 0.566 | 7.63E-10 |
| Kcnb1 | 5.94E-14 | 0.35072447 | 0.384 | 0.332 | 8.77E-10 |
| Dync2h1 | 6.42E-14 | 0.35378872 | 0.451 | 0.412 | 9.47E-10 |
| Srsf6 | 7.74E-14 | 0.35673646 | 0.411 | 0.369 | 1.14E-09 |
| Mcu | 8.53E-14 | 0.35030199 | 0.445 | 0.409 | 1.26E-09 |
| Erc1 | 9.58E-14 | 0.2853924 | 0.659 | 0.651 | 1.41E-09 |
| Slc4a7 | 1.49E-13 | 0.29019068 | 0.767 | 0.794 | 2.20E-09 |
| Maged1 | 1.52E-13 | 0.3628106 | 0.438 | 0.399 | 2.24E-09 |

| | | | | | |
|---|---|---|---|---|---|
| Cyfip2 | 1.74E-13 | 0.31562445 | 0.559 | 0.557 | 2.57E-09 |
| Hmgcr | 1.87E-13 | 0.29863716 | 0.563 | 0.538 | 2.76E-09 |
| Ythdc2 | 2.10E-13 | 0.32516861 | 0.478 | 0.445 | 3.10E-09 |
| Tmem132b | 2.17E-13 | 0.34602428 | 0.313 | 0.254 | 3.21E-09 |
| Zfp62 | 2.25E-13 | 0.32836292 | 0.286 | 0.226 | 3.32E-09 |
| Baz1b | 2.40E-13 | 0.29810981 | 0.514 | 0.488 | 3.55E-09 |
| Npas4 | 2.47E-13 | 0.38731577 | 0.295 | 0.235 | 3.65E-09 |
| Bmpr2 | 2.54E-13 | 0.34051374 | 0.343 | 0.29 | 3.75E-09 |
| Ddx6 | 2.63E-13 | 0.33397381 | 0.342 | 0.286 | 3.88E-09 |
| Atrnl1 | 3.27E-13 | 0.32923843 | 0.277 | 0.217 | 4.83E-09 |
| Nup93 | 3.37E-13 | 0.34191711 | 0.414 | 0.369 | 4.97E-09 |
| Frrs1l | 4.15E-13 | 0.30909146 | 0.624 | 0.631 | 6.12E-09 |
| Rfx3 | 4.22E-13 | 0.29446504 | 0.456 | 0.414 | 6.22E-09 |
| Slc8a2 | 4.39E-13 | 0.3215999 | 0.265 | 0.205 | 6.47E-09 |
| Sh3gl1 | 4.46E-13 | 0.30820291 | 0.687 | 0.695 | 6.59E-09 |
| Brwd1 | 5.36E-13 | 0.29360626 | 0.588 | 0.588 | 7.91E-09 |
| Dync1h1 | 6.68E-13 | 0.29993369 | 0.607 | 0.622 | 9.86E-09 |
| Unc79 | 7.54E-13 | 0.32003606 | 0.504 | 0.477 | 1.11E-08 |
| Mbnl1 | 7.98E-13 | 0.35921308 | 0.383 | 0.337 | 1.18E-08 |
| Akap11 | 8.17E-13 | 0.32974567 | 0.491 | 0.467 | 1.21E-08 |
| Ppp3cb | 1.01E-12 | 0.31593723 | 0.551 | 0.544 | 1.50E-08 |
| Hnrnpul2 | 1.06E-12 | 0.34176875 | 0.315 | 0.26 | 1.57E-08 |
| Septin4 | 1.07E-12 | -0.2519794 | 0.198 | 0.291 | 1.58E-08 |
| Ppp6r2 | 1.07E-12 | 0.33584164 | 0.382 | 0.337 | 1.58E-08 |
| Tmem131 | 1.08E-12 | 0.35222137 | 0.404 | 0.36 | 1.59E-08 |
| Pip5k1b | 1.39E-12 | 0.31608853 | 0.476 | 0.438 | 2.05E-08 |
| Ice1 | 1.75E-12 | 0.2546899 | 0.743 | 0.767 | 2.59E-08 |
| Cep83 | 1.81E-12 | 0.36824128 | 0.418 | 0.383 | 2.67E-08 |
| Adcy9 | 2.63E-12 | 0.27727673 | 0.649 | 0.656 | 3.88E-08 |
| Zdhhc17 | 2.79E-12 | 0.32326461 | 0.316 | 0.261 | 4.12E-08 |
| Slitrk4 | 2.95E-12 | 0.34819459 | 0.455 | 0.421 | 4.35E-08 |
| Ppm1a | 3.12E-12 | 0.32819268 | 0.455 | 0.421 | 4.60E-08 |
| Cap2 | 3.37E-12 | 0.32820651 | 0.439 | 0.403 | 4.97E-08 |
| Lmo3 | 4.61E-12 | 0.34949643 | 0.257 | 0.2 | 6.80E-08 |
| Ywhaz | 4.75E-12 | 0.29449263 | 0.494 | 0.473 | 7.01E-08 |
| Me3 | 4.99E-12 | 0.28512475 | 0.621 | 0.627 | 7.37E-08 |
| Cpsf7 | 5.80E-12 | 0.30550519 | 0.419 | 0.377 | 8.56E-08 |
| Tanc2 | 6.10E-12 | 0.27755042 | 0.616 | 0.631 | 9.01E-08 |
| Elavl4 | 6.13E-12 | 0.35448788 | 0.306 | 0.252 | 9.04E-08 |
| Ppfia3 | 6.37E-12 | 0.33095097 | 0.385 | 0.344 | 9.41E-08 |
| Nfix | 6.74E-12 | 0.30381829 | 0.515 | 0.499 | 9.94E-08 |
| Ctxn1 | 8.71E-12 | 0.34337475 | 0.362 | 0.319 | 1.29E-07 |
| Psip1 | 9.71E-12 | 0.260325 | 0.691 | 0.713 | 1.43E-07 |

| | | | | | |
|---|---|---|---|---|---|
| Dscaml1 | 1.08E-11 | 0.32711513 | 0.321 | 0.268 | 1.59E-07 |
| Fam120a | 1.72E-11 | 0.32106452 | 0.429 | 0.397 | 2.54E-07 |
| Pde4dip | 2.21E-11 | 0.27271184 | 0.672 | 0.682 | 3.26E-07 |
| Canx | 2.22E-11 | 0.30195284 | 0.295 | 0.243 | 3.28E-07 |
| Aftph | 2.27E-11 | 0.31060768 | 0.343 | 0.293 | 3.35E-07 |
| Eif4g1 | 2.44E-11 | 0.31847607 | 0.496 | 0.479 | 3.60E-07 |
| Git2 | 2.63E-11 | 0.30558921 | 0.533 | 0.526 | 3.88E-07 |
| Nsmaf | 3.20E-11 | 0.32261907 | 0.398 | 0.367 | 4.72E-07 |
| Ubl3 | 3.24E-11 | 0.28951108 | 0.539 | 0.534 | 4.77E-07 |
| Smarcad1 | 3.49E-11 | 0.32229149 | 0.417 | 0.388 | 5.15E-07 |
| Tmem30a | 3.52E-11 | 0.31281527 | 0.334 | 0.287 | 5.19E-07 |
| Hdgfl3 | 3.64E-11 | 0.3145948 | 0.422 | 0.389 | 5.37E-07 |
| Ar | 4.52E-11 | 0.36018758 | 0.283 | 0.229 | 6.67E-07 |
| Il16 | 4.54E-11 | 0.34160843 | 0.342 | 0.298 | 6.69E-07 |
| Lzts1 | 6.42E-11 | 0.31953357 | 0.283 | 0.231 | 9.47E-07 |
| Bicdl1 | 6.58E-11 | 0.31978682 | 0.267 | 0.216 | 9.71E-07 |
| Prpf39 | 7.81E-11 | 0.26929581 | 0.585 | 0.591 | 1.15E-06 |
| Vps13b | 8.47E-11 | 0.31124126 | 0.41 | 0.378 | 1.25E-06 |
| Pds5b | 1.05E-10 | 0.32229325 | 0.351 | 0.309 | 1.55E-06 |
| Myo6 | 1.17E-10 | 0.25317894 | 0.613 | 0.626 | 1.72E-06 |
| Ubn1 | 1.26E-10 | 0.31976238 | 0.394 | 0.358 | 1.85E-06 |
| Tmem151b | 1.30E-10 | 0.32919225 | 0.307 | 0.26 | 1.92E-06 |
| Strn3 | 1.57E-10 | 0.25068737 | 0.655 | 0.671 | 2.31E-06 |
| Ryr1 | 1.59E-10 | 0.31748825 | 0.498 | 0.476 | 2.35E-06 |
| Eml6 | 1.66E-10 | 0.33067151 | 0.325 | 0.281 | 2.44E-06 |
| Zbtb37 | 1.72E-10 | 0.31069966 | 0.259 | 0.207 | 2.54E-06 |
| Trpm4 | 1.79E-10 | 0.35614615 | 0.272 | 0.224 | 2.64E-06 |
| Asap1 | 1.90E-10 | 0.346052 | 0.282 | 0.232 | 2.80E-06 |
| Mecp2 | 2.14E-10 | 0.31848622 | 0.318 | 0.272 | 3.15E-06 |
| Sik3 | 2.65E-10 | 0.30419546 | 0.337 | 0.295 | 3.90E-06 |
| Cep350 | 2.68E-10 | 0.29850829 | 0.484 | 0.468 | 3.95E-06 |
| Pspc1 | 2.84E-10 | 0.28989925 | 0.345 | 0.301 | 4.20E-06 |
| Clstn2 | 3.22E-10 | 0.30920169 | 0.455 | 0.439 | 4.76E-06 |
| Rnf111 | 5.19E-10 | 0.28928564 | 0.484 | 0.475 | 7.66E-06 |
| Vldlr | 5.20E-10 | 0.30102691 | 0.288 | 0.243 | 7.67E-06 |
| Sh3d19 | 5.55E-10 | 0.30581996 | 0.277 | 0.23 | 8.19E-06 |
| Pan3 | 6.21E-10 | 0.26755944 | 0.485 | 0.469 | 9.17E-06 |
| Hnrnpk | 7.45E-10 | 0.29086442 | 0.396 | 0.365 | 1.10E-05 |
| Herc3 | 7.54E-10 | 0.29837851 | 0.289 | 0.246 | 1.11E-05 |
| Tspan5 | 8.23E-10 | 0.30283379 | 0.361 | 0.323 | 1.21E-05 |
| Zmym2 | 9.16E-10 | 0.28886761 | 0.365 | 0.325 | 1.35E-05 |
| Erlec1 | 1.18E-09 | 0.28831331 | 0.358 | 0.321 | 1.74E-05 |
| Myo1b | 1.26E-09 | 0.3736526 | 0.318 | 0.282 | 1.86E-05 |

| | | | | | |
|---|---|---|---|---|---|
| Cyld | 1.64E-09 | 0.293259 | 0.429 | 0.406 | 2.42E-05 |
| Ptprj | 1.66E-09 | 0.30138155 | 0.381 | 0.351 | 2.45E-05 |
| Trip12 | 1.76E-09 | 0.26033877 | 0.509 | 0.506 | 2.60E-05 |
| Usp47 | 1.99E-09 | 0.27702384 | 0.293 | 0.246 | 2.93E-05 |
| Fam227a | 2.02E-09 | 0.31277458 | 0.273 | 0.227 | 2.97E-05 |
| Epc2 | 2.20E-09 | 0.3094718 | 0.29 | 0.248 | 3.25E-05 |
| Cers6 | 2.20E-09 | 0.28152892 | 0.455 | 0.438 | 3.25E-05 |
| Zscan26 | 3.00E-09 | 0.32127211 | 0.36 | 0.324 | 4.43E-05 |
| Ppp4r3a | 3.14E-09 | 0.27096788 | 0.372 | 0.339 | 4.64E-05 |
| Smarca5 | 3.47E-09 | 0.30055529 | 0.395 | 0.364 | 5.12E-05 |
| Cacnb4 | 3.65E-09 | 0.3352799 | 0.346 | 0.31 | 5.38E-05 |
| Dixdc1 | 4.39E-09 | 0.32055287 | 0.471 | 0.457 | 6.48E-05 |
| Chrm1 | 4.71E-09 | 0.31373745 | 0.299 | 0.259 | 6.94E-05 |
| Baz2b | 5.37E-09 | 0.30641712 | 0.368 | 0.337 | 7.93E-05 |
| D130043K22 | 5.46E-09 | 0.35837776 | 0.291 | 0.254 | 8.06E-05 |
| Clcn4 | 5.99E-09 | 0.2995053 | 0.393 | 0.363 | 8.84E-05 |
| Tm9sf3 | 6.16E-09 | 0.31759019 | 0.307 | 0.27 | 9.08E-05 |
| Myo9a | 6.37E-09 | 0.28569109 | 0.544 | 0.561 | 9.41E-05 |
| Camkk1 | 7.33E-09 | 0.32175614 | 0.285 | 0.244 | 0.0001082 |
| Fbxl16 | 7.84E-09 | 0.30694922 | 0.344 | 0.312 | 0.00011567 |
| Itm2b | 7.84E-09 | 0.27126258 | 0.469 | 0.461 | 0.00011569 |
| Igsf8 | 1.02E-08 | 0.29917071 | 0.27 | 0.228 | 0.00015087 |
| Smap1 | 1.06E-08 | 0.28929125 | 0.375 | 0.35 | 0.00015665 |
| Rapgef2 | 1.78E-08 | 0.25540922 | 0.425 | 0.404 | 0.00026324 |
| Agap1 | 1.90E-08 | 0.28576045 | 0.281 | 0.239 | 0.00028081 |
| Marchf6 | 2.02E-08 | 0.2638703 | 0.273 | 0.231 | 0.00029806 |
| Pcdh20 | 2.04E-08 | 0.31450488 | 0.37 | 0.346 | 0.0003008 |
| Dhx15 | 2.51E-08 | 0.29482079 | 0.352 | 0.323 | 0.00036989 |
| Fnbp4 | 2.52E-08 | 0.25883786 | 0.483 | 0.477 | 0.00037166 |
| Man2a2 | 2.82E-08 | 0.25314668 | 0.506 | 0.499 | 0.00041561 |
| Eif3c | 2.85E-08 | 0.28226314 | 0.39 | 0.366 | 0.00042015 |
| Ankhd1 | 3.00E-08 | 0.27432676 | 0.437 | 0.422 | 0.00044201 |
| Arnt | 3.01E-08 | 0.27745984 | 0.267 | 0.223 | 0.00044373 |
| Cnot7 | 3.05E-08 | 0.27983175 | 0.371 | 0.348 | 0.00044956 |
| Mark1 | 3.16E-08 | 0.30453999 | 0.348 | 0.315 | 0.00046614 |
| Larp4 | 3.39E-08 | 0.28096844 | 0.451 | 0.435 | 0.00050081 |
| Cd47 | 3.57E-08 | 0.28078266 | 0.441 | 0.424 | 0.00052621 |
| Ppp4r2 | 3.58E-08 | 0.27593145 | 0.287 | 0.248 | 0.00052875 |
| Stxbp1 | 3.94E-08 | 0.29283967 | 0.531 | 0.545 | 0.00058087 |
| Ppip5k1 | 3.98E-08 | 0.26460071 | 0.492 | 0.487 | 0.00058718 |
| Fbxo11 | 4.65E-08 | 0.28798203 | 0.28 | 0.238 | 0.00068682 |
| Bdp1 | 5.38E-08 | 0.27661247 | 0.47 | 0.464 | 0.00079343 |
| Ptprn2 | 5.40E-08 | 0.26763106 | 0.47 | 0.466 | 0.00079615 |

| | | | | | |
|---|---|---|---|---|---|
| Dock7 | 5.70E-08 | 0.28098305 | 0.289 | 0.253 | 0.00084088 |
| Sez6l | 6.36E-08 | 0.27086977 | 0.305 | 0.266 | 0.00093878 |
| Zfp385b | 6.45E-08 | 0.26273677 | 0.323 | 0.286 | 0.00095177 |
| Wsb1 | 7.23E-08 | 0.26652011 | 0.454 | 0.447 | 0.00106659 |
| Adam22 | 7.34E-08 | 0.27952392 | 0.389 | 0.37 | 0.00108341 |
| Purg | 1.01E-07 | 0.26642795 | 0.301 | 0.266 | 0.00149534 |
| Ptbp2 | 1.63E-07 | 0.2667381 | 0.353 | 0.325 | 0.0024009 |
| Mthfr | 1.73E-07 | 0.28526258 | 0.267 | 0.232 | 0.00256013 |
| C1qtnf4 | 1.84E-07 | 0.28152862 | 0.46 | 0.45 | 0.00271113 |
| Myrip | 2.08E-07 | 0.28101373 | 0.308 | 0.274 | 0.00306951 |
| Hipk2 | 2.13E-07 | 0.27066837 | 0.3 | 0.266 | 0.00313812 |
| Acvr2a | 2.34E-07 | 0.27375516 | 0.428 | 0.415 | 0.00345262 |
| Aph1b | 3.03E-07 | 0.28071929 | 0.322 | 0.293 | 0.00447293 |
| Acsl1 | 3.52E-07 | 0.25706193 | 0.478 | 0.472 | 0.00519357 |
| Ccdc39 | 4.04E-07 | 0.2654435 | 0.459 | 0.456 | 0.00596383 |
| Peli1 | 4.16E-07 | 0.2771271 | 0.419 | 0.401 | 0.00613136 |
| Wac | 4.25E-07 | 0.26841198 | 0.437 | 0.432 | 0.00627246 |
| Glg1 | 4.34E-07 | 0.25686228 | 0.414 | 0.403 | 0.00640001 |
| Fezf2 | 4.37E-07 | 0.27492925 | 0.262 | 0.227 | 0.00644876 |
| Zfp869 | 4.60E-07 | 0.28433628 | 0.32 | 0.292 | 0.00679184 |
| Gda | 4.69E-07 | 0.29774575 | 0.402 | 0.39 | 0.00692494 |
| Ift88 | 5.52E-07 | 0.27188891 | 0.34 | 0.314 | 0.0081496 |
| Apba2 | 5.80E-07 | 0.26962302 | 0.361 | 0.337 | 0.00856309 |
| Myo5a | 6.20E-07 | 0.25506775 | 0.393 | 0.373 | 0.00914984 |
| Fcho2 | 6.76E-07 | 0.27047504 | 0.387 | 0.367 | 0.00997407 |
| Hacd3 | 7.08E-07 | 0.27155161 | 0.29 | 0.258 | 0.01045202 |
| Smc5 | 7.99E-07 | 0.26876419 | 0.301 | 0.273 | 0.01178595 |
| Agfg1 | 9.50E-07 | 0.26649738 | 0.293 | 0.264 | 0.01401238 |
| Nhsl2 | 9.81E-07 | 0.27829698 | 0.36 | 0.337 | 0.01448219 |
| Lnpep | 1.35E-06 | 0.25998145 | 0.391 | 0.376 | 0.01997648 |
| Synpo | 1.41E-06 | 0.26728951 | 0.253 | 0.219 | 0.02080844 |
| Trrap | 1.71E-06 | 0.26707811 | 0.461 | 0.466 | 0.02516166 |
| Kdm6b | 1.98E-06 | 0.25905205 | 0.314 | 0.287 | 0.02918724 |
| Ago2 | 2.08E-06 | 0.28250122 | 0.377 | 0.363 | 0.03064057 |
| Ptprg | 2.14E-06 | 0.29400494 | 0.376 | 0.368 | 0.03150903 |
| Fam131a | 2.33E-06 | 0.26546414 | 0.463 | 0.464 | 0.03444222 |
| Gpr26 | 2.55E-06 | 0.2657642 | 0.323 | 0.302 | 0.03768722 |
| Zbtb11 | 2.80E-06 | 0.26255329 | 0.309 | 0.283 | 0.04135858 |
| Spen | 2.98E-06 | 0.29440899 | 0.346 | 0.327 | 0.04398349 |
| Trim33 | 3.22E-06 | 0.26536273 | 0.298 | 0.271 | 0.04749641 |
| Trp53bp1 | 3.27E-06 | 0.25159864 | 0.424 | 0.425 | 0.04818722 |
| Nucks1 | 3.34E-06 | 0.25998016 | 0.315 | 0.288 | 0.04934333 |
| Ddx3x | 3.94E-06 | 0.2542249 | 0.332 | 0.308 | 0.05810916 |

| | | | | | |
|---|---|---|---|---|---|
| Rsbn1 | 4.27E-06 | 0.25292467 | 0.323 | 0.298 | 0.06298376 |
| Lcor | 4.57E-06 | 0.25191602 | 0.329 | 0.304 | 0.06736861 |
| Frmpd3 | 4.89E-06 | 0.29235442 | 0.293 | 0.268 | 0.07214066 |
| Tnpo1 | 4.92E-06 | 0.25132675 | 0.285 | 0.257 | 0.07261641 |
| Fto | 4.98E-06 | 0.27175287 | 0.392 | 0.382 | 0.07348124 |
| Tle1 | 5.01E-06 | 0.26706568 | 0.377 | 0.367 | 0.07396485 |
| Ppil2 | 5.22E-06 | 0.26882 | 0.37 | 0.356 | 0.07697119 |
| B3galnt2 | 7.01E-06 | 0.26669979 | 0.367 | 0.357 | 0.10342816 |
| Add3 | 7.24E-06 | 0.25984649 | 0.294 | 0.27 | 0.1068098 |
| Brk1 | 7.30E-06 | 0.2705136 | 0.3 | 0.277 | 0.10765805 |
| Tnip2 | 9.98E-06 | 0.28060029 | 0.273 | 0.247 | 0.14724882 |
| Glt8d1 | 1.17E-05 | 0.25036503 | 0.364 | 0.353 | 0.17263812 |
| Tnk2 | 1.44E-05 | 0.25780865 | 0.263 | 0.238 | 0.21215155 |
| Creb1 | 1.70E-05 | 0.26346931 | 0.253 | 0.226 | 0.25125408 |
| Pcnp | 2.42E-05 | 0.25305638 | 0.401 | 0.402 | 0.35752951 |
| Setd3 | 2.62E-05 | 0.25762453 | 0.25 | 0.222 | 0.3869687 |
| Acbd4 | 2.75E-05 | 0.28342572 | 0.251 | 0.227 | 0.40594192 |
| Ndel1 | 3.07E-05 | 0.25387975 | 0.338 | 0.324 | 0.45268236 |
| Setbp1 | 3.96E-05 | 0.26269691 | 0.286 | 0.268 | 0.58389895 |
| Ube2q2 | 6.24E-05 | 0.25065428 | 0.288 | 0.268 | 0.92117761 |
| Ep400 | 7.49E-05 | 0.25301381 | 0.37 | 0.367 | 1 |
| Slc7a4 | 0.00020973 | 0.2812744 | 0.378 | 0.378 | 1 |
| Hirip3 | 0.00024793 | 0.25863494 | 0.253 | 0.236 | 1 |
| Septin1 | 0.00051951 | 0.27297827 | 0.306 | 0.3 | 1 |
| Cep290 | 0.00102801 | 0.26476898 | 0.377 | 0.383 | 1 |
