## Supplemental table2 for "Investigation of the Fasciola Cinereum, Absent in BTBR mice, and Comparison with the Hippocampal Area CA2"

| p_val | avg_log2FC | pct.1 | pct.2 | p_val_adj | cluster | gene |
|---|---|---|---|---|---|---|
| 1.40E-50 | 0.36134685 | 0.99 | 0.966 | 2.81E-47 | Type1 | Man1a2 |
| 1.25E-48 | 0.36920333 | 0.946 | 0.871 | 2.50E-45 | Type1 | Kcnk2 |
| 3.51E-45 | 0.38337768 | 0.993 | 0.96 | 7.03E-42 | Type1 | Akap13 |
| 5.91E-43 | 0.28975687 | 0.995 | 0.959 | 1.18E-39 | Type1 | Stxbp5l |
| 2.19E-42 | 0.35766368 | 0.967 | 0.904 | 4.38E-39 | Type1 | Ptpn5 |
| 9.40E-42 | 0.27980367 | 0.997 | 0.968 | 1.88E-38 | Type1 | Ptprd |
| 1.74E-41 | 0.36196997 | 0.994 | 0.959 | 3.49E-38 | Type1 | Asph |
| 5.50E-34 | 0.31265353 | 0.991 | 0.979 | 1.10E-30 | Type1 | Nebl |
| 1.35E-32 | 0.34419544 | 0.856 | 0.689 | 2.70E-29 | Type1 | Amigo2 |
| 2.83E-32 | 0.35741189 | 0.983 | 0.964 | 5.66E-29 | Type1 | Sms |
| 2.59E-30 | 0.37115942 | 0.923 | 0.823 | 5.17E-27 | Type1 | Pygo1 |
| 1.77E-25 | 0.40938503 | 0.95 | 0.911 | 3.55E-22 | Type1 | Sox5 |
| 1.15E-23 | 0.2535278 | 0.904 | 0.826 | 2.31E-20 | Type1 | Zdhhc23 |
| 1.89E-23 | 0.25627413 | 0.986 | 0.938 | 3.79E-20 | Type1 | Srgap2 |
| 4.45E-23 | 0.27176132 | 0.932 | 0.856 | 8.90E-20 | Type1 | Rap2b |
| 2.42E-18 | 0.28022923 | 0.769 | 0.651 | 4.84E-15 | Type1 | Pycr2 |
| 4.02E-15 | 0.25344163 | 0.948 | 0.886 | 8.03E-12 | Type1 | Ablim1 |
| 8.99E-08 | 0.25942271 | 0.944 | 0.908 | 0.00017975 | Type1 | Pkp2 |
| 2.32E-07 | 0.27887308 | 0.716 | 0.639 | 0.00046361 | Type1 | mt-Cytb |
| 0.0029687 | 0.30953697 | 0.728 | 0.711 | 1 | Type1 | mt-Co3 |
| 2.11E-187 | 0.58137827 | 1 | 0.958 | 4.22E-184 | Type2 | Stxbp5l |
| 2.21E-170 | 0.71490266 | 0.96 | 0.806 | 4.41E-167 | Type2 | Rapgef5 |
| 1.78E-148 | 0.34071927 | 1 | 0.996 | 3.57E-145 | Type2 | Zbtb20 |
| 3.39E-128 | 0.55628749 | 0.997 | 0.959 | 6.79E-125 | Type2 | Asph |
| 4.61E-128 | 0.63028076 | 0.983 | 0.857 | 9.22E-125 | Type2 | Kcnk2 |
| 3.05E-112 | 0.44356331 | 0.993 | 0.959 | 6.10E-109 | Type2 | Wipf3 |
| 5.42E-83 | 0.53234523 | 0.892 | 0.75 | 1.08E-79 | Type2 | Dapk1 |
| 5.28E-78 | 0.42154521 | 0.967 | 0.912 | 1.06E-74 | Type2 | Adcy1 |
| 1.29E-74 | 0.37525314 | 0.982 | 0.899 | 2.57E-71 | Type2 | Ptpn5 |
| 4.59E-74 | 0.4617532 | 0.875 | 0.718 | 9.19E-71 | Type2 | Ntsr2 |
| 6.65E-70 | 0.32220328 | 1 | 0.967 | 1.33E-66 | Type2 | Ptprd |
| 1.97E-69 | 0.53799433 | 0.842 | 0.759 | 3.94E-66 | Type2 | Epha10 |
| 2.51E-69 | 0.35945671 | 0.985 | 0.936 | 5.01E-66 | Type2 | Slc8a1 |
| 1.13E-67 | 0.36240459 | 0.983 | 0.919 | 2.27E-64 | Type2 | Brinp2 |
| 7.65E-66 | 0.46295635 | 0.897 | 0.765 | 1.53E-62 | Type2 | Hdac7 |
| 2.31E-63 | 0.39551825 | 0.859 | 0.84 | 4.62E-60 | Type2 | Echdc2 |
| 2.98E-63 | 0.38590785 | 0.904 | 0.833 | 5.95E-60 | Type2 | Pygo1 |
| 2.34E-62 | 0.41652061 | 0.932 | 0.842 | 4.68E-59 | Type2 | Large1 |
| 2.54E-61 | 0.39406027 | 0.938 | 0.839 | 5.08E-58 | Type2 | Atp11b |
| 1.71E-60 | 0.36890779 | 0.966 | 0.857 | 3.42E-57 | Type2 | Clmn |
| 7.46E-59 | 0.39837664 | 0.925 | 0.822 | 1.49E-55 | Type2 | Fat3 |
| 6.84E-57 | 0.41730445 | 0.824 | 0.701 | 1.37E-53 | Type2 | Zbtb1 |

| | | | | | | |
|---|---|---|---|---|---|---|
| 7.44E-55 | 0.3459907 | 0.88 | 0.882 | 1.49E-51 | Type2 | Onecut2 |
| 8.58E-55 | 0.29932592 | 0.997 | 0.964 | 1.72E-51 | Type2 | Erc2 |
| 2.73E-52 | 0.31021143 | 0.987 | 0.939 | 5.46E-49 | Type2 | Srgap2 |
| 3.20E-50 | 0.27175769 | 0.988 | 0.964 | 6.41E-47 | Type2 | Dclk1 |
| 2.85E-48 | 0.29923155 | 0.997 | 0.964 | 5.71E-45 | Type2 | Man1a2 |
| 8.89E-46 | 0.31604497 | 0.836 | 0.7 | 1.78E-42 | Type2 | Amigo2 |
| 2.95E-44 | 0.27853918 | 0.983 | 0.944 | 5.91E-41 | Type2 | Negr1 |
| 3.53E-44 | 0.35719993 | 0.857 | 0.739 | 7.05E-41 | Type2 | Cdkl2 |
| 3.91E-44 | 0.33810345 | 0.759 | 0.622 | 7.82E-41 | Type2 | Flna |
| 5.45E-42 | 0.3464048 | 0.778 | 0.701 | 1.09E-38 | Type2 | Kcna4 |
| 2.07E-41 | 0.30472795 | 0.862 | 0.796 | 4.15E-38 | Type2 | Uhrf1bp1l |
| 2.84E-40 | 0.29202235 | 0.649 | 0.523 | 5.68E-37 | Type2 | Flrt2 |
| 2.98E-40 | 0.32258185 | 0.919 | 0.821 | 5.95E-37 | Type2 | Zdhhc23 |
| 2.45E-39 | 0.27513086 | 0.654 | 0.53 | 4.90E-36 | Type2 | Slitrk4 |
| 4.84E-39 | 0.45298152 | 0.687 | 0.669 | 9.67E-36 | Type2 | St8sia6 |
| 7.05E-38 | 0.38514348 | 0.816 | 0.832 | 1.41E-34 | Type2 | Fkbp5 |
| 7.10E-38 | 0.41190644 | 0.831 | 0.839 | 1.42E-34 | Type2 | Dnah9 |
| 6.28E-36 | 0.31176838 | 0.806 | 0.706 | 1.26E-32 | Type2 | Pcdh10 |
| 2.82E-35 | 0.27928481 | 0.928 | 0.859 | 5.64E-32 | Type2 | Ralgapa1 |
| 2.96E-33 | 0.26731359 | 0.955 | 0.898 | 5.92E-30 | Type2 | Gabrb1 |
| 1.43E-32 | 0.33590346 | 0.634 | 0.599 | 2.87E-29 | Type2 | Cacng5 |
| 1.85E-32 | 0.2741868 | 0.94 | 0.854 | 3.71E-29 | Type2 | Rap2b |
| 4.47E-32 | 0.25044356 | 0.977 | 0.957 | 8.93E-29 | Type2 | Vps13c |
| 6.96E-32 | 0.27598594 | 0.693 | 0.608 | 1.39E-28 | Type2 | Rgs14 |
| 1.08E-31 | 0.25632744 | 0.927 | 0.896 | 2.17E-28 | Type2 | Ablim1 |
| 1.59E-31 | 0.3185251 | 0.863 | 0.787 | 3.19E-28 | Type2 | Rapgef4 |
| 1.78E-31 | 0.27663733 | 0.934 | 0.866 | 3.56E-28 | Type2 | Specc1 |
| 2.51E-31 | 0.27440489 | 0.919 | 0.852 | 5.02E-28 | Type2 | 2010300C02F |
| 2.61E-31 | 0.27702424 | 0.889 | 0.818 | 5.22E-28 | Type2 | Pde4dip |
| 6.93E-30 | 0.28298335 | 0.868 | 0.779 | 1.39E-26 | Type2 | Trank1 |
| 1.67E-28 | 0.25885194 | 0.841 | 0.772 | 3.33E-25 | Type2 | Erc1 |
| 8.78E-28 | 0.32103195 | 0.79 | 0.82 | 1.76E-24 | Type2 | Rasl11b |
| 1.51E-26 | 0.33068004 | 0.77 | 0.814 | 3.03E-23 | Type2 | Gm2115 |
| 6.57E-26 | 0.26479343 | 0.818 | 0.85 | 1.31E-22 | Type2 | Ccdc88c |
| 2.99E-25 | 0.26932712 | 0.575 | 0.536 | 5.98E-22 | Type2 | Fgf5 |
| 3.69E-25 | 0.33287042 | 0.711 | 0.699 | 7.37E-22 | Type2 | Egfem1 |
| 2.41E-24 | 0.4590347 | 0.702 | 0.751 | 4.83E-21 | Type2 | Vcan |
| 4.51E-24 | 0.26128837 | 0.691 | 0.654 | 9.03E-21 | Type2 | Bcan |
| 4.64E-24 | 0.3719482 | 0.723 | 0.775 | 9.28E-21 | Type2 | Osbpl3 |
| 1.53E-22 | 0.28144327 | 0.591 | 0.603 | 3.06E-19 | Type2 | Osbpl10 |
| 3.70E-22 | 0.29845921 | 0.573 | 0.629 | 7.40E-19 | Type2 | Fhod3 |
| 5.13E-22 | 0.2805578 | 0.716 | 0.693 | 1.03E-18 | Type2 | Kcnt1 |
| 9.01E-22 | 0.35314008 | 0.785 | 0.837 | 1.80E-18 | Type2 | Dpyd |

| | | | | | | |
|---|---|---|---|---|---|---|
| 5.20E-19 | 0.37424598 | 0.728 | 0.781 | 1.04E-15 | Type2 | Xkr6 |
| 2.00E-18 | 0.26541464 | 0.762 | 0.8 | 4.00E-15 | Type2 | Ttc21b |
| 7.34E-15 | 0.26223647 | 0.723 | 0.75 | 1.47E-11 | Type2 | Wdfy2 |
| 9.27E-10 | 0.28280795 | 0.721 | 0.803 | 1.85E-06 | Type2 | Kcnk9 |
| 2.24E-09 | 0.28235343 | 0.695 | 0.767 | 4.48E-06 | Type2 | Fam126a |
| 1.23E-06 | 0.29519083 | 0.544 | 0.614 | 0.00246782 | Type2 | Ctsc |
| 5.67E-06 | 0.30854869 | 0.562 | 0.649 | 0.01134998 | Type2 | Hcrtr2 |
| 0.00051779 | 0.28918012 | 0.66 | 0.774 | 1 | Type2 | Cfap100 |
| 1.12E-227 | 1.24381091 | 0.999 | 0.962 | 2.24E-224 | Type3 | Opcml |
| 1.59E-208 | 1.65933033 | 0.979 | 0.846 | 3.19E-205 | Type3 | Ccnd2 |
| 2.93E-155 | 1.00322281 | 0.976 | 0.898 | 5.86E-152 | Type3 | Rnf182 |
| 5.95E-139 | 1.11243184 | 0.835 | 0.555 | 1.19E-135 | Type3 | Nectin3 |
| 1.79E-132 | 0.80049676 | 0.986 | 0.952 | 3.58E-129 | Type3 | Cpne7 |
| 4.03E-129 | 0.43198176 | 1 | 0.997 | 8.05E-126 | Type3 | Snhg11 |
| 2.75E-119 | 0.79803282 | 0.992 | 0.948 | 5.49E-116 | Type3 | Cpne6 |
| 7.54E-112 | 0.99218722 | 0.836 | 0.661 | 1.51E-108 | Type3 | Neto1 |
| 9.17E-100 | 0.68281114 | 0.991 | 0.955 | 1.83E-96 | Type3 | Gpi1 |
| 1.47E-98 | 0.83894998 | 0.798 | 0.641 | 2.93E-95 | Type3 | Slc1a2 |
| 1.33E-92 | 0.85507571 | 0.865 | 0.794 | 2.66E-89 | Type3 | Peak1 |
| 8.82E-88 | 0.90715674 | 0.827 | 0.663 | 1.76E-84 | Type3 | Cpeb1 |
| 8.67E-84 | 0.52042079 | 0.994 | 0.958 | 1.73E-80 | Type3 | Rnf112 |
| 9.04E-81 | 0.92082582 | 0.753 | 0.677 | 1.81E-77 | Type3 | Parp8 |
| 1.39E-79 | 0.97904434 | 0.713 | 0.648 | 2.77E-76 | Type3 | Nptx1 |
| 9.55E-79 | 0.6758295 | 0.959 | 0.895 | 1.91E-75 | Type3 | Rabgap1l |
| 2.73E-78 | 0.83805354 | 0.828 | 0.732 | 5.46E-75 | Type3 | Raver2 |
| 3.76E-77 | 0.69651744 | 0.723 | 0.572 | 7.53E-74 | Type3 | Cntnap2 |
| 4.70E-77 | 0.68996651 | 0.927 | 0.792 | 9.39E-74 | Type3 | Adam11 |
| 1.80E-76 | 0.93649712 | 0.728 | 0.624 | 3.61E-73 | Type3 | Cdh24 |
| 7.59E-68 | 0.67603808 | 0.904 | 0.801 | 1.52E-64 | Type3 | Srcin1 |
| 1.68E-67 | 0.61183727 | 0.939 | 0.892 | 3.36E-64 | Type3 | Col4a2 |
| 4.80E-65 | 0.96624747 | 0.676 | 0.61 | 9.60E-62 | Type3 | Slit2 |
| 4.95E-65 | 0.59979937 | 0.931 | 0.864 | 9.89E-62 | Type3 | Grik4 |
| 1.30E-61 | 0.79701505 | 0.76 | 0.633 | 2.61E-58 | Type3 | Oxr1 |
| 1.50E-58 | 0.55674555 | 0.946 | 0.867 | 2.99E-55 | Type3 | Syt7 |
| 3.43E-57 | 0.54597383 | 0.884 | 0.733 | 6.87E-54 | Type3 | Aopep |
| 8.91E-57 | 0.54890695 | 0.942 | 0.834 | 1.78E-53 | Type3 | Unc5c |
| 1.67E-56 | 0.69342127 | 0.848 | 0.779 | 3.34E-53 | Type3 | Olfm1 |
| 2.90E-54 | 0.50716069 | 0.884 | 0.859 | 5.80E-51 | Type3 | Socs2 |
| 3.45E-54 | 0.52932373 | 0.949 | 0.91 | 6.91E-51 | Type3 | Syn2 |
| 1.20E-52 | 0.4946395 | 0.982 | 0.944 | 2.40E-49 | Type3 | Tmem108 |
| 3.25E-51 | 0.5875074 | 0.79 | 0.66 | 6.50E-48 | Type3 | Islr2 |
| 7.39E-51 | 0.64263242 | 0.856 | 0.772 | 1.48E-47 | Type3 | Jak1 |
| 7.09E-50 | 0.55121594 | 0.897 | 0.775 | 1.42E-46 | Type3 | Chgb |

| | | | | | | |
|---|---|---|---|---|---|---|
| 9.40E-48 | 0.42090492 | 0.986 | 0.953 | 1.88E-44 | Type3 | Gpm6b |
| 1.21E-47 | 0.49928942 | 0.945 | 0.872 | 2.42E-44 | Type3 | Ap2a2 |
| 6.42E-47 | 0.77566929 | 0.707 | 0.665 | 1.28E-43 | Type3 | Marchf1 |
| 1.09E-46 | 0.68219727 | 0.821 | 0.792 | 2.19E-43 | Type3 | Iqsec3 |
| 1.23E-46 | 0.48805567 | 0.962 | 0.924 | 2.46E-43 | Type3 | Nav3 |
| 1.59E-46 | 0.51366208 | 0.901 | 0.795 | 3.19E-43 | Type3 | Camta1 |
| 1.70E-45 | 0.54485136 | 0.773 | 0.658 | 3.40E-42 | Type3 | Iqgap1 |
| 3.54E-44 | 0.53558069 | 0.83 | 0.719 | 7.07E-41 | Type3 | Rimbp2 |
| 5.61E-44 | 0.53606636 | 0.726 | 0.609 | 1.12E-40 | Type3 | Entpd7 |
| 8.10E-44 | 0.56191934 | 0.806 | 0.769 | 1.62E-40 | Type3 | Tenm2 |
| 2.17E-43 | 0.69560328 | 0.704 | 0.571 | 4.33E-40 | Type3 | Per3 |
| 2.77E-43 | 0.51888685 | 0.696 | 0.562 | 5.53E-40 | Type3 | Ncam1 |
| 1.90E-41 | 0.59416391 | 0.817 | 0.751 | 3.81E-38 | Type3 | Kctd4 |
| 2.63E-40 | 0.68109141 | 0.652 | 0.598 | 5.25E-37 | Type3 | Epha3 |
| 5.73E-40 | 0.41214038 | 0.96 | 0.915 | 1.15E-36 | Type3 | Zmynd8 |
| 1.13E-39 | 0.47398213 | 0.751 | 0.603 | 2.26E-36 | Type3 | Ncald |
| 2.95E-39 | 0.44454018 | 0.753 | 0.583 | 5.91E-36 | Type3 | Cacna2d1 |
| 6.37E-39 | 0.70857378 | 0.757 | 0.732 | 1.27E-35 | Type3 | Scube1 |
| 1.92E-38 | 0.45131009 | 0.824 | 0.693 | 3.85E-35 | Type3 | Spock2 |
| 2.08E-38 | 0.35335736 | 0.484 | 0.337 | 4.17E-35 | Type3 | Nol4l |
| 2.73E-38 | 0.87447522 | 0.614 | 0.633 | 5.46E-35 | Type3 | Lnx1 |
| 4.43E-38 | 0.45956067 | 0.924 | 0.889 | 8.86E-35 | Type3 | Adgrb1 |
| 4.84E-38 | 0.44724009 | 0.942 | 0.92 | 9.68E-35 | Type3 | Trhde |
| 1.98E-37 | 0.392203 | 0.534 | 0.452 | 3.95E-34 | Type3 | Ephb1 |
| 5.81E-37 | 0.46056937 | 0.913 | 0.846 | 1.16E-33 | Type3 | Wscd2 |
| 1.53E-35 | 0.62280702 | 0.712 | 0.673 | 3.06E-32 | Type3 | Magi3 |
| 2.04E-35 | 0.61060861 | 0.607 | 0.569 | 4.08E-32 | Type3 | Lrrc10b |
| 2.92E-34 | 0.4354054 | 0.688 | 0.556 | 5.84E-31 | Type3 | Nsun7 |
| 3.06E-34 | 0.37558303 | 0.511 | 0.421 | 6.11E-31 | Type3 | Mapk11 |
| 1.76E-33 | 0.42114459 | 0.761 | 0.669 | 3.53E-30 | Type3 | Nrp2 |
| 4.27E-33 | 0.62452527 | 0.679 | 0.683 | 8.55E-30 | Type3 | Prkcb |
| 7.08E-32 | 0.50891435 | 0.622 | 0.52 | 1.42E-28 | Type3 | C530008M17 |
| 7.50E-32 | 0.30595852 | 0.994 | 0.978 | 1.50E-28 | Type3 | Nos1ap |
| 1.70E-31 | 0.27729725 | 0.991 | 0.967 | 3.40E-28 | Type3 | Cadps |
| 4.17E-31 | 0.59119088 | 0.65 | 0.644 | 8.35E-28 | Type3 | Kcnc2 |
| 1.52E-30 | 0.47773458 | 0.614 | 0.482 | 3.05E-27 | Type3 | Nptxr |
| 1.89E-30 | 0.47812561 | 0.595 | 0.65 | 3.78E-27 | Type3 | Iyd |
| 3.00E-30 | 0.69669285 | 0.705 | 0.68 | 6.00E-27 | Type3 | Trps1 |
| 4.51E-30 | 0.44250339 | 0.81 | 0.785 | 9.03E-27 | Type3 | Mas1 |
| 5.84E-30 | 0.4408467 | 0.901 | 0.848 | 1.17E-26 | Type3 | Klhl7 |
| 9.04E-30 | 0.40356765 | 0.77 | 0.65 | 1.81E-26 | Type3 | Arhgap17 |
| 9.34E-30 | 0.35715472 | 0.498 | 0.399 | 1.87E-26 | Type3 | Dpf3 |
| 1.99E-29 | 0.34645423 | 0.913 | 0.825 | 3.98E-26 | Type3 | Zfp950 |

| | | | | | | |
|---|---|---|---|---|---|---|
| 2.82E-29 | 0.31405476 | 0.609 | 0.469 | 5.64E-26 | Type3 | Sntb2 |
| 3.27E-29 | 0.69555231 | 0.679 | 0.658 | 6.54E-26 | Type3 | Shisa4 |
| 5.68E-29 | 0.51694698 | 0.588 | 0.541 | 1.14E-25 | Type3 | Edil3 |
| 3.35E-28 | 0.49076216 | 0.815 | 0.735 | 6.70E-25 | Type3 | Brinp1 |
| 2.52E-27 | 0.53655874 | 0.744 | 0.73 | 5.04E-24 | Type3 | Palmd |
| 4.91E-27 | 0.36608185 | 0.696 | 0.621 | 9.82E-24 | Type3 | Ppfia4 |
| 5.17E-27 | 0.4316029 | 0.754 | 0.669 | 1.03E-23 | Type3 | Camk2n1 |
| 5.77E-27 | 0.3792024 | 0.465 | 0.38 | 1.15E-23 | Type3 | Fhl1 |
| 1.17E-26 | 0.32843818 | 0.574 | 0.443 | 2.34E-23 | Type3 | Camkk1 |
| 1.19E-26 | 0.37541263 | 0.86 | 0.86 | 2.37E-23 | Type3 | Stmn2 |
| 1.52E-26 | 0.54812702 | 0.71 | 0.693 | 3.04E-23 | Type3 | Osbpl1a |
| 1.66E-26 | 0.39687545 | 0.877 | 0.805 | 3.32E-23 | Type3 | Mast3 |
| 3.03E-26 | 0.59740956 | 0.614 | 0.626 | 6.06E-23 | Type3 | Tspan17 |
| 4.31E-26 | 0.40993849 | 0.617 | 0.576 | 8.63E-23 | Type3 | Rab15 |
| 4.70E-26 | 0.36396068 | 0.981 | 0.969 | 9.40E-23 | Type3 | Akap13 |
| 7.76E-26 | 0.33071641 | 0.535 | 0.482 | 1.55E-22 | Type3 | Cbfa2t3 |
| 8.17E-26 | 0.41828632 | 0.542 | 0.444 | 1.63E-22 | Type3 | Scn3b |
| 1.09E-25 | 0.26490737 | 0.511 | 0.493 | 2.17E-22 | Type3 | Mndal |
| 2.41E-25 | 0.42544734 | 0.748 | 0.676 | 4.82E-22 | Type3 | Klhl2 |
| 1.33E-24 | 0.37820641 | 0.784 | 0.68 | 2.66E-21 | Type3 | Golm1 |
| 2.11E-24 | 0.46840712 | 0.716 | 0.7 | 4.23E-21 | Type3 | Gas2l1 |
| 6.30E-24 | 0.45644889 | 0.752 | 0.706 | 1.26E-20 | Type3 | Mgll |
| 6.44E-24 | 0.40945622 | 0.797 | 0.774 | 1.29E-20 | Type3 | Osbpl6 |
| 1.06E-23 | 0.56101595 | 0.675 | 0.631 | 2.12E-20 | Type3 | Sh3gl2 |
| 1.40E-23 | 0.36471899 | 0.753 | 0.645 | 2.80E-20 | Type3 | Cers4 |
| 3.74E-23 | 0.35395398 | 0.828 | 0.746 | 7.49E-20 | Type3 | Epha6 |
| 6.40E-23 | 0.57118889 | 0.581 | 0.578 | 1.28E-19 | Type3 | Gm45716 |
| 6.84E-23 | 0.51719269 | 0.729 | 0.729 | 1.37E-19 | Type3 | Tenm4 |
| 1.06E-22 | 0.40497554 | 0.668 | 0.669 | 2.13E-19 | Type3 | Sgsm1 |
| 2.37E-22 | 0.26039667 | 0.951 | 0.928 | 4.75E-19 | Type3 | Kcna1 |
| 2.76E-22 | 0.46166469 | 0.536 | 0.484 | 5.52E-19 | Type3 | Hap1 |
| 3.85E-22 | 0.32443826 | 0.562 | 0.538 | 7.70E-19 | Type3 | Tead1 |
| 5.75E-22 | 0.25822308 | 0.681 | 0.604 | 1.15E-18 | Type3 | Tnr |
| 7.26E-22 | 0.51770398 | 0.731 | 0.74 | 1.45E-18 | Type3 | Slc6a7 |
| 1.00E-21 | 0.31453918 | 0.824 | 0.761 | 2.00E-18 | Type3 | Smpd4 |
| 2.55E-21 | 0.53900005 | 0.669 | 0.702 | 5.11E-18 | Type3 | Zdhhc14 |
| 1.00E-20 | 0.3290013 | 0.586 | 0.507 | 2.00E-17 | Type3 | Cxadr |
| 1.44E-20 | 0.34393218 | 0.754 | 0.717 | 2.89E-17 | Type3 | Ptpre |
| 1.70E-20 | 0.35424841 | 0.665 | 0.568 | 3.39E-17 | Type3 | Kit |
| 3.99E-20 | 0.28098608 | 0.712 | 0.633 | 7.97E-17 | Type3 | Rasa2 |
| 2.12E-19 | 0.2881467 | 0.63 | 0.543 | 4.23E-16 | Type3 | Klf7 |
| 2.48E-19 | 0.31714901 | 0.805 | 0.738 | 4.96E-16 | Type3 | Klhl3 |
| 5.42E-19 | 0.32330552 | 0.796 | 0.758 | 1.08E-15 | Type3 | Tln2 |

| | | | | | | |
|---|---|---|---|---|---|---|
| 6.54E-19 | 0.39391528 | 0.691 | 0.655 | 1.31E-15 | Type3 | Grik2 |
| 9.23E-19 | 0.58617231 | 0.56 | 0.557 | 1.85E-15 | Type3 | Bok |
| 9.32E-19 | 0.4706089 | 0.739 | 0.749 | 1.86E-15 | Type3 | Homer3 |
| 1.36E-18 | 0.34205918 | 0.58 | 0.529 | 2.72E-15 | Type3 | Gpr68 |
| 2.35E-18 | 0.40478726 | 0.789 | 0.782 | 4.69E-15 | Type3 | Agrn |
| 2.53E-18 | 0.45999124 | 0.59 | 0.627 | 5.06E-15 | Type3 | Cacng2 |
| 6.09E-18 | 0.27865002 | 0.924 | 0.883 | 1.22E-14 | Type3 | Dlg2 |
| 1.62E-17 | 0.54812334 | 0.598 | 0.618 | 3.23E-14 | Type3 | Hes1 |
| 2.79E-17 | 0.2697075 | 0.752 | 0.673 | 5.58E-14 | Type3 | Tmem178b |
| 4.29E-17 | 0.58722364 | 0.606 | 0.647 | 8.58E-14 | Type3 | Gprin3 |
| 5.77E-17 | 0.50372049 | 0.683 | 0.716 | 1.15E-13 | Type3 | Dgkb |
| 6.08E-17 | 0.26399355 | 0.735 | 0.648 | 1.22E-13 | Type3 | Nsd2 |
| 1.44E-16 | 0.26839839 | 0.821 | 0.741 | 2.88E-13 | Type3 | Raph1 |
| 3.05E-16 | 0.25823475 | 0.926 | 0.855 | 6.10E-13 | Type3 | Khdrbs3 |
| 2.77E-15 | 0.26728195 | 0.771 | 0.722 | 5.55E-12 | Type3 | Daam1 |
| 2.82E-15 | 0.42454222 | 0.76 | 0.759 | 5.63E-12 | Type3 | Grip1 |
| 2.91E-15 | 0.41886406 | 0.745 | 0.787 | 5.83E-12 | Type3 | 5031439G07 |
| 1.29E-14 | 0.37491594 | 0.795 | 0.845 | 2.57E-11 | Type3 | Cadm3 |
| 1.02E-13 | 0.2748446 | 0.808 | 0.807 | 2.04E-10 | Type3 | Mcf2l |
| 2.14E-13 | 0.27893605 | 0.862 | 0.874 | 4.28E-10 | Type3 | Kcnc1 |
| 6.35E-13 | 0.30074178 | 0.534 | 0.522 | 1.27E-09 | Type3 | Mmd |
| 9.60E-13 | 0.31895981 | 0.766 | 0.759 | 1.92E-09 | Type3 | Ogfrl1 |
| 1.59E-12 | 0.3274371 | 0.519 | 0.566 | 3.19E-09 | Type3 | Ifi203 |
| 1.72E-12 | 0.29361428 | 0.476 | 0.467 | 3.44E-09 | Type3 | Cntnap5c |
| 2.57E-12 | 0.32510621 | 0.76 | 0.769 | 5.13E-09 | Type3 | Shisa6 |
| 2.93E-12 | 0.27864841 | 0.555 | 0.623 | 5.86E-09 | Type3 | Sorcs3 |
| 3.25E-12 | 0.30539464 | 0.707 | 0.654 | 6.49E-09 | Type3 | Iqcj |
| 5.87E-12 | 0.25903623 | 0.747 | 0.732 | 1.17E-08 | Type3 | Cdh10 |
| 1.47E-11 | 0.52535274 | 0.613 | 0.675 | 2.94E-08 | Type3 | Adamts9 |
| 2.17E-11 | 0.25512902 | 0.505 | 0.507 | 4.33E-08 | Type3 | Myrf |
| 3.80E-11 | 0.46443124 | 0.614 | 0.654 | 7.61E-08 | Type3 | Clmp |
| 5.18E-11 | 0.28476529 | 0.646 | 0.628 | 1.04E-07 | Type3 | Dennd1b |
| 5.40E-11 | 0.27646536 | 0.603 | 0.565 | 1.08E-07 | Type3 | Dnajc10 |
| 1.13E-10 | 0.29842983 | 0.738 | 0.743 | 2.26E-07 | Type3 | Kcnq5 |
| 1.20E-10 | 0.45445876 | 0.714 | 0.718 | 2.40E-07 | Type3 | Lmo4 |
| 3.62E-10 | 0.42955118 | 0.683 | 0.761 | 7.24E-07 | Type3 | Chst2 |
| 6.39E-10 | 0.3709465 | 0.533 | 0.63 | 1.28E-06 | Type3 | Nkd2 |
| 9.08E-10 | 0.27547075 | 0.506 | 0.52 | 1.82E-06 | Type3 | Epb41l4b |
| 9.35E-10 | 0.56647808 | 0.561 | 0.627 | 1.87E-06 | Type3 | Iqgap2 |
| 1.13E-09 | 0.43979963 | 0.685 | 0.739 | 2.26E-06 | Type3 | Tgfb2 |
| 1.74E-09 | 0.46742188 | 0.686 | 0.706 | 3.48E-06 | Type3 | Sdk2 |
| 1.87E-09 | 0.33655087 | 0.69 | 0.693 | 3.74E-06 | Type3 | Pcdh17 |
| 4.99E-09 | 0.25930224 | 0.625 | 0.614 | 9.98E-06 | Type3 | Grm1 |

| | | | | | | |
|---|---|---|---|---|---|---|
| 1.77E-08 | 0.35223151 | 0.694 | 0.718 | 3.54E-05 | Type3 | Dkk3 |
| 1.83E-08 | 0.53184672 | 0.615 | 0.663 | 3.66E-05 | Type3 | Nnat |
| 2.89E-08 | 0.25116436 | 0.631 | 0.613 | 5.78E-05 | Type3 | Slc35f3 |
| 6.19E-08 | 0.27983043 | 0.63 | 0.637 | 0.00012382 | Type3 | Fnbp1 |
| 8.06E-08 | 0.35844266 | 0.578 | 0.626 | 0.00016111 | Type3 | Fstl4 |
| 1.63E-07 | 0.50787414 | 0.549 | 0.626 | 0.0003261 | Type3 | Car4 |
| 4.10E-07 | 0.27910052 | 0.691 | 0.727 | 0.0008193 | Type3 | Dab2ip |
| 4.38E-07 | 0.3404079 | 0.682 | 0.717 | 0.00087589 | Type3 | D430041D05 |
| 6.22E-07 | 0.39000033 | 0.879 | 0.915 | 0.00124417 | Type3 | Col11a1 |
| 1.03E-06 | 0.27789412 | 0.749 | 0.788 | 0.00205262 | Type3 | Pdcd4 |
| 2.20E-06 | 0.36448022 | 0.637 | 0.678 | 0.00439492 | Type3 | Dcc |
| 9.94E-06 | 0.44308156 | 0.55 | 0.626 | 0.0198781 | Type3 | Bicc1 |
| 1.55E-05 | 0.39268607 | 0.6 | 0.644 | 0.03100775 | Type3 | Sema3e |
| 3.38E-05 | 0.29292175 | 0.606 | 0.657 | 0.06754932 | Type3 | Dlg5 |
| 4.65E-05 | 0.31020195 | 0.629 | 0.682 | 0.0930576 | Type3 | Armh4 |
| 4.86E-05 | 0.27724984 | 0.639 | 0.663 | 0.09724355 | Type3 | Dclk2 |
| 7.29E-05 | 0.36177519 | 0.545 | 0.622 | 0.14579703 | Type3 | Dcx |
| 8.35E-05 | 0.40331804 | 0.58 | 0.658 | 0.16704887 | Type3 | Vwa5b2 |
| 9.35E-05 | 0.34122567 | 0.665 | 0.731 | 0.1869053 | Type3 | Klf3 |
| 0.0001334 | 0.39760869 | 0.544 | 0.62 | 0.26680477 | Type3 | Adgra1 |
| 0.0009359 | 0.2724426 | 0.541 | 0.569 | 1 | Type3 | Ppp2r3a |
| 0.00245547 | 0.37755931 | 0.598 | 0.677 | 1 | Type3 | Evc2 |
| 0.00350415 | 0.2501417 | 0.563 | 0.607 | 1 | Type3 | D3Ertd751e |
| 0.0068264 | 0.30304301 | 0.646 | 0.706 | 1 | Type3 | Hspa12a |
| 0.00744473 | 0.30119513 | 0.621 | 0.715 | 1 | Type3 | Ackr1 |
| 1.89E-246 | 1.71813321 | 0.912 | 0.603 | 3.79E-243 | Type4 | C1ql3 |
| 1.05E-241 | 0.69789829 | 0.872 | 0.392 | 2.11E-238 | Type4 | Cntn5 |
| 9.68E-224 | 0.57296264 | 0.872 | 0.489 | 1.94E-220 | Type4 | Stxbp6 |
| 1.21E-220 | 0.78612316 | 0.89 | 0.552 | 2.41E-217 | Type4 | Cntnap5a |
| 1.55E-205 | 0.2715927 | 0.802 | 0.261 | 3.11E-202 | Type4 | Pcsk5 |
| 7.18E-198 | 0.59418596 | 0.877 | 0.603 | 1.44E-194 | Type4 | Lingo2 |
| 1.20E-195 | 0.59314943 | 0.855 | 0.45 | 2.40E-192 | Type4 | Prox1 |
| 1.72E-195 | 0.47356004 | 0.861 | 0.533 | 3.44E-192 | Type4 | Rasl10a |
| 2.38E-189 | 0.32211632 | 0.849 | 0.516 | 4.76E-186 | Type4 | Calb1 |
| 2.18E-182 | 0.85546507 | 0.869 | 0.591 | 4.36E-179 | Type4 | Adarb2 |
| 2.85E-177 | 0.57016595 | 0.872 | 0.531 | 5.69E-174 | Type4 | Brinp3 |
| 8.74E-176 | 0.57445183 | 0.879 | 0.564 | 1.75E-172 | Type4 | AI593442 |
| 1.39E-173 | 0.26115024 | 0.676 | 0.179 | 2.77E-170 | Type4 | Cldn5 |
| 7.22E-169 | 0.78915872 | 0.877 | 0.583 | 1.44E-165 | Type4 | Dock10 |
| 1.70E-168 | 0.49968836 | 0.822 | 0.344 | 3.40E-165 | Type4 | Fnbp1l |
| 1.17E-164 | 0.30768833 | 0.831 | 0.474 | 2.34E-161 | Type4 | Slc2a13 |
| 1.17E-161 | 0.3676452 | 0.861 | 0.601 | 2.35E-158 | Type4 | Cdh9 |
| 1.92E-161 | 0.25975409 | 0.857 | 0.574 | 3.84E-158 | Type4 | Gm12216 |

| | | | | | | |
|---|---|---|---|---|---|---|
| 3.75E-148 | 0.59781679 | 0.841 | 0.526 | 7.50E-145 | Type4 | Thsd7a |
| 8.93E-147 | 0.31213333 | 0.835 | 0.415 | 1.79E-143 | Type4 | Cxcl12 |
| 6.01E-140 | 0.28900058 | 0.869 | 0.586 | 1.20E-136 | Type4 | Iqgap2 |
| 1.28E-138 | 0.34371941 | 0.787 | 0.435 | 2.55E-135 | Type4 | Gad1 |
| 6.62E-133 | 0.30525172 | 0.827 | 0.449 | 1.32E-129 | Type4 | Vav3 |
| 1.14E-132 | 0.85807177 | 0.756 | 0.292 | 2.28E-129 | Type4 | Ptchd4 |
| 3.70E-131 | 0.4197934 | 0.831 | 0.497 | 7.41E-128 | Type4 | Gnal |
| 3.09E-124 | 0.49033846 | 0.792 | 0.358 | 6.18E-121 | Type4 | Maf |
| 2.93E-120 | 0.4778338 | 0.813 | 0.508 | 5.87E-117 | Type4 | Tead1 |
| 2.06E-118 | 0.47477054 | 0.872 | 0.66 | 4.11E-115 | Type4 | Prkcb |
| 5.99E-118 | 0.3012309 | 0.739 | 0.334 | 1.20E-114 | Type4 | Rassf8 |
| 7.10E-118 | 0.53730889 | 0.871 | 0.603 | 1.42E-114 | Type4 | Scg2 |
| 9.87E-116 | 1.15237029 | 0.894 | 0.581 | 1.97E-112 | Type4 | Ntng1 |
| 9.24E-115 | 0.37481853 | 0.778 | 0.415 | 1.85E-111 | Type4 | Jun |
| 2.98E-112 | 0.42418872 | 0.831 | 0.519 | 5.96E-109 | Type4 | Cacna2d2 |
| 3.81E-112 | 0.43479129 | 0.858 | 0.602 | 7.62E-109 | Type4 | Cspg5 |
| 1.85E-109 | 0.8948396 | 0.901 | 0.551 | 3.70E-106 | Type4 | Tenm1 |
| 7.91E-107 | 0.25979859 | 0.743 | 0.343 | 1.58E-103 | Type4 | Hmcn1 |
| 1.61E-103 | 0.42758219 | 0.71 | 0.327 | 3.21E-100 | Type4 | Ndst4 |
| 8.64E-99 | 0.34486698 | 0.833 | 0.561 | 1.73E-95 | Type4 | Rora |
| 3.09E-93 | 0.25709124 | 0.66 | 0.197 | 6.17E-90 | Type4 | Gm9925 |
| 1.28E-90 | 0.32173518 | 0.792 | 0.441 | 2.55E-87 | Type4 | Zmat4 |
| 1.24E-85 | 0.40778284 | 0.806 | 0.52 | 2.48E-82 | Type4 | Camk2g |
| 4.05E-84 | 0.53941246 | 0.91 | 0.694 | 8.09E-81 | Type4 | Arhgap20 |
| 1.58E-82 | 0.26963244 | 0.844 | 0.591 | 3.16E-79 | Type4 | Npy |
| 6.82E-82 | 0.31339773 | 0.833 | 0.464 | 1.36E-78 | Type4 | Flt1 |
| 1.56E-79 | 0.2507596 | 0.721 | 0.377 | 3.12E-76 | Type4 | Col19a1 |
| 8.49E-79 | 1.05730355 | 0.828 | 0.458 | 1.70E-75 | Type4 | Mef2c |
| 3.25E-77 | 0.35295049 | 0.855 | 0.605 | 6.49E-74 | Type4 | Ppp1r16b |
| 2.25E-76 | 0.59263258 | 0.798 | 0.518 | 4.49E-73 | Type4 | Edil3 |
| 1.20E-69 | 0.46471478 | 0.772 | 0.546 | 2.40E-66 | Type4 | Magi1 |
| 7.07E-67 | 0.42314234 | 0.827 | 0.465 | 1.41E-63 | Type4 | Vtn |
| 2.95E-66 | 0.26518175 | 0.798 | 0.591 | 5.91E-63 | Type4 | Sorcs3 |
| 3.31E-66 | 0.30533468 | 0.828 | 0.539 | 6.61E-63 | Type4 | Col25a1 |
| 8.11E-65 | 0.33461746 | 0.718 | 0.375 | 1.62E-61 | Type4 | Efnb3 |
| 1.79E-61 | 0.27642396 | 0.836 | 0.583 | 3.58E-58 | Type4 | Chl1 |
| 3.63E-61 | 0.30101417 | 0.828 | 0.584 | 7.26E-58 | Type4 | Dcx |
| 4.63E-61 | 0.47014166 | 0.775 | 0.496 | 9.25E-58 | Type4 | Rgs17 |
| 6.59E-61 | 0.33416292 | 0.687 | 0.403 | 1.32E-57 | Type4 | Nxph1 |
| 4.50E-60 | 0.38341392 | 0.827 | 0.495 | 8.99E-57 | Type4 | Slco1a4 |
| 6.16E-57 | 0.58014689 | 0.701 | 0.408 | 1.23E-53 | Type4 | Grid2 |
| 6.09E-54 | 0.80458959 | 0.806 | 0.555 | 1.22E-50 | Type4 | Ncam1 |
| 5.12E-52 | 0.32331147 | 0.88 | 0.662 | 1.02E-48 | Type4 | Itpr1 |

| | | | | | | |
|---|---|---|---|---|---|---|
| 6.85E-50 | 0.56479971 | 0.739 | 0.449 | 1.37E-46 | Type4 | Hecw1 |
| 1.29E-48 | 0.30093206 | 0.597 | 0.274 | 2.59E-45 | Type4 | Slc30a3 |
| 1.38E-48 | 0.82501351 | 0.627 | 0.382 | 2.76E-45 | Type4 | Cdh13 |
| 3.14E-46 | 0.3157803 | 0.819 | 0.579 | 6.28E-43 | Type4 | Mbp |
| 8.04E-44 | 0.29426825 | 0.279 | 0.425 | 1.61E-40 | Type4 | Arhgap29 |
| 4.52E-40 | 0.36103645 | 0.877 | 0.635 | 9.03E-37 | Type4 | Shisa4 |
| 1.42E-39 | 0.57844887 | 0.715 | 0.428 | 2.85E-36 | Type4 | Scn3b |
| 2.94E-38 | 0.46039178 | 0.776 | 0.513 | 5.87E-35 | Type4 | Ccdc85a |
| 2.20E-37 | 0.3485205 | 0.857 | 0.649 | 4.40E-34 | Type4 | Marchf1 |
| 3.73E-36 | 0.6883582 | 0.865 | 0.712 | 7.45E-33 | Type4 | Syne1 |
| 9.15E-35 | 0.37228338 | 0.901 | 0.695 | 1.83E-31 | Type4 | Lmo4 |
| 2.77E-34 | 0.29209795 | 0.869 | 0.688 | 5.54E-31 | Type4 | Ankrd33b |
| 4.31E-33 | 0.30900804 | 0.8 | 0.561 | 8.63E-30 | Type4 | Psmb5 |
| 3.69E-32 | 0.74914422 | 0.841 | 0.642 | 7.38E-29 | Type4 | mt-Cytb |
| 4.73E-31 | 0.27387704 | 0.268 | 0.355 | 9.45E-28 | Type4 | Tspan18 |
| 8.28E-31 | 0.52796384 | 0.923 | 0.773 | 1.66E-27 | Type4 | Olfm1 |
| 4.58E-30 | 0.34485634 | 0.274 | 0.374 | 9.16E-27 | Type4 | Ndst3 |
| 5.15E-30 | 0.71711179 | 0.591 | 0.35 | 1.03E-26 | Type4 | Pdzd2 |
| 1.71E-29 | 0.69525155 | 0.813 | 0.643 | 3.42E-26 | Type4 | Cers4 |
| 4.19E-29 | 0.67623375 | 0.775 | 0.608 | 8.37E-26 | Type4 | Ncald |
| 1.58E-28 | 0.28769858 | 0.748 | 0.553 | 3.16E-25 | Type4 | Tmod1 |
| 1.84E-28 | 1.02470111 | 0.608 | 0.381 | 3.69E-25 | Type4 | Cit |
| 1.11E-27 | 0.35156084 | 0.62 | 0.402 | 2.21E-24 | Type4 | Fras1 |
| 3.65E-27 | 0.75556733 | 0.784 | 0.642 | 7.31E-24 | Type4 | Dgkh |
| 4.52E-27 | 1.02078052 | 0.633 | 0.416 | 9.03E-24 | Type4 | Cald1 |
| 2.19E-26 | 0.44845481 | 0.839 | 0.663 | 4.37E-23 | Type4 | Camk2n1 |
| 2.57E-25 | 0.53802666 | 0.66 | 0.473 | 5.15E-22 | Type4 | Zfpm2 |
| 4.08E-23 | 0.41065571 | 0.85 | 0.69 | 8.17E-20 | Type4 | Atp6v1d |
| 4.53E-23 | 0.35533149 | 0.751 | 0.549 | 9.07E-20 | Type4 | Sox11 |
| 6.77E-23 | 0.29535465 | 0.535 | 0.351 | 1.35E-19 | Type4 | Ly6a |
| 1.92E-22 | 0.33464191 | 0.702 | 0.46 | 3.84E-19 | Type4 | Tspan3 |
| 3.13E-22 | 0.47738485 | 0.628 | 0.415 | 6.27E-19 | Type4 | Pitpnm2 |
| 1.58E-21 | 0.6053393 | 0.701 | 0.528 | 3.16E-18 | Type4 | Sparcl1 |
| 2.00E-21 | 0.34410608 | 0.828 | 0.619 | 4.00E-18 | Type4 | Plp1 |
| 9.12E-21 | 0.30607584 | 0.893 | 0.769 | 1.82E-17 | Type4 | Pdcd4 |
| 4.77E-20 | 0.41817668 | 0.762 | 0.57 | 9.55E-17 | Type4 | Per3 |
| 1.17E-19 | 0.26093777 | 0.849 | 0.629 | 2.33E-16 | Type4 | Cst3 |
| 4.45E-19 | 0.57880302 | 0.715 | 0.548 | 8.89E-16 | Type4 | Dock3 |
| 1.23E-18 | 0.30090593 | 0.783 | 0.625 | 2.47E-15 | Type4 | Cox7b |
| 5.03E-18 | 0.50371751 | 0.88 | 0.778 | 1.01E-14 | Type4 | Cadm1 |
| 1.94E-17 | 0.31229341 | 0.589 | 0.372 | 3.87E-14 | Type4 | Ttc39c |
| 2.68E-17 | 0.76969807 | 0.871 | 0.693 | 5.36E-14 | Type4 | Bsg |
| 4.71E-17 | 0.27276986 | 0.315 | 0.377 | 9.42E-14 | Type4 | Hlf |

| | | | | | | |
|---|---|---|---|---|---|---|
| 1.74E-14 | 0.29606109 | 0.836 | 0.637 | 3.49E-11 | Type4 | Fau |
| 2.04E-14 | 0.54792962 | 0.698 | 0.568 | 4.09E-11 | Type4 | mt-Atp6 |
| 2.44E-14 | 0.25776896 | 0.306 | 0.29 | 4.88E-11 | Type4 | Pcdh8 |
| 3.18E-14 | 0.55109142 | 0.767 | 0.676 | 6.36E-11 | Type4 | mt-Co1 |
| 3.46E-14 | 0.26725047 | 0.654 | 0.546 | 6.93E-11 | Type4 | Etl4 |
| 6.48E-14 | 0.4601369 | 0.728 | 0.602 | 1.30E-10 | Type4 | Tnr |
| 9.80E-14 | 0.61726073 | 0.663 | 0.526 | 1.96E-10 | Type4 | mt-Nd4 |
| 9.94E-14 | 0.47348313 | 0.883 | 0.826 | 1.99E-10 | Type4 | Hsp90aa1 |
| 1.54E-13 | 0.26203467 | 0.753 | 0.599 | 3.08E-10 | Type4 | Pnmal2 |
| 2.81E-13 | 0.43707446 | 0.816 | 0.713 | 5.62E-10 | Type4 | Akap6 |
| 3.01E-13 | 0.46455537 | 0.85 | 0.766 | 6.01E-10 | Type4 | Cplx2 |
| 3.59E-13 | 0.44286713 | 0.852 | 0.749 | 7.19E-10 | Type4 | Hspa8 |
| 7.70E-13 | 0.65867664 | 0.743 | 0.643 | 1.54E-09 | Type4 | mt-Co2 |
| 1.92E-12 | 0.30186839 | 0.729 | 0.582 | 3.85E-09 | Type4 | Gria4 |
| 2.45E-12 | 0.49177017 | 0.792 | 0.714 | 4.91E-09 | Type4 | Ptpre |
| 2.90E-12 | 0.25385478 | 0.956 | 0.96 | 5.79E-09 | Type4 | Atp1b1 |
| 4.54E-12 | 0.52905066 | 0.71 | 0.61 | 9.08E-09 | Type4 | Il1rap |
| 4.87E-12 | 0.41832456 | 0.665 | 0.537 | 9.75E-09 | Type4 | Slc2a3 |
| 6.90E-12 | 0.34464571 | 0.86 | 0.797 | 1.38E-08 | Type4 | Atp2b2 |
| 7.47E-12 | 0.3435243 | 0.97 | 0.935 | 1.49E-08 | Type4 | Atp6v0c |
| 2.18E-11 | 0.50508498 | 0.271 | 0.156 | 4.36E-08 | Type4 | Epb41 |
| 3.70E-11 | 0.29609401 | 0.942 | 0.931 | 7.39E-08 | Type4 | Calm1 |
| 6.22E-11 | 0.47701287 | 0.704 | 0.597 | 1.24E-07 | Type4 | Cacna2d1 |
| 8.08E-11 | 0.28435493 | 0.861 | 0.734 | 1.62E-07 | Type4 | Brinp1 |
| 4.65E-10 | 0.46276496 | 0.715 | 0.615 | 9.30E-07 | Type4 | Itm2b |
| 5.11E-10 | 0.3220991 | 0.687 | 0.531 | 1.02E-06 | Type4 | Pam |
| 6.91E-10 | 0.41371712 | 0.293 | 0.25 | 1.38E-06 | Type4 | Plekha2 |
| 1.03E-09 | 0.41233461 | 0.896 | 0.823 | 2.06E-06 | Type4 | Slc25a4 |
| 1.42E-09 | 0.33994125 | 0.309 | 0.254 | 2.85E-06 | Type4 | Tac1 |
| 5.87E-09 | 2.28939886 | 0.68 | 0.523 | 1.17E-05 | Type4 | Ptgds |
| 6.29E-09 | 0.38847985 | 0.715 | 0.62 | 1.26E-05 | Type4 | Serinc1 |
| 6.74E-09 | 0.49010106 | 0.521 | 0.215 | 1.35E-05 | Type4 | Adgrl2 |
| 1.11E-08 | 0.41432868 | 0.696 | 0.661 | 2.22E-05 | Type4 | Slc1a2 |
| 2.45E-08 | 0.27497867 | 0.354 | 0.373 | 4.90E-05 | Type4 | Casd1 |
| 4.79E-08 | 0.27855067 | 0.709 | 0.553 | 9.58E-05 | Type4 | Cox5a |
| 6.79E-08 | 0.43402186 | 0.323 | 0.275 | 0.00013573 | Type4 | Cntn4 |
| 7.87E-08 | 0.34441627 | 0.609 | 0.459 | 0.00015737 | Type4 | Apod |
| 8.89E-08 | 0.32381347 | 0.765 | 0.647 | 0.00017785 | Type4 | Grik2 |
| 9.14E-08 | 0.49278756 | 0.94 | 0.888 | 0.00018286 | Type4 | Nrgn |
| 1.75E-07 | 0.51084778 | 0.655 | 0.586 | 0.00035037 | Type4 | Kcnb1 |
| 3.52E-07 | 0.34291427 | 0.608 | 0.497 | 0.00070425 | Type4 | Cblb |
| 3.56E-07 | 0.36884364 | 0.376 | 0.383 | 0.00071259 | Type4 | Satb1 |
| 4.01E-07 | 0.49142315 | 0.641 | 0.559 | 0.00080234 | Type4 | Fam135b |

| | | | | | | |
|---|---|---|---|---|---|---|
| 6.70E-07 | 0.70295857 | 0.361 | 0.25 | 0.00134003 | Type4 | Lrrtm3 |
| 7.34E-07 | 0.31831279 | 0.364 | 0.325 | 0.00146847 | Type4 | Fam81a |
| 3.35E-06 | 0.34430238 | 0.474 | 0.161 | 0.00669851 | Type4 | Arhgap6 |
| 4.48E-06 | 0.41806193 | 0.734 | 0.714 | 0.00895526 | Type4 | mt-Co3 |
| 5.47E-06 | 0.43459369 | 0.335 | 0.267 | 0.010947 | Type4 | Camk2d |
| 1.40E-05 | 0.40724079 | 0.693 | 0.62 | 0.028087 | Type4 | Scn3a |
| 1.46E-05 | 0.28955148 | 0.447 | 0.486 | 0.02928193 | Type4 | Nap1l5 |
| 1.58E-05 | 0.30049517 | 0.576 | 0.45 | 0.03160801 | Type4 | Camkk1 |
| 2.60E-05 | 0.25586461 | 0.663 | 0.555 | 0.0520101 | Type4 | Hspa4 |
| 3.04E-05 | 0.25805133 | 0.876 | 0.88 | 0.06072829 | Type4 | Syt7 |
| 3.05E-05 | 0.33251716 | 0.668 | 0.562 | 0.06098396 | Type4 | Ttyh1 |
| 3.29E-05 | 0.27099738 | 0.715 | 0.626 | 0.06579772 | Type4 | Fnbp1 |
| 3.53E-05 | 0.32089569 | 0.356 | 0.336 | 0.07058735 | Type4 | Garnl3 |
| 5.92E-05 | 0.43989271 | 0.576 | 0.465 | 0.11846718 | Type4 | Rpl24 |
| 8.48E-05 | 0.53750537 | 0.465 | 0.147 | 0.16969308 | Type4 | Zfp536 |
| 0.0001288 | 0.39041441 | 0.748 | 0.675 | 0.25759662 | Type4 | Nrp2 |
| 0.00013538 | 0.27718253 | 0.761 | 0.721 | 0.27075397 | Type4 | St6galnac5 |
| 0.00014661 | 0.34878666 | 0.564 | 0.443 | 0.2932123 | Type4 | Pard3 |
| 0.00015334 | 0.37657571 | 0.608 | 0.486 | 0.30667032 | Type4 | Ssbp2 |
| 0.00017164 | 0.54030492 | 0.913 | 0.773 | 0.3432796 | Type4 | Ntm |
| 0.00019115 | 0.33210652 | 0.32 | 0.24 | 0.38230843 | Type4 | Pcdh11x |
| 0.00051173 | 0.31409493 | 0.375 | 0.336 | 1 | Type4 | Kcnh7 |
| 0.00067413 | 0.33590711 | 0.619 | 0.507 | 1 | Type4 | Hba-a2 |
| 0.00109683 | 0.34320167 | 0.458 | 0.472 | 1 | Type4 | Ncam2 |
| 0.00136594 | 0.2861972 | 0.455 | 0.455 | 1 | Type4 | Usp29 |
| 0.00172762 | 0.28421767 | 0.409 | 0.365 | 1 | Type4 | Elmo1 |
| 0.00296145 | 0.49716935 | 0.474 | 0.332 | 1 | Type4 | Arid5b |
| 0.00344189 | 0.25427135 | 0.761 | 0.724 | 1 | Type4 | Pde10a |
| 0.00344261 | 0.33540248 | 0.595 | 0.521 | 1 | Type4 | Xpr1 |
| 0.00496537 | 0.66862539 | 0.431 | 0.286 | 1 | Type4 | Sema5a |
| 0.00880095 | 0.40948849 | 0.583 | 0.509 | 1 | Type4 | Tspan7 |
| 0.00932814 | 0.26904018 | 0.353 | 0.281 | 1 | Type4 | Btg2 |
| 0.00967043 | 0.25414476 | 0.772 | 0.702 | 1 | Type4 | Rps8 |
| 3.89E-180 | 0.57511996 | 0.89 | 0.451 | 7.78E-177 | Type5 | Gjc3 |
| 7.33E-177 | 0.29332367 | 0.848 | 0.266 | 1.47E-173 | Type5 | Ermn |
| 2.23E-172 | 0.40782763 | 0.881 | 0.402 | 4.46E-169 | Type5 | Ugt8a |
| 2.27E-168 | 0.32304606 | 0.864 | 0.413 | 4.54E-165 | Type5 | Aspa |
| 4.91E-166 | 0.40136253 | 0.883 | 0.458 | 9.82E-163 | Type5 | Tspan2 |
| 4.66E-161 | 0.3350528 | 0.881 | 0.476 | 9.32E-158 | Type5 | Cldn11 |
| 2.03E-150 | 0.31668689 | 0.813 | 0.267 | 4.05E-147 | Type5 | Opalin |
| 1.14E-147 | 0.63072632 | 0.869 | 0.519 | 2.27E-144 | Type5 | Mag |
| 2.00E-147 | 0.25427586 | 0.85 | 0.446 | 3.99E-144 | Type5 | Apod |
| 6.20E-146 | 0.41430799 | 0.881 | 0.479 | 1.24E-142 | Type5 | Mal |

| | | | | | | |
|---|---|---|---|---|---|---|
| 3.13E-142 | 0.46712937 | 0.853 | 0.452 | 6.25E-139 | Type5 | Sytl2 |
| 3.29E-142 | 0.32746794 | 0.89 | 0.586 | 6.58E-139 | Type5 | Trf |
| 2.10E-141 | 0.53894874 | 0.893 | 0.52 | 4.20E-138 | Type5 | Cnp |
| 2.42E-141 | 0.53163257 | 0.879 | 0.552 | 4.84E-138 | Type5 | Mobp |
| 8.70E-140 | 0.27343826 | 0.874 | 0.551 | 1.74E-136 | Type5 | Gatm |
| 2.91E-131 | 0.63967193 | 0.909 | 0.581 | 5.82E-128 | Type5 | Mbp |
| 5.66E-131 | 0.37527 | 0.869 | 0.56 | 1.13E-127 | Type5 | Gnb4 |
| 7.21E-130 | 0.34302051 | 0.862 | 0.559 | 1.44E-126 | Type5 | Cdk19 |
| 8.17E-129 | 0.26675939 | 0.862 | 0.534 | 1.63E-125 | Type5 | Fa2h |
| 1.14E-123 | 0.26323606 | 0.862 | 0.478 | 2.29E-120 | Type5 | Myrf |
| 6.34E-120 | 0.93658023 | 0.916 | 0.62 | 1.27E-116 | Type5 | Plp1 |
| 1.34E-110 | 0.27855262 | 0.864 | 0.595 | 2.68E-107 | Type5 | Dock10 |
| 2.32E-108 | 0.41498688 | 0.867 | 0.561 | 4.65E-105 | Type5 | Phldb1 |
| 2.23E-87 | 0.28131258 | 0.818 | 0.38 | 4.45E-84 | Type5 | Efnb3 |
| 1.48E-84 | 0.25270304 | 0.857 | 0.444 | 2.95E-81 | Type5 | Magt1 |
| 2.73E-80 | 0.69551645 | 0.9 | 0.463 | 5.46E-77 | Type5 | Car2 |
| 3.13E-78 | 0.37116926 | 0.799 | 0.367 | 6.27E-75 | Type5 | Ccp110 |
| 7.85E-69 | 0.4668185 | 0.937 | 0.64 | 1.57E-65 | Type5 | Scd2 |
| 8.10E-68 | 0.52819538 | 0.871 | 0.479 | 1.62E-64 | Type5 | Lcorl |
| 9.88E-55 | 0.38265228 | 0.949 | 0.611 | 1.98E-51 | Type5 | Fnbp1 |
| 1.52E-54 | 0.53452184 | 0.928 | 0.587 | 3.03E-51 | Type5 | Septin4 |
| 3.84E-52 | 0.29521089 | 0.942 | 0.689 | 7.68E-49 | Type5 | Rrbp1 |
| 4.87E-50 | 0.30321688 | 0.846 | 0.471 | 9.74E-47 | Type5 | Hmgcs1 |
| 9.03E-42 | 0.30949061 | 0.902 | 0.591 | 1.81E-38 | Type5 | Tmeff2 |
| 4.43E-38 | 0.25688213 | 0.855 | 0.532 | 8.86E-35 | Type5 | Xiap |
| 4.34E-35 | 0.29587324 | 0.914 | 0.615 | 8.69E-32 | Type5 | Rpl37 |
| 2.77E-30 | 0.30115592 | 0.958 | 0.721 | 5.53E-27 | Type5 | Kndc1 |
| 3.31E-29 | 0.56974088 | 0.895 | 0.615 | 6.63E-26 | Type5 | Smad7 |
| 8.90E-27 | 0.25668665 | 0.808 | 0.49 | 1.78E-23 | Type5 | 3110021N24 |
| 1.05E-23 | 0.3284602 | 0.834 | 0.562 | 2.11E-20 | Type5 | Ncam1 |
| 1.05E-22 | 0.62153343 | 0.909 | 0.629 | 2.09E-19 | Type5 | Septin7 |
| 5.00E-21 | 0.35401676 | 0.769 | 0.478 | 1.00E-17 | Type5 | Ssbp2 |
| 1.26E-16 | 0.26404922 | 0.238 | 0.256 | 2.51E-13 | Type5 | Plekha2 |
| 2.57E-15 | 0.30270082 | 0.811 | 0.526 | 5.13E-12 | Type5 | Cacnb4 |
| 1.48E-13 | 0.34443184 | 0.925 | 0.714 | 2.96E-10 | Type5 | Auh |
| 1.67E-13 | 0.29115883 | 0.979 | 0.821 | 3.35E-10 | Type5 | Dnm3 |
| 6.89E-12 | 0.2883905 | 0.243 | 0.229 | 1.38E-08 | Type5 | Kitl |
| 1.10E-11 | 0.5227051 | 0.657 | 0.485 | 2.20E-08 | Type5 | Mef2c |
| 3.65E-11 | 0.62796771 | 0.722 | 0.613 | 7.30E-08 | Type5 | Il1rap |
| 1.35E-10 | 0.29205596 | 0.318 | 0.325 | 2.71E-07 | Type5 | Sema6d |
| 1.95E-10 | 0.63396792 | 0.678 | 0.598 | 3.91E-07 | Type5 | Ptn |
| 4.12E-10 | 0.4407843 | 0.963 | 0.836 | 8.25E-07 | Type5 | Plekha1 |
| 1.03E-09 | 0.27307233 | 0.864 | 0.636 | 2.05E-06 | Type5 | Cyld |

| | | | | | | |
|---|---|---|---|---|---|---|
| 3.64E-09 | 0.25419689 | 0.785 | 0.561 | 7.27E-06 | Type5 | Smc1a |
| 5.16E-09 | 0.42921581 | 0.276 | 0.273 | 1.03E-05 | Type5 | Timp3 |
| 2.78E-08 | 0.26275162 | 0.881 | 0.703 | 5.56E-05 | Type5 | mt-Co3 |
| 2.96E-08 | 0.2934176 | 0.755 | 0.648 | 5.91E-05 | Type5 | Trpm3 |
| 3.16E-08 | 0.42014098 | 0.977 | 0.934 | 6.33E-05 | Type5 | Hsp90ab1 |
| 1.40E-07 | 0.42649767 | 0.301 | 0.218 | 0.00027982 | Type5 | Cntn3 |
| 1.51E-07 | 0.40587679 | 0.977 | 0.819 | 0.00030276 | Type5 | Mical3 |
| 3.01E-07 | 0.48061722 | 0.75 | 0.645 | 0.00060202 | Type5 | Qk |
| 3.32E-07 | 0.30366604 | 0.584 | 0.373 | 0.00066317 | Type5 | Mettl23 |
| 7.01E-07 | 0.60163556 | 0.313 | 0.256 | 0.00140124 | Type5 | Tac1 |
| 1.18E-06 | 0.39468265 | 0.846 | 0.735 | 0.00236572 | Type5 | Git2 |
| 2.28E-06 | 0.67638483 | 0.563 | 0.43 | 0.00455368 | Type5 | Cald1 |
| 2.86E-06 | 0.25053804 | 0.35 | 0.331 | 0.00571935 | Type5 | Ddo |
| 3.30E-06 | 0.29961134 | 0.776 | 0.6 | 0.00660974 | Type5 | Mbnl1 |
| 5.68E-06 | 0.32907415 | 0.956 | 0.822 | 0.01135771 | Type5 | Hsp90aa1 |
| 0.00041335 | 0.30569454 | 0.808 | 0.678 | 0.82669531 | Type5 | Klhl2 |
| 0.00147479 | 0.43442446 | 0.629 | 0.594 | 1 | Type5 | Sash1 |
| 0.00194886 | 0.26661606 | 0.902 | 0.794 | 1 | Type5 | App |
| 0.00226625 | 0.31215771 | 0.776 | 0.725 | 1 | Type5 | Tulp4 |
| 0.00305721 | 0.29430254 | 0.988 | 0.868 | 1 | Type5 | Pex5l |
| 0.00357194 | 0.35151263 | 0.729 | 0.694 | 1 | Type5 | Jph1 |
| 0.00390767 | 0.40261877 | 0.381 | 0.314 | 1 | Type5 | Igsf3 |
| 0.00401913 | 0.50348414 | 0.367 | 0.167 | 1 | Type5 | Zfp536 |
| 0.00453757 | 0.29820718 | 0.561 | 0.429 | 1 | Type5 | Usp2 |
| 0.00489921 | 0.32896468 | 0.998 | 0.949 | 1 | Type5 | Cck |
| 1.03E-68 | 1.82992584 | 0.977 | 0.72 | 2.06E-65 | Type6 | Pde10a |
| 1.18E-66 | 1.41211694 | 1 | 0.891 | 2.36E-63 | Type6 | Ntrk2 |
| 2.24E-63 | 2.00313363 | 0.869 | 0.496 | 4.48E-60 | Type6 | Nr4a2 |
| 2.55E-50 | 1.55143402 | 0.847 | 0.429 | 5.10E-47 | Type6 | Bdnf |
| 2.95E-42 | 1.21913214 | 0.79 | 0.48 | 5.90E-39 | Type6 | Nr4a3 |
| 1.51E-36 | 1.14729977 | 0.665 | 0.318 | 3.01E-33 | Type6 | Zdbf2 |
| 8.67E-31 | 0.7931877 | 1 | 0.97 | 1.73E-27 | Type6 | Dclk1 |
| 2.62E-30 | 1.16651854 | 0.938 | 0.789 | 5.23E-27 | Type6 | Chgb |
| 3.62E-29 | 0.8207266 | 0.642 | 0.432 | 7.23E-26 | Type6 | Fosl2 |
| 1.67E-27 | 1.14940264 | 0.767 | 0.536 | 3.34E-24 | Type6 | Tent5a |
| 9.04E-26 | 1.00228765 | 0.739 | 0.494 | 1.81E-22 | Type6 | Cwc25 |
| 2.11E-25 | 1.07479655 | 0.705 | 0.402 | 4.22E-22 | Type6 | Adamts1 |
| 2.18E-23 | 0.59396174 | 0.67 | 0.44 | 4.36E-20 | Type6 | Mlip |
| 8.32E-23 | 0.89406302 | 0.557 | 0.281 | 1.66E-19 | Type6 | Btg2 |
| 2.10E-22 | 0.6421436 | 0.966 | 0.92 | 4.19E-19 | Type6 | Ptprn |
| 5.77E-22 | 0.98336471 | 0.614 | 0.396 | 1.15E-18 | Type6 | Alkal2 |
| 7.13E-22 | 0.95904381 | 0.722 | 0.455 | 1.43E-18 | Type6 | Npas4 |
| 1.66E-20 | 0.59654539 | 0.557 | 0.372 | 3.33E-17 | Type6 | Fos |

| | | | | | | |
|---|---|---|---|---|---|---|
| 5.28E-20 | 0.66741885 | 0.983 | 0.933 | 1.06E-16 | Type6 | Plppr4 |
| 8.79E-19 | 0.78872798 | 0.881 | 0.781 | 1.76E-15 | Type6 | Epha10 |
| 4.23E-18 | 0.89757674 | 0.881 | 0.749 | 8.46E-15 | Type6 | Homer1 |
| 5.60E-17 | 0.45728945 | 1 | 0.97 | 1.12E-13 | Type6 | Snap25 |
| 7.64E-17 | 0.90144059 | 0.676 | 0.553 | 1.53E-13 | Type6 | Tbc1d1 |
| 1.45E-16 | 0.72915348 | 0.903 | 0.826 | 2.89E-13 | Type6 | Dmxl1 |
| 1.98E-16 | 0.72580097 | 0.886 | 0.731 | 3.96E-13 | Type6 | Hmgcr |
| 6.85E-16 | 0.67729841 | 0.886 | 0.744 | 1.37E-12 | Type6 | Myo6 |
| 1.42E-15 | 0.6874487 | 0.83 | 0.714 | 2.84E-12 | Type6 | Lmo4 |
| 3.07E-15 | 0.68860512 | 0.83 | 0.693 | 6.15E-12 | Type6 | Eprs |
| 5.71E-15 | 0.79101601 | 0.705 | 0.543 | 1.14E-11 | Type6 | Pam |
| 3.56E-14 | 0.7302083 | 0.864 | 0.78 | 7.13E-11 | Type6 | Rgs7bp |
| 4.44E-14 | 0.36379672 | 0.619 | 0.526 | 8.88E-11 | Type6 | Gadd45b |
| 5.76E-14 | 0.62575341 | 0.784 | 0.68 | 1.15E-10 | Type6 | Sik2 |
| 5.99E-14 | 0.54963911 | 0.665 | 0.416 | 1.20E-10 | Type6 | Plpp6 |
| 6.05E-14 | 0.61349645 | 0.608 | 0.491 | 1.21E-10 | Type6 | Ptgs2 |
| 7.70E-14 | 0.6510018 | 0.841 | 0.744 | 1.54E-10 | Type6 | Brinp1 |
| 1.19E-13 | 0.73540385 | 0.835 | 0.7 | 2.38E-10 | Type6 | Ina |
| 1.57E-13 | 0.68651183 | 0.733 | 0.608 | 3.14E-10 | Type6 | Kctd12 |
| 9.56E-13 | 0.45116975 | 0.989 | 0.97 | 1.91E-09 | Type6 | Asph |
| 1.63E-12 | 0.42823835 | 0.67 | 0.469 | 3.26E-09 | Type6 | Vldlr |
| 1.89E-12 | 0.37375376 | 0.545 | 0.456 | 3.77E-09 | Type6 | Mest |
| 7.73E-12 | 0.5832638 | 0.682 | 0.518 | 1.55E-08 | Type6 | Syt4 |
| 8.14E-12 | 0.54677511 | 0.886 | 0.84 | 1.63E-08 | Type6 | Errfi1 |
| 2.70E-11 | 0.4579493 | 0.949 | 0.893 | 5.41E-08 | Type6 | Atp2a2 |
| 2.71E-11 | 0.55547652 | 0.915 | 0.907 | 5.43E-08 | Type6 | Igf1r |
| 8.09E-11 | 1.07650094 | 0.636 | 0.632 | 1.62E-07 | Type6 | Scg2 |
| 1.41E-10 | 0.58571244 | 0.608 | 0.446 | 2.81E-07 | Type6 | Coq10b |
| 1.71E-10 | 0.29450597 | 0.71 | 0.551 | 3.42E-07 | Type6 | Snx30 |
| 8.16E-10 | 0.50832913 | 0.795 | 0.706 | 1.63E-06 | Type6 | Slk |
| 9.51E-10 | 0.34320436 | 0.568 | 0.38 | 1.90E-06 | Type6 | Kcnv1 |
| 1.02E-09 | 0.50772265 | 0.886 | 0.809 | 2.03E-06 | Type6 | Kdm7a |
| 1.22E-09 | 0.49288458 | 0.903 | 0.855 | 2.44E-06 | Type6 | Klhl7 |
| 1.31E-09 | 0.49775185 | 0.676 | 0.611 | 2.62E-06 | Type6 | Tmeff2 |
| 1.32E-09 | 0.54825978 | 0.915 | 0.909 | 2.64E-06 | Type6 | Col11a1 |
| 1.63E-09 | 0.47463839 | 0.739 | 0.555 | 3.25E-06 | Type6 | Flrt2 |
| 2.09E-09 | 0.4561276 | 0.653 | 0.525 | 4.18E-06 | Type6 | Hdac9 |
| 2.89E-09 | 0.46371441 | 0.83 | 0.702 | 5.78E-06 | Type6 | Fam126b |
| 5.47E-09 | 0.61437666 | 0.608 | 0.478 | 1.09E-05 | Type6 | Nap1l5 |
| 5.77E-09 | 0.64212409 | 0.71 | 0.708 | 1.15E-05 | Type6 | Ankrd33b |
| 7.27E-09 | 0.51419172 | 0.756 | 0.732 | 1.45E-05 | Type6 | Palmd |
| 1.35E-08 | 0.50456943 | 0.642 | 0.608 | 2.69E-05 | Type6 | Adgra1 |
| 2.21E-08 | 0.57755689 | 0.568 | 0.462 | 4.41E-05 | Type6 | Lats2 |

| | | | | | | | |
|---|---|---|---|---|---|---|---|
| 5.57E-08 | 0.55703585 | 0.648 | 0.548 | 0.00011147 | Type6 | | Slc2a3 |
| 1.55E-07 | 0.26824706 | 0.653 | 0.584 | 0.00030954 | Type6 | | Gadd45g |
| 1.63E-07 | 0.44748573 | 0.67 | 0.674 | 0.00032597 | Type6 | | Spag4 |
| 2.00E-07 | 0.4149976 | 0.75 | 0.613 | 0.00040074 | Type6 | | Ranbp2 |
| 2.02E-07 | 0.58596875 | 0.528 | 0.493 | 0.0004045 | Type6 | | Pcsk1 |
| 2.52E-07 | 0.45180229 | 0.96 | 0.949 | 0.00050314 | Type6 | | Tmem108 |
| 2.58E-07 | 0.45498943 | 0.733 | 0.691 | 0.00051539 | Type6 | | Pcdh17 |
| 3.17E-07 | 0.56759351 | 0.761 | 0.726 | 0.00063337 | Type6 | | Gpt2 |
| 3.32E-07 | 0.33410169 | 0.932 | 0.905 | 0.00066397 | Type6 | | Dmd |
| 5.11E-07 | 0.37489654 | 0.938 | 0.898 | 0.00102187 | Type6 | | Hnrnpa2b1 |
| 5.31E-07 | 0.39100895 | 0.733 | 0.622 | 0.001063 | Type6 | | Ncald |
| 6.30E-07 | 0.43112044 | 0.727 | 0.637 | 0.00126077 | Type6 | | Myh10 |
| 9.53E-07 | 0.41374785 | 0.909 | 0.873 | 0.00190545 | Type6 | | Peg3 |
| 1.22E-06 | 0.28059458 | 0.835 | 0.725 | 0.00244988 | Type6 | | Syne1 |
| 1.92E-06 | 0.35003484 | 0.58 | 0.51 | 0.00383914 | Type6 | | Cpeb3 |
| 2.34E-06 | 0.31655974 | 0.801 | 0.684 | 0.00468659 | Type6 | | Btbd3 |
| 2.47E-06 | 0.44679614 | 0.767 | 0.722 | 0.00494928 | Type6 | | Kcna4 |
| 2.54E-06 | 0.27400059 | 0.602 | 0.588 | 0.00508501 | Type6 | | BC049715 |
| 2.82E-06 | 0.37811918 | 0.79 | 0.716 | 0.00564439 | Type6 | | Plk2 |
| 3.15E-06 | 0.27050638 | 0.557 | 0.438 | 0.00630369 | Type6 | | Pgap1 |
| 5.41E-06 | 0.48290258 | 0.79 | 0.727 | 0.01082151 | Type6 | | Tulp4 |
| 8.41E-06 | 0.30251102 | 0.818 | 0.776 | 0.01681695 | Type6 | | Osbpl6 |
| 8.86E-06 | 0.34759807 | 0.949 | 0.92 | 0.01771728 | Type6 | | Cnrip1 |
| 1.05E-05 | 0.3444099 | 0.591 | 0.527 | 0.02108011 | Type6 | | Fbxo33 |
| 1.38E-05 | 0.29205029 | 0.915 | 0.902 | 0.02759296 | Type6 | | Efr3b |
| 1.63E-05 | 0.30540956 | 0.506 | 0.441 | 0.03266447 | Type6 | | Bcl6 |
| 1.94E-05 | 0.509342 | 0.71 | 0.731 | 0.03877946 | Type6 | | Tgfb2 |
| 2.40E-05 | 0.27599108 | 0.716 | 0.674 | 0.04795765 | Type6 | | Iqgap1 |
| 3.12E-05 | 0.29967839 | 0.756 | 0.696 | 0.06243646 | Type6 | | Armcx3 |
| 3.66E-05 | 0.2780165 | 0.938 | 0.87 | 0.0731867 | Type6 | | 2010300C02I |
| 4.71E-05 | 0.30882672 | 0.619 | 0.658 | 0.094274 | Type6 | | Galnt3 |
| 5.26E-05 | 0.30669284 | 0.847 | 0.772 | 0.10522007 | Type6 | | Fbxw7 |
| 5.97E-05 | 0.31624571 | 0.665 | 0.675 | 0.11941333 | Type6 | | Dusp5 |
| 8.95E-05 | 0.54902692 | 0.75 | 0.804 | 0.17902096 | Type6 | | Sel1l3 |
| 9.62E-05 | 0.37599735 | 0.636 | 0.589 | 0.19237469 | Type6 | | Ptprk |
| 9.70E-05 | 0.29008846 | 0.568 | 0.536 | 0.19390896 | Type6 | | Sqle |
| 9.99E-05 | 0.35153261 | 0.58 | 0.52 | 0.19977941 | Type6 | | Isca1 |
| 0.00014357 | 0.30517856 | 0.949 | 0.923 | 0.28713195 | Type6 | | Trhde |
| 0.00020013 | 0.34485763 | 0.602 | 0.573 | 0.40025814 | Type6 | | Rap2a |
| 0.00022745 | 0.30432726 | 0.54 | 0.493 | 0.4549067 | Type6 | | Nptx2 |
| 0.0002282 | 0.40410877 | 0.756 | 0.806 | 0.45639022 | Type6 | | Peak1 |
| 0.00037363 | 0.2592668 | 0.824 | 0.714 | 0.74726047 | Type6 | | Bcl11b |
| 0.00038715 | 0.38006807 | 0.733 | 0.701 | 0.77429212 | Type6 | | Arl5b |

| | | | | | | |
|---|---|---|---|---|---|---|
| 0.0004359 | 0.48617443 | 0.756 | 0.776 | 0.87179787 | Type6 | Nefl |
| 0.0007258 | 0.25961679 | 0.96 | 0.912 | 1 | Type6 | Epha4 |
| 0.00089714 | 0.37374696 | 0.693 | 0.698 | 1 | Type6 | Zdhhc14 |
| 0.00172999 | 0.26239489 | 0.705 | 0.698 | 1 | Type6 | Nrn1 |
| 0.00176363 | 0.27000976 | 0.381 | 0.346 | 1 | Type6 | Arid5b |
| 0.00206812 | 0.25472295 | 0.602 | 0.581 | 1 | Type6 | Zswim6 |
| 0.00230169 | 0.29258828 | 0.562 | 0.587 | 1 | Type6 | Ipmk |
| 0.00263349 | 0.33674983 | 0.75 | 0.728 | 1 | Type6 | Auh |
| 0.00291617 | 0.25178642 | 0.773 | 0.738 | 1 | Type6 | 4930447C04I |
| 0.00599815 | 0.39089997 | 0.636 | 0.698 | 1 | Type6 | Nefm |
| 5.14E-93 | 0.64064465 | 0.861 | 0.249 | 1.03E-89 | Type7 | Ednrb |
| 7.58E-90 | 0.8328672 | 0.879 | 0.369 | 1.52E-86 | Type7 | Sox6 |
| 3.34E-88 | 0.75115133 | 0.873 | 0.404 | 6.67E-85 | Type7 | Gjb6 |
| 3.52E-86 | 1.02133946 | 0.913 | 0.46 | 7.05E-83 | Type7 | Cpne2 |
| 8.18E-85 | 2.57025118 | 0.948 | 0.595 | 1.64E-81 | Type7 | Atp1a2 |
| 1.06E-83 | 1.01340436 | 0.896 | 0.37 | 2.11E-80 | Type7 | Tcf7l2 |
| 1.03E-82 | 0.45530303 | 0.792 | 0.09 | 2.06E-79 | Type7 | Neu4 |
| 3.26E-80 | 0.64316736 | 0.884 | 0.246 | 6.51E-77 | Type7 | Daam2 |
| 1.30E-79 | 1.59857925 | 0.908 | 0.558 | 2.60E-76 | Type7 | Slc1a3 |
| 5.01E-79 | 1.00187647 | 0.902 | 0.576 | 1.00E-75 | Type7 | Slc7a10 |
| 2.38E-78 | 0.67062194 | 0.89 | 0.516 | 4.75E-75 | Type7 | S1pr1 |
| 4.72E-78 | 0.88307995 | 0.879 | 0.363 | 9.45E-75 | Type7 | Plxnb1 |
| 1.29E-77 | 0.7620826 | 0.89 | 0.533 | 2.58E-74 | Type7 | Pbxip1 |
| 1.89E-74 | 0.33884956 | 0.855 | 0.4 | 3.78E-71 | Type7 | Zic1 |
| 4.74E-74 | 0.8461452 | 0.908 | 0.433 | 9.48E-71 | Type7 | Slc6a1 |
| 4.33E-73 | 0.47008729 | 0.879 | 0.454 | 8.66E-70 | Type7 | Plce1 |
| 6.49E-72 | 0.63897929 | 0.832 | 0.315 | 1.30E-68 | Type7 | Slc13a4 |
| 8.71E-72 | 0.27156175 | 0.838 | 0.353 | 1.74E-68 | Type7 | Lef1 |
| 1.93E-71 | 0.65452245 | 0.861 | 0.386 | 3.85E-68 | Type7 | Cxcl14 |
| 5.93E-71 | 0.47272496 | 0.861 | 0.327 | 1.19E-67 | Type7 | Ccdc114 |
| 4.47E-70 | 0.30449466 | 0.844 | 0.332 | 8.94E-67 | Type7 | Zfhx3 |
| 2.27E-69 | 0.2976852 | 0.855 | 0.346 | 4.53E-66 | Type7 | Olig2 |
| 2.02E-68 | 0.3977784 | 0.884 | 0.367 | 4.04E-65 | Type7 | Slco1c1 |
| 6.35E-68 | 3.17606388 | 0.954 | 0.65 | 1.27E-64 | Type7 | Apoe |
| 1.61E-67 | 0.32810085 | 0.85 | 0.285 | 3.22E-64 | Type7 | Mertk |
| 2.08E-66 | 0.87714464 | 0.832 | 0.245 | 4.15E-63 | Type7 | Lrrtm3 |
| 1.81E-65 | 1.06791582 | 0.879 | 0.531 | 3.61E-62 | Type7 | Pdgfra |
| 2.65E-65 | 0.62109567 | 0.861 | 0.44 | 5.30E-62 | Type7 | Cspg4 |
| 2.92E-65 | 0.36499469 | 0.873 | 0.453 | 5.83E-62 | Type7 | Fgfr2 |
| 9.23E-65 | 0.5915431 | 0.884 | 0.429 | 1.85E-61 | Type7 | Cmtm5 |
| 1.83E-64 | 0.28291337 | 0.792 | 0.256 | 3.67E-61 | Type7 | Gpr17 |
| 1.27E-63 | 0.58879134 | 0.844 | 0.453 | 2.54E-60 | Type7 | Cacng4 |
| 2.53E-63 | 0.87006121 | 0.908 | 0.519 | 5.07E-60 | Type7 | Cd9 |

| | | | | | | |
|---|---|---|---|---|---|---|
| 7.96E-61 | 0.43735052 | 0.861 | 0.381 | 1.59E-57 | Type7 | Fnbp1l |
| 5.60E-60 | 0.5416742 | 0.827 | 0.371 | 1.12E-56 | Type7 | Kank1 |
| 5.61E-60 | 0.93938599 | 0.884 | 0.545 | 1.12E-56 | Type7 | Arhgap31 |
| 7.27E-60 | 0.8744645 | 0.867 | 0.467 | 1.45E-56 | Type7 | Epb41l2 |
| 2.75E-59 | 0.60293987 | 0.867 | 0.481 | 5.50E-56 | Type7 | Eya1 |
| 8.13E-58 | 0.82946374 | 0.855 | 0.463 | 1.63E-54 | Type7 | Ppfibp1 |
| 2.52E-57 | 0.6664158 | 0.844 | 0.378 | 5.04E-54 | Type7 | Sox8 |
| 2.69E-57 | 0.70950934 | 0.867 | 0.479 | 5.39E-54 | Type7 | Slc7a11 |
| 1.05E-56 | 1.5088852 | 0.931 | 0.578 | 2.11E-53 | Type7 | Plpp3 |
| 2.02E-56 | 1.4979516 | 0.902 | 0.555 | 4.03E-53 | Type7 | Phkg1 |
| 7.34E-56 | 0.45370396 | 0.855 | 0.339 | 1.47E-52 | Type7 | Sox2 |
| 4.07E-55 | 0.25218413 | 0.803 | 0.295 | 8.14E-52 | Type7 | Tshz1 |
| 7.30E-55 | 0.29158776 | 0.821 | 0.422 | 1.46E-51 | Type7 | Amotl1 |
| 5.92E-54 | 0.25859788 | 0.815 | 0.414 | 1.18E-50 | Type7 | Ttc28 |
| 6.58E-54 | 0.4565437 | 0.867 | 0.527 | 1.32E-50 | Type7 | Tns3 |
| 1.02E-53 | 0.96316418 | 0.919 | 0.466 | 2.04E-50 | Type7 | Utp14b |
| 2.37E-53 | 0.6812571 | 0.855 | 0.503 | 4.75E-50 | Type7 | Sox10 |
| 2.57E-53 | 0.33092346 | 0.838 | 0.43 | 5.15E-50 | Type7 | Slc38a3 |
| 7.81E-53 | 0.809619 | 0.896 | 0.531 | 1.56E-49 | Type7 | Tead1 |
| 8.56E-53 | 0.26760783 | 0.844 | 0.435 | 1.71E-49 | Type7 | Myoc |
| 3.49E-52 | 0.34277537 | 0.867 | 0.426 | 6.98E-49 | Type7 | Id4 |
| 9.37E-52 | 0.68539191 | 0.873 | 0.559 | 1.87E-48 | Type7 | Brinp3 |
| 1.33E-51 | 1.73522216 | 0.913 | 0.489 | 2.66E-48 | Type7 | Clu |
| 2.65E-51 | 0.79036802 | 0.89 | 0.6 | 5.29E-48 | Type7 | Itpr2 |
| 4.71E-51 | 0.52884094 | 0.844 | 0.469 | 9.42E-48 | Type7 | Hepacam |
| 5.07E-51 | 0.34351024 | 0.815 | 0.31 | 1.01E-47 | Type7 | Otx2 |
| 5.49E-50 | 1.07085921 | 0.827 | 0.415 | 1.10E-46 | Type7 | Bcas1 |
| 8.22E-49 | 0.37436343 | 0.832 | 0.327 | 1.64E-45 | Type7 | Ptchd4 |
| 1.99E-48 | 0.29528771 | 0.867 | 0.486 | 3.98E-45 | Type7 | Erbin |
| 8.41E-48 | 0.71827155 | 0.78 | 0.176 | 1.68E-44 | Type7 | Csgalnact1 |
| 5.10E-47 | 0.52452254 | 0.884 | 0.542 | 1.02E-43 | Type7 | Nckap5 |
| 1.11E-46 | 0.36184798 | 0.855 | 0.421 | 2.22E-43 | Type7 | Nxph1 |
| 2.82E-46 | 0.55105717 | 0.838 | 0.452 | 5.64E-43 | Type7 | Pcdh15 |
| 3.53E-46 | 0.34405152 | 0.792 | 0.238 | 7.07E-43 | Type7 | Etv4 |
| 4.99E-46 | 0.39792395 | 0.85 | 0.468 | 9.98E-43 | Type7 | Sorcs1 |
| 4.24E-45 | 0.28852167 | 0.798 | 0.421 | 8.49E-42 | Type7 | Tshz2 |
| 6.61E-45 | 0.68797768 | 0.838 | 0.448 | 1.32E-41 | Type7 | Cd63 |
| 3.07E-44 | 1.371977 | 0.855 | 0.472 | 6.14E-41 | Type7 | Gjc3 |
| 2.54E-43 | 0.67009378 | 0.908 | 0.524 | 5.07E-40 | Type7 | Chd7 |
| 4.45E-43 | 1.47507051 | 0.913 | 0.556 | 8.89E-40 | Type7 | Acsl3 |
| 5.81E-43 | 0.31954827 | 0.792 | 0.23 | 1.16E-39 | Type7 | Gm9925 |
| 1.01E-42 | 0.2532666 | 0.827 | 0.286 | 2.02E-39 | Type7 | Chn2 |
| 2.10E-41 | 0.26338855 | 0.855 | 0.441 | 4.21E-38 | Type7 | Csrp1 |

| | | | | | | |
|---|---|---|---|---|---|---|
| 5.44E-41 | 0.39619592 | 0.855 | 0.494 | 1.09E-37 | Type7 | Gm2a |
| 9.15E-41 | 0.67129358 | 0.884 | 0.605 | 1.83E-37 | Type7 | Cd27 |
| 2.02E-40 | 0.28030507 | 0.803 | 0.394 | 4.04E-37 | Type7 | Cit |
| 2.26E-40 | 1.66728235 | 0.884 | 0.634 | 4.51E-37 | Type7 | Enpp2 |
| 2.91E-40 | 0.55996992 | 0.884 | 0.58 | 5.81E-37 | Type7 | Klhl5 |
| 9.33E-40 | 0.44706371 | 0.728 | 0.282 | 1.87E-36 | Type7 | Id3 |
| 2.62E-39 | 1.58944178 | 0.954 | 0.783 | 5.23E-36 | Type7 | Ntm |
| 3.16E-39 | 0.48169535 | 0.873 | 0.379 | 6.31E-36 | Type7 | Serpine2 |
| 3.76E-38 | 0.3765194 | 0.705 | 0.238 | 7.51E-35 | Type7 | Igfbp2 |
| 3.92E-38 | 0.37856723 | 0.861 | 0.465 | 7.85E-35 | Type7 | Pik3r1 |
| 1.97E-37 | 0.79012836 | 0.89 | 0.638 | 3.95E-34 | Type7 | Pla2g7 |
| 4.72E-37 | 0.79465495 | 0.798 | 0.377 | 9.43E-34 | Type7 | Prlr |
| 6.82E-37 | 0.45104988 | 0.815 | 0.377 | 1.36E-33 | Type7 | Sccpdh |
| 9.80E-37 | 0.56547528 | 0.832 | 0.556 | 1.96E-33 | Type7 | Mpp7 |
| 1.79E-36 | 0.53305843 | 0.746 | 0.26 | 3.59E-33 | Type7 | Camk2d |
| 2.83E-36 | 0.85215662 | 0.873 | 0.439 | 5.66E-33 | Type7 | Gja1 |
| 3.29E-36 | 2.03659827 | 0.931 | 0.621 | 6.59E-33 | Type7 | Cspg5 |
| 5.49E-36 | 0.70935351 | 0.89 | 0.51 | 1.10E-32 | Type7 | Col9a3 |
| 7.66E-36 | 0.27643553 | 0.879 | 0.641 | 1.53E-32 | Type7 | Slc30a10 |
| 9.03E-36 | 0.30531069 | 0.838 | 0.479 | 1.81E-32 | Type7 | Vav3 |
| 2.56E-35 | 2.78746234 | 0.954 | 0.656 | 5.13E-32 | Type7 | Slc1a2 |
| 6.23E-35 | 0.38122032 | 0.827 | 0.414 | 1.25E-31 | Type7 | Fras1 |
| 6.60E-35 | 0.69669653 | 0.855 | 0.53 | 1.32E-31 | Type7 | Ptgds |
| 1.16E-34 | 0.25754291 | 0.861 | 0.509 | 2.33E-31 | Type7 | Cgnl1 |
| 1.58E-34 | 0.44778103 | 0.85 | 0.588 | 3.16E-31 | Type7 | Mtss1 |
| 1.20E-33 | 0.4482664 | 0.827 | 0.368 | 2.40E-30 | Type7 | Maml2 |
| 1.74E-33 | 0.25560312 | 0.85 | 0.566 | 3.47E-30 | Type7 | Gatm |
| 2.52E-33 | 0.27468932 | 0.815 | 0.463 | 5.04E-30 | Type7 | Pex11a |
| 3.42E-33 | 0.36157555 | 0.861 | 0.453 | 6.83E-30 | Type7 | Tmem72 |
| 3.83E-33 | 0.31916801 | 0.821 | 0.472 | 7.65E-30 | Type7 | Ptprt |
| 1.33E-32 | 0.56741443 | 0.85 | 0.497 | 2.65E-29 | Type7 | Stard9 |
| 3.05E-32 | 0.86069839 | 0.896 | 0.615 | 6.10E-29 | Type7 | Gpr37l1 |
| 5.35E-32 | 0.49490791 | 0.879 | 0.606 | 1.07E-28 | Type7 | Dock10 |
| 6.11E-32 | 0.56998996 | 0.844 | 0.539 | 1.22E-28 | Type7 | Edil3 |
| 2.48E-30 | 0.51180348 | 0.861 | 0.623 | 4.96E-27 | Type7 | Smad1 |
| 1.46E-29 | 0.84467409 | 0.902 | 0.58 | 2.91E-26 | Type7 | Msmo1 |
| 2.28E-29 | 1.02536875 | 0.792 | 0.389 | 4.56E-26 | Type7 | Bcl2 |
| 2.33E-29 | 1.36755677 | 0.879 | 0.596 | 4.67E-26 | Type7 | Mbp |
| 4.70E-29 | 0.40153356 | 0.832 | 0.506 | 9.40E-26 | Type7 | Matn4 |
| 1.50E-28 | 0.26557885 | 0.879 | 0.519 | 3.01E-25 | Type7 | Prkrip1 |
| 1.71E-28 | 0.95380606 | 0.809 | 0.429 | 3.41E-25 | Type7 | Grid2 |
| 1.88E-28 | 0.25384626 | 0.821 | 0.469 | 3.77E-25 | Type7 | Aldh1a1 |
| 2.28E-28 | 0.44459037 | 0.809 | 0.326 | 4.56E-25 | Type7 | Tex9 |

| | | | | | | |
|---|---|---|---|---|---|---|
| 4.08E-28 | 0.40691268 | 0.879 | 0.53 | 8.15E-25 | Type7 | Tceal3 |
| 9.15E-28 | 0.26234532 | 0.63 | 0.243 | 1.83E-24 | Type7 | Pomc |
| 2.80E-27 | 0.33532578 | 0.832 | 0.526 | 5.60E-24 | Type7 | Atp1b2 |
| 7.16E-27 | 0.53349605 | 0.861 | 0.513 | 1.43E-23 | Type7 | Zmiz1 |
| 1.32E-26 | 2.7307108 | 0.948 | 0.643 | 2.64E-23 | Type7 | Cst3 |
| 1.81E-26 | 0.65895036 | 0.821 | 0.481 | 3.61E-23 | Type7 | Ccdc82 |
| 1.97E-26 | 0.27755145 | 0.827 | 0.461 | 3.94E-23 | Type7 | Chdh |
| 2.16E-26 | 0.34511993 | 0.832 | 0.581 | 4.32E-23 | Type7 | Emx2 |
| 3.97E-26 | 1.47246755 | 0.925 | 0.535 | 7.94E-23 | Type7 | Sparcl1 |
| 1.21E-25 | 0.45540611 | 0.815 | 0.396 | 2.42E-22 | Type7 | Mylk |
| 1.57E-25 | 0.53184344 | 0.769 | 0.364 | 3.14E-22 | Type7 | Pdzd2 |
| 1.22E-24 | 0.35880338 | 0.873 | 0.526 | 2.43E-21 | Type7 | Pitpnc1 |
| 2.63E-24 | 1.25894228 | 0.913 | 0.497 | 5.27E-21 | Type7 | Msi2 |
| 6.42E-24 | 1.19091034 | 0.942 | 0.653 | 1.28E-20 | Type7 | Scd2 |
| 6.69E-23 | 0.82605692 | 0.85 | 0.478 | 1.34E-19 | Type7 | Sox4 |
| 8.32E-23 | 0.64919326 | 0.723 | 0.453 | 1.66E-19 | Type7 | Aqp4 |
| 2.27E-21 | 0.26081022 | 0.15 | 0.323 | 4.55E-18 | Type7 | mt-Nd5 |
| 4.95E-21 | 0.45287039 | 0.827 | 0.516 | 9.91E-18 | Type7 | Cnst |
| 7.64E-21 | 0.31261103 | 0.162 | 0.263 | 1.53E-17 | Type7 | Tac1 |
| 1.02E-20 | 0.34363205 | 0.798 | 0.5 | 2.03E-17 | Type7 | Ap1s2 |
| 2.39E-20 | 0.26320194 | 0.786 | 0.464 | 4.79E-17 | Type7 | Cgn |
| 2.87E-20 | 0.25768085 | 0.855 | 0.561 | 5.74E-17 | Type7 | Calcrl |
| 5.45E-20 | 0.37228888 | 0.711 | 0.425 | 1.09E-16 | Type7 | Sh3pxd2b |
| 5.95E-20 | 0.54077391 | 0.873 | 0.62 | 1.19E-16 | Type7 | Agl |
| 6.68E-20 | 0.54423855 | 0.85 | 0.551 | 1.34E-16 | Type7 | Olig1 |
| 7.26E-20 | 0.81444838 | 0.89 | 0.634 | 1.45E-16 | Type7 | Plp1 |
| 5.73E-19 | 0.61584537 | 0.803 | 0.425 | 1.15E-15 | Type7 | Marcks |
| 6.00E-19 | 0.28324728 | 0.757 | 0.37 | 1.20E-15 | Type7 | Unc13c |
| 1.31E-18 | 1.93998835 | 0.821 | 0.473 | 2.61E-15 | Type7 | Ptprz1 |
| 6.33E-18 | 0.56243809 | 0.85 | 0.511 | 1.27E-14 | Type7 | Pdia3 |
| 1.57E-17 | 0.34291107 | 0.15 | 0.389 | 3.14E-14 | Type7 | Satb1 |
| 3.59E-17 | 0.28017799 | 0.855 | 0.529 | 7.19E-14 | Type7 | Tmco3 |
| 5.80E-17 | 0.90625016 | 0.873 | 0.573 | 1.16E-13 | Type7 | Ncam1 |
| 6.79E-17 | 0.38815081 | 0.798 | 0.474 | 1.36E-13 | Type7 | Dbi |
| 1.20E-16 | 0.76707793 | 0.85 | 0.529 | 2.41E-13 | Type7 | Lima1 |
| 1.68E-16 | 0.38597378 | 0.832 | 0.584 | 3.36E-13 | Type7 | Adcyap1r1 |
| 6.99E-16 | 0.2982689 | 0.26 | 0.446 | 1.40E-12 | Type7 | mt-Nd2 |
| 1.09E-15 | 0.52617424 | 0.873 | 0.551 | 2.18E-12 | Type7 | Zfp462 |
| 1.96E-15 | 0.69817434 | 0.896 | 0.62 | 3.92E-12 | Type7 | Rps24 |
| 2.08E-15 | 0.4779431 | 0.855 | 0.582 | 4.16E-12 | Type7 | Minar2 |
| 2.77E-15 | 0.44540033 | 0.855 | 0.534 | 5.55E-12 | Type7 | Mios |
| 4.51E-15 | 0.29806911 | 0.682 | 0.353 | 9.03E-12 | Type7 | Ndst3 |
| 1.44E-14 | 0.44073894 | 0.578 | 0.239 | 2.88E-11 | Type7 | Pcdh11x |

| | | | | | | |
|---|---|---|---|---|---|---|
| 1.47E-14 | 0.69444552 | 0.884 | 0.574 | 2.94E-11 | Type7 | Smarca2 |
| 3.12E-14 | 0.52175717 | 0.902 | 0.654 | 6.23E-11 | Type7 | Shisa4 |
| 5.98E-14 | 0.97013389 | 0.879 | 0.587 | 1.20E-10 | Type7 | mt-Nd1 |
| 6.13E-14 | 0.88853844 | 0.393 | 0.566 | 1.23E-10 | Type7 | Htr2c |
| 1.22E-13 | 0.65471274 | 0.584 | 0.293 | 2.43E-10 | Type7 | Grin3a |
| 1.50E-13 | 0.28332713 | 0.266 | 0.512 | 2.99E-10 | Type7 | Trpm4 |
| 2.66E-13 | 0.60955369 | 0.694 | 0.439 | 5.32E-10 | Type7 | Btbd17 |
| 6.23E-13 | 0.32301149 | 0.85 | 0.624 | 1.25E-09 | Type7 | Ccdc86 |
| 7.13E-13 | 0.67011018 | 0.89 | 0.684 | 1.43E-09 | Type7 | Cuedc1 |
| 2.13E-12 | 0.46405547 | 0.636 | 0.287 | 4.26E-09 | Type7 | B4galt1 |
| 8.48E-12 | 0.26452393 | 0.78 | 0.473 | 1.70E-08 | Type7 | Nap1l5 |
| 1.88E-11 | 0.63467727 | 0.815 | 0.485 | 3.77E-08 | Type7 | Car2 |
| 2.09E-11 | 0.28383618 | 0.723 | 0.51 | 4.17E-08 | Type7 | Fxyd6 |
| 2.52E-11 | 1.19422504 | 0.908 | 0.563 | 5.04E-08 | Type7 | Ttyh1 |
| 2.07E-10 | 0.28169878 | 0.133 | 0.21 | 4.13E-07 | Type7 | Blnk |
| 2.54E-10 | 0.42437282 | 0.838 | 0.57 | 5.08E-07 | Type7 | Smc1a |
| 5.04E-10 | 0.58213729 | 0.283 | 0.358 | 1.01E-06 | Type7 | Ikzf2 |
| 5.26E-10 | 0.35906864 | 0.844 | 0.559 | 1.05E-06 | Type7 | Fam228b |
| 7.10E-10 | 0.271443 | 0.855 | 0.538 | 1.42E-06 | Type7 | Xylt1 |
| 8.35E-10 | 0.66798406 | 0.85 | 0.526 | 1.67E-06 | Type7 | Zeb1 |
| 2.60E-09 | 0.86141946 | 0.792 | 0.508 | 5.20E-06 | Type7 | Tspan7 |
| 3.38E-09 | 0.58082593 | 0.179 | 0.276 | 6.76E-06 | Type7 | Timp3 |
| 6.08E-09 | 0.30372888 | 0.329 | 0.491 | 1.22E-05 | Type7 | Tspan3 |
| 6.96E-09 | 0.60917105 | 0.919 | 0.677 | 1.39E-05 | Type7 | Trps1 |
| 7.12E-09 | 0.32980652 | 0.838 | 0.526 | 1.42E-05 | Type7 | Lrif1 |
| 7.16E-09 | 0.29195455 | 0.318 | 0.45 | 1.43E-05 | Type7 | Soat1 |
| 9.75E-09 | 0.37267492 | 0.867 | 0.683 | 1.95E-05 | Type7 | Map1s |
| 1.29E-08 | 0.49861685 | 0.225 | 0.193 | 2.57E-05 | Type7 | Arhgap6 |
| 4.81E-08 | 0.77468309 | 0.913 | 0.701 | 9.63E-05 | Type7 | Rrbp1 |
| 4.96E-08 | 0.39122638 | 0.387 | 0.482 | 9.91E-05 | Type7 | Cyp2d22 |
| 5.04E-08 | 0.27107435 | 0.734 | 0.443 | 0.00010076 | Type7 | Zfp945 |
| 5.81E-08 | 0.44297441 | 0.89 | 0.649 | 0.00011611 | Type7 | Nnat |
| 6.35E-08 | 3.1284395 | 0.913 | 0.789 | 0.00012702 | Type7 | Ttr |
| 9.31E-08 | 0.48249939 | 0.197 | 0.277 | 0.00018613 | Type7 | Syn3 |
| 3.49E-07 | 0.3193016 | 0.168 | 0.232 | 0.00069839 | Type7 | Kitl |
| 1.09E-06 | 0.35660709 | 0.89 | 0.681 | 0.00218272 | Type7 | Tsc22d4 |
| 1.22E-06 | 0.46818781 | 0.792 | 0.55 | 0.00244337 | Type7 | 0610012G03 |
| 1.34E-06 | 0.26169181 | 0.11 | 0.257 | 0.00267437 | Type7 | Slc8a3 |
| 1.53E-06 | 0.28019149 | 0.277 | 0.235 | 0.00305621 | Type7 | Pim1 |
| 1.64E-06 | 0.63747008 | 0.902 | 0.68 | 0.00327773 | Type7 | Med13 |
| 2.03E-06 | 0.7292372 | 0.59 | 0.412 | 0.00406833 | Type7 | Gpc5 |
| 2.89E-06 | 0.37487533 | 0.538 | 0.735 | 0.00577893 | Type7 | Tulp4 |
| 2.94E-06 | 0.43938271 | 0.896 | 0.644 | 0.00588018 | Type7 | Zfp106 |

| | | | | | | | |
|---|---|---|---|---|---|---|---|
| 3.10E-06 | 0.31217949 | 0.78 | 0.524 | 0.00619407 | Type7 | | Reep3 |
| 3.47E-06 | 0.42360857 | 0.844 | 0.557 | 0.00693585 | Type7 | | Ppp2r3a |
| 3.91E-06 | 0.44172728 | 0.908 | 0.729 | 0.00782018 | Type7 | | Cdh10 |
| 4.47E-06 | 0.85666782 | 0.555 | 0.338 | 0.00893334 | Type7 | | Asap2 |
| 5.20E-06 | 0.45365711 | 0.202 | 0.225 | 0.010399 | Type7 | | Cntn3 |
| 6.51E-06 | 0.55066764 | 0.566 | 0.322 | 0.01301371 | Type7 | | Pik3ip1 |
| 9.94E-06 | 0.35908374 | 0.884 | 0.707 | 0.01987693 | Type7 | | Bsg |
| 1.00E-05 | 0.53299198 | 0.873 | 0.691 | 0.02000141 | Type7 | | Sntg1 |
| 1.38E-05 | 0.27803473 | 0.295 | 0.375 | 0.0275439 | Type7 | | Cmya5 |
| 1.64E-05 | 0.33820023 | 0.306 | 0.508 | 0.03271161 | Type7 | | Nptxr |
| 1.71E-05 | 0.2876804 | 0.272 | 0.349 | 0.03427121 | Type7 | | Atoh7 |
| 1.85E-05 | 0.3023695 | 0.509 | 0.286 | 0.03690145 | Type7 | | Stxbp3 |
| 1.85E-05 | 0.32466198 | 0.861 | 0.624 | 0.03691691 | Type7 | | Ccdc34 |
| 1.90E-05 | 0.28687459 | 0.803 | 0.557 | 0.03806864 | Type7 | | Amer1 |
| 2.08E-05 | 0.46931662 | 0.769 | 0.448 | 0.0415762 | Type7 | | Ecrg4 |
| 2.28E-05 | 0.34268143 | 0.855 | 0.593 | 0.04569035 | Type7 | | D3Ertd751e |
| 2.29E-05 | 0.67354176 | 0.335 | 0.46 | 0.0458009 | Type7 | | Sat1 |
| 2.33E-05 | 0.42299212 | 0.434 | 0.477 | 0.04663107 | Type7 | | Slc4a2 |
| 2.38E-05 | 0.52217775 | 0.191 | 0.309 | 0.0475481 | Type7 | | Inhba |
| 2.40E-05 | 0.41361183 | 0.382 | 0.409 | 0.04807718 | Type7 | | Cdh13 |
| 2.46E-05 | 1.66954168 | 0.78 | 0.649 | 0.04910092 | Type7 | | Qk |
| 2.88E-05 | 0.59806487 | 0.214 | 0.322 | 0.05764431 | Type7 | | Igsf3 |
| 3.01E-05 | 0.67548348 | 0.503 | 0.309 | 0.06017649 | Type7 | | Vangl2 |
| 5.82E-05 | 0.31925577 | 0.688 | 0.472 | 0.11640699 | Type7 | | Dph5 |
| 7.09E-05 | 0.63696062 | 0.387 | 0.514 | 0.14185528 | Type7 | | Slc4a4 |
| 8.28E-05 | 0.25310843 | 0.647 | 0.54 | 0.16566105 | Type7 | | Camk1 |
| 8.54E-05 | 0.51566058 | 0.879 | 0.642 | 0.17083178 | Type7 | | Pggt1b |
| 8.80E-05 | 0.30559199 | 0.89 | 0.798 | 0.17609707 | Type7 | | Luzp2 |
| 0.00011002 | 0.40083602 | 0.451 | 0.683 | 0.22003916 | Type7 | | Sec62 |
| 0.00011805 | 0.60976888 | 0.277 | 0.178 | 0.23609493 | Type7 | | Zfp536 |
| 0.00014283 | 0.32128416 | 0.838 | 0.649 | 0.28565701 | Type7 | | Mttp |
| 0.00017289 | 0.45200499 | 0.225 | 0.327 | 0.34578983 | Type7 | | Sema6d |
| 0.00019417 | 0.36749612 | 0.659 | 0.449 | 0.38834691 | Type7 | | Cilk1 |
| 0.00019542 | 0.25782331 | 0.769 | 0.543 | 0.39084129 | Type7 | | Zkscan1 |
| 0.00020134 | 0.39859459 | 0.474 | 0.692 | 0.40267738 | Type7 | | mt-Co1 |
| 0.00020532 | 0.43558479 | 0.827 | 0.551 | 0.41063207 | Type7 | | Ctnnal1 |
| 0.00023104 | 0.3803735 | 0.751 | 0.576 | 0.46207882 | Type7 | | Spcs2 |
| 0.00030862 | 0.64213363 | 0.844 | 0.64 | 0.61723451 | Type7 | | Rplp1 |
| 0.0003835 | 0.46090125 | 0.902 | 0.739 | 0.76700968 | Type7 | | Gkap1 |
| 0.00051306 | 0.47199293 | 0.879 | 0.652 | 1 | Type7 | | Fau |
| 0.00067406 | 0.50044979 | 0.63 | 0.449 | 1 | Type7 | | Jun |
| 0.00067621 | 0.71552801 | 0.925 | 0.711 | 1 | Type7 | | Arhgap5 |
| 0.00072594 | 0.27281391 | 0.416 | 0.541 | 1 | Type7 | | Hsp90b1 |

| | | | | | | |
|---|---|---|---|---|---|---|
| 0.0010554 | 0.35107267 | 0.734 | 0.506 | 1 | Type7 | Nucks1 |
| 0.00120895 | 0.56264335 | 0.41 | 0.545 | 1 | Type7 | mt-Nd4 |
| 0.00121149 | 0.75313797 | 0.324 | 0.246 | 1 | Type7 | Adgrl2 |
| 0.00138359 | 0.37356673 | 0.879 | 0.708 | 1 | Type7 | Mgll |
| 0.00164342 | 0.36786799 | 0.445 | 0.57 | 1 | Type7 | Dock3 |
| 0.00173734 | 0.31181239 | 0.462 | 0.664 | 1 | Type7 | Dgkh |
| 0.00179231 | 0.49192249 | 0.954 | 0.978 | 1 | Type7 | Actb |
| 0.00192885 | 0.254762 | 0.543 | 0.585 | 1 | Type7 | Emc7 |
| 0.00193821 | 0.84505535 | 0.786 | 0.65 | 1 | Type7 | mt-Co2 |
| 0.00217742 | 0.49060062 | 0.387 | 0.235 | 1 | Type7 | Samd4 |
| 0.00236035 | 0.3124773 | 0.566 | 0.785 | 1 | Type7 | Phip |
| 0.00264479 | 0.3077135 | 0.301 | 0.428 | 1 | Type7 | Kcnb2 |
| 0.00300891 | 0.49315558 | 0.647 | 0.435 | 1 | Type7 | Pou3f3 |
| 0.00359241 | 0.28021982 | 0.584 | 0.77 | 1 | Type7 | Ntsr2 |
| 0.00412643 | 0.30214983 | 0.89 | 0.923 | 1 | Type7 | Mef2a |
| 0.00435031 | 0.358439 | 0.728 | 0.723 | 1 | Type7 | Ptpre |
| 0.00450658 | 0.32238747 | 0.838 | 0.647 | 1 | Type7 | Sowaha |
| 0.00477827 | 0.44856819 | 0.855 | 0.722 | 1 | Type7 | Tgs1 |
| 0.00478172 | 0.57507435 | 0.838 | 0.646 | 1 | Type7 | Fut9 |
| 0.00492339 | 0.69479067 | 0.399 | 0.441 | 1 | Type7 | Cald1 |
| 0.00493375 | 0.29910627 | 0.734 | 0.497 | 1 | Type7 | Brwd3 |
| 0.005319 | 0.32007515 | 0.902 | 0.951 | 1 | Type7 | Tmem108 |
| 0.00539049 | 0.41389858 | 0.543 | 0.365 | 1 | Type7 | Hlf |
| 0.00706235 | 0.52911295 | 0.913 | 0.809 | 1 | Type7 | Pid1 |
| 0.00706501 | 0.62598475 | 0.919 | 0.83 | 1 | Type7 | Dnm3 |
| 0.00787029 | 0.64821589 | 0.803 | 0.61 | 1 | Type7 | Tnr |
| 0.00910238 | 0.51943517 | 0.879 | 0.779 | 1 | Type7 | Pdcd4 |
| 0.00947484 | 0.27894084 | 0.532 | 0.73 | 1 | Type7 | Klf9 |

Rik

'Rik

Rik

Rik

Rik

Rik

Rik

Rik
