## Supplemental table4 for "Investigation of the Fasciola Cinereum, Absent in BTBR mice, and Comparison with the Hippocampal Area CA2"

| | p_val | avg_log2FC | pct.1 | pct.2 | p_val_adj |
|---|---|---|---|---|---|
| Gm8797 | 7.27E-73 | 4.67696716 | 0.312 | 0.013 | 1.33E-68 |
| Cfap54 | 1.20E-60 | 2.82046635 | 0.46 | 0.146 | 2.19E-56 |
| Epb41l5 | 5.32E-59 | 3.0951617 | 0.347 | 0.056 | 9.74E-55 |
| Sav1 | 2.65E-48 | 1.85520088 | 0.364 | 0.087 | 4.86E-44 |
| Scg5 | 2.75E-43 | 1.11753903 | 0.764 | 0.491 | 5.04E-39 |
| Lpl | 1.48E-41 | 2.21277528 | 0.349 | 0.096 | 2.71E-37 |
| Mt3 | 1.84E-34 | -2.1320152 | 0.16 | 0.429 | 3.38E-30 |
| Cntn4 | 1.42E-27 | 0.71841177 | 0.477 | 0.218 | 2.60E-23 |
| Galnt1 | 2.70E-25 | -1.6008373 | 0.236 | 0.435 | 4.94E-21 |
| Timp4 | 2.46E-23 | 1.12914629 | 0.48 | 0.275 | 4.52E-19 |
| Swt1 | 1.86E-22 | 0.98730319 | 0.427 | 0.229 | 3.42E-18 |
| Nudt4 | 7.62E-21 | 1.17919843 | 0.259 | 0.095 | 1.40E-16 |
| Hacd4 | 2.52E-20 | 1.26360148 | 0.456 | 0.274 | 4.61E-16 |
| Atp2c1 | 2.89E-20 | 0.6262742 | 0.719 | 0.558 | 5.30E-16 |
| Gm4924 | 7.55E-20 | 1.33129053 | 0.267 | 0.107 | 1.38E-15 |
| Fam57b | 4.55E-18 | 1.31522243 | 0.337 | 0.176 | 8.34E-14 |
| Kcnh5 | 3.15E-16 | 1.35793857 | 0.289 | 0.139 | 5.77E-12 |
| Cpne4 | 1.19E-15 | 0.67756864 | 0.677 | 0.534 | 2.19E-11 |
| Spink8 | 1.60E-15 | -1.6137101 | 0.1 | 0.25 | 2.94E-11 |
| Wbp11 | 3.90E-15 | 0.70936534 | 0.327 | 0.169 | 7.14E-11 |
| Ttc37 | 5.03E-15 | 0.92294877 | 0.435 | 0.279 | 9.22E-11 |
| Zkscan1 | 1.04E-14 | 0.8516295 | 0.386 | 0.237 | 1.90E-10 |
| Pcdh15 | 1.08E-14 | 1.41863969 | 0.306 | 0.158 | 1.98E-10 |
| Fbxo42 | 3.46E-14 | 0.86244319 | 0.326 | 0.185 | 6.33E-10 |
| Alcam | 1.03E-13 | 0.91026824 | 0.42 | 0.256 | 1.88E-09 |
| Prkn | 2.23E-13 | -0.8288782 | 0.608 | 0.626 | 4.09E-09 |
| Rab11fip2 | 3.03E-13 | 0.50160879 | 0.539 | 0.377 | 5.56E-09 |
| Snhg11 | 3.97E-13 | -0.3876708 | 0.831 | 0.88 | 7.28E-09 |
| Map2 | 4.15E-13 | 0.3113833 | 0.872 | 0.767 | 7.60E-09 |
| Epha7 | 4.53E-13 | 0.39662008 | 0.835 | 0.743 | 8.31E-09 |
| Plp1 | 4.84E-13 | -1.1290792 | 0.483 | 0.628 | 8.87E-09 |
| Marchf1 | 6.26E-13 | -0.6934409 | 0.563 | 0.693 | 1.15E-08 |
| Mllt3 | 9.46E-13 | 0.45995857 | 0.712 | 0.55 | 1.73E-08 |
| Farp1 | 9.79E-13 | 0.89493094 | 0.398 | 0.255 | 1.80E-08 |
| Cep19 | 2.18E-12 | 0.62018534 | 0.312 | 0.18 | 4.00E-08 |
| Plcxd3 | 2.80E-12 | 1.16525984 | 0.379 | 0.251 | 5.13E-08 |
| Kcnip4 | 2.90E-12 | 0.23225418 | 0.98 | 0.978 | 5.32E-08 |
| Fam98a | 3.08E-12 | 1.29579187 | 0.25 | 0.131 | 5.65E-08 |
| Zdhhc2 | 3.42E-12 | 0.71908258 | 0.394 | 0.255 | 6.28E-08 |
| Slco3a1 | 5.34E-12 | 0.48474024 | 0.57 | 0.399 | 9.79E-08 |
| Tspan5 | 7.76E-12 | 0.36904766 | 0.756 | 0.613 | 1.42E-07 |
| Unc5d | 1.53E-11 | 1.17255021 | 0.31 | 0.185 | 2.81E-07 |

| | | | | | |
|---|---|---|---|---|---|
| Myo1b | 1.91E-11 | 1.07802872 | 0.256 | 0.139 | 3.50E-07 |
| Pla2g3 | 2.64E-11 | 0.63274517 | 0.483 | 0.33 | 4.85E-07 |
| Sidt1 | 3.16E-11 | 0.49288423 | 0.589 | 0.435 | 5.78E-07 |
| Cox7a2 | 9.63E-11 | 0.40227742 | 0.578 | 0.41 | 1.76E-06 |
| Ccdc3 | 9.69E-11 | 1.14138333 | 0.336 | 0.216 | 1.78E-06 |
| Cdk17 | 1.60E-10 | 0.40393373 | 0.544 | 0.399 | 2.94E-06 |
| Myo6 | 2.26E-10 | 0.58271937 | 0.475 | 0.343 | 4.14E-06 |
| Atp5o | 2.28E-10 | 0.42614713 | 0.632 | 0.472 | 4.18E-06 |
| Drg1 | 2.67E-10 | 0.4744847 | 0.578 | 0.434 | 4.89E-06 |
| Cmss1 | 2.89E-10 | -0.4833393 | 0.95 | 0.946 | 5.30E-06 |
| Ppp1r12b | 3.01E-10 | 0.29937029 | 0.623 | 0.47 | 5.51E-06 |
| Eif1b | 3.11E-10 | 0.5493894 | 0.529 | 0.384 | 5.69E-06 |
| Kcnip3 | 3.11E-10 | -1.3589933 | 0.131 | 0.25 | 5.69E-06 |
| Ssbp4 | 4.14E-10 | 0.555274 | 0.464 | 0.329 | 7.58E-06 |
| Rmnd1 | 4.37E-10 | 0.59505611 | 0.323 | 0.205 | 8.01E-06 |
| Tmem131l | 4.91E-10 | 0.60507806 | 0.337 | 0.218 | 9.00E-06 |
| Fstl5 | 5.13E-10 | 0.70223546 | 0.494 | 0.376 | 9.41E-06 |
| Rpl13 | 1.21E-09 | 0.2727438 | 0.76 | 0.585 | 2.22E-05 |
| Srp14 | 1.50E-09 | 0.61925905 | 0.39 | 0.262 | 2.76E-05 |
| Bend6 | 1.62E-09 | 0.76132189 | 0.341 | 0.224 | 2.98E-05 |
| Frmd4a | 1.77E-09 | 0.29096723 | 0.838 | 0.765 | 3.25E-05 |
| Stau2 | 1.94E-09 | 0.35236027 | 0.63 | 0.487 | 3.56E-05 |
| Fndc9 | 2.19E-09 | 0.39064625 | 0.584 | 0.415 | 4.02E-05 |
| Hivep2 | 2.69E-09 | -0.4335467 | 0.754 | 0.751 | 4.94E-05 |
| Pced1b | 3.55E-09 | 0.83296777 | 0.359 | 0.247 | 6.51E-05 |
| Uqcrc2 | 3.92E-09 | 0.53610114 | 0.398 | 0.272 | 7.18E-05 |
| Dcc | 4.41E-09 | -0.4138873 | 0.767 | 0.806 | 8.09E-05 |
| Prune2 | 4.42E-09 | 0.930377 | 0.255 | 0.149 | 8.10E-05 |
| Adgrb3 | 5.21E-09 | -0.3618869 | 0.825 | 0.861 | 9.56E-05 |
| Me1 | 5.74E-09 | 0.77215075 | 0.341 | 0.232 | 0.00010516 |
| Rb1 | 7.78E-09 | 0.36301287 | 0.499 | 0.386 | 0.00014264 |
| Ppfia2 | 7.91E-09 | -0.3220884 | 0.855 | 0.861 | 0.00014496 |
| Brinp2 | 9.00E-09 | 0.54137007 | 0.561 | 0.438 | 0.00016501 |
| Slc24a5 | 9.46E-09 | 0.42650488 | 0.603 | 0.46 | 0.00017333 |
| Mrtfb | 1.14E-08 | 0.35395749 | 0.712 | 0.558 | 0.00020839 |
| Son | 1.16E-08 | 0.34603054 | 0.767 | 0.627 | 0.00021327 |
| Exoc6 | 1.81E-08 | 0.58772655 | 0.4 | 0.285 | 0.00033241 |
| Rae1 | 1.96E-08 | 0.53093802 | 0.329 | 0.218 | 0.00035984 |
| Ttc8 | 2.11E-08 | -1.1641639 | 0.229 | 0.345 | 0.00038702 |
| Focad | 2.22E-08 | 0.38654805 | 0.667 | 0.523 | 0.00040697 |
| Gpr18 | 2.71E-08 | 0.77227803 | 0.321 | 0.22 | 0.00049751 |
| Ndst1 | 2.75E-08 | 0.65163766 | 0.381 | 0.275 | 0.00050446 |
| Aven | 3.02E-08 | 0.52820221 | 0.286 | 0.184 | 0.00055327 |

| | | | | | |
|---|---|---|---|---|---|
| Luc7l2 | 3.26E-08 | 0.23992127 | 0.819 | 0.711 | 0.00059714 |
| Thsd4 | 3.39E-08 | -1.048609 | 0.285 | 0.378 | 0.00062125 |
| Unc80 | 3.39E-08 | 0.26723772 | 0.746 | 0.637 | 0.00062172 |
| Lingo2 | 3.55E-08 | -0.5287546 | 0.598 | 0.695 | 0.00065008 |
| Dtd1 | 3.56E-08 | 0.3160602 | 0.449 | 0.323 | 0.00065174 |
| Sntg2 | 3.75E-08 | 0.87925977 | 0.267 | 0.17 | 0.00068695 |
| Ppm1l | 3.81E-08 | 0.41451735 | 0.584 | 0.457 | 0.00069871 |
| Atp9a | 3.91E-08 | 0.40565156 | 0.533 | 0.397 | 0.00071737 |
| Aff3 | 5.13E-08 | 0.37113209 | 0.645 | 0.504 | 0.00094049 |
| Rnft1 | 5.52E-08 | 0.87579523 | 0.311 | 0.21 | 0.00101189 |
| Galnt2l | 6.01E-08 | 0.27533181 | 0.824 | 0.754 | 0.00110113 |
| Mast4 | 6.66E-08 | 0.33352176 | 0.701 | 0.553 | 0.00122021 |
| Ncapd3 | 7.45E-08 | 0.54424023 | 0.368 | 0.262 | 0.00136538 |
| Gon4l | 7.51E-08 | 0.43516794 | 0.446 | 0.334 | 0.00137747 |
| Dapk1 | 7.53E-08 | 0.3165851 | 0.686 | 0.557 | 0.00137989 |
| Rora | 8.52E-08 | 0.37968753 | 0.741 | 0.61 | 0.00156189 |
| Rnf115 | 9.15E-08 | 0.46262408 | 0.442 | 0.323 | 0.00167647 |
| Rabgef1 | 9.25E-08 | 0.82605144 | 0.25 | 0.158 | 0.00169595 |
| Ppp2r3a | 1.00E-07 | 0.41624568 | 0.539 | 0.42 | 0.00184078 |
| Grb2 | 1.01E-07 | 0.56851137 | 0.39 | 0.284 | 0.00184877 |
| 4932438A13I | 1.10E-07 | 0.39972257 | 0.578 | 0.466 | 0.00201981 |
| Mbp | 1.15E-07 | -1.215934 | 0.244 | 0.347 | 0.00210106 |
| Tafa5 | 1.41E-07 | 0.36244758 | 0.625 | 0.495 | 0.00258346 |
| Actb | 1.47E-07 | -0.5324446 | 0.832 | 0.871 | 0.0026917 |
| Ppib | 1.53E-07 | 0.25370474 | 0.527 | 0.41 | 0.00280126 |
| Galntl6 | 1.63E-07 | -0.688709 | 0.438 | 0.552 | 0.00299256 |
| Lrtm1 | 1.68E-07 | 0.49594512 | 0.495 | 0.373 | 0.0030827 |
| Gria2 | 1.76E-07 | 0.16929127 | 0.98 | 0.944 | 0.00322513 |
| Tmem117 | 2.11E-07 | 0.51152072 | 0.435 | 0.323 | 0.0038603 |
| Sfi1 | 2.14E-07 | 0.481353 | 0.308 | 0.204 | 0.00392401 |
| Odf2 | 2.39E-07 | 0.40650416 | 0.516 | 0.404 | 0.00437212 |
| Ndrg4 | 2.51E-07 | 0.39011638 | 0.694 | 0.558 | 0.00459641 |
| Lpp | 2.58E-07 | 0.3482359 | 0.568 | 0.437 | 0.00472583 |
| Cntnap5b | 2.65E-07 | 0.64808705 | 0.506 | 0.395 | 0.00485119 |
| Ints10 | 2.86E-07 | 0.45000213 | 0.387 | 0.286 | 0.00524115 |
| Luzp1 | 2.99E-07 | 0.35142679 | 0.55 | 0.415 | 0.00547527 |
| Gabarapl2 | 3.40E-07 | -0.8179517 | 0.196 | 0.307 | 0.00622969 |
| Ggact | 3.42E-07 | 0.45058125 | 0.48 | 0.359 | 0.00627151 |
| Hunk | 3.48E-07 | 0.43214332 | 0.419 | 0.304 | 0.00638148 |
| Tmem132b | 3.56E-07 | 0.2435618 | 0.649 | 0.509 | 0.00653457 |
| Angpt1 | 3.92E-07 | 0.40403979 | 0.265 | 0.167 | 0.00718002 |
| Gabbr1 | 4.24E-07 | 0.25992829 | 0.595 | 0.475 | 0.00776788 |
| Dnm1l | 4.27E-07 | 0.30972301 | 0.518 | 0.399 | 0.0078302 |

| | | | | | |
|---|---|---|---|---|---|
| Adcy2 | 4.39E-07 | -0.5452718 | 0.415 | 0.531 | 0.00805071 |
| Itsn1 | 4.45E-07 | 0.2671875 | 0.576 | 0.443 | 0.00815407 |
| Ice1 | 4.49E-07 | 0.43909526 | 0.431 | 0.322 | 0.00823312 |
| Dpf3 | 4.54E-07 | 0.30700711 | 0.415 | 0.299 | 0.00831498 |
| Kirrel3 | 4.64E-07 | -1.0124324 | 0.33 | 0.407 | 0.0085119 |
| Slc17a7 | 4.87E-07 | 0.24068529 | 0.786 | 0.607 | 0.008935 |
| Tmem63b | 5.10E-07 | 0.51134738 | 0.447 | 0.333 | 0.00934361 |
| Tmtc1 | 5.70E-07 | 0.29460791 | 0.458 | 0.34 | 0.01045243 |
| Slmap | 5.78E-07 | 0.55678967 | 0.454 | 0.343 | 0.01059244 |
| Clmn | 6.03E-07 | 0.33621381 | 0.471 | 0.353 | 0.01104445 |
| Rbm26 | 6.11E-07 | 0.29981201 | 0.652 | 0.531 | 0.01120865 |
| Sash1 | 6.13E-07 | 0.38220523 | 0.394 | 0.301 | 0.01122942 |
| Srsf2 | 6.21E-07 | -0.9053933 | 0.256 | 0.361 | 0.01139142 |
| Slc6a6 | 6.29E-07 | 0.29591991 | 0.563 | 0.439 | 0.01152198 |
| Rab30 | 6.92E-07 | 0.44136689 | 0.422 | 0.326 | 0.01268719 |
| Bbs7 | 7.36E-07 | 0.57185311 | 0.251 | 0.164 | 0.01348647 |
| Gpr155 | 7.90E-07 | 0.30085496 | 0.393 | 0.286 | 0.0144888 |
| Gpc5 | 8.00E-07 | 0.93631276 | 0.434 | 0.326 | 0.01466576 |
| Epha3 | 8.35E-07 | 0.23073631 | 0.638 | 0.504 | 0.01530684 |
| Prkca | 8.82E-07 | 0.26956751 | 0.791 | 0.67 | 0.01617162 |
| 4430402I18R | 8.85E-07 | 0.45651955 | 0.454 | 0.339 | 0.01623154 |
| Nt5c2 | 9.06E-07 | 0.31486132 | 0.492 | 0.375 | 0.01660099 |
| Crtac1 | 9.29E-07 | -0.7912175 | 0.16 | 0.26 | 0.01702883 |
| Parp8 | 9.38E-07 | -0.6525915 | 0.254 | 0.371 | 0.0171946 |
| Strbp | 9.86E-07 | 0.24710818 | 0.808 | 0.72 | 0.01807832 |
| Cdkl5 | 9.95E-07 | 0.36264442 | 0.558 | 0.434 | 0.01823585 |
| Zzz3 | 1.03E-06 | 0.54757732 | 0.356 | 0.26 | 0.0188773 |
| Slc7a8 | 1.06E-06 | 0.40473388 | 0.338 | 0.244 | 0.01952232 |
| Tfg | 1.11E-06 | 0.83194399 | 0.263 | 0.179 | 0.02030302 |
| Stmn3 | 1.11E-06 | 0.21907296 | 0.662 | 0.531 | 0.0203823 |
| Lrrfip1 | 1.13E-06 | 0.29851594 | 0.573 | 0.442 | 0.02068861 |
| Ano3 | 1.13E-06 | 0.32541198 | 0.696 | 0.576 | 0.02076822 |
| mt-Co1 | 1.14E-06 | -0.6732639 | 0.784 | 0.788 | 0.0208786 |
| Gpcpd1 | 1.22E-06 | 0.31770197 | 0.492 | 0.379 | 0.02241575 |
| Gm37240 | 1.31E-06 | 0.30252567 | 0.607 | 0.489 | 0.02398116 |
| Pde4d | 1.39E-06 | 0.31779302 | 0.804 | 0.737 | 0.02545598 |
| Stxbp6 | 1.42E-06 | -0.9442049 | 0.241 | 0.33 | 0.02594078 |
| Il1rapl1 | 1.42E-06 | 0.35021565 | 0.846 | 0.766 | 0.02599379 |
| Angpt2 | 1.48E-06 | 0.62116497 | 0.321 | 0.234 | 0.02708965 |
| Ago3 | 1.62E-06 | 0.32887641 | 0.551 | 0.43 | 0.02975606 |
| Carmil1 | 1.65E-06 | 0.33454789 | 0.396 | 0.294 | 0.03029348 |
| Ube2e1 | 1.75E-06 | 0.48051906 | 0.353 | 0.258 | 0.03206604 |
| Slc44a2 | 2.00E-06 | 0.65618812 | 0.255 | 0.171 | 0.03658922 |

| | | | | | |
|---|---|---|---|---|---|
| D130043K22 | 2.07E-06 | 0.31694893 | 0.383 | 0.286 | 0.03791631 |
| Igf2bp3 | 2.14E-06 | 0.33064557 | 0.482 | 0.367 | 0.03913882 |
| Cnot4 | 2.19E-06 | 0.24866777 | 0.634 | 0.511 | 0.04018102 |
| Rmdn1 | 2.29E-06 | 0.39068982 | 0.543 | 0.429 | 0.04202083 |
| Chgb | 2.35E-06 | 0.25158561 | 0.761 | 0.637 | 0.04316828 |
| Nrg1 | 2.47E-06 | -0.4767846 | 0.368 | 0.473 | 0.04518741 |
| Atp5g1 | 2.49E-06 | 0.26594826 | 0.523 | 0.402 | 0.04573492 |
| Stx16 | 2.63E-06 | 0.38682987 | 0.435 | 0.336 | 0.04822726 |
| Xrcc6 | 2.70E-06 | 0.34296926 | 0.497 | 0.374 | 0.04957661 |
| Brd9 | 2.72E-06 | 0.1454148 | 0.679 | 0.526 | 0.04992658 |
| Selenow | 2.88E-06 | -0.6680626 | 0.531 | 0.595 | 0.0527534 |
| Itfg1 | 2.92E-06 | 0.31240152 | 0.48 | 0.372 | 0.05360201 |
| Cacna1c | 2.94E-06 | -0.3173484 | 0.806 | 0.787 | 0.05386214 |
| Cacna1a | 3.04E-06 | -0.4274946 | 0.772 | 0.737 | 0.05572746 |
| Pcdh7 | 3.05E-06 | 0.37879339 | 0.548 | 0.44 | 0.05582276 |
| Rufy3 | 3.07E-06 | 0.30638028 | 0.622 | 0.502 | 0.05625375 |
| Abl2 | 3.09E-06 | 0.28835939 | 0.569 | 0.452 | 0.05665205 |
| Pcsk1n | 3.11E-06 | 0.31086693 | 0.483 | 0.369 | 0.0570925 |
| Amph | 3.16E-06 | -0.5790173 | 0.6 | 0.584 | 0.05797014 |
| Kbtbd11 | 3.21E-06 | 0.59684054 | 0.338 | 0.243 | 0.05876119 |
| Pygb | 3.25E-06 | 0.43904316 | 0.345 | 0.253 | 0.05952597 |
| Rgs11 | 3.27E-06 | 0.56652357 | 0.336 | 0.246 | 0.05997364 |
| Rnps1 | 3.29E-06 | -0.6437934 | 0.192 | 0.296 | 0.06021778 |
| Col4a2 | 3.29E-06 | 0.46370889 | 0.377 | 0.285 | 0.060306 |
| Kif1c | 3.33E-06 | 0.79001193 | 0.312 | 0.231 | 0.06107717 |
| Nup93 | 3.38E-06 | 0.24312889 | 0.48 | 0.361 | 0.06200086 |
| Prdm2 | 3.50E-06 | 0.48766817 | 0.312 | 0.224 | 0.06409796 |
| P2ry14 | 3.58E-06 | 0.36063837 | 0.565 | 0.445 | 0.0655393 |
| Fbxw7 | 3.74E-06 | 0.23275072 | 0.528 | 0.406 | 0.06848973 |
| Ggt7 | 3.80E-06 | 0.40050349 | 0.42 | 0.312 | 0.06970165 |
| Stxbp1 | 3.82E-06 | 0.25513598 | 0.569 | 0.45 | 0.07008417 |
| Afg3l2 | 3.85E-06 | 0.60264356 | 0.263 | 0.182 | 0.07062412 |
| Copa | 3.93E-06 | 0.44149837 | 0.377 | 0.279 | 0.07212541 |
| 2900026A02 | 3.94E-06 | 0.35539339 | 0.492 | 0.375 | 0.07221979 |
| Dner | 4.09E-06 | 0.41105143 | 0.387 | 0.288 | 0.07499783 |
| Auh | 4.29E-06 | 0.38463253 | 0.437 | 0.333 | 0.07856024 |
| Gm11099 | 4.53E-06 | 0.5438152 | 0.356 | 0.26 | 0.08297188 |
| Atp6v0b | 4.67E-06 | 0.21482226 | 0.718 | 0.6 | 0.0856334 |
| Vwa8 | 4.68E-06 | 0.31787659 | 0.416 | 0.312 | 0.08583777 |
| Ktn1 | 4.76E-06 | 0.23771369 | 0.588 | 0.47 | 0.08720988 |
| Slc3a2 | 5.05E-06 | 0.41100578 | 0.383 | 0.288 | 0.09262811 |
| Tenm4 | 5.11E-06 | 0.28261584 | 0.689 | 0.586 | 0.09375652 |
| Caln1 | 5.28E-06 | 0.18492193 | 0.607 | 0.484 | 0.09683572 |

| | | | | | |
|---|---|---|---|---|---|
| Ptprn2 | 5.41E-06 | 0.26189816 | 0.754 | 0.643 | 0.0990892 |
| Rapgef5 | 5.52E-06 | 0.35862562 | 0.602 | 0.487 | 0.10124619 |
| Ryr2 | 5.67E-06 | -0.3445111 | 0.798 | 0.79 | 0.10395646 |
| Ssr4 | 5.85E-06 | 0.517276 | 0.338 | 0.253 | 0.10722512 |
| Riok3 | 5.94E-06 | 0.37663576 | 0.398 | 0.306 | 0.10884158 |
| Ptpn1 | 5.98E-06 | 0.26752382 | 0.445 | 0.343 | 0.10967246 |
| Trhde | 6.57E-06 | 0.31351411 | 0.598 | 0.493 | 0.12048591 |
| Ptprz1 | 6.58E-06 | 0.56691715 | 0.348 | 0.255 | 0.12062431 |
| Ap3m1 | 6.82E-06 | 0.47137727 | 0.315 | 0.229 | 0.12495964 |
| Kcnh1 | 7.06E-06 | 0.33177971 | 0.55 | 0.43 | 0.12933513 |
| Atp5g2 | 7.32E-06 | 0.232514 | 0.315 | 0.22 | 0.13427402 |
| Wnk2 | 7.49E-06 | 0.24889737 | 0.532 | 0.421 | 0.13738488 |
| Arhgef25 | 7.91E-06 | 0.25159907 | 0.547 | 0.431 | 0.1449256 |
| Gabrb1 | 8.50E-06 | 0.27362888 | 0.817 | 0.76 | 0.15579544 |
| Cntnap2 | 8.58E-06 | -0.5884208 | 0.865 | 0.898 | 0.15728772 |
| Agap2 | 8.74E-06 | 0.44291242 | 0.394 | 0.31 | 0.16020299 |
| Cox4i1 | 8.84E-06 | 0.23159681 | 0.664 | 0.543 | 0.16211373 |
| Ptprr | 8.87E-06 | 0.22106203 | 0.499 | 0.377 | 0.16253234 |
| Ip6k2 | 9.21E-06 | 0.68126961 | 0.337 | 0.257 | 0.16884735 |
| Epha5 | 9.23E-06 | 0.28162629 | 0.812 | 0.753 | 0.16924933 |
| Hspa8 | 9.29E-06 | -0.5889204 | 0.626 | 0.668 | 0.17030087 |
| Rragb | 9.58E-06 | 0.52118803 | 0.267 | 0.187 | 0.17563682 |
| Gk | 9.67E-06 | 0.45544523 | 0.325 | 0.239 | 0.1772158 |
| Ubb | 9.81E-06 | -0.6366902 | 0.602 | 0.659 | 0.17979408 |
| Sox5 | 9.84E-06 | 0.43088725 | 0.447 | 0.348 | 0.1803217 |
| Nme7 | 9.90E-06 | 0.24914908 | 0.671 | 0.555 | 0.18149909 |
| Trim46 | 9.93E-06 | 0.42083817 | 0.378 | 0.295 | 0.1820589 |
| Wwp1 | 9.97E-06 | 0.28087075 | 0.377 | 0.279 | 0.18284571 |
| Smchd1 | 9.99E-06 | 0.3455066 | 0.417 | 0.317 | 0.18310646 |
| Myef2 | 9.99E-06 | 0.11079825 | 0.548 | 0.422 | 0.18321333 |
| Scaf11 | 1.01E-05 | 0.53463389 | 0.299 | 0.214 | 0.18501166 |
| Mark2 | 1.02E-05 | 0.23335266 | 0.506 | 0.396 | 0.18616033 |
| Numa1 | 1.04E-05 | 0.34517874 | 0.322 | 0.237 | 0.18994986 |
| Hectd2 | 1.07E-05 | 0.2832395 | 0.532 | 0.42 | 0.19681341 |
| Denn2b | 1.10E-05 | 0.37373589 | 0.349 | 0.26 | 0.20159399 |
| Cog5 | 1.13E-05 | 0.23140577 | 0.559 | 0.46 | 0.20705788 |
| Cenpv | 1.15E-05 | 0.31638343 | 0.34 | 0.25 | 0.21089227 |
| Tfcp2 | 1.19E-05 | 0.46215777 | 0.254 | 0.177 | 0.21776571 |
| Csmd1 | 1.21E-05 | 0.2003413 | 0.902 | 0.885 | 0.22150945 |
| Nkain2 | 1.25E-05 | -0.3656067 | 0.862 | 0.86 | 0.2282224 |
| Usp50 | 1.29E-05 | 0.41420673 | 0.381 | 0.297 | 0.23637073 |
| Gm20696 | 1.30E-05 | 0.30112099 | 0.426 | 0.33 | 0.23910196 |
| Tfdp2 | 1.34E-05 | 0.27221815 | 0.456 | 0.355 | 0.24511549 |

| | | | | | |
|---|---|---|---|---|---|
| Thoc1 | 1.34E-05 | 0.1712945 | 0.449 | 0.348 | 0.24593177 |
| Stxbp5l | 1.34E-05 | 0.33321672 | 0.764 | 0.668 | 0.24598302 |
| Slc24a2 | 1.35E-05 | -0.2216993 | 0.885 | 0.867 | 0.24689421 |
| Rc3h1 | 1.35E-05 | 0.4230548 | 0.397 | 0.31 | 0.24776509 |
| Kctd17 | 1.38E-05 | 0.47381452 | 0.323 | 0.242 | 0.25311915 |
| Frrs1l | 1.44E-05 | -0.8053322 | 0.468 | 0.529 | 0.26355796 |
| Bicdl1 | 1.44E-05 | 0.35027175 | 0.404 | 0.313 | 0.26391039 |
| Grin1 | 1.45E-05 | 0.19929096 | 0.69 | 0.579 | 0.26502921 |
| Eif4g1 | 1.45E-05 | 0.18991824 | 0.427 | 0.325 | 0.26514211 |
| Ddx5 | 1.48E-05 | 0.14728059 | 0.836 | 0.707 | 0.27048555 |
| 3110082I17R | 1.50E-05 | 0.67672798 | 0.271 | 0.194 | 0.27458295 |
| Cox5a | 1.53E-05 | 0.21420917 | 0.477 | 0.368 | 0.28111767 |
| Mrtfa | 1.56E-05 | 0.22561566 | 0.589 | 0.464 | 0.28552118 |
| Lurap1l | 1.59E-05 | 0.25662568 | 0.48 | 0.373 | 0.29105481 |
| Lrrc4 | 1.62E-05 | 0.20708996 | 0.591 | 0.484 | 0.29639663 |
| Trpc4 | 1.62E-05 | 0.19587329 | 0.494 | 0.385 | 0.29769461 |
| Ndufb9 | 1.64E-05 | 0.3823492 | 0.432 | 0.331 | 0.30025207 |
| Pgbd5 | 1.65E-05 | 0.24637485 | 0.629 | 0.517 | 0.30211606 |
| Pde4a | 1.65E-05 | 0.27775891 | 0.505 | 0.395 | 0.30330868 |
| Hnrnpdl | 1.71E-05 | 0.18577848 | 0.525 | 0.414 | 0.31316 |
| Oxr1 | 1.71E-05 | 0.12163932 | 0.761 | 0.653 | 0.31358574 |
| Glt8d1 | 1.73E-05 | 0.30344438 | 0.329 | 0.239 | 0.31627951 |
| Megf9 | 1.74E-05 | 0.29509421 | 0.514 | 0.416 | 0.31815564 |
| Sfxn5 | 1.74E-05 | 0.54997424 | 0.352 | 0.264 | 0.31957087 |
| Gpr63 | 1.76E-05 | 0.40041569 | 0.383 | 0.294 | 0.32258335 |
| Thrb | 1.80E-05 | 0.40406205 | 0.398 | 0.304 | 0.33026044 |
| Gtf2i | 1.81E-05 | 0.28560485 | 0.538 | 0.435 | 0.33145825 |
| Plk2 | 1.83E-05 | 0.50374676 | 0.353 | 0.268 | 0.3363258 |
| Slc30a6 | 1.85E-05 | 0.26875074 | 0.269 | 0.193 | 0.3389341 |
| Naa15 | 1.87E-05 | 0.46392482 | 0.413 | 0.327 | 0.3436735 |
| Sec14l1 | 1.88E-05 | 0.31868341 | 0.471 | 0.379 | 0.34418828 |
| Mmd | 1.95E-05 | 0.31448158 | 0.405 | 0.303 | 0.35662919 |
| Ints6l | 1.97E-05 | 0.39143109 | 0.378 | 0.287 | 0.36133657 |
| Tle4 | 2.01E-05 | 0.37085939 | 0.464 | 0.361 | 0.3691173 |
| Ppil2 | 2.12E-05 | 0.60421528 | 0.321 | 0.239 | 0.38844909 |
| Fhod3 | 2.13E-05 | 0.58691101 | 0.341 | 0.261 | 0.38975062 |
| Ptpn12 | 2.21E-05 | 0.28408785 | 0.417 | 0.333 | 0.40428336 |
| Slc6a17 | 2.25E-05 | 0.27971965 | 0.424 | 0.331 | 0.41290652 |
| Opcml | 2.32E-05 | -0.1585267 | 0.937 | 0.96 | 0.42522202 |
| Lrrn1 | 2.37E-05 | 0.39085852 | 0.27 | 0.186 | 0.4338971 |
| Clec16a | 2.39E-05 | 0.30207206 | 0.432 | 0.348 | 0.43722886 |
| Pten | 2.46E-05 | 0.13424697 | 0.555 | 0.454 | 0.45176386 |
| Esf1 | 2.47E-05 | 0.27331672 | 0.396 | 0.306 | 0.452464 |

| | | | | | |
|---|---|---|---|---|---|
| Abr | 2.47E-05 | 0.20534807 | 0.678 | 0.557 | 0.4530784 |
| Ints6 | 2.53E-05 | 0.2310763 | 0.512 | 0.401 | 0.46344491 |
| Galnt7 | 2.69E-05 | 0.36251226 | 0.314 | 0.234 | 0.49375352 |
| Neto2 | 2.70E-05 | 0.60277722 | 0.271 | 0.195 | 0.49429537 |
| Zdhhc6 | 2.72E-05 | 0.49973687 | 0.292 | 0.216 | 0.49836302 |
| Shisa7 | 2.77E-05 | 0.29414643 | 0.363 | 0.279 | 0.50742356 |
| Ptpn5 | 2.78E-05 | 0.44553508 | 0.424 | 0.332 | 0.5090981 |
| Ddx17 | 2.84E-05 | 0.21656454 | 0.638 | 0.529 | 0.52086738 |
| Thoc2l | 2.88E-05 | 0.2731062 | 0.499 | 0.39 | 0.52862588 |
| Atp5j2 | 2.90E-05 | 0.32195556 | 0.443 | 0.35 | 0.53177605 |
| Rabep1 | 2.95E-05 | 0.1772146 | 0.587 | 0.468 | 0.54102316 |
| Dgkd | 2.97E-05 | 0.31159809 | 0.51 | 0.405 | 0.5448761 |
| Spata2 | 3.03E-05 | 0.49313932 | 0.323 | 0.248 | 0.55533423 |
| Edil3 | 3.41E-05 | -0.7571572 | 0.432 | 0.487 | 0.62418659 |
| Rbm4b | 3.48E-05 | 0.26179569 | 0.477 | 0.382 | 0.63793048 |
| Lrrc8b | 3.53E-05 | 0.39266533 | 0.367 | 0.282 | 0.64696244 |
| Cers6 | 3.67E-05 | 0.17305525 | 0.673 | 0.551 | 0.67293692 |
| Cox8a | 3.94E-05 | 0.19904485 | 0.731 | 0.61 | 0.72313159 |
| Uxs1 | 3.99E-05 | 0.43117211 | 0.356 | 0.276 | 0.73131999 |
| Syt13 | 4.01E-05 | 0.34973591 | 0.386 | 0.305 | 0.734449 |
| Usp12 | 4.03E-05 | 0.37111374 | 0.322 | 0.241 | 0.7380851 |
| Ash1l | 4.06E-05 | 0.13708231 | 0.649 | 0.527 | 0.7442485 |
| Ccdc88c | 4.22E-05 | 0.3631644 | 0.31 | 0.232 | 0.77428944 |
| Lsamp | 4.23E-05 | -0.118654 | 0.988 | 0.988 | 0.77559362 |
| Rpl38 | 4.26E-05 | 0.10509684 | 0.554 | 0.433 | 0.78122363 |
| Arf5 | 4.30E-05 | 0.30393783 | 0.386 | 0.297 | 0.78802713 |
| Atp5o.1 | 4.60E-05 | 0.39702045 | 0.321 | 0.235 | 0.84367193 |
| Atp6v1b2 | 4.63E-05 | 0.17549321 | 0.555 | 0.451 | 0.84785434 |
| Rps9 | 4.71E-05 | 0.41763681 | 0.393 | 0.304 | 0.86357256 |
| Crtc3 | 4.82E-05 | 0.30844267 | 0.426 | 0.342 | 0.88296198 |
| Man1a2 | 4.88E-05 | 0.18019305 | 0.623 | 0.48 | 0.89521719 |
| Camk4 | 4.92E-05 | 0.17860085 | 0.569 | 0.432 | 0.9021057 |
| Adarb1 | 4.94E-05 | 0.29653903 | 0.467 | 0.37 | 0.9060187 |
| Arglu1 | 5.05E-05 | 0.1005724 | 0.7 | 0.586 | 0.92496799 |
| Slc25a12 | 5.06E-05 | 0.1706534 | 0.483 | 0.396 | 0.92810617 |
| Peli1 | 5.16E-05 | 0.36028965 | 0.438 | 0.352 | 0.94533811 |
| Gmds | 5.18E-05 | 0.27220046 | 0.494 | 0.385 | 0.94993676 |
| Tcf25 | 5.20E-05 | 0.17332953 | 0.529 | 0.429 | 0.95360687 |
| Sorl1 | 5.22E-05 | 0.21120922 | 0.608 | 0.498 | 0.95601106 |
| Srpr | 5.28E-05 | 0.28636588 | 0.363 | 0.282 | 0.96811397 |
| Epha10 | 5.41E-05 | 0.40817172 | 0.385 | 0.297 | 0.99149741 |
| Ncam2 | 5.45E-05 | -0.422718 | 0.64 | 0.652 | 0.99920647 |
| Sub1 | 5.49E-05 | 0.27679306 | 0.471 | 0.372 | 1 |

| | | | | | |
|---|---|---|---|---|---|
| Gtf3c3 | 5.52E-05 | 0.3291497 | 0.31 | 0.231 | 1 |
| Ube2g1 | 5.56E-05 | 0.20580543 | 0.45 | 0.359 | 1 |
| Psip1 | 5.60E-05 | 0.27271851 | 0.591 | 0.488 | 1 |
| Otulin | 5.60E-05 | 0.34206162 | 0.427 | 0.343 | 1 |
| Inpp4a | 5.64E-05 | 0.14476168 | 0.431 | 0.338 | 1 |
| Dip2b | 5.73E-05 | 0.18588352 | 0.568 | 0.455 | 1 |
| Sergef | 5.93E-05 | 0.36729426 | 0.415 | 0.328 | 1 |
| Atxn7l1 | 5.94E-05 | 0.27028381 | 0.456 | 0.363 | 1 |
| Lnpep | 5.98E-05 | 0.22754204 | 0.51 | 0.416 | 1 |
| Atp6ap1 | 6.13E-05 | 0.39707942 | 0.311 | 0.234 | 1 |
| Slc23a2 | 6.24E-05 | 0.31536365 | 0.487 | 0.391 | 1 |
| Wsb1 | 6.27E-05 | 0.42852229 | 0.321 | 0.249 | 1 |
| Akap8l | 6.60E-05 | 0.14962163 | 0.589 | 0.478 | 1 |
| Angptl2 | 6.63E-05 | 0.33205964 | 0.383 | 0.297 | 1 |
| Mecr | 6.63E-05 | 0.3914069 | 0.336 | 0.256 | 1 |
| Mef2c | 6.64E-05 | 0.19842891 | 0.438 | 0.333 | 1 |
| St8sia6 | 6.70E-05 | 0.58879582 | 0.289 | 0.214 | 1 |
| Atl1 | 6.80E-05 | 0.19057913 | 0.428 | 0.345 | 1 |
| Rapgef6 | 7.08E-05 | 0.24531946 | 0.598 | 0.5 | 1 |
| Camta2 | 7.14E-05 | 0.21943725 | 0.518 | 0.412 | 1 |
| Dtnb | 7.23E-05 | 0.21981077 | 0.623 | 0.516 | 1 |
| Rap1gap | 7.27E-05 | 0.32234394 | 0.396 | 0.32 | 1 |
| Gcnt7 | 7.27E-05 | 0.37616907 | 0.263 | 0.193 | 1 |
| Sh3d19 | 7.29E-05 | 0.46123165 | 0.473 | 0.388 | 1 |
| Mfn1 | 7.29E-05 | 0.62541638 | 0.281 | 0.204 | 1 |
| 3110021N24 | 7.38E-05 | 0.27166802 | 0.443 | 0.354 | 1 |
| Taf1d | 7.39E-05 | 0.40864383 | 0.398 | 0.315 | 1 |
| Agps | 7.46E-05 | 0.46933014 | 0.295 | 0.216 | 1 |
| Nrgn | 7.55E-05 | -0.3473107 | 0.693 | 0.731 | 1 |
| Zranb2 | 7.57E-05 | 0.19166387 | 0.701 | 0.579 | 1 |
| Scamp5 | 7.99E-05 | 0.21982039 | 0.387 | 0.304 | 1 |
| Angel2 | 8.11E-05 | 0.28446211 | 0.276 | 0.2 | 1 |
| Rp9 | 8.23E-05 | 0.51864954 | 0.338 | 0.266 | 1 |
| Xylt1 | 8.45E-05 | 0.30789723 | 0.375 | 0.293 | 1 |
| Sec61a2 | 8.52E-05 | 0.14320955 | 0.442 | 0.349 | 1 |
| Setd7 | 8.58E-05 | 0.20378762 | 0.419 | 0.33 | 1 |
| Unc13b | 8.73E-05 | 0.2414227 | 0.523 | 0.42 | 1 |
| Pdzrn4 | 8.84E-05 | 0.65554236 | 0.315 | 0.24 | 1 |
| Sirpa | 8.86E-05 | 0.37920307 | 0.302 | 0.224 | 1 |
| Zcchc18 | 8.87E-05 | 0.31471517 | 0.458 | 0.367 | 1 |
| Ptbp2 | 8.89E-05 | 0.29581974 | 0.469 | 0.378 | 1 |
| Kcnip2 | 8.97E-05 | 0.2490293 | 0.46 | 0.363 | 1 |
| Mpped2 | 9.09E-05 | 0.17543444 | 0.565 | 0.457 | 1 |

| | | | | | |
|---|---|---|---|---|---|
| Vps13d | 9.24E-05 | 0.25687073 | 0.52 | 0.421 | 1 |
| Slc22a17 | 9.34E-05 | 0.1419898 | 0.607 | 0.495 | 1 |
| Snd1 | 9.41E-05 | 0.19104186 | 0.505 | 0.41 | 1 |
| Pdss2 | 9.44E-05 | 0.19700787 | 0.438 | 0.348 | 1 |
| Sv2a | 9.60E-05 | 0.32493458 | 0.442 | 0.363 | 1 |
| Nrip3 | 9.60E-05 | 0.29147556 | 0.456 | 0.367 | 1 |
| Uggt2 | 9.64E-05 | 0.20694869 | 0.435 | 0.349 | 1 |
| Oxct1 | 9.66E-05 | 0.18486921 | 0.61 | 0.497 | 1 |
| Nxpe3 | 9.70E-05 | 0.2330409 | 0.278 | 0.21 | 1 |
| Pi4ka | 9.79E-05 | 0.13245781 | 0.636 | 0.507 | 1 |
| Flnb | 9.80E-05 | 0.14155513 | 0.52 | 0.417 | 1 |
| Syndig1 | 9.83E-05 | 0.27805992 | 0.49 | 0.381 | 1 |
| Pdzd4 | 9.90E-05 | 0.23323069 | 0.456 | 0.367 | 1 |
| Atxn2 | 9.91E-05 | 0.15982147 | 0.54 | 0.437 | 1 |
| Sema6d | 1.00E-04 | 0.34557023 | 0.404 | 0.319 | 1 |
| Synj1 | 0.00010007 | 0.1281485 | 0.632 | 0.522 | 1 |
| Dtx3 | 0.00010041 | 0.2760175 | 0.396 | 0.315 | 1 |
| Sertad2 | 0.00010111 | 0.35618206 | 0.401 | 0.319 | 1 |
| Arid4a | 0.00010395 | 0.22627661 | 0.45 | 0.361 | 1 |
| Cacna1b | 0.00010497 | 0.25810413 | 0.559 | 0.461 | 1 |
| Vps50 | 0.0001059 | 0.2505165 | 0.372 | 0.289 | 1 |
| Fmn2 | 0.00010734 | 0.12451126 | 0.629 | 0.516 | 1 |
| Srgap2 | 0.00010892 | 0.20180081 | 0.613 | 0.472 | 1 |
| Slc12a5 | 0.00010966 | 0.24368005 | 0.49 | 0.395 | 1 |
| Lsm7 | 0.00011093 | 0.16897657 | 0.28 | 0.207 | 1 |
| Sfswap | 0.0001117 | 0.19355636 | 0.469 | 0.383 | 1 |
| Skil | 0.00011255 | 0.42175293 | 0.271 | 0.199 | 1 |
| Gria3 | 0.00011269 | 0.21230122 | 0.778 | 0.675 | 1 |
| Reck | 0.0001127 | 0.60685475 | 0.273 | 0.205 | 1 |
| Akap13 | 0.00011398 | 0.15000591 | 0.539 | 0.411 | 1 |
| Hbs1l | 0.00011788 | 0.39054469 | 0.34 | 0.269 | 1 |
| Atp5h | 0.00011883 | 0.26911771 | 0.503 | 0.413 | 1 |
| Trpm3 | 0.00011897 | -0.3380486 | 0.63 | 0.657 | 1 |
| Dlgap1 | 0.00011962 | -0.2092967 | 0.902 | 0.915 | 1 |
| Ubxn4 | 0.00012037 | 0.17654158 | 0.506 | 0.403 | 1 |
| Rtn1 | 0.00012161 | 0.20734707 | 0.913 | 0.833 | 1 |
| Pcnx2 | 0.00012248 | 0.28583488 | 0.43 | 0.334 | 1 |
| Chd6 | 0.00012433 | 0.19404678 | 0.563 | 0.447 | 1 |
| Rps27a | 0.00012683 | 0.25056846 | 0.613 | 0.51 | 1 |
| Rpl18a | 0.00012728 | 0.28468852 | 0.378 | 0.288 | 1 |
| Eml4 | 0.00012779 | 0.12882833 | 0.487 | 0.395 | 1 |
| Rogdi | 0.00012853 | 0.25678607 | 0.381 | 0.299 | 1 |
| Ndufaf4 | 0.00012906 | 0.44831224 | 0.285 | 0.214 | 1 |

| | | | | | |
|---|---|---|---|---|---|
| Rpl15 | 0.00013023 | 0.24172236 | 0.536 | 0.438 | 1 |
| Ubp1 | 0.00013074 | 0.2762764 | 0.372 | 0.286 | 1 |
| Nedd8 | 0.00013092 | 0.3515038 | 0.329 | 0.248 | 1 |
| Cspp1 | 0.00013203 | 0.17124605 | 0.572 | 0.469 | 1 |
| S100pbp | 0.00013287 | 0.38431721 | 0.276 | 0.202 | 1 |
| Stx12 | 0.00013359 | 0.30968743 | 0.357 | 0.282 | 1 |
| Ep400 | 0.00013464 | 0.22734552 | 0.426 | 0.349 | 1 |
| Cpt1c | 0.00013503 | 0.32843838 | 0.426 | 0.346 | 1 |
| mt-Nd3 | 0.00013529 | -0.8816017 | 0.282 | 0.354 | 1 |
| Rsrp1 | 0.00013549 | 0.17138286 | 0.703 | 0.6 | 1 |
| Rb1cc1 | 0.00013557 | 0.17173493 | 0.562 | 0.463 | 1 |
| Usp14 | 0.0001391 | 0.31428505 | 0.411 | 0.335 | 1 |
| Jmjd1c | 0.00014283 | 0.13954826 | 0.719 | 0.604 | 1 |
| Mtif2 | 0.00014461 | 0.1715356 | 0.427 | 0.338 | 1 |
| Car10 | 0.00014618 | -0.8234623 | 0.291 | 0.361 | 1 |
| Immp2l | 0.00015054 | 0.18740325 | 0.606 | 0.5 | 1 |
| Spata7 | 0.00015108 | 0.1483336 | 0.342 | 0.262 | 1 |
| Acss1 | 0.00015322 | 0.59431626 | 0.308 | 0.244 | 1 |
| Tro | 0.0001539 | 0.35743626 | 0.347 | 0.271 | 1 |
| Usp45 | 0.00015415 | 0.24424456 | 0.435 | 0.35 | 1 |
| Timm8b | 0.00016096 | 0.60468598 | 0.302 | 0.228 | 1 |
| Tmem245 | 0.00016437 | 0.64275048 | 0.291 | 0.227 | 1 |
| Gm26992 | 0.000165 | 0.12223906 | 0.363 | 0.261 | 1 |
| Crim1 | 0.00016639 | 0.3512663 | 0.441 | 0.351 | 1 |
| Hacd1 | 0.00016702 | 0.4084122 | 0.269 | 0.198 | 1 |
| Arhgap39 | 0.00016707 | 0.12301 | 0.674 | 0.54 | 1 |
| Tpp2 | 0.00017251 | 0.34653662 | 0.4 | 0.319 | 1 |
| Zfhx4 | 0.000173 | 0.34468835 | 0.291 | 0.216 | 1 |
| Fhl2 | 0.00017377 | 0.43044172 | 0.25 | 0.184 | 1 |
| Vcl | 0.00017522 | 0.37514212 | 0.296 | 0.226 | 1 |
| Klhl22 | 0.00017739 | 0.4982452 | 0.306 | 0.238 | 1 |
| Ppp4r4 | 0.00017782 | 0.26726436 | 0.374 | 0.299 | 1 |
| Supt16 | 0.00017933 | 0.47860086 | 0.254 | 0.187 | 1 |
| Sema3e | 0.00017992 | -0.6575097 | 0.377 | 0.442 | 1 |
| Rps27 | 0.00018246 | 0.27499481 | 0.528 | 0.443 | 1 |
| Arap2 | 0.00018345 | 0.17289459 | 0.484 | 0.376 | 1 |
| Mcts1 | 0.00018638 | 0.20725354 | 0.314 | 0.239 | 1 |
| 5031439G07 | 0.00019083 | 0.29735386 | 0.303 | 0.234 | 1 |
| Tgfbr3 | 0.00019631 | 0.32765437 | 0.431 | 0.35 | 1 |
| Sec24b | 0.00019926 | 0.2015689 | 0.396 | 0.311 | 1 |
| Pcca | 0.00020539 | 0.16175147 | 0.538 | 0.429 | 1 |
| Chst11 | 0.00020569 | 0.24859526 | 0.416 | 0.33 | 1 |
| Reep1 | 0.00020616 | 0.20172345 | 0.487 | 0.396 | 1 |

| | | | | | |
|---|---|---|---|---|---|
| A830018L16I | 0.00020623 | -0.6227709 | 0.488 | 0.5 | 1 |
| Tgfb2 | 0.00021077 | 0.51577671 | 0.297 | 0.229 | 1 |
| Pip5k1a | 0.00021162 | 0.19298656 | 0.503 | 0.409 | 1 |
| Rasa1 | 0.00021435 | 0.13161357 | 0.499 | 0.407 | 1 |
| Onecut2 | 0.00021622 | 0.30045562 | 0.338 | 0.266 | 1 |
| Arhgef7 | 0.00022152 | 0.18272431 | 0.428 | 0.348 | 1 |
| Rai1 | 0.00022523 | 0.13244094 | 0.446 | 0.356 | 1 |
| Arhgap26.1 | 0.00022755 | 0.25465795 | 0.454 | 0.365 | 1 |
| Mier1 | 0.00023188 | 0.11919023 | 0.382 | 0.3 | 1 |
| Sirt7 | 0.00023296 | 0.45862622 | 0.255 | 0.191 | 1 |
| Mblac2 | 0.00023533 | 0.19414092 | 0.303 | 0.226 | 1 |
| Adam15 | 0.00023602 | 0.47359932 | 0.263 | 0.199 | 1 |
| Usp34 | 0.0002399 | 0.14244545 | 0.718 | 0.602 | 1 |
| Cadm3 | 0.00024494 | 0.17226766 | 0.383 | 0.297 | 1 |
| Nfia | 0.00024989 | -0.4741852 | 0.678 | 0.668 | 1 |
| Cep83 | 0.00025023 | 0.24746388 | 0.363 | 0.285 | 1 |
| Txlng | 0.00025069 | 0.25685744 | 0.385 | 0.303 | 1 |
| Usp22 | 0.00025108 | 0.3416589 | 0.306 | 0.233 | 1 |
| Pus10 | 0.00025858 | 0.32832105 | 0.263 | 0.198 | 1 |
| Abi1 | 0.00026279 | 0.13118255 | 0.611 | 0.508 | 1 |
| Tmem243 | 0.00027372 | 0.14054549 | 0.456 | 0.368 | 1 |
| Adar | 0.00027467 | 0.31899742 | 0.375 | 0.303 | 1 |
| Kdm4c | 0.00027542 | 0.13263634 | 0.551 | 0.448 | 1 |
| Pcdh10 | 0.00027734 | 0.47180102 | 0.342 | 0.275 | 1 |
| Actn1 | 0.00028353 | 0.26744954 | 0.383 | 0.315 | 1 |
| Trio | 0.00028393 | 0.17891516 | 0.705 | 0.586 | 1 |
| Reep2 | 0.00028551 | 0.21390827 | 0.311 | 0.24 | 1 |
| Ly6h | 0.00029009 | 0.18702915 | 0.603 | 0.505 | 1 |
| Chchd2 | 0.00029139 | 0.23764063 | 0.634 | 0.534 | 1 |
| Fnbp1 | 0.00029261 | -0.9004664 | 0.337 | 0.407 | 1 |
| Trim35 | 0.00029815 | 0.26201746 | 0.51 | 0.422 | 1 |
| Sntg1 | 0.00029886 | -0.363239 | 0.742 | 0.723 | 1 |
| Smarcc1 | 0.00030535 | 0.20685432 | 0.352 | 0.279 | 1 |
| Ube2h | 0.00030549 | 0.1321208 | 0.476 | 0.392 | 1 |
| Zdhhc23 | 0.00030688 | 0.38539468 | 0.332 | 0.261 | 1 |
| Sil1 | 0.00030734 | 0.45741189 | 0.28 | 0.212 | 1 |
| Shank2 | 0.00031096 | 0.16565268 | 0.615 | 0.495 | 1 |
| Srsf7 | 0.00031429 | 0.13977201 | 0.578 | 0.478 | 1 |
| Rasgrf1 | 0.00031716 | 0.15195983 | 0.742 | 0.657 | 1 |
| Rev3l | 0.00031725 | 0.14371917 | 0.621 | 0.503 | 1 |
| Rps21 | 0.00031732 | 0.2154161 | 0.57 | 0.47 | 1 |
| Fbxl17 | 0.00031861 | 0.11593408 | 0.688 | 0.57 | 1 |
| Atrn | 0.00031951 | 0.17065214 | 0.551 | 0.444 | 1 |

| | | | | | |
|---|---|---|---|---|---|
| Sh3rf3 | 0.0003199 | 0.14894788 | 0.536 | 0.422 | 1 |
| Mvb12b | 0.00032024 | 0.11497358 | 0.347 | 0.264 | 1 |
| Ndufs4 | 0.00032689 | 0.25056208 | 0.479 | 0.391 | 1 |
| Sec22a | 0.00032903 | 0.23314624 | 0.25 | 0.18 | 1 |
| Pdlim7 | 0.00033005 | 0.28645613 | 0.381 | 0.304 | 1 |
| Sdcbp | 0.00033102 | 0.29908105 | 0.397 | 0.317 | 1 |
| Bcan | 0.00033685 | 0.54814042 | 0.276 | 0.209 | 1 |
| Ehmt1 | 0.00034019 | 0.16628123 | 0.494 | 0.418 | 1 |
| Zer1 | 0.00034212 | 0.31588235 | 0.368 | 0.3 | 1 |
| Neo1 | 0.00034342 | 0.12100793 | 0.473 | 0.393 | 1 |
| Zmynd11 | 0.00035506 | 0.14420143 | 0.578 | 0.479 | 1 |
| Susd4 | 0.0003553 | 0.20762274 | 0.529 | 0.431 | 1 |
| Ccny | 0.0003578 | 0.28749276 | 0.512 | 0.421 | 1 |
| Cenpp | 0.00035809 | 0.19768067 | 0.307 | 0.238 | 1 |
| Bri3 | 0.00035825 | 0.31203254 | 0.31 | 0.239 | 1 |
| Arl6ip5 | 0.00036042 | 0.15113236 | 0.286 | 0.216 | 1 |
| Susd6 | 0.00036256 | 0.19605206 | 0.589 | 0.483 | 1 |
| Ogdh | 0.00036386 | 0.22016649 | 0.422 | 0.347 | 1 |
| Xrn1 | 0.00037368 | 0.22470613 | 0.495 | 0.414 | 1 |
| Clint1 | 0.00037392 | 0.32926165 | 0.407 | 0.333 | 1 |
| Raf1 | 0.00037899 | 0.25790017 | 0.378 | 0.303 | 1 |
| Snx27 | 0.00039091 | 0.18852186 | 0.362 | 0.287 | 1 |
| Dnaja2 | 0.00039202 | 0.10556503 | 0.467 | 0.383 | 1 |
| Prelid3a | 0.00039875 | 0.37165385 | 0.352 | 0.283 | 1 |
| Astn2 | 0.0004104 | 0.10893996 | 0.546 | 0.457 | 1 |
| Cabin1 | 0.0004114 | 0.16893338 | 0.435 | 0.35 | 1 |
| Ksr2 | 0.00041264 | 0.11187539 | 0.584 | 0.494 | 1 |
| Bptf | 0.00041568 | 0.22670797 | 0.508 | 0.42 | 1 |
| Fgfr2 | 0.00042038 | 0.21749831 | 0.752 | 0.658 | 1 |
| Gnptg | 0.00042305 | 0.25700558 | 0.293 | 0.224 | 1 |
| Fam110b | 0.00042588 | 0.30771126 | 0.386 | 0.314 | 1 |
| Camsap2 | 0.00044114 | 0.2825784 | 0.426 | 0.345 | 1 |
| St7 | 0.00044148 | 0.14436699 | 0.565 | 0.476 | 1 |
| Rps25 | 0.00044748 | 0.34170273 | 0.383 | 0.304 | 1 |
| Ccnl1 | 0.00045951 | 0.16994549 | 0.381 | 0.308 | 1 |
| Ola1 | 0.00045976 | 0.23756402 | 0.465 | 0.391 | 1 |
| Gtf3c1 | 0.00046046 | 0.40180423 | 0.311 | 0.246 | 1 |
| Dennd4a | 0.00046812 | 0.21173913 | 0.456 | 0.367 | 1 |
| Snrnp70 | 0.00047303 | 0.14594657 | 0.744 | 0.603 | 1 |
| Wdr26 | 0.0004906 | 0.34095277 | 0.407 | 0.333 | 1 |
| Ube2v2 | 0.00049624 | 0.31091492 | 0.345 | 0.276 | 1 |
| Scai | 0.0004978 | 0.22056648 | 0.599 | 0.503 | 1 |
| Spast | 0.00049809 | 0.1530589 | 0.325 | 0.252 | 1 |

| | | | | | |
|---|---|---|---|---|---|
| Snap25 | 0.00050203 | 0.15471351 | 0.929 | 0.862 | 1 |
| Zfp654 | 0.00050798 | 0.20639155 | 0.352 | 0.28 | 1 |
| Psmd1 | 0.00051068 | 0.26190141 | 0.381 | 0.313 | 1 |
| Nab1 | 0.00051266 | 0.47897019 | 0.274 | 0.212 | 1 |
| Chic2 | 0.00051348 | 0.49026135 | 0.281 | 0.218 | 1 |
| Ocrl | 0.00051813 | 0.3143724 | 0.404 | 0.33 | 1 |
| Calm3 | 0.00052303 | 0.159948 | 0.585 | 0.494 | 1 |
| Elmod1 | 0.0005234 | 0.18707678 | 0.662 | 0.571 | 1 |
| Cetn3 | 0.00052769 | 0.23894463 | 0.299 | 0.23 | 1 |
| Tmem161b | 0.00052998 | 0.2249556 | 0.419 | 0.345 | 1 |
| Lrch3 | 0.00053915 | 0.30500662 | 0.438 | 0.365 | 1 |
| Crebzf | 0.000541 | 0.14886861 | 0.382 | 0.312 | 1 |
| Flrt3 | 0.0005423 | 0.21639935 | 0.397 | 0.329 | 1 |
| Ctnnd2 | 0.00054884 | -0.2529933 | 0.786 | 0.755 | 1 |
| Psmd4 | 0.00056919 | 0.20052363 | 0.368 | 0.298 | 1 |
| Taok1 | 0.00057162 | 0.13880688 | 0.562 | 0.477 | 1 |
| Adam11 | 0.00057311 | 0.2329665 | 0.353 | 0.283 | 1 |
| Atxn7 | 0.00057413 | 0.13402437 | 0.378 | 0.308 | 1 |
| Pspc1 | 0.00057458 | 0.37215925 | 0.394 | 0.326 | 1 |
| Ppp2r2b | 0.00058278 | 0.19075268 | 0.708 | 0.593 | 1 |
| Slc6a15 | 0.00058948 | 0.33109344 | 0.347 | 0.28 | 1 |
| Slc4a10 | 0.00059264 | 0.18410859 | 0.655 | 0.544 | 1 |
| Phf20 | 0.00059738 | 0.15630141 | 0.569 | 0.457 | 1 |
| Mosmo | 0.0005979 | 0.37183751 | 0.355 | 0.284 | 1 |
| Mib1 | 0.00060171 | 0.11756266 | 0.559 | 0.461 | 1 |
| Cdc123 | 0.0006025 | 0.44990733 | 0.265 | 0.201 | 1 |
| Mllt11 | 0.00060509 | 0.13336846 | 0.334 | 0.262 | 1 |
| Prkcz | 0.00060803 | 0.19046895 | 0.413 | 0.329 | 1 |
| Wdr47 | 0.00062303 | 0.1695224 | 0.409 | 0.334 | 1 |
| Txndc11 | 0.00062382 | 0.38484202 | 0.334 | 0.265 | 1 |
| Las1l | 0.00062663 | 0.32352895 | 0.256 | 0.194 | 1 |
| Dnah9 | 0.00062724 | 0.22738768 | 0.475 | 0.383 | 1 |
| Rngtt | 0.00062842 | 0.26611356 | 0.413 | 0.337 | 1 |
| Tent4b | 0.00064058 | 0.26681718 | 0.379 | 0.313 | 1 |
| Slc25a4 | 0.0006464 | 0.17798301 | 0.572 | 0.479 | 1 |
| Fmnl2 | 0.00065068 | 0.19193822 | 0.544 | 0.454 | 1 |
| Slain1 | 0.00065397 | 0.3061006 | 0.319 | 0.251 | 1 |
| Ankrd12 | 0.00066328 | 0.22848157 | 0.622 | 0.524 | 1 |
| Ano4 | 0.00066745 | -0.6536677 | 0.443 | 0.469 | 1 |
| Smurf1 | 0.00067274 | 0.38699062 | 0.318 | 0.254 | 1 |
| Abi2 | 0.00067561 | 0.12699491 | 0.604 | 0.514 | 1 |
| Clasrp | 0.00067843 | 0.43429582 | 0.278 | 0.217 | 1 |
| Gdi2 | 0.00068132 | 0.24112718 | 0.366 | 0.292 | 1 |

| | | | | | |
|---|---|---|---|---|---|
| Zbtb7a | 0.00068577 | 0.38241294 | 0.261 | 0.2 | 1 |
| Tsc22d2 | 0.00069137 | 0.27998399 | 0.405 | 0.328 | 1 |
| Kifc2 | 0.00069838 | 0.35138355 | 0.344 | 0.276 | 1 |
| Ndufb8 | 0.00069852 | 0.41657432 | 0.308 | 0.239 | 1 |
| Ccdc149 | 0.00070136 | 0.30794256 | 0.258 | 0.198 | 1 |
| Impact | 0.00070273 | 0.15894824 | 0.467 | 0.391 | 1 |
| Rpl12 | 0.0007038 | 0.27438065 | 0.37 | 0.291 | 1 |
| Elavl2 | 0.00070862 | 0.22871385 | 0.538 | 0.443 | 1 |
| Enc1 | 0.00071831 | 0.25976225 | 0.503 | 0.427 | 1 |
| Bsg | 0.00071951 | 0.20228765 | 0.423 | 0.355 | 1 |
| Gm10563 | 0.00072266 | 0.29773215 | 0.304 | 0.243 | 1 |
| Zfp704 | 0.00072381 | 0.12367398 | 0.379 | 0.307 | 1 |
| Slc8a1 | 0.00073105 | 0.17714544 | 0.757 | 0.683 | 1 |
| Tmem178b | 0.00073843 | 0.15330504 | 0.707 | 0.615 | 1 |
| Tuba4a | 0.00073911 | 0.233795 | 0.345 | 0.271 | 1 |
| St6galnac3 | 0.00073957 | -0.1758893 | 0.506 | 0.399 | 1 |
| Klf12 | 0.00074249 | 0.13243674 | 0.599 | 0.497 | 1 |
| Orc4 | 0.00075337 | 0.20124807 | 0.445 | 0.375 | 1 |
| Porcn | 0.0007708 | 0.38379057 | 0.329 | 0.261 | 1 |
| Braf | 0.00077095 | 0.12867676 | 0.641 | 0.542 | 1 |
| Rc3h2 | 0.00077105 | 0.20886291 | 0.461 | 0.388 | 1 |
| Plekha1 | 0.00077149 | 0.1445335 | 0.413 | 0.339 | 1 |
| Tspan7 | 0.0007877 | 0.24284062 | 0.716 | 0.615 | 1 |
| Babam2 | 0.00079513 | 0.14654397 | 0.412 | 0.333 | 1 |
| Nedd4l | 0.00079845 | 0.14066411 | 0.737 | 0.607 | 1 |
| Arhgef26 | 0.00080412 | 0.30055866 | 0.417 | 0.338 | 1 |
| Gria1 | 0.00082283 | 0.10108148 | 0.958 | 0.95 | 1 |
| Sipa1l3 | 0.00082513 | 0.11459802 | 0.632 | 0.518 | 1 |
| Slc25a26 | 0.00083146 | 0.18855816 | 0.266 | 0.204 | 1 |
| Rbm39 | 0.00084043 | 0.15096125 | 0.783 | 0.664 | 1 |
| Capzb | 0.00084053 | 0.23506332 | 0.432 | 0.358 | 1 |
| Kit | 0.00084382 | 0.40681214 | 0.267 | 0.204 | 1 |
| Ptprs | 0.00084416 | 0.15026974 | 0.705 | 0.592 | 1 |
| Rras2 | 0.00084724 | 0.11169539 | 0.322 | 0.252 | 1 |
| Ewsr1 | 0.00084943 | 0.11692009 | 0.512 | 0.418 | 1 |
| Chuk | 0.0008643 | 0.14364731 | 0.352 | 0.286 | 1 |
| Ahi1 | 0.00086981 | 0.11381826 | 0.739 | 0.619 | 1 |
| Mrps5 | 0.00087289 | 0.23041662 | 0.355 | 0.283 | 1 |
| Slc16a4 | 0.00087393 | 0.462065 | 0.27 | 0.216 | 1 |
| Fam92a | 0.00087562 | 0.68906053 | 0.291 | 0.237 | 1 |
| Nsmaf | 0.00087628 | 0.36333226 | 0.266 | 0.207 | 1 |
| Camk1d | 0.00088305 | 0.1511949 | 0.87 | 0.788 | 1 |
| Ttc7b | 0.00088827 | 0.23739433 | 0.482 | 0.402 | 1 |

| | | | | | |
|---|---|---|---|---|---|
| Tmx4 | 0.00089874 | 0.15363654 | 0.508 | 0.42 | 1 |
| Psma3 | 0.00089982 | 0.17693015 | 0.523 | 0.434 | 1 |
| Kdm5b | 0.00090374 | 0.13769621 | 0.435 | 0.351 | 1 |
| Klhdc10 | 0.00090624 | 0.18498124 | 0.523 | 0.434 | 1 |
| Cadps2 | 0.00090796 | 0.17127178 | 0.419 | 0.339 | 1 |
| Cck | 0.00091183 | 0.14468569 | 0.798 | 0.679 | 1 |
| Trpc5 | 0.00091344 | 0.28327669 | 0.587 | 0.474 | 1 |
| Nr2c2 | 0.00091807 | 0.20823824 | 0.419 | 0.346 | 1 |
| St13 | 0.00092284 | 0.18645422 | 0.387 | 0.319 | 1 |
| Selenom | 0.00092569 | 0.30895988 | 0.405 | 0.336 | 1 |
| Tln2 | 0.00095313 | 0.24644597 | 0.42 | 0.339 | 1 |
| Asb3 | 0.00096795 | 0.11816164 | 0.479 | 0.391 | 1 |
| Amy1 | 0.00097273 | 0.10133493 | 0.333 | 0.26 | 1 |
| Orc2 | 0.00097496 | 0.36848604 | 0.314 | 0.251 | 1 |
| Rplp1 | 0.00097913 | 0.11022554 | 0.779 | 0.668 | 1 |
| Tango2 | 0.00098026 | 0.14668361 | 0.404 | 0.323 | 1 |
| Lekr1 | 0.0009812 | 0.44881148 | 0.299 | 0.235 | 1 |
| Tet2 | 0.0009822 | 0.30538052 | 0.329 | 0.264 | 1 |
| Eif4g3 | 0.0009838 | 0.1156383 | 0.611 | 0.514 | 1 |
| Nphp4 | 0.00098503 | 0.37495057 | 0.329 | 0.271 | 1 |
| Htt | 0.00098562 | 0.13744639 | 0.482 | 0.392 | 1 |
| Ppp1r2 | 0.00100072 | 0.43207392 | 0.307 | 0.245 | 1 |
| Fchsd2 | 0.00101911 | 0.33054379 | 0.393 | 0.33 | 1 |
| Rhof | 0.00102173 | 0.43089796 | 0.252 | 0.198 | 1 |
| Ino80d | 0.00102907 | 0.27957077 | 0.377 | 0.305 | 1 |
| Lars | 0.00102915 | 0.19188462 | 0.396 | 0.327 | 1 |
| Wdfy2 | 0.00104727 | 0.34612597 | 0.362 | 0.295 | 1 |
| Tecr | 0.00104972 | 0.12917275 | 0.517 | 0.436 | 1 |
| Arpc3 | 0.00105353 | 0.18818618 | 0.453 | 0.382 | 1 |
| Rpsa | 0.00106244 | 0.1152815 | 0.458 | 0.37 | 1 |
| Clip4 | 0.00107386 | 0.28292145 | 0.276 | 0.215 | 1 |
| Kpna3 | 0.00107524 | 0.11220979 | 0.461 | 0.383 | 1 |
| Lamc1 | 0.00108368 | 0.21302122 | 0.261 | 0.202 | 1 |
| Nup214 | 0.00110629 | 0.36471705 | 0.297 | 0.241 | 1 |
| Mrps26 | 0.00110806 | 0.23068956 | 0.281 | 0.221 | 1 |
| Neurod2 | 0.00111103 | 0.1927186 | 0.435 | 0.361 | 1 |
| Inf2 | 0.00111477 | 0.23358212 | 0.407 | 0.332 | 1 |
| Pitpna | 0.00111981 | 0.1172768 | 0.508 | 0.429 | 1 |
| Gnl3 | 0.00113394 | 0.35162924 | 0.289 | 0.229 | 1 |
| Agk | 0.0011412 | 0.22084453 | 0.302 | 0.239 | 1 |
| Cdc40 | 0.00114146 | 0.14721123 | 0.553 | 0.457 | 1 |
| Atp1b1 | 0.00115253 | 0.16616587 | 0.892 | 0.8 | 1 |
| Maco1 | 0.00115329 | 0.21604766 | 0.412 | 0.333 | 1 |

| | | | | | |
|---|---|---|---|---|---|
| Camkv | 0.00116498 | 0.14592987 | 0.569 | 0.465 | 1 |
| Aopep | 0.00119164 | 0.17059806 | 0.637 | 0.553 | 1 |
| Ncor1 | 0.00120642 | 0.14341841 | 0.709 | 0.603 | 1 |
| Capn15 | 0.00121028 | 0.2474494 | 0.333 | 0.272 | 1 |
| Phc3 | 0.00121717 | 0.15729881 | 0.43 | 0.35 | 1 |
| Tra2a | 0.00122067 | 0.12446324 | 0.637 | 0.521 | 1 |
| Nrbp2 | 0.00122867 | 0.40323818 | 0.258 | 0.199 | 1 |
| Nptxr | 0.00123467 | 0.18315832 | 0.416 | 0.336 | 1 |
| Rpl6 | 0.00123973 | 0.19836491 | 0.468 | 0.391 | 1 |
| Ptprj | 0.00126102 | 0.20151176 | 0.693 | 0.564 | 1 |
| Iqgap1 | 0.00127725 | 0.47432059 | 0.273 | 0.215 | 1 |
| Atp5d | 0.00129956 | 0.33659233 | 0.307 | 0.241 | 1 |
| Hacd2 | 0.00132354 | 0.28503179 | 0.397 | 0.338 | 1 |
| Upf2 | 0.00132567 | 0.15368607 | 0.409 | 0.345 | 1 |
| Tmem260 | 0.0013284 | 0.26382332 | 0.263 | 0.201 | 1 |
| Cacng7 | 0.00133236 | 0.31751008 | 0.3 | 0.242 | 1 |
| Sumo1 | 0.00133305 | 0.14554275 | 0.437 | 0.361 | 1 |
| Uvrag | 0.00133425 | 0.10170703 | 0.529 | 0.434 | 1 |
| Senp2 | 0.00133426 | 0.17320067 | 0.321 | 0.256 | 1 |
| Mysm1 | 0.00133605 | 0.31432759 | 0.416 | 0.348 | 1 |
| Tacc1 | 0.00138966 | 0.14576534 | 0.4 | 0.328 | 1 |
| Kif5b | 0.00139926 | 0.10837002 | 0.547 | 0.454 | 1 |
| Ube4a | 0.00141527 | 0.21269711 | 0.352 | 0.29 | 1 |
| Dennd6a | 0.00142034 | 0.16359518 | 0.362 | 0.302 | 1 |
| Rps6kb2 | 0.00146048 | 0.37258764 | 0.273 | 0.218 | 1 |
| Bmpr1a | 0.00146647 | 0.39330283 | 0.363 | 0.304 | 1 |
| Arid4b | 0.00148395 | 0.18247208 | 0.551 | 0.476 | 1 |
| Rnpc3 | 0.00148412 | 0.28985393 | 0.315 | 0.255 | 1 |
| Ppp3cc | 0.00148928 | 0.21851678 | 0.292 | 0.232 | 1 |
| Mtor | 0.00150599 | 0.17743569 | 0.312 | 0.256 | 1 |
| Trappc8 | 0.00151769 | 0.14776543 | 0.392 | 0.321 | 1 |
| Clip1 | 0.00154401 | 0.1650739 | 0.584 | 0.481 | 1 |
| Pds5b | 0.00154666 | 0.13987112 | 0.488 | 0.4 | 1 |
| Mapk8ip3 | 0.00155089 | 0.19192258 | 0.464 | 0.385 | 1 |
| Cntn3 | 0.00155816 | 0.33605768 | 0.362 | 0.287 | 1 |
| Fam193b | 0.00156568 | 0.26602076 | 0.314 | 0.256 | 1 |
| Ddhd2 | 0.00159137 | 0.12614018 | 0.477 | 0.397 | 1 |
| Arhgef9 | 0.00161131 | 0.12531536 | 0.619 | 0.52 | 1 |
| Mindy3 | 0.00161335 | 0.13692913 | 0.411 | 0.347 | 1 |
| Spats2 | 0.00161707 | 0.17020696 | 0.326 | 0.266 | 1 |
| Higd1a | 0.00162615 | 0.47421009 | 0.252 | 0.196 | 1 |
| Supt20 | 0.00163191 | 0.21694332 | 0.37 | 0.31 | 1 |
| Runx1t1 | 0.00163823 | 0.11853644 | 0.524 | 0.435 | 1 |

| | | | | | |
|---|---|---|---|---|---|
| Ubap2 | 0.00164799 | 0.36194373 | 0.338 | 0.281 | 1 |
| Efl1 | 0.00166914 | 0.15493599 | 0.258 | 0.2 | 1 |
| Fam49b | 0.00169194 | 0.15510332 | 0.521 | 0.434 | 1 |
| Fam120a | 0.0016985 | 0.17517471 | 0.427 | 0.36 | 1 |
| Tmem175 | 0.00172165 | 0.2977638 | 0.359 | 0.295 | 1 |
| Tma7 | 0.0017522 | 0.36392504 | 0.338 | 0.277 | 1 |
| Ei24 | 0.00175413 | 0.27011497 | 0.259 | 0.2 | 1 |
| Clip3 | 0.00177976 | 0.15763746 | 0.315 | 0.254 | 1 |
| Mgat4c | 0.00178298 | 0.23700473 | 0.689 | 0.636 | 1 |
| Otud4 | 0.00178375 | 0.20491712 | 0.336 | 0.275 | 1 |
| Cntn5 | 0.0018043 | -0.624304 | 0.338 | 0.393 | 1 |
| Atp2b1 | 0.00182019 | -0.2074277 | 0.92 | 0.897 | 1 |
| Cyld | 0.00183677 | 0.16985345 | 0.409 | 0.339 | 1 |
| Atp13a3 | 0.00184266 | 0.38208553 | 0.302 | 0.247 | 1 |
| Acot7 | 0.00185932 | 0.3138057 | 0.356 | 0.301 | 1 |
| Snx24 | 0.0018663 | 0.23211071 | 0.383 | 0.325 | 1 |
| Vps29 | 0.00188429 | 0.36264053 | 0.36 | 0.299 | 1 |
| Ppp2r5c | 0.00188604 | 0.1618703 | 0.574 | 0.489 | 1 |
| Uqcr11 | 0.00189409 | 0.23179165 | 0.394 | 0.327 | 1 |
| Tpr | 0.00189449 | 0.1438668 | 0.477 | 0.405 | 1 |
| Anks3 | 0.00189461 | 0.34018568 | 0.344 | 0.291 | 1 |
| Zmat2 | 0.00191151 | 0.21593351 | 0.286 | 0.224 | 1 |
| Sez6 | 0.00191188 | 0.16257867 | 0.334 | 0.267 | 1 |
| Sacm1l | 0.00191344 | 0.15500897 | 0.408 | 0.337 | 1 |
| Gde1 | 0.00195636 | 0.1003984 | 0.419 | 0.346 | 1 |
| Paip2 | 0.00197803 | 0.12995157 | 0.503 | 0.427 | 1 |
| Plekhm3 | 0.00202 | 0.27048296 | 0.327 | 0.272 | 1 |
| Macf1 | 0.00202002 | 0.11640795 | 0.771 | 0.675 | 1 |
| Ext1 | 0.00206111 | 0.14054315 | 0.655 | 0.552 | 1 |
| Dock4 | 0.00206404 | 0.1734382 | 0.814 | 0.756 | 1 |
| Mtrex | 0.00207929 | 0.1697521 | 0.28 | 0.22 | 1 |
| Hdac4 | 0.00209059 | 0.14657204 | 0.46 | 0.375 | 1 |
| Irf2 | 0.00212643 | 0.12040505 | 0.445 | 0.372 | 1 |
| Phf8 | 0.00217019 | 0.18218792 | 0.3 | 0.241 | 1 |
| Memo1 | 0.00218968 | 0.1898908 | 0.385 | 0.326 | 1 |
| Pbrm1 | 0.00221016 | 0.12405075 | 0.461 | 0.392 | 1 |
| C530008M17 | 0.00224613 | 0.32474492 | 0.367 | 0.303 | 1 |
| Adam23 | 0.00225186 | 0.13152083 | 0.559 | 0.47 | 1 |
| Exoc2 | 0.00228271 | 0.16251335 | 0.314 | 0.258 | 1 |
| Dock10 | 0.00228374 | -0.5038001 | 0.321 | 0.378 | 1 |
| Fau | 0.0022884 | 0.25400517 | 0.494 | 0.412 | 1 |
| Tasp1 | 0.00230725 | 0.18177498 | 0.4 | 0.34 | 1 |
| Phf14 | 0.00233267 | 0.11276972 | 0.61 | 0.5 | 1 |

| | | | | | |
|---|---|---|---|---|---|
| Dph6 | 0.00233306 | 0.15760243 | 0.273 | 0.217 | 1 |
| Kcmf1 | 0.00234951 | 0.34893735 | 0.356 | 0.298 | 1 |
| Scamp1 | 0.00236078 | 0.17854072 | 0.372 | 0.311 | 1 |
| Mlf2 | 0.00236939 | 0.10122226 | 0.368 | 0.304 | 1 |
| Gns | 0.00237066 | 0.10016537 | 0.277 | 0.217 | 1 |
| Cep295 | 0.00237106 | 0.29763653 | 0.289 | 0.233 | 1 |
| Smim4 | 0.00238218 | 0.19134106 | 0.296 | 0.237 | 1 |
| Ttc4 | 0.00239547 | 0.28137807 | 0.274 | 0.218 | 1 |
| Pkp2 | 0.00240044 | 0.10888825 | 0.49 | 0.402 | 1 |
| Map3k3 | 0.00240809 | 0.12726455 | 0.446 | 0.375 | 1 |
| Mphosph9 | 0.00241447 | 0.14607101 | 0.359 | 0.297 | 1 |
| Anks1b | 0.00243849 | 0.11371199 | 0.917 | 0.9 | 1 |
| Cct2 | 0.0024461 | 0.33018478 | 0.254 | 0.198 | 1 |
| Cdkal1 | 0.0024478 | 0.11287449 | 0.468 | 0.39 | 1 |
| Prickle1 | 0.00245536 | -0.5132278 | 0.659 | 0.627 | 1 |
| Eif2s2 | 0.00245955 | 0.26290226 | 0.386 | 0.319 | 1 |
| St8sia1 | 0.00246037 | 0.20177206 | 0.315 | 0.257 | 1 |
| Tom1l2 | 0.00246339 | 0.13085005 | 0.535 | 0.451 | 1 |
| Vdac1 | 0.0024784 | 0.13546684 | 0.443 | 0.369 | 1 |
| Fbrsl1 | 0.00251005 | 0.22207024 | 0.308 | 0.251 | 1 |
| Jak1 | 0.00253693 | 0.29226644 | 0.442 | 0.382 | 1 |
| Hspa12a | 0.00254621 | 0.15252699 | 0.347 | 0.285 | 1 |
| Specc1 | 0.00256673 | 0.15297516 | 0.615 | 0.509 | 1 |
| Sf3b3 | 0.00256775 | 0.23043134 | 0.333 | 0.275 | 1 |
| Rnf187 | 0.00258835 | 0.41306139 | 0.255 | 0.196 | 1 |
| Lrpap1 | 0.00258858 | 0.12165132 | 0.308 | 0.249 | 1 |
| Pkia | 0.00261102 | 0.21544717 | 0.441 | 0.375 | 1 |
| Ube2e2 | 0.00263137 | -0.2414427 | 0.819 | 0.813 | 1 |
| Atp11b | 0.00263567 | 0.14088826 | 0.452 | 0.383 | 1 |
| Mgrn1 | 0.0026363 | 0.19260827 | 0.323 | 0.264 | 1 |
| Eprs | 0.00264351 | 0.12627147 | 0.311 | 0.251 | 1 |
| Matk | 0.00264812 | 0.14937134 | 0.524 | 0.441 | 1 |
| A630089N07 | 0.00267154 | 0.59460019 | 0.256 | 0.205 | 1 |
| Adgrl1 | 0.00269134 | 0.15007977 | 0.48 | 0.402 | 1 |
| Eefsec | 0.00272167 | 0.32137813 | 0.306 | 0.251 | 1 |
| Gm49353 | 0.00274125 | 0.21895906 | 0.364 | 0.306 | 1 |
| Ctnna2 | 0.00274305 | 0.13046747 | 0.883 | 0.839 | 1 |
| Atpif1 | 0.00274409 | 0.22680935 | 0.471 | 0.41 | 1 |
| Cenpc1 | 0.00275468 | 0.27084219 | 0.321 | 0.265 | 1 |
| Dlat | 0.00276408 | 0.10714338 | 0.367 | 0.303 | 1 |
| Rph3a | 0.00277371 | 0.20757616 | 0.296 | 0.233 | 1 |
| Trit1 | 0.00278046 | 0.26332933 | 0.27 | 0.218 | 1 |
| Fam13c | 0.00284793 | 0.14497308 | 0.491 | 0.41 | 1 |

| | | | | | |
|---|---|---|---|---|---|
| Prdm8 | 0.00288563 | 0.32696868 | 0.332 | 0.272 | 1 |
| Ube2l3 | 0.00288728 | 0.2511172 | 0.302 | 0.244 | 1 |
| Rps15a | 0.00289003 | 0.29858596 | 0.426 | 0.366 | 1 |
| Chpf2 | 0.00289514 | 0.361671 | 0.281 | 0.231 | 1 |
| Iqcb1 | 0.00290318 | 0.30656479 | 0.314 | 0.257 | 1 |
| Hnrnph1 | 0.00291 | 0.13386387 | 0.452 | 0.381 | 1 |
| Dgki | 0.00294341 | -0.349684 | 0.707 | 0.679 | 1 |
| Rcor1 | 0.00297176 | 0.16444658 | 0.351 | 0.29 | 1 |
| Eri3 | 0.00297646 | 0.22747803 | 0.416 | 0.345 | 1 |
| Ube4b | 0.00298346 | 0.15947812 | 0.415 | 0.354 | 1 |
| Ubr1 | 0.00299635 | 0.20862876 | 0.379 | 0.318 | 1 |
| Ubac2 | 0.00302665 | 0.1815754 | 0.435 | 0.363 | 1 |
| Prkacb | 0.00303374 | -0.5613826 | 0.191 | 0.251 | 1 |
| Usp7 | 0.00303935 | 0.10840404 | 0.336 | 0.273 | 1 |
| Abcc4 | 0.00307304 | 0.39350448 | 0.296 | 0.243 | 1 |
| Atcay | 0.0031142 | 0.12549835 | 0.363 | 0.301 | 1 |
| Ppfia1 | 0.003133 | 0.23443069 | 0.291 | 0.235 | 1 |
| Rab11fip4 | 0.00315949 | 0.14174554 | 0.352 | 0.289 | 1 |
| Srek1 | 0.00320155 | 0.10545283 | 0.614 | 0.516 | 1 |
| Far2 | 0.00320854 | 0.21378636 | 0.273 | 0.218 | 1 |
| Atp6v0a1 | 0.0032168 | 0.12612705 | 0.701 | 0.597 | 1 |
| Glrx2 | 0.00323331 | 0.2084095 | 0.273 | 0.218 | 1 |
| Pnisr | 0.00324187 | 0.10710264 | 0.733 | 0.618 | 1 |
| Rab3gap2 | 0.00328197 | 0.33104926 | 0.332 | 0.28 | 1 |
| Pikfyve | 0.0033418 | 0.11139419 | 0.286 | 0.233 | 1 |
| Kif3a | 0.00336324 | 0.14880868 | 0.401 | 0.335 | 1 |
| Cbl | 0.00338987 | 0.21138475 | 0.383 | 0.326 | 1 |
| Srsf4 | 0.00340236 | 0.23881403 | 0.267 | 0.216 | 1 |
| Grip1 | 0.00347044 | 0.29828412 | 0.637 | 0.544 | 1 |
| Rnf150 | 0.00347411 | -0.4215565 | 0.454 | 0.478 | 1 |
| Srrt | 0.00351074 | 0.3498134 | 0.262 | 0.212 | 1 |
| Ttc19 | 0.00351705 | 0.10027039 | 0.591 | 0.481 | 1 |
| Comt | 0.0035444 | 0.11826514 | 0.338 | 0.278 | 1 |
| Tecpr1 | 0.00358517 | 0.36715278 | 0.349 | 0.295 | 1 |
| Sdf2 | 0.0035917 | 0.14618329 | 0.266 | 0.214 | 1 |
| Herc4 | 0.00360022 | 0.12550843 | 0.318 | 0.26 | 1 |
| Strada | 0.00364557 | 0.25890161 | 0.296 | 0.243 | 1 |
| Hagh | 0.00365395 | 0.36027515 | 0.311 | 0.26 | 1 |
| Rplp2 | 0.00366831 | 0.22648114 | 0.473 | 0.406 | 1 |
| Rpl30 | 0.00367367 | 0.14973676 | 0.506 | 0.424 | 1 |
| Rims2 | 0.00368138 | 0.11668252 | 0.701 | 0.599 | 1 |
| Ddx39b | 0.00368869 | 0.15008194 | 0.383 | 0.322 | 1 |
| N4bp2l1 | 0.00372319 | 0.30472728 | 0.271 | 0.217 | 1 |

| | | | | | |
|---|---|---|---|---|---|
| Ubr3 | 0.00372543 | 0.10663601 | 0.64 | 0.545 | 1 |
| Lztfl1 | 0.00374723 | 0.28226041 | 0.392 | 0.335 | 1 |
| Plxnc1 | 0.003773 | 0.59585529 | 0.265 | 0.218 | 1 |
| Dot1l | 0.00378932 | 0.17168915 | 0.387 | 0.33 | 1 |
| Smap1 | 0.00380272 | 0.10834746 | 0.442 | 0.376 | 1 |
| Tnfrsf19 | 0.00382556 | 0.40850502 | 0.266 | 0.215 | 1 |
| Faim2 | 0.00383077 | 0.10113694 | 0.329 | 0.265 | 1 |
| Rps11 | 0.00388357 | 0.12270246 | 0.4 | 0.329 | 1 |
| Macrod2 | 0.00392372 | 0.14722003 | 0.883 | 0.837 | 1 |
| Smg1 | 0.00392825 | 0.14663294 | 0.539 | 0.459 | 1 |
| Tbce | 0.0039427 | 0.19635448 | 0.36 | 0.302 | 1 |
| Arid2 | 0.00394758 | 0.25779983 | 0.317 | 0.261 | 1 |
| Uba2 | 0.00396967 | 0.22393549 | 0.266 | 0.209 | 1 |
| B230307C23 | 0.00397095 | 0.19865203 | 0.269 | 0.217 | 1 |
| Klhl7 | 0.00397183 | -0.7512918 | 0.352 | 0.397 | 1 |
| Fam49a | 0.00399243 | 0.19284997 | 0.467 | 0.401 | 1 |
| Kif5c | 0.00400574 | 0.12557912 | 0.666 | 0.584 | 1 |
| Pgk1 | 0.00402025 | 0.16230587 | 0.295 | 0.234 | 1 |
| Zfp799 | 0.00403447 | 0.33091028 | 0.277 | 0.229 | 1 |
| Gpr107 | 0.004046 | 0.38339945 | 0.274 | 0.227 | 1 |
| Chfr | 0.0040549 | 0.11628654 | 0.423 | 0.355 | 1 |
| Mdga1 | 0.00406051 | 0.18134361 | 0.379 | 0.316 | 1 |
| Kansl1l | 0.00411101 | 0.11426699 | 0.598 | 0.502 | 1 |
| Camk2g | 0.00420481 | 0.12073514 | 0.252 | 0.194 | 1 |
| Ndufaf7 | 0.00423406 | 0.18543599 | 0.296 | 0.247 | 1 |
| Rbm3 | 0.00431422 | -0.6089391 | 0.224 | 0.286 | 1 |
| Ap2b1 | 0.00434346 | 0.13022604 | 0.559 | 0.476 | 1 |
| Cox7a2l | 0.00435789 | 0.17053668 | 0.317 | 0.258 | 1 |
| Ndufs3 | 0.0043752 | 0.35466172 | 0.333 | 0.28 | 1 |
| Rnf38 | 0.00442499 | 0.14446114 | 0.321 | 0.263 | 1 |
| Grin2a | 0.00444533 | -0.1522557 | 0.879 | 0.898 | 1 |
| Dctn1 | 0.00448227 | -0.9578351 | 0.285 | 0.337 | 1 |
| Zbtb11 | 0.00450002 | 0.1192235 | 0.409 | 0.343 | 1 |
| Nek1 | 0.00450151 | 0.18758311 | 0.392 | 0.331 | 1 |
| Zcchc7 | 0.00452448 | 0.1338721 | 0.746 | 0.653 | 1 |
| Desi1 | 0.00454985 | 0.16917225 | 0.442 | 0.374 | 1 |
| Lrfn5 | 0.00455454 | -0.1613111 | 0.784 | 0.803 | 1 |
| Nebl | 0.00455793 | 0.14739673 | 0.772 | 0.677 | 1 |
| Arnt | 0.00461203 | -0.696226 | 0.266 | 0.325 | 1 |
| Cul2 | 0.00461471 | 0.19695501 | 0.291 | 0.238 | 1 |
| Asph | 0.00465411 | 0.13113619 | 0.558 | 0.458 | 1 |
| Dcun1d2 | 0.0046542 | 0.22656791 | 0.284 | 0.234 | 1 |
| Cep78 | 0.00466485 | 0.22153014 | 0.288 | 0.232 | 1 |

| | | | | | |
|---|---|---|---|---|---|
| Foxn3 | 0.00466787 | -0.6694832 | 0.299 | 0.352 | 1 |
| Dop1a | 0.00467335 | 0.20126167 | 0.409 | 0.348 | 1 |
| Sms | 0.00471172 | 0.17044212 | 0.457 | 0.384 | 1 |
| Polr3h | 0.0047159 | 0.47168863 | 0.347 | 0.29 | 1 |
| Ccdc85a | 0.00473203 | -0.4379321 | 0.523 | 0.557 | 1 |
| Slc35f1 | 0.00476996 | 0.13069351 | 0.417 | 0.347 | 1 |
| Lrrc19 | 0.00478432 | 0.30755833 | 0.254 | 0.206 | 1 |
| Eml2 | 0.0048592 | 0.35609631 | 0.33 | 0.282 | 1 |
| Macroh2a1 | 0.00486534 | 0.16064306 | 0.296 | 0.24 | 1 |
| Meis2 | 0.00490679 | -0.3798364 | 0.22 | 0.278 | 1 |
| Lmtk2 | 0.00491706 | 0.15642716 | 0.405 | 0.344 | 1 |
| Kcng2 | 0.00495321 | 0.16759054 | 0.274 | 0.221 | 1 |
| Sorcs2 | 0.00495802 | 0.23927927 | 0.431 | 0.353 | 1 |
| Cipc | 0.00496755 | 0.18990791 | 0.276 | 0.223 | 1 |
| Khdrbs1 | 0.00499864 | 0.25875336 | 0.356 | 0.299 | 1 |
| Rsbn1 | 0.00500345 | 0.23763646 | 0.297 | 0.245 | 1 |
| 2610507B11 | 0.00500673 | 0.27047362 | 0.312 | 0.261 | 1 |
| Rcor3 | 0.00505431 | 0.45224234 | 0.353 | 0.3 | 1 |
| Rere | 0.00506828 | 0.11908787 | 0.801 | 0.734 | 1 |
| Glud1 | 0.00506952 | 0.33691483 | 0.302 | 0.253 | 1 |
| Cab39l | 0.0050805 | 0.37892097 | 0.289 | 0.239 | 1 |
| Ppp6r3 | 0.00514548 | 0.14285647 | 0.432 | 0.37 | 1 |
| Ythdc1 | 0.00516957 | 0.28784299 | 0.366 | 0.315 | 1 |
| Ssb | 0.00519351 | -0.1057542 | 0.362 | 0.3 | 1 |
| Mpp3 | 0.00526308 | 0.23222676 | 0.254 | 0.207 | 1 |
| Bod1l | 0.00529882 | 0.32627394 | 0.353 | 0.301 | 1 |
| Lgr4 | 0.00530195 | 0.12118502 | 0.278 | 0.228 | 1 |
| Phka2 | 0.00531687 | 0.21102606 | 0.332 | 0.275 | 1 |
| Brinp1 | 0.00535624 | -0.2824833 | 0.749 | 0.728 | 1 |
| Tmem167 | 0.00543152 | 0.27134267 | 0.367 | 0.313 | 1 |
| Slitrk5 | 0.00543415 | 0.21181655 | 0.341 | 0.286 | 1 |
| Tank | 0.0054671 | 0.25994785 | 0.256 | 0.207 | 1 |
| Uhrf2 | 0.00547063 | 0.1718611 | 0.355 | 0.3 | 1 |
| Kcnj3 | 0.00550709 | 0.15252147 | 0.704 | 0.62 | 1 |
| Tiam2 | 0.00551868 | 0.30225395 | 0.269 | 0.219 | 1 |
| Cltb | 0.00559872 | 0.3970314 | 0.277 | 0.228 | 1 |
| Gfm2 | 0.00560383 | 0.26683748 | 0.353 | 0.297 | 1 |
| Celf2 | 0.0056438 | -0.131195 | 0.947 | 0.957 | 1 |
| Rtl1 | 0.00565061 | 0.37694849 | 0.265 | 0.218 | 1 |
| Chic1 | 0.00570106 | 0.19309958 | 0.326 | 0.271 | 1 |
| Fbxo10 | 0.00580397 | 0.52144897 | 0.262 | 0.213 | 1 |
| Entpd6 | 0.00584868 | 0.11846244 | 0.289 | 0.241 | 1 |
| Abl1 | 0.00586686 | 0.17581422 | 0.256 | 0.206 | 1 |

| | | | | | |
|---|---|---|---|---|---|
| Copg1 | 0.00594694 | 0.13656283 | 0.278 | 0.221 | 1 |
| Trappc9 | 0.00595625 | 0.1110493 | 0.486 | 0.406 | 1 |
| Pcdh1 | 0.00596324 | 0.11690098 | 0.364 | 0.306 | 1 |
| Rpn2 | 0.00596894 | 0.24600642 | 0.255 | 0.205 | 1 |
| Reps1 | 0.00598327 | 0.18827304 | 0.366 | 0.314 | 1 |
| Ube2o | 0.00605644 | 0.1750244 | 0.262 | 0.213 | 1 |
| Rsrc1 | 0.00608571 | 0.13188279 | 0.452 | 0.382 | 1 |
| Actr3b | 0.00618239 | -0.1127037 | 0.477 | 0.408 | 1 |
| Ndufa2 | 0.00630817 | 0.10649146 | 0.265 | 0.207 | 1 |
| Tnrc18 | 0.00638535 | 0.16184859 | 0.355 | 0.299 | 1 |
| Fam118b | 0.00640099 | 0.16167796 | 0.273 | 0.223 | 1 |
| Ralgapa1 | 0.00648769 | 0.14078096 | 0.724 | 0.64 | 1 |
| Scn8a | 0.00652092 | 0.11396236 | 0.724 | 0.625 | 1 |
| Slc39a10 | 0.00653258 | 0.44879359 | 0.274 | 0.226 | 1 |
| Agfg1 | 0.0065339 | 0.14385964 | 0.337 | 0.283 | 1 |
| Map7d1 | 0.00653658 | 0.16163378 | 0.315 | 0.262 | 1 |
| Arel1 | 0.00656471 | 0.14938029 | 0.334 | 0.284 | 1 |
| Gapdh | 0.00658457 | 0.19238566 | 0.607 | 0.533 | 1 |
| Tdrd3 | 0.00661274 | 0.220031 | 0.293 | 0.244 | 1 |
| B3galt1 | 0.00667446 | -0.3334132 | 0.764 | 0.734 | 1 |
| Sec24a | 0.0066861 | 0.29867941 | 0.329 | 0.28 | 1 |
| Nectin3 | 0.00673118 | 0.14497598 | 0.357 | 0.302 | 1 |
| Acsl1 | 0.00673692 | 0.39848427 | 0.266 | 0.222 | 1 |
| Plaa | 0.00674927 | 0.2190736 | 0.285 | 0.239 | 1 |
| Mcph1 | 0.0068743 | 0.18482732 | 0.319 | 0.267 | 1 |
| Slc12a2 | 0.0068855 | -0.2181112 | 0.291 | 0.229 | 1 |
| Prkaa2 | 0.00689601 | 0.31166775 | 0.251 | 0.203 | 1 |
| Orai2 | 0.00694253 | 0.2366275 | 0.417 | 0.354 | 1 |
| Capza1 | 0.00702976 | 0.3419757 | 0.314 | 0.264 | 1 |
| D430042O09 | 0.00704861 | 0.36601695 | 0.291 | 0.248 | 1 |
| Nfat5 | 0.00709209 | 0.10886352 | 0.615 | 0.524 | 1 |
| Brsk2 | 0.00719232 | 0.18683323 | 0.295 | 0.244 | 1 |
| Zswim8 | 0.00727716 | 0.34680574 | 0.256 | 0.209 | 1 |
| Zfp398 | 0.00728307 | 0.22108745 | 0.344 | 0.289 | 1 |
| 0610010F05I | 0.00734287 | 0.24912183 | 0.336 | 0.288 | 1 |
| Adgrb2 | 0.00735372 | 0.11636212 | 0.52 | 0.445 | 1 |
| mt-Atp6 | 0.00737918 | -0.5289971 | 0.921 | 0.857 | 1 |
| Zkscan3 | 0.00739437 | 0.17144999 | 0.282 | 0.232 | 1 |
| Ndufa10 | 0.00768667 | 0.30584125 | 0.325 | 0.279 | 1 |
| Nr3c2 | 0.00775387 | 0.12817927 | 0.789 | 0.721 | 1 |
| Ndufb1-ps | 0.00782027 | 0.17711091 | 0.393 | 0.336 | 1 |
| Scfd2 | 0.00784549 | 0.12353217 | 0.36 | 0.305 | 1 |
| Rgs7 | 0.00785168 | 0.13320829 | 0.771 | 0.682 | 1 |

| | | | | | |
|---|---|---|---|---|---|
| Caly | 0.00787916 | 0.10575258 | 0.415 | 0.356 | 1 |
| Phf12 | 0.00796694 | 0.33940938 | 0.317 | 0.268 | 1 |
| Csad | 0.00797167 | 0.23469893 | 0.271 | 0.223 | 1 |
| Cct6a | 0.00806639 | 0.36397212 | 0.269 | 0.219 | 1 |
| Preb | 0.00810295 | 0.40063229 | 0.265 | 0.219 | 1 |
| Zfp771 | 0.00812215 | 0.134401 | 0.256 | 0.21 | 1 |
| Coro1c | 0.00813672 | 0.35427078 | 0.297 | 0.254 | 1 |
| Bcl9 | 0.00814152 | 0.21217095 | 0.411 | 0.353 | 1 |
| Reps2 | 0.00826192 | 0.11380779 | 0.577 | 0.484 | 1 |
| Acaca | 0.00829858 | 0.11145291 | 0.424 | 0.365 | 1 |
| Rps10 | 0.00834498 | 0.18447093 | 0.344 | 0.29 | 1 |
| Nsd3 | 0.00835151 | 0.10787367 | 0.577 | 0.495 | 1 |
| Slc39a11 | 0.00841596 | 0.12669627 | 0.311 | 0.259 | 1 |
| Baz2a | 0.00848949 | 0.15057409 | 0.37 | 0.316 | 1 |
| Gabarapl1 | 0.00855656 | 0.24868987 | 0.329 | 0.272 | 1 |
| Kmt2c | 0.00858142 | 0.10435039 | 0.606 | 0.523 | 1 |
| Frmd6 | 0.00858348 | 0.26638483 | 0.267 | 0.218 | 1 |
| Cdk5r1 | 0.00861914 | 0.16619456 | 0.262 | 0.21 | 1 |
| Syp | 0.00864443 | 0.20312623 | 0.509 | 0.448 | 1 |
| Camk2b | 0.00865158 | -0.238441 | 0.761 | 0.718 | 1 |
| Mif | 0.00869619 | 0.13404022 | 0.351 | 0.286 | 1 |
| Zdhhc21 | 0.00869999 | -0.5728234 | 0.259 | 0.314 | 1 |
| Atxn7l3b | 0.00873308 | 0.22792479 | 0.274 | 0.222 | 1 |
| Ly6e | 0.0088102 | 0.19241751 | 0.368 | 0.307 | 1 |
| Nploc4 | 0.00886962 | 0.19919849 | 0.277 | 0.226 | 1 |
| Phtf2 | 0.00894218 | 0.12654879 | 0.289 | 0.241 | 1 |
| Smpd3 | 0.00895851 | 0.14455731 | 0.547 | 0.47 | 1 |
| Naa20 | 0.00901772 | 0.31392558 | 0.299 | 0.25 | 1 |
| Gripap1 | 0.00903568 | 0.17784101 | 0.336 | 0.283 | 1 |
| Zfp827 | 0.00905622 | 0.12624861 | 0.367 | 0.306 | 1 |
| Mipep | 0.00912803 | 0.22509069 | 0.263 | 0.219 | 1 |
| Cop1 | 0.00918261 | 0.14506413 | 0.409 | 0.356 | 1 |
| Clasp1 | 0.00921632 | 0.124446 | 0.475 | 0.409 | 1 |
| Atp5pb | 0.00922641 | 0.13012027 | 0.498 | 0.428 | 1 |
| Dusp3 | 0.00936139 | 0.23595356 | 0.269 | 0.223 | 1 |
| Sugp2 | 0.00964672 | 0.23282755 | 0.336 | 0.284 | 1 |
| Pik3c2a | 0.0096665 | 0.19634922 | 0.385 | 0.335 | 1 |
| Dixdc1 | 0.00966684 | -0.163267 | 0.317 | 0.259 | 1 |
| Homer2 | 0.00968009 | 0.13276303 | 0.445 | 0.374 | 1 |
| Trip11 | 0.0096969 | 0.12037709 | 0.441 | 0.375 | 1 |
| Tnr | 0.00974185 | 0.20192469 | 0.709 | 0.616 | 1 |
| Chchd10 | 0.00975689 | 0.15628913 | 0.401 | 0.338 | 1 |
| Nfix | 0.00977024 | -0.551503 | 0.464 | 0.488 | 1 |

| | | | | | |
|---|---|---|---|---|---|
| Mfhas1 | 0.00977459 | 0.16251733 | 0.413 | 0.356 | 1 |
| Spock1 | 0.00977501 | 0.16967364 | 0.557 | 0.501 | 1 |
| Hspa5 | 0.00980575 | -0.5941116 | 0.344 | 0.399 | 1 |
| Ago2 | 0.00989009 | 0.14889477 | 0.33 | 0.279 | 1 |
| Rpl27 | 0.00999835 | 0.38497068 | 0.273 | 0.227 | 1 |
| Ptma | 0.01003682 | -0.5284203 | 0.411 | 0.457 | 1 |
| Trrap | 0.01006806 | 0.26862566 | 0.353 | 0.298 | 1 |
| Ano6 | 0.01007548 | 0.10442174 | 0.404 | 0.343 | 1 |
| Uchl1 | 0.01017787 | 0.15834172 | 0.553 | 0.484 | 1 |
| Hnrnph3 | 0.01018426 | 0.192792 | 0.326 | 0.28 | 1 |
| Herc3 | 0.01020037 | 0.10323313 | 0.473 | 0.414 | 1 |
| Cdk11b | 0.01032008 | 0.20414882 | 0.322 | 0.278 | 1 |
| Rpl31 | 0.01032934 | 0.25075429 | 0.252 | 0.201 | 1 |
| Tnpo3 | 0.01032952 | 0.1721385 | 0.398 | 0.344 | 1 |
| Swi5 | 0.0104112 | 0.15524871 | 0.263 | 0.213 | 1 |
| Nap1l1 | 0.01043315 | 0.17494497 | 0.363 | 0.311 | 1 |
| Psmd9 | 0.01045194 | 0.23336238 | 0.258 | 0.213 | 1 |
| Cep126 | 0.0104759 | 0.35479745 | 0.262 | 0.216 | 1 |
| Kantr | 0.01054679 | 0.31793373 | 0.263 | 0.216 | 1 |
| Ablim1 | 0.01058874 | 0.12657975 | 0.634 | 0.534 | 1 |
| Slc25a36 | 0.01060923 | 0.12283129 | 0.353 | 0.305 | 1 |
| Robo2 | 0.01065862 | -0.1731221 | 0.808 | 0.745 | 1 |
| Mef2d | 0.010703 | 0.11938178 | 0.363 | 0.31 | 1 |
| Pdlim5 | 0.01076293 | 0.59921119 | 0.286 | 0.243 | 1 |
| Rpl8 | 0.0109736 | 0.13424391 | 0.438 | 0.375 | 1 |
| Gpbp1 | 0.01104269 | 0.29965 | 0.259 | 0.216 | 1 |
| Cab39 | 0.01117281 | 0.11059859 | 0.379 | 0.324 | 1 |
| Gak | 0.01117352 | 0.13627384 | 0.277 | 0.231 | 1 |
| Lrp1b | 0.01130958 | 0.14809111 | 0.898 | 0.909 | 1 |
| Ap1ar | 0.01133732 | 0.25493124 | 0.251 | 0.208 | 1 |
| Eid1 | 0.01134025 | 0.10081816 | 0.432 | 0.371 | 1 |
| Mpv17l | 0.01135732 | 0.10207487 | 0.289 | 0.239 | 1 |
| Rpl11 | 0.01156446 | 0.30082617 | 0.476 | 0.424 | 1 |
| Pkn2 | 0.01159515 | 0.10190122 | 0.402 | 0.349 | 1 |
| Lrrc49 | 0.01160158 | 0.12446559 | 0.293 | 0.248 | 1 |
| Setd4 | 0.01183865 | 0.20253739 | 0.3 | 0.255 | 1 |
| Fntb | 0.01186776 | 0.22951618 | 0.296 | 0.251 | 1 |
| Por | 0.01190766 | 0.42557388 | 0.251 | 0.209 | 1 |
| Arhgef4 | 0.01191549 | 0.13499284 | 0.368 | 0.316 | 1 |
| Abhd18 | 0.01194903 | -0.1161875 | 0.417 | 0.345 | 1 |
| Gspt1 | 0.0120953 | 0.12008126 | 0.299 | 0.251 | 1 |
| Ttyh1 | 0.0121416 | 0.18652839 | 0.277 | 0.229 | 1 |
| Als2 | 0.01218683 | 0.48451198 | 0.259 | 0.217 | 1 |

| | | | | | |
|---|---|---|---|---|---|
| Rpl37a | 0.01228671 | 0.13910431 | 0.48 | 0.424 | 1 |
| Erp29 | 0.01228945 | 0.19283569 | 0.289 | 0.239 | 1 |
| Txnl1 | 0.01230538 | 0.244524 | 0.288 | 0.243 | 1 |
| Yme1l1 | 0.01233003 | 0.14518382 | 0.413 | 0.366 | 1 |
| Dnajc5 | 0.01240193 | 0.22037988 | 0.33 | 0.283 | 1 |
| Narf | 0.01244806 | 0.35258144 | 0.252 | 0.211 | 1 |
| Ndufa13 | 0.01253344 | 0.26460463 | 0.426 | 0.375 | 1 |
| Stx3 | 0.0125729 | 0.20227222 | 0.36 | 0.312 | 1 |
| Cdh10 | 0.01263781 | -0.2148696 | 0.477 | 0.511 | 1 |
| Luzp2 | 0.01265705 | -0.6578223 | 0.554 | 0.51 | 1 |
| Galnt11 | 0.01285909 | -0.5934545 | 0.314 | 0.365 | 1 |
| Fbxw8 | 0.01293922 | 0.24542443 | 0.286 | 0.243 | 1 |
| Smoc2 | 0.01297996 | 0.14770899 | 0.326 | 0.275 | 1 |
| Lrrc7 | 0.01310935 | -0.122441 | 0.805 | 0.782 | 1 |
| Smdt1 | 0.01312417 | 0.13416512 | 0.265 | 0.215 | 1 |
| Dzip1 | 0.01331594 | 0.23705718 | 0.261 | 0.216 | 1 |
| Cask | 0.01332219 | 0.14969037 | 0.435 | 0.372 | 1 |
| Cachd1 | 0.0133352 | -0.5333478 | 0.326 | 0.369 | 1 |
| Lnpk | 0.01352286 | 0.19714549 | 0.364 | 0.309 | 1 |
| Lamp1 | 0.01353123 | 0.21495393 | 0.271 | 0.224 | 1 |
| Psma1 | 0.01358193 | 0.11626455 | 0.33 | 0.283 | 1 |
| Arpc5 | 0.01359312 | 0.16724286 | 0.353 | 0.307 | 1 |
| Map3k7 | 0.01373008 | -0.7453261 | 0.211 | 0.261 | 1 |
| Uqcrq | 0.013747 | 0.16477109 | 0.37 | 0.313 | 1 |
| Chd4 | 0.01389668 | 0.15783356 | 0.51 | 0.445 | 1 |
| Grina | 0.01390653 | 0.15615105 | 0.488 | 0.427 | 1 |
| Pde10a | 0.01403309 | -0.3727203 | 0.606 | 0.591 | 1 |
| Mt1 | 0.01406496 | -0.3316105 | 0.454 | 0.5 | 1 |
| Xpo6 | 0.01407264 | 0.30979025 | 0.296 | 0.259 | 1 |
| Psmd7 | 0.01410807 | 0.29053851 | 0.297 | 0.262 | 1 |
| Zeb2 | 0.01415571 | -0.1649656 | 0.843 | 0.825 | 1 |
| Uggt1 | 0.01418779 | 0.24557682 | 0.263 | 0.224 | 1 |
| Vps37a | 0.01430388 | 0.20725295 | 0.274 | 0.234 | 1 |
| Naaladl2 | 0.01434672 | 0.24096209 | 0.379 | 0.313 | 1 |
| Pak1 | 0.01453068 | -0.4773808 | 0.311 | 0.362 | 1 |
| Ncoa5 | 0.01467325 | 0.31450207 | 0.344 | 0.302 | 1 |
| Fbxo41 | 0.01468102 | 0.15560377 | 0.312 | 0.274 | 1 |
| Ncbp1 | 0.01469661 | 0.10273041 | 0.254 | 0.211 | 1 |
| Rps24 | 0.01493516 | 0.15351499 | 0.577 | 0.511 | 1 |
| Thoc7 | 0.01497614 | 0.11051828 | 0.315 | 0.27 | 1 |
| Wapl | 0.01525163 | -0.1011473 | 0.461 | 0.395 | 1 |
| Dpp6 | 0.01555588 | -0.1391117 | 0.793 | 0.744 | 1 |
| Dtnbp1 | 0.01556077 | 0.2830287 | 0.258 | 0.215 | 1 |

| | | | | | |
|---|---|---|---|---|---|
| Arl8b | 0.01560647 | 0.264469 | 0.269 | 0.224 | 1 |
| Tbc1d14 | 0.0157523 | 0.18929918 | 0.291 | 0.244 | 1 |
| Dnajc18 | 0.01609909 | 0.20881917 | 0.269 | 0.228 | 1 |
| Cdh18 | 0.0162073 | 0.2546534 | 0.321 | 0.269 | 1 |
| Sgip1 | 0.01624118 | 0.10244394 | 0.809 | 0.706 | 1 |
| mt-Nd4 | 0.01629001 | -0.5935615 | 0.704 | 0.679 | 1 |
| Lmtk3 | 0.01629416 | 0.33592452 | 0.3 | 0.26 | 1 |
| Sod1 | 0.01631074 | 0.18106627 | 0.251 | 0.203 | 1 |
| Fgf12 | 0.01631338 | 0.12040838 | 0.756 | 0.681 | 1 |
| Ppp1r10 | 0.01636994 | 0.28148571 | 0.252 | 0.211 | 1 |
| Atp5g3 | 0.0163732 | 0.18622867 | 0.402 | 0.35 | 1 |
| Cacna2d3 | 0.0164639 | 0.10549791 | 0.806 | 0.721 | 1 |
| Matr3 | 0.01657293 | 0.12108867 | 0.427 | 0.372 | 1 |
| Prepl | 0.01675042 | -0.1009431 | 0.326 | 0.275 | 1 |
| Magi2 | 0.01699476 | 0.11386362 | 0.925 | 0.933 | 1 |
| Xpa | 0.01703681 | 0.25898327 | 0.259 | 0.219 | 1 |
| Kcnk10 | 0.01714036 | -0.2820273 | 0.265 | 0.215 | 1 |
| Echdc1 | 0.01714857 | 0.12107939 | 0.307 | 0.261 | 1 |
| Plch2 | 0.01717251 | 0.28609496 | 0.366 | 0.314 | 1 |
| Vamp4 | 0.01722894 | 0.21677606 | 0.273 | 0.232 | 1 |
| Pnpt1 | 0.0173272 | 0.19955665 | 0.312 | 0.269 | 1 |
| Myh10 | 0.01737854 | -0.1199084 | 0.408 | 0.352 | 1 |
| Atxn2l | 0.01743991 | 0.1389717 | 0.359 | 0.308 | 1 |
| Zfhx2 | 0.01764002 | 0.13600717 | 0.352 | 0.307 | 1 |
| Foxk2 | 0.0177271 | 0.15908523 | 0.28 | 0.239 | 1 |
| Tbk1 | 0.01796516 | 0.26880932 | 0.28 | 0.237 | 1 |
| Zbtb16 | 0.0183825 | 0.11781658 | 0.604 | 0.493 | 1 |
| Nbr1 | 0.01843927 | 0.12749168 | 0.318 | 0.274 | 1 |
| Supt6 | 0.01844649 | 0.14420422 | 0.386 | 0.333 | 1 |
| Jpt1 | 0.01851106 | 0.22721462 | 0.25 | 0.208 | 1 |
| Sult4a1 | 0.01856387 | 0.26876436 | 0.258 | 0.216 | 1 |
| Utp14a | 0.01858744 | 0.15150052 | 0.27 | 0.228 | 1 |
| Dctn4 | 0.01859725 | -0.1193774 | 0.411 | 0.358 | 1 |
| Phf6 | 0.01861748 | 0.13827535 | 0.303 | 0.261 | 1 |
| Lrrtm1 | 0.01880674 | 0.19502075 | 0.297 | 0.252 | 1 |
| Rnf169 | 0.01893816 | -0.5781278 | 0.362 | 0.401 | 1 |
| BC005537 | 0.01896095 | 0.31517667 | 0.352 | 0.306 | 1 |
| Mapkap1 | 0.01908892 | 0.12909457 | 0.426 | 0.369 | 1 |
| Pkd2 | 0.0191078 | 0.14746096 | 0.263 | 0.221 | 1 |
| Galnt17 | 0.01918514 | -0.1558649 | 0.786 | 0.774 | 1 |
| Trpc4ap | 0.01918872 | 0.12938315 | 0.332 | 0.284 | 1 |
| Chp1 | 0.01924848 | 0.23945558 | 0.266 | 0.229 | 1 |
| Tuba1a | 0.01937199 | 0.14530756 | 0.505 | 0.444 | 1 |

| | | | | | |
|---|---|---|---|---|---|
| Rictor | 0.0194721 | -0.1106606 | 0.355 | 0.298 | 1 |
| Dnm2 | 0.01983825 | 0.10607622 | 0.426 | 0.369 | 1 |
| Prkdc | 0.01986878 | 0.24731338 | 0.303 | 0.26 | 1 |
| Synrg | 0.0199054 | 0.22298797 | 0.325 | 0.284 | 1 |
| Srgap3 | 0.02029651 | -0.1985636 | 0.782 | 0.7 | 1 |
| Bbof1 | 0.02045641 | 0.27402454 | 0.342 | 0.302 | 1 |
| Thsd7a | 0.02046225 | 0.19202924 | 0.37 | 0.319 | 1 |
| Pcdh20 | 0.02058793 | 0.14500597 | 0.27 | 0.226 | 1 |
| Kcnc1 | 0.02062942 | 0.12446989 | 0.311 | 0.262 | 1 |
| Ndufa11 | 0.02067514 | 0.16668955 | 0.259 | 0.214 | 1 |
| Orc3 | 0.02077197 | 0.13224727 | 0.308 | 0.261 | 1 |
| Sugp1 | 0.0208904 | 0.12724687 | 0.271 | 0.229 | 1 |
| Setd2 | 0.02100498 | 0.13115492 | 0.417 | 0.364 | 1 |
| Eif4g2 | 0.02103478 | -0.5470178 | 0.327 | 0.374 | 1 |
| Vdac3 | 0.02106795 | 0.21315582 | 0.274 | 0.233 | 1 |
| Med12l | 0.02110565 | 0.1205073 | 0.55 | 0.477 | 1 |
| Ndufv2 | 0.02136879 | 0.35377068 | 0.31 | 0.269 | 1 |
| Large1 | 0.02150347 | 0.1338441 | 0.767 | 0.721 | 1 |
| Gnai1 | 0.02151662 | 0.17214248 | 0.351 | 0.305 | 1 |
| Pid1 | 0.02157411 | 0.18677083 | 0.437 | 0.38 | 1 |
| Rapgef1 | 0.02168009 | 0.1256037 | 0.315 | 0.275 | 1 |
| Srpk1 | 0.02174908 | 0.16648553 | 0.3 | 0.261 | 1 |
| Taf1 | 0.02192859 | 0.13832601 | 0.322 | 0.277 | 1 |
| Ubn1 | 0.02197908 | 0.32428757 | 0.269 | 0.227 | 1 |
| Nrip1 | 0.02213714 | 0.20165304 | 0.326 | 0.282 | 1 |
| Ddhd1 | 0.02227469 | 0.25710958 | 0.366 | 0.323 | 1 |
| Mfsd4a | 0.02233628 | 0.13337805 | 0.389 | 0.343 | 1 |
| Grk3 | 0.02249762 | 0.30338894 | 0.303 | 0.261 | 1 |
| Sptbn2 | 0.02264312 | 0.15163982 | 0.488 | 0.432 | 1 |
| Stag2 | 0.02303644 | 0.12030173 | 0.344 | 0.301 | 1 |
| Add3 | 0.02358393 | 0.19308255 | 0.452 | 0.397 | 1 |
| Hspe1 | 0.02364309 | 0.22054949 | 0.289 | 0.246 | 1 |
| Lrrc8d | 0.02373765 | 0.19654369 | 0.374 | 0.325 | 1 |
| Nkain3 | 0.0239435 | -0.2523881 | 0.697 | 0.671 | 1 |
| Pnn | 0.02400066 | 0.10321993 | 0.435 | 0.384 | 1 |
| Ralyl | 0.02403621 | -0.1852792 | 0.734 | 0.707 | 1 |
| Eif2s3y | 0.02446183 | 0.37768613 | 0.302 | 0.263 | 1 |
| Ranbp10 | 0.02504065 | 0.29467296 | 0.267 | 0.228 | 1 |
| Vps53 | 0.02510582 | 0.14853724 | 0.278 | 0.236 | 1 |
| Pygo1 | 0.0252297 | 0.10432176 | 0.355 | 0.304 | 1 |
| Ankrd13c | 0.02524502 | 0.30544741 | 0.344 | 0.299 | 1 |
| Armh4 | 0.02528266 | 0.40246239 | 0.261 | 0.223 | 1 |
| Adap1 | 0.02529759 | 0.20123647 | 0.276 | 0.241 | 1 |

| | | | | | |
|---|---|---|---|---|---|
| Rpl4 | 0.02532852 | 0.22966501 | 0.262 | 0.217 | 1 |
| Polq | 0.02542118 | 0.28043203 | 0.258 | 0.216 | 1 |
| Mob4 | 0.0255151 | 0.13304272 | 0.307 | 0.265 | 1 |
| Adarb2 | 0.0256425 | -0.4226666 | 0.337 | 0.365 | 1 |
| Myrip | 0.02582554 | -0.113825 | 0.475 | 0.391 | 1 |
| Cap1 | 0.02606902 | 0.30232378 | 0.307 | 0.266 | 1 |
| Smim13 | 0.02616188 | 0.23875308 | 0.315 | 0.275 | 1 |
| Clstn1 | 0.02689308 | 0.11444881 | 0.544 | 0.489 | 1 |
| Dok6 | 0.02703877 | -0.2568917 | 0.729 | 0.677 | 1 |
| Ywhah | 0.02755331 | -0.2671492 | 0.595 | 0.612 | 1 |
| Rufy2 | 0.02766958 | 0.12468312 | 0.442 | 0.398 | 1 |
| Eno2 | 0.02781536 | 0.12778729 | 0.401 | 0.353 | 1 |
| 5730522E02I | 0.02784972 | -0.2283642 | 0.633 | 0.608 | 1 |
| Asap1 | 0.02828852 | 0.17432642 | 0.484 | 0.425 | 1 |
| Usf3 | 0.02851141 | 0.17875777 | 0.28 | 0.242 | 1 |
| Micu2 | 0.02887689 | 0.1478388 | 0.286 | 0.241 | 1 |
| Dcaf8 | 0.02898429 | 0.10133843 | 0.325 | 0.281 | 1 |
| Ipo8 | 0.02912373 | 0.15040232 | 0.261 | 0.224 | 1 |
| Chchd6 | 0.02913541 | 0.11728905 | 0.336 | 0.289 | 1 |
| Larp4 | 0.02913819 | 0.17490617 | 0.3 | 0.262 | 1 |
| Nfasc | 0.02935261 | 0.1107491 | 0.644 | 0.549 | 1 |
| Rundc3b | 0.02963308 | 0.19051625 | 0.347 | 0.307 | 1 |
| R3hdm4 | 0.02973942 | 0.15073913 | 0.314 | 0.271 | 1 |
| Dipk1a | 0.02995983 | -0.1382301 | 0.286 | 0.244 | 1 |
| Fstl4 | 0.03046317 | -0.4932655 | 0.314 | 0.355 | 1 |
| Thrap3 | 0.03085111 | 0.17429274 | 0.34 | 0.299 | 1 |
| Map6 | 0.03088247 | -0.5086779 | 0.281 | 0.324 | 1 |
| Rasgrf2 | 0.0308921 | 0.32177135 | 0.312 | 0.277 | 1 |
| Catspere2 | 0.03126879 | 0.10567455 | 0.371 | 0.324 | 1 |
| Sh3glb2 | 0.03137131 | 0.11826644 | 0.314 | 0.275 | 1 |
| Pabpn1 | 0.03161501 | 0.10746212 | 0.45 | 0.391 | 1 |
| Cntnap5a | 0.0320872 | -0.4378323 | 0.297 | 0.335 | 1 |
| Eif4b | 0.03216221 | 0.33537716 | 0.25 | 0.215 | 1 |
| Luc7l | 0.03241435 | 0.14751298 | 0.31 | 0.268 | 1 |
| Rnf111 | 0.03241788 | 0.21495115 | 0.362 | 0.322 | 1 |
| Psma2 | 0.03270754 | 0.13057252 | 0.277 | 0.235 | 1 |
| Epc1 | 0.03277452 | 0.17089957 | 0.437 | 0.394 | 1 |
| Itgbl1 | 0.03286487 | 0.10228414 | 0.284 | 0.242 | 1 |
| Pfdn5 | 0.03304548 | -0.4322818 | 0.252 | 0.297 | 1 |
| Lrp12 | 0.03312576 | 0.13731379 | 0.315 | 0.274 | 1 |
| Filip1l | 0.03336386 | 0.22039924 | 0.265 | 0.227 | 1 |
| Syngr1 | 0.03351802 | 0.27197852 | 0.28 | 0.244 | 1 |
| Armc8 | 0.03360765 | -0.1116652 | 0.359 | 0.31 | 1 |

| | | | | | |
|---|---|---|---|---|---|
| Ncs1 | 0.0336687 | 0.17007268 | 0.368 | 0.324 | 1 |
| Brdt | 0.03386461 | 0.15045528 | 0.285 | 0.247 | 1 |
| Prpf6 | 0.03393317 | 0.16379572 | 0.263 | 0.224 | 1 |
| Ybx1 | 0.03415051 | -0.3608 | 0.236 | 0.286 | 1 |
| Mtmr6 | 0.03433571 | 0.1715692 | 0.266 | 0.231 | 1 |
| Nefl | 0.03465879 | -0.4978401 | 0.293 | 0.336 | 1 |
| Myo9b | 0.03477188 | 0.24945657 | 0.317 | 0.273 | 1 |
| Arhgef11 | 0.03486618 | 0.12722695 | 0.404 | 0.355 | 1 |
| Fnip2 | 0.03490175 | 0.38922001 | 0.254 | 0.218 | 1 |
| Suclg1 | 0.03503905 | 0.19599729 | 0.336 | 0.297 | 1 |
| Parp6 | 0.0351475 | 0.13143535 | 0.304 | 0.266 | 1 |
| Ppp1r9b | 0.03533272 | 0.16574367 | 0.304 | 0.263 | 1 |
| Foxj3 | 0.03574376 | 0.107665 | 0.311 | 0.268 | 1 |
| Rpl14 | 0.03609975 | -0.3901218 | 0.25 | 0.295 | 1 |
| Eif4e | 0.03625973 | 0.15784488 | 0.296 | 0.258 | 1 |
| Prdx2 | 0.03629686 | 0.23851718 | 0.258 | 0.219 | 1 |
| Rpl22l1 | 0.03641677 | 0.10674626 | 0.349 | 0.305 | 1 |
| Ina | 0.0369043 | -0.5114059 | 0.244 | 0.291 | 1 |
| Vps35 | 0.03697722 | -0.1177626 | 0.307 | 0.259 | 1 |
| Tacc2 | 0.03705301 | 0.30115542 | 0.286 | 0.252 | 1 |
| Esco1 | 0.03747046 | 0.11407203 | 0.333 | 0.294 | 1 |
| Atr | 0.03771455 | -0.1096241 | 0.251 | 0.211 | 1 |
| Ate1 | 0.0379761 | 0.25678188 | 0.347 | 0.313 | 1 |
| Atp8a2 | 0.03843697 | 0.10032906 | 0.454 | 0.406 | 1 |
| Kcnh7 | 0.03844872 | -0.347409 | 0.476 | 0.499 | 1 |
| Stk39 | 0.03869664 | 0.13736786 | 0.456 | 0.398 | 1 |
| Grid2 | 0.03893077 | 0.18031318 | 0.482 | 0.42 | 1 |
| Serf2 | 0.03957232 | 0.11879887 | 0.296 | 0.253 | 1 |
| Rpl23a | 0.04001715 | 0.29045411 | 0.293 | 0.254 | 1 |
| Usp3 | 0.04012644 | 0.1158546 | 0.323 | 0.284 | 1 |
| Snx2 | 0.04019752 | 0.15871024 | 0.28 | 0.24 | 1 |
| Rps26 | 0.04068992 | 0.17957374 | 0.359 | 0.319 | 1 |
| Iqsec3 | 0.0413314 | -0.1213565 | 0.362 | 0.312 | 1 |
| Ubc | 0.04161243 | 0.19228371 | 0.326 | 0.286 | 1 |
| Zwint | 0.04199736 | 0.14836005 | 0.423 | 0.376 | 1 |
| Zmat4 | 0.04213948 | 0.17774058 | 0.263 | 0.22 | 1 |
| G3bp2 | 0.04216547 | -0.1154669 | 0.366 | 0.313 | 1 |
| Ndufa12 | 0.0426125 | -0.4936551 | 0.236 | 0.279 | 1 |
| Mgat5 | 0.04301497 | 0.22375851 | 0.411 | 0.363 | 1 |
| Zdhhc20 | 0.04331194 | -0.1299545 | 0.445 | 0.387 | 1 |
| Pde4b | 0.04370394 | -0.6035877 | 0.538 | 0.525 | 1 |
| Dnajb4 | 0.04452339 | -0.1886951 | 0.271 | 0.232 | 1 |
| Mlxip | 0.04476054 | 0.42750267 | 0.263 | 0.235 | 1 |

| | | | | | |
|---|---|---|---|---|---|
| Eef1a2 | 0.04503908 | 0.11572283 | 0.299 | 0.257 | 1 |
| Arfgef2 | 0.04510543 | 0.11303037 | 0.31 | 0.268 | 1 |
| Dstyk | 0.04548646 | 0.18022963 | 0.303 | 0.268 | 1 |
| Btbd1 | 0.04599099 | 0.14163962 | 0.338 | 0.299 | 1 |
| Ireb2 | 0.04602407 | 0.14434411 | 0.276 | 0.24 | 1 |
| Acsl3 | 0.04607152 | 0.27574288 | 0.271 | 0.236 | 1 |
| Pip5k1c | 0.04611024 | 0.11928677 | 0.285 | 0.246 | 1 |
| Zfp277 | 0.04628416 | 0.24283748 | 0.284 | 0.253 | 1 |
| Stk25 | 0.04666065 | -0.5842882 | 0.216 | 0.259 | 1 |
| Phkb | 0.04687112 | 0.19365042 | 0.263 | 0.226 | 1 |
| Micu1 | 0.04716327 | 0.13791817 | 0.416 | 0.374 | 1 |
| Bdp1 | 0.04732702 | -0.1187347 | 0.366 | 0.324 | 1 |
| Plpp3 | 0.04740545 | 0.44925557 | 0.271 | 0.239 | 1 |
| Azin1 | 0.0474462 | 0.14937522 | 0.278 | 0.239 | 1 |
| Ankfy1 | 0.04752863 | 0.20308782 | 0.299 | 0.262 | 1 |
| Sesn1 | 0.04757679 | -0.3440591 | 0.274 | 0.32 | 1 |
| Tmeff2 | 0.04790273 | -0.4648651 | 0.693 | 0.654 | 1 |
| Dip2a | 0.04850115 | 0.29411605 | 0.292 | 0.258 | 1 |
| Slc38a2 | 0.04850253 | 0.18571987 | 0.456 | 0.41 | 1 |
| Cnbp | 0.04860775 | 0.21922147 | 0.251 | 0.215 | 1 |
| C1ql3 | 0.04878602 | -0.4163022 | 0.295 | 0.326 | 1 |
| Ank3 | 0.04893831 | -0.1080411 | 0.917 | 0.923 | 1 |
| Calm2 | 0.04909197 | -0.1666426 | 0.899 | 0.842 | 1 |
| Cwf19l2 | 0.04971426 | 0.20447675 | 0.262 | 0.232 | 1 |
| Gpm6a | 0.04984624 | -0.1065482 | 0.924 | 0.87 | 1 |
| Armh3 | 0.05048741 | 0.24059127 | 0.352 | 0.325 | 1 |
| Cacna2d1 | 0.05093958 | -0.3544418 | 0.798 | 0.765 | 1 |
| Pdia6 | 0.0509412 | -0.5480794 | 0.232 | 0.276 | 1 |
| Ccnt1 | 0.05095805 | 0.22899656 | 0.282 | 0.248 | 1 |
| Cfl1 | 0.05110778 | 0.24570461 | 0.367 | 0.325 | 1 |
| Ggps1 | 0.05181686 | -0.6598306 | 0.228 | 0.268 | 1 |
| Vps54 | 0.05243532 | 0.16577103 | 0.344 | 0.303 | 1 |
| AW554918 | 0.05253906 | -0.1063324 | 0.362 | 0.312 | 1 |
| Cdc73 | 0.05267007 | -0.2171252 | 0.368 | 0.319 | 1 |
| Lmbrd1 | 0.05276505 | -0.1386794 | 0.366 | 0.32 | 1 |
| Tmem126a | 0.05276529 | 0.10486826 | 0.362 | 0.324 | 1 |
| Immt | 0.05297194 | -0.1307289 | 0.286 | 0.251 | 1 |
| Dctn2 | 0.05401844 | 0.23447933 | 0.276 | 0.245 | 1 |
| Gmcl1 | 0.05453049 | 0.15692107 | 0.372 | 0.324 | 1 |
| Pdpk1 | 0.05455916 | 0.13003027 | 0.407 | 0.373 | 1 |
| Aplp1 | 0.05482889 | -0.3901117 | 0.558 | 0.574 | 1 |
| Pcdh19 | 0.0560141 | -0.3571048 | 0.211 | 0.253 | 1 |
| Fubp1 | 0.05603369 | -0.1135229 | 0.558 | 0.479 | 1 |

| | | | | | |
|---|---|---|---|---|---|
| Ino80 | 0.05659586 | -0.1060675 | 0.404 | 0.357 | 1 |
| Slc22a23 | 0.05753252 | 0.25184237 | 0.31 | 0.28 | 1 |
| Pcyt2 | 0.05756147 | 0.1774162 | 0.325 | 0.286 | 1 |
| Lpgat1 | 0.05778668 | -0.1689988 | 0.389 | 0.341 | 1 |
| Ntsr2 | 0.05779747 | -0.4490053 | 0.217 | 0.252 | 1 |
| Cnot10 | 0.05799212 | 0.16378339 | 0.293 | 0.259 | 1 |
| Hk1 | 0.05819518 | 0.25630386 | 0.326 | 0.3 | 1 |
| Mast3 | 0.058397 | -0.3569446 | 0.379 | 0.419 | 1 |
| Kdm2a | 0.0589394 | -0.1403675 | 0.454 | 0.398 | 1 |
| Ppp1r9a | 0.05896132 | -0.4221125 | 0.6 | 0.596 | 1 |
| Slc9a1 | 0.05922602 | 0.14942417 | 0.286 | 0.253 | 1 |
| Got1 | 0.05956198 | 0.10325967 | 0.281 | 0.244 | 1 |
| Olfm1 | 0.05978391 | -0.2618389 | 0.738 | 0.728 | 1 |
| Adgrl3 | 0.06009578 | -0.2074589 | 0.776 | 0.775 | 1 |
| Il16 | 0.06020329 | 0.12441732 | 0.345 | 0.311 | 1 |
| Sfmbt1 | 0.06053684 | -0.1367866 | 0.314 | 0.271 | 1 |
| Rapgef4 | 0.06084458 | -0.3217635 | 0.633 | 0.559 | 1 |
| Cox10 | 0.06097545 | -0.1006059 | 0.28 | 0.242 | 1 |
| Psme4 | 0.0615416 | -0.130719 | 0.625 | 0.522 | 1 |
| Cnrip1 | 0.06179075 | -0.4866457 | 0.37 | 0.392 | 1 |
| Syne1 | 0.06188202 | -0.1783818 | 0.659 | 0.658 | 1 |
| Cnr1 | 0.06256869 | 0.54091159 | 0.295 | 0.26 | 1 |
| Rpl37 | 0.06315759 | 0.12921341 | 0.503 | 0.444 | 1 |
| Mrps33 | 0.06398993 | 0.10894345 | 0.288 | 0.25 | 1 |
| Trps1 | 0.06401253 | 0.45041847 | 0.542 | 0.507 | 1 |
| Glmn | 0.06440719 | 0.14814544 | 0.28 | 0.245 | 1 |
| Wdpcp | 0.06559231 | -0.1754609 | 0.302 | 0.261 | 1 |
| Kif13b | 0.06564642 | -0.2350482 | 0.327 | 0.288 | 1 |
| Hsbp1 | 0.06643315 | 0.14156066 | 0.267 | 0.231 | 1 |
| Lmbr1 | 0.06654991 | -0.1215231 | 0.317 | 0.277 | 1 |
| Pcf11 | 0.06660481 | 0.12021492 | 0.27 | 0.237 | 1 |
| Pgrmc1 | 0.06748829 | 0.11092599 | 0.352 | 0.312 | 1 |
| Rps6ka3 | 0.06783356 | 0.14656564 | 0.422 | 0.38 | 1 |
| Rps2 | 0.06788357 | -0.3435715 | 0.392 | 0.43 | 1 |
| Acbd5 | 0.06799507 | 0.16979086 | 0.306 | 0.277 | 1 |
| Dhdds | 0.06827499 | -0.4705729 | 0.273 | 0.312 | 1 |
| H2az1 | 0.06831235 | -0.3571156 | 0.326 | 0.371 | 1 |
| Ssbp2 | 0.06902643 | -0.4673578 | 0.432 | 0.456 | 1 |
| Nexmif | 0.0691362 | -0.1406269 | 0.469 | 0.406 | 1 |
| Hmgcr | 0.06922162 | -0.1301194 | 0.338 | 0.299 | 1 |
| Pfdn1 | 0.0692329 | 0.11615539 | 0.273 | 0.241 | 1 |
| Ntm | 0.06955884 | 0.17167512 | 0.64 | 0.585 | 1 |
| Map3k4 | 0.07000808 | -0.172528 | 0.311 | 0.276 | 1 |

| | | | | | |
|---|---|---|---|---|---|
| Zfp346 | 0.07041967 | 0.17034959 | 0.254 | 0.223 | 1 |
| Dcaf5 | 0.07059042 | -0.1247564 | 0.375 | 0.331 | 1 |
| Cdc42bpb | 0.0708438 | 0.13232321 | 0.3 | 0.27 | 1 |
| Pnmal2 | 0.07088272 | -0.277803 | 0.256 | 0.215 | 1 |
| St3gal5 | 0.07092603 | 0.20437683 | 0.359 | 0.318 | 1 |
| Copg2 | 0.07111254 | -0.1301266 | 0.39 | 0.347 | 1 |
| Ankhd1 | 0.07150504 | -0.155731 | 0.503 | 0.44 | 1 |
| Atp5md | 0.07157853 | 0.14931013 | 0.468 | 0.42 | 1 |
| Fhit | 0.07176195 | 0.2545503 | 0.329 | 0.295 | 1 |
| Arhgap26 | 0.07199462 | -0.6191029 | 0.266 | 0.291 | 1 |
| Zfp871 | 0.07199799 | -0.1813809 | 0.322 | 0.281 | 1 |
| Actr10 | 0.0721486 | -0.4375131 | 0.231 | 0.271 | 1 |
| Dars | 0.07224539 | -0.1801908 | 0.25 | 0.216 | 1 |
| Ssu72 | 0.07235315 | 0.19958968 | 0.251 | 0.221 | 1 |
| Pabpc1 | 0.07295349 | -0.1636425 | 0.306 | 0.267 | 1 |
| Itpr1 | 0.07341078 | -0.1025216 | 0.544 | 0.485 | 1 |
| Pak2 | 0.07397278 | 0.3300258 | 0.261 | 0.23 | 1 |
| Tnfrsf21 | 0.07401008 | -0.2668783 | 0.297 | 0.342 | 1 |
| Paxbp1 | 0.07438804 | 0.11233914 | 0.372 | 0.33 | 1 |
| Etl4 | 0.07448099 | -0.1783549 | 0.319 | 0.355 | 1 |
| Dop1b | 0.0757304 | -0.1170324 | 0.393 | 0.347 | 1 |
| Tmem38a | 0.07604238 | 0.17949605 | 0.292 | 0.261 | 1 |
| Morf4l2 | 0.07619254 | -0.1497066 | 0.379 | 0.334 | 1 |
| Syt14 | 0.07627206 | 0.22427407 | 0.43 | 0.397 | 1 |
| Nav3 | 0.07653703 | -0.1190174 | 0.861 | 0.862 | 1 |
| Tcerg1 | 0.07762925 | 0.21030277 | 0.285 | 0.257 | 1 |
| Tet3 | 0.07764802 | -0.1114631 | 0.385 | 0.345 | 1 |
| Kdm7a | 0.07768409 | -0.1429534 | 0.468 | 0.42 | 1 |
| Glul | 0.07816231 | -0.4658261 | 0.281 | 0.312 | 1 |
| Plppr2 | 0.07916169 | 0.14852086 | 0.323 | 0.291 | 1 |
| Kcnq3 | 0.07924454 | -0.2908958 | 0.69 | 0.693 | 1 |
| Gucy1b1 | 0.08098905 | 0.13878169 | 0.306 | 0.271 | 1 |
| Aagab | 0.08301509 | 0.27954755 | 0.271 | 0.245 | 1 |
| Slc38a1 | 0.08390047 | 0.39528656 | 0.285 | 0.262 | 1 |
| Exd2 | 0.08403803 | 0.2935575 | 0.255 | 0.23 | 1 |
| Tulp3 | 0.08409426 | 0.10711107 | 0.263 | 0.233 | 1 |
| Rab21 | 0.08478955 | 0.11658376 | 0.261 | 0.231 | 1 |
| Sun1 | 0.08496779 | 0.20717661 | 0.277 | 0.251 | 1 |
| Chrna7 | 0.08503729 | -0.101654 | 0.393 | 0.346 | 1 |
| Atp1b3 | 0.08582526 | -0.122677 | 0.296 | 0.264 | 1 |
| Txndc15 | 0.08720034 | 0.20705396 | 0.263 | 0.235 | 1 |
| Rbbp4 | 0.08747118 | 0.23494647 | 0.261 | 0.234 | 1 |
| Frmd4b | 0.08767386 | -0.3540193 | 0.397 | 0.433 | 1 |

| | | | | | |
|---|---|---|---|---|---|
| Akap6 | 0.08792882 | -0.223964 | 0.735 | 0.694 | 1 |
| Rap1b | 0.08794414 | -0.1240418 | 0.256 | 0.224 | 1 |
| Add1 | 0.0884798 | -0.2824307 | 0.382 | 0.42 | 1 |
| Ift88 | 0.08959152 | 0.1915199 | 0.284 | 0.259 | 1 |
| Ubr2 | 0.089704 | -0.1016826 | 0.415 | 0.377 | 1 |
| Gsdme | 0.08974262 | -0.1208828 | 0.426 | 0.374 | 1 |
| Frmd5 | 0.08983038 | -0.4159681 | 0.442 | 0.46 | 1 |
| Kcnc2 | 0.08998639 | -0.1009599 | 0.477 | 0.522 | 1 |
| Usp19 | 0.0908433 | 0.12549932 | 0.311 | 0.281 | 1 |
| Psmb1 | 0.09095617 | 0.37976073 | 0.28 | 0.259 | 1 |
| Abca3 | 0.09114929 | 0.29870912 | 0.281 | 0.253 | 1 |
| Cdh2 | 0.0911681 | -0.2470746 | 0.611 | 0.581 | 1 |
| Rasa2 | 0.09163035 | -0.1323377 | 0.33 | 0.288 | 1 |
| Lrrfip2 | 0.09171194 | -0.2057342 | 0.295 | 0.26 | 1 |
| D430041D05 | 0.09402747 | -0.3280217 | 0.508 | 0.523 | 1 |
| Zyg11b | 0.09432972 | 0.15929716 | 0.377 | 0.343 | 1 |
| Qrich1 | 0.09437112 | -0.2223405 | 0.426 | 0.38 | 1 |
| Nin | 0.09438008 | -0.1042543 | 0.508 | 0.436 | 1 |
| Purb | 0.0944805 | -0.1875359 | 0.408 | 0.357 | 1 |
| Tmem63c | 0.09453913 | 0.10841832 | 0.271 | 0.245 | 1 |
| Zfr2 | 0.09484956 | 0.21353774 | 0.265 | 0.237 | 1 |
| Cntnap5c | 0.09533067 | 0.16291468 | 0.486 | 0.441 | 1 |
| Pak7 | 0.09538729 | -0.1027456 | 0.54 | 0.475 | 1 |
| Srr | 0.09583294 | -0.2068401 | 0.326 | 0.288 | 1 |
| Cep112 | 0.09655342 | -0.2102868 | 0.508 | 0.431 | 1 |
| Eef1b2 | 0.0970311 | 0.11935933 | 0.259 | 0.227 | 1 |
| mt-Cytb | 0.09728692 | -0.5188999 | 0.656 | 0.622 | 1 |
| Wdr17 | 0.10128571 | 0.13806667 | 0.486 | 0.431 | 1 |
| Vps13c | 0.10157388 | -0.3629831 | 0.596 | 0.544 | 1 |
| Arih2 | 0.10184588 | -0.3374284 | 0.303 | 0.265 | 1 |
| Tspoap1 | 0.10189831 | -0.1278355 | 0.432 | 0.369 | 1 |
| Gabra2 | 0.10207583 | -0.253664 | 0.72 | 0.635 | 1 |
| Ythdf3 | 0.10289708 | -0.1499055 | 0.359 | 0.322 | 1 |
| Tenm2 | 0.10502513 | -0.1238868 | 0.859 | 0.863 | 1 |
| Vti1a | 0.10511015 | -0.1941746 | 0.557 | 0.486 | 1 |
| Hmgcll1 | 0.10559024 | 0.14600762 | 0.322 | 0.286 | 1 |
| Rbm5 | 0.10588759 | -0.1174795 | 0.551 | 0.48 | 1 |
| Gria4 | 0.10602548 | -0.4461028 | 0.486 | 0.502 | 1 |
| Adss | 0.10651044 | 0.19849105 | 0.27 | 0.242 | 1 |
| Fam126a | 0.106531 | -0.4333404 | 0.228 | 0.257 | 1 |
| Dgkb | 0.10766191 | -0.2515376 | 0.697 | 0.66 | 1 |
| Slc4a3 | 0.10826204 | 0.21186777 | 0.299 | 0.277 | 1 |
| Arhgap32 | 0.10936267 | -0.4035048 | 0.325 | 0.363 | 1 |

| | | | | | |
|---|---|---|---|---|---|
| Rps16 | 0.11036716 | -0.1551112 | 0.393 | 0.342 | 1 |
| Chd5 | 0.11057705 | 0.21842441 | 0.28 | 0.259 | 1 |
| Khdrbs2 | 0.11069557 | -0.4114567 | 0.423 | 0.444 | 1 |
| Nr6a1 | 0.11232637 | 0.10256666 | 0.402 | 0.367 | 1 |
| Pcsk7 | 0.11260694 | 0.11817598 | 0.282 | 0.258 | 1 |
| Gnptab | 0.11279612 | -0.5716987 | 0.24 | 0.273 | 1 |
| Sars | 0.11286459 | 0.13941052 | 0.261 | 0.235 | 1 |
| Tcf4 | 0.11290335 | -0.1068448 | 0.914 | 0.923 | 1 |
| Tlk2 | 0.11292953 | -0.1311917 | 0.345 | 0.311 | 1 |
| Rgs7bp | 0.11296584 | -0.303583 | 0.686 | 0.658 | 1 |
| Fyco1 | 0.11323745 | -0.1522443 | 0.297 | 0.265 | 1 |
| Grm7 | 0.11390295 | -0.1841065 | 0.842 | 0.82 | 1 |
| Sgms1 | 0.11436994 | -0.1913304 | 0.467 | 0.41 | 1 |
| Mbnl2 | 0.11499611 | -0.219284 | 0.734 | 0.722 | 1 |
| Cacnb2 | 0.11673113 | -0.2031916 | 0.793 | 0.746 | 1 |
| Hcn1 | 0.11677314 | -0.3121511 | 0.566 | 0.532 | 1 |
| Ccnh | 0.11689218 | -0.1491908 | 0.342 | 0.309 | 1 |
| Pfkp | 0.11717272 | -0.1443783 | 0.383 | 0.342 | 1 |
| 5530401A14I | 0.11968665 | 0.19585913 | 0.254 | 0.223 | 1 |
| Rasgrp1 | 0.11989512 | -0.1134789 | 0.544 | 0.492 | 1 |
| Gabrg2 | 0.12102281 | -0.1057701 | 0.543 | 0.472 | 1 |
| Rfx7 | 0.12216947 | -0.4645576 | 0.443 | 0.448 | 1 |
| Setd3 | 0.12260231 | 0.27936883 | 0.297 | 0.273 | 1 |
| Arl3 | 0.12289897 | -0.5106165 | 0.348 | 0.375 | 1 |
| Tanc1 | 0.12325129 | -0.3750859 | 0.577 | 0.559 | 1 |
| Ncor2 | 0.12355185 | -0.1773351 | 0.415 | 0.361 | 1 |
| Ogfod1 | 0.12454327 | -0.1111934 | 0.293 | 0.264 | 1 |
| Tcf12 | 0.12467295 | -0.2044997 | 0.615 | 0.57 | 1 |
| Csnk2a2 | 0.12557121 | 0.12206972 | 0.329 | 0.296 | 1 |
| Xpr1 | 0.12558048 | 0.10702158 | 0.383 | 0.35 | 1 |
| Tenm1 | 0.12578966 | -0.2692109 | 0.381 | 0.406 | 1 |
| Chchd3 | 0.1260652 | -0.1553678 | 0.497 | 0.44 | 1 |
| Slc24a3 | 0.12634466 | -0.2202665 | 0.716 | 0.65 | 1 |
| Dmd | 0.12728093 | -0.1873175 | 0.72 | 0.682 | 1 |
| Eef1g | 0.12790304 | 0.18231862 | 0.278 | 0.253 | 1 |
| Cnot2 | 0.1280793 | -0.1588863 | 0.486 | 0.43 | 1 |
| Hnrnpa3 | 0.12824618 | -0.3327306 | 0.407 | 0.433 | 1 |
| Psmd11 | 0.1285423 | -0.1003652 | 0.364 | 0.335 | 1 |
| Selenot | 0.12914091 | 0.16105576 | 0.259 | 0.234 | 1 |
| Hace1 | 0.13114968 | -0.1224027 | 0.404 | 0.367 | 1 |
| Prex2 | 0.13135655 | 0.22731457 | 0.333 | 0.3 | 1 |
| Mdga2 | 0.13389823 | -0.1091737 | 0.859 | 0.846 | 1 |
| Rapgefl1 | 0.13475429 | -0.5986422 | 0.273 | 0.302 | 1 |

| | | | | | |
|---|---|---|---|---|---|
| Vstm2a | 0.13526911 | 0.21290121 | 0.259 | 0.234 | 1 |
| Sgsm2 | 0.1358349 | -0.2544051 | 0.289 | 0.258 | 1 |
| Smurf2 | 0.13586093 | -0.4680393 | 0.291 | 0.323 | 1 |
| Sf1 | 0.13702764 | 0.10610768 | 0.269 | 0.245 | 1 |
| Ndufa5 | 0.13962953 | -0.1813771 | 0.337 | 0.3 | 1 |
| Cox5b | 0.1407861 | -0.1235064 | 0.322 | 0.284 | 1 |
| Ube2d2a | 0.14123771 | -0.2086351 | 0.413 | 0.368 | 1 |
| Traf3ip2 | 0.14288738 | 0.17769248 | 0.266 | 0.242 | 1 |
| Brd4 | 0.14292044 | -0.1092329 | 0.467 | 0.417 | 1 |
| Tubb5 | 0.14300759 | -0.2681923 | 0.296 | 0.256 | 1 |
| Oaz1 | 0.14345368 | -0.1108064 | 0.423 | 0.382 | 1 |
| Prmt8 | 0.14481084 | -0.1553675 | 0.359 | 0.327 | 1 |
| Rbbp7 | 0.14484213 | 0.22303898 | 0.262 | 0.238 | 1 |
| Adora1 | 0.14532903 | -0.5194052 | 0.237 | 0.267 | 1 |
| Ggnbp2 | 0.14634355 | -0.1763425 | 0.437 | 0.388 | 1 |
| Dbi | 0.14661241 | -0.2090231 | 0.241 | 0.275 | 1 |
| Rimbp2 | 0.14667216 | -0.1447462 | 0.611 | 0.528 | 1 |
| Casc4 | 0.14675772 | -0.2025623 | 0.529 | 0.462 | 1 |
| Abhd17b | 0.14744288 | -0.2607773 | 0.357 | 0.32 | 1 |
| Zbtb38 | 0.14884208 | -0.2499204 | 0.357 | 0.324 | 1 |
| Zfp182 | 0.15010613 | 0.16597812 | 0.263 | 0.24 | 1 |
| mt-Nd2 | 0.15149434 | -0.4721958 | 0.629 | 0.593 | 1 |
| Fut8 | 0.1517896 | -0.1767544 | 0.528 | 0.462 | 1 |
| Rbmx | 0.15207875 | 0.10747359 | 0.258 | 0.235 | 1 |
| Dnal1 | 0.15451061 | 0.20560361 | 0.261 | 0.239 | 1 |
| Ywhaz | 0.15554114 | -0.2578875 | 0.667 | 0.653 | 1 |
| Wnk1 | 0.15606541 | -0.1369423 | 0.535 | 0.463 | 1 |
| Mitf | 0.156676 | -0.1244012 | 0.3 | 0.329 | 1 |
| Dcaf1 | 0.15708286 | -0.6834028 | 0.222 | 0.252 | 1 |
| Cry2 | 0.15747753 | 0.24872595 | 0.278 | 0.258 | 1 |
| Rprml | 0.15881564 | 0.20183576 | 0.314 | 0.296 | 1 |
| Ryk | 0.15996267 | -0.2003595 | 0.303 | 0.273 | 1 |
| Lmo4 | 0.16052018 | -0.10465 | 0.378 | 0.337 | 1 |
| Rab6b | 0.1605363 | -0.1053786 | 0.569 | 0.521 | 1 |
| Crk | 0.1616791 | 0.15036729 | 0.297 | 0.271 | 1 |
| Dock11 | 0.16186513 | -0.261358 | 0.293 | 0.316 | 1 |
| Fbxl20 | 0.16272796 | -0.1145481 | 0.39 | 0.35 | 1 |
| Usp31 | 0.16283566 | -0.1236316 | 0.3 | 0.274 | 1 |
| Nrxn1 | 0.16332756 | -0.1080063 | 0.945 | 0.943 | 1 |
| Otud7a | 0.16339199 | -0.1757595 | 0.667 | 0.617 | 1 |
| Arhgap23 | 0.16422555 | -0.2417824 | 0.349 | 0.323 | 1 |
| Syn2 | 0.16658622 | -0.1562428 | 0.791 | 0.754 | 1 |
| Dnm3 | 0.16986747 | -0.3139021 | 0.54 | 0.533 | 1 |

| | | | | | |
|---|---|---|---|---|---|
| Cdh9 | 0.16990929 | -0.2688724 | 0.37 | 0.391 | 1 |
| Strip2 | 0.17004873 | 0.22754495 | 0.292 | 0.263 | 1 |
| Hsp90b1 | 0.17038021 | -0.373753 | 0.427 | 0.454 | 1 |
| Setx | 0.17049826 | -0.1378966 | 0.385 | 0.347 | 1 |
| Glg1 | 0.17207041 | -0.1228753 | 0.497 | 0.446 | 1 |
| Bbip1 | 0.17225398 | -0.1859962 | 0.314 | 0.281 | 1 |
| Pex5l | 0.17280581 | -0.2254252 | 0.613 | 0.59 | 1 |
| St3gal3 | 0.17314387 | -0.200784 | 0.377 | 0.338 | 1 |
| Ccnc | 0.17442064 | 0.1321172 | 0.278 | 0.263 | 1 |
| Tnks2 | 0.1753488 | -0.1608929 | 0.402 | 0.366 | 1 |
| Ap1b1 | 0.17549972 | -0.1800828 | 0.267 | 0.242 | 1 |
| Qk | 0.17572069 | -0.1827088 | 0.589 | 0.526 | 1 |
| Tmem131 | 0.17577256 | -0.1485289 | 0.453 | 0.403 | 1 |
| Gpr158 | 0.17599773 | -0.2303154 | 0.708 | 0.653 | 1 |
| Rhot1 | 0.17722153 | -0.1172157 | 0.318 | 0.294 | 1 |
| Capza2 | 0.17908445 | -0.1003544 | 0.547 | 0.493 | 1 |
| Zfp280c | 0.17937251 | -0.1054995 | 0.256 | 0.232 | 1 |
| Ap3s1 | 0.17976496 | 0.15275693 | 0.333 | 0.308 | 1 |
| Hectd1 | 0.1803914 | -0.2284131 | 0.42 | 0.383 | 1 |
| Tbc1d1 | 0.18100032 | -0.1030102 | 0.259 | 0.228 | 1 |
| Stag1 | 0.1821489 | -0.3130603 | 0.644 | 0.619 | 1 |
| Smap2 | 0.18283177 | -0.1477848 | 0.344 | 0.313 | 1 |
| Tbc1d32 | 0.18314538 | -0.1048778 | 0.271 | 0.247 | 1 |
| Rfx3 | 0.18445627 | -0.2671098 | 0.707 | 0.696 | 1 |
| Cyb5b | 0.18492183 | -0.1256212 | 0.256 | 0.232 | 1 |
| Zmym4 | 0.1860477 | -0.1181406 | 0.536 | 0.482 | 1 |
| Igsf11 | 0.18692563 | -0.5173273 | 0.225 | 0.251 | 1 |
| Nfyc | 0.1879954 | -0.1365282 | 0.292 | 0.261 | 1 |
| Fkbp1a | 0.18972044 | -0.171852 | 0.465 | 0.425 | 1 |
| Zcchc17 | 0.18991521 | -0.2089125 | 0.277 | 0.248 | 1 |
| P4ha1 | 0.19023017 | -0.217597 | 0.401 | 0.364 | 1 |
| St6galnac5 | 0.1903791 | -0.2016119 | 0.69 | 0.646 | 1 |
| Ap2a2 | 0.19071452 | -0.1537993 | 0.467 | 0.417 | 1 |
| Hipk3 | 0.19099203 | -0.1400487 | 0.269 | 0.246 | 1 |
| Mga | 0.19289786 | -0.360616 | 0.322 | 0.347 | 1 |
| Micos10 | 0.1942437 | -0.1202316 | 0.315 | 0.292 | 1 |
| Clmp | 0.19578411 | 0.26217892 | 0.251 | 0.23 | 1 |
| Mical2 | 0.19711399 | -0.1753496 | 0.688 | 0.625 | 1 |
| Gsk3b | 0.19788951 | -0.1500646 | 0.602 | 0.526 | 1 |
| Grm1 | 0.19908138 | -0.2772488 | 0.69 | 0.639 | 1 |
| Hmgb1 | 0.19970952 | -0.2525753 | 0.457 | 0.482 | 1 |
| Sik2 | 0.20024953 | -0.4281381 | 0.315 | 0.339 | 1 |
| P4htm | 0.20074752 | -0.1113879 | 0.263 | 0.239 | 1 |

| | | | | | |
|---|---|---|---|---|---|
| Znrf2 | 0.20166512 | 0.16702483 | 0.258 | 0.238 | 1 |
| Osbpl10 | 0.20313741 | -0.4161729 | 0.277 | 0.287 | 1 |
| Rbm28 | 0.20420125 | -0.2627196 | 0.28 | 0.312 | 1 |
| Dhx57 | 0.20450164 | -0.1050784 | 0.293 | 0.27 | 1 |
| Aftph | 0.2053678 | -0.128526 | 0.401 | 0.37 | 1 |
| Btf3 | 0.20541941 | 0.1110716 | 0.25 | 0.227 | 1 |
| Kif1b | 0.20572741 | -0.1636605 | 0.783 | 0.728 | 1 |
| Hmg20a | 0.20775222 | -0.2503937 | 0.37 | 0.335 | 1 |
| Nipa1 | 0.20794699 | -0.5936553 | 0.252 | 0.279 | 1 |
| Man1a | 0.21000003 | -0.2258322 | 0.252 | 0.219 | 1 |
| Man2a2 | 0.2109133 | -0.1343514 | 0.27 | 0.248 | 1 |
| Slc44a5 | 0.21248812 | -0.1221414 | 0.572 | 0.48 | 1 |
| Pcp4 | 0.21303566 | -0.2779099 | 0.413 | 0.437 | 1 |
| Far1 | 0.214215 | -0.1344999 | 0.458 | 0.412 | 1 |
| Anxa11 | 0.21423683 | 0.14050625 | 0.273 | 0.257 | 1 |
| Stk3 | 0.21525967 | -0.5898448 | 0.306 | 0.319 | 1 |
| Fat3 | 0.21573111 | -0.1039554 | 0.756 | 0.669 | 1 |
| Sel1l3 | 0.21649203 | -0.5093904 | 0.256 | 0.284 | 1 |
| Nrd1 | 0.21709673 | -0.203788 | 0.461 | 0.415 | 1 |
| Cstf3 | 0.2180516 | -0.1235283 | 0.54 | 0.48 | 1 |
| Cep120 | 0.21830177 | -0.4368511 | 0.286 | 0.261 | 1 |
| Gabrg3 | 0.21839573 | 0.24750007 | 0.281 | 0.255 | 1 |
| Spock2 | 0.21874847 | -0.1235379 | 0.396 | 0.364 | 1 |
| Kctd4 | 0.21917296 | -0.3054629 | 0.261 | 0.286 | 1 |
| Ncdn | 0.2221909 | -0.1605381 | 0.577 | 0.53 | 1 |
| Nudt3 | 0.2223035 | -0.4719558 | 0.235 | 0.258 | 1 |
| Prpf39 | 0.22459075 | 0.10812162 | 0.319 | 0.303 | 1 |
| Ndufb2 | 0.22690231 | -0.1121698 | 0.277 | 0.253 | 1 |
| Cntn1 | 0.22783424 | -0.1840864 | 0.716 | 0.672 | 1 |
| Ctsb | 0.22794802 | -0.18958 | 0.337 | 0.307 | 1 |
| Cpeb3 | 0.22872546 | -0.272865 | 0.602 | 0.564 | 1 |
| Phf21a | 0.22951296 | -0.3150099 | 0.622 | 0.584 | 1 |
| Elavl1 | 0.22963154 | -0.2650753 | 0.262 | 0.237 | 1 |
| Slc24a4 | 0.23473698 | -0.2322962 | 0.33 | 0.308 | 1 |
| Atp6v0e2 | 0.23557375 | -0.1003475 | 0.261 | 0.234 | 1 |
| Camk2n1 | 0.23578584 | -0.3192981 | 0.437 | 0.45 | 1 |
| Syt4 | 0.23738454 | -0.1319437 | 0.312 | 0.285 | 1 |
| Spon1 | 0.23915718 | -0.6166895 | 0.31 | 0.321 | 1 |
| Zfand5 | 0.24298912 | -0.1141764 | 0.445 | 0.414 | 1 |
| Alkbh8 | 0.24462414 | -0.1417204 | 0.274 | 0.252 | 1 |
| Prkcb | 0.24479962 | -0.2204504 | 0.604 | 0.595 | 1 |
| Serpini1 | 0.24587432 | 0.19373968 | 0.261 | 0.243 | 1 |
| Lin7a | 0.24788075 | -0.5489457 | 0.288 | 0.304 | 1 |

| | | | | | |
|---|---|---|---|---|---|
| H3f3b | 0.24933647 | -0.2020208 | 0.484 | 0.448 | 1 |
| Xkr6 | 0.24985421 | -0.2762884 | 0.432 | 0.444 | 1 |
| Kras | 0.25133772 | -0.2320243 | 0.27 | 0.295 | 1 |
| Tnrc6a | 0.25293965 | -0.2190536 | 0.462 | 0.415 | 1 |
| Ndufab1 | 0.25441935 | 0.11873349 | 0.286 | 0.263 | 1 |
| Atp1a1 | 0.25471019 | -0.1480186 | 0.484 | 0.446 | 1 |
| Dazap1 | 0.25528127 | 0.12664171 | 0.256 | 0.238 | 1 |
| Rnf20 | 0.25626225 | -0.2030288 | 0.254 | 0.232 | 1 |
| Pura | 0.25635746 | -0.1275289 | 0.443 | 0.412 | 1 |
| Vezt | 0.25679877 | -0.1257603 | 0.353 | 0.334 | 1 |
| Cpsf6 | 0.25745075 | -0.1315233 | 0.494 | 0.448 | 1 |
| Btbd9 | 0.25963093 | -0.3216958 | 0.698 | 0.664 | 1 |
| Brd1 | 0.26099614 | 0.16567295 | 0.276 | 0.261 | 1 |
| Sppl2a | 0.26191526 | -0.1107906 | 0.266 | 0.248 | 1 |
| Esrrg | 0.26304648 | -0.2256417 | 0.299 | 0.324 | 1 |
| Pdcd4 | 0.26350619 | -0.2590008 | 0.357 | 0.325 | 1 |
| Fip1l1 | 0.2647504 | -0.102437 | 0.307 | 0.283 | 1 |
| Ap3b2 | 0.26636131 | -0.1124738 | 0.327 | 0.31 | 1 |
| Mcu | 0.26647078 | -0.1405062 | 0.544 | 0.466 | 1 |
| Tbc1d5 | 0.26761447 | -0.2866169 | 0.521 | 0.479 | 1 |
| Pld3 | 0.26922414 | 0.18201069 | 0.252 | 0.238 | 1 |
| Eif3c | 0.27044254 | -0.3297606 | 0.229 | 0.255 | 1 |
| D630045J12F | 0.27128869 | -0.2052234 | 0.342 | 0.316 | 1 |
| Nrn1 | 0.27172073 | -0.4332749 | 0.502 | 0.499 | 1 |
| Lyn | 0.27207118 | -0.2319884 | 0.228 | 0.255 | 1 |
| Bicra | 0.27209139 | -0.1621451 | 0.252 | 0.234 | 1 |
| Pcbp2 | 0.27390467 | -0.2536448 | 0.445 | 0.451 | 1 |
| Rpl22 | 0.27527605 | -0.1285471 | 0.258 | 0.238 | 1 |
| Fam189a1 | 0.27675888 | -0.2189625 | 0.66 | 0.608 | 1 |
| Pcnp | 0.27683104 | -0.1215725 | 0.337 | 0.316 | 1 |
| Gna12 | 0.27695148 | -0.1831982 | 0.271 | 0.252 | 1 |
| Dcun1d5 | 0.27732359 | -0.1924512 | 0.284 | 0.261 | 1 |
| Ube2w | 0.27738345 | -0.1404715 | 0.469 | 0.437 | 1 |
| Zfp106 | 0.27847332 | -0.3446744 | 0.326 | 0.356 | 1 |
| Eps15l1 | 0.28070856 | -0.1087462 | 0.359 | 0.335 | 1 |
| Nos1ap | 0.28257234 | -0.2490191 | 0.619 | 0.554 | 1 |
| Rps13 | 0.2826872 | -0.3732447 | 0.387 | 0.4 | 1 |
| Arpc1a | 0.28328262 | -0.2415156 | 0.393 | 0.365 | 1 |
| Tia1 | 0.28425644 | -0.1643099 | 0.565 | 0.504 | 1 |
| Rnf182 | 0.28546293 | -0.202921 | 0.404 | 0.412 | 1 |
| Lrrc4c | 0.28605619 | -0.1962587 | 0.716 | 0.663 | 1 |
| Tmem208 | 0.28882768 | -0.2034685 | 0.259 | 0.239 | 1 |
| Gphn | 0.29036291 | -0.1031462 | 0.921 | 0.88 | 1 |

| | | | | | |
|---|---|---|---|---|---|
| Eml6 | 0.29109296 | -0.2501951 | 0.653 | 0.585 | 1 |
| Dnajc3 | 0.29204678 | -0.1679904 | 0.241 | 0.271 | 1 |
| Prkce | 0.29767345 | -0.1067333 | 0.839 | 0.784 | 1 |
| Atad1 | 0.30166661 | 0.11982061 | 0.333 | 0.321 | 1 |
| Pigk | 0.30172277 | -0.2538297 | 0.674 | 0.597 | 1 |
| Gabra5 | 0.30222857 | -0.3395191 | 0.473 | 0.463 | 1 |
| Dlgap4 | 0.30238692 | -0.1917703 | 0.491 | 0.437 | 1 |
| Epha6 | 0.30325932 | -0.132998 | 0.872 | 0.857 | 1 |
| Dennd2a | 0.30366339 | -0.1759887 | 0.284 | 0.265 | 1 |
| Dst | 0.30390358 | -0.1173848 | 0.735 | 0.646 | 1 |
| Map4 | 0.30451871 | -0.1104857 | 0.538 | 0.502 | 1 |
| Acbd6 | 0.30567179 | -0.2598075 | 0.4 | 0.372 | 1 |
| Mprip | 0.30753354 | -0.126019 | 0.562 | 0.499 | 1 |
| Arhgef12 | 0.30806119 | -0.1181643 | 0.533 | 0.494 | 1 |
| Ptpn4 | 0.3089431 | -0.2431349 | 0.392 | 0.359 | 1 |
| Lrfn2 | 0.31235309 | -0.4144422 | 0.379 | 0.388 | 1 |
| Safb2 | 0.3128288 | -0.1228244 | 0.323 | 0.305 | 1 |
| Ncoa7 | 0.31308655 | -0.1242673 | 0.542 | 0.487 | 1 |
| Rab3ip | 0.31429926 | -0.4088683 | 0.277 | 0.254 | 1 |
| Cabp1 | 0.31466926 | -0.1482359 | 0.396 | 0.357 | 1 |
| Mpdz | 0.31699282 | -0.2658224 | 0.323 | 0.295 | 1 |
| Ddx10 | 0.31743228 | 0.21775141 | 0.291 | 0.276 | 1 |
| Aste1 | 0.31969216 | -0.3542267 | 0.237 | 0.256 | 1 |
| Tomm70a | 0.3217013 | -0.1730002 | 0.281 | 0.266 | 1 |
| Ddn | 0.3223846 | -0.2181153 | 0.446 | 0.4 | 1 |
| Btrc | 0.32287398 | -0.1442431 | 0.426 | 0.392 | 1 |
| Pou2f1 | 0.32304378 | -0.4121406 | 0.321 | 0.341 | 1 |
| Zfp644 | 0.32444408 | -0.1675455 | 0.445 | 0.408 | 1 |
| Psmc6 | 0.32467118 | -0.2003646 | 0.352 | 0.33 | 1 |
| Pafah1b1 | 0.32696391 | -0.2222604 | 0.703 | 0.659 | 1 |
| Esyt2 | 0.32713612 | -0.1561934 | 0.583 | 0.514 | 1 |
| Plekha5 | 0.32721516 | -0.1715328 | 0.647 | 0.573 | 1 |
| Sgk1 | 0.3283046 | -0.3174012 | 0.396 | 0.4 | 1 |
| Efr3a | 0.33033678 | -0.1072686 | 0.462 | 0.414 | 1 |
| Cdc42bpa | 0.33038964 | -0.1538862 | 0.727 | 0.69 | 1 |
| Plppr4 | 0.33283597 | -0.1048193 | 0.693 | 0.627 | 1 |
| Senp7 | 0.33393381 | -0.286405 | 0.506 | 0.465 | 1 |
| Uty | 0.33399794 | 0.16505963 | 0.258 | 0.25 | 1 |
| Naa35 | 0.3354024 | -0.1207604 | 0.286 | 0.271 | 1 |
| Pgam1 | 0.34059014 | -0.1762622 | 0.33 | 0.3 | 1 |
| Pja2 | 0.34221985 | -0.1885631 | 0.602 | 0.537 | 1 |
| Fbxl5 | 0.34351187 | -0.2140816 | 0.375 | 0.355 | 1 |
| Fkbp3 | 0.34449043 | -0.1903038 | 0.262 | 0.284 | 1 |

| | | | | | |
|---|---|---|---|---|---|
| Fam135b | 0.34470038 | -0.3749161 | 0.57 | 0.536 | 1 |
| Lemd3 | 0.34585626 | -0.2100948 | 0.236 | 0.26 | 1 |
| Mmd2 | 0.34674836 | -0.2345709 | 0.304 | 0.283 | 1 |
| Pard3 | 0.34747393 | 0.12537238 | 0.427 | 0.401 | 1 |
| Serbp1 | 0.34938173 | -0.2242344 | 0.387 | 0.415 | 1 |
| Usp40 | 0.34971426 | -0.3302899 | 0.291 | 0.313 | 1 |
| Agtpbp1 | 0.34998166 | -0.1321686 | 0.542 | 0.497 | 1 |
| Cpsf7 | 0.35213566 | -0.1921798 | 0.262 | 0.246 | 1 |
| Calr | 0.35448118 | -0.2663965 | 0.367 | 0.38 | 1 |
| Kcnab2 | 0.3552106 | -0.2170661 | 0.447 | 0.404 | 1 |
| Ppip5k1 | 0.35527784 | -0.1682947 | 0.311 | 0.293 | 1 |
| Pip4k2a | 0.35591636 | -0.2085848 | 0.349 | 0.328 | 1 |
| Raph1 | 0.35718383 | -0.2037681 | 0.447 | 0.422 | 1 |
| Sdk1 | 0.35939909 | 0.10734971 | 0.303 | 0.279 | 1 |
| Arhgap15 | 0.35975338 | -0.2140459 | 0.256 | 0.274 | 1 |
| Rars2 | 0.36083195 | -0.1320773 | 0.289 | 0.274 | 1 |
| Ncald | 0.3615076 | -0.3325564 | 0.595 | 0.578 | 1 |
| Sympk | 0.36154327 | -0.1454266 | 0.273 | 0.259 | 1 |
| Xpo7 | 0.36213592 | -0.3239811 | 0.393 | 0.359 | 1 |
| Sh3bp5 | 0.36312698 | -0.395553 | 0.435 | 0.431 | 1 |
| Smyd3 | 0.36323095 | -0.1114626 | 0.52 | 0.464 | 1 |
| Inpp4b | 0.36544022 | -0.2063244 | 0.345 | 0.312 | 1 |
| Med14 | 0.36891606 | -0.2518704 | 0.307 | 0.289 | 1 |
| Ogfrl1 | 0.36957659 | -0.4796708 | 0.317 | 0.332 | 1 |
| Secisbp2l | 0.36964725 | -0.5674124 | 0.235 | 0.25 | 1 |
| Tubb4a | 0.37022887 | -0.3048366 | 0.304 | 0.284 | 1 |
| Ski | 0.37027796 | -0.1110048 | 0.318 | 0.303 | 1 |
| Chd8 | 0.37083856 | -0.2344686 | 0.367 | 0.341 | 1 |
| Gnaq | 0.37086152 | -0.237506 | 0.73 | 0.669 | 1 |
| Stim2 | 0.37272689 | -0.2134916 | 0.591 | 0.52 | 1 |
| Cdc37l1 | 0.37289587 | -0.2098033 | 0.43 | 0.403 | 1 |
| Cltc | 0.37519705 | -0.1766101 | 0.498 | 0.471 | 1 |
| Scrn1 | 0.37539279 | -0.2813479 | 0.259 | 0.24 | 1 |
| Selenok | 0.37572161 | -0.2027629 | 0.368 | 0.344 | 1 |
| Gbf1 | 0.37650372 | -0.1275915 | 0.46 | 0.415 | 1 |
| Sec23a | 0.37698052 | -0.3133667 | 0.259 | 0.246 | 1 |
| Uqcr10 | 0.37721945 | -0.3175955 | 0.289 | 0.305 | 1 |
| Scn1a | 0.37769969 | -0.1076099 | 0.532 | 0.473 | 1 |
| Apoe | 0.3783572 | -0.1677589 | 0.634 | 0.582 | 1 |
| Btbd3 | 0.37905313 | -0.2587629 | 0.441 | 0.411 | 1 |
| Insr | 0.3795059 | -0.1160151 | 0.367 | 0.339 | 1 |
| Vapa | 0.38158464 | -0.4097841 | 0.359 | 0.375 | 1 |
| Mecp2 | 0.38174924 | 0.12064509 | 0.326 | 0.314 | 1 |

| | | | | | |
|---|---|---|---|---|---|
| Setbp1 | 0.38217997 | -0.1326498 | 0.692 | 0.597 | 1 |
| Srsf1 | 0.3832541 | -0.1415048 | 0.321 | 0.301 | 1 |
| Serinc3 | 0.38328394 | -0.149843 | 0.293 | 0.276 | 1 |
| Galnt13 | 0.38665654 | -0.1534528 | 0.411 | 0.382 | 1 |
| Etnk1 | 0.38774481 | -0.1486654 | 0.546 | 0.502 | 1 |
| Aatk | 0.3906238 | -0.1429608 | 0.338 | 0.319 | 1 |
| Peak1 | 0.39165783 | -0.1605845 | 0.447 | 0.403 | 1 |
| Pclo | 0.39201965 | -0.1136942 | 0.765 | 0.696 | 1 |
| Pknox2 | 0.39289878 | -0.2765196 | 0.258 | 0.278 | 1 |
| Slc1a1 | 0.39496498 | -0.289593 | 0.386 | 0.39 | 1 |
| Rpl10 | 0.39499306 | -0.1607072 | 0.307 | 0.286 | 1 |
| Rps3 | 0.39516995 | -0.260689 | 0.337 | 0.305 | 1 |
| Ppp4r3a | 0.39674825 | -0.209879 | 0.293 | 0.274 | 1 |
| Zfp697 | 0.39980268 | 0.1132956 | 0.258 | 0.244 | 1 |
| Ywhaq | 0.40095356 | -0.2082341 | 0.4 | 0.377 | 1 |
| Slc43a2 | 0.40234428 | -0.4974113 | 0.241 | 0.26 | 1 |
| Ranbp2 | 0.40281377 | -0.4568978 | 0.427 | 0.432 | 1 |
| Ccdc148 | 0.40488155 | -0.2397019 | 0.398 | 0.368 | 1 |
| Dscaml1 | 0.40566978 | -0.1061555 | 0.718 | 0.665 | 1 |
| Fam168b | 0.40850015 | -0.1372058 | 0.289 | 0.276 | 1 |
| Slc25a27 | 0.41022452 | -0.4264762 | 0.297 | 0.314 | 1 |
| Slc25a3 | 0.41071559 | -0.1478535 | 0.499 | 0.456 | 1 |
| Araf | 0.41488606 | -0.1348383 | 0.37 | 0.352 | 1 |
| Tmem108 | 0.41545659 | -0.1594348 | 0.677 | 0.574 | 1 |
| Ccni | 0.41629866 | -0.4662007 | 0.244 | 0.262 | 1 |
| Chrd | 0.41709455 | 0.105323 | 0.282 | 0.278 | 1 |
| Stx5a | 0.4171469 | -0.2599489 | 0.315 | 0.294 | 1 |
| Ppp2r2a | 0.41716239 | -0.2646901 | 0.393 | 0.368 | 1 |
| Eif5 | 0.41775385 | -0.2877084 | 0.259 | 0.241 | 1 |
| Hnrnpa1 | 0.4178807 | -0.2378976 | 0.255 | 0.238 | 1 |
| Aff4 | 0.41797424 | -0.1932774 | 0.55 | 0.499 | 1 |
| Rpl28 | 0.42101123 | -0.2157487 | 0.409 | 0.381 | 1 |
| Zc3h15 | 0.4218388 | -0.1671522 | 0.367 | 0.353 | 1 |
| Kmt2a | 0.42266933 | -0.1043501 | 0.613 | 0.551 | 1 |
| Ssh2 | 0.42305103 | -0.1167323 | 0.666 | 0.586 | 1 |
| Fth1 | 0.42385797 | -0.1190606 | 0.895 | 0.856 | 1 |
| Zfp608 | 0.4248421 | -0.2463662 | 0.482 | 0.449 | 1 |
| Syt16 | 0.42699237 | -0.2144696 | 0.308 | 0.284 | 1 |
| Ric1 | 0.42747263 | -0.1365094 | 0.341 | 0.325 | 1 |
| Agpat3 | 0.42767085 | -0.1333731 | 0.25 | 0.236 | 1 |
| Erlec1 | 0.42849577 | -0.1348739 | 0.308 | 0.293 | 1 |
| Ica1 | 0.43135062 | -0.2241563 | 0.591 | 0.523 | 1 |
| Mff | 0.43375196 | -0.2165806 | 0.271 | 0.258 | 1 |

| | | | | | |
|---|---|---|---|---|---|
| Mkln1 | 0.43379862 | -0.1220174 | 0.503 | 0.466 | 1 |
| Bbx | 0.43444076 | -0.2623658 | 0.345 | 0.364 | 1 |
| Tex2 | 0.43577745 | -0.1489269 | 0.296 | 0.283 | 1 |
| Prrc2c | 0.43656249 | -0.1456204 | 0.554 | 0.511 | 1 |
| Fam214a | 0.43686642 | -0.183681 | 0.349 | 0.332 | 1 |
| Supt5 | 0.43773639 | -0.2038379 | 0.285 | 0.269 | 1 |
| Ptgds | 0.43792002 | -0.6001176 | 0.292 | 0.305 | 1 |
| Prkcg | 0.43805491 | -0.2138677 | 0.508 | 0.459 | 1 |
| Kdm5c | 0.43830192 | -0.2223251 | 0.27 | 0.257 | 1 |
| Gcc2 | 0.44177908 | -0.1132353 | 0.39 | 0.367 | 1 |
| Mia3 | 0.44535828 | -0.1151844 | 0.359 | 0.343 | 1 |
| Zfp280d | 0.44924772 | -0.1718267 | 0.516 | 0.47 | 1 |
| Gpr137c | 0.45059776 | -0.3720802 | 0.28 | 0.293 | 1 |
| Myh9 | 0.45269283 | -0.1398249 | 0.325 | 0.304 | 1 |
| Elp4 | 0.45325543 | -0.2052486 | 0.357 | 0.336 | 1 |
| Ptpra | 0.45339746 | -0.150969 | 0.602 | 0.536 | 1 |
| Dnajc19 | 0.45385799 | -0.2259513 | 0.31 | 0.328 | 1 |
| Fbxw4 | 0.45465806 | -0.2851568 | 0.356 | 0.33 | 1 |
| Reep3 | 0.45578527 | -0.1657793 | 0.261 | 0.249 | 1 |
| Matn2 | 0.45705965 | -0.1381157 | 0.345 | 0.329 | 1 |
| Zfp950 | 0.45871761 | -0.4168909 | 0.375 | 0.384 | 1 |
| Bdnf | 0.46009766 | -0.5133201 | 0.244 | 0.253 | 1 |
| Ncam1 | 0.46021382 | -0.1789955 | 0.563 | 0.576 | 1 |
| Rtf1 | 0.46222278 | -0.1650346 | 0.312 | 0.293 | 1 |
| Fgfr1op2 | 0.46434251 | -0.1248526 | 0.285 | 0.305 | 1 |
| Mbnl1 | 0.46478861 | -0.1689996 | 0.364 | 0.383 | 1 |
| Mycbp2 | 0.46493919 | -0.1472807 | 0.821 | 0.775 | 1 |
| Tmem135 | 0.46548658 | -0.3179305 | 0.441 | 0.403 | 1 |
| Stim1 | 0.4657802 | -0.3491209 | 0.291 | 0.31 | 1 |
| Msi2 | 0.46863671 | -0.1549117 | 0.52 | 0.486 | 1 |
| Ppp2ca | 0.46987762 | -0.1572765 | 0.362 | 0.349 | 1 |
| Morf4l1 | 0.47007093 | -0.250233 | 0.523 | 0.479 | 1 |
| Adamts17 | 0.47160974 | -0.250198 | 0.419 | 0.39 | 1 |
| Dlg4 | 0.47642037 | -0.1549735 | 0.424 | 0.396 | 1 |
| Nol4 | 0.47751292 | -0.1163025 | 0.587 | 0.518 | 1 |
| Rtn4rl1 | 0.47760505 | -0.294858 | 0.297 | 0.282 | 1 |
| Glyr1 | 0.47922438 | -0.159057 | 0.293 | 0.314 | 1 |
| Stard5 | 0.48100491 | -0.2179863 | 0.374 | 0.33 | 1 |
| Nf2 | 0.48176493 | -0.2245841 | 0.342 | 0.324 | 1 |
| Creb1 | 0.48252291 | -0.1499692 | 0.288 | 0.277 | 1 |
| Rbm25 | 0.48364677 | -0.1147052 | 0.553 | 0.515 | 1 |
| Ube2b | 0.48507198 | -0.1807587 | 0.614 | 0.551 | 1 |
| Tsc22d1 | 0.48654844 | -0.106712 | 0.615 | 0.569 | 1 |

| | | | | | |
|---|---|---|---|---|---|
| Acvr2a | 0.48672779 | -0.1115741 | 0.405 | 0.383 | 1 |
| Stx6 | 0.48889414 | -0.4401246 | 0.25 | 0.263 | 1 |
| Tusc3 | 0.4893475 | -0.179051 | 0.42 | 0.428 | 1 |
| Plcl2 | 0.48961693 | -0.2843533 | 0.411 | 0.406 | 1 |
| Bsn | 0.49098795 | -0.3856415 | 0.529 | 0.503 | 1 |
| Nnat | 0.49440881 | -0.1923319 | 0.371 | 0.351 | 1 |
| Rpl7 | 0.49579029 | -0.4747453 | 0.289 | 0.298 | 1 |
| Mrfap1 | 0.49690301 | -0.1978976 | 0.374 | 0.353 | 1 |
| Atp2b2 | 0.49732652 | -0.1760342 | 0.708 | 0.654 | 1 |
| Ranbp9 | 0.49933508 | -0.1509279 | 0.442 | 0.42 | 1 |
| Pik3ca | 0.49965773 | -0.1661657 | 0.282 | 0.268 | 1 |
| Plxna4 | 0.50109285 | -0.1311653 | 0.805 | 0.731 | 1 |
| Map7 | 0.5039856 | -0.2320413 | 0.536 | 0.484 | 1 |
| Kcnc3 | 0.50413283 | -0.5032063 | 0.317 | 0.33 | 1 |
| Hecw2 | 0.50523251 | -0.1625445 | 0.509 | 0.466 | 1 |
| Tjp1 | 0.50966096 | -0.1724804 | 0.647 | 0.58 | 1 |
| Dnaja1 | 0.50997977 | -0.1737504 | 0.468 | 0.443 | 1 |
| Aldoa | 0.51037834 | -0.1036999 | 0.679 | 0.628 | 1 |
| Gatad2b | 0.51092646 | -0.2620625 | 0.524 | 0.487 | 1 |
| Set | 0.51104833 | -0.3144538 | 0.27 | 0.256 | 1 |
| Ppp1r14c | 0.51246725 | -0.2531846 | 0.33 | 0.311 | 1 |
| Rasal2 | 0.5153409 | -0.1293372 | 0.694 | 0.651 | 1 |
| Zc3h12b | 0.51660037 | -0.1801361 | 0.447 | 0.414 | 1 |
| Sdha | 0.51963399 | -0.2144519 | 0.352 | 0.342 | 1 |
| Slc7a6os | 0.52101113 | -0.1216214 | 0.274 | 0.268 | 1 |
| Hecw1 | 0.52168071 | -0.1135214 | 0.441 | 0.412 | 1 |
| Shank1 | 0.52259442 | -0.2050326 | 0.527 | 0.472 | 1 |
| Cwc27 | 0.522622 | -0.1183292 | 0.323 | 0.307 | 1 |
| Stam | 0.52626344 | -0.2829558 | 0.251 | 0.242 | 1 |
| Plekhg5 | 0.52873854 | -0.3982602 | 0.449 | 0.442 | 1 |
| Ankrd28 | 0.53060613 | -0.1107237 | 0.385 | 0.362 | 1 |
| Crym | 0.53097327 | -0.2634819 | 0.28 | 0.295 | 1 |
| Mtmr12 | 0.53211787 | -0.2819747 | 0.374 | 0.376 | 1 |
| Rit2 | 0.53464419 | -0.3547584 | 0.495 | 0.485 | 1 |
| Tomm20 | 0.53916265 | -0.1728806 | 0.375 | 0.353 | 1 |
| Ppargc1a | 0.54241124 | -0.2428743 | 0.278 | 0.266 | 1 |
| D5Ertd579e | 0.54321704 | -0.4360869 | 0.405 | 0.401 | 1 |
| Trank1 | 0.54527651 | -0.273387 | 0.434 | 0.409 | 1 |
| Fmn1 | 0.54603025 | -0.2585735 | 0.356 | 0.34 | 1 |
| Ctcf | 0.54773489 | -0.11075 | 0.263 | 0.256 | 1 |
| Sort1 | 0.5493067 | -0.2541682 | 0.51 | 0.494 | 1 |
| Dnajc8 | 0.55228933 | -0.1709168 | 0.25 | 0.242 | 1 |
| Ndufb3 | 0.55591428 | -0.142393 | 0.351 | 0.335 | 1 |

| | | | | | |
|---|---|---|---|---|---|
| Csnk1a1 | 0.5566662 | -0.1582736 | 0.709 | 0.63 | 1 |
| Dock3 | 0.55786874 | -0.1629832 | 0.638 | 0.579 | 1 |
| Tasor | 0.55835683 | -0.2016185 | 0.367 | 0.351 | 1 |
| Dlg1 | 0.56133444 | -0.3794222 | 0.377 | 0.384 | 1 |
| Hnrnpr | 0.56303698 | -0.1241432 | 0.417 | 0.397 | 1 |
| Mtmr1 | 0.56421467 | -0.1995131 | 0.288 | 0.28 | 1 |
| Itga8 | 0.5681029 | -0.2984932 | 0.387 | 0.365 | 1 |
| Arih1 | 0.56947936 | -0.2261085 | 0.58 | 0.524 | 1 |
| Inpp5f | 0.57288851 | -0.1113814 | 0.25 | 0.239 | 1 |
| Arfgef3 | 0.57756143 | -0.5023038 | 0.499 | 0.473 | 1 |
| Hmbox1 | 0.58073093 | -0.1102634 | 0.46 | 0.428 | 1 |
| Hook3 | 0.58167163 | -0.3010453 | 0.423 | 0.406 | 1 |
| Ppig | 0.58348054 | -0.1086941 | 0.347 | 0.343 | 1 |
| Usp46 | 0.58357044 | -0.3109055 | 0.397 | 0.392 | 1 |
| Samd12 | 0.58416202 | -0.185016 | 0.675 | 0.627 | 1 |
| Syngap1 | 0.58640593 | -0.2673022 | 0.27 | 0.258 | 1 |
| Agpat5 | 0.59220394 | -0.454195 | 0.259 | 0.27 | 1 |
| Gpt2 | 0.59337045 | 0.12815413 | 0.269 | 0.266 | 1 |
| Cdh11 | 0.59692109 | -0.1475771 | 0.614 | 0.517 | 1 |
| Cttnbp2 | 0.59759589 | -0.1092207 | 0.772 | 0.718 | 1 |
| Ppp1r16b | 0.59894042 | -0.6414491 | 0.269 | 0.273 | 1 |
| Mapt | 0.59989714 | -0.1449145 | 0.666 | 0.597 | 1 |
| Lrrc8c | 0.60132933 | -0.2567408 | 0.304 | 0.288 | 1 |
| Vps13a | 0.60249573 | -0.1904749 | 0.562 | 0.506 | 1 |
| Slc7a14 | 0.60323588 | -0.4005126 | 0.372 | 0.355 | 1 |
| Baz2b | 0.60436417 | -0.2143121 | 0.555 | 0.506 | 1 |
| Srrm2 | 0.60446135 | -0.1340829 | 0.72 | 0.643 | 1 |
| Stx1b | 0.60629489 | -0.3362355 | 0.261 | 0.268 | 1 |
| Apobec4 | 0.60688656 | -0.3463146 | 0.267 | 0.271 | 1 |
| Cdk8 | 0.60701719 | -0.1223706 | 0.764 | 0.703 | 1 |
| Srsf10 | 0.6091701 | -0.2232007 | 0.284 | 0.271 | 1 |
| Fam222b | 0.6096002 | -0.1179364 | 0.393 | 0.374 | 1 |
| Micu3 | 0.61060634 | -0.3720565 | 0.435 | 0.407 | 1 |
| Mnat1 | 0.61458668 | -0.1390196 | 0.411 | 0.389 | 1 |
| Htr4 | 0.61814477 | -0.1076175 | 0.377 | 0.378 | 1 |
| Thsd7b | 0.62056813 | -0.5016399 | 0.591 | 0.562 | 1 |
| Mta3 | 0.62095913 | -0.2660512 | 0.286 | 0.3 | 1 |
| Rab6a | 0.62258602 | -0.3401401 | 0.516 | 0.501 | 1 |
| Arf1 | 0.62260739 | -0.1484704 | 0.314 | 0.305 | 1 |
| Farsb | 0.62291654 | -0.1686636 | 0.25 | 0.267 | 1 |
| Gpi1 | 0.62298171 | -0.3377609 | 0.498 | 0.474 | 1 |
| Spen | 0.62796787 | -0.2106249 | 0.262 | 0.275 | 1 |
| Tent2 | 0.62805738 | -0.2984443 | 0.307 | 0.298 | 1 |

| | | | | | |
|---|---|---|---|---|---|
| Sntb2 | 0.63046412 | -0.4049451 | 0.352 | 0.352 | 1 |
| Synpr | 0.63581641 | -0.108639 | 0.413 | 0.386 | 1 |
| Maged1 | 0.63733108 | -0.2181663 | 0.407 | 0.416 | 1 |
| Wdr7 | 0.63823701 | -0.1478879 | 0.569 | 0.508 | 1 |
| Pebp1 | 0.6389447 | -0.1301611 | 0.58 | 0.551 | 1 |
| Unc13a | 0.63914878 | -0.2727291 | 0.479 | 0.431 | 1 |
| Herc2 | 0.63922596 | -0.1906106 | 0.54 | 0.494 | 1 |
| Gtf2f2 | 0.64041332 | -0.4997637 | 0.252 | 0.264 | 1 |
| Rprd2 | 0.64059464 | -0.1874537 | 0.327 | 0.321 | 1 |
| Map1a | 0.64181783 | -0.1012876 | 0.338 | 0.33 | 1 |
| Hdgf | 0.64303798 | -0.4087649 | 0.248 | 0.261 | 1 |
| Slc30a9 | 0.64337355 | -0.2506342 | 0.325 | 0.316 | 1 |
| Rabgap1l | 0.64344109 | -0.1271485 | 0.719 | 0.664 | 1 |
| Herc1 | 0.64361191 | -0.1116669 | 0.644 | 0.582 | 1 |
| Lims1 | 0.64569009 | -0.2280328 | 0.251 | 0.242 | 1 |
| Ubr5 | 0.64576125 | -0.1832317 | 0.63 | 0.563 | 1 |
| Fut9 | 0.64691124 | -0.1928559 | 0.498 | 0.483 | 1 |
| Fam193a | 0.64785462 | -0.2096652 | 0.501 | 0.465 | 1 |
| Zhx3 | 0.65367022 | -0.2447422 | 0.476 | 0.443 | 1 |
| Golga1 | 0.65613214 | -0.2436634 | 0.284 | 0.281 | 1 |
| Kifbp | 0.65664293 | -0.2551168 | 0.291 | 0.282 | 1 |
| D10Wsu102e | 0.65839022 | -0.3731401 | 0.293 | 0.28 | 1 |
| Smarca2 | 0.65846344 | -0.1671625 | 0.408 | 0.391 | 1 |
| Mink1 | 0.65951495 | -0.1313124 | 0.381 | 0.369 | 1 |
| Sbf1 | 0.65970287 | -0.1127008 | 0.25 | 0.246 | 1 |
| Strn | 0.66277389 | -0.435129 | 0.307 | 0.316 | 1 |
| Srpk2 | 0.66310142 | -0.1221934 | 0.591 | 0.548 | 1 |
| Aimp1 | 0.66913956 | -0.16962 | 0.25 | 0.262 | 1 |
| Gtf2h5 | 0.67058414 | -0.2294037 | 0.256 | 0.248 | 1 |
| C2cd5 | 0.67341807 | -0.2094001 | 0.533 | 0.48 | 1 |
| Rpl5 | 0.6738292 | -0.3261453 | 0.332 | 0.337 | 1 |
| Smarca4 | 0.67595447 | -0.1917772 | 0.392 | 0.378 | 1 |
| Pcbp3 | 0.67624525 | -0.3795005 | 0.274 | 0.281 | 1 |
| Depdc5 | 0.67842667 | -0.2406277 | 0.337 | 0.345 | 1 |
| Foxp1 | 0.6788294 | -0.3047332 | 0.527 | 0.493 | 1 |
| Gdap1 | 0.67901644 | -0.2066408 | 0.312 | 0.306 | 1 |
| Pum1 | 0.6794724 | -0.1405376 | 0.462 | 0.44 | 1 |
| Ppm1a | 0.68176942 | -0.2864191 | 0.317 | 0.307 | 1 |
| Lrrc28 | 0.68250892 | -0.3863432 | 0.302 | 0.29 | 1 |
| Spink10 | 0.68265584 | -0.4473222 | 0.25 | 0.258 | 1 |
| Col4a3bp | 0.68447104 | -0.2986971 | 0.293 | 0.302 | 1 |
| Homer1 | 0.68680275 | -0.4373334 | 0.495 | 0.461 | 1 |
| Zeb1 | 0.69180215 | -0.1760675 | 0.486 | 0.459 | 1 |

| | | | | | |
|---|---|---|---|---|---|
| Emc7 | 0.69420628 | -0.1888922 | 0.33 | 0.323 | 1 |
| Klhl2 | 0.69497319 | -0.3354505 | 0.607 | 0.577 | 1 |
| Ank | 0.69608202 | -0.2709671 | 0.509 | 0.473 | 1 |
| Celf3 | 0.69652485 | -0.2763334 | 0.415 | 0.404 | 1 |
| Scaper | 0.69664151 | -0.3138641 | 0.477 | 0.44 | 1 |
| Pacrg | 0.69949836 | -0.4270368 | 0.27 | 0.275 | 1 |
| Fam149b | 0.70097886 | -0.4047861 | 0.24 | 0.252 | 1 |
| Wasf1 | 0.70098602 | -0.1690263 | 0.693 | 0.634 | 1 |
| Spns2 | 0.70176424 | -0.2898034 | 0.265 | 0.26 | 1 |
| Celf5 | 0.70200402 | -0.1423612 | 0.443 | 0.42 | 1 |
| Mapk8 | 0.70301922 | -0.2436049 | 0.393 | 0.381 | 1 |
| Sbf2 | 0.70332688 | -0.1205476 | 0.495 | 0.494 | 1 |
| Car11 | 0.70675728 | 0.11322588 | 0.269 | 0.27 | 1 |
| Gda | 0.70753723 | -0.4348623 | 0.303 | 0.308 | 1 |
| Khdc4 | 0.70766969 | -0.1377377 | 0.284 | 0.28 | 1 |
| Maml2 | 0.70922725 | 0.12762034 | 0.585 | 0.581 | 1 |
| Spred1 | 0.70962411 | -0.3316515 | 0.28 | 0.293 | 1 |
| Pkp4 | 0.71115032 | -0.2012937 | 0.618 | 0.563 | 1 |
| Strn4 | 0.71332429 | -0.2657426 | 0.349 | 0.336 | 1 |
| Aplp2 | 0.71638031 | -0.1036629 | 0.535 | 0.509 | 1 |
| Lmo3 | 0.71658096 | -0.4183612 | 0.411 | 0.384 | 1 |
| Eml5 | 0.7177051 | -0.1888126 | 0.603 | 0.536 | 1 |
| Tmem178 | 0.71841545 | -0.2057616 | 0.494 | 0.443 | 1 |
| Rnf214 | 0.71916784 | -0.1282876 | 0.25 | 0.251 | 1 |
| Peg3 | 0.71952487 | -0.3198265 | 0.396 | 0.398 | 1 |
| Nsg1 | 0.72142002 | -0.2786368 | 0.295 | 0.281 | 1 |
| Ntrk2 | 0.7220605 | -0.129277 | 0.729 | 0.659 | 1 |
| Klhl32 | 0.72525056 | -0.1774921 | 0.266 | 0.272 | 1 |
| Gfod1 | 0.72616662 | -0.358168 | 0.483 | 0.457 | 1 |
| Rspry1 | 0.72650911 | -0.1568894 | 0.265 | 0.26 | 1 |
| Pdia3 | 0.72744107 | -0.2192544 | 0.323 | 0.316 | 1 |
| Epn1 | 0.72749026 | -0.2169877 | 0.33 | 0.322 | 1 |
| Samd5 | 0.72771714 | -0.2435978 | 0.396 | 0.386 | 1 |
| Smarcc2 | 0.7288767 | -0.4189228 | 0.341 | 0.342 | 1 |
| Zranb1 | 0.7289297 | -0.3469027 | 0.33 | 0.335 | 1 |
| Lrp11 | 0.72941606 | -0.2207473 | 0.318 | 0.311 | 1 |
| Fnip1 | 0.72945562 | -0.2751852 | 0.422 | 0.397 | 1 |
| Trim44 | 0.73030179 | -0.178257 | 0.446 | 0.43 | 1 |
| Glrb | 0.73626467 | -0.153398 | 0.389 | 0.382 | 1 |
| Sclt1 | 0.73662064 | -0.2783838 | 0.396 | 0.38 | 1 |
| Cfdp1 | 0.73881267 | -0.2361702 | 0.319 | 0.314 | 1 |
| Laptm4a | 0.73972234 | -0.1631728 | 0.355 | 0.346 | 1 |
| Trim2 | 0.74036479 | -0.1131922 | 0.744 | 0.68 | 1 |

| | | | | | |
|---|---|---|---|---|---|
| Pde7a | 0.74077447 | -0.1305773 | 0.317 | 0.31 | 1 |
| Usp33 | 0.74138119 | -0.1727246 | 0.377 | 0.38 | 1 |
| Zscan26 | 0.74216664 | -0.3298809 | 0.27 | 0.282 | 1 |
| Tm9sf4 | 0.74411282 | -0.1528962 | 0.292 | 0.294 | 1 |
| Dmtf1 | 0.74413197 | -0.3336351 | 0.273 | 0.27 | 1 |
| Sh3gl2 | 0.74489222 | -0.1935024 | 0.634 | 0.591 | 1 |
| Edem3 | 0.74555605 | -0.2902549 | 0.25 | 0.263 | 1 |
| Gng12 | 0.74724267 | -0.1149066 | 0.288 | 0.284 | 1 |
| Flrt2 | 0.74947676 | -0.1460936 | 0.476 | 0.43 | 1 |
| Atp6v1c1 | 0.7501131 | -0.295922 | 0.303 | 0.295 | 1 |
| Ncoa2 | 0.75299909 | -0.1648126 | 0.517 | 0.473 | 1 |
| Pls3 | 0.75375822 | -0.2000574 | 0.356 | 0.345 | 1 |
| Pitpnm2 | 0.75473468 | -0.2862963 | 0.295 | 0.294 | 1 |
| Cdkl2 | 0.75544595 | -0.1558218 | 0.329 | 0.321 | 1 |
| Csnk1g3 | 0.75734254 | -0.2652029 | 0.422 | 0.413 | 1 |
| Ephb2 | 0.75938002 | -0.1844331 | 0.363 | 0.355 | 1 |
| Mical3 | 0.75945303 | -0.2527301 | 0.495 | 0.451 | 1 |
| Akap10 | 0.76152375 | -0.2777569 | 0.312 | 0.306 | 1 |
| Nalcn | 0.76197036 | -0.1946809 | 0.576 | 0.536 | 1 |
| Rlf | 0.7635181 | -0.2789865 | 0.417 | 0.41 | 1 |
| Socs7 | 0.76573744 | -0.3229326 | 0.262 | 0.272 | 1 |
| Cacng8 | 0.76959273 | -0.3810948 | 0.441 | 0.418 | 1 |
| Cpeb4 | 0.76999399 | -0.1435344 | 0.523 | 0.491 | 1 |
| Usp29 | 0.77237314 | -0.3282625 | 0.269 | 0.269 | 1 |
| Arid1a | 0.77275629 | -0.2579874 | 0.293 | 0.302 | 1 |
| Mtch2 | 0.77281145 | -0.1916226 | 0.307 | 0.302 | 1 |
| Kcnb1 | 0.77510231 | -0.1191026 | 0.299 | 0.306 | 1 |
| mt-Co3 | 0.77625227 | -0.1380886 | 0.973 | 0.92 | 1 |
| Sh3kbp1 | 0.77769621 | -0.3100078 | 0.392 | 0.369 | 1 |
| Dhx15 | 0.78243307 | -0.1121328 | 0.271 | 0.289 | 1 |
| Virma | 0.7829933 | -0.1347903 | 0.302 | 0.298 | 1 |
| Ccbe1 | 0.78724508 | -0.2946912 | 0.52 | 0.456 | 1 |
| Agap3 | 0.78748135 | -0.2646936 | 0.311 | 0.322 | 1 |
| Syt7 | 0.78756625 | -0.1817788 | 0.55 | 0.525 | 1 |
| Abhd2 | 0.78913712 | -0.1117167 | 0.258 | 0.259 | 1 |
| Olfml2b | 0.78975654 | -0.1683148 | 0.289 | 0.292 | 1 |
| Ppm1b | 0.79059186 | -0.2724548 | 0.276 | 0.275 | 1 |
| Ldlrad4 | 0.79330478 | -0.1629716 | 0.576 | 0.514 | 1 |
| Enox1 | 0.7940819 | -0.1016732 | 0.744 | 0.708 | 1 |
| Il1rap | 0.79523909 | -0.1554715 | 0.477 | 0.457 | 1 |
| Scaf8 | 0.79551918 | -0.2195522 | 0.453 | 0.434 | 1 |
| Ptges3 | 0.7976345 | -0.1581047 | 0.263 | 0.269 | 1 |
| Aak1 | 0.79779469 | -0.2033738 | 0.685 | 0.606 | 1 |

| | | | | | |
|---|---|---|---|---|---|
| Marf1 | 0.79781322 | -0.2911486 | 0.277 | 0.273 | 1 |
| Slc38a9 | 0.79893466 | -0.3152656 | 0.329 | 0.334 | 1 |
| Sarnp | 0.80214273 | -0.2427062 | 0.341 | 0.349 | 1 |
| Ccdc88a | 0.80390221 | -0.1444048 | 0.644 | 0.585 | 1 |
| Actr3 | 0.80495643 | -0.1909778 | 0.532 | 0.49 | 1 |
| Dclk2 | 0.80540359 | -0.3013388 | 0.364 | 0.364 | 1 |
| Kcna1 | 0.80555936 | -0.2067782 | 0.321 | 0.319 | 1 |
| Nav1 | 0.80748931 | -0.3787728 | 0.487 | 0.449 | 1 |
| Arid1b | 0.81138105 | -0.1960917 | 0.709 | 0.632 | 1 |
| Cul3 | 0.81175914 | -0.2720778 | 0.411 | 0.4 | 1 |
| Ablim2 | 0.81481172 | -0.2389347 | 0.374 | 0.363 | 1 |
| Zfpm2 | 0.81501656 | -0.1479108 | 0.553 | 0.534 | 1 |
| Dlgap3 | 0.81781076 | -0.1649181 | 0.31 | 0.314 | 1 |
| Stk38 | 0.81845207 | -0.1675951 | 0.315 | 0.323 | 1 |
| Asxl1 | 0.81888697 | -0.2135614 | 0.377 | 0.373 | 1 |
| Tead1 | 0.82158607 | -0.1968182 | 0.265 | 0.261 | 1 |
| Arhgef2 | 0.82168421 | -0.2176419 | 0.303 | 0.302 | 1 |
| Atg4c | 0.82314279 | -0.1617395 | 0.33 | 0.328 | 1 |
| Atp6v1g2 | 0.82553739 | -0.3228848 | 0.262 | 0.255 | 1 |
| P2ry12 | 0.82566432 | -0.1187079 | 0.286 | 0.292 | 1 |
| Disp2 | 0.82614201 | -0.4138259 | 0.297 | 0.301 | 1 |
| Rbms3 | 0.82712152 | -0.100224 | 0.387 | 0.38 | 1 |
| Hsph1 | 0.82721869 | -0.3029576 | 0.3 | 0.296 | 1 |
| Gm21992 | 0.82737952 | -0.2436771 | 0.326 | 0.322 | 1 |
| Neto1 | 0.82834438 | -0.2066098 | 0.499 | 0.468 | 1 |
| Hdac8 | 0.83091442 | -0.1904072 | 0.452 | 0.426 | 1 |
| Phlpp1 | 0.83351872 | -0.4249514 | 0.475 | 0.458 | 1 |
| Limch1 | 0.83435186 | -0.309473 | 0.469 | 0.446 | 1 |
| Dnajc1 | 0.83496429 | -0.1388506 | 0.619 | 0.557 | 1 |
| Ankrd11 | 0.83611548 | -0.2859846 | 0.57 | 0.507 | 1 |
| Vps35l | 0.8367076 | -0.1206882 | 0.27 | 0.271 | 1 |
| Rtn4 | 0.83802278 | -0.1339553 | 0.742 | 0.667 | 1 |
| Cox6a1 | 0.84362342 | -0.128993 | 0.306 | 0.305 | 1 |
| Hpca | 0.84445002 | -0.1009422 | 0.648 | 0.619 | 1 |
| mt-Nd1 | 0.8453967 | -0.2131504 | 0.484 | 0.461 | 1 |
| Ar | 0.84565061 | -0.2065911 | 0.252 | 0.247 | 1 |
| Tbc1d24 | 0.84686362 | -0.3878774 | 0.263 | 0.269 | 1 |
| Ptpn11 | 0.84760997 | -0.1864381 | 0.277 | 0.276 | 1 |
| Dzank1 | 0.84966115 | -0.2856651 | 0.261 | 0.261 | 1 |
| Epb41l1 | 0.85181874 | -0.1419688 | 0.486 | 0.469 | 1 |
| Sptlc1 | 0.85318196 | -0.1828394 | 0.248 | 0.251 | 1 |
| Akap9 | 0.8535788 | -0.2650471 | 0.381 | 0.37 | 1 |
| Prkar1b | 0.8561408 | -0.1035912 | 0.322 | 0.321 | 1 |

| | | | | | |
|---|---|---|---|---|---|
| Slc9a9 | 0.8576702 | -0.2360625 | 0.362 | 0.355 | 1 |
| Me3 | 0.859363 | -0.258193 | 0.506 | 0.445 | 1 |
| Septin7 | 0.86436061 | -0.4562974 | 0.449 | 0.43 | 1 |
| Nrp1 | 0.8644243 | -0.162639 | 0.603 | 0.544 | 1 |
| Sybu | 0.86453753 | -0.2878285 | 0.577 | 0.537 | 1 |
| Phc2 | 0.867163 | -0.2625107 | 0.318 | 0.322 | 1 |
| Vamp2 | 0.86801711 | -0.1227325 | 0.302 | 0.309 | 1 |
| Tasor2 | 0.87491866 | -0.1789121 | 0.288 | 0.294 | 1 |
| Tpm3 | 0.87661876 | -0.1705835 | 0.341 | 0.342 | 1 |
| Tsga10 | 0.87678157 | -0.1163521 | 0.332 | 0.333 | 1 |
| Tulp4 | 0.87934803 | -0.2643409 | 0.432 | 0.417 | 1 |
| Ripor2 | 0.88147505 | -0.3157035 | 0.415 | 0.393 | 1 |
| Upp2 | 0.88385098 | -0.2506503 | 0.602 | 0.556 | 1 |
| Diaph2 | 0.88448176 | -0.2038445 | 0.565 | 0.53 | 1 |
| Pkd1 | 0.88629045 | -0.4113304 | 0.375 | 0.357 | 1 |
| Tpt1 | 0.88963778 | -0.119887 | 0.606 | 0.576 | 1 |
| Adcy5 | 0.89063779 | -0.122974 | 0.258 | 0.262 | 1 |
| Zmynd8 | 0.89088945 | -0.181594 | 0.574 | 0.534 | 1 |
| Ociad2 | 0.89125499 | -0.1520383 | 0.386 | 0.378 | 1 |
| Arhgap5 | 0.89169328 | -0.1806465 | 0.468 | 0.46 | 1 |
| Tenm3 | 0.89228152 | -0.2988436 | 0.304 | 0.294 | 1 |
| Taf15 | 0.89239329 | -0.264999 | 0.408 | 0.398 | 1 |
| Snrnp48 | 0.8934179 | -0.1107018 | 0.317 | 0.323 | 1 |
| Pank2 | 0.89385084 | -0.1353092 | 0.261 | 0.27 | 1 |
| Cpne7 | 0.89415989 | -0.1841015 | 0.34 | 0.324 | 1 |
| Myo9a | 0.89773045 | -0.1898671 | 0.63 | 0.571 | 1 |
| Acox1 | 0.89802312 | -0.2565163 | 0.255 | 0.255 | 1 |
| Atp5k | 0.9004552 | -0.1657744 | 0.345 | 0.347 | 1 |
| Nsg2 | 0.90292843 | -0.1727546 | 0.583 | 0.539 | 1 |
| Tef | 0.90326287 | -0.2262226 | 0.314 | 0.31 | 1 |
| Chordc1 | 0.9049059 | -0.289323 | 0.27 | 0.27 | 1 |
| Dennd1b | 0.90892309 | -0.177074 | 0.348 | 0.352 | 1 |
| Mapre2 | 0.91030571 | -0.3842861 | 0.389 | 0.378 | 1 |
| Wdr37 | 0.91041184 | -0.2295253 | 0.333 | 0.336 | 1 |
| Bex2 | 0.91077582 | -0.1707628 | 0.431 | 0.422 | 1 |
| Plekhm2 | 0.91293277 | -0.2399443 | 0.284 | 0.288 | 1 |
| Golga4 | 0.91578344 | -0.3062483 | 0.292 | 0.298 | 1 |
| Acvr1 | 0.91619774 | -0.2726613 | 0.381 | 0.376 | 1 |
| Dpysl2 | 0.91713588 | -0.2233927 | 0.494 | 0.481 | 1 |
| Dgkh | 0.91790995 | -0.1234736 | 0.588 | 0.56 | 1 |
| Eif1 | 0.91837994 | -0.2015726 | 0.557 | 0.53 | 1 |
| Lyst | 0.91944346 | -0.2181905 | 0.506 | 0.492 | 1 |
| Camkmt | 0.92160634 | -0.2566238 | 0.45 | 0.438 | 1 |

| | | | | | |
|---|---|---|---|---|---|
| Ralgapa2 | 0.92236419 | -0.1840097 | 0.574 | 0.536 | 1 |
| Spock3 | 0.92424288 | -0.2375266 | 0.442 | 0.431 | 1 |
| Atf7ip | 0.92495515 | -0.2099066 | 0.345 | 0.351 | 1 |
| Ndufc2 | 0.92587306 | -0.1640526 | 0.269 | 0.271 | 1 |
| Eps15 | 0.92921678 | -0.347656 | 0.442 | 0.42 | 1 |
| mt-Co2 | 0.93058617 | -0.2228183 | 0.895 | 0.82 | 1 |
| Arhgap20 | 0.93288489 | -0.4303904 | 0.261 | 0.258 | 1 |
| Ube2q2 | 0.93429335 | -0.2543328 | 0.251 | 0.252 | 1 |
| Slc9a7 | 0.9398282 | -0.2512385 | 0.311 | 0.311 | 1 |
| Fbxl16 | 0.94114546 | -0.2380135 | 0.393 | 0.382 | 1 |
| Arhgap12 | 0.94119093 | -0.2180221 | 0.261 | 0.253 | 1 |
| Nav2 | 0.94245079 | -0.2402101 | 0.714 | 0.643 | 1 |
| Hnrnpc | 0.94669407 | -0.4280824 | 0.426 | 0.411 | 1 |
| Trp53bp1 | 0.95331786 | -0.1755281 | 0.306 | 0.307 | 1 |
| Med15 | 0.95359793 | -0.2487199 | 0.293 | 0.295 | 1 |
| Akap11 | 0.9536695 | -0.2408528 | 0.505 | 0.489 | 1 |
| Ildr2 | 0.95741062 | -0.2620581 | 0.326 | 0.32 | 1 |
| Cblb | 0.95842443 | -0.1211623 | 0.342 | 0.34 | 1 |
| Hp1bp3 | 0.95892951 | -0.3251333 | 0.299 | 0.303 | 1 |
| Abhd6 | 0.95967871 | -0.1792079 | 0.297 | 0.299 | 1 |
| Ubl3 | 0.95985274 | -0.1421065 | 0.413 | 0.403 | 1 |
| Scd2 | 0.96140658 | -0.295875 | 0.438 | 0.415 | 1 |
| Ndrg3 | 0.96213652 | -0.1125533 | 0.602 | 0.542 | 1 |
| Tcp11l1 | 0.96432829 | -0.1760325 | 0.347 | 0.35 | 1 |
| Rnf112 | 0.9644266 | -0.1591798 | 0.514 | 0.492 | 1 |
| Cbx5 | 0.96557723 | -0.1850525 | 0.319 | 0.323 | 1 |
| Slf2 | 0.96635866 | -0.1946526 | 0.355 | 0.347 | 1 |
| Mon2 | 0.96694173 | -0.2594009 | 0.396 | 0.386 | 1 |
| Nckap1 | 0.96762594 | -0.1258808 | 0.739 | 0.693 | 1 |
| Snapc3 | 0.97040455 | -0.3184424 | 0.304 | 0.301 | 1 |
| Prpf40a | 0.9716007 | -0.3719938 | 0.407 | 0.387 | 1 |
| Rps7 | 0.97178922 | -0.2353947 | 0.471 | 0.466 | 1 |
| Tcaf1 | 0.97180558 | -0.275156 | 0.514 | 0.489 | 1 |
| Rps5 | 0.97227112 | -0.221398 | 0.278 | 0.279 | 1 |
| Asap2 | 0.9729871 | -0.2586319 | 0.266 | 0.264 | 1 |
| Mrps9 | 0.97354238 | -0.1329157 | 0.27 | 0.277 | 1 |
| Sik3 | 0.97358872 | -0.1992512 | 0.589 | 0.536 | 1 |
| Urgcp | 0.97785174 | -0.2467808 | 0.302 | 0.305 | 1 |
| Sf3b2 | 0.97915306 | -0.168091 | 0.289 | 0.294 | 1 |
| Ap3b1 | 0.97971698 | -0.2517567 | 0.408 | 0.398 | 1 |
| Cnih2 | 0.98118933 | -0.1335056 | 0.288 | 0.29 | 1 |
| Plcl1 | 0.98243437 | -0.1992037 | 0.696 | 0.644 | 1 |
| Dynll2 | 0.98376252 | -0.1287157 | 0.28 | 0.28 | 1 |

| | | | | | |
|---|---|---|---|---|---|
| Slc8a2 | 0.9860054 | -0.1329571 | 0.37 | 0.374 | 1 |
| Psd | 0.98860172 | -0.1614114 | 0.351 | 0.349 | 1 |
| Manf | 0.98938419 | -0.2372134 | 0.416 | 0.41 | 1 |
| Acadsb | 0.99037269 | -0.3259983 | 0.267 | 0.266 | 1 |
| Soga3 | 0.99995907 | -0.2635067 | 0.468 | 0.445 | 1 |
